# PGRMC1 phosphorylation and cell plasticity 1: glycolysis, mitochondria, tumor growth

**DOI:** 10.1101/737718

**Authors:** Bashar M. Thejer, Partho P. Adhikary, Amandeep Kaur, Sarah L. Teakel, Ashleigh Van Oosterum, Ishith Seth, Marina Pajic, Kate M. Hannan, Megan Pavy, Perlita Poh, Jalal A. Jazayeri, Thiri Zaw, Dana Pascovici, Marina Ludescher, Michael Pawlak, Juan C. Cassano, Lynne Turnbull, Mitra Jazayeri, Alexander C. James, Craig P. Coorey, Tara L. Roberts, Simon J. Kinder, Ross D. Hannan, Ellis Patrick, Mark P. Molloy, Elizabeth J. New, Tanja N. Fehm, Hans Neubauer, Ewa M. Goldys, Leslie A. Weston, Michael A. Cahill

## Abstract

Progesterone Receptor Membrane Component 1 (PGRMC1) is expressed in many cancer cells, where it is associated with detrimental patient outcomes. It contains phosphorylated tyrosines which evolutionarily preceded deuterostome gastrulation and tissue differentiation mechanisms. Here, we demonstrate that manipulating PGRMC1 phosphorylation status in MIA PaCa-2 (MP) cells imposes broad pleiotropic effects. Relative to parental cells over-expressing hemagglutinin-tagged wild-type (WT) PGRMC1-HA, cells expressing a PGRMC1-HA-S57A/S181A double mutant (DM) exhibited reduced levels of proteins involved in energy metabolism and mitochondrial function, and altered glucose metabolism suggesting modulation of the Warburg effect. This was associated with increased PI3K/Akt activity, altered cell shape, actin cytoskeleton, motility, and mitochondrial properties. An S57A/Y180F/S181A triple mutant (TM) indicated the involvement of Y180 in PI3K/Akt activation. Mutation of Y180F strongly attenuated mouse xenograft tumor growth. An accompanying paper demonstrates altered metabolism, mutation incidence, and epigenetic status in these cells, indicating that PGRMC1 phosphorylation strongly influences cancer biology.

## INTRODUCTION

Progesterone (P4) Receptor Membrane Component 1 (PGRMC1) is a cytochrome b5-related heme-binding protein with multiple functions including interaction with cytochrome P450 enzymes. Briefly, (the reader is referred to previous reviews) non-comprehensive functions briefly membrane trafficking, P4 responsiveness and steroidogenesis, fertility, lipid transport, neural axon migration, synaptic function, and anti-apoptosis. Its subcellular localization can be cytoplasmic, nuclear/nucleolar, mitochondrial, endoplasmic reticulum, cytoplasmic vesicles, or extracellular (Cahill et al., 2016a; Riad et al., 2018; Ryu et al., 2017). It is involved in cell cycle processes at the G1 checkpoint and during mitosis (Luciano et al., 2010; Luciano and Peluso, 2016; Peluso et al., 2014; Sueldo et al., 2015; Terzaghi et al., 2018; Terzaghi et al., 2016), and elevated PGRMC1 expression has been associated with poor prognosis in multiple types of cancer (Ahmed et al., 2010; Cahill et al., 2016a; Losel et al., 2008; Rohe et al., 2009; Ruan et al., 2017; Shih et al., 2019; Willibald et al., 2017).

Predicted binding site motifs for Src homology 2 (SH2) and Src homology 3 (SH3) proteins in PGRMC1 can potentially be negatively regulated by phosphorylation at adjacent casein kinase 2 (CK2) consensus sites (Cahill, 2007; Cahill et al., 2016a; Peluso et al., 2006). However, while CK2 knockdown leads to reduced phosphorylation of the corresponding C-terminal CK2 site of PGRMC2, PGRMC1 phosphorylation at S181 was unaffected by CK2 knockout in C2C12 mouse myoblast cells (Franchin et al., 2018). These and Y180 can all be phosphorylated *in vivo*, and constitute a potential regulated signaling module (Cahill et al., 2016b).

We hypothesized that PGRMC1 is a signal hub protein with wide ranging effects on cancer and general cell biology (Cahill, 2017; Cahill et al., 2016a; Cahill et al., 2016b; Cahill and Medlock, 2017). The highly conserved motif at Y180/S181 arose early in animal evolution concurrently with the embryological organizer of gastrulation (e.g. Spemann-Mangold organizer), and prior to the evolution of deuterostomes (Cahill, 2017; Hehenberger et al., 2019).

This present study was prompted by our discovery of differential PGRMC1 phosphorylation status between estrogen receptor-positive and -negative breast cancers. PGRMC1 was induced in the hypoxic zone of ductal carcinoma *in situ* breast lesions at precisely the time and place that cells require a switch to glycolytic metabolism known as the Warburg effect, leading us to predict a Warburg-mediating role for PGRMC1. Furthermore, a PGRMC1 S57A/S181A double CK2 site mutant (DM, Figure 1A) enabled the survival of peroxide treatment (Neubauer et al., 2008). Sabbir (2019) recently reported that PGRMC1 induced a P4-dependent metabolic change resembling the Warburg effect in HEK293 cells, which was associated with changes in PGRMC1 stability, post-translational modifications, and subcellular locations. PGRMC1 regulation of glucose metabolism is supported by its implicated mediation of the placental P4-dependent shift from aerobic towards anaerobic glucose metabolism in gestational diabetes (Gras et al., 2007), and association with the insulin receptor and glucose transporters (Hampton et al., 2018).

**Figure 1.**
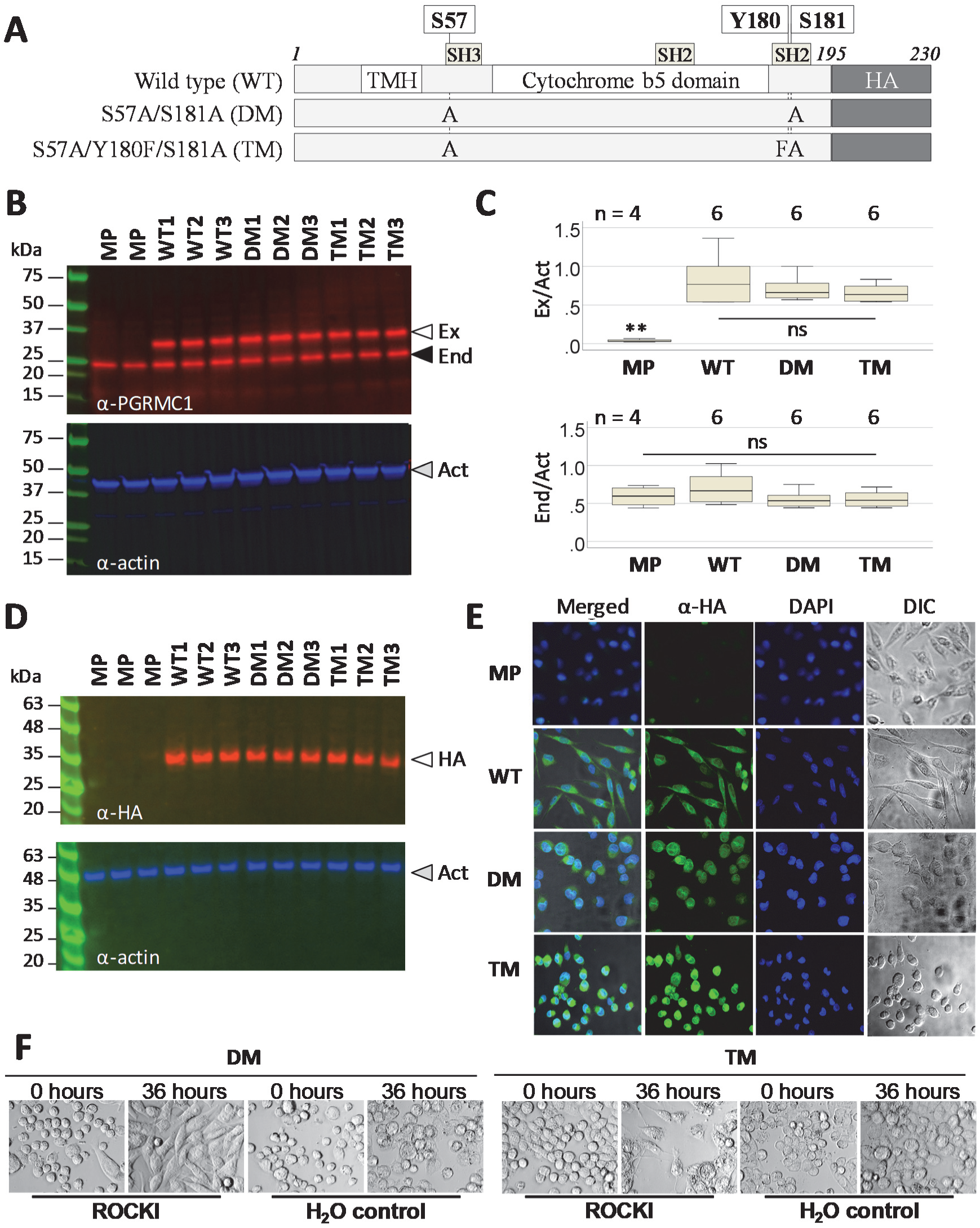
MIA PaCa-2 pancreatic cancer cells morphology is affected by PGRMC1 phosphorylation status. (A) PGRMC1-HA proteins constructed for this figure. TMH: Trans-membrane helix. HA: the C-terminal 3x hemaglutinin tag. (B) Detection of exogenous PGRMC1 expression levels by western blot (upper panel). Equal loading is controlled by quantifying beta actin (lower panel). The results show three totally independent stably transfected cell lines per plasmid from (A). Open arrow: Exogenous PGRMC1-HA (Ex.). Shaded arrow: endogenous PGRMC1 (End.). Filled arrow: beta actin. The molecular weight ladder is Bio-Rad 1610377 Dual Xtra Standards. (C) Box plots quantification of replicate gels of (B) with signals normalized to beta actin from the same respective lanes. n=4 lanes for MP and n=6 for WT, DM and TM (replicates of respective lines 1-3 per condition). There were no significant differences (ns) except for the exogenous band in MP (ANOVA, post-hoc Dunnet’s T3). (D) Western blot quantification of HA-tagged exogenous PGRMC1, following B but detecting PGRMC1 with anti-HA antibody. The molecular weight ladder is Abcam ab116028 Prestained Protein Ladder. (E) PGRMC1 mutant protein expression alters MIA PaCa-2 cell morphology. PGRMC1-HA-expressing stable cells (respective lines 1 from B) or MP cells were stained with a FITC-tagged anti-HA antibody (Anti-HA) and imaged by confocal microscopy. DNA was stained with DAPI. Cells were also imaged in differential interference contrast (DIC) microscopy mode. The respective left panels show merged images of all 3 channels. (F) The rounded phenotype of double and triple mutant (E) was reversed to elongated phenotype after 125µM ROCKI addition, but not by addition of DMSO vehicle control.

We previously observed that MIA PaCa-2 pancreatic cancer (MP) cells (Duong et al., 2013; Han et al., 2008; Iwagami et al., 2013; Yunis et al., 1977) exhibited marked morphological and metabolic changes when the DM protein was expressed (Gosnell et al., 2016b). MP cells exist in culture as a mixed adherent population of elongated “fibroblast-shaped” morphology, a minority population of rounded morphology with bleb-like protrusions, and some multicellular clumps, as well as some rounded suspension cells. They have undergone epithelial-mesenchymal transition (Gradiz et al., 2016), and can further undergo mesenchymal-amoeboid transition (MAT), which requires Rho Kinase-(ROCK)-dependent morphological change from “elongated” mesenchymal cells to rounded amoeboid cells (Fujita et al., 2011).

Here, we examined the effects of altered PGRMC1 phosphorylation status on MP cells to gain insights into PGRMC1-dependent signaling, and its role in subcutaneous mouse xenograft tumorigenesis which requires Y180. In a companion paper (Thejer et al., 2019) we describe differences in metabolism, genomic mutation rates, and epigenetic genomic CpG methylation levels associated with PGRMC1 phosphorylation status in these cells.

## RESULTS

### PGRMC1 phosphorylation status alters cellular morphology

We stably transfected MP cells with the hemagglutinin (HA) epitope-tagged PGRMC1-HA plasmids including the wild-type (WT) sequence (Suchanek et al., 2005), the S57A/S181A DM (Neubauer et al., 2008), or a novel S57A/Y180F/S181A triple mutant (TM), which removed the phosphate acceptor of Y180 (Cahill, 2007, 2017; Cahill et al., 2016b) (Figure 1A). Three independent stable cell lines from each group expressed both 32 kDa 3xHA-tagged exogenous and a 24 kDa endogenous PGRMC1 species, whereas only the 24 kDa species was present in MP cells (Figure 1B-D). Both species were present at approximately equimolar ratios, and an anti-HA antibody detected only the 32 kDa species (Figure 1D). We reason that any consistent differences between biological triplicates should be due to PGRMC1-HA mutations, rather than clonal artifacts. Subsequent experiments were performed using respective cell line triplicates 1-3 per PGRMC1-HA condition.

Like MP cells, freshly seeded WT cells exhibited predominantly elongated cell morphology with some rounded cells. DM and TM cells exhibited primarily rounded morphology (Figure 1E), which was reminiscent of the reported MAT of MP cells (Fujita et al., 2011). After 72 hours of culture the proportion of round cells in DM and TM cultures was reduced, but still elevated relative to WT or MP (not shown). Transient transfections with the DM and TM plasmids (but not WT) led to similar increased levels of cell rounding across the entire populations of cells by 24 hours after transfection (data not shown), indicating that the phosphorylation status of exogenous PGRMC1-HA affects cell morphology.

### PGRMC1-dependent altered morphology requires Rho Kinase

The ROCK pathway is required for amoeboid phenotype and migration and its inhibition reverses MAT in MP cells (Fujita et al., 2011; Matsuoka and Yashiro, 2014). ROCK inhibitor (ROCKI) reversed the rounded phenotype to elongated for DM and TM (Figure 1F), supporting the hypothesis that morphological transition involves altered actin organization. It remains unclear whether the process is truly MAT.

### PGRMC1 phosphorylation affects cell motility and invasion

To further investigate cell plasticity imposed by PGRMC1-HA phosphorylation mutants, we examined cell motility via a scratch assay (Cha et al., 1996). MP cells exhibited the lowest migration, while DM cell migration was substantially greater than other cell lines (Figure 2A-B, File S1). DM cells migrated predominantly as rounded cells, using extended filopodia and small pseudopodia, however, a minority of flattened cells exhibited more pronounced pseudopodia. Video imaging demonstrated that these cell shapes could rapidly interconvert (File S1C). Conversely, DM cells exhibited the lowest ability to invade through a Geltrex pseudo-basal membrane (Figure 2C-D).

**Figure 2.**
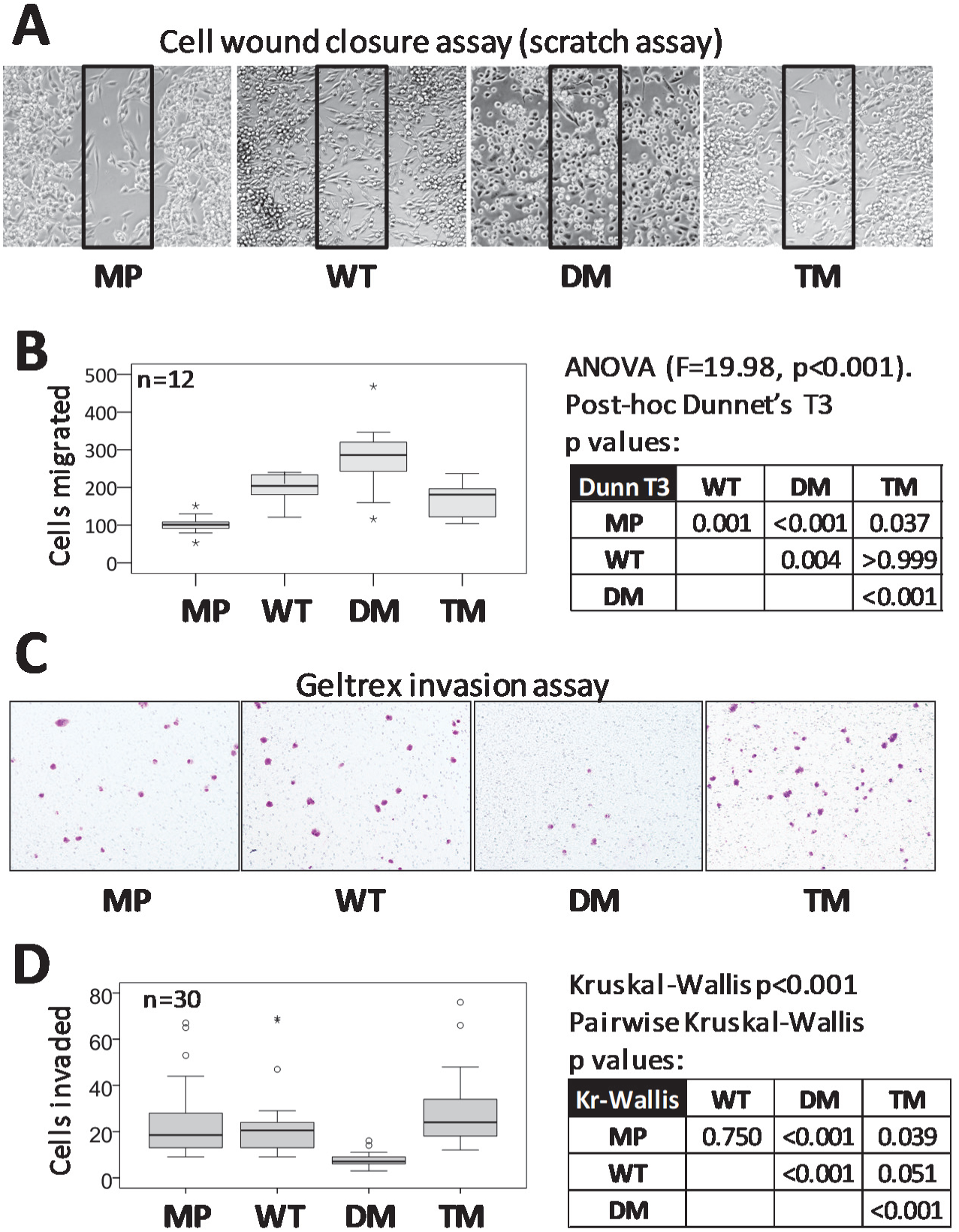
PGRMC1 phosphorylation status affects motility and invasion. (A) DM cells exhibit enhanced motility in scratch assay. Representative results for scratch assays after 36 hr of cell migration. Cells in the boxed area of the scratched void area for each image were counted. (B) Scratch assay cell migration results. Box plots of cell migration into the scratch void box areas depicted in (A) for multiple replicates. n=12 (MP), or 4 replicates each of sublines 1-3 for WT and TM (n=3x4=12). The table shows the results of 1 way ANOVA with post hoc Dunnet’s T3 *p*-values. Video files of cell migration are available in File S1. (C) DM cells exhibit reduced invasion in Geltrex invasion assay. Representative images of crystal-violet stained cells in the lower surface of the transwell insert. (D) Boxplots of invasion assay results from (C) for replicates, as produced by SPSS software. n=30 (MP), or 10 replicates each of sublines 1-3 for WT and TM (n=3x10=30). The table shows the Kruskal-Wallis *p*-values for pairwise comparisons.

### PGRMC1 phosphorylation imposes broad changes

A total of 1330 proteins were reliably identified by proteomics in at least one sample with at least 2 peptides using a combination of information dependent acquisition (IDA) and data independent SWATH-MS acquisition. Results are provided as File S2. Approximately 50% of variation was explained by two principal components (PCs) in PC analysis, which corresponded approximately to “ribosomes and translation” (PC1, separated MP&DM from WT&TM) and “mRNA splicing and processing” (PC2, separated MP&WT from DM&TM) (Figure S1A, File S3). Of the identified proteins, 243 differed by 1.5 fold or more between one or more comparisons with *p*<0.05 (t-test), and 235 of these withstood PC multiple sample correction. The heat map clustering of those 243 proteins (Figure 3, File S6) revealed a suite of proteins which strongly discriminated between the different PGRMC1-HA-induced conditions. Biological replicates clustered tightly in clades of the same cell type, with large distances between clades. We conclude that these differences are primarily specific PGRMC1-HA mutant-dependent effects.

**Figure 3.**
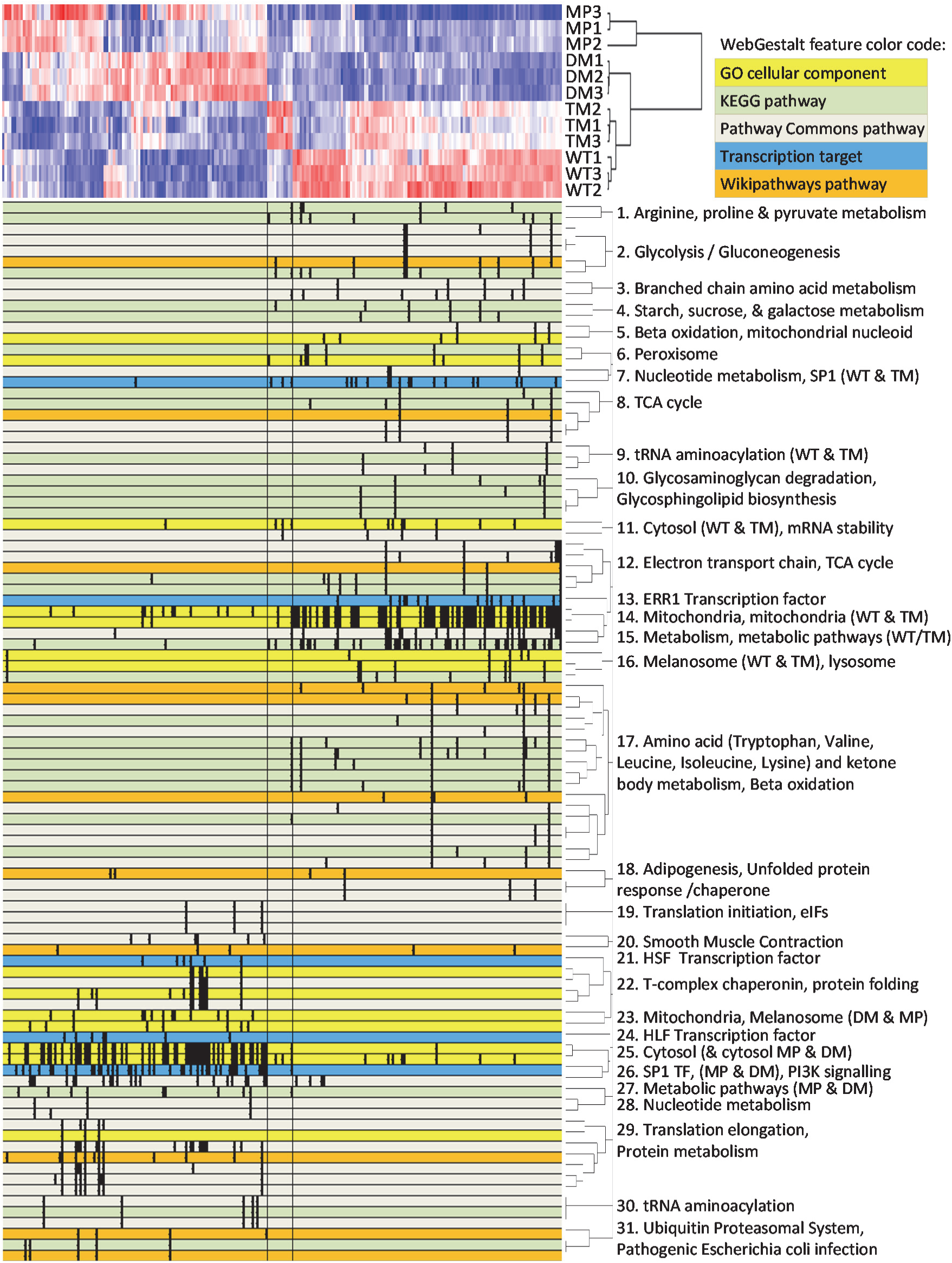
Pathways analysis of SWATH-MS proteomics results. Pathways significantly enriched at the adjP>0.001 level between all 6 comparisons of “red” and “blue” differential proteins (red = higher abundance, blue = lower abundance, white = equal abundance). Top left: the proteomic heat map of 243 significantly differential proteins. A color code for WebGestalt pathways is given at top right. Bottom: WebGestalt pathways mapping. This image is derived from File S6, which contains all protein and WebGestalt pathway identities.

Results from those six comparisons of protein abundance between the four sample types [1) MP v. WT, 2) MP v. DM, 3) MP v. TM, 4) WT v. DM, 5) WT v. TM, and 6) DM v. TM] provide lists of significantly differentially abundant proteins in each pair-wise comparison (File S2). WebGestalt enrichment analyses were performed to identify pathways or features either significantly more or less abundant (respectively the “red” and “blue” lists of proteins for each comparison from columns B of File S4) between each of the respective six comparisons at the Benjamini–Hochberg adjusted *p*-value (adjP)<0.001 level in at least one pairwise comparison. WebGestalt mapped features (File S5) are plotted against the heat map in File S6 (for primary WebGestalt data see File S5 and File S6). The results are schematically mapped against the 243 protein heat map in Figure 3, using all pathways that were detected by WebGestalt in any of the inter-sample comparisons at the adjP<0.001 level, and including all proteins detected in any of those pathways across all 12 comparisons from File S8 at the adjP<0.1 level for respective red and blue protein lists from File S4.

WT PGRMC1-HA protein induced the elevated abundance of many proteins involved in energy metabolism, including proteasomal components involved in protein degradation, and pathways for amino acid, carbohydrate, and fatty acid catabolism (Figure 3). These proteins were annotated as both cytoplasmic and mitochondrial. Peroxisomal and lysosomal proteins were also upregulated in WT cells. A suite of proteins putatively involved in the recognition of mRNA by ribosomes, tRNA aminoacylation, ribosomal protein translation, and chaperone-mediated protein folding, was generally less abundant in WT and TM than MP and DM cells (Figure 3, File S6). Many of the changes in fatty acid and glucose metabolism enzymes resemble the effects of the insulin/glucagon system of metabolic regulation.

Many of the above proteins were inversely regulated in the comparison of WT and DM cells (Figure 3). The energy metabolizing suite of proteins and some proteins associated with translation were less abundant in DM, whereas the T-complex chaperone complex as well as some nuclear exportins and importins were elevated in DM cells (Figure S1D-E). Some differentially enriched pathways were specific for the WT vs. DM comparison. Different enzymes associated with heme metabolism were both up-and down-regulated while enzymes associated with glycosaminoglycan metabolism were up-regulated in WT relative to DM (Figure 3, File S6).

Mitochondrial proteins accounted for a large percentage of the proteins more abundant in WT than DM cells. Intriguingly, many cytoplasmic proteins were more abundant in WT cells than DM (Figure 3, File S6). We also noted higher abundance components of ATP synthase in WT and TM cells (Figure S1C), changes of proteins involved in chaperonin and microtubule function (Figure S1D), and a group of proteins involved in major histocompatibility complex antigen processing and presentation, and proteolysis (Figure S1F). The latter are reduced in DM cells and may be associated with their reduced invasiveness (Figure 2C-D), which requires future confirmation.

Consideration of the extreme (highest and lowest abundance) differential proteins for each cell type offers useful insight into biology (Figure S2). WT and TM cells exhibited overlap in the subset of most abundant proteins, which included PSIP1 transcriptional coactivator, TOM40 mitochondrial import channel, as well as CDIPT which catalyzes the biosynthesis of phosphatidylinositol (circles in Figure S2). One of the WT abundant proteins was phosphofructokinase (UniProt P08237), which catalyzes the rate limiting reaction and first committed step of glycolysis. The most abundant DM proteins included keratin 19, ubiquitin-associated protein 2-like, which is involved in stem cell maintenance (Bordeleau et al., 2014), and methyl-CpG-binding protein 2, which was more abundant in WT and TM cells, suggesting PGRMC1-mediated changes in genomic methylation (see accompanying paper (Thejer et al., 2019)). The least abundant proteins shared a surprising mixed overlap between cell types (triangles in Figure S2). TM and WT cells shared low levels of Ephrin type-A receptor 2, a tyrosine kinase receptor, which was higher in DM (and MP) cells without being a top abundant protein in those cells. WT exhibited low levels of signal recognition particle 54 kDa protein, suggesting altered translation of endoplasmic reticulum proteins, and AL1A1 retinal dehydrogenase. This was notable because both DM and TM exhibited low levels of AL1A3 NAD-dependent aldehyde dehydrogenase involved in the formation of retinoic acid, suggesting alterations in retinoic acid metabolism by mutating S57/S181. DM and TM also shared low levels of ApoC3 and ApoA1 (Figure S2), probably reflecting common lower lipoprotein synthesis by those cells. Taken together, our proteomics analysis revealed significant differences in the abundance of enzymes involved in diverse cell processes, many of which are directly implicated in cancer biology. The resemblance of WT and TM differential proteomics profiles suggests that the DM mutation activates signaling processes that are largely dependent upon Y180. (The sole difference between DM and TM proteins is the phosphate acceptor oxygen of Y180). Overall, this study indicated that PGRMC1 phosphorylation status exerts higher order effects in MP cells.

### ERR1 activity is not directly affected by PGRMC1

Some mitochondrial proteins associated with energy metabolism were predicted by pathways enrichment analysis to be regulated by estrogen receptor related 1 (ERR1) transcription factor (Figure S3A) in the comparisons of DM cells with both WT (adjP=0.004) and TM (adjP=0.04) (File S4). Since ERR1 is a steroid receptor, we investigated any potential link between the biology of PGRMC1 and ERR1 by attenuating ERR1 levels in WT cells by shRNA. This changed cell morphology from predominantly elongated to rounded cells (Figure S3B-D). SWATH-MS proteomics revealed that ERR1 indeed regulated genes differentially abundant between WT and DM cells observed in Figure S3A and Figure 3. However, PGRMC1 phosphorylation status affected the abundance of only a subset of ERR1-driven proteins.

### PGRMC1 phosphorylation affects PI3K/Akt signaling

Strikingly, proteins associated with PI3K/Akt activity from Figure 3 and File S6 were revealed by File S5 to exhibit lower abundance in TM cells (Figure S1B) relative to MP (adjP=0.0063), WT (adjP=0.0001) and DM (adjP=0.0002) cells. We assayed the phosphorylation status of two Akt substrates by reverse phase protein array (RPPA). Bad is phosphorylated by both PKA at S112 (Harada et al., 1999), and at S136 by Akt (Hayakawa et al., 2000). Whereas there was no significant difference between Bad S112 phosphorylation between DM and TM (Figure 4A), phosphorylated S136 levels were lower in TM cells than in all other cells (*p*<0.001, Figure 4B). It was not possible to reliably quantify these levels relative to Bad itself since Bad signals were too low (not shown). One of the best attested substrates of Akt is glycogen synthase kinase 3 beta (GSK3β), which is phosphorylated on S9 by Akt leading to inactivation of GSK3β (Cross et al., 1995). Levels of phosphorylated GSK3β S9 were elevated in WT over MP cells, elevated once more by removing the inhibitory CK2 sites in the DM mutation, and reduced by the further mutation of Y180 in TM mutant cells (Figure 4C-E). These results strengthen the model that PI3K/Akt pathway which is activated in DM cells requires phosphorylated PGRMC1 Y180, and is therefore attenuated in TM cells.

**Figure 4.**
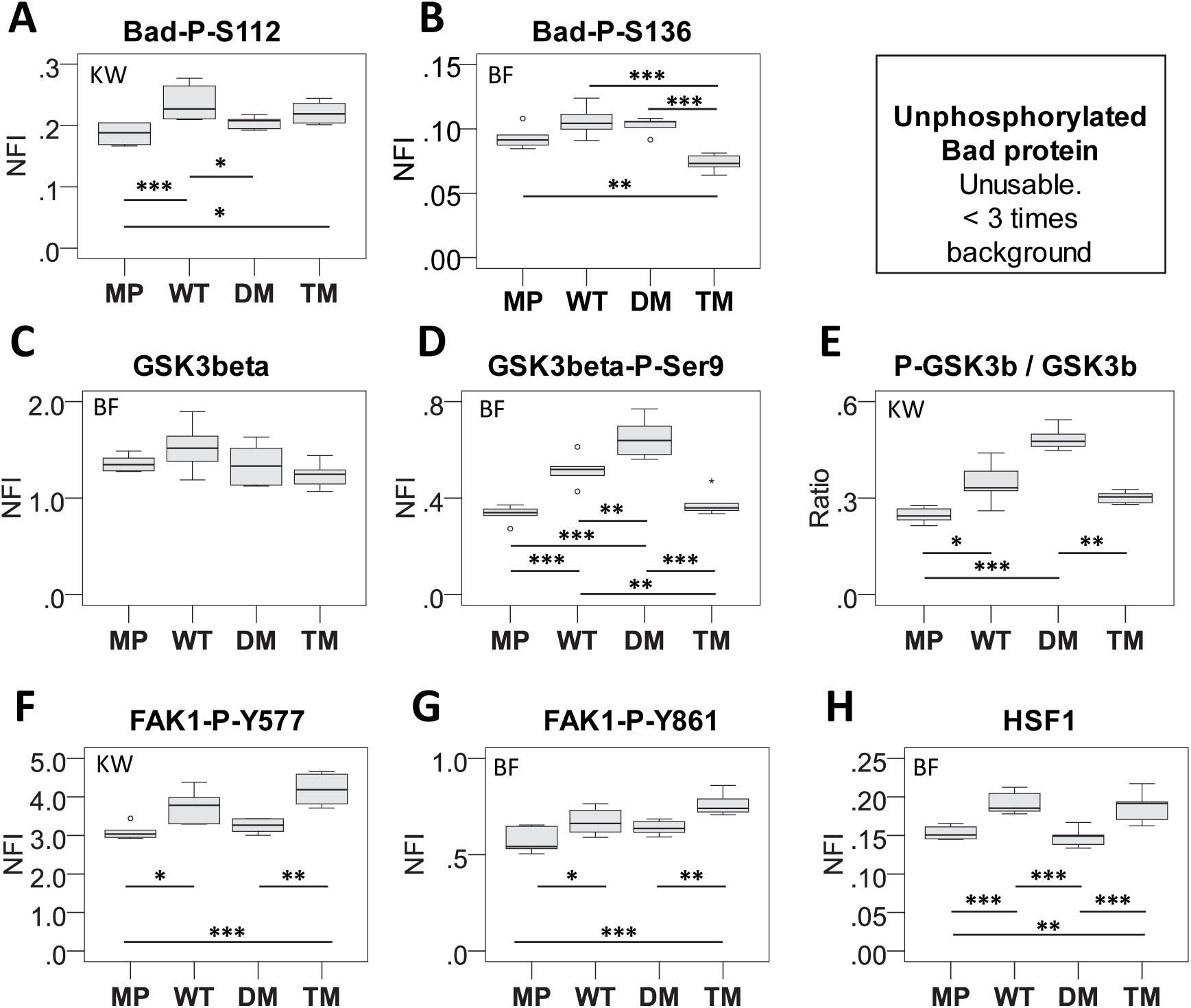
RPPA measurements of protein and phosphorylated protein levels. (A-D, F-H) Average reverse phase protein array (RPPA) normalized fluorescent intensity (NFI) from the indicated antibodies (described in supplemental methods) is plotted from 6 replicate measurements. NFI is normalized to protein content. Statistical calculations for normally distributed data were made using one way ANOVA, and post-hoc Bonferroni (BF) for equal variances (all variances were equal). For non-parametric data, Kruskall-Wallis (KW) pairwise comparisons were calculated for 24 unrelated samples. * *p*<0.05; ** *p*<0.01, *** *p*<0.001. Non-phosphorylated Bad levels could not be accurately determined because signal values were less than three times background. (E) The ratio of average NFI of D relative to C. Labels follow the above.

### PGRMC1 phosphorylation affects FAK activation and HSF levels

Pathways mapping (File S5) suggested that transcription factor HSF1 activity could be involved in the difference between WT v. DM (adjP=0.006). HSF1 has been linked with Focal Adhesion Kinase (FAK) activity (Antonietti et al., 2017), and FAK activity is dependent upon Rho/ROCK signaling which influences focal adhesion dynamics and tumor cell migration and invasion (Joshi et al., 2008). Reverse Phase Protein Array (RPPA) measurements showed that FAK1 tyrosine phosphorylation and increased HSF1 levels were all significantly elevated in WT and TM (Figure 4F-H). Notably, this profile resembled the differential proteomics profile of Figure 3, rather than the ROCK-dependent rounded morphology of Figure 1D.

### Enhanced DM cell motility requires vinculin

Proteins of the actin cytoskeleton were more abundant in DM cells (Figure S4A), one of which was vinculin (Figure S1B), an actin filament-binding protein associated with cell differentiation status, locomotion, and PI3K/Akt, E-cadherin, and β-catenin-regulated Wnt signaling in colon carcinoma (Le Clainche et al., 2010; Pal et al., 2019). We attenuated vinculin levels in MP, WT, and DM cells via shRNA. Scrambled shRNA control (shScr) DM cells exhibited elevated scratch assay motility relative to MP cells. However in anti-vinculin shRNA (shVCL) cells, both MP and DM cell motility was reduced (Figure S4). These results are consistent with elevated levels of proteins involved in the actin cytoskeleton (Figure S4A) contributing directly to the enhanced motility of DM cells (Figure 1F-G). However, that hypothesis remains untested except for vinculin.

### PGRMC1 affects glucose metabolism

Figure 3 predicted altered glycolysis activity, which we investigated by glucose uptake and lactate production assays. Expression of all PGRMC1-HA proteins (WT, DM, and TM) led to significantly lower levels of both measures relative to MP, with DM cells exhibiting the lowest levels (Figure 5A-B). PGRMC1 phosphorylation status regulates both features, consistent with recently reported regulation of Warburg metabolism by PGRMC1 (Sabbir, 2019), which we confirm is regulated by PGRMC1 phosphorylation status.

**Figure 5.**
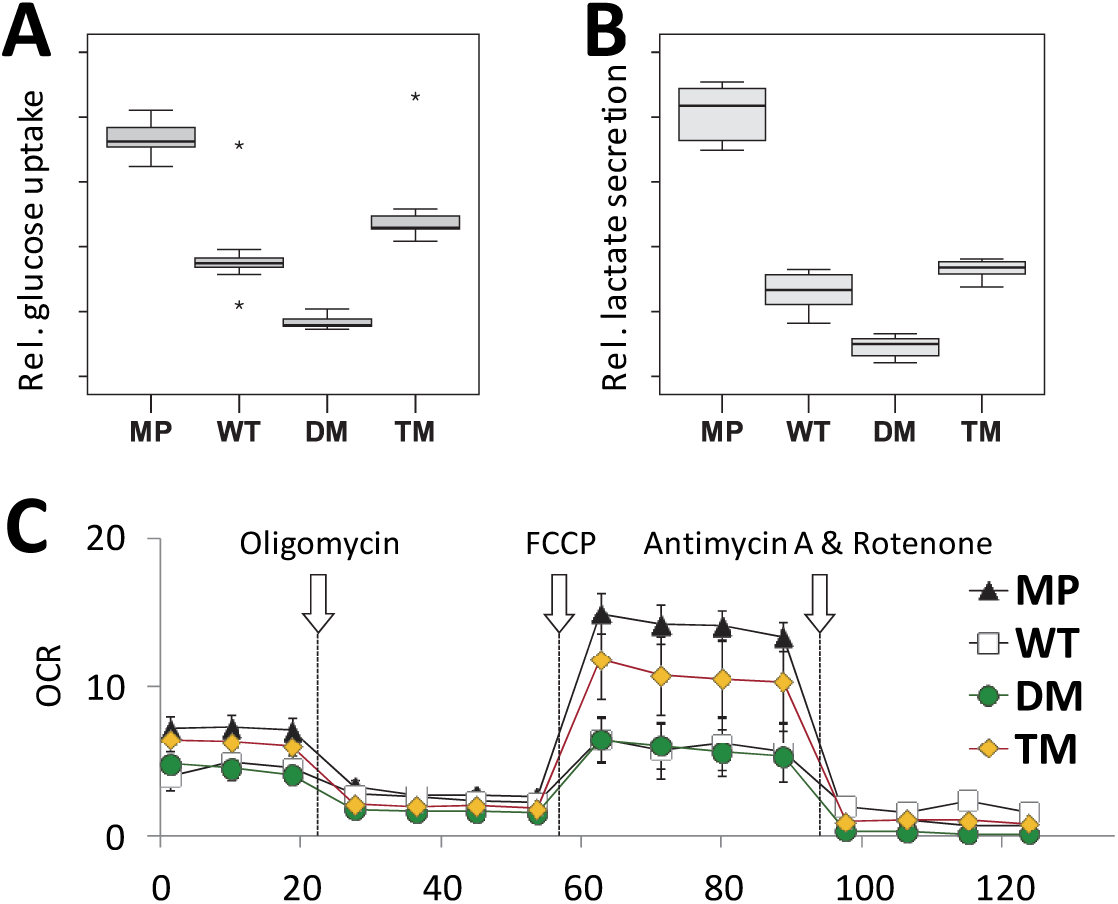
PGRMC1 phosphorylation status affects glucose uptake, lactate secretion, and mitochondrial function. (A) Glucose uptake by cell lines using the Cayman “Glycolysis” kit. The boxplots represent four technical replicates each for each of independent stably transfected sub-lines 1-3 of each condition WT, DM and TM (lines from Figure 1). i.e. n=4x3 = 12 per condition. For MP cells, 12 replicates of the MP cell line were performed. There was significant difference between all means (Kruskal-Wallis *p*<0.0001, pairwise two-sample Kolmogorov-Smirnov Tests *p*<0.0002). (B) Lactate secretion by cell lines. Details follow (A), except duplicates of each stable cell line were measured (n=3x2 = 6 per condition). Inter cell-type comparison tests revealed that the means of all pairwise comparisons were significantly different from one another (ANOVA, post-hoc Dunnet’s T3, *p*<0.003), except the WT-TM comparison which was not significant (*p*=0.211). (C) The maximal respiratory capacity of mitochondria is affected by PGRMC1 phosphorylation status. Mitochondrial oxygen consumption of respective independent clonal stable lines C1 of each PGRMC1-HA mutant condition WT, DM and TM. Arrows indicate the time of addition of ATP synthase inhibitor oligomycin, Δψm uncoupler FCCP, and electron transport chain inhibitors rotenone & antimycin A. OCR: oxygen consumption rate (pmol/min normalized per μg protein), n=5; +/-s.d.

### PGRMC1 phosphorylation affects mitochondrial function

Figure 3 also implied that mitochondria may be affected by PGRMC1 phosphorylation status. Naphthalimide-flavin redox sensor 2 (NpFR2) is a fluorophore targeted to the mitochondrial matrix. Its fluorescence is elevated approximately 100-fold when oxidized, providing an assay for mitochondrial matrix redox state (Kaur et al., 2015). NpFR2 revealed that the matrix of WT and TM cells was more oxidizing than MP and DM cells (Figure S5A, B), which corresponded with the elevated expression of many nuclear-encoded mitochondrial proteins in Figure 3.

We then examined mitochondria using the fluorescent marker MitoTracker, whose affinity for mitochondria is affected by mitochondrial membrane potential (Δψm) (Perry et al., 2011). Flow cytometry revealed the presence of two populations of MitoTracker-binding cells in each cell type: low and high MitoTracker-binding (Figure S5C-E). These populations were about equal for MP, WT, and TM cells, however, DM cells exhibited overall lower relative fluorescence level in each population (Figure S5C-D) and a higher proportion of cells with higher MitoTracker binding (Figure S5C,E). Notably, higher levels of mitochondrial proteins in WT and to some extent TM cells apparently did not correspond with higher Δψm caused by actively respiring mitochondria.

Relative to MP cells, the maximal respiratory rate was reduced (between 2 and 3 fold in Figure 5C) by expression of DM or WT PGRMC1-HA, but not by TM cells (Figure 5C). The relative profiles of basal (Figure S5F) and maximal (Figure S5G) respiratory rates for WT and DM were similar, with TM exhibiting rates intermediate to those of MP. This profile was observed on three independent comparisons WT/DM/TM. The single experiment including MP is shown. We conclude that the altered abundance of mitochondrial proteins due to PGRMC1 phosphorylation status detected in Figure 3 was accompanied by altered mitochondrial function. However, the relationship is not as simple as lower glucose uptake being associated with higher mitochondrial oxygen consumption, and may involve alterations in mitochondrial permeability to protons or other uncoupling mechanisms, for instance by altered cholesterol content (Cahill and Medlock, 2017).

### PGRMC1 phosphorylation affects mitochondrial morphology and function

Mitochondria were explored by measuring mitochondrial content (area per cell), size (perimeter), and morphology, or form factor (FF). FF is a parameter derived from individual mitochondrial area and perimeter, where higher values correspond to a greater degree of filamentous than fragmented mitochondria (Gosnell et al., 2016a; Koopman et al., 2006). Representative images of mitochondria are shown in Figure 6A. Numbers of mitochondria per cell varied greatly, with no significant differences detected between cell types (not shown). Over the entire data set, elongated cells exhibited greater mitochondrial area, larger mitochondria, and greater average FF (avFF) (Kolmogorov-Smirnov p<0.0001; not shown). When analyzed according to PGRMC1 status (cell type), MP and WT cells were predominantly elongated, and DM and TM were predominantly rounded, as expected (Figure 6A-B). We detected no significant differences in average mitochondrial area, perimeter or avFF between cell types for elongated cells, however rounded cell types exhibited significant differences between the cell types for area, perimeter, and avFF (Figure 6B). All cells with avFF < 2.2 exhibited rounded cell shape, while all cells with avFF > 2.6 exhibited elongated shape (Figure 6C). The observed avFF-associated transition from round to elongated cell shape was discrete for all cells except WT, occurring at avFF= 2.4 (MP), 2.2-2.6 (WT), 2.7 (DM) and 2.6 (TM). Notably, the single elongated TM cell also exhibited the highest avFF value for TM (Figure 6C). Holo-tomographic time-lapse videos (Ali et al., 2016) for each cell type show live mitochondria (File S9). PGRMC1 phosphorylation status probably influences mitochondrial content, size, and FF (degree of filamentation) by the same mechanisms that affect cell shape, consistent with the proposed influence of cytoskeleton on mitochondrial morphology and function (Anesti and Scorrano, 2006).

**Figure 6.**
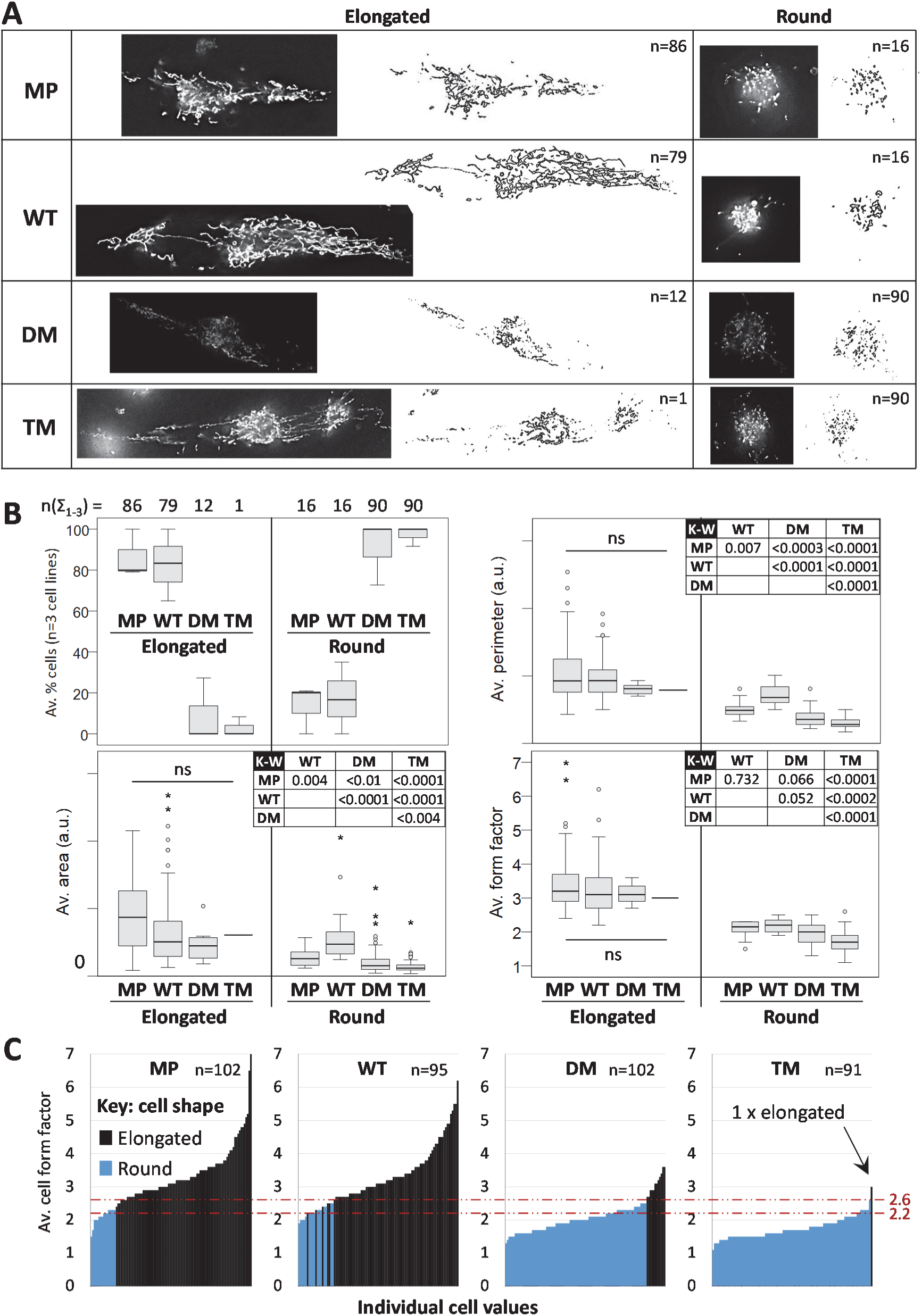
PGRMC1 phosphorylation influences mitochondrial and cell morphology together. (A) Images show cells with close to average form factor (avFF) values for elongate or rounded cells of each cell type. The numbers of cells scored in each class are indicated. All cell images are reproduced to the same scale. (B) Elongated and Round cells from (B) examined according to cell type. The top left panel shows average percentage elongated cells scored for three independent biological replicate cell lines per cell type across all 3 replicates (n(Σ_1-3_)) corresponding to the given n values in (A). The remaining panels showing the distribution of cells in each shape and type category of Area, Perimeter and FF. Kruskal-Wallis analysis revealed no significant differences at the *p*<0.05 level between any elongated cell type comparisons (ns), whereas all cell types exhibited significant differences for round cells (Kruskal-Wallis, *P*<0.001). The accompanying Kruskal-Wallis *post-hoc* pairwise comparison *p-* values for round cells are given in the respective tables. (C) AvFF per cell plotted including cell shape. The values where cells transition between round and elongated morphology (avFF 2.2-2.6) are indicated by dotted lines.

### PGRMC1 Y180 is required for subcutaneous mouse xenograft tumor growth

No significant differences in cell proliferation between cell types were observed in culture IncuCyte imaging (Figure 7A), or repeated MTT assays (not shown). We established subcutaneous xenograft tumors in replicate mice carrying each of sub-lines 1-3 of WT, DM and TM (e.g. 4x line 1, 3x line 2, 3x line 3, n=13 per PGRMC1-HA condition), as well as n=5 mice with cells expressing the single Y180F mutant (Neubauer et al., 2008), or MP cells. Tumors produced by both TM and Y180F cells were significantly smaller than those produced by WT or DM cells (Figure 7B-C), indicating that PGRMC1 Y180 was required for optimal tumor growth, and demonstrating that the cellular responses to altered PGRMC1 phosphorylation strongly influence cancer biology. All WT, DM, TM and Y180F tumor tissue expressed the PGRMC1-HA proteins (not shown), and therefore arose from the injected cells. There were no obvious differences in histology between cell types based upon hematoxylin and eosin staining (not shown).

**Figure 7.**
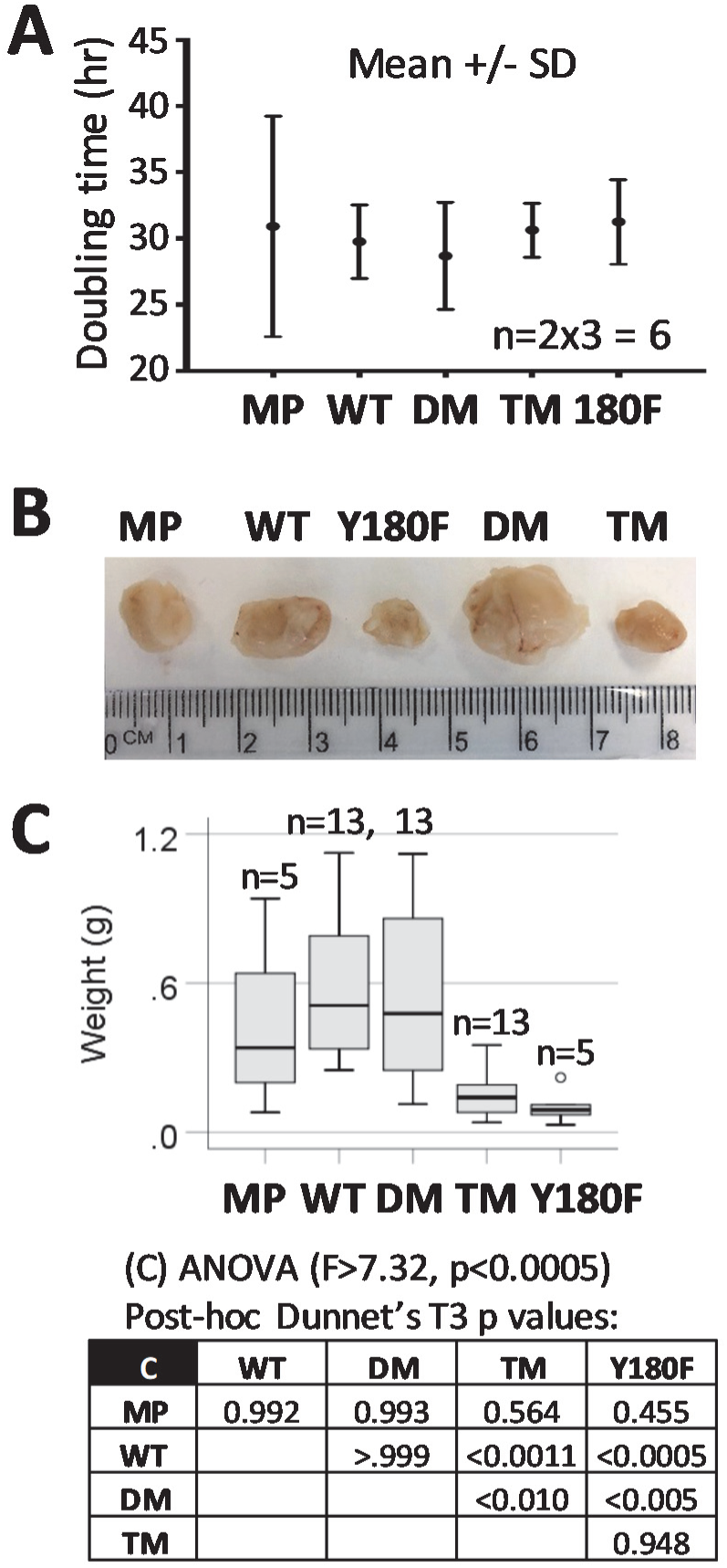
PGRMC1 Y180 contributed to growth of subcutaneous mouse xenograft tumors. (A) There are no differences in cell doubling time in culture. Replicates of stable cell lines 1-3 for each condition were measure to give n=6. There were no significant differences between cell conditions. (B) Typical tumors produced by each respective class of cell. (C) Box plot of the distribution of tumor sizes among mice injected with 2×10^6^ cells of each of the cell lines. For WT, DM and TM the results depict 4x each of lines 2 and 3, and 5 mice from line 1. MP and Y180F each represent 5 replicates of a single cell line. The box shows pairwise post-hoc Dunnet’s T3 *p*-values after one way ANOVA.

## DISCUSSION

We report new biology associated with the phosphorylation status of PGRMC1-HA proteins from Figure 1A profoundly affected cell morphology and migratory behavior. The morphotypic change from WT to DM resembled MAT in MP cells, being sensitive to ROCKI (Figure 1D). The DM and TM altered morphology is dependent upon activated ROCK, which leads to stiffening of cortical actomyosin (Sahai and Marshall, 2003). In human glioma cells over-expression of CD99 is implicated in MAT, resulting in rounded morphology, increased Rho activity, and enhanced migration (Seol et al., 2012). These properties superficially resemble the phenotype of our DM cells (Figure 1E-F), however DM cell migration involved pseudopodia and cell adhesion as evidenced in cell migration videos (File S1). Very little else is known at the molecular level about the events that promote MAT and altered cell motility (Friedl, 2004; Friedl and Wolf, 2010; Parri et al., 2009; Taddei et al., 2014). We provide here a global expression study of a possibly MAT-related process, and show that PGRMC1 phosphorylation status dramatically affects mitochondrial morphology and function.

PGRMC1 also influenced mitochondrial function and morphology (Figure 6). The elongated cell morphology that predominated in MP and WT cells was associated with a higher index of filamentous rather than fragmented mitochondria. Cells with rounded morphology and more fragmented mitochondria predominated in DM and TM cells (Figure 6). Such changes in mitochondrial function are driven by altered relative rates of mitochondrial fission and fusion, leading to mitochondrial fragmentation or elongated hypertubulation, respectively (Wai and Langer, 2016). Fragmented mitochondria are associated with pathological conditions including cardiovascular and neuromuscular disorders, cancer, obesity, and the process of aging, associated largely with altered cell differentiation (Wai and Langer, 2016). One of the proteins more abundant in WT and TM cells was Opa1 (O60313) (File S6), a protein known specifically to regulate mitochondrial fission/fusion, and one that has been reported to interact directly with PGRMC1 (Piel et al., 2016).

The strongest driver of mitochondrial morphology appears to have been cell shape, or *vice versa* (Figure 6). The cytoskeleton is thought to influence mitochondrial function and morphology (Anesti and Scorrano, 2006), and proteomics pathways mapping suggested cytoskeletal changes. WT cells had elevated levels of Tubulin 1alpha (Q71U36), and decreased levels of Tubulin alpha-1C (Q9BQE3), Tubulin beta-2A chain (Q13885), Tubulin beta-4B chain (P68371) (Figure S1E) relative to DM cells. Rounded DM cells exhibited more abundant type II cytoskeletal keratin 8 (P05787), type I cytoskeletal Keratin 18 (P05783), and type I cytoskeletal Keratin 19 (P08727) relative to MP cells (which are epithelial markers (Kim et al., 2015)), with WT and TM cells exhibiting intermediate keratin levels (see File S6 associated with Figure 3). These keratin abundance profiles paralleled the elevated levels of proteins in an enriched actin cytoskeleton pathway in DM cells (Figure S1B).

DM cells also displayed elevated levels of proteins in the T-complex protein-1 ring complex (TRiC, also known as CCT) (Figure S1E), which contributes to folding of proteins including actin and microtubules, and influences deregulated growth control, apoptosis, and genomic instability (Boudiaf-Benmammar et al., 2013; Roh et al., 2015). TRiC additionally contributes obligatory growth/survival functions in breast (Guest et al., 2015) and liver (Zhang et al., 2016) cancers. This complex is highly likely to contribute to the altered cytoskeletal properties and rounded phenotype of DM cells. TCP1 (P17987, Figure S1E) expression is driven by oncogenic PI3K signaling in breast cancer (Guest et al., 2015), and we observe both elevated PGRMC1-dependent PI3K/Akt activity and TRiC abundance in DM cells (Figure S1E).

In summary, many of the mitochondrial differences observed could be attributable to altered cytoskeletal properties. However, the differential mitochondrial functions of Figure 5 and Figure S5 did not correspond well with cell shape, indicating that PGRMC1 also changes complex causative processes driven by more than mitochondrial morphology. Results depicted in Figure 3 (as presented in File S5 and File S6) revealed that both the ATP synthase subunit beta of the F1 catalytic domain as well as the F0 proton pore domain were up-regulated in WT cells relative to DM cells (Figure S1C). It is possible that the higher Δψm of DM cells is related to low levels of F0/F1 ATPase proton channel (Figure S1C), resulting in relatively inefficient proton gradient clearance. Mitochondrial cholesterol decreases the permeability of the inner membrane to protons, increasing the efficiency of electron transport chain yield (Cahill and Medlock, 2017). Further work will be required to explain the mechanisms underlying the observed responses, which we are currently exploring.

Loss of PGRMC1 affects the SREBP-1/fatty acid homeostasis system (Lee et al., 2018), and PGRMC1 influences cell surface localization of insulin receptor and glucose transporters (Hampton et al., 2018). Chemical proteomics showed that PGRMC2 but not PGRMC1 promotes adipogenesis in 3T3-L1 preadipocytes following a gain of function interaction with a novel small molecule which displaced heme (Parker et al., 2017). It will be interesting to examine whether that treatment mimics the effects of phosphorylation. (PGRMC2 possesses cognates to PGRMC1 Y180 and S181, as well as heme-chelating Y113 (Cahill, 2017).)

Sabbir recently demonstrated the presence of SUMOylated PGRMC1 primarily in nuclear cell fractions (Sabbir, 2019), and Terzaghi et al. (2018) confirmed a nucleolar localization for PGRMC1, where it was responsible for nuclear localization of nucleolin which they proposed was associated with stress response. The zebrafish knockout of PGRMC1 results in elevated levels of mPRα mRNA, but decreased levels of the corresponding protein (Wu et al., 2018), suggesting that PGRMC1 can indeed affect the translational efficiency of certain mRNAs by ribosomes, which is consistent with our pathways analysis results, and especially the principal components analysis of Figure S1A which predicts that ribosomes and translation contributed most to the differences between cells.

Our results indicate that PI3K/Akt signaling in DM cells required PGRMC1 Y180, which was the sole difference to TM cells. PGRMC1 has long been recognized as a modulator of Akt activity, with cell type-specific effects (Hampton et al., 2018; Hand and Craven, 2003; He et al., 2018; Liu et al., 2009; Neubauer et al., 2008; Zhu et al., 2013; Zhu et al., 2017). This predicted activation of signals by removal of the putative inhibitory CK2 consensus sites in the DM protein (Cahill et al., 2016b; Neubauer et al., 2008) was dependent upon Y180 because TM (which differs from DM by a single oxygen atom) exhibited a protein expression profile that was more similar to WT than DM (Figure 3). Furthermore, Y180 is very important for the growth of subcutaneous tumors (Figure 7). In a methylomics study of these cells in an accompanying paper, the most significantly down-regulated KEGG pathway in the TM/DM comparison was PI3K-Akt (Thejer et al., 2019), consistent with PI3K/Akt activation requiring PGRMC1 Y180, which was required for tumor growth (Figure 7).

Interestingly, PGRMC1 knockdown in human pluripotent stem cells (hPSCs) led to an increase in GSK3β inhibitory phosphorylation (Kim et al., 2018). Examination of PGRMC1 phosphorylation status in that system is merited, where PGRMC1 suppressed the p53 and Wnt pathways to maintain hPSC pluripotency. Similarly to our results, those authors concluded that “that PGRMC1 is able to suppress broad networks necessary for multi-lineage fate specification.” Our hypothesis suggests that PGRMC1 Y180 phosphorylation and PI3K/Akt activity could be associated with elevated GSK-3β Ser9 phosphorylation and β-catenin signaling in some cancers (Pal et al., 2019).

The stem cell-like zygote (most similar animal cells to the unicellular animal ancestor) expresses cross-phylum conserved genes involved in processes such as cell cycle, mitosis, and chromatin structure (Yanai, 2018). All of these processes can be influenced by PGRMC1 (Cahill et al., 2016a). During animal development later embryological stages involve the induction of conserved germ layer-specific genes such as those for muscle (Yanai, 2018). This may be related to DM actin biology seen in our system, and suggests the hypothesis that CK2-like-site mediated negative regulation of PGRMC1 could be involved in these embryological processes, which merits further investigation.

Our initial hypothesis related to differential phosphorylation of PGRMC1, being potentially spatially and temporally associated with the onset of the Warburg effect (Neubauer et al., 2008). It is notable that the Warburg effect resembles a reversion to stem-cell-like metabolism (Riester et al., 2018). While this manuscript was in preparation, Sabbir showed that PGRMC1 post-translational modification status in HEK293 cells responds to P4 treatment, which was accompanied by a PGRMC1-dependent increase in glycolysis (Sabbir, 2019). Because the phospho-acceptor amino acid Y180 has been conserved in PGRMC1 proteins since the evolutionary appearance of the differentiation-inducing Spemann-organizer (Hehenberger et al., 2019), we believe it likely that PGRMC1 Y180-regulated modulation of metabolic and growth control that we have manipulated could represent a major newly identified foundational axis of animal cell biology, whose perturbation is inconsistent with the maintenance of differentiated states acquired during the subsequent evolution of complex body plans.

Although we can confidently deduce the existence of a PGRMC1 signal network, as yet we have identified neither immediate upstream PGRMC1 effectors nor downstream targets. In an accompanying paper (Thejer et al., 2019), we show that the cells characterized in this paper differ dramatically in genomic methylation and mutation rates. Future studies should urgently explore the relationship between PGRMC1 signaling and diseases such as cancer, diabetes, Alzheimer’s disease, and others (Cahill et al., 2016a).

## ACKNOWLEDGEMENTS

MP cells were obtained from Dr. Patsy Soon, Kolling Institute of Medical Research, Sydney. Jean Yang facilitated bioinformatics analysis. Research was primarily supported by Charles Sturt University internal funds to M.A.C and J.A.J, and by collaborating labs. Open access publication fees were jointly supported by CSU’s Faculty of Science, School of Biomedical Sciences, and Research Office. B.M.T. was supported by a PhD scholarship from the Ministry of Higher Education and Research, through the University of Wasit, Iraq. A.K. acknowledges the University of Sydney for a World Scholars Scholarship. E.M.G. acknowledges partial support of Australian Research Council award CE140100003. E.J.N. acknowledges the support of the Australian Research Council (DE120102687) and the Ramaciotti Foundation (ES2012/0051). T.L.R is supported by a Cancer Institute New South Wales Future Research Leader Fellowship. M.P. is supported by a NHMRC RD Wright Biomedical Career Development Fellowship and the Cancer Institute NSW Career Development Fellowship.

## AUTHOR CONTRIBUTIONS

Conceptualization, M.A.C.; Methodology, M.A.C, P.P.A, A.V.O., M.P., E.J.N, S.J.K. and M.P.M; Software, D.P. (proteomics); Formal Analysis, P.P.A., D.P., M.J., E.P. M.A.C., and M.J.; Investigation, B.M.T., P.P.A., S.L.T., A.V.O., A.K., I.S., N.C., X.S., C.P.C., D.P., L.T., and J.C.C.; Resources, B.M.T., M.P.M., M.P., E.J.N., H.N., T.F., and J.C.Q.; Writing – Original Draft, M.A.C.; Writing – Review & Editing, M.P.M., M.A.C., P.P.A., E.M.G., C.P.C., B.M.T., M.P., E.M.G, and X.S.; Visualization, M.A.C.; Supervision, M.A.C, J.A.J., E.M.G., E.J.N., T.L.R, and M.P.M.; Project Administration, M.A.C.; Funding Acquisition, M.A.C., J.A.J., J.C.Q., E.M.G., M.P.M., T.L.R., and E.J.N.

## DECLARATION OF INTERESTS

M.A.C. is a shareholder of Cognition Therapeutics Inc. (Pittsburgh) and a member of its scientific advisory board. The authors declare no other competing interests.

## STAR METHODS

### Plasmid preparation

Plasmids pcDNA3.1-PGRMC1-HA (wild type: WT), pcDNA3.1-PGRMC1-HA_Y180F (Y180F), pcDNA3.1-PGRMC1-HA_S57A/S181A (double mutant, DM) and pcDNA3.1-PGRMC1-HA_Y180F (Y180F) (Neubauer et al., 2008) have been described. The triple mutant S57A/Y180F/S181 (TM) was constructed by Genscript (Honk Kong) using the DM plasmid as the template and introducing codon TTC for F180. PGRMC1-HA open reading frames were reconfirmed by DNA sequencing at Monash Micromon DNA Sequencing Facility (Clayton, Vic., Australia) using the 5′ T7 and 3′ BGH sequencing primers specific for the parental vector. PGRMC1-HA plasmids were transformed into *Escherichia coli* Top10 strain, and cultured overnight at 37°C on 1% agar plates containing Luria broth (LB) media (Invitrogen) and 50 μg/mL ampicillin. A single colony was picked and bacteria were grown in 250 ml culture by aeration overnight at 37°C in LB media. Plasmid DNA was isolated by GeneJet Maxiprep Kit (ThermoScientific) following the manufacturer’s protocol. Plasmid DNA concentration was measured by using Nanodrop (Thermo Scientific).

### Cell culture

MIA PaCa-2 (MP) cell identity was verified as MIA PaCa-2 (ATCC CRL-1420) by the MHTP Medical Genomics Facility (Monash University, Melbourne) following the ATCC Standards Development Organization document ASN-0002 for cell line identification via short tandem repeat profiling. MP cells were maintained in Dulbecco’s Modified Eagle’s medium (DMEM-high glucose, Sigma-Aldrich, D5796) supplemented with 10% Foetal bovine calf serum (Sigma-Aldrich, F9423) and 1% penicillin-streptomycin (Sigma-Aldrich, P4333) (complete DMEM) at 37°C and 5% CO2 in a 150i CO_2_ incubator (Heracell, Lane Cove NSW). Cell doubling times were estimated by 3-(4,5-dimethylthiazolyl-2)-2,5-diphenyltetrazolium bromide (MTT) assay or by IncuCyte, as described (Quin et al., 2016). Mitochondrial respiratory capacity was measured using a Seahorse Extracellular Flux analyzer XF24 (Seahorse Biosciences).

### Transfection and stable cell line generation

Under our culture conditions, MP cells exist in culture as flattened adherent cells of mesenchymal shape, a minority of rounded adherent cells, and a small population of rounded suspension cells. Plating the suspension cells regenerates a similar population distribution (not shown). On the day before transfection, 2×10^6^ MP cells were seeded onto a 6-well plate. The cells were transfected at 70-80% confluency. Before transfection, cells were washed with Dulbecco’s phosphate-buffered saline (PBS, Sigma-Aldrich, D8537) and maintained in antibiotic-free Dulbecco’s Modified eagle Medium high glucose (DMEM, Sigma-Aldrich, D5796) containing 10% bovine calf serum (Sigma-Aldrich, 12133C) and 1% penicillin/streptomycin (Sigma-Aldrich, P4458) (complete DMEM). In separate transfections, 4μg each of respective PGRMC1-HA plasmids (WT, DM or TM) and Lipofectamine 2000 (Life Technologies, 11668-019) were mixed at 1:2 ratio and incubated for 25 min at room temperature. The mixture was then added drop-wise to the wells of the culture plate. After 6 hours of incubation, cells were washed with PBS and cultured at 37°C and 5% CO2 in complete DMEM for 48 hours, after which cells were harvested and plated in three fold limiting dilution in complete DMEM containing 50 μg/ml Hygromycin B (EMD Millipore, 400052) in 96 well plates. Cells were cultured at 37°C and 5% CO2 for 2 weeks, with regular media changes containing complete DMEM with Hygromycin B every 3 days to select for stable integration events. Typically 8 independent stably transfected cell lines were expanded for each of PGRMC1-HA WT, DM and TM and 3 lines with similar levels of PGRMC1-HA expression were selected by Western blot.

Cells were frozen 0.5 – 1.0 mL at -−0°C in Bambanker (Novachem, #306-14684) at 2×10^6^ cells/mL. Frozen cells were introduced back into culture by thawing at 37°C for 20 seconds followed by addition to 5 ml of complete media and low speed centrifugation at 180 x g for 3 mins at 25°C. Pelleted cells were resuspended in 6mL fresh complete media and seeded in 25 cm^2^ flasks.

Because of the dramatic effects observed, MP cells are included in our experiments as a literature reference point. MP differ from WT cells by not having undergone hygromycin selection, and by lack of overexpression of PGRMC1-HA. Therefore we cannot ascribe differences between MP and WT cells to PGRMC1-HA expression. The effects of the DM and TM PGRMC1 mutations are assessed relative to WT control levels.

### shRNA lentiviral production

Lentiviral-delivered shRNAs were constructed using Mission TRC2-pLKO-Puro series lentiplasmids (SHCLND, Sigma-Aldrich) targeting ERR1/ESRRA (TRCN0000330191, GAGAGGAGTATGTTCTACTAA), vinculin (shVCL) (TRCN0000290168, CGGTTGGTACTGCTAATAAAT) or non-target scrambled shRNA (shScr) (SHC202; CAACAAGATGAAGAGCACCAA). MP and DM shVCL cells were obtained at first attempt, however, despite three attempts, WT cells could not be established. To generate virus particles, we co-transfected HEK293 cells with the shRNA plasmids and helper plasmids using Lipofectamine 2000 (Invitrogen, 11668-027). Prior to transfection, 6 well plates were treated with 50µg/mL D-Lysine overnight. Next day, 1×10^6^ HEK293 cells were seeded per well and incubated overnight at 37^°^C in complete medium. Transfection mixture A contained 4 μg plasmid mixture consisting of 2.5 μg target or shScr shRNA lentiplasmid, 0.75 μg Pax, 0.3μg Rev and 0.45μg VSV-G helper plasmids (Gurusinghe et al., 2015) in 250 μL antibiotic-free medium. Transfection mixture B contained 8 μL Lipofectamine and 242 μL antibiotic-free medium. After 5 min incubation at 25^°^C, mixtures A and B were gently mixed and incubated for 25 min at 25^°^C. HEK293 culture medium was removed, cells were washed with PBS, and 2 mL fresh antibiotic-free medium was added followed by addition of combined transfection mixture to the cells, dropwise with gentle shaking, followed by incubation for 6 h at 37^°^C. After incubation, the medium was replaced with complete medium overnight at 37^°^C. Virus particles were harvested by collecting culture medium followed by the addition of new medium every 24 h for 72 h. Collected media for each culture were pooled, filtered through a 22 μM filter, aliquoted into 1 mL fractions, and frozen at -−0^°^C.

### shRNA lentiviral transduction

Briefly, 1×10^5^ MP, WT, or DM cells per 24 plate well were seeded in 1 mL complete DMEM medium and grown to 60% confluency. The medium was removed, and replaced by 1 mL of medium per well, containing 2-fold serially diluted virus particles in adjacent wells, plus 5µg/mL Polybrene (hexadimethrine bromide, Sigma-Aldrich 107689) to enhance viral transduction. After incubation for 24 hours, the medium was removed and the cells were washed twice with PBS after which fresh medium was added supplemented with 1.5 µg/mL Puromycin, which was replaced every 48 h for 1 week. Cells from wells transduced with the lowest dilutions of respective virus particles that survived selection were expanded, and stocks frozen at −80°C in Bambanker.

### Scratch migration assay

MP cells or stable transfected monoclonal MP cell lines expressing PGRMC1-HA WT, DM or TM proteins (1×10^4^ cells) were seeded in a 24 well plate. The monolayer of cells at more than 90% confluency was subjected to serum starvation for 2 hours. A scratch was created in the middle of the monolayer by a sterile p200 tip and washed twice with PBS to remove floating cells. Complete media was then added. The cell monolayer was incubated for 36 hours to allow cell migration into the scratched area. Photographic images were taken at 0 and 36 hours using an inverted phase microscope (Nikon Eclipse Ti-U). Cells in the boxed areas of Figure 2A were manually scored from printed images. Cell treatments included 125 nM Y-27632 dihydrochloride (Abcam, ab120129, Rho Kinase inhibitor: ROCKI) or vehicle control (DMSO). For video files the cells were incubated at 37°C and 5% CO_2_ for 36 hr in a stage top electrically heated chamber (Okolab H301-NIKON-NZ100/200/500-N) including transparent heated lid (H301-EC-HG-LID), with a 24-well Nunc/Greiner plate base adapter (24MW-NUNC) and a chamber riser, for a working distance of 28 mm. The chamber was regulated by a Control Unit (Okolab H301-TC1-HMTC) with Digital CO2 controller (Okolab DGT-CO2 BX) and Air Pump (Okolab, OKO-AP) and was inserted to a Nano-Z100-N Piezo stage (Mad City Labs) on a motorized XY stage (Nikon TI-S-ER). Images were taken every 10 minutes for 36 hours with a 10× (0.45 NA) Plan Apo objective using the transmitted light detector (TD) on a Nikon Ti Eclipse Confocal microscope controlled by NIS Elements V4.10 software (Nikon).

### Proteomics sample preparation

Three independent stable transfected lines of each PGRMC1-HA-expressing cell type, as well as triplicates of the MP parental cell line, were measured in technical replicate data-dependent and independent data acquisition SWATH-MS modes on a 5600 TripleTof™ mass spectrometer (ABSciex). Global proteomics analysis was carried out at the Australian Proteome Analysis Facility (APAF). Cells were grown in Wagga Wagga to 80% confluency in 75 cm^2^ flasks. Three separate cultures of MP cells (passages 8, 9 & 11) and three lines of each PGRMC1-HA WT, DM and TM cells were used (independent biological triplicates). Cells were harvested and frozen cell pellets were shipped on dry ice to APAF for Mass spectrometric analysis. Cell pellets were lysed using 200 μL of sodium deoxycholate buffer (1% in 0.03M triethyl ammonium bicarbonate), and DNA digested using 0.5 μg of benzonase. Direct detect assay (EMD Millipore, DDAC00010-8P) was performed on the samples and 100 μg of each sample was taken for digestion. Samples were reduced with dithiothreitol (5 mM), alkylated with iodoacetamide (10 mM) and then digested with 4 μg trypsin for 16 hours at 37°C. The digested sample was acidified and centrifuged to remove the sodium deoxycholate. Samples were then dried and resuspended in 100 μL of loading buffer (2% acetonitrile 0.1% formic acid). Individual samples for SWATH analyses were diluted 1:4 into loading buffer and transferred to a vial. Each sample was measured in technical replicate. For IDA runs a pool was made for each group (MP, WT, DM, and TM) by taking equal portions from each biological replicate and diluting 1:4 in loading buffer.

### Proteomic Information dependent acquisition

Tryptic peptides were analyzed on a 5600 TripleTof™ mass spectrometer (ABSciex). Chromatographic separation of peptides was performed on a NanoLC-2Dplus HPLC system (Eksigent, Dublin, CA) coupled to a self-packed analytical column (Halo C18, 160Å, 2.7 μm, 75 μm x 10cm). Peptide samples (4 μg of total peptide amount) were loaded onto a peptide trap (Opti-trap Cap 0.5mm x 1.3mm, Optimize Technologies) for pre-concentration and desalted with 0.1% formic acid, 2% ACN, at 10 µL/min for 5 minutes. The peptide trap was then switched into line with the analytical column and peptides were eluted from the column using linear solvent gradients with steps, from 98% Buffer A (0.1% formic acid) and 2% Buffer B (99.9% acetonitrile, 0.1% formic acid) to 90% Buffer A and 10 % Buffer B for 10 minutes, then to 65% Buffer A and 35% Buffer B at 500 nL/min over a 78 min period. After peptide elution, the column was cleaned with 95% Buffer B for 15 minutes and then equilibrated with 98% Buffer A for 15 minutes before the next sample injection. The reverse phase nanoLC eluent was subject to positive ion nanoflow electrospray analysis in an IDA mode. Sample analysis order for LC/MS was DM1, DM2, DM2, DM3, DM3, MP2, MP2, MP3, MP3, TM2, TM2, TM3, TM3, WT1, WT1, WT2, WT2, WT3, WT3, DM1, MP1, MP1, TM1, TM1.

In the IDA mode a TOFMS survey scan was acquired (m/z 350 - 1500, 0.25 second), with the 10 most intense multiply charged ions (2+ - 5+; counts >150) in the survey scan sequentially subjected to MS/MS analysis. The selected precursors were then added to a dynamic exclusion list for 20s. MS/MS spectra were accumulated for 50 milliseconds in the mass range m/z 100 – 1500 with rolling collision energy.

### Proteomic Data independent acquisition (SWATH)

Samples were analyzed by Sequential Window Acquisition of all Theoretical Mass Spectrometry (SWATH-MS) (Gillet et al., 2012) proteomics profiling in duplicate, with the chromatographic conditions as for IDA analysis above. The reverse phase nanoLC eluent was subject to positive ion nanoflow electrospray analysis in a data independent acquisition mode (SWATH). For SWATH MS, m/z window sizes were determined based on precursor m/z frequencies (m/z 400 – 1250) in previous IDA data (SWATH variable window acquisition, 60 windows in total). In SWATH mode, first a TOFMS survey scan was acquired (m/z 350-1500, 0.05 sec) then the 60 predefined m/z ranges were sequentially subjected to MS/MS analysis. MS/MS spectra were accumulated for 96 milliseconds in the mass range m/z 350-1500 with rolling collision energy optimized for lower m/z in m/z window +10%. To minimize instrument condition caused bias, SWATH data were acquired in random order for the samples with one blank run between every sample.

### SWATH library generation

The LC-MS/MS data of the IDA data were searched using ProteinPilot (version 4.2) (Sciex) and combined into a single search report file. The files were searched against Human entries in the Swissprot 2014_04 database (released 16/04/2015, containing 545,388 entries). The search parameters were selected as follows: iodoacetamide cysteine alkylation, trypsin digestion, Triple TOF 5600 instrumentation, biological modifications, thorough search and false discovery rate enabled.

### SWATH data processing

SWATH data were extracted using PeakView (version 2.1, Sciex) with the following parameters: Top 6 most intense fragments of each peptide were extracted from the SWATH data sets (75 ppm mass tolerance, 10 min retention time window). Shared peptides were excluded. After data processing, peptides (max 50 peptides per protein) with confidence ≥ 99% and FDR ≤1% (based on chromatographic feature after fragment extraction) were used for quantitation. The extracted SWATH protein peak areas were normalized to the total peak area for each run and subjected to t-test to compare relative protein peak area between the samples. Protein t-test with *p*-value smaller than 0.05 and fold change larger than 1.5 were highlighted as differentially expressed. The analysis of four different cell types was treated as six separate paired comparisons: 1) MP vs. WT, 2) MP vs. DM, 3) MP Vs. TM, 4) WT vs. DM, 5) WT vs. TM, and 6) DM vs. TM.

Additionally, a similar method for determining differential expression was run at the peptide level, with the peptide level fold changes then averaged for each protein. The peptide level analysis is more conservative, as peptide level *p*-values are only generated when a protein was identified by at least two proteins. Peptide level data were used for this study. The mass spectrometry proteomics data have been deposited to the ProteomeXchange Consortium via the PRIDE (Perez-Riverol et al., 2019) partner repository with the dataset identifiers PXD014716 (Figure 3) and PXD014789 (Figure S3). Identical methods were employed to quantify effects of the shRNA-mediated attenuation of ERR1 versus scramble shRNA expression.

### WebGestalt enrichment analyses

Gene Ontology (GO) and pathway enrichment analysis were conducted on differentially abundant proteins from Figure S2D using the WEB-based GEne SeT AnaLysis Toolkit (WebGestalt) platform (http://bioinfo.vanderbilt.edu/webgestalt/) (Wang et al., 2013). Figures or Supplementary Figures depicting or relying on WebGestalt results describe supplementary data files available online that contain the respective original WebGestalt analysis files. Attempting to identify the most important driving contributions to the phenotypic alterations observed, a complementary WebGestalt analysis was also performed at the adjP<0.001% level, but employing the subsets of proteins which were detected to be either significantly up- or down-regulated (“red” and “blue” lists of proteins for each comparison from File S1). The UniProt protein IDs from respective proteomics comparisons were uploaded as text files which accompany the respective WebGestalt supplementary data files. KEGG, Pathway Commons, and Wiki Pathways were analyzed at the 5% level. GO and transcription Factor analyses were also performed for comparisons of differentially up- or down-regulated proteins for each pair-wise comparison between cell types at the 0.1% and 10% levels. The following WebGestalt settings were employed: Organism: hsapiens; gene Id Type: entrezgene; Reference Set for Enrichment Analysis; entrezgene_protein-coding; Significance Level: (variable see individual analysis descriptions), Statistical Method: Hypergeometric, Multiple Test Adjustment: Benjamini-Hochberg (BH), Minimum Number of Genes for a Category: 3.

Pathway enrichment analyses can consider either a) all differential proteins together (including both up- and down-regulated proteins in the one analysis), or b) can examine the higher abundance and lower abundance proteins in separate analyses for each comparison (“red” or “blue” analyses for each comparison). Our WebGestalt pathway analysis strategy of Figure 3 pursued the second of these alternatives. In the first analysis, all proteins found to be differential (both up- and down-regulated) between two samples in any six of the analyses were entered as a protein list to WebGestalt, and pathway enrichment analysis was performed with multiple sample correction at the Benjamini and Hochberg (BH) adjusted p (adjP) significance level of 0.001 for KEGG, Pathway Commons, and Wikipathways for each of the six basic cell-type comparisons.

Twelve separate WebGestalt analyses were performed for Figure 3, with enrichment analyses including KEGG, PC, WikiPathways, as well as Transcription Factor and GO cellular component. The schematic representation of Figure 3 shows pathways identified as significantly enriched for at least one of 12 comparisons (6x more abundant “red” and 6x less abundant “blue”) between 1) MP vs. WT, 2) MP vs. DM, 3) MP vs. TM, 4) WT vs. DM, 5) WT vs. TM, and 6) DM vs. TM. The original WebGestalt analyses at adjP<0.001 are available as File S7. The pathway mapping for all significantly detected features between all comparisons detected at adjP<0.001 is available as File S5A. Another WebGestalt analysis was performed at adjP<0.1 (available as File S8) and the significances of each pathway identified in File S5A at the adjP<0.001 level were then recorded for all comparisons at the adjP<0.1 level in File S5B. The analysis results are presented for reference as File S4B as drawn from File S7. That information was used to assign statistical significance in the pathways map for all features that were identified at the adjP<0.001 level in any one comparison, across all 12 comparisons at the adjP<0.1 level. These data were then used to map all proteins from all pathways and all comparisons of File S5B to produce the original image of Figure 3 in Microsoft Excel, which is available as File S6 and contains all protein and pathway identities.

### Mapping pathways to the expression heat map

The matrix of protein membership to pathway or functional group category is a resulting sparse matrix with 0/1 indicating that the respective protein is/is not present in the respective category. This matrix was clustered using the hclust implementation in the R Base Package (www.r-project.org/), using a binary distance and complete linkage, to reorder the columns (pathways in this case) according to the proportion of shared proteins. The resulting cladogram including overlapping features identified by all WebGestalt analyses appears to the right of pathways in Figure 3, and with complete accompanying protein and pathway identities in File S5.

### Principal Components Analysis on Proteomics results

Principal component analysis was used to examine the largest contributions to variation in the protein measurements. Wilcoxon rank-sum tests were used to identify the pathways that were positively or negatively associated with the principal component scores.

### Sample Preparation for Western blots

Approximately 70% confluent cells in a T75 flask were washed twice with chilled PBS buffer and incubated with 500 µL radio immunoprecipitation assay buffer (RIPA buffer) (Sigma-Aldrich, R0278) supplemented with protease and phosphatase inhibitor cocktail (Thermoscientific, 88668) following manufacturer’s recommendations. After scraping, the lysate was centrifuged at 8000g for 20 minutes (Hermle Centrifuge Z233 M-2) at 4°C. Protein concentration was determined using the Pierce BCA protein assay kit (ThermoFisher, 23225) following the manufacturer’s instructions. 20 µg cell lysates were each mixed with 2x Laemmli loading buffer (Sigma-Aldrich, S3401) at a 1:1 ratio to give final volume 20 μL, followed by denaturation at 95°C for 5 minutes in a digital dry bath heater. Lysates were loaded immediately to a 10% SDS-PAGE gel. For HA western we used 17 well gels (Life technologies, NW04127), for PGRMC1 Western blots we used 15 well gels (Bio Rad. 456-1069), for vinculin and ERR1 Western blots we used 10 well gels (Bio Rad 456-1096). Electrophoresis was at 150V for 45 min. Protein was transferred onto PVDF membranes (Bio-Rad, 1620174) with a Trans-Blot Turbo transfer system (Bio-Rad, Gladesville NSW) for 7 min by Trans-Blot® Turbo RTA Mini LF PVDF Transfer Kit (Bio-Rad, 1704274) or wet transferred in 25 mM Tris, 192 mM glycine, 20% (v/v) methanol (pH 8.3) (1x Towbin buffer) at 20V for 2.5 hours on mini trans-blot cell (Bio Rad, 1703930) cooled on ice.

### Western blots

Membranes were blocked with TBS-T (0.1% Tween-20 in 1× Tris-buffered saline) containing 5% Woolworths Instant Skim Milk Powder (Woolworths, Wagga Wagga, NSW, Australia) for 1 hr and incubated overnight at 4°C with primary (1°) antibody. After washing 3 times with TBS-T, blots were incubated with secondary (2°) antibody for 1 hr at room temperature. Proteins were detected by the following methods. For chemiluminescence detection the membranes were incubated with Clarity Max Western ECL Substrate (Bio-Rad, #1705062) for 5 min for detection by enhanced chemiluminescence using a Bio-Rad ChemiDoc MP imaging system (Bio-Rad, Gladesville NSW). Fluorescence detection was performed on the ChemiDoc (at the indicated wavelength). Molecular weight standard proteins for gels imaged for fluorescence or chemiluminescence were detected on the ChemiDoc using the IRDy680 channel. Colorimetric detection (Vinculin and ERR1 Western blots) was performed by incubation of membranes with 3 mL Tetramethylbenzidine (TMB) (Sigma-Aldrich, T0565) for 5 minutes. These images were captured by Molecular Imager Gel Doc XR+ System (Bio-Rad, Gladesville NSW). Multi-channel ChemiDoc images were generated with the Bio-Rad Image Lab Software. Some dual channels images were manipulated in Adobe Photoshop CC 2018 (Adobe Systems Inc.) by reducing intensity in either red or green channel to lower background in the published image. Adjustments were applied identically over all image pixels so as to not alter the relative intensities of any bands.

The following primary (1°) and secondary (2°) antibody pairs were used (at the specified dilutions) with the indicated detection methods. For PGRMC1 Western: 1° goat anti-PGRMC1 antibody (Abcam, ab48012) (1:1000) and 2° rabbit anti-goat secondary antibody (Abcam, ab6741) (1:4000) detected by chemiluminescence. After detection, membranes were blocked with TBS-T overnight and then incubated with 1° mouse anti-beta actin (Sigma-Aldrich, A5541) (1:2000) and 2° goat anti-mouse IgG H&L (IRDye 800CW) (1:5000) detected by fluorescence (IRDye 800CW). For HA epitope Western: 1° mouse anti-HA (Sigma-Aldrich, H3663) (1:2000) and 2° goat anti-mouse IgG H&L (IRDye 800CW) (dilution 1:5000) detected by fluorescence (IRDye 800CW), as well as 1° rabbit anti-beta-actin (Cell Signaling, 4967) (1:2000) and 2° donkey anti-rabbit IgG (Abcam, ab16284) (1:2000) detected by chemiluminescence.

For vinculin Western: 1° anti-vinculin (E1E9V) XP (Cell Signaling Technology, 13901) (1:1000) and 2° donkey anti-rabbit IgG (Abcam, ab16284) (1:2000) detected colorimetrically. For ERR1 Western: 1° rabbit anti-ERRα (E1G1J) (Cell Signaling, 13826S) (1:1000) and 2° donkey anti-rabbit IgG (Abcam, ab16284) (1:2000) detected colorimetrically.

### Reverse Phase Protein Array analysis

Reverse Phase Protein Arrays (RPPA) using Zeptosens technology (Bayer AG, Leverkusen, Germany) were used for analysis of signaling protein expression and activity profiling as described (Bader et al., 2015; Kaistha et al., 2016; Pawlak et al., 2002; Pirnia et al., 2009). For the analysis, flash frozen cell pellets were lysed by incubation with 100 µl cell lysis buffer CLB1 (Bayer, Germany) for 30 minutes at room temperature. Total protein concentrations of the lysate supernatants were determined by Bradford Assay (Coomassie Plus, Thermo Scientific). Cell lysate samples were adjusted to uniform protein concentration in CLB1, diluted 10-fold in RPPA spotting buffer CSBL1 (Bayer) and subsequently printed as series of four dilutions (starting concentration at 0.3 µg/µl plus 1.6-fold dilutions) and in two replicates each. All samples were printed as replicate microarrays onto Zeptosens hydrophobic chips (Bayer) using a NanoPlotter 2 (GeSim, Grosserkmannsdorf, Germany) applying single droplet depositions (0.4 nL volume per spot). After printing, the microarrays were blocked with 3% w/v albumin, washed thoroughly with double distilled H2O, dried in a stream of nitrogen and stored in the dark at 4°C until further use.

Protein expression and activity levels were measured using a direct two-step sequential immunoassay and sensitive, quantitative fluorescence read-out. A single array was probed for each protein. Highly specific and upfront validated primary antibodies were incubated at the respective dilution in Zeptosens assay buffer overnight (15 hours) at room temperature. Arrays were washed once in assay buffer and incubated for 45 minutes with Alexa647-labeled anti-species secondary antibody (Invitrogen, Paisley, UK). Arrays were then washed as before and imaged using a ZeptoREADER instrument (Bayer) in the red laser channel. Typically, six fluorescence images were recorded for each array at exposure times of between 0.5 and 16 seconds. Negative control assays incubated in the absence of primary antibody (blank assays) were also performed to measure the non-specific signal contributions of the secondary antibody. In addition, one chip out of the print series was stained to measure the relative amount of immobilized protein per spot (protein stain assay). The following primary antibodies (provider and reagent number, dilution) were used: Bad (CST 9239, 1:200), Bad-P-Ser112 (CST 5284, 1:100), Bad-P-Ser136 (CST 4366, 1:100), FAK1-P-Tyr577 (Invitrogen 44-614ZG, 1:100), FAK1-P-Tyr861 (Epitomics 2153-1, 1:100), GSK3beta (CST 9315, 1:200), GSK3beta-P-Ser9 (CST 9336, 1:100), HSF1 (Epitomics 2043-1, 1:1000).

After assay measurements (one protein per array), array images and data were analyzed with the software ZeptoVIEW 3.1 (Bayer). For each array/antibody, the image taken at the longest exposure time without showing any saturation effect was analyzed with the spot diameters set to 160 µm. Mean fluorescence signal intensity (MFI) of each sample was calculated from referenced, background-corrected mean intensities of the single spots (eight spots per sample) applying a linear fit and interpolating to the mean of the four printed protein concentrations. Blank-corrected MFI signals of the samples were normalized for the relative protein concentration printed on the chip to obtain normalized fluorescence intensity signals (NFI). NFI values were used for all subsequent statistical analyses.

### Glucose uptake & Lactate production assay

Glucose uptake and lactate production assays were performed by using commercially available kits from Cayman chemical (#600470, #700510) following manufacturer’s protocols. Glucose uptake was measured with a Fluostar Omega fluorescence microplate reader (BMG Labtech, Ortenberg, Germany) and lactate production was quantified with a Molecular Devices Spectra Max 190 microplate reader (Bio-Strategy P/L, Campbellfield, Vic., Australia).

### NpFR2 redox assay

Intramitochondrial redox status was measured by naphthalimide flavin redox sensor 2 (NpFR2) (Kaur et al., 2015). Mia PaCa-2 and PGRMC1-HA-expressing stable cells (1×10^6^) were suspended in 2 mL complete media and seeded in six well plates and cultured for 24 hr at 37°C and 5% CO_2_. Cells were washed with PBS, trypsinsed, harvested, and resuspended in 1 mL of fresh media containing 25μM NpFR2 in a 1.5 mL microcentrifuge tube, followed by incubation for 20 min at 37°C. Cells were then centrifuged in a microcentrifuge at 180 x g, the pellet was resuspended once with 1 mL PBS followed by recentrifugation, and the washed pellet was again resuspended in 1 mL PBS. 500 μL cell suspension was loaded to a Gallios Flow Cytometer (Beckman Coulter) and fluorescence of 2×10^4^ cells was detected using FL1 (green) channel.

### Immunofluorescence microscopy

To detect the expression of exogenous HA tagged PGRMC1 in Figure 1E, cells were seeded on coverslips on a six well plate. The cells were washed with ice-cold PBS, mildly fixed with 3.7% formaldehyde for 5 minutes at 4°C. The cells were then permeabilized with ice-cold 100% methanol for 10 minutes at -20°C, followed by overnight incubation with anti-HA tag antibody (Sigma, H3663). The cells were washed extensively and incubated with FITC conjugated secondary antibody (Sigma, F8521) in dark for 1 hour at 4°C. Cells were washed three times with PBS and counterstained with DAPI mounting solution. Images were captured using a Nikon Ti Eclipse Confocal microscope (Nikon Australia Pty Ltd).

### Analysis of mitochondrial morphology

Mitochondria were quantified for cell shape (elongated/round), mitochondrial content (sum of mitochondrial area/cell), mitochondrial size (average perimeter/cell), and mitochondrial morphology or Formfactor (FF): a measure where higher values correspond to a greater level of filamentous mitochondria and lower values correspond to more highly fragmented mitochondria (Koopman et al., 2006). Formfactor (calculated as the P^2^/4πA) measures mitochondrial morphology based on the perimeter and area of shape. The calculation takes in to account not only changes in length, but also the degree of branching, making at an ideal form of measurement for the quantification of mitochondrial morphology.

To measure form factor, 1×10^5^ cells were seeded onto Nunc 176740 four well plates with a 22×22mm #1.5 glass coverslip on the bottom. Cells were fixed and permeabilized as above, then incubated with Abcam mouse anti-mitochondrial IgG1 antibody (Abcam ab3298) and then with FITC-conjugated goat anti-mouse secondary antibody (Sigma-Aldrich F4018) and DAPI, followed phalloidin red staining and imaged with 3D-Structured Illumination Microscopy (SIM) on a DeltaVisionOMX Blaze microscope as described (Strauss et al., 2012). Images were processed using Fiji/ImageJ software (Schindelin et al., 2012), and Area and Perimeter values were extracted to calculate form factor. Cell morphology was scored as either ‘round’ or ‘elongated’ by JCC as part of the mitochondrial quantification process.

### Holo-tomographic imaging

Holo-tomographic video imaging was performed on a NanoLive (Switzerland) 3D Cell Explorer fluo (AXT Pty Ltd, Warriewood, NSW) equipped with a NanoLive live cell incubator (AXT Pty Ltd). 1×10^4^ cells were seeded into a FluoroDish cell culture dish 35mm, 23mm well (World Precision Instruments, FD35) and maintained in phenol red free DMEM medium (Sigma-Aldrich, D1145) supplemented with 10% fetal bovine calf serum (Sigma-Aldrich, F9423), 2 mM glutamine (Sigma-Aldrich, G7513) and 1% penicillin-streptomycin for 48 hours. Immediately prior to imaging the medium was removed and replaced with 400 μL of the same medium, followed by transfer to the live cell incubator chamber of the 3D Cell Explorer. Cells were incubated at 37°C, 5% CO_2_ and 100% humidity for the duration of the time-lapse. Three dimensional holo-tomographic images were captured every 20 seconds for the duration of the time-lapse using the Nanolive STEVE software. For File S9 the center plane of each 96 slice stack was exported after capture using the built in STEVE export wizard as an .avi movie file. These files were exported at 5 frames per second (100x actual speed) to visualize cellular dynamics.

### Subcutaneous mouse xenograft tumors

Cells were expanded in culture for a maximum of 2 weeks before injection. Cells were trypsinized, pelleted, washed with PBS and stored on ice. Cell count was determined using a hemocytometer and trypan blue. Two million cells were resuspended in 100 µL 50:50 Matrigel:PBS for sub-cutaneous injection into the left flank of female Nod-Skid gamma mice (8-12 weeks) via a 27-gauge needle. Mice were monitored and once the tumor was palpable (∼2.5 -4 weeks), tumor growth was measured 3 times a week using calipers until tumors reach 1 cm^3^ in volume. Once a group in the cohort reached the maximum size of tumor all tumors were harvested, weighed, photographed and fixed with formalin. Mouse experiments were approved by The Australian National University Animal Experimentation Ethics Committee ethics protocol ANU A2017/16, and by Charles Sturt University Animal Care and Ethics Committee ethics protocol CSU A17046.

### Statistical Analyses

Unless specified otherwise, statistical analysis was performed using the SPSS package (IBM). For boxplot data depictions, whiskers represent quartiles 1 and 4 (with maximum and minimum values). Boxes represent quartiles 2 and 3, separated by the median, as generated by SPSS analyze data function. Datasets conforming with normal distribution were analyzed by ANOVA and post-hoc Bonferroni or Tukey HSD test (equal variance) or post-hoc Dunnett’s T3 Test (unequal variance). Statistical differences between divergent treatments of different cell lines were calculated using two way ANOVA and post-hoc pairwise comparisons. For non-parametric data sets Kruskal-Wallis or Kolmogorov-Smirnov tests were performed, as indicated in relevant figure legends.

**Figure S1.**
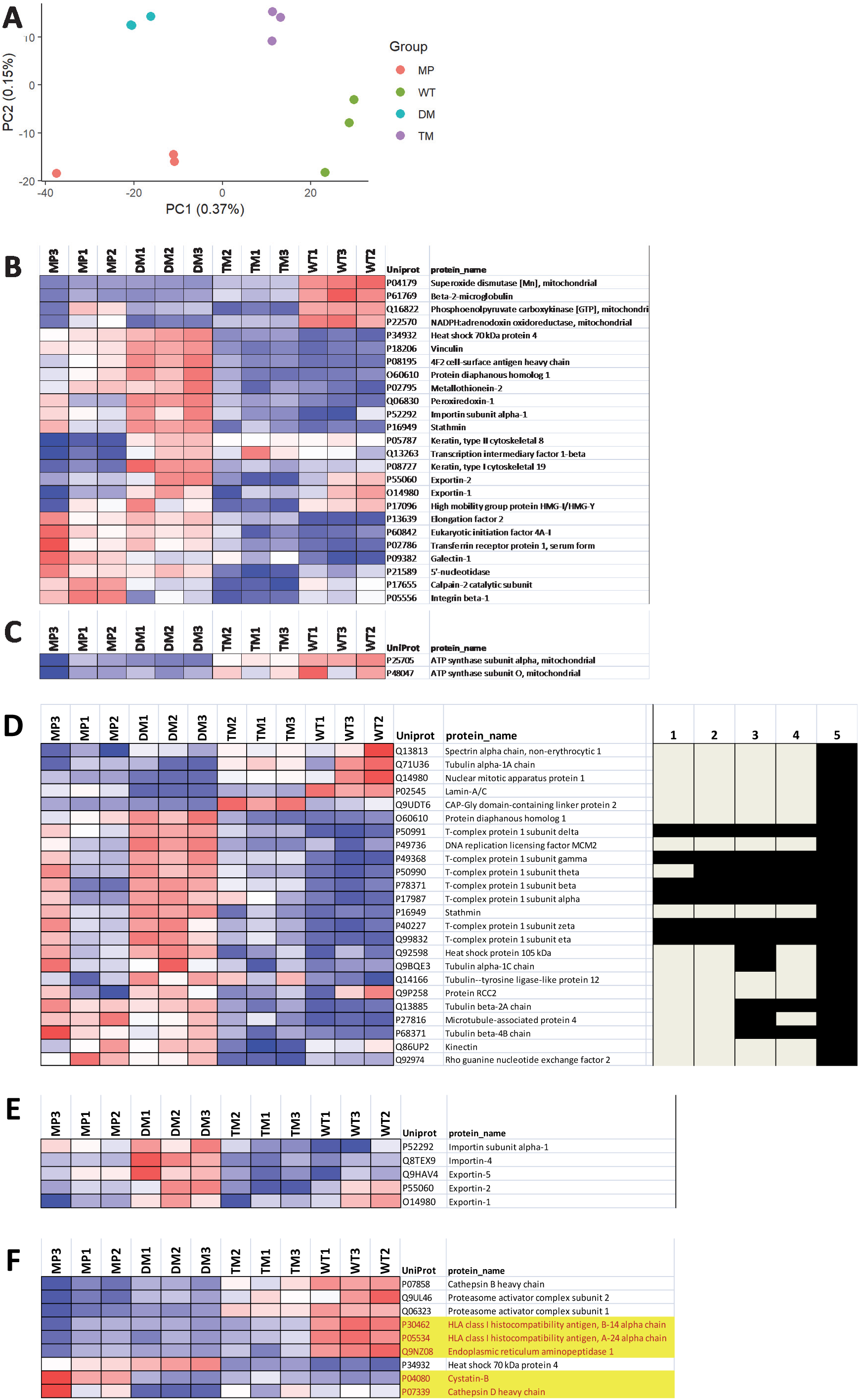
Detailed views of selected pathways identified by WebGestalt analyses. Related to Figure 3. All panels are adapted from File S6. Heat map colors follow Figure 3. (A) Principal component (PC) analysis of SWATH-MS proteomics results showing distribution of PC1 and PC2. PC1 corresponded to pathways associated with ribosomes and translation, while PC2 corresponded to pathways associated with mRNA splicing processing (see File S3). (B) Proteins associated with PI3K/Akt activity (WebGestalt Database: PC, DB_ID:1648, “Class I PI3K signaling events mediated by Akt”) are less abundant in TM cells. (C) F1/F0 ATPase subunits elevated in WT and TM cells. (D) Abundances of proteins associated with protein folding and microtubule function are altered by PGRMC1 phosphorylation status. Proteins detected in any of the following WebGestalt pathways or functions (1-4) or a manual search (5) are mapped against their expression profiles. 1) cellular component chaperonin-containing T-complex GO:0005832. 2) PC pathway Chaperonin-mediated protein folding DB_ID:710. 3) cellular component microtubule GO:0005874. 4) PC pathway Protein folding DB_ID:712. 5) Description from the list of 243 proteins (File S6) contains keywords “tubulin” or “microtubule” (manual search) (Adapted from File S6). 2-tailed t-test *p*-values for all sample comparisons are available in File S4. (E) Proteins associated with nuclear import/export that are elevated in DM cells. (F) Antigen processing and presentation enzymes are affected by PGRMC1 phosphorylation status. Manual additions to KEGG pathway ID:04612 “Antigen processing and presentation” (no yellow shading: from File S6 and File S5) are indicated with yellow highlighting.

**Figure S2.**
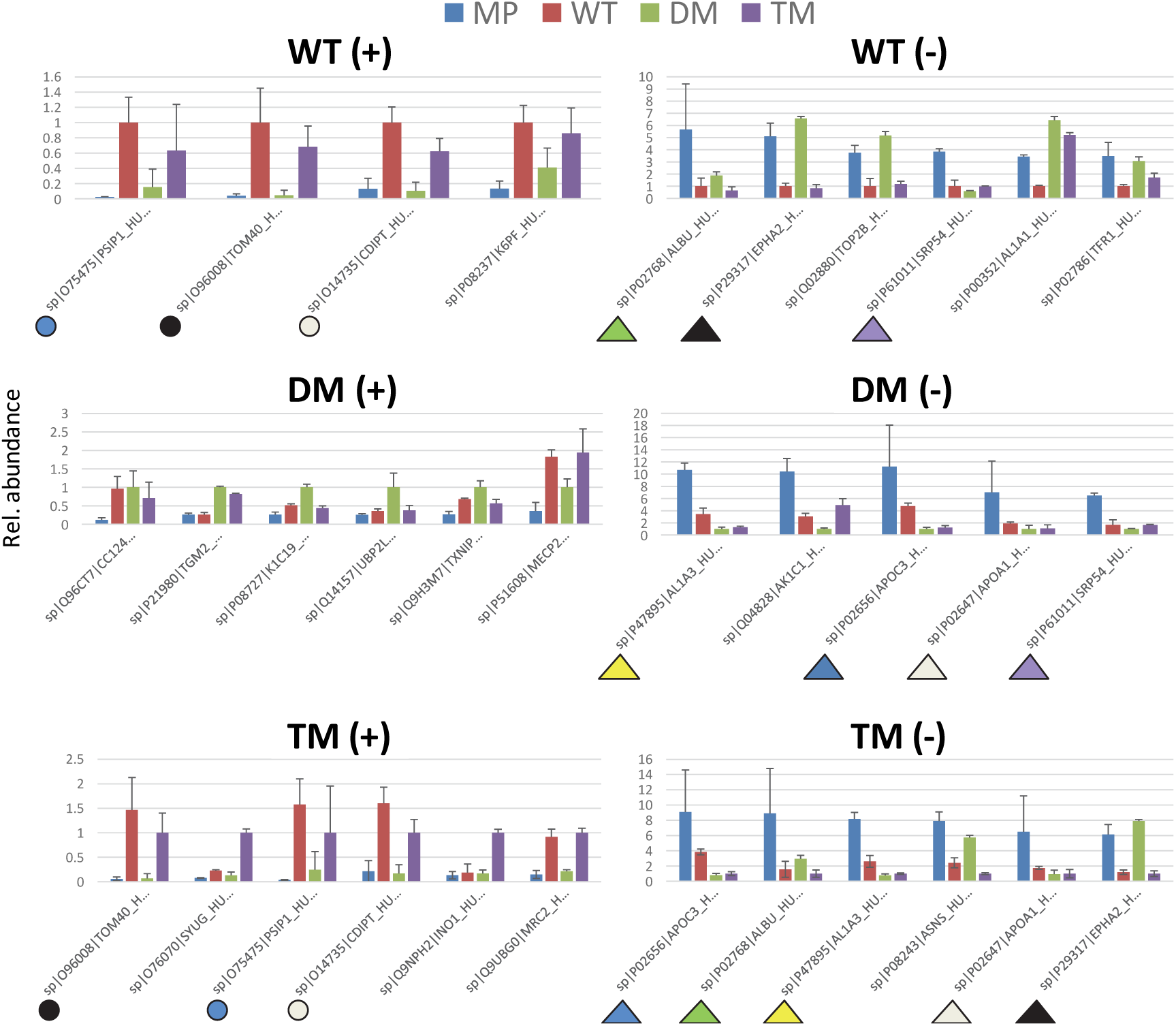
Highest and lowest differentially abundant proteins. Panels show the six most (+) and least (-) abundant proteins for each cell type that were significantly differentially abundant between cell types. Related to Figure 3. Identical colored symbols depict the same protein in different cell types, where circles represent high abundance and triangles represent low abundance. E.g. Mitochondrial import receptor subunit TOM40 (O96008), CDP-diacylglycerol-inositol 3-phosphatidyltransferase (CDIPT, O14735) (phosphatidylinositol synthesis) and transcriptional coactivator PSIP1 (O75475) are more abundant in WT and TM. Aldehyde dehydrogenase 1A3 (P47895), APOC3 (P02656) and APOA1 (P02647) are less abundant in DM and TM, whereas the receptor tyrosine kinase ephrin type-A receptor 2 (EPHA2, P29317) is among the lowest abundance differential proteins in WT and TM. Protein abundances (measured ion intensities) for all proteins are available in File S2.

**Figure S3.**
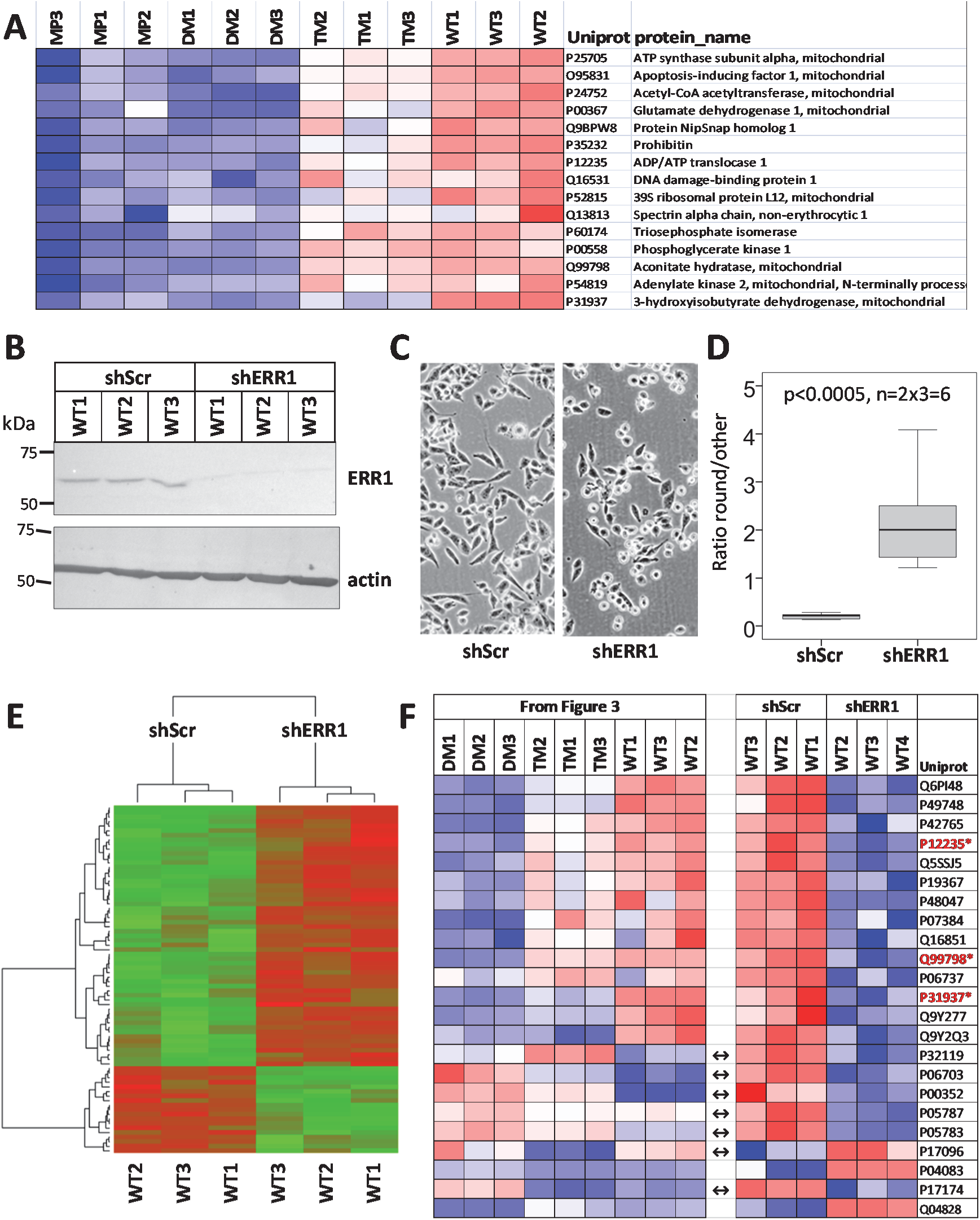
Proteins associated with predicted Estrogen Receptor Related 1 (ERR1) transcription factor activity are more abundant in WT and TM cells. Related to Figure 3 and File S6. Data are available via ProteomeXchange with identifier PXD014789. (A) Proteins quantified by SWATH-MS and predicted by WebGestalt Transcription Factor target enrichment analysis (WebGestalt Database: Transcription Target, Name: hsa_TGACCTY_V$ERR1_Q2, ID:DB_ID:2414) to be dependent upon ERR1 transcription (Adapted from File S6). Heat map colors follow Figure 3. (B) Western blot of shRNA attenuation of ERR1 (top panel) in WT cells, compared to scramble shRNA control. Lower panel: beta actin loading control. (C) ERR1 attenuation by shRNA induces morphological changes to WT cells. (D) Ratio of rounded to other cells as scored for replicate images from each of the cell lines from B. *p*<0.0005, 1 tailed t-test. (E) SWATH MS quantification of proteins significantly differentially abundant between cells expressing scramble shRNA or anti-ERR1 shRNA (according to the selection criteria of File S4). 57 proteins became more abundant after shRNA depletion of ERR1, and 19 proteins became less abundant. In this panel green depicts lower abundance and red depicts higher abundance. Brown depicts similar abundance. (F) Proteins present in (E) which are also present in the 243 protein list of File S6. Heat map colors follow Figure 3. The left panel shows expression in the original result of File S6. The right panel shows expression in the presence of scramble shRNA or anti-ESSR1 shRNA. All proteins from Figure S3A except Q9BPW8 were detected in the shRNA experiment. Double headed arrows indicate proteins which differ in expression tendency between WT (left) and WT-scramble shRNA cells (right). This may be caused by the puromycin selection of both sh-scr and shERR1 cells but not the parental WT cells, however this requires further investigation. Proteins with Uniprot ID highlighted by asterisk (bold red) are those both originally predicted by WebGestalt to be associated with ERR1 transcription factor (A), and which exhibited significantly altered abundance after shRNA attenuation of ERR1 protein.

**Figure S4.**
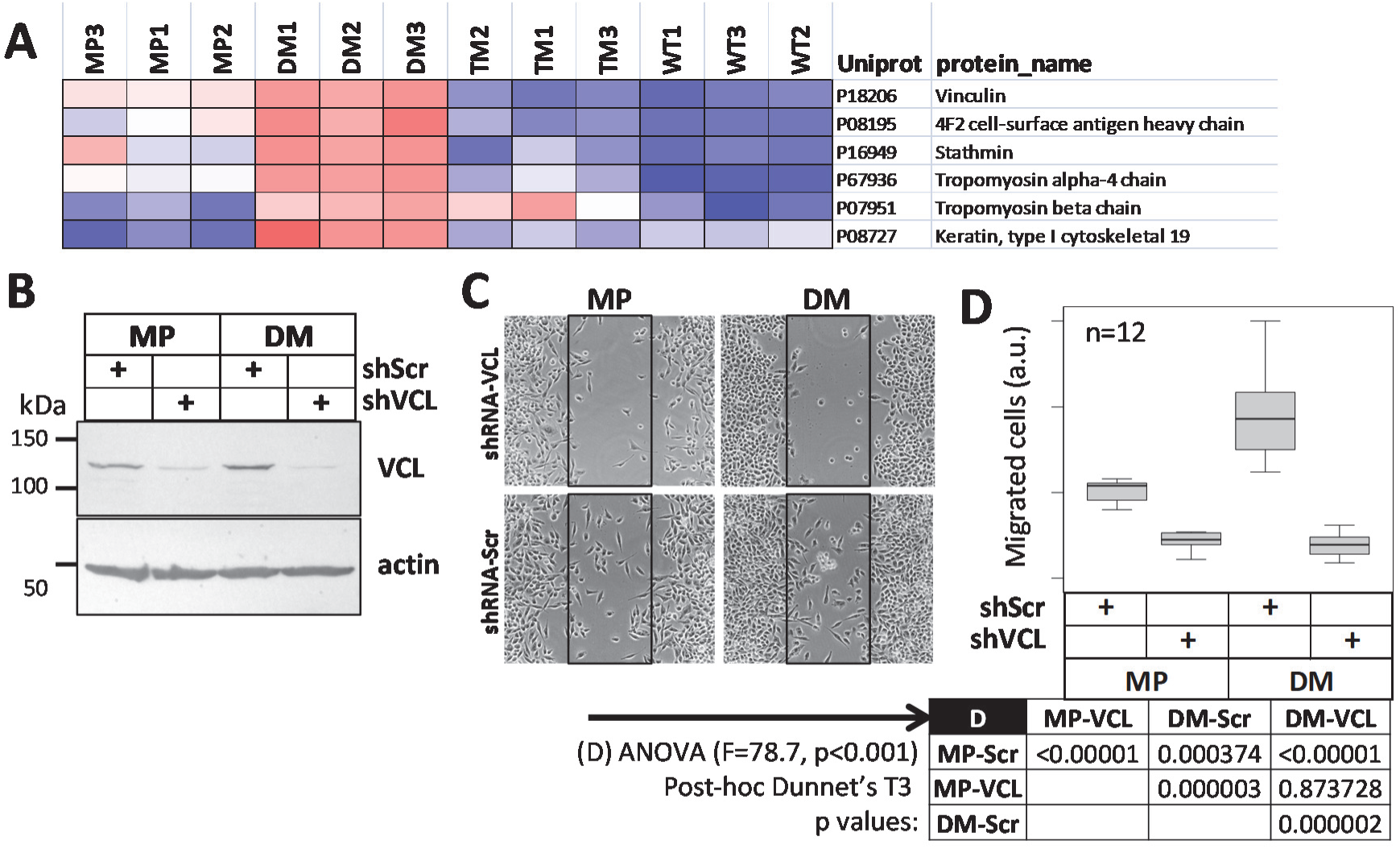
The DM migratory phenotype is dependent upon vinculin. Related to Figure 3. (A) Proteins associated with actin-myosin contraction (WebGestalt Database: PC, DB_ID:144, “Smooth Muscle Contraction”) are more abundant in DM cells. Heat map colors follow Figure 3. (B) Western blot of Vinculin (VCL) and actin levels in cells expressing scramble control (shScr) or anti Vinclulin (shVNC) shRNA. (C) Scratch assays as indicated were performed following the methods of Figure 1E-F. (D) Boxplot showing results of 12 replicates of (C). ANOVA gave F=78.7, p <0.00001. The table shows post-hoc Dunnet’s T3 *p*-values.

**Figure S5.**
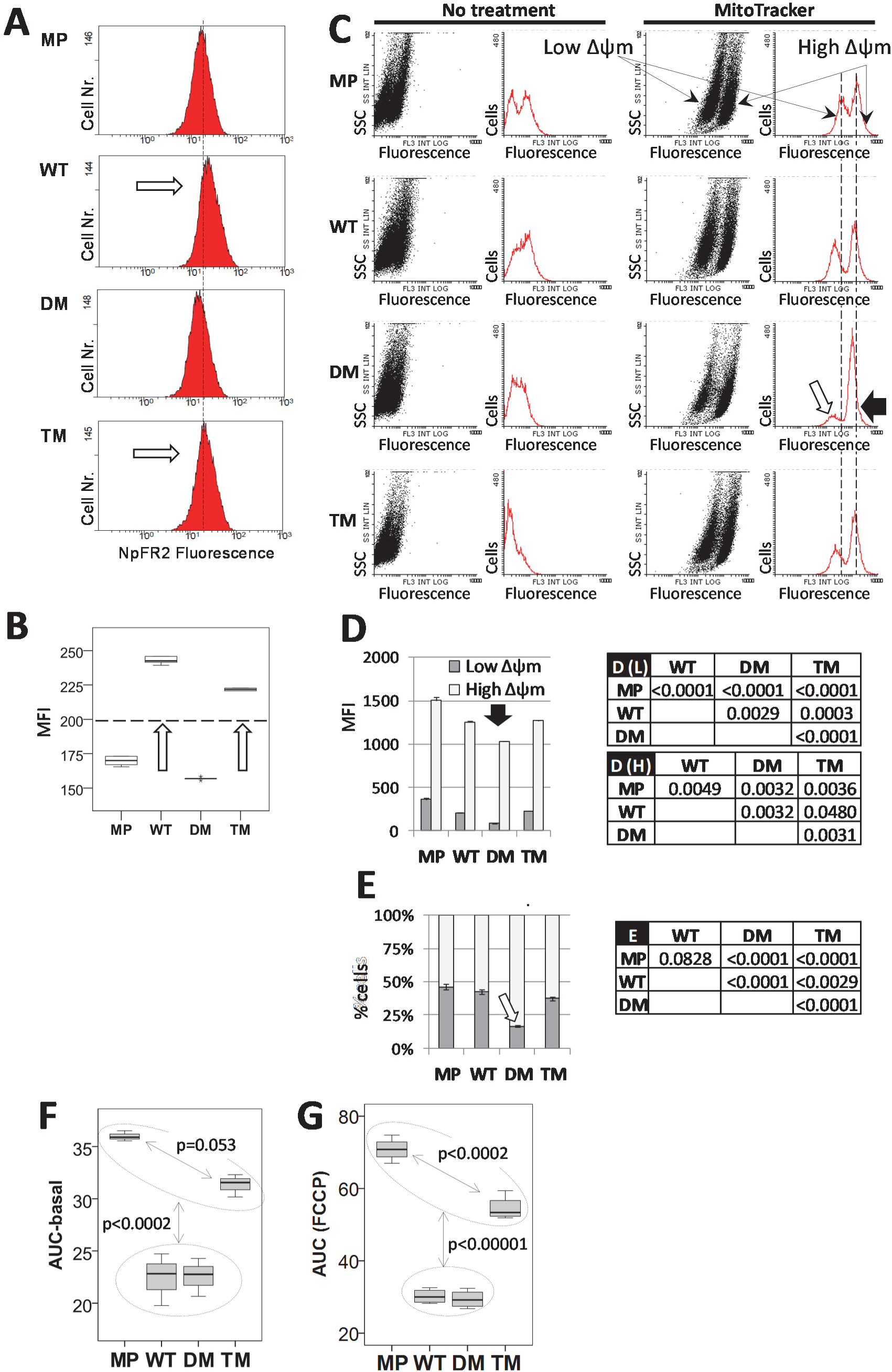
PGRMC1 phosphorylation status affects mitochondrial function. Related to Figure 5. (A) Representative flow cytometry results of cells labeled with NpFR2. The percentage of the cell population to the right of the dashed reference line (interval labeled “B”, marked by the dotted line) is quantified for each measurement. White arrows indicate more oxidized NpFR2 fluorescence in WT and TM cells. (B) Boxplots of the percentage of cells exhibiting >10 fluorescent intensity units to the right of the reference line in (A). n=6 for each cell type, being 6 replicates of MP cells, or duplicate measurements of each of 3 independent lines 1-3 (n=3x2=6) for WT, DM and TM cells. White arrows indicate the same differences as in (A). There was significant difference between the means (Kruskal-Wallis Test *p*<0.0001). Independent sample median tests revealed that all medians were significantly different from one another (*p*<0.001). (C) Representative flow cytometry results of cells labeled with MitoTracker. The respective panels depict fluorescence intensity (x axis) plotted against either side scatter (left/black plots) or cell number (right/red plots), with or without the addition of MitoTracker as indicated. MitoTracker affinity for mitochondria is increased with higher mitochondrial membrane potential (Δψm), such that respective left MitoTracker-treated populations/peaks represent low Δψm populations, and respective right MitoTracker-treated populations/peaks represent high ψm populations. Vertical dotted lines for Mitotracker-treated cells depict the MFI for MP cell low and high Δψm populations. The black arrow for DM cells with Mitotracker highlights reduced MFI of both the low and high Δψm DM cell populations. The white arrow highlights the reduced proportion of DM cells in the low Δψm peak. The same arrows are depicted for replicates in (D) and (E) respectively. (D) Median Fluorescent Intensity (MFI) for each of the cell populations observed from (C). n=6 measurements per cell type, as per (B). Error bars depict standard deviation. The low and high Δψm populations correspond to the respective populations from (C). For low Δψm cells the means were significantly different between cell types (ANOVA, post-hoc Dunnet’s T3, *p*<0.003). For high Δψm cells WT vs. TM (*p*<0.05) and other comparisons (*p*<0.004) were significantly different (Kruskall-Wallis). Tables D (L) and D(H) show pairwise comparison *p*-values for low Δψm (F(L)) and high Δψm (F(H)) cells. (E) The percentage of cells in each of the low and high Δψm populations from (C) and (D), from the same data. Standard deviation error bars for the respective high Δψm (upper error bar) and low Δψm (lower error bar) cell populations are given for each cell type. Color-coding and all other terminology follows (C). For low Δψm cells, Shapiro-Wilk’s test showed normal distributions, and Levene’s test revealed non-homogenous variance (p=0.04). Table E shows *p*-values for ANOVA/post hoc Dunnet’s T3 test pairwise comparisons. (F, G) Area under curve (AUC) plots for basal and FCCP curves from (Figure 5C) showing one way ANOVA/post-hoc Bonferroni *p*-values between samples in the respective indicated regions.

### Supporting Information files

**File S1**. A zip archive containing time lapse mp4 movies of migrating cells in scratch assays. Related to Figure 1. Images were taken at 10 minute intervals over 36 hours and are replayed over 65 seconds at 9 frames per second (2000x real time). The presented 32.4 MB MP4 files were generated from the original 666MB (6000x real time) .avi files (length 21 sec, frame width x height 1024×1024, Data rate/bit rate 287462 kbps, frame rate 100 fps) by processing in Adobe Premiere Pro 2017 using the following settings: Preset “custom”; Width: 1024; Height 1024; Aspect Square Pixels (1.0); Field Order: progressive; Profile: high; Target Bitrate [Mbps] 4.08; Maximum Bitrate [Mbps] 4.39; 9 fps; No Audio; Output as standard .mp4; Time Interpolation: Frame Sampling; Stream Compatibility: Standard; Variable bit rate, single pass; Number of frames 00:01:04:09. The zip archive contains 4 files with filenames representing cell type and date of measurement (yyyymmdd).

(A) A_MP_20170807.mp4

(B) B_WT_20170809.mp4

(C) C_DM_20170815.mp4

(D) D_TM_20170817.mp4

**File S2**. An Excel file showing experimental design, normalized ion intensities for 1330 proteins identified by SWATH-MS proteomics, and six pairwise comparisons between the 4 sample types [1) MP v. WT, 2) MP v. DM, 3) MP v. TM, 4) WT v. DM, 5) WT v. TM, and 6) DM v. TM]. Related to Figure 3. The first tab contains a detailed descriptive legend. Data are available via ProteomeXchange with identifier PXD014716.

**File S3**. Principal component analysis results for pathways associated with SWATH-MS proteomics results.

**File S4.** An excel file containing proteomics results for 243 proteins which fulfil stringency criteria of t-test *p*-value of less than 0.05, and a fold change greater than 1.5 by both the protein and peptide approaches from File S2. Related to Figure 3. Column B shows “red” (more abundant in comparative sample 1) and “blue” (less abundant in sample 1) significantly differential proteins for each pair wise comparison which were later used for “red” and ‘blue” WebGestalt pathways enrichment analysis (File S5). Comparisons follow File S2.

**File S5.** WebGestalt pathway mapping excel results file for red and blue proteins from File S4. Related to Figure 3.

(A) WebGestalt Features (GO, KEGG Pathways, Wikipathways, Pathway Commons, Transcription Factors) significant at the adjP≤0.001 level between any 2 comparisons.

(B) Features from A, viewed at the adjP<0.05 level for each red and blue comparison, and showing adjusted *p*-values (adjP) for each comparison where adjP<0.05 (or adjP<0.1 for two comparisons as indicated by paler coloring). Blue means that proteins associated with a given the feature were less abundant in that cell line, with red indicating higher abundance. Because separate WebGestalt analyses were performed for red and blue lists of proteins from File S4, some features were significant for both red and blue. In that case the color and adjP for the most significant analysis is given, with the other cell being colored black.

**File S6.** An excel file with heat map protein IDs and pathways for red vs. blue pathways adjP<0.001. Related to Figure 3. This file is derived from the results of File S5, and is the source file for Figure 3. Proteins suggested by clustering by inferred models of evolution (CLIME) analysis to co-evolve with PGRMC1 with log likelihood ratio greater than 12 (Cahill and Medlock, 2017) are present in the list, marked yellow for mitochondrial localization (WebGestalt GO:0005739) or green for cytoplasmic (P00387).

**File S7.** Original WebGestalt results files for the analysis of Figure 3 and File S5. Related to Figure 3. WebGestalt results files for red and blue comparison pathways adjP<0.001.

**File S8.** Original WebGestalt results files for red and blue comparison pathways where adjP<0.1. Related to Figure 3.

**File S9.** A zip archive containing time lapse Holo-tomographic video .avi files of cells. These images are based upon differences in refractive index (Ali et al., 2016), and are provided for the dynamic visualization of mitochondria. Prominent visible features include small white lipid droplets and cholesterol-rich mitochondria (Cahill and Medlock, 2017), as well as nuclear membrane and nucleoli. The previously described MIA PaCa-2 cell bleb-like protrusions (Gradiz et al., 2016) are apparent as highly dynamic rearrangements of the cytoplasmic membrane, which may contribute to intercellular communication.

(A) MP cells

(B) WT cells

(C) DM cells

(D) TM cells

## LIST OF ABBREVIATIONS

adjP: Benjamini-Hochberg adjusted *p-*value
avFF: average mitochondrial form factor
CK2: Casein kinase 2
DM: hemagglutinin-tagged PGRMC1 S57A/S181 double mutant
EMT: epithelial-mesenchymal transition
ERR1: estrogen receptor related 1
FF: mitochondrial form factor
hPSCs: human pluripotent stem cells
IDA: information dependent acquisition
LDLR: low density lipoprotein receptor
MAT: mesenchymal-amoeboid transition
MP: MIA PaCa-2 pancreatic cancer
MS/MS: tandem mass spectrometry
MTT: 3-(4,5-dimethylthiazolyl-2)-2,5-diphenyltetrazolium bromide
NpFR2: Naphthalimide-flavin redox sensor 2
P4: progesterone
PAQR7: progestin and adipoQ receptor 7
PGRMC1: Progesterone Receptor Membrane Component 1
PGRMC2: Progesterone Receptor Membrane Component 2
ROCK: Rho kinase
ROCKI: Rho kinase inhibitor
RPPA: Reverse Phase Protein Array
S2R: Sigma-2 receptor
SH2: Src homology 2
SH3: Src homology 3
SWATH-MS: Sequential Window Acquisition of all Theoretical Mass Spectrometry
TM: hemagglutinin-tagged PGRMC1 S57A/Y180F/S181 triple mutant
TRiC: T-complex protein-1 ring complex
WT: hemagglutinin-tagged PGRMC1 wild type
Δψm: mitochondrial membrane potential

## Notes

#### Summary of Updates

Manuscript was reformatted, with re-arrangement of some figure panels and editing of text, and shortening of title. The data and conclusions are unaltered.

