## Supplementary material for "PGRMC1 phosphorylation and cell plasticity 1: glycolysis, mitochondria, tumor growth": File S3

Proteomics analysis


Code 

- Show All Code
- Hide All Code

### Proteomics analysis

###### *Ellis Patrick*

###### *February 18, 2019*

### 1 Basic QC

#### 1.1 PCA

If we look at the first PCs, all the biological differences appear very strong so technical artifacts are likely to have little influence. When I perform further normalization I see no drastic change in the PCA plots .

```
pca = prcomp(log2(t(data)),scale = TRUE)
summary(pca)
```

```
## Importance of components:
##                            PC1     PC2     PC3      PC4     PC5     PC6
## Standard deviation     22.1978 14.3094 12.9749 11.04819 9.70028 8.61227
## Proportion of Variance  0.3705  0.1540  0.1266  0.09178 0.07075 0.05577
## Cumulative Proportion   0.3705  0.5244  0.6510  0.74279 0.81354 0.86931
##                            PC7     PC8     PC9    PC10    PC11      PC12
## Standard deviation     7.14468 6.70831 5.76817 5.05142 4.35730 1.543e-13
## Proportion of Variance 0.03838 0.03384 0.02502 0.01919 0.01428 0.000e+00
## Cumulative Proportion  0.90769 0.94152 0.96654 0.98572 1.00000 1.000e+00
```

```
prop = signif(summary(pca)$importance['Proportion of Variance',],2)

df = data.frame(pca$x, design[colnames(data),])
p1 = ggplot(df, aes(x = PC1, y = PC2, colour = Group)) + geom_point(size = 3) + theme_classic() + xlab(paste('PC1 (',prop['PC1'],'%)',sep = ''))+ ylab(paste('PC2 (',prop['PC2'],'%)',sep = ''))
p2 = ggplot(df, aes(x = PC3, y = PC4, colour = Group)) + geom_point(size = 3) + theme_classic() + xlab(paste('PC3 (',prop['PC3'],'%)',sep = ''))+ ylab(paste('PC4 (',prop['PC4'],'%)',sep = ''))

library(gridExtra)
grid.arrange(p1,p2)
```

```
library(org.Hs.eg.db)
#Read in pathways
keys = unlist(lapply(strsplit(rownames(data),'\\|'),function(x)x[2]))
names(keys) = rownames(data)
keysR = rownames(data)
names(keysR) = keys
mapKeys = mapIds(org.Hs.eg.db, keys = keys,column = 'ENTREZID', keytype = "UNIPROT",multiVals="CharacterList")
mapKeysRev = unlist(reverseSplit(mapKeys))
#map = unlist(lapply(mapKeys,function(x)x[1]))
#mapR = names(map)
#names(mapR) = map

Uniprot2prot = unlist(lapply(strsplit(rownames(data),'\\|'),function(x)x[3]))
Uniprot2prot = unlist(lapply(strsplit(Uniprot2prot,'_'),function(x)x[1]))
names(Uniprot2prot) = rownames(data)
```

```
library(org.Hs.eg.db)
file = read.delim('homo-sapiens-9606.txt',sep = '$',col.names=FALSE)
file = apply(file,1,function(x)unlist(strsplit(x,'\t')))
def = unlist(lapply(file,function(x)x[2]))
names(def) = unlist(lapply(file,function(x)x[1]))
names(file) = unlist(lapply(file,function(x)x[1]))

file = lapply(file,function(x)unlist(strsplit(unlist(strsplit(x,':')),',')))
#file = lapply(file,intersect,keys)
#PC2symbol = lapply(file,function(x)keysR[x])

u = unique(keys(org.Hs.eg.db,"UNIPROT"))
keysRU = keysR[u]
names(keysRU) = u
keysRU[is.na(keysRU)] = 1:sum(is.na(keysRU))

file = lapply(file,intersect,keys(org.Hs.eg.db,"UNIPROT"))
PC2symbol = lapply(file,function(x)keysRU[x])


PClen = unlist(lapply(PC2symbol,length))
```

```
vec = pca$rot[,1]
wtest.1 = unlist(lapply(PC2symbol[PClen>5&PClen<5000],function(x)wilcoxGST(names(vec)%in%x,vec,alternative =  'mixed')))

vec = pca$rot[,2]
wtest.2 = unlist(lapply(PC2symbol[PClen>5&PClen<5000],function(x)wilcoxGST(names(vec)%in%x,vec,alternative =  'mixed')))


#plot(-log10(wtest.1),-log10(wtest.2))


u = names(wtest.1)
OR = data.frame(Pathway = u,Pvalue.PC1 = signif(wtest.1[u],2), AdjustedPvalue.PC1 = signif(p.adjust(wtest.1[u],'BH'),2),Pvalue.PC2 = signif(wtest.2[u],2), AdjustedPvalue.PC2 = signif(p.adjust(wtest.2[u],'BH'),2), NumberInPathway = PClen[u])
OR = OR[order(OR$Pvalue.PC1),]
```

We find 33 Pathway Commons pathways are significant changed for PC1 and 5 changed for PC2.

```
DT::datatable(OR,  escape = FALSE,
                            extensions = 'Buttons',

                            options = list(
                                paging = TRUE,
                                searching = TRUE,
                                fixedColumns = TRUE,
                                autoWidth = TRUE,
                                ordering = TRUE,
                                dom = 'Bfrtip',
                                buttons = c('copy', 'csv', 'excel')
                            )
                       )
```
