## Supplementary material for "PGRMC1 phosphorylation and cell plasticity 1: glycolysis, mitochondria, tumor growth": File S7: final_drug_geneset_file_1438481923.html

Anchored HTML File of EIDs

|  |  |
| --- | --- |
|  | WEB-based GEne SeT AnaLysis Toolkit |
|  | ***Translating gene lists into biological insights...*** |

---

  

| **Database:drug      &nbspName:cefacetrile      &nbspID:DB\_ID:PA164776752** | | | | | | |
| --- | --- | --- | --- | --- | --- | --- |
| C=100; O=12; E=0.53; R=22.54; rawP=1.97e-13; adjP=1.84e-12 | | | | | | |
| Index | UserID | Value | Gene Symbol | Gene Name | EntrezGene | Ensembl |
| 1 | Q99798 | NA | ACO2 | aconitase 2, mitochondrial | 50 | ENSG00000100412 |
| 2 | P00558 | NA | PGK1 | phosphoglycerate kinase 1 | 5230 | ENSG00000102144 |
| 3 | P02768 | NA | ALB | albumin | 213 | ENSG00000163631 |
| 4 | P40926 | NA | MDH2 | malate dehydrogenase 2, NAD (mitochondrial) | 4191 | ENSG00000146701 |
| 5 | P35232 | NA | PHB | prohibitin | 5245 | ENSG00000167085 |
| 6 | P30084 | NA | ECHS1 | enoyl CoA hydratase, short chain, 1, mitochondrial | 1892 | ENSG00000127884 |
| 7 | P09382 | NA | LGALS1 | lectin, galactoside-binding, soluble, 1 | 3956 | ENSG00000100097 |
| 8 | P40227 | NA | CCT6A | chaperonin containing TCP1, subunit 6A (zeta 1) | 908 | ENSG00000146731 |
| 9 | P25705 | NA | ATP5A1 | ATP synthase, H+ transporting, mitochondrial F1 complex, alpha subunit 1, cardiac muscle | 498 | ENSG00000152234 |
| 10 | P21796 | NA | VDAC1 | voltage-dependent anion channel 1 | 7416 | ENSG00000213585 |
| 11 | P60174 | NA | TPI1 | triosephosphate isomerase 1 | 7167 | ENSG00000111669 |
| 12 | P04083 | NA | ANXA1 | annexin A1 | 301 | ENSG00000135046 |

  
  

| **Database:drug      &nbspName:netilmicin      &nbspID:DB\_ID:PA164754913** | | | | | | |
| --- | --- | --- | --- | --- | --- | --- |
| C=97; O=12; E=0.52; R=23.23; rawP=1.36e-13; adjP=1.84e-12 | | | | | | |
| Index | UserID | Value | Gene Symbol | Gene Name | EntrezGene | Ensembl |
| 1 | Q99798 | NA | ACO2 | aconitase 2, mitochondrial | 50 | ENSG00000100412 |
| 2 | P00558 | NA | PGK1 | phosphoglycerate kinase 1 | 5230 | ENSG00000102144 |
| 3 | P02768 | NA | ALB | albumin | 213 | ENSG00000163631 |
| 4 | P40926 | NA | MDH2 | malate dehydrogenase 2, NAD (mitochondrial) | 4191 | ENSG00000146701 |
| 5 | P35232 | NA | PHB | prohibitin | 5245 | ENSG00000167085 |
| 6 | P30084 | NA | ECHS1 | enoyl CoA hydratase, short chain, 1, mitochondrial | 1892 | ENSG00000127884 |
| 7 | P09382 | NA | LGALS1 | lectin, galactoside-binding, soluble, 1 | 3956 | ENSG00000100097 |
| 8 | P40227 | NA | CCT6A | chaperonin containing TCP1, subunit 6A (zeta 1) | 908 | ENSG00000146731 |
| 9 | P25705 | NA | ATP5A1 | ATP synthase, H+ transporting, mitochondrial F1 complex, alpha subunit 1, cardiac muscle | 498 | ENSG00000152234 |
| 10 | P21796 | NA | VDAC1 | voltage-dependent anion channel 1 | 7416 | ENSG00000213585 |
| 11 | P60174 | NA | TPI1 | triosephosphate isomerase 1 | 7167 | ENSG00000111669 |
| 12 | P04083 | NA | ANXA1 | annexin A1 | 301 | ENSG00000135046 |

  
  

| **Database:drug      &nbspName:cefotaxime      &nbspID:DB\_ID:PA448852** | | | | | | |
| --- | --- | --- | --- | --- | --- | --- |
| C=100; O=12; E=0.53; R=22.54; rawP=1.97e-13; adjP=1.84e-12 | | | | | | |
| Index | UserID | Value | Gene Symbol | Gene Name | EntrezGene | Ensembl |
| 1 | Q99798 | NA | ACO2 | aconitase 2, mitochondrial | 50 | ENSG00000100412 |
| 2 | P00558 | NA | PGK1 | phosphoglycerate kinase 1 | 5230 | ENSG00000102144 |
| 3 | P02768 | NA | ALB | albumin | 213 | ENSG00000163631 |
| 4 | P40926 | NA | MDH2 | malate dehydrogenase 2, NAD (mitochondrial) | 4191 | ENSG00000146701 |
| 5 | P35232 | NA | PHB | prohibitin | 5245 | ENSG00000167085 |
| 6 | P30084 | NA | ECHS1 | enoyl CoA hydratase, short chain, 1, mitochondrial | 1892 | ENSG00000127884 |
| 7 | P09382 | NA | LGALS1 | lectin, galactoside-binding, soluble, 1 | 3956 | ENSG00000100097 |
| 8 | P40227 | NA | CCT6A | chaperonin containing TCP1, subunit 6A (zeta 1) | 908 | ENSG00000146731 |
| 9 | P25705 | NA | ATP5A1 | ATP synthase, H+ transporting, mitochondrial F1 complex, alpha subunit 1, cardiac muscle | 498 | ENSG00000152234 |
| 10 | P21796 | NA | VDAC1 | voltage-dependent anion channel 1 | 7416 | ENSG00000213585 |
| 11 | P60174 | NA | TPI1 | triosephosphate isomerase 1 | 7167 | ENSG00000111669 |
| 12 | P04083 | NA | ANXA1 | annexin A1 | 301 | ENSG00000135046 |

  
  

| **Database:drug      &nbspName:ciprofloxacin      &nbspID:DB\_ID:PA449009** | | | | | | |
| --- | --- | --- | --- | --- | --- | --- |
| C=111; O=12; E=0.59; R=20.30; rawP=7.06e-13; adjP=4.94e-12 | | | | | | |
| Index | UserID | Value | Gene Symbol | Gene Name | EntrezGene | Ensembl |
| 1 | Q99798 | NA | ACO2 | aconitase 2, mitochondrial | 50 | ENSG00000100412 |
| 2 | P00558 | NA | PGK1 | phosphoglycerate kinase 1 | 5230 | ENSG00000102144 |
| 3 | P02768 | NA | ALB | albumin | 213 | ENSG00000163631 |
| 4 | P40926 | NA | MDH2 | malate dehydrogenase 2, NAD (mitochondrial) | 4191 | ENSG00000146701 |
| 5 | P35232 | NA | PHB | prohibitin | 5245 | ENSG00000167085 |
| 6 | P30084 | NA | ECHS1 | enoyl CoA hydratase, short chain, 1, mitochondrial | 1892 | ENSG00000127884 |
| 7 | P09382 | NA | LGALS1 | lectin, galactoside-binding, soluble, 1 | 3956 | ENSG00000100097 |
| 8 | P40227 | NA | CCT6A | chaperonin containing TCP1, subunit 6A (zeta 1) | 908 | ENSG00000146731 |
| 9 | P25705 | NA | ATP5A1 | ATP synthase, H+ transporting, mitochondrial F1 complex, alpha subunit 1, cardiac muscle | 498 | ENSG00000152234 |
| 10 | P21796 | NA | VDAC1 | voltage-dependent anion channel 1 | 7416 | ENSG00000213585 |
| 11 | P60174 | NA | TPI1 | triosephosphate isomerase 1 | 7167 | ENSG00000111669 |
| 12 | P04083 | NA | ANXA1 | annexin A1 | 301 | ENSG00000135046 |

  
  

| **Database:drug      &nbspName:gentamicin      &nbspID:DB\_ID:PA449753** | | | | | | |
| --- | --- | --- | --- | --- | --- | --- |
| C=69; O=8; E=0.37; R=21.77; rawP=3.17e-09; adjP=1.78e-08 | | | | | | |
| Index | UserID | Value | Gene Symbol | Gene Name | EntrezGene | Ensembl |
| 1 | P40227 | NA | CCT6A | chaperonin containing TCP1, subunit 6A (zeta 1) | 908 | ENSG00000146731 |
| 2 | Q99798 | NA | ACO2 | aconitase 2, mitochondrial | 50 | ENSG00000100412 |
| 3 | P25705 | NA | ATP5A1 | ATP synthase, H+ transporting, mitochondrial F1 complex, alpha subunit 1, cardiac muscle | 498 | ENSG00000152234 |
| 4 | P00558 | NA | PGK1 | phosphoglycerate kinase 1 | 5230 | ENSG00000102144 |
| 5 | P02768 | NA | ALB | albumin | 213 | ENSG00000163631 |
| 6 | P40926 | NA | MDH2 | malate dehydrogenase 2, NAD (mitochondrial) | 4191 | ENSG00000146701 |
| 7 | P60174 | NA | TPI1 | triosephosphate isomerase 1 | 7167 | ENSG00000111669 |
| 8 | P30084 | NA | ECHS1 | enoyl CoA hydratase, short chain, 1, mitochondrial | 1892 | ENSG00000127884 |

  
  

| **Database:drug      &nbspName:adenine      &nbspID:DB\_ID:PA448048** | | | | | | |
| --- | --- | --- | --- | --- | --- | --- |
| C=147; O=10; E=0.78; R=12.78; rawP=6.20e-09; adjP=2.89e-08 | | | | | | |
| Index | UserID | Value | Gene Symbol | Gene Name | EntrezGene | Ensembl |
| 1 | P00491 | NA | PNP | purine nucleoside phosphorylase | 4860 | ENSG00000198805 |
| 2 | P12235 | NA | SLC25A4 | solute carrier family 25 (mitochondrial carrier; adenine nucleotide translocator), member 4 | 291 | ENSG00000151729 |
| 3 | O95831 | NA | AIFM1 | apoptosis-inducing factor, mitochondrion-associated, 1 | 9131 | ENSG00000156709 |
| 4 | P13804 | NA | ETFA | electron-transfer-flavoprotein, alpha polypeptide | 2108 | ENSG00000140374 |
| 5 | Q9Y277 | NA | VDAC3 | voltage-dependent anion channel 3 | 7419 | ENSG00000078668 |
| 6 | P48735 | NA | IDH2 | isocitrate dehydrogenase 2 (NADP+), mitochondrial | 3418 | ENSG00000182054 |
| 7 | P07741 | NA | APRT | adenine phosphoribosyltransferase | 353 | ENSG00000198931 |
| 8 | P21796 | NA | VDAC1 | voltage-dependent anion channel 1 | 7416 | ENSG00000213585 |
| 9 | P38117 | NA | ETFB | electron-transfer-flavoprotein, beta polypeptide | 2109 | ENSG00000105379 |
| 10 | P43490 | NA | NAMPT | nicotinamide phosphoribosyltransferase | 10135 | ENSG00000105835 |

  
  

| **Database:drug      &nbspName:nadh      &nbspID:DB\_ID:PA164755085** | | | | | | |
| --- | --- | --- | --- | --- | --- | --- |
| C=223; O=10; E=1.19; R=8.42; rawP=3.19e-07; adjP=1.28e-06 | | | | | | |
| Index | UserID | Value | Gene Symbol | Gene Name | EntrezGene | Ensembl |
| 1 | P51649 | NA | ALDH5A1 | aldehyde dehydrogenase 5 family, member A1 | 7915 | ENSG00000112294 |
| 2 | P40926 | NA | MDH2 | malate dehydrogenase 2, NAD (mitochondrial) | 4191 | ENSG00000146701 |
| 3 | O95831 | NA | AIFM1 | apoptosis-inducing factor, mitochondrion-associated, 1 | 9131 | ENSG00000156709 |
| 4 | Q99714 | NA | HSD17B10 | hydroxysteroid (17-beta) dehydrogenase 10 | 3028 | ENSG00000072506 |
| 5 | P04179 | NA | SOD2 | superoxide dismutase 2, mitochondrial | 6648 | ENSG00000112096 |
| 6 | P00367 | NA | GLUD1 | glutamate dehydrogenase 1 | 2746 | ENSG00000148672 |
| 7 | P21796 | NA | VDAC1 | voltage-dependent anion channel 1 | 7416 | ENSG00000213585 |
| 8 | P00352 | NA | ALDH1A1 | aldehyde dehydrogenase 1 family, member A1 | 216 | ENSG00000165092 |
| 9 | P43490 | NA | NAMPT | nicotinamide phosphoribosyltransferase | 10135 | ENSG00000105835 |
| 10 | P32322 | NA | PYCR1 | pyrroline-5-carboxylate reductase 1 | 5831 | ENSG00000183010 |

  
  

| **Database:drug      &nbspName:iron      &nbspID:DB\_ID:PA450087** | | | | | | |
| --- | --- | --- | --- | --- | --- | --- |
| C=142; O=8; E=0.76; R=10.58; rawP=9.24e-07; adjP=3.23e-06 | | | | | | |
| Index | UserID | Value | Gene Symbol | Gene Name | EntrezGene | Ensembl |
| 1 | O75439 | NA | PMPCB | peptidase (mitochondrial processing) beta | 9512 | ENSG00000105819 |
| 2 | P02792 | NA | FTL | ferritin, light polypeptide | 2512 | ENSG00000087086 |
| 3 | Q99798 | NA | ACO2 | aconitase 2, mitochondrial | 50 | ENSG00000100412 |
| 4 | P13804 | NA | ETFA | electron-transfer-flavoprotein, alpha polypeptide | 2108 | ENSG00000140374 |
| 5 | P04179 | NA | SOD2 | superoxide dismutase 2, mitochondrial | 6648 | ENSG00000112096 |
| 6 | P31040 | NA | SDHA | succinate dehydrogenase complex, subunit A, flavoprotein (Fp) | 6389 | ENSG00000073578 |
| 7 | P02786 | NA | TFRC | transferrin receptor (p90, CD71) | 7037 | ENSG00000072274 |
| 8 | P38117 | NA | ETFB | electron-transfer-flavoprotein, beta polypeptide | 2109 | ENSG00000105379 |

  
  

| **Database:drug      &nbspName:adenosine triphosphate      &nbspID:DB\_ID:PA164743471** | | | | | | |
| --- | --- | --- | --- | --- | --- | --- |
| C=284; O=10; E=1.51; R=6.61; rawP=2.86e-06; adjP=8.90e-06 | | | | | | |
| Index | UserID | Value | Gene Symbol | Gene Name | EntrezGene | Ensembl |
| 1 | Q99798 | NA | ACO2 | aconitase 2, mitochondrial | 50 | ENSG00000100412 |
| 2 | P12235 | NA | SLC25A4 | solute carrier family 25 (mitochondrial carrier; adenine nucleotide translocator), member 4 | 291 | ENSG00000151729 |
| 3 | P00558 | NA | PGK1 | phosphoglycerate kinase 1 | 5230 | ENSG00000102144 |
| 4 | O95831 | NA | AIFM1 | apoptosis-inducing factor, mitochondrion-associated, 1 | 9131 | ENSG00000156709 |
| 5 | P00367 | NA | GLUD1 | glutamate dehydrogenase 1 | 2746 | ENSG00000148672 |
| 6 | P60842 | NA | EIF4A1 | eukaryotic translation initiation factor 4A1 | 1973 | ENSG00000161960 |
| 7 | P19367 | NA | HK1 | hexokinase 1 | 3098 | ENSG00000156515 |
| 8 | P25705 | NA | ATP5A1 | ATP synthase, H+ transporting, mitochondrial F1 complex, alpha subunit 1, cardiac muscle | 498 | ENSG00000152234 |
| 9 | P21796 | NA | VDAC1 | voltage-dependent anion channel 1 | 7416 | ENSG00000213585 |
| 10 | O60313 | NA | OPA1 | optic atrophy 1 (autosomal dominant) | 4976 | ENSG00000198836 |

  
  

| **Database:drug      &nbspName:clonazepam      &nbspID:DB\_ID:PA449050** | | | | | | |
| --- | --- | --- | --- | --- | --- | --- |
| C=27; O=4; E=0.14; R=27.82; rawP=1.21e-05; adjP=3.39e-05 | | | | | | |
| Index | UserID | Value | Gene Symbol | Gene Name | EntrezGene | Ensembl |
| 1 | P02792 | NA | FTL | ferritin, light polypeptide | 2512 | ENSG00000087086 |
| 2 | P04080 | NA | CSTB | cystatin B (stefin B) | 1476 | ENSG00000160213 |
| 3 | P07108 | NA | DBI | diazepam binding inhibitor (GABA receptor modulator, acyl-CoA binding protein) | 1622 | ENSG00000155368 |
| 4 | Q9UJZ1 | NA | STOML2 | stomatin (EPB72)-like 2 | 30968 | ENSG00000165283 |

  
  

| **Database:drug      &nbspName:betamethasone      &nbspID:DB\_ID:PA164754818** | | | | | | |
| --- | --- | --- | --- | --- | --- | --- |
| C=10; O=3; E=0.05; R=56.34; rawP=1.71e-05; adjP=4.35e-05 | | | | | | |
| Index | UserID | Value | Gene Symbol | Gene Name | EntrezGene | Ensembl |
| 1 | P04040 | NA | CAT | catalase | 847 | ENSG00000121691 |
| 2 | P04083 | NA | ANXA1 | annexin A1 | 301 | ENSG00000135046 |
| 3 | P09382 | NA | LGALS1 | lectin, galactoside-binding, soluble, 1 | 3956 | ENSG00000100097 |

  
  

| **Database:drug      &nbspName:miglustat      &nbspID:DB\_ID:PA10140** | | | | | | |
| --- | --- | --- | --- | --- | --- | --- |
| C=13; O=3; E=0.07; R=43.34; rawP=4.04e-05; adjP=9.43e-05 | | | | | | |
| Index | UserID | Value | Gene Symbol | Gene Name | EntrezGene | Ensembl |
| 1 | P10253 | NA | GAA | glucosidase, alpha; acid | 2548 | ENSG00000171298 |
| 2 | Q14697 | NA | GANAB | glucosidase, alpha; neutral AB | 23193 | ENSG00000089597 |
| 3 | Q13724 | NA | MOGS | mannosyl-oligosaccharide glucosidase | 7841 | ENSG00000115275 |

  
  

| **Database:drug      &nbspName:creatine      &nbspID:DB\_ID:PA164778930** | | | | | | |
| --- | --- | --- | --- | --- | --- | --- |
| C=89; O=5; E=0.47; R=10.55; rawP=0.0001; adjP=0.0002 | | | | | | |
| Index | UserID | Value | Gene Symbol | Gene Name | EntrezGene | Ensembl |
| 1 | P17174 | NA | GOT1 | glutamic-oxaloacetic transaminase 1, soluble (aspartate aminotransferase 1) | 2805 | ENSG00000120053 |
| 2 | P21796 | NA | VDAC1 | voltage-dependent anion channel 1 | 7416 | ENSG00000213585 |
| 3 | P49748 | NA | ACADVL | acyl-CoA dehydrogenase, very long chain | 37 | ENSG00000072778 |
| 4 | P60174 | NA | TPI1 | triosephosphate isomerase 1 | 7167 | ENSG00000111669 |
| 5 | P31937 | NA | HIBADH | 3-hydroxyisobutyrate dehydrogenase | 11112 | ENSG00000106049 |

  
  
  
  

---

WebGestalt is currently developed and maintained by Jing Wang and Bing Zhang at the  Zhang Lab. Other people who have made significant contribution to the project include Dexter Duncan, Stefan Kirov, Zhiao Shi, and Jay Snoddy.  
  
**Funding credits:** NIH/NIAAA (U01 AA016662, U01 AA013512); NIH/NIDA (P01 DA015027); NIH/NIMH (P50 MH078028, P50 MH096972); NIH/NCI (U24 CA159988); NIH/NIGMS (R01 GM088822).
