## Supplementary material for "PGRMC1 phosphorylation and cell plasticity 1: glycolysis, mitochondria, tumor growth": File S7: final_sig_file_1438480126.html

Anchored HTML File of EIDs

|  |  |
| --- | --- |
|  | WEB-based GEne SeT AnaLysis Toolkit |
|  | ***Translating gene lists into biological insights...*** |

---

  

| **Database:biological process      &nbspName:small molecule metabolic process      &nbspID:GO:0044281** | | | | | | |
| --- | --- | --- | --- | --- | --- | --- |
| C=2500; O=52; E=18.03; R=2.88; rawP=2.01e-14; adjP=1.33e-11 | | | | | | |
| Index | UserID | Value | Gene Symbol | Gene Name | EntrezGene | Ensembl |
| 1 | P49589 | NA | CARS | cysteinyl-tRNA synthetase | 833 | ENSG00000110619 |
| 2 | P54819 | NA | AK2 | adenylate kinase 2 | 204 | ENSG00000004455 |
| 3 | Q9UL46 | NA | PSME2 | proteasome (prosome, macropain) activator subunit 2 (PA28 beta) | 5721 | ENSG00000100911 |
| 4 | P40926 | NA | MDH2 | malate dehydrogenase 2, NAD (mitochondrial) | 4191 | ENSG00000146701 |
| 5 | Q99714 | NA | HSD17B10 | hydroxysteroid (17-beta) dehydrogenase 10 | 3028 | ENSG00000072506 |
| 6 | P04179 | NA | SOD2 | superoxide dismutase 2, mitochondrial | 6648 | ENSG00000112096 |
| 7 | P07741 | NA | APRT | adenine phosphoribosyltransferase | 353 | ENSG00000198931 |
| 8 | P00367 | NA | GLUD1 | glutamate dehydrogenase 1 | 2746 | ENSG00000148672 |
| 9 | Q06323 | NA | PSME1 | proteasome (prosome, macropain) activator subunit 1 (PA28 alpha) | 5720 | ENSG00000092010 |
| 10 | P48735 | NA | IDH2 | isocitrate dehydrogenase 2 (NADP+), mitochondrial | 3418 | ENSG00000182054 |
| 11 | O60313 | NA | OPA1 | optic atrophy 1 (autosomal dominant) | 4976 | ENSG00000198836 |
| 12 | P35270 | NA | SPR | sepiapterin reductase (7,8-dihydrobiopterin:NADP+ oxidoreductase) | 6697 | ENSG00000116096 |
| 13 | P31040 | NA | SDHA | succinate dehydrogenase complex, subunit A, flavoprotein (Fp) | 6389 | ENSG00000073578 |
| 14 | P38117 | NA | ETFB | electron-transfer-flavoprotein, beta polypeptide | 2109 | ENSG00000105379 |
| 15 | P43490 | NA | NAMPT | nicotinamide phosphoribosyltransferase | 10135 | ENSG00000105835 |
| 16 | P22102 | NA | GART | phosphoribosylglycinamide formyltransferase, phosphoribosylglycinamide synthetase, phosphoribosylaminoimidazole synthetase | 2618 | ENSG00000159131 |
| 17 | P47895 | NA | ALDH1A3 | aldehyde dehydrogenase 1 family, member A3 | 220 | ENSG00000184254 |
| 18 | P13804 | NA | ETFA | electron-transfer-flavoprotein, alpha polypeptide | 2108 | ENSG00000140374 |
| 19 | P42765 | NA | ACAA2 | acetyl-CoA acyltransferase 2 | 10449 | ENSG00000167315 |
| 20 | O75521 | NA | ECI2 | enoyl-CoA delta isomerase 2 | 10455 | ENSG00000198721 |
| 21 | P49419 | NA | ALDH7A1 | aldehyde dehydrogenase 7 family, member A1 | 501 | ENSG00000164904 |
| 22 | P17174 | NA | GOT1 | glutamic-oxaloacetic transaminase 1, soluble (aspartate aminotransferase 1) | 2805 | ENSG00000120053 |
| 23 | P00491 | NA | PNP | purine nucleoside phosphorylase | 4860 | ENSG00000198805 |
| 24 | P21953 | NA | BCKDHB | branched chain keto acid dehydrogenase E1, beta polypeptide | 594 | ENSG00000083123 |
| 25 | P40939 | NA | HADHA | hydroxyacyl-CoA dehydrogenase/3-ketoacyl-CoA thiolase/enoyl-CoA hydratase (trifunctional protein), alpha subunit | 3030 | ENSG00000084754 |
| 26 | P30084 | NA | ECHS1 | enoyl CoA hydratase, short chain, 1, mitochondrial | 1892 | ENSG00000127884 |
| 27 | P38571 | NA | LIPA | lipase A, lysosomal acid, cholesterol esterase | 3988 | ENSG00000107798 |
| 28 | Q13011 | NA | ECH1 | enoyl CoA hydratase 1, peroxisomal | 1891 | ENSG00000104823 |
| 29 | P04083 | NA | ANXA1 | annexin A1 | 301 | ENSG00000135046 |
| 30 | P32322 | NA | PYCR1 | pyrroline-5-carboxylate reductase 1 | 5831 | ENSG00000183010 |
| 31 | P04040 | NA | CAT | catalase | 847 | ENSG00000121691 |
| 32 | Q99798 | NA | ACO2 | aconitase 2, mitochondrial | 50 | ENSG00000100412 |
| 33 | P51688 | NA | SGSH | N-sulfoglucosamine sulfohydrolase | 6448 | ENSG00000181523 |
| 34 | P51649 | NA | ALDH5A1 | aldehyde dehydrogenase 5 family, member A1 | 7915 | ENSG00000112294 |
| 35 | P12235 | NA | SLC25A4 | solute carrier family 25 (mitochondrial carrier; adenine nucleotide translocator), member 4 | 291 | ENSG00000151729 |
| 36 | P00558 | NA | PGK1 | phosphoglycerate kinase 1 | 5230 | ENSG00000102144 |
| 37 | P41091 | NA | EIF2S3 | eukaryotic translation initiation factor 2, subunit 3 gamma, 52kDa | 1968 | ENSG00000130741 |
| 38 | P02768 | NA | ALB | albumin | 213 | ENSG00000163631 |
| 39 | Q9NSE4 | NA | IARS2 | isoleucyl-tRNA synthetase 2, mitochondrial | 55699 | ENSG00000067704 |
| 40 | Q6PI48 | NA | DARS2 | aspartyl-tRNA synthetase 2, mitochondrial | 55157 | ENSG00000117593 |
| 41 | P24752 | NA | ACAT1 | acetyl-CoA acetyltransferase 1 | 38 | ENSG00000075239 |
| 42 | P13674 | NA | P4HA1 | prolyl 4-hydroxylase, alpha polypeptide I | 5033 | ENSG00000122884 |
| 43 | Q02978 | NA | SLC25A11 | solute carrier family 25 (mitochondrial carrier; oxoglutarate carrier), member 11 | 8402 | ENSG00000108528 |
| 44 | P19367 | NA | HK1 | hexokinase 1 | 3098 | ENSG00000156515 |
| 45 | P25705 | NA | ATP5A1 | ATP synthase, H+ transporting, mitochondrial F1 complex, alpha subunit 1, cardiac muscle | 498 | ENSG00000152234 |
| 46 | Q9BQ69 | NA | MACROD1 | MACRO domain containing 1 | 28992 | ENSG00000133315 |
| 47 | P06865 | NA | HEXA | hexosaminidase A (alpha polypeptide) | 3073 | ENSG00000213614 |
| 48 | P49748 | NA | ACADVL | acyl-CoA dehydrogenase, very long chain | 37 | ENSG00000072778 |
| 49 | P00352 | NA | ALDH1A1 | aldehyde dehydrogenase 1 family, member A1 | 216 | ENSG00000165092 |
| 50 | P48047 | NA | ATP5O | ATP synthase, H+ transporting, mitochondrial F1 complex, O subunit | 539 | ENSG00000241837 |
| 51 | P60174 | NA | TPI1 | triosephosphate isomerase 1 | 7167 | ENSG00000111669 |
| 52 | P31937 | NA | HIBADH | 3-hydroxyisobutyrate dehydrogenase | 11112 | ENSG00000106049 |

  
  

| **Database:biological process      &nbspName:organic acid metabolic process      &nbspID:GO:0006082** | | | | | | |
| --- | --- | --- | --- | --- | --- | --- |
| C=967; O=32; E=6.97; R=4.59; rawP=1.01e-13; adjP=2.43e-11 | | | | | | |
| Index | UserID | Value | Gene Symbol | Gene Name | EntrezGene | Ensembl |
| 1 | P49589 | NA | CARS | cysteinyl-tRNA synthetase | 833 | ENSG00000110619 |
| 2 | P00491 | NA | PNP | purine nucleoside phosphorylase | 4860 | ENSG00000198805 |
| 3 | P21953 | NA | BCKDHB | branched chain keto acid dehydrogenase E1, beta polypeptide | 594 | ENSG00000083123 |
| 4 | P40939 | NA | HADHA | hydroxyacyl-CoA dehydrogenase/3-ketoacyl-CoA thiolase/enoyl-CoA hydratase (trifunctional protein), alpha subunit | 3030 | ENSG00000084754 |
| 5 | Q9UL46 | NA | PSME2 | proteasome (prosome, macropain) activator subunit 2 (PA28 beta) | 5721 | ENSG00000100911 |
| 6 | P40926 | NA | MDH2 | malate dehydrogenase 2, NAD (mitochondrial) | 4191 | ENSG00000146701 |
| 7 | P30084 | NA | ECHS1 | enoyl CoA hydratase, short chain, 1, mitochondrial | 1892 | ENSG00000127884 |
| 8 | Q99714 | NA | HSD17B10 | hydroxysteroid (17-beta) dehydrogenase 10 | 3028 | ENSG00000072506 |
| 9 | P04179 | NA | SOD2 | superoxide dismutase 2, mitochondrial | 6648 | ENSG00000112096 |
| 10 | P00367 | NA | GLUD1 | glutamate dehydrogenase 1 | 2746 | ENSG00000148672 |
| 11 | Q06323 | NA | PSME1 | proteasome (prosome, macropain) activator subunit 1 (PA28 alpha) | 5720 | ENSG00000092010 |
| 12 | P48735 | NA | IDH2 | isocitrate dehydrogenase 2 (NADP+), mitochondrial | 3418 | ENSG00000182054 |
| 13 | P38571 | NA | LIPA | lipase A, lysosomal acid, cholesterol esterase | 3988 | ENSG00000107798 |
| 14 | P31040 | NA | SDHA | succinate dehydrogenase complex, subunit A, flavoprotein (Fp) | 6389 | ENSG00000073578 |
| 15 | Q13011 | NA | ECH1 | enoyl CoA hydratase 1, peroxisomal | 1891 | ENSG00000104823 |
| 16 | P04083 | NA | ANXA1 | annexin A1 | 301 | ENSG00000135046 |
| 17 | P32322 | NA | PYCR1 | pyrroline-5-carboxylate reductase 1 | 5831 | ENSG00000183010 |
| 18 | Q99798 | NA | ACO2 | aconitase 2, mitochondrial | 50 | ENSG00000100412 |
| 19 | P51649 | NA | ALDH5A1 | aldehyde dehydrogenase 5 family, member A1 | 7915 | ENSG00000112294 |
| 20 | P02768 | NA | ALB | albumin | 213 | ENSG00000163631 |
| 21 | P47895 | NA | ALDH1A3 | aldehyde dehydrogenase 1 family, member A3 | 220 | ENSG00000184254 |
| 22 | Q6PI48 | NA | DARS2 | aspartyl-tRNA synthetase 2, mitochondrial | 55157 | ENSG00000117593 |
| 23 | P42765 | NA | ACAA2 | acetyl-CoA acyltransferase 2 | 10449 | ENSG00000167315 |
| 24 | Q9NSE4 | NA | IARS2 | isoleucyl-tRNA synthetase 2, mitochondrial | 55699 | ENSG00000067704 |
| 25 | P24752 | NA | ACAT1 | acetyl-CoA acetyltransferase 1 | 38 | ENSG00000075239 |
| 26 | O75521 | NA | ECI2 | enoyl-CoA delta isomerase 2 | 10455 | ENSG00000198721 |
| 27 | P13674 | NA | P4HA1 | prolyl 4-hydroxylase, alpha polypeptide I | 5033 | ENSG00000122884 |
| 28 | P49419 | NA | ALDH7A1 | aldehyde dehydrogenase 7 family, member A1 | 501 | ENSG00000164904 |
| 29 | P17174 | NA | GOT1 | glutamic-oxaloacetic transaminase 1, soluble (aspartate aminotransferase 1) | 2805 | ENSG00000120053 |
| 30 | P06865 | NA | HEXA | hexosaminidase A (alpha polypeptide) | 3073 | ENSG00000213614 |
| 31 | P49748 | NA | ACADVL | acyl-CoA dehydrogenase, very long chain | 37 | ENSG00000072778 |
| 32 | P31937 | NA | HIBADH | 3-hydroxyisobutyrate dehydrogenase | 11112 | ENSG00000106049 |

  
  

| **Database:biological process      &nbspName:carboxylic acid metabolic process      &nbspID:GO:0019752** | | | | | | |
| --- | --- | --- | --- | --- | --- | --- |
| C=842; O=30; E=6.07; R=4.94; rawP=1.10e-13; adjP=2.43e-11 | | | | | | |
| Index | UserID | Value | Gene Symbol | Gene Name | EntrezGene | Ensembl |
| 1 | P49589 | NA | CARS | cysteinyl-tRNA synthetase | 833 | ENSG00000110619 |
| 2 | P21953 | NA | BCKDHB | branched chain keto acid dehydrogenase E1, beta polypeptide | 594 | ENSG00000083123 |
| 3 | P40939 | NA | HADHA | hydroxyacyl-CoA dehydrogenase/3-ketoacyl-CoA thiolase/enoyl-CoA hydratase (trifunctional protein), alpha subunit | 3030 | ENSG00000084754 |
| 4 | Q9UL46 | NA | PSME2 | proteasome (prosome, macropain) activator subunit 2 (PA28 beta) | 5721 | ENSG00000100911 |
| 5 | P40926 | NA | MDH2 | malate dehydrogenase 2, NAD (mitochondrial) | 4191 | ENSG00000146701 |
| 6 | P30084 | NA | ECHS1 | enoyl CoA hydratase, short chain, 1, mitochondrial | 1892 | ENSG00000127884 |
| 7 | Q99714 | NA | HSD17B10 | hydroxysteroid (17-beta) dehydrogenase 10 | 3028 | ENSG00000072506 |
| 8 | P04179 | NA | SOD2 | superoxide dismutase 2, mitochondrial | 6648 | ENSG00000112096 |
| 9 | P00367 | NA | GLUD1 | glutamate dehydrogenase 1 | 2746 | ENSG00000148672 |
| 10 | Q06323 | NA | PSME1 | proteasome (prosome, macropain) activator subunit 1 (PA28 alpha) | 5720 | ENSG00000092010 |
| 11 | P48735 | NA | IDH2 | isocitrate dehydrogenase 2 (NADP+), mitochondrial | 3418 | ENSG00000182054 |
| 12 | P38571 | NA | LIPA | lipase A, lysosomal acid, cholesterol esterase | 3988 | ENSG00000107798 |
| 13 | P31040 | NA | SDHA | succinate dehydrogenase complex, subunit A, flavoprotein (Fp) | 6389 | ENSG00000073578 |
| 14 | Q13011 | NA | ECH1 | enoyl CoA hydratase 1, peroxisomal | 1891 | ENSG00000104823 |
| 15 | P04083 | NA | ANXA1 | annexin A1 | 301 | ENSG00000135046 |
| 16 | P32322 | NA | PYCR1 | pyrroline-5-carboxylate reductase 1 | 5831 | ENSG00000183010 |
| 17 | Q99798 | NA | ACO2 | aconitase 2, mitochondrial | 50 | ENSG00000100412 |
| 18 | P51649 | NA | ALDH5A1 | aldehyde dehydrogenase 5 family, member A1 | 7915 | ENSG00000112294 |
| 19 | P02768 | NA | ALB | albumin | 213 | ENSG00000163631 |
| 20 | P47895 | NA | ALDH1A3 | aldehyde dehydrogenase 1 family, member A3 | 220 | ENSG00000184254 |
| 21 | Q6PI48 | NA | DARS2 | aspartyl-tRNA synthetase 2, mitochondrial | 55157 | ENSG00000117593 |
| 22 | P42765 | NA | ACAA2 | acetyl-CoA acyltransferase 2 | 10449 | ENSG00000167315 |
| 23 | Q9NSE4 | NA | IARS2 | isoleucyl-tRNA synthetase 2, mitochondrial | 55699 | ENSG00000067704 |
| 24 | P24752 | NA | ACAT1 | acetyl-CoA acetyltransferase 1 | 38 | ENSG00000075239 |
| 25 | O75521 | NA | ECI2 | enoyl-CoA delta isomerase 2 | 10455 | ENSG00000198721 |
| 26 | P13674 | NA | P4HA1 | prolyl 4-hydroxylase, alpha polypeptide I | 5033 | ENSG00000122884 |
| 27 | P49419 | NA | ALDH7A1 | aldehyde dehydrogenase 7 family, member A1 | 501 | ENSG00000164904 |
| 28 | P17174 | NA | GOT1 | glutamic-oxaloacetic transaminase 1, soluble (aspartate aminotransferase 1) | 2805 | ENSG00000120053 |
| 29 | P49748 | NA | ACADVL | acyl-CoA dehydrogenase, very long chain | 37 | ENSG00000072778 |
| 30 | P31937 | NA | HIBADH | 3-hydroxyisobutyrate dehydrogenase | 11112 | ENSG00000106049 |

  
  

| **Database:biological process      &nbspName:oxoacid metabolic process      &nbspID:GO:0043436** | | | | | | |
| --- | --- | --- | --- | --- | --- | --- |
| C=949; O=31; E=6.84; R=4.53; rawP=3.89e-13; adjP=6.44e-11 | | | | | | |
| Index | UserID | Value | Gene Symbol | Gene Name | EntrezGene | Ensembl |
| 1 | P49589 | NA | CARS | cysteinyl-tRNA synthetase | 833 | ENSG00000110619 |
| 2 | P21953 | NA | BCKDHB | branched chain keto acid dehydrogenase E1, beta polypeptide | 594 | ENSG00000083123 |
| 3 | P40939 | NA | HADHA | hydroxyacyl-CoA dehydrogenase/3-ketoacyl-CoA thiolase/enoyl-CoA hydratase (trifunctional protein), alpha subunit | 3030 | ENSG00000084754 |
| 4 | Q9UL46 | NA | PSME2 | proteasome (prosome, macropain) activator subunit 2 (PA28 beta) | 5721 | ENSG00000100911 |
| 5 | P40926 | NA | MDH2 | malate dehydrogenase 2, NAD (mitochondrial) | 4191 | ENSG00000146701 |
| 6 | P30084 | NA | ECHS1 | enoyl CoA hydratase, short chain, 1, mitochondrial | 1892 | ENSG00000127884 |
| 7 | Q99714 | NA | HSD17B10 | hydroxysteroid (17-beta) dehydrogenase 10 | 3028 | ENSG00000072506 |
| 8 | P04179 | NA | SOD2 | superoxide dismutase 2, mitochondrial | 6648 | ENSG00000112096 |
| 9 | P00367 | NA | GLUD1 | glutamate dehydrogenase 1 | 2746 | ENSG00000148672 |
| 10 | Q06323 | NA | PSME1 | proteasome (prosome, macropain) activator subunit 1 (PA28 alpha) | 5720 | ENSG00000092010 |
| 11 | P48735 | NA | IDH2 | isocitrate dehydrogenase 2 (NADP+), mitochondrial | 3418 | ENSG00000182054 |
| 12 | P38571 | NA | LIPA | lipase A, lysosomal acid, cholesterol esterase | 3988 | ENSG00000107798 |
| 13 | P31040 | NA | SDHA | succinate dehydrogenase complex, subunit A, flavoprotein (Fp) | 6389 | ENSG00000073578 |
| 14 | Q13011 | NA | ECH1 | enoyl CoA hydratase 1, peroxisomal | 1891 | ENSG00000104823 |
| 15 | P04083 | NA | ANXA1 | annexin A1 | 301 | ENSG00000135046 |
| 16 | P32322 | NA | PYCR1 | pyrroline-5-carboxylate reductase 1 | 5831 | ENSG00000183010 |
| 17 | Q99798 | NA | ACO2 | aconitase 2, mitochondrial | 50 | ENSG00000100412 |
| 18 | P51649 | NA | ALDH5A1 | aldehyde dehydrogenase 5 family, member A1 | 7915 | ENSG00000112294 |
| 19 | P02768 | NA | ALB | albumin | 213 | ENSG00000163631 |
| 20 | P47895 | NA | ALDH1A3 | aldehyde dehydrogenase 1 family, member A3 | 220 | ENSG00000184254 |
| 21 | Q6PI48 | NA | DARS2 | aspartyl-tRNA synthetase 2, mitochondrial | 55157 | ENSG00000117593 |
| 22 | P42765 | NA | ACAA2 | acetyl-CoA acyltransferase 2 | 10449 | ENSG00000167315 |
| 23 | Q9NSE4 | NA | IARS2 | isoleucyl-tRNA synthetase 2, mitochondrial | 55699 | ENSG00000067704 |
| 24 | P24752 | NA | ACAT1 | acetyl-CoA acetyltransferase 1 | 38 | ENSG00000075239 |
| 25 | O75521 | NA | ECI2 | enoyl-CoA delta isomerase 2 | 10455 | ENSG00000198721 |
| 26 | P13674 | NA | P4HA1 | prolyl 4-hydroxylase, alpha polypeptide I | 5033 | ENSG00000122884 |
| 27 | P49419 | NA | ALDH7A1 | aldehyde dehydrogenase 7 family, member A1 | 501 | ENSG00000164904 |
| 28 | P17174 | NA | GOT1 | glutamic-oxaloacetic transaminase 1, soluble (aspartate aminotransferase 1) | 2805 | ENSG00000120053 |
| 29 | P06865 | NA | HEXA | hexosaminidase A (alpha polypeptide) | 3073 | ENSG00000213614 |
| 30 | P49748 | NA | ACADVL | acyl-CoA dehydrogenase, very long chain | 37 | ENSG00000072778 |
| 31 | P31937 | NA | HIBADH | 3-hydroxyisobutyrate dehydrogenase | 11112 | ENSG00000106049 |

  
  

| **Database:biological process      &nbspName:oxidation-reduction process      &nbspID:GO:0055114** | | | | | | |
| --- | --- | --- | --- | --- | --- | --- |
| C=554; O=23; E=4.00; R=5.76; rawP=6.62e-12; adjP=8.76e-10 | | | | | | |
| Index | UserID | Value | Gene Symbol | Gene Name | EntrezGene | Ensembl |
| 1 | P10253 | NA | GAA | glucosidase, alpha; acid | 2548 | ENSG00000171298 |
| 2 | P40939 | NA | HADHA | hydroxyacyl-CoA dehydrogenase/3-ketoacyl-CoA thiolase/enoyl-CoA hydratase (trifunctional protein), alpha subunit | 3030 | ENSG00000084754 |
| 3 | P40926 | NA | MDH2 | malate dehydrogenase 2, NAD (mitochondrial) | 4191 | ENSG00000146701 |
| 4 | P30084 | NA | ECHS1 | enoyl CoA hydratase, short chain, 1, mitochondrial | 1892 | ENSG00000127884 |
| 5 | P04179 | NA | SOD2 | superoxide dismutase 2, mitochondrial | 6648 | ENSG00000112096 |
| 6 | P48735 | NA | IDH2 | isocitrate dehydrogenase 2 (NADP+), mitochondrial | 3418 | ENSG00000182054 |
| 7 | P53701 | NA | HCCS | holocytochrome c synthase | 3052 | ENSG00000004961 |
| 8 | P35270 | NA | SPR | sepiapterin reductase (7,8-dihydrobiopterin:NADP+ oxidoreductase) | 6697 | ENSG00000116096 |
| 9 | P31040 | NA | SDHA | succinate dehydrogenase complex, subunit A, flavoprotein (Fp) | 6389 | ENSG00000073578 |
| 10 | Q13011 | NA | ECH1 | enoyl CoA hydratase 1, peroxisomal | 1891 | ENSG00000104823 |
| 11 | P38117 | NA | ETFB | electron-transfer-flavoprotein, beta polypeptide | 2109 | ENSG00000105379 |
| 12 | P04040 | NA | CAT | catalase | 847 | ENSG00000121691 |
| 13 | Q99798 | NA | ACO2 | aconitase 2, mitochondrial | 50 | ENSG00000100412 |
| 14 | P51649 | NA | ALDH5A1 | aldehyde dehydrogenase 5 family, member A1 | 7915 | ENSG00000112294 |
| 15 | P12235 | NA | SLC25A4 | solute carrier family 25 (mitochondrial carrier; adenine nucleotide translocator), member 4 | 291 | ENSG00000151729 |
| 16 | P13804 | NA | ETFA | electron-transfer-flavoprotein, alpha polypeptide | 2108 | ENSG00000140374 |
| 17 | P42765 | NA | ACAA2 | acetyl-CoA acyltransferase 2 | 10449 | ENSG00000167315 |
| 18 | P25705 | NA | ATP5A1 | ATP synthase, H+ transporting, mitochondrial F1 complex, alpha subunit 1, cardiac muscle | 498 | ENSG00000152234 |
| 19 | P49748 | NA | ACADVL | acyl-CoA dehydrogenase, very long chain | 37 | ENSG00000072778 |
| 20 | P00352 | NA | ALDH1A1 | aldehyde dehydrogenase 1 family, member A1 | 216 | ENSG00000165092 |
| 21 | P48047 | NA | ATP5O | ATP synthase, H+ transporting, mitochondrial F1 complex, O subunit | 539 | ENSG00000241837 |
| 22 | P60174 | NA | TPI1 | triosephosphate isomerase 1 | 7167 | ENSG00000111669 |
| 23 | P31937 | NA | HIBADH | 3-hydroxyisobutyrate dehydrogenase | 11112 | ENSG00000106049 |

  
  

| **Database:biological process      &nbspName:carboxylic acid catabolic process      &nbspID:GO:0046395** | | | | | | |
| --- | --- | --- | --- | --- | --- | --- |
| C=198; O=14; E=1.43; R=9.81; rawP=1.44e-10; adjP=1.21e-08 | | | | | | |
| Index | UserID | Value | Gene Symbol | Gene Name | EntrezGene | Ensembl |
| 1 | P51649 | NA | ALDH5A1 | aldehyde dehydrogenase 5 family, member A1 | 7915 | ENSG00000112294 |
| 2 | P21953 | NA | BCKDHB | branched chain keto acid dehydrogenase E1, beta polypeptide | 594 | ENSG00000083123 |
| 3 | P40939 | NA | HADHA | hydroxyacyl-CoA dehydrogenase/3-ketoacyl-CoA thiolase/enoyl-CoA hydratase (trifunctional protein), alpha subunit | 3030 | ENSG00000084754 |
| 4 | P30084 | NA | ECHS1 | enoyl CoA hydratase, short chain, 1, mitochondrial | 1892 | ENSG00000127884 |
| 5 | Q99714 | NA | HSD17B10 | hydroxysteroid (17-beta) dehydrogenase 10 | 3028 | ENSG00000072506 |
| 6 | P42765 | NA | ACAA2 | acetyl-CoA acyltransferase 2 | 10449 | ENSG00000167315 |
| 7 | P24752 | NA | ACAT1 | acetyl-CoA acetyltransferase 1 | 38 | ENSG00000075239 |
| 8 | O75521 | NA | ECI2 | enoyl-CoA delta isomerase 2 | 10455 | ENSG00000198721 |
| 9 | P00367 | NA | GLUD1 | glutamate dehydrogenase 1 | 2746 | ENSG00000148672 |
| 10 | P17174 | NA | GOT1 | glutamic-oxaloacetic transaminase 1, soluble (aspartate aminotransferase 1) | 2805 | ENSG00000120053 |
| 11 | P49419 | NA | ALDH7A1 | aldehyde dehydrogenase 7 family, member A1 | 501 | ENSG00000164904 |
| 12 | P49748 | NA | ACADVL | acyl-CoA dehydrogenase, very long chain | 37 | ENSG00000072778 |
| 13 | Q13011 | NA | ECH1 | enoyl CoA hydratase 1, peroxisomal | 1891 | ENSG00000104823 |
| 14 | P31937 | NA | HIBADH | 3-hydroxyisobutyrate dehydrogenase | 11112 | ENSG00000106049 |

  
  

| **Database:biological process      &nbspName:organic acid catabolic process      &nbspID:GO:0016054** | | | | | | |
| --- | --- | --- | --- | --- | --- | --- |
| C=198; O=14; E=1.43; R=9.81; rawP=1.44e-10; adjP=1.21e-08 | | | | | | |
| Index | UserID | Value | Gene Symbol | Gene Name | EntrezGene | Ensembl |
| 1 | P51649 | NA | ALDH5A1 | aldehyde dehydrogenase 5 family, member A1 | 7915 | ENSG00000112294 |
| 2 | P21953 | NA | BCKDHB | branched chain keto acid dehydrogenase E1, beta polypeptide | 594 | ENSG00000083123 |
| 3 | P40939 | NA | HADHA | hydroxyacyl-CoA dehydrogenase/3-ketoacyl-CoA thiolase/enoyl-CoA hydratase (trifunctional protein), alpha subunit | 3030 | ENSG00000084754 |
| 4 | P30084 | NA | ECHS1 | enoyl CoA hydratase, short chain, 1, mitochondrial | 1892 | ENSG00000127884 |
| 5 | Q99714 | NA | HSD17B10 | hydroxysteroid (17-beta) dehydrogenase 10 | 3028 | ENSG00000072506 |
| 6 | P42765 | NA | ACAA2 | acetyl-CoA acyltransferase 2 | 10449 | ENSG00000167315 |
| 7 | P24752 | NA | ACAT1 | acetyl-CoA acetyltransferase 1 | 38 | ENSG00000075239 |
| 8 | O75521 | NA | ECI2 | enoyl-CoA delta isomerase 2 | 10455 | ENSG00000198721 |
| 9 | P00367 | NA | GLUD1 | glutamate dehydrogenase 1 | 2746 | ENSG00000148672 |
| 10 | P17174 | NA | GOT1 | glutamic-oxaloacetic transaminase 1, soluble (aspartate aminotransferase 1) | 2805 | ENSG00000120053 |
| 11 | P49419 | NA | ALDH7A1 | aldehyde dehydrogenase 7 family, member A1 | 501 | ENSG00000164904 |
| 12 | P49748 | NA | ACADVL | acyl-CoA dehydrogenase, very long chain | 37 | ENSG00000072778 |
| 13 | Q13011 | NA | ECH1 | enoyl CoA hydratase 1, peroxisomal | 1891 | ENSG00000104823 |
| 14 | P31937 | NA | HIBADH | 3-hydroxyisobutyrate dehydrogenase | 11112 | ENSG00000106049 |

  
  

| **Database:biological process      &nbspName:organic substance catabolic process      &nbspID:GO:1901575** | | | | | | |
| --- | --- | --- | --- | --- | --- | --- |
| C=1851; O=39; E=13.35; R=2.92; rawP=1.46e-10; adjP=1.21e-08 | | | | | | |
| Index | UserID | Value | Gene Symbol | Gene Name | EntrezGene | Ensembl |
| 1 | P10253 | NA | GAA | glucosidase, alpha; acid | 2548 | ENSG00000171298 |
| 2 | Q9UL46 | NA | PSME2 | proteasome (prosome, macropain) activator subunit 2 (PA28 beta) | 5721 | ENSG00000100911 |
| 3 | P40926 | NA | MDH2 | malate dehydrogenase 2, NAD (mitochondrial) | 4191 | ENSG00000146701 |
| 4 | O95831 | NA | AIFM1 | apoptosis-inducing factor, mitochondrion-associated, 1 | 9131 | ENSG00000156709 |
| 5 | Q99714 | NA | HSD17B10 | hydroxysteroid (17-beta) dehydrogenase 10 | 3028 | ENSG00000072506 |
| 6 | P61313 | NA | RPL15 | ribosomal protein L15 | 6138 | ENSG00000174748 |
| 7 | P00367 | NA | GLUD1 | glutamate dehydrogenase 1 | 2746 | ENSG00000148672 |
| 8 | Q06323 | NA | PSME1 | proteasome (prosome, macropain) activator subunit 1 (PA28 alpha) | 5720 | ENSG00000092010 |
| 9 | P48735 | NA | IDH2 | isocitrate dehydrogenase 2 (NADP+), mitochondrial | 3418 | ENSG00000182054 |
| 10 | O60313 | NA | OPA1 | optic atrophy 1 (autosomal dominant) | 4976 | ENSG00000198836 |
| 11 | P31040 | NA | SDHA | succinate dehydrogenase complex, subunit A, flavoprotein (Fp) | 6389 | ENSG00000073578 |
| 12 | O14773 | NA | TPP1 | tripeptidyl peptidase I | 1200 | ENSG00000166340 |
| 13 | P42765 | NA | ACAA2 | acetyl-CoA acyltransferase 2 | 10449 | ENSG00000167315 |
| 14 | O75521 | NA | ECI2 | enoyl-CoA delta isomerase 2 | 10455 | ENSG00000198721 |
| 15 | P49419 | NA | ALDH7A1 | aldehyde dehydrogenase 7 family, member A1 | 501 | ENSG00000164904 |
| 16 | P17174 | NA | GOT1 | glutamic-oxaloacetic transaminase 1, soluble (aspartate aminotransferase 1) | 2805 | ENSG00000120053 |
| 17 | Q6PIU2 | NA | NCEH1 | neutral cholesterol ester hydrolase 1 | 57552 | ENSG00000144959 |
| 18 | P00491 | NA | PNP | purine nucleoside phosphorylase | 4860 | ENSG00000198805 |
| 19 | P21953 | NA | BCKDHB | branched chain keto acid dehydrogenase E1, beta polypeptide | 594 | ENSG00000083123 |
| 20 | P40939 | NA | HADHA | hydroxyacyl-CoA dehydrogenase/3-ketoacyl-CoA thiolase/enoyl-CoA hydratase (trifunctional protein), alpha subunit | 3030 | ENSG00000084754 |
| 21 | P30084 | NA | ECHS1 | enoyl CoA hydratase, short chain, 1, mitochondrial | 1892 | ENSG00000127884 |
| 22 | P60842 | NA | EIF4A1 | eukaryotic translation initiation factor 4A1 | 1973 | ENSG00000161960 |
| 23 | P38571 | NA | LIPA | lipase A, lysosomal acid, cholesterol esterase | 3988 | ENSG00000107798 |
| 24 | Q13011 | NA | ECH1 | enoyl CoA hydratase 1, peroxisomal | 1891 | ENSG00000104823 |
| 25 | P04040 | NA | CAT | catalase | 847 | ENSG00000121691 |
| 26 | Q99798 | NA | ACO2 | aconitase 2, mitochondrial | 50 | ENSG00000100412 |
| 27 | P51649 | NA | ALDH5A1 | aldehyde dehydrogenase 5 family, member A1 | 7915 | ENSG00000112294 |
| 28 | P51688 | NA | SGSH | N-sulfoglucosamine sulfohydrolase | 6448 | ENSG00000181523 |
| 29 | P41091 | NA | EIF2S3 | eukaryotic translation initiation factor 2, subunit 3 gamma, 52kDa | 1968 | ENSG00000130741 |
| 30 | P00558 | NA | PGK1 | phosphoglycerate kinase 1 | 5230 | ENSG00000102144 |
| 31 | P24752 | NA | ACAT1 | acetyl-CoA acetyltransferase 1 | 38 | ENSG00000075239 |
| 32 | P19367 | NA | HK1 | hexokinase 1 | 3098 | ENSG00000156515 |
| 33 | P06865 | NA | HEXA | hexosaminidase A (alpha polypeptide) | 3073 | ENSG00000213614 |
| 34 | P49748 | NA | ACADVL | acyl-CoA dehydrogenase, very long chain | 37 | ENSG00000072778 |
| 35 | P00352 | NA | ALDH1A1 | aldehyde dehydrogenase 1 family, member A1 | 216 | ENSG00000165092 |
| 36 | P48047 | NA | ATP5O | ATP synthase, H+ transporting, mitochondrial F1 complex, O subunit | 539 | ENSG00000241837 |
| 37 | Q16531 | NA | DDB1 | damage-specific DNA binding protein 1, 127kDa | 1642 | ENSG00000167986 |
| 38 | P60174 | NA | TPI1 | triosephosphate isomerase 1 | 7167 | ENSG00000111669 |
| 39 | P31937 | NA | HIBADH | 3-hydroxyisobutyrate dehydrogenase | 11112 | ENSG00000106049 |

  
  

| **Database:biological process      &nbspName:catabolic process      &nbspID:GO:0009056** | | | | | | |
| --- | --- | --- | --- | --- | --- | --- |
| C=1991; O=40; E=14.36; R=2.79; rawP=3.24e-10; adjP=2.38e-08 | | | | | | |
| Index | UserID | Value | Gene Symbol | Gene Name | EntrezGene | Ensembl |
| 1 | P10253 | NA | GAA | glucosidase, alpha; acid | 2548 | ENSG00000171298 |
| 2 | Q9UL46 | NA | PSME2 | proteasome (prosome, macropain) activator subunit 2 (PA28 beta) | 5721 | ENSG00000100911 |
| 3 | P40926 | NA | MDH2 | malate dehydrogenase 2, NAD (mitochondrial) | 4191 | ENSG00000146701 |
| 4 | O95831 | NA | AIFM1 | apoptosis-inducing factor, mitochondrion-associated, 1 | 9131 | ENSG00000156709 |
| 5 | Q99714 | NA | HSD17B10 | hydroxysteroid (17-beta) dehydrogenase 10 | 3028 | ENSG00000072506 |
| 6 | P61313 | NA | RPL15 | ribosomal protein L15 | 6138 | ENSG00000174748 |
| 7 | P00367 | NA | GLUD1 | glutamate dehydrogenase 1 | 2746 | ENSG00000148672 |
| 8 | Q06323 | NA | PSME1 | proteasome (prosome, macropain) activator subunit 1 (PA28 alpha) | 5720 | ENSG00000092010 |
| 9 | P48735 | NA | IDH2 | isocitrate dehydrogenase 2 (NADP+), mitochondrial | 3418 | ENSG00000182054 |
| 10 | O60313 | NA | OPA1 | optic atrophy 1 (autosomal dominant) | 4976 | ENSG00000198836 |
| 11 | P31040 | NA | SDHA | succinate dehydrogenase complex, subunit A, flavoprotein (Fp) | 6389 | ENSG00000073578 |
| 12 | O14773 | NA | TPP1 | tripeptidyl peptidase I | 1200 | ENSG00000166340 |
| 13 | P42765 | NA | ACAA2 | acetyl-CoA acyltransferase 2 | 10449 | ENSG00000167315 |
| 14 | O75521 | NA | ECI2 | enoyl-CoA delta isomerase 2 | 10455 | ENSG00000198721 |
| 15 | P49419 | NA | ALDH7A1 | aldehyde dehydrogenase 7 family, member A1 | 501 | ENSG00000164904 |
| 16 | P17174 | NA | GOT1 | glutamic-oxaloacetic transaminase 1, soluble (aspartate aminotransferase 1) | 2805 | ENSG00000120053 |
| 17 | Q6PIU2 | NA | NCEH1 | neutral cholesterol ester hydrolase 1 | 57552 | ENSG00000144959 |
| 18 | P00491 | NA | PNP | purine nucleoside phosphorylase | 4860 | ENSG00000198805 |
| 19 | P21953 | NA | BCKDHB | branched chain keto acid dehydrogenase E1, beta polypeptide | 594 | ENSG00000083123 |
| 20 | P40939 | NA | HADHA | hydroxyacyl-CoA dehydrogenase/3-ketoacyl-CoA thiolase/enoyl-CoA hydratase (trifunctional protein), alpha subunit | 3030 | ENSG00000084754 |
| 21 | P30084 | NA | ECHS1 | enoyl CoA hydratase, short chain, 1, mitochondrial | 1892 | ENSG00000127884 |
| 22 | P60842 | NA | EIF4A1 | eukaryotic translation initiation factor 4A1 | 1973 | ENSG00000161960 |
| 23 | P38571 | NA | LIPA | lipase A, lysosomal acid, cholesterol esterase | 3988 | ENSG00000107798 |
| 24 | Q13011 | NA | ECH1 | enoyl CoA hydratase 1, peroxisomal | 1891 | ENSG00000104823 |
| 25 | P04040 | NA | CAT | catalase | 847 | ENSG00000121691 |
| 26 | Q99798 | NA | ACO2 | aconitase 2, mitochondrial | 50 | ENSG00000100412 |
| 27 | P51649 | NA | ALDH5A1 | aldehyde dehydrogenase 5 family, member A1 | 7915 | ENSG00000112294 |
| 28 | P51688 | NA | SGSH | N-sulfoglucosamine sulfohydrolase | 6448 | ENSG00000181523 |
| 29 | P41091 | NA | EIF2S3 | eukaryotic translation initiation factor 2, subunit 3 gamma, 52kDa | 1968 | ENSG00000130741 |
| 30 | P30048 | NA | PRDX3 | peroxiredoxin 3 | 10935 | ENSG00000165672 |
| 31 | P00558 | NA | PGK1 | phosphoglycerate kinase 1 | 5230 | ENSG00000102144 |
| 32 | P24752 | NA | ACAT1 | acetyl-CoA acetyltransferase 1 | 38 | ENSG00000075239 |
| 33 | P19367 | NA | HK1 | hexokinase 1 | 3098 | ENSG00000156515 |
| 34 | P06865 | NA | HEXA | hexosaminidase A (alpha polypeptide) | 3073 | ENSG00000213614 |
| 35 | P49748 | NA | ACADVL | acyl-CoA dehydrogenase, very long chain | 37 | ENSG00000072778 |
| 36 | P00352 | NA | ALDH1A1 | aldehyde dehydrogenase 1 family, member A1 | 216 | ENSG00000165092 |
| 37 | P48047 | NA | ATP5O | ATP synthase, H+ transporting, mitochondrial F1 complex, O subunit | 539 | ENSG00000241837 |
| 38 | Q16531 | NA | DDB1 | damage-specific DNA binding protein 1, 127kDa | 1642 | ENSG00000167986 |
| 39 | P60174 | NA | TPI1 | triosephosphate isomerase 1 | 7167 | ENSG00000111669 |
| 40 | P31937 | NA | HIBADH | 3-hydroxyisobutyrate dehydrogenase | 11112 | ENSG00000106049 |

  
  

| **Database:biological process      &nbspName:small molecule catabolic process      &nbspID:GO:0044282** | | | | | | |
| --- | --- | --- | --- | --- | --- | --- |
| C=258; O=14; E=1.86; R=7.52; rawP=4.58e-09; adjP=2.90e-07 | | | | | | |
| Index | UserID | Value | Gene Symbol | Gene Name | EntrezGene | Ensembl |
| 1 | P51649 | NA | ALDH5A1 | aldehyde dehydrogenase 5 family, member A1 | 7915 | ENSG00000112294 |
| 2 | P21953 | NA | BCKDHB | branched chain keto acid dehydrogenase E1, beta polypeptide | 594 | ENSG00000083123 |
| 3 | P40939 | NA | HADHA | hydroxyacyl-CoA dehydrogenase/3-ketoacyl-CoA thiolase/enoyl-CoA hydratase (trifunctional protein), alpha subunit | 3030 | ENSG00000084754 |
| 4 | P30084 | NA | ECHS1 | enoyl CoA hydratase, short chain, 1, mitochondrial | 1892 | ENSG00000127884 |
| 5 | Q99714 | NA | HSD17B10 | hydroxysteroid (17-beta) dehydrogenase 10 | 3028 | ENSG00000072506 |
| 6 | P42765 | NA | ACAA2 | acetyl-CoA acyltransferase 2 | 10449 | ENSG00000167315 |
| 7 | P24752 | NA | ACAT1 | acetyl-CoA acetyltransferase 1 | 38 | ENSG00000075239 |
| 8 | O75521 | NA | ECI2 | enoyl-CoA delta isomerase 2 | 10455 | ENSG00000198721 |
| 9 | P00367 | NA | GLUD1 | glutamate dehydrogenase 1 | 2746 | ENSG00000148672 |
| 10 | P17174 | NA | GOT1 | glutamic-oxaloacetic transaminase 1, soluble (aspartate aminotransferase 1) | 2805 | ENSG00000120053 |
| 11 | P49419 | NA | ALDH7A1 | aldehyde dehydrogenase 7 family, member A1 | 501 | ENSG00000164904 |
| 12 | P49748 | NA | ACADVL | acyl-CoA dehydrogenase, very long chain | 37 | ENSG00000072778 |
| 13 | Q13011 | NA | ECH1 | enoyl CoA hydratase 1, peroxisomal | 1891 | ENSG00000104823 |
| 14 | P31937 | NA | HIBADH | 3-hydroxyisobutyrate dehydrogenase | 11112 | ENSG00000106049 |

  
  

| **Database:biological process      &nbspName:single-organism catabolic process      &nbspID:GO:0044712** | | | | | | |
| --- | --- | --- | --- | --- | --- | --- |
| C=259; O=14; E=1.87; R=7.50; rawP=4.82e-09; adjP=2.90e-07 | | | | | | |
| Index | UserID | Value | Gene Symbol | Gene Name | EntrezGene | Ensembl |
| 1 | P51649 | NA | ALDH5A1 | aldehyde dehydrogenase 5 family, member A1 | 7915 | ENSG00000112294 |
| 2 | P21953 | NA | BCKDHB | branched chain keto acid dehydrogenase E1, beta polypeptide | 594 | ENSG00000083123 |
| 3 | P40939 | NA | HADHA | hydroxyacyl-CoA dehydrogenase/3-ketoacyl-CoA thiolase/enoyl-CoA hydratase (trifunctional protein), alpha subunit | 3030 | ENSG00000084754 |
| 4 | P30084 | NA | ECHS1 | enoyl CoA hydratase, short chain, 1, mitochondrial | 1892 | ENSG00000127884 |
| 5 | Q99714 | NA | HSD17B10 | hydroxysteroid (17-beta) dehydrogenase 10 | 3028 | ENSG00000072506 |
| 6 | P42765 | NA | ACAA2 | acetyl-CoA acyltransferase 2 | 10449 | ENSG00000167315 |
| 7 | P24752 | NA | ACAT1 | acetyl-CoA acetyltransferase 1 | 38 | ENSG00000075239 |
| 8 | O75521 | NA | ECI2 | enoyl-CoA delta isomerase 2 | 10455 | ENSG00000198721 |
| 9 | P00367 | NA | GLUD1 | glutamate dehydrogenase 1 | 2746 | ENSG00000148672 |
| 10 | P17174 | NA | GOT1 | glutamic-oxaloacetic transaminase 1, soluble (aspartate aminotransferase 1) | 2805 | ENSG00000120053 |
| 11 | P49419 | NA | ALDH7A1 | aldehyde dehydrogenase 7 family, member A1 | 501 | ENSG00000164904 |
| 12 | P49748 | NA | ACADVL | acyl-CoA dehydrogenase, very long chain | 37 | ENSG00000072778 |
| 13 | Q13011 | NA | ECH1 | enoyl CoA hydratase 1, peroxisomal | 1891 | ENSG00000104823 |
| 14 | P31937 | NA | HIBADH | 3-hydroxyisobutyrate dehydrogenase | 11112 | ENSG00000106049 |

  
  

| **Database:biological process      &nbspName:cellular respiration      &nbspID:GO:0045333** | | | | | | |
| --- | --- | --- | --- | --- | --- | --- |
| C=159; O=11; E=1.15; R=9.59; rawP=1.91e-08; adjP=9.73e-07 | | | | | | |
| Index | UserID | Value | Gene Symbol | Gene Name | EntrezGene | Ensembl |
| 1 | P04040 | NA | CAT | catalase | 847 | ENSG00000121691 |
| 2 | Q99798 | NA | ACO2 | aconitase 2, mitochondrial | 50 | ENSG00000100412 |
| 3 | P51649 | NA | ALDH5A1 | aldehyde dehydrogenase 5 family, member A1 | 7915 | ENSG00000112294 |
| 4 | P40926 | NA | MDH2 | malate dehydrogenase 2, NAD (mitochondrial) | 4191 | ENSG00000146701 |
| 5 | P13804 | NA | ETFA | electron-transfer-flavoprotein, alpha polypeptide | 2108 | ENSG00000140374 |
| 6 | P04179 | NA | SOD2 | superoxide dismutase 2, mitochondrial | 6648 | ENSG00000112096 |
| 7 | P48735 | NA | IDH2 | isocitrate dehydrogenase 2 (NADP+), mitochondrial | 3418 | ENSG00000182054 |
| 8 | P25705 | NA | ATP5A1 | ATP synthase, H+ transporting, mitochondrial F1 complex, alpha subunit 1, cardiac muscle | 498 | ENSG00000152234 |
| 9 | P48047 | NA | ATP5O | ATP synthase, H+ transporting, mitochondrial F1 complex, O subunit | 539 | ENSG00000241837 |
| 10 | P31040 | NA | SDHA | succinate dehydrogenase complex, subunit A, flavoprotein (Fp) | 6389 | ENSG00000073578 |
| 11 | P38117 | NA | ETFB | electron-transfer-flavoprotein, beta polypeptide | 2109 | ENSG00000105379 |

  
  

| **Database:biological process      &nbspName:generation of precursor metabolites and energy      &nbspID:GO:0006091** | | | | | | |
| --- | --- | --- | --- | --- | --- | --- |
| C=447; O=17; E=3.22; R=5.27; rawP=1.91e-08; adjP=9.73e-07 | | | | | | |
| Index | UserID | Value | Gene Symbol | Gene Name | EntrezGene | Ensembl |
| 1 | P10253 | NA | GAA | glucosidase, alpha; acid | 2548 | ENSG00000171298 |
| 2 | P40926 | NA | MDH2 | malate dehydrogenase 2, NAD (mitochondrial) | 4191 | ENSG00000146701 |
| 3 | P04179 | NA | SOD2 | superoxide dismutase 2, mitochondrial | 6648 | ENSG00000112096 |
| 4 | P48735 | NA | IDH2 | isocitrate dehydrogenase 2 (NADP+), mitochondrial | 3418 | ENSG00000182054 |
| 5 | P31040 | NA | SDHA | succinate dehydrogenase complex, subunit A, flavoprotein (Fp) | 6389 | ENSG00000073578 |
| 6 | P38117 | NA | ETFB | electron-transfer-flavoprotein, beta polypeptide | 2109 | ENSG00000105379 |
| 7 | P04040 | NA | CAT | catalase | 847 | ENSG00000121691 |
| 8 | Q99798 | NA | ACO2 | aconitase 2, mitochondrial | 50 | ENSG00000100412 |
| 9 | P51649 | NA | ALDH5A1 | aldehyde dehydrogenase 5 family, member A1 | 7915 | ENSG00000112294 |
| 10 | P12235 | NA | SLC25A4 | solute carrier family 25 (mitochondrial carrier; adenine nucleotide translocator), member 4 | 291 | ENSG00000151729 |
| 11 | P00558 | NA | PGK1 | phosphoglycerate kinase 1 | 5230 | ENSG00000102144 |
| 12 | P13804 | NA | ETFA | electron-transfer-flavoprotein, alpha polypeptide | 2108 | ENSG00000140374 |
| 13 | P25705 | NA | ATP5A1 | ATP synthase, H+ transporting, mitochondrial F1 complex, alpha subunit 1, cardiac muscle | 498 | ENSG00000152234 |
| 14 | P19367 | NA | HK1 | hexokinase 1 | 3098 | ENSG00000156515 |
| 15 | P49748 | NA | ACADVL | acyl-CoA dehydrogenase, very long chain | 37 | ENSG00000072778 |
| 16 | P48047 | NA | ATP5O | ATP synthase, H+ transporting, mitochondrial F1 complex, O subunit | 539 | ENSG00000241837 |
| 17 | P60174 | NA | TPI1 | triosephosphate isomerase 1 | 7167 | ENSG00000111669 |

  
  

| **Database:biological process      &nbspName:cellular catabolic process      &nbspID:GO:0044248** | | | | | | |
| --- | --- | --- | --- | --- | --- | --- |
| C=1665; O=33; E=12.01; R=2.75; rawP=3.11e-08; adjP=1.47e-06 | | | | | | |
| Index | UserID | Value | Gene Symbol | Gene Name | EntrezGene | Ensembl |
| 1 | P00491 | NA | PNP | purine nucleoside phosphorylase | 4860 | ENSG00000198805 |
| 2 | P21953 | NA | BCKDHB | branched chain keto acid dehydrogenase E1, beta polypeptide | 594 | ENSG00000083123 |
| 3 | P40939 | NA | HADHA | hydroxyacyl-CoA dehydrogenase/3-ketoacyl-CoA thiolase/enoyl-CoA hydratase (trifunctional protein), alpha subunit | 3030 | ENSG00000084754 |
| 4 | Q9UL46 | NA | PSME2 | proteasome (prosome, macropain) activator subunit 2 (PA28 beta) | 5721 | ENSG00000100911 |
| 5 | P40926 | NA | MDH2 | malate dehydrogenase 2, NAD (mitochondrial) | 4191 | ENSG00000146701 |
| 6 | P30084 | NA | ECHS1 | enoyl CoA hydratase, short chain, 1, mitochondrial | 1892 | ENSG00000127884 |
| 7 | O95831 | NA | AIFM1 | apoptosis-inducing factor, mitochondrion-associated, 1 | 9131 | ENSG00000156709 |
| 8 | Q99714 | NA | HSD17B10 | hydroxysteroid (17-beta) dehydrogenase 10 | 3028 | ENSG00000072506 |
| 9 | P61313 | NA | RPL15 | ribosomal protein L15 | 6138 | ENSG00000174748 |
| 10 | P60842 | NA | EIF4A1 | eukaryotic translation initiation factor 4A1 | 1973 | ENSG00000161960 |
| 11 | P00367 | NA | GLUD1 | glutamate dehydrogenase 1 | 2746 | ENSG00000148672 |
| 12 | Q06323 | NA | PSME1 | proteasome (prosome, macropain) activator subunit 1 (PA28 alpha) | 5720 | ENSG00000092010 |
| 13 | P48735 | NA | IDH2 | isocitrate dehydrogenase 2 (NADP+), mitochondrial | 3418 | ENSG00000182054 |
| 14 | O60313 | NA | OPA1 | optic atrophy 1 (autosomal dominant) | 4976 | ENSG00000198836 |
| 15 | P31040 | NA | SDHA | succinate dehydrogenase complex, subunit A, flavoprotein (Fp) | 6389 | ENSG00000073578 |
| 16 | Q13011 | NA | ECH1 | enoyl CoA hydratase 1, peroxisomal | 1891 | ENSG00000104823 |
| 17 | P04040 | NA | CAT | catalase | 847 | ENSG00000121691 |
| 18 | O14773 | NA | TPP1 | tripeptidyl peptidase I | 1200 | ENSG00000166340 |
| 19 | Q99798 | NA | ACO2 | aconitase 2, mitochondrial | 50 | ENSG00000100412 |
| 20 | P51649 | NA | ALDH5A1 | aldehyde dehydrogenase 5 family, member A1 | 7915 | ENSG00000112294 |
| 21 | P30048 | NA | PRDX3 | peroxiredoxin 3 | 10935 | ENSG00000165672 |
| 22 | P41091 | NA | EIF2S3 | eukaryotic translation initiation factor 2, subunit 3 gamma, 52kDa | 1968 | ENSG00000130741 |
| 23 | P42765 | NA | ACAA2 | acetyl-CoA acyltransferase 2 | 10449 | ENSG00000167315 |
| 24 | P24752 | NA | ACAT1 | acetyl-CoA acetyltransferase 1 | 38 | ENSG00000075239 |
| 25 | O75521 | NA | ECI2 | enoyl-CoA delta isomerase 2 | 10455 | ENSG00000198721 |
| 26 | P49419 | NA | ALDH7A1 | aldehyde dehydrogenase 7 family, member A1 | 501 | ENSG00000164904 |
| 27 | P17174 | NA | GOT1 | glutamic-oxaloacetic transaminase 1, soluble (aspartate aminotransferase 1) | 2805 | ENSG00000120053 |
| 28 | P06865 | NA | HEXA | hexosaminidase A (alpha polypeptide) | 3073 | ENSG00000213614 |
| 29 | P49748 | NA | ACADVL | acyl-CoA dehydrogenase, very long chain | 37 | ENSG00000072778 |
| 30 | P00352 | NA | ALDH1A1 | aldehyde dehydrogenase 1 family, member A1 | 216 | ENSG00000165092 |
| 31 | P48047 | NA | ATP5O | ATP synthase, H+ transporting, mitochondrial F1 complex, O subunit | 539 | ENSG00000241837 |
| 32 | Q16531 | NA | DDB1 | damage-specific DNA binding protein 1, 127kDa | 1642 | ENSG00000167986 |
| 33 | P31937 | NA | HIBADH | 3-hydroxyisobutyrate dehydrogenase | 11112 | ENSG00000106049 |

  
  

| **Database:biological process      &nbspName:organonitrogen compound metabolic process      &nbspID:GO:1901564** | | | | | | |
| --- | --- | --- | --- | --- | --- | --- |
| C=1673; O=33; E=12.06; R=2.74; rawP=3.49e-08; adjP=1.54e-06 | | | | | | |
| Index | UserID | Value | Gene Symbol | Gene Name | EntrezGene | Ensembl |
| 1 | P49589 | NA | CARS | cysteinyl-tRNA synthetase | 833 | ENSG00000110619 |
| 2 | P00491 | NA | PNP | purine nucleoside phosphorylase | 4860 | ENSG00000198805 |
| 3 | P21953 | NA | BCKDHB | branched chain keto acid dehydrogenase E1, beta polypeptide | 594 | ENSG00000083123 |
| 4 | Q9UL46 | NA | PSME2 | proteasome (prosome, macropain) activator subunit 2 (PA28 beta) | 5721 | ENSG00000100911 |
| 5 | P40926 | NA | MDH2 | malate dehydrogenase 2, NAD (mitochondrial) | 4191 | ENSG00000146701 |
| 6 | Q99714 | NA | HSD17B10 | hydroxysteroid (17-beta) dehydrogenase 10 | 3028 | ENSG00000072506 |
| 7 | P04179 | NA | SOD2 | superoxide dismutase 2, mitochondrial | 6648 | ENSG00000112096 |
| 8 | P07741 | NA | APRT | adenine phosphoribosyltransferase | 353 | ENSG00000198931 |
| 9 | P00367 | NA | GLUD1 | glutamate dehydrogenase 1 | 2746 | ENSG00000148672 |
| 10 | Q06323 | NA | PSME1 | proteasome (prosome, macropain) activator subunit 1 (PA28 alpha) | 5720 | ENSG00000092010 |
| 11 | O60313 | NA | OPA1 | optic atrophy 1 (autosomal dominant) | 4976 | ENSG00000198836 |
| 12 | P35270 | NA | SPR | sepiapterin reductase (7,8-dihydrobiopterin:NADP+ oxidoreductase) | 6697 | ENSG00000116096 |
| 13 | P43490 | NA | NAMPT | nicotinamide phosphoribosyltransferase | 10135 | ENSG00000105835 |
| 14 | P32322 | NA | PYCR1 | pyrroline-5-carboxylate reductase 1 | 5831 | ENSG00000183010 |
| 15 | P04040 | NA | CAT | catalase | 847 | ENSG00000121691 |
| 16 | O14773 | NA | TPP1 | tripeptidyl peptidase I | 1200 | ENSG00000166340 |
| 17 | P51688 | NA | SGSH | N-sulfoglucosamine sulfohydrolase | 6448 | ENSG00000181523 |
| 18 | P22102 | NA | GART | phosphoribosylglycinamide formyltransferase, phosphoribosylglycinamide synthetase, phosphoribosylaminoimidazole synthetase | 2618 | ENSG00000159131 |
| 19 | P51649 | NA | ALDH5A1 | aldehyde dehydrogenase 5 family, member A1 | 7915 | ENSG00000112294 |
| 20 | P41091 | NA | EIF2S3 | eukaryotic translation initiation factor 2, subunit 3 gamma, 52kDa | 1968 | ENSG00000130741 |
| 21 | Q9NSE4 | NA | IARS2 | isoleucyl-tRNA synthetase 2, mitochondrial | 55699 | ENSG00000067704 |
| 22 | Q6PI48 | NA | DARS2 | aspartyl-tRNA synthetase 2, mitochondrial | 55157 | ENSG00000117593 |
| 23 | P24752 | NA | ACAT1 | acetyl-CoA acetyltransferase 1 | 38 | ENSG00000075239 |
| 24 | P13674 | NA | P4HA1 | prolyl 4-hydroxylase, alpha polypeptide I | 5033 | ENSG00000122884 |
| 25 | P49419 | NA | ALDH7A1 | aldehyde dehydrogenase 7 family, member A1 | 501 | ENSG00000164904 |
| 26 | P17174 | NA | GOT1 | glutamic-oxaloacetic transaminase 1, soluble (aspartate aminotransferase 1) | 2805 | ENSG00000120053 |
| 27 | P25705 | NA | ATP5A1 | ATP synthase, H+ transporting, mitochondrial F1 complex, alpha subunit 1, cardiac muscle | 498 | ENSG00000152234 |
| 28 | Q9BQ69 | NA | MACROD1 | MACRO domain containing 1 | 28992 | ENSG00000133315 |
| 29 | P06865 | NA | HEXA | hexosaminidase A (alpha polypeptide) | 3073 | ENSG00000213614 |
| 30 | P00352 | NA | ALDH1A1 | aldehyde dehydrogenase 1 family, member A1 | 216 | ENSG00000165092 |
| 31 | P48047 | NA | ATP5O | ATP synthase, H+ transporting, mitochondrial F1 complex, O subunit | 539 | ENSG00000241837 |
| 32 | P60174 | NA | TPI1 | triosephosphate isomerase 1 | 7167 | ENSG00000111669 |
| 33 | P31937 | NA | HIBADH | 3-hydroxyisobutyrate dehydrogenase | 11112 | ENSG00000106049 |

  
  

| **Database:biological process      &nbspName:energy derivation by oxidation of organic compounds      &nbspID:GO:0015980** | | | | | | |
| --- | --- | --- | --- | --- | --- | --- |
| C=326; O=14; E=2.35; R=5.96; rawP=8.75e-08; adjP=3.62e-06 | | | | | | |
| Index | UserID | Value | Gene Symbol | Gene Name | EntrezGene | Ensembl |
| 1 | P04040 | NA | CAT | catalase | 847 | ENSG00000121691 |
| 2 | Q99798 | NA | ACO2 | aconitase 2, mitochondrial | 50 | ENSG00000100412 |
| 3 | P10253 | NA | GAA | glucosidase, alpha; acid | 2548 | ENSG00000171298 |
| 4 | P51649 | NA | ALDH5A1 | aldehyde dehydrogenase 5 family, member A1 | 7915 | ENSG00000112294 |
| 5 | P12235 | NA | SLC25A4 | solute carrier family 25 (mitochondrial carrier; adenine nucleotide translocator), member 4 | 291 | ENSG00000151729 |
| 6 | P40926 | NA | MDH2 | malate dehydrogenase 2, NAD (mitochondrial) | 4191 | ENSG00000146701 |
| 7 | P13804 | NA | ETFA | electron-transfer-flavoprotein, alpha polypeptide | 2108 | ENSG00000140374 |
| 8 | P04179 | NA | SOD2 | superoxide dismutase 2, mitochondrial | 6648 | ENSG00000112096 |
| 9 | P48735 | NA | IDH2 | isocitrate dehydrogenase 2 (NADP+), mitochondrial | 3418 | ENSG00000182054 |
| 10 | P25705 | NA | ATP5A1 | ATP synthase, H+ transporting, mitochondrial F1 complex, alpha subunit 1, cardiac muscle | 498 | ENSG00000152234 |
| 11 | P49748 | NA | ACADVL | acyl-CoA dehydrogenase, very long chain | 37 | ENSG00000072778 |
| 12 | P48047 | NA | ATP5O | ATP synthase, H+ transporting, mitochondrial F1 complex, O subunit | 539 | ENSG00000241837 |
| 13 | P31040 | NA | SDHA | succinate dehydrogenase complex, subunit A, flavoprotein (Fp) | 6389 | ENSG00000073578 |
| 14 | P38117 | NA | ETFB | electron-transfer-flavoprotein, beta polypeptide | 2109 | ENSG00000105379 |

  
  

| **Database:biological process      &nbspName:cellular amino acid metabolic process      &nbspID:GO:0006520** | | | | | | |
| --- | --- | --- | --- | --- | --- | --- |
| C=340; O=14; E=2.45; R=5.71; rawP=1.47e-07; adjP=5.72e-06 | | | | | | |
| Index | UserID | Value | Gene Symbol | Gene Name | EntrezGene | Ensembl |
| 1 | P49589 | NA | CARS | cysteinyl-tRNA synthetase | 833 | ENSG00000110619 |
| 2 | P51649 | NA | ALDH5A1 | aldehyde dehydrogenase 5 family, member A1 | 7915 | ENSG00000112294 |
| 3 | P21953 | NA | BCKDHB | branched chain keto acid dehydrogenase E1, beta polypeptide | 594 | ENSG00000083123 |
| 4 | Q9UL46 | NA | PSME2 | proteasome (prosome, macropain) activator subunit 2 (PA28 beta) | 5721 | ENSG00000100911 |
| 5 | Q99714 | NA | HSD17B10 | hydroxysteroid (17-beta) dehydrogenase 10 | 3028 | ENSG00000072506 |
| 6 | Q6PI48 | NA | DARS2 | aspartyl-tRNA synthetase 2, mitochondrial | 55157 | ENSG00000117593 |
| 7 | Q9NSE4 | NA | IARS2 | isoleucyl-tRNA synthetase 2, mitochondrial | 55699 | ENSG00000067704 |
| 8 | P24752 | NA | ACAT1 | acetyl-CoA acetyltransferase 1 | 38 | ENSG00000075239 |
| 9 | P49419 | NA | ALDH7A1 | aldehyde dehydrogenase 7 family, member A1 | 501 | ENSG00000164904 |
| 10 | P00367 | NA | GLUD1 | glutamate dehydrogenase 1 | 2746 | ENSG00000148672 |
| 11 | Q06323 | NA | PSME1 | proteasome (prosome, macropain) activator subunit 1 (PA28 alpha) | 5720 | ENSG00000092010 |
| 12 | P17174 | NA | GOT1 | glutamic-oxaloacetic transaminase 1, soluble (aspartate aminotransferase 1) | 2805 | ENSG00000120053 |
| 13 | P31937 | NA | HIBADH | 3-hydroxyisobutyrate dehydrogenase | 11112 | ENSG00000106049 |
| 14 | P32322 | NA | PYCR1 | pyrroline-5-carboxylate reductase 1 | 5831 | ENSG00000183010 |

  
  

| **Database:biological process      &nbspName:cellular amino acid catabolic process      &nbspID:GO:0009063** | | | | | | |
| --- | --- | --- | --- | --- | --- | --- |
| C=104; O=8; E=0.75; R=10.67; rawP=8.10e-07; adjP=2.98e-05 | | | | | | |
| Index | UserID | Value | Gene Symbol | Gene Name | EntrezGene | Ensembl |
| 1 | P51649 | NA | ALDH5A1 | aldehyde dehydrogenase 5 family, member A1 | 7915 | ENSG00000112294 |
| 2 | P21953 | NA | BCKDHB | branched chain keto acid dehydrogenase E1, beta polypeptide | 594 | ENSG00000083123 |
| 3 | Q99714 | NA | HSD17B10 | hydroxysteroid (17-beta) dehydrogenase 10 | 3028 | ENSG00000072506 |
| 4 | P24752 | NA | ACAT1 | acetyl-CoA acetyltransferase 1 | 38 | ENSG00000075239 |
| 5 | P00367 | NA | GLUD1 | glutamate dehydrogenase 1 | 2746 | ENSG00000148672 |
| 6 | P17174 | NA | GOT1 | glutamic-oxaloacetic transaminase 1, soluble (aspartate aminotransferase 1) | 2805 | ENSG00000120053 |
| 7 | P49419 | NA | ALDH7A1 | aldehyde dehydrogenase 7 family, member A1 | 501 | ENSG00000164904 |
| 8 | P31937 | NA | HIBADH | 3-hydroxyisobutyrate dehydrogenase | 11112 | ENSG00000106049 |

  
  

| **Database:biological process      &nbspName:fatty acid catabolic process      &nbspID:GO:0009062** | | | | | | |
| --- | --- | --- | --- | --- | --- | --- |
| C=75; O=7; E=0.54; R=12.94; rawP=1.10e-06; adjP=3.83e-05 | | | | | | |
| Index | UserID | Value | Gene Symbol | Gene Name | EntrezGene | Ensembl |
| 1 | O75521 | NA | ECI2 | enoyl-CoA delta isomerase 2 | 10455 | ENSG00000198721 |
| 2 | P51649 | NA | ALDH5A1 | aldehyde dehydrogenase 5 family, member A1 | 7915 | ENSG00000112294 |
| 3 | P49748 | NA | ACADVL | acyl-CoA dehydrogenase, very long chain | 37 | ENSG00000072778 |
| 4 | P40939 | NA | HADHA | hydroxyacyl-CoA dehydrogenase/3-ketoacyl-CoA thiolase/enoyl-CoA hydratase (trifunctional protein), alpha subunit | 3030 | ENSG00000084754 |
| 5 | Q13011 | NA | ECH1 | enoyl CoA hydratase 1, peroxisomal | 1891 | ENSG00000104823 |
| 6 | P30084 | NA | ECHS1 | enoyl CoA hydratase, short chain, 1, mitochondrial | 1892 | ENSG00000127884 |
| 7 | P42765 | NA | ACAA2 | acetyl-CoA acyltransferase 2 | 10449 | ENSG00000167315 |

  
  

| **Database:biological process      &nbspName:monocarboxylic acid metabolic process      &nbspID:GO:0032787** | | | | | | |
| --- | --- | --- | --- | --- | --- | --- |
| C=405; O=14; E=2.92; R=4.79; rawP=1.21e-06; adjP=4.01e-05 | | | | | | |
| Index | UserID | Value | Gene Symbol | Gene Name | EntrezGene | Ensembl |
| 1 | P51649 | NA | ALDH5A1 | aldehyde dehydrogenase 5 family, member A1 | 7915 | ENSG00000112294 |
| 2 | P02768 | NA | ALB | albumin | 213 | ENSG00000163631 |
| 3 | P40939 | NA | HADHA | hydroxyacyl-CoA dehydrogenase/3-ketoacyl-CoA thiolase/enoyl-CoA hydratase (trifunctional protein), alpha subunit | 3030 | ENSG00000084754 |
| 4 | P47895 | NA | ALDH1A3 | aldehyde dehydrogenase 1 family, member A3 | 220 | ENSG00000184254 |
| 5 | P30084 | NA | ECHS1 | enoyl CoA hydratase, short chain, 1, mitochondrial | 1892 | ENSG00000127884 |
| 6 | P42765 | NA | ACAA2 | acetyl-CoA acyltransferase 2 | 10449 | ENSG00000167315 |
| 7 | P13674 | NA | P4HA1 | prolyl 4-hydroxylase, alpha polypeptide I | 5033 | ENSG00000122884 |
| 8 | O75521 | NA | ECI2 | enoyl-CoA delta isomerase 2 | 10455 | ENSG00000198721 |
| 9 | P48735 | NA | IDH2 | isocitrate dehydrogenase 2 (NADP+), mitochondrial | 3418 | ENSG00000182054 |
| 10 | P17174 | NA | GOT1 | glutamic-oxaloacetic transaminase 1, soluble (aspartate aminotransferase 1) | 2805 | ENSG00000120053 |
| 11 | P49748 | NA | ACADVL | acyl-CoA dehydrogenase, very long chain | 37 | ENSG00000072778 |
| 12 | P38571 | NA | LIPA | lipase A, lysosomal acid, cholesterol esterase | 3988 | ENSG00000107798 |
| 13 | Q13011 | NA | ECH1 | enoyl CoA hydratase 1, peroxisomal | 1891 | ENSG00000104823 |
| 14 | P04083 | NA | ANXA1 | annexin A1 | 301 | ENSG00000135046 |

  
  

| **Database:biological process      &nbspName:coenzyme metabolic process      &nbspID:GO:0006732** | | | | | | |
| --- | --- | --- | --- | --- | --- | --- |
| C=213; O=10; E=1.54; R=6.51; rawP=3.12e-06; adjP=9.84e-05 | | | | | | |
| Index | UserID | Value | Gene Symbol | Gene Name | EntrezGene | Ensembl |
| 1 | Q99798 | NA | ACO2 | aconitase 2, mitochondrial | 50 | ENSG00000100412 |
| 2 | P00491 | NA | PNP | purine nucleoside phosphorylase | 4860 | ENSG00000198805 |
| 3 | P40926 | NA | MDH2 | malate dehydrogenase 2, NAD (mitochondrial) | 4191 | ENSG00000146701 |
| 4 | P42765 | NA | ACAA2 | acetyl-CoA acyltransferase 2 | 10449 | ENSG00000167315 |
| 5 | P48735 | NA | IDH2 | isocitrate dehydrogenase 2 (NADP+), mitochondrial | 3418 | ENSG00000182054 |
| 6 | P35270 | NA | SPR | sepiapterin reductase (7,8-dihydrobiopterin:NADP+ oxidoreductase) | 6697 | ENSG00000116096 |
| 7 | P31040 | NA | SDHA | succinate dehydrogenase complex, subunit A, flavoprotein (Fp) | 6389 | ENSG00000073578 |
| 8 | P60174 | NA | TPI1 | triosephosphate isomerase 1 | 7167 | ENSG00000111669 |
| 9 | P31937 | NA | HIBADH | 3-hydroxyisobutyrate dehydrogenase | 11112 | ENSG00000106049 |
| 10 | P43490 | NA | NAMPT | nicotinamide phosphoribosyltransferase | 10135 | ENSG00000105835 |

  
  

| **Database:biological process      &nbspName:monocarboxylic acid catabolic process      &nbspID:GO:0072329** | | | | | | |
| --- | --- | --- | --- | --- | --- | --- |
| C=89; O=7; E=0.64; R=10.91; rawP=3.51e-06; adjP=0.0001 | | | | | | |
| Index | UserID | Value | Gene Symbol | Gene Name | EntrezGene | Ensembl |
| 1 | O75521 | NA | ECI2 | enoyl-CoA delta isomerase 2 | 10455 | ENSG00000198721 |
| 2 | P51649 | NA | ALDH5A1 | aldehyde dehydrogenase 5 family, member A1 | 7915 | ENSG00000112294 |
| 3 | P49748 | NA | ACADVL | acyl-CoA dehydrogenase, very long chain | 37 | ENSG00000072778 |
| 4 | P40939 | NA | HADHA | hydroxyacyl-CoA dehydrogenase/3-ketoacyl-CoA thiolase/enoyl-CoA hydratase (trifunctional protein), alpha subunit | 3030 | ENSG00000084754 |
| 5 | Q13011 | NA | ECH1 | enoyl CoA hydratase 1, peroxisomal | 1891 | ENSG00000104823 |
| 6 | P30084 | NA | ECHS1 | enoyl CoA hydratase, short chain, 1, mitochondrial | 1892 | ENSG00000127884 |
| 7 | P42765 | NA | ACAA2 | acetyl-CoA acyltransferase 2 | 10449 | ENSG00000167315 |

  
  

| **Database:biological process      &nbspName:aerobic respiration      &nbspID:GO:0009060** | | | | | | |
| --- | --- | --- | --- | --- | --- | --- |
| C=44; O=5; E=0.32; R=15.76; rawP=1.54e-05; adjP=0.0003 | | | | | | |
| Index | UserID | Value | Gene Symbol | Gene Name | EntrezGene | Ensembl |
| 1 | P04040 | NA | CAT | catalase | 847 | ENSG00000121691 |
| 2 | Q99798 | NA | ACO2 | aconitase 2, mitochondrial | 50 | ENSG00000100412 |
| 3 | P48735 | NA | IDH2 | isocitrate dehydrogenase 2 (NADP+), mitochondrial | 3418 | ENSG00000182054 |
| 4 | P31040 | NA | SDHA | succinate dehydrogenase complex, subunit A, flavoprotein (Fp) | 6389 | ENSG00000073578 |
| 5 | P40926 | NA | MDH2 | malate dehydrogenase 2, NAD (mitochondrial) | 4191 | ENSG00000146701 |

  
  

| **Database:biological process      &nbspName:cellular lipid catabolic process      &nbspID:GO:0044242** | | | | | | |
| --- | --- | --- | --- | --- | --- | --- |
| C=147; O=8; E=1.06; R=7.55; rawP=1.09e-05; adjP=0.0003 | | | | | | |
| Index | UserID | Value | Gene Symbol | Gene Name | EntrezGene | Ensembl |
| 1 | P51649 | NA | ALDH5A1 | aldehyde dehydrogenase 5 family, member A1 | 7915 | ENSG00000112294 |
| 2 | P40939 | NA | HADHA | hydroxyacyl-CoA dehydrogenase/3-ketoacyl-CoA thiolase/enoyl-CoA hydratase (trifunctional protein), alpha subunit | 3030 | ENSG00000084754 |
| 3 | P30084 | NA | ECHS1 | enoyl CoA hydratase, short chain, 1, mitochondrial | 1892 | ENSG00000127884 |
| 4 | P42765 | NA | ACAA2 | acetyl-CoA acyltransferase 2 | 10449 | ENSG00000167315 |
| 5 | O75521 | NA | ECI2 | enoyl-CoA delta isomerase 2 | 10455 | ENSG00000198721 |
| 6 | P49748 | NA | ACADVL | acyl-CoA dehydrogenase, very long chain | 37 | ENSG00000072778 |
| 7 | P06865 | NA | HEXA | hexosaminidase A (alpha polypeptide) | 3073 | ENSG00000213614 |
| 8 | Q13011 | NA | ECH1 | enoyl CoA hydratase 1, peroxisomal | 1891 | ENSG00000104823 |

  
  

| **Database:biological process      &nbspName:organonitrogen compound catabolic process      &nbspID:GO:1901565** | | | | | | |
| --- | --- | --- | --- | --- | --- | --- |
| C=711; O=17; E=5.13; R=3.32; rawP=1.22e-05; adjP=0.0003 | | | | | | |
| Index | UserID | Value | Gene Symbol | Gene Name | EntrezGene | Ensembl |
| 1 | P04040 | NA | CAT | catalase | 847 | ENSG00000121691 |
| 2 | O14773 | NA | TPP1 | tripeptidyl peptidase I | 1200 | ENSG00000166340 |
| 3 | P51688 | NA | SGSH | N-sulfoglucosamine sulfohydrolase | 6448 | ENSG00000181523 |
| 4 | P51649 | NA | ALDH5A1 | aldehyde dehydrogenase 5 family, member A1 | 7915 | ENSG00000112294 |
| 5 | P00491 | NA | PNP | purine nucleoside phosphorylase | 4860 | ENSG00000198805 |
| 6 | P41091 | NA | EIF2S3 | eukaryotic translation initiation factor 2, subunit 3 gamma, 52kDa | 1968 | ENSG00000130741 |
| 7 | P21953 | NA | BCKDHB | branched chain keto acid dehydrogenase E1, beta polypeptide | 594 | ENSG00000083123 |
| 8 | Q99714 | NA | HSD17B10 | hydroxysteroid (17-beta) dehydrogenase 10 | 3028 | ENSG00000072506 |
| 9 | P24752 | NA | ACAT1 | acetyl-CoA acetyltransferase 1 | 38 | ENSG00000075239 |
| 10 | P00367 | NA | GLUD1 | glutamate dehydrogenase 1 | 2746 | ENSG00000148672 |
| 11 | P17174 | NA | GOT1 | glutamic-oxaloacetic transaminase 1, soluble (aspartate aminotransferase 1) | 2805 | ENSG00000120053 |
| 12 | P49419 | NA | ALDH7A1 | aldehyde dehydrogenase 7 family, member A1 | 501 | ENSG00000164904 |
| 13 | O60313 | NA | OPA1 | optic atrophy 1 (autosomal dominant) | 4976 | ENSG00000198836 |
| 14 | P06865 | NA | HEXA | hexosaminidase A (alpha polypeptide) | 3073 | ENSG00000213614 |
| 15 | P00352 | NA | ALDH1A1 | aldehyde dehydrogenase 1 family, member A1 | 216 | ENSG00000165092 |
| 16 | P48047 | NA | ATP5O | ATP synthase, H+ transporting, mitochondrial F1 complex, O subunit | 539 | ENSG00000241837 |
| 17 | P31937 | NA | HIBADH | 3-hydroxyisobutyrate dehydrogenase | 11112 | ENSG00000106049 |

  
  

| **Database:biological process      &nbspName:branched-chain amino acid catabolic process      &nbspID:GO:0009083** | | | | | | |
| --- | --- | --- | --- | --- | --- | --- |
| C=19; O=4; E=0.14; R=29.19; rawP=9.10e-06; adjP=0.0003 | | | | | | |
| Index | UserID | Value | Gene Symbol | Gene Name | EntrezGene | Ensembl |
| 1 | P24752 | NA | ACAT1 | acetyl-CoA acetyltransferase 1 | 38 | ENSG00000075239 |
| 2 | P21953 | NA | BCKDHB | branched chain keto acid dehydrogenase E1, beta polypeptide | 594 | ENSG00000083123 |
| 3 | P31937 | NA | HIBADH | 3-hydroxyisobutyrate dehydrogenase | 11112 | ENSG00000106049 |
| 4 | Q99714 | NA | HSD17B10 | hydroxysteroid (17-beta) dehydrogenase 10 | 3028 | ENSG00000072506 |

  
  

| **Database:biological process      &nbspName:cofactor metabolic process      &nbspID:GO:0051186** | | | | | | |
| --- | --- | --- | --- | --- | --- | --- |
| C=255; O=10; E=1.84; R=5.44; rawP=1.52e-05; adjP=0.0003 | | | | | | |
| Index | UserID | Value | Gene Symbol | Gene Name | EntrezGene | Ensembl |
| 1 | Q99798 | NA | ACO2 | aconitase 2, mitochondrial | 50 | ENSG00000100412 |
| 2 | P00491 | NA | PNP | purine nucleoside phosphorylase | 4860 | ENSG00000198805 |
| 3 | P40926 | NA | MDH2 | malate dehydrogenase 2, NAD (mitochondrial) | 4191 | ENSG00000146701 |
| 4 | P42765 | NA | ACAA2 | acetyl-CoA acyltransferase 2 | 10449 | ENSG00000167315 |
| 5 | P48735 | NA | IDH2 | isocitrate dehydrogenase 2 (NADP+), mitochondrial | 3418 | ENSG00000182054 |
| 6 | P35270 | NA | SPR | sepiapterin reductase (7,8-dihydrobiopterin:NADP+ oxidoreductase) | 6697 | ENSG00000116096 |
| 7 | P31040 | NA | SDHA | succinate dehydrogenase complex, subunit A, flavoprotein (Fp) | 6389 | ENSG00000073578 |
| 8 | P60174 | NA | TPI1 | triosephosphate isomerase 1 | 7167 | ENSG00000111669 |
| 9 | P31937 | NA | HIBADH | 3-hydroxyisobutyrate dehydrogenase | 11112 | ENSG00000106049 |
| 10 | P43490 | NA | NAMPT | nicotinamide phosphoribosyltransferase | 10135 | ENSG00000105835 |

  
  

| **Database:biological process      &nbspName:respiratory electron transport chain      &nbspID:GO:0022904** | | | | | | |
| --- | --- | --- | --- | --- | --- | --- |
| C=110; O=7; E=0.79; R=8.82; rawP=1.43e-05; adjP=0.0003 | | | | | | |
| Index | UserID | Value | Gene Symbol | Gene Name | EntrezGene | Ensembl |
| 1 | P25705 | NA | ATP5A1 | ATP synthase, H+ transporting, mitochondrial F1 complex, alpha subunit 1, cardiac muscle | 498 | ENSG00000152234 |
| 2 | P51649 | NA | ALDH5A1 | aldehyde dehydrogenase 5 family, member A1 | 7915 | ENSG00000112294 |
| 3 | P48047 | NA | ATP5O | ATP synthase, H+ transporting, mitochondrial F1 complex, O subunit | 539 | ENSG00000241837 |
| 4 | P31040 | NA | SDHA | succinate dehydrogenase complex, subunit A, flavoprotein (Fp) | 6389 | ENSG00000073578 |
| 5 | P38117 | NA | ETFB | electron-transfer-flavoprotein, beta polypeptide | 2109 | ENSG00000105379 |
| 6 | P13804 | NA | ETFA | electron-transfer-flavoprotein, alpha polypeptide | 2108 | ENSG00000140374 |
| 7 | P04179 | NA | SOD2 | superoxide dismutase 2, mitochondrial | 6648 | ENSG00000112096 |

  
  

| **Database:biological process      &nbspName:cellular aldehyde metabolic process      &nbspID:GO:0006081** | | | | | | |
| --- | --- | --- | --- | --- | --- | --- |
| C=40; O=5; E=0.29; R=17.33; rawP=9.53e-06; adjP=0.0003 | | | | | | |
| Index | UserID | Value | Gene Symbol | Gene Name | EntrezGene | Ensembl |
| 1 | P49419 | NA | ALDH7A1 | aldehyde dehydrogenase 7 family, member A1 | 501 | ENSG00000164904 |
| 2 | P48735 | NA | IDH2 | isocitrate dehydrogenase 2 (NADP+), mitochondrial | 3418 | ENSG00000182054 |
| 3 | P00352 | NA | ALDH1A1 | aldehyde dehydrogenase 1 family, member A1 | 216 | ENSG00000165092 |
| 4 | P60174 | NA | TPI1 | triosephosphate isomerase 1 | 7167 | ENSG00000111669 |
| 5 | P47895 | NA | ALDH1A3 | aldehyde dehydrogenase 1 family, member A3 | 220 | ENSG00000184254 |

  
  

| **Database:biological process      &nbspName:lipid catabolic process      &nbspID:GO:0016042** | | | | | | |
| --- | --- | --- | --- | --- | --- | --- |
| C=252; O=10; E=1.82; R=5.50; rawP=1.37e-05; adjP=0.0003 | | | | | | |
| Index | UserID | Value | Gene Symbol | Gene Name | EntrezGene | Ensembl |
| 1 | P51649 | NA | ALDH5A1 | aldehyde dehydrogenase 5 family, member A1 | 7915 | ENSG00000112294 |
| 2 | P40939 | NA | HADHA | hydroxyacyl-CoA dehydrogenase/3-ketoacyl-CoA thiolase/enoyl-CoA hydratase (trifunctional protein), alpha subunit | 3030 | ENSG00000084754 |
| 3 | P30084 | NA | ECHS1 | enoyl CoA hydratase, short chain, 1, mitochondrial | 1892 | ENSG00000127884 |
| 4 | P42765 | NA | ACAA2 | acetyl-CoA acyltransferase 2 | 10449 | ENSG00000167315 |
| 5 | O75521 | NA | ECI2 | enoyl-CoA delta isomerase 2 | 10455 | ENSG00000198721 |
| 6 | Q6PIU2 | NA | NCEH1 | neutral cholesterol ester hydrolase 1 | 57552 | ENSG00000144959 |
| 7 | P06865 | NA | HEXA | hexosaminidase A (alpha polypeptide) | 3073 | ENSG00000213614 |
| 8 | P38571 | NA | LIPA | lipase A, lysosomal acid, cholesterol esterase | 3988 | ENSG00000107798 |
| 9 | P49748 | NA | ACADVL | acyl-CoA dehydrogenase, very long chain | 37 | ENSG00000072778 |
| 10 | Q13011 | NA | ECH1 | enoyl CoA hydratase 1, peroxisomal | 1891 | ENSG00000104823 |

  
  

| **Database:biological process      &nbspName:metabolic process      &nbspID:GO:0008152** | | | | | | |
| --- | --- | --- | --- | --- | --- | --- |
| C=9363; O=87; E=67.52; R=1.29; rawP=1.74e-05; adjP=0.0004 | | | | | | |
| Index | UserID | Value | Gene Symbol | Gene Name | EntrezGene | Ensembl |
| 1 | P10253 | NA | GAA | glucosidase, alpha; acid | 2548 | ENSG00000171298 |
| 2 | P68371 | NA | TUBB4B | tubulin, beta 4B class IVb | 10383 | ENSG00000188229 |
| 3 | Q9UL46 | NA | PSME2 | proteasome (prosome, macropain) activator subunit 2 (PA28 beta) | 5721 | ENSG00000100911 |
| 4 | P40926 | NA | MDH2 | malate dehydrogenase 2, NAD (mitochondrial) | 4191 | ENSG00000146701 |
| 5 | P04179 | NA | SOD2 | superoxide dismutase 2, mitochondrial | 6648 | ENSG00000112096 |
| 6 | Q13724 | NA | MOGS | mannosyl-oligosaccharide glucosidase | 7841 | ENSG00000115275 |
| 7 | P48735 | NA | IDH2 | isocitrate dehydrogenase 2 (NADP+), mitochondrial | 3418 | ENSG00000182054 |
| 8 | P00367 | NA | GLUD1 | glutamate dehydrogenase 1 | 2746 | ENSG00000148672 |
| 9 | Q06323 | NA | PSME1 | proteasome (prosome, macropain) activator subunit 1 (PA28 alpha) | 5720 | ENSG00000092010 |
| 10 | P53701 | NA | HCCS | holocytochrome c synthase | 3052 | ENSG00000004961 |
| 11 | P31040 | NA | SDHA | succinate dehydrogenase complex, subunit A, flavoprotein (Fp) | 6389 | ENSG00000073578 |
| 12 | P24534 | NA | EEF1B2 | eukaryotic translation elongation factor 1 beta 2 | 1933 | ENSG00000114942 |
| 13 | O14773 | NA | TPP1 | tripeptidyl peptidase I | 1200 | ENSG00000166340 |
| 14 | P47895 | NA | ALDH1A3 | aldehyde dehydrogenase 1 family, member A3 | 220 | ENSG00000184254 |
| 15 | P13804 | NA | ETFA | electron-transfer-flavoprotein, alpha polypeptide | 2108 | ENSG00000140374 |
| 16 | P42765 | NA | ACAA2 | acetyl-CoA acyltransferase 2 | 10449 | ENSG00000167315 |
| 17 | P17174 | NA | GOT1 | glutamic-oxaloacetic transaminase 1, soluble (aspartate aminotransferase 1) | 2805 | ENSG00000120053 |
| 18 | Q04837 | NA | SSBP1 | single-stranded DNA binding protein 1, mitochondrial | 6742 | ENSG00000106028 |
| 19 | P50454 | NA | SERPINH1 | serpin peptidase inhibitor, clade H (heat shock protein 47), member 1, (collagen binding protein 1) | 871 | ENSG00000149257 |
| 20 | Q14697 | NA | GANAB | glucosidase, alpha; neutral AB | 23193 | ENSG00000089597 |
| 21 | P08195 | NA | SLC3A2 | solute carrier family 3 (activators of dibasic and neutral amino acid transport), member 2 | 6520 | ENSG00000168003 |
| 22 | P40939 | NA | HADHA | hydroxyacyl-CoA dehydrogenase/3-ketoacyl-CoA thiolase/enoyl-CoA hydratase (trifunctional protein), alpha subunit | 3030 | ENSG00000084754 |
| 23 | P52815 | NA | MRPL12 | mitochondrial ribosomal protein L12 | 6182 | ENSG00000262814 |
| 24 | P04080 | NA | CSTB | cystatin B (stefin B) | 1476 | ENSG00000160213 |
| 25 | P38571 | NA | LIPA | lipase A, lysosomal acid, cholesterol esterase | 3988 | ENSG00000107798 |
| 26 | P32322 | NA | PYCR1 | pyrroline-5-carboxylate reductase 1 | 5831 | ENSG00000183010 |
| 27 | Q99798 | NA | ACO2 | aconitase 2, mitochondrial | 50 | ENSG00000100412 |
| 28 | P51649 | NA | ALDH5A1 | aldehyde dehydrogenase 5 family, member A1 | 7915 | ENSG00000112294 |
| 29 | P12235 | NA | SLC25A4 | solute carrier family 25 (mitochondrial carrier; adenine nucleotide translocator), member 4 | 291 | ENSG00000151729 |
| 30 | P05556 | NA | ITGB1 | integrin, beta 1 (fibronectin receptor, beta polypeptide, antigen CD29 includes MDF2, MSK12) | 3688 | ENSG00000150093 |
| 31 | Q9NSE4 | NA | IARS2 | isoleucyl-tRNA synthetase 2, mitochondrial | 55699 | ENSG00000067704 |
| 32 | Q6PI48 | NA | DARS2 | aspartyl-tRNA synthetase 2, mitochondrial | 55157 | ENSG00000117593 |
| 33 | P13674 | NA | P4HA1 | prolyl 4-hydroxylase, alpha polypeptide I | 5033 | ENSG00000122884 |
| 34 | P40227 | NA | CCT6A | chaperonin containing TCP1, subunit 6A (zeta 1) | 908 | ENSG00000146731 |
| 35 | Q02978 | NA | SLC25A11 | solute carrier family 25 (mitochondrial carrier; oxoglutarate carrier), member 11 | 8402 | ENSG00000108528 |
| 36 | P60174 | NA | TPI1 | triosephosphate isomerase 1 | 7167 | ENSG00000111669 |
| 37 | P49589 | NA | CARS | cysteinyl-tRNA synthetase | 833 | ENSG00000110619 |
| 38 | P54819 | NA | AK2 | adenylate kinase 2 | 204 | ENSG00000004455 |
| 39 | P17931 | NA | LGALS3 | lectin, galactoside-binding, soluble, 3 | 3958 | ENSG00000131981 |
| 40 | Q99623 | NA | PHB2 | prohibitin 2 | 11331 | ENSG00000215021 |
| 41 | P50453 | NA | SERPINB9 | serpin peptidase inhibitor, clade B (ovalbumin), member 9 | 5272 | ENSG00000170542 |
| 42 | P35232 | NA | PHB | prohibitin | 5245 | ENSG00000167085 |
| 43 | Q99714 | NA | HSD17B10 | hydroxysteroid (17-beta) dehydrogenase 10 | 3028 | ENSG00000072506 |
| 44 | O95831 | NA | AIFM1 | apoptosis-inducing factor, mitochondrion-associated, 1 | 9131 | ENSG00000156709 |
| 45 | P61313 | NA | RPL15 | ribosomal protein L15 | 6138 | ENSG00000174748 |
| 46 | P07741 | NA | APRT | adenine phosphoribosyltransferase | 353 | ENSG00000198931 |
| 47 | O60313 | NA | OPA1 | optic atrophy 1 (autosomal dominant) | 4976 | ENSG00000198836 |
| 48 | P35270 | NA | SPR | sepiapterin reductase (7,8-dihydrobiopterin:NADP+ oxidoreductase) | 6697 | ENSG00000116096 |
| 49 | P68104 | NA | EEF1A1 | eukaryotic translation elongation factor 1 alpha 1 | 1915 | ENSG00000156508 |
| 50 | P02786 | NA | TFRC | transferrin receptor (p90, CD71) | 7037 | ENSG00000072274 |
| 51 | P38117 | NA | ETFB | electron-transfer-flavoprotein, beta polypeptide | 2109 | ENSG00000105379 |
| 52 | P43490 | NA | NAMPT | nicotinamide phosphoribosyltransferase | 10135 | ENSG00000105835 |
| 53 | P22102 | NA | GART | phosphoribosylglycinamide formyltransferase, phosphoribosylglycinamide synthetase, phosphoribosylaminoimidazole synthetase | 2618 | ENSG00000159131 |
| 54 | P13639 | NA | EEF2 | eukaryotic translation elongation factor 2 | 1938 | ENSG00000167658 |
| 55 | O75367 | NA | H2AFY | H2A histone family, member Y | 9555 | ENSG00000113648 |
| 56 | O75521 | NA | ECI2 | enoyl-CoA delta isomerase 2 | 10455 | ENSG00000198721 |
| 57 | P67809 | NA | YBX1 | Y box binding protein 1 | 4904 | ENSG00000065978 |
| 58 | P49419 | NA | ALDH7A1 | aldehyde dehydrogenase 7 family, member A1 | 501 | ENSG00000164904 |
| 59 | Q6PIU2 | NA | NCEH1 | neutral cholesterol ester hydrolase 1 | 57552 | ENSG00000144959 |
| 60 | P29692 | NA | EEF1D | eukaryotic translation elongation factor 1 delta (guanine nucleotide exchange protein) | 1936 | ENSG00000104529 |
| 61 | P11387 | NA | TOP1 | topoisomerase (DNA) I | 7150 | ENSG00000198900 |
| 62 | Q5SSJ5 | NA | HP1BP3 | heterochromatin protein 1, binding protein 3 | 50809 | ENSG00000127483 |
| 63 | O75439 | NA | PMPCB | peptidase (mitochondrial processing) beta | 9512 | ENSG00000105819 |
| 64 | P00491 | NA | PNP | purine nucleoside phosphorylase | 4860 | ENSG00000198805 |
| 65 | P21953 | NA | BCKDHB | branched chain keto acid dehydrogenase E1, beta polypeptide | 594 | ENSG00000083123 |
| 66 | P30084 | NA | ECHS1 | enoyl CoA hydratase, short chain, 1, mitochondrial | 1892 | ENSG00000127884 |
| 67 | P60842 | NA | EIF4A1 | eukaryotic translation initiation factor 4A1 | 1973 | ENSG00000161960 |
| 68 | Q13011 | NA | ECH1 | enoyl CoA hydratase 1, peroxisomal | 1891 | ENSG00000104823 |
| 69 | P04083 | NA | ANXA1 | annexin A1 | 301 | ENSG00000135046 |
| 70 | P04040 | NA | CAT | catalase | 847 | ENSG00000121691 |
| 71 | O75340 | NA | PDCD6 | programmed cell death 6 | 10016 | ENSG00000249915 |
| 72 | P51688 | NA | SGSH | N-sulfoglucosamine sulfohydrolase | 6448 | ENSG00000181523 |
| 73 | P00558 | NA | PGK1 | phosphoglycerate kinase 1 | 5230 | ENSG00000102144 |
| 74 | P30048 | NA | PRDX3 | peroxiredoxin 3 | 10935 | ENSG00000165672 |
| 75 | P41091 | NA | EIF2S3 | eukaryotic translation initiation factor 2, subunit 3 gamma, 52kDa | 1968 | ENSG00000130741 |
| 76 | P02768 | NA | ALB | albumin | 213 | ENSG00000163631 |
| 77 | P07108 | NA | DBI | diazepam binding inhibitor (GABA receptor modulator, acyl-CoA binding protein) | 1622 | ENSG00000155368 |
| 78 | P24752 | NA | ACAT1 | acetyl-CoA acetyltransferase 1 | 38 | ENSG00000075239 |
| 79 | P19367 | NA | HK1 | hexokinase 1 | 3098 | ENSG00000156515 |
| 80 | P25705 | NA | ATP5A1 | ATP synthase, H+ transporting, mitochondrial F1 complex, alpha subunit 1, cardiac muscle | 498 | ENSG00000152234 |
| 81 | Q9BQ69 | NA | MACROD1 | MACRO domain containing 1 | 28992 | ENSG00000133315 |
| 82 | P06865 | NA | HEXA | hexosaminidase A (alpha polypeptide) | 3073 | ENSG00000213614 |
| 83 | P49748 | NA | ACADVL | acyl-CoA dehydrogenase, very long chain | 37 | ENSG00000072778 |
| 84 | P00352 | NA | ALDH1A1 | aldehyde dehydrogenase 1 family, member A1 | 216 | ENSG00000165092 |
| 85 | P48047 | NA | ATP5O | ATP synthase, H+ transporting, mitochondrial F1 complex, O subunit | 539 | ENSG00000241837 |
| 86 | Q16531 | NA | DDB1 | damage-specific DNA binding protein 1, 127kDa | 1642 | ENSG00000167986 |
| 87 | P31937 | NA | HIBADH | 3-hydroxyisobutyrate dehydrogenase | 11112 | ENSG00000106049 |

  
  

| **Database:biological process      &nbspName:branched-chain amino acid metabolic process      &nbspID:GO:0009081** | | | | | | |
| --- | --- | --- | --- | --- | --- | --- |
| C=23; O=4; E=0.17; R=24.12; rawP=2.03e-05; adjP=0.0004 | | | | | | |
| Index | UserID | Value | Gene Symbol | Gene Name | EntrezGene | Ensembl |
| 1 | P24752 | NA | ACAT1 | acetyl-CoA acetyltransferase 1 | 38 | ENSG00000075239 |
| 2 | P21953 | NA | BCKDHB | branched chain keto acid dehydrogenase E1, beta polypeptide | 594 | ENSG00000083123 |
| 3 | P31937 | NA | HIBADH | 3-hydroxyisobutyrate dehydrogenase | 11112 | ENSG00000106049 |
| 4 | Q99714 | NA | HSD17B10 | hydroxysteroid (17-beta) dehydrogenase 10 | 3028 | ENSG00000072506 |

  
  

| **Database:biological process      &nbspName:nicotinamide nucleotide metabolic process      &nbspID:GO:0046496** | | | | | | |
| --- | --- | --- | --- | --- | --- | --- |
| C=49; O=5; E=0.35; R=14.15; rawP=2.62e-05; adjP=0.0005 | | | | | | |
| Index | UserID | Value | Gene Symbol | Gene Name | EntrezGene | Ensembl |
| 1 | P00491 | NA | PNP | purine nucleoside phosphorylase | 4860 | ENSG00000198805 |
| 2 | P40926 | NA | MDH2 | malate dehydrogenase 2, NAD (mitochondrial) | 4191 | ENSG00000146701 |
| 3 | P60174 | NA | TPI1 | triosephosphate isomerase 1 | 7167 | ENSG00000111669 |
| 4 | P31937 | NA | HIBADH | 3-hydroxyisobutyrate dehydrogenase | 11112 | ENSG00000106049 |
| 5 | P43490 | NA | NAMPT | nicotinamide phosphoribosyltransferase | 10135 | ENSG00000105835 |

  
  

| **Database:biological process      &nbspName:single-organism metabolic process      &nbspID:GO:0044710** | | | | | | |
| --- | --- | --- | --- | --- | --- | --- |
| C=9109; O=85; E=65.69; R=1.29; rawP=2.81e-05; adjP=0.0005 | | | | | | |
| Index | UserID | Value | Gene Symbol | Gene Name | EntrezGene | Ensembl |
| 1 | P10253 | NA | GAA | glucosidase, alpha; acid | 2548 | ENSG00000171298 |
| 2 | P68371 | NA | TUBB4B | tubulin, beta 4B class IVb | 10383 | ENSG00000188229 |
| 3 | Q9UL46 | NA | PSME2 | proteasome (prosome, macropain) activator subunit 2 (PA28 beta) | 5721 | ENSG00000100911 |
| 4 | P40926 | NA | MDH2 | malate dehydrogenase 2, NAD (mitochondrial) | 4191 | ENSG00000146701 |
| 5 | P04179 | NA | SOD2 | superoxide dismutase 2, mitochondrial | 6648 | ENSG00000112096 |
| 6 | Q13724 | NA | MOGS | mannosyl-oligosaccharide glucosidase | 7841 | ENSG00000115275 |
| 7 | P48735 | NA | IDH2 | isocitrate dehydrogenase 2 (NADP+), mitochondrial | 3418 | ENSG00000182054 |
| 8 | P00367 | NA | GLUD1 | glutamate dehydrogenase 1 | 2746 | ENSG00000148672 |
| 9 | Q06323 | NA | PSME1 | proteasome (prosome, macropain) activator subunit 1 (PA28 alpha) | 5720 | ENSG00000092010 |
| 10 | P53701 | NA | HCCS | holocytochrome c synthase | 3052 | ENSG00000004961 |
| 11 | P31040 | NA | SDHA | succinate dehydrogenase complex, subunit A, flavoprotein (Fp) | 6389 | ENSG00000073578 |
| 12 | P24534 | NA | EEF1B2 | eukaryotic translation elongation factor 1 beta 2 | 1933 | ENSG00000114942 |
| 13 | O14773 | NA | TPP1 | tripeptidyl peptidase I | 1200 | ENSG00000166340 |
| 14 | P47895 | NA | ALDH1A3 | aldehyde dehydrogenase 1 family, member A3 | 220 | ENSG00000184254 |
| 15 | P13804 | NA | ETFA | electron-transfer-flavoprotein, alpha polypeptide | 2108 | ENSG00000140374 |
| 16 | P42765 | NA | ACAA2 | acetyl-CoA acyltransferase 2 | 10449 | ENSG00000167315 |
| 17 | P17174 | NA | GOT1 | glutamic-oxaloacetic transaminase 1, soluble (aspartate aminotransferase 1) | 2805 | ENSG00000120053 |
| 18 | Q04837 | NA | SSBP1 | single-stranded DNA binding protein 1, mitochondrial | 6742 | ENSG00000106028 |
| 19 | P50454 | NA | SERPINH1 | serpin peptidase inhibitor, clade H (heat shock protein 47), member 1, (collagen binding protein 1) | 871 | ENSG00000149257 |
| 20 | Q14697 | NA | GANAB | glucosidase, alpha; neutral AB | 23193 | ENSG00000089597 |
| 21 | P08195 | NA | SLC3A2 | solute carrier family 3 (activators of dibasic and neutral amino acid transport), member 2 | 6520 | ENSG00000168003 |
| 22 | P40939 | NA | HADHA | hydroxyacyl-CoA dehydrogenase/3-ketoacyl-CoA thiolase/enoyl-CoA hydratase (trifunctional protein), alpha subunit | 3030 | ENSG00000084754 |
| 23 | P52815 | NA | MRPL12 | mitochondrial ribosomal protein L12 | 6182 | ENSG00000262814 |
| 24 | P38571 | NA | LIPA | lipase A, lysosomal acid, cholesterol esterase | 3988 | ENSG00000107798 |
| 25 | P32322 | NA | PYCR1 | pyrroline-5-carboxylate reductase 1 | 5831 | ENSG00000183010 |
| 26 | Q99798 | NA | ACO2 | aconitase 2, mitochondrial | 50 | ENSG00000100412 |
| 27 | P51649 | NA | ALDH5A1 | aldehyde dehydrogenase 5 family, member A1 | 7915 | ENSG00000112294 |
| 28 | P12235 | NA | SLC25A4 | solute carrier family 25 (mitochondrial carrier; adenine nucleotide translocator), member 4 | 291 | ENSG00000151729 |
| 29 | P05556 | NA | ITGB1 | integrin, beta 1 (fibronectin receptor, beta polypeptide, antigen CD29 includes MDF2, MSK12) | 3688 | ENSG00000150093 |
| 30 | Q9NSE4 | NA | IARS2 | isoleucyl-tRNA synthetase 2, mitochondrial | 55699 | ENSG00000067704 |
| 31 | Q6PI48 | NA | DARS2 | aspartyl-tRNA synthetase 2, mitochondrial | 55157 | ENSG00000117593 |
| 32 | P40227 | NA | CCT6A | chaperonin containing TCP1, subunit 6A (zeta 1) | 908 | ENSG00000146731 |
| 33 | P13674 | NA | P4HA1 | prolyl 4-hydroxylase, alpha polypeptide I | 5033 | ENSG00000122884 |
| 34 | Q02978 | NA | SLC25A11 | solute carrier family 25 (mitochondrial carrier; oxoglutarate carrier), member 11 | 8402 | ENSG00000108528 |
| 35 | P60174 | NA | TPI1 | triosephosphate isomerase 1 | 7167 | ENSG00000111669 |
| 36 | P49589 | NA | CARS | cysteinyl-tRNA synthetase | 833 | ENSG00000110619 |
| 37 | P54819 | NA | AK2 | adenylate kinase 2 | 204 | ENSG00000004455 |
| 38 | P17931 | NA | LGALS3 | lectin, galactoside-binding, soluble, 3 | 3958 | ENSG00000131981 |
| 39 | Q99623 | NA | PHB2 | prohibitin 2 | 11331 | ENSG00000215021 |
| 40 | P50453 | NA | SERPINB9 | serpin peptidase inhibitor, clade B (ovalbumin), member 9 | 5272 | ENSG00000170542 |
| 41 | P35232 | NA | PHB | prohibitin | 5245 | ENSG00000167085 |
| 42 | Q99714 | NA | HSD17B10 | hydroxysteroid (17-beta) dehydrogenase 10 | 3028 | ENSG00000072506 |
| 43 | O95831 | NA | AIFM1 | apoptosis-inducing factor, mitochondrion-associated, 1 | 9131 | ENSG00000156709 |
| 44 | P61313 | NA | RPL15 | ribosomal protein L15 | 6138 | ENSG00000174748 |
| 45 | P07741 | NA | APRT | adenine phosphoribosyltransferase | 353 | ENSG00000198931 |
| 46 | O60313 | NA | OPA1 | optic atrophy 1 (autosomal dominant) | 4976 | ENSG00000198836 |
| 47 | P35270 | NA | SPR | sepiapterin reductase (7,8-dihydrobiopterin:NADP+ oxidoreductase) | 6697 | ENSG00000116096 |
| 48 | P68104 | NA | EEF1A1 | eukaryotic translation elongation factor 1 alpha 1 | 1915 | ENSG00000156508 |
| 49 | P02786 | NA | TFRC | transferrin receptor (p90, CD71) | 7037 | ENSG00000072274 |
| 50 | P38117 | NA | ETFB | electron-transfer-flavoprotein, beta polypeptide | 2109 | ENSG00000105379 |
| 51 | P43490 | NA | NAMPT | nicotinamide phosphoribosyltransferase | 10135 | ENSG00000105835 |
| 52 | P22102 | NA | GART | phosphoribosylglycinamide formyltransferase, phosphoribosylglycinamide synthetase, phosphoribosylaminoimidazole synthetase | 2618 | ENSG00000159131 |
| 53 | P13639 | NA | EEF2 | eukaryotic translation elongation factor 2 | 1938 | ENSG00000167658 |
| 54 | O75367 | NA | H2AFY | H2A histone family, member Y | 9555 | ENSG00000113648 |
| 55 | O75521 | NA | ECI2 | enoyl-CoA delta isomerase 2 | 10455 | ENSG00000198721 |
| 56 | P67809 | NA | YBX1 | Y box binding protein 1 | 4904 | ENSG00000065978 |
| 57 | P49419 | NA | ALDH7A1 | aldehyde dehydrogenase 7 family, member A1 | 501 | ENSG00000164904 |
| 58 | Q6PIU2 | NA | NCEH1 | neutral cholesterol ester hydrolase 1 | 57552 | ENSG00000144959 |
| 59 | P29692 | NA | EEF1D | eukaryotic translation elongation factor 1 delta (guanine nucleotide exchange protein) | 1936 | ENSG00000104529 |
| 60 | P11387 | NA | TOP1 | topoisomerase (DNA) I | 7150 | ENSG00000198900 |
| 61 | Q5SSJ5 | NA | HP1BP3 | heterochromatin protein 1, binding protein 3 | 50809 | ENSG00000127483 |
| 62 | O75439 | NA | PMPCB | peptidase (mitochondrial processing) beta | 9512 | ENSG00000105819 |
| 63 | P00491 | NA | PNP | purine nucleoside phosphorylase | 4860 | ENSG00000198805 |
| 64 | P21953 | NA | BCKDHB | branched chain keto acid dehydrogenase E1, beta polypeptide | 594 | ENSG00000083123 |
| 65 | P30084 | NA | ECHS1 | enoyl CoA hydratase, short chain, 1, mitochondrial | 1892 | ENSG00000127884 |
| 66 | P60842 | NA | EIF4A1 | eukaryotic translation initiation factor 4A1 | 1973 | ENSG00000161960 |
| 67 | Q13011 | NA | ECH1 | enoyl CoA hydratase 1, peroxisomal | 1891 | ENSG00000104823 |
| 68 | P04083 | NA | ANXA1 | annexin A1 | 301 | ENSG00000135046 |
| 69 | P04040 | NA | CAT | catalase | 847 | ENSG00000121691 |
| 70 | P51688 | NA | SGSH | N-sulfoglucosamine sulfohydrolase | 6448 | ENSG00000181523 |
| 71 | P00558 | NA | PGK1 | phosphoglycerate kinase 1 | 5230 | ENSG00000102144 |
| 72 | P30048 | NA | PRDX3 | peroxiredoxin 3 | 10935 | ENSG00000165672 |
| 73 | P41091 | NA | EIF2S3 | eukaryotic translation initiation factor 2, subunit 3 gamma, 52kDa | 1968 | ENSG00000130741 |
| 74 | P02768 | NA | ALB | albumin | 213 | ENSG00000163631 |
| 75 | P07108 | NA | DBI | diazepam binding inhibitor (GABA receptor modulator, acyl-CoA binding protein) | 1622 | ENSG00000155368 |
| 76 | P24752 | NA | ACAT1 | acetyl-CoA acetyltransferase 1 | 38 | ENSG00000075239 |
| 77 | P19367 | NA | HK1 | hexokinase 1 | 3098 | ENSG00000156515 |
| 78 | P25705 | NA | ATP5A1 | ATP synthase, H+ transporting, mitochondrial F1 complex, alpha subunit 1, cardiac muscle | 498 | ENSG00000152234 |
| 79 | Q9BQ69 | NA | MACROD1 | MACRO domain containing 1 | 28992 | ENSG00000133315 |
| 80 | P06865 | NA | HEXA | hexosaminidase A (alpha polypeptide) | 3073 | ENSG00000213614 |
| 81 | P49748 | NA | ACADVL | acyl-CoA dehydrogenase, very long chain | 37 | ENSG00000072778 |
| 82 | P00352 | NA | ALDH1A1 | aldehyde dehydrogenase 1 family, member A1 | 216 | ENSG00000165092 |
| 83 | P48047 | NA | ATP5O | ATP synthase, H+ transporting, mitochondrial F1 complex, O subunit | 539 | ENSG00000241837 |
| 84 | Q16531 | NA | DDB1 | damage-specific DNA binding protein 1, 127kDa | 1642 | ENSG00000167986 |
| 85 | P31937 | NA | HIBADH | 3-hydroxyisobutyrate dehydrogenase | 11112 | ENSG00000106049 |

  
  

| **Database:biological process      &nbspName:pyridine nucleotide metabolic process      &nbspID:GO:0019362** | | | | | | |
| --- | --- | --- | --- | --- | --- | --- |
| C=49; O=5; E=0.35; R=14.15; rawP=2.62e-05; adjP=0.0005 | | | | | | |
| Index | UserID | Value | Gene Symbol | Gene Name | EntrezGene | Ensembl |
| 1 | P00491 | NA | PNP | purine nucleoside phosphorylase | 4860 | ENSG00000198805 |
| 2 | P40926 | NA | MDH2 | malate dehydrogenase 2, NAD (mitochondrial) | 4191 | ENSG00000146701 |
| 3 | P60174 | NA | TPI1 | triosephosphate isomerase 1 | 7167 | ENSG00000111669 |
| 4 | P31937 | NA | HIBADH | 3-hydroxyisobutyrate dehydrogenase | 11112 | ENSG00000106049 |
| 5 | P43490 | NA | NAMPT | nicotinamide phosphoribosyltransferase | 10135 | ENSG00000105835 |

  
  

| **Database:biological process      &nbspName:dicarboxylic acid metabolic process      &nbspID:GO:0043648** | | | | | | |
| --- | --- | --- | --- | --- | --- | --- |
| C=81; O=6; E=0.58; R=10.27; rawP=2.55e-05; adjP=0.0005 | | | | | | |
| Index | UserID | Value | Gene Symbol | Gene Name | EntrezGene | Ensembl |
| 1 | P17174 | NA | GOT1 | glutamic-oxaloacetic transaminase 1, soluble (aspartate aminotransferase 1) | 2805 | ENSG00000120053 |
| 2 | P48735 | NA | IDH2 | isocitrate dehydrogenase 2 (NADP+), mitochondrial | 3418 | ENSG00000182054 |
| 3 | P00367 | NA | GLUD1 | glutamate dehydrogenase 1 | 2746 | ENSG00000148672 |
| 4 | P51649 | NA | ALDH5A1 | aldehyde dehydrogenase 5 family, member A1 | 7915 | ENSG00000112294 |
| 5 | P31040 | NA | SDHA | succinate dehydrogenase complex, subunit A, flavoprotein (Fp) | 6389 | ENSG00000073578 |
| 6 | P40926 | NA | MDH2 | malate dehydrogenase 2, NAD (mitochondrial) | 4191 | ENSG00000146701 |

  
  

| **Database:biological process      &nbspName:glucose metabolic process      &nbspID:GO:0006006** | | | | | | |
| --- | --- | --- | --- | --- | --- | --- |
| C=213; O=9; E=1.54; R=5.86; rawP=2.31e-05; adjP=0.0005 | | | | | | |
| Index | UserID | Value | Gene Symbol | Gene Name | EntrezGene | Ensembl |
| 1 | P10253 | NA | GAA | glucosidase, alpha; acid | 2548 | ENSG00000171298 |
| 2 | P51649 | NA | ALDH5A1 | aldehyde dehydrogenase 5 family, member A1 | 7915 | ENSG00000112294 |
| 3 | P00558 | NA | PGK1 | phosphoglycerate kinase 1 | 5230 | ENSG00000102144 |
| 4 | P40926 | NA | MDH2 | malate dehydrogenase 2, NAD (mitochondrial) | 4191 | ENSG00000146701 |
| 5 | P17174 | NA | GOT1 | glutamic-oxaloacetic transaminase 1, soluble (aspartate aminotransferase 1) | 2805 | ENSG00000120053 |
| 6 | Q02978 | NA | SLC25A11 | solute carrier family 25 (mitochondrial carrier; oxoglutarate carrier), member 11 | 8402 | ENSG00000108528 |
| 7 | P19367 | NA | HK1 | hexokinase 1 | 3098 | ENSG00000156515 |
| 8 | P60174 | NA | TPI1 | triosephosphate isomerase 1 | 7167 | ENSG00000111669 |
| 9 | P31937 | NA | HIBADH | 3-hydroxyisobutyrate dehydrogenase | 11112 | ENSG00000106049 |

  
  

| **Database:biological process      &nbspName:acetyl-CoA catabolic process      &nbspID:GO:0046356** | | | | | | |
| --- | --- | --- | --- | --- | --- | --- |
| C=27; O=4; E=0.19; R=20.54; rawP=3.94e-05; adjP=0.0006 | | | | | | |
| Index | UserID | Value | Gene Symbol | Gene Name | EntrezGene | Ensembl |
| 1 | Q99798 | NA | ACO2 | aconitase 2, mitochondrial | 50 | ENSG00000100412 |
| 2 | P48735 | NA | IDH2 | isocitrate dehydrogenase 2 (NADP+), mitochondrial | 3418 | ENSG00000182054 |
| 3 | P31040 | NA | SDHA | succinate dehydrogenase complex, subunit A, flavoprotein (Fp) | 6389 | ENSG00000073578 |
| 4 | P40926 | NA | MDH2 | malate dehydrogenase 2, NAD (mitochondrial) | 4191 | ENSG00000146701 |

  
  

| **Database:biological process      &nbspName:fatty acid metabolic process      &nbspID:GO:0006631** | | | | | | |
| --- | --- | --- | --- | --- | --- | --- |
| C=281; O=10; E=2.03; R=4.93; rawP=3.50e-05; adjP=0.0006 | | | | | | |
| Index | UserID | Value | Gene Symbol | Gene Name | EntrezGene | Ensembl |
| 1 | P51649 | NA | ALDH5A1 | aldehyde dehydrogenase 5 family, member A1 | 7915 | ENSG00000112294 |
| 2 | P40939 | NA | HADHA | hydroxyacyl-CoA dehydrogenase/3-ketoacyl-CoA thiolase/enoyl-CoA hydratase (trifunctional protein), alpha subunit | 3030 | ENSG00000084754 |
| 3 | P30084 | NA | ECHS1 | enoyl CoA hydratase, short chain, 1, mitochondrial | 1892 | ENSG00000127884 |
| 4 | P42765 | NA | ACAA2 | acetyl-CoA acyltransferase 2 | 10449 | ENSG00000167315 |
| 5 | O75521 | NA | ECI2 | enoyl-CoA delta isomerase 2 | 10455 | ENSG00000198721 |
| 6 | P17174 | NA | GOT1 | glutamic-oxaloacetic transaminase 1, soluble (aspartate aminotransferase 1) | 2805 | ENSG00000120053 |
| 7 | P49748 | NA | ACADVL | acyl-CoA dehydrogenase, very long chain | 37 | ENSG00000072778 |
| 8 | P38571 | NA | LIPA | lipase A, lysosomal acid, cholesterol esterase | 3988 | ENSG00000107798 |
| 9 | Q13011 | NA | ECH1 | enoyl CoA hydratase 1, peroxisomal | 1891 | ENSG00000104823 |
| 10 | P04083 | NA | ANXA1 | annexin A1 | 301 | ENSG00000135046 |

  
  

| **Database:biological process      &nbspName:mitochondrion organization      &nbspID:GO:0007005** | | | | | | |
| --- | --- | --- | --- | --- | --- | --- |
| C=223; O=9; E=1.61; R=5.60; rawP=3.31e-05; adjP=0.0006 | | | | | | |
| Index | UserID | Value | Gene Symbol | Gene Name | EntrezGene | Ensembl |
| 1 | O75439 | NA | PMPCB | peptidase (mitochondrial processing) beta | 9512 | ENSG00000105819 |
| 2 | Q04837 | NA | SSBP1 | single-stranded DNA binding protein 1, mitochondrial | 6742 | ENSG00000106028 |
| 3 | O60313 | NA | OPA1 | optic atrophy 1 (autosomal dominant) | 4976 | ENSG00000198836 |
| 4 | P12235 | NA | SLC25A4 | solute carrier family 25 (mitochondrial carrier; adenine nucleotide translocator), member 4 | 291 | ENSG00000151729 |
| 5 | P30048 | NA | PRDX3 | peroxiredoxin 3 | 10935 | ENSG00000165672 |
| 6 | O95831 | NA | AIFM1 | apoptosis-inducing factor, mitochondrion-associated, 1 | 9131 | ENSG00000156709 |
| 7 | Q6PI48 | NA | DARS2 | aspartyl-tRNA synthetase 2, mitochondrial | 55157 | ENSG00000117593 |
| 8 | P52815 | NA | MRPL12 | mitochondrial ribosomal protein L12 | 6182 | ENSG00000262814 |
| 9 | P04179 | NA | SOD2 | superoxide dismutase 2, mitochondrial | 6648 | ENSG00000112096 |

  
  

| **Database:molecular function      &nbspName:coenzyme binding      &nbspID:GO:0050662** | | | | | | |
| --- | --- | --- | --- | --- | --- | --- |
| C=184; O=15; E=1.28; R=11.74; rawP=2.27e-12; adjP=1.50e-10 | | | | | | |
| Index | UserID | Value | Gene Symbol | Gene Name | EntrezGene | Ensembl |
| 1 | P04040 | NA | CAT | catalase | 847 | ENSG00000121691 |
| 2 | P40939 | NA | HADHA | hydroxyacyl-CoA dehydrogenase/3-ketoacyl-CoA thiolase/enoyl-CoA hydratase (trifunctional protein), alpha subunit | 3030 | ENSG00000084754 |
| 3 | P07108 | NA | DBI | diazepam binding inhibitor (GABA receptor modulator, acyl-CoA binding protein) | 1622 | ENSG00000155368 |
| 4 | P47895 | NA | ALDH1A3 | aldehyde dehydrogenase 1 family, member A3 | 220 | ENSG00000184254 |
| 5 | O95831 | NA | AIFM1 | apoptosis-inducing factor, mitochondrion-associated, 1 | 9131 | ENSG00000156709 |
| 6 | Q99714 | NA | HSD17B10 | hydroxysteroid (17-beta) dehydrogenase 10 | 3028 | ENSG00000072506 |
| 7 | P13804 | NA | ETFA | electron-transfer-flavoprotein, alpha polypeptide | 2108 | ENSG00000140374 |
| 8 | P24752 | NA | ACAT1 | acetyl-CoA acetyltransferase 1 | 38 | ENSG00000075239 |
| 9 | O75521 | NA | ECI2 | enoyl-CoA delta isomerase 2 | 10455 | ENSG00000198721 |
| 10 | P00367 | NA | GLUD1 | glutamate dehydrogenase 1 | 2746 | ENSG00000148672 |
| 11 | P48735 | NA | IDH2 | isocitrate dehydrogenase 2 (NADP+), mitochondrial | 3418 | ENSG00000182054 |
| 12 | P49748 | NA | ACADVL | acyl-CoA dehydrogenase, very long chain | 37 | ENSG00000072778 |
| 13 | P35270 | NA | SPR | sepiapterin reductase (7,8-dihydrobiopterin:NADP+ oxidoreductase) | 6697 | ENSG00000116096 |
| 14 | P31040 | NA | SDHA | succinate dehydrogenase complex, subunit A, flavoprotein (Fp) | 6389 | ENSG00000073578 |
| 15 | P31937 | NA | HIBADH | 3-hydroxyisobutyrate dehydrogenase | 11112 | ENSG00000106049 |

  
  

| **Database:molecular function      &nbspName:cofactor binding      &nbspID:GO:0048037** | | | | | | |
| --- | --- | --- | --- | --- | --- | --- |
| C=259; O=17; E=1.80; R=9.45; rawP=2.35e-12; adjP=1.50e-10 | | | | | | |
| Index | UserID | Value | Gene Symbol | Gene Name | EntrezGene | Ensembl |
| 1 | P40939 | NA | HADHA | hydroxyacyl-CoA dehydrogenase/3-ketoacyl-CoA thiolase/enoyl-CoA hydratase (trifunctional protein), alpha subunit | 3030 | ENSG00000084754 |
| 2 | O95831 | NA | AIFM1 | apoptosis-inducing factor, mitochondrion-associated, 1 | 9131 | ENSG00000156709 |
| 3 | Q99714 | NA | HSD17B10 | hydroxysteroid (17-beta) dehydrogenase 10 | 3028 | ENSG00000072506 |
| 4 | P00367 | NA | GLUD1 | glutamate dehydrogenase 1 | 2746 | ENSG00000148672 |
| 5 | P48735 | NA | IDH2 | isocitrate dehydrogenase 2 (NADP+), mitochondrial | 3418 | ENSG00000182054 |
| 6 | P35270 | NA | SPR | sepiapterin reductase (7,8-dihydrobiopterin:NADP+ oxidoreductase) | 6697 | ENSG00000116096 |
| 7 | P31040 | NA | SDHA | succinate dehydrogenase complex, subunit A, flavoprotein (Fp) | 6389 | ENSG00000073578 |
| 8 | P04040 | NA | CAT | catalase | 847 | ENSG00000121691 |
| 9 | P02768 | NA | ALB | albumin | 213 | ENSG00000163631 |
| 10 | P47895 | NA | ALDH1A3 | aldehyde dehydrogenase 1 family, member A3 | 220 | ENSG00000184254 |
| 11 | P07108 | NA | DBI | diazepam binding inhibitor (GABA receptor modulator, acyl-CoA binding protein) | 1622 | ENSG00000155368 |
| 12 | P13804 | NA | ETFA | electron-transfer-flavoprotein, alpha polypeptide | 2108 | ENSG00000140374 |
| 13 | O75521 | NA | ECI2 | enoyl-CoA delta isomerase 2 | 10455 | ENSG00000198721 |
| 14 | P24752 | NA | ACAT1 | acetyl-CoA acetyltransferase 1 | 38 | ENSG00000075239 |
| 15 | P17174 | NA | GOT1 | glutamic-oxaloacetic transaminase 1, soluble (aspartate aminotransferase 1) | 2805 | ENSG00000120053 |
| 16 | P49748 | NA | ACADVL | acyl-CoA dehydrogenase, very long chain | 37 | ENSG00000072778 |
| 17 | P31937 | NA | HIBADH | 3-hydroxyisobutyrate dehydrogenase | 11112 | ENSG00000106049 |

  
  

| **Database:molecular function      &nbspName:oxidoreductase activity      &nbspID:GO:0016491** | | | | | | |
| --- | --- | --- | --- | --- | --- | --- |
| C=697; O=22; E=4.84; R=4.55; rawP=1.86e-09; adjP=7.94e-08 | | | | | | |
| Index | UserID | Value | Gene Symbol | Gene Name | EntrezGene | Ensembl |
| 1 | P02792 | NA | FTL | ferritin, light polypeptide | 2512 | ENSG00000087086 |
| 2 | P21953 | NA | BCKDHB | branched chain keto acid dehydrogenase E1, beta polypeptide | 594 | ENSG00000083123 |
| 3 | P40939 | NA | HADHA | hydroxyacyl-CoA dehydrogenase/3-ketoacyl-CoA thiolase/enoyl-CoA hydratase (trifunctional protein), alpha subunit | 3030 | ENSG00000084754 |
| 4 | P40926 | NA | MDH2 | malate dehydrogenase 2, NAD (mitochondrial) | 4191 | ENSG00000146701 |
| 5 | O95831 | NA | AIFM1 | apoptosis-inducing factor, mitochondrion-associated, 1 | 9131 | ENSG00000156709 |
| 6 | Q99714 | NA | HSD17B10 | hydroxysteroid (17-beta) dehydrogenase 10 | 3028 | ENSG00000072506 |
| 7 | P04179 | NA | SOD2 | superoxide dismutase 2, mitochondrial | 6648 | ENSG00000112096 |
| 8 | P00367 | NA | GLUD1 | glutamate dehydrogenase 1 | 2746 | ENSG00000148672 |
| 9 | P48735 | NA | IDH2 | isocitrate dehydrogenase 2 (NADP+), mitochondrial | 3418 | ENSG00000182054 |
| 10 | P35270 | NA | SPR | sepiapterin reductase (7,8-dihydrobiopterin:NADP+ oxidoreductase) | 6697 | ENSG00000116096 |
| 11 | P31040 | NA | SDHA | succinate dehydrogenase complex, subunit A, flavoprotein (Fp) | 6389 | ENSG00000073578 |
| 12 | P32322 | NA | PYCR1 | pyrroline-5-carboxylate reductase 1 | 5831 | ENSG00000183010 |
| 13 | P04040 | NA | CAT | catalase | 847 | ENSG00000121691 |
| 14 | P51649 | NA | ALDH5A1 | aldehyde dehydrogenase 5 family, member A1 | 7915 | ENSG00000112294 |
| 15 | P30048 | NA | PRDX3 | peroxiredoxin 3 | 10935 | ENSG00000165672 |
| 16 | P47895 | NA | ALDH1A3 | aldehyde dehydrogenase 1 family, member A3 | 220 | ENSG00000184254 |
| 17 | P13804 | NA | ETFA | electron-transfer-flavoprotein, alpha polypeptide | 2108 | ENSG00000140374 |
| 18 | P13674 | NA | P4HA1 | prolyl 4-hydroxylase, alpha polypeptide I | 5033 | ENSG00000122884 |
| 19 | P49419 | NA | ALDH7A1 | aldehyde dehydrogenase 7 family, member A1 | 501 | ENSG00000164904 |
| 20 | P49748 | NA | ACADVL | acyl-CoA dehydrogenase, very long chain | 37 | ENSG00000072778 |
| 21 | P00352 | NA | ALDH1A1 | aldehyde dehydrogenase 1 family, member A1 | 216 | ENSG00000165092 |
| 22 | P31937 | NA | HIBADH | 3-hydroxyisobutyrate dehydrogenase | 11112 | ENSG00000106049 |

  
  

| **Database:molecular function      &nbspName:catalytic activity      &nbspID:GO:0003824** | | | | | | |
| --- | --- | --- | --- | --- | --- | --- |
| C=5299; O=65; E=36.80; R=1.77; rawP=1.50e-08; adjP=4.80e-07 | | | | | | |
| Index | UserID | Value | Gene Symbol | Gene Name | EntrezGene | Ensembl |
| 1 | P49589 | NA | CARS | cysteinyl-tRNA synthetase | 833 | ENSG00000110619 |
| 2 | P10253 | NA | GAA | glucosidase, alpha; acid | 2548 | ENSG00000171298 |
| 3 | P54819 | NA | AK2 | adenylate kinase 2 | 204 | ENSG00000004455 |
| 4 | P68371 | NA | TUBB4B | tubulin, beta 4B class IVb | 10383 | ENSG00000188229 |
| 5 | P40926 | NA | MDH2 | malate dehydrogenase 2, NAD (mitochondrial) | 4191 | ENSG00000146701 |
| 6 | O95831 | NA | AIFM1 | apoptosis-inducing factor, mitochondrion-associated, 1 | 9131 | ENSG00000156709 |
| 7 | Q99714 | NA | HSD17B10 | hydroxysteroid (17-beta) dehydrogenase 10 | 3028 | ENSG00000072506 |
| 8 | P04179 | NA | SOD2 | superoxide dismutase 2, mitochondrial | 6648 | ENSG00000112096 |
| 9 | Q13724 | NA | MOGS | mannosyl-oligosaccharide glucosidase | 7841 | ENSG00000115275 |
| 10 | P07741 | NA | APRT | adenine phosphoribosyltransferase | 353 | ENSG00000198931 |
| 11 | P00367 | NA | GLUD1 | glutamate dehydrogenase 1 | 2746 | ENSG00000148672 |
| 12 | P48735 | NA | IDH2 | isocitrate dehydrogenase 2 (NADP+), mitochondrial | 3418 | ENSG00000182054 |
| 13 | O60313 | NA | OPA1 | optic atrophy 1 (autosomal dominant) | 4976 | ENSG00000198836 |
| 14 | P53701 | NA | HCCS | holocytochrome c synthase | 3052 | ENSG00000004961 |
| 15 | P35270 | NA | SPR | sepiapterin reductase (7,8-dihydrobiopterin:NADP+ oxidoreductase) | 6697 | ENSG00000116096 |
| 16 | P35580 | NA | MYH10 | myosin, heavy chain 10, non-muscle | 4628 | ENSG00000133026 |
| 17 | P68104 | NA | EEF1A1 | eukaryotic translation elongation factor 1 alpha 1 | 1915 | ENSG00000156508 |
| 18 | P31040 | NA | SDHA | succinate dehydrogenase complex, subunit A, flavoprotein (Fp) | 6389 | ENSG00000073578 |
| 19 | P02786 | NA | TFRC | transferrin receptor (p90, CD71) | 7037 | ENSG00000072274 |
| 20 | P43490 | NA | NAMPT | nicotinamide phosphoribosyltransferase | 10135 | ENSG00000105835 |
| 21 | O14773 | NA | TPP1 | tripeptidyl peptidase I | 1200 | ENSG00000166340 |
| 22 | P22102 | NA | GART | phosphoribosylglycinamide formyltransferase, phosphoribosylglycinamide synthetase, phosphoribosylaminoimidazole synthetase | 2618 | ENSG00000159131 |
| 23 | P13639 | NA | EEF2 | eukaryotic translation elongation factor 2 | 1938 | ENSG00000167658 |
| 24 | P47895 | NA | ALDH1A3 | aldehyde dehydrogenase 1 family, member A3 | 220 | ENSG00000184254 |
| 25 | P13804 | NA | ETFA | electron-transfer-flavoprotein, alpha polypeptide | 2108 | ENSG00000140374 |
| 26 | P42765 | NA | ACAA2 | acetyl-CoA acyltransferase 2 | 10449 | ENSG00000167315 |
| 27 | O75521 | NA | ECI2 | enoyl-CoA delta isomerase 2 | 10455 | ENSG00000198721 |
| 28 | P49419 | NA | ALDH7A1 | aldehyde dehydrogenase 7 family, member A1 | 501 | ENSG00000164904 |
| 29 | P17174 | NA | GOT1 | glutamic-oxaloacetic transaminase 1, soluble (aspartate aminotransferase 1) | 2805 | ENSG00000120053 |
| 30 | Q6PIU2 | NA | NCEH1 | neutral cholesterol ester hydrolase 1 | 57552 | ENSG00000144959 |
| 31 | Q14697 | NA | GANAB | glucosidase, alpha; neutral AB | 23193 | ENSG00000089597 |
| 32 | P11387 | NA | TOP1 | topoisomerase (DNA) I | 7150 | ENSG00000198900 |
| 33 | P02792 | NA | FTL | ferritin, light polypeptide | 2512 | ENSG00000087086 |
| 34 | O75439 | NA | PMPCB | peptidase (mitochondrial processing) beta | 9512 | ENSG00000105819 |
| 35 | Q9NR30 | NA | DDX21 | DEAD (Asp-Glu-Ala-Asp) box helicase 21 | 9188 | ENSG00000165732 |
| 36 | P08195 | NA | SLC3A2 | solute carrier family 3 (activators of dibasic and neutral amino acid transport), member 2 | 6520 | ENSG00000168003 |
| 37 | P00491 | NA | PNP | purine nucleoside phosphorylase | 4860 | ENSG00000198805 |
| 38 | P21953 | NA | BCKDHB | branched chain keto acid dehydrogenase E1, beta polypeptide | 594 | ENSG00000083123 |
| 39 | P40939 | NA | HADHA | hydroxyacyl-CoA dehydrogenase/3-ketoacyl-CoA thiolase/enoyl-CoA hydratase (trifunctional protein), alpha subunit | 3030 | ENSG00000084754 |
| 40 | P30084 | NA | ECHS1 | enoyl CoA hydratase, short chain, 1, mitochondrial | 1892 | ENSG00000127884 |
| 41 | P60842 | NA | EIF4A1 | eukaryotic translation initiation factor 4A1 | 1973 | ENSG00000161960 |
| 42 | Q9BXW7 | NA | CECR5 | cat eye syndrome chromosome region, candidate 5 | 27440 | ENSG00000069998 |
| 43 | P38571 | NA | LIPA | lipase A, lysosomal acid, cholesterol esterase | 3988 | ENSG00000107798 |
| 44 | Q13011 | NA | ECH1 | enoyl CoA hydratase 1, peroxisomal | 1891 | ENSG00000104823 |
| 45 | P32322 | NA | PYCR1 | pyrroline-5-carboxylate reductase 1 | 5831 | ENSG00000183010 |
| 46 | P04040 | NA | CAT | catalase | 847 | ENSG00000121691 |
| 47 | Q99798 | NA | ACO2 | aconitase 2, mitochondrial | 50 | ENSG00000100412 |
| 48 | P51688 | NA | SGSH | N-sulfoglucosamine sulfohydrolase | 6448 | ENSG00000181523 |
| 49 | P51649 | NA | ALDH5A1 | aldehyde dehydrogenase 5 family, member A1 | 7915 | ENSG00000112294 |
| 50 | P00558 | NA | PGK1 | phosphoglycerate kinase 1 | 5230 | ENSG00000102144 |
| 51 | P30048 | NA | PRDX3 | peroxiredoxin 3 | 10935 | ENSG00000165672 |
| 52 | P41091 | NA | EIF2S3 | eukaryotic translation initiation factor 2, subunit 3 gamma, 52kDa | 1968 | ENSG00000130741 |
| 53 | Q9NSE4 | NA | IARS2 | isoleucyl-tRNA synthetase 2, mitochondrial | 55699 | ENSG00000067704 |
| 54 | Q6PI48 | NA | DARS2 | aspartyl-tRNA synthetase 2, mitochondrial | 55157 | ENSG00000117593 |
| 55 | P24752 | NA | ACAT1 | acetyl-CoA acetyltransferase 1 | 38 | ENSG00000075239 |
| 56 | P13674 | NA | P4HA1 | prolyl 4-hydroxylase, alpha polypeptide I | 5033 | ENSG00000122884 |
| 57 | P19367 | NA | HK1 | hexokinase 1 | 3098 | ENSG00000156515 |
| 58 | P25705 | NA | ATP5A1 | ATP synthase, H+ transporting, mitochondrial F1 complex, alpha subunit 1, cardiac muscle | 498 | ENSG00000152234 |
| 59 | Q9BQ69 | NA | MACROD1 | MACRO domain containing 1 | 28992 | ENSG00000133315 |
| 60 | P06865 | NA | HEXA | hexosaminidase A (alpha polypeptide) | 3073 | ENSG00000213614 |
| 61 | P49748 | NA | ACADVL | acyl-CoA dehydrogenase, very long chain | 37 | ENSG00000072778 |
| 62 | P00352 | NA | ALDH1A1 | aldehyde dehydrogenase 1 family, member A1 | 216 | ENSG00000165092 |
| 63 | P48047 | NA | ATP5O | ATP synthase, H+ transporting, mitochondrial F1 complex, O subunit | 539 | ENSG00000241837 |
| 64 | P60174 | NA | TPI1 | triosephosphate isomerase 1 | 7167 | ENSG00000111669 |
| 65 | P31937 | NA | HIBADH | 3-hydroxyisobutyrate dehydrogenase | 11112 | ENSG00000106049 |

  
  

| **Database:molecular function      &nbspName:NAD binding      &nbspID:GO:0051287** | | | | | | |
| --- | --- | --- | --- | --- | --- | --- |
| C=49; O=6; E=0.34; R=17.63; rawP=1.06e-06; adjP=2.71e-05 | | | | | | |
| Index | UserID | Value | Gene Symbol | Gene Name | EntrezGene | Ensembl |
| 1 | P48735 | NA | IDH2 | isocitrate dehydrogenase 2 (NADP+), mitochondrial | 3418 | ENSG00000182054 |
| 2 | P00367 | NA | GLUD1 | glutamate dehydrogenase 1 | 2746 | ENSG00000148672 |
| 3 | P40939 | NA | HADHA | hydroxyacyl-CoA dehydrogenase/3-ketoacyl-CoA thiolase/enoyl-CoA hydratase (trifunctional protein), alpha subunit | 3030 | ENSG00000084754 |
| 4 | P47895 | NA | ALDH1A3 | aldehyde dehydrogenase 1 family, member A3 | 220 | ENSG00000184254 |
| 5 | P31937 | NA | HIBADH | 3-hydroxyisobutyrate dehydrogenase | 11112 | ENSG00000106049 |
| 6 | Q99714 | NA | HSD17B10 | hydroxysteroid (17-beta) dehydrogenase 10 | 3028 | ENSG00000072506 |

  
  

| **Database:molecular function      &nbspName:small molecule binding      &nbspID:GO:0036094** | | | | | | |
| --- | --- | --- | --- | --- | --- | --- |
| C=2595; O=38; E=18.02; R=2.11; rawP=2.09e-06; adjP=4.46e-05 | | | | | | |
| Index | UserID | Value | Gene Symbol | Gene Name | EntrezGene | Ensembl |
| 1 | P49589 | NA | CARS | cysteinyl-tRNA synthetase | 833 | ENSG00000110619 |
| 2 | P54819 | NA | AK2 | adenylate kinase 2 | 204 | ENSG00000004455 |
| 3 | P68371 | NA | TUBB4B | tubulin, beta 4B class IVb | 10383 | ENSG00000188229 |
| 4 | P40926 | NA | MDH2 | malate dehydrogenase 2, NAD (mitochondrial) | 4191 | ENSG00000146701 |
| 5 | O95831 | NA | AIFM1 | apoptosis-inducing factor, mitochondrion-associated, 1 | 9131 | ENSG00000156709 |
| 6 | Q99714 | NA | HSD17B10 | hydroxysteroid (17-beta) dehydrogenase 10 | 3028 | ENSG00000072506 |
| 7 | P07741 | NA | APRT | adenine phosphoribosyltransferase | 353 | ENSG00000198931 |
| 8 | P00367 | NA | GLUD1 | glutamate dehydrogenase 1 | 2746 | ENSG00000148672 |
| 9 | P48735 | NA | IDH2 | isocitrate dehydrogenase 2 (NADP+), mitochondrial | 3418 | ENSG00000182054 |
| 10 | O60313 | NA | OPA1 | optic atrophy 1 (autosomal dominant) | 4976 | ENSG00000198836 |
| 11 | P35270 | NA | SPR | sepiapterin reductase (7,8-dihydrobiopterin:NADP+ oxidoreductase) | 6697 | ENSG00000116096 |
| 12 | P35580 | NA | MYH10 | myosin, heavy chain 10, non-muscle | 4628 | ENSG00000133026 |
| 13 | P68104 | NA | EEF1A1 | eukaryotic translation elongation factor 1 alpha 1 | 1915 | ENSG00000156508 |
| 14 | P31040 | NA | SDHA | succinate dehydrogenase complex, subunit A, flavoprotein (Fp) | 6389 | ENSG00000073578 |
| 15 | P22102 | NA | GART | phosphoribosylglycinamide formyltransferase, phosphoribosylglycinamide synthetase, phosphoribosylaminoimidazole synthetase | 2618 | ENSG00000159131 |
| 16 | P13639 | NA | EEF2 | eukaryotic translation elongation factor 2 | 1938 | ENSG00000167658 |
| 17 | P47895 | NA | ALDH1A3 | aldehyde dehydrogenase 1 family, member A3 | 220 | ENSG00000184254 |
| 18 | P13804 | NA | ETFA | electron-transfer-flavoprotein, alpha polypeptide | 2108 | ENSG00000140374 |
| 19 | Q9Y277 | NA | VDAC3 | voltage-dependent anion channel 3 | 7419 | ENSG00000078668 |
| 20 | P17174 | NA | GOT1 | glutamic-oxaloacetic transaminase 1, soluble (aspartate aminotransferase 1) | 2805 | ENSG00000120053 |
| 21 | P11387 | NA | TOP1 | topoisomerase (DNA) I | 7150 | ENSG00000198900 |
| 22 | Q9NR30 | NA | DDX21 | DEAD (Asp-Glu-Ala-Asp) box helicase 21 | 9188 | ENSG00000165732 |
| 23 | P00491 | NA | PNP | purine nucleoside phosphorylase | 4860 | ENSG00000198805 |
| 24 | P40939 | NA | HADHA | hydroxyacyl-CoA dehydrogenase/3-ketoacyl-CoA thiolase/enoyl-CoA hydratase (trifunctional protein), alpha subunit | 3030 | ENSG00000084754 |
| 25 | P60842 | NA | EIF4A1 | eukaryotic translation initiation factor 4A1 | 1973 | ENSG00000161960 |
| 26 | P32322 | NA | PYCR1 | pyrroline-5-carboxylate reductase 1 | 5831 | ENSG00000183010 |
| 27 | P04040 | NA | CAT | catalase | 847 | ENSG00000121691 |
| 28 | P00558 | NA | PGK1 | phosphoglycerate kinase 1 | 5230 | ENSG00000102144 |
| 29 | P02768 | NA | ALB | albumin | 213 | ENSG00000163631 |
| 30 | P41091 | NA | EIF2S3 | eukaryotic translation initiation factor 2, subunit 3 gamma, 52kDa | 1968 | ENSG00000130741 |
| 31 | Q6PI48 | NA | DARS2 | aspartyl-tRNA synthetase 2, mitochondrial | 55157 | ENSG00000117593 |
| 32 | Q9NSE4 | NA | IARS2 | isoleucyl-tRNA synthetase 2, mitochondrial | 55699 | ENSG00000067704 |
| 33 | P13674 | NA | P4HA1 | prolyl 4-hydroxylase, alpha polypeptide I | 5033 | ENSG00000122884 |
| 34 | P40227 | NA | CCT6A | chaperonin containing TCP1, subunit 6A (zeta 1) | 908 | ENSG00000146731 |
| 35 | P19367 | NA | HK1 | hexokinase 1 | 3098 | ENSG00000156515 |
| 36 | P25705 | NA | ATP5A1 | ATP synthase, H+ transporting, mitochondrial F1 complex, alpha subunit 1, cardiac muscle | 498 | ENSG00000152234 |
| 37 | P49748 | NA | ACADVL | acyl-CoA dehydrogenase, very long chain | 37 | ENSG00000072778 |
| 38 | P31937 | NA | HIBADH | 3-hydroxyisobutyrate dehydrogenase | 11112 | ENSG00000106049 |

  
  

| **Database:molecular function      &nbspName:translation elongation factor activity      &nbspID:GO:0003746** | | | | | | |
| --- | --- | --- | --- | --- | --- | --- |
| C=17; O=4; E=0.12; R=33.88; rawP=4.87e-06; adjP=8.91e-05 | | | | | | |
| Index | UserID | Value | Gene Symbol | Gene Name | EntrezGene | Ensembl |
| 1 | P13639 | NA | EEF2 | eukaryotic translation elongation factor 2 | 1938 | ENSG00000167658 |
| 2 | P68104 | NA | EEF1A1 | eukaryotic translation elongation factor 1 alpha 1 | 1915 | ENSG00000156508 |
| 3 | P29692 | NA | EEF1D | eukaryotic translation elongation factor 1 delta (guanine nucleotide exchange protein) | 1936 | ENSG00000104529 |
| 4 | P24534 | NA | EEF1B2 | eukaryotic translation elongation factor 1 beta 2 | 1933 | ENSG00000114942 |

  
  

| **Database:molecular function      &nbspName:acetyl-CoA C-acyltransferase activity      &nbspID:GO:0003988** | | | | | | |
| --- | --- | --- | --- | --- | --- | --- |
| C=6; O=3; E=0.04; R=72.00; rawP=6.41e-06; adjP=0.0001 | | | | | | |
| Index | UserID | Value | Gene Symbol | Gene Name | EntrezGene | Ensembl |
| 1 | P24752 | NA | ACAT1 | acetyl-CoA acetyltransferase 1 | 38 | ENSG00000075239 |
| 2 | P40939 | NA | HADHA | hydroxyacyl-CoA dehydrogenase/3-ketoacyl-CoA thiolase/enoyl-CoA hydratase (trifunctional protein), alpha subunit | 3030 | ENSG00000084754 |
| 3 | P42765 | NA | ACAA2 | acetyl-CoA acyltransferase 2 | 10449 | ENSG00000167315 |

  
  

| **Database:molecular function      &nbspName:oxidoreductase activity, acting on the aldehyde or oxo group of donors      &nbspID:GO:0016903** | | | | | | |
| --- | --- | --- | --- | --- | --- | --- |
| C=40; O=5; E=0.28; R=18.00; rawP=7.95e-06; adjP=0.0001 | | | | | | |
| Index | UserID | Value | Gene Symbol | Gene Name | EntrezGene | Ensembl |
| 1 | P49419 | NA | ALDH7A1 | aldehyde dehydrogenase 7 family, member A1 | 501 | ENSG00000164904 |
| 2 | P51649 | NA | ALDH5A1 | aldehyde dehydrogenase 5 family, member A1 | 7915 | ENSG00000112294 |
| 3 | P00352 | NA | ALDH1A1 | aldehyde dehydrogenase 1 family, member A1 | 216 | ENSG00000165092 |
| 4 | P21953 | NA | BCKDHB | branched chain keto acid dehydrogenase E1, beta polypeptide | 594 | ENSG00000083123 |
| 5 | P47895 | NA | ALDH1A3 | aldehyde dehydrogenase 1 family, member A3 | 220 | ENSG00000184254 |

  
  

| **Database:molecular function      &nbspName:fatty-acyl-CoA binding      &nbspID:GO:0000062** | | | | | | |
| --- | --- | --- | --- | --- | --- | --- |
| C=21; O=4; E=0.15; R=27.43; rawP=1.20e-05; adjP=0.0002 | | | | | | |
| Index | UserID | Value | Gene Symbol | Gene Name | EntrezGene | Ensembl |
| 1 | O75521 | NA | ECI2 | enoyl-CoA delta isomerase 2 | 10455 | ENSG00000198721 |
| 2 | P49748 | NA | ACADVL | acyl-CoA dehydrogenase, very long chain | 37 | ENSG00000072778 |
| 3 | P40939 | NA | HADHA | hydroxyacyl-CoA dehydrogenase/3-ketoacyl-CoA thiolase/enoyl-CoA hydratase (trifunctional protein), alpha subunit | 3030 | ENSG00000084754 |
| 4 | P07108 | NA | DBI | diazepam binding inhibitor (GABA receptor modulator, acyl-CoA binding protein) | 1622 | ENSG00000155368 |

  
  

| **Database:molecular function      &nbspName:translation factor activity, nucleic acid binding      &nbspID:GO:0008135** | | | | | | |
| --- | --- | --- | --- | --- | --- | --- |
| C=79; O=6; E=0.55; R=10.94; rawP=1.79e-05; adjP=0.0002 | | | | | | |
| Index | UserID | Value | Gene Symbol | Gene Name | EntrezGene | Ensembl |
| 1 | P60842 | NA | EIF4A1 | eukaryotic translation initiation factor 4A1 | 1973 | ENSG00000161960 |
| 2 | P13639 | NA | EEF2 | eukaryotic translation elongation factor 2 | 1938 | ENSG00000167658 |
| 3 | P41091 | NA | EIF2S3 | eukaryotic translation initiation factor 2, subunit 3 gamma, 52kDa | 1968 | ENSG00000130741 |
| 4 | P68104 | NA | EEF1A1 | eukaryotic translation elongation factor 1 alpha 1 | 1915 | ENSG00000156508 |
| 5 | P29692 | NA | EEF1D | eukaryotic translation elongation factor 1 delta (guanine nucleotide exchange protein) | 1936 | ENSG00000104529 |
| 6 | P24534 | NA | EEF1B2 | eukaryotic translation elongation factor 1 beta 2 | 1933 | ENSG00000114942 |

  
  

| **Database:molecular function      &nbspName:protein binding      &nbspID:GO:0005515** | | | | | | |
| --- | --- | --- | --- | --- | --- | --- |
| C=7301; O=72; E=50.70; R=1.42; rawP=1.77e-05; adjP=0.0002 | | | | | | |
| Index | UserID | Value | Gene Symbol | Gene Name | EntrezGene | Ensembl |
| 1 | P68371 | NA | TUBB4B | tubulin, beta 4B class IVb | 10383 | ENSG00000188229 |
| 2 | P40926 | NA | MDH2 | malate dehydrogenase 2, NAD (mitochondrial) | 4191 | ENSG00000146701 |
| 3 | P04179 | NA | SOD2 | superoxide dismutase 2, mitochondrial | 6648 | ENSG00000112096 |
| 4 | P00367 | NA | GLUD1 | glutamate dehydrogenase 1 | 2746 | ENSG00000148672 |
| 5 | P35580 | NA | MYH10 | myosin, heavy chain 10, non-muscle | 4628 | ENSG00000133026 |
| 6 | P31040 | NA | SDHA | succinate dehydrogenase complex, subunit A, flavoprotein (Fp) | 6389 | ENSG00000073578 |
| 7 | O75369 | NA | FLNB | filamin B, beta | 2317 | ENSG00000136068 |
| 8 | P24534 | NA | EEF1B2 | eukaryotic translation elongation factor 1 beta 2 | 1933 | ENSG00000114942 |
| 9 | O14773 | NA | TPP1 | tripeptidyl peptidase I | 1200 | ENSG00000166340 |
| 10 | P47895 | NA | ALDH1A3 | aldehyde dehydrogenase 1 family, member A3 | 220 | ENSG00000184254 |
| 11 | P40121 | NA | CAPG | capping protein (actin filament), gelsolin-like | 822 | ENSG00000042493 |
| 12 | P42765 | NA | ACAA2 | acetyl-CoA acyltransferase 2 | 10449 | ENSG00000167315 |
| 13 | Q04837 | NA | SSBP1 | single-stranded DNA binding protein 1, mitochondrial | 6742 | ENSG00000106028 |
| 14 | P08727 | NA | KRT19 | keratin 19 | 3880 | ENSG00000171345 |
| 15 | P50454 | NA | SERPINH1 | serpin peptidase inhibitor, clade H (heat shock protein 47), member 1, (collagen binding protein 1) | 871 | ENSG00000149257 |
| 16 | P18206 | NA | VCL | vinculin | 7414 | ENSG00000035403 |
| 17 | P08195 | NA | SLC3A2 | solute carrier family 3 (activators of dibasic and neutral amino acid transport), member 2 | 6520 | ENSG00000168003 |
| 18 | P40939 | NA | HADHA | hydroxyacyl-CoA dehydrogenase/3-ketoacyl-CoA thiolase/enoyl-CoA hydratase (trifunctional protein), alpha subunit | 3030 | ENSG00000084754 |
| 19 | P09382 | NA | LGALS1 | lectin, galactoside-binding, soluble, 1 | 3956 | ENSG00000100097 |
| 20 | P52815 | NA | MRPL12 | mitochondrial ribosomal protein L12 | 6182 | ENSG00000262814 |
| 21 | P21796 | NA | VDAC1 | voltage-dependent anion channel 1 | 7416 | ENSG00000213585 |
| 22 | P04080 | NA | CSTB | cystatin B (stefin B) | 1476 | ENSG00000160213 |
| 23 | P32322 | NA | PYCR1 | pyrroline-5-carboxylate reductase 1 | 5831 | ENSG00000183010 |
| 24 | Q9UJZ1 | NA | STOML2 | stomatin (EPB72)-like 2 | 30968 | ENSG00000165283 |
| 25 | Q14980 | NA | NUMA1 | nuclear mitotic apparatus protein 1 | 4926 | ENSG00000137497 |
| 26 | P51649 | NA | ALDH5A1 | aldehyde dehydrogenase 5 family, member A1 | 7915 | ENSG00000112294 |
| 27 | P12235 | NA | SLC25A4 | solute carrier family 25 (mitochondrial carrier; adenine nucleotide translocator), member 4 | 291 | ENSG00000151729 |
| 28 | P27816 | NA | MAP4 | microtubule-associated protein 4 | 4134 | ENSG00000047849 |
| 29 | P05556 | NA | ITGB1 | integrin, beta 1 (fibronectin receptor, beta polypeptide, antigen CD29 includes MDF2, MSK12) | 3688 | ENSG00000150093 |
| 30 | Q6PI48 | NA | DARS2 | aspartyl-tRNA synthetase 2, mitochondrial | 55157 | ENSG00000117593 |
| 31 | P40227 | NA | CCT6A | chaperonin containing TCP1, subunit 6A (zeta 1) | 908 | ENSG00000146731 |
| 32 | P49589 | NA | CARS | cysteinyl-tRNA synthetase | 833 | ENSG00000110619 |
| 33 | P17931 | NA | LGALS3 | lectin, galactoside-binding, soluble, 3 | 3958 | ENSG00000131981 |
| 34 | P05787 | NA | KRT8 | keratin 8 | 3856 | ENSG00000170421 |
| 35 | Q99623 | NA | PHB2 | prohibitin 2 | 11331 | ENSG00000215021 |
| 36 | P50453 | NA | SERPINB9 | serpin peptidase inhibitor, clade B (ovalbumin), member 9 | 5272 | ENSG00000170542 |
| 37 | P35232 | NA | PHB | prohibitin | 5245 | ENSG00000167085 |
| 38 | Q99714 | NA | HSD17B10 | hydroxysteroid (17-beta) dehydrogenase 10 | 3028 | ENSG00000072506 |
| 39 | O95831 | NA | AIFM1 | apoptosis-inducing factor, mitochondrion-associated, 1 | 9131 | ENSG00000156709 |
| 40 | P61313 | NA | RPL15 | ribosomal protein L15 | 6138 | ENSG00000174748 |
| 41 | O60313 | NA | OPA1 | optic atrophy 1 (autosomal dominant) | 4976 | ENSG00000198836 |
| 42 | P68104 | NA | EEF1A1 | eukaryotic translation elongation factor 1 alpha 1 | 1915 | ENSG00000156508 |
| 43 | P02786 | NA | TFRC | transferrin receptor (p90, CD71) | 7037 | ENSG00000072274 |
| 44 | P43490 | NA | NAMPT | nicotinamide phosphoribosyltransferase | 10135 | ENSG00000105835 |
| 45 | O75367 | NA | H2AFY | H2A histone family, member Y | 9555 | ENSG00000113648 |
| 46 | O75521 | NA | ECI2 | enoyl-CoA delta isomerase 2 | 10455 | ENSG00000198721 |
| 47 | P67809 | NA | YBX1 | Y box binding protein 1 | 4904 | ENSG00000065978 |
| 48 | P49419 | NA | ALDH7A1 | aldehyde dehydrogenase 7 family, member A1 | 501 | ENSG00000164904 |
| 49 | Q14764 | NA | MVP | major vault protein | 9961 | ENSG00000013364 |
| 50 | P29692 | NA | EEF1D | eukaryotic translation elongation factor 1 delta (guanine nucleotide exchange protein) | 1936 | ENSG00000104529 |
| 51 | P11387 | NA | TOP1 | topoisomerase (DNA) I | 7150 | ENSG00000198900 |
| 52 | Q13813 | NA | SPTAN1 | spectrin, alpha, non-erythrocytic 1 | 6709 | ENSG00000197694 |
| 53 | P02792 | NA | FTL | ferritin, light polypeptide | 2512 | ENSG00000087086 |
| 54 | P21953 | NA | BCKDHB | branched chain keto acid dehydrogenase E1, beta polypeptide | 594 | ENSG00000083123 |
| 55 | P30084 | NA | ECHS1 | enoyl CoA hydratase, short chain, 1, mitochondrial | 1892 | ENSG00000127884 |
| 56 | P60842 | NA | EIF4A1 | eukaryotic translation initiation factor 4A1 | 1973 | ENSG00000161960 |
| 57 | Q13011 | NA | ECH1 | enoyl CoA hydratase 1, peroxisomal | 1891 | ENSG00000104823 |
| 58 | P04083 | NA | ANXA1 | annexin A1 | 301 | ENSG00000135046 |
| 59 | P04040 | NA | CAT | catalase | 847 | ENSG00000121691 |
| 60 | O75340 | NA | PDCD6 | programmed cell death 6 | 10016 | ENSG00000249915 |
| 61 | P02768 | NA | ALB | albumin | 213 | ENSG00000163631 |
| 62 | P41091 | NA | EIF2S3 | eukaryotic translation initiation factor 2, subunit 3 gamma, 52kDa | 1968 | ENSG00000130741 |
| 63 | P30048 | NA | PRDX3 | peroxiredoxin 3 | 10935 | ENSG00000165672 |
| 64 | P07108 | NA | DBI | diazepam binding inhibitor (GABA receptor modulator, acyl-CoA binding protein) | 1622 | ENSG00000155368 |
| 65 | P24752 | NA | ACAT1 | acetyl-CoA acetyltransferase 1 | 38 | ENSG00000075239 |
| 66 | P19367 | NA | HK1 | hexokinase 1 | 3098 | ENSG00000156515 |
| 67 | P25705 | NA | ATP5A1 | ATP synthase, H+ transporting, mitochondrial F1 complex, alpha subunit 1, cardiac muscle | 498 | ENSG00000152234 |
| 68 | Q9BQ69 | NA | MACROD1 | MACRO domain containing 1 | 28992 | ENSG00000133315 |
| 69 | P06865 | NA | HEXA | hexosaminidase A (alpha polypeptide) | 3073 | ENSG00000213614 |
| 70 | P05783 | NA | KRT18 | keratin 18 | 3875 | ENSG00000111057 |
| 71 | P48047 | NA | ATP5O | ATP synthase, H+ transporting, mitochondrial F1 complex, O subunit | 539 | ENSG00000241837 |
| 72 | Q16531 | NA | DDB1 | damage-specific DNA binding protein 1, 127kDa | 1642 | ENSG00000167986 |

  
  

| **Database:molecular function      &nbspName:nucleotide binding      &nbspID:GO:0000166** | | | | | | |
| --- | --- | --- | --- | --- | --- | --- |
| C=2403; O=34; E=16.69; R=2.04; rawP=1.99e-05; adjP=0.0002 | | | | | | |
| Index | UserID | Value | Gene Symbol | Gene Name | EntrezGene | Ensembl |
| 1 | P49589 | NA | CARS | cysteinyl-tRNA synthetase | 833 | ENSG00000110619 |
| 2 | P54819 | NA | AK2 | adenylate kinase 2 | 204 | ENSG00000004455 |
| 3 | P68371 | NA | TUBB4B | tubulin, beta 4B class IVb | 10383 | ENSG00000188229 |
| 4 | P40926 | NA | MDH2 | malate dehydrogenase 2, NAD (mitochondrial) | 4191 | ENSG00000146701 |
| 5 | O95831 | NA | AIFM1 | apoptosis-inducing factor, mitochondrion-associated, 1 | 9131 | ENSG00000156709 |
| 6 | Q99714 | NA | HSD17B10 | hydroxysteroid (17-beta) dehydrogenase 10 | 3028 | ENSG00000072506 |
| 7 | P07741 | NA | APRT | adenine phosphoribosyltransferase | 353 | ENSG00000198931 |
| 8 | P00367 | NA | GLUD1 | glutamate dehydrogenase 1 | 2746 | ENSG00000148672 |
| 9 | P48735 | NA | IDH2 | isocitrate dehydrogenase 2 (NADP+), mitochondrial | 3418 | ENSG00000182054 |
| 10 | O60313 | NA | OPA1 | optic atrophy 1 (autosomal dominant) | 4976 | ENSG00000198836 |
| 11 | P35270 | NA | SPR | sepiapterin reductase (7,8-dihydrobiopterin:NADP+ oxidoreductase) | 6697 | ENSG00000116096 |
| 12 | P35580 | NA | MYH10 | myosin, heavy chain 10, non-muscle | 4628 | ENSG00000133026 |
| 13 | P68104 | NA | EEF1A1 | eukaryotic translation elongation factor 1 alpha 1 | 1915 | ENSG00000156508 |
| 14 | P31040 | NA | SDHA | succinate dehydrogenase complex, subunit A, flavoprotein (Fp) | 6389 | ENSG00000073578 |
| 15 | P22102 | NA | GART | phosphoribosylglycinamide formyltransferase, phosphoribosylglycinamide synthetase, phosphoribosylaminoimidazole synthetase | 2618 | ENSG00000159131 |
| 16 | P13639 | NA | EEF2 | eukaryotic translation elongation factor 2 | 1938 | ENSG00000167658 |
| 17 | P47895 | NA | ALDH1A3 | aldehyde dehydrogenase 1 family, member A3 | 220 | ENSG00000184254 |
| 18 | P13804 | NA | ETFA | electron-transfer-flavoprotein, alpha polypeptide | 2108 | ENSG00000140374 |
| 19 | Q9Y277 | NA | VDAC3 | voltage-dependent anion channel 3 | 7419 | ENSG00000078668 |
| 20 | P11387 | NA | TOP1 | topoisomerase (DNA) I | 7150 | ENSG00000198900 |
| 21 | Q9NR30 | NA | DDX21 | DEAD (Asp-Glu-Ala-Asp) box helicase 21 | 9188 | ENSG00000165732 |
| 22 | P40939 | NA | HADHA | hydroxyacyl-CoA dehydrogenase/3-ketoacyl-CoA thiolase/enoyl-CoA hydratase (trifunctional protein), alpha subunit | 3030 | ENSG00000084754 |
| 23 | P60842 | NA | EIF4A1 | eukaryotic translation initiation factor 4A1 | 1973 | ENSG00000161960 |
| 24 | P32322 | NA | PYCR1 | pyrroline-5-carboxylate reductase 1 | 5831 | ENSG00000183010 |
| 25 | P04040 | NA | CAT | catalase | 847 | ENSG00000121691 |
| 26 | P41091 | NA | EIF2S3 | eukaryotic translation initiation factor 2, subunit 3 gamma, 52kDa | 1968 | ENSG00000130741 |
| 27 | P00558 | NA | PGK1 | phosphoglycerate kinase 1 | 5230 | ENSG00000102144 |
| 28 | Q6PI48 | NA | DARS2 | aspartyl-tRNA synthetase 2, mitochondrial | 55157 | ENSG00000117593 |
| 29 | Q9NSE4 | NA | IARS2 | isoleucyl-tRNA synthetase 2, mitochondrial | 55699 | ENSG00000067704 |
| 30 | P40227 | NA | CCT6A | chaperonin containing TCP1, subunit 6A (zeta 1) | 908 | ENSG00000146731 |
| 31 | P25705 | NA | ATP5A1 | ATP synthase, H+ transporting, mitochondrial F1 complex, alpha subunit 1, cardiac muscle | 498 | ENSG00000152234 |
| 32 | P19367 | NA | HK1 | hexokinase 1 | 3098 | ENSG00000156515 |
| 33 | P49748 | NA | ACADVL | acyl-CoA dehydrogenase, very long chain | 37 | ENSG00000072778 |
| 34 | P31937 | NA | HIBADH | 3-hydroxyisobutyrate dehydrogenase | 11112 | ENSG00000106049 |

  
  

| **Database:molecular function      &nbspName:nucleoside phosphate binding      &nbspID:GO:1901265** | | | | | | |
| --- | --- | --- | --- | --- | --- | --- |
| C=2404; O=34; E=16.69; R=2.04; rawP=2.01e-05; adjP=0.0002 | | | | | | |
| Index | UserID | Value | Gene Symbol | Gene Name | EntrezGene | Ensembl |
| 1 | P49589 | NA | CARS | cysteinyl-tRNA synthetase | 833 | ENSG00000110619 |
| 2 | P54819 | NA | AK2 | adenylate kinase 2 | 204 | ENSG00000004455 |
| 3 | P68371 | NA | TUBB4B | tubulin, beta 4B class IVb | 10383 | ENSG00000188229 |
| 4 | P40926 | NA | MDH2 | malate dehydrogenase 2, NAD (mitochondrial) | 4191 | ENSG00000146701 |
| 5 | O95831 | NA | AIFM1 | apoptosis-inducing factor, mitochondrion-associated, 1 | 9131 | ENSG00000156709 |
| 6 | Q99714 | NA | HSD17B10 | hydroxysteroid (17-beta) dehydrogenase 10 | 3028 | ENSG00000072506 |
| 7 | P07741 | NA | APRT | adenine phosphoribosyltransferase | 353 | ENSG00000198931 |
| 8 | P00367 | NA | GLUD1 | glutamate dehydrogenase 1 | 2746 | ENSG00000148672 |
| 9 | P48735 | NA | IDH2 | isocitrate dehydrogenase 2 (NADP+), mitochondrial | 3418 | ENSG00000182054 |
| 10 | O60313 | NA | OPA1 | optic atrophy 1 (autosomal dominant) | 4976 | ENSG00000198836 |
| 11 | P35270 | NA | SPR | sepiapterin reductase (7,8-dihydrobiopterin:NADP+ oxidoreductase) | 6697 | ENSG00000116096 |
| 12 | P35580 | NA | MYH10 | myosin, heavy chain 10, non-muscle | 4628 | ENSG00000133026 |
| 13 | P68104 | NA | EEF1A1 | eukaryotic translation elongation factor 1 alpha 1 | 1915 | ENSG00000156508 |
| 14 | P31040 | NA | SDHA | succinate dehydrogenase complex, subunit A, flavoprotein (Fp) | 6389 | ENSG00000073578 |
| 15 | P22102 | NA | GART | phosphoribosylglycinamide formyltransferase, phosphoribosylglycinamide synthetase, phosphoribosylaminoimidazole synthetase | 2618 | ENSG00000159131 |
| 16 | P13639 | NA | EEF2 | eukaryotic translation elongation factor 2 | 1938 | ENSG00000167658 |
| 17 | P47895 | NA | ALDH1A3 | aldehyde dehydrogenase 1 family, member A3 | 220 | ENSG00000184254 |
| 18 | P13804 | NA | ETFA | electron-transfer-flavoprotein, alpha polypeptide | 2108 | ENSG00000140374 |
| 19 | Q9Y277 | NA | VDAC3 | voltage-dependent anion channel 3 | 7419 | ENSG00000078668 |
| 20 | P11387 | NA | TOP1 | topoisomerase (DNA) I | 7150 | ENSG00000198900 |
| 21 | Q9NR30 | NA | DDX21 | DEAD (Asp-Glu-Ala-Asp) box helicase 21 | 9188 | ENSG00000165732 |
| 22 | P40939 | NA | HADHA | hydroxyacyl-CoA dehydrogenase/3-ketoacyl-CoA thiolase/enoyl-CoA hydratase (trifunctional protein), alpha subunit | 3030 | ENSG00000084754 |
| 23 | P60842 | NA | EIF4A1 | eukaryotic translation initiation factor 4A1 | 1973 | ENSG00000161960 |
| 24 | P32322 | NA | PYCR1 | pyrroline-5-carboxylate reductase 1 | 5831 | ENSG00000183010 |
| 25 | P04040 | NA | CAT | catalase | 847 | ENSG00000121691 |
| 26 | P41091 | NA | EIF2S3 | eukaryotic translation initiation factor 2, subunit 3 gamma, 52kDa | 1968 | ENSG00000130741 |
| 27 | P00558 | NA | PGK1 | phosphoglycerate kinase 1 | 5230 | ENSG00000102144 |
| 28 | Q6PI48 | NA | DARS2 | aspartyl-tRNA synthetase 2, mitochondrial | 55157 | ENSG00000117593 |
| 29 | Q9NSE4 | NA | IARS2 | isoleucyl-tRNA synthetase 2, mitochondrial | 55699 | ENSG00000067704 |
| 30 | P40227 | NA | CCT6A | chaperonin containing TCP1, subunit 6A (zeta 1) | 908 | ENSG00000146731 |
| 31 | P25705 | NA | ATP5A1 | ATP synthase, H+ transporting, mitochondrial F1 complex, alpha subunit 1, cardiac muscle | 498 | ENSG00000152234 |
| 32 | P19367 | NA | HK1 | hexokinase 1 | 3098 | ENSG00000156515 |
| 33 | P49748 | NA | ACADVL | acyl-CoA dehydrogenase, very long chain | 37 | ENSG00000072778 |
| 34 | P31937 | NA | HIBADH | 3-hydroxyisobutyrate dehydrogenase | 11112 | ENSG00000106049 |

  
  

| **Database:molecular function      &nbspName:aldehyde dehydrogenase (NAD) activity      &nbspID:GO:0004029** | | | | | | |
| --- | --- | --- | --- | --- | --- | --- |
| C=10; O=3; E=0.07; R=43.20; rawP=3.77e-05; adjP=0.0003 | | | | | | |
| Index | UserID | Value | Gene Symbol | Gene Name | EntrezGene | Ensembl |
| 1 | P49419 | NA | ALDH7A1 | aldehyde dehydrogenase 7 family, member A1 | 501 | ENSG00000164904 |
| 2 | P00352 | NA | ALDH1A1 | aldehyde dehydrogenase 1 family, member A1 | 216 | ENSG00000165092 |
| 3 | P47895 | NA | ALDH1A3 | aldehyde dehydrogenase 1 family, member A3 | 220 | ENSG00000184254 |

  
  

| **Database:molecular function      &nbspName:glucosidase activity      &nbspID:GO:0015926** | | | | | | |
| --- | --- | --- | --- | --- | --- | --- |
| C=10; O=3; E=0.07; R=43.20; rawP=3.77e-05; adjP=0.0003 | | | | | | |
| Index | UserID | Value | Gene Symbol | Gene Name | EntrezGene | Ensembl |
| 1 | P10253 | NA | GAA | glucosidase, alpha; acid | 2548 | ENSG00000171298 |
| 2 | Q14697 | NA | GANAB | glucosidase, alpha; neutral AB | 23193 | ENSG00000089597 |
| 3 | Q13724 | NA | MOGS | mannosyl-oligosaccharide glucosidase | 7841 | ENSG00000115275 |

  
  

| **Database:molecular function      &nbspName:oxidoreductase activity, acting on the aldehyde or oxo group of donors, NAD or NADP as acceptor      &nbspID:GO:0016620** | | | | | | |
| --- | --- | --- | --- | --- | --- | --- |
| C=29; O=4; E=0.20; R=19.86; rawP=4.56e-05; adjP=0.0003 | | | | | | |
| Index | UserID | Value | Gene Symbol | Gene Name | EntrezGene | Ensembl |
| 1 | P49419 | NA | ALDH7A1 | aldehyde dehydrogenase 7 family, member A1 | 501 | ENSG00000164904 |
| 2 | P51649 | NA | ALDH5A1 | aldehyde dehydrogenase 5 family, member A1 | 7915 | ENSG00000112294 |
| 3 | P00352 | NA | ALDH1A1 | aldehyde dehydrogenase 1 family, member A1 | 216 | ENSG00000165092 |
| 4 | P47895 | NA | ALDH1A3 | aldehyde dehydrogenase 1 family, member A3 | 220 | ENSG00000184254 |

  
  

| **Database:molecular function      &nbspName:C-acyltransferase activity      &nbspID:GO:0016408** | | | | | | |
| --- | --- | --- | --- | --- | --- | --- |
| C=15; O=3; E=0.10; R=28.80; rawP=0.0001; adjP=0.0006 | | | | | | |
| Index | UserID | Value | Gene Symbol | Gene Name | EntrezGene | Ensembl |
| 1 | P24752 | NA | ACAT1 | acetyl-CoA acetyltransferase 1 | 38 | ENSG00000075239 |
| 2 | P40939 | NA | HADHA | hydroxyacyl-CoA dehydrogenase/3-ketoacyl-CoA thiolase/enoyl-CoA hydratase (trifunctional protein), alpha subunit | 3030 | ENSG00000084754 |
| 3 | P42765 | NA | ACAA2 | acetyl-CoA acyltransferase 2 | 10449 | ENSG00000167315 |

  
  

| **Database:molecular function      &nbspName:binding      &nbspID:GO:0005488** | | | | | | |
| --- | --- | --- | --- | --- | --- | --- |
| C=11779; O=96; E=81.79; R=1.17; rawP=0.0001; adjP=0.0006 | | | | | | |
| Index | UserID | Value | Gene Symbol | Gene Name | EntrezGene | Ensembl |
| 1 | P10253 | NA | GAA | glucosidase, alpha; acid | 2548 | ENSG00000171298 |
| 2 | P68371 | NA | TUBB4B | tubulin, beta 4B class IVb | 10383 | ENSG00000188229 |
| 3 | P40926 | NA | MDH2 | malate dehydrogenase 2, NAD (mitochondrial) | 4191 | ENSG00000146701 |
| 4 | P04179 | NA | SOD2 | superoxide dismutase 2, mitochondrial | 6648 | ENSG00000112096 |
| 5 | P00367 | NA | GLUD1 | glutamate dehydrogenase 1 | 2746 | ENSG00000148672 |
| 6 | P48735 | NA | IDH2 | isocitrate dehydrogenase 2 (NADP+), mitochondrial | 3418 | ENSG00000182054 |
| 7 | P53701 | NA | HCCS | holocytochrome c synthase | 3052 | ENSG00000004961 |
| 8 | P35580 | NA | MYH10 | myosin, heavy chain 10, non-muscle | 4628 | ENSG00000133026 |
| 9 | P31040 | NA | SDHA | succinate dehydrogenase complex, subunit A, flavoprotein (Fp) | 6389 | ENSG00000073578 |
| 10 | O75369 | NA | FLNB | filamin B, beta | 2317 | ENSG00000136068 |
| 11 | P24534 | NA | EEF1B2 | eukaryotic translation elongation factor 1 beta 2 | 1933 | ENSG00000114942 |
| 12 | O14773 | NA | TPP1 | tripeptidyl peptidase I | 1200 | ENSG00000166340 |
| 13 | P47895 | NA | ALDH1A3 | aldehyde dehydrogenase 1 family, member A3 | 220 | ENSG00000184254 |
| 14 | P13804 | NA | ETFA | electron-transfer-flavoprotein, alpha polypeptide | 2108 | ENSG00000140374 |
| 15 | P40121 | NA | CAPG | capping protein (actin filament), gelsolin-like | 822 | ENSG00000042493 |
| 16 | P42765 | NA | ACAA2 | acetyl-CoA acyltransferase 2 | 10449 | ENSG00000167315 |
| 17 | Q9Y277 | NA | VDAC3 | voltage-dependent anion channel 3 | 7419 | ENSG00000078668 |
| 18 | P17174 | NA | GOT1 | glutamic-oxaloacetic transaminase 1, soluble (aspartate aminotransferase 1) | 2805 | ENSG00000120053 |
| 19 | P08727 | NA | KRT19 | keratin 19 | 3880 | ENSG00000171345 |
| 20 | Q04837 | NA | SSBP1 | single-stranded DNA binding protein 1, mitochondrial | 6742 | ENSG00000106028 |
| 21 | P50454 | NA | SERPINH1 | serpin peptidase inhibitor, clade H (heat shock protein 47), member 1, (collagen binding protein 1) | 871 | ENSG00000149257 |
| 22 | Q14697 | NA | GANAB | glucosidase, alpha; neutral AB | 23193 | ENSG00000089597 |
| 23 | P18206 | NA | VCL | vinculin | 7414 | ENSG00000035403 |
| 24 | P08195 | NA | SLC3A2 | solute carrier family 3 (activators of dibasic and neutral amino acid transport), member 2 | 6520 | ENSG00000168003 |
| 25 | P40939 | NA | HADHA | hydroxyacyl-CoA dehydrogenase/3-ketoacyl-CoA thiolase/enoyl-CoA hydratase (trifunctional protein), alpha subunit | 3030 | ENSG00000084754 |
| 26 | P09382 | NA | LGALS1 | lectin, galactoside-binding, soluble, 1 | 3956 | ENSG00000100097 |
| 27 | P52815 | NA | MRPL12 | mitochondrial ribosomal protein L12 | 6182 | ENSG00000262814 |
| 28 | P21796 | NA | VDAC1 | voltage-dependent anion channel 1 | 7416 | ENSG00000213585 |
| 29 | P04080 | NA | CSTB | cystatin B (stefin B) | 1476 | ENSG00000160213 |
| 30 | P32322 | NA | PYCR1 | pyrroline-5-carboxylate reductase 1 | 5831 | ENSG00000183010 |
| 31 | Q9UJZ1 | NA | STOML2 | stomatin (EPB72)-like 2 | 30968 | ENSG00000165283 |
| 32 | Q14980 | NA | NUMA1 | nuclear mitotic apparatus protein 1 | 4926 | ENSG00000137497 |
| 33 | Q99798 | NA | ACO2 | aconitase 2, mitochondrial | 50 | ENSG00000100412 |
| 34 | P51649 | NA | ALDH5A1 | aldehyde dehydrogenase 5 family, member A1 | 7915 | ENSG00000112294 |
| 35 | P12235 | NA | SLC25A4 | solute carrier family 25 (mitochondrial carrier; adenine nucleotide translocator), member 4 | 291 | ENSG00000151729 |
| 36 | P27816 | NA | MAP4 | microtubule-associated protein 4 | 4134 | ENSG00000047849 |
| 37 | Q6PI48 | NA | DARS2 | aspartyl-tRNA synthetase 2, mitochondrial | 55157 | ENSG00000117593 |
| 38 | Q9NSE4 | NA | IARS2 | isoleucyl-tRNA synthetase 2, mitochondrial | 55699 | ENSG00000067704 |
| 39 | P05556 | NA | ITGB1 | integrin, beta 1 (fibronectin receptor, beta polypeptide, antigen CD29 includes MDF2, MSK12) | 3688 | ENSG00000150093 |
| 40 | P13674 | NA | P4HA1 | prolyl 4-hydroxylase, alpha polypeptide I | 5033 | ENSG00000122884 |
| 41 | P40227 | NA | CCT6A | chaperonin containing TCP1, subunit 6A (zeta 1) | 908 | ENSG00000146731 |
| 42 | P49589 | NA | CARS | cysteinyl-tRNA synthetase | 833 | ENSG00000110619 |
| 43 | P54819 | NA | AK2 | adenylate kinase 2 | 204 | ENSG00000004455 |
| 44 | P17931 | NA | LGALS3 | lectin, galactoside-binding, soluble, 3 | 3958 | ENSG00000131981 |
| 45 | P05787 | NA | KRT8 | keratin 8 | 3856 | ENSG00000170421 |
| 46 | Q99623 | NA | PHB2 | prohibitin 2 | 11331 | ENSG00000215021 |
| 47 | P50453 | NA | SERPINB9 | serpin peptidase inhibitor, clade B (ovalbumin), member 9 | 5272 | ENSG00000170542 |
| 48 | P35232 | NA | PHB | prohibitin | 5245 | ENSG00000167085 |
| 49 | Q99714 | NA | HSD17B10 | hydroxysteroid (17-beta) dehydrogenase 10 | 3028 | ENSG00000072506 |
| 50 | O95831 | NA | AIFM1 | apoptosis-inducing factor, mitochondrion-associated, 1 | 9131 | ENSG00000156709 |
| 51 | P61313 | NA | RPL15 | ribosomal protein L15 | 6138 | ENSG00000174748 |
| 52 | P07741 | NA | APRT | adenine phosphoribosyltransferase | 353 | ENSG00000198931 |
| 53 | O60313 | NA | OPA1 | optic atrophy 1 (autosomal dominant) | 4976 | ENSG00000198836 |
| 54 | P35270 | NA | SPR | sepiapterin reductase (7,8-dihydrobiopterin:NADP+ oxidoreductase) | 6697 | ENSG00000116096 |
| 55 | P68104 | NA | EEF1A1 | eukaryotic translation elongation factor 1 alpha 1 | 1915 | ENSG00000156508 |
| 56 | P02786 | NA | TFRC | transferrin receptor (p90, CD71) | 7037 | ENSG00000072274 |
| 57 | P43490 | NA | NAMPT | nicotinamide phosphoribosyltransferase | 10135 | ENSG00000105835 |
| 58 | P22102 | NA | GART | phosphoribosylglycinamide formyltransferase, phosphoribosylglycinamide synthetase, phosphoribosylaminoimidazole synthetase | 2618 | ENSG00000159131 |
| 59 | P13639 | NA | EEF2 | eukaryotic translation elongation factor 2 | 1938 | ENSG00000167658 |
| 60 | O75367 | NA | H2AFY | H2A histone family, member Y | 9555 | ENSG00000113648 |
| 61 | O75521 | NA | ECI2 | enoyl-CoA delta isomerase 2 | 10455 | ENSG00000198721 |
| 62 | P67809 | NA | YBX1 | Y box binding protein 1 | 4904 | ENSG00000065978 |
| 63 | P49419 | NA | ALDH7A1 | aldehyde dehydrogenase 7 family, member A1 | 501 | ENSG00000164904 |
| 64 | Q14764 | NA | MVP | major vault protein | 9961 | ENSG00000013364 |
| 65 | P29692 | NA | EEF1D | eukaryotic translation elongation factor 1 delta (guanine nucleotide exchange protein) | 1936 | ENSG00000104529 |
| 66 | Q13813 | NA | SPTAN1 | spectrin, alpha, non-erythrocytic 1 | 6709 | ENSG00000197694 |
| 67 | Q5SSJ5 | NA | HP1BP3 | heterochromatin protein 1, binding protein 3 | 50809 | ENSG00000127483 |
| 68 | P11387 | NA | TOP1 | topoisomerase (DNA) I | 7150 | ENSG00000198900 |
| 69 | P02792 | NA | FTL | ferritin, light polypeptide | 2512 | ENSG00000087086 |
| 70 | O75439 | NA | PMPCB | peptidase (mitochondrial processing) beta | 9512 | ENSG00000105819 |
| 71 | Q9NR30 | NA | DDX21 | DEAD (Asp-Glu-Ala-Asp) box helicase 21 | 9188 | ENSG00000165732 |
| 72 | P00491 | NA | PNP | purine nucleoside phosphorylase | 4860 | ENSG00000198805 |
| 73 | P21953 | NA | BCKDHB | branched chain keto acid dehydrogenase E1, beta polypeptide | 594 | ENSG00000083123 |
| 74 | P30084 | NA | ECHS1 | enoyl CoA hydratase, short chain, 1, mitochondrial | 1892 | ENSG00000127884 |
| 75 | P60842 | NA | EIF4A1 | eukaryotic translation initiation factor 4A1 | 1973 | ENSG00000161960 |
| 76 | Q13011 | NA | ECH1 | enoyl CoA hydratase 1, peroxisomal | 1891 | ENSG00000104823 |
| 77 | P04083 | NA | ANXA1 | annexin A1 | 301 | ENSG00000135046 |
| 78 | P04040 | NA | CAT | catalase | 847 | ENSG00000121691 |
| 79 | O75340 | NA | PDCD6 | programmed cell death 6 | 10016 | ENSG00000249915 |
| 80 | P51688 | NA | SGSH | N-sulfoglucosamine sulfohydrolase | 6448 | ENSG00000181523 |
| 81 | P00558 | NA | PGK1 | phosphoglycerate kinase 1 | 5230 | ENSG00000102144 |
| 82 | P30048 | NA | PRDX3 | peroxiredoxin 3 | 10935 | ENSG00000165672 |
| 83 | P41091 | NA | EIF2S3 | eukaryotic translation initiation factor 2, subunit 3 gamma, 52kDa | 1968 | ENSG00000130741 |
| 84 | P02768 | NA | ALB | albumin | 213 | ENSG00000163631 |
| 85 | P07108 | NA | DBI | diazepam binding inhibitor (GABA receptor modulator, acyl-CoA binding protein) | 1622 | ENSG00000155368 |
| 86 | P24752 | NA | ACAT1 | acetyl-CoA acetyltransferase 1 | 38 | ENSG00000075239 |
| 87 | P19367 | NA | HK1 | hexokinase 1 | 3098 | ENSG00000156515 |
| 88 | P25705 | NA | ATP5A1 | ATP synthase, H+ transporting, mitochondrial F1 complex, alpha subunit 1, cardiac muscle | 498 | ENSG00000152234 |
| 89 | Q9BQ69 | NA | MACROD1 | MACRO domain containing 1 | 28992 | ENSG00000133315 |
| 90 | P06865 | NA | HEXA | hexosaminidase A (alpha polypeptide) | 3073 | ENSG00000213614 |
| 91 | P49748 | NA | ACADVL | acyl-CoA dehydrogenase, very long chain | 37 | ENSG00000072778 |
| 92 | P05783 | NA | KRT18 | keratin 18 | 3875 | ENSG00000111057 |
| 93 | P00352 | NA | ALDH1A1 | aldehyde dehydrogenase 1 family, member A1 | 216 | ENSG00000165092 |
| 94 | P48047 | NA | ATP5O | ATP synthase, H+ transporting, mitochondrial F1 complex, O subunit | 539 | ENSG00000241837 |
| 95 | Q16531 | NA | DDB1 | damage-specific DNA binding protein 1, 127kDa | 1642 | ENSG00000167986 |
| 96 | P31937 | NA | HIBADH | 3-hydroxyisobutyrate dehydrogenase | 11112 | ENSG00000106049 |

  
  

| **Database:molecular function      &nbspName:anion binding      &nbspID:GO:0043168** | | | | | | |
| --- | --- | --- | --- | --- | --- | --- |
| C=2371; O=32; E=16.46; R=1.94; rawP=9.91e-05; adjP=0.0006 | | | | | | |
| Index | UserID | Value | Gene Symbol | Gene Name | EntrezGene | Ensembl |
| 1 | P49589 | NA | CARS | cysteinyl-tRNA synthetase | 833 | ENSG00000110619 |
| 2 | Q9NR30 | NA | DDX21 | DEAD (Asp-Glu-Ala-Asp) box helicase 21 | 9188 | ENSG00000165732 |
| 3 | P54819 | NA | AK2 | adenylate kinase 2 | 204 | ENSG00000004455 |
| 4 | P00491 | NA | PNP | purine nucleoside phosphorylase | 4860 | ENSG00000198805 |
| 5 | P68371 | NA | TUBB4B | tubulin, beta 4B class IVb | 10383 | ENSG00000188229 |
| 6 | P40939 | NA | HADHA | hydroxyacyl-CoA dehydrogenase/3-ketoacyl-CoA thiolase/enoyl-CoA hydratase (trifunctional protein), alpha subunit | 3030 | ENSG00000084754 |
| 7 | O95831 | NA | AIFM1 | apoptosis-inducing factor, mitochondrion-associated, 1 | 9131 | ENSG00000156709 |
| 8 | P60842 | NA | EIF4A1 | eukaryotic translation initiation factor 4A1 | 1973 | ENSG00000161960 |
| 9 | P07741 | NA | APRT | adenine phosphoribosyltransferase | 353 | ENSG00000198931 |
| 10 | P00367 | NA | GLUD1 | glutamate dehydrogenase 1 | 2746 | ENSG00000148672 |
| 11 | O60313 | NA | OPA1 | optic atrophy 1 (autosomal dominant) | 4976 | ENSG00000198836 |
| 12 | P35580 | NA | MYH10 | myosin, heavy chain 10, non-muscle | 4628 | ENSG00000133026 |
| 13 | P68104 | NA | EEF1A1 | eukaryotic translation elongation factor 1 alpha 1 | 1915 | ENSG00000156508 |
| 14 | P31040 | NA | SDHA | succinate dehydrogenase complex, subunit A, flavoprotein (Fp) | 6389 | ENSG00000073578 |
| 15 | P22102 | NA | GART | phosphoribosylglycinamide formyltransferase, phosphoribosylglycinamide synthetase, phosphoribosylaminoimidazole synthetase | 2618 | ENSG00000159131 |
| 16 | P13639 | NA | EEF2 | eukaryotic translation elongation factor 2 | 1938 | ENSG00000167658 |
| 17 | P00558 | NA | PGK1 | phosphoglycerate kinase 1 | 5230 | ENSG00000102144 |
| 18 | P41091 | NA | EIF2S3 | eukaryotic translation initiation factor 2, subunit 3 gamma, 52kDa | 1968 | ENSG00000130741 |
| 19 | P02768 | NA | ALB | albumin | 213 | ENSG00000163631 |
| 20 | P07108 | NA | DBI | diazepam binding inhibitor (GABA receptor modulator, acyl-CoA binding protein) | 1622 | ENSG00000155368 |
| 21 | P47895 | NA | ALDH1A3 | aldehyde dehydrogenase 1 family, member A3 | 220 | ENSG00000184254 |
| 22 | P13804 | NA | ETFA | electron-transfer-flavoprotein, alpha polypeptide | 2108 | ENSG00000140374 |
| 23 | Q9NSE4 | NA | IARS2 | isoleucyl-tRNA synthetase 2, mitochondrial | 55699 | ENSG00000067704 |
| 24 | Q6PI48 | NA | DARS2 | aspartyl-tRNA synthetase 2, mitochondrial | 55157 | ENSG00000117593 |
| 25 | O75521 | NA | ECI2 | enoyl-CoA delta isomerase 2 | 10455 | ENSG00000198721 |
| 26 | P40227 | NA | CCT6A | chaperonin containing TCP1, subunit 6A (zeta 1) | 908 | ENSG00000146731 |
| 27 | P13674 | NA | P4HA1 | prolyl 4-hydroxylase, alpha polypeptide I | 5033 | ENSG00000122884 |
| 28 | P17174 | NA | GOT1 | glutamic-oxaloacetic transaminase 1, soluble (aspartate aminotransferase 1) | 2805 | ENSG00000120053 |
| 29 | P19367 | NA | HK1 | hexokinase 1 | 3098 | ENSG00000156515 |
| 30 | P25705 | NA | ATP5A1 | ATP synthase, H+ transporting, mitochondrial F1 complex, alpha subunit 1, cardiac muscle | 498 | ENSG00000152234 |
| 31 | P49748 | NA | ACADVL | acyl-CoA dehydrogenase, very long chain | 37 | ENSG00000072778 |
| 32 | P11387 | NA | TOP1 | topoisomerase (DNA) I | 7150 | ENSG00000198900 |

  
  

| **Database:cellular component      &nbspName:mitochondrial part      &nbspID:GO:0044429** | | | | | | |
| --- | --- | --- | --- | --- | --- | --- |
| C=740; O=42; E=4.79; R=8.77; rawP=5.23e-29; adjP=6.12e-27 | | | | | | |
| Index | UserID | Value | Gene Symbol | Gene Name | EntrezGene | Ensembl |
| 1 | P54819 | NA | AK2 | adenylate kinase 2 | 204 | ENSG00000004455 |
| 2 | P17931 | NA | LGALS3 | lectin, galactoside-binding, soluble, 3 | 3958 | ENSG00000131981 |
| 3 | Q99623 | NA | PHB2 | prohibitin 2 | 11331 | ENSG00000215021 |
| 4 | P40926 | NA | MDH2 | malate dehydrogenase 2, NAD (mitochondrial) | 4191 | ENSG00000146701 |
| 5 | P35232 | NA | PHB | prohibitin | 5245 | ENSG00000167085 |
| 6 | Q99714 | NA | HSD17B10 | hydroxysteroid (17-beta) dehydrogenase 10 | 3028 | ENSG00000072506 |
| 7 | O95831 | NA | AIFM1 | apoptosis-inducing factor, mitochondrion-associated, 1 | 9131 | ENSG00000156709 |
| 8 | P04179 | NA | SOD2 | superoxide dismutase 2, mitochondrial | 6648 | ENSG00000112096 |
| 9 | P00367 | NA | GLUD1 | glutamate dehydrogenase 1 | 2746 | ENSG00000148672 |
| 10 | P48735 | NA | IDH2 | isocitrate dehydrogenase 2 (NADP+), mitochondrial | 3418 | ENSG00000182054 |
| 11 | O60313 | NA | OPA1 | optic atrophy 1 (autosomal dominant) | 4976 | ENSG00000198836 |
| 12 | P53701 | NA | HCCS | holocytochrome c synthase | 3052 | ENSG00000004961 |
| 13 | P31040 | NA | SDHA | succinate dehydrogenase complex, subunit A, flavoprotein (Fp) | 6389 | ENSG00000073578 |
| 14 | P38117 | NA | ETFB | electron-transfer-flavoprotein, beta polypeptide | 2109 | ENSG00000105379 |
| 15 | P13804 | NA | ETFA | electron-transfer-flavoprotein, alpha polypeptide | 2108 | ENSG00000140374 |
| 16 | P42765 | NA | ACAA2 | acetyl-CoA acyltransferase 2 | 10449 | ENSG00000167315 |
| 17 | Q9Y277 | NA | VDAC3 | voltage-dependent anion channel 3 | 7419 | ENSG00000078668 |
| 18 | P49419 | NA | ALDH7A1 | aldehyde dehydrogenase 7 family, member A1 | 501 | ENSG00000164904 |
| 19 | Q04837 | NA | SSBP1 | single-stranded DNA binding protein 1, mitochondrial | 6742 | ENSG00000106028 |
| 20 | Q9BPW8 | NA | NIPSNAP1 | nipsnap homolog 1 (C. elegans) | 8508 | ENSG00000184117 |
| 21 | O75439 | NA | PMPCB | peptidase (mitochondrial processing) beta | 9512 | ENSG00000105819 |
| 22 | P21953 | NA | BCKDHB | branched chain keto acid dehydrogenase E1, beta polypeptide | 594 | ENSG00000083123 |
| 23 | P40939 | NA | HADHA | hydroxyacyl-CoA dehydrogenase/3-ketoacyl-CoA thiolase/enoyl-CoA hydratase (trifunctional protein), alpha subunit | 3030 | ENSG00000084754 |
| 24 | P30084 | NA | ECHS1 | enoyl CoA hydratase, short chain, 1, mitochondrial | 1892 | ENSG00000127884 |
| 25 | P52815 | NA | MRPL12 | mitochondrial ribosomal protein L12 | 6182 | ENSG00000262814 |
| 26 | P21796 | NA | VDAC1 | voltage-dependent anion channel 1 | 7416 | ENSG00000213585 |
| 27 | P04083 | NA | ANXA1 | annexin A1 | 301 | ENSG00000135046 |
| 28 | P32322 | NA | PYCR1 | pyrroline-5-carboxylate reductase 1 | 5831 | ENSG00000183010 |
| 29 | Q9UJZ1 | NA | STOML2 | stomatin (EPB72)-like 2 | 30968 | ENSG00000165283 |
| 30 | P04040 | NA | CAT | catalase | 847 | ENSG00000121691 |
| 31 | Q99798 | NA | ACO2 | aconitase 2, mitochondrial | 50 | ENSG00000100412 |
| 32 | P51649 | NA | ALDH5A1 | aldehyde dehydrogenase 5 family, member A1 | 7915 | ENSG00000112294 |
| 33 | P12235 | NA | SLC25A4 | solute carrier family 25 (mitochondrial carrier; adenine nucleotide translocator), member 4 | 291 | ENSG00000151729 |
| 34 | Q9NSE4 | NA | IARS2 | isoleucyl-tRNA synthetase 2, mitochondrial | 55699 | ENSG00000067704 |
| 35 | Q6PI48 | NA | DARS2 | aspartyl-tRNA synthetase 2, mitochondrial | 55157 | ENSG00000117593 |
| 36 | P24752 | NA | ACAT1 | acetyl-CoA acetyltransferase 1 | 38 | ENSG00000075239 |
| 37 | Q02978 | NA | SLC25A11 | solute carrier family 25 (mitochondrial carrier; oxoglutarate carrier), member 11 | 8402 | ENSG00000108528 |
| 38 | P19367 | NA | HK1 | hexokinase 1 | 3098 | ENSG00000156515 |
| 39 | P25705 | NA | ATP5A1 | ATP synthase, H+ transporting, mitochondrial F1 complex, alpha subunit 1, cardiac muscle | 498 | ENSG00000152234 |
| 40 | P49748 | NA | ACADVL | acyl-CoA dehydrogenase, very long chain | 37 | ENSG00000072778 |
| 41 | P48047 | NA | ATP5O | ATP synthase, H+ transporting, mitochondrial F1 complex, O subunit | 539 | ENSG00000241837 |
| 42 | P31937 | NA | HIBADH | 3-hydroxyisobutyrate dehydrogenase | 11112 | ENSG00000106049 |

  
  

| **Database:cellular component      &nbspName:cytoplasmic part      &nbspID:GO:0044444** | | | | | | |
| --- | --- | --- | --- | --- | --- | --- |
| C=6728; O=97; E=43.56; R=2.23; rawP=4.99e-27; adjP=2.92e-25 | | | | | | |
| Index | UserID | Value | Gene Symbol | Gene Name | EntrezGene | Ensembl |
| 1 | P10253 | NA | GAA | glucosidase, alpha; acid | 2548 | ENSG00000171298 |
| 2 | P68371 | NA | TUBB4B | tubulin, beta 4B class IVb | 10383 | ENSG00000188229 |
| 3 | Q9UL46 | NA | PSME2 | proteasome (prosome, macropain) activator subunit 2 (PA28 beta) | 5721 | ENSG00000100911 |
| 4 | P40926 | NA | MDH2 | malate dehydrogenase 2, NAD (mitochondrial) | 4191 | ENSG00000146701 |
| 5 | P04179 | NA | SOD2 | superoxide dismutase 2, mitochondrial | 6648 | ENSG00000112096 |
| 6 | Q13724 | NA | MOGS | mannosyl-oligosaccharide glucosidase | 7841 | ENSG00000115275 |
| 7 | P48735 | NA | IDH2 | isocitrate dehydrogenase 2 (NADP+), mitochondrial | 3418 | ENSG00000182054 |
| 8 | P00367 | NA | GLUD1 | glutamate dehydrogenase 1 | 2746 | ENSG00000148672 |
| 9 | Q06323 | NA | PSME1 | proteasome (prosome, macropain) activator subunit 1 (PA28 alpha) | 5720 | ENSG00000092010 |
| 10 | P53701 | NA | HCCS | holocytochrome c synthase | 3052 | ENSG00000004961 |
| 11 | P35580 | NA | MYH10 | myosin, heavy chain 10, non-muscle | 4628 | ENSG00000133026 |
| 12 | P31040 | NA | SDHA | succinate dehydrogenase complex, subunit A, flavoprotein (Fp) | 6389 | ENSG00000073578 |
| 13 | O75369 | NA | FLNB | filamin B, beta | 2317 | ENSG00000136068 |
| 14 | P24534 | NA | EEF1B2 | eukaryotic translation elongation factor 1 beta 2 | 1933 | ENSG00000114942 |
| 15 | O14773 | NA | TPP1 | tripeptidyl peptidase I | 1200 | ENSG00000166340 |
| 16 | P42765 | NA | ACAA2 | acetyl-CoA acyltransferase 2 | 10449 | ENSG00000167315 |
| 17 | P40121 | NA | CAPG | capping protein (actin filament), gelsolin-like | 822 | ENSG00000042493 |
| 18 | P13804 | NA | ETFA | electron-transfer-flavoprotein, alpha polypeptide | 2108 | ENSG00000140374 |
| 19 | Q9Y277 | NA | VDAC3 | voltage-dependent anion channel 3 | 7419 | ENSG00000078668 |
| 20 | P17174 | NA | GOT1 | glutamic-oxaloacetic transaminase 1, soluble (aspartate aminotransferase 1) | 2805 | ENSG00000120053 |
| 21 | P08727 | NA | KRT19 | keratin 19 | 3880 | ENSG00000171345 |
| 22 | Q04837 | NA | SSBP1 | single-stranded DNA binding protein 1, mitochondrial | 6742 | ENSG00000106028 |
| 23 | P50454 | NA | SERPINH1 | serpin peptidase inhibitor, clade H (heat shock protein 47), member 1, (collagen binding protein 1) | 871 | ENSG00000149257 |
| 24 | Q14697 | NA | GANAB | glucosidase, alpha; neutral AB | 23193 | ENSG00000089597 |
| 25 | P18206 | NA | VCL | vinculin | 7414 | ENSG00000035403 |
| 26 | P08195 | NA | SLC3A2 | solute carrier family 3 (activators of dibasic and neutral amino acid transport), member 2 | 6520 | ENSG00000168003 |
| 27 | P40939 | NA | HADHA | hydroxyacyl-CoA dehydrogenase/3-ketoacyl-CoA thiolase/enoyl-CoA hydratase (trifunctional protein), alpha subunit | 3030 | ENSG00000084754 |
| 28 | P52815 | NA | MRPL12 | mitochondrial ribosomal protein L12 | 6182 | ENSG00000262814 |
| 29 | Q9BXW7 | NA | CECR5 | cat eye syndrome chromosome region, candidate 5 | 27440 | ENSG00000069998 |
| 30 | P21796 | NA | VDAC1 | voltage-dependent anion channel 1 | 7416 | ENSG00000213585 |
| 31 | P38571 | NA | LIPA | lipase A, lysosomal acid, cholesterol esterase | 3988 | ENSG00000107798 |
| 32 | Q9UJZ1 | NA | STOML2 | stomatin (EPB72)-like 2 | 30968 | ENSG00000165283 |
| 33 | P32322 | NA | PYCR1 | pyrroline-5-carboxylate reductase 1 | 5831 | ENSG00000183010 |
| 34 | Q14980 | NA | NUMA1 | nuclear mitotic apparatus protein 1 | 4926 | ENSG00000137497 |
| 35 | Q99798 | NA | ACO2 | aconitase 2, mitochondrial | 50 | ENSG00000100412 |
| 36 | P51649 | NA | ALDH5A1 | aldehyde dehydrogenase 5 family, member A1 | 7915 | ENSG00000112294 |
| 37 | P12235 | NA | SLC25A4 | solute carrier family 25 (mitochondrial carrier; adenine nucleotide translocator), member 4 | 291 | ENSG00000151729 |
| 38 | Q6PI48 | NA | DARS2 | aspartyl-tRNA synthetase 2, mitochondrial | 55157 | ENSG00000117593 |
| 39 | Q9NSE4 | NA | IARS2 | isoleucyl-tRNA synthetase 2, mitochondrial | 55699 | ENSG00000067704 |
| 40 | P05556 | NA | ITGB1 | integrin, beta 1 (fibronectin receptor, beta polypeptide, antigen CD29 includes MDF2, MSK12) | 3688 | ENSG00000150093 |
| 41 | P13674 | NA | P4HA1 | prolyl 4-hydroxylase, alpha polypeptide I | 5033 | ENSG00000122884 |
| 42 | P40227 | NA | CCT6A | chaperonin containing TCP1, subunit 6A (zeta 1) | 908 | ENSG00000146731 |
| 43 | Q02978 | NA | SLC25A11 | solute carrier family 25 (mitochondrial carrier; oxoglutarate carrier), member 11 | 8402 | ENSG00000108528 |
| 44 | P60174 | NA | TPI1 | triosephosphate isomerase 1 | 7167 | ENSG00000111669 |
| 45 | P30042 | NA | C21orf33 | chromosome 21 open reading frame 33 | 8209 | ENSG00000160221 |
| 46 | P49589 | NA | CARS | cysteinyl-tRNA synthetase | 833 | ENSG00000110619 |
| 47 | P54819 | NA | AK2 | adenylate kinase 2 | 204 | ENSG00000004455 |
| 48 | P17931 | NA | LGALS3 | lectin, galactoside-binding, soluble, 3 | 3958 | ENSG00000131981 |
| 49 | Q99623 | NA | PHB2 | prohibitin 2 | 11331 | ENSG00000215021 |
| 50 | P50453 | NA | SERPINB9 | serpin peptidase inhibitor, clade B (ovalbumin), member 9 | 5272 | ENSG00000170542 |
| 51 | P35232 | NA | PHB | prohibitin | 5245 | ENSG00000167085 |
| 52 | O95831 | NA | AIFM1 | apoptosis-inducing factor, mitochondrion-associated, 1 | 9131 | ENSG00000156709 |
| 53 | Q99714 | NA | HSD17B10 | hydroxysteroid (17-beta) dehydrogenase 10 | 3028 | ENSG00000072506 |
| 54 | P61313 | NA | RPL15 | ribosomal protein L15 | 6138 | ENSG00000174748 |
| 55 | P07741 | NA | APRT | adenine phosphoribosyltransferase | 353 | ENSG00000198931 |
| 56 | O60313 | NA | OPA1 | optic atrophy 1 (autosomal dominant) | 4976 | ENSG00000198836 |
| 57 | P35270 | NA | SPR | sepiapterin reductase (7,8-dihydrobiopterin:NADP+ oxidoreductase) | 6697 | ENSG00000116096 |
| 58 | P68104 | NA | EEF1A1 | eukaryotic translation elongation factor 1 alpha 1 | 1915 | ENSG00000156508 |
| 59 | P02786 | NA | TFRC | transferrin receptor (p90, CD71) | 7037 | ENSG00000072274 |
| 60 | P38117 | NA | ETFB | electron-transfer-flavoprotein, beta polypeptide | 2109 | ENSG00000105379 |
| 61 | P43490 | NA | NAMPT | nicotinamide phosphoribosyltransferase | 10135 | ENSG00000105835 |
| 62 | P22102 | NA | GART | phosphoribosylglycinamide formyltransferase, phosphoribosylglycinamide synthetase, phosphoribosylaminoimidazole synthetase | 2618 | ENSG00000159131 |
| 63 | P13639 | NA | EEF2 | eukaryotic translation elongation factor 2 | 1938 | ENSG00000167658 |
| 64 | O75521 | NA | ECI2 | enoyl-CoA delta isomerase 2 | 10455 | ENSG00000198721 |
| 65 | P49419 | NA | ALDH7A1 | aldehyde dehydrogenase 7 family, member A1 | 501 | ENSG00000164904 |
| 66 | P67809 | NA | YBX1 | Y box binding protein 1 | 4904 | ENSG00000065978 |
| 67 | Q6PIU2 | NA | NCEH1 | neutral cholesterol ester hydrolase 1 | 57552 | ENSG00000144959 |
| 68 | P29692 | NA | EEF1D | eukaryotic translation elongation factor 1 delta (guanine nucleotide exchange protein) | 1936 | ENSG00000104529 |
| 69 | Q13813 | NA | SPTAN1 | spectrin, alpha, non-erythrocytic 1 | 6709 | ENSG00000197694 |
| 70 | Q9BPW8 | NA | NIPSNAP1 | nipsnap homolog 1 (C. elegans) | 8508 | ENSG00000184117 |
| 71 | P11387 | NA | TOP1 | topoisomerase (DNA) I | 7150 | ENSG00000198900 |
| 72 | P02792 | NA | FTL | ferritin, light polypeptide | 2512 | ENSG00000087086 |
| 73 | O75439 | NA | PMPCB | peptidase (mitochondrial processing) beta | 9512 | ENSG00000105819 |
| 74 | P00491 | NA | PNP | purine nucleoside phosphorylase | 4860 | ENSG00000198805 |
| 75 | P21953 | NA | BCKDHB | branched chain keto acid dehydrogenase E1, beta polypeptide | 594 | ENSG00000083123 |
| 76 | P30084 | NA | ECHS1 | enoyl CoA hydratase, short chain, 1, mitochondrial | 1892 | ENSG00000127884 |
| 77 | P60842 | NA | EIF4A1 | eukaryotic translation initiation factor 4A1 | 1973 | ENSG00000161960 |
| 78 | Q13011 | NA | ECH1 | enoyl CoA hydratase 1, peroxisomal | 1891 | ENSG00000104823 |
| 79 | P04083 | NA | ANXA1 | annexin A1 | 301 | ENSG00000135046 |
| 80 | P04040 | NA | CAT | catalase | 847 | ENSG00000121691 |
| 81 | O75340 | NA | PDCD6 | programmed cell death 6 | 10016 | ENSG00000249915 |
| 82 | P51688 | NA | SGSH | N-sulfoglucosamine sulfohydrolase | 6448 | ENSG00000181523 |
| 83 | P00558 | NA | PGK1 | phosphoglycerate kinase 1 | 5230 | ENSG00000102144 |
| 84 | P30048 | NA | PRDX3 | peroxiredoxin 3 | 10935 | ENSG00000165672 |
| 85 | P41091 | NA | EIF2S3 | eukaryotic translation initiation factor 2, subunit 3 gamma, 52kDa | 1968 | ENSG00000130741 |
| 86 | P02768 | NA | ALB | albumin | 213 | ENSG00000163631 |
| 87 | P07108 | NA | DBI | diazepam binding inhibitor (GABA receptor modulator, acyl-CoA binding protein) | 1622 | ENSG00000155368 |
| 88 | P24752 | NA | ACAT1 | acetyl-CoA acetyltransferase 1 | 38 | ENSG00000075239 |
| 89 | P19367 | NA | HK1 | hexokinase 1 | 3098 | ENSG00000156515 |
| 90 | P25705 | NA | ATP5A1 | ATP synthase, H+ transporting, mitochondrial F1 complex, alpha subunit 1, cardiac muscle | 498 | ENSG00000152234 |
| 91 | Q9BQ69 | NA | MACROD1 | MACRO domain containing 1 | 28992 | ENSG00000133315 |
| 92 | P05783 | NA | KRT18 | keratin 18 | 3875 | ENSG00000111057 |
| 93 | P06865 | NA | HEXA | hexosaminidase A (alpha polypeptide) | 3073 | ENSG00000213614 |
| 94 | P49748 | NA | ACADVL | acyl-CoA dehydrogenase, very long chain | 37 | ENSG00000072778 |
| 95 | P00352 | NA | ALDH1A1 | aldehyde dehydrogenase 1 family, member A1 | 216 | ENSG00000165092 |
| 96 | P48047 | NA | ATP5O | ATP synthase, H+ transporting, mitochondrial F1 complex, O subunit | 539 | ENSG00000241837 |
| 97 | P31937 | NA | HIBADH | 3-hydroxyisobutyrate dehydrogenase | 11112 | ENSG00000106049 |

  
  

| **Database:cellular component      &nbspName:mitochondrion      &nbspID:GO:0005739** | | | | | | |
| --- | --- | --- | --- | --- | --- | --- |
| C=1515; O=52; E=9.81; R=5.30; rawP=4.38e-26; adjP=1.71e-24 | | | | | | |
| Index | UserID | Value | Gene Symbol | Gene Name | EntrezGene | Ensembl |
| 1 | P54819 | NA | AK2 | adenylate kinase 2 | 204 | ENSG00000004455 |
| 2 | P17931 | NA | LGALS3 | lectin, galactoside-binding, soluble, 3 | 3958 | ENSG00000131981 |
| 3 | Q99623 | NA | PHB2 | prohibitin 2 | 11331 | ENSG00000215021 |
| 4 | P40926 | NA | MDH2 | malate dehydrogenase 2, NAD (mitochondrial) | 4191 | ENSG00000146701 |
| 5 | P35232 | NA | PHB | prohibitin | 5245 | ENSG00000167085 |
| 6 | Q99714 | NA | HSD17B10 | hydroxysteroid (17-beta) dehydrogenase 10 | 3028 | ENSG00000072506 |
| 7 | O95831 | NA | AIFM1 | apoptosis-inducing factor, mitochondrion-associated, 1 | 9131 | ENSG00000156709 |
| 8 | P04179 | NA | SOD2 | superoxide dismutase 2, mitochondrial | 6648 | ENSG00000112096 |
| 9 | P00367 | NA | GLUD1 | glutamate dehydrogenase 1 | 2746 | ENSG00000148672 |
| 10 | P48735 | NA | IDH2 | isocitrate dehydrogenase 2 (NADP+), mitochondrial | 3418 | ENSG00000182054 |
| 11 | O60313 | NA | OPA1 | optic atrophy 1 (autosomal dominant) | 4976 | ENSG00000198836 |
| 12 | P53701 | NA | HCCS | holocytochrome c synthase | 3052 | ENSG00000004961 |
| 13 | P31040 | NA | SDHA | succinate dehydrogenase complex, subunit A, flavoprotein (Fp) | 6389 | ENSG00000073578 |
| 14 | P38117 | NA | ETFB | electron-transfer-flavoprotein, beta polypeptide | 2109 | ENSG00000105379 |
| 15 | P02786 | NA | TFRC | transferrin receptor (p90, CD71) | 7037 | ENSG00000072274 |
| 16 | O14773 | NA | TPP1 | tripeptidyl peptidase I | 1200 | ENSG00000166340 |
| 17 | P13804 | NA | ETFA | electron-transfer-flavoprotein, alpha polypeptide | 2108 | ENSG00000140374 |
| 18 | P42765 | NA | ACAA2 | acetyl-CoA acyltransferase 2 | 10449 | ENSG00000167315 |
| 19 | O75521 | NA | ECI2 | enoyl-CoA delta isomerase 2 | 10455 | ENSG00000198721 |
| 20 | Q9Y277 | NA | VDAC3 | voltage-dependent anion channel 3 | 7419 | ENSG00000078668 |
| 21 | P49419 | NA | ALDH7A1 | aldehyde dehydrogenase 7 family, member A1 | 501 | ENSG00000164904 |
| 22 | Q04837 | NA | SSBP1 | single-stranded DNA binding protein 1, mitochondrial | 6742 | ENSG00000106028 |
| 23 | Q9BPW8 | NA | NIPSNAP1 | nipsnap homolog 1 (C. elegans) | 8508 | ENSG00000184117 |
| 24 | O75439 | NA | PMPCB | peptidase (mitochondrial processing) beta | 9512 | ENSG00000105819 |
| 25 | P21953 | NA | BCKDHB | branched chain keto acid dehydrogenase E1, beta polypeptide | 594 | ENSG00000083123 |
| 26 | P40939 | NA | HADHA | hydroxyacyl-CoA dehydrogenase/3-ketoacyl-CoA thiolase/enoyl-CoA hydratase (trifunctional protein), alpha subunit | 3030 | ENSG00000084754 |
| 27 | P30084 | NA | ECHS1 | enoyl CoA hydratase, short chain, 1, mitochondrial | 1892 | ENSG00000127884 |
| 28 | P52815 | NA | MRPL12 | mitochondrial ribosomal protein L12 | 6182 | ENSG00000262814 |
| 29 | Q9BXW7 | NA | CECR5 | cat eye syndrome chromosome region, candidate 5 | 27440 | ENSG00000069998 |
| 30 | P21796 | NA | VDAC1 | voltage-dependent anion channel 1 | 7416 | ENSG00000213585 |
| 31 | Q13011 | NA | ECH1 | enoyl CoA hydratase 1, peroxisomal | 1891 | ENSG00000104823 |
| 32 | P04083 | NA | ANXA1 | annexin A1 | 301 | ENSG00000135046 |
| 33 | P32322 | NA | PYCR1 | pyrroline-5-carboxylate reductase 1 | 5831 | ENSG00000183010 |
| 34 | Q9UJZ1 | NA | STOML2 | stomatin (EPB72)-like 2 | 30968 | ENSG00000165283 |
| 35 | P04040 | NA | CAT | catalase | 847 | ENSG00000121691 |
| 36 | Q99798 | NA | ACO2 | aconitase 2, mitochondrial | 50 | ENSG00000100412 |
| 37 | P51649 | NA | ALDH5A1 | aldehyde dehydrogenase 5 family, member A1 | 7915 | ENSG00000112294 |
| 38 | P12235 | NA | SLC25A4 | solute carrier family 25 (mitochondrial carrier; adenine nucleotide translocator), member 4 | 291 | ENSG00000151729 |
| 39 | P30048 | NA | PRDX3 | peroxiredoxin 3 | 10935 | ENSG00000165672 |
| 40 | P07108 | NA | DBI | diazepam binding inhibitor (GABA receptor modulator, acyl-CoA binding protein) | 1622 | ENSG00000155368 |
| 41 | Q9NSE4 | NA | IARS2 | isoleucyl-tRNA synthetase 2, mitochondrial | 55699 | ENSG00000067704 |
| 42 | Q6PI48 | NA | DARS2 | aspartyl-tRNA synthetase 2, mitochondrial | 55157 | ENSG00000117593 |
| 43 | P24752 | NA | ACAT1 | acetyl-CoA acetyltransferase 1 | 38 | ENSG00000075239 |
| 44 | P13674 | NA | P4HA1 | prolyl 4-hydroxylase, alpha polypeptide I | 5033 | ENSG00000122884 |
| 45 | Q02978 | NA | SLC25A11 | solute carrier family 25 (mitochondrial carrier; oxoglutarate carrier), member 11 | 8402 | ENSG00000108528 |
| 46 | P19367 | NA | HK1 | hexokinase 1 | 3098 | ENSG00000156515 |
| 47 | P25705 | NA | ATP5A1 | ATP synthase, H+ transporting, mitochondrial F1 complex, alpha subunit 1, cardiac muscle | 498 | ENSG00000152234 |
| 48 | Q9BQ69 | NA | MACROD1 | MACRO domain containing 1 | 28992 | ENSG00000133315 |
| 49 | P49748 | NA | ACADVL | acyl-CoA dehydrogenase, very long chain | 37 | ENSG00000072778 |
| 50 | P48047 | NA | ATP5O | ATP synthase, H+ transporting, mitochondrial F1 complex, O subunit | 539 | ENSG00000241837 |
| 51 | P31937 | NA | HIBADH | 3-hydroxyisobutyrate dehydrogenase | 11112 | ENSG00000106049 |
| 52 | P30042 | NA | C21orf33 | chromosome 21 open reading frame 33 | 8209 | ENSG00000160221 |

  
  

| **Database:cellular component      &nbspName:cytoplasm      &nbspID:GO:0005737** | | | | | | |
| --- | --- | --- | --- | --- | --- | --- |
| C=9051; O=104; E=58.60; R=1.77; rawP=4.62e-23; adjP=1.35e-21 | | | | | | |
| Index | UserID | Value | Gene Symbol | Gene Name | EntrezGene | Ensembl |
| 1 | P10253 | NA | GAA | glucosidase, alpha; acid | 2548 | ENSG00000171298 |
| 2 | P68371 | NA | TUBB4B | tubulin, beta 4B class IVb | 10383 | ENSG00000188229 |
| 3 | Q9UL46 | NA | PSME2 | proteasome (prosome, macropain) activator subunit 2 (PA28 beta) | 5721 | ENSG00000100911 |
| 4 | P40926 | NA | MDH2 | malate dehydrogenase 2, NAD (mitochondrial) | 4191 | ENSG00000146701 |
| 5 | P04179 | NA | SOD2 | superoxide dismutase 2, mitochondrial | 6648 | ENSG00000112096 |
| 6 | Q13724 | NA | MOGS | mannosyl-oligosaccharide glucosidase | 7841 | ENSG00000115275 |
| 7 | P48735 | NA | IDH2 | isocitrate dehydrogenase 2 (NADP+), mitochondrial | 3418 | ENSG00000182054 |
| 8 | P00367 | NA | GLUD1 | glutamate dehydrogenase 1 | 2746 | ENSG00000148672 |
| 9 | Q06323 | NA | PSME1 | proteasome (prosome, macropain) activator subunit 1 (PA28 alpha) | 5720 | ENSG00000092010 |
| 10 | P53701 | NA | HCCS | holocytochrome c synthase | 3052 | ENSG00000004961 |
| 11 | P35580 | NA | MYH10 | myosin, heavy chain 10, non-muscle | 4628 | ENSG00000133026 |
| 12 | P31040 | NA | SDHA | succinate dehydrogenase complex, subunit A, flavoprotein (Fp) | 6389 | ENSG00000073578 |
| 13 | O75369 | NA | FLNB | filamin B, beta | 2317 | ENSG00000136068 |
| 14 | P24534 | NA | EEF1B2 | eukaryotic translation elongation factor 1 beta 2 | 1933 | ENSG00000114942 |
| 15 | O14773 | NA | TPP1 | tripeptidyl peptidase I | 1200 | ENSG00000166340 |
| 16 | P47895 | NA | ALDH1A3 | aldehyde dehydrogenase 1 family, member A3 | 220 | ENSG00000184254 |
| 17 | P42765 | NA | ACAA2 | acetyl-CoA acyltransferase 2 | 10449 | ENSG00000167315 |
| 18 | P40121 | NA | CAPG | capping protein (actin filament), gelsolin-like | 822 | ENSG00000042493 |
| 19 | P13804 | NA | ETFA | electron-transfer-flavoprotein, alpha polypeptide | 2108 | ENSG00000140374 |
| 20 | Q9Y277 | NA | VDAC3 | voltage-dependent anion channel 3 | 7419 | ENSG00000078668 |
| 21 | P17174 | NA | GOT1 | glutamic-oxaloacetic transaminase 1, soluble (aspartate aminotransferase 1) | 2805 | ENSG00000120053 |
| 22 | P08727 | NA | KRT19 | keratin 19 | 3880 | ENSG00000171345 |
| 23 | Q04837 | NA | SSBP1 | single-stranded DNA binding protein 1, mitochondrial | 6742 | ENSG00000106028 |
| 24 | P50454 | NA | SERPINH1 | serpin peptidase inhibitor, clade H (heat shock protein 47), member 1, (collagen binding protein 1) | 871 | ENSG00000149257 |
| 25 | Q14697 | NA | GANAB | glucosidase, alpha; neutral AB | 23193 | ENSG00000089597 |
| 26 | P18206 | NA | VCL | vinculin | 7414 | ENSG00000035403 |
| 27 | P08195 | NA | SLC3A2 | solute carrier family 3 (activators of dibasic and neutral amino acid transport), member 2 | 6520 | ENSG00000168003 |
| 28 | P40939 | NA | HADHA | hydroxyacyl-CoA dehydrogenase/3-ketoacyl-CoA thiolase/enoyl-CoA hydratase (trifunctional protein), alpha subunit | 3030 | ENSG00000084754 |
| 29 | P09382 | NA | LGALS1 | lectin, galactoside-binding, soluble, 1 | 3956 | ENSG00000100097 |
| 30 | P52815 | NA | MRPL12 | mitochondrial ribosomal protein L12 | 6182 | ENSG00000262814 |
| 31 | Q9BXW7 | NA | CECR5 | cat eye syndrome chromosome region, candidate 5 | 27440 | ENSG00000069998 |
| 32 | P21796 | NA | VDAC1 | voltage-dependent anion channel 1 | 7416 | ENSG00000213585 |
| 33 | P38571 | NA | LIPA | lipase A, lysosomal acid, cholesterol esterase | 3988 | ENSG00000107798 |
| 34 | P04080 | NA | CSTB | cystatin B (stefin B) | 1476 | ENSG00000160213 |
| 35 | Q9UJZ1 | NA | STOML2 | stomatin (EPB72)-like 2 | 30968 | ENSG00000165283 |
| 36 | P32322 | NA | PYCR1 | pyrroline-5-carboxylate reductase 1 | 5831 | ENSG00000183010 |
| 37 | Q14980 | NA | NUMA1 | nuclear mitotic apparatus protein 1 | 4926 | ENSG00000137497 |
| 38 | Q99798 | NA | ACO2 | aconitase 2, mitochondrial | 50 | ENSG00000100412 |
| 39 | P51649 | NA | ALDH5A1 | aldehyde dehydrogenase 5 family, member A1 | 7915 | ENSG00000112294 |
| 40 | P12235 | NA | SLC25A4 | solute carrier family 25 (mitochondrial carrier; adenine nucleotide translocator), member 4 | 291 | ENSG00000151729 |
| 41 | P27816 | NA | MAP4 | microtubule-associated protein 4 | 4134 | ENSG00000047849 |
| 42 | Q6PI48 | NA | DARS2 | aspartyl-tRNA synthetase 2, mitochondrial | 55157 | ENSG00000117593 |
| 43 | Q9NSE4 | NA | IARS2 | isoleucyl-tRNA synthetase 2, mitochondrial | 55699 | ENSG00000067704 |
| 44 | P05556 | NA | ITGB1 | integrin, beta 1 (fibronectin receptor, beta polypeptide, antigen CD29 includes MDF2, MSK12) | 3688 | ENSG00000150093 |
| 45 | P13674 | NA | P4HA1 | prolyl 4-hydroxylase, alpha polypeptide I | 5033 | ENSG00000122884 |
| 46 | P40227 | NA | CCT6A | chaperonin containing TCP1, subunit 6A (zeta 1) | 908 | ENSG00000146731 |
| 47 | Q02978 | NA | SLC25A11 | solute carrier family 25 (mitochondrial carrier; oxoglutarate carrier), member 11 | 8402 | ENSG00000108528 |
| 48 | P60174 | NA | TPI1 | triosephosphate isomerase 1 | 7167 | ENSG00000111669 |
| 49 | P30042 | NA | C21orf33 | chromosome 21 open reading frame 33 | 8209 | ENSG00000160221 |
| 50 | P49589 | NA | CARS | cysteinyl-tRNA synthetase | 833 | ENSG00000110619 |
| 51 | P54819 | NA | AK2 | adenylate kinase 2 | 204 | ENSG00000004455 |
| 52 | P17931 | NA | LGALS3 | lectin, galactoside-binding, soluble, 3 | 3958 | ENSG00000131981 |
| 53 | P05787 | NA | KRT8 | keratin 8 | 3856 | ENSG00000170421 |
| 54 | Q99623 | NA | PHB2 | prohibitin 2 | 11331 | ENSG00000215021 |
| 55 | P50453 | NA | SERPINB9 | serpin peptidase inhibitor, clade B (ovalbumin), member 9 | 5272 | ENSG00000170542 |
| 56 | P35232 | NA | PHB | prohibitin | 5245 | ENSG00000167085 |
| 57 | O95831 | NA | AIFM1 | apoptosis-inducing factor, mitochondrion-associated, 1 | 9131 | ENSG00000156709 |
| 58 | Q99714 | NA | HSD17B10 | hydroxysteroid (17-beta) dehydrogenase 10 | 3028 | ENSG00000072506 |
| 59 | P61313 | NA | RPL15 | ribosomal protein L15 | 6138 | ENSG00000174748 |
| 60 | P07741 | NA | APRT | adenine phosphoribosyltransferase | 353 | ENSG00000198931 |
| 61 | O60313 | NA | OPA1 | optic atrophy 1 (autosomal dominant) | 4976 | ENSG00000198836 |
| 62 | P35270 | NA | SPR | sepiapterin reductase (7,8-dihydrobiopterin:NADP+ oxidoreductase) | 6697 | ENSG00000116096 |
| 63 | P68104 | NA | EEF1A1 | eukaryotic translation elongation factor 1 alpha 1 | 1915 | ENSG00000156508 |
| 64 | P38117 | NA | ETFB | electron-transfer-flavoprotein, beta polypeptide | 2109 | ENSG00000105379 |
| 65 | P02786 | NA | TFRC | transferrin receptor (p90, CD71) | 7037 | ENSG00000072274 |
| 66 | P43490 | NA | NAMPT | nicotinamide phosphoribosyltransferase | 10135 | ENSG00000105835 |
| 67 | P22102 | NA | GART | phosphoribosylglycinamide formyltransferase, phosphoribosylglycinamide synthetase, phosphoribosylaminoimidazole synthetase | 2618 | ENSG00000159131 |
| 68 | P13639 | NA | EEF2 | eukaryotic translation elongation factor 2 | 1938 | ENSG00000167658 |
| 69 | O75521 | NA | ECI2 | enoyl-CoA delta isomerase 2 | 10455 | ENSG00000198721 |
| 70 | P49419 | NA | ALDH7A1 | aldehyde dehydrogenase 7 family, member A1 | 501 | ENSG00000164904 |
| 71 | P67809 | NA | YBX1 | Y box binding protein 1 | 4904 | ENSG00000065978 |
| 72 | Q14764 | NA | MVP | major vault protein | 9961 | ENSG00000013364 |
| 73 | Q6PIU2 | NA | NCEH1 | neutral cholesterol ester hydrolase 1 | 57552 | ENSG00000144959 |
| 74 | P29692 | NA | EEF1D | eukaryotic translation elongation factor 1 delta (guanine nucleotide exchange protein) | 1936 | ENSG00000104529 |
| 75 | Q13813 | NA | SPTAN1 | spectrin, alpha, non-erythrocytic 1 | 6709 | ENSG00000197694 |
| 76 | Q9BPW8 | NA | NIPSNAP1 | nipsnap homolog 1 (C. elegans) | 8508 | ENSG00000184117 |
| 77 | P11387 | NA | TOP1 | topoisomerase (DNA) I | 7150 | ENSG00000198900 |
| 78 | P02792 | NA | FTL | ferritin, light polypeptide | 2512 | ENSG00000087086 |
| 79 | O75439 | NA | PMPCB | peptidase (mitochondrial processing) beta | 9512 | ENSG00000105819 |
| 80 | P00491 | NA | PNP | purine nucleoside phosphorylase | 4860 | ENSG00000198805 |
| 81 | P21953 | NA | BCKDHB | branched chain keto acid dehydrogenase E1, beta polypeptide | 594 | ENSG00000083123 |
| 82 | P30084 | NA | ECHS1 | enoyl CoA hydratase, short chain, 1, mitochondrial | 1892 | ENSG00000127884 |
| 83 | P60842 | NA | EIF4A1 | eukaryotic translation initiation factor 4A1 | 1973 | ENSG00000161960 |
| 84 | Q13011 | NA | ECH1 | enoyl CoA hydratase 1, peroxisomal | 1891 | ENSG00000104823 |
| 85 | P04083 | NA | ANXA1 | annexin A1 | 301 | ENSG00000135046 |
| 86 | P04040 | NA | CAT | catalase | 847 | ENSG00000121691 |
| 87 | O75340 | NA | PDCD6 | programmed cell death 6 | 10016 | ENSG00000249915 |
| 88 | P51688 | NA | SGSH | N-sulfoglucosamine sulfohydrolase | 6448 | ENSG00000181523 |
| 89 | P00558 | NA | PGK1 | phosphoglycerate kinase 1 | 5230 | ENSG00000102144 |
| 90 | P30048 | NA | PRDX3 | peroxiredoxin 3 | 10935 | ENSG00000165672 |
| 91 | P41091 | NA | EIF2S3 | eukaryotic translation initiation factor 2, subunit 3 gamma, 52kDa | 1968 | ENSG00000130741 |
| 92 | P02768 | NA | ALB | albumin | 213 | ENSG00000163631 |
| 93 | P07108 | NA | DBI | diazepam binding inhibitor (GABA receptor modulator, acyl-CoA binding protein) | 1622 | ENSG00000155368 |
| 94 | P24752 | NA | ACAT1 | acetyl-CoA acetyltransferase 1 | 38 | ENSG00000075239 |
| 95 | P19367 | NA | HK1 | hexokinase 1 | 3098 | ENSG00000156515 |
| 96 | P25705 | NA | ATP5A1 | ATP synthase, H+ transporting, mitochondrial F1 complex, alpha subunit 1, cardiac muscle | 498 | ENSG00000152234 |
| 97 | Q9BQ69 | NA | MACROD1 | MACRO domain containing 1 | 28992 | ENSG00000133315 |
| 98 | P05783 | NA | KRT18 | keratin 18 | 3875 | ENSG00000111057 |
| 99 | P06865 | NA | HEXA | hexosaminidase A (alpha polypeptide) | 3073 | ENSG00000213614 |
| 100 | P49748 | NA | ACADVL | acyl-CoA dehydrogenase, very long chain | 37 | ENSG00000072778 |
| 101 | P00352 | NA | ALDH1A1 | aldehyde dehydrogenase 1 family, member A1 | 216 | ENSG00000165092 |
| 102 | P48047 | NA | ATP5O | ATP synthase, H+ transporting, mitochondrial F1 complex, O subunit | 539 | ENSG00000241837 |
| 103 | Q16531 | NA | DDB1 | damage-specific DNA binding protein 1, 127kDa | 1642 | ENSG00000167986 |
| 104 | P31937 | NA | HIBADH | 3-hydroxyisobutyrate dehydrogenase | 11112 | ENSG00000106049 |

  
  

| **Database:cellular component      &nbspName:mitochondrial matrix      &nbspID:GO:0005759** | | | | | | |
| --- | --- | --- | --- | --- | --- | --- |
| C=278; O=24; E=1.80; R=13.33; rawP=1.43e-20; adjP=3.35e-19 | | | | | | |
| Index | UserID | Value | Gene Symbol | Gene Name | EntrezGene | Ensembl |
| 1 | O75439 | NA | PMPCB | peptidase (mitochondrial processing) beta | 9512 | ENSG00000105819 |
| 2 | P21953 | NA | BCKDHB | branched chain keto acid dehydrogenase E1, beta polypeptide | 594 | ENSG00000083123 |
| 3 | P40939 | NA | HADHA | hydroxyacyl-CoA dehydrogenase/3-ketoacyl-CoA thiolase/enoyl-CoA hydratase (trifunctional protein), alpha subunit | 3030 | ENSG00000084754 |
| 4 | P40926 | NA | MDH2 | malate dehydrogenase 2, NAD (mitochondrial) | 4191 | ENSG00000146701 |
| 5 | P30084 | NA | ECHS1 | enoyl CoA hydratase, short chain, 1, mitochondrial | 1892 | ENSG00000127884 |
| 6 | Q99714 | NA | HSD17B10 | hydroxysteroid (17-beta) dehydrogenase 10 | 3028 | ENSG00000072506 |
| 7 | P52815 | NA | MRPL12 | mitochondrial ribosomal protein L12 | 6182 | ENSG00000262814 |
| 8 | P04179 | NA | SOD2 | superoxide dismutase 2, mitochondrial | 6648 | ENSG00000112096 |
| 9 | P00367 | NA | GLUD1 | glutamate dehydrogenase 1 | 2746 | ENSG00000148672 |
| 10 | P48735 | NA | IDH2 | isocitrate dehydrogenase 2 (NADP+), mitochondrial | 3418 | ENSG00000182054 |
| 11 | P21796 | NA | VDAC1 | voltage-dependent anion channel 1 | 7416 | ENSG00000213585 |
| 12 | P38117 | NA | ETFB | electron-transfer-flavoprotein, beta polypeptide | 2109 | ENSG00000105379 |
| 13 | P32322 | NA | PYCR1 | pyrroline-5-carboxylate reductase 1 | 5831 | ENSG00000183010 |
| 14 | Q99798 | NA | ACO2 | aconitase 2, mitochondrial | 50 | ENSG00000100412 |
| 15 | P51649 | NA | ALDH5A1 | aldehyde dehydrogenase 5 family, member A1 | 7915 | ENSG00000112294 |
| 16 | P13804 | NA | ETFA | electron-transfer-flavoprotein, alpha polypeptide | 2108 | ENSG00000140374 |
| 17 | Q9NSE4 | NA | IARS2 | isoleucyl-tRNA synthetase 2, mitochondrial | 55699 | ENSG00000067704 |
| 18 | Q6PI48 | NA | DARS2 | aspartyl-tRNA synthetase 2, mitochondrial | 55157 | ENSG00000117593 |
| 19 | P24752 | NA | ACAT1 | acetyl-CoA acetyltransferase 1 | 38 | ENSG00000075239 |
| 20 | P49419 | NA | ALDH7A1 | aldehyde dehydrogenase 7 family, member A1 | 501 | ENSG00000164904 |
| 21 | P25705 | NA | ATP5A1 | ATP synthase, H+ transporting, mitochondrial F1 complex, alpha subunit 1, cardiac muscle | 498 | ENSG00000152234 |
| 22 | Q04837 | NA | SSBP1 | single-stranded DNA binding protein 1, mitochondrial | 6742 | ENSG00000106028 |
| 23 | P49748 | NA | ACADVL | acyl-CoA dehydrogenase, very long chain | 37 | ENSG00000072778 |
| 24 | P31937 | NA | HIBADH | 3-hydroxyisobutyrate dehydrogenase | 11112 | ENSG00000106049 |

  
  

| **Database:cellular component      &nbspName:mitochondrial envelope      &nbspID:GO:0005740** | | | | | | |
| --- | --- | --- | --- | --- | --- | --- |
| C=526; O=28; E=3.41; R=8.22; rawP=2.64e-18; adjP=5.15e-17 | | | | | | |
| Index | UserID | Value | Gene Symbol | Gene Name | EntrezGene | Ensembl |
| 1 | O75439 | NA | PMPCB | peptidase (mitochondrial processing) beta | 9512 | ENSG00000105819 |
| 2 | P54819 | NA | AK2 | adenylate kinase 2 | 204 | ENSG00000004455 |
| 3 | P17931 | NA | LGALS3 | lectin, galactoside-binding, soluble, 3 | 3958 | ENSG00000131981 |
| 4 | Q99623 | NA | PHB2 | prohibitin 2 | 11331 | ENSG00000215021 |
| 5 | P40939 | NA | HADHA | hydroxyacyl-CoA dehydrogenase/3-ketoacyl-CoA thiolase/enoyl-CoA hydratase (trifunctional protein), alpha subunit | 3030 | ENSG00000084754 |
| 6 | P40926 | NA | MDH2 | malate dehydrogenase 2, NAD (mitochondrial) | 4191 | ENSG00000146701 |
| 7 | P35232 | NA | PHB | prohibitin | 5245 | ENSG00000167085 |
| 8 | Q99714 | NA | HSD17B10 | hydroxysteroid (17-beta) dehydrogenase 10 | 3028 | ENSG00000072506 |
| 9 | O95831 | NA | AIFM1 | apoptosis-inducing factor, mitochondrion-associated, 1 | 9131 | ENSG00000156709 |
| 10 | P04179 | NA | SOD2 | superoxide dismutase 2, mitochondrial | 6648 | ENSG00000112096 |
| 11 | P48735 | NA | IDH2 | isocitrate dehydrogenase 2 (NADP+), mitochondrial | 3418 | ENSG00000182054 |
| 12 | O60313 | NA | OPA1 | optic atrophy 1 (autosomal dominant) | 4976 | ENSG00000198836 |
| 13 | P21796 | NA | VDAC1 | voltage-dependent anion channel 1 | 7416 | ENSG00000213585 |
| 14 | P53701 | NA | HCCS | holocytochrome c synthase | 3052 | ENSG00000004961 |
| 15 | P31040 | NA | SDHA | succinate dehydrogenase complex, subunit A, flavoprotein (Fp) | 6389 | ENSG00000073578 |
| 16 | P04083 | NA | ANXA1 | annexin A1 | 301 | ENSG00000135046 |
| 17 | Q9UJZ1 | NA | STOML2 | stomatin (EPB72)-like 2 | 30968 | ENSG00000165283 |
| 18 | P04040 | NA | CAT | catalase | 847 | ENSG00000121691 |
| 19 | P12235 | NA | SLC25A4 | solute carrier family 25 (mitochondrial carrier; adenine nucleotide translocator), member 4 | 291 | ENSG00000151729 |
| 20 | P42765 | NA | ACAA2 | acetyl-CoA acyltransferase 2 | 10449 | ENSG00000167315 |
| 21 | P24752 | NA | ACAT1 | acetyl-CoA acetyltransferase 1 | 38 | ENSG00000075239 |
| 22 | Q9Y277 | NA | VDAC3 | voltage-dependent anion channel 3 | 7419 | ENSG00000078668 |
| 23 | Q02978 | NA | SLC25A11 | solute carrier family 25 (mitochondrial carrier; oxoglutarate carrier), member 11 | 8402 | ENSG00000108528 |
| 24 | P19367 | NA | HK1 | hexokinase 1 | 3098 | ENSG00000156515 |
| 25 | P25705 | NA | ATP5A1 | ATP synthase, H+ transporting, mitochondrial F1 complex, alpha subunit 1, cardiac muscle | 498 | ENSG00000152234 |
| 26 | P49748 | NA | ACADVL | acyl-CoA dehydrogenase, very long chain | 37 | ENSG00000072778 |
| 27 | P48047 | NA | ATP5O | ATP synthase, H+ transporting, mitochondrial F1 complex, O subunit | 539 | ENSG00000241837 |
| 28 | Q9BPW8 | NA | NIPSNAP1 | nipsnap homolog 1 (C. elegans) | 8508 | ENSG00000184117 |

  
  

| **Database:cellular component      &nbspName:mitochondrial inner membrane      &nbspID:GO:0005743** | | | | | | |
| --- | --- | --- | --- | --- | --- | --- |
| C=351; O=24; E=2.27; R=10.56; rawP=3.35e-18; adjP=5.60e-17 | | | | | | |
| Index | UserID | Value | Gene Symbol | Gene Name | EntrezGene | Ensembl |
| 1 | O75439 | NA | PMPCB | peptidase (mitochondrial processing) beta | 9512 | ENSG00000105819 |
| 2 | P54819 | NA | AK2 | adenylate kinase 2 | 204 | ENSG00000004455 |
| 3 | P17931 | NA | LGALS3 | lectin, galactoside-binding, soluble, 3 | 3958 | ENSG00000131981 |
| 4 | Q99623 | NA | PHB2 | prohibitin 2 | 11331 | ENSG00000215021 |
| 5 | P40939 | NA | HADHA | hydroxyacyl-CoA dehydrogenase/3-ketoacyl-CoA thiolase/enoyl-CoA hydratase (trifunctional protein), alpha subunit | 3030 | ENSG00000084754 |
| 6 | P40926 | NA | MDH2 | malate dehydrogenase 2, NAD (mitochondrial) | 4191 | ENSG00000146701 |
| 7 | P35232 | NA | PHB | prohibitin | 5245 | ENSG00000167085 |
| 8 | Q99714 | NA | HSD17B10 | hydroxysteroid (17-beta) dehydrogenase 10 | 3028 | ENSG00000072506 |
| 9 | O95831 | NA | AIFM1 | apoptosis-inducing factor, mitochondrion-associated, 1 | 9131 | ENSG00000156709 |
| 10 | P04179 | NA | SOD2 | superoxide dismutase 2, mitochondrial | 6648 | ENSG00000112096 |
| 11 | P48735 | NA | IDH2 | isocitrate dehydrogenase 2 (NADP+), mitochondrial | 3418 | ENSG00000182054 |
| 12 | O60313 | NA | OPA1 | optic atrophy 1 (autosomal dominant) | 4976 | ENSG00000198836 |
| 13 | P53701 | NA | HCCS | holocytochrome c synthase | 3052 | ENSG00000004961 |
| 14 | P21796 | NA | VDAC1 | voltage-dependent anion channel 1 | 7416 | ENSG00000213585 |
| 15 | P31040 | NA | SDHA | succinate dehydrogenase complex, subunit A, flavoprotein (Fp) | 6389 | ENSG00000073578 |
| 16 | Q9UJZ1 | NA | STOML2 | stomatin (EPB72)-like 2 | 30968 | ENSG00000165283 |
| 17 | P12235 | NA | SLC25A4 | solute carrier family 25 (mitochondrial carrier; adenine nucleotide translocator), member 4 | 291 | ENSG00000151729 |
| 18 | P42765 | NA | ACAA2 | acetyl-CoA acyltransferase 2 | 10449 | ENSG00000167315 |
| 19 | P24752 | NA | ACAT1 | acetyl-CoA acetyltransferase 1 | 38 | ENSG00000075239 |
| 20 | Q02978 | NA | SLC25A11 | solute carrier family 25 (mitochondrial carrier; oxoglutarate carrier), member 11 | 8402 | ENSG00000108528 |
| 21 | P25705 | NA | ATP5A1 | ATP synthase, H+ transporting, mitochondrial F1 complex, alpha subunit 1, cardiac muscle | 498 | ENSG00000152234 |
| 22 | P49748 | NA | ACADVL | acyl-CoA dehydrogenase, very long chain | 37 | ENSG00000072778 |
| 23 | P48047 | NA | ATP5O | ATP synthase, H+ transporting, mitochondrial F1 complex, O subunit | 539 | ENSG00000241837 |
| 24 | Q9BPW8 | NA | NIPSNAP1 | nipsnap homolog 1 (C. elegans) | 8508 | ENSG00000184117 |

  
  

| **Database:cellular component      &nbspName:mitochondrial membrane      &nbspID:GO:0031966** | | | | | | |
| --- | --- | --- | --- | --- | --- | --- |
| C=505; O=27; E=3.27; R=8.26; rawP=1.06e-17; adjP=1.55e-16 | | | | | | |
| Index | UserID | Value | Gene Symbol | Gene Name | EntrezGene | Ensembl |
| 1 | O75439 | NA | PMPCB | peptidase (mitochondrial processing) beta | 9512 | ENSG00000105819 |
| 2 | P54819 | NA | AK2 | adenylate kinase 2 | 204 | ENSG00000004455 |
| 3 | P17931 | NA | LGALS3 | lectin, galactoside-binding, soluble, 3 | 3958 | ENSG00000131981 |
| 4 | Q99623 | NA | PHB2 | prohibitin 2 | 11331 | ENSG00000215021 |
| 5 | P40939 | NA | HADHA | hydroxyacyl-CoA dehydrogenase/3-ketoacyl-CoA thiolase/enoyl-CoA hydratase (trifunctional protein), alpha subunit | 3030 | ENSG00000084754 |
| 6 | P40926 | NA | MDH2 | malate dehydrogenase 2, NAD (mitochondrial) | 4191 | ENSG00000146701 |
| 7 | P35232 | NA | PHB | prohibitin | 5245 | ENSG00000167085 |
| 8 | Q99714 | NA | HSD17B10 | hydroxysteroid (17-beta) dehydrogenase 10 | 3028 | ENSG00000072506 |
| 9 | O95831 | NA | AIFM1 | apoptosis-inducing factor, mitochondrion-associated, 1 | 9131 | ENSG00000156709 |
| 10 | P04179 | NA | SOD2 | superoxide dismutase 2, mitochondrial | 6648 | ENSG00000112096 |
| 11 | P48735 | NA | IDH2 | isocitrate dehydrogenase 2 (NADP+), mitochondrial | 3418 | ENSG00000182054 |
| 12 | O60313 | NA | OPA1 | optic atrophy 1 (autosomal dominant) | 4976 | ENSG00000198836 |
| 13 | P21796 | NA | VDAC1 | voltage-dependent anion channel 1 | 7416 | ENSG00000213585 |
| 14 | P53701 | NA | HCCS | holocytochrome c synthase | 3052 | ENSG00000004961 |
| 15 | P31040 | NA | SDHA | succinate dehydrogenase complex, subunit A, flavoprotein (Fp) | 6389 | ENSG00000073578 |
| 16 | P04083 | NA | ANXA1 | annexin A1 | 301 | ENSG00000135046 |
| 17 | Q9UJZ1 | NA | STOML2 | stomatin (EPB72)-like 2 | 30968 | ENSG00000165283 |
| 18 | P12235 | NA | SLC25A4 | solute carrier family 25 (mitochondrial carrier; adenine nucleotide translocator), member 4 | 291 | ENSG00000151729 |
| 19 | P42765 | NA | ACAA2 | acetyl-CoA acyltransferase 2 | 10449 | ENSG00000167315 |
| 20 | P24752 | NA | ACAT1 | acetyl-CoA acetyltransferase 1 | 38 | ENSG00000075239 |
| 21 | Q9Y277 | NA | VDAC3 | voltage-dependent anion channel 3 | 7419 | ENSG00000078668 |
| 22 | Q02978 | NA | SLC25A11 | solute carrier family 25 (mitochondrial carrier; oxoglutarate carrier), member 11 | 8402 | ENSG00000108528 |
| 23 | P19367 | NA | HK1 | hexokinase 1 | 3098 | ENSG00000156515 |
| 24 | P25705 | NA | ATP5A1 | ATP synthase, H+ transporting, mitochondrial F1 complex, alpha subunit 1, cardiac muscle | 498 | ENSG00000152234 |
| 25 | P49748 | NA | ACADVL | acyl-CoA dehydrogenase, very long chain | 37 | ENSG00000072778 |
| 26 | P48047 | NA | ATP5O | ATP synthase, H+ transporting, mitochondrial F1 complex, O subunit | 539 | ENSG00000241837 |
| 27 | Q9BPW8 | NA | NIPSNAP1 | nipsnap homolog 1 (C. elegans) | 8508 | ENSG00000184117 |

  
  

| **Database:cellular component      &nbspName:organelle inner membrane      &nbspID:GO:0019866** | | | | | | |
| --- | --- | --- | --- | --- | --- | --- |
| C=381; O=24; E=2.47; R=9.73; rawP=2.21e-17; adjP=2.87e-16 | | | | | | |
| Index | UserID | Value | Gene Symbol | Gene Name | EntrezGene | Ensembl |
| 1 | O75439 | NA | PMPCB | peptidase (mitochondrial processing) beta | 9512 | ENSG00000105819 |
| 2 | P54819 | NA | AK2 | adenylate kinase 2 | 204 | ENSG00000004455 |
| 3 | P17931 | NA | LGALS3 | lectin, galactoside-binding, soluble, 3 | 3958 | ENSG00000131981 |
| 4 | Q99623 | NA | PHB2 | prohibitin 2 | 11331 | ENSG00000215021 |
| 5 | P40939 | NA | HADHA | hydroxyacyl-CoA dehydrogenase/3-ketoacyl-CoA thiolase/enoyl-CoA hydratase (trifunctional protein), alpha subunit | 3030 | ENSG00000084754 |
| 6 | P40926 | NA | MDH2 | malate dehydrogenase 2, NAD (mitochondrial) | 4191 | ENSG00000146701 |
| 7 | P35232 | NA | PHB | prohibitin | 5245 | ENSG00000167085 |
| 8 | Q99714 | NA | HSD17B10 | hydroxysteroid (17-beta) dehydrogenase 10 | 3028 | ENSG00000072506 |
| 9 | O95831 | NA | AIFM1 | apoptosis-inducing factor, mitochondrion-associated, 1 | 9131 | ENSG00000156709 |
| 10 | P04179 | NA | SOD2 | superoxide dismutase 2, mitochondrial | 6648 | ENSG00000112096 |
| 11 | P48735 | NA | IDH2 | isocitrate dehydrogenase 2 (NADP+), mitochondrial | 3418 | ENSG00000182054 |
| 12 | O60313 | NA | OPA1 | optic atrophy 1 (autosomal dominant) | 4976 | ENSG00000198836 |
| 13 | P53701 | NA | HCCS | holocytochrome c synthase | 3052 | ENSG00000004961 |
| 14 | P21796 | NA | VDAC1 | voltage-dependent anion channel 1 | 7416 | ENSG00000213585 |
| 15 | P31040 | NA | SDHA | succinate dehydrogenase complex, subunit A, flavoprotein (Fp) | 6389 | ENSG00000073578 |
| 16 | Q9UJZ1 | NA | STOML2 | stomatin (EPB72)-like 2 | 30968 | ENSG00000165283 |
| 17 | P12235 | NA | SLC25A4 | solute carrier family 25 (mitochondrial carrier; adenine nucleotide translocator), member 4 | 291 | ENSG00000151729 |
| 18 | P42765 | NA | ACAA2 | acetyl-CoA acyltransferase 2 | 10449 | ENSG00000167315 |
| 19 | P24752 | NA | ACAT1 | acetyl-CoA acetyltransferase 1 | 38 | ENSG00000075239 |
| 20 | Q02978 | NA | SLC25A11 | solute carrier family 25 (mitochondrial carrier; oxoglutarate carrier), member 11 | 8402 | ENSG00000108528 |
| 21 | P25705 | NA | ATP5A1 | ATP synthase, H+ transporting, mitochondrial F1 complex, alpha subunit 1, cardiac muscle | 498 | ENSG00000152234 |
| 22 | P49748 | NA | ACADVL | acyl-CoA dehydrogenase, very long chain | 37 | ENSG00000072778 |
| 23 | P48047 | NA | ATP5O | ATP synthase, H+ transporting, mitochondrial F1 complex, O subunit | 539 | ENSG00000241837 |
| 24 | Q9BPW8 | NA | NIPSNAP1 | nipsnap homolog 1 (C. elegans) | 8508 | ENSG00000184117 |

  
  

| **Database:cellular component      &nbspName:organelle envelope      &nbspID:GO:0031967** | | | | | | |
| --- | --- | --- | --- | --- | --- | --- |
| C=817; O=31; E=5.29; R=5.86; rawP=4.27e-16; adjP=5.00e-15 | | | | | | |
| Index | UserID | Value | Gene Symbol | Gene Name | EntrezGene | Ensembl |
| 1 | O75439 | NA | PMPCB | peptidase (mitochondrial processing) beta | 9512 | ENSG00000105819 |
| 2 | P54819 | NA | AK2 | adenylate kinase 2 | 204 | ENSG00000004455 |
| 3 | P17931 | NA | LGALS3 | lectin, galactoside-binding, soluble, 3 | 3958 | ENSG00000131981 |
| 4 | Q99623 | NA | PHB2 | prohibitin 2 | 11331 | ENSG00000215021 |
| 5 | P40939 | NA | HADHA | hydroxyacyl-CoA dehydrogenase/3-ketoacyl-CoA thiolase/enoyl-CoA hydratase (trifunctional protein), alpha subunit | 3030 | ENSG00000084754 |
| 6 | P40926 | NA | MDH2 | malate dehydrogenase 2, NAD (mitochondrial) | 4191 | ENSG00000146701 |
| 7 | P35232 | NA | PHB | prohibitin | 5245 | ENSG00000167085 |
| 8 | Q99714 | NA | HSD17B10 | hydroxysteroid (17-beta) dehydrogenase 10 | 3028 | ENSG00000072506 |
| 9 | O95831 | NA | AIFM1 | apoptosis-inducing factor, mitochondrion-associated, 1 | 9131 | ENSG00000156709 |
| 10 | P04179 | NA | SOD2 | superoxide dismutase 2, mitochondrial | 6648 | ENSG00000112096 |
| 11 | P48735 | NA | IDH2 | isocitrate dehydrogenase 2 (NADP+), mitochondrial | 3418 | ENSG00000182054 |
| 12 | O60313 | NA | OPA1 | optic atrophy 1 (autosomal dominant) | 4976 | ENSG00000198836 |
| 13 | P21796 | NA | VDAC1 | voltage-dependent anion channel 1 | 7416 | ENSG00000213585 |
| 14 | P53701 | NA | HCCS | holocytochrome c synthase | 3052 | ENSG00000004961 |
| 15 | P31040 | NA | SDHA | succinate dehydrogenase complex, subunit A, flavoprotein (Fp) | 6389 | ENSG00000073578 |
| 16 | P04083 | NA | ANXA1 | annexin A1 | 301 | ENSG00000135046 |
| 17 | Q9UJZ1 | NA | STOML2 | stomatin (EPB72)-like 2 | 30968 | ENSG00000165283 |
| 18 | P04040 | NA | CAT | catalase | 847 | ENSG00000121691 |
| 19 | O75340 | NA | PDCD6 | programmed cell death 6 | 10016 | ENSG00000249915 |
| 20 | P12235 | NA | SLC25A4 | solute carrier family 25 (mitochondrial carrier; adenine nucleotide translocator), member 4 | 291 | ENSG00000151729 |
| 21 | P40121 | NA | CAPG | capping protein (actin filament), gelsolin-like | 822 | ENSG00000042493 |
| 22 | P42765 | NA | ACAA2 | acetyl-CoA acyltransferase 2 | 10449 | ENSG00000167315 |
| 23 | P24752 | NA | ACAT1 | acetyl-CoA acetyltransferase 1 | 38 | ENSG00000075239 |
| 24 | Q9Y277 | NA | VDAC3 | voltage-dependent anion channel 3 | 7419 | ENSG00000078668 |
| 25 | Q02978 | NA | SLC25A11 | solute carrier family 25 (mitochondrial carrier; oxoglutarate carrier), member 11 | 8402 | ENSG00000108528 |
| 26 | P19367 | NA | HK1 | hexokinase 1 | 3098 | ENSG00000156515 |
| 27 | P25705 | NA | ATP5A1 | ATP synthase, H+ transporting, mitochondrial F1 complex, alpha subunit 1, cardiac muscle | 498 | ENSG00000152234 |
| 28 | Q14764 | NA | MVP | major vault protein | 9961 | ENSG00000013364 |
| 29 | P49748 | NA | ACADVL | acyl-CoA dehydrogenase, very long chain | 37 | ENSG00000072778 |
| 30 | P48047 | NA | ATP5O | ATP synthase, H+ transporting, mitochondrial F1 complex, O subunit | 539 | ENSG00000241837 |
| 31 | Q9BPW8 | NA | NIPSNAP1 | nipsnap homolog 1 (C. elegans) | 8508 | ENSG00000184117 |

  
  

| **Database:cellular component      &nbspName:envelope      &nbspID:GO:0031975** | | | | | | |
| --- | --- | --- | --- | --- | --- | --- |
| C=829; O=31; E=5.37; R=5.78; rawP=6.40e-16; adjP=6.81e-15 | | | | | | |
| Index | UserID | Value | Gene Symbol | Gene Name | EntrezGene | Ensembl |
| 1 | O75439 | NA | PMPCB | peptidase (mitochondrial processing) beta | 9512 | ENSG00000105819 |
| 2 | P54819 | NA | AK2 | adenylate kinase 2 | 204 | ENSG00000004455 |
| 3 | P17931 | NA | LGALS3 | lectin, galactoside-binding, soluble, 3 | 3958 | ENSG00000131981 |
| 4 | Q99623 | NA | PHB2 | prohibitin 2 | 11331 | ENSG00000215021 |
| 5 | P40939 | NA | HADHA | hydroxyacyl-CoA dehydrogenase/3-ketoacyl-CoA thiolase/enoyl-CoA hydratase (trifunctional protein), alpha subunit | 3030 | ENSG00000084754 |
| 6 | P40926 | NA | MDH2 | malate dehydrogenase 2, NAD (mitochondrial) | 4191 | ENSG00000146701 |
| 7 | P35232 | NA | PHB | prohibitin | 5245 | ENSG00000167085 |
| 8 | Q99714 | NA | HSD17B10 | hydroxysteroid (17-beta) dehydrogenase 10 | 3028 | ENSG00000072506 |
| 9 | O95831 | NA | AIFM1 | apoptosis-inducing factor, mitochondrion-associated, 1 | 9131 | ENSG00000156709 |
| 10 | P04179 | NA | SOD2 | superoxide dismutase 2, mitochondrial | 6648 | ENSG00000112096 |
| 11 | P48735 | NA | IDH2 | isocitrate dehydrogenase 2 (NADP+), mitochondrial | 3418 | ENSG00000182054 |
| 12 | O60313 | NA | OPA1 | optic atrophy 1 (autosomal dominant) | 4976 | ENSG00000198836 |
| 13 | P21796 | NA | VDAC1 | voltage-dependent anion channel 1 | 7416 | ENSG00000213585 |
| 14 | P53701 | NA | HCCS | holocytochrome c synthase | 3052 | ENSG00000004961 |
| 15 | P31040 | NA | SDHA | succinate dehydrogenase complex, subunit A, flavoprotein (Fp) | 6389 | ENSG00000073578 |
| 16 | P04083 | NA | ANXA1 | annexin A1 | 301 | ENSG00000135046 |
| 17 | Q9UJZ1 | NA | STOML2 | stomatin (EPB72)-like 2 | 30968 | ENSG00000165283 |
| 18 | P04040 | NA | CAT | catalase | 847 | ENSG00000121691 |
| 19 | O75340 | NA | PDCD6 | programmed cell death 6 | 10016 | ENSG00000249915 |
| 20 | P12235 | NA | SLC25A4 | solute carrier family 25 (mitochondrial carrier; adenine nucleotide translocator), member 4 | 291 | ENSG00000151729 |
| 21 | P40121 | NA | CAPG | capping protein (actin filament), gelsolin-like | 822 | ENSG00000042493 |
| 22 | P42765 | NA | ACAA2 | acetyl-CoA acyltransferase 2 | 10449 | ENSG00000167315 |
| 23 | P24752 | NA | ACAT1 | acetyl-CoA acetyltransferase 1 | 38 | ENSG00000075239 |
| 24 | Q9Y277 | NA | VDAC3 | voltage-dependent anion channel 3 | 7419 | ENSG00000078668 |
| 25 | Q02978 | NA | SLC25A11 | solute carrier family 25 (mitochondrial carrier; oxoglutarate carrier), member 11 | 8402 | ENSG00000108528 |
| 26 | P19367 | NA | HK1 | hexokinase 1 | 3098 | ENSG00000156515 |
| 27 | P25705 | NA | ATP5A1 | ATP synthase, H+ transporting, mitochondrial F1 complex, alpha subunit 1, cardiac muscle | 498 | ENSG00000152234 |
| 28 | Q14764 | NA | MVP | major vault protein | 9961 | ENSG00000013364 |
| 29 | P49748 | NA | ACADVL | acyl-CoA dehydrogenase, very long chain | 37 | ENSG00000072778 |
| 30 | P48047 | NA | ATP5O | ATP synthase, H+ transporting, mitochondrial F1 complex, O subunit | 539 | ENSG00000241837 |
| 31 | Q9BPW8 | NA | NIPSNAP1 | nipsnap homolog 1 (C. elegans) | 8508 | ENSG00000184117 |

  
  

| **Database:cellular component      &nbspName:intracellular part      &nbspID:GO:0044424** | | | | | | |
| --- | --- | --- | --- | --- | --- | --- |
| C=12096; O=107; E=78.31; R=1.37; rawP=3.10e-14; adjP=3.02e-13 | | | | | | |
| Index | UserID | Value | Gene Symbol | Gene Name | EntrezGene | Ensembl |
| 1 | P10253 | NA | GAA | glucosidase, alpha; acid | 2548 | ENSG00000171298 |
| 2 | P68371 | NA | TUBB4B | tubulin, beta 4B class IVb | 10383 | ENSG00000188229 |
| 3 | Q9UL46 | NA | PSME2 | proteasome (prosome, macropain) activator subunit 2 (PA28 beta) | 5721 | ENSG00000100911 |
| 4 | P40926 | NA | MDH2 | malate dehydrogenase 2, NAD (mitochondrial) | 4191 | ENSG00000146701 |
| 5 | P04179 | NA | SOD2 | superoxide dismutase 2, mitochondrial | 6648 | ENSG00000112096 |
| 6 | Q13724 | NA | MOGS | mannosyl-oligosaccharide glucosidase | 7841 | ENSG00000115275 |
| 7 | P48735 | NA | IDH2 | isocitrate dehydrogenase 2 (NADP+), mitochondrial | 3418 | ENSG00000182054 |
| 8 | P00367 | NA | GLUD1 | glutamate dehydrogenase 1 | 2746 | ENSG00000148672 |
| 9 | Q06323 | NA | PSME1 | proteasome (prosome, macropain) activator subunit 1 (PA28 alpha) | 5720 | ENSG00000092010 |
| 10 | P53701 | NA | HCCS | holocytochrome c synthase | 3052 | ENSG00000004961 |
| 11 | P35580 | NA | MYH10 | myosin, heavy chain 10, non-muscle | 4628 | ENSG00000133026 |
| 12 | P31040 | NA | SDHA | succinate dehydrogenase complex, subunit A, flavoprotein (Fp) | 6389 | ENSG00000073578 |
| 13 | O75369 | NA | FLNB | filamin B, beta | 2317 | ENSG00000136068 |
| 14 | P24534 | NA | EEF1B2 | eukaryotic translation elongation factor 1 beta 2 | 1933 | ENSG00000114942 |
| 15 | O14773 | NA | TPP1 | tripeptidyl peptidase I | 1200 | ENSG00000166340 |
| 16 | P47895 | NA | ALDH1A3 | aldehyde dehydrogenase 1 family, member A3 | 220 | ENSG00000184254 |
| 17 | P42765 | NA | ACAA2 | acetyl-CoA acyltransferase 2 | 10449 | ENSG00000167315 |
| 18 | P40121 | NA | CAPG | capping protein (actin filament), gelsolin-like | 822 | ENSG00000042493 |
| 19 | P13804 | NA | ETFA | electron-transfer-flavoprotein, alpha polypeptide | 2108 | ENSG00000140374 |
| 20 | Q9Y277 | NA | VDAC3 | voltage-dependent anion channel 3 | 7419 | ENSG00000078668 |
| 21 | P17174 | NA | GOT1 | glutamic-oxaloacetic transaminase 1, soluble (aspartate aminotransferase 1) | 2805 | ENSG00000120053 |
| 22 | P08727 | NA | KRT19 | keratin 19 | 3880 | ENSG00000171345 |
| 23 | Q04837 | NA | SSBP1 | single-stranded DNA binding protein 1, mitochondrial | 6742 | ENSG00000106028 |
| 24 | P50454 | NA | SERPINH1 | serpin peptidase inhibitor, clade H (heat shock protein 47), member 1, (collagen binding protein 1) | 871 | ENSG00000149257 |
| 25 | Q14697 | NA | GANAB | glucosidase, alpha; neutral AB | 23193 | ENSG00000089597 |
| 26 | P18206 | NA | VCL | vinculin | 7414 | ENSG00000035403 |
| 27 | P08195 | NA | SLC3A2 | solute carrier family 3 (activators of dibasic and neutral amino acid transport), member 2 | 6520 | ENSG00000168003 |
| 28 | P40939 | NA | HADHA | hydroxyacyl-CoA dehydrogenase/3-ketoacyl-CoA thiolase/enoyl-CoA hydratase (trifunctional protein), alpha subunit | 3030 | ENSG00000084754 |
| 29 | P09382 | NA | LGALS1 | lectin, galactoside-binding, soluble, 1 | 3956 | ENSG00000100097 |
| 30 | P52815 | NA | MRPL12 | mitochondrial ribosomal protein L12 | 6182 | ENSG00000262814 |
| 31 | Q9BXW7 | NA | CECR5 | cat eye syndrome chromosome region, candidate 5 | 27440 | ENSG00000069998 |
| 32 | P21796 | NA | VDAC1 | voltage-dependent anion channel 1 | 7416 | ENSG00000213585 |
| 33 | P38571 | NA | LIPA | lipase A, lysosomal acid, cholesterol esterase | 3988 | ENSG00000107798 |
| 34 | P04080 | NA | CSTB | cystatin B (stefin B) | 1476 | ENSG00000160213 |
| 35 | Q9UJZ1 | NA | STOML2 | stomatin (EPB72)-like 2 | 30968 | ENSG00000165283 |
| 36 | P32322 | NA | PYCR1 | pyrroline-5-carboxylate reductase 1 | 5831 | ENSG00000183010 |
| 37 | Q14980 | NA | NUMA1 | nuclear mitotic apparatus protein 1 | 4926 | ENSG00000137497 |
| 38 | Q99798 | NA | ACO2 | aconitase 2, mitochondrial | 50 | ENSG00000100412 |
| 39 | P51649 | NA | ALDH5A1 | aldehyde dehydrogenase 5 family, member A1 | 7915 | ENSG00000112294 |
| 40 | P12235 | NA | SLC25A4 | solute carrier family 25 (mitochondrial carrier; adenine nucleotide translocator), member 4 | 291 | ENSG00000151729 |
| 41 | P27816 | NA | MAP4 | microtubule-associated protein 4 | 4134 | ENSG00000047849 |
| 42 | Q6PI48 | NA | DARS2 | aspartyl-tRNA synthetase 2, mitochondrial | 55157 | ENSG00000117593 |
| 43 | Q9NSE4 | NA | IARS2 | isoleucyl-tRNA synthetase 2, mitochondrial | 55699 | ENSG00000067704 |
| 44 | P05556 | NA | ITGB1 | integrin, beta 1 (fibronectin receptor, beta polypeptide, antigen CD29 includes MDF2, MSK12) | 3688 | ENSG00000150093 |
| 45 | P13674 | NA | P4HA1 | prolyl 4-hydroxylase, alpha polypeptide I | 5033 | ENSG00000122884 |
| 46 | P40227 | NA | CCT6A | chaperonin containing TCP1, subunit 6A (zeta 1) | 908 | ENSG00000146731 |
| 47 | Q02978 | NA | SLC25A11 | solute carrier family 25 (mitochondrial carrier; oxoglutarate carrier), member 11 | 8402 | ENSG00000108528 |
| 48 | P60174 | NA | TPI1 | triosephosphate isomerase 1 | 7167 | ENSG00000111669 |
| 49 | P30042 | NA | C21orf33 | chromosome 21 open reading frame 33 | 8209 | ENSG00000160221 |
| 50 | P49589 | NA | CARS | cysteinyl-tRNA synthetase | 833 | ENSG00000110619 |
| 51 | P54819 | NA | AK2 | adenylate kinase 2 | 204 | ENSG00000004455 |
| 52 | P17931 | NA | LGALS3 | lectin, galactoside-binding, soluble, 3 | 3958 | ENSG00000131981 |
| 53 | P05787 | NA | KRT8 | keratin 8 | 3856 | ENSG00000170421 |
| 54 | Q99623 | NA | PHB2 | prohibitin 2 | 11331 | ENSG00000215021 |
| 55 | P50453 | NA | SERPINB9 | serpin peptidase inhibitor, clade B (ovalbumin), member 9 | 5272 | ENSG00000170542 |
| 56 | P35232 | NA | PHB | prohibitin | 5245 | ENSG00000167085 |
| 57 | O95831 | NA | AIFM1 | apoptosis-inducing factor, mitochondrion-associated, 1 | 9131 | ENSG00000156709 |
| 58 | Q99714 | NA | HSD17B10 | hydroxysteroid (17-beta) dehydrogenase 10 | 3028 | ENSG00000072506 |
| 59 | P61313 | NA | RPL15 | ribosomal protein L15 | 6138 | ENSG00000174748 |
| 60 | P07741 | NA | APRT | adenine phosphoribosyltransferase | 353 | ENSG00000198931 |
| 61 | O60313 | NA | OPA1 | optic atrophy 1 (autosomal dominant) | 4976 | ENSG00000198836 |
| 62 | P35270 | NA | SPR | sepiapterin reductase (7,8-dihydrobiopterin:NADP+ oxidoreductase) | 6697 | ENSG00000116096 |
| 63 | P68104 | NA | EEF1A1 | eukaryotic translation elongation factor 1 alpha 1 | 1915 | ENSG00000156508 |
| 64 | P38117 | NA | ETFB | electron-transfer-flavoprotein, beta polypeptide | 2109 | ENSG00000105379 |
| 65 | P02786 | NA | TFRC | transferrin receptor (p90, CD71) | 7037 | ENSG00000072274 |
| 66 | P43490 | NA | NAMPT | nicotinamide phosphoribosyltransferase | 10135 | ENSG00000105835 |
| 67 | P22102 | NA | GART | phosphoribosylglycinamide formyltransferase, phosphoribosylglycinamide synthetase, phosphoribosylaminoimidazole synthetase | 2618 | ENSG00000159131 |
| 68 | P13639 | NA | EEF2 | eukaryotic translation elongation factor 2 | 1938 | ENSG00000167658 |
| 69 | O75521 | NA | ECI2 | enoyl-CoA delta isomerase 2 | 10455 | ENSG00000198721 |
| 70 | O75367 | NA | H2AFY | H2A histone family, member Y | 9555 | ENSG00000113648 |
| 71 | P49419 | NA | ALDH7A1 | aldehyde dehydrogenase 7 family, member A1 | 501 | ENSG00000164904 |
| 72 | P67809 | NA | YBX1 | Y box binding protein 1 | 4904 | ENSG00000065978 |
| 73 | Q14764 | NA | MVP | major vault protein | 9961 | ENSG00000013364 |
| 74 | Q6PIU2 | NA | NCEH1 | neutral cholesterol ester hydrolase 1 | 57552 | ENSG00000144959 |
| 75 | P29692 | NA | EEF1D | eukaryotic translation elongation factor 1 delta (guanine nucleotide exchange protein) | 1936 | ENSG00000104529 |
| 76 | Q13813 | NA | SPTAN1 | spectrin, alpha, non-erythrocytic 1 | 6709 | ENSG00000197694 |
| 77 | Q5SSJ5 | NA | HP1BP3 | heterochromatin protein 1, binding protein 3 | 50809 | ENSG00000127483 |
| 78 | Q9BPW8 | NA | NIPSNAP1 | nipsnap homolog 1 (C. elegans) | 8508 | ENSG00000184117 |
| 79 | P11387 | NA | TOP1 | topoisomerase (DNA) I | 7150 | ENSG00000198900 |
| 80 | P02792 | NA | FTL | ferritin, light polypeptide | 2512 | ENSG00000087086 |
| 81 | O75439 | NA | PMPCB | peptidase (mitochondrial processing) beta | 9512 | ENSG00000105819 |
| 82 | Q9NR30 | NA | DDX21 | DEAD (Asp-Glu-Ala-Asp) box helicase 21 | 9188 | ENSG00000165732 |
| 83 | P00491 | NA | PNP | purine nucleoside phosphorylase | 4860 | ENSG00000198805 |
| 84 | P21953 | NA | BCKDHB | branched chain keto acid dehydrogenase E1, beta polypeptide | 594 | ENSG00000083123 |
| 85 | P30084 | NA | ECHS1 | enoyl CoA hydratase, short chain, 1, mitochondrial | 1892 | ENSG00000127884 |
| 86 | P60842 | NA | EIF4A1 | eukaryotic translation initiation factor 4A1 | 1973 | ENSG00000161960 |
| 87 | Q13011 | NA | ECH1 | enoyl CoA hydratase 1, peroxisomal | 1891 | ENSG00000104823 |
| 88 | P04083 | NA | ANXA1 | annexin A1 | 301 | ENSG00000135046 |
| 89 | P04040 | NA | CAT | catalase | 847 | ENSG00000121691 |
| 90 | O75340 | NA | PDCD6 | programmed cell death 6 | 10016 | ENSG00000249915 |
| 91 | P51688 | NA | SGSH | N-sulfoglucosamine sulfohydrolase | 6448 | ENSG00000181523 |
| 92 | P00558 | NA | PGK1 | phosphoglycerate kinase 1 | 5230 | ENSG00000102144 |
| 93 | P30048 | NA | PRDX3 | peroxiredoxin 3 | 10935 | ENSG00000165672 |
| 94 | P41091 | NA | EIF2S3 | eukaryotic translation initiation factor 2, subunit 3 gamma, 52kDa | 1968 | ENSG00000130741 |
| 95 | P02768 | NA | ALB | albumin | 213 | ENSG00000163631 |
| 96 | P07108 | NA | DBI | diazepam binding inhibitor (GABA receptor modulator, acyl-CoA binding protein) | 1622 | ENSG00000155368 |
| 97 | P24752 | NA | ACAT1 | acetyl-CoA acetyltransferase 1 | 38 | ENSG00000075239 |
| 98 | P19367 | NA | HK1 | hexokinase 1 | 3098 | ENSG00000156515 |
| 99 | P25705 | NA | ATP5A1 | ATP synthase, H+ transporting, mitochondrial F1 complex, alpha subunit 1, cardiac muscle | 498 | ENSG00000152234 |
| 100 | Q9BQ69 | NA | MACROD1 | MACRO domain containing 1 | 28992 | ENSG00000133315 |
| 101 | P05783 | NA | KRT18 | keratin 18 | 3875 | ENSG00000111057 |
| 102 | P06865 | NA | HEXA | hexosaminidase A (alpha polypeptide) | 3073 | ENSG00000213614 |
| 103 | P49748 | NA | ACADVL | acyl-CoA dehydrogenase, very long chain | 37 | ENSG00000072778 |
| 104 | P00352 | NA | ALDH1A1 | aldehyde dehydrogenase 1 family, member A1 | 216 | ENSG00000165092 |
| 105 | P48047 | NA | ATP5O | ATP synthase, H+ transporting, mitochondrial F1 complex, O subunit | 539 | ENSG00000241837 |
| 106 | Q16531 | NA | DDB1 | damage-specific DNA binding protein 1, 127kDa | 1642 | ENSG00000167986 |
| 107 | P31937 | NA | HIBADH | 3-hydroxyisobutyrate dehydrogenase | 11112 | ENSG00000106049 |

  
  

| **Database:cellular component      &nbspName:intracellular      &nbspID:GO:0005622** | | | | | | |
| --- | --- | --- | --- | --- | --- | --- |
| C=12412; O=107; E=80.36; R=1.33; rawP=4.63e-13; adjP=4.17e-12 | | | | | | |
| Index | UserID | Value | Gene Symbol | Gene Name | EntrezGene | Ensembl |
| 1 | P10253 | NA | GAA | glucosidase, alpha; acid | 2548 | ENSG00000171298 |
| 2 | P68371 | NA | TUBB4B | tubulin, beta 4B class IVb | 10383 | ENSG00000188229 |
| 3 | Q9UL46 | NA | PSME2 | proteasome (prosome, macropain) activator subunit 2 (PA28 beta) | 5721 | ENSG00000100911 |
| 4 | P40926 | NA | MDH2 | malate dehydrogenase 2, NAD (mitochondrial) | 4191 | ENSG00000146701 |
| 5 | P04179 | NA | SOD2 | superoxide dismutase 2, mitochondrial | 6648 | ENSG00000112096 |
| 6 | Q13724 | NA | MOGS | mannosyl-oligosaccharide glucosidase | 7841 | ENSG00000115275 |
| 7 | P48735 | NA | IDH2 | isocitrate dehydrogenase 2 (NADP+), mitochondrial | 3418 | ENSG00000182054 |
| 8 | P00367 | NA | GLUD1 | glutamate dehydrogenase 1 | 2746 | ENSG00000148672 |
| 9 | Q06323 | NA | PSME1 | proteasome (prosome, macropain) activator subunit 1 (PA28 alpha) | 5720 | ENSG00000092010 |
| 10 | P53701 | NA | HCCS | holocytochrome c synthase | 3052 | ENSG00000004961 |
| 11 | P35580 | NA | MYH10 | myosin, heavy chain 10, non-muscle | 4628 | ENSG00000133026 |
| 12 | P31040 | NA | SDHA | succinate dehydrogenase complex, subunit A, flavoprotein (Fp) | 6389 | ENSG00000073578 |
| 13 | O75369 | NA | FLNB | filamin B, beta | 2317 | ENSG00000136068 |
| 14 | P24534 | NA | EEF1B2 | eukaryotic translation elongation factor 1 beta 2 | 1933 | ENSG00000114942 |
| 15 | O14773 | NA | TPP1 | tripeptidyl peptidase I | 1200 | ENSG00000166340 |
| 16 | P47895 | NA | ALDH1A3 | aldehyde dehydrogenase 1 family, member A3 | 220 | ENSG00000184254 |
| 17 | P42765 | NA | ACAA2 | acetyl-CoA acyltransferase 2 | 10449 | ENSG00000167315 |
| 18 | P40121 | NA | CAPG | capping protein (actin filament), gelsolin-like | 822 | ENSG00000042493 |
| 19 | P13804 | NA | ETFA | electron-transfer-flavoprotein, alpha polypeptide | 2108 | ENSG00000140374 |
| 20 | Q9Y277 | NA | VDAC3 | voltage-dependent anion channel 3 | 7419 | ENSG00000078668 |
| 21 | P17174 | NA | GOT1 | glutamic-oxaloacetic transaminase 1, soluble (aspartate aminotransferase 1) | 2805 | ENSG00000120053 |
| 22 | P08727 | NA | KRT19 | keratin 19 | 3880 | ENSG00000171345 |
| 23 | Q04837 | NA | SSBP1 | single-stranded DNA binding protein 1, mitochondrial | 6742 | ENSG00000106028 |
| 24 | P50454 | NA | SERPINH1 | serpin peptidase inhibitor, clade H (heat shock protein 47), member 1, (collagen binding protein 1) | 871 | ENSG00000149257 |
| 25 | Q14697 | NA | GANAB | glucosidase, alpha; neutral AB | 23193 | ENSG00000089597 |
| 26 | P18206 | NA | VCL | vinculin | 7414 | ENSG00000035403 |
| 27 | P08195 | NA | SLC3A2 | solute carrier family 3 (activators of dibasic and neutral amino acid transport), member 2 | 6520 | ENSG00000168003 |
| 28 | P40939 | NA | HADHA | hydroxyacyl-CoA dehydrogenase/3-ketoacyl-CoA thiolase/enoyl-CoA hydratase (trifunctional protein), alpha subunit | 3030 | ENSG00000084754 |
| 29 | P09382 | NA | LGALS1 | lectin, galactoside-binding, soluble, 1 | 3956 | ENSG00000100097 |
| 30 | P52815 | NA | MRPL12 | mitochondrial ribosomal protein L12 | 6182 | ENSG00000262814 |
| 31 | Q9BXW7 | NA | CECR5 | cat eye syndrome chromosome region, candidate 5 | 27440 | ENSG00000069998 |
| 32 | P21796 | NA | VDAC1 | voltage-dependent anion channel 1 | 7416 | ENSG00000213585 |
| 33 | P38571 | NA | LIPA | lipase A, lysosomal acid, cholesterol esterase | 3988 | ENSG00000107798 |
| 34 | P04080 | NA | CSTB | cystatin B (stefin B) | 1476 | ENSG00000160213 |
| 35 | Q9UJZ1 | NA | STOML2 | stomatin (EPB72)-like 2 | 30968 | ENSG00000165283 |
| 36 | P32322 | NA | PYCR1 | pyrroline-5-carboxylate reductase 1 | 5831 | ENSG00000183010 |
| 37 | Q14980 | NA | NUMA1 | nuclear mitotic apparatus protein 1 | 4926 | ENSG00000137497 |
| 38 | Q99798 | NA | ACO2 | aconitase 2, mitochondrial | 50 | ENSG00000100412 |
| 39 | P51649 | NA | ALDH5A1 | aldehyde dehydrogenase 5 family, member A1 | 7915 | ENSG00000112294 |
| 40 | P12235 | NA | SLC25A4 | solute carrier family 25 (mitochondrial carrier; adenine nucleotide translocator), member 4 | 291 | ENSG00000151729 |
| 41 | P27816 | NA | MAP4 | microtubule-associated protein 4 | 4134 | ENSG00000047849 |
| 42 | Q6PI48 | NA | DARS2 | aspartyl-tRNA synthetase 2, mitochondrial | 55157 | ENSG00000117593 |
| 43 | Q9NSE4 | NA | IARS2 | isoleucyl-tRNA synthetase 2, mitochondrial | 55699 | ENSG00000067704 |
| 44 | P05556 | NA | ITGB1 | integrin, beta 1 (fibronectin receptor, beta polypeptide, antigen CD29 includes MDF2, MSK12) | 3688 | ENSG00000150093 |
| 45 | P13674 | NA | P4HA1 | prolyl 4-hydroxylase, alpha polypeptide I | 5033 | ENSG00000122884 |
| 46 | P40227 | NA | CCT6A | chaperonin containing TCP1, subunit 6A (zeta 1) | 908 | ENSG00000146731 |
| 47 | Q02978 | NA | SLC25A11 | solute carrier family 25 (mitochondrial carrier; oxoglutarate carrier), member 11 | 8402 | ENSG00000108528 |
| 48 | P60174 | NA | TPI1 | triosephosphate isomerase 1 | 7167 | ENSG00000111669 |
| 49 | P30042 | NA | C21orf33 | chromosome 21 open reading frame 33 | 8209 | ENSG00000160221 |
| 50 | P49589 | NA | CARS | cysteinyl-tRNA synthetase | 833 | ENSG00000110619 |
| 51 | P54819 | NA | AK2 | adenylate kinase 2 | 204 | ENSG00000004455 |
| 52 | P17931 | NA | LGALS3 | lectin, galactoside-binding, soluble, 3 | 3958 | ENSG00000131981 |
| 53 | P05787 | NA | KRT8 | keratin 8 | 3856 | ENSG00000170421 |
| 54 | Q99623 | NA | PHB2 | prohibitin 2 | 11331 | ENSG00000215021 |
| 55 | P50453 | NA | SERPINB9 | serpin peptidase inhibitor, clade B (ovalbumin), member 9 | 5272 | ENSG00000170542 |
| 56 | P35232 | NA | PHB | prohibitin | 5245 | ENSG00000167085 |
| 57 | O95831 | NA | AIFM1 | apoptosis-inducing factor, mitochondrion-associated, 1 | 9131 | ENSG00000156709 |
| 58 | Q99714 | NA | HSD17B10 | hydroxysteroid (17-beta) dehydrogenase 10 | 3028 | ENSG00000072506 |
| 59 | P61313 | NA | RPL15 | ribosomal protein L15 | 6138 | ENSG00000174748 |
| 60 | P07741 | NA | APRT | adenine phosphoribosyltransferase | 353 | ENSG00000198931 |
| 61 | O60313 | NA | OPA1 | optic atrophy 1 (autosomal dominant) | 4976 | ENSG00000198836 |
| 62 | P35270 | NA | SPR | sepiapterin reductase (7,8-dihydrobiopterin:NADP+ oxidoreductase) | 6697 | ENSG00000116096 |
| 63 | P68104 | NA | EEF1A1 | eukaryotic translation elongation factor 1 alpha 1 | 1915 | ENSG00000156508 |
| 64 | P38117 | NA | ETFB | electron-transfer-flavoprotein, beta polypeptide | 2109 | ENSG00000105379 |
| 65 | P02786 | NA | TFRC | transferrin receptor (p90, CD71) | 7037 | ENSG00000072274 |
| 66 | P43490 | NA | NAMPT | nicotinamide phosphoribosyltransferase | 10135 | ENSG00000105835 |
| 67 | P22102 | NA | GART | phosphoribosylglycinamide formyltransferase, phosphoribosylglycinamide synthetase, phosphoribosylaminoimidazole synthetase | 2618 | ENSG00000159131 |
| 68 | P13639 | NA | EEF2 | eukaryotic translation elongation factor 2 | 1938 | ENSG00000167658 |
| 69 | O75521 | NA | ECI2 | enoyl-CoA delta isomerase 2 | 10455 | ENSG00000198721 |
| 70 | O75367 | NA | H2AFY | H2A histone family, member Y | 9555 | ENSG00000113648 |
| 71 | P49419 | NA | ALDH7A1 | aldehyde dehydrogenase 7 family, member A1 | 501 | ENSG00000164904 |
| 72 | P67809 | NA | YBX1 | Y box binding protein 1 | 4904 | ENSG00000065978 |
| 73 | Q14764 | NA | MVP | major vault protein | 9961 | ENSG00000013364 |
| 74 | Q6PIU2 | NA | NCEH1 | neutral cholesterol ester hydrolase 1 | 57552 | ENSG00000144959 |
| 75 | P29692 | NA | EEF1D | eukaryotic translation elongation factor 1 delta (guanine nucleotide exchange protein) | 1936 | ENSG00000104529 |
| 76 | Q13813 | NA | SPTAN1 | spectrin, alpha, non-erythrocytic 1 | 6709 | ENSG00000197694 |
| 77 | Q5SSJ5 | NA | HP1BP3 | heterochromatin protein 1, binding protein 3 | 50809 | ENSG00000127483 |
| 78 | Q9BPW8 | NA | NIPSNAP1 | nipsnap homolog 1 (C. elegans) | 8508 | ENSG00000184117 |
| 79 | P11387 | NA | TOP1 | topoisomerase (DNA) I | 7150 | ENSG00000198900 |
| 80 | P02792 | NA | FTL | ferritin, light polypeptide | 2512 | ENSG00000087086 |
| 81 | O75439 | NA | PMPCB | peptidase (mitochondrial processing) beta | 9512 | ENSG00000105819 |
| 82 | Q9NR30 | NA | DDX21 | DEAD (Asp-Glu-Ala-Asp) box helicase 21 | 9188 | ENSG00000165732 |
| 83 | P00491 | NA | PNP | purine nucleoside phosphorylase | 4860 | ENSG00000198805 |
| 84 | P21953 | NA | BCKDHB | branched chain keto acid dehydrogenase E1, beta polypeptide | 594 | ENSG00000083123 |
| 85 | P30084 | NA | ECHS1 | enoyl CoA hydratase, short chain, 1, mitochondrial | 1892 | ENSG00000127884 |
| 86 | P60842 | NA | EIF4A1 | eukaryotic translation initiation factor 4A1 | 1973 | ENSG00000161960 |
| 87 | Q13011 | NA | ECH1 | enoyl CoA hydratase 1, peroxisomal | 1891 | ENSG00000104823 |
| 88 | P04083 | NA | ANXA1 | annexin A1 | 301 | ENSG00000135046 |
| 89 | P04040 | NA | CAT | catalase | 847 | ENSG00000121691 |
| 90 | O75340 | NA | PDCD6 | programmed cell death 6 | 10016 | ENSG00000249915 |
| 91 | P51688 | NA | SGSH | N-sulfoglucosamine sulfohydrolase | 6448 | ENSG00000181523 |
| 92 | P00558 | NA | PGK1 | phosphoglycerate kinase 1 | 5230 | ENSG00000102144 |
| 93 | P30048 | NA | PRDX3 | peroxiredoxin 3 | 10935 | ENSG00000165672 |
| 94 | P41091 | NA | EIF2S3 | eukaryotic translation initiation factor 2, subunit 3 gamma, 52kDa | 1968 | ENSG00000130741 |
| 95 | P02768 | NA | ALB | albumin | 213 | ENSG00000163631 |
| 96 | P07108 | NA | DBI | diazepam binding inhibitor (GABA receptor modulator, acyl-CoA binding protein) | 1622 | ENSG00000155368 |
| 97 | P24752 | NA | ACAT1 | acetyl-CoA acetyltransferase 1 | 38 | ENSG00000075239 |
| 98 | P19367 | NA | HK1 | hexokinase 1 | 3098 | ENSG00000156515 |
| 99 | P25705 | NA | ATP5A1 | ATP synthase, H+ transporting, mitochondrial F1 complex, alpha subunit 1, cardiac muscle | 498 | ENSG00000152234 |
| 100 | Q9BQ69 | NA | MACROD1 | MACRO domain containing 1 | 28992 | ENSG00000133315 |
| 101 | P05783 | NA | KRT18 | keratin 18 | 3875 | ENSG00000111057 |
| 102 | P06865 | NA | HEXA | hexosaminidase A (alpha polypeptide) | 3073 | ENSG00000213614 |
| 103 | P49748 | NA | ACADVL | acyl-CoA dehydrogenase, very long chain | 37 | ENSG00000072778 |
| 104 | P00352 | NA | ALDH1A1 | aldehyde dehydrogenase 1 family, member A1 | 216 | ENSG00000165092 |
| 105 | P48047 | NA | ATP5O | ATP synthase, H+ transporting, mitochondrial F1 complex, O subunit | 539 | ENSG00000241837 |
| 106 | Q16531 | NA | DDB1 | damage-specific DNA binding protein 1, 127kDa | 1642 | ENSG00000167986 |
| 107 | P31937 | NA | HIBADH | 3-hydroxyisobutyrate dehydrogenase | 11112 | ENSG00000106049 |

  
  

| **Database:cellular component      &nbspName:intracellular organelle part      &nbspID:GO:0044446** | | | | | | |
| --- | --- | --- | --- | --- | --- | --- |
| C=6690; O=79; E=43.31; R=1.82; rawP=2.95e-12; adjP=2.47e-11 | | | | | | |
| Index | UserID | Value | Gene Symbol | Gene Name | EntrezGene | Ensembl |
| 1 | P10253 | NA | GAA | glucosidase, alpha; acid | 2548 | ENSG00000171298 |
| 2 | P68371 | NA | TUBB4B | tubulin, beta 4B class IVb | 10383 | ENSG00000188229 |
| 3 | Q9UL46 | NA | PSME2 | proteasome (prosome, macropain) activator subunit 2 (PA28 beta) | 5721 | ENSG00000100911 |
| 4 | P40926 | NA | MDH2 | malate dehydrogenase 2, NAD (mitochondrial) | 4191 | ENSG00000146701 |
| 5 | P04179 | NA | SOD2 | superoxide dismutase 2, mitochondrial | 6648 | ENSG00000112096 |
| 6 | Q13724 | NA | MOGS | mannosyl-oligosaccharide glucosidase | 7841 | ENSG00000115275 |
| 7 | P48735 | NA | IDH2 | isocitrate dehydrogenase 2 (NADP+), mitochondrial | 3418 | ENSG00000182054 |
| 8 | P00367 | NA | GLUD1 | glutamate dehydrogenase 1 | 2746 | ENSG00000148672 |
| 9 | Q06323 | NA | PSME1 | proteasome (prosome, macropain) activator subunit 1 (PA28 alpha) | 5720 | ENSG00000092010 |
| 10 | P53701 | NA | HCCS | holocytochrome c synthase | 3052 | ENSG00000004961 |
| 11 | P35580 | NA | MYH10 | myosin, heavy chain 10, non-muscle | 4628 | ENSG00000133026 |
| 12 | P31040 | NA | SDHA | succinate dehydrogenase complex, subunit A, flavoprotein (Fp) | 6389 | ENSG00000073578 |
| 13 | O75369 | NA | FLNB | filamin B, beta | 2317 | ENSG00000136068 |
| 14 | O14773 | NA | TPP1 | tripeptidyl peptidase I | 1200 | ENSG00000166340 |
| 15 | P42765 | NA | ACAA2 | acetyl-CoA acyltransferase 2 | 10449 | ENSG00000167315 |
| 16 | P13804 | NA | ETFA | electron-transfer-flavoprotein, alpha polypeptide | 2108 | ENSG00000140374 |
| 17 | P40121 | NA | CAPG | capping protein (actin filament), gelsolin-like | 822 | ENSG00000042493 |
| 18 | Q9Y277 | NA | VDAC3 | voltage-dependent anion channel 3 | 7419 | ENSG00000078668 |
| 19 | P08727 | NA | KRT19 | keratin 19 | 3880 | ENSG00000171345 |
| 20 | Q04837 | NA | SSBP1 | single-stranded DNA binding protein 1, mitochondrial | 6742 | ENSG00000106028 |
| 21 | P50454 | NA | SERPINH1 | serpin peptidase inhibitor, clade H (heat shock protein 47), member 1, (collagen binding protein 1) | 871 | ENSG00000149257 |
| 22 | Q14697 | NA | GANAB | glucosidase, alpha; neutral AB | 23193 | ENSG00000089597 |
| 23 | P18206 | NA | VCL | vinculin | 7414 | ENSG00000035403 |
| 24 | P40939 | NA | HADHA | hydroxyacyl-CoA dehydrogenase/3-ketoacyl-CoA thiolase/enoyl-CoA hydratase (trifunctional protein), alpha subunit | 3030 | ENSG00000084754 |
| 25 | P52815 | NA | MRPL12 | mitochondrial ribosomal protein L12 | 6182 | ENSG00000262814 |
| 26 | P21796 | NA | VDAC1 | voltage-dependent anion channel 1 | 7416 | ENSG00000213585 |
| 27 | P04080 | NA | CSTB | cystatin B (stefin B) | 1476 | ENSG00000160213 |
| 28 | P32322 | NA | PYCR1 | pyrroline-5-carboxylate reductase 1 | 5831 | ENSG00000183010 |
| 29 | Q9UJZ1 | NA | STOML2 | stomatin (EPB72)-like 2 | 30968 | ENSG00000165283 |
| 30 | Q14980 | NA | NUMA1 | nuclear mitotic apparatus protein 1 | 4926 | ENSG00000137497 |
| 31 | Q99798 | NA | ACO2 | aconitase 2, mitochondrial | 50 | ENSG00000100412 |
| 32 | P51649 | NA | ALDH5A1 | aldehyde dehydrogenase 5 family, member A1 | 7915 | ENSG00000112294 |
| 33 | P12235 | NA | SLC25A4 | solute carrier family 25 (mitochondrial carrier; adenine nucleotide translocator), member 4 | 291 | ENSG00000151729 |
| 34 | P27816 | NA | MAP4 | microtubule-associated protein 4 | 4134 | ENSG00000047849 |
| 35 | Q6PI48 | NA | DARS2 | aspartyl-tRNA synthetase 2, mitochondrial | 55157 | ENSG00000117593 |
| 36 | Q9NSE4 | NA | IARS2 | isoleucyl-tRNA synthetase 2, mitochondrial | 55699 | ENSG00000067704 |
| 37 | P13674 | NA | P4HA1 | prolyl 4-hydroxylase, alpha polypeptide I | 5033 | ENSG00000122884 |
| 38 | P40227 | NA | CCT6A | chaperonin containing TCP1, subunit 6A (zeta 1) | 908 | ENSG00000146731 |
| 39 | Q02978 | NA | SLC25A11 | solute carrier family 25 (mitochondrial carrier; oxoglutarate carrier), member 11 | 8402 | ENSG00000108528 |
| 40 | P54819 | NA | AK2 | adenylate kinase 2 | 204 | ENSG00000004455 |
| 41 | P17931 | NA | LGALS3 | lectin, galactoside-binding, soluble, 3 | 3958 | ENSG00000131981 |
| 42 | P05787 | NA | KRT8 | keratin 8 | 3856 | ENSG00000170421 |
| 43 | Q99623 | NA | PHB2 | prohibitin 2 | 11331 | ENSG00000215021 |
| 44 | P35232 | NA | PHB | prohibitin | 5245 | ENSG00000167085 |
| 45 | O95831 | NA | AIFM1 | apoptosis-inducing factor, mitochondrion-associated, 1 | 9131 | ENSG00000156709 |
| 46 | Q99714 | NA | HSD17B10 | hydroxysteroid (17-beta) dehydrogenase 10 | 3028 | ENSG00000072506 |
| 47 | P61313 | NA | RPL15 | ribosomal protein L15 | 6138 | ENSG00000174748 |
| 48 | P07741 | NA | APRT | adenine phosphoribosyltransferase | 353 | ENSG00000198931 |
| 49 | O60313 | NA | OPA1 | optic atrophy 1 (autosomal dominant) | 4976 | ENSG00000198836 |
| 50 | P35270 | NA | SPR | sepiapterin reductase (7,8-dihydrobiopterin:NADP+ oxidoreductase) | 6697 | ENSG00000116096 |
| 51 | P02786 | NA | TFRC | transferrin receptor (p90, CD71) | 7037 | ENSG00000072274 |
| 52 | P38117 | NA | ETFB | electron-transfer-flavoprotein, beta polypeptide | 2109 | ENSG00000105379 |
| 53 | O75367 | NA | H2AFY | H2A histone family, member Y | 9555 | ENSG00000113648 |
| 54 | O75521 | NA | ECI2 | enoyl-CoA delta isomerase 2 | 10455 | ENSG00000198721 |
| 55 | P67809 | NA | YBX1 | Y box binding protein 1 | 4904 | ENSG00000065978 |
| 56 | P49419 | NA | ALDH7A1 | aldehyde dehydrogenase 7 family, member A1 | 501 | ENSG00000164904 |
| 57 | Q14764 | NA | MVP | major vault protein | 9961 | ENSG00000013364 |
| 58 | P11387 | NA | TOP1 | topoisomerase (DNA) I | 7150 | ENSG00000198900 |
| 59 | Q9BPW8 | NA | NIPSNAP1 | nipsnap homolog 1 (C. elegans) | 8508 | ENSG00000184117 |
| 60 | Q5SSJ5 | NA | HP1BP3 | heterochromatin protein 1, binding protein 3 | 50809 | ENSG00000127483 |
| 61 | Q13813 | NA | SPTAN1 | spectrin, alpha, non-erythrocytic 1 | 6709 | ENSG00000197694 |
| 62 | Q9NR30 | NA | DDX21 | DEAD (Asp-Glu-Ala-Asp) box helicase 21 | 9188 | ENSG00000165732 |
| 63 | O75439 | NA | PMPCB | peptidase (mitochondrial processing) beta | 9512 | ENSG00000105819 |
| 64 | P21953 | NA | BCKDHB | branched chain keto acid dehydrogenase E1, beta polypeptide | 594 | ENSG00000083123 |
| 65 | P30084 | NA | ECHS1 | enoyl CoA hydratase, short chain, 1, mitochondrial | 1892 | ENSG00000127884 |
| 66 | P04083 | NA | ANXA1 | annexin A1 | 301 | ENSG00000135046 |
| 67 | P04040 | NA | CAT | catalase | 847 | ENSG00000121691 |
| 68 | O75340 | NA | PDCD6 | programmed cell death 6 | 10016 | ENSG00000249915 |
| 69 | P51688 | NA | SGSH | N-sulfoglucosamine sulfohydrolase | 6448 | ENSG00000181523 |
| 70 | P02768 | NA | ALB | albumin | 213 | ENSG00000163631 |
| 71 | P24752 | NA | ACAT1 | acetyl-CoA acetyltransferase 1 | 38 | ENSG00000075239 |
| 72 | P19367 | NA | HK1 | hexokinase 1 | 3098 | ENSG00000156515 |
| 73 | P25705 | NA | ATP5A1 | ATP synthase, H+ transporting, mitochondrial F1 complex, alpha subunit 1, cardiac muscle | 498 | ENSG00000152234 |
| 74 | P05783 | NA | KRT18 | keratin 18 | 3875 | ENSG00000111057 |
| 75 | P06865 | NA | HEXA | hexosaminidase A (alpha polypeptide) | 3073 | ENSG00000213614 |
| 76 | P49748 | NA | ACADVL | acyl-CoA dehydrogenase, very long chain | 37 | ENSG00000072778 |
| 77 | P48047 | NA | ATP5O | ATP synthase, H+ transporting, mitochondrial F1 complex, O subunit | 539 | ENSG00000241837 |
| 78 | Q16531 | NA | DDB1 | damage-specific DNA binding protein 1, 127kDa | 1642 | ENSG00000167986 |
| 79 | P31937 | NA | HIBADH | 3-hydroxyisobutyrate dehydrogenase | 11112 | ENSG00000106049 |

  
  

| **Database:cellular component      &nbspName:organelle part      &nbspID:GO:0044422** | | | | | | |
| --- | --- | --- | --- | --- | --- | --- |
| C=6777; O=79; E=43.87; R=1.80; rawP=6.43e-12; adjP=5.02e-11 | | | | | | |
| Index | UserID | Value | Gene Symbol | Gene Name | EntrezGene | Ensembl |
| 1 | P10253 | NA | GAA | glucosidase, alpha; acid | 2548 | ENSG00000171298 |
| 2 | P68371 | NA | TUBB4B | tubulin, beta 4B class IVb | 10383 | ENSG00000188229 |
| 3 | Q9UL46 | NA | PSME2 | proteasome (prosome, macropain) activator subunit 2 (PA28 beta) | 5721 | ENSG00000100911 |
| 4 | P40926 | NA | MDH2 | malate dehydrogenase 2, NAD (mitochondrial) | 4191 | ENSG00000146701 |
| 5 | P04179 | NA | SOD2 | superoxide dismutase 2, mitochondrial | 6648 | ENSG00000112096 |
| 6 | Q13724 | NA | MOGS | mannosyl-oligosaccharide glucosidase | 7841 | ENSG00000115275 |
| 7 | P48735 | NA | IDH2 | isocitrate dehydrogenase 2 (NADP+), mitochondrial | 3418 | ENSG00000182054 |
| 8 | P00367 | NA | GLUD1 | glutamate dehydrogenase 1 | 2746 | ENSG00000148672 |
| 9 | Q06323 | NA | PSME1 | proteasome (prosome, macropain) activator subunit 1 (PA28 alpha) | 5720 | ENSG00000092010 |
| 10 | P53701 | NA | HCCS | holocytochrome c synthase | 3052 | ENSG00000004961 |
| 11 | P35580 | NA | MYH10 | myosin, heavy chain 10, non-muscle | 4628 | ENSG00000133026 |
| 12 | P31040 | NA | SDHA | succinate dehydrogenase complex, subunit A, flavoprotein (Fp) | 6389 | ENSG00000073578 |
| 13 | O75369 | NA | FLNB | filamin B, beta | 2317 | ENSG00000136068 |
| 14 | O14773 | NA | TPP1 | tripeptidyl peptidase I | 1200 | ENSG00000166340 |
| 15 | P42765 | NA | ACAA2 | acetyl-CoA acyltransferase 2 | 10449 | ENSG00000167315 |
| 16 | P13804 | NA | ETFA | electron-transfer-flavoprotein, alpha polypeptide | 2108 | ENSG00000140374 |
| 17 | P40121 | NA | CAPG | capping protein (actin filament), gelsolin-like | 822 | ENSG00000042493 |
| 18 | Q9Y277 | NA | VDAC3 | voltage-dependent anion channel 3 | 7419 | ENSG00000078668 |
| 19 | P08727 | NA | KRT19 | keratin 19 | 3880 | ENSG00000171345 |
| 20 | Q04837 | NA | SSBP1 | single-stranded DNA binding protein 1, mitochondrial | 6742 | ENSG00000106028 |
| 21 | P50454 | NA | SERPINH1 | serpin peptidase inhibitor, clade H (heat shock protein 47), member 1, (collagen binding protein 1) | 871 | ENSG00000149257 |
| 22 | Q14697 | NA | GANAB | glucosidase, alpha; neutral AB | 23193 | ENSG00000089597 |
| 23 | P18206 | NA | VCL | vinculin | 7414 | ENSG00000035403 |
| 24 | P40939 | NA | HADHA | hydroxyacyl-CoA dehydrogenase/3-ketoacyl-CoA thiolase/enoyl-CoA hydratase (trifunctional protein), alpha subunit | 3030 | ENSG00000084754 |
| 25 | P52815 | NA | MRPL12 | mitochondrial ribosomal protein L12 | 6182 | ENSG00000262814 |
| 26 | P21796 | NA | VDAC1 | voltage-dependent anion channel 1 | 7416 | ENSG00000213585 |
| 27 | P04080 | NA | CSTB | cystatin B (stefin B) | 1476 | ENSG00000160213 |
| 28 | P32322 | NA | PYCR1 | pyrroline-5-carboxylate reductase 1 | 5831 | ENSG00000183010 |
| 29 | Q9UJZ1 | NA | STOML2 | stomatin (EPB72)-like 2 | 30968 | ENSG00000165283 |
| 30 | Q14980 | NA | NUMA1 | nuclear mitotic apparatus protein 1 | 4926 | ENSG00000137497 |
| 31 | Q99798 | NA | ACO2 | aconitase 2, mitochondrial | 50 | ENSG00000100412 |
| 32 | P51649 | NA | ALDH5A1 | aldehyde dehydrogenase 5 family, member A1 | 7915 | ENSG00000112294 |
| 33 | P12235 | NA | SLC25A4 | solute carrier family 25 (mitochondrial carrier; adenine nucleotide translocator), member 4 | 291 | ENSG00000151729 |
| 34 | P27816 | NA | MAP4 | microtubule-associated protein 4 | 4134 | ENSG00000047849 |
| 35 | Q6PI48 | NA | DARS2 | aspartyl-tRNA synthetase 2, mitochondrial | 55157 | ENSG00000117593 |
| 36 | Q9NSE4 | NA | IARS2 | isoleucyl-tRNA synthetase 2, mitochondrial | 55699 | ENSG00000067704 |
| 37 | P13674 | NA | P4HA1 | prolyl 4-hydroxylase, alpha polypeptide I | 5033 | ENSG00000122884 |
| 38 | P40227 | NA | CCT6A | chaperonin containing TCP1, subunit 6A (zeta 1) | 908 | ENSG00000146731 |
| 39 | Q02978 | NA | SLC25A11 | solute carrier family 25 (mitochondrial carrier; oxoglutarate carrier), member 11 | 8402 | ENSG00000108528 |
| 40 | P54819 | NA | AK2 | adenylate kinase 2 | 204 | ENSG00000004455 |
| 41 | P17931 | NA | LGALS3 | lectin, galactoside-binding, soluble, 3 | 3958 | ENSG00000131981 |
| 42 | P05787 | NA | KRT8 | keratin 8 | 3856 | ENSG00000170421 |
| 43 | Q99623 | NA | PHB2 | prohibitin 2 | 11331 | ENSG00000215021 |
| 44 | P35232 | NA | PHB | prohibitin | 5245 | ENSG00000167085 |
| 45 | O95831 | NA | AIFM1 | apoptosis-inducing factor, mitochondrion-associated, 1 | 9131 | ENSG00000156709 |
| 46 | Q99714 | NA | HSD17B10 | hydroxysteroid (17-beta) dehydrogenase 10 | 3028 | ENSG00000072506 |
| 47 | P61313 | NA | RPL15 | ribosomal protein L15 | 6138 | ENSG00000174748 |
| 48 | P07741 | NA | APRT | adenine phosphoribosyltransferase | 353 | ENSG00000198931 |
| 49 | O60313 | NA | OPA1 | optic atrophy 1 (autosomal dominant) | 4976 | ENSG00000198836 |
| 50 | P35270 | NA | SPR | sepiapterin reductase (7,8-dihydrobiopterin:NADP+ oxidoreductase) | 6697 | ENSG00000116096 |
| 51 | P02786 | NA | TFRC | transferrin receptor (p90, CD71) | 7037 | ENSG00000072274 |
| 52 | P38117 | NA | ETFB | electron-transfer-flavoprotein, beta polypeptide | 2109 | ENSG00000105379 |
| 53 | O75367 | NA | H2AFY | H2A histone family, member Y | 9555 | ENSG00000113648 |
| 54 | O75521 | NA | ECI2 | enoyl-CoA delta isomerase 2 | 10455 | ENSG00000198721 |
| 55 | P67809 | NA | YBX1 | Y box binding protein 1 | 4904 | ENSG00000065978 |
| 56 | P49419 | NA | ALDH7A1 | aldehyde dehydrogenase 7 family, member A1 | 501 | ENSG00000164904 |
| 57 | Q14764 | NA | MVP | major vault protein | 9961 | ENSG00000013364 |
| 58 | P11387 | NA | TOP1 | topoisomerase (DNA) I | 7150 | ENSG00000198900 |
| 59 | Q9BPW8 | NA | NIPSNAP1 | nipsnap homolog 1 (C. elegans) | 8508 | ENSG00000184117 |
| 60 | Q5SSJ5 | NA | HP1BP3 | heterochromatin protein 1, binding protein 3 | 50809 | ENSG00000127483 |
| 61 | Q13813 | NA | SPTAN1 | spectrin, alpha, non-erythrocytic 1 | 6709 | ENSG00000197694 |
| 62 | Q9NR30 | NA | DDX21 | DEAD (Asp-Glu-Ala-Asp) box helicase 21 | 9188 | ENSG00000165732 |
| 63 | O75439 | NA | PMPCB | peptidase (mitochondrial processing) beta | 9512 | ENSG00000105819 |
| 64 | P21953 | NA | BCKDHB | branched chain keto acid dehydrogenase E1, beta polypeptide | 594 | ENSG00000083123 |
| 65 | P30084 | NA | ECHS1 | enoyl CoA hydratase, short chain, 1, mitochondrial | 1892 | ENSG00000127884 |
| 66 | P04083 | NA | ANXA1 | annexin A1 | 301 | ENSG00000135046 |
| 67 | P04040 | NA | CAT | catalase | 847 | ENSG00000121691 |
| 68 | O75340 | NA | PDCD6 | programmed cell death 6 | 10016 | ENSG00000249915 |
| 69 | P51688 | NA | SGSH | N-sulfoglucosamine sulfohydrolase | 6448 | ENSG00000181523 |
| 70 | P02768 | NA | ALB | albumin | 213 | ENSG00000163631 |
| 71 | P24752 | NA | ACAT1 | acetyl-CoA acetyltransferase 1 | 38 | ENSG00000075239 |
| 72 | P19367 | NA | HK1 | hexokinase 1 | 3098 | ENSG00000156515 |
| 73 | P25705 | NA | ATP5A1 | ATP synthase, H+ transporting, mitochondrial F1 complex, alpha subunit 1, cardiac muscle | 498 | ENSG00000152234 |
| 74 | P05783 | NA | KRT18 | keratin 18 | 3875 | ENSG00000111057 |
| 75 | P06865 | NA | HEXA | hexosaminidase A (alpha polypeptide) | 3073 | ENSG00000213614 |
| 76 | P49748 | NA | ACADVL | acyl-CoA dehydrogenase, very long chain | 37 | ENSG00000072778 |
| 77 | P48047 | NA | ATP5O | ATP synthase, H+ transporting, mitochondrial F1 complex, O subunit | 539 | ENSG00000241837 |
| 78 | Q16531 | NA | DDB1 | damage-specific DNA binding protein 1, 127kDa | 1642 | ENSG00000167986 |
| 79 | P31937 | NA | HIBADH | 3-hydroxyisobutyrate dehydrogenase | 11112 | ENSG00000106049 |

  
  

| **Database:cellular component      &nbspName:membrane-enclosed lumen      &nbspID:GO:0031974** | | | | | | |
| --- | --- | --- | --- | --- | --- | --- |
| C=3375; O=52; E=21.85; R=2.38; rawP=6.92e-11; adjP=5.06e-10 | | | | | | |
| Index | UserID | Value | Gene Symbol | Gene Name | EntrezGene | Ensembl |
| 1 | P54819 | NA | AK2 | adenylate kinase 2 | 204 | ENSG00000004455 |
| 2 | P05787 | NA | KRT8 | keratin 8 | 3856 | ENSG00000170421 |
| 3 | Q9UL46 | NA | PSME2 | proteasome (prosome, macropain) activator subunit 2 (PA28 beta) | 5721 | ENSG00000100911 |
| 4 | P40926 | NA | MDH2 | malate dehydrogenase 2, NAD (mitochondrial) | 4191 | ENSG00000146701 |
| 5 | P35232 | NA | PHB | prohibitin | 5245 | ENSG00000167085 |
| 6 | Q99714 | NA | HSD17B10 | hydroxysteroid (17-beta) dehydrogenase 10 | 3028 | ENSG00000072506 |
| 7 | O95831 | NA | AIFM1 | apoptosis-inducing factor, mitochondrion-associated, 1 | 9131 | ENSG00000156709 |
| 8 | P04179 | NA | SOD2 | superoxide dismutase 2, mitochondrial | 6648 | ENSG00000112096 |
| 9 | P07741 | NA | APRT | adenine phosphoribosyltransferase | 353 | ENSG00000198931 |
| 10 | P00367 | NA | GLUD1 | glutamate dehydrogenase 1 | 2746 | ENSG00000148672 |
| 11 | Q06323 | NA | PSME1 | proteasome (prosome, macropain) activator subunit 1 (PA28 alpha) | 5720 | ENSG00000092010 |
| 12 | P48735 | NA | IDH2 | isocitrate dehydrogenase 2 (NADP+), mitochondrial | 3418 | ENSG00000182054 |
| 13 | O60313 | NA | OPA1 | optic atrophy 1 (autosomal dominant) | 4976 | ENSG00000198836 |
| 14 | P35270 | NA | SPR | sepiapterin reductase (7,8-dihydrobiopterin:NADP+ oxidoreductase) | 6697 | ENSG00000116096 |
| 15 | P38117 | NA | ETFB | electron-transfer-flavoprotein, beta polypeptide | 2109 | ENSG00000105379 |
| 16 | O14773 | NA | TPP1 | tripeptidyl peptidase I | 1200 | ENSG00000166340 |
| 17 | P13804 | NA | ETFA | electron-transfer-flavoprotein, alpha polypeptide | 2108 | ENSG00000140374 |
| 18 | P40121 | NA | CAPG | capping protein (actin filament), gelsolin-like | 822 | ENSG00000042493 |
| 19 | O75521 | NA | ECI2 | enoyl-CoA delta isomerase 2 | 10455 | ENSG00000198721 |
| 20 | O75367 | NA | H2AFY | H2A histone family, member Y | 9555 | ENSG00000113648 |
| 21 | P49419 | NA | ALDH7A1 | aldehyde dehydrogenase 7 family, member A1 | 501 | ENSG00000164904 |
| 22 | P67809 | NA | YBX1 | Y box binding protein 1 | 4904 | ENSG00000065978 |
| 23 | Q04837 | NA | SSBP1 | single-stranded DNA binding protein 1, mitochondrial | 6742 | ENSG00000106028 |
| 24 | P50454 | NA | SERPINH1 | serpin peptidase inhibitor, clade H (heat shock protein 47), member 1, (collagen binding protein 1) | 871 | ENSG00000149257 |
| 25 | Q14697 | NA | GANAB | glucosidase, alpha; neutral AB | 23193 | ENSG00000089597 |
| 26 | P11387 | NA | TOP1 | topoisomerase (DNA) I | 7150 | ENSG00000198900 |
| 27 | O75439 | NA | PMPCB | peptidase (mitochondrial processing) beta | 9512 | ENSG00000105819 |
| 28 | Q9NR30 | NA | DDX21 | DEAD (Asp-Glu-Ala-Asp) box helicase 21 | 9188 | ENSG00000165732 |
| 29 | P21953 | NA | BCKDHB | branched chain keto acid dehydrogenase E1, beta polypeptide | 594 | ENSG00000083123 |
| 30 | P40939 | NA | HADHA | hydroxyacyl-CoA dehydrogenase/3-ketoacyl-CoA thiolase/enoyl-CoA hydratase (trifunctional protein), alpha subunit | 3030 | ENSG00000084754 |
| 31 | P30084 | NA | ECHS1 | enoyl CoA hydratase, short chain, 1, mitochondrial | 1892 | ENSG00000127884 |
| 32 | P52815 | NA | MRPL12 | mitochondrial ribosomal protein L12 | 6182 | ENSG00000262814 |
| 33 | P21796 | NA | VDAC1 | voltage-dependent anion channel 1 | 7416 | ENSG00000213585 |
| 34 | P04080 | NA | CSTB | cystatin B (stefin B) | 1476 | ENSG00000160213 |
| 35 | P32322 | NA | PYCR1 | pyrroline-5-carboxylate reductase 1 | 5831 | ENSG00000183010 |
| 36 | P04040 | NA | CAT | catalase | 847 | ENSG00000121691 |
| 37 | Q14980 | NA | NUMA1 | nuclear mitotic apparatus protein 1 | 4926 | ENSG00000137497 |
| 38 | Q99798 | NA | ACO2 | aconitase 2, mitochondrial | 50 | ENSG00000100412 |
| 39 | P51688 | NA | SGSH | N-sulfoglucosamine sulfohydrolase | 6448 | ENSG00000181523 |
| 40 | P51649 | NA | ALDH5A1 | aldehyde dehydrogenase 5 family, member A1 | 7915 | ENSG00000112294 |
| 41 | P02768 | NA | ALB | albumin | 213 | ENSG00000163631 |
| 42 | Q9NSE4 | NA | IARS2 | isoleucyl-tRNA synthetase 2, mitochondrial | 55699 | ENSG00000067704 |
| 43 | Q6PI48 | NA | DARS2 | aspartyl-tRNA synthetase 2, mitochondrial | 55157 | ENSG00000117593 |
| 44 | P24752 | NA | ACAT1 | acetyl-CoA acetyltransferase 1 | 38 | ENSG00000075239 |
| 45 | P13674 | NA | P4HA1 | prolyl 4-hydroxylase, alpha polypeptide I | 5033 | ENSG00000122884 |
| 46 | P19367 | NA | HK1 | hexokinase 1 | 3098 | ENSG00000156515 |
| 47 | P25705 | NA | ATP5A1 | ATP synthase, H+ transporting, mitochondrial F1 complex, alpha subunit 1, cardiac muscle | 498 | ENSG00000152234 |
| 48 | P05783 | NA | KRT18 | keratin 18 | 3875 | ENSG00000111057 |
| 49 | P06865 | NA | HEXA | hexosaminidase A (alpha polypeptide) | 3073 | ENSG00000213614 |
| 50 | P49748 | NA | ACADVL | acyl-CoA dehydrogenase, very long chain | 37 | ENSG00000072778 |
| 51 | Q16531 | NA | DDB1 | damage-specific DNA binding protein 1, 127kDa | 1642 | ENSG00000167986 |
| 52 | P31937 | NA | HIBADH | 3-hydroxyisobutyrate dehydrogenase | 11112 | ENSG00000106049 |

  
  

| **Database:cellular component      &nbspName:organelle lumen      &nbspID:GO:0043233** | | | | | | |
| --- | --- | --- | --- | --- | --- | --- |
| C=3331; O=49; E=21.57; R=2.27; rawP=1.96e-09; adjP=1.35e-08 | | | | | | |
| Index | UserID | Value | Gene Symbol | Gene Name | EntrezGene | Ensembl |
| 1 | P05787 | NA | KRT8 | keratin 8 | 3856 | ENSG00000170421 |
| 2 | Q9UL46 | NA | PSME2 | proteasome (prosome, macropain) activator subunit 2 (PA28 beta) | 5721 | ENSG00000100911 |
| 3 | P40926 | NA | MDH2 | malate dehydrogenase 2, NAD (mitochondrial) | 4191 | ENSG00000146701 |
| 4 | P35232 | NA | PHB | prohibitin | 5245 | ENSG00000167085 |
| 5 | Q99714 | NA | HSD17B10 | hydroxysteroid (17-beta) dehydrogenase 10 | 3028 | ENSG00000072506 |
| 6 | P04179 | NA | SOD2 | superoxide dismutase 2, mitochondrial | 6648 | ENSG00000112096 |
| 7 | P07741 | NA | APRT | adenine phosphoribosyltransferase | 353 | ENSG00000198931 |
| 8 | P00367 | NA | GLUD1 | glutamate dehydrogenase 1 | 2746 | ENSG00000148672 |
| 9 | Q06323 | NA | PSME1 | proteasome (prosome, macropain) activator subunit 1 (PA28 alpha) | 5720 | ENSG00000092010 |
| 10 | P48735 | NA | IDH2 | isocitrate dehydrogenase 2 (NADP+), mitochondrial | 3418 | ENSG00000182054 |
| 11 | P35270 | NA | SPR | sepiapterin reductase (7,8-dihydrobiopterin:NADP+ oxidoreductase) | 6697 | ENSG00000116096 |
| 12 | P38117 | NA | ETFB | electron-transfer-flavoprotein, beta polypeptide | 2109 | ENSG00000105379 |
| 13 | O14773 | NA | TPP1 | tripeptidyl peptidase I | 1200 | ENSG00000166340 |
| 14 | P13804 | NA | ETFA | electron-transfer-flavoprotein, alpha polypeptide | 2108 | ENSG00000140374 |
| 15 | P40121 | NA | CAPG | capping protein (actin filament), gelsolin-like | 822 | ENSG00000042493 |
| 16 | O75521 | NA | ECI2 | enoyl-CoA delta isomerase 2 | 10455 | ENSG00000198721 |
| 17 | O75367 | NA | H2AFY | H2A histone family, member Y | 9555 | ENSG00000113648 |
| 18 | P49419 | NA | ALDH7A1 | aldehyde dehydrogenase 7 family, member A1 | 501 | ENSG00000164904 |
| 19 | P67809 | NA | YBX1 | Y box binding protein 1 | 4904 | ENSG00000065978 |
| 20 | Q04837 | NA | SSBP1 | single-stranded DNA binding protein 1, mitochondrial | 6742 | ENSG00000106028 |
| 21 | P50454 | NA | SERPINH1 | serpin peptidase inhibitor, clade H (heat shock protein 47), member 1, (collagen binding protein 1) | 871 | ENSG00000149257 |
| 22 | Q14697 | NA | GANAB | glucosidase, alpha; neutral AB | 23193 | ENSG00000089597 |
| 23 | P11387 | NA | TOP1 | topoisomerase (DNA) I | 7150 | ENSG00000198900 |
| 24 | O75439 | NA | PMPCB | peptidase (mitochondrial processing) beta | 9512 | ENSG00000105819 |
| 25 | Q9NR30 | NA | DDX21 | DEAD (Asp-Glu-Ala-Asp) box helicase 21 | 9188 | ENSG00000165732 |
| 26 | P21953 | NA | BCKDHB | branched chain keto acid dehydrogenase E1, beta polypeptide | 594 | ENSG00000083123 |
| 27 | P40939 | NA | HADHA | hydroxyacyl-CoA dehydrogenase/3-ketoacyl-CoA thiolase/enoyl-CoA hydratase (trifunctional protein), alpha subunit | 3030 | ENSG00000084754 |
| 28 | P30084 | NA | ECHS1 | enoyl CoA hydratase, short chain, 1, mitochondrial | 1892 | ENSG00000127884 |
| 29 | P52815 | NA | MRPL12 | mitochondrial ribosomal protein L12 | 6182 | ENSG00000262814 |
| 30 | P21796 | NA | VDAC1 | voltage-dependent anion channel 1 | 7416 | ENSG00000213585 |
| 31 | P04080 | NA | CSTB | cystatin B (stefin B) | 1476 | ENSG00000160213 |
| 32 | P32322 | NA | PYCR1 | pyrroline-5-carboxylate reductase 1 | 5831 | ENSG00000183010 |
| 33 | P04040 | NA | CAT | catalase | 847 | ENSG00000121691 |
| 34 | Q14980 | NA | NUMA1 | nuclear mitotic apparatus protein 1 | 4926 | ENSG00000137497 |
| 35 | Q99798 | NA | ACO2 | aconitase 2, mitochondrial | 50 | ENSG00000100412 |
| 36 | P51688 | NA | SGSH | N-sulfoglucosamine sulfohydrolase | 6448 | ENSG00000181523 |
| 37 | P51649 | NA | ALDH5A1 | aldehyde dehydrogenase 5 family, member A1 | 7915 | ENSG00000112294 |
| 38 | P02768 | NA | ALB | albumin | 213 | ENSG00000163631 |
| 39 | Q9NSE4 | NA | IARS2 | isoleucyl-tRNA synthetase 2, mitochondrial | 55699 | ENSG00000067704 |
| 40 | Q6PI48 | NA | DARS2 | aspartyl-tRNA synthetase 2, mitochondrial | 55157 | ENSG00000117593 |
| 41 | P24752 | NA | ACAT1 | acetyl-CoA acetyltransferase 1 | 38 | ENSG00000075239 |
| 42 | P13674 | NA | P4HA1 | prolyl 4-hydroxylase, alpha polypeptide I | 5033 | ENSG00000122884 |
| 43 | P19367 | NA | HK1 | hexokinase 1 | 3098 | ENSG00000156515 |
| 44 | P25705 | NA | ATP5A1 | ATP synthase, H+ transporting, mitochondrial F1 complex, alpha subunit 1, cardiac muscle | 498 | ENSG00000152234 |
| 45 | P05783 | NA | KRT18 | keratin 18 | 3875 | ENSG00000111057 |
| 46 | P06865 | NA | HEXA | hexosaminidase A (alpha polypeptide) | 3073 | ENSG00000213614 |
| 47 | P49748 | NA | ACADVL | acyl-CoA dehydrogenase, very long chain | 37 | ENSG00000072778 |
| 48 | Q16531 | NA | DDB1 | damage-specific DNA binding protein 1, 127kDa | 1642 | ENSG00000167986 |
| 49 | P31937 | NA | HIBADH | 3-hydroxyisobutyrate dehydrogenase | 11112 | ENSG00000106049 |

  
  

| **Database:cellular component      &nbspName:intracellular organelle lumen      &nbspID:GO:0070013** | | | | | | |
| --- | --- | --- | --- | --- | --- | --- |
| C=3285; O=48; E=21.27; R=2.26; rawP=4.10e-09; adjP=2.67e-08 | | | | | | |
| Index | UserID | Value | Gene Symbol | Gene Name | EntrezGene | Ensembl |
| 1 | P05787 | NA | KRT8 | keratin 8 | 3856 | ENSG00000170421 |
| 2 | Q9UL46 | NA | PSME2 | proteasome (prosome, macropain) activator subunit 2 (PA28 beta) | 5721 | ENSG00000100911 |
| 3 | P40926 | NA | MDH2 | malate dehydrogenase 2, NAD (mitochondrial) | 4191 | ENSG00000146701 |
| 4 | P35232 | NA | PHB | prohibitin | 5245 | ENSG00000167085 |
| 5 | Q99714 | NA | HSD17B10 | hydroxysteroid (17-beta) dehydrogenase 10 | 3028 | ENSG00000072506 |
| 6 | P04179 | NA | SOD2 | superoxide dismutase 2, mitochondrial | 6648 | ENSG00000112096 |
| 7 | P07741 | NA | APRT | adenine phosphoribosyltransferase | 353 | ENSG00000198931 |
| 8 | P00367 | NA | GLUD1 | glutamate dehydrogenase 1 | 2746 | ENSG00000148672 |
| 9 | Q06323 | NA | PSME1 | proteasome (prosome, macropain) activator subunit 1 (PA28 alpha) | 5720 | ENSG00000092010 |
| 10 | P48735 | NA | IDH2 | isocitrate dehydrogenase 2 (NADP+), mitochondrial | 3418 | ENSG00000182054 |
| 11 | P35270 | NA | SPR | sepiapterin reductase (7,8-dihydrobiopterin:NADP+ oxidoreductase) | 6697 | ENSG00000116096 |
| 12 | P38117 | NA | ETFB | electron-transfer-flavoprotein, beta polypeptide | 2109 | ENSG00000105379 |
| 13 | O14773 | NA | TPP1 | tripeptidyl peptidase I | 1200 | ENSG00000166340 |
| 14 | P13804 | NA | ETFA | electron-transfer-flavoprotein, alpha polypeptide | 2108 | ENSG00000140374 |
| 15 | P40121 | NA | CAPG | capping protein (actin filament), gelsolin-like | 822 | ENSG00000042493 |
| 16 | O75521 | NA | ECI2 | enoyl-CoA delta isomerase 2 | 10455 | ENSG00000198721 |
| 17 | O75367 | NA | H2AFY | H2A histone family, member Y | 9555 | ENSG00000113648 |
| 18 | P49419 | NA | ALDH7A1 | aldehyde dehydrogenase 7 family, member A1 | 501 | ENSG00000164904 |
| 19 | P67809 | NA | YBX1 | Y box binding protein 1 | 4904 | ENSG00000065978 |
| 20 | Q04837 | NA | SSBP1 | single-stranded DNA binding protein 1, mitochondrial | 6742 | ENSG00000106028 |
| 21 | P50454 | NA | SERPINH1 | serpin peptidase inhibitor, clade H (heat shock protein 47), member 1, (collagen binding protein 1) | 871 | ENSG00000149257 |
| 22 | Q14697 | NA | GANAB | glucosidase, alpha; neutral AB | 23193 | ENSG00000089597 |
| 23 | P11387 | NA | TOP1 | topoisomerase (DNA) I | 7150 | ENSG00000198900 |
| 24 | O75439 | NA | PMPCB | peptidase (mitochondrial processing) beta | 9512 | ENSG00000105819 |
| 25 | Q9NR30 | NA | DDX21 | DEAD (Asp-Glu-Ala-Asp) box helicase 21 | 9188 | ENSG00000165732 |
| 26 | P21953 | NA | BCKDHB | branched chain keto acid dehydrogenase E1, beta polypeptide | 594 | ENSG00000083123 |
| 27 | P40939 | NA | HADHA | hydroxyacyl-CoA dehydrogenase/3-ketoacyl-CoA thiolase/enoyl-CoA hydratase (trifunctional protein), alpha subunit | 3030 | ENSG00000084754 |
| 28 | P30084 | NA | ECHS1 | enoyl CoA hydratase, short chain, 1, mitochondrial | 1892 | ENSG00000127884 |
| 29 | P52815 | NA | MRPL12 | mitochondrial ribosomal protein L12 | 6182 | ENSG00000262814 |
| 30 | P21796 | NA | VDAC1 | voltage-dependent anion channel 1 | 7416 | ENSG00000213585 |
| 31 | P04080 | NA | CSTB | cystatin B (stefin B) | 1476 | ENSG00000160213 |
| 32 | P32322 | NA | PYCR1 | pyrroline-5-carboxylate reductase 1 | 5831 | ENSG00000183010 |
| 33 | P04040 | NA | CAT | catalase | 847 | ENSG00000121691 |
| 34 | Q14980 | NA | NUMA1 | nuclear mitotic apparatus protein 1 | 4926 | ENSG00000137497 |
| 35 | Q99798 | NA | ACO2 | aconitase 2, mitochondrial | 50 | ENSG00000100412 |
| 36 | P51688 | NA | SGSH | N-sulfoglucosamine sulfohydrolase | 6448 | ENSG00000181523 |
| 37 | P51649 | NA | ALDH5A1 | aldehyde dehydrogenase 5 family, member A1 | 7915 | ENSG00000112294 |
| 38 | Q9NSE4 | NA | IARS2 | isoleucyl-tRNA synthetase 2, mitochondrial | 55699 | ENSG00000067704 |
| 39 | Q6PI48 | NA | DARS2 | aspartyl-tRNA synthetase 2, mitochondrial | 55157 | ENSG00000117593 |
| 40 | P24752 | NA | ACAT1 | acetyl-CoA acetyltransferase 1 | 38 | ENSG00000075239 |
| 41 | P13674 | NA | P4HA1 | prolyl 4-hydroxylase, alpha polypeptide I | 5033 | ENSG00000122884 |
| 42 | P19367 | NA | HK1 | hexokinase 1 | 3098 | ENSG00000156515 |
| 43 | P25705 | NA | ATP5A1 | ATP synthase, H+ transporting, mitochondrial F1 complex, alpha subunit 1, cardiac muscle | 498 | ENSG00000152234 |
| 44 | P05783 | NA | KRT18 | keratin 18 | 3875 | ENSG00000111057 |
| 45 | P06865 | NA | HEXA | hexosaminidase A (alpha polypeptide) | 3073 | ENSG00000213614 |
| 46 | P49748 | NA | ACADVL | acyl-CoA dehydrogenase, very long chain | 37 | ENSG00000072778 |
| 47 | Q16531 | NA | DDB1 | damage-specific DNA binding protein 1, 127kDa | 1642 | ENSG00000167986 |
| 48 | P31937 | NA | HIBADH | 3-hydroxyisobutyrate dehydrogenase | 11112 | ENSG00000106049 |

  
  

| **Database:cellular component      &nbspName:intracellular organelle      &nbspID:GO:0043229** | | | | | | |
| --- | --- | --- | --- | --- | --- | --- |
| C=10521; O=95; E=68.11; R=1.39; rawP=5.53e-09; adjP=3.41e-08 | | | | | | |
| Index | UserID | Value | Gene Symbol | Gene Name | EntrezGene | Ensembl |
| 1 | P10253 | NA | GAA | glucosidase, alpha; acid | 2548 | ENSG00000171298 |
| 2 | P68371 | NA | TUBB4B | tubulin, beta 4B class IVb | 10383 | ENSG00000188229 |
| 3 | Q9UL46 | NA | PSME2 | proteasome (prosome, macropain) activator subunit 2 (PA28 beta) | 5721 | ENSG00000100911 |
| 4 | P40926 | NA | MDH2 | malate dehydrogenase 2, NAD (mitochondrial) | 4191 | ENSG00000146701 |
| 5 | P04179 | NA | SOD2 | superoxide dismutase 2, mitochondrial | 6648 | ENSG00000112096 |
| 6 | Q13724 | NA | MOGS | mannosyl-oligosaccharide glucosidase | 7841 | ENSG00000115275 |
| 7 | P48735 | NA | IDH2 | isocitrate dehydrogenase 2 (NADP+), mitochondrial | 3418 | ENSG00000182054 |
| 8 | P00367 | NA | GLUD1 | glutamate dehydrogenase 1 | 2746 | ENSG00000148672 |
| 9 | Q06323 | NA | PSME1 | proteasome (prosome, macropain) activator subunit 1 (PA28 alpha) | 5720 | ENSG00000092010 |
| 10 | P53701 | NA | HCCS | holocytochrome c synthase | 3052 | ENSG00000004961 |
| 11 | P35580 | NA | MYH10 | myosin, heavy chain 10, non-muscle | 4628 | ENSG00000133026 |
| 12 | P31040 | NA | SDHA | succinate dehydrogenase complex, subunit A, flavoprotein (Fp) | 6389 | ENSG00000073578 |
| 13 | O75369 | NA | FLNB | filamin B, beta | 2317 | ENSG00000136068 |
| 14 | O14773 | NA | TPP1 | tripeptidyl peptidase I | 1200 | ENSG00000166340 |
| 15 | P42765 | NA | ACAA2 | acetyl-CoA acyltransferase 2 | 10449 | ENSG00000167315 |
| 16 | P13804 | NA | ETFA | electron-transfer-flavoprotein, alpha polypeptide | 2108 | ENSG00000140374 |
| 17 | P40121 | NA | CAPG | capping protein (actin filament), gelsolin-like | 822 | ENSG00000042493 |
| 18 | Q9Y277 | NA | VDAC3 | voltage-dependent anion channel 3 | 7419 | ENSG00000078668 |
| 19 | P17174 | NA | GOT1 | glutamic-oxaloacetic transaminase 1, soluble (aspartate aminotransferase 1) | 2805 | ENSG00000120053 |
| 20 | P08727 | NA | KRT19 | keratin 19 | 3880 | ENSG00000171345 |
| 21 | Q04837 | NA | SSBP1 | single-stranded DNA binding protein 1, mitochondrial | 6742 | ENSG00000106028 |
| 22 | P50454 | NA | SERPINH1 | serpin peptidase inhibitor, clade H (heat shock protein 47), member 1, (collagen binding protein 1) | 871 | ENSG00000149257 |
| 23 | Q14697 | NA | GANAB | glucosidase, alpha; neutral AB | 23193 | ENSG00000089597 |
| 24 | P18206 | NA | VCL | vinculin | 7414 | ENSG00000035403 |
| 25 | P08195 | NA | SLC3A2 | solute carrier family 3 (activators of dibasic and neutral amino acid transport), member 2 | 6520 | ENSG00000168003 |
| 26 | P40939 | NA | HADHA | hydroxyacyl-CoA dehydrogenase/3-ketoacyl-CoA thiolase/enoyl-CoA hydratase (trifunctional protein), alpha subunit | 3030 | ENSG00000084754 |
| 27 | P09382 | NA | LGALS1 | lectin, galactoside-binding, soluble, 1 | 3956 | ENSG00000100097 |
| 28 | P52815 | NA | MRPL12 | mitochondrial ribosomal protein L12 | 6182 | ENSG00000262814 |
| 29 | Q9BXW7 | NA | CECR5 | cat eye syndrome chromosome region, candidate 5 | 27440 | ENSG00000069998 |
| 30 | P21796 | NA | VDAC1 | voltage-dependent anion channel 1 | 7416 | ENSG00000213585 |
| 31 | P04080 | NA | CSTB | cystatin B (stefin B) | 1476 | ENSG00000160213 |
| 32 | P38571 | NA | LIPA | lipase A, lysosomal acid, cholesterol esterase | 3988 | ENSG00000107798 |
| 33 | Q9UJZ1 | NA | STOML2 | stomatin (EPB72)-like 2 | 30968 | ENSG00000165283 |
| 34 | P32322 | NA | PYCR1 | pyrroline-5-carboxylate reductase 1 | 5831 | ENSG00000183010 |
| 35 | Q14980 | NA | NUMA1 | nuclear mitotic apparatus protein 1 | 4926 | ENSG00000137497 |
| 36 | Q99798 | NA | ACO2 | aconitase 2, mitochondrial | 50 | ENSG00000100412 |
| 37 | P51649 | NA | ALDH5A1 | aldehyde dehydrogenase 5 family, member A1 | 7915 | ENSG00000112294 |
| 38 | P12235 | NA | SLC25A4 | solute carrier family 25 (mitochondrial carrier; adenine nucleotide translocator), member 4 | 291 | ENSG00000151729 |
| 39 | P27816 | NA | MAP4 | microtubule-associated protein 4 | 4134 | ENSG00000047849 |
| 40 | Q6PI48 | NA | DARS2 | aspartyl-tRNA synthetase 2, mitochondrial | 55157 | ENSG00000117593 |
| 41 | Q9NSE4 | NA | IARS2 | isoleucyl-tRNA synthetase 2, mitochondrial | 55699 | ENSG00000067704 |
| 42 | P05556 | NA | ITGB1 | integrin, beta 1 (fibronectin receptor, beta polypeptide, antigen CD29 includes MDF2, MSK12) | 3688 | ENSG00000150093 |
| 43 | P13674 | NA | P4HA1 | prolyl 4-hydroxylase, alpha polypeptide I | 5033 | ENSG00000122884 |
| 44 | P40227 | NA | CCT6A | chaperonin containing TCP1, subunit 6A (zeta 1) | 908 | ENSG00000146731 |
| 45 | Q02978 | NA | SLC25A11 | solute carrier family 25 (mitochondrial carrier; oxoglutarate carrier), member 11 | 8402 | ENSG00000108528 |
| 46 | P60174 | NA | TPI1 | triosephosphate isomerase 1 | 7167 | ENSG00000111669 |
| 47 | P30042 | NA | C21orf33 | chromosome 21 open reading frame 33 | 8209 | ENSG00000160221 |
| 48 | P54819 | NA | AK2 | adenylate kinase 2 | 204 | ENSG00000004455 |
| 49 | P17931 | NA | LGALS3 | lectin, galactoside-binding, soluble, 3 | 3958 | ENSG00000131981 |
| 50 | P05787 | NA | KRT8 | keratin 8 | 3856 | ENSG00000170421 |
| 51 | Q99623 | NA | PHB2 | prohibitin 2 | 11331 | ENSG00000215021 |
| 52 | P50453 | NA | SERPINB9 | serpin peptidase inhibitor, clade B (ovalbumin), member 9 | 5272 | ENSG00000170542 |
| 53 | P35232 | NA | PHB | prohibitin | 5245 | ENSG00000167085 |
| 54 | O95831 | NA | AIFM1 | apoptosis-inducing factor, mitochondrion-associated, 1 | 9131 | ENSG00000156709 |
| 55 | Q99714 | NA | HSD17B10 | hydroxysteroid (17-beta) dehydrogenase 10 | 3028 | ENSG00000072506 |
| 56 | P61313 | NA | RPL15 | ribosomal protein L15 | 6138 | ENSG00000174748 |
| 57 | P07741 | NA | APRT | adenine phosphoribosyltransferase | 353 | ENSG00000198931 |
| 58 | O60313 | NA | OPA1 | optic atrophy 1 (autosomal dominant) | 4976 | ENSG00000198836 |
| 59 | P35270 | NA | SPR | sepiapterin reductase (7,8-dihydrobiopterin:NADP+ oxidoreductase) | 6697 | ENSG00000116096 |
| 60 | P68104 | NA | EEF1A1 | eukaryotic translation elongation factor 1 alpha 1 | 1915 | ENSG00000156508 |
| 61 | P02786 | NA | TFRC | transferrin receptor (p90, CD71) | 7037 | ENSG00000072274 |
| 62 | P38117 | NA | ETFB | electron-transfer-flavoprotein, beta polypeptide | 2109 | ENSG00000105379 |
| 63 | O75367 | NA | H2AFY | H2A histone family, member Y | 9555 | ENSG00000113648 |
| 64 | O75521 | NA | ECI2 | enoyl-CoA delta isomerase 2 | 10455 | ENSG00000198721 |
| 65 | P49419 | NA | ALDH7A1 | aldehyde dehydrogenase 7 family, member A1 | 501 | ENSG00000164904 |
| 66 | P67809 | NA | YBX1 | Y box binding protein 1 | 4904 | ENSG00000065978 |
| 67 | Q14764 | NA | MVP | major vault protein | 9961 | ENSG00000013364 |
| 68 | Q6PIU2 | NA | NCEH1 | neutral cholesterol ester hydrolase 1 | 57552 | ENSG00000144959 |
| 69 | Q13813 | NA | SPTAN1 | spectrin, alpha, non-erythrocytic 1 | 6709 | ENSG00000197694 |
| 70 | Q5SSJ5 | NA | HP1BP3 | heterochromatin protein 1, binding protein 3 | 50809 | ENSG00000127483 |
| 71 | Q9BPW8 | NA | NIPSNAP1 | nipsnap homolog 1 (C. elegans) | 8508 | ENSG00000184117 |
| 72 | P11387 | NA | TOP1 | topoisomerase (DNA) I | 7150 | ENSG00000198900 |
| 73 | O75439 | NA | PMPCB | peptidase (mitochondrial processing) beta | 9512 | ENSG00000105819 |
| 74 | Q9NR30 | NA | DDX21 | DEAD (Asp-Glu-Ala-Asp) box helicase 21 | 9188 | ENSG00000165732 |
| 75 | P00491 | NA | PNP | purine nucleoside phosphorylase | 4860 | ENSG00000198805 |
| 76 | P21953 | NA | BCKDHB | branched chain keto acid dehydrogenase E1, beta polypeptide | 594 | ENSG00000083123 |
| 77 | P30084 | NA | ECHS1 | enoyl CoA hydratase, short chain, 1, mitochondrial | 1892 | ENSG00000127884 |
| 78 | Q13011 | NA | ECH1 | enoyl CoA hydratase 1, peroxisomal | 1891 | ENSG00000104823 |
| 79 | P04083 | NA | ANXA1 | annexin A1 | 301 | ENSG00000135046 |
| 80 | P04040 | NA | CAT | catalase | 847 | ENSG00000121691 |
| 81 | O75340 | NA | PDCD6 | programmed cell death 6 | 10016 | ENSG00000249915 |
| 82 | P51688 | NA | SGSH | N-sulfoglucosamine sulfohydrolase | 6448 | ENSG00000181523 |
| 83 | P30048 | NA | PRDX3 | peroxiredoxin 3 | 10935 | ENSG00000165672 |
| 84 | P02768 | NA | ALB | albumin | 213 | ENSG00000163631 |
| 85 | P07108 | NA | DBI | diazepam binding inhibitor (GABA receptor modulator, acyl-CoA binding protein) | 1622 | ENSG00000155368 |
| 86 | P24752 | NA | ACAT1 | acetyl-CoA acetyltransferase 1 | 38 | ENSG00000075239 |
| 87 | P19367 | NA | HK1 | hexokinase 1 | 3098 | ENSG00000156515 |
| 88 | P25705 | NA | ATP5A1 | ATP synthase, H+ transporting, mitochondrial F1 complex, alpha subunit 1, cardiac muscle | 498 | ENSG00000152234 |
| 89 | Q9BQ69 | NA | MACROD1 | MACRO domain containing 1 | 28992 | ENSG00000133315 |
| 90 | P05783 | NA | KRT18 | keratin 18 | 3875 | ENSG00000111057 |
| 91 | P06865 | NA | HEXA | hexosaminidase A (alpha polypeptide) | 3073 | ENSG00000213614 |
| 92 | P49748 | NA | ACADVL | acyl-CoA dehydrogenase, very long chain | 37 | ENSG00000072778 |
| 93 | P48047 | NA | ATP5O | ATP synthase, H+ transporting, mitochondrial F1 complex, O subunit | 539 | ENSG00000241837 |
| 94 | Q16531 | NA | DDB1 | damage-specific DNA binding protein 1, 127kDa | 1642 | ENSG00000167986 |
| 95 | P31937 | NA | HIBADH | 3-hydroxyisobutyrate dehydrogenase | 11112 | ENSG00000106049 |

  
  

| **Database:cellular component      &nbspName:organelle      &nbspID:GO:0043226** | | | | | | |
| --- | --- | --- | --- | --- | --- | --- |
| C=10536; O=95; E=68.21; R=1.39; rawP=6.14e-09; adjP=3.59e-08 | | | | | | |
| Index | UserID | Value | Gene Symbol | Gene Name | EntrezGene | Ensembl |
| 1 | P10253 | NA | GAA | glucosidase, alpha; acid | 2548 | ENSG00000171298 |
| 2 | P68371 | NA | TUBB4B | tubulin, beta 4B class IVb | 10383 | ENSG00000188229 |
| 3 | Q9UL46 | NA | PSME2 | proteasome (prosome, macropain) activator subunit 2 (PA28 beta) | 5721 | ENSG00000100911 |
| 4 | P40926 | NA | MDH2 | malate dehydrogenase 2, NAD (mitochondrial) | 4191 | ENSG00000146701 |
| 5 | P04179 | NA | SOD2 | superoxide dismutase 2, mitochondrial | 6648 | ENSG00000112096 |
| 6 | Q13724 | NA | MOGS | mannosyl-oligosaccharide glucosidase | 7841 | ENSG00000115275 |
| 7 | P48735 | NA | IDH2 | isocitrate dehydrogenase 2 (NADP+), mitochondrial | 3418 | ENSG00000182054 |
| 8 | P00367 | NA | GLUD1 | glutamate dehydrogenase 1 | 2746 | ENSG00000148672 |
| 9 | Q06323 | NA | PSME1 | proteasome (prosome, macropain) activator subunit 1 (PA28 alpha) | 5720 | ENSG00000092010 |
| 10 | P53701 | NA | HCCS | holocytochrome c synthase | 3052 | ENSG00000004961 |
| 11 | P35580 | NA | MYH10 | myosin, heavy chain 10, non-muscle | 4628 | ENSG00000133026 |
| 12 | P31040 | NA | SDHA | succinate dehydrogenase complex, subunit A, flavoprotein (Fp) | 6389 | ENSG00000073578 |
| 13 | O75369 | NA | FLNB | filamin B, beta | 2317 | ENSG00000136068 |
| 14 | O14773 | NA | TPP1 | tripeptidyl peptidase I | 1200 | ENSG00000166340 |
| 15 | P42765 | NA | ACAA2 | acetyl-CoA acyltransferase 2 | 10449 | ENSG00000167315 |
| 16 | P13804 | NA | ETFA | electron-transfer-flavoprotein, alpha polypeptide | 2108 | ENSG00000140374 |
| 17 | P40121 | NA | CAPG | capping protein (actin filament), gelsolin-like | 822 | ENSG00000042493 |
| 18 | Q9Y277 | NA | VDAC3 | voltage-dependent anion channel 3 | 7419 | ENSG00000078668 |
| 19 | P17174 | NA | GOT1 | glutamic-oxaloacetic transaminase 1, soluble (aspartate aminotransferase 1) | 2805 | ENSG00000120053 |
| 20 | P08727 | NA | KRT19 | keratin 19 | 3880 | ENSG00000171345 |
| 21 | Q04837 | NA | SSBP1 | single-stranded DNA binding protein 1, mitochondrial | 6742 | ENSG00000106028 |
| 22 | P50454 | NA | SERPINH1 | serpin peptidase inhibitor, clade H (heat shock protein 47), member 1, (collagen binding protein 1) | 871 | ENSG00000149257 |
| 23 | Q14697 | NA | GANAB | glucosidase, alpha; neutral AB | 23193 | ENSG00000089597 |
| 24 | P18206 | NA | VCL | vinculin | 7414 | ENSG00000035403 |
| 25 | P08195 | NA | SLC3A2 | solute carrier family 3 (activators of dibasic and neutral amino acid transport), member 2 | 6520 | ENSG00000168003 |
| 26 | P40939 | NA | HADHA | hydroxyacyl-CoA dehydrogenase/3-ketoacyl-CoA thiolase/enoyl-CoA hydratase (trifunctional protein), alpha subunit | 3030 | ENSG00000084754 |
| 27 | P09382 | NA | LGALS1 | lectin, galactoside-binding, soluble, 1 | 3956 | ENSG00000100097 |
| 28 | P52815 | NA | MRPL12 | mitochondrial ribosomal protein L12 | 6182 | ENSG00000262814 |
| 29 | Q9BXW7 | NA | CECR5 | cat eye syndrome chromosome region, candidate 5 | 27440 | ENSG00000069998 |
| 30 | P21796 | NA | VDAC1 | voltage-dependent anion channel 1 | 7416 | ENSG00000213585 |
| 31 | P04080 | NA | CSTB | cystatin B (stefin B) | 1476 | ENSG00000160213 |
| 32 | P38571 | NA | LIPA | lipase A, lysosomal acid, cholesterol esterase | 3988 | ENSG00000107798 |
| 33 | Q9UJZ1 | NA | STOML2 | stomatin (EPB72)-like 2 | 30968 | ENSG00000165283 |
| 34 | P32322 | NA | PYCR1 | pyrroline-5-carboxylate reductase 1 | 5831 | ENSG00000183010 |
| 35 | Q14980 | NA | NUMA1 | nuclear mitotic apparatus protein 1 | 4926 | ENSG00000137497 |
| 36 | Q99798 | NA | ACO2 | aconitase 2, mitochondrial | 50 | ENSG00000100412 |
| 37 | P51649 | NA | ALDH5A1 | aldehyde dehydrogenase 5 family, member A1 | 7915 | ENSG00000112294 |
| 38 | P12235 | NA | SLC25A4 | solute carrier family 25 (mitochondrial carrier; adenine nucleotide translocator), member 4 | 291 | ENSG00000151729 |
| 39 | P27816 | NA | MAP4 | microtubule-associated protein 4 | 4134 | ENSG00000047849 |
| 40 | Q6PI48 | NA | DARS2 | aspartyl-tRNA synthetase 2, mitochondrial | 55157 | ENSG00000117593 |
| 41 | Q9NSE4 | NA | IARS2 | isoleucyl-tRNA synthetase 2, mitochondrial | 55699 | ENSG00000067704 |
| 42 | P05556 | NA | ITGB1 | integrin, beta 1 (fibronectin receptor, beta polypeptide, antigen CD29 includes MDF2, MSK12) | 3688 | ENSG00000150093 |
| 43 | P13674 | NA | P4HA1 | prolyl 4-hydroxylase, alpha polypeptide I | 5033 | ENSG00000122884 |
| 44 | P40227 | NA | CCT6A | chaperonin containing TCP1, subunit 6A (zeta 1) | 908 | ENSG00000146731 |
| 45 | Q02978 | NA | SLC25A11 | solute carrier family 25 (mitochondrial carrier; oxoglutarate carrier), member 11 | 8402 | ENSG00000108528 |
| 46 | P60174 | NA | TPI1 | triosephosphate isomerase 1 | 7167 | ENSG00000111669 |
| 47 | P30042 | NA | C21orf33 | chromosome 21 open reading frame 33 | 8209 | ENSG00000160221 |
| 48 | P54819 | NA | AK2 | adenylate kinase 2 | 204 | ENSG00000004455 |
| 49 | P17931 | NA | LGALS3 | lectin, galactoside-binding, soluble, 3 | 3958 | ENSG00000131981 |
| 50 | P05787 | NA | KRT8 | keratin 8 | 3856 | ENSG00000170421 |
| 51 | Q99623 | NA | PHB2 | prohibitin 2 | 11331 | ENSG00000215021 |
| 52 | P50453 | NA | SERPINB9 | serpin peptidase inhibitor, clade B (ovalbumin), member 9 | 5272 | ENSG00000170542 |
| 53 | P35232 | NA | PHB | prohibitin | 5245 | ENSG00000167085 |
| 54 | O95831 | NA | AIFM1 | apoptosis-inducing factor, mitochondrion-associated, 1 | 9131 | ENSG00000156709 |
| 55 | Q99714 | NA | HSD17B10 | hydroxysteroid (17-beta) dehydrogenase 10 | 3028 | ENSG00000072506 |
| 56 | P61313 | NA | RPL15 | ribosomal protein L15 | 6138 | ENSG00000174748 |
| 57 | P07741 | NA | APRT | adenine phosphoribosyltransferase | 353 | ENSG00000198931 |
| 58 | O60313 | NA | OPA1 | optic atrophy 1 (autosomal dominant) | 4976 | ENSG00000198836 |
| 59 | P35270 | NA | SPR | sepiapterin reductase (7,8-dihydrobiopterin:NADP+ oxidoreductase) | 6697 | ENSG00000116096 |
| 60 | P68104 | NA | EEF1A1 | eukaryotic translation elongation factor 1 alpha 1 | 1915 | ENSG00000156508 |
| 61 | P02786 | NA | TFRC | transferrin receptor (p90, CD71) | 7037 | ENSG00000072274 |
| 62 | P38117 | NA | ETFB | electron-transfer-flavoprotein, beta polypeptide | 2109 | ENSG00000105379 |
| 63 | O75367 | NA | H2AFY | H2A histone family, member Y | 9555 | ENSG00000113648 |
| 64 | O75521 | NA | ECI2 | enoyl-CoA delta isomerase 2 | 10455 | ENSG00000198721 |
| 65 | P49419 | NA | ALDH7A1 | aldehyde dehydrogenase 7 family, member A1 | 501 | ENSG00000164904 |
| 66 | P67809 | NA | YBX1 | Y box binding protein 1 | 4904 | ENSG00000065978 |
| 67 | Q14764 | NA | MVP | major vault protein | 9961 | ENSG00000013364 |
| 68 | Q6PIU2 | NA | NCEH1 | neutral cholesterol ester hydrolase 1 | 57552 | ENSG00000144959 |
| 69 | Q13813 | NA | SPTAN1 | spectrin, alpha, non-erythrocytic 1 | 6709 | ENSG00000197694 |
| 70 | Q5SSJ5 | NA | HP1BP3 | heterochromatin protein 1, binding protein 3 | 50809 | ENSG00000127483 |
| 71 | Q9BPW8 | NA | NIPSNAP1 | nipsnap homolog 1 (C. elegans) | 8508 | ENSG00000184117 |
| 72 | P11387 | NA | TOP1 | topoisomerase (DNA) I | 7150 | ENSG00000198900 |
| 73 | O75439 | NA | PMPCB | peptidase (mitochondrial processing) beta | 9512 | ENSG00000105819 |
| 74 | Q9NR30 | NA | DDX21 | DEAD (Asp-Glu-Ala-Asp) box helicase 21 | 9188 | ENSG00000165732 |
| 75 | P00491 | NA | PNP | purine nucleoside phosphorylase | 4860 | ENSG00000198805 |
| 76 | P21953 | NA | BCKDHB | branched chain keto acid dehydrogenase E1, beta polypeptide | 594 | ENSG00000083123 |
| 77 | P30084 | NA | ECHS1 | enoyl CoA hydratase, short chain, 1, mitochondrial | 1892 | ENSG00000127884 |
| 78 | Q13011 | NA | ECH1 | enoyl CoA hydratase 1, peroxisomal | 1891 | ENSG00000104823 |
| 79 | P04083 | NA | ANXA1 | annexin A1 | 301 | ENSG00000135046 |
| 80 | P04040 | NA | CAT | catalase | 847 | ENSG00000121691 |
| 81 | O75340 | NA | PDCD6 | programmed cell death 6 | 10016 | ENSG00000249915 |
| 82 | P51688 | NA | SGSH | N-sulfoglucosamine sulfohydrolase | 6448 | ENSG00000181523 |
| 83 | P30048 | NA | PRDX3 | peroxiredoxin 3 | 10935 | ENSG00000165672 |
| 84 | P02768 | NA | ALB | albumin | 213 | ENSG00000163631 |
| 85 | P07108 | NA | DBI | diazepam binding inhibitor (GABA receptor modulator, acyl-CoA binding protein) | 1622 | ENSG00000155368 |
| 86 | P24752 | NA | ACAT1 | acetyl-CoA acetyltransferase 1 | 38 | ENSG00000075239 |
| 87 | P19367 | NA | HK1 | hexokinase 1 | 3098 | ENSG00000156515 |
| 88 | P25705 | NA | ATP5A1 | ATP synthase, H+ transporting, mitochondrial F1 complex, alpha subunit 1, cardiac muscle | 498 | ENSG00000152234 |
| 89 | Q9BQ69 | NA | MACROD1 | MACRO domain containing 1 | 28992 | ENSG00000133315 |
| 90 | P05783 | NA | KRT18 | keratin 18 | 3875 | ENSG00000111057 |
| 91 | P06865 | NA | HEXA | hexosaminidase A (alpha polypeptide) | 3073 | ENSG00000213614 |
| 92 | P49748 | NA | ACADVL | acyl-CoA dehydrogenase, very long chain | 37 | ENSG00000072778 |
| 93 | P48047 | NA | ATP5O | ATP synthase, H+ transporting, mitochondrial F1 complex, O subunit | 539 | ENSG00000241837 |
| 94 | Q16531 | NA | DDB1 | damage-specific DNA binding protein 1, 127kDa | 1642 | ENSG00000167986 |
| 95 | P31937 | NA | HIBADH | 3-hydroxyisobutyrate dehydrogenase | 11112 | ENSG00000106049 |

  
  

| **Database:cellular component      &nbspName:cell part      &nbspID:GO:0044464** | | | | | | |
| --- | --- | --- | --- | --- | --- | --- |
| C=14405; O=108; E=93.26; R=1.16; rawP=1.24e-07; adjP=6.65e-07 | | | | | | |
| Index | UserID | Value | Gene Symbol | Gene Name | EntrezGene | Ensembl |
| 1 | P10253 | NA | GAA | glucosidase, alpha; acid | 2548 | ENSG00000171298 |
| 2 | P68371 | NA | TUBB4B | tubulin, beta 4B class IVb | 10383 | ENSG00000188229 |
| 3 | Q9UL46 | NA | PSME2 | proteasome (prosome, macropain) activator subunit 2 (PA28 beta) | 5721 | ENSG00000100911 |
| 4 | P40926 | NA | MDH2 | malate dehydrogenase 2, NAD (mitochondrial) | 4191 | ENSG00000146701 |
| 5 | P04179 | NA | SOD2 | superoxide dismutase 2, mitochondrial | 6648 | ENSG00000112096 |
| 6 | Q13724 | NA | MOGS | mannosyl-oligosaccharide glucosidase | 7841 | ENSG00000115275 |
| 7 | P48735 | NA | IDH2 | isocitrate dehydrogenase 2 (NADP+), mitochondrial | 3418 | ENSG00000182054 |
| 8 | P00367 | NA | GLUD1 | glutamate dehydrogenase 1 | 2746 | ENSG00000148672 |
| 9 | Q06323 | NA | PSME1 | proteasome (prosome, macropain) activator subunit 1 (PA28 alpha) | 5720 | ENSG00000092010 |
| 10 | P53701 | NA | HCCS | holocytochrome c synthase | 3052 | ENSG00000004961 |
| 11 | P35580 | NA | MYH10 | myosin, heavy chain 10, non-muscle | 4628 | ENSG00000133026 |
| 12 | P31040 | NA | SDHA | succinate dehydrogenase complex, subunit A, flavoprotein (Fp) | 6389 | ENSG00000073578 |
| 13 | O75369 | NA | FLNB | filamin B, beta | 2317 | ENSG00000136068 |
| 14 | P24534 | NA | EEF1B2 | eukaryotic translation elongation factor 1 beta 2 | 1933 | ENSG00000114942 |
| 15 | O14773 | NA | TPP1 | tripeptidyl peptidase I | 1200 | ENSG00000166340 |
| 16 | P47895 | NA | ALDH1A3 | aldehyde dehydrogenase 1 family, member A3 | 220 | ENSG00000184254 |
| 17 | P42765 | NA | ACAA2 | acetyl-CoA acyltransferase 2 | 10449 | ENSG00000167315 |
| 18 | P40121 | NA | CAPG | capping protein (actin filament), gelsolin-like | 822 | ENSG00000042493 |
| 19 | P13804 | NA | ETFA | electron-transfer-flavoprotein, alpha polypeptide | 2108 | ENSG00000140374 |
| 20 | Q96QD8 | NA | SLC38A2 | solute carrier family 38, member 2 | 54407 | ENSG00000134294 |
| 21 | Q9Y277 | NA | VDAC3 | voltage-dependent anion channel 3 | 7419 | ENSG00000078668 |
| 22 | P17174 | NA | GOT1 | glutamic-oxaloacetic transaminase 1, soluble (aspartate aminotransferase 1) | 2805 | ENSG00000120053 |
| 23 | P08727 | NA | KRT19 | keratin 19 | 3880 | ENSG00000171345 |
| 24 | Q04837 | NA | SSBP1 | single-stranded DNA binding protein 1, mitochondrial | 6742 | ENSG00000106028 |
| 25 | P50454 | NA | SERPINH1 | serpin peptidase inhibitor, clade H (heat shock protein 47), member 1, (collagen binding protein 1) | 871 | ENSG00000149257 |
| 26 | Q14697 | NA | GANAB | glucosidase, alpha; neutral AB | 23193 | ENSG00000089597 |
| 27 | P18206 | NA | VCL | vinculin | 7414 | ENSG00000035403 |
| 28 | P08195 | NA | SLC3A2 | solute carrier family 3 (activators of dibasic and neutral amino acid transport), member 2 | 6520 | ENSG00000168003 |
| 29 | P40939 | NA | HADHA | hydroxyacyl-CoA dehydrogenase/3-ketoacyl-CoA thiolase/enoyl-CoA hydratase (trifunctional protein), alpha subunit | 3030 | ENSG00000084754 |
| 30 | P09382 | NA | LGALS1 | lectin, galactoside-binding, soluble, 1 | 3956 | ENSG00000100097 |
| 31 | P52815 | NA | MRPL12 | mitochondrial ribosomal protein L12 | 6182 | ENSG00000262814 |
| 32 | Q9BXW7 | NA | CECR5 | cat eye syndrome chromosome region, candidate 5 | 27440 | ENSG00000069998 |
| 33 | P21796 | NA | VDAC1 | voltage-dependent anion channel 1 | 7416 | ENSG00000213585 |
| 34 | P38571 | NA | LIPA | lipase A, lysosomal acid, cholesterol esterase | 3988 | ENSG00000107798 |
| 35 | P04080 | NA | CSTB | cystatin B (stefin B) | 1476 | ENSG00000160213 |
| 36 | Q9UJZ1 | NA | STOML2 | stomatin (EPB72)-like 2 | 30968 | ENSG00000165283 |
| 37 | P32322 | NA | PYCR1 | pyrroline-5-carboxylate reductase 1 | 5831 | ENSG00000183010 |
| 38 | Q14980 | NA | NUMA1 | nuclear mitotic apparatus protein 1 | 4926 | ENSG00000137497 |
| 39 | Q99798 | NA | ACO2 | aconitase 2, mitochondrial | 50 | ENSG00000100412 |
| 40 | P51649 | NA | ALDH5A1 | aldehyde dehydrogenase 5 family, member A1 | 7915 | ENSG00000112294 |
| 41 | P12235 | NA | SLC25A4 | solute carrier family 25 (mitochondrial carrier; adenine nucleotide translocator), member 4 | 291 | ENSG00000151729 |
| 42 | P27816 | NA | MAP4 | microtubule-associated protein 4 | 4134 | ENSG00000047849 |
| 43 | Q6PI48 | NA | DARS2 | aspartyl-tRNA synthetase 2, mitochondrial | 55157 | ENSG00000117593 |
| 44 | Q9NSE4 | NA | IARS2 | isoleucyl-tRNA synthetase 2, mitochondrial | 55699 | ENSG00000067704 |
| 45 | P05556 | NA | ITGB1 | integrin, beta 1 (fibronectin receptor, beta polypeptide, antigen CD29 includes MDF2, MSK12) | 3688 | ENSG00000150093 |
| 46 | P13674 | NA | P4HA1 | prolyl 4-hydroxylase, alpha polypeptide I | 5033 | ENSG00000122884 |
| 47 | P40227 | NA | CCT6A | chaperonin containing TCP1, subunit 6A (zeta 1) | 908 | ENSG00000146731 |
| 48 | Q02978 | NA | SLC25A11 | solute carrier family 25 (mitochondrial carrier; oxoglutarate carrier), member 11 | 8402 | ENSG00000108528 |
| 49 | P60174 | NA | TPI1 | triosephosphate isomerase 1 | 7167 | ENSG00000111669 |
| 50 | P30042 | NA | C21orf33 | chromosome 21 open reading frame 33 | 8209 | ENSG00000160221 |
| 51 | P49589 | NA | CARS | cysteinyl-tRNA synthetase | 833 | ENSG00000110619 |
| 52 | P54819 | NA | AK2 | adenylate kinase 2 | 204 | ENSG00000004455 |
| 53 | P17931 | NA | LGALS3 | lectin, galactoside-binding, soluble, 3 | 3958 | ENSG00000131981 |
| 54 | P05787 | NA | KRT8 | keratin 8 | 3856 | ENSG00000170421 |
| 55 | Q99623 | NA | PHB2 | prohibitin 2 | 11331 | ENSG00000215021 |
| 56 | P50453 | NA | SERPINB9 | serpin peptidase inhibitor, clade B (ovalbumin), member 9 | 5272 | ENSG00000170542 |
| 57 | P35232 | NA | PHB | prohibitin | 5245 | ENSG00000167085 |
| 58 | O95831 | NA | AIFM1 | apoptosis-inducing factor, mitochondrion-associated, 1 | 9131 | ENSG00000156709 |
| 59 | Q99714 | NA | HSD17B10 | hydroxysteroid (17-beta) dehydrogenase 10 | 3028 | ENSG00000072506 |
| 60 | P61313 | NA | RPL15 | ribosomal protein L15 | 6138 | ENSG00000174748 |
| 61 | P07741 | NA | APRT | adenine phosphoribosyltransferase | 353 | ENSG00000198931 |
| 62 | O60313 | NA | OPA1 | optic atrophy 1 (autosomal dominant) | 4976 | ENSG00000198836 |
| 63 | P35270 | NA | SPR | sepiapterin reductase (7,8-dihydrobiopterin:NADP+ oxidoreductase) | 6697 | ENSG00000116096 |
| 64 | P68104 | NA | EEF1A1 | eukaryotic translation elongation factor 1 alpha 1 | 1915 | ENSG00000156508 |
| 65 | P38117 | NA | ETFB | electron-transfer-flavoprotein, beta polypeptide | 2109 | ENSG00000105379 |
| 66 | P02786 | NA | TFRC | transferrin receptor (p90, CD71) | 7037 | ENSG00000072274 |
| 67 | P43490 | NA | NAMPT | nicotinamide phosphoribosyltransferase | 10135 | ENSG00000105835 |
| 68 | P22102 | NA | GART | phosphoribosylglycinamide formyltransferase, phosphoribosylglycinamide synthetase, phosphoribosylaminoimidazole synthetase | 2618 | ENSG00000159131 |
| 69 | P13639 | NA | EEF2 | eukaryotic translation elongation factor 2 | 1938 | ENSG00000167658 |
| 70 | O75521 | NA | ECI2 | enoyl-CoA delta isomerase 2 | 10455 | ENSG00000198721 |
| 71 | O75367 | NA | H2AFY | H2A histone family, member Y | 9555 | ENSG00000113648 |
| 72 | P49419 | NA | ALDH7A1 | aldehyde dehydrogenase 7 family, member A1 | 501 | ENSG00000164904 |
| 73 | P67809 | NA | YBX1 | Y box binding protein 1 | 4904 | ENSG00000065978 |
| 74 | Q14764 | NA | MVP | major vault protein | 9961 | ENSG00000013364 |
| 75 | Q6PIU2 | NA | NCEH1 | neutral cholesterol ester hydrolase 1 | 57552 | ENSG00000144959 |
| 76 | P29692 | NA | EEF1D | eukaryotic translation elongation factor 1 delta (guanine nucleotide exchange protein) | 1936 | ENSG00000104529 |
| 77 | Q13813 | NA | SPTAN1 | spectrin, alpha, non-erythrocytic 1 | 6709 | ENSG00000197694 |
| 78 | Q5SSJ5 | NA | HP1BP3 | heterochromatin protein 1, binding protein 3 | 50809 | ENSG00000127483 |
| 79 | Q9BPW8 | NA | NIPSNAP1 | nipsnap homolog 1 (C. elegans) | 8508 | ENSG00000184117 |
| 80 | P11387 | NA | TOP1 | topoisomerase (DNA) I | 7150 | ENSG00000198900 |
| 81 | P02792 | NA | FTL | ferritin, light polypeptide | 2512 | ENSG00000087086 |
| 82 | O75439 | NA | PMPCB | peptidase (mitochondrial processing) beta | 9512 | ENSG00000105819 |
| 83 | Q9NR30 | NA | DDX21 | DEAD (Asp-Glu-Ala-Asp) box helicase 21 | 9188 | ENSG00000165732 |
| 84 | P00491 | NA | PNP | purine nucleoside phosphorylase | 4860 | ENSG00000198805 |
| 85 | P21953 | NA | BCKDHB | branched chain keto acid dehydrogenase E1, beta polypeptide | 594 | ENSG00000083123 |
| 86 | P30084 | NA | ECHS1 | enoyl CoA hydratase, short chain, 1, mitochondrial | 1892 | ENSG00000127884 |
| 87 | P60842 | NA | EIF4A1 | eukaryotic translation initiation factor 4A1 | 1973 | ENSG00000161960 |
| 88 | Q13011 | NA | ECH1 | enoyl CoA hydratase 1, peroxisomal | 1891 | ENSG00000104823 |
| 89 | P04083 | NA | ANXA1 | annexin A1 | 301 | ENSG00000135046 |
| 90 | P04040 | NA | CAT | catalase | 847 | ENSG00000121691 |
| 91 | O75340 | NA | PDCD6 | programmed cell death 6 | 10016 | ENSG00000249915 |
| 92 | P51688 | NA | SGSH | N-sulfoglucosamine sulfohydrolase | 6448 | ENSG00000181523 |
| 93 | P00558 | NA | PGK1 | phosphoglycerate kinase 1 | 5230 | ENSG00000102144 |
| 94 | P30048 | NA | PRDX3 | peroxiredoxin 3 | 10935 | ENSG00000165672 |
| 95 | P41091 | NA | EIF2S3 | eukaryotic translation initiation factor 2, subunit 3 gamma, 52kDa | 1968 | ENSG00000130741 |
| 96 | P02768 | NA | ALB | albumin | 213 | ENSG00000163631 |
| 97 | P07108 | NA | DBI | diazepam binding inhibitor (GABA receptor modulator, acyl-CoA binding protein) | 1622 | ENSG00000155368 |
| 98 | P24752 | NA | ACAT1 | acetyl-CoA acetyltransferase 1 | 38 | ENSG00000075239 |
| 99 | P19367 | NA | HK1 | hexokinase 1 | 3098 | ENSG00000156515 |
| 100 | P25705 | NA | ATP5A1 | ATP synthase, H+ transporting, mitochondrial F1 complex, alpha subunit 1, cardiac muscle | 498 | ENSG00000152234 |
| 101 | Q9BQ69 | NA | MACROD1 | MACRO domain containing 1 | 28992 | ENSG00000133315 |
| 102 | P05783 | NA | KRT18 | keratin 18 | 3875 | ENSG00000111057 |
| 103 | P06865 | NA | HEXA | hexosaminidase A (alpha polypeptide) | 3073 | ENSG00000213614 |
| 104 | P49748 | NA | ACADVL | acyl-CoA dehydrogenase, very long chain | 37 | ENSG00000072778 |
| 105 | P00352 | NA | ALDH1A1 | aldehyde dehydrogenase 1 family, member A1 | 216 | ENSG00000165092 |
| 106 | P48047 | NA | ATP5O | ATP synthase, H+ transporting, mitochondrial F1 complex, O subunit | 539 | ENSG00000241837 |
| 107 | Q16531 | NA | DDB1 | damage-specific DNA binding protein 1, 127kDa | 1642 | ENSG00000167986 |
| 108 | P31937 | NA | HIBADH | 3-hydroxyisobutyrate dehydrogenase | 11112 | ENSG00000106049 |

  
  

| **Database:cellular component      &nbspName:cell      &nbspID:GO:0005623** | | | | | | |
| --- | --- | --- | --- | --- | --- | --- |
| C=14406; O=108; E=93.27; R=1.16; rawP=1.25e-07; adjP=6.65e-07 | | | | | | |
| Index | UserID | Value | Gene Symbol | Gene Name | EntrezGene | Ensembl |
| 1 | P10253 | NA | GAA | glucosidase, alpha; acid | 2548 | ENSG00000171298 |
| 2 | P68371 | NA | TUBB4B | tubulin, beta 4B class IVb | 10383 | ENSG00000188229 |
| 3 | Q9UL46 | NA | PSME2 | proteasome (prosome, macropain) activator subunit 2 (PA28 beta) | 5721 | ENSG00000100911 |
| 4 | P40926 | NA | MDH2 | malate dehydrogenase 2, NAD (mitochondrial) | 4191 | ENSG00000146701 |
| 5 | P04179 | NA | SOD2 | superoxide dismutase 2, mitochondrial | 6648 | ENSG00000112096 |
| 6 | Q13724 | NA | MOGS | mannosyl-oligosaccharide glucosidase | 7841 | ENSG00000115275 |
| 7 | P48735 | NA | IDH2 | isocitrate dehydrogenase 2 (NADP+), mitochondrial | 3418 | ENSG00000182054 |
| 8 | P00367 | NA | GLUD1 | glutamate dehydrogenase 1 | 2746 | ENSG00000148672 |
| 9 | Q06323 | NA | PSME1 | proteasome (prosome, macropain) activator subunit 1 (PA28 alpha) | 5720 | ENSG00000092010 |
| 10 | P53701 | NA | HCCS | holocytochrome c synthase | 3052 | ENSG00000004961 |
| 11 | P35580 | NA | MYH10 | myosin, heavy chain 10, non-muscle | 4628 | ENSG00000133026 |
| 12 | P31040 | NA | SDHA | succinate dehydrogenase complex, subunit A, flavoprotein (Fp) | 6389 | ENSG00000073578 |
| 13 | O75369 | NA | FLNB | filamin B, beta | 2317 | ENSG00000136068 |
| 14 | P24534 | NA | EEF1B2 | eukaryotic translation elongation factor 1 beta 2 | 1933 | ENSG00000114942 |
| 15 | O14773 | NA | TPP1 | tripeptidyl peptidase I | 1200 | ENSG00000166340 |
| 16 | P47895 | NA | ALDH1A3 | aldehyde dehydrogenase 1 family, member A3 | 220 | ENSG00000184254 |
| 17 | P42765 | NA | ACAA2 | acetyl-CoA acyltransferase 2 | 10449 | ENSG00000167315 |
| 18 | P40121 | NA | CAPG | capping protein (actin filament), gelsolin-like | 822 | ENSG00000042493 |
| 19 | P13804 | NA | ETFA | electron-transfer-flavoprotein, alpha polypeptide | 2108 | ENSG00000140374 |
| 20 | Q96QD8 | NA | SLC38A2 | solute carrier family 38, member 2 | 54407 | ENSG00000134294 |
| 21 | Q9Y277 | NA | VDAC3 | voltage-dependent anion channel 3 | 7419 | ENSG00000078668 |
| 22 | P17174 | NA | GOT1 | glutamic-oxaloacetic transaminase 1, soluble (aspartate aminotransferase 1) | 2805 | ENSG00000120053 |
| 23 | P08727 | NA | KRT19 | keratin 19 | 3880 | ENSG00000171345 |
| 24 | Q04837 | NA | SSBP1 | single-stranded DNA binding protein 1, mitochondrial | 6742 | ENSG00000106028 |
| 25 | P50454 | NA | SERPINH1 | serpin peptidase inhibitor, clade H (heat shock protein 47), member 1, (collagen binding protein 1) | 871 | ENSG00000149257 |
| 26 | Q14697 | NA | GANAB | glucosidase, alpha; neutral AB | 23193 | ENSG00000089597 |
| 27 | P18206 | NA | VCL | vinculin | 7414 | ENSG00000035403 |
| 28 | P08195 | NA | SLC3A2 | solute carrier family 3 (activators of dibasic and neutral amino acid transport), member 2 | 6520 | ENSG00000168003 |
| 29 | P40939 | NA | HADHA | hydroxyacyl-CoA dehydrogenase/3-ketoacyl-CoA thiolase/enoyl-CoA hydratase (trifunctional protein), alpha subunit | 3030 | ENSG00000084754 |
| 30 | P09382 | NA | LGALS1 | lectin, galactoside-binding, soluble, 1 | 3956 | ENSG00000100097 |
| 31 | P52815 | NA | MRPL12 | mitochondrial ribosomal protein L12 | 6182 | ENSG00000262814 |
| 32 | Q9BXW7 | NA | CECR5 | cat eye syndrome chromosome region, candidate 5 | 27440 | ENSG00000069998 |
| 33 | P21796 | NA | VDAC1 | voltage-dependent anion channel 1 | 7416 | ENSG00000213585 |
| 34 | P38571 | NA | LIPA | lipase A, lysosomal acid, cholesterol esterase | 3988 | ENSG00000107798 |
| 35 | P04080 | NA | CSTB | cystatin B (stefin B) | 1476 | ENSG00000160213 |
| 36 | Q9UJZ1 | NA | STOML2 | stomatin (EPB72)-like 2 | 30968 | ENSG00000165283 |
| 37 | P32322 | NA | PYCR1 | pyrroline-5-carboxylate reductase 1 | 5831 | ENSG00000183010 |
| 38 | Q14980 | NA | NUMA1 | nuclear mitotic apparatus protein 1 | 4926 | ENSG00000137497 |
| 39 | Q99798 | NA | ACO2 | aconitase 2, mitochondrial | 50 | ENSG00000100412 |
| 40 | P51649 | NA | ALDH5A1 | aldehyde dehydrogenase 5 family, member A1 | 7915 | ENSG00000112294 |
| 41 | P12235 | NA | SLC25A4 | solute carrier family 25 (mitochondrial carrier; adenine nucleotide translocator), member 4 | 291 | ENSG00000151729 |
| 42 | P27816 | NA | MAP4 | microtubule-associated protein 4 | 4134 | ENSG00000047849 |
| 43 | Q6PI48 | NA | DARS2 | aspartyl-tRNA synthetase 2, mitochondrial | 55157 | ENSG00000117593 |
| 44 | Q9NSE4 | NA | IARS2 | isoleucyl-tRNA synthetase 2, mitochondrial | 55699 | ENSG00000067704 |
| 45 | P05556 | NA | ITGB1 | integrin, beta 1 (fibronectin receptor, beta polypeptide, antigen CD29 includes MDF2, MSK12) | 3688 | ENSG00000150093 |
| 46 | P13674 | NA | P4HA1 | prolyl 4-hydroxylase, alpha polypeptide I | 5033 | ENSG00000122884 |
| 47 | P40227 | NA | CCT6A | chaperonin containing TCP1, subunit 6A (zeta 1) | 908 | ENSG00000146731 |
| 48 | Q02978 | NA | SLC25A11 | solute carrier family 25 (mitochondrial carrier; oxoglutarate carrier), member 11 | 8402 | ENSG00000108528 |
| 49 | P60174 | NA | TPI1 | triosephosphate isomerase 1 | 7167 | ENSG00000111669 |
| 50 | P30042 | NA | C21orf33 | chromosome 21 open reading frame 33 | 8209 | ENSG00000160221 |
| 51 | P49589 | NA | CARS | cysteinyl-tRNA synthetase | 833 | ENSG00000110619 |
| 52 | P54819 | NA | AK2 | adenylate kinase 2 | 204 | ENSG00000004455 |
| 53 | P17931 | NA | LGALS3 | lectin, galactoside-binding, soluble, 3 | 3958 | ENSG00000131981 |
| 54 | P05787 | NA | KRT8 | keratin 8 | 3856 | ENSG00000170421 |
| 55 | Q99623 | NA | PHB2 | prohibitin 2 | 11331 | ENSG00000215021 |
| 56 | P50453 | NA | SERPINB9 | serpin peptidase inhibitor, clade B (ovalbumin), member 9 | 5272 | ENSG00000170542 |
| 57 | P35232 | NA | PHB | prohibitin | 5245 | ENSG00000167085 |
| 58 | O95831 | NA | AIFM1 | apoptosis-inducing factor, mitochondrion-associated, 1 | 9131 | ENSG00000156709 |
| 59 | Q99714 | NA | HSD17B10 | hydroxysteroid (17-beta) dehydrogenase 10 | 3028 | ENSG00000072506 |
| 60 | P61313 | NA | RPL15 | ribosomal protein L15 | 6138 | ENSG00000174748 |
| 61 | P07741 | NA | APRT | adenine phosphoribosyltransferase | 353 | ENSG00000198931 |
| 62 | O60313 | NA | OPA1 | optic atrophy 1 (autosomal dominant) | 4976 | ENSG00000198836 |
| 63 | P35270 | NA | SPR | sepiapterin reductase (7,8-dihydrobiopterin:NADP+ oxidoreductase) | 6697 | ENSG00000116096 |
| 64 | P68104 | NA | EEF1A1 | eukaryotic translation elongation factor 1 alpha 1 | 1915 | ENSG00000156508 |
| 65 | P38117 | NA | ETFB | electron-transfer-flavoprotein, beta polypeptide | 2109 | ENSG00000105379 |
| 66 | P02786 | NA | TFRC | transferrin receptor (p90, CD71) | 7037 | ENSG00000072274 |
| 67 | P43490 | NA | NAMPT | nicotinamide phosphoribosyltransferase | 10135 | ENSG00000105835 |
| 68 | P22102 | NA | GART | phosphoribosylglycinamide formyltransferase, phosphoribosylglycinamide synthetase, phosphoribosylaminoimidazole synthetase | 2618 | ENSG00000159131 |
| 69 | P13639 | NA | EEF2 | eukaryotic translation elongation factor 2 | 1938 | ENSG00000167658 |
| 70 | O75521 | NA | ECI2 | enoyl-CoA delta isomerase 2 | 10455 | ENSG00000198721 |
| 71 | O75367 | NA | H2AFY | H2A histone family, member Y | 9555 | ENSG00000113648 |
| 72 | P49419 | NA | ALDH7A1 | aldehyde dehydrogenase 7 family, member A1 | 501 | ENSG00000164904 |
| 73 | P67809 | NA | YBX1 | Y box binding protein 1 | 4904 | ENSG00000065978 |
| 74 | Q14764 | NA | MVP | major vault protein | 9961 | ENSG00000013364 |
| 75 | Q6PIU2 | NA | NCEH1 | neutral cholesterol ester hydrolase 1 | 57552 | ENSG00000144959 |
| 76 | P29692 | NA | EEF1D | eukaryotic translation elongation factor 1 delta (guanine nucleotide exchange protein) | 1936 | ENSG00000104529 |
| 77 | Q13813 | NA | SPTAN1 | spectrin, alpha, non-erythrocytic 1 | 6709 | ENSG00000197694 |
| 78 | Q5SSJ5 | NA | HP1BP3 | heterochromatin protein 1, binding protein 3 | 50809 | ENSG00000127483 |
| 79 | Q9BPW8 | NA | NIPSNAP1 | nipsnap homolog 1 (C. elegans) | 8508 | ENSG00000184117 |
| 80 | P11387 | NA | TOP1 | topoisomerase (DNA) I | 7150 | ENSG00000198900 |
| 81 | P02792 | NA | FTL | ferritin, light polypeptide | 2512 | ENSG00000087086 |
| 82 | O75439 | NA | PMPCB | peptidase (mitochondrial processing) beta | 9512 | ENSG00000105819 |
| 83 | Q9NR30 | NA | DDX21 | DEAD (Asp-Glu-Ala-Asp) box helicase 21 | 9188 | ENSG00000165732 |
| 84 | P00491 | NA | PNP | purine nucleoside phosphorylase | 4860 | ENSG00000198805 |
| 85 | P21953 | NA | BCKDHB | branched chain keto acid dehydrogenase E1, beta polypeptide | 594 | ENSG00000083123 |
| 86 | P30084 | NA | ECHS1 | enoyl CoA hydratase, short chain, 1, mitochondrial | 1892 | ENSG00000127884 |
| 87 | P60842 | NA | EIF4A1 | eukaryotic translation initiation factor 4A1 | 1973 | ENSG00000161960 |
| 88 | Q13011 | NA | ECH1 | enoyl CoA hydratase 1, peroxisomal | 1891 | ENSG00000104823 |
| 89 | P04083 | NA | ANXA1 | annexin A1 | 301 | ENSG00000135046 |
| 90 | P04040 | NA | CAT | catalase | 847 | ENSG00000121691 |
| 91 | O75340 | NA | PDCD6 | programmed cell death 6 | 10016 | ENSG00000249915 |
| 92 | P51688 | NA | SGSH | N-sulfoglucosamine sulfohydrolase | 6448 | ENSG00000181523 |
| 93 | P00558 | NA | PGK1 | phosphoglycerate kinase 1 | 5230 | ENSG00000102144 |
| 94 | P30048 | NA | PRDX3 | peroxiredoxin 3 | 10935 | ENSG00000165672 |
| 95 | P41091 | NA | EIF2S3 | eukaryotic translation initiation factor 2, subunit 3 gamma, 52kDa | 1968 | ENSG00000130741 |
| 96 | P02768 | NA | ALB | albumin | 213 | ENSG00000163631 |
| 97 | P07108 | NA | DBI | diazepam binding inhibitor (GABA receptor modulator, acyl-CoA binding protein) | 1622 | ENSG00000155368 |
| 98 | P24752 | NA | ACAT1 | acetyl-CoA acetyltransferase 1 | 38 | ENSG00000075239 |
| 99 | P19367 | NA | HK1 | hexokinase 1 | 3098 | ENSG00000156515 |
| 100 | P25705 | NA | ATP5A1 | ATP synthase, H+ transporting, mitochondrial F1 complex, alpha subunit 1, cardiac muscle | 498 | ENSG00000152234 |
| 101 | Q9BQ69 | NA | MACROD1 | MACRO domain containing 1 | 28992 | ENSG00000133315 |
| 102 | P05783 | NA | KRT18 | keratin 18 | 3875 | ENSG00000111057 |
| 103 | P06865 | NA | HEXA | hexosaminidase A (alpha polypeptide) | 3073 | ENSG00000213614 |
| 104 | P49748 | NA | ACADVL | acyl-CoA dehydrogenase, very long chain | 37 | ENSG00000072778 |
| 105 | P00352 | NA | ALDH1A1 | aldehyde dehydrogenase 1 family, member A1 | 216 | ENSG00000165092 |
| 106 | P48047 | NA | ATP5O | ATP synthase, H+ transporting, mitochondrial F1 complex, O subunit | 539 | ENSG00000241837 |
| 107 | Q16531 | NA | DDB1 | damage-specific DNA binding protein 1, 127kDa | 1642 | ENSG00000167986 |
| 108 | P31937 | NA | HIBADH | 3-hydroxyisobutyrate dehydrogenase | 11112 | ENSG00000106049 |

  
  

| **Database:cellular component      &nbspName:intracellular membrane-bounded organelle      &nbspID:GO:0043231** | | | | | | |
| --- | --- | --- | --- | --- | --- | --- |
| C=9484; O=87; E=61.40; R=1.42; rawP=1.63e-07; adjP=8.29e-07 | | | | | | |
| Index | UserID | Value | Gene Symbol | Gene Name | EntrezGene | Ensembl |
| 1 | P10253 | NA | GAA | glucosidase, alpha; acid | 2548 | ENSG00000171298 |
| 2 | P68371 | NA | TUBB4B | tubulin, beta 4B class IVb | 10383 | ENSG00000188229 |
| 3 | Q9UL46 | NA | PSME2 | proteasome (prosome, macropain) activator subunit 2 (PA28 beta) | 5721 | ENSG00000100911 |
| 4 | P40926 | NA | MDH2 | malate dehydrogenase 2, NAD (mitochondrial) | 4191 | ENSG00000146701 |
| 5 | P04179 | NA | SOD2 | superoxide dismutase 2, mitochondrial | 6648 | ENSG00000112096 |
| 6 | Q13724 | NA | MOGS | mannosyl-oligosaccharide glucosidase | 7841 | ENSG00000115275 |
| 7 | P48735 | NA | IDH2 | isocitrate dehydrogenase 2 (NADP+), mitochondrial | 3418 | ENSG00000182054 |
| 8 | P00367 | NA | GLUD1 | glutamate dehydrogenase 1 | 2746 | ENSG00000148672 |
| 9 | Q06323 | NA | PSME1 | proteasome (prosome, macropain) activator subunit 1 (PA28 alpha) | 5720 | ENSG00000092010 |
| 10 | P53701 | NA | HCCS | holocytochrome c synthase | 3052 | ENSG00000004961 |
| 11 | P31040 | NA | SDHA | succinate dehydrogenase complex, subunit A, flavoprotein (Fp) | 6389 | ENSG00000073578 |
| 12 | O14773 | NA | TPP1 | tripeptidyl peptidase I | 1200 | ENSG00000166340 |
| 13 | P13804 | NA | ETFA | electron-transfer-flavoprotein, alpha polypeptide | 2108 | ENSG00000140374 |
| 14 | P40121 | NA | CAPG | capping protein (actin filament), gelsolin-like | 822 | ENSG00000042493 |
| 15 | P42765 | NA | ACAA2 | acetyl-CoA acyltransferase 2 | 10449 | ENSG00000167315 |
| 16 | Q9Y277 | NA | VDAC3 | voltage-dependent anion channel 3 | 7419 | ENSG00000078668 |
| 17 | P17174 | NA | GOT1 | glutamic-oxaloacetic transaminase 1, soluble (aspartate aminotransferase 1) | 2805 | ENSG00000120053 |
| 18 | Q04837 | NA | SSBP1 | single-stranded DNA binding protein 1, mitochondrial | 6742 | ENSG00000106028 |
| 19 | P50454 | NA | SERPINH1 | serpin peptidase inhibitor, clade H (heat shock protein 47), member 1, (collagen binding protein 1) | 871 | ENSG00000149257 |
| 20 | Q14697 | NA | GANAB | glucosidase, alpha; neutral AB | 23193 | ENSG00000089597 |
| 21 | P08195 | NA | SLC3A2 | solute carrier family 3 (activators of dibasic and neutral amino acid transport), member 2 | 6520 | ENSG00000168003 |
| 22 | P40939 | NA | HADHA | hydroxyacyl-CoA dehydrogenase/3-ketoacyl-CoA thiolase/enoyl-CoA hydratase (trifunctional protein), alpha subunit | 3030 | ENSG00000084754 |
| 23 | P09382 | NA | LGALS1 | lectin, galactoside-binding, soluble, 1 | 3956 | ENSG00000100097 |
| 24 | P52815 | NA | MRPL12 | mitochondrial ribosomal protein L12 | 6182 | ENSG00000262814 |
| 25 | Q9BXW7 | NA | CECR5 | cat eye syndrome chromosome region, candidate 5 | 27440 | ENSG00000069998 |
| 26 | P21796 | NA | VDAC1 | voltage-dependent anion channel 1 | 7416 | ENSG00000213585 |
| 27 | P04080 | NA | CSTB | cystatin B (stefin B) | 1476 | ENSG00000160213 |
| 28 | P38571 | NA | LIPA | lipase A, lysosomal acid, cholesterol esterase | 3988 | ENSG00000107798 |
| 29 | P32322 | NA | PYCR1 | pyrroline-5-carboxylate reductase 1 | 5831 | ENSG00000183010 |
| 30 | Q9UJZ1 | NA | STOML2 | stomatin (EPB72)-like 2 | 30968 | ENSG00000165283 |
| 31 | Q14980 | NA | NUMA1 | nuclear mitotic apparatus protein 1 | 4926 | ENSG00000137497 |
| 32 | Q99798 | NA | ACO2 | aconitase 2, mitochondrial | 50 | ENSG00000100412 |
| 33 | P51649 | NA | ALDH5A1 | aldehyde dehydrogenase 5 family, member A1 | 7915 | ENSG00000112294 |
| 34 | P12235 | NA | SLC25A4 | solute carrier family 25 (mitochondrial carrier; adenine nucleotide translocator), member 4 | 291 | ENSG00000151729 |
| 35 | Q6PI48 | NA | DARS2 | aspartyl-tRNA synthetase 2, mitochondrial | 55157 | ENSG00000117593 |
| 36 | Q9NSE4 | NA | IARS2 | isoleucyl-tRNA synthetase 2, mitochondrial | 55699 | ENSG00000067704 |
| 37 | P05556 | NA | ITGB1 | integrin, beta 1 (fibronectin receptor, beta polypeptide, antigen CD29 includes MDF2, MSK12) | 3688 | ENSG00000150093 |
| 38 | P13674 | NA | P4HA1 | prolyl 4-hydroxylase, alpha polypeptide I | 5033 | ENSG00000122884 |
| 39 | Q02978 | NA | SLC25A11 | solute carrier family 25 (mitochondrial carrier; oxoglutarate carrier), member 11 | 8402 | ENSG00000108528 |
| 40 | P60174 | NA | TPI1 | triosephosphate isomerase 1 | 7167 | ENSG00000111669 |
| 41 | P30042 | NA | C21orf33 | chromosome 21 open reading frame 33 | 8209 | ENSG00000160221 |
| 42 | P54819 | NA | AK2 | adenylate kinase 2 | 204 | ENSG00000004455 |
| 43 | P17931 | NA | LGALS3 | lectin, galactoside-binding, soluble, 3 | 3958 | ENSG00000131981 |
| 44 | P05787 | NA | KRT8 | keratin 8 | 3856 | ENSG00000170421 |
| 45 | Q99623 | NA | PHB2 | prohibitin 2 | 11331 | ENSG00000215021 |
| 46 | P50453 | NA | SERPINB9 | serpin peptidase inhibitor, clade B (ovalbumin), member 9 | 5272 | ENSG00000170542 |
| 47 | P35232 | NA | PHB | prohibitin | 5245 | ENSG00000167085 |
| 48 | O95831 | NA | AIFM1 | apoptosis-inducing factor, mitochondrion-associated, 1 | 9131 | ENSG00000156709 |
| 49 | Q99714 | NA | HSD17B10 | hydroxysteroid (17-beta) dehydrogenase 10 | 3028 | ENSG00000072506 |
| 50 | P07741 | NA | APRT | adenine phosphoribosyltransferase | 353 | ENSG00000198931 |
| 51 | O60313 | NA | OPA1 | optic atrophy 1 (autosomal dominant) | 4976 | ENSG00000198836 |
| 52 | P35270 | NA | SPR | sepiapterin reductase (7,8-dihydrobiopterin:NADP+ oxidoreductase) | 6697 | ENSG00000116096 |
| 53 | P68104 | NA | EEF1A1 | eukaryotic translation elongation factor 1 alpha 1 | 1915 | ENSG00000156508 |
| 54 | P02786 | NA | TFRC | transferrin receptor (p90, CD71) | 7037 | ENSG00000072274 |
| 55 | P38117 | NA | ETFB | electron-transfer-flavoprotein, beta polypeptide | 2109 | ENSG00000105379 |
| 56 | O75367 | NA | H2AFY | H2A histone family, member Y | 9555 | ENSG00000113648 |
| 57 | O75521 | NA | ECI2 | enoyl-CoA delta isomerase 2 | 10455 | ENSG00000198721 |
| 58 | P67809 | NA | YBX1 | Y box binding protein 1 | 4904 | ENSG00000065978 |
| 59 | P49419 | NA | ALDH7A1 | aldehyde dehydrogenase 7 family, member A1 | 501 | ENSG00000164904 |
| 60 | Q6PIU2 | NA | NCEH1 | neutral cholesterol ester hydrolase 1 | 57552 | ENSG00000144959 |
| 61 | Q14764 | NA | MVP | major vault protein | 9961 | ENSG00000013364 |
| 62 | Q13813 | NA | SPTAN1 | spectrin, alpha, non-erythrocytic 1 | 6709 | ENSG00000197694 |
| 63 | P11387 | NA | TOP1 | topoisomerase (DNA) I | 7150 | ENSG00000198900 |
| 64 | Q9BPW8 | NA | NIPSNAP1 | nipsnap homolog 1 (C. elegans) | 8508 | ENSG00000184117 |
| 65 | Q5SSJ5 | NA | HP1BP3 | heterochromatin protein 1, binding protein 3 | 50809 | ENSG00000127483 |
| 66 | O75439 | NA | PMPCB | peptidase (mitochondrial processing) beta | 9512 | ENSG00000105819 |
| 67 | Q9NR30 | NA | DDX21 | DEAD (Asp-Glu-Ala-Asp) box helicase 21 | 9188 | ENSG00000165732 |
| 68 | P21953 | NA | BCKDHB | branched chain keto acid dehydrogenase E1, beta polypeptide | 594 | ENSG00000083123 |
| 69 | P30084 | NA | ECHS1 | enoyl CoA hydratase, short chain, 1, mitochondrial | 1892 | ENSG00000127884 |
| 70 | Q13011 | NA | ECH1 | enoyl CoA hydratase 1, peroxisomal | 1891 | ENSG00000104823 |
| 71 | P04083 | NA | ANXA1 | annexin A1 | 301 | ENSG00000135046 |
| 72 | P04040 | NA | CAT | catalase | 847 | ENSG00000121691 |
| 73 | O75340 | NA | PDCD6 | programmed cell death 6 | 10016 | ENSG00000249915 |
| 74 | P51688 | NA | SGSH | N-sulfoglucosamine sulfohydrolase | 6448 | ENSG00000181523 |
| 75 | P30048 | NA | PRDX3 | peroxiredoxin 3 | 10935 | ENSG00000165672 |
| 76 | P02768 | NA | ALB | albumin | 213 | ENSG00000163631 |
| 77 | P07108 | NA | DBI | diazepam binding inhibitor (GABA receptor modulator, acyl-CoA binding protein) | 1622 | ENSG00000155368 |
| 78 | P24752 | NA | ACAT1 | acetyl-CoA acetyltransferase 1 | 38 | ENSG00000075239 |
| 79 | P19367 | NA | HK1 | hexokinase 1 | 3098 | ENSG00000156515 |
| 80 | P25705 | NA | ATP5A1 | ATP synthase, H+ transporting, mitochondrial F1 complex, alpha subunit 1, cardiac muscle | 498 | ENSG00000152234 |
| 81 | Q9BQ69 | NA | MACROD1 | MACRO domain containing 1 | 28992 | ENSG00000133315 |
| 82 | P05783 | NA | KRT18 | keratin 18 | 3875 | ENSG00000111057 |
| 83 | P06865 | NA | HEXA | hexosaminidase A (alpha polypeptide) | 3073 | ENSG00000213614 |
| 84 | P49748 | NA | ACADVL | acyl-CoA dehydrogenase, very long chain | 37 | ENSG00000072778 |
| 85 | P48047 | NA | ATP5O | ATP synthase, H+ transporting, mitochondrial F1 complex, O subunit | 539 | ENSG00000241837 |
| 86 | Q16531 | NA | DDB1 | damage-specific DNA binding protein 1, 127kDa | 1642 | ENSG00000167986 |
| 87 | P31937 | NA | HIBADH | 3-hydroxyisobutyrate dehydrogenase | 11112 | ENSG00000106049 |

  
  

| **Database:cellular component      &nbspName:membrane-bounded organelle      &nbspID:GO:0043227** | | | | | | |
| --- | --- | --- | --- | --- | --- | --- |
| C=9495; O=87; E=61.47; R=1.42; rawP=1.74e-07; adjP=8.48e-07 | | | | | | |
| Index | UserID | Value | Gene Symbol | Gene Name | EntrezGene | Ensembl |
| 1 | P10253 | NA | GAA | glucosidase, alpha; acid | 2548 | ENSG00000171298 |
| 2 | P68371 | NA | TUBB4B | tubulin, beta 4B class IVb | 10383 | ENSG00000188229 |
| 3 | Q9UL46 | NA | PSME2 | proteasome (prosome, macropain) activator subunit 2 (PA28 beta) | 5721 | ENSG00000100911 |
| 4 | P40926 | NA | MDH2 | malate dehydrogenase 2, NAD (mitochondrial) | 4191 | ENSG00000146701 |
| 5 | P04179 | NA | SOD2 | superoxide dismutase 2, mitochondrial | 6648 | ENSG00000112096 |
| 6 | Q13724 | NA | MOGS | mannosyl-oligosaccharide glucosidase | 7841 | ENSG00000115275 |
| 7 | P48735 | NA | IDH2 | isocitrate dehydrogenase 2 (NADP+), mitochondrial | 3418 | ENSG00000182054 |
| 8 | P00367 | NA | GLUD1 | glutamate dehydrogenase 1 | 2746 | ENSG00000148672 |
| 9 | Q06323 | NA | PSME1 | proteasome (prosome, macropain) activator subunit 1 (PA28 alpha) | 5720 | ENSG00000092010 |
| 10 | P53701 | NA | HCCS | holocytochrome c synthase | 3052 | ENSG00000004961 |
| 11 | P31040 | NA | SDHA | succinate dehydrogenase complex, subunit A, flavoprotein (Fp) | 6389 | ENSG00000073578 |
| 12 | O14773 | NA | TPP1 | tripeptidyl peptidase I | 1200 | ENSG00000166340 |
| 13 | P13804 | NA | ETFA | electron-transfer-flavoprotein, alpha polypeptide | 2108 | ENSG00000140374 |
| 14 | P40121 | NA | CAPG | capping protein (actin filament), gelsolin-like | 822 | ENSG00000042493 |
| 15 | P42765 | NA | ACAA2 | acetyl-CoA acyltransferase 2 | 10449 | ENSG00000167315 |
| 16 | Q9Y277 | NA | VDAC3 | voltage-dependent anion channel 3 | 7419 | ENSG00000078668 |
| 17 | P17174 | NA | GOT1 | glutamic-oxaloacetic transaminase 1, soluble (aspartate aminotransferase 1) | 2805 | ENSG00000120053 |
| 18 | Q04837 | NA | SSBP1 | single-stranded DNA binding protein 1, mitochondrial | 6742 | ENSG00000106028 |
| 19 | P50454 | NA | SERPINH1 | serpin peptidase inhibitor, clade H (heat shock protein 47), member 1, (collagen binding protein 1) | 871 | ENSG00000149257 |
| 20 | Q14697 | NA | GANAB | glucosidase, alpha; neutral AB | 23193 | ENSG00000089597 |
| 21 | P08195 | NA | SLC3A2 | solute carrier family 3 (activators of dibasic and neutral amino acid transport), member 2 | 6520 | ENSG00000168003 |
| 22 | P40939 | NA | HADHA | hydroxyacyl-CoA dehydrogenase/3-ketoacyl-CoA thiolase/enoyl-CoA hydratase (trifunctional protein), alpha subunit | 3030 | ENSG00000084754 |
| 23 | P09382 | NA | LGALS1 | lectin, galactoside-binding, soluble, 1 | 3956 | ENSG00000100097 |
| 24 | P52815 | NA | MRPL12 | mitochondrial ribosomal protein L12 | 6182 | ENSG00000262814 |
| 25 | Q9BXW7 | NA | CECR5 | cat eye syndrome chromosome region, candidate 5 | 27440 | ENSG00000069998 |
| 26 | P21796 | NA | VDAC1 | voltage-dependent anion channel 1 | 7416 | ENSG00000213585 |
| 27 | P04080 | NA | CSTB | cystatin B (stefin B) | 1476 | ENSG00000160213 |
| 28 | P38571 | NA | LIPA | lipase A, lysosomal acid, cholesterol esterase | 3988 | ENSG00000107798 |
| 29 | P32322 | NA | PYCR1 | pyrroline-5-carboxylate reductase 1 | 5831 | ENSG00000183010 |
| 30 | Q9UJZ1 | NA | STOML2 | stomatin (EPB72)-like 2 | 30968 | ENSG00000165283 |
| 31 | Q14980 | NA | NUMA1 | nuclear mitotic apparatus protein 1 | 4926 | ENSG00000137497 |
| 32 | Q99798 | NA | ACO2 | aconitase 2, mitochondrial | 50 | ENSG00000100412 |
| 33 | P51649 | NA | ALDH5A1 | aldehyde dehydrogenase 5 family, member A1 | 7915 | ENSG00000112294 |
| 34 | P12235 | NA | SLC25A4 | solute carrier family 25 (mitochondrial carrier; adenine nucleotide translocator), member 4 | 291 | ENSG00000151729 |
| 35 | Q6PI48 | NA | DARS2 | aspartyl-tRNA synthetase 2, mitochondrial | 55157 | ENSG00000117593 |
| 36 | Q9NSE4 | NA | IARS2 | isoleucyl-tRNA synthetase 2, mitochondrial | 55699 | ENSG00000067704 |
| 37 | P05556 | NA | ITGB1 | integrin, beta 1 (fibronectin receptor, beta polypeptide, antigen CD29 includes MDF2, MSK12) | 3688 | ENSG00000150093 |
| 38 | P13674 | NA | P4HA1 | prolyl 4-hydroxylase, alpha polypeptide I | 5033 | ENSG00000122884 |
| 39 | Q02978 | NA | SLC25A11 | solute carrier family 25 (mitochondrial carrier; oxoglutarate carrier), member 11 | 8402 | ENSG00000108528 |
| 40 | P60174 | NA | TPI1 | triosephosphate isomerase 1 | 7167 | ENSG00000111669 |
| 41 | P30042 | NA | C21orf33 | chromosome 21 open reading frame 33 | 8209 | ENSG00000160221 |
| 42 | P54819 | NA | AK2 | adenylate kinase 2 | 204 | ENSG00000004455 |
| 43 | P17931 | NA | LGALS3 | lectin, galactoside-binding, soluble, 3 | 3958 | ENSG00000131981 |
| 44 | P05787 | NA | KRT8 | keratin 8 | 3856 | ENSG00000170421 |
| 45 | Q99623 | NA | PHB2 | prohibitin 2 | 11331 | ENSG00000215021 |
| 46 | P50453 | NA | SERPINB9 | serpin peptidase inhibitor, clade B (ovalbumin), member 9 | 5272 | ENSG00000170542 |
| 47 | P35232 | NA | PHB | prohibitin | 5245 | ENSG00000167085 |
| 48 | O95831 | NA | AIFM1 | apoptosis-inducing factor, mitochondrion-associated, 1 | 9131 | ENSG00000156709 |
| 49 | Q99714 | NA | HSD17B10 | hydroxysteroid (17-beta) dehydrogenase 10 | 3028 | ENSG00000072506 |
| 50 | P07741 | NA | APRT | adenine phosphoribosyltransferase | 353 | ENSG00000198931 |
| 51 | O60313 | NA | OPA1 | optic atrophy 1 (autosomal dominant) | 4976 | ENSG00000198836 |
| 52 | P35270 | NA | SPR | sepiapterin reductase (7,8-dihydrobiopterin:NADP+ oxidoreductase) | 6697 | ENSG00000116096 |
| 53 | P68104 | NA | EEF1A1 | eukaryotic translation elongation factor 1 alpha 1 | 1915 | ENSG00000156508 |
| 54 | P02786 | NA | TFRC | transferrin receptor (p90, CD71) | 7037 | ENSG00000072274 |
| 55 | P38117 | NA | ETFB | electron-transfer-flavoprotein, beta polypeptide | 2109 | ENSG00000105379 |
| 56 | O75367 | NA | H2AFY | H2A histone family, member Y | 9555 | ENSG00000113648 |
| 57 | O75521 | NA | ECI2 | enoyl-CoA delta isomerase 2 | 10455 | ENSG00000198721 |
| 58 | P67809 | NA | YBX1 | Y box binding protein 1 | 4904 | ENSG00000065978 |
| 59 | P49419 | NA | ALDH7A1 | aldehyde dehydrogenase 7 family, member A1 | 501 | ENSG00000164904 |
| 60 | Q6PIU2 | NA | NCEH1 | neutral cholesterol ester hydrolase 1 | 57552 | ENSG00000144959 |
| 61 | Q14764 | NA | MVP | major vault protein | 9961 | ENSG00000013364 |
| 62 | Q13813 | NA | SPTAN1 | spectrin, alpha, non-erythrocytic 1 | 6709 | ENSG00000197694 |
| 63 | P11387 | NA | TOP1 | topoisomerase (DNA) I | 7150 | ENSG00000198900 |
| 64 | Q9BPW8 | NA | NIPSNAP1 | nipsnap homolog 1 (C. elegans) | 8508 | ENSG00000184117 |
| 65 | Q5SSJ5 | NA | HP1BP3 | heterochromatin protein 1, binding protein 3 | 50809 | ENSG00000127483 |
| 66 | O75439 | NA | PMPCB | peptidase (mitochondrial processing) beta | 9512 | ENSG00000105819 |
| 67 | Q9NR30 | NA | DDX21 | DEAD (Asp-Glu-Ala-Asp) box helicase 21 | 9188 | ENSG00000165732 |
| 68 | P21953 | NA | BCKDHB | branched chain keto acid dehydrogenase E1, beta polypeptide | 594 | ENSG00000083123 |
| 69 | P30084 | NA | ECHS1 | enoyl CoA hydratase, short chain, 1, mitochondrial | 1892 | ENSG00000127884 |
| 70 | Q13011 | NA | ECH1 | enoyl CoA hydratase 1, peroxisomal | 1891 | ENSG00000104823 |
| 71 | P04083 | NA | ANXA1 | annexin A1 | 301 | ENSG00000135046 |
| 72 | P04040 | NA | CAT | catalase | 847 | ENSG00000121691 |
| 73 | O75340 | NA | PDCD6 | programmed cell death 6 | 10016 | ENSG00000249915 |
| 74 | P51688 | NA | SGSH | N-sulfoglucosamine sulfohydrolase | 6448 | ENSG00000181523 |
| 75 | P30048 | NA | PRDX3 | peroxiredoxin 3 | 10935 | ENSG00000165672 |
| 76 | P02768 | NA | ALB | albumin | 213 | ENSG00000163631 |
| 77 | P07108 | NA | DBI | diazepam binding inhibitor (GABA receptor modulator, acyl-CoA binding protein) | 1622 | ENSG00000155368 |
| 78 | P24752 | NA | ACAT1 | acetyl-CoA acetyltransferase 1 | 38 | ENSG00000075239 |
| 79 | P19367 | NA | HK1 | hexokinase 1 | 3098 | ENSG00000156515 |
| 80 | P25705 | NA | ATP5A1 | ATP synthase, H+ transporting, mitochondrial F1 complex, alpha subunit 1, cardiac muscle | 498 | ENSG00000152234 |
| 81 | Q9BQ69 | NA | MACROD1 | MACRO domain containing 1 | 28992 | ENSG00000133315 |
| 82 | P05783 | NA | KRT18 | keratin 18 | 3875 | ENSG00000111057 |
| 83 | P06865 | NA | HEXA | hexosaminidase A (alpha polypeptide) | 3073 | ENSG00000213614 |
| 84 | P49748 | NA | ACADVL | acyl-CoA dehydrogenase, very long chain | 37 | ENSG00000072778 |
| 85 | P48047 | NA | ATP5O | ATP synthase, H+ transporting, mitochondrial F1 complex, O subunit | 539 | ENSG00000241837 |
| 86 | Q16531 | NA | DDB1 | damage-specific DNA binding protein 1, 127kDa | 1642 | ENSG00000167986 |
| 87 | P31937 | NA | HIBADH | 3-hydroxyisobutyrate dehydrogenase | 11112 | ENSG00000106049 |

  
  

| **Database:cellular component      &nbspName:eukaryotic translation elongation factor 1 complex      &nbspID:GO:0005853** | | | | | | |
| --- | --- | --- | --- | --- | --- | --- |
| C=5; O=3; E=0.03; R=92.68; rawP=2.61e-06; adjP=1.22e-05 | | | | | | |
| Index | UserID | Value | Gene Symbol | Gene Name | EntrezGene | Ensembl |
| 1 | P68104 | NA | EEF1A1 | eukaryotic translation elongation factor 1 alpha 1 | 1915 | ENSG00000156508 |
| 2 | P29692 | NA | EEF1D | eukaryotic translation elongation factor 1 delta (guanine nucleotide exchange protein) | 1936 | ENSG00000104529 |
| 3 | P24534 | NA | EEF1B2 | eukaryotic translation elongation factor 1 beta 2 | 1933 | ENSG00000114942 |

  
  

| **Database:cellular component      &nbspName:mitochondrial nucleoid      &nbspID:GO:0042645** | | | | | | |
| --- | --- | --- | --- | --- | --- | --- |
| C=37; O=5; E=0.24; R=20.87; rawP=3.83e-06; adjP=1.72e-05 | | | | | | |
| Index | UserID | Value | Gene Symbol | Gene Name | EntrezGene | Ensembl |
| 1 | Q04837 | NA | SSBP1 | single-stranded DNA binding protein 1, mitochondrial | 6742 | ENSG00000106028 |
| 2 | P21796 | NA | VDAC1 | voltage-dependent anion channel 1 | 7416 | ENSG00000213585 |
| 3 | P49748 | NA | ACADVL | acyl-CoA dehydrogenase, very long chain | 37 | ENSG00000072778 |
| 4 | P40939 | NA | HADHA | hydroxyacyl-CoA dehydrogenase/3-ketoacyl-CoA thiolase/enoyl-CoA hydratase (trifunctional protein), alpha subunit | 3030 | ENSG00000084754 |
| 5 | P04179 | NA | SOD2 | superoxide dismutase 2, mitochondrial | 6648 | ENSG00000112096 |

  
  

| **Database:cellular component      &nbspName:nucleoid      &nbspID:GO:0009295** | | | | | | |
| --- | --- | --- | --- | --- | --- | --- |
| C=40; O=5; E=0.26; R=19.31; rawP=5.69e-06; adjP=2.47e-05 | | | | | | |
| Index | UserID | Value | Gene Symbol | Gene Name | EntrezGene | Ensembl |
| 1 | Q04837 | NA | SSBP1 | single-stranded DNA binding protein 1, mitochondrial | 6742 | ENSG00000106028 |
| 2 | P21796 | NA | VDAC1 | voltage-dependent anion channel 1 | 7416 | ENSG00000213585 |
| 3 | P49748 | NA | ACADVL | acyl-CoA dehydrogenase, very long chain | 37 | ENSG00000072778 |
| 4 | P40939 | NA | HADHA | hydroxyacyl-CoA dehydrogenase/3-ketoacyl-CoA thiolase/enoyl-CoA hydratase (trifunctional protein), alpha subunit | 3030 | ENSG00000084754 |
| 5 | P04179 | NA | SOD2 | superoxide dismutase 2, mitochondrial | 6648 | ENSG00000112096 |

  
  

| **Database:cellular component      &nbspName:organelle membrane      &nbspID:GO:0031090** | | | | | | |
| --- | --- | --- | --- | --- | --- | --- |
| C=2361; O=33; E=15.29; R=2.16; rawP=8.99e-06; adjP=3.76e-05 | | | | | | |
| Index | UserID | Value | Gene Symbol | Gene Name | EntrezGene | Ensembl |
| 1 | O75439 | NA | PMPCB | peptidase (mitochondrial processing) beta | 9512 | ENSG00000105819 |
| 2 | P10253 | NA | GAA | glucosidase, alpha; acid | 2548 | ENSG00000171298 |
| 3 | P54819 | NA | AK2 | adenylate kinase 2 | 204 | ENSG00000004455 |
| 4 | P17931 | NA | LGALS3 | lectin, galactoside-binding, soluble, 3 | 3958 | ENSG00000131981 |
| 5 | Q99623 | NA | PHB2 | prohibitin 2 | 11331 | ENSG00000215021 |
| 6 | P40939 | NA | HADHA | hydroxyacyl-CoA dehydrogenase/3-ketoacyl-CoA thiolase/enoyl-CoA hydratase (trifunctional protein), alpha subunit | 3030 | ENSG00000084754 |
| 7 | P40926 | NA | MDH2 | malate dehydrogenase 2, NAD (mitochondrial) | 4191 | ENSG00000146701 |
| 8 | P35232 | NA | PHB | prohibitin | 5245 | ENSG00000167085 |
| 9 | Q99714 | NA | HSD17B10 | hydroxysteroid (17-beta) dehydrogenase 10 | 3028 | ENSG00000072506 |
| 10 | O95831 | NA | AIFM1 | apoptosis-inducing factor, mitochondrion-associated, 1 | 9131 | ENSG00000156709 |
| 11 | P04179 | NA | SOD2 | superoxide dismutase 2, mitochondrial | 6648 | ENSG00000112096 |
| 12 | Q13724 | NA | MOGS | mannosyl-oligosaccharide glucosidase | 7841 | ENSG00000115275 |
| 13 | P48735 | NA | IDH2 | isocitrate dehydrogenase 2 (NADP+), mitochondrial | 3418 | ENSG00000182054 |
| 14 | O60313 | NA | OPA1 | optic atrophy 1 (autosomal dominant) | 4976 | ENSG00000198836 |
| 15 | P21796 | NA | VDAC1 | voltage-dependent anion channel 1 | 7416 | ENSG00000213585 |
| 16 | P53701 | NA | HCCS | holocytochrome c synthase | 3052 | ENSG00000004961 |
| 17 | P31040 | NA | SDHA | succinate dehydrogenase complex, subunit A, flavoprotein (Fp) | 6389 | ENSG00000073578 |
| 18 | P02786 | NA | TFRC | transferrin receptor (p90, CD71) | 7037 | ENSG00000072274 |
| 19 | P04083 | NA | ANXA1 | annexin A1 | 301 | ENSG00000135046 |
| 20 | Q9UJZ1 | NA | STOML2 | stomatin (EPB72)-like 2 | 30968 | ENSG00000165283 |
| 21 | P04040 | NA | CAT | catalase | 847 | ENSG00000121691 |
| 22 | O75340 | NA | PDCD6 | programmed cell death 6 | 10016 | ENSG00000249915 |
| 23 | P12235 | NA | SLC25A4 | solute carrier family 25 (mitochondrial carrier; adenine nucleotide translocator), member 4 | 291 | ENSG00000151729 |
| 24 | P40121 | NA | CAPG | capping protein (actin filament), gelsolin-like | 822 | ENSG00000042493 |
| 25 | P42765 | NA | ACAA2 | acetyl-CoA acyltransferase 2 | 10449 | ENSG00000167315 |
| 26 | P24752 | NA | ACAT1 | acetyl-CoA acetyltransferase 1 | 38 | ENSG00000075239 |
| 27 | Q9Y277 | NA | VDAC3 | voltage-dependent anion channel 3 | 7419 | ENSG00000078668 |
| 28 | Q02978 | NA | SLC25A11 | solute carrier family 25 (mitochondrial carrier; oxoglutarate carrier), member 11 | 8402 | ENSG00000108528 |
| 29 | P19367 | NA | HK1 | hexokinase 1 | 3098 | ENSG00000156515 |
| 30 | P25705 | NA | ATP5A1 | ATP synthase, H+ transporting, mitochondrial F1 complex, alpha subunit 1, cardiac muscle | 498 | ENSG00000152234 |
| 31 | P49748 | NA | ACADVL | acyl-CoA dehydrogenase, very long chain | 37 | ENSG00000072778 |
| 32 | P48047 | NA | ATP5O | ATP synthase, H+ transporting, mitochondrial F1 complex, O subunit | 539 | ENSG00000241837 |
| 33 | Q9BPW8 | NA | NIPSNAP1 | nipsnap homolog 1 (C. elegans) | 8508 | ENSG00000184117 |

  
  

| **Database:cellular component      &nbspName:pigment granule      &nbspID:GO:0048770** | | | | | | |
| --- | --- | --- | --- | --- | --- | --- |
| C=93; O=6; E=0.60; R=9.97; rawP=3.09e-05; adjP=0.0001 | | | | | | |
| Index | UserID | Value | Gene Symbol | Gene Name | EntrezGene | Ensembl |
| 1 | O14773 | NA | TPP1 | tripeptidyl peptidase I | 1200 | ENSG00000166340 |
| 2 | P08195 | NA | SLC3A2 | solute carrier family 3 (activators of dibasic and neutral amino acid transport), member 2 | 6520 | ENSG00000168003 |
| 3 | Q14697 | NA | GANAB | glucosidase, alpha; neutral AB | 23193 | ENSG00000089597 |
| 4 | P02786 | NA | TFRC | transferrin receptor (p90, CD71) | 7037 | ENSG00000072274 |
| 5 | P05556 | NA | ITGB1 | integrin, beta 1 (fibronectin receptor, beta polypeptide, antigen CD29 includes MDF2, MSK12) | 3688 | ENSG00000150093 |
| 6 | P40121 | NA | CAPG | capping protein (actin filament), gelsolin-like | 822 | ENSG00000042493 |

  
  

| **Database:cellular component      &nbspName:cytosol      &nbspID:GO:0005829** | | | | | | |
| --- | --- | --- | --- | --- | --- | --- |
| C=2367; O=32; E=15.32; R=2.09; rawP=2.57e-05; adjP=0.0001 | | | | | | |
| Index | UserID | Value | Gene Symbol | Gene Name | EntrezGene | Ensembl |
| 1 | P49589 | NA | CARS | cysteinyl-tRNA synthetase | 833 | ENSG00000110619 |
| 2 | P68371 | NA | TUBB4B | tubulin, beta 4B class IVb | 10383 | ENSG00000188229 |
| 3 | Q9UL46 | NA | PSME2 | proteasome (prosome, macropain) activator subunit 2 (PA28 beta) | 5721 | ENSG00000100911 |
| 4 | P50453 | NA | SERPINB9 | serpin peptidase inhibitor, clade B (ovalbumin), member 9 | 5272 | ENSG00000170542 |
| 5 | O95831 | NA | AIFM1 | apoptosis-inducing factor, mitochondrion-associated, 1 | 9131 | ENSG00000156709 |
| 6 | P61313 | NA | RPL15 | ribosomal protein L15 | 6138 | ENSG00000174748 |
| 7 | P07741 | NA | APRT | adenine phosphoribosyltransferase | 353 | ENSG00000198931 |
| 8 | Q06323 | NA | PSME1 | proteasome (prosome, macropain) activator subunit 1 (PA28 alpha) | 5720 | ENSG00000092010 |
| 9 | P35270 | NA | SPR | sepiapterin reductase (7,8-dihydrobiopterin:NADP+ oxidoreductase) | 6697 | ENSG00000116096 |
| 10 | P68104 | NA | EEF1A1 | eukaryotic translation elongation factor 1 alpha 1 | 1915 | ENSG00000156508 |
| 11 | O75369 | NA | FLNB | filamin B, beta | 2317 | ENSG00000136068 |
| 12 | P43490 | NA | NAMPT | nicotinamide phosphoribosyltransferase | 10135 | ENSG00000105835 |
| 13 | P24534 | NA | EEF1B2 | eukaryotic translation elongation factor 1 beta 2 | 1933 | ENSG00000114942 |
| 14 | P22102 | NA | GART | phosphoribosylglycinamide formyltransferase, phosphoribosylglycinamide synthetase, phosphoribosylaminoimidazole synthetase | 2618 | ENSG00000159131 |
| 15 | P13639 | NA | EEF2 | eukaryotic translation elongation factor 2 | 1938 | ENSG00000167658 |
| 16 | P49419 | NA | ALDH7A1 | aldehyde dehydrogenase 7 family, member A1 | 501 | ENSG00000164904 |
| 17 | P17174 | NA | GOT1 | glutamic-oxaloacetic transaminase 1, soluble (aspartate aminotransferase 1) | 2805 | ENSG00000120053 |
| 18 | P18206 | NA | VCL | vinculin | 7414 | ENSG00000035403 |
| 19 | P29692 | NA | EEF1D | eukaryotic translation elongation factor 1 delta (guanine nucleotide exchange protein) | 1936 | ENSG00000104529 |
| 20 | Q13813 | NA | SPTAN1 | spectrin, alpha, non-erythrocytic 1 | 6709 | ENSG00000197694 |
| 21 | P02792 | NA | FTL | ferritin, light polypeptide | 2512 | ENSG00000087086 |
| 22 | P00491 | NA | PNP | purine nucleoside phosphorylase | 4860 | ENSG00000198805 |
| 23 | P60842 | NA | EIF4A1 | eukaryotic translation initiation factor 4A1 | 1973 | ENSG00000161960 |
| 24 | Q14980 | NA | NUMA1 | nuclear mitotic apparatus protein 1 | 4926 | ENSG00000137497 |
| 25 | P04040 | NA | CAT | catalase | 847 | ENSG00000121691 |
| 26 | P41091 | NA | EIF2S3 | eukaryotic translation initiation factor 2, subunit 3 gamma, 52kDa | 1968 | ENSG00000130741 |
| 27 | P30048 | NA | PRDX3 | peroxiredoxin 3 | 10935 | ENSG00000165672 |
| 28 | P00558 | NA | PGK1 | phosphoglycerate kinase 1 | 5230 | ENSG00000102144 |
| 29 | P40227 | NA | CCT6A | chaperonin containing TCP1, subunit 6A (zeta 1) | 908 | ENSG00000146731 |
| 30 | P19367 | NA | HK1 | hexokinase 1 | 3098 | ENSG00000156515 |
| 31 | P00352 | NA | ALDH1A1 | aldehyde dehydrogenase 1 family, member A1 | 216 | ENSG00000165092 |
| 32 | P60174 | NA | TPI1 | triosephosphate isomerase 1 | 7167 | ENSG00000111669 |

  
  

| **Database:cellular component      &nbspName:melanosome      &nbspID:GO:0042470** | | | | | | |
| --- | --- | --- | --- | --- | --- | --- |
| C=93; O=6; E=0.60; R=9.97; rawP=3.09e-05; adjP=0.0001 | | | | | | |
| Index | UserID | Value | Gene Symbol | Gene Name | EntrezGene | Ensembl |
| 1 | O14773 | NA | TPP1 | tripeptidyl peptidase I | 1200 | ENSG00000166340 |
| 2 | P08195 | NA | SLC3A2 | solute carrier family 3 (activators of dibasic and neutral amino acid transport), member 2 | 6520 | ENSG00000168003 |
| 3 | Q14697 | NA | GANAB | glucosidase, alpha; neutral AB | 23193 | ENSG00000089597 |
| 4 | P02786 | NA | TFRC | transferrin receptor (p90, CD71) | 7037 | ENSG00000072274 |
| 5 | P05556 | NA | ITGB1 | integrin, beta 1 (fibronectin receptor, beta polypeptide, antigen CD29 includes MDF2, MSK12) | 3688 | ENSG00000150093 |
| 6 | P40121 | NA | CAPG | capping protein (actin filament), gelsolin-like | 822 | ENSG00000042493 |

  
  

| **Database:cellular component      &nbspName:mitochondrial intermembrane space      &nbspID:GO:0005758** | | | | | | |
| --- | --- | --- | --- | --- | --- | --- |
| C=44; O=4; E=0.28; R=14.04; rawP=0.0002; adjP=0.0007 | | | | | | |
| Index | UserID | Value | Gene Symbol | Gene Name | EntrezGene | Ensembl |
| 1 | P04040 | NA | CAT | catalase | 847 | ENSG00000121691 |
| 2 | P54819 | NA | AK2 | adenylate kinase 2 | 204 | ENSG00000004455 |
| 3 | O60313 | NA | OPA1 | optic atrophy 1 (autosomal dominant) | 4976 | ENSG00000198836 |
| 4 | O95831 | NA | AIFM1 | apoptosis-inducing factor, mitochondrion-associated, 1 | 9131 | ENSG00000156709 |

  
  
  
  

---

WebGestalt is currently developed and maintained by Jing Wang and Bing Zhang at the  Zhang Lab. Other people who have made significant contribution to the project include Dexter Duncan, Stefan Kirov, Zhiao Shi, and Jay Snoddy.  
  
**Funding credits:** NIH/NIAAA (U01 AA016662, U01 AA013512); NIH/NIDA (P01 DA015027); NIH/NIMH (P50 MH078028, P50 MH096972); NIH/NCI (U24 CA159988); NIH/NIGMS (R01 GM088822).
