## Supplementary material for "PGRMC1 phosphorylation and cell plasticity 1: glycolysis, mitochondria, tumor growth": File S7: final_pc_geneset_file_1438480126.html

Anchored HTML File of EIDs

|  |  |
| --- | --- |
|  | WEB-based GEne SeT AnaLysis Toolkit |
|  | ***Translating gene lists into biological insights...*** |

---

  

| **Database:Pathway Commons pathway      &nbspName:Metabolism      &nbspID:DB\_ID:634** | | | | | | |
| --- | --- | --- | --- | --- | --- | --- |
| C=824; O=28; E=4.39; R=6.38; rawP=1.96e-15; adjP=2.25e-13 | | | | | | |
| Index | UserID | Value | Gene Symbol | Gene Name | EntrezGene | Ensembl |
| 1 | P54819 | NA | AK2 | adenylate kinase 2 | 204 | ENSG00000004455 |
| 2 | P00491 | NA | PNP | purine nucleoside phosphorylase | 4860 | ENSG00000198805 |
| 3 | P21953 | NA | BCKDHB | branched chain keto acid dehydrogenase E1, beta polypeptide | 594 | ENSG00000083123 |
| 4 | Q9UL46 | NA | PSME2 | proteasome (prosome, macropain) activator subunit 2 (PA28 beta) | 5721 | ENSG00000100911 |
| 5 | P40926 | NA | MDH2 | malate dehydrogenase 2, NAD (mitochondrial) | 4191 | ENSG00000146701 |
| 6 | Q99714 | NA | HSD17B10 | hydroxysteroid (17-beta) dehydrogenase 10 | 3028 | ENSG00000072506 |
| 7 | P61313 | NA | RPL15 | ribosomal protein L15 | 6138 | ENSG00000174748 |
| 8 | P60842 | NA | EIF4A1 | eukaryotic translation initiation factor 4A1 | 1973 | ENSG00000161960 |
| 9 | P07741 | NA | APRT | adenine phosphoribosyltransferase | 353 | ENSG00000198931 |
| 10 | P00367 | NA | GLUD1 | glutamate dehydrogenase 1 | 2746 | ENSG00000148672 |
| 11 | Q06323 | NA | PSME1 | proteasome (prosome, macropain) activator subunit 1 (PA28 alpha) | 5720 | ENSG00000092010 |
| 12 | P48735 | NA | IDH2 | isocitrate dehydrogenase 2 (NADP+), mitochondrial | 3418 | ENSG00000182054 |
| 13 | P35270 | NA | SPR | sepiapterin reductase (7,8-dihydrobiopterin:NADP+ oxidoreductase) | 6697 | ENSG00000116096 |
| 14 | P31040 | NA | SDHA | succinate dehydrogenase complex, subunit A, flavoprotein (Fp) | 6389 | ENSG00000073578 |
| 15 | P38117 | NA | ETFB | electron-transfer-flavoprotein, beta polypeptide | 2109 | ENSG00000105379 |
| 16 | P43490 | NA | NAMPT | nicotinamide phosphoribosyltransferase | 10135 | ENSG00000105835 |
| 17 | P32322 | NA | PYCR1 | pyrroline-5-carboxylate reductase 1 | 5831 | ENSG00000183010 |
| 18 | P04040 | NA | CAT | catalase | 847 | ENSG00000121691 |
| 19 | Q99798 | NA | ACO2 | aconitase 2, mitochondrial | 50 | ENSG00000100412 |
| 20 | P22102 | NA | GART | phosphoribosylglycinamide formyltransferase, phosphoribosylglycinamide synthetase, phosphoribosylaminoimidazole synthetase | 2618 | ENSG00000159131 |
| 21 | P13804 | NA | ETFA | electron-transfer-flavoprotein, alpha polypeptide | 2108 | ENSG00000140374 |
| 22 | P24752 | NA | ACAT1 | acetyl-CoA acetyltransferase 1 | 38 | ENSG00000075239 |
| 23 | P49419 | NA | ALDH7A1 | aldehyde dehydrogenase 7 family, member A1 | 501 | ENSG00000164904 |
| 24 | P17174 | NA | GOT1 | glutamic-oxaloacetic transaminase 1, soluble (aspartate aminotransferase 1) | 2805 | ENSG00000120053 |
| 25 | P25705 | NA | ATP5A1 | ATP synthase, H+ transporting, mitochondrial F1 complex, alpha subunit 1, cardiac muscle | 498 | ENSG00000152234 |
| 26 | P00352 | NA | ALDH1A1 | aldehyde dehydrogenase 1 family, member A1 | 216 | ENSG00000165092 |
| 27 | P48047 | NA | ATP5O | ATP synthase, H+ transporting, mitochondrial F1 complex, O subunit | 539 | ENSG00000241837 |
| 28 | P31937 | NA | HIBADH | 3-hydroxyisobutyrate dehydrogenase | 11112 | ENSG00000106049 |

  
  

| **Database:Pathway Commons pathway      &nbspName:Gluconeogenesis      &nbspID:DB\_ID:929** | | | | | | |
| --- | --- | --- | --- | --- | --- | --- |
| C=20; O=5; E=0.11; R=46.95; rawP=5.67e-08; adjP=1.87e-06 | | | | | | |
| Index | UserID | Value | Gene Symbol | Gene Name | EntrezGene | Ensembl |
| 1 | Q02978 | NA | SLC25A11 | solute carrier family 25 (mitochondrial carrier; oxoglutarate carrier), member 11 | 8402 | ENSG00000108528 |
| 2 | P17174 | NA | GOT1 | glutamic-oxaloacetic transaminase 1, soluble (aspartate aminotransferase 1) | 2805 | ENSG00000120053 |
| 3 | P00558 | NA | PGK1 | phosphoglycerate kinase 1 | 5230 | ENSG00000102144 |
| 4 | P40926 | NA | MDH2 | malate dehydrogenase 2, NAD (mitochondrial) | 4191 | ENSG00000146701 |
| 5 | P60174 | NA | TPI1 | triosephosphate isomerase 1 | 7167 | ENSG00000111669 |

  
  

| **Database:Pathway Commons pathway      &nbspName:Metabolism of amino acids and derivatives      &nbspID:DB\_ID:875** | | | | | | |
| --- | --- | --- | --- | --- | --- | --- |
| C=188; O=10; E=1.00; R=9.99; rawP=6.50e-08; adjP=1.87e-06 | | | | | | |
| Index | UserID | Value | Gene Symbol | Gene Name | EntrezGene | Ensembl |
| 1 | P21953 | NA | BCKDHB | branched chain keto acid dehydrogenase E1, beta polypeptide | 594 | ENSG00000083123 |
| 2 | Q9UL46 | NA | PSME2 | proteasome (prosome, macropain) activator subunit 2 (PA28 beta) | 5721 | ENSG00000100911 |
| 3 | Q99714 | NA | HSD17B10 | hydroxysteroid (17-beta) dehydrogenase 10 | 3028 | ENSG00000072506 |
| 4 | P24752 | NA | ACAT1 | acetyl-CoA acetyltransferase 1 | 38 | ENSG00000075239 |
| 5 | P00367 | NA | GLUD1 | glutamate dehydrogenase 1 | 2746 | ENSG00000148672 |
| 6 | Q06323 | NA | PSME1 | proteasome (prosome, macropain) activator subunit 1 (PA28 alpha) | 5720 | ENSG00000092010 |
| 7 | P17174 | NA | GOT1 | glutamic-oxaloacetic transaminase 1, soluble (aspartate aminotransferase 1) | 2805 | ENSG00000120053 |
| 8 | P49419 | NA | ALDH7A1 | aldehyde dehydrogenase 7 family, member A1 | 501 | ENSG00000164904 |
| 9 | P32322 | NA | PYCR1 | pyrroline-5-carboxylate reductase 1 | 5831 | ENSG00000183010 |
| 10 | P31937 | NA | HIBADH | 3-hydroxyisobutyrate dehydrogenase | 11112 | ENSG00000106049 |

  
  

| **Database:Pathway Commons pathway      &nbspName:fatty acid beta-oxidation I      &nbspID:DB\_ID:1395** | | | | | | |
| --- | --- | --- | --- | --- | --- | --- |
| C=19; O=5; E=0.10; R=49.42; rawP=4.27e-08; adjP=1.87e-06 | | | | | | |
| Index | UserID | Value | Gene Symbol | Gene Name | EntrezGene | Ensembl |
| 1 | O75521 | NA | ECI2 | enoyl-CoA delta isomerase 2 | 10455 | ENSG00000198721 |
| 2 | P40939 | NA | HADHA | hydroxyacyl-CoA dehydrogenase/3-ketoacyl-CoA thiolase/enoyl-CoA hydratase (trifunctional protein), alpha subunit | 3030 | ENSG00000084754 |
| 3 | P30084 | NA | ECHS1 | enoyl CoA hydratase, short chain, 1, mitochondrial | 1892 | ENSG00000127884 |
| 4 | Q99714 | NA | HSD17B10 | hydroxysteroid (17-beta) dehydrogenase 10 | 3028 | ENSG00000072506 |
| 5 | P42765 | NA | ACAA2 | acetyl-CoA acyltransferase 2 | 10449 | ENSG00000167315 |

  
  

| **Database:Pathway Commons pathway      &nbspName:Metabolism of proteins      &nbspID:DB\_ID:728** | | | | | | |
| --- | --- | --- | --- | --- | --- | --- |
| C=260; O=11; E=1.38; R=7.95; rawP=1.43e-07; adjP=3.29e-06 | | | | | | |
| Index | UserID | Value | Gene Symbol | Gene Name | EntrezGene | Ensembl |
| 1 | P13639 | NA | EEF2 | eukaryotic translation elongation factor 2 | 1938 | ENSG00000167658 |
| 2 | P41091 | NA | EIF2S3 | eukaryotic translation initiation factor 2, subunit 3 gamma, 52kDa | 1968 | ENSG00000130741 |
| 3 | P68371 | NA | TUBB4B | tubulin, beta 4B class IVb | 10383 | ENSG00000188229 |
| 4 | Q13724 | NA | MOGS | mannosyl-oligosaccharide glucosidase | 7841 | ENSG00000115275 |
| 5 | P40227 | NA | CCT6A | chaperonin containing TCP1, subunit 6A (zeta 1) | 908 | ENSG00000146731 |
| 6 | P61313 | NA | RPL15 | ribosomal protein L15 | 6138 | ENSG00000174748 |
| 7 | P60842 | NA | EIF4A1 | eukaryotic translation initiation factor 4A1 | 1973 | ENSG00000161960 |
| 8 | Q14697 | NA | GANAB | glucosidase, alpha; neutral AB | 23193 | ENSG00000089597 |
| 9 | P68104 | NA | EEF1A1 | eukaryotic translation elongation factor 1 alpha 1 | 1915 | ENSG00000156508 |
| 10 | P29692 | NA | EEF1D | eukaryotic translation elongation factor 1 delta (guanine nucleotide exchange protein) | 1936 | ENSG00000104529 |
| 11 | P24534 | NA | EEF1B2 | eukaryotic translation elongation factor 1 beta 2 | 1933 | ENSG00000114942 |

  
  

| **Database:Pathway Commons pathway      &nbspName:The citric acid (TCA) cycle and respiratory electron transport      &nbspID:DB\_ID:547** | | | | | | |
| --- | --- | --- | --- | --- | --- | --- |
| C=118; O=8; E=0.63; R=12.73; rawP=2.24e-07; adjP=4.01e-06 | | | | | | |
| Index | UserID | Value | Gene Symbol | Gene Name | EntrezGene | Ensembl |
| 1 | Q99798 | NA | ACO2 | aconitase 2, mitochondrial | 50 | ENSG00000100412 |
| 2 | P40926 | NA | MDH2 | malate dehydrogenase 2, NAD (mitochondrial) | 4191 | ENSG00000146701 |
| 3 | P13804 | NA | ETFA | electron-transfer-flavoprotein, alpha polypeptide | 2108 | ENSG00000140374 |
| 4 | P48735 | NA | IDH2 | isocitrate dehydrogenase 2 (NADP+), mitochondrial | 3418 | ENSG00000182054 |
| 5 | P25705 | NA | ATP5A1 | ATP synthase, H+ transporting, mitochondrial F1 complex, alpha subunit 1, cardiac muscle | 498 | ENSG00000152234 |
| 6 | P31040 | NA | SDHA | succinate dehydrogenase complex, subunit A, flavoprotein (Fp) | 6389 | ENSG00000073578 |
| 7 | P48047 | NA | ATP5O | ATP synthase, H+ transporting, mitochondrial F1 complex, O subunit | 539 | ENSG00000241837 |
| 8 | P38117 | NA | ETFB | electron-transfer-flavoprotein, beta polypeptide | 2109 | ENSG00000105379 |

  
  

| **Database:Pathway Commons pathway      &nbspName:isoleucine degradation I      &nbspID:DB\_ID:1352** | | | | | | |
| --- | --- | --- | --- | --- | --- | --- |
| C=11; O=4; E=0.06; R=68.29; rawP=2.44e-07; adjP=4.01e-06 | | | | | | |
| Index | UserID | Value | Gene Symbol | Gene Name | EntrezGene | Ensembl |
| 1 | P24752 | NA | ACAT1 | acetyl-CoA acetyltransferase 1 | 38 | ENSG00000075239 |
| 2 | P40939 | NA | HADHA | hydroxyacyl-CoA dehydrogenase/3-ketoacyl-CoA thiolase/enoyl-CoA hydratase (trifunctional protein), alpha subunit | 3030 | ENSG00000084754 |
| 3 | P30084 | NA | ECHS1 | enoyl CoA hydratase, short chain, 1, mitochondrial | 1892 | ENSG00000127884 |
| 4 | Q99714 | NA | HSD17B10 | hydroxysteroid (17-beta) dehydrogenase 10 | 3028 | ENSG00000072506 |

  
  

| **Database:Pathway Commons pathway      &nbspName:Glucose metabolism      &nbspID:DB\_ID:898** | | | | | | |
| --- | --- | --- | --- | --- | --- | --- |
| C=36; O=5; E=0.19; R=26.08; rawP=1.29e-06; adjP=1.85e-05 | | | | | | |
| Index | UserID | Value | Gene Symbol | Gene Name | EntrezGene | Ensembl |
| 1 | Q02978 | NA | SLC25A11 | solute carrier family 25 (mitochondrial carrier; oxoglutarate carrier), member 11 | 8402 | ENSG00000108528 |
| 2 | P17174 | NA | GOT1 | glutamic-oxaloacetic transaminase 1, soluble (aspartate aminotransferase 1) | 2805 | ENSG00000120053 |
| 3 | P00558 | NA | PGK1 | phosphoglycerate kinase 1 | 5230 | ENSG00000102144 |
| 4 | P40926 | NA | MDH2 | malate dehydrogenase 2, NAD (mitochondrial) | 4191 | ENSG00000146701 |
| 5 | P60174 | NA | TPI1 | triosephosphate isomerase 1 | 7167 | ENSG00000111669 |

  
  

| **Database:Pathway Commons pathway      &nbspName:Branched-chain amino acid catabolism      &nbspID:DB\_ID:878** | | | | | | |
| --- | --- | --- | --- | --- | --- | --- |
| C=17; O=4; E=0.09; R=44.19; rawP=1.72e-06; adjP=2.20e-05 | | | | | | |
| Index | UserID | Value | Gene Symbol | Gene Name | EntrezGene | Ensembl |
| 1 | P24752 | NA | ACAT1 | acetyl-CoA acetyltransferase 1 | 38 | ENSG00000075239 |
| 2 | P21953 | NA | BCKDHB | branched chain keto acid dehydrogenase E1, beta polypeptide | 594 | ENSG00000083123 |
| 3 | P31937 | NA | HIBADH | 3-hydroxyisobutyrate dehydrogenase | 11112 | ENSG00000106049 |
| 4 | Q99714 | NA | HSD17B10 | hydroxysteroid (17-beta) dehydrogenase 10 | 3028 | ENSG00000072506 |

  
  

| **Database:Pathway Commons pathway      &nbspName:Citric acid cycle (TCA cycle)      &nbspID:DB\_ID:872** | | | | | | |
| --- | --- | --- | --- | --- | --- | --- |
| C=19; O=4; E=0.10; R=39.54; rawP=2.77e-06; adjP=3.19e-05 | | | | | | |
| Index | UserID | Value | Gene Symbol | Gene Name | EntrezGene | Ensembl |
| 1 | Q99798 | NA | ACO2 | aconitase 2, mitochondrial | 50 | ENSG00000100412 |
| 2 | P48735 | NA | IDH2 | isocitrate dehydrogenase 2 (NADP+), mitochondrial | 3418 | ENSG00000182054 |
| 3 | P31040 | NA | SDHA | succinate dehydrogenase complex, subunit A, flavoprotein (Fp) | 6389 | ENSG00000073578 |
| 4 | P40926 | NA | MDH2 | malate dehydrogenase 2, NAD (mitochondrial) | 4191 | ENSG00000146701 |

  
  

| **Database:Pathway Commons pathway      &nbspName:Translation      &nbspID:DB\_ID:722** | | | | | | |
| --- | --- | --- | --- | --- | --- | --- |
| C=118; O=7; E=0.63; R=11.14; rawP=3.24e-06; adjP=3.39e-05 | | | | | | |
| Index | UserID | Value | Gene Symbol | Gene Name | EntrezGene | Ensembl |
| 1 | P61313 | NA | RPL15 | ribosomal protein L15 | 6138 | ENSG00000174748 |
| 2 | P60842 | NA | EIF4A1 | eukaryotic translation initiation factor 4A1 | 1973 | ENSG00000161960 |
| 3 | P13639 | NA | EEF2 | eukaryotic translation elongation factor 2 | 1938 | ENSG00000167658 |
| 4 | P41091 | NA | EIF2S3 | eukaryotic translation initiation factor 2, subunit 3 gamma, 52kDa | 1968 | ENSG00000130741 |
| 5 | P68104 | NA | EEF1A1 | eukaryotic translation elongation factor 1 alpha 1 | 1915 | ENSG00000156508 |
| 6 | P29692 | NA | EEF1D | eukaryotic translation elongation factor 1 delta (guanine nucleotide exchange protein) | 1936 | ENSG00000104529 |
| 7 | P24534 | NA | EEF1B2 | eukaryotic translation elongation factor 1 beta 2 | 1933 | ENSG00000114942 |

  
  

| **Database:Pathway Commons pathway      &nbspName:Gene Expression      &nbspID:DB\_ID:531** | | | | | | |
| --- | --- | --- | --- | --- | --- | --- |
| C=379; O=11; E=2.02; R=5.45; rawP=5.73e-06; adjP=5.49e-05 | | | | | | |
| Index | UserID | Value | Gene Symbol | Gene Name | EntrezGene | Ensembl |
| 1 | P49589 | NA | CARS | cysteinyl-tRNA synthetase | 833 | ENSG00000110619 |
| 2 | P13639 | NA | EEF2 | eukaryotic translation elongation factor 2 | 1938 | ENSG00000167658 |
| 3 | P41091 | NA | EIF2S3 | eukaryotic translation initiation factor 2, subunit 3 gamma, 52kDa | 1968 | ENSG00000130741 |
| 4 | Q6PI48 | NA | DARS2 | aspartyl-tRNA synthetase 2, mitochondrial | 55157 | ENSG00000117593 |
| 5 | Q9NSE4 | NA | IARS2 | isoleucyl-tRNA synthetase 2, mitochondrial | 55699 | ENSG00000067704 |
| 6 | P61313 | NA | RPL15 | ribosomal protein L15 | 6138 | ENSG00000174748 |
| 7 | P67809 | NA | YBX1 | Y box binding protein 1 | 4904 | ENSG00000065978 |
| 8 | P60842 | NA | EIF4A1 | eukaryotic translation initiation factor 4A1 | 1973 | ENSG00000161960 |
| 9 | P68104 | NA | EEF1A1 | eukaryotic translation elongation factor 1 alpha 1 | 1915 | ENSG00000156508 |
| 10 | P29692 | NA | EEF1D | eukaryotic translation elongation factor 1 delta (guanine nucleotide exchange protein) | 1936 | ENSG00000104529 |
| 11 | P24534 | NA | EEF1B2 | eukaryotic translation elongation factor 1 beta 2 | 1933 | ENSG00000114942 |

  
  

| **Database:Pathway Commons pathway      &nbspName:Purine metabolism      &nbspID:DB\_ID:638** | | | | | | |
| --- | --- | --- | --- | --- | --- | --- |
| C=24; O=4; E=0.13; R=31.30; rawP=7.44e-06; adjP=6.58e-05 | | | | | | |
| Index | UserID | Value | Gene Symbol | Gene Name | EntrezGene | Ensembl |
| 1 | P04040 | NA | CAT | catalase | 847 | ENSG00000121691 |
| 2 | P07741 | NA | APRT | adenine phosphoribosyltransferase | 353 | ENSG00000198931 |
| 3 | P22102 | NA | GART | phosphoribosylglycinamide formyltransferase, phosphoribosylglycinamide synthetase, phosphoribosylaminoimidazole synthetase | 2618 | ENSG00000159131 |
| 4 | P00491 | NA | PNP | purine nucleoside phosphorylase | 4860 | ENSG00000198805 |

  
  

| **Database:Pathway Commons pathway      &nbspName:mitochondrial fatty acid beta-oxidation of saturated fatty acids      &nbspID:DB\_ID:911** | | | | | | |
| --- | --- | --- | --- | --- | --- | --- |
| C=8; O=3; E=0.04; R=70.43; rawP=8.06e-06; adjP=6.62e-05 | | | | | | |
| Index | UserID | Value | Gene Symbol | Gene Name | EntrezGene | Ensembl |
| 1 | P49748 | NA | ACADVL | acyl-CoA dehydrogenase, very long chain | 37 | ENSG00000072778 |
| 2 | P40939 | NA | HADHA | hydroxyacyl-CoA dehydrogenase/3-ketoacyl-CoA thiolase/enoyl-CoA hydratase (trifunctional protein), alpha subunit | 3030 | ENSG00000084754 |
| 3 | P30084 | NA | ECHS1 | enoyl CoA hydratase, short chain, 1, mitochondrial | 1892 | ENSG00000127884 |

  
  

| **Database:Pathway Commons pathway      &nbspName:Metabolism of carbohydrates      &nbspID:DB\_ID:905** | | | | | | |
| --- | --- | --- | --- | --- | --- | --- |
| C=92; O=6; E=0.49; R=12.25; rawP=9.75e-06; adjP=7.48e-05 | | | | | | |
| Index | UserID | Value | Gene Symbol | Gene Name | EntrezGene | Ensembl |
| 1 | Q02978 | NA | SLC25A11 | solute carrier family 25 (mitochondrial carrier; oxoglutarate carrier), member 11 | 8402 | ENSG00000108528 |
| 2 | P17174 | NA | GOT1 | glutamic-oxaloacetic transaminase 1, soluble (aspartate aminotransferase 1) | 2805 | ENSG00000120053 |
| 3 | P19367 | NA | HK1 | hexokinase 1 | 3098 | ENSG00000156515 |
| 4 | P00558 | NA | PGK1 | phosphoglycerate kinase 1 | 5230 | ENSG00000102144 |
| 5 | P40926 | NA | MDH2 | malate dehydrogenase 2, NAD (mitochondrial) | 4191 | ENSG00000146701 |
| 6 | P60174 | NA | TPI1 | triosephosphate isomerase 1 | 7167 | ENSG00000111669 |

  
  

| **Database:Pathway Commons pathway      &nbspName:TRAIL signaling pathway      &nbspID:DB\_ID:1480** | | | | | | |
| --- | --- | --- | --- | --- | --- | --- |
| C=1328; O=20; E=7.07; R=2.83; rawP=2.09e-05; adjP=0.0001 | | | | | | |
| Index | UserID | Value | Gene Symbol | Gene Name | EntrezGene | Ensembl |
| 1 | P08195 | NA | SLC3A2 | solute carrier family 3 (activators of dibasic and neutral amino acid transport), member 2 | 6520 | ENSG00000168003 |
| 2 | P05787 | NA | KRT8 | keratin 8 | 3856 | ENSG00000170421 |
| 3 | O95831 | NA | AIFM1 | apoptosis-inducing factor, mitochondrion-associated, 1 | 9131 | ENSG00000156709 |
| 4 | P09382 | NA | LGALS1 | lectin, galactoside-binding, soluble, 1 | 3956 | ENSG00000100097 |
| 5 | P04179 | NA | SOD2 | superoxide dismutase 2, mitochondrial | 6648 | ENSG00000112096 |
| 6 | P60842 | NA | EIF4A1 | eukaryotic translation initiation factor 4A1 | 1973 | ENSG00000161960 |
| 7 | P02786 | NA | TFRC | transferrin receptor (p90, CD71) | 7037 | ENSG00000072274 |
| 8 | Q14980 | NA | NUMA1 | nuclear mitotic apparatus protein 1 | 4926 | ENSG00000137497 |
| 9 | P04040 | NA | CAT | catalase | 847 | ENSG00000121691 |
| 10 | P13639 | NA | EEF2 | eukaryotic translation elongation factor 2 | 1938 | ENSG00000167658 |
| 11 | P30048 | NA | PRDX3 | peroxiredoxin 3 | 10935 | ENSG00000165672 |
| 12 | P00558 | NA | PGK1 | phosphoglycerate kinase 1 | 5230 | ENSG00000102144 |
| 13 | P05556 | NA | ITGB1 | integrin, beta 1 (fibronectin receptor, beta polypeptide, antigen CD29 includes MDF2, MSK12) | 3688 | ENSG00000150093 |
| 14 | O75367 | NA | H2AFY | H2A histone family, member Y | 9555 | ENSG00000113648 |
| 15 | P19367 | NA | HK1 | hexokinase 1 | 3098 | ENSG00000156515 |
| 16 | P08727 | NA | KRT19 | keratin 19 | 3880 | ENSG00000171345 |
| 17 | P05783 | NA | KRT18 | keratin 18 | 3875 | ENSG00000111057 |
| 18 | P18206 | NA | VCL | vinculin | 7414 | ENSG00000035403 |
| 19 | Q13813 | NA | SPTAN1 | spectrin, alpha, non-erythrocytic 1 | 6709 | ENSG00000197694 |
| 20 | P11387 | NA | TOP1 | topoisomerase (DNA) I | 7150 | ENSG00000198900 |

  
  

| **Database:Pathway Commons pathway      &nbspName:Metabolism of nucleotides      &nbspID:DB\_ID:639** | | | | | | |
| --- | --- | --- | --- | --- | --- | --- |
| C=58; O=5; E=0.31; R=16.19; rawP=1.43e-05; adjP=0.0001 | | | | | | |
| Index | UserID | Value | Gene Symbol | Gene Name | EntrezGene | Ensembl |
| 1 | P04040 | NA | CAT | catalase | 847 | ENSG00000121691 |
| 2 | P07741 | NA | APRT | adenine phosphoribosyltransferase | 353 | ENSG00000198931 |
| 3 | P22102 | NA | GART | phosphoribosylglycinamide formyltransferase, phosphoribosylglycinamide synthetase, phosphoribosylaminoimidazole synthetase | 2618 | ENSG00000159131 |
| 4 | P54819 | NA | AK2 | adenylate kinase 2 | 204 | ENSG00000004455 |
| 5 | P00491 | NA | PNP | purine nucleoside phosphorylase | 4860 | ENSG00000198805 |

  
  

| **Database:Pathway Commons pathway      &nbspName:Amino acid synthesis and interconversion (transamination)      &nbspID:DB\_ID:879** | | | | | | |
| --- | --- | --- | --- | --- | --- | --- |
| C=10; O=3; E=0.05; R=56.34; rawP=1.71e-05; adjP=0.0001 | | | | | | |
| Index | UserID | Value | Gene Symbol | Gene Name | EntrezGene | Ensembl |
| 1 | P17174 | NA | GOT1 | glutamic-oxaloacetic transaminase 1, soluble (aspartate aminotransferase 1) | 2805 | ENSG00000120053 |
| 2 | P00367 | NA | GLUD1 | glutamate dehydrogenase 1 | 2746 | ENSG00000148672 |
| 3 | P32322 | NA | PYCR1 | pyrroline-5-carboxylate reductase 1 | 5831 | ENSG00000183010 |

  
  

| **Database:Pathway Commons pathway      &nbspName:Pyruvate metabolism and Citric Acid (TCA) cycle      &nbspID:DB\_ID:546** | | | | | | |
| --- | --- | --- | --- | --- | --- | --- |
| C=31; O=4; E=0.17; R=24.23; rawP=2.14e-05; adjP=0.0001 | | | | | | |
| Index | UserID | Value | Gene Symbol | Gene Name | EntrezGene | Ensembl |
| 1 | Q99798 | NA | ACO2 | aconitase 2, mitochondrial | 50 | ENSG00000100412 |
| 2 | P48735 | NA | IDH2 | isocitrate dehydrogenase 2 (NADP+), mitochondrial | 3418 | ENSG00000182054 |
| 3 | P31040 | NA | SDHA | succinate dehydrogenase complex, subunit A, flavoprotein (Fp) | 6389 | ENSG00000073578 |
| 4 | P40926 | NA | MDH2 | malate dehydrogenase 2, NAD (mitochondrial) | 4191 | ENSG00000146701 |

  
  

| **Database:Pathway Commons pathway      &nbspName:valine degradation I      &nbspID:DB\_ID:1369** | | | | | | |
| --- | --- | --- | --- | --- | --- | --- |
| C=12; O=3; E=0.06; R=46.95; rawP=3.12e-05; adjP=0.0002 | | | | | | |
| Index | UserID | Value | Gene Symbol | Gene Name | EntrezGene | Ensembl |
| 1 | P40939 | NA | HADHA | hydroxyacyl-CoA dehydrogenase/3-ketoacyl-CoA thiolase/enoyl-CoA hydratase (trifunctional protein), alpha subunit | 3030 | ENSG00000084754 |
| 2 | P31937 | NA | HIBADH | 3-hydroxyisobutyrate dehydrogenase | 11112 | ENSG00000106049 |
| 3 | P30084 | NA | ECHS1 | enoyl CoA hydratase, short chain, 1, mitochondrial | 1892 | ENSG00000127884 |

  
  

| **Database:Pathway Commons pathway      &nbspName:Mitochondrial Fatty Acid Beta-Oxidation      &nbspID:DB\_ID:912** | | | | | | |
| --- | --- | --- | --- | --- | --- | --- |
| C=14; O=3; E=0.07; R=40.24; rawP=5.12e-05; adjP=0.0003 | | | | | | |
| Index | UserID | Value | Gene Symbol | Gene Name | EntrezGene | Ensembl |
| 1 | P49748 | NA | ACADVL | acyl-CoA dehydrogenase, very long chain | 37 | ENSG00000072778 |
| 2 | P40939 | NA | HADHA | hydroxyacyl-CoA dehydrogenase/3-ketoacyl-CoA thiolase/enoyl-CoA hydratase (trifunctional protein), alpha subunit | 3030 | ENSG00000084754 |
| 3 | P30084 | NA | ECHS1 | enoyl CoA hydratase, short chain, 1, mitochondrial | 1892 | ENSG00000127884 |

  
  

| **Database:Pathway Commons pathway      &nbspName:Eukaryotic Translation Elongation      &nbspID:DB\_ID:713** | | | | | | |
| --- | --- | --- | --- | --- | --- | --- |
| C=86; O=5; E=0.46; R=10.92; rawP=9.64e-05; adjP=0.0005 | | | | | | |
| Index | UserID | Value | Gene Symbol | Gene Name | EntrezGene | Ensembl |
| 1 | P61313 | NA | RPL15 | ribosomal protein L15 | 6138 | ENSG00000174748 |
| 2 | P13639 | NA | EEF2 | eukaryotic translation elongation factor 2 | 1938 | ENSG00000167658 |
| 3 | P68104 | NA | EEF1A1 | eukaryotic translation elongation factor 1 alpha 1 | 1915 | ENSG00000156508 |
| 4 | P29692 | NA | EEF1D | eukaryotic translation elongation factor 1 delta (guanine nucleotide exchange protein) | 1936 | ENSG00000104529 |
| 5 | P24534 | NA | EEF1B2 | eukaryotic translation elongation factor 1 beta 2 | 1933 | ENSG00000114942 |

  
  

| **Database:Pathway Commons pathway      &nbspName:Respiratory electron transport, ATP synthesis by chemiosmotic coupling, and heat production by uncoupling proteins.      &nbspID:DB\_ID:874** | | | | | | |
| --- | --- | --- | --- | --- | --- | --- |
| C=91; O=5; E=0.48; R=10.32; rawP=0.0001; adjP=0.0005 | | | | | | |
| Index | UserID | Value | Gene Symbol | Gene Name | EntrezGene | Ensembl |
| 1 | P25705 | NA | ATP5A1 | ATP synthase, H+ transporting, mitochondrial F1 complex, alpha subunit 1, cardiac muscle | 498 | ENSG00000152234 |
| 2 | P48047 | NA | ATP5O | ATP synthase, H+ transporting, mitochondrial F1 complex, O subunit | 539 | ENSG00000241837 |
| 3 | P31040 | NA | SDHA | succinate dehydrogenase complex, subunit A, flavoprotein (Fp) | 6389 | ENSG00000073578 |
| 4 | P38117 | NA | ETFB | electron-transfer-flavoprotein, beta polypeptide | 2109 | ENSG00000105379 |
| 5 | P13804 | NA | ETFA | electron-transfer-flavoprotein, alpha polypeptide | 2108 | ENSG00000140374 |

  
  

| **Database:Pathway Commons pathway      &nbspName:Caspase cascade in apoptosis      &nbspID:DB\_ID:1501** | | | | | | |
| --- | --- | --- | --- | --- | --- | --- |
| C=52; O=4; E=0.28; R=14.45; rawP=0.0002; adjP=0.0009 | | | | | | |
| Index | UserID | Value | Gene Symbol | Gene Name | EntrezGene | Ensembl |
| 1 | Q14980 | NA | NUMA1 | nuclear mitotic apparatus protein 1 | 4926 | ENSG00000137497 |
| 2 | P05783 | NA | KRT18 | keratin 18 | 3875 | ENSG00000111057 |
| 3 | P11387 | NA | TOP1 | topoisomerase (DNA) I | 7150 | ENSG00000198900 |
| 4 | Q13813 | NA | SPTAN1 | spectrin, alpha, non-erythrocytic 1 | 6709 | ENSG00000197694 |

  
  

| **Database:Pathway Commons pathway      &nbspName:TNF receptor signaling pathway      &nbspID:DB\_ID:1600** | | | | | | |
| --- | --- | --- | --- | --- | --- | --- |
| C=299; O=8; E=1.59; R=5.02; rawP=0.0002; adjP=0.0009 | | | | | | |
| Index | UserID | Value | Gene Symbol | Gene Name | EntrezGene | Ensembl |
| 1 | Q14980 | NA | NUMA1 | nuclear mitotic apparatus protein 1 | 4926 | ENSG00000137497 |
| 2 | P08195 | NA | SLC3A2 | solute carrier family 3 (activators of dibasic and neutral amino acid transport), member 2 | 6520 | ENSG00000168003 |
| 3 | P05787 | NA | KRT8 | keratin 8 | 3856 | ENSG00000170421 |
| 4 | O95831 | NA | AIFM1 | apoptosis-inducing factor, mitochondrion-associated, 1 | 9131 | ENSG00000156709 |
| 5 | P08727 | NA | KRT19 | keratin 19 | 3880 | ENSG00000171345 |
| 6 | P05783 | NA | KRT18 | keratin 18 | 3875 | ENSG00000111057 |
| 7 | Q13813 | NA | SPTAN1 | spectrin, alpha, non-erythrocytic 1 | 6709 | ENSG00000197694 |
| 8 | P11387 | NA | TOP1 | topoisomerase (DNA) I | 7150 | ENSG00000198900 |

  
  
  
  

---

WebGestalt is currently developed and maintained by Jing Wang and Bing Zhang at the  Zhang Lab. Other people who have made significant contribution to the project include Dexter Duncan, Stefan Kirov, Zhiao Shi, and Jay Snoddy.  
  
**Funding credits:** NIH/NIAAA (U01 AA016662, U01 AA013512); NIH/NIDA (P01 DA015027); NIH/NIMH (P50 MH078028, P50 MH096972); NIH/NCI (U24 CA159988); NIH/NIGMS (R01 GM088822).
