## Supplementary material for "PGRMC1 phosphorylation and cell plasticity 1: glycolysis, mitochondria, tumor growth": File S7: datasource.txt

The current version of WebGestalt was updated on 1/30/2013. It supports eight organisms including human, mouse, rat, worm, fly, yeast, dog, and zebrafish. Information in this version was collected from the following resources: 

ID mapping: NCBI Gene (10/26/2012), NCBI GEO (12/10/2012), Ensembl (version 68, 11/23/2012), MGI (10/21/2012), SGD (10/21/2012), Affymetrix (1/11/2013), Illumina (1/11/2013), IPI (11/21/2011), NCBI dbSNP (12/06/2012). 

Gene Ontology: Gene Ontology (version 1.2, 11/11/2012). 

Pathways: KEGG (03/21/2011), Pathway Commons (11/11/2012), WikiPathways (11/11/2012). 

Regulatory modules (Motif gene sets): MSigDB (11/11/2012). 

Protein-protein interaction modules: Protein-protein interaction data were downloaded from HPRD (11/11/2012), BioGrid(11/11/2012), BOND(11/11/2012), DIP(11/11/2012), IntAct(11/11/2012), MINT(11/11/2012), and Reactome(11/11/2012). Only interactions with publication support were retained. Protein-protein interaction modules at different hierarchical levels were identified by iterative partitioning of the integrated protein-protein interaction network using the Walktrap algorithm. 

Disease and drug associated genes: disease and drug terms were downloaded from PharmGKB (1/26/2013), genes associated with individual disease and drug terms were inferred using GLAD4U (1/26/2013). 

Cytogenetic band: NCBI Gene (10/26/2012). 

Phenotype: Mammalian Phenotype Ontology (04/10/2013) and Human Phenotype Ontology (04/10/2013).

PheWAS: Human PheWAS Ontology (05/19/2014)
