## Supplementary material for "PGRMC1 phosphorylation and cell plasticity 1: glycolysis, mitochondria, tumor growth": File S7: final_disease_geneset_file_1438486678.html

Anchored HTML File of EIDs

|  |  |
| --- | --- |
|  | WEB-based GEne SeT AnaLysis Toolkit |
|  | ***Translating gene lists into biological insights...*** |

---

  

| **Database:disease      &nbspName:Metabolic Diseases      &nbspID:DB\_ID:PA444938** | | | | | | |
| --- | --- | --- | --- | --- | --- | --- |
| C=574; O=12; E=1.16; R=10.34; rawP=8.79e-10; adjP=5.01e-08 | | | | | | |
| Index | UserID | Value | Gene Symbol | Gene Name | EntrezGene | Ensembl |
| 1 | O14773 | NA | TPP1 | tripeptidyl peptidase I | 1200 | ENSG00000166340 |
| 2 | P02792 | NA | FTL | ferritin, light polypeptide | 2512 | ENSG00000087086 |
| 3 | P10253 | NA | GAA | glucosidase, alpha; acid | 2548 | ENSG00000171298 |
| 4 | P51688 | NA | SGSH | N-sulfoglucosamine sulfohydrolase | 6448 | ENSG00000181523 |
| 5 | P61769 | NA | B2M | beta-2-microglobulin | 567 | ENSG00000166710 |
| 6 | P21980 | NA | TGM2 | transglutaminase 2 (C polypeptide, protein-glutamine-gamma-glutamyltransferase) | 7052 | ENSG00000198959 |
| 7 | Q16822 | NA | PCK2 | phosphoenolpyruvate carboxykinase 2 (mitochondrial) | 5106 | ENSG00000100889 |
| 8 | P49748 | NA | ACADVL | acyl-CoA dehydrogenase, very long chain | 37 | ENSG00000072778 |
| 9 | P51648 | NA | ALDH3A2 | aldehyde dehydrogenase 3 family, member A2 | 224 | ENSG00000072210 |
| 10 | P02545 | NA | LMNA | lamin A/C | 4000 | ENSG00000160789 |
| 11 | P02786 | NA | TFRC | transferrin receptor (p90, CD71) | 7037 | ENSG00000072274 |
| 12 | P43490 | NA | NAMPT | nicotinamide phosphoribosyltransferase | 10135 | ENSG00000105835 |

  
  

| **Database:disease      &nbspName:Metabolism, Inborn Errors      &nbspID:DB\_ID:PA444939** | | | | | | |
| --- | --- | --- | --- | --- | --- | --- |
| C=335; O=8; E=0.68; R=11.81; rawP=3.03e-07; adjP=8.64e-06 | | | | | | |
| Index | UserID | Value | Gene Symbol | Gene Name | EntrezGene | Ensembl |
| 1 | O14773 | NA | TPP1 | tripeptidyl peptidase I | 1200 | ENSG00000166340 |
| 2 | P10253 | NA | GAA | glucosidase, alpha; acid | 2548 | ENSG00000171298 |
| 3 | P51688 | NA | SGSH | N-sulfoglucosamine sulfohydrolase | 6448 | ENSG00000181523 |
| 4 | P51659 | NA | HSD17B4 | hydroxysteroid (17-beta) dehydrogenase 4 | 3295 | ENSG00000133835 |
| 5 | P49748 | NA | ACADVL | acyl-CoA dehydrogenase, very long chain | 37 | ENSG00000072778 |
| 6 | P51648 | NA | ALDH3A2 | aldehyde dehydrogenase 3 family, member A2 | 224 | ENSG00000072210 |
| 7 | P02786 | NA | TFRC | transferrin receptor (p90, CD71) | 7037 | ENSG00000072274 |
| 8 | P02545 | NA | LMNA | lamin A/C | 4000 | ENSG00000160789 |

  
  

| **Database:disease      &nbspName:Iron Overload      &nbspID:DB\_ID:PA446812** | | | | | | |
| --- | --- | --- | --- | --- | --- | --- |
| C=37; O=4; E=0.07; R=53.48; rawP=9.04e-07; adjP=1.72e-05 | | | | | | |
| Index | UserID | Value | Gene Symbol | Gene Name | EntrezGene | Ensembl |
| 1 | P02792 | NA | FTL | ferritin, light polypeptide | 2512 | ENSG00000087086 |
| 2 | P61769 | NA | B2M | beta-2-microglobulin | 567 | ENSG00000166710 |
| 3 | P02786 | NA | TFRC | transferrin receptor (p90, CD71) | 7037 | ENSG00000072274 |
| 4 | P04179 | NA | SOD2 | superoxide dismutase 2, mitochondrial | 6648 | ENSG00000112096 |

  
  

| **Database:disease      &nbspName:Hemochromatosis      &nbspID:DB\_ID:PA444405** | | | | | | |
| --- | --- | --- | --- | --- | --- | --- |
| C=29; O=3; E=0.06; R=51.18; rawP=2.70e-05; adjP=0.0004 | | | | | | |
| Index | UserID | Value | Gene Symbol | Gene Name | EntrezGene | Ensembl |
| 1 | P02792 | NA | FTL | ferritin, light polypeptide | 2512 | ENSG00000087086 |
| 2 | P61769 | NA | B2M | beta-2-microglobulin | 567 | ENSG00000166710 |
| 3 | P02786 | NA | TFRC | transferrin receptor (p90, CD71) | 7037 | ENSG00000072274 |

  
  
  
  

---

WebGestalt is currently developed and maintained by Jing Wang and Bing Zhang at the  Zhang Lab. Other people who have made significant contribution to the project include Dexter Duncan, Stefan Kirov, Zhiao Shi, and Jay Snoddy.  
  
**Funding credits:** NIH/NIAAA (U01 AA016662, U01 AA013512); NIH/NIDA (P01 DA015027); NIH/NIMH (P50 MH078028, P50 MH096972); NIH/NCI (U24 CA159988); NIH/NIGMS (R01 GM088822).
