## Supplementary material for "PGRMC1 phosphorylation and cell plasticity 1: glycolysis, mitochondria, tumor growth": File S7: final_sig_file_1438486678.html

Anchored HTML File of EIDs

|  |  |
| --- | --- |
|  | WEB-based GEne SeT AnaLysis Toolkit |
|  | ***Translating gene lists into biological insights...*** |

---

  

| **Database:molecular function      &nbspName:oxidoreductase activity      &nbspID:GO:0016491** | | | | | | |
| --- | --- | --- | --- | --- | --- | --- |
| C=697; O=12; E=1.86; R=6.45; rawP=1.58e-07; adjP=7.43e-06 | | | | | | |
| Index | UserID | Value | Gene Symbol | Gene Name | EntrezGene | Ensembl |
| 1 | P02792 | NA | FTL | ferritin, light polypeptide | 2512 | ENSG00000087086 |
| 2 | Q9Y2Q3 | NA | GSTK1 | glutathione S-transferase kappa 1 | 373156 | ENSG00000197448 |
| 3 | P22570 | NA | FDXR | ferredoxin reductase | 2232 | ENSG00000161513 |
| 4 | P04179 | NA | SOD2 | superoxide dismutase 2, mitochondrial | 6648 | ENSG00000112096 |
| 5 | P48735 | NA | IDH2 | isocitrate dehydrogenase 2 (NADP+), mitochondrial | 3418 | ENSG00000182054 |
| 6 | P49419 | NA | ALDH7A1 | aldehyde dehydrogenase 7 family, member A1 | 501 | ENSG00000164904 |
| 7 | P51659 | NA | HSD17B4 | hydroxysteroid (17-beta) dehydrogenase 4 | 3295 | ENSG00000133835 |
| 8 | P49748 | NA | ACADVL | acyl-CoA dehydrogenase, very long chain | 37 | ENSG00000072778 |
| 9 | P00352 | NA | ALDH1A1 | aldehyde dehydrogenase 1 family, member A1 | 216 | ENSG00000165092 |
| 10 | P51648 | NA | ALDH3A2 | aldehyde dehydrogenase 3 family, member A2 | 224 | ENSG00000072210 |
| 11 | P31937 | NA | HIBADH | 3-hydroxyisobutyrate dehydrogenase | 11112 | ENSG00000106049 |
| 12 | P32322 | NA | PYCR1 | pyrroline-5-carboxylate reductase 1 | 5831 | ENSG00000183010 |

  
  

| **Database:molecular function      &nbspName:catalytic activity      &nbspID:GO:0003824** | | | | | | |
| --- | --- | --- | --- | --- | --- | --- |
| C=5299; O=29; E=14.15; R=2.05; rawP=1.87e-06; adjP=3.27e-05 | | | | | | |
| Index | UserID | Value | Gene Symbol | Gene Name | EntrezGene | Ensembl |
| 1 | P02792 | NA | FTL | ferritin, light polypeptide | 2512 | ENSG00000087086 |
| 2 | P10253 | NA | GAA | glucosidase, alpha; acid | 2548 | ENSG00000171298 |
| 3 | Q7Z2K6 | NA | ERMP1 | endoplasmic reticulum metallopeptidase 1 | 79956 | ENSG00000099219 |
| 4 | Q13724 | NA | MOGS | mannosyl-oligosaccharide glucosidase | 7841 | ENSG00000115275 |
| 5 | P04179 | NA | SOD2 | superoxide dismutase 2, mitochondrial | 6648 | ENSG00000112096 |
| 6 | Q8NFV4 | NA | ABHD11 | abhydrolase domain containing 11 | 83451 | ENSG00000106077 |
| 7 | P48735 | NA | IDH2 | isocitrate dehydrogenase 2 (NADP+), mitochondrial | 3418 | ENSG00000182054 |
| 8 | P51659 | NA | HSD17B4 | hydroxysteroid (17-beta) dehydrogenase 4 | 3295 | ENSG00000133835 |
| 9 | P49588 | NA | AARS | alanyl-tRNA synthetase | 16 | ENSG00000090861 |
| 10 | P02786 | NA | TFRC | transferrin receptor (p90, CD71) | 7037 | ENSG00000072274 |
| 11 | P32322 | NA | PYCR1 | pyrroline-5-carboxylate reductase 1 | 5831 | ENSG00000183010 |
| 12 | P43490 | NA | NAMPT | nicotinamide phosphoribosyltransferase | 10135 | ENSG00000105835 |
| 13 | P17655 | NA | CAPN2 | calpain 2, (m/II) large subunit | 824 | ENSG00000162909 |
| 14 | O14773 | NA | TPP1 | tripeptidyl peptidase I | 1200 | ENSG00000166340 |
| 15 | Q9Y2Q3 | NA | GSTK1 | glutathione S-transferase kappa 1 | 373156 | ENSG00000197448 |
| 16 | P51688 | NA | SGSH | N-sulfoglucosamine sulfohydrolase | 6448 | ENSG00000181523 |
| 17 | P13639 | NA | EEF2 | eukaryotic translation elongation factor 2 | 1938 | ENSG00000167658 |
| 18 | Q16555 | NA | DPYSL2 | dihydropyrimidinase-like 2 | 1808 | ENSG00000092964 |
| 19 | P22570 | NA | FDXR | ferredoxin reductase | 2232 | ENSG00000161513 |
| 20 | O75521 | NA | ECI2 | enoyl-CoA delta isomerase 2 | 10455 | ENSG00000198721 |
| 21 | P49419 | NA | ALDH7A1 | aldehyde dehydrogenase 7 family, member A1 | 501 | ENSG00000164904 |
| 22 | P17096 | NA | HMGA1 | high mobility group AT-hook 1 | 3159 | ENSG00000137309 |
| 23 | P21980 | NA | TGM2 | transglutaminase 2 (C polypeptide, protein-glutamine-gamma-glutamyltransferase) | 7052 | ENSG00000198959 |
| 24 | Q16822 | NA | PCK2 | phosphoenolpyruvate carboxykinase 2 (mitochondrial) | 5106 | ENSG00000100889 |
| 25 | P49748 | NA | ACADVL | acyl-CoA dehydrogenase, very long chain | 37 | ENSG00000072778 |
| 26 | P00352 | NA | ALDH1A1 | aldehyde dehydrogenase 1 family, member A1 | 216 | ENSG00000165092 |
| 27 | P51648 | NA | ALDH3A2 | aldehyde dehydrogenase 3 family, member A2 | 224 | ENSG00000072210 |
| 28 | P31937 | NA | HIBADH | 3-hydroxyisobutyrate dehydrogenase | 11112 | ENSG00000106049 |
| 29 | P21589 | NA | NT5E | 5'-nucleotidase, ecto (CD73) | 4907 | ENSG00000135318 |

  
  

| **Database:molecular function      &nbspName:aldehyde dehydrogenase (NAD) activity      &nbspID:GO:0004029** | | | | | | |
| --- | --- | --- | --- | --- | --- | --- |
| C=10; O=3; E=0.03; R=112.33; rawP=2.09e-06; adjP=3.27e-05 | | | | | | |
| Index | UserID | Value | Gene Symbol | Gene Name | EntrezGene | Ensembl |
| 1 | P49419 | NA | ALDH7A1 | aldehyde dehydrogenase 7 family, member A1 | 501 | ENSG00000164904 |
| 2 | P00352 | NA | ALDH1A1 | aldehyde dehydrogenase 1 family, member A1 | 216 | ENSG00000165092 |
| 3 | P51648 | NA | ALDH3A2 | aldehyde dehydrogenase 3 family, member A2 | 224 | ENSG00000072210 |

  
  

| **Database:molecular function      &nbspName:oxidoreductase activity, acting on the aldehyde or oxo group of donors, NAD or NADP as acceptor      &nbspID:GO:0016620** | | | | | | |
| --- | --- | --- | --- | --- | --- | --- |
| C=29; O=3; E=0.08; R=38.73; rawP=6.15e-05; adjP=0.0007 | | | | | | |
| Index | UserID | Value | Gene Symbol | Gene Name | EntrezGene | Ensembl |
| 1 | P49419 | NA | ALDH7A1 | aldehyde dehydrogenase 7 family, member A1 | 501 | ENSG00000164904 |
| 2 | P00352 | NA | ALDH1A1 | aldehyde dehydrogenase 1 family, member A1 | 216 | ENSG00000165092 |
| 3 | P51648 | NA | ALDH3A2 | aldehyde dehydrogenase 3 family, member A2 | 224 | ENSG00000072210 |

  
  

| **Database:cellular component      &nbspName:cytoplasm      &nbspID:GO:0005737** | | | | | | |
| --- | --- | --- | --- | --- | --- | --- |
| C=9051; O=41; E=22.24; R=1.84; rawP=1.24e-11; adjP=8.93e-10 | | | | | | |
| Index | UserID | Value | Gene Symbol | Gene Name | EntrezGene | Ensembl |
| 1 | P10253 | NA | GAA | glucosidase, alpha; acid | 2548 | ENSG00000171298 |
| 2 | P55060 | NA | CSE1L | CSE1 chromosome segregation 1-like (yeast) | 1434 | ENSG00000124207 |
| 3 | Q13724 | NA | MOGS | mannosyl-oligosaccharide glucosidase | 7841 | ENSG00000115275 |
| 4 | P04179 | NA | SOD2 | superoxide dismutase 2, mitochondrial | 6648 | ENSG00000112096 |
| 5 | P61769 | NA | B2M | beta-2-microglobulin | 567 | ENSG00000166710 |
| 6 | P48735 | NA | IDH2 | isocitrate dehydrogenase 2 (NADP+), mitochondrial | 3418 | ENSG00000182054 |
| 7 | P49588 | NA | AARS | alanyl-tRNA synthetase | 16 | ENSG00000090861 |
| 8 | Q86UP2 | NA | KTN1 | kinectin 1 (kinesin receptor) | 3895 | ENSG00000126777 |
| 9 | P02786 | NA | TFRC | transferrin receptor (p90, CD71) | 7037 | ENSG00000072274 |
| 10 | P02545 | NA | LMNA | lamin A/C | 4000 | ENSG00000160789 |
| 11 | P43490 | NA | NAMPT | nicotinamide phosphoribosyltransferase | 10135 | ENSG00000105835 |
| 12 | P17655 | NA | CAPN2 | calpain 2, (m/II) large subunit | 824 | ENSG00000162909 |
| 13 | O14773 | NA | TPP1 | tripeptidyl peptidase I | 1200 | ENSG00000166340 |
| 14 | Q9Y2Q3 | NA | GSTK1 | glutathione S-transferase kappa 1 | 373156 | ENSG00000197448 |
| 15 | P13639 | NA | EEF2 | eukaryotic translation elongation factor 2 | 1938 | ENSG00000167658 |
| 16 | Q16555 | NA | DPYSL2 | dihydropyrimidinase-like 2 | 1808 | ENSG00000092964 |
| 17 | P22570 | NA | FDXR | ferredoxin reductase | 2232 | ENSG00000161513 |
| 18 | O75521 | NA | ECI2 | enoyl-CoA delta isomerase 2 | 10455 | ENSG00000198721 |
| 19 | P49419 | NA | ALDH7A1 | aldehyde dehydrogenase 7 family, member A1 | 501 | ENSG00000164904 |
| 20 | Q04837 | NA | SSBP1 | single-stranded DNA binding protein 1, mitochondrial | 6742 | ENSG00000106028 |
| 21 | P21980 | NA | TGM2 | transglutaminase 2 (C polypeptide, protein-glutamine-gamma-glutamyltransferase) | 7052 | ENSG00000198959 |
| 22 | Q16822 | NA | PCK2 | phosphoenolpyruvate carboxykinase 2 (mitochondrial) | 5106 | ENSG00000100889 |
| 23 | Q9Y3E0 | NA | GOLT1B | golgi transport 1B | 51026 | ENSG00000111711 |
| 24 | P51648 | NA | ALDH3A2 | aldehyde dehydrogenase 3 family, member A2 | 224 | ENSG00000072210 |
| 25 | P21589 | NA | NT5E | 5'-nucleotidase, ecto (CD73) | 4907 | ENSG00000135318 |
| 26 | P02792 | NA | FTL | ferritin, light polypeptide | 2512 | ENSG00000087086 |
| 27 | Q7Z2K6 | NA | ERMP1 | endoplasmic reticulum metallopeptidase 1 | 79956 | ENSG00000099219 |
| 28 | P26447 | NA | S100A4 | S100 calcium binding protein A4 | 6275 | ENSG00000196154 |
| 29 | Q8NFV4 | NA | ABHD11 | abhydrolase domain containing 11 | 83451 | ENSG00000106077 |
| 30 | P51659 | NA | HSD17B4 | hydroxysteroid (17-beta) dehydrogenase 4 | 3295 | ENSG00000133835 |
| 31 | P21796 | NA | VDAC1 | voltage-dependent anion channel 1 | 7416 | ENSG00000213585 |
| 32 | P32322 | NA | PYCR1 | pyrroline-5-carboxylate reductase 1 | 5831 | ENSG00000183010 |
| 33 | P51688 | NA | SGSH | N-sulfoglucosamine sulfohydrolase | 6448 | ENSG00000181523 |
| 34 | P17987 | NA | TCP1 | t-complex 1 | 6950 | ENSG00000120438 |
| 35 | P05556 | NA | ITGB1 | integrin, beta 1 (fibronectin receptor, beta polypeptide, antigen CD29 includes MDF2, MSK12) | 3688 | ENSG00000150093 |
| 36 | P17096 | NA | HMGA1 | high mobility group AT-hook 1 | 3159 | ENSG00000137309 |
| 37 | P49748 | NA | ACADVL | acyl-CoA dehydrogenase, very long chain | 37 | ENSG00000072778 |
| 38 | O14979 | NA | HNRPDL | heterogeneous nuclear ribonucleoprotein D-like | 9987 | ENSG00000152795 |
| 39 | P00352 | NA | ALDH1A1 | aldehyde dehydrogenase 1 family, member A1 | 216 | ENSG00000165092 |
| 40 | P31937 | NA | HIBADH | 3-hydroxyisobutyrate dehydrogenase | 11112 | ENSG00000106049 |
| 41 | P30042 | NA | C21orf33 | chromosome 21 open reading frame 33 | 8209 | ENSG00000160221 |

  
  

| **Database:cellular component      &nbspName:cytoplasmic part      &nbspID:GO:0044444** | | | | | | |
| --- | --- | --- | --- | --- | --- | --- |
| C=6728; O=37; E=16.54; R=2.24; rawP=3.36e-11; adjP=1.21e-09 | | | | | | |
| Index | UserID | Value | Gene Symbol | Gene Name | EntrezGene | Ensembl |
| 1 | P10253 | NA | GAA | glucosidase, alpha; acid | 2548 | ENSG00000171298 |
| 2 | Q13724 | NA | MOGS | mannosyl-oligosaccharide glucosidase | 7841 | ENSG00000115275 |
| 3 | P04179 | NA | SOD2 | superoxide dismutase 2, mitochondrial | 6648 | ENSG00000112096 |
| 4 | P61769 | NA | B2M | beta-2-microglobulin | 567 | ENSG00000166710 |
| 5 | P48735 | NA | IDH2 | isocitrate dehydrogenase 2 (NADP+), mitochondrial | 3418 | ENSG00000182054 |
| 6 | P49588 | NA | AARS | alanyl-tRNA synthetase | 16 | ENSG00000090861 |
| 7 | Q86UP2 | NA | KTN1 | kinectin 1 (kinesin receptor) | 3895 | ENSG00000126777 |
| 8 | P02545 | NA | LMNA | lamin A/C | 4000 | ENSG00000160789 |
| 9 | P02786 | NA | TFRC | transferrin receptor (p90, CD71) | 7037 | ENSG00000072274 |
| 10 | P43490 | NA | NAMPT | nicotinamide phosphoribosyltransferase | 10135 | ENSG00000105835 |
| 11 | O14773 | NA | TPP1 | tripeptidyl peptidase I | 1200 | ENSG00000166340 |
| 12 | Q9Y2Q3 | NA | GSTK1 | glutathione S-transferase kappa 1 | 373156 | ENSG00000197448 |
| 13 | P13639 | NA | EEF2 | eukaryotic translation elongation factor 2 | 1938 | ENSG00000167658 |
| 14 | Q16555 | NA | DPYSL2 | dihydropyrimidinase-like 2 | 1808 | ENSG00000092964 |
| 15 | P22570 | NA | FDXR | ferredoxin reductase | 2232 | ENSG00000161513 |
| 16 | O75521 | NA | ECI2 | enoyl-CoA delta isomerase 2 | 10455 | ENSG00000198721 |
| 17 | P49419 | NA | ALDH7A1 | aldehyde dehydrogenase 7 family, member A1 | 501 | ENSG00000164904 |
| 18 | Q04837 | NA | SSBP1 | single-stranded DNA binding protein 1, mitochondrial | 6742 | ENSG00000106028 |
| 19 | P21980 | NA | TGM2 | transglutaminase 2 (C polypeptide, protein-glutamine-gamma-glutamyltransferase) | 7052 | ENSG00000198959 |
| 20 | Q16822 | NA | PCK2 | phosphoenolpyruvate carboxykinase 2 (mitochondrial) | 5106 | ENSG00000100889 |
| 21 | Q9Y3E0 | NA | GOLT1B | golgi transport 1B | 51026 | ENSG00000111711 |
| 22 | P51648 | NA | ALDH3A2 | aldehyde dehydrogenase 3 family, member A2 | 224 | ENSG00000072210 |
| 23 | P02792 | NA | FTL | ferritin, light polypeptide | 2512 | ENSG00000087086 |
| 24 | Q7Z2K6 | NA | ERMP1 | endoplasmic reticulum metallopeptidase 1 | 79956 | ENSG00000099219 |
| 25 | P26447 | NA | S100A4 | S100 calcium binding protein A4 | 6275 | ENSG00000196154 |
| 26 | Q8NFV4 | NA | ABHD11 | abhydrolase domain containing 11 | 83451 | ENSG00000106077 |
| 27 | P51659 | NA | HSD17B4 | hydroxysteroid (17-beta) dehydrogenase 4 | 3295 | ENSG00000133835 |
| 28 | P21796 | NA | VDAC1 | voltage-dependent anion channel 1 | 7416 | ENSG00000213585 |
| 29 | P32322 | NA | PYCR1 | pyrroline-5-carboxylate reductase 1 | 5831 | ENSG00000183010 |
| 30 | P17987 | NA | TCP1 | t-complex 1 | 6950 | ENSG00000120438 |
| 31 | P51688 | NA | SGSH | N-sulfoglucosamine sulfohydrolase | 6448 | ENSG00000181523 |
| 32 | P05556 | NA | ITGB1 | integrin, beta 1 (fibronectin receptor, beta polypeptide, antigen CD29 includes MDF2, MSK12) | 3688 | ENSG00000150093 |
| 33 | P17096 | NA | HMGA1 | high mobility group AT-hook 1 | 3159 | ENSG00000137309 |
| 34 | P49748 | NA | ACADVL | acyl-CoA dehydrogenase, very long chain | 37 | ENSG00000072778 |
| 35 | P00352 | NA | ALDH1A1 | aldehyde dehydrogenase 1 family, member A1 | 216 | ENSG00000165092 |
| 36 | P31937 | NA | HIBADH | 3-hydroxyisobutyrate dehydrogenase | 11112 | ENSG00000106049 |
| 37 | P30042 | NA | C21orf33 | chromosome 21 open reading frame 33 | 8209 | ENSG00000160221 |

  
  

| **Database:cellular component      &nbspName:mitochondrial matrix      &nbspID:GO:0005759** | | | | | | |
| --- | --- | --- | --- | --- | --- | --- |
| C=278; O=10; E=0.68; R=14.64; rawP=9.99e-10; adjP=2.40e-08 | | | | | | |
| Index | UserID | Value | Gene Symbol | Gene Name | EntrezGene | Ensembl |
| 1 | P22570 | NA | FDXR | ferredoxin reductase | 2232 | ENSG00000161513 |
| 2 | P04179 | NA | SOD2 | superoxide dismutase 2, mitochondrial | 6648 | ENSG00000112096 |
| 3 | P48735 | NA | IDH2 | isocitrate dehydrogenase 2 (NADP+), mitochondrial | 3418 | ENSG00000182054 |
| 4 | P49419 | NA | ALDH7A1 | aldehyde dehydrogenase 7 family, member A1 | 501 | ENSG00000164904 |
| 5 | Q04837 | NA | SSBP1 | single-stranded DNA binding protein 1, mitochondrial | 6742 | ENSG00000106028 |
| 6 | Q16822 | NA | PCK2 | phosphoenolpyruvate carboxykinase 2 (mitochondrial) | 5106 | ENSG00000100889 |
| 7 | P21796 | NA | VDAC1 | voltage-dependent anion channel 1 | 7416 | ENSG00000213585 |
| 8 | P49748 | NA | ACADVL | acyl-CoA dehydrogenase, very long chain | 37 | ENSG00000072778 |
| 9 | P31937 | NA | HIBADH | 3-hydroxyisobutyrate dehydrogenase | 11112 | ENSG00000106049 |
| 10 | P32322 | NA | PYCR1 | pyrroline-5-carboxylate reductase 1 | 5831 | ENSG00000183010 |

  
  

| **Database:cellular component      &nbspName:mitochondrion      &nbspID:GO:0005739** | | | | | | |
| --- | --- | --- | --- | --- | --- | --- |
| C=1515; O=18; E=3.72; R=4.83; rawP=4.23e-09; adjP=7.61e-08 | | | | | | |
| Index | UserID | Value | Gene Symbol | Gene Name | EntrezGene | Ensembl |
| 1 | P04179 | NA | SOD2 | superoxide dismutase 2, mitochondrial | 6648 | ENSG00000112096 |
| 2 | Q8NFV4 | NA | ABHD11 | abhydrolase domain containing 11 | 83451 | ENSG00000106077 |
| 3 | P48735 | NA | IDH2 | isocitrate dehydrogenase 2 (NADP+), mitochondrial | 3418 | ENSG00000182054 |
| 4 | P21796 | NA | VDAC1 | voltage-dependent anion channel 1 | 7416 | ENSG00000213585 |
| 5 | P02786 | NA | TFRC | transferrin receptor (p90, CD71) | 7037 | ENSG00000072274 |
| 6 | P32322 | NA | PYCR1 | pyrroline-5-carboxylate reductase 1 | 5831 | ENSG00000183010 |
| 7 | O14773 | NA | TPP1 | tripeptidyl peptidase I | 1200 | ENSG00000166340 |
| 8 | Q9Y2Q3 | NA | GSTK1 | glutathione S-transferase kappa 1 | 373156 | ENSG00000197448 |
| 9 | Q16555 | NA | DPYSL2 | dihydropyrimidinase-like 2 | 1808 | ENSG00000092964 |
| 10 | P22570 | NA | FDXR | ferredoxin reductase | 2232 | ENSG00000161513 |
| 11 | O75521 | NA | ECI2 | enoyl-CoA delta isomerase 2 | 10455 | ENSG00000198721 |
| 12 | P49419 | NA | ALDH7A1 | aldehyde dehydrogenase 7 family, member A1 | 501 | ENSG00000164904 |
| 13 | P21980 | NA | TGM2 | transglutaminase 2 (C polypeptide, protein-glutamine-gamma-glutamyltransferase) | 7052 | ENSG00000198959 |
| 14 | Q04837 | NA | SSBP1 | single-stranded DNA binding protein 1, mitochondrial | 6742 | ENSG00000106028 |
| 15 | Q16822 | NA | PCK2 | phosphoenolpyruvate carboxykinase 2 (mitochondrial) | 5106 | ENSG00000100889 |
| 16 | P49748 | NA | ACADVL | acyl-CoA dehydrogenase, very long chain | 37 | ENSG00000072778 |
| 17 | P31937 | NA | HIBADH | 3-hydroxyisobutyrate dehydrogenase | 11112 | ENSG00000106049 |
| 18 | P30042 | NA | C21orf33 | chromosome 21 open reading frame 33 | 8209 | ENSG00000160221 |

  
  

| **Database:cellular component      &nbspName:mitochondrial part      &nbspID:GO:0044429** | | | | | | |
| --- | --- | --- | --- | --- | --- | --- |
| C=740; O=11; E=1.82; R=6.05; rawP=1.13e-06; adjP=1.63e-05 | | | | | | |
| Index | UserID | Value | Gene Symbol | Gene Name | EntrezGene | Ensembl |
| 1 | Q9Y2Q3 | NA | GSTK1 | glutathione S-transferase kappa 1 | 373156 | ENSG00000197448 |
| 2 | P22570 | NA | FDXR | ferredoxin reductase | 2232 | ENSG00000161513 |
| 3 | P04179 | NA | SOD2 | superoxide dismutase 2, mitochondrial | 6648 | ENSG00000112096 |
| 4 | P48735 | NA | IDH2 | isocitrate dehydrogenase 2 (NADP+), mitochondrial | 3418 | ENSG00000182054 |
| 5 | P49419 | NA | ALDH7A1 | aldehyde dehydrogenase 7 family, member A1 | 501 | ENSG00000164904 |
| 6 | Q04837 | NA | SSBP1 | single-stranded DNA binding protein 1, mitochondrial | 6742 | ENSG00000106028 |
| 7 | Q16822 | NA | PCK2 | phosphoenolpyruvate carboxykinase 2 (mitochondrial) | 5106 | ENSG00000100889 |
| 8 | P21796 | NA | VDAC1 | voltage-dependent anion channel 1 | 7416 | ENSG00000213585 |
| 9 | P49748 | NA | ACADVL | acyl-CoA dehydrogenase, very long chain | 37 | ENSG00000072778 |
| 10 | P31937 | NA | HIBADH | 3-hydroxyisobutyrate dehydrogenase | 11112 | ENSG00000106049 |
| 11 | P32322 | NA | PYCR1 | pyrroline-5-carboxylate reductase 1 | 5831 | ENSG00000183010 |

  
  

| **Database:cellular component      &nbspName:mitochondrial nucleoid      &nbspID:GO:0042645** | | | | | | |
| --- | --- | --- | --- | --- | --- | --- |
| C=37; O=4; E=0.09; R=43.99; rawP=1.96e-06; adjP=2.02e-05 | | | | | | |
| Index | UserID | Value | Gene Symbol | Gene Name | EntrezGene | Ensembl |
| 1 | Q04837 | NA | SSBP1 | single-stranded DNA binding protein 1, mitochondrial | 6742 | ENSG00000106028 |
| 2 | P21796 | NA | VDAC1 | voltage-dependent anion channel 1 | 7416 | ENSG00000213585 |
| 3 | P49748 | NA | ACADVL | acyl-CoA dehydrogenase, very long chain | 37 | ENSG00000072778 |
| 4 | P04179 | NA | SOD2 | superoxide dismutase 2, mitochondrial | 6648 | ENSG00000112096 |

  
  

| **Database:cellular component      &nbspName:intracellular part      &nbspID:GO:0044424** | | | | | | |
| --- | --- | --- | --- | --- | --- | --- |
| C=12096; O=41; E=29.73; R=1.38; rawP=1.85e-06; adjP=2.02e-05 | | | | | | |
| Index | UserID | Value | Gene Symbol | Gene Name | EntrezGene | Ensembl |
| 1 | P10253 | NA | GAA | glucosidase, alpha; acid | 2548 | ENSG00000171298 |
| 2 | P55060 | NA | CSE1L | CSE1 chromosome segregation 1-like (yeast) | 1434 | ENSG00000124207 |
| 3 | Q13724 | NA | MOGS | mannosyl-oligosaccharide glucosidase | 7841 | ENSG00000115275 |
| 4 | P04179 | NA | SOD2 | superoxide dismutase 2, mitochondrial | 6648 | ENSG00000112096 |
| 5 | P61769 | NA | B2M | beta-2-microglobulin | 567 | ENSG00000166710 |
| 6 | P48735 | NA | IDH2 | isocitrate dehydrogenase 2 (NADP+), mitochondrial | 3418 | ENSG00000182054 |
| 7 | P49588 | NA | AARS | alanyl-tRNA synthetase | 16 | ENSG00000090861 |
| 8 | Q86UP2 | NA | KTN1 | kinectin 1 (kinesin receptor) | 3895 | ENSG00000126777 |
| 9 | P02786 | NA | TFRC | transferrin receptor (p90, CD71) | 7037 | ENSG00000072274 |
| 10 | P02545 | NA | LMNA | lamin A/C | 4000 | ENSG00000160789 |
| 11 | P43490 | NA | NAMPT | nicotinamide phosphoribosyltransferase | 10135 | ENSG00000105835 |
| 12 | P17655 | NA | CAPN2 | calpain 2, (m/II) large subunit | 824 | ENSG00000162909 |
| 13 | O14773 | NA | TPP1 | tripeptidyl peptidase I | 1200 | ENSG00000166340 |
| 14 | Q9Y2Q3 | NA | GSTK1 | glutathione S-transferase kappa 1 | 373156 | ENSG00000197448 |
| 15 | P13639 | NA | EEF2 | eukaryotic translation elongation factor 2 | 1938 | ENSG00000167658 |
| 16 | Q16555 | NA | DPYSL2 | dihydropyrimidinase-like 2 | 1808 | ENSG00000092964 |
| 17 | P22570 | NA | FDXR | ferredoxin reductase | 2232 | ENSG00000161513 |
| 18 | O75521 | NA | ECI2 | enoyl-CoA delta isomerase 2 | 10455 | ENSG00000198721 |
| 19 | P49419 | NA | ALDH7A1 | aldehyde dehydrogenase 7 family, member A1 | 501 | ENSG00000164904 |
| 20 | Q04837 | NA | SSBP1 | single-stranded DNA binding protein 1, mitochondrial | 6742 | ENSG00000106028 |
| 21 | P21980 | NA | TGM2 | transglutaminase 2 (C polypeptide, protein-glutamine-gamma-glutamyltransferase) | 7052 | ENSG00000198959 |
| 22 | Q16822 | NA | PCK2 | phosphoenolpyruvate carboxykinase 2 (mitochondrial) | 5106 | ENSG00000100889 |
| 23 | Q9Y3E0 | NA | GOLT1B | golgi transport 1B | 51026 | ENSG00000111711 |
| 24 | P51648 | NA | ALDH3A2 | aldehyde dehydrogenase 3 family, member A2 | 224 | ENSG00000072210 |
| 25 | P21589 | NA | NT5E | 5'-nucleotidase, ecto (CD73) | 4907 | ENSG00000135318 |
| 26 | P02792 | NA | FTL | ferritin, light polypeptide | 2512 | ENSG00000087086 |
| 27 | Q7Z2K6 | NA | ERMP1 | endoplasmic reticulum metallopeptidase 1 | 79956 | ENSG00000099219 |
| 28 | P26447 | NA | S100A4 | S100 calcium binding protein A4 | 6275 | ENSG00000196154 |
| 29 | Q8NFV4 | NA | ABHD11 | abhydrolase domain containing 11 | 83451 | ENSG00000106077 |
| 30 | P51659 | NA | HSD17B4 | hydroxysteroid (17-beta) dehydrogenase 4 | 3295 | ENSG00000133835 |
| 31 | P21796 | NA | VDAC1 | voltage-dependent anion channel 1 | 7416 | ENSG00000213585 |
| 32 | P32322 | NA | PYCR1 | pyrroline-5-carboxylate reductase 1 | 5831 | ENSG00000183010 |
| 33 | P51688 | NA | SGSH | N-sulfoglucosamine sulfohydrolase | 6448 | ENSG00000181523 |
| 34 | P17987 | NA | TCP1 | t-complex 1 | 6950 | ENSG00000120438 |
| 35 | P05556 | NA | ITGB1 | integrin, beta 1 (fibronectin receptor, beta polypeptide, antigen CD29 includes MDF2, MSK12) | 3688 | ENSG00000150093 |
| 36 | P17096 | NA | HMGA1 | high mobility group AT-hook 1 | 3159 | ENSG00000137309 |
| 37 | P49748 | NA | ACADVL | acyl-CoA dehydrogenase, very long chain | 37 | ENSG00000072778 |
| 38 | O14979 | NA | HNRPDL | heterogeneous nuclear ribonucleoprotein D-like | 9987 | ENSG00000152795 |
| 39 | P00352 | NA | ALDH1A1 | aldehyde dehydrogenase 1 family, member A1 | 216 | ENSG00000165092 |
| 40 | P31937 | NA | HIBADH | 3-hydroxyisobutyrate dehydrogenase | 11112 | ENSG00000106049 |
| 41 | P30042 | NA | C21orf33 | chromosome 21 open reading frame 33 | 8209 | ENSG00000160221 |

  
  

| **Database:cellular component      &nbspName:nucleoid      &nbspID:GO:0009295** | | | | | | |
| --- | --- | --- | --- | --- | --- | --- |
| C=40; O=4; E=0.10; R=40.69; rawP=2.69e-06; adjP=2.42e-05 | | | | | | |
| Index | UserID | Value | Gene Symbol | Gene Name | EntrezGene | Ensembl |
| 1 | Q04837 | NA | SSBP1 | single-stranded DNA binding protein 1, mitochondrial | 6742 | ENSG00000106028 |
| 2 | P21796 | NA | VDAC1 | voltage-dependent anion channel 1 | 7416 | ENSG00000213585 |
| 3 | P49748 | NA | ACADVL | acyl-CoA dehydrogenase, very long chain | 37 | ENSG00000072778 |
| 4 | P04179 | NA | SOD2 | superoxide dismutase 2, mitochondrial | 6648 | ENSG00000112096 |

  
  

| **Database:cellular component      &nbspName:intracellular      &nbspID:GO:0005622** | | | | | | |
| --- | --- | --- | --- | --- | --- | --- |
| C=12412; O=41; E=30.51; R=1.34; rawP=5.35e-06; adjP=4.28e-05 | | | | | | |
| Index | UserID | Value | Gene Symbol | Gene Name | EntrezGene | Ensembl |
| 1 | P10253 | NA | GAA | glucosidase, alpha; acid | 2548 | ENSG00000171298 |
| 2 | P55060 | NA | CSE1L | CSE1 chromosome segregation 1-like (yeast) | 1434 | ENSG00000124207 |
| 3 | Q13724 | NA | MOGS | mannosyl-oligosaccharide glucosidase | 7841 | ENSG00000115275 |
| 4 | P04179 | NA | SOD2 | superoxide dismutase 2, mitochondrial | 6648 | ENSG00000112096 |
| 5 | P61769 | NA | B2M | beta-2-microglobulin | 567 | ENSG00000166710 |
| 6 | P48735 | NA | IDH2 | isocitrate dehydrogenase 2 (NADP+), mitochondrial | 3418 | ENSG00000182054 |
| 7 | P49588 | NA | AARS | alanyl-tRNA synthetase | 16 | ENSG00000090861 |
| 8 | Q86UP2 | NA | KTN1 | kinectin 1 (kinesin receptor) | 3895 | ENSG00000126777 |
| 9 | P02786 | NA | TFRC | transferrin receptor (p90, CD71) | 7037 | ENSG00000072274 |
| 10 | P02545 | NA | LMNA | lamin A/C | 4000 | ENSG00000160789 |
| 11 | P43490 | NA | NAMPT | nicotinamide phosphoribosyltransferase | 10135 | ENSG00000105835 |
| 12 | P17655 | NA | CAPN2 | calpain 2, (m/II) large subunit | 824 | ENSG00000162909 |
| 13 | O14773 | NA | TPP1 | tripeptidyl peptidase I | 1200 | ENSG00000166340 |
| 14 | Q9Y2Q3 | NA | GSTK1 | glutathione S-transferase kappa 1 | 373156 | ENSG00000197448 |
| 15 | P13639 | NA | EEF2 | eukaryotic translation elongation factor 2 | 1938 | ENSG00000167658 |
| 16 | Q16555 | NA | DPYSL2 | dihydropyrimidinase-like 2 | 1808 | ENSG00000092964 |
| 17 | P22570 | NA | FDXR | ferredoxin reductase | 2232 | ENSG00000161513 |
| 18 | O75521 | NA | ECI2 | enoyl-CoA delta isomerase 2 | 10455 | ENSG00000198721 |
| 19 | P49419 | NA | ALDH7A1 | aldehyde dehydrogenase 7 family, member A1 | 501 | ENSG00000164904 |
| 20 | Q04837 | NA | SSBP1 | single-stranded DNA binding protein 1, mitochondrial | 6742 | ENSG00000106028 |
| 21 | P21980 | NA | TGM2 | transglutaminase 2 (C polypeptide, protein-glutamine-gamma-glutamyltransferase) | 7052 | ENSG00000198959 |
| 22 | Q16822 | NA | PCK2 | phosphoenolpyruvate carboxykinase 2 (mitochondrial) | 5106 | ENSG00000100889 |
| 23 | Q9Y3E0 | NA | GOLT1B | golgi transport 1B | 51026 | ENSG00000111711 |
| 24 | P51648 | NA | ALDH3A2 | aldehyde dehydrogenase 3 family, member A2 | 224 | ENSG00000072210 |
| 25 | P21589 | NA | NT5E | 5'-nucleotidase, ecto (CD73) | 4907 | ENSG00000135318 |
| 26 | P02792 | NA | FTL | ferritin, light polypeptide | 2512 | ENSG00000087086 |
| 27 | Q7Z2K6 | NA | ERMP1 | endoplasmic reticulum metallopeptidase 1 | 79956 | ENSG00000099219 |
| 28 | P26447 | NA | S100A4 | S100 calcium binding protein A4 | 6275 | ENSG00000196154 |
| 29 | Q8NFV4 | NA | ABHD11 | abhydrolase domain containing 11 | 83451 | ENSG00000106077 |
| 30 | P51659 | NA | HSD17B4 | hydroxysteroid (17-beta) dehydrogenase 4 | 3295 | ENSG00000133835 |
| 31 | P21796 | NA | VDAC1 | voltage-dependent anion channel 1 | 7416 | ENSG00000213585 |
| 32 | P32322 | NA | PYCR1 | pyrroline-5-carboxylate reductase 1 | 5831 | ENSG00000183010 |
| 33 | P51688 | NA | SGSH | N-sulfoglucosamine sulfohydrolase | 6448 | ENSG00000181523 |
| 34 | P17987 | NA | TCP1 | t-complex 1 | 6950 | ENSG00000120438 |
| 35 | P05556 | NA | ITGB1 | integrin, beta 1 (fibronectin receptor, beta polypeptide, antigen CD29 includes MDF2, MSK12) | 3688 | ENSG00000150093 |
| 36 | P17096 | NA | HMGA1 | high mobility group AT-hook 1 | 3159 | ENSG00000137309 |
| 37 | P49748 | NA | ACADVL | acyl-CoA dehydrogenase, very long chain | 37 | ENSG00000072778 |
| 38 | O14979 | NA | HNRPDL | heterogeneous nuclear ribonucleoprotein D-like | 9987 | ENSG00000152795 |
| 39 | P00352 | NA | ALDH1A1 | aldehyde dehydrogenase 1 family, member A1 | 216 | ENSG00000165092 |
| 40 | P31937 | NA | HIBADH | 3-hydroxyisobutyrate dehydrogenase | 11112 | ENSG00000106049 |
| 41 | P30042 | NA | C21orf33 | chromosome 21 open reading frame 33 | 8209 | ENSG00000160221 |

  
  

| **Database:cellular component      &nbspName:intracellular membrane-bounded organelle      &nbspID:GO:0043231** | | | | | | |
| --- | --- | --- | --- | --- | --- | --- |
| C=9484; O=35; E=23.31; R=1.50; rawP=9.41e-05; adjP=0.0006 | | | | | | |
| Index | UserID | Value | Gene Symbol | Gene Name | EntrezGene | Ensembl |
| 1 | P10253 | NA | GAA | glucosidase, alpha; acid | 2548 | ENSG00000171298 |
| 2 | P55060 | NA | CSE1L | CSE1 chromosome segregation 1-like (yeast) | 1434 | ENSG00000124207 |
| 3 | Q13724 | NA | MOGS | mannosyl-oligosaccharide glucosidase | 7841 | ENSG00000115275 |
| 4 | P04179 | NA | SOD2 | superoxide dismutase 2, mitochondrial | 6648 | ENSG00000112096 |
| 5 | P61769 | NA | B2M | beta-2-microglobulin | 567 | ENSG00000166710 |
| 6 | P48735 | NA | IDH2 | isocitrate dehydrogenase 2 (NADP+), mitochondrial | 3418 | ENSG00000182054 |
| 7 | Q86UP2 | NA | KTN1 | kinectin 1 (kinesin receptor) | 3895 | ENSG00000126777 |
| 8 | P02545 | NA | LMNA | lamin A/C | 4000 | ENSG00000160789 |
| 9 | P02786 | NA | TFRC | transferrin receptor (p90, CD71) | 7037 | ENSG00000072274 |
| 10 | P17655 | NA | CAPN2 | calpain 2, (m/II) large subunit | 824 | ENSG00000162909 |
| 11 | O14773 | NA | TPP1 | tripeptidyl peptidase I | 1200 | ENSG00000166340 |
| 12 | Q9Y2Q3 | NA | GSTK1 | glutathione S-transferase kappa 1 | 373156 | ENSG00000197448 |
| 13 | Q16555 | NA | DPYSL2 | dihydropyrimidinase-like 2 | 1808 | ENSG00000092964 |
| 14 | P22570 | NA | FDXR | ferredoxin reductase | 2232 | ENSG00000161513 |
| 15 | O75521 | NA | ECI2 | enoyl-CoA delta isomerase 2 | 10455 | ENSG00000198721 |
| 16 | P49419 | NA | ALDH7A1 | aldehyde dehydrogenase 7 family, member A1 | 501 | ENSG00000164904 |
| 17 | Q04837 | NA | SSBP1 | single-stranded DNA binding protein 1, mitochondrial | 6742 | ENSG00000106028 |
| 18 | P21980 | NA | TGM2 | transglutaminase 2 (C polypeptide, protein-glutamine-gamma-glutamyltransferase) | 7052 | ENSG00000198959 |
| 19 | Q16822 | NA | PCK2 | phosphoenolpyruvate carboxykinase 2 (mitochondrial) | 5106 | ENSG00000100889 |
| 20 | Q9Y3E0 | NA | GOLT1B | golgi transport 1B | 51026 | ENSG00000111711 |
| 21 | P51648 | NA | ALDH3A2 | aldehyde dehydrogenase 3 family, member A2 | 224 | ENSG00000072210 |
| 22 | Q7Z2K6 | NA | ERMP1 | endoplasmic reticulum metallopeptidase 1 | 79956 | ENSG00000099219 |
| 23 | P26447 | NA | S100A4 | S100 calcium binding protein A4 | 6275 | ENSG00000196154 |
| 24 | Q8NFV4 | NA | ABHD11 | abhydrolase domain containing 11 | 83451 | ENSG00000106077 |
| 25 | P51659 | NA | HSD17B4 | hydroxysteroid (17-beta) dehydrogenase 4 | 3295 | ENSG00000133835 |
| 26 | P21796 | NA | VDAC1 | voltage-dependent anion channel 1 | 7416 | ENSG00000213585 |
| 27 | P32322 | NA | PYCR1 | pyrroline-5-carboxylate reductase 1 | 5831 | ENSG00000183010 |
| 28 | P17987 | NA | TCP1 | t-complex 1 | 6950 | ENSG00000120438 |
| 29 | P51688 | NA | SGSH | N-sulfoglucosamine sulfohydrolase | 6448 | ENSG00000181523 |
| 30 | P05556 | NA | ITGB1 | integrin, beta 1 (fibronectin receptor, beta polypeptide, antigen CD29 includes MDF2, MSK12) | 3688 | ENSG00000150093 |
| 31 | P17096 | NA | HMGA1 | high mobility group AT-hook 1 | 3159 | ENSG00000137309 |
| 32 | P49748 | NA | ACADVL | acyl-CoA dehydrogenase, very long chain | 37 | ENSG00000072778 |
| 33 | O14979 | NA | HNRPDL | heterogeneous nuclear ribonucleoprotein D-like | 9987 | ENSG00000152795 |
| 34 | P31937 | NA | HIBADH | 3-hydroxyisobutyrate dehydrogenase | 11112 | ENSG00000106049 |
| 35 | P30042 | NA | C21orf33 | chromosome 21 open reading frame 33 | 8209 | ENSG00000160221 |

  
  

| **Database:cellular component      &nbspName:membrane-bounded organelle      &nbspID:GO:0043227** | | | | | | |
| --- | --- | --- | --- | --- | --- | --- |
| C=9495; O=35; E=23.34; R=1.50; rawP=9.72e-05; adjP=0.0006 | | | | | | |
| Index | UserID | Value | Gene Symbol | Gene Name | EntrezGene | Ensembl |
| 1 | P10253 | NA | GAA | glucosidase, alpha; acid | 2548 | ENSG00000171298 |
| 2 | P55060 | NA | CSE1L | CSE1 chromosome segregation 1-like (yeast) | 1434 | ENSG00000124207 |
| 3 | Q13724 | NA | MOGS | mannosyl-oligosaccharide glucosidase | 7841 | ENSG00000115275 |
| 4 | P04179 | NA | SOD2 | superoxide dismutase 2, mitochondrial | 6648 | ENSG00000112096 |
| 5 | P61769 | NA | B2M | beta-2-microglobulin | 567 | ENSG00000166710 |
| 6 | P48735 | NA | IDH2 | isocitrate dehydrogenase 2 (NADP+), mitochondrial | 3418 | ENSG00000182054 |
| 7 | Q86UP2 | NA | KTN1 | kinectin 1 (kinesin receptor) | 3895 | ENSG00000126777 |
| 8 | P02545 | NA | LMNA | lamin A/C | 4000 | ENSG00000160789 |
| 9 | P02786 | NA | TFRC | transferrin receptor (p90, CD71) | 7037 | ENSG00000072274 |
| 10 | P17655 | NA | CAPN2 | calpain 2, (m/II) large subunit | 824 | ENSG00000162909 |
| 11 | O14773 | NA | TPP1 | tripeptidyl peptidase I | 1200 | ENSG00000166340 |
| 12 | Q9Y2Q3 | NA | GSTK1 | glutathione S-transferase kappa 1 | 373156 | ENSG00000197448 |
| 13 | Q16555 | NA | DPYSL2 | dihydropyrimidinase-like 2 | 1808 | ENSG00000092964 |
| 14 | P22570 | NA | FDXR | ferredoxin reductase | 2232 | ENSG00000161513 |
| 15 | O75521 | NA | ECI2 | enoyl-CoA delta isomerase 2 | 10455 | ENSG00000198721 |
| 16 | P49419 | NA | ALDH7A1 | aldehyde dehydrogenase 7 family, member A1 | 501 | ENSG00000164904 |
| 17 | Q04837 | NA | SSBP1 | single-stranded DNA binding protein 1, mitochondrial | 6742 | ENSG00000106028 |
| 18 | P21980 | NA | TGM2 | transglutaminase 2 (C polypeptide, protein-glutamine-gamma-glutamyltransferase) | 7052 | ENSG00000198959 |
| 19 | Q16822 | NA | PCK2 | phosphoenolpyruvate carboxykinase 2 (mitochondrial) | 5106 | ENSG00000100889 |
| 20 | Q9Y3E0 | NA | GOLT1B | golgi transport 1B | 51026 | ENSG00000111711 |
| 21 | P51648 | NA | ALDH3A2 | aldehyde dehydrogenase 3 family, member A2 | 224 | ENSG00000072210 |
| 22 | Q7Z2K6 | NA | ERMP1 | endoplasmic reticulum metallopeptidase 1 | 79956 | ENSG00000099219 |
| 23 | P26447 | NA | S100A4 | S100 calcium binding protein A4 | 6275 | ENSG00000196154 |
| 24 | Q8NFV4 | NA | ABHD11 | abhydrolase domain containing 11 | 83451 | ENSG00000106077 |
| 25 | P51659 | NA | HSD17B4 | hydroxysteroid (17-beta) dehydrogenase 4 | 3295 | ENSG00000133835 |
| 26 | P21796 | NA | VDAC1 | voltage-dependent anion channel 1 | 7416 | ENSG00000213585 |
| 27 | P32322 | NA | PYCR1 | pyrroline-5-carboxylate reductase 1 | 5831 | ENSG00000183010 |
| 28 | P17987 | NA | TCP1 | t-complex 1 | 6950 | ENSG00000120438 |
| 29 | P51688 | NA | SGSH | N-sulfoglucosamine sulfohydrolase | 6448 | ENSG00000181523 |
| 30 | P05556 | NA | ITGB1 | integrin, beta 1 (fibronectin receptor, beta polypeptide, antigen CD29 includes MDF2, MSK12) | 3688 | ENSG00000150093 |
| 31 | P17096 | NA | HMGA1 | high mobility group AT-hook 1 | 3159 | ENSG00000137309 |
| 32 | P49748 | NA | ACADVL | acyl-CoA dehydrogenase, very long chain | 37 | ENSG00000072778 |
| 33 | O14979 | NA | HNRPDL | heterogeneous nuclear ribonucleoprotein D-like | 9987 | ENSG00000152795 |
| 34 | P31937 | NA | HIBADH | 3-hydroxyisobutyrate dehydrogenase | 11112 | ENSG00000106049 |
| 35 | P30042 | NA | C21orf33 | chromosome 21 open reading frame 33 | 8209 | ENSG00000160221 |

  
  
  
  

---

WebGestalt is currently developed and maintained by Jing Wang and Bing Zhang at the  Zhang Lab. Other people who have made significant contribution to the project include Dexter Duncan, Stefan Kirov, Zhiao Shi, and Jay Snoddy.  
  
**Funding credits:** NIH/NIAAA (U01 AA016662, U01 AA013512); NIH/NIDA (P01 DA015027); NIH/NIMH (P50 MH078028, P50 MH096972); NIH/NCI (U24 CA159988); NIH/NIGMS (R01 GM088822).
