## Supplementary material for "PGRMC1 phosphorylation and cell plasticity 1: glycolysis, mitochondria, tumor growth": File S7: README.txt

Open the DAG_timestamp.html file in your browser.  This is an image map file.
Click on the red nodes in the DAG to view your genes of interest with descriptions
and links to ensembl.

The files directory contains the gif file and the html file with anchors. The
final_sig_file_timestamp.html may be opened also in your browser for viewing.

The final_params_timestamp.txt file has the parameters used for the analysis.
