## Supplementary material for "PGRMC1 phosphorylation and cell plasticity 1: glycolysis, mitochondria, tumor growth": File S7: README.txt

Open final_KEGG_file_# in a browser to view the same results one would find by clicking on KEGG Enrichment in the results page.

The first column is the name of the pathway with a link to KEGG with your ids highlighted.
The second column is the number of your uniq ids that are in the pathway with a link to the final_sig_kegg_file_# that has
the user ids, values if uploaded, Ensebml ids with link to Ensembl, Entrez Ids with link to Entrez, Symbol, and Description.
The fourth column has the entrez ids with links to Entrez for the respective user entrez ids.
The fifth column has the Stats for the Enrichment Analysis.

The final_params_timestamp.txt file has the parameters used for the analysis.
