## Supplementary material for "PGRMC1 phosphorylation and cell plasticity 1: glycolysis, mitochondria, tumor growth": File S7: final_pc_geneset_file_1438486678.html

Anchored HTML File of EIDs

|  |  |
| --- | --- |
|  | WEB-based GEne SeT AnaLysis Toolkit |
|  | ***Translating gene lists into biological insights...*** |

---

  

| **Database:Pathway Commons pathway      &nbspName:Sphingosine 1-phosphate (S1P) pathway      &nbspID:DB\_ID:1635** | | | | | | |
| --- | --- | --- | --- | --- | --- | --- |
| C=1311; O=11; E=2.65; R=4.15; rawP=4.11e-05; adjP=5.54e-05 | | | | | | |
| Index | UserID | Value | Gene Symbol | Gene Name | EntrezGene | Ensembl |
| 1 | P17655 | NA | CAPN2 | calpain 2, (m/II) large subunit | 824 | ENSG00000162909 |
| 2 | P13639 | NA | EEF2 | eukaryotic translation elongation factor 2 | 1938 | ENSG00000167658 |
| 3 | P22570 | NA | FDXR | ferredoxin reductase | 2232 | ENSG00000161513 |
| 4 | P05556 | NA | ITGB1 | integrin, beta 1 (fibronectin receptor, beta polypeptide, antigen CD29 includes MDF2, MSK12) | 3688 | ENSG00000150093 |
| 5 | P55060 | NA | CSE1L | CSE1 chromosome segregation 1-like (yeast) | 1434 | ENSG00000124207 |
| 6 | P04179 | NA | SOD2 | superoxide dismutase 2, mitochondrial | 6648 | ENSG00000112096 |
| 7 | P61769 | NA | B2M | beta-2-microglobulin | 567 | ENSG00000166710 |
| 8 | P17096 | NA | HMGA1 | high mobility group AT-hook 1 | 3159 | ENSG00000137309 |
| 9 | Q16822 | NA | PCK2 | phosphoenolpyruvate carboxykinase 2 (mitochondrial) | 5106 | ENSG00000100889 |
| 10 | P02786 | NA | TFRC | transferrin receptor (p90, CD71) | 7037 | ENSG00000072274 |
| 11 | P21589 | NA | NT5E | 5'-nucleotidase, ecto (CD73) | 4907 | ENSG00000135318 |

  
  

| **Database:Pathway Commons pathway      &nbspName:Thrombin/protease-activated receptor (PAR) pathway      &nbspID:DB\_ID:1552** | | | | | | |
| --- | --- | --- | --- | --- | --- | --- |
| C=1300; O=11; E=2.63; R=4.19; rawP=3.80e-05; adjP=5.54e-05 | | | | | | |
| Index | UserID | Value | Gene Symbol | Gene Name | EntrezGene | Ensembl |
| 1 | P17655 | NA | CAPN2 | calpain 2, (m/II) large subunit | 824 | ENSG00000162909 |
| 2 | P13639 | NA | EEF2 | eukaryotic translation elongation factor 2 | 1938 | ENSG00000167658 |
| 3 | P22570 | NA | FDXR | ferredoxin reductase | 2232 | ENSG00000161513 |
| 4 | P05556 | NA | ITGB1 | integrin, beta 1 (fibronectin receptor, beta polypeptide, antigen CD29 includes MDF2, MSK12) | 3688 | ENSG00000150093 |
| 5 | P55060 | NA | CSE1L | CSE1 chromosome segregation 1-like (yeast) | 1434 | ENSG00000124207 |
| 6 | P04179 | NA | SOD2 | superoxide dismutase 2, mitochondrial | 6648 | ENSG00000112096 |
| 7 | P61769 | NA | B2M | beta-2-microglobulin | 567 | ENSG00000166710 |
| 8 | P17096 | NA | HMGA1 | high mobility group AT-hook 1 | 3159 | ENSG00000137309 |
| 9 | Q16822 | NA | PCK2 | phosphoenolpyruvate carboxykinase 2 (mitochondrial) | 5106 | ENSG00000100889 |
| 10 | P02786 | NA | TFRC | transferrin receptor (p90, CD71) | 7037 | ENSG00000072274 |
| 11 | P21589 | NA | NT5E | 5'-nucleotidase, ecto (CD73) | 4907 | ENSG00000135318 |

  
  

| **Database:Pathway Commons pathway      &nbspName:ErbB1 downstream signaling      &nbspID:DB\_ID:1602** | | | | | | |
| --- | --- | --- | --- | --- | --- | --- |
| C=1288; O=11; E=2.60; R=4.22; rawP=3.49e-05; adjP=5.54e-05 | | | | | | |
| Index | UserID | Value | Gene Symbol | Gene Name | EntrezGene | Ensembl |
| 1 | P17655 | NA | CAPN2 | calpain 2, (m/II) large subunit | 824 | ENSG00000162909 |
| 2 | P13639 | NA | EEF2 | eukaryotic translation elongation factor 2 | 1938 | ENSG00000167658 |
| 3 | P22570 | NA | FDXR | ferredoxin reductase | 2232 | ENSG00000161513 |
| 4 | P05556 | NA | ITGB1 | integrin, beta 1 (fibronectin receptor, beta polypeptide, antigen CD29 includes MDF2, MSK12) | 3688 | ENSG00000150093 |
| 5 | P55060 | NA | CSE1L | CSE1 chromosome segregation 1-like (yeast) | 1434 | ENSG00000124207 |
| 6 | P04179 | NA | SOD2 | superoxide dismutase 2, mitochondrial | 6648 | ENSG00000112096 |
| 7 | P61769 | NA | B2M | beta-2-microglobulin | 567 | ENSG00000166710 |
| 8 | P17096 | NA | HMGA1 | high mobility group AT-hook 1 | 3159 | ENSG00000137309 |
| 9 | Q16822 | NA | PCK2 | phosphoenolpyruvate carboxykinase 2 (mitochondrial) | 5106 | ENSG00000100889 |
| 10 | P02786 | NA | TFRC | transferrin receptor (p90, CD71) | 7037 | ENSG00000072274 |
| 11 | P21589 | NA | NT5E | 5'-nucleotidase, ecto (CD73) | 4907 | ENSG00000135318 |

  
  

| **Database:Pathway Commons pathway      &nbspName:Arf6 trafficking events      &nbspID:DB\_ID:1615** | | | | | | |
| --- | --- | --- | --- | --- | --- | --- |
| C=1288; O=11; E=2.60; R=4.22; rawP=3.49e-05; adjP=5.54e-05 | | | | | | |
| Index | UserID | Value | Gene Symbol | Gene Name | EntrezGene | Ensembl |
| 1 | P17655 | NA | CAPN2 | calpain 2, (m/II) large subunit | 824 | ENSG00000162909 |
| 2 | P13639 | NA | EEF2 | eukaryotic translation elongation factor 2 | 1938 | ENSG00000167658 |
| 3 | P22570 | NA | FDXR | ferredoxin reductase | 2232 | ENSG00000161513 |
| 4 | P05556 | NA | ITGB1 | integrin, beta 1 (fibronectin receptor, beta polypeptide, antigen CD29 includes MDF2, MSK12) | 3688 | ENSG00000150093 |
| 5 | P55060 | NA | CSE1L | CSE1 chromosome segregation 1-like (yeast) | 1434 | ENSG00000124207 |
| 6 | P04179 | NA | SOD2 | superoxide dismutase 2, mitochondrial | 6648 | ENSG00000112096 |
| 7 | P61769 | NA | B2M | beta-2-microglobulin | 567 | ENSG00000166710 |
| 8 | P17096 | NA | HMGA1 | high mobility group AT-hook 1 | 3159 | ENSG00000137309 |
| 9 | Q16822 | NA | PCK2 | phosphoenolpyruvate carboxykinase 2 (mitochondrial) | 5106 | ENSG00000100889 |
| 10 | P02786 | NA | TFRC | transferrin receptor (p90, CD71) | 7037 | ENSG00000072274 |
| 11 | P21589 | NA | NT5E | 5'-nucleotidase, ecto (CD73) | 4907 | ENSG00000135318 |

  
  

| **Database:Pathway Commons pathway      &nbspName:Alpha9 beta1 integrin signaling events      &nbspID:DB\_ID:1578** | | | | | | |
| --- | --- | --- | --- | --- | --- | --- |
| C=1305; O=12; E=2.64; R=4.55; rawP=6.58e-06; adjP=5.54e-05 | | | | | | |
| Index | UserID | Value | Gene Symbol | Gene Name | EntrezGene | Ensembl |
| 1 | P17655 | NA | CAPN2 | calpain 2, (m/II) large subunit | 824 | ENSG00000162909 |
| 2 | P13639 | NA | EEF2 | eukaryotic translation elongation factor 2 | 1938 | ENSG00000167658 |
| 3 | P22570 | NA | FDXR | ferredoxin reductase | 2232 | ENSG00000161513 |
| 4 | P05556 | NA | ITGB1 | integrin, beta 1 (fibronectin receptor, beta polypeptide, antigen CD29 includes MDF2, MSK12) | 3688 | ENSG00000150093 |
| 5 | P55060 | NA | CSE1L | CSE1 chromosome segregation 1-like (yeast) | 1434 | ENSG00000124207 |
| 6 | P04179 | NA | SOD2 | superoxide dismutase 2, mitochondrial | 6648 | ENSG00000112096 |
| 7 | P61769 | NA | B2M | beta-2-microglobulin | 567 | ENSG00000166710 |
| 8 | P17096 | NA | HMGA1 | high mobility group AT-hook 1 | 3159 | ENSG00000137309 |
| 9 | P21980 | NA | TGM2 | transglutaminase 2 (C polypeptide, protein-glutamine-gamma-glutamyltransferase) | 7052 | ENSG00000198959 |
| 10 | Q16822 | NA | PCK2 | phosphoenolpyruvate carboxykinase 2 (mitochondrial) | 5106 | ENSG00000100889 |
| 11 | P02786 | NA | TFRC | transferrin receptor (p90, CD71) | 7037 | ENSG00000072274 |
| 12 | P21589 | NA | NT5E | 5'-nucleotidase, ecto (CD73) | 4907 | ENSG00000135318 |

  
  

| **Database:Pathway Commons pathway      &nbspName:Arf6 signaling events      &nbspID:DB\_ID:1554** | | | | | | |
| --- | --- | --- | --- | --- | --- | --- |
| C=1288; O=11; E=2.60; R=4.22; rawP=3.49e-05; adjP=5.54e-05 | | | | | | |
| Index | UserID | Value | Gene Symbol | Gene Name | EntrezGene | Ensembl |
| 1 | P17655 | NA | CAPN2 | calpain 2, (m/II) large subunit | 824 | ENSG00000162909 |
| 2 | P13639 | NA | EEF2 | eukaryotic translation elongation factor 2 | 1938 | ENSG00000167658 |
| 3 | P22570 | NA | FDXR | ferredoxin reductase | 2232 | ENSG00000161513 |
| 4 | P05556 | NA | ITGB1 | integrin, beta 1 (fibronectin receptor, beta polypeptide, antigen CD29 includes MDF2, MSK12) | 3688 | ENSG00000150093 |
| 5 | P55060 | NA | CSE1L | CSE1 chromosome segregation 1-like (yeast) | 1434 | ENSG00000124207 |
| 6 | P04179 | NA | SOD2 | superoxide dismutase 2, mitochondrial | 6648 | ENSG00000112096 |
| 7 | P61769 | NA | B2M | beta-2-microglobulin | 567 | ENSG00000166710 |
| 8 | P17096 | NA | HMGA1 | high mobility group AT-hook 1 | 3159 | ENSG00000137309 |
| 9 | Q16822 | NA | PCK2 | phosphoenolpyruvate carboxykinase 2 (mitochondrial) | 5106 | ENSG00000100889 |
| 10 | P02786 | NA | TFRC | transferrin receptor (p90, CD71) | 7037 | ENSG00000072274 |
| 11 | P21589 | NA | NT5E | 5'-nucleotidase, ecto (CD73) | 4907 | ENSG00000135318 |

  
  

| **Database:Pathway Commons pathway      &nbspName:Arf6 downstream pathway      &nbspID:DB\_ID:1585** | | | | | | |
| --- | --- | --- | --- | --- | --- | --- |
| C=1288; O=11; E=2.60; R=4.22; rawP=3.49e-05; adjP=5.54e-05 | | | | | | |
| Index | UserID | Value | Gene Symbol | Gene Name | EntrezGene | Ensembl |
| 1 | P17655 | NA | CAPN2 | calpain 2, (m/II) large subunit | 824 | ENSG00000162909 |
| 2 | P13639 | NA | EEF2 | eukaryotic translation elongation factor 2 | 1938 | ENSG00000167658 |
| 3 | P22570 | NA | FDXR | ferredoxin reductase | 2232 | ENSG00000161513 |
| 4 | P05556 | NA | ITGB1 | integrin, beta 1 (fibronectin receptor, beta polypeptide, antigen CD29 includes MDF2, MSK12) | 3688 | ENSG00000150093 |
| 5 | P55060 | NA | CSE1L | CSE1 chromosome segregation 1-like (yeast) | 1434 | ENSG00000124207 |
| 6 | P04179 | NA | SOD2 | superoxide dismutase 2, mitochondrial | 6648 | ENSG00000112096 |
| 7 | P61769 | NA | B2M | beta-2-microglobulin | 567 | ENSG00000166710 |
| 8 | P17096 | NA | HMGA1 | high mobility group AT-hook 1 | 3159 | ENSG00000137309 |
| 9 | Q16822 | NA | PCK2 | phosphoenolpyruvate carboxykinase 2 (mitochondrial) | 5106 | ENSG00000100889 |
| 10 | P02786 | NA | TFRC | transferrin receptor (p90, CD71) | 7037 | ENSG00000072274 |
| 11 | P21589 | NA | NT5E | 5'-nucleotidase, ecto (CD73) | 4907 | ENSG00000135318 |

  
  

| **Database:Pathway Commons pathway      &nbspName:IFN-gamma pathway      &nbspID:DB\_ID:1529** | | | | | | |
| --- | --- | --- | --- | --- | --- | --- |
| C=1296; O=11; E=2.62; R=4.20; rawP=3.69e-05; adjP=5.54e-05 | | | | | | |
| Index | UserID | Value | Gene Symbol | Gene Name | EntrezGene | Ensembl |
| 1 | P17655 | NA | CAPN2 | calpain 2, (m/II) large subunit | 824 | ENSG00000162909 |
| 2 | P13639 | NA | EEF2 | eukaryotic translation elongation factor 2 | 1938 | ENSG00000167658 |
| 3 | P22570 | NA | FDXR | ferredoxin reductase | 2232 | ENSG00000161513 |
| 4 | P05556 | NA | ITGB1 | integrin, beta 1 (fibronectin receptor, beta polypeptide, antigen CD29 includes MDF2, MSK12) | 3688 | ENSG00000150093 |
| 5 | P55060 | NA | CSE1L | CSE1 chromosome segregation 1-like (yeast) | 1434 | ENSG00000124207 |
| 6 | P04179 | NA | SOD2 | superoxide dismutase 2, mitochondrial | 6648 | ENSG00000112096 |
| 7 | P61769 | NA | B2M | beta-2-microglobulin | 567 | ENSG00000166710 |
| 8 | P17096 | NA | HMGA1 | high mobility group AT-hook 1 | 3159 | ENSG00000137309 |
| 9 | Q16822 | NA | PCK2 | phosphoenolpyruvate carboxykinase 2 (mitochondrial) | 5106 | ENSG00000100889 |
| 10 | P02786 | NA | TFRC | transferrin receptor (p90, CD71) | 7037 | ENSG00000072274 |
| 11 | P21589 | NA | NT5E | 5'-nucleotidase, ecto (CD73) | 4907 | ENSG00000135318 |

  
  

| **Database:Pathway Commons pathway      &nbspName:Nectin adhesion pathway      &nbspID:DB\_ID:1472** | | | | | | |
| --- | --- | --- | --- | --- | --- | --- |
| C=1295; O=11; E=2.62; R=4.20; rawP=3.67e-05; adjP=5.54e-05 | | | | | | |
| Index | UserID | Value | Gene Symbol | Gene Name | EntrezGene | Ensembl |
| 1 | P17655 | NA | CAPN2 | calpain 2, (m/II) large subunit | 824 | ENSG00000162909 |
| 2 | P13639 | NA | EEF2 | eukaryotic translation elongation factor 2 | 1938 | ENSG00000167658 |
| 3 | P22570 | NA | FDXR | ferredoxin reductase | 2232 | ENSG00000161513 |
| 4 | P05556 | NA | ITGB1 | integrin, beta 1 (fibronectin receptor, beta polypeptide, antigen CD29 includes MDF2, MSK12) | 3688 | ENSG00000150093 |
| 5 | P55060 | NA | CSE1L | CSE1 chromosome segregation 1-like (yeast) | 1434 | ENSG00000124207 |
| 6 | P04179 | NA | SOD2 | superoxide dismutase 2, mitochondrial | 6648 | ENSG00000112096 |
| 7 | P61769 | NA | B2M | beta-2-microglobulin | 567 | ENSG00000166710 |
| 8 | P17096 | NA | HMGA1 | high mobility group AT-hook 1 | 3159 | ENSG00000137309 |
| 9 | Q16822 | NA | PCK2 | phosphoenolpyruvate carboxykinase 2 (mitochondrial) | 5106 | ENSG00000100889 |
| 10 | P02786 | NA | TFRC | transferrin receptor (p90, CD71) | 7037 | ENSG00000072274 |
| 11 | P21589 | NA | NT5E | 5'-nucleotidase, ecto (CD73) | 4907 | ENSG00000135318 |

  
  

| **Database:Pathway Commons pathway      &nbspName:S1P1 pathway      &nbspID:DB\_ID:1594** | | | | | | |
| --- | --- | --- | --- | --- | --- | --- |
| C=1288; O=11; E=2.60; R=4.22; rawP=3.49e-05; adjP=5.54e-05 | | | | | | |
| Index | UserID | Value | Gene Symbol | Gene Name | EntrezGene | Ensembl |
| 1 | P17655 | NA | CAPN2 | calpain 2, (m/II) large subunit | 824 | ENSG00000162909 |
| 2 | P13639 | NA | EEF2 | eukaryotic translation elongation factor 2 | 1938 | ENSG00000167658 |
| 3 | P22570 | NA | FDXR | ferredoxin reductase | 2232 | ENSG00000161513 |
| 4 | P05556 | NA | ITGB1 | integrin, beta 1 (fibronectin receptor, beta polypeptide, antigen CD29 includes MDF2, MSK12) | 3688 | ENSG00000150093 |
| 5 | P55060 | NA | CSE1L | CSE1 chromosome segregation 1-like (yeast) | 1434 | ENSG00000124207 |
| 6 | P04179 | NA | SOD2 | superoxide dismutase 2, mitochondrial | 6648 | ENSG00000112096 |
| 7 | P61769 | NA | B2M | beta-2-microglobulin | 567 | ENSG00000166710 |
| 8 | P17096 | NA | HMGA1 | high mobility group AT-hook 1 | 3159 | ENSG00000137309 |
| 9 | Q16822 | NA | PCK2 | phosphoenolpyruvate carboxykinase 2 (mitochondrial) | 5106 | ENSG00000100889 |
| 10 | P02786 | NA | TFRC | transferrin receptor (p90, CD71) | 7037 | ENSG00000072274 |
| 11 | P21589 | NA | NT5E | 5'-nucleotidase, ecto (CD73) | 4907 | ENSG00000135318 |

  
  

| **Database:Pathway Commons pathway      &nbspName:Signaling events mediated by focal adhesion kinase      &nbspID:DB\_ID:1574** | | | | | | |
| --- | --- | --- | --- | --- | --- | --- |
| C=1288; O=11; E=2.60; R=4.22; rawP=3.49e-05; adjP=5.54e-05 | | | | | | |
| Index | UserID | Value | Gene Symbol | Gene Name | EntrezGene | Ensembl |
| 1 | P17655 | NA | CAPN2 | calpain 2, (m/II) large subunit | 824 | ENSG00000162909 |
| 2 | P13639 | NA | EEF2 | eukaryotic translation elongation factor 2 | 1938 | ENSG00000167658 |
| 3 | P22570 | NA | FDXR | ferredoxin reductase | 2232 | ENSG00000161513 |
| 4 | P05556 | NA | ITGB1 | integrin, beta 1 (fibronectin receptor, beta polypeptide, antigen CD29 includes MDF2, MSK12) | 3688 | ENSG00000150093 |
| 5 | P55060 | NA | CSE1L | CSE1 chromosome segregation 1-like (yeast) | 1434 | ENSG00000124207 |
| 6 | P04179 | NA | SOD2 | superoxide dismutase 2, mitochondrial | 6648 | ENSG00000112096 |
| 7 | P61769 | NA | B2M | beta-2-microglobulin | 567 | ENSG00000166710 |
| 8 | P17096 | NA | HMGA1 | high mobility group AT-hook 1 | 3159 | ENSG00000137309 |
| 9 | Q16822 | NA | PCK2 | phosphoenolpyruvate carboxykinase 2 (mitochondrial) | 5106 | ENSG00000100889 |
| 10 | P02786 | NA | TFRC | transferrin receptor (p90, CD71) | 7037 | ENSG00000072274 |
| 11 | P21589 | NA | NT5E | 5'-nucleotidase, ecto (CD73) | 4907 | ENSG00000135318 |

  
  

| **Database:Pathway Commons pathway      &nbspName:EGFR-dependent Endothelin signaling events      &nbspID:DB\_ID:1603** | | | | | | |
| --- | --- | --- | --- | --- | --- | --- |
| C=1289; O=11; E=2.61; R=4.22; rawP=3.51e-05; adjP=5.54e-05 | | | | | | |
| Index | UserID | Value | Gene Symbol | Gene Name | EntrezGene | Ensembl |
| 1 | P17655 | NA | CAPN2 | calpain 2, (m/II) large subunit | 824 | ENSG00000162909 |
| 2 | P13639 | NA | EEF2 | eukaryotic translation elongation factor 2 | 1938 | ENSG00000167658 |
| 3 | P22570 | NA | FDXR | ferredoxin reductase | 2232 | ENSG00000161513 |
| 4 | P05556 | NA | ITGB1 | integrin, beta 1 (fibronectin receptor, beta polypeptide, antigen CD29 includes MDF2, MSK12) | 3688 | ENSG00000150093 |
| 5 | P55060 | NA | CSE1L | CSE1 chromosome segregation 1-like (yeast) | 1434 | ENSG00000124207 |
| 6 | P04179 | NA | SOD2 | superoxide dismutase 2, mitochondrial | 6648 | ENSG00000112096 |
| 7 | P61769 | NA | B2M | beta-2-microglobulin | 567 | ENSG00000166710 |
| 8 | P17096 | NA | HMGA1 | high mobility group AT-hook 1 | 3159 | ENSG00000137309 |
| 9 | Q16822 | NA | PCK2 | phosphoenolpyruvate carboxykinase 2 (mitochondrial) | 5106 | ENSG00000100889 |
| 10 | P02786 | NA | TFRC | transferrin receptor (p90, CD71) | 7037 | ENSG00000072274 |
| 11 | P21589 | NA | NT5E | 5'-nucleotidase, ecto (CD73) | 4907 | ENSG00000135318 |

  
  

| **Database:Pathway Commons pathway      &nbspName:Internalization of ErbB1      &nbspID:DB\_ID:1509** | | | | | | |
| --- | --- | --- | --- | --- | --- | --- |
| C=1288; O=11; E=2.60; R=4.22; rawP=3.49e-05; adjP=5.54e-05 | | | | | | |
| Index | UserID | Value | Gene Symbol | Gene Name | EntrezGene | Ensembl |
| 1 | P17655 | NA | CAPN2 | calpain 2, (m/II) large subunit | 824 | ENSG00000162909 |
| 2 | P13639 | NA | EEF2 | eukaryotic translation elongation factor 2 | 1938 | ENSG00000167658 |
| 3 | P22570 | NA | FDXR | ferredoxin reductase | 2232 | ENSG00000161513 |
| 4 | P05556 | NA | ITGB1 | integrin, beta 1 (fibronectin receptor, beta polypeptide, antigen CD29 includes MDF2, MSK12) | 3688 | ENSG00000150093 |
| 5 | P55060 | NA | CSE1L | CSE1 chromosome segregation 1-like (yeast) | 1434 | ENSG00000124207 |
| 6 | P04179 | NA | SOD2 | superoxide dismutase 2, mitochondrial | 6648 | ENSG00000112096 |
| 7 | P61769 | NA | B2M | beta-2-microglobulin | 567 | ENSG00000166710 |
| 8 | P17096 | NA | HMGA1 | high mobility group AT-hook 1 | 3159 | ENSG00000137309 |
| 9 | Q16822 | NA | PCK2 | phosphoenolpyruvate carboxykinase 2 (mitochondrial) | 5106 | ENSG00000100889 |
| 10 | P02786 | NA | TFRC | transferrin receptor (p90, CD71) | 7037 | ENSG00000072274 |
| 11 | P21589 | NA | NT5E | 5'-nucleotidase, ecto (CD73) | 4907 | ENSG00000135318 |

  
  

| **Database:Pathway Commons pathway      &nbspName:Signaling events mediated by VEGFR1 and VEGFR2      &nbspID:DB\_ID:1516** | | | | | | |
| --- | --- | --- | --- | --- | --- | --- |
| C=1296; O=11; E=2.62; R=4.20; rawP=3.69e-05; adjP=5.54e-05 | | | | | | |
| Index | UserID | Value | Gene Symbol | Gene Name | EntrezGene | Ensembl |
| 1 | P17655 | NA | CAPN2 | calpain 2, (m/II) large subunit | 824 | ENSG00000162909 |
| 2 | P13639 | NA | EEF2 | eukaryotic translation elongation factor 2 | 1938 | ENSG00000167658 |
| 3 | P22570 | NA | FDXR | ferredoxin reductase | 2232 | ENSG00000161513 |
| 4 | P05556 | NA | ITGB1 | integrin, beta 1 (fibronectin receptor, beta polypeptide, antigen CD29 includes MDF2, MSK12) | 3688 | ENSG00000150093 |
| 5 | P55060 | NA | CSE1L | CSE1 chromosome segregation 1-like (yeast) | 1434 | ENSG00000124207 |
| 6 | P04179 | NA | SOD2 | superoxide dismutase 2, mitochondrial | 6648 | ENSG00000112096 |
| 7 | P61769 | NA | B2M | beta-2-microglobulin | 567 | ENSG00000166710 |
| 8 | P17096 | NA | HMGA1 | high mobility group AT-hook 1 | 3159 | ENSG00000137309 |
| 9 | Q16822 | NA | PCK2 | phosphoenolpyruvate carboxykinase 2 (mitochondrial) | 5106 | ENSG00000100889 |
| 10 | P02786 | NA | TFRC | transferrin receptor (p90, CD71) | 7037 | ENSG00000072274 |
| 11 | P21589 | NA | NT5E | 5'-nucleotidase, ecto (CD73) | 4907 | ENSG00000135318 |

  
  

| **Database:Pathway Commons pathway      &nbspName:Beta1 integrin cell surface interactions      &nbspID:DB\_ID:1517** | | | | | | |
| --- | --- | --- | --- | --- | --- | --- |
| C=1351; O=12; E=2.73; R=4.39; rawP=9.37e-06; adjP=5.54e-05 | | | | | | |
| Index | UserID | Value | Gene Symbol | Gene Name | EntrezGene | Ensembl |
| 1 | P17655 | NA | CAPN2 | calpain 2, (m/II) large subunit | 824 | ENSG00000162909 |
| 2 | P13639 | NA | EEF2 | eukaryotic translation elongation factor 2 | 1938 | ENSG00000167658 |
| 3 | P22570 | NA | FDXR | ferredoxin reductase | 2232 | ENSG00000161513 |
| 4 | P05556 | NA | ITGB1 | integrin, beta 1 (fibronectin receptor, beta polypeptide, antigen CD29 includes MDF2, MSK12) | 3688 | ENSG00000150093 |
| 5 | P55060 | NA | CSE1L | CSE1 chromosome segregation 1-like (yeast) | 1434 | ENSG00000124207 |
| 6 | P04179 | NA | SOD2 | superoxide dismutase 2, mitochondrial | 6648 | ENSG00000112096 |
| 7 | P61769 | NA | B2M | beta-2-microglobulin | 567 | ENSG00000166710 |
| 8 | P17096 | NA | HMGA1 | high mobility group AT-hook 1 | 3159 | ENSG00000137309 |
| 9 | P21980 | NA | TGM2 | transglutaminase 2 (C polypeptide, protein-glutamine-gamma-glutamyltransferase) | 7052 | ENSG00000198959 |
| 10 | Q16822 | NA | PCK2 | phosphoenolpyruvate carboxykinase 2 (mitochondrial) | 5106 | ENSG00000100889 |
| 11 | P02786 | NA | TFRC | transferrin receptor (p90, CD71) | 7037 | ENSG00000072274 |
| 12 | P21589 | NA | NT5E | 5'-nucleotidase, ecto (CD73) | 4907 | ENSG00000135318 |

  
  

| **Database:Pathway Commons pathway      &nbspName:IL3-mediated signaling events      &nbspID:DB\_ID:1564** | | | | | | |
| --- | --- | --- | --- | --- | --- | --- |
| C=1295; O=11; E=2.62; R=4.20; rawP=3.67e-05; adjP=5.54e-05 | | | | | | |
| Index | UserID | Value | Gene Symbol | Gene Name | EntrezGene | Ensembl |
| 1 | P17655 | NA | CAPN2 | calpain 2, (m/II) large subunit | 824 | ENSG00000162909 |
| 2 | P13639 | NA | EEF2 | eukaryotic translation elongation factor 2 | 1938 | ENSG00000167658 |
| 3 | P22570 | NA | FDXR | ferredoxin reductase | 2232 | ENSG00000161513 |
| 4 | P05556 | NA | ITGB1 | integrin, beta 1 (fibronectin receptor, beta polypeptide, antigen CD29 includes MDF2, MSK12) | 3688 | ENSG00000150093 |
| 5 | P55060 | NA | CSE1L | CSE1 chromosome segregation 1-like (yeast) | 1434 | ENSG00000124207 |
| 6 | P04179 | NA | SOD2 | superoxide dismutase 2, mitochondrial | 6648 | ENSG00000112096 |
| 7 | P61769 | NA | B2M | beta-2-microglobulin | 567 | ENSG00000166710 |
| 8 | P17096 | NA | HMGA1 | high mobility group AT-hook 1 | 3159 | ENSG00000137309 |
| 9 | Q16822 | NA | PCK2 | phosphoenolpyruvate carboxykinase 2 (mitochondrial) | 5106 | ENSG00000100889 |
| 10 | P02786 | NA | TFRC | transferrin receptor (p90, CD71) | 7037 | ENSG00000072274 |
| 11 | P21589 | NA | NT5E | 5'-nucleotidase, ecto (CD73) | 4907 | ENSG00000135318 |

  
  

| **Database:Pathway Commons pathway      &nbspName:GMCSF-mediated signaling events      &nbspID:DB\_ID:1461** | | | | | | |
| --- | --- | --- | --- | --- | --- | --- |
| C=1292; O=11; E=2.61; R=4.21; rawP=3.59e-05; adjP=5.54e-05 | | | | | | |
| Index | UserID | Value | Gene Symbol | Gene Name | EntrezGene | Ensembl |
| 1 | P17655 | NA | CAPN2 | calpain 2, (m/II) large subunit | 824 | ENSG00000162909 |
| 2 | P13639 | NA | EEF2 | eukaryotic translation elongation factor 2 | 1938 | ENSG00000167658 |
| 3 | P22570 | NA | FDXR | ferredoxin reductase | 2232 | ENSG00000161513 |
| 4 | P05556 | NA | ITGB1 | integrin, beta 1 (fibronectin receptor, beta polypeptide, antigen CD29 includes MDF2, MSK12) | 3688 | ENSG00000150093 |
| 5 | P55060 | NA | CSE1L | CSE1 chromosome segregation 1-like (yeast) | 1434 | ENSG00000124207 |
| 6 | P04179 | NA | SOD2 | superoxide dismutase 2, mitochondrial | 6648 | ENSG00000112096 |
| 7 | P61769 | NA | B2M | beta-2-microglobulin | 567 | ENSG00000166710 |
| 8 | P17096 | NA | HMGA1 | high mobility group AT-hook 1 | 3159 | ENSG00000137309 |
| 9 | Q16822 | NA | PCK2 | phosphoenolpyruvate carboxykinase 2 (mitochondrial) | 5106 | ENSG00000100889 |
| 10 | P02786 | NA | TFRC | transferrin receptor (p90, CD71) | 7037 | ENSG00000072274 |
| 11 | P21589 | NA | NT5E | 5'-nucleotidase, ecto (CD73) | 4907 | ENSG00000135318 |

  
  

| **Database:Pathway Commons pathway      &nbspName:Plasma membrane estrogen receptor signaling      &nbspID:DB\_ID:1556** | | | | | | |
| --- | --- | --- | --- | --- | --- | --- |
| C=1301; O=11; E=2.63; R=4.18; rawP=3.83e-05; adjP=5.54e-05 | | | | | | |
| Index | UserID | Value | Gene Symbol | Gene Name | EntrezGene | Ensembl |
| 1 | P17655 | NA | CAPN2 | calpain 2, (m/II) large subunit | 824 | ENSG00000162909 |
| 2 | P13639 | NA | EEF2 | eukaryotic translation elongation factor 2 | 1938 | ENSG00000167658 |
| 3 | P22570 | NA | FDXR | ferredoxin reductase | 2232 | ENSG00000161513 |
| 4 | P05556 | NA | ITGB1 | integrin, beta 1 (fibronectin receptor, beta polypeptide, antigen CD29 includes MDF2, MSK12) | 3688 | ENSG00000150093 |
| 5 | P55060 | NA | CSE1L | CSE1 chromosome segregation 1-like (yeast) | 1434 | ENSG00000124207 |
| 6 | P04179 | NA | SOD2 | superoxide dismutase 2, mitochondrial | 6648 | ENSG00000112096 |
| 7 | P61769 | NA | B2M | beta-2-microglobulin | 567 | ENSG00000166710 |
| 8 | P17096 | NA | HMGA1 | high mobility group AT-hook 1 | 3159 | ENSG00000137309 |
| 9 | Q16822 | NA | PCK2 | phosphoenolpyruvate carboxykinase 2 (mitochondrial) | 5106 | ENSG00000100889 |
| 10 | P02786 | NA | TFRC | transferrin receptor (p90, CD71) | 7037 | ENSG00000072274 |
| 11 | P21589 | NA | NT5E | 5'-nucleotidase, ecto (CD73) | 4907 | ENSG00000135318 |

  
  

| **Database:Pathway Commons pathway      &nbspName:TRAIL signaling pathway      &nbspID:DB\_ID:1480** | | | | | | |
| --- | --- | --- | --- | --- | --- | --- |
| C=1328; O=12; E=2.68; R=4.47; rawP=7.87e-06; adjP=5.54e-05 | | | | | | |
| Index | UserID | Value | Gene Symbol | Gene Name | EntrezGene | Ensembl |
| 1 | P17655 | NA | CAPN2 | calpain 2, (m/II) large subunit | 824 | ENSG00000162909 |
| 2 | P13639 | NA | EEF2 | eukaryotic translation elongation factor 2 | 1938 | ENSG00000167658 |
| 3 | P22570 | NA | FDXR | ferredoxin reductase | 2232 | ENSG00000161513 |
| 4 | P05556 | NA | ITGB1 | integrin, beta 1 (fibronectin receptor, beta polypeptide, antigen CD29 includes MDF2, MSK12) | 3688 | ENSG00000150093 |
| 5 | P55060 | NA | CSE1L | CSE1 chromosome segregation 1-like (yeast) | 1434 | ENSG00000124207 |
| 6 | P04179 | NA | SOD2 | superoxide dismutase 2, mitochondrial | 6648 | ENSG00000112096 |
| 7 | P61769 | NA | B2M | beta-2-microglobulin | 567 | ENSG00000166710 |
| 8 | P17096 | NA | HMGA1 | high mobility group AT-hook 1 | 3159 | ENSG00000137309 |
| 9 | Q16822 | NA | PCK2 | phosphoenolpyruvate carboxykinase 2 (mitochondrial) | 5106 | ENSG00000100889 |
| 10 | P02545 | NA | LMNA | lamin A/C | 4000 | ENSG00000160789 |
| 11 | P02786 | NA | TFRC | transferrin receptor (p90, CD71) | 7037 | ENSG00000072274 |
| 12 | P21589 | NA | NT5E | 5'-nucleotidase, ecto (CD73) | 4907 | ENSG00000135318 |

  
  

| **Database:Pathway Commons pathway      &nbspName:Insulin Pathway      &nbspID:DB\_ID:1466** | | | | | | |
| --- | --- | --- | --- | --- | --- | --- |
| C=1288; O=11; E=2.60; R=4.22; rawP=3.49e-05; adjP=5.54e-05 | | | | | | |
| Index | UserID | Value | Gene Symbol | Gene Name | EntrezGene | Ensembl |
| 1 | P17655 | NA | CAPN2 | calpain 2, (m/II) large subunit | 824 | ENSG00000162909 |
| 2 | P13639 | NA | EEF2 | eukaryotic translation elongation factor 2 | 1938 | ENSG00000167658 |
| 3 | P22570 | NA | FDXR | ferredoxin reductase | 2232 | ENSG00000161513 |
| 4 | P05556 | NA | ITGB1 | integrin, beta 1 (fibronectin receptor, beta polypeptide, antigen CD29 includes MDF2, MSK12) | 3688 | ENSG00000150093 |
| 5 | P55060 | NA | CSE1L | CSE1 chromosome segregation 1-like (yeast) | 1434 | ENSG00000124207 |
| 6 | P04179 | NA | SOD2 | superoxide dismutase 2, mitochondrial | 6648 | ENSG00000112096 |
| 7 | P61769 | NA | B2M | beta-2-microglobulin | 567 | ENSG00000166710 |
| 8 | P17096 | NA | HMGA1 | high mobility group AT-hook 1 | 3159 | ENSG00000137309 |
| 9 | Q16822 | NA | PCK2 | phosphoenolpyruvate carboxykinase 2 (mitochondrial) | 5106 | ENSG00000100889 |
| 10 | P02786 | NA | TFRC | transferrin receptor (p90, CD71) | 7037 | ENSG00000072274 |
| 11 | P21589 | NA | NT5E | 5'-nucleotidase, ecto (CD73) | 4907 | ENSG00000135318 |

  
  

| **Database:Pathway Commons pathway      &nbspName:Integrin family cell surface interactions      &nbspID:DB\_ID:1499** | | | | | | |
| --- | --- | --- | --- | --- | --- | --- |
| C=1378; O=12; E=2.79; R=4.31; rawP=1.15e-05; adjP=5.54e-05 | | | | | | |
| Index | UserID | Value | Gene Symbol | Gene Name | EntrezGene | Ensembl |
| 1 | P17655 | NA | CAPN2 | calpain 2, (m/II) large subunit | 824 | ENSG00000162909 |
| 2 | P13639 | NA | EEF2 | eukaryotic translation elongation factor 2 | 1938 | ENSG00000167658 |
| 3 | P22570 | NA | FDXR | ferredoxin reductase | 2232 | ENSG00000161513 |
| 4 | P05556 | NA | ITGB1 | integrin, beta 1 (fibronectin receptor, beta polypeptide, antigen CD29 includes MDF2, MSK12) | 3688 | ENSG00000150093 |
| 5 | P55060 | NA | CSE1L | CSE1 chromosome segregation 1-like (yeast) | 1434 | ENSG00000124207 |
| 6 | P04179 | NA | SOD2 | superoxide dismutase 2, mitochondrial | 6648 | ENSG00000112096 |
| 7 | P61769 | NA | B2M | beta-2-microglobulin | 567 | ENSG00000166710 |
| 8 | P17096 | NA | HMGA1 | high mobility group AT-hook 1 | 3159 | ENSG00000137309 |
| 9 | P21980 | NA | TGM2 | transglutaminase 2 (C polypeptide, protein-glutamine-gamma-glutamyltransferase) | 7052 | ENSG00000198959 |
| 10 | Q16822 | NA | PCK2 | phosphoenolpyruvate carboxykinase 2 (mitochondrial) | 5106 | ENSG00000100889 |
| 11 | P02786 | NA | TFRC | transferrin receptor (p90, CD71) | 7037 | ENSG00000072274 |
| 12 | P21589 | NA | NT5E | 5'-nucleotidase, ecto (CD73) | 4907 | ENSG00000135318 |

  
  

| **Database:Pathway Commons pathway      &nbspName:EGF receptor (ErbB1) signaling pathway      &nbspID:DB\_ID:1550** | | | | | | |
| --- | --- | --- | --- | --- | --- | --- |
| C=1288; O=11; E=2.60; R=4.22; rawP=3.49e-05; adjP=5.54e-05 | | | | | | |
| Index | UserID | Value | Gene Symbol | Gene Name | EntrezGene | Ensembl |
| 1 | P17655 | NA | CAPN2 | calpain 2, (m/II) large subunit | 824 | ENSG00000162909 |
| 2 | P13639 | NA | EEF2 | eukaryotic translation elongation factor 2 | 1938 | ENSG00000167658 |
| 3 | P22570 | NA | FDXR | ferredoxin reductase | 2232 | ENSG00000161513 |
| 4 | P05556 | NA | ITGB1 | integrin, beta 1 (fibronectin receptor, beta polypeptide, antigen CD29 includes MDF2, MSK12) | 3688 | ENSG00000150093 |
| 5 | P55060 | NA | CSE1L | CSE1 chromosome segregation 1-like (yeast) | 1434 | ENSG00000124207 |
| 6 | P04179 | NA | SOD2 | superoxide dismutase 2, mitochondrial | 6648 | ENSG00000112096 |
| 7 | P61769 | NA | B2M | beta-2-microglobulin | 567 | ENSG00000166710 |
| 8 | P17096 | NA | HMGA1 | high mobility group AT-hook 1 | 3159 | ENSG00000137309 |
| 9 | Q16822 | NA | PCK2 | phosphoenolpyruvate carboxykinase 2 (mitochondrial) | 5106 | ENSG00000100889 |
| 10 | P02786 | NA | TFRC | transferrin receptor (p90, CD71) | 7037 | ENSG00000072274 |
| 11 | P21589 | NA | NT5E | 5'-nucleotidase, ecto (CD73) | 4907 | ENSG00000135318 |

  
  

| **Database:Pathway Commons pathway      &nbspName:Syndecan-1-mediated signaling events      &nbspID:DB\_ID:1454** | | | | | | |
| --- | --- | --- | --- | --- | --- | --- |
| C=1300; O=11; E=2.63; R=4.19; rawP=3.80e-05; adjP=5.54e-05 | | | | | | |
| Index | UserID | Value | Gene Symbol | Gene Name | EntrezGene | Ensembl |
| 1 | P17655 | NA | CAPN2 | calpain 2, (m/II) large subunit | 824 | ENSG00000162909 |
| 2 | P13639 | NA | EEF2 | eukaryotic translation elongation factor 2 | 1938 | ENSG00000167658 |
| 3 | P22570 | NA | FDXR | ferredoxin reductase | 2232 | ENSG00000161513 |
| 4 | P05556 | NA | ITGB1 | integrin, beta 1 (fibronectin receptor, beta polypeptide, antigen CD29 includes MDF2, MSK12) | 3688 | ENSG00000150093 |
| 5 | P55060 | NA | CSE1L | CSE1 chromosome segregation 1-like (yeast) | 1434 | ENSG00000124207 |
| 6 | P04179 | NA | SOD2 | superoxide dismutase 2, mitochondrial | 6648 | ENSG00000112096 |
| 7 | P61769 | NA | B2M | beta-2-microglobulin | 567 | ENSG00000166710 |
| 8 | P17096 | NA | HMGA1 | high mobility group AT-hook 1 | 3159 | ENSG00000137309 |
| 9 | Q16822 | NA | PCK2 | phosphoenolpyruvate carboxykinase 2 (mitochondrial) | 5106 | ENSG00000100889 |
| 10 | P02786 | NA | TFRC | transferrin receptor (p90, CD71) | 7037 | ENSG00000072274 |
| 11 | P21589 | NA | NT5E | 5'-nucleotidase, ecto (CD73) | 4907 | ENSG00000135318 |

  
  

| **Database:Pathway Commons pathway      &nbspName:Class I PI3K signaling events mediated by Akt      &nbspID:DB\_ID:1648** | | | | | | |
| --- | --- | --- | --- | --- | --- | --- |
| C=1288; O=11; E=2.60; R=4.22; rawP=3.49e-05; adjP=5.54e-05 | | | | | | |
| Index | UserID | Value | Gene Symbol | Gene Name | EntrezGene | Ensembl |
| 1 | P17655 | NA | CAPN2 | calpain 2, (m/II) large subunit | 824 | ENSG00000162909 |
| 2 | P13639 | NA | EEF2 | eukaryotic translation elongation factor 2 | 1938 | ENSG00000167658 |
| 3 | P22570 | NA | FDXR | ferredoxin reductase | 2232 | ENSG00000161513 |
| 4 | P05556 | NA | ITGB1 | integrin, beta 1 (fibronectin receptor, beta polypeptide, antigen CD29 includes MDF2, MSK12) | 3688 | ENSG00000150093 |
| 5 | P55060 | NA | CSE1L | CSE1 chromosome segregation 1-like (yeast) | 1434 | ENSG00000124207 |
| 6 | P04179 | NA | SOD2 | superoxide dismutase 2, mitochondrial | 6648 | ENSG00000112096 |
| 7 | P61769 | NA | B2M | beta-2-microglobulin | 567 | ENSG00000166710 |
| 8 | P17096 | NA | HMGA1 | high mobility group AT-hook 1 | 3159 | ENSG00000137309 |
| 9 | Q16822 | NA | PCK2 | phosphoenolpyruvate carboxykinase 2 (mitochondrial) | 5106 | ENSG00000100889 |
| 10 | P02786 | NA | TFRC | transferrin receptor (p90, CD71) | 7037 | ENSG00000072274 |
| 11 | P21589 | NA | NT5E | 5'-nucleotidase, ecto (CD73) | 4907 | ENSG00000135318 |

  
  

| **Database:Pathway Commons pathway      &nbspName:Urokinase-type plasminogen activator (uPA) and uPAR-mediated signaling      &nbspID:DB\_ID:1519** | | | | | | |
| --- | --- | --- | --- | --- | --- | --- |
| C=1288; O=11; E=2.60; R=4.22; rawP=3.49e-05; adjP=5.54e-05 | | | | | | |
| Index | UserID | Value | Gene Symbol | Gene Name | EntrezGene | Ensembl |
| 1 | P17655 | NA | CAPN2 | calpain 2, (m/II) large subunit | 824 | ENSG00000162909 |
| 2 | P13639 | NA | EEF2 | eukaryotic translation elongation factor 2 | 1938 | ENSG00000167658 |
| 3 | P22570 | NA | FDXR | ferredoxin reductase | 2232 | ENSG00000161513 |
| 4 | P05556 | NA | ITGB1 | integrin, beta 1 (fibronectin receptor, beta polypeptide, antigen CD29 includes MDF2, MSK12) | 3688 | ENSG00000150093 |
| 5 | P55060 | NA | CSE1L | CSE1 chromosome segregation 1-like (yeast) | 1434 | ENSG00000124207 |
| 6 | P04179 | NA | SOD2 | superoxide dismutase 2, mitochondrial | 6648 | ENSG00000112096 |
| 7 | P61769 | NA | B2M | beta-2-microglobulin | 567 | ENSG00000166710 |
| 8 | P17096 | NA | HMGA1 | high mobility group AT-hook 1 | 3159 | ENSG00000137309 |
| 9 | Q16822 | NA | PCK2 | phosphoenolpyruvate carboxykinase 2 (mitochondrial) | 5106 | ENSG00000100889 |
| 10 | P02786 | NA | TFRC | transferrin receptor (p90, CD71) | 7037 | ENSG00000072274 |
| 11 | P21589 | NA | NT5E | 5'-nucleotidase, ecto (CD73) | 4907 | ENSG00000135318 |

  
  

| **Database:Pathway Commons pathway      &nbspName:PAR1-mediated thrombin signaling events      &nbspID:DB\_ID:1531** | | | | | | |
| --- | --- | --- | --- | --- | --- | --- |
| C=1299; O=11; E=2.63; R=4.19; rawP=3.77e-05; adjP=5.54e-05 | | | | | | |
| Index | UserID | Value | Gene Symbol | Gene Name | EntrezGene | Ensembl |
| 1 | P17655 | NA | CAPN2 | calpain 2, (m/II) large subunit | 824 | ENSG00000162909 |
| 2 | P13639 | NA | EEF2 | eukaryotic translation elongation factor 2 | 1938 | ENSG00000167658 |
| 3 | P22570 | NA | FDXR | ferredoxin reductase | 2232 | ENSG00000161513 |
| 4 | P05556 | NA | ITGB1 | integrin, beta 1 (fibronectin receptor, beta polypeptide, antigen CD29 includes MDF2, MSK12) | 3688 | ENSG00000150093 |
| 5 | P55060 | NA | CSE1L | CSE1 chromosome segregation 1-like (yeast) | 1434 | ENSG00000124207 |
| 6 | P04179 | NA | SOD2 | superoxide dismutase 2, mitochondrial | 6648 | ENSG00000112096 |
| 7 | P61769 | NA | B2M | beta-2-microglobulin | 567 | ENSG00000166710 |
| 8 | P17096 | NA | HMGA1 | high mobility group AT-hook 1 | 3159 | ENSG00000137309 |
| 9 | Q16822 | NA | PCK2 | phosphoenolpyruvate carboxykinase 2 (mitochondrial) | 5106 | ENSG00000100889 |
| 10 | P02786 | NA | TFRC | transferrin receptor (p90, CD71) | 7037 | ENSG00000072274 |
| 11 | P21589 | NA | NT5E | 5'-nucleotidase, ecto (CD73) | 4907 | ENSG00000135318 |

  
  

| **Database:Pathway Commons pathway      &nbspName:ErbB receptor signaling network      &nbspID:DB\_ID:1573** | | | | | | |
| --- | --- | --- | --- | --- | --- | --- |
| C=1312; O=11; E=2.65; R=4.15; rawP=4.13e-05; adjP=5.54e-05 | | | | | | |
| Index | UserID | Value | Gene Symbol | Gene Name | EntrezGene | Ensembl |
| 1 | P17655 | NA | CAPN2 | calpain 2, (m/II) large subunit | 824 | ENSG00000162909 |
| 2 | P13639 | NA | EEF2 | eukaryotic translation elongation factor 2 | 1938 | ENSG00000167658 |
| 3 | P22570 | NA | FDXR | ferredoxin reductase | 2232 | ENSG00000161513 |
| 4 | P05556 | NA | ITGB1 | integrin, beta 1 (fibronectin receptor, beta polypeptide, antigen CD29 includes MDF2, MSK12) | 3688 | ENSG00000150093 |
| 5 | P55060 | NA | CSE1L | CSE1 chromosome segregation 1-like (yeast) | 1434 | ENSG00000124207 |
| 6 | P04179 | NA | SOD2 | superoxide dismutase 2, mitochondrial | 6648 | ENSG00000112096 |
| 7 | P61769 | NA | B2M | beta-2-microglobulin | 567 | ENSG00000166710 |
| 8 | P17096 | NA | HMGA1 | high mobility group AT-hook 1 | 3159 | ENSG00000137309 |
| 9 | Q16822 | NA | PCK2 | phosphoenolpyruvate carboxykinase 2 (mitochondrial) | 5106 | ENSG00000100889 |
| 10 | P02786 | NA | TFRC | transferrin receptor (p90, CD71) | 7037 | ENSG00000072274 |
| 11 | P21589 | NA | NT5E | 5'-nucleotidase, ecto (CD73) | 4907 | ENSG00000135318 |

  
  

| **Database:Pathway Commons pathway      &nbspName:Glypican 1 network      &nbspID:DB\_ID:1492** | | | | | | |
| --- | --- | --- | --- | --- | --- | --- |
| C=1299; O=11; E=2.63; R=4.19; rawP=3.77e-05; adjP=5.54e-05 | | | | | | |
| Index | UserID | Value | Gene Symbol | Gene Name | EntrezGene | Ensembl |
| 1 | P17655 | NA | CAPN2 | calpain 2, (m/II) large subunit | 824 | ENSG00000162909 |
| 2 | P13639 | NA | EEF2 | eukaryotic translation elongation factor 2 | 1938 | ENSG00000167658 |
| 3 | P22570 | NA | FDXR | ferredoxin reductase | 2232 | ENSG00000161513 |
| 4 | P05556 | NA | ITGB1 | integrin, beta 1 (fibronectin receptor, beta polypeptide, antigen CD29 includes MDF2, MSK12) | 3688 | ENSG00000150093 |
| 5 | P55060 | NA | CSE1L | CSE1 chromosome segregation 1-like (yeast) | 1434 | ENSG00000124207 |
| 6 | P04179 | NA | SOD2 | superoxide dismutase 2, mitochondrial | 6648 | ENSG00000112096 |
| 7 | P61769 | NA | B2M | beta-2-microglobulin | 567 | ENSG00000166710 |
| 8 | P17096 | NA | HMGA1 | high mobility group AT-hook 1 | 3159 | ENSG00000137309 |
| 9 | Q16822 | NA | PCK2 | phosphoenolpyruvate carboxykinase 2 (mitochondrial) | 5106 | ENSG00000100889 |
| 10 | P02786 | NA | TFRC | transferrin receptor (p90, CD71) | 7037 | ENSG00000072274 |
| 11 | P21589 | NA | NT5E | 5'-nucleotidase, ecto (CD73) | 4907 | ENSG00000135318 |

  
  

| **Database:Pathway Commons pathway      &nbspName:PDGF receptor signaling network      &nbspID:DB\_ID:1497** | | | | | | |
| --- | --- | --- | --- | --- | --- | --- |
| C=1293; O=11; E=2.61; R=4.21; rawP=3.62e-05; adjP=5.54e-05 | | | | | | |
| Index | UserID | Value | Gene Symbol | Gene Name | EntrezGene | Ensembl |
| 1 | P17655 | NA | CAPN2 | calpain 2, (m/II) large subunit | 824 | ENSG00000162909 |
| 2 | P13639 | NA | EEF2 | eukaryotic translation elongation factor 2 | 1938 | ENSG00000167658 |
| 3 | P22570 | NA | FDXR | ferredoxin reductase | 2232 | ENSG00000161513 |
| 4 | P05556 | NA | ITGB1 | integrin, beta 1 (fibronectin receptor, beta polypeptide, antigen CD29 includes MDF2, MSK12) | 3688 | ENSG00000150093 |
| 5 | P55060 | NA | CSE1L | CSE1 chromosome segregation 1-like (yeast) | 1434 | ENSG00000124207 |
| 6 | P04179 | NA | SOD2 | superoxide dismutase 2, mitochondrial | 6648 | ENSG00000112096 |
| 7 | P61769 | NA | B2M | beta-2-microglobulin | 567 | ENSG00000166710 |
| 8 | P17096 | NA | HMGA1 | high mobility group AT-hook 1 | 3159 | ENSG00000137309 |
| 9 | Q16822 | NA | PCK2 | phosphoenolpyruvate carboxykinase 2 (mitochondrial) | 5106 | ENSG00000100889 |
| 10 | P02786 | NA | TFRC | transferrin receptor (p90, CD71) | 7037 | ENSG00000072274 |
| 11 | P21589 | NA | NT5E | 5'-nucleotidase, ecto (CD73) | 4907 | ENSG00000135318 |

  
  

| **Database:Pathway Commons pathway      &nbspName:Signaling events mediated by Hepatocyte Growth Factor Receptor (c-Met)      &nbspID:DB\_ID:1491** | | | | | | |
| --- | --- | --- | --- | --- | --- | --- |
| C=1293; O=11; E=2.61; R=4.21; rawP=3.62e-05; adjP=5.54e-05 | | | | | | |
| Index | UserID | Value | Gene Symbol | Gene Name | EntrezGene | Ensembl |
| 1 | P17655 | NA | CAPN2 | calpain 2, (m/II) large subunit | 824 | ENSG00000162909 |
| 2 | P13639 | NA | EEF2 | eukaryotic translation elongation factor 2 | 1938 | ENSG00000167658 |
| 3 | P22570 | NA | FDXR | ferredoxin reductase | 2232 | ENSG00000161513 |
| 4 | P05556 | NA | ITGB1 | integrin, beta 1 (fibronectin receptor, beta polypeptide, antigen CD29 includes MDF2, MSK12) | 3688 | ENSG00000150093 |
| 5 | P55060 | NA | CSE1L | CSE1 chromosome segregation 1-like (yeast) | 1434 | ENSG00000124207 |
| 6 | P04179 | NA | SOD2 | superoxide dismutase 2, mitochondrial | 6648 | ENSG00000112096 |
| 7 | P61769 | NA | B2M | beta-2-microglobulin | 567 | ENSG00000166710 |
| 8 | P17096 | NA | HMGA1 | high mobility group AT-hook 1 | 3159 | ENSG00000137309 |
| 9 | Q16822 | NA | PCK2 | phosphoenolpyruvate carboxykinase 2 (mitochondrial) | 5106 | ENSG00000100889 |
| 10 | P02786 | NA | TFRC | transferrin receptor (p90, CD71) | 7037 | ENSG00000072274 |
| 11 | P21589 | NA | NT5E | 5'-nucleotidase, ecto (CD73) | 4907 | ENSG00000135318 |

  
  

| **Database:Pathway Commons pathway      &nbspName:mTOR signaling pathway      &nbspID:DB\_ID:1571** | | | | | | |
| --- | --- | --- | --- | --- | --- | --- |
| C=1288; O=11; E=2.60; R=4.22; rawP=3.49e-05; adjP=5.54e-05 | | | | | | |
| Index | UserID | Value | Gene Symbol | Gene Name | EntrezGene | Ensembl |
| 1 | P17655 | NA | CAPN2 | calpain 2, (m/II) large subunit | 824 | ENSG00000162909 |
| 2 | P13639 | NA | EEF2 | eukaryotic translation elongation factor 2 | 1938 | ENSG00000167658 |
| 3 | P22570 | NA | FDXR | ferredoxin reductase | 2232 | ENSG00000161513 |
| 4 | P05556 | NA | ITGB1 | integrin, beta 1 (fibronectin receptor, beta polypeptide, antigen CD29 includes MDF2, MSK12) | 3688 | ENSG00000150093 |
| 5 | P55060 | NA | CSE1L | CSE1 chromosome segregation 1-like (yeast) | 1434 | ENSG00000124207 |
| 6 | P04179 | NA | SOD2 | superoxide dismutase 2, mitochondrial | 6648 | ENSG00000112096 |
| 7 | P61769 | NA | B2M | beta-2-microglobulin | 567 | ENSG00000166710 |
| 8 | P17096 | NA | HMGA1 | high mobility group AT-hook 1 | 3159 | ENSG00000137309 |
| 9 | Q16822 | NA | PCK2 | phosphoenolpyruvate carboxykinase 2 (mitochondrial) | 5106 | ENSG00000100889 |
| 10 | P02786 | NA | TFRC | transferrin receptor (p90, CD71) | 7037 | ENSG00000072274 |
| 11 | P21589 | NA | NT5E | 5'-nucleotidase, ecto (CD73) | 4907 | ENSG00000135318 |

  
  

| **Database:Pathway Commons pathway      &nbspName:IGF1 pathway      &nbspID:DB\_ID:1482** | | | | | | |
| --- | --- | --- | --- | --- | --- | --- |
| C=1291; O=11; E=2.61; R=4.22; rawP=3.56e-05; adjP=5.54e-05 | | | | | | |
| Index | UserID | Value | Gene Symbol | Gene Name | EntrezGene | Ensembl |
| 1 | P17655 | NA | CAPN2 | calpain 2, (m/II) large subunit | 824 | ENSG00000162909 |
| 2 | P13639 | NA | EEF2 | eukaryotic translation elongation factor 2 | 1938 | ENSG00000167658 |
| 3 | P22570 | NA | FDXR | ferredoxin reductase | 2232 | ENSG00000161513 |
| 4 | P05556 | NA | ITGB1 | integrin, beta 1 (fibronectin receptor, beta polypeptide, antigen CD29 includes MDF2, MSK12) | 3688 | ENSG00000150093 |
| 5 | P55060 | NA | CSE1L | CSE1 chromosome segregation 1-like (yeast) | 1434 | ENSG00000124207 |
| 6 | P04179 | NA | SOD2 | superoxide dismutase 2, mitochondrial | 6648 | ENSG00000112096 |
| 7 | P61769 | NA | B2M | beta-2-microglobulin | 567 | ENSG00000166710 |
| 8 | P17096 | NA | HMGA1 | high mobility group AT-hook 1 | 3159 | ENSG00000137309 |
| 9 | Q16822 | NA | PCK2 | phosphoenolpyruvate carboxykinase 2 (mitochondrial) | 5106 | ENSG00000100889 |
| 10 | P02786 | NA | TFRC | transferrin receptor (p90, CD71) | 7037 | ENSG00000072274 |
| 11 | P21589 | NA | NT5E | 5'-nucleotidase, ecto (CD73) | 4907 | ENSG00000135318 |

  
  

| **Database:Pathway Commons pathway      &nbspName:VEGF and VEGFR signaling network      &nbspID:DB\_ID:1575** | | | | | | |
| --- | --- | --- | --- | --- | --- | --- |
| C=1304; O=11; E=2.64; R=4.17; rawP=3.91e-05; adjP=5.54e-05 | | | | | | |
| Index | UserID | Value | Gene Symbol | Gene Name | EntrezGene | Ensembl |
| 1 | P17655 | NA | CAPN2 | calpain 2, (m/II) large subunit | 824 | ENSG00000162909 |
| 2 | P13639 | NA | EEF2 | eukaryotic translation elongation factor 2 | 1938 | ENSG00000167658 |
| 3 | P22570 | NA | FDXR | ferredoxin reductase | 2232 | ENSG00000161513 |
| 4 | P05556 | NA | ITGB1 | integrin, beta 1 (fibronectin receptor, beta polypeptide, antigen CD29 includes MDF2, MSK12) | 3688 | ENSG00000150093 |
| 5 | P55060 | NA | CSE1L | CSE1 chromosome segregation 1-like (yeast) | 1434 | ENSG00000124207 |
| 6 | P04179 | NA | SOD2 | superoxide dismutase 2, mitochondrial | 6648 | ENSG00000112096 |
| 7 | P61769 | NA | B2M | beta-2-microglobulin | 567 | ENSG00000166710 |
| 8 | P17096 | NA | HMGA1 | high mobility group AT-hook 1 | 3159 | ENSG00000137309 |
| 9 | Q16822 | NA | PCK2 | phosphoenolpyruvate carboxykinase 2 (mitochondrial) | 5106 | ENSG00000100889 |
| 10 | P02786 | NA | TFRC | transferrin receptor (p90, CD71) | 7037 | ENSG00000072274 |
| 11 | P21589 | NA | NT5E | 5'-nucleotidase, ecto (CD73) | 4907 | ENSG00000135318 |

  
  

| **Database:Pathway Commons pathway      &nbspName:Endothelins      &nbspID:DB\_ID:1619** | | | | | | |
| --- | --- | --- | --- | --- | --- | --- |
| C=1307; O=11; E=2.64; R=4.16; rawP=3.99e-05; adjP=5.54e-05 | | | | | | |
| Index | UserID | Value | Gene Symbol | Gene Name | EntrezGene | Ensembl |
| 1 | P17655 | NA | CAPN2 | calpain 2, (m/II) large subunit | 824 | ENSG00000162909 |
| 2 | P13639 | NA | EEF2 | eukaryotic translation elongation factor 2 | 1938 | ENSG00000167658 |
| 3 | P22570 | NA | FDXR | ferredoxin reductase | 2232 | ENSG00000161513 |
| 4 | P05556 | NA | ITGB1 | integrin, beta 1 (fibronectin receptor, beta polypeptide, antigen CD29 includes MDF2, MSK12) | 3688 | ENSG00000150093 |
| 5 | P55060 | NA | CSE1L | CSE1 chromosome segregation 1-like (yeast) | 1434 | ENSG00000124207 |
| 6 | P04179 | NA | SOD2 | superoxide dismutase 2, mitochondrial | 6648 | ENSG00000112096 |
| 7 | P61769 | NA | B2M | beta-2-microglobulin | 567 | ENSG00000166710 |
| 8 | P17096 | NA | HMGA1 | high mobility group AT-hook 1 | 3159 | ENSG00000137309 |
| 9 | Q16822 | NA | PCK2 | phosphoenolpyruvate carboxykinase 2 (mitochondrial) | 5106 | ENSG00000100889 |
| 10 | P02786 | NA | TFRC | transferrin receptor (p90, CD71) | 7037 | ENSG00000072274 |
| 11 | P21589 | NA | NT5E | 5'-nucleotidase, ecto (CD73) | 4907 | ENSG00000135318 |

  
  

| **Database:Pathway Commons pathway      &nbspName:IL5-mediated signaling events      &nbspID:DB\_ID:1627** | | | | | | |
| --- | --- | --- | --- | --- | --- | --- |
| C=1292; O=11; E=2.61; R=4.21; rawP=3.59e-05; adjP=5.54e-05 | | | | | | |
| Index | UserID | Value | Gene Symbol | Gene Name | EntrezGene | Ensembl |
| 1 | P17655 | NA | CAPN2 | calpain 2, (m/II) large subunit | 824 | ENSG00000162909 |
| 2 | P13639 | NA | EEF2 | eukaryotic translation elongation factor 2 | 1938 | ENSG00000167658 |
| 3 | P22570 | NA | FDXR | ferredoxin reductase | 2232 | ENSG00000161513 |
| 4 | P05556 | NA | ITGB1 | integrin, beta 1 (fibronectin receptor, beta polypeptide, antigen CD29 includes MDF2, MSK12) | 3688 | ENSG00000150093 |
| 5 | P55060 | NA | CSE1L | CSE1 chromosome segregation 1-like (yeast) | 1434 | ENSG00000124207 |
| 6 | P04179 | NA | SOD2 | superoxide dismutase 2, mitochondrial | 6648 | ENSG00000112096 |
| 7 | P61769 | NA | B2M | beta-2-microglobulin | 567 | ENSG00000166710 |
| 8 | P17096 | NA | HMGA1 | high mobility group AT-hook 1 | 3159 | ENSG00000137309 |
| 9 | Q16822 | NA | PCK2 | phosphoenolpyruvate carboxykinase 2 (mitochondrial) | 5106 | ENSG00000100889 |
| 10 | P02786 | NA | TFRC | transferrin receptor (p90, CD71) | 7037 | ENSG00000072274 |
| 11 | P21589 | NA | NT5E | 5'-nucleotidase, ecto (CD73) | 4907 | ENSG00000135318 |

  
  

| **Database:Pathway Commons pathway      &nbspName:LKB1 signaling events      &nbspID:DB\_ID:1649** | | | | | | |
| --- | --- | --- | --- | --- | --- | --- |
| C=1308; O=11; E=2.64; R=4.16; rawP=4.02e-05; adjP=5.54e-05 | | | | | | |
| Index | UserID | Value | Gene Symbol | Gene Name | EntrezGene | Ensembl |
| 1 | P17655 | NA | CAPN2 | calpain 2, (m/II) large subunit | 824 | ENSG00000162909 |
| 2 | P13639 | NA | EEF2 | eukaryotic translation elongation factor 2 | 1938 | ENSG00000167658 |
| 3 | P22570 | NA | FDXR | ferredoxin reductase | 2232 | ENSG00000161513 |
| 4 | P05556 | NA | ITGB1 | integrin, beta 1 (fibronectin receptor, beta polypeptide, antigen CD29 includes MDF2, MSK12) | 3688 | ENSG00000150093 |
| 5 | P55060 | NA | CSE1L | CSE1 chromosome segregation 1-like (yeast) | 1434 | ENSG00000124207 |
| 6 | P04179 | NA | SOD2 | superoxide dismutase 2, mitochondrial | 6648 | ENSG00000112096 |
| 7 | P61769 | NA | B2M | beta-2-microglobulin | 567 | ENSG00000166710 |
| 8 | P17096 | NA | HMGA1 | high mobility group AT-hook 1 | 3159 | ENSG00000137309 |
| 9 | Q16822 | NA | PCK2 | phosphoenolpyruvate carboxykinase 2 (mitochondrial) | 5106 | ENSG00000100889 |
| 10 | P02786 | NA | TFRC | transferrin receptor (p90, CD71) | 7037 | ENSG00000072274 |
| 11 | P21589 | NA | NT5E | 5'-nucleotidase, ecto (CD73) | 4907 | ENSG00000135318 |

  
  

| **Database:Pathway Commons pathway      &nbspName:PDGFR-beta signaling pathway      &nbspID:DB\_ID:1540** | | | | | | |
| --- | --- | --- | --- | --- | --- | --- |
| C=1288; O=11; E=2.60; R=4.22; rawP=3.49e-05; adjP=5.54e-05 | | | | | | |
| Index | UserID | Value | Gene Symbol | Gene Name | EntrezGene | Ensembl |
| 1 | P17655 | NA | CAPN2 | calpain 2, (m/II) large subunit | 824 | ENSG00000162909 |
| 2 | P13639 | NA | EEF2 | eukaryotic translation elongation factor 2 | 1938 | ENSG00000167658 |
| 3 | P22570 | NA | FDXR | ferredoxin reductase | 2232 | ENSG00000161513 |
| 4 | P05556 | NA | ITGB1 | integrin, beta 1 (fibronectin receptor, beta polypeptide, antigen CD29 includes MDF2, MSK12) | 3688 | ENSG00000150093 |
| 5 | P55060 | NA | CSE1L | CSE1 chromosome segregation 1-like (yeast) | 1434 | ENSG00000124207 |
| 6 | P04179 | NA | SOD2 | superoxide dismutase 2, mitochondrial | 6648 | ENSG00000112096 |
| 7 | P61769 | NA | B2M | beta-2-microglobulin | 567 | ENSG00000166710 |
| 8 | P17096 | NA | HMGA1 | high mobility group AT-hook 1 | 3159 | ENSG00000137309 |
| 9 | Q16822 | NA | PCK2 | phosphoenolpyruvate carboxykinase 2 (mitochondrial) | 5106 | ENSG00000100889 |
| 10 | P02786 | NA | TFRC | transferrin receptor (p90, CD71) | 7037 | ENSG00000072274 |
| 11 | P21589 | NA | NT5E | 5'-nucleotidase, ecto (CD73) | 4907 | ENSG00000135318 |

  
  

| **Database:Pathway Commons pathway      &nbspName:Class I PI3K signaling events      &nbspID:DB\_ID:1553** | | | | | | |
| --- | --- | --- | --- | --- | --- | --- |
| C=1288; O=11; E=2.60; R=4.22; rawP=3.49e-05; adjP=5.54e-05 | | | | | | |
| Index | UserID | Value | Gene Symbol | Gene Name | EntrezGene | Ensembl |
| 1 | P17655 | NA | CAPN2 | calpain 2, (m/II) large subunit | 824 | ENSG00000162909 |
| 2 | P13639 | NA | EEF2 | eukaryotic translation elongation factor 2 | 1938 | ENSG00000167658 |
| 3 | P22570 | NA | FDXR | ferredoxin reductase | 2232 | ENSG00000161513 |
| 4 | P05556 | NA | ITGB1 | integrin, beta 1 (fibronectin receptor, beta polypeptide, antigen CD29 includes MDF2, MSK12) | 3688 | ENSG00000150093 |
| 5 | P55060 | NA | CSE1L | CSE1 chromosome segregation 1-like (yeast) | 1434 | ENSG00000124207 |
| 6 | P04179 | NA | SOD2 | superoxide dismutase 2, mitochondrial | 6648 | ENSG00000112096 |
| 7 | P61769 | NA | B2M | beta-2-microglobulin | 567 | ENSG00000166710 |
| 8 | P17096 | NA | HMGA1 | high mobility group AT-hook 1 | 3159 | ENSG00000137309 |
| 9 | Q16822 | NA | PCK2 | phosphoenolpyruvate carboxykinase 2 (mitochondrial) | 5106 | ENSG00000100889 |
| 10 | P02786 | NA | TFRC | transferrin receptor (p90, CD71) | 7037 | ENSG00000072274 |
| 11 | P21589 | NA | NT5E | 5'-nucleotidase, ecto (CD73) | 4907 | ENSG00000135318 |

  
  

| **Database:Pathway Commons pathway      &nbspName:Glypican pathway      &nbspID:DB\_ID:1459** | | | | | | |
| --- | --- | --- | --- | --- | --- | --- |
| C=1338; O=11; E=2.70; R=4.07; rawP=4.95e-05; adjP=6.47e-05 | | | | | | |
| Index | UserID | Value | Gene Symbol | Gene Name | EntrezGene | Ensembl |
| 1 | P17655 | NA | CAPN2 | calpain 2, (m/II) large subunit | 824 | ENSG00000162909 |
| 2 | P13639 | NA | EEF2 | eukaryotic translation elongation factor 2 | 1938 | ENSG00000167658 |
| 3 | P22570 | NA | FDXR | ferredoxin reductase | 2232 | ENSG00000161513 |
| 4 | P05556 | NA | ITGB1 | integrin, beta 1 (fibronectin receptor, beta polypeptide, antigen CD29 includes MDF2, MSK12) | 3688 | ENSG00000150093 |
| 5 | P55060 | NA | CSE1L | CSE1 chromosome segregation 1-like (yeast) | 1434 | ENSG00000124207 |
| 6 | P04179 | NA | SOD2 | superoxide dismutase 2, mitochondrial | 6648 | ENSG00000112096 |
| 7 | P61769 | NA | B2M | beta-2-microglobulin | 567 | ENSG00000166710 |
| 8 | P17096 | NA | HMGA1 | high mobility group AT-hook 1 | 3159 | ENSG00000137309 |
| 9 | Q16822 | NA | PCK2 | phosphoenolpyruvate carboxykinase 2 (mitochondrial) | 5106 | ENSG00000100889 |
| 10 | P02786 | NA | TFRC | transferrin receptor (p90, CD71) | 7037 | ENSG00000072274 |
| 11 | P21589 | NA | NT5E | 5'-nucleotidase, ecto (CD73) | 4907 | ENSG00000135318 |

  
  

| **Database:Pathway Commons pathway      &nbspName:Proteoglycan syndecan-mediated signaling events      &nbspID:DB\_ID:1637** | | | | | | |
| --- | --- | --- | --- | --- | --- | --- |
| C=1345; O=11; E=2.72; R=4.05; rawP=5.19e-05; adjP=6.62e-05 | | | | | | |
| Index | UserID | Value | Gene Symbol | Gene Name | EntrezGene | Ensembl |
| 1 | P17655 | NA | CAPN2 | calpain 2, (m/II) large subunit | 824 | ENSG00000162909 |
| 2 | P13639 | NA | EEF2 | eukaryotic translation elongation factor 2 | 1938 | ENSG00000167658 |
| 3 | P22570 | NA | FDXR | ferredoxin reductase | 2232 | ENSG00000161513 |
| 4 | P05556 | NA | ITGB1 | integrin, beta 1 (fibronectin receptor, beta polypeptide, antigen CD29 includes MDF2, MSK12) | 3688 | ENSG00000150093 |
| 5 | P55060 | NA | CSE1L | CSE1 chromosome segregation 1-like (yeast) | 1434 | ENSG00000124207 |
| 6 | P04179 | NA | SOD2 | superoxide dismutase 2, mitochondrial | 6648 | ENSG00000112096 |
| 7 | P61769 | NA | B2M | beta-2-microglobulin | 567 | ENSG00000166710 |
| 8 | P17096 | NA | HMGA1 | high mobility group AT-hook 1 | 3159 | ENSG00000137309 |
| 9 | Q16822 | NA | PCK2 | phosphoenolpyruvate carboxykinase 2 (mitochondrial) | 5106 | ENSG00000100889 |
| 10 | P02786 | NA | TFRC | transferrin receptor (p90, CD71) | 7037 | ENSG00000072274 |
| 11 | P21589 | NA | NT5E | 5'-nucleotidase, ecto (CD73) | 4907 | ENSG00000135318 |

  
  

| **Database:Pathway Commons pathway      &nbspName:Activation of Chaperones by IRE1alpha      &nbspID:DB\_ID:411** | | | | | | |
| --- | --- | --- | --- | --- | --- | --- |
| C=45; O=3; E=0.09; R=32.98; rawP=0.0001; adjP=0.0001 | | | | | | |
| Index | UserID | Value | Gene Symbol | Gene Name | EntrezGene | Ensembl |
| 1 | O14773 | NA | TPP1 | tripeptidyl peptidase I | 1200 | ENSG00000166340 |
| 2 | P49748 | NA | ACADVL | acyl-CoA dehydrogenase, very long chain | 37 | ENSG00000072778 |
| 3 | P02545 | NA | LMNA | lamin A/C | 4000 | ENSG00000160789 |

  
  

| **Database:Pathway Commons pathway      &nbspName:Unfolded Protein Response      &nbspID:DB\_ID:409** | | | | | | |
| --- | --- | --- | --- | --- | --- | --- |
| C=63; O=3; E=0.13; R=23.56; rawP=0.0003; adjP=0.0004 | | | | | | |
| Index | UserID | Value | Gene Symbol | Gene Name | EntrezGene | Ensembl |
| 1 | O14773 | NA | TPP1 | tripeptidyl peptidase I | 1200 | ENSG00000166340 |
| 2 | P49748 | NA | ACADVL | acyl-CoA dehydrogenase, very long chain | 37 | ENSG00000072778 |
| 3 | P02545 | NA | LMNA | lamin A/C | 4000 | ENSG00000160789 |

  
  
  
  

---

WebGestalt is currently developed and maintained by Jing Wang and Bing Zhang at the  Zhang Lab. Other people who have made significant contribution to the project include Dexter Duncan, Stefan Kirov, Zhiao Shi, and Jay Snoddy.  
  
**Funding credits:** NIH/NIAAA (U01 AA016662, U01 AA013512); NIH/NIDA (P01 DA015027); NIH/NIMH (P50 MH078028, P50 MH096972); NIH/NCI (U24 CA159988); NIH/NIGMS (R01 GM088822).
