## Supplementary material for "PGRMC1 phosphorylation and cell plasticity 1: glycolysis, mitochondria, tumor growth": File S7: NO-WebGestalt Protein Inter...pdf

### WEB-based GEne SeT AnaLysis Toolkit

WebGestalt

*Translating gene lists into biological insights...*

**Your analysis is complete. Thank you for waiting.**

Analysis parameters: Data: 5_WT-v-TM.txt, Organism: hsapiens, Id Type: uniprot_swissprot_accession, Ref Set: entrezgene_protein-coding, Statistic: Hypergeometric, Signficance Level: .001, MTC: BH, Minimum: 3

There were either no significant results for your gene set of interest using the chosen options such as p-value, cufoff etc.. Or please be sure to choose the correct Id Type when you upload your User Id file. Then either upload a User Reference set and choose the proper Id Type for this Reference set or choose a Reference set from the gene set drop down list. The program does take the User Uploaded Reference set as priority over the gene set Reference set list just in case the user inadvertantly chooses a gene set Reference set and also uploads a User Reference Set. Please be sure to have valid User Ids and Reference sets and/or resubmit your analysis by chosing for example a higher p-value significance cutoff. Choosing Top10 as a signicance level should return results.

Close

WebGestalt is currently developed and maintained by Jing Wang and Bing Zhang at the Zhang Lab. Other people who have made significant contribution to the project include Dexter Duncan, Stefan Kirov, Zhiao Shi, and Jay Snoddy.

**Funding credits:** NIH/NIAAA (U01 AA016662, U01 AA013512); NIH/NIDA (P01 DA015027); NIH/NIMH (P50 MH078028, P50 MH096972); NIH/NCI (U24 CA159988); NIH/NIGMS (R01 GM088822).
