## Supplementary material for "PGRMC1 phosphorylation and cell plasticity 1: glycolysis, mitochondria, tumor growth": File S7: final_sig_file_1438753413.html

Anchored HTML File of EIDs

|  |  |
| --- | --- |
|  | WEB-based GEne SeT AnaLysis Toolkit |
|  | ***Translating gene lists into biological insights...*** |

---

  

| **Database:molecular function      &nbspName:oxidoreductase activity      &nbspID:GO:0016491** | | | | | | |
| --- | --- | --- | --- | --- | --- | --- |
| C=697; O=10; E=1.40; R=7.16; rawP=5.75e-07; adjP=1.55e-05 | | | | | | |
| Index | UserID | Value | Gene Symbol | Gene Name | EntrezGene | Ensembl |
| 1 | Q9Y2Q3 | NA | GSTK1 | glutathione S-transferase kappa 1 | 373156 | ENSG00000197448 |
| 2 | P02792 | NA | FTL | ferritin, light polypeptide | 2512 | ENSG00000087086 |
| 3 | P22570 | NA | FDXR | ferredoxin reductase | 2232 | ENSG00000161513 |
| 4 | P04179 | NA | SOD2 | superoxide dismutase 2, mitochondrial | 6648 | ENSG00000112096 |
| 5 | P48735 | NA | IDH2 | isocitrate dehydrogenase 2 (NADP+), mitochondrial | 3418 | ENSG00000182054 |
| 6 | P51659 | NA | HSD17B4 | hydroxysteroid (17-beta) dehydrogenase 4 | 3295 | ENSG00000133835 |
| 7 | P49748 | NA | ACADVL | acyl-CoA dehydrogenase, very long chain | 37 | ENSG00000072778 |
| 8 | P51648 | NA | ALDH3A2 | aldehyde dehydrogenase 3 family, member A2 | 224 | ENSG00000072210 |
| 9 | P32322 | NA | PYCR1 | pyrroline-5-carboxylate reductase 1 | 5831 | ENSG00000183010 |
| 10 | P31937 | NA | HIBADH | 3-hydroxyisobutyrate dehydrogenase | 11112 | ENSG00000106049 |

  
  

| **Database:molecular function      &nbspName:catalytic activity      &nbspID:GO:0003824** | | | | | | |
| --- | --- | --- | --- | --- | --- | --- |
| C=5299; O=22; E=10.61; R=2.07; rawP=2.53e-05; adjP=0.0003 | | | | | | |
| Index | UserID | Value | Gene Symbol | Gene Name | EntrezGene | Ensembl |
| 1 | P10253 | NA | GAA | glucosidase, alpha; acid | 2548 | ENSG00000171298 |
| 2 | P02792 | NA | FTL | ferritin, light polypeptide | 2512 | ENSG00000087086 |
| 3 | Q13724 | NA | MOGS | mannosyl-oligosaccharide glucosidase | 7841 | ENSG00000115275 |
| 4 | P04179 | NA | SOD2 | superoxide dismutase 2, mitochondrial | 6648 | ENSG00000112096 |
| 5 | Q8NFV4 | NA | ABHD11 | abhydrolase domain containing 11 | 83451 | ENSG00000106077 |
| 6 | P48735 | NA | IDH2 | isocitrate dehydrogenase 2 (NADP+), mitochondrial | 3418 | ENSG00000182054 |
| 7 | P51659 | NA | HSD17B4 | hydroxysteroid (17-beta) dehydrogenase 4 | 3295 | ENSG00000133835 |
| 8 | P49588 | NA | AARS | alanyl-tRNA synthetase | 16 | ENSG00000090861 |
| 9 | P43490 | NA | NAMPT | nicotinamide phosphoribosyltransferase | 10135 | ENSG00000105835 |
| 10 | P32322 | NA | PYCR1 | pyrroline-5-carboxylate reductase 1 | 5831 | ENSG00000183010 |
| 11 | P17655 | NA | CAPN2 | calpain 2, (m/II) large subunit | 824 | ENSG00000162909 |
| 12 | O14773 | NA | TPP1 | tripeptidyl peptidase I | 1200 | ENSG00000166340 |
| 13 | Q9Y2Q3 | NA | GSTK1 | glutathione S-transferase kappa 1 | 373156 | ENSG00000197448 |
| 14 | P51688 | NA | SGSH | N-sulfoglucosamine sulfohydrolase | 6448 | ENSG00000181523 |
| 15 | P22570 | NA | FDXR | ferredoxin reductase | 2232 | ENSG00000161513 |
| 16 | O75521 | NA | ECI2 | enoyl-CoA delta isomerase 2 | 10455 | ENSG00000198721 |
| 17 | P17096 | NA | HMGA1 | high mobility group AT-hook 1 | 3159 | ENSG00000137309 |
| 18 | Q16822 | NA | PCK2 | phosphoenolpyruvate carboxykinase 2 (mitochondrial) | 5106 | ENSG00000100889 |
| 19 | P49748 | NA | ACADVL | acyl-CoA dehydrogenase, very long chain | 37 | ENSG00000072778 |
| 20 | P51648 | NA | ALDH3A2 | aldehyde dehydrogenase 3 family, member A2 | 224 | ENSG00000072210 |
| 21 | P21589 | NA | NT5E | 5'-nucleotidase, ecto (CD73) | 4907 | ENSG00000135318 |
| 22 | P31937 | NA | HIBADH | 3-hydroxyisobutyrate dehydrogenase | 11112 | ENSG00000106049 |

  
  

| **Database:molecular function      &nbspName:oxidoreductase activity, acting on CH-OH group of donors      &nbspID:GO:0016614** | | | | | | |
| --- | --- | --- | --- | --- | --- | --- |
| C=128; O=4; E=0.26; R=15.60; rawP=0.0001; adjP=0.0009 | | | | | | |
| Index | UserID | Value | Gene Symbol | Gene Name | EntrezGene | Ensembl |
| 1 | P48735 | NA | IDH2 | isocitrate dehydrogenase 2 (NADP+), mitochondrial | 3418 | ENSG00000182054 |
| 2 | P51659 | NA | HSD17B4 | hydroxysteroid (17-beta) dehydrogenase 4 | 3295 | ENSG00000133835 |
| 3 | P51648 | NA | ALDH3A2 | aldehyde dehydrogenase 3 family, member A2 | 224 | ENSG00000072210 |
| 4 | P31937 | NA | HIBADH | 3-hydroxyisobutyrate dehydrogenase | 11112 | ENSG00000106049 |

  
  

| **Database:cellular component      &nbspName:mitochondrial matrix      &nbspID:GO:0005759** | | | | | | |
| --- | --- | --- | --- | --- | --- | --- |
| C=278; O=9; E=0.52; R=17.42; rawP=1.27e-09; adjP=7.11e-08 | | | | | | |
| Index | UserID | Value | Gene Symbol | Gene Name | EntrezGene | Ensembl |
| 1 | P22570 | NA | FDXR | ferredoxin reductase | 2232 | ENSG00000161513 |
| 2 | P04179 | NA | SOD2 | superoxide dismutase 2, mitochondrial | 6648 | ENSG00000112096 |
| 3 | P48735 | NA | IDH2 | isocitrate dehydrogenase 2 (NADP+), mitochondrial | 3418 | ENSG00000182054 |
| 4 | Q04837 | NA | SSBP1 | single-stranded DNA binding protein 1, mitochondrial | 6742 | ENSG00000106028 |
| 5 | P21796 | NA | VDAC1 | voltage-dependent anion channel 1 | 7416 | ENSG00000213585 |
| 6 | Q16822 | NA | PCK2 | phosphoenolpyruvate carboxykinase 2 (mitochondrial) | 5106 | ENSG00000100889 |
| 7 | P49748 | NA | ACADVL | acyl-CoA dehydrogenase, very long chain | 37 | ENSG00000072778 |
| 8 | P32322 | NA | PYCR1 | pyrroline-5-carboxylate reductase 1 | 5831 | ENSG00000183010 |
| 9 | P31937 | NA | HIBADH | 3-hydroxyisobutyrate dehydrogenase | 11112 | ENSG00000106049 |

  
  

| **Database:cellular component      &nbspName:cytoplasm      &nbspID:GO:0005737** | | | | | | |
| --- | --- | --- | --- | --- | --- | --- |
| C=9051; O=31; E=16.82; R=1.84; rawP=5.72e-09; adjP=1.60e-07 | | | | | | |
| Index | UserID | Value | Gene Symbol | Gene Name | EntrezGene | Ensembl |
| 1 | P10253 | NA | GAA | glucosidase, alpha; acid | 2548 | ENSG00000171298 |
| 2 | P02792 | NA | FTL | ferritin, light polypeptide | 2512 | ENSG00000087086 |
| 3 | P55060 | NA | CSE1L | CSE1 chromosome segregation 1-like (yeast) | 1434 | ENSG00000124207 |
| 4 | Q13724 | NA | MOGS | mannosyl-oligosaccharide glucosidase | 7841 | ENSG00000115275 |
| 5 | P04179 | NA | SOD2 | superoxide dismutase 2, mitochondrial | 6648 | ENSG00000112096 |
| 6 | P61769 | NA | B2M | beta-2-microglobulin | 567 | ENSG00000166710 |
| 7 | Q8NFV4 | NA | ABHD11 | abhydrolase domain containing 11 | 83451 | ENSG00000106077 |
| 8 | P48735 | NA | IDH2 | isocitrate dehydrogenase 2 (NADP+), mitochondrial | 3418 | ENSG00000182054 |
| 9 | P51659 | NA | HSD17B4 | hydroxysteroid (17-beta) dehydrogenase 4 | 3295 | ENSG00000133835 |
| 10 | P21796 | NA | VDAC1 | voltage-dependent anion channel 1 | 7416 | ENSG00000213585 |
| 11 | Q86UP2 | NA | KTN1 | kinectin 1 (kinesin receptor) | 3895 | ENSG00000126777 |
| 12 | P49588 | NA | AARS | alanyl-tRNA synthetase | 16 | ENSG00000090861 |
| 13 | P02545 | NA | LMNA | lamin A/C | 4000 | ENSG00000160789 |
| 14 | P43490 | NA | NAMPT | nicotinamide phosphoribosyltransferase | 10135 | ENSG00000105835 |
| 15 | P32322 | NA | PYCR1 | pyrroline-5-carboxylate reductase 1 | 5831 | ENSG00000183010 |
| 16 | P17655 | NA | CAPN2 | calpain 2, (m/II) large subunit | 824 | ENSG00000162909 |
| 17 | O14773 | NA | TPP1 | tripeptidyl peptidase I | 1200 | ENSG00000166340 |
| 18 | Q9Y2Q3 | NA | GSTK1 | glutathione S-transferase kappa 1 | 373156 | ENSG00000197448 |
| 19 | P51688 | NA | SGSH | N-sulfoglucosamine sulfohydrolase | 6448 | ENSG00000181523 |
| 20 | P22570 | NA | FDXR | ferredoxin reductase | 2232 | ENSG00000161513 |
| 21 | P05556 | NA | ITGB1 | integrin, beta 1 (fibronectin receptor, beta polypeptide, antigen CD29 includes MDF2, MSK12) | 3688 | ENSG00000150093 |
| 22 | O75521 | NA | ECI2 | enoyl-CoA delta isomerase 2 | 10455 | ENSG00000198721 |
| 23 | Q04837 | NA | SSBP1 | single-stranded DNA binding protein 1, mitochondrial | 6742 | ENSG00000106028 |
| 24 | P17096 | NA | HMGA1 | high mobility group AT-hook 1 | 3159 | ENSG00000137309 |
| 25 | Q16822 | NA | PCK2 | phosphoenolpyruvate carboxykinase 2 (mitochondrial) | 5106 | ENSG00000100889 |
| 26 | P49748 | NA | ACADVL | acyl-CoA dehydrogenase, very long chain | 37 | ENSG00000072778 |
| 27 | Q9Y3E0 | NA | GOLT1B | golgi transport 1B | 51026 | ENSG00000111711 |
| 28 | P51648 | NA | ALDH3A2 | aldehyde dehydrogenase 3 family, member A2 | 224 | ENSG00000072210 |
| 29 | P30042 | NA | C21orf33 | chromosome 21 open reading frame 33 | 8209 | ENSG00000160221 |
| 30 | P21589 | NA | NT5E | 5'-nucleotidase, ecto (CD73) | 4907 | ENSG00000135318 |
| 31 | P31937 | NA | HIBADH | 3-hydroxyisobutyrate dehydrogenase | 11112 | ENSG00000106049 |

  
  

| **Database:cellular component      &nbspName:cytoplasmic part      &nbspID:GO:0044444** | | | | | | |
| --- | --- | --- | --- | --- | --- | --- |
| C=6728; O=28; E=12.50; R=2.24; rawP=9.03e-09; adjP=1.69e-07 | | | | | | |
| Index | UserID | Value | Gene Symbol | Gene Name | EntrezGene | Ensembl |
| 1 | P10253 | NA | GAA | glucosidase, alpha; acid | 2548 | ENSG00000171298 |
| 2 | P02792 | NA | FTL | ferritin, light polypeptide | 2512 | ENSG00000087086 |
| 3 | Q13724 | NA | MOGS | mannosyl-oligosaccharide glucosidase | 7841 | ENSG00000115275 |
| 4 | P04179 | NA | SOD2 | superoxide dismutase 2, mitochondrial | 6648 | ENSG00000112096 |
| 5 | P61769 | NA | B2M | beta-2-microglobulin | 567 | ENSG00000166710 |
| 6 | Q8NFV4 | NA | ABHD11 | abhydrolase domain containing 11 | 83451 | ENSG00000106077 |
| 7 | P48735 | NA | IDH2 | isocitrate dehydrogenase 2 (NADP+), mitochondrial | 3418 | ENSG00000182054 |
| 8 | P51659 | NA | HSD17B4 | hydroxysteroid (17-beta) dehydrogenase 4 | 3295 | ENSG00000133835 |
| 9 | P21796 | NA | VDAC1 | voltage-dependent anion channel 1 | 7416 | ENSG00000213585 |
| 10 | P49588 | NA | AARS | alanyl-tRNA synthetase | 16 | ENSG00000090861 |
| 11 | Q86UP2 | NA | KTN1 | kinectin 1 (kinesin receptor) | 3895 | ENSG00000126777 |
| 12 | P02545 | NA | LMNA | lamin A/C | 4000 | ENSG00000160789 |
| 13 | P43490 | NA | NAMPT | nicotinamide phosphoribosyltransferase | 10135 | ENSG00000105835 |
| 14 | P32322 | NA | PYCR1 | pyrroline-5-carboxylate reductase 1 | 5831 | ENSG00000183010 |
| 15 | O14773 | NA | TPP1 | tripeptidyl peptidase I | 1200 | ENSG00000166340 |
| 16 | Q9Y2Q3 | NA | GSTK1 | glutathione S-transferase kappa 1 | 373156 | ENSG00000197448 |
| 17 | P51688 | NA | SGSH | N-sulfoglucosamine sulfohydrolase | 6448 | ENSG00000181523 |
| 18 | P22570 | NA | FDXR | ferredoxin reductase | 2232 | ENSG00000161513 |
| 19 | P05556 | NA | ITGB1 | integrin, beta 1 (fibronectin receptor, beta polypeptide, antigen CD29 includes MDF2, MSK12) | 3688 | ENSG00000150093 |
| 20 | O75521 | NA | ECI2 | enoyl-CoA delta isomerase 2 | 10455 | ENSG00000198721 |
| 21 | Q04837 | NA | SSBP1 | single-stranded DNA binding protein 1, mitochondrial | 6742 | ENSG00000106028 |
| 22 | P17096 | NA | HMGA1 | high mobility group AT-hook 1 | 3159 | ENSG00000137309 |
| 23 | Q16822 | NA | PCK2 | phosphoenolpyruvate carboxykinase 2 (mitochondrial) | 5106 | ENSG00000100889 |
| 24 | P49748 | NA | ACADVL | acyl-CoA dehydrogenase, very long chain | 37 | ENSG00000072778 |
| 25 | Q9Y3E0 | NA | GOLT1B | golgi transport 1B | 51026 | ENSG00000111711 |
| 26 | P51648 | NA | ALDH3A2 | aldehyde dehydrogenase 3 family, member A2 | 224 | ENSG00000072210 |
| 27 | P30042 | NA | C21orf33 | chromosome 21 open reading frame 33 | 8209 | ENSG00000160221 |
| 28 | P31937 | NA | HIBADH | 3-hydroxyisobutyrate dehydrogenase | 11112 | ENSG00000106049 |

  
  

| **Database:cellular component      &nbspName:mitochondrion      &nbspID:GO:0005739** | | | | | | |
| --- | --- | --- | --- | --- | --- | --- |
| C=1515; O=14; E=2.82; R=4.97; rawP=1.47e-07; adjP=2.06e-06 | | | | | | |
| Index | UserID | Value | Gene Symbol | Gene Name | EntrezGene | Ensembl |
| 1 | O14773 | NA | TPP1 | tripeptidyl peptidase I | 1200 | ENSG00000166340 |
| 2 | Q9Y2Q3 | NA | GSTK1 | glutathione S-transferase kappa 1 | 373156 | ENSG00000197448 |
| 3 | P22570 | NA | FDXR | ferredoxin reductase | 2232 | ENSG00000161513 |
| 4 | P04179 | NA | SOD2 | superoxide dismutase 2, mitochondrial | 6648 | ENSG00000112096 |
| 5 | O75521 | NA | ECI2 | enoyl-CoA delta isomerase 2 | 10455 | ENSG00000198721 |
| 6 | Q8NFV4 | NA | ABHD11 | abhydrolase domain containing 11 | 83451 | ENSG00000106077 |
| 7 | P48735 | NA | IDH2 | isocitrate dehydrogenase 2 (NADP+), mitochondrial | 3418 | ENSG00000182054 |
| 8 | Q04837 | NA | SSBP1 | single-stranded DNA binding protein 1, mitochondrial | 6742 | ENSG00000106028 |
| 9 | Q16822 | NA | PCK2 | phosphoenolpyruvate carboxykinase 2 (mitochondrial) | 5106 | ENSG00000100889 |
| 10 | P21796 | NA | VDAC1 | voltage-dependent anion channel 1 | 7416 | ENSG00000213585 |
| 11 | P49748 | NA | ACADVL | acyl-CoA dehydrogenase, very long chain | 37 | ENSG00000072778 |
| 12 | P30042 | NA | C21orf33 | chromosome 21 open reading frame 33 | 8209 | ENSG00000160221 |
| 13 | P31937 | NA | HIBADH | 3-hydroxyisobutyrate dehydrogenase | 11112 | ENSG00000106049 |
| 14 | P32322 | NA | PYCR1 | pyrroline-5-carboxylate reductase 1 | 5831 | ENSG00000183010 |

  
  

| **Database:cellular component      &nbspName:intracellular part      &nbspID:GO:0044424** | | | | | | |
| --- | --- | --- | --- | --- | --- | --- |
| C=12096; O=31; E=22.48; R=1.38; rawP=4.65e-05; adjP=0.0003 | | | | | | |
| Index | UserID | Value | Gene Symbol | Gene Name | EntrezGene | Ensembl |
| 1 | P10253 | NA | GAA | glucosidase, alpha; acid | 2548 | ENSG00000171298 |
| 2 | P02792 | NA | FTL | ferritin, light polypeptide | 2512 | ENSG00000087086 |
| 3 | P55060 | NA | CSE1L | CSE1 chromosome segregation 1-like (yeast) | 1434 | ENSG00000124207 |
| 4 | Q13724 | NA | MOGS | mannosyl-oligosaccharide glucosidase | 7841 | ENSG00000115275 |
| 5 | P04179 | NA | SOD2 | superoxide dismutase 2, mitochondrial | 6648 | ENSG00000112096 |
| 6 | P61769 | NA | B2M | beta-2-microglobulin | 567 | ENSG00000166710 |
| 7 | Q8NFV4 | NA | ABHD11 | abhydrolase domain containing 11 | 83451 | ENSG00000106077 |
| 8 | P48735 | NA | IDH2 | isocitrate dehydrogenase 2 (NADP+), mitochondrial | 3418 | ENSG00000182054 |
| 9 | P51659 | NA | HSD17B4 | hydroxysteroid (17-beta) dehydrogenase 4 | 3295 | ENSG00000133835 |
| 10 | P21796 | NA | VDAC1 | voltage-dependent anion channel 1 | 7416 | ENSG00000213585 |
| 11 | Q86UP2 | NA | KTN1 | kinectin 1 (kinesin receptor) | 3895 | ENSG00000126777 |
| 12 | P49588 | NA | AARS | alanyl-tRNA synthetase | 16 | ENSG00000090861 |
| 13 | P02545 | NA | LMNA | lamin A/C | 4000 | ENSG00000160789 |
| 14 | P43490 | NA | NAMPT | nicotinamide phosphoribosyltransferase | 10135 | ENSG00000105835 |
| 15 | P32322 | NA | PYCR1 | pyrroline-5-carboxylate reductase 1 | 5831 | ENSG00000183010 |
| 16 | P17655 | NA | CAPN2 | calpain 2, (m/II) large subunit | 824 | ENSG00000162909 |
| 17 | O14773 | NA | TPP1 | tripeptidyl peptidase I | 1200 | ENSG00000166340 |
| 18 | Q9Y2Q3 | NA | GSTK1 | glutathione S-transferase kappa 1 | 373156 | ENSG00000197448 |
| 19 | P51688 | NA | SGSH | N-sulfoglucosamine sulfohydrolase | 6448 | ENSG00000181523 |
| 20 | P22570 | NA | FDXR | ferredoxin reductase | 2232 | ENSG00000161513 |
| 21 | P05556 | NA | ITGB1 | integrin, beta 1 (fibronectin receptor, beta polypeptide, antigen CD29 includes MDF2, MSK12) | 3688 | ENSG00000150093 |
| 22 | O75521 | NA | ECI2 | enoyl-CoA delta isomerase 2 | 10455 | ENSG00000198721 |
| 23 | Q04837 | NA | SSBP1 | single-stranded DNA binding protein 1, mitochondrial | 6742 | ENSG00000106028 |
| 24 | P17096 | NA | HMGA1 | high mobility group AT-hook 1 | 3159 | ENSG00000137309 |
| 25 | Q16822 | NA | PCK2 | phosphoenolpyruvate carboxykinase 2 (mitochondrial) | 5106 | ENSG00000100889 |
| 26 | P49748 | NA | ACADVL | acyl-CoA dehydrogenase, very long chain | 37 | ENSG00000072778 |
| 27 | Q9Y3E0 | NA | GOLT1B | golgi transport 1B | 51026 | ENSG00000111711 |
| 28 | P51648 | NA | ALDH3A2 | aldehyde dehydrogenase 3 family, member A2 | 224 | ENSG00000072210 |
| 29 | P30042 | NA | C21orf33 | chromosome 21 open reading frame 33 | 8209 | ENSG00000160221 |
| 30 | P21589 | NA | NT5E | 5'-nucleotidase, ecto (CD73) | 4907 | ENSG00000135318 |
| 31 | P31937 | NA | HIBADH | 3-hydroxyisobutyrate dehydrogenase | 11112 | ENSG00000106049 |

  
  

| **Database:cellular component      &nbspName:organelle lumen      &nbspID:GO:0043233** | | | | | | |
| --- | --- | --- | --- | --- | --- | --- |
| C=3331; O=16; E=6.19; R=2.58; rawP=8.49e-05; adjP=0.0004 | | | | | | |
| Index | UserID | Value | Gene Symbol | Gene Name | EntrezGene | Ensembl |
| 1 | P04179 | NA | SOD2 | superoxide dismutase 2, mitochondrial | 6648 | ENSG00000112096 |
| 2 | P61769 | NA | B2M | beta-2-microglobulin | 567 | ENSG00000166710 |
| 3 | P48735 | NA | IDH2 | isocitrate dehydrogenase 2 (NADP+), mitochondrial | 3418 | ENSG00000182054 |
| 4 | P51659 | NA | HSD17B4 | hydroxysteroid (17-beta) dehydrogenase 4 | 3295 | ENSG00000133835 |
| 5 | P21796 | NA | VDAC1 | voltage-dependent anion channel 1 | 7416 | ENSG00000213585 |
| 6 | P02545 | NA | LMNA | lamin A/C | 4000 | ENSG00000160789 |
| 7 | P32322 | NA | PYCR1 | pyrroline-5-carboxylate reductase 1 | 5831 | ENSG00000183010 |
| 8 | O14773 | NA | TPP1 | tripeptidyl peptidase I | 1200 | ENSG00000166340 |
| 9 | P51688 | NA | SGSH | N-sulfoglucosamine sulfohydrolase | 6448 | ENSG00000181523 |
| 10 | P22570 | NA | FDXR | ferredoxin reductase | 2232 | ENSG00000161513 |
| 11 | O75521 | NA | ECI2 | enoyl-CoA delta isomerase 2 | 10455 | ENSG00000198721 |
| 12 | P17096 | NA | HMGA1 | high mobility group AT-hook 1 | 3159 | ENSG00000137309 |
| 13 | Q04837 | NA | SSBP1 | single-stranded DNA binding protein 1, mitochondrial | 6742 | ENSG00000106028 |
| 14 | Q16822 | NA | PCK2 | phosphoenolpyruvate carboxykinase 2 (mitochondrial) | 5106 | ENSG00000100889 |
| 15 | P49748 | NA | ACADVL | acyl-CoA dehydrogenase, very long chain | 37 | ENSG00000072778 |
| 16 | P31937 | NA | HIBADH | 3-hydroxyisobutyrate dehydrogenase | 11112 | ENSG00000106049 |

  
  

| **Database:cellular component      &nbspName:microbody      &nbspID:GO:0042579** | | | | | | |
| --- | --- | --- | --- | --- | --- | --- |
| C=122; O=4; E=0.23; R=17.64; rawP=7.35e-05; adjP=0.0004 | | | | | | |
| Index | UserID | Value | Gene Symbol | Gene Name | EntrezGene | Ensembl |
| 1 | O75521 | NA | ECI2 | enoyl-CoA delta isomerase 2 | 10455 | ENSG00000198721 |
| 2 | Q9Y2Q3 | NA | GSTK1 | glutathione S-transferase kappa 1 | 373156 | ENSG00000197448 |
| 3 | P51659 | NA | HSD17B4 | hydroxysteroid (17-beta) dehydrogenase 4 | 3295 | ENSG00000133835 |
| 4 | P51648 | NA | ALDH3A2 | aldehyde dehydrogenase 3 family, member A2 | 224 | ENSG00000072210 |

  
  

| **Database:cellular component      &nbspName:membrane-enclosed lumen      &nbspID:GO:0031974** | | | | | | |
| --- | --- | --- | --- | --- | --- | --- |
| C=3375; O=16; E=6.27; R=2.55; rawP=0.0001; adjP=0.0004 | | | | | | |
| Index | UserID | Value | Gene Symbol | Gene Name | EntrezGene | Ensembl |
| 1 | P04179 | NA | SOD2 | superoxide dismutase 2, mitochondrial | 6648 | ENSG00000112096 |
| 2 | P61769 | NA | B2M | beta-2-microglobulin | 567 | ENSG00000166710 |
| 3 | P48735 | NA | IDH2 | isocitrate dehydrogenase 2 (NADP+), mitochondrial | 3418 | ENSG00000182054 |
| 4 | P51659 | NA | HSD17B4 | hydroxysteroid (17-beta) dehydrogenase 4 | 3295 | ENSG00000133835 |
| 5 | P21796 | NA | VDAC1 | voltage-dependent anion channel 1 | 7416 | ENSG00000213585 |
| 6 | P02545 | NA | LMNA | lamin A/C | 4000 | ENSG00000160789 |
| 7 | P32322 | NA | PYCR1 | pyrroline-5-carboxylate reductase 1 | 5831 | ENSG00000183010 |
| 8 | O14773 | NA | TPP1 | tripeptidyl peptidase I | 1200 | ENSG00000166340 |
| 9 | P51688 | NA | SGSH | N-sulfoglucosamine sulfohydrolase | 6448 | ENSG00000181523 |
| 10 | P22570 | NA | FDXR | ferredoxin reductase | 2232 | ENSG00000161513 |
| 11 | O75521 | NA | ECI2 | enoyl-CoA delta isomerase 2 | 10455 | ENSG00000198721 |
| 12 | P17096 | NA | HMGA1 | high mobility group AT-hook 1 | 3159 | ENSG00000137309 |
| 13 | Q04837 | NA | SSBP1 | single-stranded DNA binding protein 1, mitochondrial | 6742 | ENSG00000106028 |
| 14 | Q16822 | NA | PCK2 | phosphoenolpyruvate carboxykinase 2 (mitochondrial) | 5106 | ENSG00000100889 |
| 15 | P49748 | NA | ACADVL | acyl-CoA dehydrogenase, very long chain | 37 | ENSG00000072778 |
| 16 | P31937 | NA | HIBADH | 3-hydroxyisobutyrate dehydrogenase | 11112 | ENSG00000106049 |

  
  

| **Database:cellular component      &nbspName:peroxisome      &nbspID:GO:0005777** | | | | | | |
| --- | --- | --- | --- | --- | --- | --- |
| C=122; O=4; E=0.23; R=17.64; rawP=7.35e-05; adjP=0.0004 | | | | | | |
| Index | UserID | Value | Gene Symbol | Gene Name | EntrezGene | Ensembl |
| 1 | O75521 | NA | ECI2 | enoyl-CoA delta isomerase 2 | 10455 | ENSG00000198721 |
| 2 | Q9Y2Q3 | NA | GSTK1 | glutathione S-transferase kappa 1 | 373156 | ENSG00000197448 |
| 3 | P51659 | NA | HSD17B4 | hydroxysteroid (17-beta) dehydrogenase 4 | 3295 | ENSG00000133835 |
| 4 | P51648 | NA | ALDH3A2 | aldehyde dehydrogenase 3 family, member A2 | 224 | ENSG00000072210 |

  
  

| **Database:cellular component      &nbspName:intracellular      &nbspID:GO:0005622** | | | | | | |
| --- | --- | --- | --- | --- | --- | --- |
| C=12412; O=31; E=23.07; R=1.34; rawP=0.0001; adjP=0.0004 | | | | | | |
| Index | UserID | Value | Gene Symbol | Gene Name | EntrezGene | Ensembl |
| 1 | P10253 | NA | GAA | glucosidase, alpha; acid | 2548 | ENSG00000171298 |
| 2 | P02792 | NA | FTL | ferritin, light polypeptide | 2512 | ENSG00000087086 |
| 3 | P55060 | NA | CSE1L | CSE1 chromosome segregation 1-like (yeast) | 1434 | ENSG00000124207 |
| 4 | Q13724 | NA | MOGS | mannosyl-oligosaccharide glucosidase | 7841 | ENSG00000115275 |
| 5 | P04179 | NA | SOD2 | superoxide dismutase 2, mitochondrial | 6648 | ENSG00000112096 |
| 6 | P61769 | NA | B2M | beta-2-microglobulin | 567 | ENSG00000166710 |
| 7 | Q8NFV4 | NA | ABHD11 | abhydrolase domain containing 11 | 83451 | ENSG00000106077 |
| 8 | P48735 | NA | IDH2 | isocitrate dehydrogenase 2 (NADP+), mitochondrial | 3418 | ENSG00000182054 |
| 9 | P51659 | NA | HSD17B4 | hydroxysteroid (17-beta) dehydrogenase 4 | 3295 | ENSG00000133835 |
| 10 | P21796 | NA | VDAC1 | voltage-dependent anion channel 1 | 7416 | ENSG00000213585 |
| 11 | Q86UP2 | NA | KTN1 | kinectin 1 (kinesin receptor) | 3895 | ENSG00000126777 |
| 12 | P49588 | NA | AARS | alanyl-tRNA synthetase | 16 | ENSG00000090861 |
| 13 | P02545 | NA | LMNA | lamin A/C | 4000 | ENSG00000160789 |
| 14 | P43490 | NA | NAMPT | nicotinamide phosphoribosyltransferase | 10135 | ENSG00000105835 |
| 15 | P32322 | NA | PYCR1 | pyrroline-5-carboxylate reductase 1 | 5831 | ENSG00000183010 |
| 16 | P17655 | NA | CAPN2 | calpain 2, (m/II) large subunit | 824 | ENSG00000162909 |
| 17 | O14773 | NA | TPP1 | tripeptidyl peptidase I | 1200 | ENSG00000166340 |
| 18 | Q9Y2Q3 | NA | GSTK1 | glutathione S-transferase kappa 1 | 373156 | ENSG00000197448 |
| 19 | P51688 | NA | SGSH | N-sulfoglucosamine sulfohydrolase | 6448 | ENSG00000181523 |
| 20 | P22570 | NA | FDXR | ferredoxin reductase | 2232 | ENSG00000161513 |
| 21 | P05556 | NA | ITGB1 | integrin, beta 1 (fibronectin receptor, beta polypeptide, antigen CD29 includes MDF2, MSK12) | 3688 | ENSG00000150093 |
| 22 | O75521 | NA | ECI2 | enoyl-CoA delta isomerase 2 | 10455 | ENSG00000198721 |
| 23 | Q04837 | NA | SSBP1 | single-stranded DNA binding protein 1, mitochondrial | 6742 | ENSG00000106028 |
| 24 | P17096 | NA | HMGA1 | high mobility group AT-hook 1 | 3159 | ENSG00000137309 |
| 25 | Q16822 | NA | PCK2 | phosphoenolpyruvate carboxykinase 2 (mitochondrial) | 5106 | ENSG00000100889 |
| 26 | P49748 | NA | ACADVL | acyl-CoA dehydrogenase, very long chain | 37 | ENSG00000072778 |
| 27 | Q9Y3E0 | NA | GOLT1B | golgi transport 1B | 51026 | ENSG00000111711 |
| 28 | P51648 | NA | ALDH3A2 | aldehyde dehydrogenase 3 family, member A2 | 224 | ENSG00000072210 |
| 29 | P30042 | NA | C21orf33 | chromosome 21 open reading frame 33 | 8209 | ENSG00000160221 |
| 30 | P21589 | NA | NT5E | 5'-nucleotidase, ecto (CD73) | 4907 | ENSG00000135318 |
| 31 | P31937 | NA | HIBADH | 3-hydroxyisobutyrate dehydrogenase | 11112 | ENSG00000106049 |

  
  

| **Database:cellular component      &nbspName:intracellular organelle lumen      &nbspID:GO:0070013** | | | | | | |
| --- | --- | --- | --- | --- | --- | --- |
| C=3285; O=16; E=6.10; R=2.62; rawP=7.12e-05; adjP=0.0004 | | | | | | |
| Index | UserID | Value | Gene Symbol | Gene Name | EntrezGene | Ensembl |
| 1 | P04179 | NA | SOD2 | superoxide dismutase 2, mitochondrial | 6648 | ENSG00000112096 |
| 2 | P61769 | NA | B2M | beta-2-microglobulin | 567 | ENSG00000166710 |
| 3 | P48735 | NA | IDH2 | isocitrate dehydrogenase 2 (NADP+), mitochondrial | 3418 | ENSG00000182054 |
| 4 | P51659 | NA | HSD17B4 | hydroxysteroid (17-beta) dehydrogenase 4 | 3295 | ENSG00000133835 |
| 5 | P21796 | NA | VDAC1 | voltage-dependent anion channel 1 | 7416 | ENSG00000213585 |
| 6 | P02545 | NA | LMNA | lamin A/C | 4000 | ENSG00000160789 |
| 7 | P32322 | NA | PYCR1 | pyrroline-5-carboxylate reductase 1 | 5831 | ENSG00000183010 |
| 8 | O14773 | NA | TPP1 | tripeptidyl peptidase I | 1200 | ENSG00000166340 |
| 9 | P51688 | NA | SGSH | N-sulfoglucosamine sulfohydrolase | 6448 | ENSG00000181523 |
| 10 | P22570 | NA | FDXR | ferredoxin reductase | 2232 | ENSG00000161513 |
| 11 | O75521 | NA | ECI2 | enoyl-CoA delta isomerase 2 | 10455 | ENSG00000198721 |
| 12 | P17096 | NA | HMGA1 | high mobility group AT-hook 1 | 3159 | ENSG00000137309 |
| 13 | Q04837 | NA | SSBP1 | single-stranded DNA binding protein 1, mitochondrial | 6742 | ENSG00000106028 |
| 14 | Q16822 | NA | PCK2 | phosphoenolpyruvate carboxykinase 2 (mitochondrial) | 5106 | ENSG00000100889 |
| 15 | P49748 | NA | ACADVL | acyl-CoA dehydrogenase, very long chain | 37 | ENSG00000072778 |
| 16 | P31937 | NA | HIBADH | 3-hydroxyisobutyrate dehydrogenase | 11112 | ENSG00000106049 |

  
  

| **Database:cellular component      &nbspName:intracellular organelle part      &nbspID:GO:0044446** | | | | | | |
| --- | --- | --- | --- | --- | --- | --- |
| C=6690; O=23; E=12.43; R=1.85; rawP=0.0001; adjP=0.0004 | | | | | | |
| Index | UserID | Value | Gene Symbol | Gene Name | EntrezGene | Ensembl |
| 1 | P10253 | NA | GAA | glucosidase, alpha; acid | 2548 | ENSG00000171298 |
| 2 | Q13724 | NA | MOGS | mannosyl-oligosaccharide glucosidase | 7841 | ENSG00000115275 |
| 3 | P04179 | NA | SOD2 | superoxide dismutase 2, mitochondrial | 6648 | ENSG00000112096 |
| 4 | P61769 | NA | B2M | beta-2-microglobulin | 567 | ENSG00000166710 |
| 5 | P48735 | NA | IDH2 | isocitrate dehydrogenase 2 (NADP+), mitochondrial | 3418 | ENSG00000182054 |
| 6 | P51659 | NA | HSD17B4 | hydroxysteroid (17-beta) dehydrogenase 4 | 3295 | ENSG00000133835 |
| 7 | P21796 | NA | VDAC1 | voltage-dependent anion channel 1 | 7416 | ENSG00000213585 |
| 8 | Q86UP2 | NA | KTN1 | kinectin 1 (kinesin receptor) | 3895 | ENSG00000126777 |
| 9 | P02545 | NA | LMNA | lamin A/C | 4000 | ENSG00000160789 |
| 10 | P32322 | NA | PYCR1 | pyrroline-5-carboxylate reductase 1 | 5831 | ENSG00000183010 |
| 11 | P17655 | NA | CAPN2 | calpain 2, (m/II) large subunit | 824 | ENSG00000162909 |
| 12 | O14773 | NA | TPP1 | tripeptidyl peptidase I | 1200 | ENSG00000166340 |
| 13 | Q9Y2Q3 | NA | GSTK1 | glutathione S-transferase kappa 1 | 373156 | ENSG00000197448 |
| 14 | P51688 | NA | SGSH | N-sulfoglucosamine sulfohydrolase | 6448 | ENSG00000181523 |
| 15 | P22570 | NA | FDXR | ferredoxin reductase | 2232 | ENSG00000161513 |
| 16 | O75521 | NA | ECI2 | enoyl-CoA delta isomerase 2 | 10455 | ENSG00000198721 |
| 17 | Q04837 | NA | SSBP1 | single-stranded DNA binding protein 1, mitochondrial | 6742 | ENSG00000106028 |
| 18 | P17096 | NA | HMGA1 | high mobility group AT-hook 1 | 3159 | ENSG00000137309 |
| 19 | Q16822 | NA | PCK2 | phosphoenolpyruvate carboxykinase 2 (mitochondrial) | 5106 | ENSG00000100889 |
| 20 | P49748 | NA | ACADVL | acyl-CoA dehydrogenase, very long chain | 37 | ENSG00000072778 |
| 21 | Q9Y3E0 | NA | GOLT1B | golgi transport 1B | 51026 | ENSG00000111711 |
| 22 | P51648 | NA | ALDH3A2 | aldehyde dehydrogenase 3 family, member A2 | 224 | ENSG00000072210 |
| 23 | P31937 | NA | HIBADH | 3-hydroxyisobutyrate dehydrogenase | 11112 | ENSG00000106049 |

  
  

| **Database:cellular component      &nbspName:organelle part      &nbspID:GO:0044422** | | | | | | |
| --- | --- | --- | --- | --- | --- | --- |
| C=6777; O=23; E=12.59; R=1.83; rawP=0.0002; adjP=0.0007 | | | | | | |
| Index | UserID | Value | Gene Symbol | Gene Name | EntrezGene | Ensembl |
| 1 | P10253 | NA | GAA | glucosidase, alpha; acid | 2548 | ENSG00000171298 |
| 2 | Q13724 | NA | MOGS | mannosyl-oligosaccharide glucosidase | 7841 | ENSG00000115275 |
| 3 | P04179 | NA | SOD2 | superoxide dismutase 2, mitochondrial | 6648 | ENSG00000112096 |
| 4 | P61769 | NA | B2M | beta-2-microglobulin | 567 | ENSG00000166710 |
| 5 | P48735 | NA | IDH2 | isocitrate dehydrogenase 2 (NADP+), mitochondrial | 3418 | ENSG00000182054 |
| 6 | P51659 | NA | HSD17B4 | hydroxysteroid (17-beta) dehydrogenase 4 | 3295 | ENSG00000133835 |
| 7 | P21796 | NA | VDAC1 | voltage-dependent anion channel 1 | 7416 | ENSG00000213585 |
| 8 | Q86UP2 | NA | KTN1 | kinectin 1 (kinesin receptor) | 3895 | ENSG00000126777 |
| 9 | P02545 | NA | LMNA | lamin A/C | 4000 | ENSG00000160789 |
| 10 | P32322 | NA | PYCR1 | pyrroline-5-carboxylate reductase 1 | 5831 | ENSG00000183010 |
| 11 | P17655 | NA | CAPN2 | calpain 2, (m/II) large subunit | 824 | ENSG00000162909 |
| 12 | O14773 | NA | TPP1 | tripeptidyl peptidase I | 1200 | ENSG00000166340 |
| 13 | Q9Y2Q3 | NA | GSTK1 | glutathione S-transferase kappa 1 | 373156 | ENSG00000197448 |
| 14 | P51688 | NA | SGSH | N-sulfoglucosamine sulfohydrolase | 6448 | ENSG00000181523 |
| 15 | P22570 | NA | FDXR | ferredoxin reductase | 2232 | ENSG00000161513 |
| 16 | O75521 | NA | ECI2 | enoyl-CoA delta isomerase 2 | 10455 | ENSG00000198721 |
| 17 | Q04837 | NA | SSBP1 | single-stranded DNA binding protein 1, mitochondrial | 6742 | ENSG00000106028 |
| 18 | P17096 | NA | HMGA1 | high mobility group AT-hook 1 | 3159 | ENSG00000137309 |
| 19 | Q16822 | NA | PCK2 | phosphoenolpyruvate carboxykinase 2 (mitochondrial) | 5106 | ENSG00000100889 |
| 20 | P49748 | NA | ACADVL | acyl-CoA dehydrogenase, very long chain | 37 | ENSG00000072778 |
| 21 | Q9Y3E0 | NA | GOLT1B | golgi transport 1B | 51026 | ENSG00000111711 |
| 22 | P51648 | NA | ALDH3A2 | aldehyde dehydrogenase 3 family, member A2 | 224 | ENSG00000072210 |
| 23 | P31937 | NA | HIBADH | 3-hydroxyisobutyrate dehydrogenase | 11112 | ENSG00000106049 |

  
  

| **Database:cellular component      &nbspName:intracellular membrane-bounded organelle      &nbspID:GO:0043231** | | | | | | |
| --- | --- | --- | --- | --- | --- | --- |
| C=9484; O=27; E=17.62; R=1.53; rawP=0.0003; adjP=0.0009 | | | | | | |
| Index | UserID | Value | Gene Symbol | Gene Name | EntrezGene | Ensembl |
| 1 | P10253 | NA | GAA | glucosidase, alpha; acid | 2548 | ENSG00000171298 |
| 2 | P55060 | NA | CSE1L | CSE1 chromosome segregation 1-like (yeast) | 1434 | ENSG00000124207 |
| 3 | Q13724 | NA | MOGS | mannosyl-oligosaccharide glucosidase | 7841 | ENSG00000115275 |
| 4 | P04179 | NA | SOD2 | superoxide dismutase 2, mitochondrial | 6648 | ENSG00000112096 |
| 5 | P61769 | NA | B2M | beta-2-microglobulin | 567 | ENSG00000166710 |
| 6 | Q8NFV4 | NA | ABHD11 | abhydrolase domain containing 11 | 83451 | ENSG00000106077 |
| 7 | P48735 | NA | IDH2 | isocitrate dehydrogenase 2 (NADP+), mitochondrial | 3418 | ENSG00000182054 |
| 8 | P51659 | NA | HSD17B4 | hydroxysteroid (17-beta) dehydrogenase 4 | 3295 | ENSG00000133835 |
| 9 | P21796 | NA | VDAC1 | voltage-dependent anion channel 1 | 7416 | ENSG00000213585 |
| 10 | Q86UP2 | NA | KTN1 | kinectin 1 (kinesin receptor) | 3895 | ENSG00000126777 |
| 11 | P02545 | NA | LMNA | lamin A/C | 4000 | ENSG00000160789 |
| 12 | P32322 | NA | PYCR1 | pyrroline-5-carboxylate reductase 1 | 5831 | ENSG00000183010 |
| 13 | P17655 | NA | CAPN2 | calpain 2, (m/II) large subunit | 824 | ENSG00000162909 |
| 14 | O14773 | NA | TPP1 | tripeptidyl peptidase I | 1200 | ENSG00000166340 |
| 15 | Q9Y2Q3 | NA | GSTK1 | glutathione S-transferase kappa 1 | 373156 | ENSG00000197448 |
| 16 | P51688 | NA | SGSH | N-sulfoglucosamine sulfohydrolase | 6448 | ENSG00000181523 |
| 17 | P22570 | NA | FDXR | ferredoxin reductase | 2232 | ENSG00000161513 |
| 18 | P05556 | NA | ITGB1 | integrin, beta 1 (fibronectin receptor, beta polypeptide, antigen CD29 includes MDF2, MSK12) | 3688 | ENSG00000150093 |
| 19 | O75521 | NA | ECI2 | enoyl-CoA delta isomerase 2 | 10455 | ENSG00000198721 |
| 20 | Q04837 | NA | SSBP1 | single-stranded DNA binding protein 1, mitochondrial | 6742 | ENSG00000106028 |
| 21 | P17096 | NA | HMGA1 | high mobility group AT-hook 1 | 3159 | ENSG00000137309 |
| 22 | Q16822 | NA | PCK2 | phosphoenolpyruvate carboxykinase 2 (mitochondrial) | 5106 | ENSG00000100889 |
| 23 | P49748 | NA | ACADVL | acyl-CoA dehydrogenase, very long chain | 37 | ENSG00000072778 |
| 24 | Q9Y3E0 | NA | GOLT1B | golgi transport 1B | 51026 | ENSG00000111711 |
| 25 | P51648 | NA | ALDH3A2 | aldehyde dehydrogenase 3 family, member A2 | 224 | ENSG00000072210 |
| 26 | P30042 | NA | C21orf33 | chromosome 21 open reading frame 33 | 8209 | ENSG00000160221 |
| 27 | P31937 | NA | HIBADH | 3-hydroxyisobutyrate dehydrogenase | 11112 | ENSG00000106049 |

  
  

| **Database:cellular component      &nbspName:membrane-bounded organelle      &nbspID:GO:0043227** | | | | | | |
| --- | --- | --- | --- | --- | --- | --- |
| C=9495; O=27; E=17.64; R=1.53; rawP=0.0003; adjP=0.0009 | | | | | | |
| Index | UserID | Value | Gene Symbol | Gene Name | EntrezGene | Ensembl |
| 1 | P10253 | NA | GAA | glucosidase, alpha; acid | 2548 | ENSG00000171298 |
| 2 | P55060 | NA | CSE1L | CSE1 chromosome segregation 1-like (yeast) | 1434 | ENSG00000124207 |
| 3 | Q13724 | NA | MOGS | mannosyl-oligosaccharide glucosidase | 7841 | ENSG00000115275 |
| 4 | P04179 | NA | SOD2 | superoxide dismutase 2, mitochondrial | 6648 | ENSG00000112096 |
| 5 | P61769 | NA | B2M | beta-2-microglobulin | 567 | ENSG00000166710 |
| 6 | Q8NFV4 | NA | ABHD11 | abhydrolase domain containing 11 | 83451 | ENSG00000106077 |
| 7 | P48735 | NA | IDH2 | isocitrate dehydrogenase 2 (NADP+), mitochondrial | 3418 | ENSG00000182054 |
| 8 | P51659 | NA | HSD17B4 | hydroxysteroid (17-beta) dehydrogenase 4 | 3295 | ENSG00000133835 |
| 9 | P21796 | NA | VDAC1 | voltage-dependent anion channel 1 | 7416 | ENSG00000213585 |
| 10 | Q86UP2 | NA | KTN1 | kinectin 1 (kinesin receptor) | 3895 | ENSG00000126777 |
| 11 | P02545 | NA | LMNA | lamin A/C | 4000 | ENSG00000160789 |
| 12 | P32322 | NA | PYCR1 | pyrroline-5-carboxylate reductase 1 | 5831 | ENSG00000183010 |
| 13 | P17655 | NA | CAPN2 | calpain 2, (m/II) large subunit | 824 | ENSG00000162909 |
| 14 | O14773 | NA | TPP1 | tripeptidyl peptidase I | 1200 | ENSG00000166340 |
| 15 | Q9Y2Q3 | NA | GSTK1 | glutathione S-transferase kappa 1 | 373156 | ENSG00000197448 |
| 16 | P51688 | NA | SGSH | N-sulfoglucosamine sulfohydrolase | 6448 | ENSG00000181523 |
| 17 | P22570 | NA | FDXR | ferredoxin reductase | 2232 | ENSG00000161513 |
| 18 | P05556 | NA | ITGB1 | integrin, beta 1 (fibronectin receptor, beta polypeptide, antigen CD29 includes MDF2, MSK12) | 3688 | ENSG00000150093 |
| 19 | O75521 | NA | ECI2 | enoyl-CoA delta isomerase 2 | 10455 | ENSG00000198721 |
| 20 | Q04837 | NA | SSBP1 | single-stranded DNA binding protein 1, mitochondrial | 6742 | ENSG00000106028 |
| 21 | P17096 | NA | HMGA1 | high mobility group AT-hook 1 | 3159 | ENSG00000137309 |
| 22 | Q16822 | NA | PCK2 | phosphoenolpyruvate carboxykinase 2 (mitochondrial) | 5106 | ENSG00000100889 |
| 23 | P49748 | NA | ACADVL | acyl-CoA dehydrogenase, very long chain | 37 | ENSG00000072778 |
| 24 | Q9Y3E0 | NA | GOLT1B | golgi transport 1B | 51026 | ENSG00000111711 |
| 25 | P51648 | NA | ALDH3A2 | aldehyde dehydrogenase 3 family, member A2 | 224 | ENSG00000072210 |
| 26 | P30042 | NA | C21orf33 | chromosome 21 open reading frame 33 | 8209 | ENSG00000160221 |
| 27 | P31937 | NA | HIBADH | 3-hydroxyisobutyrate dehydrogenase | 11112 | ENSG00000106049 |

  
  
  
  

---

WebGestalt is currently developed and maintained by Jing Wang and Bing Zhang at the  Zhang Lab. Other people who have made significant contribution to the project include Dexter Duncan, Stefan Kirov, Zhiao Shi, and Jay Snoddy.  
  
**Funding credits:** NIH/NIAAA (U01 AA016662, U01 AA013512); NIH/NIDA (P01 DA015027); NIH/NIMH (P50 MH078028, P50 MH096972); NIH/NCI (U24 CA159988); NIH/NIGMS (R01 GM088822).
