## Supplementary material for "PGRMC1 phosphorylation and cell plasticity 1: glycolysis, mitochondria, tumor growth": File S7: final_wiki_geneset_file_1438753413.html

Anchored HTML File of EIDs

|  |  |
| --- | --- |
|  | WEB-based GEne SeT AnaLysis Toolkit |
|  | ***Translating gene lists into biological insights...*** |

---

  

| **Database:Wikipathways pathway      &nbspName:Adipogenesis      &nbspID:WP236** | | | | | | |
| --- | --- | --- | --- | --- | --- | --- |
| C=130; O=4; E=0.20; R=20.13; rawP=4.43e-05; adjP=8.86e-05 | | | | | | |
| Index | UserID | Value | Gene Symbol | Gene Name | EntrezGene | Ensembl |
| 1 | P17096 | NA | HMGA1 | high mobility group AT-hook 1 | 3159 | ENSG00000137309 |
| 2 | Q16822 | NA | PCK2 | phosphoenolpyruvate carboxykinase 2 (mitochondrial) | 5106 | ENSG00000100889 |
| 3 | P02545 | NA | LMNA | lamin A/C | 4000 | ENSG00000160789 |
| 4 | P43490 | NA | NAMPT | nicotinamide phosphoribosyltransferase | 10135 | ENSG00000105835 |

  
  
  
  

---

WebGestalt is currently developed and maintained by Jing Wang and Bing Zhang at the  Zhang Lab. Other people who have made significant contribution to the project include Dexter Duncan, Stefan Kirov, Zhiao Shi, and Jay Snoddy.  
  
**Funding credits:** NIH/NIAAA (U01 AA016662, U01 AA013512); NIH/NIDA (P01 DA015027); NIH/NIMH (P50 MH078028, P50 MH096972); NIH/NCI (U24 CA159988); NIH/NIGMS (R01 GM088822).
