## Supplementary material for "PGRMC1 phosphorylation and cell plasticity 1: glycolysis, mitochondria, tumor growth": File S7: final_disease_geneset_file_1438487532.html

Anchored HTML File of EIDs

|  |  |
| --- | --- |
|  | WEB-based GEne SeT AnaLysis Toolkit |
|  | ***Translating gene lists into biological insights...*** |

---

  

| **Database:disease      &nbspName:Metabolism, Inborn Errors      &nbspID:DB\_ID:PA444939** | | | | | | |
| --- | --- | --- | --- | --- | --- | --- |
| C=335; O=11; E=1.42; R=7.74; rawP=1.72e-07; adjP=2.24e-05 | | | | | | |
| Index | UserID | Value | Gene Symbol | Gene Name | EntrezGene | Ensembl |
| 1 | O14773 | NA | TPP1 | tripeptidyl peptidase I | 1200 | ENSG00000166340 |
| 2 | Q04446 | NA | GBE1 | glucan (1,4-alpha-), branching enzyme 1 | 2632 | ENSG00000114480 |
| 3 | Q9UBQ7 | NA | GRHPR | glyoxylate reductase/hydroxypyruvate reductase | 9380 | ENSG00000137106 |
| 4 | P10253 | NA | GAA | glucosidase, alpha; acid | 2548 | ENSG00000171298 |
| 5 | P51649 | NA | ALDH5A1 | aldehyde dehydrogenase 5 family, member A1 | 7915 | ENSG00000112294 |
| 6 | P24752 | NA | ACAT1 | acetyl-CoA acetyltransferase 1 | 38 | ENSG00000075239 |
| 7 | P16278 | NA | GLB1 | galactosidase, beta 1 | 2720 | ENSG00000170266 |
| 8 | P07602 | NA | PSAP | prosaposin | 5660 | ENSG00000197746 |
| 9 | P49748 | NA | ACADVL | acyl-CoA dehydrogenase, very long chain | 37 | ENSG00000072778 |
| 10 | P38571 | NA | LIPA | lipase A, lysosomal acid, cholesterol esterase | 3988 | ENSG00000107798 |
| 11 | P02786 | NA | TFRC | transferrin receptor (p90, CD71) | 7037 | ENSG00000072274 |

  
  

| **Database:disease      &nbspName:Metabolic Diseases      &nbspID:DB\_ID:PA444938** | | | | | | |
| --- | --- | --- | --- | --- | --- | --- |
| C=574; O=13; E=2.43; R=5.34; rawP=8.56e-07; adjP=3.61e-05 | | | | | | |
| Index | UserID | Value | Gene Symbol | Gene Name | EntrezGene | Ensembl |
| 1 | O14773 | NA | TPP1 | tripeptidyl peptidase I | 1200 | ENSG00000166340 |
| 2 | Q04446 | NA | GBE1 | glucan (1,4-alpha-), branching enzyme 1 | 2632 | ENSG00000114480 |
| 3 | P02792 | NA | FTL | ferritin, light polypeptide | 2512 | ENSG00000087086 |
| 4 | Q9UBQ7 | NA | GRHPR | glyoxylate reductase/hydroxypyruvate reductase | 9380 | ENSG00000137106 |
| 5 | P10253 | NA | GAA | glucosidase, alpha; acid | 2548 | ENSG00000171298 |
| 6 | P24752 | NA | ACAT1 | acetyl-CoA acetyltransferase 1 | 38 | ENSG00000075239 |
| 7 | P16278 | NA | GLB1 | galactosidase, beta 1 | 2720 | ENSG00000170266 |
| 8 | P00367 | NA | GLUD1 | glutamate dehydrogenase 1 | 2746 | ENSG00000148672 |
| 9 | P07602 | NA | PSAP | prosaposin | 5660 | ENSG00000197746 |
| 10 | P49748 | NA | ACADVL | acyl-CoA dehydrogenase, very long chain | 37 | ENSG00000072778 |
| 11 | P38571 | NA | LIPA | lipase A, lysosomal acid, cholesterol esterase | 3988 | ENSG00000107798 |
| 12 | P02786 | NA | TFRC | transferrin receptor (p90, CD71) | 7037 | ENSG00000072274 |
| 13 | P43490 | NA | NAMPT | nicotinamide phosphoribosyltransferase | 10135 | ENSG00000105835 |

  
  

| **Database:disease      &nbspName:Arsenic Poisoning      &nbspID:DB\_ID:PA446977** | | | | | | |
| --- | --- | --- | --- | --- | --- | --- |
| C=19; O=4; E=0.08; R=49.65; rawP=1.11e-06; adjP=3.61e-05 | | | | | | |
| Index | UserID | Value | Gene Symbol | Gene Name | EntrezGene | Ensembl |
| 1 | P04040 | NA | CAT | catalase | 847 | ENSG00000121691 |
| 2 | Q16881 | NA | TXNRD1 | thioredoxin reductase 1 | 7296 | ENSG00000198431 |
| 3 | P78417 | NA | GSTO1 | glutathione S-transferase omega 1 | 9446 | ENSG00000148834 |
| 4 | P09211 | NA | GSTP1 | glutathione S-transferase pi 1 | 2950 | ENSG00000084207 |

  
  

| **Database:disease      &nbspName:Neoplasm Metastasis      &nbspID:DB\_ID:PA445058** | | | | | | |
| --- | --- | --- | --- | --- | --- | --- |
| C=302; O=10; E=1.28; R=7.81; rawP=6.02e-07; adjP=3.61e-05 | | | | | | |
| Index | UserID | Value | Gene Symbol | Gene Name | EntrezGene | Ensembl |
| 1 | P17931 | NA | LGALS3 | lectin, galactoside-binding, soluble, 3 | 3958 | ENSG00000131981 |
| 2 | P30086 | NA | PEBP1 | phosphatidylethanolamine binding protein 1 | 5037 | ENSG00000089220 |
| 3 | P05556 | NA | ITGB1 | integrin, beta 1 (fibronectin receptor, beta polypeptide, antigen CD29 includes MDF2, MSK12) | 3688 | ENSG00000150093 |
| 4 | P40121 | NA | CAPG | capping protein (actin filament), gelsolin-like | 822 | ENSG00000042493 |
| 5 | P06744 | NA | GPI | glucose-6-phosphate isomerase | 2821 | ENSG00000105220 |
| 6 | P08727 | NA | KRT19 | keratin 19 | 3880 | ENSG00000171345 |
| 7 | P07339 | NA | CTSD | cathepsin D | 1509 | ENSG00000117984 |
| 8 | P16949 | NA | STMN1 | stathmin 1 | 3925 | ENSG00000117632 |
| 9 | P04083 | NA | ANXA1 | annexin A1 | 301 | ENSG00000135046 |
| 10 | O00425 | NA | IGF2BP3 | insulin-like growth factor 2 mRNA binding protein 3 | 10643 | ENSG00000136231 |

  
  

| **Database:disease      &nbspName:Carcinoma, Large Cell      &nbspID:DB\_ID:PA446660** | | | | | | |
| --- | --- | --- | --- | --- | --- | --- |
| C=99; O=6; E=0.42; R=14.29; rawP=3.98e-06; adjP=0.0001 | | | | | | |
| Index | UserID | Value | Gene Symbol | Gene Name | EntrezGene | Ensembl |
| 1 | P37802 | NA | TAGLN2 | transgelin 2 | 8407 | ENSG00000158710 |
| 2 | P08195 | NA | SLC3A2 | solute carrier family 3 (activators of dibasic and neutral amino acid transport), member 2 | 6520 | ENSG00000168003 |
| 3 | P08727 | NA | KRT19 | keratin 19 | 3880 | ENSG00000171345 |
| 4 | P49736 | NA | MCM2 | minichromosome maintenance complex component 2 | 4171 | ENSG00000073111 |
| 5 | P09211 | NA | GSTP1 | glutathione S-transferase pi 1 | 2950 | ENSG00000084207 |
| 6 | O00425 | NA | IGF2BP3 | insulin-like growth factor 2 mRNA binding protein 3 | 10643 | ENSG00000136231 |

  
  

| **Database:disease      &nbspName:Neoplastic Processes      &nbspID:DB\_ID:PA445078** | | | | | | |
| --- | --- | --- | --- | --- | --- | --- |
| C=370; O=9; E=1.57; R=5.74; rawP=2.71e-05; adjP=0.0004 | | | | | | |
| Index | UserID | Value | Gene Symbol | Gene Name | EntrezGene | Ensembl |
| 1 | P17931 | NA | LGALS3 | lectin, galactoside-binding, soluble, 3 | 3958 | ENSG00000131981 |
| 2 | P30086 | NA | PEBP1 | phosphatidylethanolamine binding protein 1 | 5037 | ENSG00000089220 |
| 3 | P05556 | NA | ITGB1 | integrin, beta 1 (fibronectin receptor, beta polypeptide, antigen CD29 includes MDF2, MSK12) | 3688 | ENSG00000150093 |
| 4 | P08727 | NA | KRT19 | keratin 19 | 3880 | ENSG00000171345 |
| 5 | P07339 | NA | CTSD | cathepsin D | 1509 | ENSG00000117984 |
| 6 | P16949 | NA | STMN1 | stathmin 1 | 3925 | ENSG00000117632 |
| 7 | Q9P0J0 | NA | NDUFA13 | NADH dehydrogenase (ubiquinone) 1 alpha subcomplex, 13 | 51079 | ENSG00000186010 |
| 8 | P04083 | NA | ANXA1 | annexin A1 | 301 | ENSG00000135046 |
| 9 | O00425 | NA | IGF2BP3 | insulin-like growth factor 2 mRNA binding protein 3 | 10643 | ENSG00000136231 |

  
  

| **Database:disease      &nbspName:Stress      &nbspID:DB\_ID:PA445752** | | | | | | |
| --- | --- | --- | --- | --- | --- | --- |
| C=459; O=10; E=1.95; R=5.14; rawP=2.43e-05; adjP=0.0004 | | | | | | |
| Index | UserID | Value | Gene Symbol | Gene Name | EntrezGene | Ensembl |
| 1 | P04040 | NA | CAT | catalase | 847 | ENSG00000121691 |
| 2 | Q16881 | NA | TXNRD1 | thioredoxin reductase 1 | 7296 | ENSG00000198431 |
| 3 | P17655 | NA | CAPN2 | calpain 2, (m/II) large subunit | 824 | ENSG00000162909 |
| 4 | P02792 | NA | FTL | ferritin, light polypeptide | 2512 | ENSG00000087086 |
| 5 | Q92598 | NA | HSPH1 | heat shock 105kDa/110kDa protein 1 | 10808 | ENSG00000120694 |
| 6 | P30044 | NA | PRDX5 | peroxiredoxin 5 | 25824 | ENSG00000126432 |
| 7 | P00338 | NA | LDHA | lactate dehydrogenase A | 3939 | ENSG00000134333 |
| 8 | P78417 | NA | GSTO1 | glutathione S-transferase omega 1 | 9446 | ENSG00000148834 |
| 9 | P09211 | NA | GSTP1 | glutathione S-transferase pi 1 | 2950 | ENSG00000084207 |
| 10 | Q06830 | NA | PRDX1 | peroxiredoxin 1 | 5052 | ENSG00000117450 |

  
  

| **Database:disease      &nbspName:Protein Deficiency      &nbspID:DB\_ID:PA445428** | | | | | | |
| --- | --- | --- | --- | --- | --- | --- |
| C=354; O=9; E=1.50; R=6.00; rawP=1.91e-05; adjP=0.0004 | | | | | | |
| Index | UserID | Value | Gene Symbol | Gene Name | EntrezGene | Ensembl |
| 1 | O14773 | NA | TPP1 | tripeptidyl peptidase I | 1200 | ENSG00000166340 |
| 2 | P10253 | NA | GAA | glucosidase, alpha; acid | 2548 | ENSG00000171298 |
| 3 | P51649 | NA | ALDH5A1 | aldehyde dehydrogenase 5 family, member A1 | 7915 | ENSG00000112294 |
| 4 | P00491 | NA | PNP | purine nucleoside phosphorylase | 4860 | ENSG00000198805 |
| 5 | P24752 | NA | ACAT1 | acetyl-CoA acetyltransferase 1 | 38 | ENSG00000075239 |
| 6 | P49419 | NA | ALDH7A1 | aldehyde dehydrogenase 7 family, member A1 | 501 | ENSG00000164904 |
| 7 | P16278 | NA | GLB1 | galactosidase, beta 1 | 2720 | ENSG00000170266 |
| 8 | P07602 | NA | PSAP | prosaposin | 5660 | ENSG00000197746 |
| 9 | P49748 | NA | ACADVL | acyl-CoA dehydrogenase, very long chain | 37 | ENSG00000072778 |

  
  

| **Database:disease      &nbspName:Adenocarcinoma      &nbspID:DB\_ID:PA443265** | | | | | | |
| --- | --- | --- | --- | --- | --- | --- |
| C=348; O=9; E=1.48; R=6.10; rawP=1.67e-05; adjP=0.0004 | | | | | | |
| Index | UserID | Value | Gene Symbol | Gene Name | EntrezGene | Ensembl |
| 1 | P17931 | NA | LGALS3 | lectin, galactoside-binding, soluble, 3 | 3958 | ENSG00000131981 |
| 2 | Q05639 | NA | EEF1A2 | eukaryotic translation elongation factor 1 alpha 2 | 1917 | ENSG00000101210 |
| 3 | P30086 | NA | PEBP1 | phosphatidylethanolamine binding protein 1 | 5037 | ENSG00000089220 |
| 4 | P08727 | NA | KRT19 | keratin 19 | 3880 | ENSG00000171345 |
| 5 | P49736 | NA | MCM2 | minichromosome maintenance complex component 2 | 4171 | ENSG00000073111 |
| 6 | P09211 | NA | GSTP1 | glutathione S-transferase pi 1 | 2950 | ENSG00000084207 |
| 7 | Q06830 | NA | PRDX1 | peroxiredoxin 1 | 5052 | ENSG00000117450 |
| 8 | P04083 | NA | ANXA1 | annexin A1 | 301 | ENSG00000135046 |
| 9 | O00425 | NA | IGF2BP3 | insulin-like growth factor 2 mRNA binding protein 3 | 10643 | ENSG00000136231 |

  
  

| **Database:disease      &nbspName:Lipidoses      &nbspID:DB\_ID:PA444787** | | | | | | |
| --- | --- | --- | --- | --- | --- | --- |
| C=45; O=4; E=0.19; R=20.96; rawP=3.93e-05; adjP=0.0005 | | | | | | |
| Index | UserID | Value | Gene Symbol | Gene Name | EntrezGene | Ensembl |
| 1 | O14773 | NA | TPP1 | tripeptidyl peptidase I | 1200 | ENSG00000166340 |
| 2 | P16278 | NA | GLB1 | galactosidase, beta 1 | 2720 | ENSG00000170266 |
| 3 | P07602 | NA | PSAP | prosaposin | 5660 | ENSG00000197746 |
| 4 | P38571 | NA | LIPA | lipase A, lysosomal acid, cholesterol esterase | 3988 | ENSG00000107798 |

  
  

| **Database:disease      &nbspName:Protein deficiency disease      &nbspID:DB\_ID:PA165108904** | | | | | | |
| --- | --- | --- | --- | --- | --- | --- |
| C=233; O=7; E=0.99; R=7.09; rawP=6.00e-05; adjP=0.0007 | | | | | | |
| Index | UserID | Value | Gene Symbol | Gene Name | EntrezGene | Ensembl |
| 1 | O14773 | NA | TPP1 | tripeptidyl peptidase I | 1200 | ENSG00000166340 |
| 2 | P16278 | NA | GLB1 | galactosidase, beta 1 | 2720 | ENSG00000170266 |
| 3 | P49419 | NA | ALDH7A1 | aldehyde dehydrogenase 7 family, member A1 | 501 | ENSG00000164904 |
| 4 | P10253 | NA | GAA | glucosidase, alpha; acid | 2548 | ENSG00000171298 |
| 5 | P07602 | NA | PSAP | prosaposin | 5660 | ENSG00000197746 |
| 6 | P51649 | NA | ALDH5A1 | aldehyde dehydrogenase 5 family, member A1 | 7915 | ENSG00000112294 |
| 7 | P38571 | NA | LIPA | lipase A, lysosomal acid, cholesterol esterase | 3988 | ENSG00000107798 |

  
  
  
  

---

WebGestalt is currently developed and maintained by Jing Wang and Bing Zhang at the  Zhang Lab. Other people who have made significant contribution to the project include Dexter Duncan, Stefan Kirov, Zhiao Shi, and Jay Snoddy.  
  
**Funding credits:** NIH/NIAAA (U01 AA016662, U01 AA013512); NIH/NIDA (P01 DA015027); NIH/NIMH (P50 MH078028, P50 MH096972); NIH/NCI (U24 CA159988); NIH/NIGMS (R01 GM088822).
