## Supplementary material for "PGRMC1 phosphorylation and cell plasticity 1: glycolysis, mitochondria, tumor growth": File S7: final_sig_file_1438487532.html

Anchored HTML File of EIDs

|  |  |
| --- | --- |
|  | WEB-based GEne SeT AnaLysis Toolkit |
|  | ***Translating gene lists into biological insights...*** |

---

  

| **Database:biological process      &nbspName:organic acid metabolic process      &nbspID:GO:0006082** | | | | | | |
| --- | --- | --- | --- | --- | --- | --- |
| C=967; O=25; E=5.63; R=4.44; rawP=1.14e-10; adjP=6.97e-08 | | | | | | |
| Index | UserID | Value | Gene Symbol | Gene Name | EntrezGene | Ensembl |
| 1 | P49591 | NA | SARS | seryl-tRNA synthetase | 6301 | ENSG00000031698 |
| 2 | P00491 | NA | PNP | purine nucleoside phosphorylase | 4860 | ENSG00000198805 |
| 3 | Q16851 | NA | UGP2 | UDP-glucose pyrophosphorylase 2 | 7360 | ENSG00000169764 |
| 4 | P00338 | NA | LDHA | lactate dehydrogenase A | 3939 | ENSG00000134333 |
| 5 | P56192 | NA | MARS | methionyl-tRNA synthetase | 4141 | ENSG00000166986 |
| 6 | P00367 | NA | GLUD1 | glutamate dehydrogenase 1 | 2746 | ENSG00000148672 |
| 7 | Q06323 | NA | PSME1 | proteasome (prosome, macropain) activator subunit 1 (PA28 alpha) | 5720 | ENSG00000092010 |
| 8 | P48735 | NA | IDH2 | isocitrate dehydrogenase 2 (NADP+), mitochondrial | 3418 | ENSG00000182054 |
| 9 | P78417 | NA | GSTO1 | glutathione S-transferase omega 1 | 9446 | ENSG00000148834 |
| 10 | P09211 | NA | GSTP1 | glutathione S-transferase pi 1 | 2950 | ENSG00000084207 |
| 11 | P38571 | NA | LIPA | lipase A, lysosomal acid, cholesterol esterase | 3988 | ENSG00000107798 |
| 12 | P49588 | NA | AARS | alanyl-tRNA synthetase | 16 | ENSG00000090861 |
| 13 | Q13011 | NA | ECH1 | enoyl CoA hydratase 1, peroxisomal | 1891 | ENSG00000104823 |
| 14 | P04083 | NA | ANXA1 | annexin A1 | 301 | ENSG00000135046 |
| 15 | P23381 | NA | WARS | tryptophanyl-tRNA synthetase | 7453 | ENSG00000140105 |
| 16 | Q9UBQ7 | NA | GRHPR | glyoxylate reductase/hydroxypyruvate reductase | 9380 | ENSG00000137106 |
| 17 | P51649 | NA | ALDH5A1 | aldehyde dehydrogenase 5 family, member A1 | 7915 | ENSG00000112294 |
| 18 | P02768 | NA | ALB | albumin | 213 | ENSG00000163631 |
| 19 | P42765 | NA | ACAA2 | acetyl-CoA acyltransferase 2 | 10449 | ENSG00000167315 |
| 20 | P24752 | NA | ACAT1 | acetyl-CoA acetyltransferase 1 | 38 | ENSG00000075239 |
| 21 | P13674 | NA | P4HA1 | prolyl 4-hydroxylase, alpha polypeptide I | 5033 | ENSG00000122884 |
| 22 | P16278 | NA | GLB1 | galactosidase, beta 1 | 2720 | ENSG00000170266 |
| 23 | P17174 | NA | GOT1 | glutamic-oxaloacetic transaminase 1, soluble (aspartate aminotransferase 1) | 2805 | ENSG00000120053 |
| 24 | P49419 | NA | ALDH7A1 | aldehyde dehydrogenase 7 family, member A1 | 501 | ENSG00000164904 |
| 25 | P49748 | NA | ACADVL | acyl-CoA dehydrogenase, very long chain | 37 | ENSG00000072778 |

  
  

| **Database:biological process      &nbspName:carboxylic acid metabolic process      &nbspID:GO:0019752** | | | | | | |
| --- | --- | --- | --- | --- | --- | --- |
| C=842; O=23; E=4.90; R=4.69; rawP=2.63e-10; adjP=8.03e-08 | | | | | | |
| Index | UserID | Value | Gene Symbol | Gene Name | EntrezGene | Ensembl |
| 1 | P49591 | NA | SARS | seryl-tRNA synthetase | 6301 | ENSG00000031698 |
| 2 | Q16851 | NA | UGP2 | UDP-glucose pyrophosphorylase 2 | 7360 | ENSG00000169764 |
| 3 | P00338 | NA | LDHA | lactate dehydrogenase A | 3939 | ENSG00000134333 |
| 4 | P56192 | NA | MARS | methionyl-tRNA synthetase | 4141 | ENSG00000166986 |
| 5 | P00367 | NA | GLUD1 | glutamate dehydrogenase 1 | 2746 | ENSG00000148672 |
| 6 | Q06323 | NA | PSME1 | proteasome (prosome, macropain) activator subunit 1 (PA28 alpha) | 5720 | ENSG00000092010 |
| 7 | P48735 | NA | IDH2 | isocitrate dehydrogenase 2 (NADP+), mitochondrial | 3418 | ENSG00000182054 |
| 8 | P78417 | NA | GSTO1 | glutathione S-transferase omega 1 | 9446 | ENSG00000148834 |
| 9 | P09211 | NA | GSTP1 | glutathione S-transferase pi 1 | 2950 | ENSG00000084207 |
| 10 | P38571 | NA | LIPA | lipase A, lysosomal acid, cholesterol esterase | 3988 | ENSG00000107798 |
| 11 | P49588 | NA | AARS | alanyl-tRNA synthetase | 16 | ENSG00000090861 |
| 12 | Q13011 | NA | ECH1 | enoyl CoA hydratase 1, peroxisomal | 1891 | ENSG00000104823 |
| 13 | P04083 | NA | ANXA1 | annexin A1 | 301 | ENSG00000135046 |
| 14 | P23381 | NA | WARS | tryptophanyl-tRNA synthetase | 7453 | ENSG00000140105 |
| 15 | Q9UBQ7 | NA | GRHPR | glyoxylate reductase/hydroxypyruvate reductase | 9380 | ENSG00000137106 |
| 16 | P51649 | NA | ALDH5A1 | aldehyde dehydrogenase 5 family, member A1 | 7915 | ENSG00000112294 |
| 17 | P02768 | NA | ALB | albumin | 213 | ENSG00000163631 |
| 18 | P42765 | NA | ACAA2 | acetyl-CoA acyltransferase 2 | 10449 | ENSG00000167315 |
| 19 | P24752 | NA | ACAT1 | acetyl-CoA acetyltransferase 1 | 38 | ENSG00000075239 |
| 20 | P13674 | NA | P4HA1 | prolyl 4-hydroxylase, alpha polypeptide I | 5033 | ENSG00000122884 |
| 21 | P17174 | NA | GOT1 | glutamic-oxaloacetic transaminase 1, soluble (aspartate aminotransferase 1) | 2805 | ENSG00000120053 |
| 22 | P49419 | NA | ALDH7A1 | aldehyde dehydrogenase 7 family, member A1 | 501 | ENSG00000164904 |
| 23 | P49748 | NA | ACADVL | acyl-CoA dehydrogenase, very long chain | 37 | ENSG00000072778 |

  
  

| **Database:biological process      &nbspName:oxoacid metabolic process      &nbspID:GO:0043436** | | | | | | |
| --- | --- | --- | --- | --- | --- | --- |
| C=949; O=24; E=5.53; R=4.34; rawP=4.68e-10; adjP=9.27e-08 | | | | | | |
| Index | UserID | Value | Gene Symbol | Gene Name | EntrezGene | Ensembl |
| 1 | P49591 | NA | SARS | seryl-tRNA synthetase | 6301 | ENSG00000031698 |
| 2 | Q16851 | NA | UGP2 | UDP-glucose pyrophosphorylase 2 | 7360 | ENSG00000169764 |
| 3 | P00338 | NA | LDHA | lactate dehydrogenase A | 3939 | ENSG00000134333 |
| 4 | P56192 | NA | MARS | methionyl-tRNA synthetase | 4141 | ENSG00000166986 |
| 5 | P00367 | NA | GLUD1 | glutamate dehydrogenase 1 | 2746 | ENSG00000148672 |
| 6 | Q06323 | NA | PSME1 | proteasome (prosome, macropain) activator subunit 1 (PA28 alpha) | 5720 | ENSG00000092010 |
| 7 | P48735 | NA | IDH2 | isocitrate dehydrogenase 2 (NADP+), mitochondrial | 3418 | ENSG00000182054 |
| 8 | P78417 | NA | GSTO1 | glutathione S-transferase omega 1 | 9446 | ENSG00000148834 |
| 9 | P09211 | NA | GSTP1 | glutathione S-transferase pi 1 | 2950 | ENSG00000084207 |
| 10 | P38571 | NA | LIPA | lipase A, lysosomal acid, cholesterol esterase | 3988 | ENSG00000107798 |
| 11 | P49588 | NA | AARS | alanyl-tRNA synthetase | 16 | ENSG00000090861 |
| 12 | Q13011 | NA | ECH1 | enoyl CoA hydratase 1, peroxisomal | 1891 | ENSG00000104823 |
| 13 | P04083 | NA | ANXA1 | annexin A1 | 301 | ENSG00000135046 |
| 14 | P23381 | NA | WARS | tryptophanyl-tRNA synthetase | 7453 | ENSG00000140105 |
| 15 | Q9UBQ7 | NA | GRHPR | glyoxylate reductase/hydroxypyruvate reductase | 9380 | ENSG00000137106 |
| 16 | P51649 | NA | ALDH5A1 | aldehyde dehydrogenase 5 family, member A1 | 7915 | ENSG00000112294 |
| 17 | P02768 | NA | ALB | albumin | 213 | ENSG00000163631 |
| 18 | P42765 | NA | ACAA2 | acetyl-CoA acyltransferase 2 | 10449 | ENSG00000167315 |
| 19 | P24752 | NA | ACAT1 | acetyl-CoA acetyltransferase 1 | 38 | ENSG00000075239 |
| 20 | P13674 | NA | P4HA1 | prolyl 4-hydroxylase, alpha polypeptide I | 5033 | ENSG00000122884 |
| 21 | P16278 | NA | GLB1 | galactosidase, beta 1 | 2720 | ENSG00000170266 |
| 22 | P17174 | NA | GOT1 | glutamic-oxaloacetic transaminase 1, soluble (aspartate aminotransferase 1) | 2805 | ENSG00000120053 |
| 23 | P49419 | NA | ALDH7A1 | aldehyde dehydrogenase 7 family, member A1 | 501 | ENSG00000164904 |
| 24 | P49748 | NA | ACADVL | acyl-CoA dehydrogenase, very long chain | 37 | ENSG00000072778 |

  
  

| **Database:biological process      &nbspName:small molecule metabolic process      &nbspID:GO:0044281** | | | | | | |
| --- | --- | --- | --- | --- | --- | --- |
| C=2500; O=39; E=14.56; R=2.68; rawP=6.07e-10; adjP=9.27e-08 | | | | | | |
| Index | UserID | Value | Gene Symbol | Gene Name | EntrezGene | Ensembl |
| 1 | Q04446 | NA | GBE1 | glucan (1,4-alpha-), branching enzyme 1 | 2632 | ENSG00000114480 |
| 2 | P49591 | NA | SARS | seryl-tRNA synthetase | 6301 | ENSG00000031698 |
| 3 | Q16851 | NA | UGP2 | UDP-glucose pyrophosphorylase 2 | 7360 | ENSG00000169764 |
| 4 | P00367 | NA | GLUD1 | glutamate dehydrogenase 1 | 2746 | ENSG00000148672 |
| 5 | Q06323 | NA | PSME1 | proteasome (prosome, macropain) activator subunit 1 (PA28 alpha) | 5720 | ENSG00000092010 |
| 6 | P48735 | NA | IDH2 | isocitrate dehydrogenase 2 (NADP+), mitochondrial | 3418 | ENSG00000182054 |
| 7 | P09211 | NA | GSTP1 | glutathione S-transferase pi 1 | 2950 | ENSG00000084207 |
| 8 | Q9P0J0 | NA | NDUFA13 | NADH dehydrogenase (ubiquinone) 1 alpha subcomplex, 13 | 51079 | ENSG00000186010 |
| 9 | P49588 | NA | AARS | alanyl-tRNA synthetase | 16 | ENSG00000090861 |
| 10 | P43490 | NA | NAMPT | nicotinamide phosphoribosyltransferase | 10135 | ENSG00000105835 |
| 11 | P22102 | NA | GART | phosphoribosylglycinamide formyltransferase, phosphoribosylglycinamide synthetase, phosphoribosylaminoimidazole synthetase | 2618 | ENSG00000159131 |
| 12 | P16615 | NA | ATP2A2 | ATPase, Ca++ transporting, cardiac muscle, slow twitch 2 | 488 | ENSG00000174437 |
| 13 | P22570 | NA | FDXR | ferredoxin reductase | 2232 | ENSG00000161513 |
| 14 | P42765 | NA | ACAA2 | acetyl-CoA acyltransferase 2 | 10449 | ENSG00000167315 |
| 15 | P49419 | NA | ALDH7A1 | aldehyde dehydrogenase 7 family, member A1 | 501 | ENSG00000164904 |
| 16 | P17174 | NA | GOT1 | glutamic-oxaloacetic transaminase 1, soluble (aspartate aminotransferase 1) | 2805 | ENSG00000120053 |
| 17 | P53007 | NA | SLC25A1 | solute carrier family 25 (mitochondrial carrier; citrate transporter), member 1 | 6576 | ENSG00000100075 |
| 18 | P21589 | NA | NT5E | 5'-nucleotidase, ecto (CD73) | 4907 | ENSG00000135318 |
| 19 | Q16881 | NA | TXNRD1 | thioredoxin reductase 1 | 7296 | ENSG00000198431 |
| 20 | P00491 | NA | PNP | purine nucleoside phosphorylase | 4860 | ENSG00000198805 |
| 21 | P00338 | NA | LDHA | lactate dehydrogenase A | 3939 | ENSG00000134333 |
| 22 | P78417 | NA | GSTO1 | glutathione S-transferase omega 1 | 9446 | ENSG00000148834 |
| 23 | P56192 | NA | MARS | methionyl-tRNA synthetase | 4141 | ENSG00000166986 |
| 24 | P38571 | NA | LIPA | lipase A, lysosomal acid, cholesterol esterase | 3988 | ENSG00000107798 |
| 25 | Q13011 | NA | ECH1 | enoyl CoA hydratase 1, peroxisomal | 1891 | ENSG00000104823 |
| 26 | P23381 | NA | WARS | tryptophanyl-tRNA synthetase | 7453 | ENSG00000140105 |
| 27 | P04083 | NA | ANXA1 | annexin A1 | 301 | ENSG00000135046 |
| 28 | P04040 | NA | CAT | catalase | 847 | ENSG00000121691 |
| 29 | Q9UBQ7 | NA | GRHPR | glyoxylate reductase/hydroxypyruvate reductase | 9380 | ENSG00000137106 |
| 30 | P51649 | NA | ALDH5A1 | aldehyde dehydrogenase 5 family, member A1 | 7915 | ENSG00000112294 |
| 31 | P02768 | NA | ALB | albumin | 213 | ENSG00000163631 |
| 32 | P41091 | NA | EIF2S3 | eukaryotic translation initiation factor 2, subunit 3 gamma, 52kDa | 1968 | ENSG00000130741 |
| 33 | P30044 | NA | PRDX5 | peroxiredoxin 5 | 25824 | ENSG00000126432 |
| 34 | P24752 | NA | ACAT1 | acetyl-CoA acetyltransferase 1 | 38 | ENSG00000075239 |
| 35 | P13674 | NA | P4HA1 | prolyl 4-hydroxylase, alpha polypeptide I | 5033 | ENSG00000122884 |
| 36 | P16278 | NA | GLB1 | galactosidase, beta 1 | 2720 | ENSG00000170266 |
| 37 | P07602 | NA | PSAP | prosaposin | 5660 | ENSG00000197746 |
| 38 | P06744 | NA | GPI | glucose-6-phosphate isomerase | 2821 | ENSG00000105220 |
| 39 | P49748 | NA | ACADVL | acyl-CoA dehydrogenase, very long chain | 37 | ENSG00000072778 |

  
  

| **Database:biological process      &nbspName:monocarboxylic acid metabolic process      &nbspID:GO:0032787** | | | | | | |
| --- | --- | --- | --- | --- | --- | --- |
| C=405; O=13; E=2.36; R=5.51; rawP=5.72e-07; adjP=6.99e-05 | | | | | | |
| Index | UserID | Value | Gene Symbol | Gene Name | EntrezGene | Ensembl |
| 1 | Q9UBQ7 | NA | GRHPR | glyoxylate reductase/hydroxypyruvate reductase | 9380 | ENSG00000137106 |
| 2 | P51649 | NA | ALDH5A1 | aldehyde dehydrogenase 5 family, member A1 | 7915 | ENSG00000112294 |
| 3 | Q16851 | NA | UGP2 | UDP-glucose pyrophosphorylase 2 | 7360 | ENSG00000169764 |
| 4 | P02768 | NA | ALB | albumin | 213 | ENSG00000163631 |
| 5 | P00338 | NA | LDHA | lactate dehydrogenase A | 3939 | ENSG00000134333 |
| 6 | P42765 | NA | ACAA2 | acetyl-CoA acyltransferase 2 | 10449 | ENSG00000167315 |
| 7 | P13674 | NA | P4HA1 | prolyl 4-hydroxylase, alpha polypeptide I | 5033 | ENSG00000122884 |
| 8 | P48735 | NA | IDH2 | isocitrate dehydrogenase 2 (NADP+), mitochondrial | 3418 | ENSG00000182054 |
| 9 | P17174 | NA | GOT1 | glutamic-oxaloacetic transaminase 1, soluble (aspartate aminotransferase 1) | 2805 | ENSG00000120053 |
| 10 | P49748 | NA | ACADVL | acyl-CoA dehydrogenase, very long chain | 37 | ENSG00000072778 |
| 11 | P38571 | NA | LIPA | lipase A, lysosomal acid, cholesterol esterase | 3988 | ENSG00000107798 |
| 12 | Q13011 | NA | ECH1 | enoyl CoA hydratase 1, peroxisomal | 1891 | ENSG00000104823 |
| 13 | P04083 | NA | ANXA1 | annexin A1 | 301 | ENSG00000135046 |

  
  

| **Database:biological process      &nbspName:response to inorganic substance      &nbspID:GO:0010035** | | | | | | |
| --- | --- | --- | --- | --- | --- | --- |
| C=423; O=13; E=2.46; R=5.28; rawP=9.33e-07; adjP=9.50e-05 | | | | | | |
| Index | UserID | Value | Gene Symbol | Gene Name | EntrezGene | Ensembl |
| 1 | P04040 | NA | CAT | catalase | 847 | ENSG00000121691 |
| 2 | Q16881 | NA | TXNRD1 | thioredoxin reductase 1 | 7296 | ENSG00000198431 |
| 3 | P22102 | NA | GART | phosphoribosylglycinamide formyltransferase, phosphoribosylglycinamide synthetase, phosphoribosylaminoimidazole synthetase | 2618 | ENSG00000159131 |
| 4 | P02768 | NA | ALB | albumin | 213 | ENSG00000163631 |
| 5 | Q05639 | NA | EEF1A2 | eukaryotic translation elongation factor 1 alpha 2 | 1917 | ENSG00000101210 |
| 6 | P30044 | NA | PRDX5 | peroxiredoxin 5 | 25824 | ENSG00000126432 |
| 7 | P30086 | NA | PEBP1 | phosphatidylethanolamine binding protein 1 | 5037 | ENSG00000089220 |
| 8 | P78417 | NA | GSTO1 | glutathione S-transferase omega 1 | 9446 | ENSG00000148834 |
| 9 | P09211 | NA | GSTP1 | glutathione S-transferase pi 1 | 2950 | ENSG00000084207 |
| 10 | O60610 | NA | DIAPH1 | diaphanous homolog 1 (Drosophila) | 1729 | ENSG00000131504 |
| 11 | P02786 | NA | TFRC | transferrin receptor (p90, CD71) | 7037 | ENSG00000072274 |
| 12 | Q06830 | NA | PRDX1 | peroxiredoxin 1 | 5052 | ENSG00000117450 |
| 13 | P04083 | NA | ANXA1 | annexin A1 | 301 | ENSG00000135046 |

  
  

| **Database:biological process      &nbspName:oxidation-reduction process      &nbspID:GO:0055114** | | | | | | |
| --- | --- | --- | --- | --- | --- | --- |
| C=554; O=14; E=3.23; R=4.34; rawP=3.44e-06; adjP=0.0003 | | | | | | |
| Index | UserID | Value | Gene Symbol | Gene Name | EntrezGene | Ensembl |
| 1 | P04040 | NA | CAT | catalase | 847 | ENSG00000121691 |
| 2 | Q16881 | NA | TXNRD1 | thioredoxin reductase 1 | 7296 | ENSG00000198431 |
| 3 | Q04446 | NA | GBE1 | glucan (1,4-alpha-), branching enzyme 1 | 2632 | ENSG00000114480 |
| 4 | Q9UBQ7 | NA | GRHPR | glyoxylate reductase/hydroxypyruvate reductase | 9380 | ENSG00000137106 |
| 5 | P10253 | NA | GAA | glucosidase, alpha; acid | 2548 | ENSG00000171298 |
| 6 | P51649 | NA | ALDH5A1 | aldehyde dehydrogenase 5 family, member A1 | 7915 | ENSG00000112294 |
| 7 | Q16851 | NA | UGP2 | UDP-glucose pyrophosphorylase 2 | 7360 | ENSG00000169764 |
| 8 | P22570 | NA | FDXR | ferredoxin reductase | 2232 | ENSG00000161513 |
| 9 | P30044 | NA | PRDX5 | peroxiredoxin 5 | 25824 | ENSG00000126432 |
| 10 | P42765 | NA | ACAA2 | acetyl-CoA acyltransferase 2 | 10449 | ENSG00000167315 |
| 11 | P48735 | NA | IDH2 | isocitrate dehydrogenase 2 (NADP+), mitochondrial | 3418 | ENSG00000182054 |
| 12 | P49748 | NA | ACADVL | acyl-CoA dehydrogenase, very long chain | 37 | ENSG00000072778 |
| 13 | Q9P0J0 | NA | NDUFA13 | NADH dehydrogenase (ubiquinone) 1 alpha subcomplex, 13 | 51079 | ENSG00000186010 |
| 14 | Q13011 | NA | ECH1 | enoyl CoA hydratase 1, peroxisomal | 1891 | ENSG00000104823 |

  
  

| **Database:biological process      &nbspName:monosaccharide metabolic process      &nbspID:GO:0005996** | | | | | | |
| --- | --- | --- | --- | --- | --- | --- |
| C=279; O=10; E=1.63; R=6.15; rawP=4.89e-06; adjP=0.0004 | | | | | | |
| Index | UserID | Value | Gene Symbol | Gene Name | EntrezGene | Ensembl |
| 1 | Q04446 | NA | GBE1 | glucan (1,4-alpha-), branching enzyme 1 | 2632 | ENSG00000114480 |
| 2 | P10253 | NA | GAA | glucosidase, alpha; acid | 2548 | ENSG00000171298 |
| 3 | P51649 | NA | ALDH5A1 | aldehyde dehydrogenase 5 family, member A1 | 7915 | ENSG00000112294 |
| 4 | Q16851 | NA | UGP2 | UDP-glucose pyrophosphorylase 2 | 7360 | ENSG00000169764 |
| 5 | P00338 | NA | LDHA | lactate dehydrogenase A | 3939 | ENSG00000134333 |
| 6 | P16278 | NA | GLB1 | galactosidase, beta 1 | 2720 | ENSG00000170266 |
| 7 | P17174 | NA | GOT1 | glutamic-oxaloacetic transaminase 1, soluble (aspartate aminotransferase 1) | 2805 | ENSG00000120053 |
| 8 | P78417 | NA | GSTO1 | glutathione S-transferase omega 1 | 9446 | ENSG00000148834 |
| 9 | P06744 | NA | GPI | glucose-6-phosphate isomerase | 2821 | ENSG00000105220 |
| 10 | P53007 | NA | SLC25A1 | solute carrier family 25 (mitochondrial carrier; citrate transporter), member 1 | 6576 | ENSG00000100075 |

  
  

| **Database:biological process      &nbspName:hydrogen peroxide catabolic process      &nbspID:GO:0042744** | | | | | | |
| --- | --- | --- | --- | --- | --- | --- |
| C=21; O=4; E=0.12; R=32.70; rawP=5.94e-06; adjP=0.0004 | | | | | | |
| Index | UserID | Value | Gene Symbol | Gene Name | EntrezGene | Ensembl |
| 1 | P04040 | NA | CAT | catalase | 847 | ENSG00000121691 |
| 2 | Q16881 | NA | TXNRD1 | thioredoxin reductase 1 | 7296 | ENSG00000198431 |
| 3 | Q06830 | NA | PRDX1 | peroxiredoxin 1 | 5052 | ENSG00000117450 |
| 4 | P30044 | NA | PRDX5 | peroxiredoxin 5 | 25824 | ENSG00000126432 |

  
  

| **Database:biological process      &nbspName:cellular response to hydrogen peroxide      &nbspID:GO:0070301** | | | | | | |
| --- | --- | --- | --- | --- | --- | --- |
| C=47; O=5; E=0.27; R=18.26; rawP=7.52e-06; adjP=0.0005 | | | | | | |
| Index | UserID | Value | Gene Symbol | Gene Name | EntrezGene | Ensembl |
| 1 | P04040 | NA | CAT | catalase | 847 | ENSG00000121691 |
| 2 | Q16881 | NA | TXNRD1 | thioredoxin reductase 1 | 7296 | ENSG00000198431 |
| 3 | Q06830 | NA | PRDX1 | peroxiredoxin 1 | 5052 | ENSG00000117450 |
| 4 | P30044 | NA | PRDX5 | peroxiredoxin 5 | 25824 | ENSG00000126432 |
| 5 | P04083 | NA | ANXA1 | annexin A1 | 301 | ENSG00000135046 |

  
  

| **Database:biological process      &nbspName:generation of precursor metabolites and energy      &nbspID:GO:0006091** | | | | | | |
| --- | --- | --- | --- | --- | --- | --- |
| C=447; O=12; E=2.60; R=4.61; rawP=1.01e-05; adjP=0.0006 | | | | | | |
| Index | UserID | Value | Gene Symbol | Gene Name | EntrezGene | Ensembl |
| 1 | P04040 | NA | CAT | catalase | 847 | ENSG00000121691 |
| 2 | Q16881 | NA | TXNRD1 | thioredoxin reductase 1 | 7296 | ENSG00000198431 |
| 3 | Q04446 | NA | GBE1 | glucan (1,4-alpha-), branching enzyme 1 | 2632 | ENSG00000114480 |
| 4 | P10253 | NA | GAA | glucosidase, alpha; acid | 2548 | ENSG00000171298 |
| 5 | P51649 | NA | ALDH5A1 | aldehyde dehydrogenase 5 family, member A1 | 7915 | ENSG00000112294 |
| 6 | Q16851 | NA | UGP2 | UDP-glucose pyrophosphorylase 2 | 7360 | ENSG00000169764 |
| 7 | P22570 | NA | FDXR | ferredoxin reductase | 2232 | ENSG00000161513 |
| 8 | P00338 | NA | LDHA | lactate dehydrogenase A | 3939 | ENSG00000134333 |
| 9 | P48735 | NA | IDH2 | isocitrate dehydrogenase 2 (NADP+), mitochondrial | 3418 | ENSG00000182054 |
| 10 | P06744 | NA | GPI | glucose-6-phosphate isomerase | 2821 | ENSG00000105220 |
| 11 | P49748 | NA | ACADVL | acyl-CoA dehydrogenase, very long chain | 37 | ENSG00000072778 |
| 12 | Q9P0J0 | NA | NDUFA13 | NADH dehydrogenase (ubiquinone) 1 alpha subcomplex, 13 | 51079 | ENSG00000186010 |

  
  

| **Database:biological process      &nbspName:hexose metabolic process      &nbspID:GO:0019318** | | | | | | |
| --- | --- | --- | --- | --- | --- | --- |
| C=249; O=9; E=1.45; R=6.21; rawP=1.41e-05; adjP=0.0007 | | | | | | |
| Index | UserID | Value | Gene Symbol | Gene Name | EntrezGene | Ensembl |
| 1 | Q04446 | NA | GBE1 | glucan (1,4-alpha-), branching enzyme 1 | 2632 | ENSG00000114480 |
| 2 | P10253 | NA | GAA | glucosidase, alpha; acid | 2548 | ENSG00000171298 |
| 3 | P51649 | NA | ALDH5A1 | aldehyde dehydrogenase 5 family, member A1 | 7915 | ENSG00000112294 |
| 4 | Q16851 | NA | UGP2 | UDP-glucose pyrophosphorylase 2 | 7360 | ENSG00000169764 |
| 5 | P00338 | NA | LDHA | lactate dehydrogenase A | 3939 | ENSG00000134333 |
| 6 | P17174 | NA | GOT1 | glutamic-oxaloacetic transaminase 1, soluble (aspartate aminotransferase 1) | 2805 | ENSG00000120053 |
| 7 | P16278 | NA | GLB1 | galactosidase, beta 1 | 2720 | ENSG00000170266 |
| 8 | P06744 | NA | GPI | glucose-6-phosphate isomerase | 2821 | ENSG00000105220 |
| 9 | P53007 | NA | SLC25A1 | solute carrier family 25 (mitochondrial carrier; citrate transporter), member 1 | 6576 | ENSG00000100075 |

  
  

| **Database:biological process      &nbspName:carboxylic acid catabolic process      &nbspID:GO:0046395** | | | | | | |
| --- | --- | --- | --- | --- | --- | --- |
| C=198; O=8; E=1.15; R=6.94; rawP=1.96e-05; adjP=0.0008 | | | | | | |
| Index | UserID | Value | Gene Symbol | Gene Name | EntrezGene | Ensembl |
| 1 | P24752 | NA | ACAT1 | acetyl-CoA acetyltransferase 1 | 38 | ENSG00000075239 |
| 2 | P49419 | NA | ALDH7A1 | aldehyde dehydrogenase 7 family, member A1 | 501 | ENSG00000164904 |
| 3 | P17174 | NA | GOT1 | glutamic-oxaloacetic transaminase 1, soluble (aspartate aminotransferase 1) | 2805 | ENSG00000120053 |
| 4 | P00367 | NA | GLUD1 | glutamate dehydrogenase 1 | 2746 | ENSG00000148672 |
| 5 | P51649 | NA | ALDH5A1 | aldehyde dehydrogenase 5 family, member A1 | 7915 | ENSG00000112294 |
| 6 | P49748 | NA | ACADVL | acyl-CoA dehydrogenase, very long chain | 37 | ENSG00000072778 |
| 7 | Q13011 | NA | ECH1 | enoyl CoA hydratase 1, peroxisomal | 1891 | ENSG00000104823 |
| 8 | P42765 | NA | ACAA2 | acetyl-CoA acyltransferase 2 | 10449 | ENSG00000167315 |

  
  

| **Database:biological process      &nbspName:organonitrogen compound metabolic process      &nbspID:GO:1901564** | | | | | | |
| --- | --- | --- | --- | --- | --- | --- |
| C=1673; O=24; E=9.74; R=2.46; rawP=1.87e-05; adjP=0.0008 | | | | | | |
| Index | UserID | Value | Gene Symbol | Gene Name | EntrezGene | Ensembl |
| 1 | P49591 | NA | SARS | seryl-tRNA synthetase | 6301 | ENSG00000031698 |
| 2 | P00491 | NA | PNP | purine nucleoside phosphorylase | 4860 | ENSG00000198805 |
| 3 | P30086 | NA | PEBP1 | phosphatidylethanolamine binding protein 1 | 5037 | ENSG00000089220 |
| 4 | P56192 | NA | MARS | methionyl-tRNA synthetase | 4141 | ENSG00000166986 |
| 5 | P00367 | NA | GLUD1 | glutamate dehydrogenase 1 | 2746 | ENSG00000148672 |
| 6 | Q06323 | NA | PSME1 | proteasome (prosome, macropain) activator subunit 1 (PA28 alpha) | 5720 | ENSG00000092010 |
| 7 | P09211 | NA | GSTP1 | glutathione S-transferase pi 1 | 2950 | ENSG00000084207 |
| 8 | P49588 | NA | AARS | alanyl-tRNA synthetase | 16 | ENSG00000090861 |
| 9 | P43490 | NA | NAMPT | nicotinamide phosphoribosyltransferase | 10135 | ENSG00000105835 |
| 10 | P23381 | NA | WARS | tryptophanyl-tRNA synthetase | 7453 | ENSG00000140105 |
| 11 | P04040 | NA | CAT | catalase | 847 | ENSG00000121691 |
| 12 | O14773 | NA | TPP1 | tripeptidyl peptidase I | 1200 | ENSG00000166340 |
| 13 | P51649 | NA | ALDH5A1 | aldehyde dehydrogenase 5 family, member A1 | 7915 | ENSG00000112294 |
| 14 | P22102 | NA | GART | phosphoribosylglycinamide formyltransferase, phosphoribosylglycinamide synthetase, phosphoribosylaminoimidazole synthetase | 2618 | ENSG00000159131 |
| 15 | P16615 | NA | ATP2A2 | ATPase, Ca++ transporting, cardiac muscle, slow twitch 2 | 488 | ENSG00000174437 |
| 16 | P41091 | NA | EIF2S3 | eukaryotic translation initiation factor 2, subunit 3 gamma, 52kDa | 1968 | ENSG00000130741 |
| 17 | P30044 | NA | PRDX5 | peroxiredoxin 5 | 25824 | ENSG00000126432 |
| 18 | P24752 | NA | ACAT1 | acetyl-CoA acetyltransferase 1 | 38 | ENSG00000075239 |
| 19 | P13674 | NA | P4HA1 | prolyl 4-hydroxylase, alpha polypeptide I | 5033 | ENSG00000122884 |
| 20 | P16278 | NA | GLB1 | galactosidase, beta 1 | 2720 | ENSG00000170266 |
| 21 | P17174 | NA | GOT1 | glutamic-oxaloacetic transaminase 1, soluble (aspartate aminotransferase 1) | 2805 | ENSG00000120053 |
| 22 | P49419 | NA | ALDH7A1 | aldehyde dehydrogenase 7 family, member A1 | 501 | ENSG00000164904 |
| 23 | P07602 | NA | PSAP | prosaposin | 5660 | ENSG00000197746 |
| 24 | P21589 | NA | NT5E | 5'-nucleotidase, ecto (CD73) | 4907 | ENSG00000135318 |

  
  

| **Database:biological process      &nbspName:organic acid catabolic process      &nbspID:GO:0016054** | | | | | | |
| --- | --- | --- | --- | --- | --- | --- |
| C=198; O=8; E=1.15; R=6.94; rawP=1.96e-05; adjP=0.0008 | | | | | | |
| Index | UserID | Value | Gene Symbol | Gene Name | EntrezGene | Ensembl |
| 1 | P24752 | NA | ACAT1 | acetyl-CoA acetyltransferase 1 | 38 | ENSG00000075239 |
| 2 | P49419 | NA | ALDH7A1 | aldehyde dehydrogenase 7 family, member A1 | 501 | ENSG00000164904 |
| 3 | P17174 | NA | GOT1 | glutamic-oxaloacetic transaminase 1, soluble (aspartate aminotransferase 1) | 2805 | ENSG00000120053 |
| 4 | P00367 | NA | GLUD1 | glutamate dehydrogenase 1 | 2746 | ENSG00000148672 |
| 5 | P51649 | NA | ALDH5A1 | aldehyde dehydrogenase 5 family, member A1 | 7915 | ENSG00000112294 |
| 6 | P49748 | NA | ACADVL | acyl-CoA dehydrogenase, very long chain | 37 | ENSG00000072778 |
| 7 | Q13011 | NA | ECH1 | enoyl CoA hydratase 1, peroxisomal | 1891 | ENSG00000104823 |
| 8 | P42765 | NA | ACAA2 | acetyl-CoA acyltransferase 2 | 10449 | ENSG00000167315 |

  
  

| **Database:molecular function      &nbspName:catalytic activity      &nbspID:GO:0003824** | | | | | | |
| --- | --- | --- | --- | --- | --- | --- |
| C=5299; O=53; E=29.01; R=1.83; rawP=6.25e-08; adjP=3.39e-06 | | | | | | |
| Index | UserID | Value | Gene Symbol | Gene Name | EntrezGene | Ensembl |
| 1 | Q04446 | NA | GBE1 | glucan (1,4-alpha-), branching enzyme 1 | 2632 | ENSG00000114480 |
| 2 | P49591 | NA | SARS | seryl-tRNA synthetase | 6301 | ENSG00000031698 |
| 3 | P10253 | NA | GAA | glucosidase, alpha; acid | 2548 | ENSG00000171298 |
| 4 | Q16851 | NA | UGP2 | UDP-glucose pyrophosphorylase 2 | 7360 | ENSG00000169764 |
| 5 | P61225 | NA | RAP2B | RAP2B, member of RAS oncogene family | 5912 | ENSG00000181467 |
| 6 | P00367 | NA | GLUD1 | glutamate dehydrogenase 1 | 2746 | ENSG00000148672 |
| 7 | P48735 | NA | IDH2 | isocitrate dehydrogenase 2 (NADP+), mitochondrial | 3418 | ENSG00000182054 |
| 8 | P33993 | NA | MCM7 | minichromosome maintenance complex component 7 | 4176 | ENSG00000166508 |
| 9 | P09211 | NA | GSTP1 | glutathione S-transferase pi 1 | 2950 | ENSG00000084207 |
| 10 | Q9P0J0 | NA | NDUFA13 | NADH dehydrogenase (ubiquinone) 1 alpha subcomplex, 13 | 51079 | ENSG00000186010 |
| 11 | P68104 | NA | EEF1A1 | eukaryotic translation elongation factor 1 alpha 1 | 1915 | ENSG00000156508 |
| 12 | P49588 | NA | AARS | alanyl-tRNA synthetase | 16 | ENSG00000090861 |
| 13 | P02786 | NA | TFRC | transferrin receptor (p90, CD71) | 7037 | ENSG00000072274 |
| 14 | P43490 | NA | NAMPT | nicotinamide phosphoribosyltransferase | 10135 | ENSG00000105835 |
| 15 | P17655 | NA | CAPN2 | calpain 2, (m/II) large subunit | 824 | ENSG00000162909 |
| 16 | O14773 | NA | TPP1 | tripeptidyl peptidase I | 1200 | ENSG00000166340 |
| 17 | P22102 | NA | GART | phosphoribosylglycinamide formyltransferase, phosphoribosylglycinamide synthetase, phosphoribosylaminoimidazole synthetase | 2618 | ENSG00000159131 |
| 18 | P16615 | NA | ATP2A2 | ATPase, Ca++ transporting, cardiac muscle, slow twitch 2 | 488 | ENSG00000174437 |
| 19 | Q05639 | NA | EEF1A2 | eukaryotic translation elongation factor 1 alpha 2 | 1917 | ENSG00000101210 |
| 20 | P22570 | NA | FDXR | ferredoxin reductase | 2232 | ENSG00000161513 |
| 21 | P42765 | NA | ACAA2 | acetyl-CoA acyltransferase 2 | 10449 | ENSG00000167315 |
| 22 | P49419 | NA | ALDH7A1 | aldehyde dehydrogenase 7 family, member A1 | 501 | ENSG00000164904 |
| 23 | P17174 | NA | GOT1 | glutamic-oxaloacetic transaminase 1, soluble (aspartate aminotransferase 1) | 2805 | ENSG00000120053 |
| 24 | Q6PIU2 | NA | NCEH1 | neutral cholesterol ester hydrolase 1 | 57552 | ENSG00000144959 |
| 25 | P07339 | NA | CTSD | cathepsin D | 1509 | ENSG00000117984 |
| 26 | Q14697 | NA | GANAB | glucosidase, alpha; neutral AB | 23193 | ENSG00000089597 |
| 27 | P11387 | NA | TOP1 | topoisomerase (DNA) I | 7150 | ENSG00000198900 |
| 28 | P21589 | NA | NT5E | 5'-nucleotidase, ecto (CD73) | 4907 | ENSG00000135318 |
| 29 | Q16881 | NA | TXNRD1 | thioredoxin reductase 1 | 7296 | ENSG00000198431 |
| 30 | P02792 | NA | FTL | ferritin, light polypeptide | 2512 | ENSG00000087086 |
| 31 | Q9NR30 | NA | DDX21 | DEAD (Asp-Glu-Ala-Asp) box helicase 21 | 9188 | ENSG00000165732 |
| 32 | P08195 | NA | SLC3A2 | solute carrier family 3 (activators of dibasic and neutral amino acid transport), member 2 | 6520 | ENSG00000168003 |
| 33 | P00491 | NA | PNP | purine nucleoside phosphorylase | 4860 | ENSG00000198805 |
| 34 | Q71U36 | NA | TUBA1A | tubulin, alpha 1a | 7846 | ENSG00000167552 |
| 35 | P00338 | NA | LDHA | lactate dehydrogenase A | 3939 | ENSG00000134333 |
| 36 | P56192 | NA | MARS | methionyl-tRNA synthetase | 4141 | ENSG00000166986 |
| 37 | P78417 | NA | GSTO1 | glutathione S-transferase omega 1 | 9446 | ENSG00000148834 |
| 38 | P49736 | NA | MCM2 | minichromosome maintenance complex component 2 | 4171 | ENSG00000073111 |
| 39 | P38571 | NA | LIPA | lipase A, lysosomal acid, cholesterol esterase | 3988 | ENSG00000107798 |
| 40 | Q06830 | NA | PRDX1 | peroxiredoxin 1 | 5052 | ENSG00000117450 |
| 41 | Q13011 | NA | ECH1 | enoyl CoA hydratase 1, peroxisomal | 1891 | ENSG00000104823 |
| 42 | P23381 | NA | WARS | tryptophanyl-tRNA synthetase | 7453 | ENSG00000140105 |
| 43 | P04040 | NA | CAT | catalase | 847 | ENSG00000121691 |
| 44 | Q9UBQ7 | NA | GRHPR | glyoxylate reductase/hydroxypyruvate reductase | 9380 | ENSG00000137106 |
| 45 | P51649 | NA | ALDH5A1 | aldehyde dehydrogenase 5 family, member A1 | 7915 | ENSG00000112294 |
| 46 | P41091 | NA | EIF2S3 | eukaryotic translation initiation factor 2, subunit 3 gamma, 52kDa | 1968 | ENSG00000130741 |
| 47 | P30044 | NA | PRDX5 | peroxiredoxin 5 | 25824 | ENSG00000126432 |
| 48 | P24752 | NA | ACAT1 | acetyl-CoA acetyltransferase 1 | 38 | ENSG00000075239 |
| 49 | P13674 | NA | P4HA1 | prolyl 4-hydroxylase, alpha polypeptide I | 5033 | ENSG00000122884 |
| 50 | P16278 | NA | GLB1 | galactosidase, beta 1 | 2720 | ENSG00000170266 |
| 51 | P06744 | NA | GPI | glucose-6-phosphate isomerase | 2821 | ENSG00000105220 |
| 52 | P49748 | NA | ACADVL | acyl-CoA dehydrogenase, very long chain | 37 | ENSG00000072778 |
| 53 | P07384 | NA | CAPN1 | calpain 1, (mu/I) large subunit | 823 | ENSG00000014216 |

  
  

| **Database:molecular function      &nbspName:small molecule binding      &nbspID:GO:0036094** | | | | | | |
| --- | --- | --- | --- | --- | --- | --- |
| C=2595; O=35; E=14.21; R=2.46; rawP=6.52e-08; adjP=3.39e-06 | | | | | | |
| Index | UserID | Value | Gene Symbol | Gene Name | EntrezGene | Ensembl |
| 1 | Q92598 | NA | HSPH1 | heat shock 105kDa/110kDa protein 1 | 10808 | ENSG00000120694 |
| 2 | P49591 | NA | SARS | seryl-tRNA synthetase | 6301 | ENSG00000031698 |
| 3 | Q16851 | NA | UGP2 | UDP-glucose pyrophosphorylase 2 | 7360 | ENSG00000169764 |
| 4 | P30086 | NA | PEBP1 | phosphatidylethanolamine binding protein 1 | 5037 | ENSG00000089220 |
| 5 | P61225 | NA | RAP2B | RAP2B, member of RAS oncogene family | 5912 | ENSG00000181467 |
| 6 | P00367 | NA | GLUD1 | glutamate dehydrogenase 1 | 2746 | ENSG00000148672 |
| 7 | P48735 | NA | IDH2 | isocitrate dehydrogenase 2 (NADP+), mitochondrial | 3418 | ENSG00000182054 |
| 8 | Q15019 | NA | SEPT2 | septin 2 | 4735 | ENSG00000168385 |
| 9 | P33993 | NA | MCM7 | minichromosome maintenance complex component 7 | 4176 | ENSG00000166508 |
| 10 | Q9P0J0 | NA | NDUFA13 | NADH dehydrogenase (ubiquinone) 1 alpha subcomplex, 13 | 51079 | ENSG00000186010 |
| 11 | P68104 | NA | EEF1A1 | eukaryotic translation elongation factor 1 alpha 1 | 1915 | ENSG00000156508 |
| 12 | P49588 | NA | AARS | alanyl-tRNA synthetase | 16 | ENSG00000090861 |
| 13 | O00425 | NA | IGF2BP3 | insulin-like growth factor 2 mRNA binding protein 3 | 10643 | ENSG00000136231 |
| 14 | P22102 | NA | GART | phosphoribosylglycinamide formyltransferase, phosphoribosylglycinamide synthetase, phosphoribosylaminoimidazole synthetase | 2618 | ENSG00000159131 |
| 15 | P16615 | NA | ATP2A2 | ATPase, Ca++ transporting, cardiac muscle, slow twitch 2 | 488 | ENSG00000174437 |
| 16 | Q05639 | NA | EEF1A2 | eukaryotic translation elongation factor 1 alpha 2 | 1917 | ENSG00000101210 |
| 17 | P22570 | NA | FDXR | ferredoxin reductase | 2232 | ENSG00000161513 |
| 18 | P17174 | NA | GOT1 | glutamic-oxaloacetic transaminase 1, soluble (aspartate aminotransferase 1) | 2805 | ENSG00000120053 |
| 19 | P11387 | NA | TOP1 | topoisomerase (DNA) I | 7150 | ENSG00000198900 |
| 20 | P21589 | NA | NT5E | 5'-nucleotidase, ecto (CD73) | 4907 | ENSG00000135318 |
| 21 | Q16881 | NA | TXNRD1 | thioredoxin reductase 1 | 7296 | ENSG00000198431 |
| 22 | Q9NR30 | NA | DDX21 | DEAD (Asp-Glu-Ala-Asp) box helicase 21 | 9188 | ENSG00000165732 |
| 23 | Q71U36 | NA | TUBA1A | tubulin, alpha 1a | 7846 | ENSG00000167552 |
| 24 | P00491 | NA | PNP | purine nucleoside phosphorylase | 4860 | ENSG00000198805 |
| 25 | P00338 | NA | LDHA | lactate dehydrogenase A | 3939 | ENSG00000134333 |
| 26 | P56192 | NA | MARS | methionyl-tRNA synthetase | 4141 | ENSG00000166986 |
| 27 | P49736 | NA | MCM2 | minichromosome maintenance complex component 2 | 4171 | ENSG00000073111 |
| 28 | Q15056 | NA | EIF4H | eukaryotic translation initiation factor 4H | 7458 | ENSG00000106682 |
| 29 | P23381 | NA | WARS | tryptophanyl-tRNA synthetase | 7453 | ENSG00000140105 |
| 30 | P04040 | NA | CAT | catalase | 847 | ENSG00000121691 |
| 31 | Q9UBQ7 | NA | GRHPR | glyoxylate reductase/hydroxypyruvate reductase | 9380 | ENSG00000137106 |
| 32 | P02768 | NA | ALB | albumin | 213 | ENSG00000163631 |
| 33 | P41091 | NA | EIF2S3 | eukaryotic translation initiation factor 2, subunit 3 gamma, 52kDa | 1968 | ENSG00000130741 |
| 34 | P13674 | NA | P4HA1 | prolyl 4-hydroxylase, alpha polypeptide I | 5033 | ENSG00000122884 |
| 35 | P49748 | NA | ACADVL | acyl-CoA dehydrogenase, very long chain | 37 | ENSG00000072778 |

  
  

| **Database:molecular function      &nbspName:oxidoreductase activity      &nbspID:GO:0016491** | | | | | | |
| --- | --- | --- | --- | --- | --- | --- |
| C=697; O=16; E=3.82; R=4.19; rawP=9.62e-07; adjP=2.54e-05 | | | | | | |
| Index | UserID | Value | Gene Symbol | Gene Name | EntrezGene | Ensembl |
| 1 | Q16881 | NA | TXNRD1 | thioredoxin reductase 1 | 7296 | ENSG00000198431 |
| 2 | P02792 | NA | FTL | ferritin, light polypeptide | 2512 | ENSG00000087086 |
| 3 | P00338 | NA | LDHA | lactate dehydrogenase A | 3939 | ENSG00000134333 |
| 4 | P00367 | NA | GLUD1 | glutamate dehydrogenase 1 | 2746 | ENSG00000148672 |
| 5 | P48735 | NA | IDH2 | isocitrate dehydrogenase 2 (NADP+), mitochondrial | 3418 | ENSG00000182054 |
| 6 | P78417 | NA | GSTO1 | glutathione S-transferase omega 1 | 9446 | ENSG00000148834 |
| 7 | Q9P0J0 | NA | NDUFA13 | NADH dehydrogenase (ubiquinone) 1 alpha subcomplex, 13 | 51079 | ENSG00000186010 |
| 8 | Q06830 | NA | PRDX1 | peroxiredoxin 1 | 5052 | ENSG00000117450 |
| 9 | P04040 | NA | CAT | catalase | 847 | ENSG00000121691 |
| 10 | Q9UBQ7 | NA | GRHPR | glyoxylate reductase/hydroxypyruvate reductase | 9380 | ENSG00000137106 |
| 11 | P51649 | NA | ALDH5A1 | aldehyde dehydrogenase 5 family, member A1 | 7915 | ENSG00000112294 |
| 12 | P30044 | NA | PRDX5 | peroxiredoxin 5 | 25824 | ENSG00000126432 |
| 13 | P22570 | NA | FDXR | ferredoxin reductase | 2232 | ENSG00000161513 |
| 14 | P13674 | NA | P4HA1 | prolyl 4-hydroxylase, alpha polypeptide I | 5033 | ENSG00000122884 |
| 15 | P49419 | NA | ALDH7A1 | aldehyde dehydrogenase 7 family, member A1 | 501 | ENSG00000164904 |
| 16 | P49748 | NA | ACADVL | acyl-CoA dehydrogenase, very long chain | 37 | ENSG00000072778 |

  
  

| **Database:molecular function      &nbspName:antioxidant activity      &nbspID:GO:0016209** | | | | | | |
| --- | --- | --- | --- | --- | --- | --- |
| C=61; O=6; E=0.33; R=17.97; rawP=9.77e-07; adjP=2.54e-05 | | | | | | |
| Index | UserID | Value | Gene Symbol | Gene Name | EntrezGene | Ensembl |
| 1 | P04040 | NA | CAT | catalase | 847 | ENSG00000121691 |
| 2 | Q16881 | NA | TXNRD1 | thioredoxin reductase 1 | 7296 | ENSG00000198431 |
| 3 | P78417 | NA | GSTO1 | glutathione S-transferase omega 1 | 9446 | ENSG00000148834 |
| 4 | P02768 | NA | ALB | albumin | 213 | ENSG00000163631 |
| 5 | Q06830 | NA | PRDX1 | peroxiredoxin 1 | 5052 | ENSG00000117450 |
| 6 | P30044 | NA | PRDX5 | peroxiredoxin 5 | 25824 | ENSG00000126432 |

  
  

| **Database:molecular function      &nbspName:nucleotide binding      &nbspID:GO:0000166** | | | | | | |
| --- | --- | --- | --- | --- | --- | --- |
| C=2403; O=31; E=13.16; R=2.36; rawP=1.53e-06; adjP=2.67e-05 | | | | | | |
| Index | UserID | Value | Gene Symbol | Gene Name | EntrezGene | Ensembl |
| 1 | Q16881 | NA | TXNRD1 | thioredoxin reductase 1 | 7296 | ENSG00000198431 |
| 2 | Q9NR30 | NA | DDX21 | DEAD (Asp-Glu-Ala-Asp) box helicase 21 | 9188 | ENSG00000165732 |
| 3 | Q92598 | NA | HSPH1 | heat shock 105kDa/110kDa protein 1 | 10808 | ENSG00000120694 |
| 4 | P49591 | NA | SARS | seryl-tRNA synthetase | 6301 | ENSG00000031698 |
| 5 | Q71U36 | NA | TUBA1A | tubulin, alpha 1a | 7846 | ENSG00000167552 |
| 6 | Q16851 | NA | UGP2 | UDP-glucose pyrophosphorylase 2 | 7360 | ENSG00000169764 |
| 7 | P30086 | NA | PEBP1 | phosphatidylethanolamine binding protein 1 | 5037 | ENSG00000089220 |
| 8 | P00338 | NA | LDHA | lactate dehydrogenase A | 3939 | ENSG00000134333 |
| 9 | P61225 | NA | RAP2B | RAP2B, member of RAS oncogene family | 5912 | ENSG00000181467 |
| 10 | P56192 | NA | MARS | methionyl-tRNA synthetase | 4141 | ENSG00000166986 |
| 11 | P00367 | NA | GLUD1 | glutamate dehydrogenase 1 | 2746 | ENSG00000148672 |
| 12 | P48735 | NA | IDH2 | isocitrate dehydrogenase 2 (NADP+), mitochondrial | 3418 | ENSG00000182054 |
| 13 | Q15019 | NA | SEPT2 | septin 2 | 4735 | ENSG00000168385 |
| 14 | P49736 | NA | MCM2 | minichromosome maintenance complex component 2 | 4171 | ENSG00000073111 |
| 15 | Q15056 | NA | EIF4H | eukaryotic translation initiation factor 4H | 7458 | ENSG00000106682 |
| 16 | P33993 | NA | MCM7 | minichromosome maintenance complex component 7 | 4176 | ENSG00000166508 |
| 17 | Q9P0J0 | NA | NDUFA13 | NADH dehydrogenase (ubiquinone) 1 alpha subcomplex, 13 | 51079 | ENSG00000186010 |
| 18 | P68104 | NA | EEF1A1 | eukaryotic translation elongation factor 1 alpha 1 | 1915 | ENSG00000156508 |
| 19 | P49588 | NA | AARS | alanyl-tRNA synthetase | 16 | ENSG00000090861 |
| 20 | O00425 | NA | IGF2BP3 | insulin-like growth factor 2 mRNA binding protein 3 | 10643 | ENSG00000136231 |
| 21 | P23381 | NA | WARS | tryptophanyl-tRNA synthetase | 7453 | ENSG00000140105 |
| 22 | P04040 | NA | CAT | catalase | 847 | ENSG00000121691 |
| 23 | Q9UBQ7 | NA | GRHPR | glyoxylate reductase/hydroxypyruvate reductase | 9380 | ENSG00000137106 |
| 24 | P22102 | NA | GART | phosphoribosylglycinamide formyltransferase, phosphoribosylglycinamide synthetase, phosphoribosylaminoimidazole synthetase | 2618 | ENSG00000159131 |
| 25 | P16615 | NA | ATP2A2 | ATPase, Ca++ transporting, cardiac muscle, slow twitch 2 | 488 | ENSG00000174437 |
| 26 | P41091 | NA | EIF2S3 | eukaryotic translation initiation factor 2, subunit 3 gamma, 52kDa | 1968 | ENSG00000130741 |
| 27 | Q05639 | NA | EEF1A2 | eukaryotic translation elongation factor 1 alpha 2 | 1917 | ENSG00000101210 |
| 28 | P22570 | NA | FDXR | ferredoxin reductase | 2232 | ENSG00000161513 |
| 29 | P49748 | NA | ACADVL | acyl-CoA dehydrogenase, very long chain | 37 | ENSG00000072778 |
| 30 | P11387 | NA | TOP1 | topoisomerase (DNA) I | 7150 | ENSG00000198900 |
| 31 | P21589 | NA | NT5E | 5'-nucleotidase, ecto (CD73) | 4907 | ENSG00000135318 |

  
  

| **Database:molecular function      &nbspName:nucleoside phosphate binding      &nbspID:GO:1901265** | | | | | | |
| --- | --- | --- | --- | --- | --- | --- |
| C=2404; O=31; E=13.16; R=2.36; rawP=1.54e-06; adjP=2.67e-05 | | | | | | |
| Index | UserID | Value | Gene Symbol | Gene Name | EntrezGene | Ensembl |
| 1 | Q16881 | NA | TXNRD1 | thioredoxin reductase 1 | 7296 | ENSG00000198431 |
| 2 | Q9NR30 | NA | DDX21 | DEAD (Asp-Glu-Ala-Asp) box helicase 21 | 9188 | ENSG00000165732 |
| 3 | Q92598 | NA | HSPH1 | heat shock 105kDa/110kDa protein 1 | 10808 | ENSG00000120694 |
| 4 | P49591 | NA | SARS | seryl-tRNA synthetase | 6301 | ENSG00000031698 |
| 5 | Q71U36 | NA | TUBA1A | tubulin, alpha 1a | 7846 | ENSG00000167552 |
| 6 | Q16851 | NA | UGP2 | UDP-glucose pyrophosphorylase 2 | 7360 | ENSG00000169764 |
| 7 | P30086 | NA | PEBP1 | phosphatidylethanolamine binding protein 1 | 5037 | ENSG00000089220 |
| 8 | P00338 | NA | LDHA | lactate dehydrogenase A | 3939 | ENSG00000134333 |
| 9 | P61225 | NA | RAP2B | RAP2B, member of RAS oncogene family | 5912 | ENSG00000181467 |
| 10 | P56192 | NA | MARS | methionyl-tRNA synthetase | 4141 | ENSG00000166986 |
| 11 | P00367 | NA | GLUD1 | glutamate dehydrogenase 1 | 2746 | ENSG00000148672 |
| 12 | P48735 | NA | IDH2 | isocitrate dehydrogenase 2 (NADP+), mitochondrial | 3418 | ENSG00000182054 |
| 13 | Q15019 | NA | SEPT2 | septin 2 | 4735 | ENSG00000168385 |
| 14 | P49736 | NA | MCM2 | minichromosome maintenance complex component 2 | 4171 | ENSG00000073111 |
| 15 | Q15056 | NA | EIF4H | eukaryotic translation initiation factor 4H | 7458 | ENSG00000106682 |
| 16 | P33993 | NA | MCM7 | minichromosome maintenance complex component 7 | 4176 | ENSG00000166508 |
| 17 | Q9P0J0 | NA | NDUFA13 | NADH dehydrogenase (ubiquinone) 1 alpha subcomplex, 13 | 51079 | ENSG00000186010 |
| 18 | P68104 | NA | EEF1A1 | eukaryotic translation elongation factor 1 alpha 1 | 1915 | ENSG00000156508 |
| 19 | P49588 | NA | AARS | alanyl-tRNA synthetase | 16 | ENSG00000090861 |
| 20 | O00425 | NA | IGF2BP3 | insulin-like growth factor 2 mRNA binding protein 3 | 10643 | ENSG00000136231 |
| 21 | P23381 | NA | WARS | tryptophanyl-tRNA synthetase | 7453 | ENSG00000140105 |
| 22 | P04040 | NA | CAT | catalase | 847 | ENSG00000121691 |
| 23 | Q9UBQ7 | NA | GRHPR | glyoxylate reductase/hydroxypyruvate reductase | 9380 | ENSG00000137106 |
| 24 | P22102 | NA | GART | phosphoribosylglycinamide formyltransferase, phosphoribosylglycinamide synthetase, phosphoribosylaminoimidazole synthetase | 2618 | ENSG00000159131 |
| 25 | P16615 | NA | ATP2A2 | ATPase, Ca++ transporting, cardiac muscle, slow twitch 2 | 488 | ENSG00000174437 |
| 26 | P41091 | NA | EIF2S3 | eukaryotic translation initiation factor 2, subunit 3 gamma, 52kDa | 1968 | ENSG00000130741 |
| 27 | Q05639 | NA | EEF1A2 | eukaryotic translation elongation factor 1 alpha 2 | 1917 | ENSG00000101210 |
| 28 | P22570 | NA | FDXR | ferredoxin reductase | 2232 | ENSG00000161513 |
| 29 | P49748 | NA | ACADVL | acyl-CoA dehydrogenase, very long chain | 37 | ENSG00000072778 |
| 30 | P11387 | NA | TOP1 | topoisomerase (DNA) I | 7150 | ENSG00000198900 |
| 31 | P21589 | NA | NT5E | 5'-nucleotidase, ecto (CD73) | 4907 | ENSG00000135318 |

  
  

| **Database:molecular function      &nbspName:translation factor activity, nucleic acid binding      &nbspID:GO:0008135** | | | | | | |
| --- | --- | --- | --- | --- | --- | --- |
| C=79; O=6; E=0.43; R=13.87; rawP=4.52e-06; adjP=6.72e-05 | | | | | | |
| Index | UserID | Value | Gene Symbol | Gene Name | EntrezGene | Ensembl |
| 1 | Q92616 | NA | GCN1L1 | GCN1 general control of amino-acid synthesis 1-like 1 (yeast) | 10985 | ENSG00000089154 |
| 2 | P20042 | NA | EIF2S2 | eukaryotic translation initiation factor 2, subunit 2 beta, 38kDa | 8894 | ENSG00000125977 |
| 3 | P41091 | NA | EIF2S3 | eukaryotic translation initiation factor 2, subunit 3 gamma, 52kDa | 1968 | ENSG00000130741 |
| 4 | Q15056 | NA | EIF4H | eukaryotic translation initiation factor 4H | 7458 | ENSG00000106682 |
| 5 | P68104 | NA | EEF1A1 | eukaryotic translation elongation factor 1 alpha 1 | 1915 | ENSG00000156508 |
| 6 | Q05639 | NA | EEF1A2 | eukaryotic translation elongation factor 1 alpha 2 | 1917 | ENSG00000101210 |

  
  

| **Database:molecular function      &nbspName:cofactor binding      &nbspID:GO:0048037** | | | | | | |
| --- | --- | --- | --- | --- | --- | --- |
| C=259; O=9; E=1.42; R=6.35; rawP=1.17e-05; adjP=0.0002 | | | | | | |
| Index | UserID | Value | Gene Symbol | Gene Name | EntrezGene | Ensembl |
| 1 | P04040 | NA | CAT | catalase | 847 | ENSG00000121691 |
| 2 | Q16881 | NA | TXNRD1 | thioredoxin reductase 1 | 7296 | ENSG00000198431 |
| 3 | Q9UBQ7 | NA | GRHPR | glyoxylate reductase/hydroxypyruvate reductase | 9380 | ENSG00000137106 |
| 4 | P02768 | NA | ALB | albumin | 213 | ENSG00000163631 |
| 5 | P24752 | NA | ACAT1 | acetyl-CoA acetyltransferase 1 | 38 | ENSG00000075239 |
| 6 | P00367 | NA | GLUD1 | glutamate dehydrogenase 1 | 2746 | ENSG00000148672 |
| 7 | P48735 | NA | IDH2 | isocitrate dehydrogenase 2 (NADP+), mitochondrial | 3418 | ENSG00000182054 |
| 8 | P17174 | NA | GOT1 | glutamic-oxaloacetic transaminase 1, soluble (aspartate aminotransferase 1) | 2805 | ENSG00000120053 |
| 9 | P49748 | NA | ACADVL | acyl-CoA dehydrogenase, very long chain | 37 | ENSG00000072778 |

  
  

| **Database:molecular function      &nbspName:coenzyme binding      &nbspID:GO:0050662** | | | | | | |
| --- | --- | --- | --- | --- | --- | --- |
| C=184; O=7; E=1.01; R=6.95; rawP=6.59e-05; adjP=0.0007 | | | | | | |
| Index | UserID | Value | Gene Symbol | Gene Name | EntrezGene | Ensembl |
| 1 | P24752 | NA | ACAT1 | acetyl-CoA acetyltransferase 1 | 38 | ENSG00000075239 |
| 2 | P04040 | NA | CAT | catalase | 847 | ENSG00000121691 |
| 3 | Q16881 | NA | TXNRD1 | thioredoxin reductase 1 | 7296 | ENSG00000198431 |
| 4 | Q9UBQ7 | NA | GRHPR | glyoxylate reductase/hydroxypyruvate reductase | 9380 | ENSG00000137106 |
| 5 | P48735 | NA | IDH2 | isocitrate dehydrogenase 2 (NADP+), mitochondrial | 3418 | ENSG00000182054 |
| 6 | P00367 | NA | GLUD1 | glutamate dehydrogenase 1 | 2746 | ENSG00000148672 |
| 7 | P49748 | NA | ACADVL | acyl-CoA dehydrogenase, very long chain | 37 | ENSG00000072778 |

  
  

| **Database:molecular function      &nbspName:protein binding      &nbspID:GO:0005515** | | | | | | |
| --- | --- | --- | --- | --- | --- | --- |
| C=7301; O=57; E=39.97; R=1.43; rawP=0.0001; adjP=0.0007 | | | | | | |
| Index | UserID | Value | Gene Symbol | Gene Name | EntrezGene | Ensembl |
| 1 | Q92598 | NA | HSPH1 | heat shock 105kDa/110kDa protein 1 | 10808 | ENSG00000120694 |
| 2 | P08133 | NA | ANXA6 | annexin A6 | 309 | ENSG00000197043 |
| 3 | P17931 | NA | LGALS3 | lectin, galactoside-binding, soluble, 3 | 3958 | ENSG00000131981 |
| 4 | Q16851 | NA | UGP2 | UDP-glucose pyrophosphorylase 2 | 7360 | ENSG00000169764 |
| 5 | P30086 | NA | PEBP1 | phosphatidylethanolamine binding protein 1 | 5037 | ENSG00000089220 |
| 6 | P61225 | NA | RAP2B | RAP2B, member of RAS oncogene family | 5912 | ENSG00000181467 |
| 7 | P55060 | NA | CSE1L | CSE1 chromosome segregation 1-like (yeast) | 1434 | ENSG00000124207 |
| 8 | Q9NQC3 | NA | RTN4 | reticulon 4 | 57142 | ENSG00000115310 |
| 9 | P00367 | NA | GLUD1 | glutamate dehydrogenase 1 | 2746 | ENSG00000148672 |
| 10 | Q15019 | NA | SEPT2 | septin 2 | 4735 | ENSG00000168385 |
| 11 | P33993 | NA | MCM7 | minichromosome maintenance complex component 7 | 4176 | ENSG00000166508 |
| 12 | P09211 | NA | GSTP1 | glutathione S-transferase pi 1 | 2950 | ENSG00000084207 |
| 13 | P16949 | NA | STMN1 | stathmin 1 | 3925 | ENSG00000117632 |
| 14 | Q9P0J0 | NA | NDUFA13 | NADH dehydrogenase (ubiquinone) 1 alpha subcomplex, 13 | 51079 | ENSG00000186010 |
| 15 | P68104 | NA | EEF1A1 | eukaryotic translation elongation factor 1 alpha 1 | 1915 | ENSG00000156508 |
| 16 | Q86UP2 | NA | KTN1 | kinectin 1 (kinesin receptor) | 3895 | ENSG00000126777 |
| 17 | P02786 | NA | TFRC | transferrin receptor (p90, CD71) | 7037 | ENSG00000072274 |
| 18 | O75369 | NA | FLNB | filamin B, beta | 2317 | ENSG00000136068 |
| 19 | P43490 | NA | NAMPT | nicotinamide phosphoribosyltransferase | 10135 | ENSG00000105835 |
| 20 | O00425 | NA | IGF2BP3 | insulin-like growth factor 2 mRNA binding protein 3 | 10643 | ENSG00000136231 |
| 21 | P17655 | NA | CAPN2 | calpain 2, (m/II) large subunit | 824 | ENSG00000162909 |
| 22 | O14773 | NA | TPP1 | tripeptidyl peptidase I | 1200 | ENSG00000166340 |
| 23 | P20042 | NA | EIF2S2 | eukaryotic translation initiation factor 2, subunit 2 beta, 38kDa | 8894 | ENSG00000125977 |
| 24 | P16615 | NA | ATP2A2 | ATPase, Ca++ transporting, cardiac muscle, slow twitch 2 | 488 | ENSG00000174437 |
| 25 | P22570 | NA | FDXR | ferredoxin reductase | 2232 | ENSG00000161513 |
| 26 | P40121 | NA | CAPG | capping protein (actin filament), gelsolin-like | 822 | ENSG00000042493 |
| 27 | P42765 | NA | ACAA2 | acetyl-CoA acyltransferase 2 | 10449 | ENSG00000167315 |
| 28 | P49419 | NA | ALDH7A1 | aldehyde dehydrogenase 7 family, member A1 | 501 | ENSG00000164904 |
| 29 | P37802 | NA | TAGLN2 | transgelin 2 | 8407 | ENSG00000158710 |
| 30 | P08727 | NA | KRT19 | keratin 19 | 3880 | ENSG00000171345 |
| 31 | Q14764 | NA | MVP | major vault protein | 9961 | ENSG00000013364 |
| 32 | P18206 | NA | VCL | vinculin | 7414 | ENSG00000035403 |
| 33 | P11387 | NA | TOP1 | topoisomerase (DNA) I | 7150 | ENSG00000198900 |
| 34 | P02792 | NA | FTL | ferritin, light polypeptide | 2512 | ENSG00000087086 |
| 35 | P08195 | NA | SLC3A2 | solute carrier family 3 (activators of dibasic and neutral amino acid transport), member 2 | 6520 | ENSG00000168003 |
| 36 | P49321 | NA | NASP | nuclear autoantigenic sperm protein (histone-binding) | 4678 | ENSG00000132780 |
| 37 | Q71U36 | NA | TUBA1A | tubulin, alpha 1a | 7846 | ENSG00000167552 |
| 38 | Q9HAV4 | NA | XPO5 | exportin 5 | 57510 | ENSG00000124571 |
| 39 | P49736 | NA | MCM2 | minichromosome maintenance complex component 2 | 4171 | ENSG00000073111 |
| 40 | Q15056 | NA | EIF4H | eukaryotic translation initiation factor 4H | 7458 | ENSG00000106682 |
| 41 | O60610 | NA | DIAPH1 | diaphanous homolog 1 (Drosophila) | 1729 | ENSG00000131504 |
| 42 | Q06830 | NA | PRDX1 | peroxiredoxin 1 | 5052 | ENSG00000117450 |
| 43 | Q13011 | NA | ECH1 | enoyl CoA hydratase 1, peroxisomal | 1891 | ENSG00000104823 |
| 44 | P04083 | NA | ANXA1 | annexin A1 | 301 | ENSG00000135046 |
| 45 | P23381 | NA | WARS | tryptophanyl-tRNA synthetase | 7453 | ENSG00000140105 |
| 46 | P04040 | NA | CAT | catalase | 847 | ENSG00000121691 |
| 47 | Q14980 | NA | NUMA1 | nuclear mitotic apparatus protein 1 | 4926 | ENSG00000137497 |
| 48 | Q9UBQ7 | NA | GRHPR | glyoxylate reductase/hydroxypyruvate reductase | 9380 | ENSG00000137106 |
| 49 | P51649 | NA | ALDH5A1 | aldehyde dehydrogenase 5 family, member A1 | 7915 | ENSG00000112294 |
| 50 | P41091 | NA | EIF2S3 | eukaryotic translation initiation factor 2, subunit 3 gamma, 52kDa | 1968 | ENSG00000130741 |
| 51 | P02768 | NA | ALB | albumin | 213 | ENSG00000163631 |
| 52 | P30044 | NA | PRDX5 | peroxiredoxin 5 | 25824 | ENSG00000126432 |
| 53 | P05556 | NA | ITGB1 | integrin, beta 1 (fibronectin receptor, beta polypeptide, antigen CD29 includes MDF2, MSK12) | 3688 | ENSG00000150093 |
| 54 | P24752 | NA | ACAT1 | acetyl-CoA acetyltransferase 1 | 38 | ENSG00000075239 |
| 55 | P16278 | NA | GLB1 | galactosidase, beta 1 | 2720 | ENSG00000170266 |
| 56 | P06744 | NA | GPI | glucose-6-phosphate isomerase | 2821 | ENSG00000105220 |
| 57 | P07384 | NA | CAPN1 | calpain 1, (mu/I) large subunit | 823 | ENSG00000014216 |

  
  

| **Database:molecular function      &nbspName:ligase activity, forming aminoacyl-tRNA and related compounds      &nbspID:GO:0016876** | | | | | | |
| --- | --- | --- | --- | --- | --- | --- |
| C=44; O=4; E=0.24; R=16.60; rawP=9.59e-05; adjP=0.0007 | | | | | | |
| Index | UserID | Value | Gene Symbol | Gene Name | EntrezGene | Ensembl |
| 1 | P56192 | NA | MARS | methionyl-tRNA synthetase | 4141 | ENSG00000166986 |
| 2 | P49591 | NA | SARS | seryl-tRNA synthetase | 6301 | ENSG00000031698 |
| 3 | P49588 | NA | AARS | alanyl-tRNA synthetase | 16 | ENSG00000090861 |
| 4 | P23381 | NA | WARS | tryptophanyl-tRNA synthetase | 7453 | ENSG00000140105 |

  
  

| **Database:molecular function      &nbspName:aminoacyl-tRNA ligase activity      &nbspID:GO:0004812** | | | | | | |
| --- | --- | --- | --- | --- | --- | --- |
| C=44; O=4; E=0.24; R=16.60; rawP=9.59e-05; adjP=0.0007 | | | | | | |
| Index | UserID | Value | Gene Symbol | Gene Name | EntrezGene | Ensembl |
| 1 | P56192 | NA | MARS | methionyl-tRNA synthetase | 4141 | ENSG00000166986 |
| 2 | P49591 | NA | SARS | seryl-tRNA synthetase | 6301 | ENSG00000031698 |
| 3 | P49588 | NA | AARS | alanyl-tRNA synthetase | 16 | ENSG00000090861 |
| 4 | P23381 | NA | WARS | tryptophanyl-tRNA synthetase | 7453 | ENSG00000140105 |

  
  

| **Database:molecular function      &nbspName:anion binding      &nbspID:GO:0043168** | | | | | | |
| --- | --- | --- | --- | --- | --- | --- |
| C=2371; O=27; E=12.98; R=2.08; rawP=9.52e-05; adjP=0.0007 | | | | | | |
| Index | UserID | Value | Gene Symbol | Gene Name | EntrezGene | Ensembl |
| 1 | Q16881 | NA | TXNRD1 | thioredoxin reductase 1 | 7296 | ENSG00000198431 |
| 2 | Q9NR30 | NA | DDX21 | DEAD (Asp-Glu-Ala-Asp) box helicase 21 | 9188 | ENSG00000165732 |
| 3 | Q92598 | NA | HSPH1 | heat shock 105kDa/110kDa protein 1 | 10808 | ENSG00000120694 |
| 4 | P49591 | NA | SARS | seryl-tRNA synthetase | 6301 | ENSG00000031698 |
| 5 | P00491 | NA | PNP | purine nucleoside phosphorylase | 4860 | ENSG00000198805 |
| 6 | Q71U36 | NA | TUBA1A | tubulin, alpha 1a | 7846 | ENSG00000167552 |
| 7 | P30086 | NA | PEBP1 | phosphatidylethanolamine binding protein 1 | 5037 | ENSG00000089220 |
| 8 | P61225 | NA | RAP2B | RAP2B, member of RAS oncogene family | 5912 | ENSG00000181467 |
| 9 | P56192 | NA | MARS | methionyl-tRNA synthetase | 4141 | ENSG00000166986 |
| 10 | P00367 | NA | GLUD1 | glutamate dehydrogenase 1 | 2746 | ENSG00000148672 |
| 11 | Q15019 | NA | SEPT2 | septin 2 | 4735 | ENSG00000168385 |
| 12 | P49736 | NA | MCM2 | minichromosome maintenance complex component 2 | 4171 | ENSG00000073111 |
| 13 | P33993 | NA | MCM7 | minichromosome maintenance complex component 7 | 4176 | ENSG00000166508 |
| 14 | Q9P0J0 | NA | NDUFA13 | NADH dehydrogenase (ubiquinone) 1 alpha subcomplex, 13 | 51079 | ENSG00000186010 |
| 15 | P68104 | NA | EEF1A1 | eukaryotic translation elongation factor 1 alpha 1 | 1915 | ENSG00000156508 |
| 16 | P49588 | NA | AARS | alanyl-tRNA synthetase | 16 | ENSG00000090861 |
| 17 | P23381 | NA | WARS | tryptophanyl-tRNA synthetase | 7453 | ENSG00000140105 |
| 18 | Q9UBQ7 | NA | GRHPR | glyoxylate reductase/hydroxypyruvate reductase | 9380 | ENSG00000137106 |
| 19 | P22102 | NA | GART | phosphoribosylglycinamide formyltransferase, phosphoribosylglycinamide synthetase, phosphoribosylaminoimidazole synthetase | 2618 | ENSG00000159131 |
| 20 | P16615 | NA | ATP2A2 | ATPase, Ca++ transporting, cardiac muscle, slow twitch 2 | 488 | ENSG00000174437 |
| 21 | P41091 | NA | EIF2S3 | eukaryotic translation initiation factor 2, subunit 3 gamma, 52kDa | 1968 | ENSG00000130741 |
| 22 | P02768 | NA | ALB | albumin | 213 | ENSG00000163631 |
| 23 | Q05639 | NA | EEF1A2 | eukaryotic translation elongation factor 1 alpha 2 | 1917 | ENSG00000101210 |
| 24 | P13674 | NA | P4HA1 | prolyl 4-hydroxylase, alpha polypeptide I | 5033 | ENSG00000122884 |
| 25 | P17174 | NA | GOT1 | glutamic-oxaloacetic transaminase 1, soluble (aspartate aminotransferase 1) | 2805 | ENSG00000120053 |
| 26 | P49748 | NA | ACADVL | acyl-CoA dehydrogenase, very long chain | 37 | ENSG00000072778 |
| 27 | P11387 | NA | TOP1 | topoisomerase (DNA) I | 7150 | ENSG00000198900 |

  
  

| **Database:molecular function      &nbspName:ligase activity, forming carbon-oxygen bonds      &nbspID:GO:0016875** | | | | | | |
| --- | --- | --- | --- | --- | --- | --- |
| C=44; O=4; E=0.24; R=16.60; rawP=9.59e-05; adjP=0.0007 | | | | | | |
| Index | UserID | Value | Gene Symbol | Gene Name | EntrezGene | Ensembl |
| 1 | P56192 | NA | MARS | methionyl-tRNA synthetase | 4141 | ENSG00000166986 |
| 2 | P49591 | NA | SARS | seryl-tRNA synthetase | 6301 | ENSG00000031698 |
| 3 | P49588 | NA | AARS | alanyl-tRNA synthetase | 16 | ENSG00000090861 |
| 4 | P23381 | NA | WARS | tryptophanyl-tRNA synthetase | 7453 | ENSG00000140105 |

  
  

| **Database:cellular component      &nbspName:cytoplasmic part      &nbspID:GO:0044444** | | | | | | |
| --- | --- | --- | --- | --- | --- | --- |
| C=6728; O=73; E=34.68; R=2.10; rawP=1.63e-17; adjP=1.70e-15 | | | | | | |
| Index | UserID | Value | Gene Symbol | Gene Name | EntrezGene | Ensembl |
| 1 | Q04446 | NA | GBE1 | glucan (1,4-alpha-), branching enzyme 1 | 2632 | ENSG00000114480 |
| 2 | P10253 | NA | GAA | glucosidase, alpha; acid | 2548 | ENSG00000171298 |
| 3 | Q16851 | NA | UGP2 | UDP-glucose pyrophosphorylase 2 | 7360 | ENSG00000169764 |
| 4 | P30086 | NA | PEBP1 | phosphatidylethanolamine binding protein 1 | 5037 | ENSG00000089220 |
| 5 | P00367 | NA | GLUD1 | glutamate dehydrogenase 1 | 2746 | ENSG00000148672 |
| 6 | Q06323 | NA | PSME1 | proteasome (prosome, macropain) activator subunit 1 (PA28 alpha) | 5720 | ENSG00000092010 |
| 7 | P48735 | NA | IDH2 | isocitrate dehydrogenase 2 (NADP+), mitochondrial | 3418 | ENSG00000182054 |
| 8 | P16949 | NA | STMN1 | stathmin 1 | 3925 | ENSG00000117632 |
| 9 | Q9P0J0 | NA | NDUFA13 | NADH dehydrogenase (ubiquinone) 1 alpha subcomplex, 13 | 51079 | ENSG00000186010 |
| 10 | Q86UP2 | NA | KTN1 | kinectin 1 (kinesin receptor) | 3895 | ENSG00000126777 |
| 11 | P49588 | NA | AARS | alanyl-tRNA synthetase | 16 | ENSG00000090861 |
| 12 | O75369 | NA | FLNB | filamin B, beta | 2317 | ENSG00000136068 |
| 13 | O14773 | NA | TPP1 | tripeptidyl peptidase I | 1200 | ENSG00000166340 |
| 14 | P16615 | NA | ATP2A2 | ATPase, Ca++ transporting, cardiac muscle, slow twitch 2 | 488 | ENSG00000174437 |
| 15 | P40121 | NA | CAPG | capping protein (actin filament), gelsolin-like | 822 | ENSG00000042493 |
| 16 | P42765 | NA | ACAA2 | acetyl-CoA acyltransferase 2 | 10449 | ENSG00000167315 |
| 17 | P17174 | NA | GOT1 | glutamic-oxaloacetic transaminase 1, soluble (aspartate aminotransferase 1) | 2805 | ENSG00000120053 |
| 18 | P08727 | NA | KRT19 | keratin 19 | 3880 | ENSG00000171345 |
| 19 | Q14697 | NA | GANAB | glucosidase, alpha; neutral AB | 23193 | ENSG00000089597 |
| 20 | P18206 | NA | VCL | vinculin | 7414 | ENSG00000035403 |
| 21 | P08195 | NA | SLC3A2 | solute carrier family 3 (activators of dibasic and neutral amino acid transport), member 2 | 6520 | ENSG00000168003 |
| 22 | Q71U36 | NA | TUBA1A | tubulin, alpha 1a | 7846 | ENSG00000167552 |
| 23 | P78417 | NA | GSTO1 | glutathione S-transferase omega 1 | 9446 | ENSG00000148834 |
| 24 | P38571 | NA | LIPA | lipase A, lysosomal acid, cholesterol esterase | 3988 | ENSG00000107798 |
| 25 | P23381 | NA | WARS | tryptophanyl-tRNA synthetase | 7453 | ENSG00000140105 |
| 26 | Q14980 | NA | NUMA1 | nuclear mitotic apparatus protein 1 | 4926 | ENSG00000137497 |
| 27 | Q9UBQ7 | NA | GRHPR | glyoxylate reductase/hydroxypyruvate reductase | 9380 | ENSG00000137106 |
| 28 | P51649 | NA | ALDH5A1 | aldehyde dehydrogenase 5 family, member A1 | 7915 | ENSG00000112294 |
| 29 | P30044 | NA | PRDX5 | peroxiredoxin 5 | 25824 | ENSG00000126432 |
| 30 | P05556 | NA | ITGB1 | integrin, beta 1 (fibronectin receptor, beta polypeptide, antigen CD29 includes MDF2, MSK12) | 3688 | ENSG00000150093 |
| 31 | P13674 | NA | P4HA1 | prolyl 4-hydroxylase, alpha polypeptide I | 5033 | ENSG00000122884 |
| 32 | P49591 | NA | SARS | seryl-tRNA synthetase | 6301 | ENSG00000031698 |
| 33 | P17931 | NA | LGALS3 | lectin, galactoside-binding, soluble, 3 | 3958 | ENSG00000131981 |
| 34 | P08133 | NA | ANXA6 | annexin A6 | 309 | ENSG00000197043 |
| 35 | P61225 | NA | RAP2B | RAP2B, member of RAS oncogene family | 5912 | ENSG00000181467 |
| 36 | Q9NQC3 | NA | RTN4 | reticulon 4 | 57142 | ENSG00000115310 |
| 37 | Q15019 | NA | SEPT2 | septin 2 | 4735 | ENSG00000168385 |
| 38 | P09211 | NA | GSTP1 | glutathione S-transferase pi 1 | 2950 | ENSG00000084207 |
| 39 | P68104 | NA | EEF1A1 | eukaryotic translation elongation factor 1 alpha 1 | 1915 | ENSG00000156508 |
| 40 | P02786 | NA | TFRC | transferrin receptor (p90, CD71) | 7037 | ENSG00000072274 |
| 41 | O00425 | NA | IGF2BP3 | insulin-like growth factor 2 mRNA binding protein 3 | 10643 | ENSG00000136231 |
| 42 | P43490 | NA | NAMPT | nicotinamide phosphoribosyltransferase | 10135 | ENSG00000105835 |
| 43 | Q9P258 | NA | RCC2 | regulator of chromosome condensation 2 | 55920 | ENSG00000179051 |
| 44 | P20042 | NA | EIF2S2 | eukaryotic translation initiation factor 2, subunit 2 beta, 38kDa | 8894 | ENSG00000125977 |
| 45 | P22102 | NA | GART | phosphoribosylglycinamide formyltransferase, phosphoribosylglycinamide synthetase, phosphoribosylaminoimidazole synthetase | 2618 | ENSG00000159131 |
| 46 | P22570 | NA | FDXR | ferredoxin reductase | 2232 | ENSG00000161513 |
| 47 | Q05639 | NA | EEF1A2 | eukaryotic translation elongation factor 1 alpha 2 | 1917 | ENSG00000101210 |
| 48 | P49419 | NA | ALDH7A1 | aldehyde dehydrogenase 7 family, member A1 | 501 | ENSG00000164904 |
| 49 | Q92616 | NA | GCN1L1 | GCN1 general control of amino-acid synthesis 1-like 1 (yeast) | 10985 | ENSG00000089154 |
| 50 | P07339 | NA | CTSD | cathepsin D | 1509 | ENSG00000117984 |
| 51 | Q6PIU2 | NA | NCEH1 | neutral cholesterol ester hydrolase 1 | 57552 | ENSG00000144959 |
| 52 | P11387 | NA | TOP1 | topoisomerase (DNA) I | 7150 | ENSG00000198900 |
| 53 | P53007 | NA | SLC25A1 | solute carrier family 25 (mitochondrial carrier; citrate transporter), member 1 | 6576 | ENSG00000100075 |
| 54 | Q9BPW8 | NA | NIPSNAP1 | nipsnap homolog 1 (C. elegans) | 8508 | ENSG00000184117 |
| 55 | Q16881 | NA | TXNRD1 | thioredoxin reductase 1 | 7296 | ENSG00000198431 |
| 56 | P02792 | NA | FTL | ferritin, light polypeptide | 2512 | ENSG00000087086 |
| 57 | P00491 | NA | PNP | purine nucleoside phosphorylase | 4860 | ENSG00000198805 |
| 58 | Q9HAV4 | NA | XPO5 | exportin 5 | 57510 | ENSG00000124571 |
| 59 | P00338 | NA | LDHA | lactate dehydrogenase A | 3939 | ENSG00000134333 |
| 60 | P56192 | NA | MARS | methionyl-tRNA synthetase | 4141 | ENSG00000166986 |
| 61 | Q15056 | NA | EIF4H | eukaryotic translation initiation factor 4H | 7458 | ENSG00000106682 |
| 62 | Q13011 | NA | ECH1 | enoyl CoA hydratase 1, peroxisomal | 1891 | ENSG00000104823 |
| 63 | Q06830 | NA | PRDX1 | peroxiredoxin 1 | 5052 | ENSG00000117450 |
| 64 | P04083 | NA | ANXA1 | annexin A1 | 301 | ENSG00000135046 |
| 65 | P04040 | NA | CAT | catalase | 847 | ENSG00000121691 |
| 66 | P41091 | NA | EIF2S3 | eukaryotic translation initiation factor 2, subunit 3 gamma, 52kDa | 1968 | ENSG00000130741 |
| 67 | P02768 | NA | ALB | albumin | 213 | ENSG00000163631 |
| 68 | P24752 | NA | ACAT1 | acetyl-CoA acetyltransferase 1 | 38 | ENSG00000075239 |
| 69 | P16278 | NA | GLB1 | galactosidase, beta 1 | 2720 | ENSG00000170266 |
| 70 | P07602 | NA | PSAP | prosaposin | 5660 | ENSG00000197746 |
| 71 | P06744 | NA | GPI | glucose-6-phosphate isomerase | 2821 | ENSG00000105220 |
| 72 | Q15758 | NA | SLC1A5 | solute carrier family 1 (neutral amino acid transporter), member 5 | 6510 | ENSG00000105281 |
| 73 | P49748 | NA | ACADVL | acyl-CoA dehydrogenase, very long chain | 37 | ENSG00000072778 |

  
  

| **Database:cellular component      &nbspName:cytoplasm      &nbspID:GO:0005737** | | | | | | |
| --- | --- | --- | --- | --- | --- | --- |
| C=9051; O=81; E=46.66; R=1.74; rawP=2.02e-16; adjP=1.05e-14 | | | | | | |
| Index | UserID | Value | Gene Symbol | Gene Name | EntrezGene | Ensembl |
| 1 | Q04446 | NA | GBE1 | glucan (1,4-alpha-), branching enzyme 1 | 2632 | ENSG00000114480 |
| 2 | P10253 | NA | GAA | glucosidase, alpha; acid | 2548 | ENSG00000171298 |
| 3 | Q16851 | NA | UGP2 | UDP-glucose pyrophosphorylase 2 | 7360 | ENSG00000169764 |
| 4 | P30086 | NA | PEBP1 | phosphatidylethanolamine binding protein 1 | 5037 | ENSG00000089220 |
| 5 | P55060 | NA | CSE1L | CSE1 chromosome segregation 1-like (yeast) | 1434 | ENSG00000124207 |
| 6 | P00367 | NA | GLUD1 | glutamate dehydrogenase 1 | 2746 | ENSG00000148672 |
| 7 | Q06323 | NA | PSME1 | proteasome (prosome, macropain) activator subunit 1 (PA28 alpha) | 5720 | ENSG00000092010 |
| 8 | P48735 | NA | IDH2 | isocitrate dehydrogenase 2 (NADP+), mitochondrial | 3418 | ENSG00000182054 |
| 9 | Q9P0J0 | NA | NDUFA13 | NADH dehydrogenase (ubiquinone) 1 alpha subcomplex, 13 | 51079 | ENSG00000186010 |
| 10 | P16949 | NA | STMN1 | stathmin 1 | 3925 | ENSG00000117632 |
| 11 | Q86UP2 | NA | KTN1 | kinectin 1 (kinesin receptor) | 3895 | ENSG00000126777 |
| 12 | P49588 | NA | AARS | alanyl-tRNA synthetase | 16 | ENSG00000090861 |
| 13 | O75369 | NA | FLNB | filamin B, beta | 2317 | ENSG00000136068 |
| 14 | O14773 | NA | TPP1 | tripeptidyl peptidase I | 1200 | ENSG00000166340 |
| 15 | P16615 | NA | ATP2A2 | ATPase, Ca++ transporting, cardiac muscle, slow twitch 2 | 488 | ENSG00000174437 |
| 16 | P42765 | NA | ACAA2 | acetyl-CoA acyltransferase 2 | 10449 | ENSG00000167315 |
| 17 | P40121 | NA | CAPG | capping protein (actin filament), gelsolin-like | 822 | ENSG00000042493 |
| 18 | P17174 | NA | GOT1 | glutamic-oxaloacetic transaminase 1, soluble (aspartate aminotransferase 1) | 2805 | ENSG00000120053 |
| 19 | P08727 | NA | KRT19 | keratin 19 | 3880 | ENSG00000171345 |
| 20 | Q14697 | NA | GANAB | glucosidase, alpha; neutral AB | 23193 | ENSG00000089597 |
| 21 | P18206 | NA | VCL | vinculin | 7414 | ENSG00000035403 |
| 22 | P08195 | NA | SLC3A2 | solute carrier family 3 (activators of dibasic and neutral amino acid transport), member 2 | 6520 | ENSG00000168003 |
| 23 | Q71U36 | NA | TUBA1A | tubulin, alpha 1a | 7846 | ENSG00000167552 |
| 24 | P78417 | NA | GSTO1 | glutathione S-transferase omega 1 | 9446 | ENSG00000148834 |
| 25 | P38571 | NA | LIPA | lipase A, lysosomal acid, cholesterol esterase | 3988 | ENSG00000107798 |
| 26 | P23381 | NA | WARS | tryptophanyl-tRNA synthetase | 7453 | ENSG00000140105 |
| 27 | Q14980 | NA | NUMA1 | nuclear mitotic apparatus protein 1 | 4926 | ENSG00000137497 |
| 28 | Q9UBQ7 | NA | GRHPR | glyoxylate reductase/hydroxypyruvate reductase | 9380 | ENSG00000137106 |
| 29 | P51649 | NA | ALDH5A1 | aldehyde dehydrogenase 5 family, member A1 | 7915 | ENSG00000112294 |
| 30 | P30044 | NA | PRDX5 | peroxiredoxin 5 | 25824 | ENSG00000126432 |
| 31 | P05556 | NA | ITGB1 | integrin, beta 1 (fibronectin receptor, beta polypeptide, antigen CD29 includes MDF2, MSK12) | 3688 | ENSG00000150093 |
| 32 | P13674 | NA | P4HA1 | prolyl 4-hydroxylase, alpha polypeptide I | 5033 | ENSG00000122884 |
| 33 | P07384 | NA | CAPN1 | calpain 1, (mu/I) large subunit | 823 | ENSG00000014216 |
| 34 | P49591 | NA | SARS | seryl-tRNA synthetase | 6301 | ENSG00000031698 |
| 35 | Q92598 | NA | HSPH1 | heat shock 105kDa/110kDa protein 1 | 10808 | ENSG00000120694 |
| 36 | P17931 | NA | LGALS3 | lectin, galactoside-binding, soluble, 3 | 3958 | ENSG00000131981 |
| 37 | P08133 | NA | ANXA6 | annexin A6 | 309 | ENSG00000197043 |
| 38 | P61225 | NA | RAP2B | RAP2B, member of RAS oncogene family | 5912 | ENSG00000181467 |
| 39 | Q9NQC3 | NA | RTN4 | reticulon 4 | 57142 | ENSG00000115310 |
| 40 | Q15019 | NA | SEPT2 | septin 2 | 4735 | ENSG00000168385 |
| 41 | P09211 | NA | GSTP1 | glutathione S-transferase pi 1 | 2950 | ENSG00000084207 |
| 42 | P68104 | NA | EEF1A1 | eukaryotic translation elongation factor 1 alpha 1 | 1915 | ENSG00000156508 |
| 43 | P02786 | NA | TFRC | transferrin receptor (p90, CD71) | 7037 | ENSG00000072274 |
| 44 | O00425 | NA | IGF2BP3 | insulin-like growth factor 2 mRNA binding protein 3 | 10643 | ENSG00000136231 |
| 45 | P43490 | NA | NAMPT | nicotinamide phosphoribosyltransferase | 10135 | ENSG00000105835 |
| 46 | P17655 | NA | CAPN2 | calpain 2, (m/II) large subunit | 824 | ENSG00000162909 |
| 47 | Q9P258 | NA | RCC2 | regulator of chromosome condensation 2 | 55920 | ENSG00000179051 |
| 48 | P20042 | NA | EIF2S2 | eukaryotic translation initiation factor 2, subunit 2 beta, 38kDa | 8894 | ENSG00000125977 |
| 49 | P22102 | NA | GART | phosphoribosylglycinamide formyltransferase, phosphoribosylglycinamide synthetase, phosphoribosylaminoimidazole synthetase | 2618 | ENSG00000159131 |
| 50 | P22570 | NA | FDXR | ferredoxin reductase | 2232 | ENSG00000161513 |
| 51 | Q05639 | NA | EEF1A2 | eukaryotic translation elongation factor 1 alpha 2 | 1917 | ENSG00000101210 |
| 52 | P49419 | NA | ALDH7A1 | aldehyde dehydrogenase 7 family, member A1 | 501 | ENSG00000164904 |
| 53 | Q92616 | NA | GCN1L1 | GCN1 general control of amino-acid synthesis 1-like 1 (yeast) | 10985 | ENSG00000089154 |
| 54 | P07339 | NA | CTSD | cathepsin D | 1509 | ENSG00000117984 |
| 55 | Q14764 | NA | MVP | major vault protein | 9961 | ENSG00000013364 |
| 56 | Q6PIU2 | NA | NCEH1 | neutral cholesterol ester hydrolase 1 | 57552 | ENSG00000144959 |
| 57 | P21589 | NA | NT5E | 5'-nucleotidase, ecto (CD73) | 4907 | ENSG00000135318 |
| 58 | P11387 | NA | TOP1 | topoisomerase (DNA) I | 7150 | ENSG00000198900 |
| 59 | P53007 | NA | SLC25A1 | solute carrier family 25 (mitochondrial carrier; citrate transporter), member 1 | 6576 | ENSG00000100075 |
| 60 | Q9BPW8 | NA | NIPSNAP1 | nipsnap homolog 1 (C. elegans) | 8508 | ENSG00000184117 |
| 61 | Q16881 | NA | TXNRD1 | thioredoxin reductase 1 | 7296 | ENSG00000198431 |
| 62 | P02792 | NA | FTL | ferritin, light polypeptide | 2512 | ENSG00000087086 |
| 63 | P00491 | NA | PNP | purine nucleoside phosphorylase | 4860 | ENSG00000198805 |
| 64 | P49321 | NA | NASP | nuclear autoantigenic sperm protein (histone-binding) | 4678 | ENSG00000132780 |
| 65 | Q9HAV4 | NA | XPO5 | exportin 5 | 57510 | ENSG00000124571 |
| 66 | P00338 | NA | LDHA | lactate dehydrogenase A | 3939 | ENSG00000134333 |
| 67 | P56192 | NA | MARS | methionyl-tRNA synthetase | 4141 | ENSG00000166986 |
| 68 | Q15056 | NA | EIF4H | eukaryotic translation initiation factor 4H | 7458 | ENSG00000106682 |
| 69 | O60610 | NA | DIAPH1 | diaphanous homolog 1 (Drosophila) | 1729 | ENSG00000131504 |
| 70 | Q06830 | NA | PRDX1 | peroxiredoxin 1 | 5052 | ENSG00000117450 |
| 71 | Q13011 | NA | ECH1 | enoyl CoA hydratase 1, peroxisomal | 1891 | ENSG00000104823 |
| 72 | P04083 | NA | ANXA1 | annexin A1 | 301 | ENSG00000135046 |
| 73 | P04040 | NA | CAT | catalase | 847 | ENSG00000121691 |
| 74 | P41091 | NA | EIF2S3 | eukaryotic translation initiation factor 2, subunit 3 gamma, 52kDa | 1968 | ENSG00000130741 |
| 75 | P02768 | NA | ALB | albumin | 213 | ENSG00000163631 |
| 76 | P24752 | NA | ACAT1 | acetyl-CoA acetyltransferase 1 | 38 | ENSG00000075239 |
| 77 | P16278 | NA | GLB1 | galactosidase, beta 1 | 2720 | ENSG00000170266 |
| 78 | P07602 | NA | PSAP | prosaposin | 5660 | ENSG00000197746 |
| 79 | P06744 | NA | GPI | glucose-6-phosphate isomerase | 2821 | ENSG00000105220 |
| 80 | Q15758 | NA | SLC1A5 | solute carrier family 1 (neutral amino acid transporter), member 5 | 6510 | ENSG00000105281 |
| 81 | P49748 | NA | ACADVL | acyl-CoA dehydrogenase, very long chain | 37 | ENSG00000072778 |

  
  

| **Database:cellular component      &nbspName:intracellular part      &nbspID:GO:0044424** | | | | | | |
| --- | --- | --- | --- | --- | --- | --- |
| C=12096; O=86; E=62.36; R=1.38; rawP=9.07e-13; adjP=3.14e-11 | | | | | | |
| Index | UserID | Value | Gene Symbol | Gene Name | EntrezGene | Ensembl |
| 1 | Q04446 | NA | GBE1 | glucan (1,4-alpha-), branching enzyme 1 | 2632 | ENSG00000114480 |
| 2 | P10253 | NA | GAA | glucosidase, alpha; acid | 2548 | ENSG00000171298 |
| 3 | Q16851 | NA | UGP2 | UDP-glucose pyrophosphorylase 2 | 7360 | ENSG00000169764 |
| 4 | P30086 | NA | PEBP1 | phosphatidylethanolamine binding protein 1 | 5037 | ENSG00000089220 |
| 5 | P55060 | NA | CSE1L | CSE1 chromosome segregation 1-like (yeast) | 1434 | ENSG00000124207 |
| 6 | P00367 | NA | GLUD1 | glutamate dehydrogenase 1 | 2746 | ENSG00000148672 |
| 7 | Q06323 | NA | PSME1 | proteasome (prosome, macropain) activator subunit 1 (PA28 alpha) | 5720 | ENSG00000092010 |
| 8 | P48735 | NA | IDH2 | isocitrate dehydrogenase 2 (NADP+), mitochondrial | 3418 | ENSG00000182054 |
| 9 | Q9P0J0 | NA | NDUFA13 | NADH dehydrogenase (ubiquinone) 1 alpha subcomplex, 13 | 51079 | ENSG00000186010 |
| 10 | P16949 | NA | STMN1 | stathmin 1 | 3925 | ENSG00000117632 |
| 11 | Q86UP2 | NA | KTN1 | kinectin 1 (kinesin receptor) | 3895 | ENSG00000126777 |
| 12 | P49588 | NA | AARS | alanyl-tRNA synthetase | 16 | ENSG00000090861 |
| 13 | O75369 | NA | FLNB | filamin B, beta | 2317 | ENSG00000136068 |
| 14 | O14773 | NA | TPP1 | tripeptidyl peptidase I | 1200 | ENSG00000166340 |
| 15 | P16615 | NA | ATP2A2 | ATPase, Ca++ transporting, cardiac muscle, slow twitch 2 | 488 | ENSG00000174437 |
| 16 | P42765 | NA | ACAA2 | acetyl-CoA acyltransferase 2 | 10449 | ENSG00000167315 |
| 17 | P40121 | NA | CAPG | capping protein (actin filament), gelsolin-like | 822 | ENSG00000042493 |
| 18 | P17174 | NA | GOT1 | glutamic-oxaloacetic transaminase 1, soluble (aspartate aminotransferase 1) | 2805 | ENSG00000120053 |
| 19 | P08727 | NA | KRT19 | keratin 19 | 3880 | ENSG00000171345 |
| 20 | Q14697 | NA | GANAB | glucosidase, alpha; neutral AB | 23193 | ENSG00000089597 |
| 21 | P18206 | NA | VCL | vinculin | 7414 | ENSG00000035403 |
| 22 | Q9UK76 | NA | HN1 | hematological and neurological expressed 1 | 51155 | ENSG00000189159 |
| 23 | P08195 | NA | SLC3A2 | solute carrier family 3 (activators of dibasic and neutral amino acid transport), member 2 | 6520 | ENSG00000168003 |
| 24 | Q71U36 | NA | TUBA1A | tubulin, alpha 1a | 7846 | ENSG00000167552 |
| 25 | P78417 | NA | GSTO1 | glutathione S-transferase omega 1 | 9446 | ENSG00000148834 |
| 26 | P49736 | NA | MCM2 | minichromosome maintenance complex component 2 | 4171 | ENSG00000073111 |
| 27 | P38571 | NA | LIPA | lipase A, lysosomal acid, cholesterol esterase | 3988 | ENSG00000107798 |
| 28 | P23381 | NA | WARS | tryptophanyl-tRNA synthetase | 7453 | ENSG00000140105 |
| 29 | Q14980 | NA | NUMA1 | nuclear mitotic apparatus protein 1 | 4926 | ENSG00000137497 |
| 30 | Q9UBQ7 | NA | GRHPR | glyoxylate reductase/hydroxypyruvate reductase | 9380 | ENSG00000137106 |
| 31 | P51649 | NA | ALDH5A1 | aldehyde dehydrogenase 5 family, member A1 | 7915 | ENSG00000112294 |
| 32 | P30044 | NA | PRDX5 | peroxiredoxin 5 | 25824 | ENSG00000126432 |
| 33 | P05556 | NA | ITGB1 | integrin, beta 1 (fibronectin receptor, beta polypeptide, antigen CD29 includes MDF2, MSK12) | 3688 | ENSG00000150093 |
| 34 | P13674 | NA | P4HA1 | prolyl 4-hydroxylase, alpha polypeptide I | 5033 | ENSG00000122884 |
| 35 | P07384 | NA | CAPN1 | calpain 1, (mu/I) large subunit | 823 | ENSG00000014216 |
| 36 | P49591 | NA | SARS | seryl-tRNA synthetase | 6301 | ENSG00000031698 |
| 37 | Q92598 | NA | HSPH1 | heat shock 105kDa/110kDa protein 1 | 10808 | ENSG00000120694 |
| 38 | P17931 | NA | LGALS3 | lectin, galactoside-binding, soluble, 3 | 3958 | ENSG00000131981 |
| 39 | P08133 | NA | ANXA6 | annexin A6 | 309 | ENSG00000197043 |
| 40 | P61225 | NA | RAP2B | RAP2B, member of RAS oncogene family | 5912 | ENSG00000181467 |
| 41 | Q9NQC3 | NA | RTN4 | reticulon 4 | 57142 | ENSG00000115310 |
| 42 | Q15019 | NA | SEPT2 | septin 2 | 4735 | ENSG00000168385 |
| 43 | P09211 | NA | GSTP1 | glutathione S-transferase pi 1 | 2950 | ENSG00000084207 |
| 44 | P33993 | NA | MCM7 | minichromosome maintenance complex component 7 | 4176 | ENSG00000166508 |
| 45 | P68104 | NA | EEF1A1 | eukaryotic translation elongation factor 1 alpha 1 | 1915 | ENSG00000156508 |
| 46 | P02786 | NA | TFRC | transferrin receptor (p90, CD71) | 7037 | ENSG00000072274 |
| 47 | O00425 | NA | IGF2BP3 | insulin-like growth factor 2 mRNA binding protein 3 | 10643 | ENSG00000136231 |
| 48 | P43490 | NA | NAMPT | nicotinamide phosphoribosyltransferase | 10135 | ENSG00000105835 |
| 49 | P17655 | NA | CAPN2 | calpain 2, (m/II) large subunit | 824 | ENSG00000162909 |
| 50 | Q9P258 | NA | RCC2 | regulator of chromosome condensation 2 | 55920 | ENSG00000179051 |
| 51 | P20042 | NA | EIF2S2 | eukaryotic translation initiation factor 2, subunit 2 beta, 38kDa | 8894 | ENSG00000125977 |
| 52 | P22102 | NA | GART | phosphoribosylglycinamide formyltransferase, phosphoribosylglycinamide synthetase, phosphoribosylaminoimidazole synthetase | 2618 | ENSG00000159131 |
| 53 | P22570 | NA | FDXR | ferredoxin reductase | 2232 | ENSG00000161513 |
| 54 | Q05639 | NA | EEF1A2 | eukaryotic translation elongation factor 1 alpha 2 | 1917 | ENSG00000101210 |
| 55 | P49419 | NA | ALDH7A1 | aldehyde dehydrogenase 7 family, member A1 | 501 | ENSG00000164904 |
| 56 | Q92616 | NA | GCN1L1 | GCN1 general control of amino-acid synthesis 1-like 1 (yeast) | 10985 | ENSG00000089154 |
| 57 | P37802 | NA | TAGLN2 | transgelin 2 | 8407 | ENSG00000158710 |
| 58 | P07339 | NA | CTSD | cathepsin D | 1509 | ENSG00000117984 |
| 59 | Q14764 | NA | MVP | major vault protein | 9961 | ENSG00000013364 |
| 60 | Q6PIU2 | NA | NCEH1 | neutral cholesterol ester hydrolase 1 | 57552 | ENSG00000144959 |
| 61 | P21589 | NA | NT5E | 5'-nucleotidase, ecto (CD73) | 4907 | ENSG00000135318 |
| 62 | P11387 | NA | TOP1 | topoisomerase (DNA) I | 7150 | ENSG00000198900 |
| 63 | P53007 | NA | SLC25A1 | solute carrier family 25 (mitochondrial carrier; citrate transporter), member 1 | 6576 | ENSG00000100075 |
| 64 | Q9BPW8 | NA | NIPSNAP1 | nipsnap homolog 1 (C. elegans) | 8508 | ENSG00000184117 |
| 65 | Q16881 | NA | TXNRD1 | thioredoxin reductase 1 | 7296 | ENSG00000198431 |
| 66 | P02792 | NA | FTL | ferritin, light polypeptide | 2512 | ENSG00000087086 |
| 67 | Q9NR30 | NA | DDX21 | DEAD (Asp-Glu-Ala-Asp) box helicase 21 | 9188 | ENSG00000165732 |
| 68 | P49321 | NA | NASP | nuclear autoantigenic sperm protein (histone-binding) | 4678 | ENSG00000132780 |
| 69 | P00491 | NA | PNP | purine nucleoside phosphorylase | 4860 | ENSG00000198805 |
| 70 | Q9HAV4 | NA | XPO5 | exportin 5 | 57510 | ENSG00000124571 |
| 71 | P00338 | NA | LDHA | lactate dehydrogenase A | 3939 | ENSG00000134333 |
| 72 | P56192 | NA | MARS | methionyl-tRNA synthetase | 4141 | ENSG00000166986 |
| 73 | Q15056 | NA | EIF4H | eukaryotic translation initiation factor 4H | 7458 | ENSG00000106682 |
| 74 | O60610 | NA | DIAPH1 | diaphanous homolog 1 (Drosophila) | 1729 | ENSG00000131504 |
| 75 | Q06830 | NA | PRDX1 | peroxiredoxin 1 | 5052 | ENSG00000117450 |
| 76 | Q13011 | NA | ECH1 | enoyl CoA hydratase 1, peroxisomal | 1891 | ENSG00000104823 |
| 77 | P04083 | NA | ANXA1 | annexin A1 | 301 | ENSG00000135046 |
| 78 | P04040 | NA | CAT | catalase | 847 | ENSG00000121691 |
| 79 | P41091 | NA | EIF2S3 | eukaryotic translation initiation factor 2, subunit 3 gamma, 52kDa | 1968 | ENSG00000130741 |
| 80 | P02768 | NA | ALB | albumin | 213 | ENSG00000163631 |
| 81 | P24752 | NA | ACAT1 | acetyl-CoA acetyltransferase 1 | 38 | ENSG00000075239 |
| 82 | P16278 | NA | GLB1 | galactosidase, beta 1 | 2720 | ENSG00000170266 |
| 83 | P07602 | NA | PSAP | prosaposin | 5660 | ENSG00000197746 |
| 84 | P06744 | NA | GPI | glucose-6-phosphate isomerase | 2821 | ENSG00000105220 |
| 85 | Q15758 | NA | SLC1A5 | solute carrier family 1 (neutral amino acid transporter), member 5 | 6510 | ENSG00000105281 |
| 86 | P49748 | NA | ACADVL | acyl-CoA dehydrogenase, very long chain | 37 | ENSG00000072778 |

  
  

| **Database:cellular component      &nbspName:intracellular      &nbspID:GO:0005622** | | | | | | |
| --- | --- | --- | --- | --- | --- | --- |
| C=12412; O=86; E=63.99; R=1.34; rawP=8.40e-12; adjP=2.18e-10 | | | | | | |
| Index | UserID | Value | Gene Symbol | Gene Name | EntrezGene | Ensembl |
| 1 | Q04446 | NA | GBE1 | glucan (1,4-alpha-), branching enzyme 1 | 2632 | ENSG00000114480 |
| 2 | P10253 | NA | GAA | glucosidase, alpha; acid | 2548 | ENSG00000171298 |
| 3 | Q16851 | NA | UGP2 | UDP-glucose pyrophosphorylase 2 | 7360 | ENSG00000169764 |
| 4 | P30086 | NA | PEBP1 | phosphatidylethanolamine binding protein 1 | 5037 | ENSG00000089220 |
| 5 | P55060 | NA | CSE1L | CSE1 chromosome segregation 1-like (yeast) | 1434 | ENSG00000124207 |
| 6 | P00367 | NA | GLUD1 | glutamate dehydrogenase 1 | 2746 | ENSG00000148672 |
| 7 | Q06323 | NA | PSME1 | proteasome (prosome, macropain) activator subunit 1 (PA28 alpha) | 5720 | ENSG00000092010 |
| 8 | P48735 | NA | IDH2 | isocitrate dehydrogenase 2 (NADP+), mitochondrial | 3418 | ENSG00000182054 |
| 9 | Q9P0J0 | NA | NDUFA13 | NADH dehydrogenase (ubiquinone) 1 alpha subcomplex, 13 | 51079 | ENSG00000186010 |
| 10 | P16949 | NA | STMN1 | stathmin 1 | 3925 | ENSG00000117632 |
| 11 | Q86UP2 | NA | KTN1 | kinectin 1 (kinesin receptor) | 3895 | ENSG00000126777 |
| 12 | P49588 | NA | AARS | alanyl-tRNA synthetase | 16 | ENSG00000090861 |
| 13 | O75369 | NA | FLNB | filamin B, beta | 2317 | ENSG00000136068 |
| 14 | O14773 | NA | TPP1 | tripeptidyl peptidase I | 1200 | ENSG00000166340 |
| 15 | P16615 | NA | ATP2A2 | ATPase, Ca++ transporting, cardiac muscle, slow twitch 2 | 488 | ENSG00000174437 |
| 16 | P42765 | NA | ACAA2 | acetyl-CoA acyltransferase 2 | 10449 | ENSG00000167315 |
| 17 | P40121 | NA | CAPG | capping protein (actin filament), gelsolin-like | 822 | ENSG00000042493 |
| 18 | P17174 | NA | GOT1 | glutamic-oxaloacetic transaminase 1, soluble (aspartate aminotransferase 1) | 2805 | ENSG00000120053 |
| 19 | P08727 | NA | KRT19 | keratin 19 | 3880 | ENSG00000171345 |
| 20 | Q14697 | NA | GANAB | glucosidase, alpha; neutral AB | 23193 | ENSG00000089597 |
| 21 | P18206 | NA | VCL | vinculin | 7414 | ENSG00000035403 |
| 22 | Q9UK76 | NA | HN1 | hematological and neurological expressed 1 | 51155 | ENSG00000189159 |
| 23 | P08195 | NA | SLC3A2 | solute carrier family 3 (activators of dibasic and neutral amino acid transport), member 2 | 6520 | ENSG00000168003 |
| 24 | Q71U36 | NA | TUBA1A | tubulin, alpha 1a | 7846 | ENSG00000167552 |
| 25 | P78417 | NA | GSTO1 | glutathione S-transferase omega 1 | 9446 | ENSG00000148834 |
| 26 | P49736 | NA | MCM2 | minichromosome maintenance complex component 2 | 4171 | ENSG00000073111 |
| 27 | P38571 | NA | LIPA | lipase A, lysosomal acid, cholesterol esterase | 3988 | ENSG00000107798 |
| 28 | P23381 | NA | WARS | tryptophanyl-tRNA synthetase | 7453 | ENSG00000140105 |
| 29 | Q14980 | NA | NUMA1 | nuclear mitotic apparatus protein 1 | 4926 | ENSG00000137497 |
| 30 | Q9UBQ7 | NA | GRHPR | glyoxylate reductase/hydroxypyruvate reductase | 9380 | ENSG00000137106 |
| 31 | P51649 | NA | ALDH5A1 | aldehyde dehydrogenase 5 family, member A1 | 7915 | ENSG00000112294 |
| 32 | P30044 | NA | PRDX5 | peroxiredoxin 5 | 25824 | ENSG00000126432 |
| 33 | P05556 | NA | ITGB1 | integrin, beta 1 (fibronectin receptor, beta polypeptide, antigen CD29 includes MDF2, MSK12) | 3688 | ENSG00000150093 |
| 34 | P13674 | NA | P4HA1 | prolyl 4-hydroxylase, alpha polypeptide I | 5033 | ENSG00000122884 |
| 35 | P07384 | NA | CAPN1 | calpain 1, (mu/I) large subunit | 823 | ENSG00000014216 |
| 36 | P49591 | NA | SARS | seryl-tRNA synthetase | 6301 | ENSG00000031698 |
| 37 | Q92598 | NA | HSPH1 | heat shock 105kDa/110kDa protein 1 | 10808 | ENSG00000120694 |
| 38 | P17931 | NA | LGALS3 | lectin, galactoside-binding, soluble, 3 | 3958 | ENSG00000131981 |
| 39 | P08133 | NA | ANXA6 | annexin A6 | 309 | ENSG00000197043 |
| 40 | P61225 | NA | RAP2B | RAP2B, member of RAS oncogene family | 5912 | ENSG00000181467 |
| 41 | Q9NQC3 | NA | RTN4 | reticulon 4 | 57142 | ENSG00000115310 |
| 42 | Q15019 | NA | SEPT2 | septin 2 | 4735 | ENSG00000168385 |
| 43 | P09211 | NA | GSTP1 | glutathione S-transferase pi 1 | 2950 | ENSG00000084207 |
| 44 | P33993 | NA | MCM7 | minichromosome maintenance complex component 7 | 4176 | ENSG00000166508 |
| 45 | P68104 | NA | EEF1A1 | eukaryotic translation elongation factor 1 alpha 1 | 1915 | ENSG00000156508 |
| 46 | P02786 | NA | TFRC | transferrin receptor (p90, CD71) | 7037 | ENSG00000072274 |
| 47 | O00425 | NA | IGF2BP3 | insulin-like growth factor 2 mRNA binding protein 3 | 10643 | ENSG00000136231 |
| 48 | P43490 | NA | NAMPT | nicotinamide phosphoribosyltransferase | 10135 | ENSG00000105835 |
| 49 | P17655 | NA | CAPN2 | calpain 2, (m/II) large subunit | 824 | ENSG00000162909 |
| 50 | Q9P258 | NA | RCC2 | regulator of chromosome condensation 2 | 55920 | ENSG00000179051 |
| 51 | P20042 | NA | EIF2S2 | eukaryotic translation initiation factor 2, subunit 2 beta, 38kDa | 8894 | ENSG00000125977 |
| 52 | P22102 | NA | GART | phosphoribosylglycinamide formyltransferase, phosphoribosylglycinamide synthetase, phosphoribosylaminoimidazole synthetase | 2618 | ENSG00000159131 |
| 53 | P22570 | NA | FDXR | ferredoxin reductase | 2232 | ENSG00000161513 |
| 54 | Q05639 | NA | EEF1A2 | eukaryotic translation elongation factor 1 alpha 2 | 1917 | ENSG00000101210 |
| 55 | P49419 | NA | ALDH7A1 | aldehyde dehydrogenase 7 family, member A1 | 501 | ENSG00000164904 |
| 56 | Q92616 | NA | GCN1L1 | GCN1 general control of amino-acid synthesis 1-like 1 (yeast) | 10985 | ENSG00000089154 |
| 57 | P37802 | NA | TAGLN2 | transgelin 2 | 8407 | ENSG00000158710 |
| 58 | P07339 | NA | CTSD | cathepsin D | 1509 | ENSG00000117984 |
| 59 | Q14764 | NA | MVP | major vault protein | 9961 | ENSG00000013364 |
| 60 | Q6PIU2 | NA | NCEH1 | neutral cholesterol ester hydrolase 1 | 57552 | ENSG00000144959 |
| 61 | P21589 | NA | NT5E | 5'-nucleotidase, ecto (CD73) | 4907 | ENSG00000135318 |
| 62 | P11387 | NA | TOP1 | topoisomerase (DNA) I | 7150 | ENSG00000198900 |
| 63 | P53007 | NA | SLC25A1 | solute carrier family 25 (mitochondrial carrier; citrate transporter), member 1 | 6576 | ENSG00000100075 |
| 64 | Q9BPW8 | NA | NIPSNAP1 | nipsnap homolog 1 (C. elegans) | 8508 | ENSG00000184117 |
| 65 | Q16881 | NA | TXNRD1 | thioredoxin reductase 1 | 7296 | ENSG00000198431 |
| 66 | P02792 | NA | FTL | ferritin, light polypeptide | 2512 | ENSG00000087086 |
| 67 | Q9NR30 | NA | DDX21 | DEAD (Asp-Glu-Ala-Asp) box helicase 21 | 9188 | ENSG00000165732 |
| 68 | P49321 | NA | NASP | nuclear autoantigenic sperm protein (histone-binding) | 4678 | ENSG00000132780 |
| 69 | P00491 | NA | PNP | purine nucleoside phosphorylase | 4860 | ENSG00000198805 |
| 70 | Q9HAV4 | NA | XPO5 | exportin 5 | 57510 | ENSG00000124571 |
| 71 | P00338 | NA | LDHA | lactate dehydrogenase A | 3939 | ENSG00000134333 |
| 72 | P56192 | NA | MARS | methionyl-tRNA synthetase | 4141 | ENSG00000166986 |
| 73 | Q15056 | NA | EIF4H | eukaryotic translation initiation factor 4H | 7458 | ENSG00000106682 |
| 74 | O60610 | NA | DIAPH1 | diaphanous homolog 1 (Drosophila) | 1729 | ENSG00000131504 |
| 75 | Q06830 | NA | PRDX1 | peroxiredoxin 1 | 5052 | ENSG00000117450 |
| 76 | Q13011 | NA | ECH1 | enoyl CoA hydratase 1, peroxisomal | 1891 | ENSG00000104823 |
| 77 | P04083 | NA | ANXA1 | annexin A1 | 301 | ENSG00000135046 |
| 78 | P04040 | NA | CAT | catalase | 847 | ENSG00000121691 |
| 79 | P41091 | NA | EIF2S3 | eukaryotic translation initiation factor 2, subunit 3 gamma, 52kDa | 1968 | ENSG00000130741 |
| 80 | P02768 | NA | ALB | albumin | 213 | ENSG00000163631 |
| 81 | P24752 | NA | ACAT1 | acetyl-CoA acetyltransferase 1 | 38 | ENSG00000075239 |
| 82 | P16278 | NA | GLB1 | galactosidase, beta 1 | 2720 | ENSG00000170266 |
| 83 | P07602 | NA | PSAP | prosaposin | 5660 | ENSG00000197746 |
| 84 | P06744 | NA | GPI | glucose-6-phosphate isomerase | 2821 | ENSG00000105220 |
| 85 | Q15758 | NA | SLC1A5 | solute carrier family 1 (neutral amino acid transporter), member 5 | 6510 | ENSG00000105281 |
| 86 | P49748 | NA | ACADVL | acyl-CoA dehydrogenase, very long chain | 37 | ENSG00000072778 |

  
  

| **Database:cellular component      &nbspName:pigment granule      &nbspID:GO:0048770** | | | | | | |
| --- | --- | --- | --- | --- | --- | --- |
| C=93; O=10; E=0.48; R=20.86; rawP=4.42e-11; adjP=7.66e-10 | | | | | | |
| Index | UserID | Value | Gene Symbol | Gene Name | EntrezGene | Ensembl |
| 1 | O14773 | NA | TPP1 | tripeptidyl peptidase I | 1200 | ENSG00000166340 |
| 2 | P08195 | NA | SLC3A2 | solute carrier family 3 (activators of dibasic and neutral amino acid transport), member 2 | 6520 | ENSG00000168003 |
| 3 | P08133 | NA | ANXA6 | annexin A6 | 309 | ENSG00000197043 |
| 4 | P05556 | NA | ITGB1 | integrin, beta 1 (fibronectin receptor, beta polypeptide, antigen CD29 includes MDF2, MSK12) | 3688 | ENSG00000150093 |
| 5 | P40121 | NA | CAPG | capping protein (actin filament), gelsolin-like | 822 | ENSG00000042493 |
| 6 | Q15758 | NA | SLC1A5 | solute carrier family 1 (neutral amino acid transporter), member 5 | 6510 | ENSG00000105281 |
| 7 | P07339 | NA | CTSD | cathepsin D | 1509 | ENSG00000117984 |
| 8 | Q14697 | NA | GANAB | glucosidase, alpha; neutral AB | 23193 | ENSG00000089597 |
| 9 | Q06830 | NA | PRDX1 | peroxiredoxin 1 | 5052 | ENSG00000117450 |
| 10 | P02786 | NA | TFRC | transferrin receptor (p90, CD71) | 7037 | ENSG00000072274 |

  
  

| **Database:cellular component      &nbspName:melanosome      &nbspID:GO:0042470** | | | | | | |
| --- | --- | --- | --- | --- | --- | --- |
| C=93; O=10; E=0.48; R=20.86; rawP=4.42e-11; adjP=7.66e-10 | | | | | | |
| Index | UserID | Value | Gene Symbol | Gene Name | EntrezGene | Ensembl |
| 1 | O14773 | NA | TPP1 | tripeptidyl peptidase I | 1200 | ENSG00000166340 |
| 2 | P08195 | NA | SLC3A2 | solute carrier family 3 (activators of dibasic and neutral amino acid transport), member 2 | 6520 | ENSG00000168003 |
| 3 | P08133 | NA | ANXA6 | annexin A6 | 309 | ENSG00000197043 |
| 4 | P05556 | NA | ITGB1 | integrin, beta 1 (fibronectin receptor, beta polypeptide, antigen CD29 includes MDF2, MSK12) | 3688 | ENSG00000150093 |
| 5 | P40121 | NA | CAPG | capping protein (actin filament), gelsolin-like | 822 | ENSG00000042493 |
| 6 | Q15758 | NA | SLC1A5 | solute carrier family 1 (neutral amino acid transporter), member 5 | 6510 | ENSG00000105281 |
| 7 | P07339 | NA | CTSD | cathepsin D | 1509 | ENSG00000117984 |
| 8 | Q14697 | NA | GANAB | glucosidase, alpha; neutral AB | 23193 | ENSG00000089597 |
| 9 | Q06830 | NA | PRDX1 | peroxiredoxin 1 | 5052 | ENSG00000117450 |
| 10 | P02786 | NA | TFRC | transferrin receptor (p90, CD71) | 7037 | ENSG00000072274 |

  
  

| **Database:cellular component      &nbspName:mitochondrion      &nbspID:GO:0005739** | | | | | | |
| --- | --- | --- | --- | --- | --- | --- |
| C=1515; O=27; E=7.81; R=3.46; rawP=4.79e-09; adjP=7.12e-08 | | | | | | |
| Index | UserID | Value | Gene Symbol | Gene Name | EntrezGene | Ensembl |
| 1 | Q16881 | NA | TXNRD1 | thioredoxin reductase 1 | 7296 | ENSG00000198431 |
| 2 | P49591 | NA | SARS | seryl-tRNA synthetase | 6301 | ENSG00000031698 |
| 3 | P17931 | NA | LGALS3 | lectin, galactoside-binding, soluble, 3 | 3958 | ENSG00000131981 |
| 4 | P30086 | NA | PEBP1 | phosphatidylethanolamine binding protein 1 | 5037 | ENSG00000089220 |
| 5 | P00338 | NA | LDHA | lactate dehydrogenase A | 3939 | ENSG00000134333 |
| 6 | P56192 | NA | MARS | methionyl-tRNA synthetase | 4141 | ENSG00000166986 |
| 7 | P00367 | NA | GLUD1 | glutamate dehydrogenase 1 | 2746 | ENSG00000148672 |
| 8 | P48735 | NA | IDH2 | isocitrate dehydrogenase 2 (NADP+), mitochondrial | 3418 | ENSG00000182054 |
| 9 | P09211 | NA | GSTP1 | glutathione S-transferase pi 1 | 2950 | ENSG00000084207 |
| 10 | Q9P0J0 | NA | NDUFA13 | NADH dehydrogenase (ubiquinone) 1 alpha subcomplex, 13 | 51079 | ENSG00000186010 |
| 11 | Q06830 | NA | PRDX1 | peroxiredoxin 1 | 5052 | ENSG00000117450 |
| 12 | Q13011 | NA | ECH1 | enoyl CoA hydratase 1, peroxisomal | 1891 | ENSG00000104823 |
| 13 | P02786 | NA | TFRC | transferrin receptor (p90, CD71) | 7037 | ENSG00000072274 |
| 14 | P04083 | NA | ANXA1 | annexin A1 | 301 | ENSG00000135046 |
| 15 | P04040 | NA | CAT | catalase | 847 | ENSG00000121691 |
| 16 | O14773 | NA | TPP1 | tripeptidyl peptidase I | 1200 | ENSG00000166340 |
| 17 | P51649 | NA | ALDH5A1 | aldehyde dehydrogenase 5 family, member A1 | 7915 | ENSG00000112294 |
| 18 | P30044 | NA | PRDX5 | peroxiredoxin 5 | 25824 | ENSG00000126432 |
| 19 | P22570 | NA | FDXR | ferredoxin reductase | 2232 | ENSG00000161513 |
| 20 | P42765 | NA | ACAA2 | acetyl-CoA acyltransferase 2 | 10449 | ENSG00000167315 |
| 21 | P24752 | NA | ACAT1 | acetyl-CoA acetyltransferase 1 | 38 | ENSG00000075239 |
| 22 | P13674 | NA | P4HA1 | prolyl 4-hydroxylase, alpha polypeptide I | 5033 | ENSG00000122884 |
| 23 | P49419 | NA | ALDH7A1 | aldehyde dehydrogenase 7 family, member A1 | 501 | ENSG00000164904 |
| 24 | P07602 | NA | PSAP | prosaposin | 5660 | ENSG00000197746 |
| 25 | P49748 | NA | ACADVL | acyl-CoA dehydrogenase, very long chain | 37 | ENSG00000072778 |
| 26 | P53007 | NA | SLC25A1 | solute carrier family 25 (mitochondrial carrier; citrate transporter), member 1 | 6576 | ENSG00000100075 |
| 27 | Q9BPW8 | NA | NIPSNAP1 | nipsnap homolog 1 (C. elegans) | 8508 | ENSG00000184117 |

  
  

| **Database:cellular component      &nbspName:cytosol      &nbspID:GO:0005829** | | | | | | |
| --- | --- | --- | --- | --- | --- | --- |
| C=2367; O=33; E=12.20; R=2.70; rawP=2.45e-08; adjP=3.18e-07 | | | | | | |
| Index | UserID | Value | Gene Symbol | Gene Name | EntrezGene | Ensembl |
| 1 | Q16881 | NA | TXNRD1 | thioredoxin reductase 1 | 7296 | ENSG00000198431 |
| 2 | Q04446 | NA | GBE1 | glucan (1,4-alpha-), branching enzyme 1 | 2632 | ENSG00000114480 |
| 3 | P02792 | NA | FTL | ferritin, light polypeptide | 2512 | ENSG00000087086 |
| 4 | P49591 | NA | SARS | seryl-tRNA synthetase | 6301 | ENSG00000031698 |
| 5 | P00491 | NA | PNP | purine nucleoside phosphorylase | 4860 | ENSG00000198805 |
| 6 | Q71U36 | NA | TUBA1A | tubulin, alpha 1a | 7846 | ENSG00000167552 |
| 7 | Q16851 | NA | UGP2 | UDP-glucose pyrophosphorylase 2 | 7360 | ENSG00000169764 |
| 8 | Q9HAV4 | NA | XPO5 | exportin 5 | 57510 | ENSG00000124571 |
| 9 | P00338 | NA | LDHA | lactate dehydrogenase A | 3939 | ENSG00000134333 |
| 10 | P61225 | NA | RAP2B | RAP2B, member of RAS oncogene family | 5912 | ENSG00000181467 |
| 11 | P56192 | NA | MARS | methionyl-tRNA synthetase | 4141 | ENSG00000166986 |
| 12 | P78417 | NA | GSTO1 | glutathione S-transferase omega 1 | 9446 | ENSG00000148834 |
| 13 | Q06323 | NA | PSME1 | proteasome (prosome, macropain) activator subunit 1 (PA28 alpha) | 5720 | ENSG00000092010 |
| 14 | Q15056 | NA | EIF4H | eukaryotic translation initiation factor 4H | 7458 | ENSG00000106682 |
| 15 | P09211 | NA | GSTP1 | glutathione S-transferase pi 1 | 2950 | ENSG00000084207 |
| 16 | P16949 | NA | STMN1 | stathmin 1 | 3925 | ENSG00000117632 |
| 17 | P68104 | NA | EEF1A1 | eukaryotic translation elongation factor 1 alpha 1 | 1915 | ENSG00000156508 |
| 18 | P49588 | NA | AARS | alanyl-tRNA synthetase | 16 | ENSG00000090861 |
| 19 | O75369 | NA | FLNB | filamin B, beta | 2317 | ENSG00000136068 |
| 20 | P43490 | NA | NAMPT | nicotinamide phosphoribosyltransferase | 10135 | ENSG00000105835 |
| 21 | O00425 | NA | IGF2BP3 | insulin-like growth factor 2 mRNA binding protein 3 | 10643 | ENSG00000136231 |
| 22 | P23381 | NA | WARS | tryptophanyl-tRNA synthetase | 7453 | ENSG00000140105 |
| 23 | P04040 | NA | CAT | catalase | 847 | ENSG00000121691 |
| 24 | Q14980 | NA | NUMA1 | nuclear mitotic apparatus protein 1 | 4926 | ENSG00000137497 |
| 25 | P22102 | NA | GART | phosphoribosylglycinamide formyltransferase, phosphoribosylglycinamide synthetase, phosphoribosylaminoimidazole synthetase | 2618 | ENSG00000159131 |
| 26 | P20042 | NA | EIF2S2 | eukaryotic translation initiation factor 2, subunit 2 beta, 38kDa | 8894 | ENSG00000125977 |
| 27 | Q9P258 | NA | RCC2 | regulator of chromosome condensation 2 | 55920 | ENSG00000179051 |
| 28 | P41091 | NA | EIF2S3 | eukaryotic translation initiation factor 2, subunit 3 gamma, 52kDa | 1968 | ENSG00000130741 |
| 29 | P30044 | NA | PRDX5 | peroxiredoxin 5 | 25824 | ENSG00000126432 |
| 30 | P49419 | NA | ALDH7A1 | aldehyde dehydrogenase 7 family, member A1 | 501 | ENSG00000164904 |
| 31 | P17174 | NA | GOT1 | glutamic-oxaloacetic transaminase 1, soluble (aspartate aminotransferase 1) | 2805 | ENSG00000120053 |
| 32 | P06744 | NA | GPI | glucose-6-phosphate isomerase | 2821 | ENSG00000105220 |
| 33 | P18206 | NA | VCL | vinculin | 7414 | ENSG00000035403 |

  
  

| **Database:cellular component      &nbspName:cell part      &nbspID:GO:0044464** | | | | | | |
| --- | --- | --- | --- | --- | --- | --- |
| C=14405; O=86; E=74.26; R=1.16; rawP=3.19e-06; adjP=3.34e-05 | | | | | | |
| Index | UserID | Value | Gene Symbol | Gene Name | EntrezGene | Ensembl |
| 1 | Q04446 | NA | GBE1 | glucan (1,4-alpha-), branching enzyme 1 | 2632 | ENSG00000114480 |
| 2 | P10253 | NA | GAA | glucosidase, alpha; acid | 2548 | ENSG00000171298 |
| 3 | Q16851 | NA | UGP2 | UDP-glucose pyrophosphorylase 2 | 7360 | ENSG00000169764 |
| 4 | P30086 | NA | PEBP1 | phosphatidylethanolamine binding protein 1 | 5037 | ENSG00000089220 |
| 5 | P55060 | NA | CSE1L | CSE1 chromosome segregation 1-like (yeast) | 1434 | ENSG00000124207 |
| 6 | P00367 | NA | GLUD1 | glutamate dehydrogenase 1 | 2746 | ENSG00000148672 |
| 7 | Q06323 | NA | PSME1 | proteasome (prosome, macropain) activator subunit 1 (PA28 alpha) | 5720 | ENSG00000092010 |
| 8 | P48735 | NA | IDH2 | isocitrate dehydrogenase 2 (NADP+), mitochondrial | 3418 | ENSG00000182054 |
| 9 | Q9P0J0 | NA | NDUFA13 | NADH dehydrogenase (ubiquinone) 1 alpha subcomplex, 13 | 51079 | ENSG00000186010 |
| 10 | P16949 | NA | STMN1 | stathmin 1 | 3925 | ENSG00000117632 |
| 11 | Q86UP2 | NA | KTN1 | kinectin 1 (kinesin receptor) | 3895 | ENSG00000126777 |
| 12 | P49588 | NA | AARS | alanyl-tRNA synthetase | 16 | ENSG00000090861 |
| 13 | O75369 | NA | FLNB | filamin B, beta | 2317 | ENSG00000136068 |
| 14 | O14773 | NA | TPP1 | tripeptidyl peptidase I | 1200 | ENSG00000166340 |
| 15 | P16615 | NA | ATP2A2 | ATPase, Ca++ transporting, cardiac muscle, slow twitch 2 | 488 | ENSG00000174437 |
| 16 | P42765 | NA | ACAA2 | acetyl-CoA acyltransferase 2 | 10449 | ENSG00000167315 |
| 17 | P40121 | NA | CAPG | capping protein (actin filament), gelsolin-like | 822 | ENSG00000042493 |
| 18 | P17174 | NA | GOT1 | glutamic-oxaloacetic transaminase 1, soluble (aspartate aminotransferase 1) | 2805 | ENSG00000120053 |
| 19 | P08727 | NA | KRT19 | keratin 19 | 3880 | ENSG00000171345 |
| 20 | Q14697 | NA | GANAB | glucosidase, alpha; neutral AB | 23193 | ENSG00000089597 |
| 21 | P18206 | NA | VCL | vinculin | 7414 | ENSG00000035403 |
| 22 | Q9UK76 | NA | HN1 | hematological and neurological expressed 1 | 51155 | ENSG00000189159 |
| 23 | P08195 | NA | SLC3A2 | solute carrier family 3 (activators of dibasic and neutral amino acid transport), member 2 | 6520 | ENSG00000168003 |
| 24 | Q71U36 | NA | TUBA1A | tubulin, alpha 1a | 7846 | ENSG00000167552 |
| 25 | P78417 | NA | GSTO1 | glutathione S-transferase omega 1 | 9446 | ENSG00000148834 |
| 26 | P49736 | NA | MCM2 | minichromosome maintenance complex component 2 | 4171 | ENSG00000073111 |
| 27 | P38571 | NA | LIPA | lipase A, lysosomal acid, cholesterol esterase | 3988 | ENSG00000107798 |
| 28 | P23381 | NA | WARS | tryptophanyl-tRNA synthetase | 7453 | ENSG00000140105 |
| 29 | Q14980 | NA | NUMA1 | nuclear mitotic apparatus protein 1 | 4926 | ENSG00000137497 |
| 30 | Q9UBQ7 | NA | GRHPR | glyoxylate reductase/hydroxypyruvate reductase | 9380 | ENSG00000137106 |
| 31 | P51649 | NA | ALDH5A1 | aldehyde dehydrogenase 5 family, member A1 | 7915 | ENSG00000112294 |
| 32 | P30044 | NA | PRDX5 | peroxiredoxin 5 | 25824 | ENSG00000126432 |
| 33 | P05556 | NA | ITGB1 | integrin, beta 1 (fibronectin receptor, beta polypeptide, antigen CD29 includes MDF2, MSK12) | 3688 | ENSG00000150093 |
| 34 | P13674 | NA | P4HA1 | prolyl 4-hydroxylase, alpha polypeptide I | 5033 | ENSG00000122884 |
| 35 | P07384 | NA | CAPN1 | calpain 1, (mu/I) large subunit | 823 | ENSG00000014216 |
| 36 | P49591 | NA | SARS | seryl-tRNA synthetase | 6301 | ENSG00000031698 |
| 37 | Q92598 | NA | HSPH1 | heat shock 105kDa/110kDa protein 1 | 10808 | ENSG00000120694 |
| 38 | P17931 | NA | LGALS3 | lectin, galactoside-binding, soluble, 3 | 3958 | ENSG00000131981 |
| 39 | P08133 | NA | ANXA6 | annexin A6 | 309 | ENSG00000197043 |
| 40 | P61225 | NA | RAP2B | RAP2B, member of RAS oncogene family | 5912 | ENSG00000181467 |
| 41 | Q9NQC3 | NA | RTN4 | reticulon 4 | 57142 | ENSG00000115310 |
| 42 | Q15019 | NA | SEPT2 | septin 2 | 4735 | ENSG00000168385 |
| 43 | P09211 | NA | GSTP1 | glutathione S-transferase pi 1 | 2950 | ENSG00000084207 |
| 44 | P33993 | NA | MCM7 | minichromosome maintenance complex component 7 | 4176 | ENSG00000166508 |
| 45 | P68104 | NA | EEF1A1 | eukaryotic translation elongation factor 1 alpha 1 | 1915 | ENSG00000156508 |
| 46 | P02786 | NA | TFRC | transferrin receptor (p90, CD71) | 7037 | ENSG00000072274 |
| 47 | O00425 | NA | IGF2BP3 | insulin-like growth factor 2 mRNA binding protein 3 | 10643 | ENSG00000136231 |
| 48 | P43490 | NA | NAMPT | nicotinamide phosphoribosyltransferase | 10135 | ENSG00000105835 |
| 49 | P17655 | NA | CAPN2 | calpain 2, (m/II) large subunit | 824 | ENSG00000162909 |
| 50 | Q9P258 | NA | RCC2 | regulator of chromosome condensation 2 | 55920 | ENSG00000179051 |
| 51 | P20042 | NA | EIF2S2 | eukaryotic translation initiation factor 2, subunit 2 beta, 38kDa | 8894 | ENSG00000125977 |
| 52 | P22102 | NA | GART | phosphoribosylglycinamide formyltransferase, phosphoribosylglycinamide synthetase, phosphoribosylaminoimidazole synthetase | 2618 | ENSG00000159131 |
| 53 | P22570 | NA | FDXR | ferredoxin reductase | 2232 | ENSG00000161513 |
| 54 | Q05639 | NA | EEF1A2 | eukaryotic translation elongation factor 1 alpha 2 | 1917 | ENSG00000101210 |
| 55 | P49419 | NA | ALDH7A1 | aldehyde dehydrogenase 7 family, member A1 | 501 | ENSG00000164904 |
| 56 | Q92616 | NA | GCN1L1 | GCN1 general control of amino-acid synthesis 1-like 1 (yeast) | 10985 | ENSG00000089154 |
| 57 | P37802 | NA | TAGLN2 | transgelin 2 | 8407 | ENSG00000158710 |
| 58 | P07339 | NA | CTSD | cathepsin D | 1509 | ENSG00000117984 |
| 59 | Q14764 | NA | MVP | major vault protein | 9961 | ENSG00000013364 |
| 60 | Q6PIU2 | NA | NCEH1 | neutral cholesterol ester hydrolase 1 | 57552 | ENSG00000144959 |
| 61 | P21589 | NA | NT5E | 5'-nucleotidase, ecto (CD73) | 4907 | ENSG00000135318 |
| 62 | P11387 | NA | TOP1 | topoisomerase (DNA) I | 7150 | ENSG00000198900 |
| 63 | P53007 | NA | SLC25A1 | solute carrier family 25 (mitochondrial carrier; citrate transporter), member 1 | 6576 | ENSG00000100075 |
| 64 | Q9BPW8 | NA | NIPSNAP1 | nipsnap homolog 1 (C. elegans) | 8508 | ENSG00000184117 |
| 65 | Q16881 | NA | TXNRD1 | thioredoxin reductase 1 | 7296 | ENSG00000198431 |
| 66 | P02792 | NA | FTL | ferritin, light polypeptide | 2512 | ENSG00000087086 |
| 67 | Q9NR30 | NA | DDX21 | DEAD (Asp-Glu-Ala-Asp) box helicase 21 | 9188 | ENSG00000165732 |
| 68 | P49321 | NA | NASP | nuclear autoantigenic sperm protein (histone-binding) | 4678 | ENSG00000132780 |
| 69 | P00491 | NA | PNP | purine nucleoside phosphorylase | 4860 | ENSG00000198805 |
| 70 | Q9HAV4 | NA | XPO5 | exportin 5 | 57510 | ENSG00000124571 |
| 71 | P00338 | NA | LDHA | lactate dehydrogenase A | 3939 | ENSG00000134333 |
| 72 | P56192 | NA | MARS | methionyl-tRNA synthetase | 4141 | ENSG00000166986 |
| 73 | Q15056 | NA | EIF4H | eukaryotic translation initiation factor 4H | 7458 | ENSG00000106682 |
| 74 | O60610 | NA | DIAPH1 | diaphanous homolog 1 (Drosophila) | 1729 | ENSG00000131504 |
| 75 | Q06830 | NA | PRDX1 | peroxiredoxin 1 | 5052 | ENSG00000117450 |
| 76 | Q13011 | NA | ECH1 | enoyl CoA hydratase 1, peroxisomal | 1891 | ENSG00000104823 |
| 77 | P04083 | NA | ANXA1 | annexin A1 | 301 | ENSG00000135046 |
| 78 | P04040 | NA | CAT | catalase | 847 | ENSG00000121691 |
| 79 | P41091 | NA | EIF2S3 | eukaryotic translation initiation factor 2, subunit 3 gamma, 52kDa | 1968 | ENSG00000130741 |
| 80 | P02768 | NA | ALB | albumin | 213 | ENSG00000163631 |
| 81 | P24752 | NA | ACAT1 | acetyl-CoA acetyltransferase 1 | 38 | ENSG00000075239 |
| 82 | P16278 | NA | GLB1 | galactosidase, beta 1 | 2720 | ENSG00000170266 |
| 83 | P07602 | NA | PSAP | prosaposin | 5660 | ENSG00000197746 |
| 84 | P06744 | NA | GPI | glucose-6-phosphate isomerase | 2821 | ENSG00000105220 |
| 85 | Q15758 | NA | SLC1A5 | solute carrier family 1 (neutral amino acid transporter), member 5 | 6510 | ENSG00000105281 |
| 86 | P49748 | NA | ACADVL | acyl-CoA dehydrogenase, very long chain | 37 | ENSG00000072778 |

  
  

| **Database:cellular component      &nbspName:cell      &nbspID:GO:0005623** | | | | | | |
| --- | --- | --- | --- | --- | --- | --- |
| C=14406; O=86; E=74.27; R=1.16; rawP=3.21e-06; adjP=3.34e-05 | | | | | | |
| Index | UserID | Value | Gene Symbol | Gene Name | EntrezGene | Ensembl |
| 1 | Q04446 | NA | GBE1 | glucan (1,4-alpha-), branching enzyme 1 | 2632 | ENSG00000114480 |
| 2 | P10253 | NA | GAA | glucosidase, alpha; acid | 2548 | ENSG00000171298 |
| 3 | Q16851 | NA | UGP2 | UDP-glucose pyrophosphorylase 2 | 7360 | ENSG00000169764 |
| 4 | P30086 | NA | PEBP1 | phosphatidylethanolamine binding protein 1 | 5037 | ENSG00000089220 |
| 5 | P55060 | NA | CSE1L | CSE1 chromosome segregation 1-like (yeast) | 1434 | ENSG00000124207 |
| 6 | P00367 | NA | GLUD1 | glutamate dehydrogenase 1 | 2746 | ENSG00000148672 |
| 7 | Q06323 | NA | PSME1 | proteasome (prosome, macropain) activator subunit 1 (PA28 alpha) | 5720 | ENSG00000092010 |
| 8 | P48735 | NA | IDH2 | isocitrate dehydrogenase 2 (NADP+), mitochondrial | 3418 | ENSG00000182054 |
| 9 | Q9P0J0 | NA | NDUFA13 | NADH dehydrogenase (ubiquinone) 1 alpha subcomplex, 13 | 51079 | ENSG00000186010 |
| 10 | P16949 | NA | STMN1 | stathmin 1 | 3925 | ENSG00000117632 |
| 11 | Q86UP2 | NA | KTN1 | kinectin 1 (kinesin receptor) | 3895 | ENSG00000126777 |
| 12 | P49588 | NA | AARS | alanyl-tRNA synthetase | 16 | ENSG00000090861 |
| 13 | O75369 | NA | FLNB | filamin B, beta | 2317 | ENSG00000136068 |
| 14 | O14773 | NA | TPP1 | tripeptidyl peptidase I | 1200 | ENSG00000166340 |
| 15 | P16615 | NA | ATP2A2 | ATPase, Ca++ transporting, cardiac muscle, slow twitch 2 | 488 | ENSG00000174437 |
| 16 | P42765 | NA | ACAA2 | acetyl-CoA acyltransferase 2 | 10449 | ENSG00000167315 |
| 17 | P40121 | NA | CAPG | capping protein (actin filament), gelsolin-like | 822 | ENSG00000042493 |
| 18 | P17174 | NA | GOT1 | glutamic-oxaloacetic transaminase 1, soluble (aspartate aminotransferase 1) | 2805 | ENSG00000120053 |
| 19 | P08727 | NA | KRT19 | keratin 19 | 3880 | ENSG00000171345 |
| 20 | Q14697 | NA | GANAB | glucosidase, alpha; neutral AB | 23193 | ENSG00000089597 |
| 21 | P18206 | NA | VCL | vinculin | 7414 | ENSG00000035403 |
| 22 | Q9UK76 | NA | HN1 | hematological and neurological expressed 1 | 51155 | ENSG00000189159 |
| 23 | P08195 | NA | SLC3A2 | solute carrier family 3 (activators of dibasic and neutral amino acid transport), member 2 | 6520 | ENSG00000168003 |
| 24 | Q71U36 | NA | TUBA1A | tubulin, alpha 1a | 7846 | ENSG00000167552 |
| 25 | P78417 | NA | GSTO1 | glutathione S-transferase omega 1 | 9446 | ENSG00000148834 |
| 26 | P49736 | NA | MCM2 | minichromosome maintenance complex component 2 | 4171 | ENSG00000073111 |
| 27 | P38571 | NA | LIPA | lipase A, lysosomal acid, cholesterol esterase | 3988 | ENSG00000107798 |
| 28 | P23381 | NA | WARS | tryptophanyl-tRNA synthetase | 7453 | ENSG00000140105 |
| 29 | Q14980 | NA | NUMA1 | nuclear mitotic apparatus protein 1 | 4926 | ENSG00000137497 |
| 30 | Q9UBQ7 | NA | GRHPR | glyoxylate reductase/hydroxypyruvate reductase | 9380 | ENSG00000137106 |
| 31 | P51649 | NA | ALDH5A1 | aldehyde dehydrogenase 5 family, member A1 | 7915 | ENSG00000112294 |
| 32 | P30044 | NA | PRDX5 | peroxiredoxin 5 | 25824 | ENSG00000126432 |
| 33 | P05556 | NA | ITGB1 | integrin, beta 1 (fibronectin receptor, beta polypeptide, antigen CD29 includes MDF2, MSK12) | 3688 | ENSG00000150093 |
| 34 | P13674 | NA | P4HA1 | prolyl 4-hydroxylase, alpha polypeptide I | 5033 | ENSG00000122884 |
| 35 | P07384 | NA | CAPN1 | calpain 1, (mu/I) large subunit | 823 | ENSG00000014216 |
| 36 | P49591 | NA | SARS | seryl-tRNA synthetase | 6301 | ENSG00000031698 |
| 37 | Q92598 | NA | HSPH1 | heat shock 105kDa/110kDa protein 1 | 10808 | ENSG00000120694 |
| 38 | P17931 | NA | LGALS3 | lectin, galactoside-binding, soluble, 3 | 3958 | ENSG00000131981 |
| 39 | P08133 | NA | ANXA6 | annexin A6 | 309 | ENSG00000197043 |
| 40 | P61225 | NA | RAP2B | RAP2B, member of RAS oncogene family | 5912 | ENSG00000181467 |
| 41 | Q9NQC3 | NA | RTN4 | reticulon 4 | 57142 | ENSG00000115310 |
| 42 | Q15019 | NA | SEPT2 | septin 2 | 4735 | ENSG00000168385 |
| 43 | P09211 | NA | GSTP1 | glutathione S-transferase pi 1 | 2950 | ENSG00000084207 |
| 44 | P33993 | NA | MCM7 | minichromosome maintenance complex component 7 | 4176 | ENSG00000166508 |
| 45 | P68104 | NA | EEF1A1 | eukaryotic translation elongation factor 1 alpha 1 | 1915 | ENSG00000156508 |
| 46 | P02786 | NA | TFRC | transferrin receptor (p90, CD71) | 7037 | ENSG00000072274 |
| 47 | O00425 | NA | IGF2BP3 | insulin-like growth factor 2 mRNA binding protein 3 | 10643 | ENSG00000136231 |
| 48 | P43490 | NA | NAMPT | nicotinamide phosphoribosyltransferase | 10135 | ENSG00000105835 |
| 49 | P17655 | NA | CAPN2 | calpain 2, (m/II) large subunit | 824 | ENSG00000162909 |
| 50 | Q9P258 | NA | RCC2 | regulator of chromosome condensation 2 | 55920 | ENSG00000179051 |
| 51 | P20042 | NA | EIF2S2 | eukaryotic translation initiation factor 2, subunit 2 beta, 38kDa | 8894 | ENSG00000125977 |
| 52 | P22102 | NA | GART | phosphoribosylglycinamide formyltransferase, phosphoribosylglycinamide synthetase, phosphoribosylaminoimidazole synthetase | 2618 | ENSG00000159131 |
| 53 | P22570 | NA | FDXR | ferredoxin reductase | 2232 | ENSG00000161513 |
| 54 | Q05639 | NA | EEF1A2 | eukaryotic translation elongation factor 1 alpha 2 | 1917 | ENSG00000101210 |
| 55 | P49419 | NA | ALDH7A1 | aldehyde dehydrogenase 7 family, member A1 | 501 | ENSG00000164904 |
| 56 | Q92616 | NA | GCN1L1 | GCN1 general control of amino-acid synthesis 1-like 1 (yeast) | 10985 | ENSG00000089154 |
| 57 | P37802 | NA | TAGLN2 | transgelin 2 | 8407 | ENSG00000158710 |
| 58 | P07339 | NA | CTSD | cathepsin D | 1509 | ENSG00000117984 |
| 59 | Q14764 | NA | MVP | major vault protein | 9961 | ENSG00000013364 |
| 60 | Q6PIU2 | NA | NCEH1 | neutral cholesterol ester hydrolase 1 | 57552 | ENSG00000144959 |
| 61 | P21589 | NA | NT5E | 5'-nucleotidase, ecto (CD73) | 4907 | ENSG00000135318 |
| 62 | P11387 | NA | TOP1 | topoisomerase (DNA) I | 7150 | ENSG00000198900 |
| 63 | P53007 | NA | SLC25A1 | solute carrier family 25 (mitochondrial carrier; citrate transporter), member 1 | 6576 | ENSG00000100075 |
| 64 | Q9BPW8 | NA | NIPSNAP1 | nipsnap homolog 1 (C. elegans) | 8508 | ENSG00000184117 |
| 65 | Q16881 | NA | TXNRD1 | thioredoxin reductase 1 | 7296 | ENSG00000198431 |
| 66 | P02792 | NA | FTL | ferritin, light polypeptide | 2512 | ENSG00000087086 |
| 67 | Q9NR30 | NA | DDX21 | DEAD (Asp-Glu-Ala-Asp) box helicase 21 | 9188 | ENSG00000165732 |
| 68 | P49321 | NA | NASP | nuclear autoantigenic sperm protein (histone-binding) | 4678 | ENSG00000132780 |
| 69 | P00491 | NA | PNP | purine nucleoside phosphorylase | 4860 | ENSG00000198805 |
| 70 | Q9HAV4 | NA | XPO5 | exportin 5 | 57510 | ENSG00000124571 |
| 71 | P00338 | NA | LDHA | lactate dehydrogenase A | 3939 | ENSG00000134333 |
| 72 | P56192 | NA | MARS | methionyl-tRNA synthetase | 4141 | ENSG00000166986 |
| 73 | Q15056 | NA | EIF4H | eukaryotic translation initiation factor 4H | 7458 | ENSG00000106682 |
| 74 | O60610 | NA | DIAPH1 | diaphanous homolog 1 (Drosophila) | 1729 | ENSG00000131504 |
| 75 | Q06830 | NA | PRDX1 | peroxiredoxin 1 | 5052 | ENSG00000117450 |
| 76 | Q13011 | NA | ECH1 | enoyl CoA hydratase 1, peroxisomal | 1891 | ENSG00000104823 |
| 77 | P04083 | NA | ANXA1 | annexin A1 | 301 | ENSG00000135046 |
| 78 | P04040 | NA | CAT | catalase | 847 | ENSG00000121691 |
| 79 | P41091 | NA | EIF2S3 | eukaryotic translation initiation factor 2, subunit 3 gamma, 52kDa | 1968 | ENSG00000130741 |
| 80 | P02768 | NA | ALB | albumin | 213 | ENSG00000163631 |
| 81 | P24752 | NA | ACAT1 | acetyl-CoA acetyltransferase 1 | 38 | ENSG00000075239 |
| 82 | P16278 | NA | GLB1 | galactosidase, beta 1 | 2720 | ENSG00000170266 |
| 83 | P07602 | NA | PSAP | prosaposin | 5660 | ENSG00000197746 |
| 84 | P06744 | NA | GPI | glucose-6-phosphate isomerase | 2821 | ENSG00000105220 |
| 85 | Q15758 | NA | SLC1A5 | solute carrier family 1 (neutral amino acid transporter), member 5 | 6510 | ENSG00000105281 |
| 86 | P49748 | NA | ACADVL | acyl-CoA dehydrogenase, very long chain | 37 | ENSG00000072778 |

  
  

| **Database:cellular component      &nbspName:mitochondrial part      &nbspID:GO:0044429** | | | | | | |
| --- | --- | --- | --- | --- | --- | --- |
| C=740; O=15; E=3.81; R=3.93; rawP=5.08e-06; adjP=4.80e-05 | | | | | | |
| Index | UserID | Value | Gene Symbol | Gene Name | EntrezGene | Ensembl |
| 1 | P04040 | NA | CAT | catalase | 847 | ENSG00000121691 |
| 2 | P51649 | NA | ALDH5A1 | aldehyde dehydrogenase 5 family, member A1 | 7915 | ENSG00000112294 |
| 3 | P17931 | NA | LGALS3 | lectin, galactoside-binding, soluble, 3 | 3958 | ENSG00000131981 |
| 4 | P22570 | NA | FDXR | ferredoxin reductase | 2232 | ENSG00000161513 |
| 5 | P30086 | NA | PEBP1 | phosphatidylethanolamine binding protein 1 | 5037 | ENSG00000089220 |
| 6 | P42765 | NA | ACAA2 | acetyl-CoA acyltransferase 2 | 10449 | ENSG00000167315 |
| 7 | P24752 | NA | ACAT1 | acetyl-CoA acetyltransferase 1 | 38 | ENSG00000075239 |
| 8 | P49419 | NA | ALDH7A1 | aldehyde dehydrogenase 7 family, member A1 | 501 | ENSG00000164904 |
| 9 | P00367 | NA | GLUD1 | glutamate dehydrogenase 1 | 2746 | ENSG00000148672 |
| 10 | P48735 | NA | IDH2 | isocitrate dehydrogenase 2 (NADP+), mitochondrial | 3418 | ENSG00000182054 |
| 11 | P49748 | NA | ACADVL | acyl-CoA dehydrogenase, very long chain | 37 | ENSG00000072778 |
| 12 | Q9P0J0 | NA | NDUFA13 | NADH dehydrogenase (ubiquinone) 1 alpha subcomplex, 13 | 51079 | ENSG00000186010 |
| 13 | P04083 | NA | ANXA1 | annexin A1 | 301 | ENSG00000135046 |
| 14 | Q9BPW8 | NA | NIPSNAP1 | nipsnap homolog 1 (C. elegans) | 8508 | ENSG00000184117 |
| 15 | P53007 | NA | SLC25A1 | solute carrier family 25 (mitochondrial carrier; citrate transporter), member 1 | 6576 | ENSG00000100075 |

  
  

| **Database:cellular component      &nbspName:intracellular organelle      &nbspID:GO:0043229** | | | | | | |
| --- | --- | --- | --- | --- | --- | --- |
| C=10521; O=73; E=54.24; R=1.35; rawP=6.77e-06; adjP=5.83e-05 | | | | | | |
| Index | UserID | Value | Gene Symbol | Gene Name | EntrezGene | Ensembl |
| 1 | P10253 | NA | GAA | glucosidase, alpha; acid | 2548 | ENSG00000171298 |
| 2 | P30086 | NA | PEBP1 | phosphatidylethanolamine binding protein 1 | 5037 | ENSG00000089220 |
| 3 | P55060 | NA | CSE1L | CSE1 chromosome segregation 1-like (yeast) | 1434 | ENSG00000124207 |
| 4 | P00367 | NA | GLUD1 | glutamate dehydrogenase 1 | 2746 | ENSG00000148672 |
| 5 | Q06323 | NA | PSME1 | proteasome (prosome, macropain) activator subunit 1 (PA28 alpha) | 5720 | ENSG00000092010 |
| 6 | P48735 | NA | IDH2 | isocitrate dehydrogenase 2 (NADP+), mitochondrial | 3418 | ENSG00000182054 |
| 7 | P16949 | NA | STMN1 | stathmin 1 | 3925 | ENSG00000117632 |
| 8 | Q9P0J0 | NA | NDUFA13 | NADH dehydrogenase (ubiquinone) 1 alpha subcomplex, 13 | 51079 | ENSG00000186010 |
| 9 | Q86UP2 | NA | KTN1 | kinectin 1 (kinesin receptor) | 3895 | ENSG00000126777 |
| 10 | O75369 | NA | FLNB | filamin B, beta | 2317 | ENSG00000136068 |
| 11 | O14773 | NA | TPP1 | tripeptidyl peptidase I | 1200 | ENSG00000166340 |
| 12 | P16615 | NA | ATP2A2 | ATPase, Ca++ transporting, cardiac muscle, slow twitch 2 | 488 | ENSG00000174437 |
| 13 | P40121 | NA | CAPG | capping protein (actin filament), gelsolin-like | 822 | ENSG00000042493 |
| 14 | P42765 | NA | ACAA2 | acetyl-CoA acyltransferase 2 | 10449 | ENSG00000167315 |
| 15 | P17174 | NA | GOT1 | glutamic-oxaloacetic transaminase 1, soluble (aspartate aminotransferase 1) | 2805 | ENSG00000120053 |
| 16 | P08727 | NA | KRT19 | keratin 19 | 3880 | ENSG00000171345 |
| 17 | Q14697 | NA | GANAB | glucosidase, alpha; neutral AB | 23193 | ENSG00000089597 |
| 18 | P18206 | NA | VCL | vinculin | 7414 | ENSG00000035403 |
| 19 | Q9UK76 | NA | HN1 | hematological and neurological expressed 1 | 51155 | ENSG00000189159 |
| 20 | P08195 | NA | SLC3A2 | solute carrier family 3 (activators of dibasic and neutral amino acid transport), member 2 | 6520 | ENSG00000168003 |
| 21 | Q71U36 | NA | TUBA1A | tubulin, alpha 1a | 7846 | ENSG00000167552 |
| 22 | P49736 | NA | MCM2 | minichromosome maintenance complex component 2 | 4171 | ENSG00000073111 |
| 23 | P38571 | NA | LIPA | lipase A, lysosomal acid, cholesterol esterase | 3988 | ENSG00000107798 |
| 24 | Q14980 | NA | NUMA1 | nuclear mitotic apparatus protein 1 | 4926 | ENSG00000137497 |
| 25 | Q9UBQ7 | NA | GRHPR | glyoxylate reductase/hydroxypyruvate reductase | 9380 | ENSG00000137106 |
| 26 | P51649 | NA | ALDH5A1 | aldehyde dehydrogenase 5 family, member A1 | 7915 | ENSG00000112294 |
| 27 | P30044 | NA | PRDX5 | peroxiredoxin 5 | 25824 | ENSG00000126432 |
| 28 | P05556 | NA | ITGB1 | integrin, beta 1 (fibronectin receptor, beta polypeptide, antigen CD29 includes MDF2, MSK12) | 3688 | ENSG00000150093 |
| 29 | P13674 | NA | P4HA1 | prolyl 4-hydroxylase, alpha polypeptide I | 5033 | ENSG00000122884 |
| 30 | P49591 | NA | SARS | seryl-tRNA synthetase | 6301 | ENSG00000031698 |
| 31 | Q92598 | NA | HSPH1 | heat shock 105kDa/110kDa protein 1 | 10808 | ENSG00000120694 |
| 32 | P17931 | NA | LGALS3 | lectin, galactoside-binding, soluble, 3 | 3958 | ENSG00000131981 |
| 33 | P08133 | NA | ANXA6 | annexin A6 | 309 | ENSG00000197043 |
| 34 | P61225 | NA | RAP2B | RAP2B, member of RAS oncogene family | 5912 | ENSG00000181467 |
| 35 | Q9NQC3 | NA | RTN4 | reticulon 4 | 57142 | ENSG00000115310 |
| 36 | Q15019 | NA | SEPT2 | septin 2 | 4735 | ENSG00000168385 |
| 37 | P09211 | NA | GSTP1 | glutathione S-transferase pi 1 | 2950 | ENSG00000084207 |
| 38 | P33993 | NA | MCM7 | minichromosome maintenance complex component 7 | 4176 | ENSG00000166508 |
| 39 | P68104 | NA | EEF1A1 | eukaryotic translation elongation factor 1 alpha 1 | 1915 | ENSG00000156508 |
| 40 | P02786 | NA | TFRC | transferrin receptor (p90, CD71) | 7037 | ENSG00000072274 |
| 41 | O00425 | NA | IGF2BP3 | insulin-like growth factor 2 mRNA binding protein 3 | 10643 | ENSG00000136231 |
| 42 | P17655 | NA | CAPN2 | calpain 2, (m/II) large subunit | 824 | ENSG00000162909 |
| 43 | Q9P258 | NA | RCC2 | regulator of chromosome condensation 2 | 55920 | ENSG00000179051 |
| 44 | P22570 | NA | FDXR | ferredoxin reductase | 2232 | ENSG00000161513 |
| 45 | Q05639 | NA | EEF1A2 | eukaryotic translation elongation factor 1 alpha 2 | 1917 | ENSG00000101210 |
| 46 | P49419 | NA | ALDH7A1 | aldehyde dehydrogenase 7 family, member A1 | 501 | ENSG00000164904 |
| 47 | Q92616 | NA | GCN1L1 | GCN1 general control of amino-acid synthesis 1-like 1 (yeast) | 10985 | ENSG00000089154 |
| 48 | P37802 | NA | TAGLN2 | transgelin 2 | 8407 | ENSG00000158710 |
| 49 | P07339 | NA | CTSD | cathepsin D | 1509 | ENSG00000117984 |
| 50 | Q14764 | NA | MVP | major vault protein | 9961 | ENSG00000013364 |
| 51 | Q6PIU2 | NA | NCEH1 | neutral cholesterol ester hydrolase 1 | 57552 | ENSG00000144959 |
| 52 | P11387 | NA | TOP1 | topoisomerase (DNA) I | 7150 | ENSG00000198900 |
| 53 | P53007 | NA | SLC25A1 | solute carrier family 25 (mitochondrial carrier; citrate transporter), member 1 | 6576 | ENSG00000100075 |
| 54 | Q9BPW8 | NA | NIPSNAP1 | nipsnap homolog 1 (C. elegans) | 8508 | ENSG00000184117 |
| 55 | Q16881 | NA | TXNRD1 | thioredoxin reductase 1 | 7296 | ENSG00000198431 |
| 56 | Q9NR30 | NA | DDX21 | DEAD (Asp-Glu-Ala-Asp) box helicase 21 | 9188 | ENSG00000165732 |
| 57 | P00491 | NA | PNP | purine nucleoside phosphorylase | 4860 | ENSG00000198805 |
| 58 | P49321 | NA | NASP | nuclear autoantigenic sperm protein (histone-binding) | 4678 | ENSG00000132780 |
| 59 | Q9HAV4 | NA | XPO5 | exportin 5 | 57510 | ENSG00000124571 |
| 60 | P00338 | NA | LDHA | lactate dehydrogenase A | 3939 | ENSG00000134333 |
| 61 | P56192 | NA | MARS | methionyl-tRNA synthetase | 4141 | ENSG00000166986 |
| 62 | O60610 | NA | DIAPH1 | diaphanous homolog 1 (Drosophila) | 1729 | ENSG00000131504 |
| 63 | Q13011 | NA | ECH1 | enoyl CoA hydratase 1, peroxisomal | 1891 | ENSG00000104823 |
| 64 | Q06830 | NA | PRDX1 | peroxiredoxin 1 | 5052 | ENSG00000117450 |
| 65 | P04083 | NA | ANXA1 | annexin A1 | 301 | ENSG00000135046 |
| 66 | P04040 | NA | CAT | catalase | 847 | ENSG00000121691 |
| 67 | P02768 | NA | ALB | albumin | 213 | ENSG00000163631 |
| 68 | P24752 | NA | ACAT1 | acetyl-CoA acetyltransferase 1 | 38 | ENSG00000075239 |
| 69 | P16278 | NA | GLB1 | galactosidase, beta 1 | 2720 | ENSG00000170266 |
| 70 | P07602 | NA | PSAP | prosaposin | 5660 | ENSG00000197746 |
| 71 | P06744 | NA | GPI | glucose-6-phosphate isomerase | 2821 | ENSG00000105220 |
| 72 | Q15758 | NA | SLC1A5 | solute carrier family 1 (neutral amino acid transporter), member 5 | 6510 | ENSG00000105281 |
| 73 | P49748 | NA | ACADVL | acyl-CoA dehydrogenase, very long chain | 37 | ENSG00000072778 |

  
  

| **Database:cellular component      &nbspName:organelle      &nbspID:GO:0043226** | | | | | | |
| --- | --- | --- | --- | --- | --- | --- |
| C=10536; O=73; E=54.32; R=1.34; rawP=7.29e-06; adjP=5.83e-05 | | | | | | |
| Index | UserID | Value | Gene Symbol | Gene Name | EntrezGene | Ensembl |
| 1 | P10253 | NA | GAA | glucosidase, alpha; acid | 2548 | ENSG00000171298 |
| 2 | P30086 | NA | PEBP1 | phosphatidylethanolamine binding protein 1 | 5037 | ENSG00000089220 |
| 3 | P55060 | NA | CSE1L | CSE1 chromosome segregation 1-like (yeast) | 1434 | ENSG00000124207 |
| 4 | P00367 | NA | GLUD1 | glutamate dehydrogenase 1 | 2746 | ENSG00000148672 |
| 5 | Q06323 | NA | PSME1 | proteasome (prosome, macropain) activator subunit 1 (PA28 alpha) | 5720 | ENSG00000092010 |
| 6 | P48735 | NA | IDH2 | isocitrate dehydrogenase 2 (NADP+), mitochondrial | 3418 | ENSG00000182054 |
| 7 | P16949 | NA | STMN1 | stathmin 1 | 3925 | ENSG00000117632 |
| 8 | Q9P0J0 | NA | NDUFA13 | NADH dehydrogenase (ubiquinone) 1 alpha subcomplex, 13 | 51079 | ENSG00000186010 |
| 9 | Q86UP2 | NA | KTN1 | kinectin 1 (kinesin receptor) | 3895 | ENSG00000126777 |
| 10 | O75369 | NA | FLNB | filamin B, beta | 2317 | ENSG00000136068 |
| 11 | O14773 | NA | TPP1 | tripeptidyl peptidase I | 1200 | ENSG00000166340 |
| 12 | P16615 | NA | ATP2A2 | ATPase, Ca++ transporting, cardiac muscle, slow twitch 2 | 488 | ENSG00000174437 |
| 13 | P40121 | NA | CAPG | capping protein (actin filament), gelsolin-like | 822 | ENSG00000042493 |
| 14 | P42765 | NA | ACAA2 | acetyl-CoA acyltransferase 2 | 10449 | ENSG00000167315 |
| 15 | P17174 | NA | GOT1 | glutamic-oxaloacetic transaminase 1, soluble (aspartate aminotransferase 1) | 2805 | ENSG00000120053 |
| 16 | P08727 | NA | KRT19 | keratin 19 | 3880 | ENSG00000171345 |
| 17 | Q14697 | NA | GANAB | glucosidase, alpha; neutral AB | 23193 | ENSG00000089597 |
| 18 | P18206 | NA | VCL | vinculin | 7414 | ENSG00000035403 |
| 19 | Q9UK76 | NA | HN1 | hematological and neurological expressed 1 | 51155 | ENSG00000189159 |
| 20 | P08195 | NA | SLC3A2 | solute carrier family 3 (activators of dibasic and neutral amino acid transport), member 2 | 6520 | ENSG00000168003 |
| 21 | Q71U36 | NA | TUBA1A | tubulin, alpha 1a | 7846 | ENSG00000167552 |
| 22 | P49736 | NA | MCM2 | minichromosome maintenance complex component 2 | 4171 | ENSG00000073111 |
| 23 | P38571 | NA | LIPA | lipase A, lysosomal acid, cholesterol esterase | 3988 | ENSG00000107798 |
| 24 | Q14980 | NA | NUMA1 | nuclear mitotic apparatus protein 1 | 4926 | ENSG00000137497 |
| 25 | Q9UBQ7 | NA | GRHPR | glyoxylate reductase/hydroxypyruvate reductase | 9380 | ENSG00000137106 |
| 26 | P51649 | NA | ALDH5A1 | aldehyde dehydrogenase 5 family, member A1 | 7915 | ENSG00000112294 |
| 27 | P30044 | NA | PRDX5 | peroxiredoxin 5 | 25824 | ENSG00000126432 |
| 28 | P05556 | NA | ITGB1 | integrin, beta 1 (fibronectin receptor, beta polypeptide, antigen CD29 includes MDF2, MSK12) | 3688 | ENSG00000150093 |
| 29 | P13674 | NA | P4HA1 | prolyl 4-hydroxylase, alpha polypeptide I | 5033 | ENSG00000122884 |
| 30 | P49591 | NA | SARS | seryl-tRNA synthetase | 6301 | ENSG00000031698 |
| 31 | Q92598 | NA | HSPH1 | heat shock 105kDa/110kDa protein 1 | 10808 | ENSG00000120694 |
| 32 | P17931 | NA | LGALS3 | lectin, galactoside-binding, soluble, 3 | 3958 | ENSG00000131981 |
| 33 | P08133 | NA | ANXA6 | annexin A6 | 309 | ENSG00000197043 |
| 34 | P61225 | NA | RAP2B | RAP2B, member of RAS oncogene family | 5912 | ENSG00000181467 |
| 35 | Q9NQC3 | NA | RTN4 | reticulon 4 | 57142 | ENSG00000115310 |
| 36 | Q15019 | NA | SEPT2 | septin 2 | 4735 | ENSG00000168385 |
| 37 | P09211 | NA | GSTP1 | glutathione S-transferase pi 1 | 2950 | ENSG00000084207 |
| 38 | P33993 | NA | MCM7 | minichromosome maintenance complex component 7 | 4176 | ENSG00000166508 |
| 39 | P68104 | NA | EEF1A1 | eukaryotic translation elongation factor 1 alpha 1 | 1915 | ENSG00000156508 |
| 40 | P02786 | NA | TFRC | transferrin receptor (p90, CD71) | 7037 | ENSG00000072274 |
| 41 | O00425 | NA | IGF2BP3 | insulin-like growth factor 2 mRNA binding protein 3 | 10643 | ENSG00000136231 |
| 42 | P17655 | NA | CAPN2 | calpain 2, (m/II) large subunit | 824 | ENSG00000162909 |
| 43 | Q9P258 | NA | RCC2 | regulator of chromosome condensation 2 | 55920 | ENSG00000179051 |
| 44 | P22570 | NA | FDXR | ferredoxin reductase | 2232 | ENSG00000161513 |
| 45 | Q05639 | NA | EEF1A2 | eukaryotic translation elongation factor 1 alpha 2 | 1917 | ENSG00000101210 |
| 46 | P49419 | NA | ALDH7A1 | aldehyde dehydrogenase 7 family, member A1 | 501 | ENSG00000164904 |
| 47 | Q92616 | NA | GCN1L1 | GCN1 general control of amino-acid synthesis 1-like 1 (yeast) | 10985 | ENSG00000089154 |
| 48 | P37802 | NA | TAGLN2 | transgelin 2 | 8407 | ENSG00000158710 |
| 49 | P07339 | NA | CTSD | cathepsin D | 1509 | ENSG00000117984 |
| 50 | Q14764 | NA | MVP | major vault protein | 9961 | ENSG00000013364 |
| 51 | Q6PIU2 | NA | NCEH1 | neutral cholesterol ester hydrolase 1 | 57552 | ENSG00000144959 |
| 52 | P11387 | NA | TOP1 | topoisomerase (DNA) I | 7150 | ENSG00000198900 |
| 53 | P53007 | NA | SLC25A1 | solute carrier family 25 (mitochondrial carrier; citrate transporter), member 1 | 6576 | ENSG00000100075 |
| 54 | Q9BPW8 | NA | NIPSNAP1 | nipsnap homolog 1 (C. elegans) | 8508 | ENSG00000184117 |
| 55 | Q16881 | NA | TXNRD1 | thioredoxin reductase 1 | 7296 | ENSG00000198431 |
| 56 | Q9NR30 | NA | DDX21 | DEAD (Asp-Glu-Ala-Asp) box helicase 21 | 9188 | ENSG00000165732 |
| 57 | P00491 | NA | PNP | purine nucleoside phosphorylase | 4860 | ENSG00000198805 |
| 58 | P49321 | NA | NASP | nuclear autoantigenic sperm protein (histone-binding) | 4678 | ENSG00000132780 |
| 59 | Q9HAV4 | NA | XPO5 | exportin 5 | 57510 | ENSG00000124571 |
| 60 | P00338 | NA | LDHA | lactate dehydrogenase A | 3939 | ENSG00000134333 |
| 61 | P56192 | NA | MARS | methionyl-tRNA synthetase | 4141 | ENSG00000166986 |
| 62 | O60610 | NA | DIAPH1 | diaphanous homolog 1 (Drosophila) | 1729 | ENSG00000131504 |
| 63 | Q13011 | NA | ECH1 | enoyl CoA hydratase 1, peroxisomal | 1891 | ENSG00000104823 |
| 64 | Q06830 | NA | PRDX1 | peroxiredoxin 1 | 5052 | ENSG00000117450 |
| 65 | P04083 | NA | ANXA1 | annexin A1 | 301 | ENSG00000135046 |
| 66 | P04040 | NA | CAT | catalase | 847 | ENSG00000121691 |
| 67 | P02768 | NA | ALB | albumin | 213 | ENSG00000163631 |
| 68 | P24752 | NA | ACAT1 | acetyl-CoA acetyltransferase 1 | 38 | ENSG00000075239 |
| 69 | P16278 | NA | GLB1 | galactosidase, beta 1 | 2720 | ENSG00000170266 |
| 70 | P07602 | NA | PSAP | prosaposin | 5660 | ENSG00000197746 |
| 71 | P06744 | NA | GPI | glucose-6-phosphate isomerase | 2821 | ENSG00000105220 |
| 72 | Q15758 | NA | SLC1A5 | solute carrier family 1 (neutral amino acid transporter), member 5 | 6510 | ENSG00000105281 |
| 73 | P49748 | NA | ACADVL | acyl-CoA dehydrogenase, very long chain | 37 | ENSG00000072778 |

  
  

| **Database:cellular component      &nbspName:envelope      &nbspID:GO:0031975** | | | | | | |
| --- | --- | --- | --- | --- | --- | --- |
| C=829; O=15; E=4.27; R=3.51; rawP=1.96e-05; adjP=0.0001 | | | | | | |
| Index | UserID | Value | Gene Symbol | Gene Name | EntrezGene | Ensembl |
| 1 | P04040 | NA | CAT | catalase | 847 | ENSG00000121691 |
| 2 | P17931 | NA | LGALS3 | lectin, galactoside-binding, soluble, 3 | 3958 | ENSG00000131981 |
| 3 | P30086 | NA | PEBP1 | phosphatidylethanolamine binding protein 1 | 5037 | ENSG00000089220 |
| 4 | P42765 | NA | ACAA2 | acetyl-CoA acyltransferase 2 | 10449 | ENSG00000167315 |
| 5 | P40121 | NA | CAPG | capping protein (actin filament), gelsolin-like | 822 | ENSG00000042493 |
| 6 | P24752 | NA | ACAT1 | acetyl-CoA acetyltransferase 1 | 38 | ENSG00000075239 |
| 7 | P48735 | NA | IDH2 | isocitrate dehydrogenase 2 (NADP+), mitochondrial | 3418 | ENSG00000182054 |
| 8 | Q9NQC3 | NA | RTN4 | reticulon 4 | 57142 | ENSG00000115310 |
| 9 | P37802 | NA | TAGLN2 | transgelin 2 | 8407 | ENSG00000158710 |
| 10 | Q14764 | NA | MVP | major vault protein | 9961 | ENSG00000013364 |
| 11 | P49748 | NA | ACADVL | acyl-CoA dehydrogenase, very long chain | 37 | ENSG00000072778 |
| 12 | Q9P0J0 | NA | NDUFA13 | NADH dehydrogenase (ubiquinone) 1 alpha subcomplex, 13 | 51079 | ENSG00000186010 |
| 13 | P04083 | NA | ANXA1 | annexin A1 | 301 | ENSG00000135046 |
| 14 | Q9BPW8 | NA | NIPSNAP1 | nipsnap homolog 1 (C. elegans) | 8508 | ENSG00000184117 |
| 15 | P53007 | NA | SLC25A1 | solute carrier family 25 (mitochondrial carrier; citrate transporter), member 1 | 6576 | ENSG00000100075 |

  
  

| **Database:cellular component      &nbspName:organelle envelope      &nbspID:GO:0031967** | | | | | | |
| --- | --- | --- | --- | --- | --- | --- |
| C=817; O=15; E=4.21; R=3.56; rawP=1.66e-05; adjP=0.0001 | | | | | | |
| Index | UserID | Value | Gene Symbol | Gene Name | EntrezGene | Ensembl |
| 1 | P04040 | NA | CAT | catalase | 847 | ENSG00000121691 |
| 2 | P17931 | NA | LGALS3 | lectin, galactoside-binding, soluble, 3 | 3958 | ENSG00000131981 |
| 3 | P30086 | NA | PEBP1 | phosphatidylethanolamine binding protein 1 | 5037 | ENSG00000089220 |
| 4 | P42765 | NA | ACAA2 | acetyl-CoA acyltransferase 2 | 10449 | ENSG00000167315 |
| 5 | P40121 | NA | CAPG | capping protein (actin filament), gelsolin-like | 822 | ENSG00000042493 |
| 6 | P24752 | NA | ACAT1 | acetyl-CoA acetyltransferase 1 | 38 | ENSG00000075239 |
| 7 | P48735 | NA | IDH2 | isocitrate dehydrogenase 2 (NADP+), mitochondrial | 3418 | ENSG00000182054 |
| 8 | Q9NQC3 | NA | RTN4 | reticulon 4 | 57142 | ENSG00000115310 |
| 9 | P37802 | NA | TAGLN2 | transgelin 2 | 8407 | ENSG00000158710 |
| 10 | Q14764 | NA | MVP | major vault protein | 9961 | ENSG00000013364 |
| 11 | P49748 | NA | ACADVL | acyl-CoA dehydrogenase, very long chain | 37 | ENSG00000072778 |
| 12 | Q9P0J0 | NA | NDUFA13 | NADH dehydrogenase (ubiquinone) 1 alpha subcomplex, 13 | 51079 | ENSG00000186010 |
| 13 | P04083 | NA | ANXA1 | annexin A1 | 301 | ENSG00000135046 |
| 14 | Q9BPW8 | NA | NIPSNAP1 | nipsnap homolog 1 (C. elegans) | 8508 | ENSG00000184117 |
| 15 | P53007 | NA | SLC25A1 | solute carrier family 25 (mitochondrial carrier; citrate transporter), member 1 | 6576 | ENSG00000100075 |

  
  

| **Database:cellular component      &nbspName:mitochondrial envelope      &nbspID:GO:0005740** | | | | | | |
| --- | --- | --- | --- | --- | --- | --- |
| C=526; O=11; E=2.71; R=4.06; rawP=7.96e-05; adjP=0.0005 | | | | | | |
| Index | UserID | Value | Gene Symbol | Gene Name | EntrezGene | Ensembl |
| 1 | P04040 | NA | CAT | catalase | 847 | ENSG00000121691 |
| 2 | P17931 | NA | LGALS3 | lectin, galactoside-binding, soluble, 3 | 3958 | ENSG00000131981 |
| 3 | P30086 | NA | PEBP1 | phosphatidylethanolamine binding protein 1 | 5037 | ENSG00000089220 |
| 4 | P42765 | NA | ACAA2 | acetyl-CoA acyltransferase 2 | 10449 | ENSG00000167315 |
| 5 | P24752 | NA | ACAT1 | acetyl-CoA acetyltransferase 1 | 38 | ENSG00000075239 |
| 6 | P48735 | NA | IDH2 | isocitrate dehydrogenase 2 (NADP+), mitochondrial | 3418 | ENSG00000182054 |
| 7 | P49748 | NA | ACADVL | acyl-CoA dehydrogenase, very long chain | 37 | ENSG00000072778 |
| 8 | Q9P0J0 | NA | NDUFA13 | NADH dehydrogenase (ubiquinone) 1 alpha subcomplex, 13 | 51079 | ENSG00000186010 |
| 9 | P53007 | NA | SLC25A1 | solute carrier family 25 (mitochondrial carrier; citrate transporter), member 1 | 6576 | ENSG00000100075 |
| 10 | Q9BPW8 | NA | NIPSNAP1 | nipsnap homolog 1 (C. elegans) | 8508 | ENSG00000184117 |
| 11 | P04083 | NA | ANXA1 | annexin A1 | 301 | ENSG00000135046 |

  
  

| **Database:cellular component      &nbspName:intracellular organelle part      &nbspID:GO:0044446** | | | | | | |
| --- | --- | --- | --- | --- | --- | --- |
| C=6690; O=52; E=34.49; R=1.51; rawP=0.0001; adjP=0.0006 | | | | | | |
| Index | UserID | Value | Gene Symbol | Gene Name | EntrezGene | Ensembl |
| 1 | Q92598 | NA | HSPH1 | heat shock 105kDa/110kDa protein 1 | 10808 | ENSG00000120694 |
| 2 | P10253 | NA | GAA | glucosidase, alpha; acid | 2548 | ENSG00000171298 |
| 3 | P17931 | NA | LGALS3 | lectin, galactoside-binding, soluble, 3 | 3958 | ENSG00000131981 |
| 4 | P30086 | NA | PEBP1 | phosphatidylethanolamine binding protein 1 | 5037 | ENSG00000089220 |
| 5 | P61225 | NA | RAP2B | RAP2B, member of RAS oncogene family | 5912 | ENSG00000181467 |
| 6 | Q9NQC3 | NA | RTN4 | reticulon 4 | 57142 | ENSG00000115310 |
| 7 | P00367 | NA | GLUD1 | glutamate dehydrogenase 1 | 2746 | ENSG00000148672 |
| 8 | Q06323 | NA | PSME1 | proteasome (prosome, macropain) activator subunit 1 (PA28 alpha) | 5720 | ENSG00000092010 |
| 9 | P48735 | NA | IDH2 | isocitrate dehydrogenase 2 (NADP+), mitochondrial | 3418 | ENSG00000182054 |
| 10 | Q15019 | NA | SEPT2 | septin 2 | 4735 | ENSG00000168385 |
| 11 | P33993 | NA | MCM7 | minichromosome maintenance complex component 7 | 4176 | ENSG00000166508 |
| 12 | P16949 | NA | STMN1 | stathmin 1 | 3925 | ENSG00000117632 |
| 13 | Q9P0J0 | NA | NDUFA13 | NADH dehydrogenase (ubiquinone) 1 alpha subcomplex, 13 | 51079 | ENSG00000186010 |
| 14 | Q86UP2 | NA | KTN1 | kinectin 1 (kinesin receptor) | 3895 | ENSG00000126777 |
| 15 | P02786 | NA | TFRC | transferrin receptor (p90, CD71) | 7037 | ENSG00000072274 |
| 16 | O75369 | NA | FLNB | filamin B, beta | 2317 | ENSG00000136068 |
| 17 | P17655 | NA | CAPN2 | calpain 2, (m/II) large subunit | 824 | ENSG00000162909 |
| 18 | O14773 | NA | TPP1 | tripeptidyl peptidase I | 1200 | ENSG00000166340 |
| 19 | Q9P258 | NA | RCC2 | regulator of chromosome condensation 2 | 55920 | ENSG00000179051 |
| 20 | P16615 | NA | ATP2A2 | ATPase, Ca++ transporting, cardiac muscle, slow twitch 2 | 488 | ENSG00000174437 |
| 21 | P22570 | NA | FDXR | ferredoxin reductase | 2232 | ENSG00000161513 |
| 22 | P40121 | NA | CAPG | capping protein (actin filament), gelsolin-like | 822 | ENSG00000042493 |
| 23 | P42765 | NA | ACAA2 | acetyl-CoA acyltransferase 2 | 10449 | ENSG00000167315 |
| 24 | P49419 | NA | ALDH7A1 | aldehyde dehydrogenase 7 family, member A1 | 501 | ENSG00000164904 |
| 25 | P37802 | NA | TAGLN2 | transgelin 2 | 8407 | ENSG00000158710 |
| 26 | P08727 | NA | KRT19 | keratin 19 | 3880 | ENSG00000171345 |
| 27 | Q14764 | NA | MVP | major vault protein | 9961 | ENSG00000013364 |
| 28 | P07339 | NA | CTSD | cathepsin D | 1509 | ENSG00000117984 |
| 29 | P18206 | NA | VCL | vinculin | 7414 | ENSG00000035403 |
| 30 | Q14697 | NA | GANAB | glucosidase, alpha; neutral AB | 23193 | ENSG00000089597 |
| 31 | Q9BPW8 | NA | NIPSNAP1 | nipsnap homolog 1 (C. elegans) | 8508 | ENSG00000184117 |
| 32 | P53007 | NA | SLC25A1 | solute carrier family 25 (mitochondrial carrier; citrate transporter), member 1 | 6576 | ENSG00000100075 |
| 33 | P11387 | NA | TOP1 | topoisomerase (DNA) I | 7150 | ENSG00000198900 |
| 34 | Q16881 | NA | TXNRD1 | thioredoxin reductase 1 | 7296 | ENSG00000198431 |
| 35 | Q9NR30 | NA | DDX21 | DEAD (Asp-Glu-Ala-Asp) box helicase 21 | 9188 | ENSG00000165732 |
| 36 | Q71U36 | NA | TUBA1A | tubulin, alpha 1a | 7846 | ENSG00000167552 |
| 37 | Q9HAV4 | NA | XPO5 | exportin 5 | 57510 | ENSG00000124571 |
| 38 | P49736 | NA | MCM2 | minichromosome maintenance complex component 2 | 4171 | ENSG00000073111 |
| 39 | O60610 | NA | DIAPH1 | diaphanous homolog 1 (Drosophila) | 1729 | ENSG00000131504 |
| 40 | P04083 | NA | ANXA1 | annexin A1 | 301 | ENSG00000135046 |
| 41 | P04040 | NA | CAT | catalase | 847 | ENSG00000121691 |
| 42 | Q14980 | NA | NUMA1 | nuclear mitotic apparatus protein 1 | 4926 | ENSG00000137497 |
| 43 | Q9UBQ7 | NA | GRHPR | glyoxylate reductase/hydroxypyruvate reductase | 9380 | ENSG00000137106 |
| 44 | P51649 | NA | ALDH5A1 | aldehyde dehydrogenase 5 family, member A1 | 7915 | ENSG00000112294 |
| 45 | P02768 | NA | ALB | albumin | 213 | ENSG00000163631 |
| 46 | P30044 | NA | PRDX5 | peroxiredoxin 5 | 25824 | ENSG00000126432 |
| 47 | P24752 | NA | ACAT1 | acetyl-CoA acetyltransferase 1 | 38 | ENSG00000075239 |
| 48 | P13674 | NA | P4HA1 | prolyl 4-hydroxylase, alpha polypeptide I | 5033 | ENSG00000122884 |
| 49 | P16278 | NA | GLB1 | galactosidase, beta 1 | 2720 | ENSG00000170266 |
| 50 | P07602 | NA | PSAP | prosaposin | 5660 | ENSG00000197746 |
| 51 | P06744 | NA | GPI | glucose-6-phosphate isomerase | 2821 | ENSG00000105220 |
| 52 | P49748 | NA | ACADVL | acyl-CoA dehydrogenase, very long chain | 37 | ENSG00000072778 |

  
  
  
  

---

WebGestalt is currently developed and maintained by Jing Wang and Bing Zhang at the  Zhang Lab. Other people who have made significant contribution to the project include Dexter Duncan, Stefan Kirov, Zhiao Shi, and Jay Snoddy.  
  
**Funding credits:** NIH/NIAAA (U01 AA016662, U01 AA013512); NIH/NIDA (P01 DA015027); NIH/NIMH (P50 MH078028, P50 MH096972); NIH/NCI (U24 CA159988); NIH/NIGMS (R01 GM088822).
