## Supplementary material for "PGRMC1 phosphorylation and cell plasticity 1: glycolysis, mitochondria, tumor growth": File S7: final_sig_kegg_file_1438487532.html

Anchored HTML File of EIDs

|  |  |
| --- | --- |
|  | WEB-based GEne SeT AnaLysis Toolkit |
|  | ***Translating gene lists into biological insights...*** |

---

  

| **Database:KEGG pathway      &nbspName:Metabolic pathways      &nbspID:01100** | | | | | | |
| --- | --- | --- | --- | --- | --- | --- |
| C=1130; O=21; E=4.79; R=4.38; rawP=6.87e-09; adjP=1.72e-07 | | | | | | |
| Index | UserID | Value | Gene Symbol | Gene Name | EntrezGene | Ensembl |
| 1 | Q04446 | NA | GBE1 | glucan (1,4-alpha-), branching enzyme 1 | 2632 | ENSG00000114480 |
| 2 | P10253 | NA | GAA | glucosidase, alpha; acid | 2548 | ENSG00000171298 |
| 3 | P00491 | NA | PNP | purine nucleoside phosphorylase | 4860 | ENSG00000198805 |
| 4 | Q16851 | NA | UGP2 | UDP-glucose pyrophosphorylase 2 | 7360 | ENSG00000169764 |
| 5 | P00338 | NA | LDHA | lactate dehydrogenase A | 3939 | ENSG00000134333 |
| 6 | P00367 | NA | GLUD1 | glutamate dehydrogenase 1 | 2746 | ENSG00000148672 |
| 7 | P48735 | NA | IDH2 | isocitrate dehydrogenase 2 (NADP+), mitochondrial | 3418 | ENSG00000182054 |
| 8 | P04040 | NA | CAT | catalase | 847 | ENSG00000121691 |
| 9 | Q9UBQ7 | NA | GRHPR | glyoxylate reductase/hydroxypyruvate reductase | 9380 | ENSG00000137106 |
| 10 | P22102 | NA | GART | phosphoribosylglycinamide formyltransferase, phosphoribosylglycinamide synthetase, phosphoribosylaminoimidazole synthetase | 2618 | ENSG00000159131 |
| 11 | P51649 | NA | ALDH5A1 | aldehyde dehydrogenase 5 family, member A1 | 7915 | ENSG00000112294 |
| 12 | P42765 | NA | ACAA2 | acetyl-CoA acyltransferase 2 | 10449 | ENSG00000167315 |
| 13 | P13674 | NA | P4HA1 | prolyl 4-hydroxylase, alpha polypeptide I | 5033 | ENSG00000122884 |
| 14 | P24752 | NA | ACAT1 | acetyl-CoA acetyltransferase 1 | 38 | ENSG00000075239 |
| 15 | P16278 | NA | GLB1 | galactosidase, beta 1 | 2720 | ENSG00000170266 |
| 16 | P17174 | NA | GOT1 | glutamic-oxaloacetic transaminase 1, soluble (aspartate aminotransferase 1) | 2805 | ENSG00000120053 |
| 17 | P49419 | NA | ALDH7A1 | aldehyde dehydrogenase 7 family, member A1 | 501 | ENSG00000164904 |
| 18 | P06744 | NA | GPI | glucose-6-phosphate isomerase | 2821 | ENSG00000105220 |
| 19 | P49748 | NA | ACADVL | acyl-CoA dehydrogenase, very long chain | 37 | ENSG00000072778 |
| 20 | Q14697 | NA | GANAB | glucosidase, alpha; neutral AB | 23193 | ENSG00000089597 |
| 21 | P21589 | NA | NT5E | 5'-nucleotidase, ecto (CD73) | 4907 | ENSG00000135318 |

  
  

| **Database:KEGG pathway      &nbspName:Pyruvate metabolism      &nbspID:00620** | | | | | | |
| --- | --- | --- | --- | --- | --- | --- |
| C=40; O=4; E=0.17; R=23.58; rawP=2.45e-05; adjP=0.0001 | | | | | | |
| Index | UserID | Value | Gene Symbol | Gene Name | EntrezGene | Ensembl |
| 1 | P24752 | NA | ACAT1 | acetyl-CoA acetyltransferase 1 | 38 | ENSG00000075239 |
| 2 | Q9UBQ7 | NA | GRHPR | glyoxylate reductase/hydroxypyruvate reductase | 9380 | ENSG00000137106 |
| 3 | P49419 | NA | ALDH7A1 | aldehyde dehydrogenase 7 family, member A1 | 501 | ENSG00000164904 |
| 4 | P00338 | NA | LDHA | lactate dehydrogenase A | 3939 | ENSG00000134333 |

  
  

| **Database:KEGG pathway      &nbspName:Lysosome      &nbspID:04142** | | | | | | |
| --- | --- | --- | --- | --- | --- | --- |
| C=121; O=6; E=0.51; R=11.69; rawP=1.27e-05; adjP=0.0001 | | | | | | |
| Index | UserID | Value | Gene Symbol | Gene Name | EntrezGene | Ensembl |
| 1 | O14773 | NA | TPP1 | tripeptidyl peptidase I | 1200 | ENSG00000166340 |
| 2 | P16278 | NA | GLB1 | galactosidase, beta 1 | 2720 | ENSG00000170266 |
| 3 | P10253 | NA | GAA | glucosidase, alpha; acid | 2548 | ENSG00000171298 |
| 4 | P07602 | NA | PSAP | prosaposin | 5660 | ENSG00000197746 |
| 5 | P07339 | NA | CTSD | cathepsin D | 1509 | ENSG00000117984 |
| 6 | P38571 | NA | LIPA | lipase A, lysosomal acid, cholesterol esterase | 3988 | ENSG00000107798 |

  
  

| **Database:KEGG pathway      &nbspName:Fatty acid metabolism      &nbspID:00071** | | | | | | |
| --- | --- | --- | --- | --- | --- | --- |
| C=43; O=4; E=0.18; R=21.94; rawP=3.28e-05; adjP=0.0001 | | | | | | |
| Index | UserID | Value | Gene Symbol | Gene Name | EntrezGene | Ensembl |
| 1 | P24752 | NA | ACAT1 | acetyl-CoA acetyltransferase 1 | 38 | ENSG00000075239 |
| 2 | P49419 | NA | ALDH7A1 | aldehyde dehydrogenase 7 family, member A1 | 501 | ENSG00000164904 |
| 3 | P49748 | NA | ACADVL | acyl-CoA dehydrogenase, very long chain | 37 | ENSG00000072778 |
| 4 | P42765 | NA | ACAA2 | acetyl-CoA acyltransferase 2 | 10449 | ENSG00000167315 |

  
  

| **Database:KEGG pathway      &nbspName:Tryptophan metabolism      &nbspID:00380** | | | | | | |
| --- | --- | --- | --- | --- | --- | --- |
| C=42; O=4; E=0.18; R=22.46; rawP=2.98e-05; adjP=0.0001 | | | | | | |
| Index | UserID | Value | Gene Symbol | Gene Name | EntrezGene | Ensembl |
| 1 | P24752 | NA | ACAT1 | acetyl-CoA acetyltransferase 1 | 38 | ENSG00000075239 |
| 2 | P04040 | NA | CAT | catalase | 847 | ENSG00000121691 |
| 3 | P49419 | NA | ALDH7A1 | aldehyde dehydrogenase 7 family, member A1 | 501 | ENSG00000164904 |
| 4 | P23381 | NA | WARS | tryptophanyl-tRNA synthetase | 7453 | ENSG00000140105 |

  
  

| **Database:KEGG pathway      &nbspName:Peroxisome      &nbspID:04146** | | | | | | |
| --- | --- | --- | --- | --- | --- | --- |
| C=79; O=5; E=0.33; R=14.93; rawP=2.15e-05; adjP=0.0001 | | | | | | |
| Index | UserID | Value | Gene Symbol | Gene Name | EntrezGene | Ensembl |
| 1 | P04040 | NA | CAT | catalase | 847 | ENSG00000121691 |
| 2 | P48735 | NA | IDH2 | isocitrate dehydrogenase 2 (NADP+), mitochondrial | 3418 | ENSG00000182054 |
| 3 | P30044 | NA | PRDX5 | peroxiredoxin 5 | 25824 | ENSG00000126432 |
| 4 | Q13011 | NA | ECH1 | enoyl CoA hydratase 1, peroxisomal | 1891 | ENSG00000104823 |
| 5 | Q06830 | NA | PRDX1 | peroxiredoxin 1 | 5052 | ENSG00000117450 |

  
  

| **Database:KEGG pathway      &nbspName:Aminoacyl-tRNA biosynthesis      &nbspID:00970** | | | | | | |
| --- | --- | --- | --- | --- | --- | --- |
| C=41; O=4; E=0.17; R=23.01; rawP=2.71e-05; adjP=0.0001 | | | | | | |
| Index | UserID | Value | Gene Symbol | Gene Name | EntrezGene | Ensembl |
| 1 | P56192 | NA | MARS | methionyl-tRNA synthetase | 4141 | ENSG00000166986 |
| 2 | P49591 | NA | SARS | seryl-tRNA synthetase | 6301 | ENSG00000031698 |
| 3 | P49588 | NA | AARS | alanyl-tRNA synthetase | 16 | ENSG00000090861 |
| 4 | P23381 | NA | WARS | tryptophanyl-tRNA synthetase | 7453 | ENSG00000140105 |

  
  

| **Database:KEGG pathway      &nbspName:Starch and sucrose metabolism      &nbspID:00500** | | | | | | |
| --- | --- | --- | --- | --- | --- | --- |
| C=54; O=4; E=0.23; R=17.47; rawP=8.10e-05; adjP=0.0002 | | | | | | |
| Index | UserID | Value | Gene Symbol | Gene Name | EntrezGene | Ensembl |
| 1 | Q04446 | NA | GBE1 | glucan (1,4-alpha-), branching enzyme 1 | 2632 | ENSG00000114480 |
| 2 | P10253 | NA | GAA | glucosidase, alpha; acid | 2548 | ENSG00000171298 |
| 3 | P06744 | NA | GPI | glucose-6-phosphate isomerase | 2821 | ENSG00000105220 |
| 4 | Q16851 | NA | UGP2 | UDP-glucose pyrophosphorylase 2 | 7360 | ENSG00000169764 |

  
  

| **Database:KEGG pathway      &nbspName:Arginine and proline metabolism      &nbspID:00330** | | | | | | |
| --- | --- | --- | --- | --- | --- | --- |
| C=54; O=4; E=0.23; R=17.47; rawP=8.10e-05; adjP=0.0002 | | | | | | |
| Index | UserID | Value | Gene Symbol | Gene Name | EntrezGene | Ensembl |
| 1 | P13674 | NA | P4HA1 | prolyl 4-hydroxylase, alpha polypeptide I | 5033 | ENSG00000122884 |
| 2 | P49419 | NA | ALDH7A1 | aldehyde dehydrogenase 7 family, member A1 | 501 | ENSG00000164904 |
| 3 | P17174 | NA | GOT1 | glutamic-oxaloacetic transaminase 1, soluble (aspartate aminotransferase 1) | 2805 | ENSG00000120053 |
| 4 | P00367 | NA | GLUD1 | glutamate dehydrogenase 1 | 2746 | ENSG00000148672 |

  
  

| **Database:KEGG pathway      &nbspName:Nicotinate and nicotinamide metabolism      &nbspID:00760** | | | | | | |
| --- | --- | --- | --- | --- | --- | --- |
| C=24; O=3; E=0.10; R=29.48; rawP=0.0001; adjP=0.0003 | | | | | | |
| Index | UserID | Value | Gene Symbol | Gene Name | EntrezGene | Ensembl |
| 1 | P00491 | NA | PNP | purine nucleoside phosphorylase | 4860 | ENSG00000198805 |
| 2 | P21589 | NA | NT5E | 5'-nucleotidase, ecto (CD73) | 4907 | ENSG00000135318 |
| 3 | P43490 | NA | NAMPT | nicotinamide phosphoribosyltransferase | 10135 | ENSG00000105835 |

  
  

| **Database:KEGG pathway      &nbspName:Galactose metabolism      &nbspID:00052** | | | | | | |
| --- | --- | --- | --- | --- | --- | --- |
| C=27; O=3; E=0.11; R=26.21; rawP=0.0002; adjP=0.0005 | | | | | | |
| Index | UserID | Value | Gene Symbol | Gene Name | EntrezGene | Ensembl |
| 1 | P16278 | NA | GLB1 | galactosidase, beta 1 | 2720 | ENSG00000170266 |
| 2 | P10253 | NA | GAA | glucosidase, alpha; acid | 2548 | ENSG00000171298 |
| 3 | Q16851 | NA | UGP2 | UDP-glucose pyrophosphorylase 2 | 7360 | ENSG00000169764 |

  
  

| **Database:KEGG pathway      &nbspName:Propanoate metabolism      &nbspID:00640** | | | | | | |
| --- | --- | --- | --- | --- | --- | --- |
| C=32; O=3; E=0.14; R=22.11; rawP=0.0003; adjP=0.0006 | | | | | | |
| Index | UserID | Value | Gene Symbol | Gene Name | EntrezGene | Ensembl |
| 1 | P24752 | NA | ACAT1 | acetyl-CoA acetyltransferase 1 | 38 | ENSG00000075239 |
| 2 | P49419 | NA | ALDH7A1 | aldehyde dehydrogenase 7 family, member A1 | 501 | ENSG00000164904 |
| 3 | P00338 | NA | LDHA | lactate dehydrogenase A | 3939 | ENSG00000134333 |

  
  

| **Database:KEGG pathway      &nbspName:Alanine, aspartate and glutamate metabolism      &nbspID:00250** | | | | | | |
| --- | --- | --- | --- | --- | --- | --- |
| C=32; O=3; E=0.14; R=22.11; rawP=0.0003; adjP=0.0006 | | | | | | |
| Index | UserID | Value | Gene Symbol | Gene Name | EntrezGene | Ensembl |
| 1 | P17174 | NA | GOT1 | glutamic-oxaloacetic transaminase 1, soluble (aspartate aminotransferase 1) | 2805 | ENSG00000120053 |
| 2 | P00367 | NA | GLUD1 | glutamate dehydrogenase 1 | 2746 | ENSG00000148672 |
| 3 | P51649 | NA | ALDH5A1 | aldehyde dehydrogenase 5 family, member A1 | 7915 | ENSG00000112294 |

  
  

| **Database:KEGG pathway      &nbspName:RNA transport      &nbspID:03013** | | | | | | |
| --- | --- | --- | --- | --- | --- | --- |
| C=151; O=5; E=0.64; R=7.81; rawP=0.0005; adjP=0.0009 | | | | | | |
| Index | UserID | Value | Gene Symbol | Gene Name | EntrezGene | Ensembl |
| 1 | P20042 | NA | EIF2S2 | eukaryotic translation initiation factor 2, subunit 2 beta, 38kDa | 8894 | ENSG00000125977 |
| 2 | P41091 | NA | EIF2S3 | eukaryotic translation initiation factor 2, subunit 3 gamma, 52kDa | 1968 | ENSG00000130741 |
| 3 | P68104 | NA | EEF1A1 | eukaryotic translation elongation factor 1 alpha 1 | 1915 | ENSG00000156508 |
| 4 | Q9HAV4 | NA | XPO5 | exportin 5 | 57510 | ENSG00000124571 |
| 5 | Q05639 | NA | EEF1A2 | eukaryotic translation elongation factor 1 alpha 2 | 1917 | ENSG00000101210 |

  
  

  
  

---

WebGestalt is currently developed and maintained by Jing Wang and Bing Zhang at the  Zhang Lab. Other people who have made significant contribution to the project include Dexter Duncan, Stefan Kirov, Zhiao Shi, and Jay Snoddy.  
  
**Funding credits:** NIH/NIAAA (U01 AA016662, U01 AA013512); NIH/NIDA (P01 DA015027); NIH/NIMH (P50 MH078028, P50 MH096972); NIH/NCI (U24 CA159988); NIH/NIGMS (R01 GM088822).
