## Supplementary material for "PGRMC1 phosphorylation and cell plasticity 1: glycolysis, mitochondria, tumor growth": File S7: final_pc_geneset_file_1438487532.html

Anchored HTML File of EIDs

|  |  |
| --- | --- |
|  | WEB-based GEne SeT AnaLysis Toolkit |
|  | ***Translating gene lists into biological insights...*** |

---

  

| **Database:Pathway Commons pathway      &nbspName:Metabolism      &nbspID:DB\_ID:634** | | | | | | |
| --- | --- | --- | --- | --- | --- | --- |
| C=824; O=17; E=3.49; R=4.87; rawP=5.43e-08; adjP=6.35e-06 | | | | | | |
| Index | UserID | Value | Gene Symbol | Gene Name | EntrezGene | Ensembl |
| 1 | Q16881 | NA | TXNRD1 | thioredoxin reductase 1 | 7296 | ENSG00000198431 |
| 2 | P00491 | NA | PNP | purine nucleoside phosphorylase | 4860 | ENSG00000198805 |
| 3 | Q16851 | NA | UGP2 | UDP-glucose pyrophosphorylase 2 | 7360 | ENSG00000169764 |
| 4 | P00367 | NA | GLUD1 | glutamate dehydrogenase 1 | 2746 | ENSG00000148672 |
| 5 | Q06323 | NA | PSME1 | proteasome (prosome, macropain) activator subunit 1 (PA28 alpha) | 5720 | ENSG00000092010 |
| 6 | P48735 | NA | IDH2 | isocitrate dehydrogenase 2 (NADP+), mitochondrial | 3418 | ENSG00000182054 |
| 7 | P78417 | NA | GSTO1 | glutathione S-transferase omega 1 | 9446 | ENSG00000148834 |
| 8 | P09211 | NA | GSTP1 | glutathione S-transferase pi 1 | 2950 | ENSG00000084207 |
| 9 | Q9P0J0 | NA | NDUFA13 | NADH dehydrogenase (ubiquinone) 1 alpha subcomplex, 13 | 51079 | ENSG00000186010 |
| 10 | P43490 | NA | NAMPT | nicotinamide phosphoribosyltransferase | 10135 | ENSG00000105835 |
| 11 | P04040 | NA | CAT | catalase | 847 | ENSG00000121691 |
| 12 | Q9UBQ7 | NA | GRHPR | glyoxylate reductase/hydroxypyruvate reductase | 9380 | ENSG00000137106 |
| 13 | P22102 | NA | GART | phosphoribosylglycinamide formyltransferase, phosphoribosylglycinamide synthetase, phosphoribosylaminoimidazole synthetase | 2618 | ENSG00000159131 |
| 14 | P24752 | NA | ACAT1 | acetyl-CoA acetyltransferase 1 | 38 | ENSG00000075239 |
| 15 | P17174 | NA | GOT1 | glutamic-oxaloacetic transaminase 1, soluble (aspartate aminotransferase 1) | 2805 | ENSG00000120053 |
| 16 | P49419 | NA | ALDH7A1 | aldehyde dehydrogenase 7 family, member A1 | 501 | ENSG00000164904 |
| 17 | P21589 | NA | NT5E | 5'-nucleotidase, ecto (CD73) | 4907 | ENSG00000135318 |

  
  

| **Database:Pathway Commons pathway      &nbspName:Glucose metabolism      &nbspID:DB\_ID:898** | | | | | | |
| --- | --- | --- | --- | --- | --- | --- |
| C=36; O=5; E=0.15; R=32.76; rawP=4.14e-07; adjP=2.42e-05 | | | | | | |
| Index | UserID | Value | Gene Symbol | Gene Name | EntrezGene | Ensembl |
| 1 | Q04446 | NA | GBE1 | glucan (1,4-alpha-), branching enzyme 1 | 2632 | ENSG00000114480 |
| 2 | P17174 | NA | GOT1 | glutamic-oxaloacetic transaminase 1, soluble (aspartate aminotransferase 1) | 2805 | ENSG00000120053 |
| 3 | P06744 | NA | GPI | glucose-6-phosphate isomerase | 2821 | ENSG00000105220 |
| 4 | Q16851 | NA | UGP2 | UDP-glucose pyrophosphorylase 2 | 7360 | ENSG00000169764 |
| 5 | P53007 | NA | SLC25A1 | solute carrier family 25 (mitochondrial carrier; citrate transporter), member 1 | 6576 | ENSG00000100075 |

  
  

| **Database:Pathway Commons pathway      &nbspName:Purine metabolism      &nbspID:DB\_ID:638** | | | | | | |
| --- | --- | --- | --- | --- | --- | --- |
| C=24; O=4; E=0.10; R=39.31; rawP=3.00e-06; adjP=8.78e-05 | | | | | | |
| Index | UserID | Value | Gene Symbol | Gene Name | EntrezGene | Ensembl |
| 1 | P04040 | NA | CAT | catalase | 847 | ENSG00000121691 |
| 2 | P22102 | NA | GART | phosphoribosylglycinamide formyltransferase, phosphoribosylglycinamide synthetase, phosphoribosylaminoimidazole synthetase | 2618 | ENSG00000159131 |
| 3 | P00491 | NA | PNP | purine nucleoside phosphorylase | 4860 | ENSG00000198805 |
| 4 | P21589 | NA | NT5E | 5'-nucleotidase, ecto (CD73) | 4907 | ENSG00000135318 |

  
  

| **Database:Pathway Commons pathway      &nbspName:Cytosolic tRNA aminoacylation      &nbspID:DB\_ID:520** | | | | | | |
| --- | --- | --- | --- | --- | --- | --- |
| C=24; O=4; E=0.10; R=39.31; rawP=3.00e-06; adjP=8.78e-05 | | | | | | |
| Index | UserID | Value | Gene Symbol | Gene Name | EntrezGene | Ensembl |
| 1 | P56192 | NA | MARS | methionyl-tRNA synthetase | 4141 | ENSG00000166986 |
| 2 | P49591 | NA | SARS | seryl-tRNA synthetase | 6301 | ENSG00000031698 |
| 3 | P49588 | NA | AARS | alanyl-tRNA synthetase | 16 | ENSG00000090861 |
| 4 | P23381 | NA | WARS | tryptophanyl-tRNA synthetase | 7453 | ENSG00000140105 |

  
  

| **Database:Pathway Commons pathway      &nbspName:Metabolism of nucleotides      &nbspID:DB\_ID:639** | | | | | | |
| --- | --- | --- | --- | --- | --- | --- |
| C=58; O=5; E=0.25; R=20.33; rawP=4.68e-06; adjP=0.0001 | | | | | | |
| Index | UserID | Value | Gene Symbol | Gene Name | EntrezGene | Ensembl |
| 1 | P04040 | NA | CAT | catalase | 847 | ENSG00000121691 |
| 2 | Q16881 | NA | TXNRD1 | thioredoxin reductase 1 | 7296 | ENSG00000198431 |
| 3 | P22102 | NA | GART | phosphoribosylglycinamide formyltransferase, phosphoribosylglycinamide synthetase, phosphoribosylaminoimidazole synthetase | 2618 | ENSG00000159131 |
| 4 | P00491 | NA | PNP | purine nucleoside phosphorylase | 4860 | ENSG00000198805 |
| 5 | P21589 | NA | NT5E | 5'-nucleotidase, ecto (CD73) | 4907 | ENSG00000135318 |

  
  

| **Database:Pathway Commons pathway      &nbspName:Metabolism of carbohydrates      &nbspID:DB\_ID:905** | | | | | | |
| --- | --- | --- | --- | --- | --- | --- |
| C=92; O=5; E=0.39; R=12.82; rawP=4.48e-05; adjP=0.0002 | | | | | | |
| Index | UserID | Value | Gene Symbol | Gene Name | EntrezGene | Ensembl |
| 1 | Q04446 | NA | GBE1 | glucan (1,4-alpha-), branching enzyme 1 | 2632 | ENSG00000114480 |
| 2 | P17174 | NA | GOT1 | glutamic-oxaloacetic transaminase 1, soluble (aspartate aminotransferase 1) | 2805 | ENSG00000120053 |
| 3 | P06744 | NA | GPI | glucose-6-phosphate isomerase | 2821 | ENSG00000105220 |
| 4 | Q16851 | NA | UGP2 | UDP-glucose pyrophosphorylase 2 | 7360 | ENSG00000169764 |
| 5 | P53007 | NA | SLC25A1 | solute carrier family 25 (mitochondrial carrier; citrate transporter), member 1 | 6576 | ENSG00000100075 |

  
  

| **Database:Pathway Commons pathway      &nbspName:Arf6 downstream pathway      &nbspID:DB\_ID:1585** | | | | | | |
| --- | --- | --- | --- | --- | --- | --- |
| C=1288; O=16; E=5.46; R=2.93; rawP=9.00e-05; adjP=0.0002 | | | | | | |
| Index | UserID | Value | Gene Symbol | Gene Name | EntrezGene | Ensembl |
| 1 | P08195 | NA | SLC3A2 | solute carrier family 3 (activators of dibasic and neutral amino acid transport), member 2 | 6520 | ENSG00000168003 |
| 2 | P30086 | NA | PEBP1 | phosphatidylethanolamine binding protein 1 | 5037 | ENSG00000089220 |
| 3 | P00338 | NA | LDHA | lactate dehydrogenase A | 3939 | ENSG00000134333 |
| 4 | P55060 | NA | CSE1L | CSE1 chromosome segregation 1-like (yeast) | 1434 | ENSG00000124207 |
| 5 | P16949 | NA | STMN1 | stathmin 1 | 3925 | ENSG00000117632 |
| 6 | O60610 | NA | DIAPH1 | diaphanous homolog 1 (Drosophila) | 1729 | ENSG00000131504 |
| 7 | Q06830 | NA | PRDX1 | peroxiredoxin 1 | 5052 | ENSG00000117450 |
| 8 | P02786 | NA | TFRC | transferrin receptor (p90, CD71) | 7037 | ENSG00000072274 |
| 9 | P17655 | NA | CAPN2 | calpain 2, (m/II) large subunit | 824 | ENSG00000162909 |
| 10 | P04040 | NA | CAT | catalase | 847 | ENSG00000121691 |
| 11 | P22570 | NA | FDXR | ferredoxin reductase | 2232 | ENSG00000161513 |
| 12 | P05556 | NA | ITGB1 | integrin, beta 1 (fibronectin receptor, beta polypeptide, antigen CD29 includes MDF2, MSK12) | 3688 | ENSG00000150093 |
| 13 | P08727 | NA | KRT19 | keratin 19 | 3880 | ENSG00000171345 |
| 14 | P07339 | NA | CTSD | cathepsin D | 1509 | ENSG00000117984 |
| 15 | P18206 | NA | VCL | vinculin | 7414 | ENSG00000035403 |
| 16 | P21589 | NA | NT5E | 5'-nucleotidase, ecto (CD73) | 4907 | ENSG00000135318 |

  
  

| **Database:Pathway Commons pathway      &nbspName:IFN-gamma pathway      &nbspID:DB\_ID:1529** | | | | | | |
| --- | --- | --- | --- | --- | --- | --- |
| C=1296; O=16; E=5.50; R=2.91; rawP=9.68e-05; adjP=0.0002 | | | | | | |
| Index | UserID | Value | Gene Symbol | Gene Name | EntrezGene | Ensembl |
| 1 | P08195 | NA | SLC3A2 | solute carrier family 3 (activators of dibasic and neutral amino acid transport), member 2 | 6520 | ENSG00000168003 |
| 2 | P30086 | NA | PEBP1 | phosphatidylethanolamine binding protein 1 | 5037 | ENSG00000089220 |
| 3 | P00338 | NA | LDHA | lactate dehydrogenase A | 3939 | ENSG00000134333 |
| 4 | P55060 | NA | CSE1L | CSE1 chromosome segregation 1-like (yeast) | 1434 | ENSG00000124207 |
| 5 | P16949 | NA | STMN1 | stathmin 1 | 3925 | ENSG00000117632 |
| 6 | O60610 | NA | DIAPH1 | diaphanous homolog 1 (Drosophila) | 1729 | ENSG00000131504 |
| 7 | Q06830 | NA | PRDX1 | peroxiredoxin 1 | 5052 | ENSG00000117450 |
| 8 | P02786 | NA | TFRC | transferrin receptor (p90, CD71) | 7037 | ENSG00000072274 |
| 9 | P17655 | NA | CAPN2 | calpain 2, (m/II) large subunit | 824 | ENSG00000162909 |
| 10 | P04040 | NA | CAT | catalase | 847 | ENSG00000121691 |
| 11 | P22570 | NA | FDXR | ferredoxin reductase | 2232 | ENSG00000161513 |
| 12 | P05556 | NA | ITGB1 | integrin, beta 1 (fibronectin receptor, beta polypeptide, antigen CD29 includes MDF2, MSK12) | 3688 | ENSG00000150093 |
| 13 | P08727 | NA | KRT19 | keratin 19 | 3880 | ENSG00000171345 |
| 14 | P07339 | NA | CTSD | cathepsin D | 1509 | ENSG00000117984 |
| 15 | P18206 | NA | VCL | vinculin | 7414 | ENSG00000035403 |
| 16 | P21589 | NA | NT5E | 5'-nucleotidase, ecto (CD73) | 4907 | ENSG00000135318 |

  
  

| **Database:Pathway Commons pathway      &nbspName:IL5-mediated signaling events      &nbspID:DB\_ID:1627** | | | | | | |
| --- | --- | --- | --- | --- | --- | --- |
| C=1292; O=16; E=5.48; R=2.92; rawP=9.34e-05; adjP=0.0002 | | | | | | |
| Index | UserID | Value | Gene Symbol | Gene Name | EntrezGene | Ensembl |
| 1 | P08195 | NA | SLC3A2 | solute carrier family 3 (activators of dibasic and neutral amino acid transport), member 2 | 6520 | ENSG00000168003 |
| 2 | P30086 | NA | PEBP1 | phosphatidylethanolamine binding protein 1 | 5037 | ENSG00000089220 |
| 3 | P00338 | NA | LDHA | lactate dehydrogenase A | 3939 | ENSG00000134333 |
| 4 | P55060 | NA | CSE1L | CSE1 chromosome segregation 1-like (yeast) | 1434 | ENSG00000124207 |
| 5 | P16949 | NA | STMN1 | stathmin 1 | 3925 | ENSG00000117632 |
| 6 | O60610 | NA | DIAPH1 | diaphanous homolog 1 (Drosophila) | 1729 | ENSG00000131504 |
| 7 | Q06830 | NA | PRDX1 | peroxiredoxin 1 | 5052 | ENSG00000117450 |
| 8 | P02786 | NA | TFRC | transferrin receptor (p90, CD71) | 7037 | ENSG00000072274 |
| 9 | P17655 | NA | CAPN2 | calpain 2, (m/II) large subunit | 824 | ENSG00000162909 |
| 10 | P04040 | NA | CAT | catalase | 847 | ENSG00000121691 |
| 11 | P22570 | NA | FDXR | ferredoxin reductase | 2232 | ENSG00000161513 |
| 12 | P05556 | NA | ITGB1 | integrin, beta 1 (fibronectin receptor, beta polypeptide, antigen CD29 includes MDF2, MSK12) | 3688 | ENSG00000150093 |
| 13 | P08727 | NA | KRT19 | keratin 19 | 3880 | ENSG00000171345 |
| 14 | P07339 | NA | CTSD | cathepsin D | 1509 | ENSG00000117984 |
| 15 | P18206 | NA | VCL | vinculin | 7414 | ENSG00000035403 |
| 16 | P21589 | NA | NT5E | 5'-nucleotidase, ecto (CD73) | 4907 | ENSG00000135318 |

  
  

| **Database:Pathway Commons pathway      &nbspName:Thrombin/protease-activated receptor (PAR) pathway      &nbspID:DB\_ID:1552** | | | | | | |
| --- | --- | --- | --- | --- | --- | --- |
| C=1300; O=16; E=5.51; R=2.90; rawP=0.0001; adjP=0.0002 | | | | | | |
| Index | UserID | Value | Gene Symbol | Gene Name | EntrezGene | Ensembl |
| 1 | P08195 | NA | SLC3A2 | solute carrier family 3 (activators of dibasic and neutral amino acid transport), member 2 | 6520 | ENSG00000168003 |
| 2 | P30086 | NA | PEBP1 | phosphatidylethanolamine binding protein 1 | 5037 | ENSG00000089220 |
| 3 | P00338 | NA | LDHA | lactate dehydrogenase A | 3939 | ENSG00000134333 |
| 4 | P55060 | NA | CSE1L | CSE1 chromosome segregation 1-like (yeast) | 1434 | ENSG00000124207 |
| 5 | P16949 | NA | STMN1 | stathmin 1 | 3925 | ENSG00000117632 |
| 6 | O60610 | NA | DIAPH1 | diaphanous homolog 1 (Drosophila) | 1729 | ENSG00000131504 |
| 7 | Q06830 | NA | PRDX1 | peroxiredoxin 1 | 5052 | ENSG00000117450 |
| 8 | P02786 | NA | TFRC | transferrin receptor (p90, CD71) | 7037 | ENSG00000072274 |
| 9 | P17655 | NA | CAPN2 | calpain 2, (m/II) large subunit | 824 | ENSG00000162909 |
| 10 | P04040 | NA | CAT | catalase | 847 | ENSG00000121691 |
| 11 | P22570 | NA | FDXR | ferredoxin reductase | 2232 | ENSG00000161513 |
| 12 | P05556 | NA | ITGB1 | integrin, beta 1 (fibronectin receptor, beta polypeptide, antigen CD29 includes MDF2, MSK12) | 3688 | ENSG00000150093 |
| 13 | P08727 | NA | KRT19 | keratin 19 | 3880 | ENSG00000171345 |
| 14 | P07339 | NA | CTSD | cathepsin D | 1509 | ENSG00000117984 |
| 15 | P18206 | NA | VCL | vinculin | 7414 | ENSG00000035403 |
| 16 | P21589 | NA | NT5E | 5'-nucleotidase, ecto (CD73) | 4907 | ENSG00000135318 |

  
  

| **Database:Pathway Commons pathway      &nbspName:PDGFR-beta signaling pathway      &nbspID:DB\_ID:1540** | | | | | | |
| --- | --- | --- | --- | --- | --- | --- |
| C=1288; O=16; E=5.46; R=2.93; rawP=9.00e-05; adjP=0.0002 | | | | | | |
| Index | UserID | Value | Gene Symbol | Gene Name | EntrezGene | Ensembl |
| 1 | P08195 | NA | SLC3A2 | solute carrier family 3 (activators of dibasic and neutral amino acid transport), member 2 | 6520 | ENSG00000168003 |
| 2 | P30086 | NA | PEBP1 | phosphatidylethanolamine binding protein 1 | 5037 | ENSG00000089220 |
| 3 | P00338 | NA | LDHA | lactate dehydrogenase A | 3939 | ENSG00000134333 |
| 4 | P55060 | NA | CSE1L | CSE1 chromosome segregation 1-like (yeast) | 1434 | ENSG00000124207 |
| 5 | P16949 | NA | STMN1 | stathmin 1 | 3925 | ENSG00000117632 |
| 6 | O60610 | NA | DIAPH1 | diaphanous homolog 1 (Drosophila) | 1729 | ENSG00000131504 |
| 7 | Q06830 | NA | PRDX1 | peroxiredoxin 1 | 5052 | ENSG00000117450 |
| 8 | P02786 | NA | TFRC | transferrin receptor (p90, CD71) | 7037 | ENSG00000072274 |
| 9 | P17655 | NA | CAPN2 | calpain 2, (m/II) large subunit | 824 | ENSG00000162909 |
| 10 | P04040 | NA | CAT | catalase | 847 | ENSG00000121691 |
| 11 | P22570 | NA | FDXR | ferredoxin reductase | 2232 | ENSG00000161513 |
| 12 | P05556 | NA | ITGB1 | integrin, beta 1 (fibronectin receptor, beta polypeptide, antigen CD29 includes MDF2, MSK12) | 3688 | ENSG00000150093 |
| 13 | P08727 | NA | KRT19 | keratin 19 | 3880 | ENSG00000171345 |
| 14 | P07339 | NA | CTSD | cathepsin D | 1509 | ENSG00000117984 |
| 15 | P18206 | NA | VCL | vinculin | 7414 | ENSG00000035403 |
| 16 | P21589 | NA | NT5E | 5'-nucleotidase, ecto (CD73) | 4907 | ENSG00000135318 |

  
  

| **Database:Pathway Commons pathway      &nbspName:Arf6 signaling events      &nbspID:DB\_ID:1554** | | | | | | |
| --- | --- | --- | --- | --- | --- | --- |
| C=1288; O=16; E=5.46; R=2.93; rawP=9.00e-05; adjP=0.0002 | | | | | | |
| Index | UserID | Value | Gene Symbol | Gene Name | EntrezGene | Ensembl |
| 1 | P08195 | NA | SLC3A2 | solute carrier family 3 (activators of dibasic and neutral amino acid transport), member 2 | 6520 | ENSG00000168003 |
| 2 | P30086 | NA | PEBP1 | phosphatidylethanolamine binding protein 1 | 5037 | ENSG00000089220 |
| 3 | P00338 | NA | LDHA | lactate dehydrogenase A | 3939 | ENSG00000134333 |
| 4 | P55060 | NA | CSE1L | CSE1 chromosome segregation 1-like (yeast) | 1434 | ENSG00000124207 |
| 5 | P16949 | NA | STMN1 | stathmin 1 | 3925 | ENSG00000117632 |
| 6 | O60610 | NA | DIAPH1 | diaphanous homolog 1 (Drosophila) | 1729 | ENSG00000131504 |
| 7 | Q06830 | NA | PRDX1 | peroxiredoxin 1 | 5052 | ENSG00000117450 |
| 8 | P02786 | NA | TFRC | transferrin receptor (p90, CD71) | 7037 | ENSG00000072274 |
| 9 | P17655 | NA | CAPN2 | calpain 2, (m/II) large subunit | 824 | ENSG00000162909 |
| 10 | P04040 | NA | CAT | catalase | 847 | ENSG00000121691 |
| 11 | P22570 | NA | FDXR | ferredoxin reductase | 2232 | ENSG00000161513 |
| 12 | P05556 | NA | ITGB1 | integrin, beta 1 (fibronectin receptor, beta polypeptide, antigen CD29 includes MDF2, MSK12) | 3688 | ENSG00000150093 |
| 13 | P08727 | NA | KRT19 | keratin 19 | 3880 | ENSG00000171345 |
| 14 | P07339 | NA | CTSD | cathepsin D | 1509 | ENSG00000117984 |
| 15 | P18206 | NA | VCL | vinculin | 7414 | ENSG00000035403 |
| 16 | P21589 | NA | NT5E | 5'-nucleotidase, ecto (CD73) | 4907 | ENSG00000135318 |

  
  

| **Database:Pathway Commons pathway      &nbspName:Syndecan-1-mediated signaling events      &nbspID:DB\_ID:1454** | | | | | | |
| --- | --- | --- | --- | --- | --- | --- |
| C=1300; O=16; E=5.51; R=2.90; rawP=0.0001; adjP=0.0002 | | | | | | |
| Index | UserID | Value | Gene Symbol | Gene Name | EntrezGene | Ensembl |
| 1 | P08195 | NA | SLC3A2 | solute carrier family 3 (activators of dibasic and neutral amino acid transport), member 2 | 6520 | ENSG00000168003 |
| 2 | P30086 | NA | PEBP1 | phosphatidylethanolamine binding protein 1 | 5037 | ENSG00000089220 |
| 3 | P00338 | NA | LDHA | lactate dehydrogenase A | 3939 | ENSG00000134333 |
| 4 | P55060 | NA | CSE1L | CSE1 chromosome segregation 1-like (yeast) | 1434 | ENSG00000124207 |
| 5 | P16949 | NA | STMN1 | stathmin 1 | 3925 | ENSG00000117632 |
| 6 | O60610 | NA | DIAPH1 | diaphanous homolog 1 (Drosophila) | 1729 | ENSG00000131504 |
| 7 | Q06830 | NA | PRDX1 | peroxiredoxin 1 | 5052 | ENSG00000117450 |
| 8 | P02786 | NA | TFRC | transferrin receptor (p90, CD71) | 7037 | ENSG00000072274 |
| 9 | P17655 | NA | CAPN2 | calpain 2, (m/II) large subunit | 824 | ENSG00000162909 |
| 10 | P04040 | NA | CAT | catalase | 847 | ENSG00000121691 |
| 11 | P22570 | NA | FDXR | ferredoxin reductase | 2232 | ENSG00000161513 |
| 12 | P05556 | NA | ITGB1 | integrin, beta 1 (fibronectin receptor, beta polypeptide, antigen CD29 includes MDF2, MSK12) | 3688 | ENSG00000150093 |
| 13 | P08727 | NA | KRT19 | keratin 19 | 3880 | ENSG00000171345 |
| 14 | P07339 | NA | CTSD | cathepsin D | 1509 | ENSG00000117984 |
| 15 | P18206 | NA | VCL | vinculin | 7414 | ENSG00000035403 |
| 16 | P21589 | NA | NT5E | 5'-nucleotidase, ecto (CD73) | 4907 | ENSG00000135318 |

  
  

| **Database:Pathway Commons pathway      &nbspName:Nectin adhesion pathway      &nbspID:DB\_ID:1472** | | | | | | |
| --- | --- | --- | --- | --- | --- | --- |
| C=1295; O=16; E=5.49; R=2.91; rawP=9.60e-05; adjP=0.0002 | | | | | | |
| Index | UserID | Value | Gene Symbol | Gene Name | EntrezGene | Ensembl |
| 1 | P08195 | NA | SLC3A2 | solute carrier family 3 (activators of dibasic and neutral amino acid transport), member 2 | 6520 | ENSG00000168003 |
| 2 | P30086 | NA | PEBP1 | phosphatidylethanolamine binding protein 1 | 5037 | ENSG00000089220 |
| 3 | P00338 | NA | LDHA | lactate dehydrogenase A | 3939 | ENSG00000134333 |
| 4 | P55060 | NA | CSE1L | CSE1 chromosome segregation 1-like (yeast) | 1434 | ENSG00000124207 |
| 5 | P16949 | NA | STMN1 | stathmin 1 | 3925 | ENSG00000117632 |
| 6 | O60610 | NA | DIAPH1 | diaphanous homolog 1 (Drosophila) | 1729 | ENSG00000131504 |
| 7 | Q06830 | NA | PRDX1 | peroxiredoxin 1 | 5052 | ENSG00000117450 |
| 8 | P02786 | NA | TFRC | transferrin receptor (p90, CD71) | 7037 | ENSG00000072274 |
| 9 | P17655 | NA | CAPN2 | calpain 2, (m/II) large subunit | 824 | ENSG00000162909 |
| 10 | P04040 | NA | CAT | catalase | 847 | ENSG00000121691 |
| 11 | P22570 | NA | FDXR | ferredoxin reductase | 2232 | ENSG00000161513 |
| 12 | P05556 | NA | ITGB1 | integrin, beta 1 (fibronectin receptor, beta polypeptide, antigen CD29 includes MDF2, MSK12) | 3688 | ENSG00000150093 |
| 13 | P08727 | NA | KRT19 | keratin 19 | 3880 | ENSG00000171345 |
| 14 | P07339 | NA | CTSD | cathepsin D | 1509 | ENSG00000117984 |
| 15 | P18206 | NA | VCL | vinculin | 7414 | ENSG00000035403 |
| 16 | P21589 | NA | NT5E | 5'-nucleotidase, ecto (CD73) | 4907 | ENSG00000135318 |

  
  

| **Database:Pathway Commons pathway      &nbspName:Signaling events mediated by focal adhesion kinase      &nbspID:DB\_ID:1574** | | | | | | |
| --- | --- | --- | --- | --- | --- | --- |
| C=1288; O=16; E=5.46; R=2.93; rawP=9.00e-05; adjP=0.0002 | | | | | | |
| Index | UserID | Value | Gene Symbol | Gene Name | EntrezGene | Ensembl |
| 1 | P08195 | NA | SLC3A2 | solute carrier family 3 (activators of dibasic and neutral amino acid transport), member 2 | 6520 | ENSG00000168003 |
| 2 | P30086 | NA | PEBP1 | phosphatidylethanolamine binding protein 1 | 5037 | ENSG00000089220 |
| 3 | P00338 | NA | LDHA | lactate dehydrogenase A | 3939 | ENSG00000134333 |
| 4 | P55060 | NA | CSE1L | CSE1 chromosome segregation 1-like (yeast) | 1434 | ENSG00000124207 |
| 5 | P16949 | NA | STMN1 | stathmin 1 | 3925 | ENSG00000117632 |
| 6 | O60610 | NA | DIAPH1 | diaphanous homolog 1 (Drosophila) | 1729 | ENSG00000131504 |
| 7 | Q06830 | NA | PRDX1 | peroxiredoxin 1 | 5052 | ENSG00000117450 |
| 8 | P02786 | NA | TFRC | transferrin receptor (p90, CD71) | 7037 | ENSG00000072274 |
| 9 | P17655 | NA | CAPN2 | calpain 2, (m/II) large subunit | 824 | ENSG00000162909 |
| 10 | P04040 | NA | CAT | catalase | 847 | ENSG00000121691 |
| 11 | P22570 | NA | FDXR | ferredoxin reductase | 2232 | ENSG00000161513 |
| 12 | P05556 | NA | ITGB1 | integrin, beta 1 (fibronectin receptor, beta polypeptide, antigen CD29 includes MDF2, MSK12) | 3688 | ENSG00000150093 |
| 13 | P08727 | NA | KRT19 | keratin 19 | 3880 | ENSG00000171345 |
| 14 | P07339 | NA | CTSD | cathepsin D | 1509 | ENSG00000117984 |
| 15 | P18206 | NA | VCL | vinculin | 7414 | ENSG00000035403 |
| 16 | P21589 | NA | NT5E | 5'-nucleotidase, ecto (CD73) | 4907 | ENSG00000135318 |

  
  

| **Database:Pathway Commons pathway      &nbspName:LKB1 signaling events      &nbspID:DB\_ID:1649** | | | | | | |
| --- | --- | --- | --- | --- | --- | --- |
| C=1308; O=16; E=5.55; R=2.89; rawP=0.0001; adjP=0.0002 | | | | | | |
| Index | UserID | Value | Gene Symbol | Gene Name | EntrezGene | Ensembl |
| 1 | P08195 | NA | SLC3A2 | solute carrier family 3 (activators of dibasic and neutral amino acid transport), member 2 | 6520 | ENSG00000168003 |
| 2 | P30086 | NA | PEBP1 | phosphatidylethanolamine binding protein 1 | 5037 | ENSG00000089220 |
| 3 | P00338 | NA | LDHA | lactate dehydrogenase A | 3939 | ENSG00000134333 |
| 4 | P55060 | NA | CSE1L | CSE1 chromosome segregation 1-like (yeast) | 1434 | ENSG00000124207 |
| 5 | P16949 | NA | STMN1 | stathmin 1 | 3925 | ENSG00000117632 |
| 6 | O60610 | NA | DIAPH1 | diaphanous homolog 1 (Drosophila) | 1729 | ENSG00000131504 |
| 7 | Q06830 | NA | PRDX1 | peroxiredoxin 1 | 5052 | ENSG00000117450 |
| 8 | P02786 | NA | TFRC | transferrin receptor (p90, CD71) | 7037 | ENSG00000072274 |
| 9 | P17655 | NA | CAPN2 | calpain 2, (m/II) large subunit | 824 | ENSG00000162909 |
| 10 | P04040 | NA | CAT | catalase | 847 | ENSG00000121691 |
| 11 | P22570 | NA | FDXR | ferredoxin reductase | 2232 | ENSG00000161513 |
| 12 | P05556 | NA | ITGB1 | integrin, beta 1 (fibronectin receptor, beta polypeptide, antigen CD29 includes MDF2, MSK12) | 3688 | ENSG00000150093 |
| 13 | P08727 | NA | KRT19 | keratin 19 | 3880 | ENSG00000171345 |
| 14 | P07339 | NA | CTSD | cathepsin D | 1509 | ENSG00000117984 |
| 15 | P18206 | NA | VCL | vinculin | 7414 | ENSG00000035403 |
| 16 | P21589 | NA | NT5E | 5'-nucleotidase, ecto (CD73) | 4907 | ENSG00000135318 |

  
  

| **Database:Pathway Commons pathway      &nbspName:Class I PI3K signaling events mediated by Akt      &nbspID:DB\_ID:1648** | | | | | | |
| --- | --- | --- | --- | --- | --- | --- |
| C=1288; O=16; E=5.46; R=2.93; rawP=9.00e-05; adjP=0.0002 | | | | | | |
| Index | UserID | Value | Gene Symbol | Gene Name | EntrezGene | Ensembl |
| 1 | P08195 | NA | SLC3A2 | solute carrier family 3 (activators of dibasic and neutral amino acid transport), member 2 | 6520 | ENSG00000168003 |
| 2 | P30086 | NA | PEBP1 | phosphatidylethanolamine binding protein 1 | 5037 | ENSG00000089220 |
| 3 | P00338 | NA | LDHA | lactate dehydrogenase A | 3939 | ENSG00000134333 |
| 4 | P55060 | NA | CSE1L | CSE1 chromosome segregation 1-like (yeast) | 1434 | ENSG00000124207 |
| 5 | P16949 | NA | STMN1 | stathmin 1 | 3925 | ENSG00000117632 |
| 6 | O60610 | NA | DIAPH1 | diaphanous homolog 1 (Drosophila) | 1729 | ENSG00000131504 |
| 7 | Q06830 | NA | PRDX1 | peroxiredoxin 1 | 5052 | ENSG00000117450 |
| 8 | P02786 | NA | TFRC | transferrin receptor (p90, CD71) | 7037 | ENSG00000072274 |
| 9 | P17655 | NA | CAPN2 | calpain 2, (m/II) large subunit | 824 | ENSG00000162909 |
| 10 | P04040 | NA | CAT | catalase | 847 | ENSG00000121691 |
| 11 | P22570 | NA | FDXR | ferredoxin reductase | 2232 | ENSG00000161513 |
| 12 | P05556 | NA | ITGB1 | integrin, beta 1 (fibronectin receptor, beta polypeptide, antigen CD29 includes MDF2, MSK12) | 3688 | ENSG00000150093 |
| 13 | P08727 | NA | KRT19 | keratin 19 | 3880 | ENSG00000171345 |
| 14 | P07339 | NA | CTSD | cathepsin D | 1509 | ENSG00000117984 |
| 15 | P18206 | NA | VCL | vinculin | 7414 | ENSG00000035403 |
| 16 | P21589 | NA | NT5E | 5'-nucleotidase, ecto (CD73) | 4907 | ENSG00000135318 |

  
  

| **Database:Pathway Commons pathway      &nbspName:GMCSF-mediated signaling events      &nbspID:DB\_ID:1461** | | | | | | |
| --- | --- | --- | --- | --- | --- | --- |
| C=1292; O=16; E=5.48; R=2.92; rawP=9.34e-05; adjP=0.0002 | | | | | | |
| Index | UserID | Value | Gene Symbol | Gene Name | EntrezGene | Ensembl |
| 1 | P08195 | NA | SLC3A2 | solute carrier family 3 (activators of dibasic and neutral amino acid transport), member 2 | 6520 | ENSG00000168003 |
| 2 | P30086 | NA | PEBP1 | phosphatidylethanolamine binding protein 1 | 5037 | ENSG00000089220 |
| 3 | P00338 | NA | LDHA | lactate dehydrogenase A | 3939 | ENSG00000134333 |
| 4 | P55060 | NA | CSE1L | CSE1 chromosome segregation 1-like (yeast) | 1434 | ENSG00000124207 |
| 5 | P16949 | NA | STMN1 | stathmin 1 | 3925 | ENSG00000117632 |
| 6 | O60610 | NA | DIAPH1 | diaphanous homolog 1 (Drosophila) | 1729 | ENSG00000131504 |
| 7 | Q06830 | NA | PRDX1 | peroxiredoxin 1 | 5052 | ENSG00000117450 |
| 8 | P02786 | NA | TFRC | transferrin receptor (p90, CD71) | 7037 | ENSG00000072274 |
| 9 | P17655 | NA | CAPN2 | calpain 2, (m/II) large subunit | 824 | ENSG00000162909 |
| 10 | P04040 | NA | CAT | catalase | 847 | ENSG00000121691 |
| 11 | P22570 | NA | FDXR | ferredoxin reductase | 2232 | ENSG00000161513 |
| 12 | P05556 | NA | ITGB1 | integrin, beta 1 (fibronectin receptor, beta polypeptide, antigen CD29 includes MDF2, MSK12) | 3688 | ENSG00000150093 |
| 13 | P08727 | NA | KRT19 | keratin 19 | 3880 | ENSG00000171345 |
| 14 | P07339 | NA | CTSD | cathepsin D | 1509 | ENSG00000117984 |
| 15 | P18206 | NA | VCL | vinculin | 7414 | ENSG00000035403 |
| 16 | P21589 | NA | NT5E | 5'-nucleotidase, ecto (CD73) | 4907 | ENSG00000135318 |

  
  

| **Database:Pathway Commons pathway      &nbspName:Endothelins      &nbspID:DB\_ID:1619** | | | | | | |
| --- | --- | --- | --- | --- | --- | --- |
| C=1307; O=16; E=5.54; R=2.89; rawP=0.0001; adjP=0.0002 | | | | | | |
| Index | UserID | Value | Gene Symbol | Gene Name | EntrezGene | Ensembl |
| 1 | P08195 | NA | SLC3A2 | solute carrier family 3 (activators of dibasic and neutral amino acid transport), member 2 | 6520 | ENSG00000168003 |
| 2 | P30086 | NA | PEBP1 | phosphatidylethanolamine binding protein 1 | 5037 | ENSG00000089220 |
| 3 | P00338 | NA | LDHA | lactate dehydrogenase A | 3939 | ENSG00000134333 |
| 4 | P55060 | NA | CSE1L | CSE1 chromosome segregation 1-like (yeast) | 1434 | ENSG00000124207 |
| 5 | P16949 | NA | STMN1 | stathmin 1 | 3925 | ENSG00000117632 |
| 6 | O60610 | NA | DIAPH1 | diaphanous homolog 1 (Drosophila) | 1729 | ENSG00000131504 |
| 7 | Q06830 | NA | PRDX1 | peroxiredoxin 1 | 5052 | ENSG00000117450 |
| 8 | P02786 | NA | TFRC | transferrin receptor (p90, CD71) | 7037 | ENSG00000072274 |
| 9 | P17655 | NA | CAPN2 | calpain 2, (m/II) large subunit | 824 | ENSG00000162909 |
| 10 | P04040 | NA | CAT | catalase | 847 | ENSG00000121691 |
| 11 | P22570 | NA | FDXR | ferredoxin reductase | 2232 | ENSG00000161513 |
| 12 | P05556 | NA | ITGB1 | integrin, beta 1 (fibronectin receptor, beta polypeptide, antigen CD29 includes MDF2, MSK12) | 3688 | ENSG00000150093 |
| 13 | P08727 | NA | KRT19 | keratin 19 | 3880 | ENSG00000171345 |
| 14 | P07339 | NA | CTSD | cathepsin D | 1509 | ENSG00000117984 |
| 15 | P18206 | NA | VCL | vinculin | 7414 | ENSG00000035403 |
| 16 | P21589 | NA | NT5E | 5'-nucleotidase, ecto (CD73) | 4907 | ENSG00000135318 |

  
  

| **Database:Pathway Commons pathway      &nbspName:mTOR signaling pathway      &nbspID:DB\_ID:1571** | | | | | | |
| --- | --- | --- | --- | --- | --- | --- |
| C=1288; O=16; E=5.46; R=2.93; rawP=9.00e-05; adjP=0.0002 | | | | | | |
| Index | UserID | Value | Gene Symbol | Gene Name | EntrezGene | Ensembl |
| 1 | P08195 | NA | SLC3A2 | solute carrier family 3 (activators of dibasic and neutral amino acid transport), member 2 | 6520 | ENSG00000168003 |
| 2 | P30086 | NA | PEBP1 | phosphatidylethanolamine binding protein 1 | 5037 | ENSG00000089220 |
| 3 | P00338 | NA | LDHA | lactate dehydrogenase A | 3939 | ENSG00000134333 |
| 4 | P55060 | NA | CSE1L | CSE1 chromosome segregation 1-like (yeast) | 1434 | ENSG00000124207 |
| 5 | P16949 | NA | STMN1 | stathmin 1 | 3925 | ENSG00000117632 |
| 6 | O60610 | NA | DIAPH1 | diaphanous homolog 1 (Drosophila) | 1729 | ENSG00000131504 |
| 7 | Q06830 | NA | PRDX1 | peroxiredoxin 1 | 5052 | ENSG00000117450 |
| 8 | P02786 | NA | TFRC | transferrin receptor (p90, CD71) | 7037 | ENSG00000072274 |
| 9 | P17655 | NA | CAPN2 | calpain 2, (m/II) large subunit | 824 | ENSG00000162909 |
| 10 | P04040 | NA | CAT | catalase | 847 | ENSG00000121691 |
| 11 | P22570 | NA | FDXR | ferredoxin reductase | 2232 | ENSG00000161513 |
| 12 | P05556 | NA | ITGB1 | integrin, beta 1 (fibronectin receptor, beta polypeptide, antigen CD29 includes MDF2, MSK12) | 3688 | ENSG00000150093 |
| 13 | P08727 | NA | KRT19 | keratin 19 | 3880 | ENSG00000171345 |
| 14 | P07339 | NA | CTSD | cathepsin D | 1509 | ENSG00000117984 |
| 15 | P18206 | NA | VCL | vinculin | 7414 | ENSG00000035403 |
| 16 | P21589 | NA | NT5E | 5'-nucleotidase, ecto (CD73) | 4907 | ENSG00000135318 |

  
  

| **Database:Pathway Commons pathway      &nbspName:PDGF receptor signaling network      &nbspID:DB\_ID:1497** | | | | | | |
| --- | --- | --- | --- | --- | --- | --- |
| C=1293; O=16; E=5.48; R=2.92; rawP=9.42e-05; adjP=0.0002 | | | | | | |
| Index | UserID | Value | Gene Symbol | Gene Name | EntrezGene | Ensembl |
| 1 | P08195 | NA | SLC3A2 | solute carrier family 3 (activators of dibasic and neutral amino acid transport), member 2 | 6520 | ENSG00000168003 |
| 2 | P30086 | NA | PEBP1 | phosphatidylethanolamine binding protein 1 | 5037 | ENSG00000089220 |
| 3 | P00338 | NA | LDHA | lactate dehydrogenase A | 3939 | ENSG00000134333 |
| 4 | P55060 | NA | CSE1L | CSE1 chromosome segregation 1-like (yeast) | 1434 | ENSG00000124207 |
| 5 | P16949 | NA | STMN1 | stathmin 1 | 3925 | ENSG00000117632 |
| 6 | O60610 | NA | DIAPH1 | diaphanous homolog 1 (Drosophila) | 1729 | ENSG00000131504 |
| 7 | Q06830 | NA | PRDX1 | peroxiredoxin 1 | 5052 | ENSG00000117450 |
| 8 | P02786 | NA | TFRC | transferrin receptor (p90, CD71) | 7037 | ENSG00000072274 |
| 9 | P17655 | NA | CAPN2 | calpain 2, (m/II) large subunit | 824 | ENSG00000162909 |
| 10 | P04040 | NA | CAT | catalase | 847 | ENSG00000121691 |
| 11 | P22570 | NA | FDXR | ferredoxin reductase | 2232 | ENSG00000161513 |
| 12 | P05556 | NA | ITGB1 | integrin, beta 1 (fibronectin receptor, beta polypeptide, antigen CD29 includes MDF2, MSK12) | 3688 | ENSG00000150093 |
| 13 | P08727 | NA | KRT19 | keratin 19 | 3880 | ENSG00000171345 |
| 14 | P07339 | NA | CTSD | cathepsin D | 1509 | ENSG00000117984 |
| 15 | P18206 | NA | VCL | vinculin | 7414 | ENSG00000035403 |
| 16 | P21589 | NA | NT5E | 5'-nucleotidase, ecto (CD73) | 4907 | ENSG00000135318 |

  
  

| **Database:Pathway Commons pathway      &nbspName:Proteoglycan syndecan-mediated signaling events      &nbspID:DB\_ID:1637** | | | | | | |
| --- | --- | --- | --- | --- | --- | --- |
| C=1345; O=16; E=5.70; R=2.81; rawP=0.0001; adjP=0.0002 | | | | | | |
| Index | UserID | Value | Gene Symbol | Gene Name | EntrezGene | Ensembl |
| 1 | P08195 | NA | SLC3A2 | solute carrier family 3 (activators of dibasic and neutral amino acid transport), member 2 | 6520 | ENSG00000168003 |
| 2 | P30086 | NA | PEBP1 | phosphatidylethanolamine binding protein 1 | 5037 | ENSG00000089220 |
| 3 | P00338 | NA | LDHA | lactate dehydrogenase A | 3939 | ENSG00000134333 |
| 4 | P55060 | NA | CSE1L | CSE1 chromosome segregation 1-like (yeast) | 1434 | ENSG00000124207 |
| 5 | P16949 | NA | STMN1 | stathmin 1 | 3925 | ENSG00000117632 |
| 6 | O60610 | NA | DIAPH1 | diaphanous homolog 1 (Drosophila) | 1729 | ENSG00000131504 |
| 7 | Q06830 | NA | PRDX1 | peroxiredoxin 1 | 5052 | ENSG00000117450 |
| 8 | P02786 | NA | TFRC | transferrin receptor (p90, CD71) | 7037 | ENSG00000072274 |
| 9 | P17655 | NA | CAPN2 | calpain 2, (m/II) large subunit | 824 | ENSG00000162909 |
| 10 | P04040 | NA | CAT | catalase | 847 | ENSG00000121691 |
| 11 | P22570 | NA | FDXR | ferredoxin reductase | 2232 | ENSG00000161513 |
| 12 | P05556 | NA | ITGB1 | integrin, beta 1 (fibronectin receptor, beta polypeptide, antigen CD29 includes MDF2, MSK12) | 3688 | ENSG00000150093 |
| 13 | P08727 | NA | KRT19 | keratin 19 | 3880 | ENSG00000171345 |
| 14 | P07339 | NA | CTSD | cathepsin D | 1509 | ENSG00000117984 |
| 15 | P18206 | NA | VCL | vinculin | 7414 | ENSG00000035403 |
| 16 | P21589 | NA | NT5E | 5'-nucleotidase, ecto (CD73) | 4907 | ENSG00000135318 |

  
  

| **Database:Pathway Commons pathway      &nbspName:VEGF and VEGFR signaling network      &nbspID:DB\_ID:1575** | | | | | | |
| --- | --- | --- | --- | --- | --- | --- |
| C=1304; O=16; E=5.53; R=2.89; rawP=0.0001; adjP=0.0002 | | | | | | |
| Index | UserID | Value | Gene Symbol | Gene Name | EntrezGene | Ensembl |
| 1 | P08195 | NA | SLC3A2 | solute carrier family 3 (activators of dibasic and neutral amino acid transport), member 2 | 6520 | ENSG00000168003 |
| 2 | P30086 | NA | PEBP1 | phosphatidylethanolamine binding protein 1 | 5037 | ENSG00000089220 |
| 3 | P00338 | NA | LDHA | lactate dehydrogenase A | 3939 | ENSG00000134333 |
| 4 | P55060 | NA | CSE1L | CSE1 chromosome segregation 1-like (yeast) | 1434 | ENSG00000124207 |
| 5 | P16949 | NA | STMN1 | stathmin 1 | 3925 | ENSG00000117632 |
| 6 | O60610 | NA | DIAPH1 | diaphanous homolog 1 (Drosophila) | 1729 | ENSG00000131504 |
| 7 | Q06830 | NA | PRDX1 | peroxiredoxin 1 | 5052 | ENSG00000117450 |
| 8 | P02786 | NA | TFRC | transferrin receptor (p90, CD71) | 7037 | ENSG00000072274 |
| 9 | P17655 | NA | CAPN2 | calpain 2, (m/II) large subunit | 824 | ENSG00000162909 |
| 10 | P04040 | NA | CAT | catalase | 847 | ENSG00000121691 |
| 11 | P22570 | NA | FDXR | ferredoxin reductase | 2232 | ENSG00000161513 |
| 12 | P05556 | NA | ITGB1 | integrin, beta 1 (fibronectin receptor, beta polypeptide, antigen CD29 includes MDF2, MSK12) | 3688 | ENSG00000150093 |
| 13 | P08727 | NA | KRT19 | keratin 19 | 3880 | ENSG00000171345 |
| 14 | P07339 | NA | CTSD | cathepsin D | 1509 | ENSG00000117984 |
| 15 | P18206 | NA | VCL | vinculin | 7414 | ENSG00000035403 |
| 16 | P21589 | NA | NT5E | 5'-nucleotidase, ecto (CD73) | 4907 | ENSG00000135318 |

  
  

| **Database:Pathway Commons pathway      &nbspName:Internalization of ErbB1      &nbspID:DB\_ID:1509** | | | | | | |
| --- | --- | --- | --- | --- | --- | --- |
| C=1288; O=16; E=5.46; R=2.93; rawP=9.00e-05; adjP=0.0002 | | | | | | |
| Index | UserID | Value | Gene Symbol | Gene Name | EntrezGene | Ensembl |
| 1 | P08195 | NA | SLC3A2 | solute carrier family 3 (activators of dibasic and neutral amino acid transport), member 2 | 6520 | ENSG00000168003 |
| 2 | P30086 | NA | PEBP1 | phosphatidylethanolamine binding protein 1 | 5037 | ENSG00000089220 |
| 3 | P00338 | NA | LDHA | lactate dehydrogenase A | 3939 | ENSG00000134333 |
| 4 | P55060 | NA | CSE1L | CSE1 chromosome segregation 1-like (yeast) | 1434 | ENSG00000124207 |
| 5 | P16949 | NA | STMN1 | stathmin 1 | 3925 | ENSG00000117632 |
| 6 | O60610 | NA | DIAPH1 | diaphanous homolog 1 (Drosophila) | 1729 | ENSG00000131504 |
| 7 | Q06830 | NA | PRDX1 | peroxiredoxin 1 | 5052 | ENSG00000117450 |
| 8 | P02786 | NA | TFRC | transferrin receptor (p90, CD71) | 7037 | ENSG00000072274 |
| 9 | P17655 | NA | CAPN2 | calpain 2, (m/II) large subunit | 824 | ENSG00000162909 |
| 10 | P04040 | NA | CAT | catalase | 847 | ENSG00000121691 |
| 11 | P22570 | NA | FDXR | ferredoxin reductase | 2232 | ENSG00000161513 |
| 12 | P05556 | NA | ITGB1 | integrin, beta 1 (fibronectin receptor, beta polypeptide, antigen CD29 includes MDF2, MSK12) | 3688 | ENSG00000150093 |
| 13 | P08727 | NA | KRT19 | keratin 19 | 3880 | ENSG00000171345 |
| 14 | P07339 | NA | CTSD | cathepsin D | 1509 | ENSG00000117984 |
| 15 | P18206 | NA | VCL | vinculin | 7414 | ENSG00000035403 |
| 16 | P21589 | NA | NT5E | 5'-nucleotidase, ecto (CD73) | 4907 | ENSG00000135318 |

  
  

| **Database:Pathway Commons pathway      &nbspName:Sphingosine 1-phosphate (S1P) pathway      &nbspID:DB\_ID:1635** | | | | | | |
| --- | --- | --- | --- | --- | --- | --- |
| C=1311; O=16; E=5.56; R=2.88; rawP=0.0001; adjP=0.0002 | | | | | | |
| Index | UserID | Value | Gene Symbol | Gene Name | EntrezGene | Ensembl |
| 1 | P08195 | NA | SLC3A2 | solute carrier family 3 (activators of dibasic and neutral amino acid transport), member 2 | 6520 | ENSG00000168003 |
| 2 | P30086 | NA | PEBP1 | phosphatidylethanolamine binding protein 1 | 5037 | ENSG00000089220 |
| 3 | P00338 | NA | LDHA | lactate dehydrogenase A | 3939 | ENSG00000134333 |
| 4 | P55060 | NA | CSE1L | CSE1 chromosome segregation 1-like (yeast) | 1434 | ENSG00000124207 |
| 5 | P16949 | NA | STMN1 | stathmin 1 | 3925 | ENSG00000117632 |
| 6 | O60610 | NA | DIAPH1 | diaphanous homolog 1 (Drosophila) | 1729 | ENSG00000131504 |
| 7 | Q06830 | NA | PRDX1 | peroxiredoxin 1 | 5052 | ENSG00000117450 |
| 8 | P02786 | NA | TFRC | transferrin receptor (p90, CD71) | 7037 | ENSG00000072274 |
| 9 | P17655 | NA | CAPN2 | calpain 2, (m/II) large subunit | 824 | ENSG00000162909 |
| 10 | P04040 | NA | CAT | catalase | 847 | ENSG00000121691 |
| 11 | P22570 | NA | FDXR | ferredoxin reductase | 2232 | ENSG00000161513 |
| 12 | P05556 | NA | ITGB1 | integrin, beta 1 (fibronectin receptor, beta polypeptide, antigen CD29 includes MDF2, MSK12) | 3688 | ENSG00000150093 |
| 13 | P08727 | NA | KRT19 | keratin 19 | 3880 | ENSG00000171345 |
| 14 | P07339 | NA | CTSD | cathepsin D | 1509 | ENSG00000117984 |
| 15 | P18206 | NA | VCL | vinculin | 7414 | ENSG00000035403 |
| 16 | P21589 | NA | NT5E | 5'-nucleotidase, ecto (CD73) | 4907 | ENSG00000135318 |

  
  

| **Database:Pathway Commons pathway      &nbspName:Metabolism of amino acids and derivatives      &nbspID:DB\_ID:875** | | | | | | |
| --- | --- | --- | --- | --- | --- | --- |
| C=188; O=6; E=0.80; R=7.53; rawP=0.0001; adjP=0.0002 | | | | | | |
| Index | UserID | Value | Gene Symbol | Gene Name | EntrezGene | Ensembl |
| 1 | P24752 | NA | ACAT1 | acetyl-CoA acetyltransferase 1 | 38 | ENSG00000075239 |
| 2 | P49419 | NA | ALDH7A1 | aldehyde dehydrogenase 7 family, member A1 | 501 | ENSG00000164904 |
| 3 | Q9UBQ7 | NA | GRHPR | glyoxylate reductase/hydroxypyruvate reductase | 9380 | ENSG00000137106 |
| 4 | P17174 | NA | GOT1 | glutamic-oxaloacetic transaminase 1, soluble (aspartate aminotransferase 1) | 2805 | ENSG00000120053 |
| 5 | Q06323 | NA | PSME1 | proteasome (prosome, macropain) activator subunit 1 (PA28 alpha) | 5720 | ENSG00000092010 |
| 6 | P00367 | NA | GLUD1 | glutamate dehydrogenase 1 | 2746 | ENSG00000148672 |

  
  

| **Database:Pathway Commons pathway      &nbspName:TRAIL signaling pathway      &nbspID:DB\_ID:1480** | | | | | | |
| --- | --- | --- | --- | --- | --- | --- |
| C=1328; O=18; E=5.63; R=3.20; rawP=9.33e-06; adjP=0.0002 | | | | | | |
| Index | UserID | Value | Gene Symbol | Gene Name | EntrezGene | Ensembl |
| 1 | P08195 | NA | SLC3A2 | solute carrier family 3 (activators of dibasic and neutral amino acid transport), member 2 | 6520 | ENSG00000168003 |
| 2 | P30086 | NA | PEBP1 | phosphatidylethanolamine binding protein 1 | 5037 | ENSG00000089220 |
| 3 | P00338 | NA | LDHA | lactate dehydrogenase A | 3939 | ENSG00000134333 |
| 4 | P55060 | NA | CSE1L | CSE1 chromosome segregation 1-like (yeast) | 1434 | ENSG00000124207 |
| 5 | P16949 | NA | STMN1 | stathmin 1 | 3925 | ENSG00000117632 |
| 6 | O60610 | NA | DIAPH1 | diaphanous homolog 1 (Drosophila) | 1729 | ENSG00000131504 |
| 7 | Q06830 | NA | PRDX1 | peroxiredoxin 1 | 5052 | ENSG00000117450 |
| 8 | P02786 | NA | TFRC | transferrin receptor (p90, CD71) | 7037 | ENSG00000072274 |
| 9 | P04040 | NA | CAT | catalase | 847 | ENSG00000121691 |
| 10 | Q14980 | NA | NUMA1 | nuclear mitotic apparatus protein 1 | 4926 | ENSG00000137497 |
| 11 | P17655 | NA | CAPN2 | calpain 2, (m/II) large subunit | 824 | ENSG00000162909 |
| 12 | P22570 | NA | FDXR | ferredoxin reductase | 2232 | ENSG00000161513 |
| 13 | P05556 | NA | ITGB1 | integrin, beta 1 (fibronectin receptor, beta polypeptide, antigen CD29 includes MDF2, MSK12) | 3688 | ENSG00000150093 |
| 14 | P08727 | NA | KRT19 | keratin 19 | 3880 | ENSG00000171345 |
| 15 | P07339 | NA | CTSD | cathepsin D | 1509 | ENSG00000117984 |
| 16 | P18206 | NA | VCL | vinculin | 7414 | ENSG00000035403 |
| 17 | P21589 | NA | NT5E | 5'-nucleotidase, ecto (CD73) | 4907 | ENSG00000135318 |
| 18 | P11387 | NA | TOP1 | topoisomerase (DNA) I | 7150 | ENSG00000198900 |

  
  

| **Database:Pathway Commons pathway      &nbspName:PAR1-mediated thrombin signaling events      &nbspID:DB\_ID:1531** | | | | | | |
| --- | --- | --- | --- | --- | --- | --- |
| C=1299; O=16; E=5.51; R=2.90; rawP=9.95e-05; adjP=0.0002 | | | | | | |
| Index | UserID | Value | Gene Symbol | Gene Name | EntrezGene | Ensembl |
| 1 | P08195 | NA | SLC3A2 | solute carrier family 3 (activators of dibasic and neutral amino acid transport), member 2 | 6520 | ENSG00000168003 |
| 2 | P30086 | NA | PEBP1 | phosphatidylethanolamine binding protein 1 | 5037 | ENSG00000089220 |
| 3 | P00338 | NA | LDHA | lactate dehydrogenase A | 3939 | ENSG00000134333 |
| 4 | P55060 | NA | CSE1L | CSE1 chromosome segregation 1-like (yeast) | 1434 | ENSG00000124207 |
| 5 | P16949 | NA | STMN1 | stathmin 1 | 3925 | ENSG00000117632 |
| 6 | O60610 | NA | DIAPH1 | diaphanous homolog 1 (Drosophila) | 1729 | ENSG00000131504 |
| 7 | Q06830 | NA | PRDX1 | peroxiredoxin 1 | 5052 | ENSG00000117450 |
| 8 | P02786 | NA | TFRC | transferrin receptor (p90, CD71) | 7037 | ENSG00000072274 |
| 9 | P17655 | NA | CAPN2 | calpain 2, (m/II) large subunit | 824 | ENSG00000162909 |
| 10 | P04040 | NA | CAT | catalase | 847 | ENSG00000121691 |
| 11 | P22570 | NA | FDXR | ferredoxin reductase | 2232 | ENSG00000161513 |
| 12 | P05556 | NA | ITGB1 | integrin, beta 1 (fibronectin receptor, beta polypeptide, antigen CD29 includes MDF2, MSK12) | 3688 | ENSG00000150093 |
| 13 | P08727 | NA | KRT19 | keratin 19 | 3880 | ENSG00000171345 |
| 14 | P07339 | NA | CTSD | cathepsin D | 1509 | ENSG00000117984 |
| 15 | P18206 | NA | VCL | vinculin | 7414 | ENSG00000035403 |
| 16 | P21589 | NA | NT5E | 5'-nucleotidase, ecto (CD73) | 4907 | ENSG00000135318 |

  
  

| **Database:Pathway Commons pathway      &nbspName:Glypican pathway      &nbspID:DB\_ID:1459** | | | | | | |
| --- | --- | --- | --- | --- | --- | --- |
| C=1338; O=16; E=5.67; R=2.82; rawP=0.0001; adjP=0.0002 | | | | | | |
| Index | UserID | Value | Gene Symbol | Gene Name | EntrezGene | Ensembl |
| 1 | P08195 | NA | SLC3A2 | solute carrier family 3 (activators of dibasic and neutral amino acid transport), member 2 | 6520 | ENSG00000168003 |
| 2 | P30086 | NA | PEBP1 | phosphatidylethanolamine binding protein 1 | 5037 | ENSG00000089220 |
| 3 | P00338 | NA | LDHA | lactate dehydrogenase A | 3939 | ENSG00000134333 |
| 4 | P55060 | NA | CSE1L | CSE1 chromosome segregation 1-like (yeast) | 1434 | ENSG00000124207 |
| 5 | P16949 | NA | STMN1 | stathmin 1 | 3925 | ENSG00000117632 |
| 6 | O60610 | NA | DIAPH1 | diaphanous homolog 1 (Drosophila) | 1729 | ENSG00000131504 |
| 7 | Q06830 | NA | PRDX1 | peroxiredoxin 1 | 5052 | ENSG00000117450 |
| 8 | P02786 | NA | TFRC | transferrin receptor (p90, CD71) | 7037 | ENSG00000072274 |
| 9 | P17655 | NA | CAPN2 | calpain 2, (m/II) large subunit | 824 | ENSG00000162909 |
| 10 | P04040 | NA | CAT | catalase | 847 | ENSG00000121691 |
| 11 | P22570 | NA | FDXR | ferredoxin reductase | 2232 | ENSG00000161513 |
| 12 | P05556 | NA | ITGB1 | integrin, beta 1 (fibronectin receptor, beta polypeptide, antigen CD29 includes MDF2, MSK12) | 3688 | ENSG00000150093 |
| 13 | P08727 | NA | KRT19 | keratin 19 | 3880 | ENSG00000171345 |
| 14 | P07339 | NA | CTSD | cathepsin D | 1509 | ENSG00000117984 |
| 15 | P18206 | NA | VCL | vinculin | 7414 | ENSG00000035403 |
| 16 | P21589 | NA | NT5E | 5'-nucleotidase, ecto (CD73) | 4907 | ENSG00000135318 |

  
  

| **Database:Pathway Commons pathway      &nbspName:ErbB receptor signaling network      &nbspID:DB\_ID:1573** | | | | | | |
| --- | --- | --- | --- | --- | --- | --- |
| C=1312; O=16; E=5.56; R=2.88; rawP=0.0001; adjP=0.0002 | | | | | | |
| Index | UserID | Value | Gene Symbol | Gene Name | EntrezGene | Ensembl |
| 1 | P08195 | NA | SLC3A2 | solute carrier family 3 (activators of dibasic and neutral amino acid transport), member 2 | 6520 | ENSG00000168003 |
| 2 | P30086 | NA | PEBP1 | phosphatidylethanolamine binding protein 1 | 5037 | ENSG00000089220 |
| 3 | P00338 | NA | LDHA | lactate dehydrogenase A | 3939 | ENSG00000134333 |
| 4 | P55060 | NA | CSE1L | CSE1 chromosome segregation 1-like (yeast) | 1434 | ENSG00000124207 |
| 5 | P16949 | NA | STMN1 | stathmin 1 | 3925 | ENSG00000117632 |
| 6 | O60610 | NA | DIAPH1 | diaphanous homolog 1 (Drosophila) | 1729 | ENSG00000131504 |
| 7 | Q06830 | NA | PRDX1 | peroxiredoxin 1 | 5052 | ENSG00000117450 |
| 8 | P02786 | NA | TFRC | transferrin receptor (p90, CD71) | 7037 | ENSG00000072274 |
| 9 | P17655 | NA | CAPN2 | calpain 2, (m/II) large subunit | 824 | ENSG00000162909 |
| 10 | P04040 | NA | CAT | catalase | 847 | ENSG00000121691 |
| 11 | P22570 | NA | FDXR | ferredoxin reductase | 2232 | ENSG00000161513 |
| 12 | P05556 | NA | ITGB1 | integrin, beta 1 (fibronectin receptor, beta polypeptide, antigen CD29 includes MDF2, MSK12) | 3688 | ENSG00000150093 |
| 13 | P08727 | NA | KRT19 | keratin 19 | 3880 | ENSG00000171345 |
| 14 | P07339 | NA | CTSD | cathepsin D | 1509 | ENSG00000117984 |
| 15 | P18206 | NA | VCL | vinculin | 7414 | ENSG00000035403 |
| 16 | P21589 | NA | NT5E | 5'-nucleotidase, ecto (CD73) | 4907 | ENSG00000135318 |

  
  

| **Database:Pathway Commons pathway      &nbspName:Insulin Pathway      &nbspID:DB\_ID:1466** | | | | | | |
| --- | --- | --- | --- | --- | --- | --- |
| C=1288; O=16; E=5.46; R=2.93; rawP=9.00e-05; adjP=0.0002 | | | | | | |
| Index | UserID | Value | Gene Symbol | Gene Name | EntrezGene | Ensembl |
| 1 | P08195 | NA | SLC3A2 | solute carrier family 3 (activators of dibasic and neutral amino acid transport), member 2 | 6520 | ENSG00000168003 |
| 2 | P30086 | NA | PEBP1 | phosphatidylethanolamine binding protein 1 | 5037 | ENSG00000089220 |
| 3 | P00338 | NA | LDHA | lactate dehydrogenase A | 3939 | ENSG00000134333 |
| 4 | P55060 | NA | CSE1L | CSE1 chromosome segregation 1-like (yeast) | 1434 | ENSG00000124207 |
| 5 | P16949 | NA | STMN1 | stathmin 1 | 3925 | ENSG00000117632 |
| 6 | O60610 | NA | DIAPH1 | diaphanous homolog 1 (Drosophila) | 1729 | ENSG00000131504 |
| 7 | Q06830 | NA | PRDX1 | peroxiredoxin 1 | 5052 | ENSG00000117450 |
| 8 | P02786 | NA | TFRC | transferrin receptor (p90, CD71) | 7037 | ENSG00000072274 |
| 9 | P17655 | NA | CAPN2 | calpain 2, (m/II) large subunit | 824 | ENSG00000162909 |
| 10 | P04040 | NA | CAT | catalase | 847 | ENSG00000121691 |
| 11 | P22570 | NA | FDXR | ferredoxin reductase | 2232 | ENSG00000161513 |
| 12 | P05556 | NA | ITGB1 | integrin, beta 1 (fibronectin receptor, beta polypeptide, antigen CD29 includes MDF2, MSK12) | 3688 | ENSG00000150093 |
| 13 | P08727 | NA | KRT19 | keratin 19 | 3880 | ENSG00000171345 |
| 14 | P07339 | NA | CTSD | cathepsin D | 1509 | ENSG00000117984 |
| 15 | P18206 | NA | VCL | vinculin | 7414 | ENSG00000035403 |
| 16 | P21589 | NA | NT5E | 5'-nucleotidase, ecto (CD73) | 4907 | ENSG00000135318 |

  
  

| **Database:Pathway Commons pathway      &nbspName:Alpha9 beta1 integrin signaling events      &nbspID:DB\_ID:1578** | | | | | | |
| --- | --- | --- | --- | --- | --- | --- |
| C=1305; O=16; E=5.53; R=2.89; rawP=0.0001; adjP=0.0002 | | | | | | |
| Index | UserID | Value | Gene Symbol | Gene Name | EntrezGene | Ensembl |
| 1 | P08195 | NA | SLC3A2 | solute carrier family 3 (activators of dibasic and neutral amino acid transport), member 2 | 6520 | ENSG00000168003 |
| 2 | P30086 | NA | PEBP1 | phosphatidylethanolamine binding protein 1 | 5037 | ENSG00000089220 |
| 3 | P00338 | NA | LDHA | lactate dehydrogenase A | 3939 | ENSG00000134333 |
| 4 | P55060 | NA | CSE1L | CSE1 chromosome segregation 1-like (yeast) | 1434 | ENSG00000124207 |
| 5 | P16949 | NA | STMN1 | stathmin 1 | 3925 | ENSG00000117632 |
| 6 | O60610 | NA | DIAPH1 | diaphanous homolog 1 (Drosophila) | 1729 | ENSG00000131504 |
| 7 | Q06830 | NA | PRDX1 | peroxiredoxin 1 | 5052 | ENSG00000117450 |
| 8 | P02786 | NA | TFRC | transferrin receptor (p90, CD71) | 7037 | ENSG00000072274 |
| 9 | P17655 | NA | CAPN2 | calpain 2, (m/II) large subunit | 824 | ENSG00000162909 |
| 10 | P04040 | NA | CAT | catalase | 847 | ENSG00000121691 |
| 11 | P22570 | NA | FDXR | ferredoxin reductase | 2232 | ENSG00000161513 |
| 12 | P05556 | NA | ITGB1 | integrin, beta 1 (fibronectin receptor, beta polypeptide, antigen CD29 includes MDF2, MSK12) | 3688 | ENSG00000150093 |
| 13 | P08727 | NA | KRT19 | keratin 19 | 3880 | ENSG00000171345 |
| 14 | P07339 | NA | CTSD | cathepsin D | 1509 | ENSG00000117984 |
| 15 | P18206 | NA | VCL | vinculin | 7414 | ENSG00000035403 |
| 16 | P21589 | NA | NT5E | 5'-nucleotidase, ecto (CD73) | 4907 | ENSG00000135318 |

  
  

| **Database:Pathway Commons pathway      &nbspName:IL3-mediated signaling events      &nbspID:DB\_ID:1564** | | | | | | |
| --- | --- | --- | --- | --- | --- | --- |
| C=1295; O=16; E=5.49; R=2.91; rawP=9.60e-05; adjP=0.0002 | | | | | | |
| Index | UserID | Value | Gene Symbol | Gene Name | EntrezGene | Ensembl |
| 1 | P08195 | NA | SLC3A2 | solute carrier family 3 (activators of dibasic and neutral amino acid transport), member 2 | 6520 | ENSG00000168003 |
| 2 | P30086 | NA | PEBP1 | phosphatidylethanolamine binding protein 1 | 5037 | ENSG00000089220 |
| 3 | P00338 | NA | LDHA | lactate dehydrogenase A | 3939 | ENSG00000134333 |
| 4 | P55060 | NA | CSE1L | CSE1 chromosome segregation 1-like (yeast) | 1434 | ENSG00000124207 |
| 5 | P16949 | NA | STMN1 | stathmin 1 | 3925 | ENSG00000117632 |
| 6 | O60610 | NA | DIAPH1 | diaphanous homolog 1 (Drosophila) | 1729 | ENSG00000131504 |
| 7 | Q06830 | NA | PRDX1 | peroxiredoxin 1 | 5052 | ENSG00000117450 |
| 8 | P02786 | NA | TFRC | transferrin receptor (p90, CD71) | 7037 | ENSG00000072274 |
| 9 | P17655 | NA | CAPN2 | calpain 2, (m/II) large subunit | 824 | ENSG00000162909 |
| 10 | P04040 | NA | CAT | catalase | 847 | ENSG00000121691 |
| 11 | P22570 | NA | FDXR | ferredoxin reductase | 2232 | ENSG00000161513 |
| 12 | P05556 | NA | ITGB1 | integrin, beta 1 (fibronectin receptor, beta polypeptide, antigen CD29 includes MDF2, MSK12) | 3688 | ENSG00000150093 |
| 13 | P08727 | NA | KRT19 | keratin 19 | 3880 | ENSG00000171345 |
| 14 | P07339 | NA | CTSD | cathepsin D | 1509 | ENSG00000117984 |
| 15 | P18206 | NA | VCL | vinculin | 7414 | ENSG00000035403 |
| 16 | P21589 | NA | NT5E | 5'-nucleotidase, ecto (CD73) | 4907 | ENSG00000135318 |

  
  

| **Database:Pathway Commons pathway      &nbspName:EGFR-dependent Endothelin signaling events      &nbspID:DB\_ID:1603** | | | | | | |
| --- | --- | --- | --- | --- | --- | --- |
| C=1289; O=16; E=5.47; R=2.93; rawP=9.09e-05; adjP=0.0002 | | | | | | |
| Index | UserID | Value | Gene Symbol | Gene Name | EntrezGene | Ensembl |
| 1 | P08195 | NA | SLC3A2 | solute carrier family 3 (activators of dibasic and neutral amino acid transport), member 2 | 6520 | ENSG00000168003 |
| 2 | P30086 | NA | PEBP1 | phosphatidylethanolamine binding protein 1 | 5037 | ENSG00000089220 |
| 3 | P00338 | NA | LDHA | lactate dehydrogenase A | 3939 | ENSG00000134333 |
| 4 | P55060 | NA | CSE1L | CSE1 chromosome segregation 1-like (yeast) | 1434 | ENSG00000124207 |
| 5 | P16949 | NA | STMN1 | stathmin 1 | 3925 | ENSG00000117632 |
| 6 | O60610 | NA | DIAPH1 | diaphanous homolog 1 (Drosophila) | 1729 | ENSG00000131504 |
| 7 | Q06830 | NA | PRDX1 | peroxiredoxin 1 | 5052 | ENSG00000117450 |
| 8 | P02786 | NA | TFRC | transferrin receptor (p90, CD71) | 7037 | ENSG00000072274 |
| 9 | P17655 | NA | CAPN2 | calpain 2, (m/II) large subunit | 824 | ENSG00000162909 |
| 10 | P04040 | NA | CAT | catalase | 847 | ENSG00000121691 |
| 11 | P22570 | NA | FDXR | ferredoxin reductase | 2232 | ENSG00000161513 |
| 12 | P05556 | NA | ITGB1 | integrin, beta 1 (fibronectin receptor, beta polypeptide, antigen CD29 includes MDF2, MSK12) | 3688 | ENSG00000150093 |
| 13 | P08727 | NA | KRT19 | keratin 19 | 3880 | ENSG00000171345 |
| 14 | P07339 | NA | CTSD | cathepsin D | 1509 | ENSG00000117984 |
| 15 | P18206 | NA | VCL | vinculin | 7414 | ENSG00000035403 |
| 16 | P21589 | NA | NT5E | 5'-nucleotidase, ecto (CD73) | 4907 | ENSG00000135318 |

  
  

| **Database:Pathway Commons pathway      &nbspName:Signaling events mediated by Hepatocyte Growth Factor Receptor (c-Met)      &nbspID:DB\_ID:1491** | | | | | | |
| --- | --- | --- | --- | --- | --- | --- |
| C=1293; O=16; E=5.48; R=2.92; rawP=9.42e-05; adjP=0.0002 | | | | | | |
| Index | UserID | Value | Gene Symbol | Gene Name | EntrezGene | Ensembl |
| 1 | P08195 | NA | SLC3A2 | solute carrier family 3 (activators of dibasic and neutral amino acid transport), member 2 | 6520 | ENSG00000168003 |
| 2 | P30086 | NA | PEBP1 | phosphatidylethanolamine binding protein 1 | 5037 | ENSG00000089220 |
| 3 | P00338 | NA | LDHA | lactate dehydrogenase A | 3939 | ENSG00000134333 |
| 4 | P55060 | NA | CSE1L | CSE1 chromosome segregation 1-like (yeast) | 1434 | ENSG00000124207 |
| 5 | P16949 | NA | STMN1 | stathmin 1 | 3925 | ENSG00000117632 |
| 6 | O60610 | NA | DIAPH1 | diaphanous homolog 1 (Drosophila) | 1729 | ENSG00000131504 |
| 7 | Q06830 | NA | PRDX1 | peroxiredoxin 1 | 5052 | ENSG00000117450 |
| 8 | P02786 | NA | TFRC | transferrin receptor (p90, CD71) | 7037 | ENSG00000072274 |
| 9 | P17655 | NA | CAPN2 | calpain 2, (m/II) large subunit | 824 | ENSG00000162909 |
| 10 | P04040 | NA | CAT | catalase | 847 | ENSG00000121691 |
| 11 | P22570 | NA | FDXR | ferredoxin reductase | 2232 | ENSG00000161513 |
| 12 | P05556 | NA | ITGB1 | integrin, beta 1 (fibronectin receptor, beta polypeptide, antigen CD29 includes MDF2, MSK12) | 3688 | ENSG00000150093 |
| 13 | P08727 | NA | KRT19 | keratin 19 | 3880 | ENSG00000171345 |
| 14 | P07339 | NA | CTSD | cathepsin D | 1509 | ENSG00000117984 |
| 15 | P18206 | NA | VCL | vinculin | 7414 | ENSG00000035403 |
| 16 | P21589 | NA | NT5E | 5'-nucleotidase, ecto (CD73) | 4907 | ENSG00000135318 |

  
  

| **Database:Pathway Commons pathway      &nbspName:Gluconeogenesis      &nbspID:DB\_ID:929** | | | | | | |
| --- | --- | --- | --- | --- | --- | --- |
| C=20; O=3; E=0.08; R=35.38; rawP=7.96e-05; adjP=0.0002 | | | | | | |
| Index | UserID | Value | Gene Symbol | Gene Name | EntrezGene | Ensembl |
| 1 | P17174 | NA | GOT1 | glutamic-oxaloacetic transaminase 1, soluble (aspartate aminotransferase 1) | 2805 | ENSG00000120053 |
| 2 | P06744 | NA | GPI | glucose-6-phosphate isomerase | 2821 | ENSG00000105220 |
| 3 | P53007 | NA | SLC25A1 | solute carrier family 25 (mitochondrial carrier; citrate transporter), member 1 | 6576 | ENSG00000100075 |

  
  

| **Database:Pathway Commons pathway      &nbspName:IGF1 pathway      &nbspID:DB\_ID:1482** | | | | | | |
| --- | --- | --- | --- | --- | --- | --- |
| C=1291; O=16; E=5.47; R=2.92; rawP=9.25e-05; adjP=0.0002 | | | | | | |
| Index | UserID | Value | Gene Symbol | Gene Name | EntrezGene | Ensembl |
| 1 | P08195 | NA | SLC3A2 | solute carrier family 3 (activators of dibasic and neutral amino acid transport), member 2 | 6520 | ENSG00000168003 |
| 2 | P30086 | NA | PEBP1 | phosphatidylethanolamine binding protein 1 | 5037 | ENSG00000089220 |
| 3 | P00338 | NA | LDHA | lactate dehydrogenase A | 3939 | ENSG00000134333 |
| 4 | P55060 | NA | CSE1L | CSE1 chromosome segregation 1-like (yeast) | 1434 | ENSG00000124207 |
| 5 | P16949 | NA | STMN1 | stathmin 1 | 3925 | ENSG00000117632 |
| 6 | O60610 | NA | DIAPH1 | diaphanous homolog 1 (Drosophila) | 1729 | ENSG00000131504 |
| 7 | Q06830 | NA | PRDX1 | peroxiredoxin 1 | 5052 | ENSG00000117450 |
| 8 | P02786 | NA | TFRC | transferrin receptor (p90, CD71) | 7037 | ENSG00000072274 |
| 9 | P17655 | NA | CAPN2 | calpain 2, (m/II) large subunit | 824 | ENSG00000162909 |
| 10 | P04040 | NA | CAT | catalase | 847 | ENSG00000121691 |
| 11 | P22570 | NA | FDXR | ferredoxin reductase | 2232 | ENSG00000161513 |
| 12 | P05556 | NA | ITGB1 | integrin, beta 1 (fibronectin receptor, beta polypeptide, antigen CD29 includes MDF2, MSK12) | 3688 | ENSG00000150093 |
| 13 | P08727 | NA | KRT19 | keratin 19 | 3880 | ENSG00000171345 |
| 14 | P07339 | NA | CTSD | cathepsin D | 1509 | ENSG00000117984 |
| 15 | P18206 | NA | VCL | vinculin | 7414 | ENSG00000035403 |
| 16 | P21589 | NA | NT5E | 5'-nucleotidase, ecto (CD73) | 4907 | ENSG00000135318 |

  
  

| **Database:Pathway Commons pathway      &nbspName:Signaling events mediated by VEGFR1 and VEGFR2      &nbspID:DB\_ID:1516** | | | | | | |
| --- | --- | --- | --- | --- | --- | --- |
| C=1296; O=16; E=5.50; R=2.91; rawP=9.68e-05; adjP=0.0002 | | | | | | |
| Index | UserID | Value | Gene Symbol | Gene Name | EntrezGene | Ensembl |
| 1 | P08195 | NA | SLC3A2 | solute carrier family 3 (activators of dibasic and neutral amino acid transport), member 2 | 6520 | ENSG00000168003 |
| 2 | P30086 | NA | PEBP1 | phosphatidylethanolamine binding protein 1 | 5037 | ENSG00000089220 |
| 3 | P00338 | NA | LDHA | lactate dehydrogenase A | 3939 | ENSG00000134333 |
| 4 | P55060 | NA | CSE1L | CSE1 chromosome segregation 1-like (yeast) | 1434 | ENSG00000124207 |
| 5 | P16949 | NA | STMN1 | stathmin 1 | 3925 | ENSG00000117632 |
| 6 | O60610 | NA | DIAPH1 | diaphanous homolog 1 (Drosophila) | 1729 | ENSG00000131504 |
| 7 | Q06830 | NA | PRDX1 | peroxiredoxin 1 | 5052 | ENSG00000117450 |
| 8 | P02786 | NA | TFRC | transferrin receptor (p90, CD71) | 7037 | ENSG00000072274 |
| 9 | P17655 | NA | CAPN2 | calpain 2, (m/II) large subunit | 824 | ENSG00000162909 |
| 10 | P04040 | NA | CAT | catalase | 847 | ENSG00000121691 |
| 11 | P22570 | NA | FDXR | ferredoxin reductase | 2232 | ENSG00000161513 |
| 12 | P05556 | NA | ITGB1 | integrin, beta 1 (fibronectin receptor, beta polypeptide, antigen CD29 includes MDF2, MSK12) | 3688 | ENSG00000150093 |
| 13 | P08727 | NA | KRT19 | keratin 19 | 3880 | ENSG00000171345 |
| 14 | P07339 | NA | CTSD | cathepsin D | 1509 | ENSG00000117984 |
| 15 | P18206 | NA | VCL | vinculin | 7414 | ENSG00000035403 |
| 16 | P21589 | NA | NT5E | 5'-nucleotidase, ecto (CD73) | 4907 | ENSG00000135318 |

  
  

| **Database:Pathway Commons pathway      &nbspName:Plasma membrane estrogen receptor signaling      &nbspID:DB\_ID:1556** | | | | | | |
| --- | --- | --- | --- | --- | --- | --- |
| C=1301; O=16; E=5.52; R=2.90; rawP=0.0001; adjP=0.0002 | | | | | | |
| Index | UserID | Value | Gene Symbol | Gene Name | EntrezGene | Ensembl |
| 1 | P08195 | NA | SLC3A2 | solute carrier family 3 (activators of dibasic and neutral amino acid transport), member 2 | 6520 | ENSG00000168003 |
| 2 | P30086 | NA | PEBP1 | phosphatidylethanolamine binding protein 1 | 5037 | ENSG00000089220 |
| 3 | P00338 | NA | LDHA | lactate dehydrogenase A | 3939 | ENSG00000134333 |
| 4 | P55060 | NA | CSE1L | CSE1 chromosome segregation 1-like (yeast) | 1434 | ENSG00000124207 |
| 5 | P16949 | NA | STMN1 | stathmin 1 | 3925 | ENSG00000117632 |
| 6 | O60610 | NA | DIAPH1 | diaphanous homolog 1 (Drosophila) | 1729 | ENSG00000131504 |
| 7 | Q06830 | NA | PRDX1 | peroxiredoxin 1 | 5052 | ENSG00000117450 |
| 8 | P02786 | NA | TFRC | transferrin receptor (p90, CD71) | 7037 | ENSG00000072274 |
| 9 | P17655 | NA | CAPN2 | calpain 2, (m/II) large subunit | 824 | ENSG00000162909 |
| 10 | P04040 | NA | CAT | catalase | 847 | ENSG00000121691 |
| 11 | P22570 | NA | FDXR | ferredoxin reductase | 2232 | ENSG00000161513 |
| 12 | P05556 | NA | ITGB1 | integrin, beta 1 (fibronectin receptor, beta polypeptide, antigen CD29 includes MDF2, MSK12) | 3688 | ENSG00000150093 |
| 13 | P08727 | NA | KRT19 | keratin 19 | 3880 | ENSG00000171345 |
| 14 | P07339 | NA | CTSD | cathepsin D | 1509 | ENSG00000117984 |
| 15 | P18206 | NA | VCL | vinculin | 7414 | ENSG00000035403 |
| 16 | P21589 | NA | NT5E | 5'-nucleotidase, ecto (CD73) | 4907 | ENSG00000135318 |

  
  

| **Database:Pathway Commons pathway      &nbspName:S1P1 pathway      &nbspID:DB\_ID:1594** | | | | | | |
| --- | --- | --- | --- | --- | --- | --- |
| C=1288; O=16; E=5.46; R=2.93; rawP=9.00e-05; adjP=0.0002 | | | | | | |
| Index | UserID | Value | Gene Symbol | Gene Name | EntrezGene | Ensembl |
| 1 | P08195 | NA | SLC3A2 | solute carrier family 3 (activators of dibasic and neutral amino acid transport), member 2 | 6520 | ENSG00000168003 |
| 2 | P30086 | NA | PEBP1 | phosphatidylethanolamine binding protein 1 | 5037 | ENSG00000089220 |
| 3 | P00338 | NA | LDHA | lactate dehydrogenase A | 3939 | ENSG00000134333 |
| 4 | P55060 | NA | CSE1L | CSE1 chromosome segregation 1-like (yeast) | 1434 | ENSG00000124207 |
| 5 | P16949 | NA | STMN1 | stathmin 1 | 3925 | ENSG00000117632 |
| 6 | O60610 | NA | DIAPH1 | diaphanous homolog 1 (Drosophila) | 1729 | ENSG00000131504 |
| 7 | Q06830 | NA | PRDX1 | peroxiredoxin 1 | 5052 | ENSG00000117450 |
| 8 | P02786 | NA | TFRC | transferrin receptor (p90, CD71) | 7037 | ENSG00000072274 |
| 9 | P17655 | NA | CAPN2 | calpain 2, (m/II) large subunit | 824 | ENSG00000162909 |
| 10 | P04040 | NA | CAT | catalase | 847 | ENSG00000121691 |
| 11 | P22570 | NA | FDXR | ferredoxin reductase | 2232 | ENSG00000161513 |
| 12 | P05556 | NA | ITGB1 | integrin, beta 1 (fibronectin receptor, beta polypeptide, antigen CD29 includes MDF2, MSK12) | 3688 | ENSG00000150093 |
| 13 | P08727 | NA | KRT19 | keratin 19 | 3880 | ENSG00000171345 |
| 14 | P07339 | NA | CTSD | cathepsin D | 1509 | ENSG00000117984 |
| 15 | P18206 | NA | VCL | vinculin | 7414 | ENSG00000035403 |
| 16 | P21589 | NA | NT5E | 5'-nucleotidase, ecto (CD73) | 4907 | ENSG00000135318 |

  
  

| **Database:Pathway Commons pathway      &nbspName:Purine catabolism      &nbspID:DB\_ID:555** | | | | | | |
| --- | --- | --- | --- | --- | --- | --- |
| C=11; O=3; E=0.05; R=64.32; rawP=1.18e-05; adjP=0.0002 | | | | | | |
| Index | UserID | Value | Gene Symbol | Gene Name | EntrezGene | Ensembl |
| 1 | P04040 | NA | CAT | catalase | 847 | ENSG00000121691 |
| 2 | P00491 | NA | PNP | purine nucleoside phosphorylase | 4860 | ENSG00000198805 |
| 3 | P21589 | NA | NT5E | 5'-nucleotidase, ecto (CD73) | 4907 | ENSG00000135318 |

  
  

| **Database:Pathway Commons pathway      &nbspName:Class I PI3K signaling events      &nbspID:DB\_ID:1553** | | | | | | |
| --- | --- | --- | --- | --- | --- | --- |
| C=1288; O=16; E=5.46; R=2.93; rawP=9.00e-05; adjP=0.0002 | | | | | | |
| Index | UserID | Value | Gene Symbol | Gene Name | EntrezGene | Ensembl |
| 1 | P08195 | NA | SLC3A2 | solute carrier family 3 (activators of dibasic and neutral amino acid transport), member 2 | 6520 | ENSG00000168003 |
| 2 | P30086 | NA | PEBP1 | phosphatidylethanolamine binding protein 1 | 5037 | ENSG00000089220 |
| 3 | P00338 | NA | LDHA | lactate dehydrogenase A | 3939 | ENSG00000134333 |
| 4 | P55060 | NA | CSE1L | CSE1 chromosome segregation 1-like (yeast) | 1434 | ENSG00000124207 |
| 5 | P16949 | NA | STMN1 | stathmin 1 | 3925 | ENSG00000117632 |
| 6 | O60610 | NA | DIAPH1 | diaphanous homolog 1 (Drosophila) | 1729 | ENSG00000131504 |
| 7 | Q06830 | NA | PRDX1 | peroxiredoxin 1 | 5052 | ENSG00000117450 |
| 8 | P02786 | NA | TFRC | transferrin receptor (p90, CD71) | 7037 | ENSG00000072274 |
| 9 | P17655 | NA | CAPN2 | calpain 2, (m/II) large subunit | 824 | ENSG00000162909 |
| 10 | P04040 | NA | CAT | catalase | 847 | ENSG00000121691 |
| 11 | P22570 | NA | FDXR | ferredoxin reductase | 2232 | ENSG00000161513 |
| 12 | P05556 | NA | ITGB1 | integrin, beta 1 (fibronectin receptor, beta polypeptide, antigen CD29 includes MDF2, MSK12) | 3688 | ENSG00000150093 |
| 13 | P08727 | NA | KRT19 | keratin 19 | 3880 | ENSG00000171345 |
| 14 | P07339 | NA | CTSD | cathepsin D | 1509 | ENSG00000117984 |
| 15 | P18206 | NA | VCL | vinculin | 7414 | ENSG00000035403 |
| 16 | P21589 | NA | NT5E | 5'-nucleotidase, ecto (CD73) | 4907 | ENSG00000135318 |

  
  

| **Database:Pathway Commons pathway      &nbspName:Urokinase-type plasminogen activator (uPA) and uPAR-mediated signaling      &nbspID:DB\_ID:1519** | | | | | | |
| --- | --- | --- | --- | --- | --- | --- |
| C=1288; O=16; E=5.46; R=2.93; rawP=9.00e-05; adjP=0.0002 | | | | | | |
| Index | UserID | Value | Gene Symbol | Gene Name | EntrezGene | Ensembl |
| 1 | P08195 | NA | SLC3A2 | solute carrier family 3 (activators of dibasic and neutral amino acid transport), member 2 | 6520 | ENSG00000168003 |
| 2 | P30086 | NA | PEBP1 | phosphatidylethanolamine binding protein 1 | 5037 | ENSG00000089220 |
| 3 | P00338 | NA | LDHA | lactate dehydrogenase A | 3939 | ENSG00000134333 |
| 4 | P55060 | NA | CSE1L | CSE1 chromosome segregation 1-like (yeast) | 1434 | ENSG00000124207 |
| 5 | P16949 | NA | STMN1 | stathmin 1 | 3925 | ENSG00000117632 |
| 6 | O60610 | NA | DIAPH1 | diaphanous homolog 1 (Drosophila) | 1729 | ENSG00000131504 |
| 7 | Q06830 | NA | PRDX1 | peroxiredoxin 1 | 5052 | ENSG00000117450 |
| 8 | P02786 | NA | TFRC | transferrin receptor (p90, CD71) | 7037 | ENSG00000072274 |
| 9 | P17655 | NA | CAPN2 | calpain 2, (m/II) large subunit | 824 | ENSG00000162909 |
| 10 | P04040 | NA | CAT | catalase | 847 | ENSG00000121691 |
| 11 | P22570 | NA | FDXR | ferredoxin reductase | 2232 | ENSG00000161513 |
| 12 | P05556 | NA | ITGB1 | integrin, beta 1 (fibronectin receptor, beta polypeptide, antigen CD29 includes MDF2, MSK12) | 3688 | ENSG00000150093 |
| 13 | P08727 | NA | KRT19 | keratin 19 | 3880 | ENSG00000171345 |
| 14 | P07339 | NA | CTSD | cathepsin D | 1509 | ENSG00000117984 |
| 15 | P18206 | NA | VCL | vinculin | 7414 | ENSG00000035403 |
| 16 | P21589 | NA | NT5E | 5'-nucleotidase, ecto (CD73) | 4907 | ENSG00000135318 |

  
  

| **Database:Pathway Commons pathway      &nbspName:Glypican 1 network      &nbspID:DB\_ID:1492** | | | | | | |
| --- | --- | --- | --- | --- | --- | --- |
| C=1299; O=16; E=5.51; R=2.90; rawP=9.95e-05; adjP=0.0002 | | | | | | |
| Index | UserID | Value | Gene Symbol | Gene Name | EntrezGene | Ensembl |
| 1 | P08195 | NA | SLC3A2 | solute carrier family 3 (activators of dibasic and neutral amino acid transport), member 2 | 6520 | ENSG00000168003 |
| 2 | P30086 | NA | PEBP1 | phosphatidylethanolamine binding protein 1 | 5037 | ENSG00000089220 |
| 3 | P00338 | NA | LDHA | lactate dehydrogenase A | 3939 | ENSG00000134333 |
| 4 | P55060 | NA | CSE1L | CSE1 chromosome segregation 1-like (yeast) | 1434 | ENSG00000124207 |
| 5 | P16949 | NA | STMN1 | stathmin 1 | 3925 | ENSG00000117632 |
| 6 | O60610 | NA | DIAPH1 | diaphanous homolog 1 (Drosophila) | 1729 | ENSG00000131504 |
| 7 | Q06830 | NA | PRDX1 | peroxiredoxin 1 | 5052 | ENSG00000117450 |
| 8 | P02786 | NA | TFRC | transferrin receptor (p90, CD71) | 7037 | ENSG00000072274 |
| 9 | P17655 | NA | CAPN2 | calpain 2, (m/II) large subunit | 824 | ENSG00000162909 |
| 10 | P04040 | NA | CAT | catalase | 847 | ENSG00000121691 |
| 11 | P22570 | NA | FDXR | ferredoxin reductase | 2232 | ENSG00000161513 |
| 12 | P05556 | NA | ITGB1 | integrin, beta 1 (fibronectin receptor, beta polypeptide, antigen CD29 includes MDF2, MSK12) | 3688 | ENSG00000150093 |
| 13 | P08727 | NA | KRT19 | keratin 19 | 3880 | ENSG00000171345 |
| 14 | P07339 | NA | CTSD | cathepsin D | 1509 | ENSG00000117984 |
| 15 | P18206 | NA | VCL | vinculin | 7414 | ENSG00000035403 |
| 16 | P21589 | NA | NT5E | 5'-nucleotidase, ecto (CD73) | 4907 | ENSG00000135318 |

  
  

| **Database:Pathway Commons pathway      &nbspName:ErbB1 downstream signaling      &nbspID:DB\_ID:1602** | | | | | | |
| --- | --- | --- | --- | --- | --- | --- |
| C=1288; O=16; E=5.46; R=2.93; rawP=9.00e-05; adjP=0.0002 | | | | | | |
| Index | UserID | Value | Gene Symbol | Gene Name | EntrezGene | Ensembl |
| 1 | P08195 | NA | SLC3A2 | solute carrier family 3 (activators of dibasic and neutral amino acid transport), member 2 | 6520 | ENSG00000168003 |
| 2 | P30086 | NA | PEBP1 | phosphatidylethanolamine binding protein 1 | 5037 | ENSG00000089220 |
| 3 | P00338 | NA | LDHA | lactate dehydrogenase A | 3939 | ENSG00000134333 |
| 4 | P55060 | NA | CSE1L | CSE1 chromosome segregation 1-like (yeast) | 1434 | ENSG00000124207 |
| 5 | P16949 | NA | STMN1 | stathmin 1 | 3925 | ENSG00000117632 |
| 6 | O60610 | NA | DIAPH1 | diaphanous homolog 1 (Drosophila) | 1729 | ENSG00000131504 |
| 7 | Q06830 | NA | PRDX1 | peroxiredoxin 1 | 5052 | ENSG00000117450 |
| 8 | P02786 | NA | TFRC | transferrin receptor (p90, CD71) | 7037 | ENSG00000072274 |
| 9 | P17655 | NA | CAPN2 | calpain 2, (m/II) large subunit | 824 | ENSG00000162909 |
| 10 | P04040 | NA | CAT | catalase | 847 | ENSG00000121691 |
| 11 | P22570 | NA | FDXR | ferredoxin reductase | 2232 | ENSG00000161513 |
| 12 | P05556 | NA | ITGB1 | integrin, beta 1 (fibronectin receptor, beta polypeptide, antigen CD29 includes MDF2, MSK12) | 3688 | ENSG00000150093 |
| 13 | P08727 | NA | KRT19 | keratin 19 | 3880 | ENSG00000171345 |
| 14 | P07339 | NA | CTSD | cathepsin D | 1509 | ENSG00000117984 |
| 15 | P18206 | NA | VCL | vinculin | 7414 | ENSG00000035403 |
| 16 | P21589 | NA | NT5E | 5'-nucleotidase, ecto (CD73) | 4907 | ENSG00000135318 |

  
  

| **Database:Pathway Commons pathway      &nbspName:EGF receptor (ErbB1) signaling pathway      &nbspID:DB\_ID:1550** | | | | | | |
| --- | --- | --- | --- | --- | --- | --- |
| C=1288; O=16; E=5.46; R=2.93; rawP=9.00e-05; adjP=0.0002 | | | | | | |
| Index | UserID | Value | Gene Symbol | Gene Name | EntrezGene | Ensembl |
| 1 | P08195 | NA | SLC3A2 | solute carrier family 3 (activators of dibasic and neutral amino acid transport), member 2 | 6520 | ENSG00000168003 |
| 2 | P30086 | NA | PEBP1 | phosphatidylethanolamine binding protein 1 | 5037 | ENSG00000089220 |
| 3 | P00338 | NA | LDHA | lactate dehydrogenase A | 3939 | ENSG00000134333 |
| 4 | P55060 | NA | CSE1L | CSE1 chromosome segregation 1-like (yeast) | 1434 | ENSG00000124207 |
| 5 | P16949 | NA | STMN1 | stathmin 1 | 3925 | ENSG00000117632 |
| 6 | O60610 | NA | DIAPH1 | diaphanous homolog 1 (Drosophila) | 1729 | ENSG00000131504 |
| 7 | Q06830 | NA | PRDX1 | peroxiredoxin 1 | 5052 | ENSG00000117450 |
| 8 | P02786 | NA | TFRC | transferrin receptor (p90, CD71) | 7037 | ENSG00000072274 |
| 9 | P17655 | NA | CAPN2 | calpain 2, (m/II) large subunit | 824 | ENSG00000162909 |
| 10 | P04040 | NA | CAT | catalase | 847 | ENSG00000121691 |
| 11 | P22570 | NA | FDXR | ferredoxin reductase | 2232 | ENSG00000161513 |
| 12 | P05556 | NA | ITGB1 | integrin, beta 1 (fibronectin receptor, beta polypeptide, antigen CD29 includes MDF2, MSK12) | 3688 | ENSG00000150093 |
| 13 | P08727 | NA | KRT19 | keratin 19 | 3880 | ENSG00000171345 |
| 14 | P07339 | NA | CTSD | cathepsin D | 1509 | ENSG00000117984 |
| 15 | P18206 | NA | VCL | vinculin | 7414 | ENSG00000035403 |
| 16 | P21589 | NA | NT5E | 5'-nucleotidase, ecto (CD73) | 4907 | ENSG00000135318 |

  
  

| **Database:Pathway Commons pathway      &nbspName:Arf6 trafficking events      &nbspID:DB\_ID:1615** | | | | | | |
| --- | --- | --- | --- | --- | --- | --- |
| C=1288; O=16; E=5.46; R=2.93; rawP=9.00e-05; adjP=0.0002 | | | | | | |
| Index | UserID | Value | Gene Symbol | Gene Name | EntrezGene | Ensembl |
| 1 | P08195 | NA | SLC3A2 | solute carrier family 3 (activators of dibasic and neutral amino acid transport), member 2 | 6520 | ENSG00000168003 |
| 2 | P30086 | NA | PEBP1 | phosphatidylethanolamine binding protein 1 | 5037 | ENSG00000089220 |
| 3 | P00338 | NA | LDHA | lactate dehydrogenase A | 3939 | ENSG00000134333 |
| 4 | P55060 | NA | CSE1L | CSE1 chromosome segregation 1-like (yeast) | 1434 | ENSG00000124207 |
| 5 | P16949 | NA | STMN1 | stathmin 1 | 3925 | ENSG00000117632 |
| 6 | O60610 | NA | DIAPH1 | diaphanous homolog 1 (Drosophila) | 1729 | ENSG00000131504 |
| 7 | Q06830 | NA | PRDX1 | peroxiredoxin 1 | 5052 | ENSG00000117450 |
| 8 | P02786 | NA | TFRC | transferrin receptor (p90, CD71) | 7037 | ENSG00000072274 |
| 9 | P17655 | NA | CAPN2 | calpain 2, (m/II) large subunit | 824 | ENSG00000162909 |
| 10 | P04040 | NA | CAT | catalase | 847 | ENSG00000121691 |
| 11 | P22570 | NA | FDXR | ferredoxin reductase | 2232 | ENSG00000161513 |
| 12 | P05556 | NA | ITGB1 | integrin, beta 1 (fibronectin receptor, beta polypeptide, antigen CD29 includes MDF2, MSK12) | 3688 | ENSG00000150093 |
| 13 | P08727 | NA | KRT19 | keratin 19 | 3880 | ENSG00000171345 |
| 14 | P07339 | NA | CTSD | cathepsin D | 1509 | ENSG00000117984 |
| 15 | P18206 | NA | VCL | vinculin | 7414 | ENSG00000035403 |
| 16 | P21589 | NA | NT5E | 5'-nucleotidase, ecto (CD73) | 4907 | ENSG00000135318 |

  
  

| **Database:Pathway Commons pathway      &nbspName:tRNA Aminoacylation      &nbspID:DB\_ID:519** | | | | | | |
| --- | --- | --- | --- | --- | --- | --- |
| C=42; O=4; E=0.18; R=22.46; rawP=2.98e-05; adjP=0.0002 | | | | | | |
| Index | UserID | Value | Gene Symbol | Gene Name | EntrezGene | Ensembl |
| 1 | P56192 | NA | MARS | methionyl-tRNA synthetase | 4141 | ENSG00000166986 |
| 2 | P49591 | NA | SARS | seryl-tRNA synthetase | 6301 | ENSG00000031698 |
| 3 | P49588 | NA | AARS | alanyl-tRNA synthetase | 16 | ENSG00000090861 |
| 4 | P23381 | NA | WARS | tryptophanyl-tRNA synthetase | 7453 | ENSG00000140105 |

  
  

| **Database:Pathway Commons pathway      &nbspName:Integrin family cell surface interactions      &nbspID:DB\_ID:1499** | | | | | | |
| --- | --- | --- | --- | --- | --- | --- |
| C=1378; O=16; E=5.84; R=2.74; rawP=0.0002; adjP=0.0005 | | | | | | |
| Index | UserID | Value | Gene Symbol | Gene Name | EntrezGene | Ensembl |
| 1 | P08195 | NA | SLC3A2 | solute carrier family 3 (activators of dibasic and neutral amino acid transport), member 2 | 6520 | ENSG00000168003 |
| 2 | P30086 | NA | PEBP1 | phosphatidylethanolamine binding protein 1 | 5037 | ENSG00000089220 |
| 3 | P00338 | NA | LDHA | lactate dehydrogenase A | 3939 | ENSG00000134333 |
| 4 | P55060 | NA | CSE1L | CSE1 chromosome segregation 1-like (yeast) | 1434 | ENSG00000124207 |
| 5 | P16949 | NA | STMN1 | stathmin 1 | 3925 | ENSG00000117632 |
| 6 | O60610 | NA | DIAPH1 | diaphanous homolog 1 (Drosophila) | 1729 | ENSG00000131504 |
| 7 | Q06830 | NA | PRDX1 | peroxiredoxin 1 | 5052 | ENSG00000117450 |
| 8 | P02786 | NA | TFRC | transferrin receptor (p90, CD71) | 7037 | ENSG00000072274 |
| 9 | P17655 | NA | CAPN2 | calpain 2, (m/II) large subunit | 824 | ENSG00000162909 |
| 10 | P04040 | NA | CAT | catalase | 847 | ENSG00000121691 |
| 11 | P22570 | NA | FDXR | ferredoxin reductase | 2232 | ENSG00000161513 |
| 12 | P05556 | NA | ITGB1 | integrin, beta 1 (fibronectin receptor, beta polypeptide, antigen CD29 includes MDF2, MSK12) | 3688 | ENSG00000150093 |
| 13 | P08727 | NA | KRT19 | keratin 19 | 3880 | ENSG00000171345 |
| 14 | P07339 | NA | CTSD | cathepsin D | 1509 | ENSG00000117984 |
| 15 | P18206 | NA | VCL | vinculin | 7414 | ENSG00000035403 |
| 16 | P21589 | NA | NT5E | 5'-nucleotidase, ecto (CD73) | 4907 | ENSG00000135318 |

  
  

| **Database:Pathway Commons pathway      &nbspName:Beta1 integrin cell surface interactions      &nbspID:DB\_ID:1517** | | | | | | |
| --- | --- | --- | --- | --- | --- | --- |
| C=1351; O=16; E=5.73; R=2.79; rawP=0.0002; adjP=0.0005 | | | | | | |
| Index | UserID | Value | Gene Symbol | Gene Name | EntrezGene | Ensembl |
| 1 | P08195 | NA | SLC3A2 | solute carrier family 3 (activators of dibasic and neutral amino acid transport), member 2 | 6520 | ENSG00000168003 |
| 2 | P30086 | NA | PEBP1 | phosphatidylethanolamine binding protein 1 | 5037 | ENSG00000089220 |
| 3 | P00338 | NA | LDHA | lactate dehydrogenase A | 3939 | ENSG00000134333 |
| 4 | P55060 | NA | CSE1L | CSE1 chromosome segregation 1-like (yeast) | 1434 | ENSG00000124207 |
| 5 | P16949 | NA | STMN1 | stathmin 1 | 3925 | ENSG00000117632 |
| 6 | O60610 | NA | DIAPH1 | diaphanous homolog 1 (Drosophila) | 1729 | ENSG00000131504 |
| 7 | Q06830 | NA | PRDX1 | peroxiredoxin 1 | 5052 | ENSG00000117450 |
| 8 | P02786 | NA | TFRC | transferrin receptor (p90, CD71) | 7037 | ENSG00000072274 |
| 9 | P17655 | NA | CAPN2 | calpain 2, (m/II) large subunit | 824 | ENSG00000162909 |
| 10 | P04040 | NA | CAT | catalase | 847 | ENSG00000121691 |
| 11 | P22570 | NA | FDXR | ferredoxin reductase | 2232 | ENSG00000161513 |
| 12 | P05556 | NA | ITGB1 | integrin, beta 1 (fibronectin receptor, beta polypeptide, antigen CD29 includes MDF2, MSK12) | 3688 | ENSG00000150093 |
| 13 | P08727 | NA | KRT19 | keratin 19 | 3880 | ENSG00000171345 |
| 14 | P07339 | NA | CTSD | cathepsin D | 1509 | ENSG00000117984 |
| 15 | P18206 | NA | VCL | vinculin | 7414 | ENSG00000035403 |
| 16 | P21589 | NA | NT5E | 5'-nucleotidase, ecto (CD73) | 4907 | ENSG00000135318 |

  
  

| **Database:Pathway Commons pathway      &nbspName:Gene Expression      &nbspID:DB\_ID:531** | | | | | | |
| --- | --- | --- | --- | --- | --- | --- |
| C=379; O=8; E=1.61; R=4.98; rawP=0.0002; adjP=0.0005 | | | | | | |
| Index | UserID | Value | Gene Symbol | Gene Name | EntrezGene | Ensembl |
| 1 | P49591 | NA | SARS | seryl-tRNA synthetase | 6301 | ENSG00000031698 |
| 2 | P20042 | NA | EIF2S2 | eukaryotic translation initiation factor 2, subunit 2 beta, 38kDa | 8894 | ENSG00000125977 |
| 3 | P41091 | NA | EIF2S3 | eukaryotic translation initiation factor 2, subunit 3 gamma, 52kDa | 1968 | ENSG00000130741 |
| 4 | P56192 | NA | MARS | methionyl-tRNA synthetase | 4141 | ENSG00000166986 |
| 5 | Q15056 | NA | EIF4H | eukaryotic translation initiation factor 4H | 7458 | ENSG00000106682 |
| 6 | P49588 | NA | AARS | alanyl-tRNA synthetase | 16 | ENSG00000090861 |
| 7 | P68104 | NA | EEF1A1 | eukaryotic translation elongation factor 1 alpha 1 | 1915 | ENSG00000156508 |
| 8 | P23381 | NA | WARS | tryptophanyl-tRNA synthetase | 7453 | ENSG00000140105 |

  
  

| **Database:Pathway Commons pathway      &nbspName:CDC42 signaling events      &nbspID:DB\_ID:1488** | | | | | | |
| --- | --- | --- | --- | --- | --- | --- |
| C=757; O=11; E=3.21; R=3.43; rawP=0.0003; adjP=0.0007 | | | | | | |
| Index | UserID | Value | Gene Symbol | Gene Name | EntrezGene | Ensembl |
| 1 | P08195 | NA | SLC3A2 | solute carrier family 3 (activators of dibasic and neutral amino acid transport), member 2 | 6520 | ENSG00000168003 |
| 2 | P00338 | NA | LDHA | lactate dehydrogenase A | 3939 | ENSG00000134333 |
| 3 | P05556 | NA | ITGB1 | integrin, beta 1 (fibronectin receptor, beta polypeptide, antigen CD29 includes MDF2, MSK12) | 3688 | ENSG00000150093 |
| 4 | P08727 | NA | KRT19 | keratin 19 | 3880 | ENSG00000171345 |
| 5 | Q15019 | NA | SEPT2 | septin 2 | 4735 | ENSG00000168385 |
| 6 | P16949 | NA | STMN1 | stathmin 1 | 3925 | ENSG00000117632 |
| 7 | P18206 | NA | VCL | vinculin | 7414 | ENSG00000035403 |
| 8 | O60610 | NA | DIAPH1 | diaphanous homolog 1 (Drosophila) | 1729 | ENSG00000131504 |
| 9 | Q06830 | NA | PRDX1 | peroxiredoxin 1 | 5052 | ENSG00000117450 |
| 10 | P02786 | NA | TFRC | transferrin receptor (p90, CD71) | 7037 | ENSG00000072274 |
| 11 | P21589 | NA | NT5E | 5'-nucleotidase, ecto (CD73) | 4907 | ENSG00000135318 |

  
  

| **Database:Pathway Commons pathway      &nbspName:Regulation of CDC42 activity      &nbspID:DB\_ID:1456** | | | | | | |
| --- | --- | --- | --- | --- | --- | --- |
| C=770; O=11; E=3.26; R=3.37; rawP=0.0004; adjP=0.0009 | | | | | | |
| Index | UserID | Value | Gene Symbol | Gene Name | EntrezGene | Ensembl |
| 1 | P08195 | NA | SLC3A2 | solute carrier family 3 (activators of dibasic and neutral amino acid transport), member 2 | 6520 | ENSG00000168003 |
| 2 | P00338 | NA | LDHA | lactate dehydrogenase A | 3939 | ENSG00000134333 |
| 3 | P05556 | NA | ITGB1 | integrin, beta 1 (fibronectin receptor, beta polypeptide, antigen CD29 includes MDF2, MSK12) | 3688 | ENSG00000150093 |
| 4 | P08727 | NA | KRT19 | keratin 19 | 3880 | ENSG00000171345 |
| 5 | Q15019 | NA | SEPT2 | septin 2 | 4735 | ENSG00000168385 |
| 6 | P16949 | NA | STMN1 | stathmin 1 | 3925 | ENSG00000117632 |
| 7 | P18206 | NA | VCL | vinculin | 7414 | ENSG00000035403 |
| 8 | O60610 | NA | DIAPH1 | diaphanous homolog 1 (Drosophila) | 1729 | ENSG00000131504 |
| 9 | Q06830 | NA | PRDX1 | peroxiredoxin 1 | 5052 | ENSG00000117450 |
| 10 | P02786 | NA | TFRC | transferrin receptor (p90, CD71) | 7037 | ENSG00000072274 |
| 11 | P21589 | NA | NT5E | 5'-nucleotidase, ecto (CD73) | 4907 | ENSG00000135318 |

  
  

| **Database:Pathway Commons pathway      &nbspName:Fatty acid, triacylglycerol, and ketone body metabolism      &nbspID:DB\_ID:919** | | | | | | |
| --- | --- | --- | --- | --- | --- | --- |
| C=83; O=4; E=0.35; R=11.37; rawP=0.0004; adjP=0.0009 | | | | | | |
| Index | UserID | Value | Gene Symbol | Gene Name | EntrezGene | Ensembl |
| 1 | P24752 | NA | ACAT1 | acetyl-CoA acetyltransferase 1 | 38 | ENSG00000075239 |
| 2 | Q16881 | NA | TXNRD1 | thioredoxin reductase 1 | 7296 | ENSG00000198431 |
| 3 | P49748 | NA | ACADVL | acyl-CoA dehydrogenase, very long chain | 37 | ENSG00000072778 |
| 4 | P53007 | NA | SLC25A1 | solute carrier family 25 (mitochondrial carrier; citrate transporter), member 1 | 6576 | ENSG00000100075 |

  
  
  
  

---

WebGestalt is currently developed and maintained by Jing Wang and Bing Zhang at the  Zhang Lab. Other people who have made significant contribution to the project include Dexter Duncan, Stefan Kirov, Zhiao Shi, and Jay Snoddy.  
  
**Funding credits:** NIH/NIAAA (U01 AA016662, U01 AA013512); NIH/NIDA (P01 DA015027); NIH/NIMH (P50 MH078028, P50 MH096972); NIH/NCI (U24 CA159988); NIH/NIGMS (R01 GM088822).
