## Supplementary material for "PGRMC1 phosphorylation and cell plasticity 1: glycolysis, mitochondria, tumor growth": File S7: final_ppi_sig_file_1438487532.html

Anchored HTML File of EIDs

|  |  |
| --- | --- |
|  | WEB-based GEne SeT AnaLysis Toolkit |
|  | ***Translating gene lists into biological insights...*** |

---

  

| **Database:PPI      &nbspName:Hsapiens\_Module\_493      &nbspID:DB\_ID:493** **Related Function \|**  **Network View** | | | | | | |
| --- | --- | --- | --- | --- | --- | --- |
| C=277; O=10; E=1.50; R=6.68; rawP=2.40e-06; adjP=5.52e-05 | | | | | | |
| Index | UserID | Value | Gene Symbol | Gene Name | EntrezGene | Ensembl |
| 1 | Q04446 | NA | GBE1 | glucan (1,4-alpha-), branching enzyme 1 | 2632 | ENSG00000114480 |
| 2 | P22102 | NA | GART | phosphoribosylglycinamide formyltransferase, phosphoribosylglycinamide synthetase, phosphoribosylaminoimidazole synthetase | 2618 | ENSG00000159131 |
| 3 | P00491 | NA | PNP | purine nucleoside phosphorylase | 4860 | ENSG00000198805 |
| 4 | P55060 | NA | CSE1L | CSE1 chromosome segregation 1-like (yeast) | 1434 | ENSG00000124207 |
| 5 | P00367 | NA | GLUD1 | glutamate dehydrogenase 1 | 2746 | ENSG00000148672 |
| 6 | P06744 | NA | GPI | glucose-6-phosphate isomerase | 2821 | ENSG00000105220 |
| 7 | P37802 | NA | TAGLN2 | transgelin 2 | 8407 | ENSG00000158710 |
| 8 | Q14764 | NA | MVP | major vault protein | 9961 | ENSG00000013364 |
| 9 | Q14697 | NA | GANAB | glucosidase, alpha; neutral AB | 23193 | ENSG00000089597 |
| 10 | Q13011 | NA | ECH1 | enoyl CoA hydratase 1, peroxisomal | 1891 | ENSG00000104823 |

  
  

| **Database:PPI      &nbspName:Hsapiens\_Module\_414      &nbspID:DB\_ID:414** **Related Function \|**  **Network View** | | | | | | |
| --- | --- | --- | --- | --- | --- | --- |
| C=362; O=10; E=1.96; R=5.11; rawP=2.50e-05; adjP=0.0002 | | | | | | |
| Index | UserID | Value | Gene Symbol | Gene Name | EntrezGene | Ensembl |
| 1 | Q14980 | NA | NUMA1 | nuclear mitotic apparatus protein 1 | 4926 | ENSG00000137497 |
| 2 | P08133 | NA | ANXA6 | annexin A6 | 309 | ENSG00000197043 |
| 3 | P17931 | NA | LGALS3 | lectin, galactoside-binding, soluble, 3 | 3958 | ENSG00000131981 |
| 4 | P42765 | NA | ACAA2 | acetyl-CoA acyltransferase 2 | 10449 | ENSG00000167315 |
| 5 | P49748 | NA | ACADVL | acyl-CoA dehydrogenase, very long chain | 37 | ENSG00000072778 |
| 6 | P16949 | NA | STMN1 | stathmin 1 | 3925 | ENSG00000117632 |
| 7 | P09211 | NA | GSTP1 | glutathione S-transferase pi 1 | 2950 | ENSG00000084207 |
| 8 | Q06830 | NA | PRDX1 | peroxiredoxin 1 | 5052 | ENSG00000117450 |
| 9 | O75369 | NA | FLNB | filamin B, beta | 2317 | ENSG00000136068 |
| 10 | Q9BPW8 | NA | NIPSNAP1 | nipsnap homolog 1 (C. elegans) | 8508 | ENSG00000184117 |

  
  

| **Database:PPI      &nbspName:Hsapiens\_Module\_808      &nbspID:DB\_ID:808** **Related Function \|**  **Network View** | | | | | | |
| --- | --- | --- | --- | --- | --- | --- |
| C=67; O=5; E=0.36; R=13.80; rawP=3.05e-05; adjP=0.0002 | | | | | | |
| Index | UserID | Value | Gene Symbol | Gene Name | EntrezGene | Ensembl |
| 1 | P00367 | NA | GLUD1 | glutamate dehydrogenase 1 | 2746 | ENSG00000148672 |
| 2 | P37802 | NA | TAGLN2 | transgelin 2 | 8407 | ENSG00000158710 |
| 3 | P22102 | NA | GART | phosphoribosylglycinamide formyltransferase, phosphoribosylglycinamide synthetase, phosphoribosylaminoimidazole synthetase | 2618 | ENSG00000159131 |
| 4 | P00491 | NA | PNP | purine nucleoside phosphorylase | 4860 | ENSG00000198805 |
| 5 | Q13011 | NA | ECH1 | enoyl CoA hydratase 1, peroxisomal | 1891 | ENSG00000104823 |

  
  

| **Database:PPI      &nbspName:Hsapiens\_Module\_763      &nbspID:DB\_ID:763** **Related Function \|**  **Network View** | | | | | | |
| --- | --- | --- | --- | --- | --- | --- |
| C=85; O=5; E=0.46; R=10.88; rawP=9.59e-05; adjP=0.0006 | | | | | | |
| Index | UserID | Value | Gene Symbol | Gene Name | EntrezGene | Ensembl |
| 1 | Q14980 | NA | NUMA1 | nuclear mitotic apparatus protein 1 | 4926 | ENSG00000137497 |
| 2 | P08133 | NA | ANXA6 | annexin A6 | 309 | ENSG00000197043 |
| 3 | P49748 | NA | ACADVL | acyl-CoA dehydrogenase, very long chain | 37 | ENSG00000072778 |
| 4 | P16949 | NA | STMN1 | stathmin 1 | 3925 | ENSG00000117632 |
| 5 | Q9BPW8 | NA | NIPSNAP1 | nipsnap homolog 1 (C. elegans) | 8508 | ENSG00000184117 |

  
  

| **Database:PPI      &nbspName:Hsapiens\_Module\_715      &nbspID:DB\_ID:715** **Related Function \|**  **Network View** | | | | | | |
| --- | --- | --- | --- | --- | --- | --- |
| C=20; O=3; E=0.11; R=27.74; rawP=0.0002; adjP=0.0009 | | | | | | |
| Index | UserID | Value | Gene Symbol | Gene Name | EntrezGene | Ensembl |
| 1 | Q92598 | NA | HSPH1 | heat shock 105kDa/110kDa protein 1 | 10808 | ENSG00000120694 |
| 2 | P49588 | NA | AARS | alanyl-tRNA synthetase | 16 | ENSG00000090861 |
| 3 | P00338 | NA | LDHA | lactate dehydrogenase A | 3939 | ENSG00000134333 |

  
  
  
  

---

WebGestalt is currently developed and maintained by Jing Wang and Bing Zhang at the  Zhang Lab. Other people who have made significant contribution to the project include Dexter Duncan, Stefan Kirov, Zhiao Shi, and Jay Snoddy.  
  
**Funding credits:** NIH/NIAAA (U01 AA016662, U01 AA013512); NIH/NIDA (P01 DA015027); NIH/NIMH (P50 MH078028, P50 MH096972); NIH/NCI (U24 CA159988); NIH/NIGMS (R01 GM088822).
