## Supplementary material for "PGRMC1 phosphorylation and cell plasticity 1: glycolysis, mitochondria, tumor growth": File S7: final_wiki_geneset_file_1438487532.html

Anchored HTML File of EIDs

|  |  |
| --- | --- |
|  | WEB-based GEne SeT AnaLysis Toolkit |
|  | ***Translating gene lists into biological insights...*** |

---

  

| **Database:Wikipathways pathway      &nbspName:Leukocyte TarBase      &nbspID:WP2003** | | | | | | |
| --- | --- | --- | --- | --- | --- | --- |
| C=160; O=8; E=0.68; R=11.79; rawP=3.96e-07; adjP=5.54e-06 | | | | | | |
| Index | UserID | Value | Gene Symbol | Gene Name | EntrezGene | Ensembl |
| 1 | Q16881 | NA | TXNRD1 | thioredoxin reductase 1 | 7296 | ENSG00000198431 |
| 2 | Q71U36 | NA | TUBA1A | tubulin, alpha 1a | 7846 | ENSG00000167552 |
| 3 | P00491 | NA | PNP | purine nucleoside phosphorylase | 4860 | ENSG00000198805 |
| 4 | P16615 | NA | ATP2A2 | ATPase, Ca++ transporting, cardiac muscle, slow twitch 2 | 488 | ENSG00000174437 |
| 5 | P40121 | NA | CAPG | capping protein (actin filament), gelsolin-like | 822 | ENSG00000042493 |
| 6 | P42765 | NA | ACAA2 | acetyl-CoA acyltransferase 2 | 10449 | ENSG00000167315 |
| 7 | Q9NQC3 | NA | RTN4 | reticulon 4 | 57142 | ENSG00000115310 |
| 8 | P53007 | NA | SLC25A1 | solute carrier family 25 (mitochondrial carrier; citrate transporter), member 1 | 6576 | ENSG00000100075 |

  
  

| **Database:Wikipathways pathway      &nbspName:Translation Factors      &nbspID:WP107** | | | | | | |
| --- | --- | --- | --- | --- | --- | --- |
| C=51; O=5; E=0.22; R=23.12; rawP=2.45e-06; adjP=1.72e-05 | | | | | | |
| Index | UserID | Value | Gene Symbol | Gene Name | EntrezGene | Ensembl |
| 1 | P20042 | NA | EIF2S2 | eukaryotic translation initiation factor 2, subunit 2 beta, 38kDa | 8894 | ENSG00000125977 |
| 2 | P41091 | NA | EIF2S3 | eukaryotic translation initiation factor 2, subunit 3 gamma, 52kDa | 1968 | ENSG00000130741 |
| 3 | Q15056 | NA | EIF4H | eukaryotic translation initiation factor 4H | 7458 | ENSG00000106682 |
| 4 | P68104 | NA | EEF1A1 | eukaryotic translation elongation factor 1 alpha 1 | 1915 | ENSG00000156508 |
| 5 | Q05639 | NA | EEF1A2 | eukaryotic translation elongation factor 1 alpha 2 | 1917 | ENSG00000101210 |

  
  

| **Database:Wikipathways pathway      &nbspName:Squamous cell TarBase      &nbspID:WP2006** | | | | | | |
| --- | --- | --- | --- | --- | --- | --- |
| C=150; O=6; E=0.64; R=9.43; rawP=4.28e-05; adjP=0.0002 | | | | | | |
| Index | UserID | Value | Gene Symbol | Gene Name | EntrezGene | Ensembl |
| 1 | Q16881 | NA | TXNRD1 | thioredoxin reductase 1 | 7296 | ENSG00000198431 |
| 2 | Q9NQC3 | NA | RTN4 | reticulon 4 | 57142 | ENSG00000115310 |
| 3 | Q71U36 | NA | TUBA1A | tubulin, alpha 1a | 7846 | ENSG00000167552 |
| 4 | P00491 | NA | PNP | purine nucleoside phosphorylase | 4860 | ENSG00000198805 |
| 5 | P16615 | NA | ATP2A2 | ATPase, Ca++ transporting, cardiac muscle, slow twitch 2 | 488 | ENSG00000174437 |
| 6 | P42765 | NA | ACAA2 | acetyl-CoA acyltransferase 2 | 10449 | ENSG00000167315 |

  
  

| **Database:Wikipathways pathway      &nbspName:Selenium Pathway      &nbspID:WP15** | | | | | | |
| --- | --- | --- | --- | --- | --- | --- |
| C=107; O=5; E=0.45; R=11.02; rawP=9.22e-05; adjP=0.0002 | | | | | | |
| Index | UserID | Value | Gene Symbol | Gene Name | EntrezGene | Ensembl |
| 1 | P04040 | NA | CAT | catalase | 847 | ENSG00000121691 |
| 2 | Q16881 | NA | TXNRD1 | thioredoxin reductase 1 | 7296 | ENSG00000198431 |
| 3 | P02768 | NA | ALB | albumin | 213 | ENSG00000163631 |
| 4 | P30044 | NA | PRDX5 | peroxiredoxin 5 | 25824 | ENSG00000126432 |
| 5 | Q06830 | NA | PRDX1 | peroxiredoxin 1 | 5052 | ENSG00000117450 |

  
  

| **Database:Wikipathways pathway      &nbspName:Lymphocyte TarBase      &nbspID:WP2004** | | | | | | |
| --- | --- | --- | --- | --- | --- | --- |
| C=533; O=10; E=2.26; R=4.42; rawP=8.53e-05; adjP=0.0002 | | | | | | |
| Index | UserID | Value | Gene Symbol | Gene Name | EntrezGene | Ensembl |
| 1 | Q16881 | NA | TXNRD1 | thioredoxin reductase 1 | 7296 | ENSG00000198431 |
| 2 | Q71U36 | NA | TUBA1A | tubulin, alpha 1a | 7846 | ENSG00000167552 |
| 3 | P00491 | NA | PNP | purine nucleoside phosphorylase | 4860 | ENSG00000198805 |
| 4 | P16615 | NA | ATP2A2 | ATPase, Ca++ transporting, cardiac muscle, slow twitch 2 | 488 | ENSG00000174437 |
| 5 | P40121 | NA | CAPG | capping protein (actin filament), gelsolin-like | 822 | ENSG00000042493 |
| 6 | P42765 | NA | ACAA2 | acetyl-CoA acyltransferase 2 | 10449 | ENSG00000167315 |
| 7 | Q9NQC3 | NA | RTN4 | reticulon 4 | 57142 | ENSG00000115310 |
| 8 | Q6PIU2 | NA | NCEH1 | neutral cholesterol ester hydrolase 1 | 57552 | ENSG00000144959 |
| 9 | P21589 | NA | NT5E | 5'-nucleotidase, ecto (CD73) | 4907 | ENSG00000135318 |
| 10 | P53007 | NA | SLC25A1 | solute carrier family 25 (mitochondrial carrier; citrate transporter), member 1 | 6576 | ENSG00000100075 |

  
  

| **Database:Wikipathways pathway      &nbspName:Epithelium TarBase      &nbspID:WP2002** | | | | | | |
| --- | --- | --- | --- | --- | --- | --- |
| C=340; O=8; E=1.44; R=5.55; rawP=9.78e-05; adjP=0.0002 | | | | | | |
| Index | UserID | Value | Gene Symbol | Gene Name | EntrezGene | Ensembl |
| 1 | Q71U36 | NA | TUBA1A | tubulin, alpha 1a | 7846 | ENSG00000167552 |
| 2 | P00491 | NA | PNP | purine nucleoside phosphorylase | 4860 | ENSG00000198805 |
| 3 | P16615 | NA | ATP2A2 | ATPase, Ca++ transporting, cardiac muscle, slow twitch 2 | 488 | ENSG00000174437 |
| 4 | P40121 | NA | CAPG | capping protein (actin filament), gelsolin-like | 822 | ENSG00000042493 |
| 5 | P42765 | NA | ACAA2 | acetyl-CoA acyltransferase 2 | 10449 | ENSG00000167315 |
| 6 | Q9NQC3 | NA | RTN4 | reticulon 4 | 57142 | ENSG00000115310 |
| 7 | P21589 | NA | NT5E | 5'-nucleotidase, ecto (CD73) | 4907 | ENSG00000135318 |
| 8 | P53007 | NA | SLC25A1 | solute carrier family 25 (mitochondrial carrier; citrate transporter), member 1 | 6576 | ENSG00000100075 |

  
  

| **Database:Wikipathways pathway      &nbspName:Muscle cell TarBase      &nbspID:WP2005** | | | | | | |
| --- | --- | --- | --- | --- | --- | --- |
| C=424; O=8; E=1.80; R=4.45; rawP=0.0004; adjP=0.0008 | | | | | | |
| Index | UserID | Value | Gene Symbol | Gene Name | EntrezGene | Ensembl |
| 1 | Q16881 | NA | TXNRD1 | thioredoxin reductase 1 | 7296 | ENSG00000198431 |
| 2 | Q9NQC3 | NA | RTN4 | reticulon 4 | 57142 | ENSG00000115310 |
| 3 | Q6PIU2 | NA | NCEH1 | neutral cholesterol ester hydrolase 1 | 57552 | ENSG00000144959 |
| 4 | P16615 | NA | ATP2A2 | ATPase, Ca++ transporting, cardiac muscle, slow twitch 2 | 488 | ENSG00000174437 |
| 5 | P53007 | NA | SLC25A1 | solute carrier family 25 (mitochondrial carrier; citrate transporter), member 1 | 6576 | ENSG00000100075 |
| 6 | P21589 | NA | NT5E | 5'-nucleotidase, ecto (CD73) | 4907 | ENSG00000135318 |
| 7 | P40121 | NA | CAPG | capping protein (actin filament), gelsolin-like | 822 | ENSG00000042493 |
| 8 | P42765 | NA | ACAA2 | acetyl-CoA acyltransferase 2 | 10449 | ENSG00000167315 |

  
  
  
  

---

WebGestalt is currently developed and maintained by Jing Wang and Bing Zhang at the  Zhang Lab. Other people who have made significant contribution to the project include Dexter Duncan, Stefan Kirov, Zhiao Shi, and Jay Snoddy.  
  
**Funding credits:** NIH/NIAAA (U01 AA016662, U01 AA013512); NIH/NIDA (P01 DA015027); NIH/NIMH (P50 MH078028, P50 MH096972); NIH/NCI (U24 CA159988); NIH/NIGMS (R01 GM088822).
