## Supplementary material for "PGRMC1 phosphorylation and cell plasticity 1: glycolysis, mitochondria, tumor growth": File S7: final_sig_file_1438746600.html

Anchored HTML File of EIDs

|  |  |
| --- | --- |
|  | WEB-based GEne SeT AnaLysis Toolkit |
|  | ***Translating gene lists into biological insights...*** |

---

  

| **Database:biological process      &nbspName:oxoacid metabolic process      &nbspID:GO:0043436** | | | | | | |
| --- | --- | --- | --- | --- | --- | --- |
| C=949; O=17; E=2.57; R=6.62; rawP=9.08e-11; adjP=1.63e-08 | | | | | | |
| Index | UserID | Value | Gene Symbol | Gene Name | EntrezGene | Ensembl |
| 1 | Q16851 | NA | UGP2 | UDP-glucose pyrophosphorylase 2 | 7360 | ENSG00000169764 |
| 2 | P00338 | NA | LDHA | lactate dehydrogenase A | 3939 | ENSG00000134333 |
| 3 | P00367 | NA | GLUD1 | glutamate dehydrogenase 1 | 2746 | ENSG00000148672 |
| 4 | Q06323 | NA | PSME1 | proteasome (prosome, macropain) activator subunit 1 (PA28 alpha) | 5720 | ENSG00000092010 |
| 5 | P48735 | NA | IDH2 | isocitrate dehydrogenase 2 (NADP+), mitochondrial | 3418 | ENSG00000182054 |
| 6 | P38571 | NA | LIPA | lipase A, lysosomal acid, cholesterol esterase | 3988 | ENSG00000107798 |
| 7 | P09211 | NA | GSTP1 | glutathione S-transferase pi 1 | 2950 | ENSG00000084207 |
| 8 | Q13011 | NA | ECH1 | enoyl CoA hydratase 1, peroxisomal | 1891 | ENSG00000104823 |
| 9 | P23381 | NA | WARS | tryptophanyl-tRNA synthetase | 7453 | ENSG00000140105 |
| 10 | Q9UBQ7 | NA | GRHPR | glyoxylate reductase/hydroxypyruvate reductase | 9380 | ENSG00000137106 |
| 11 | P51649 | NA | ALDH5A1 | aldehyde dehydrogenase 5 family, member A1 | 7915 | ENSG00000112294 |
| 12 | P42765 | NA | ACAA2 | acetyl-CoA acyltransferase 2 | 10449 | ENSG00000167315 |
| 13 | P24752 | NA | ACAT1 | acetyl-CoA acetyltransferase 1 | 38 | ENSG00000075239 |
| 14 | P13674 | NA | P4HA1 | prolyl 4-hydroxylase, alpha polypeptide I | 5033 | ENSG00000122884 |
| 15 | P49419 | NA | ALDH7A1 | aldehyde dehydrogenase 7 family, member A1 | 501 | ENSG00000164904 |
| 16 | P16278 | NA | GLB1 | galactosidase, beta 1 | 2720 | ENSG00000170266 |
| 17 | P49748 | NA | ACADVL | acyl-CoA dehydrogenase, very long chain | 37 | ENSG00000072778 |

  
  

| **Database:biological process      &nbspName:carboxylic acid metabolic process      &nbspID:GO:0019752** | | | | | | |
| --- | --- | --- | --- | --- | --- | --- |
| C=842; O=16; E=2.28; R=7.03; rawP=1.68e-10; adjP=1.63e-08 | | | | | | |
| Index | UserID | Value | Gene Symbol | Gene Name | EntrezGene | Ensembl |
| 1 | Q16851 | NA | UGP2 | UDP-glucose pyrophosphorylase 2 | 7360 | ENSG00000169764 |
| 2 | P00338 | NA | LDHA | lactate dehydrogenase A | 3939 | ENSG00000134333 |
| 3 | P00367 | NA | GLUD1 | glutamate dehydrogenase 1 | 2746 | ENSG00000148672 |
| 4 | Q06323 | NA | PSME1 | proteasome (prosome, macropain) activator subunit 1 (PA28 alpha) | 5720 | ENSG00000092010 |
| 5 | P48735 | NA | IDH2 | isocitrate dehydrogenase 2 (NADP+), mitochondrial | 3418 | ENSG00000182054 |
| 6 | P38571 | NA | LIPA | lipase A, lysosomal acid, cholesterol esterase | 3988 | ENSG00000107798 |
| 7 | P09211 | NA | GSTP1 | glutathione S-transferase pi 1 | 2950 | ENSG00000084207 |
| 8 | Q13011 | NA | ECH1 | enoyl CoA hydratase 1, peroxisomal | 1891 | ENSG00000104823 |
| 9 | P23381 | NA | WARS | tryptophanyl-tRNA synthetase | 7453 | ENSG00000140105 |
| 10 | Q9UBQ7 | NA | GRHPR | glyoxylate reductase/hydroxypyruvate reductase | 9380 | ENSG00000137106 |
| 11 | P51649 | NA | ALDH5A1 | aldehyde dehydrogenase 5 family, member A1 | 7915 | ENSG00000112294 |
| 12 | P42765 | NA | ACAA2 | acetyl-CoA acyltransferase 2 | 10449 | ENSG00000167315 |
| 13 | P24752 | NA | ACAT1 | acetyl-CoA acetyltransferase 1 | 38 | ENSG00000075239 |
| 14 | P13674 | NA | P4HA1 | prolyl 4-hydroxylase, alpha polypeptide I | 5033 | ENSG00000122884 |
| 15 | P49419 | NA | ALDH7A1 | aldehyde dehydrogenase 7 family, member A1 | 501 | ENSG00000164904 |
| 16 | P49748 | NA | ACADVL | acyl-CoA dehydrogenase, very long chain | 37 | ENSG00000072778 |

  
  

| **Database:biological process      &nbspName:organic acid metabolic process      &nbspID:GO:0006082** | | | | | | |
| --- | --- | --- | --- | --- | --- | --- |
| C=967; O=17; E=2.61; R=6.50; rawP=1.22e-10; adjP=1.63e-08 | | | | | | |
| Index | UserID | Value | Gene Symbol | Gene Name | EntrezGene | Ensembl |
| 1 | Q16851 | NA | UGP2 | UDP-glucose pyrophosphorylase 2 | 7360 | ENSG00000169764 |
| 2 | P00338 | NA | LDHA | lactate dehydrogenase A | 3939 | ENSG00000134333 |
| 3 | P00367 | NA | GLUD1 | glutamate dehydrogenase 1 | 2746 | ENSG00000148672 |
| 4 | Q06323 | NA | PSME1 | proteasome (prosome, macropain) activator subunit 1 (PA28 alpha) | 5720 | ENSG00000092010 |
| 5 | P48735 | NA | IDH2 | isocitrate dehydrogenase 2 (NADP+), mitochondrial | 3418 | ENSG00000182054 |
| 6 | P38571 | NA | LIPA | lipase A, lysosomal acid, cholesterol esterase | 3988 | ENSG00000107798 |
| 7 | P09211 | NA | GSTP1 | glutathione S-transferase pi 1 | 2950 | ENSG00000084207 |
| 8 | Q13011 | NA | ECH1 | enoyl CoA hydratase 1, peroxisomal | 1891 | ENSG00000104823 |
| 9 | P23381 | NA | WARS | tryptophanyl-tRNA synthetase | 7453 | ENSG00000140105 |
| 10 | Q9UBQ7 | NA | GRHPR | glyoxylate reductase/hydroxypyruvate reductase | 9380 | ENSG00000137106 |
| 11 | P51649 | NA | ALDH5A1 | aldehyde dehydrogenase 5 family, member A1 | 7915 | ENSG00000112294 |
| 12 | P42765 | NA | ACAA2 | acetyl-CoA acyltransferase 2 | 10449 | ENSG00000167315 |
| 13 | P24752 | NA | ACAT1 | acetyl-CoA acetyltransferase 1 | 38 | ENSG00000075239 |
| 14 | P13674 | NA | P4HA1 | prolyl 4-hydroxylase, alpha polypeptide I | 5033 | ENSG00000122884 |
| 15 | P49419 | NA | ALDH7A1 | aldehyde dehydrogenase 7 family, member A1 | 501 | ENSG00000164904 |
| 16 | P16278 | NA | GLB1 | galactosidase, beta 1 | 2720 | ENSG00000170266 |
| 17 | P49748 | NA | ACADVL | acyl-CoA dehydrogenase, very long chain | 37 | ENSG00000072778 |

  
  

| **Database:biological process      &nbspName:small molecule metabolic process      &nbspID:GO:0044281** | | | | | | |
| --- | --- | --- | --- | --- | --- | --- |
| C=2500; O=24; E=6.76; R=3.55; rawP=8.35e-10; adjP=6.07e-08 | | | | | | |
| Index | UserID | Value | Gene Symbol | Gene Name | EntrezGene | Ensembl |
| 1 | Q16851 | NA | UGP2 | UDP-glucose pyrophosphorylase 2 | 7360 | ENSG00000169764 |
| 2 | P00338 | NA | LDHA | lactate dehydrogenase A | 3939 | ENSG00000134333 |
| 3 | P00367 | NA | GLUD1 | glutamate dehydrogenase 1 | 2746 | ENSG00000148672 |
| 4 | Q06323 | NA | PSME1 | proteasome (prosome, macropain) activator subunit 1 (PA28 alpha) | 5720 | ENSG00000092010 |
| 5 | P48735 | NA | IDH2 | isocitrate dehydrogenase 2 (NADP+), mitochondrial | 3418 | ENSG00000182054 |
| 6 | P38571 | NA | LIPA | lipase A, lysosomal acid, cholesterol esterase | 3988 | ENSG00000107798 |
| 7 | P09211 | NA | GSTP1 | glutathione S-transferase pi 1 | 2950 | ENSG00000084207 |
| 8 | Q9P0J0 | NA | NDUFA13 | NADH dehydrogenase (ubiquinone) 1 alpha subcomplex, 13 | 51079 | ENSG00000186010 |
| 9 | Q13011 | NA | ECH1 | enoyl CoA hydratase 1, peroxisomal | 1891 | ENSG00000104823 |
| 10 | P23381 | NA | WARS | tryptophanyl-tRNA synthetase | 7453 | ENSG00000140105 |
| 11 | P04040 | NA | CAT | catalase | 847 | ENSG00000121691 |
| 12 | Q9UBQ7 | NA | GRHPR | glyoxylate reductase/hydroxypyruvate reductase | 9380 | ENSG00000137106 |
| 13 | P51649 | NA | ALDH5A1 | aldehyde dehydrogenase 5 family, member A1 | 7915 | ENSG00000112294 |
| 14 | P22570 | NA | FDXR | ferredoxin reductase | 2232 | ENSG00000161513 |
| 15 | P30044 | NA | PRDX5 | peroxiredoxin 5 | 25824 | ENSG00000126432 |
| 16 | P42765 | NA | ACAA2 | acetyl-CoA acyltransferase 2 | 10449 | ENSG00000167315 |
| 17 | P13674 | NA | P4HA1 | prolyl 4-hydroxylase, alpha polypeptide I | 5033 | ENSG00000122884 |
| 18 | P24752 | NA | ACAT1 | acetyl-CoA acetyltransferase 1 | 38 | ENSG00000075239 |
| 19 | P16278 | NA | GLB1 | galactosidase, beta 1 | 2720 | ENSG00000170266 |
| 20 | P49419 | NA | ALDH7A1 | aldehyde dehydrogenase 7 family, member A1 | 501 | ENSG00000164904 |
| 21 | P06744 | NA | GPI | glucose-6-phosphate isomerase | 2821 | ENSG00000105220 |
| 22 | P07602 | NA | PSAP | prosaposin | 5660 | ENSG00000197746 |
| 23 | P49748 | NA | ACADVL | acyl-CoA dehydrogenase, very long chain | 37 | ENSG00000072778 |
| 24 | P53007 | NA | SLC25A1 | solute carrier family 25 (mitochondrial carrier; citrate transporter), member 1 | 6576 | ENSG00000100075 |

  
  

| **Database:biological process      &nbspName:oxidation-reduction process      &nbspID:GO:0055114** | | | | | | |
| --- | --- | --- | --- | --- | --- | --- |
| C=554; O=12; E=1.50; R=8.01; rawP=1.39e-08; adjP=8.09e-07 | | | | | | |
| Index | UserID | Value | Gene Symbol | Gene Name | EntrezGene | Ensembl |
| 1 | P04040 | NA | CAT | catalase | 847 | ENSG00000121691 |
| 2 | Q9UBQ7 | NA | GRHPR | glyoxylate reductase/hydroxypyruvate reductase | 9380 | ENSG00000137106 |
| 3 | P10253 | NA | GAA | glucosidase, alpha; acid | 2548 | ENSG00000171298 |
| 4 | P51649 | NA | ALDH5A1 | aldehyde dehydrogenase 5 family, member A1 | 7915 | ENSG00000112294 |
| 5 | Q16851 | NA | UGP2 | UDP-glucose pyrophosphorylase 2 | 7360 | ENSG00000169764 |
| 6 | P30044 | NA | PRDX5 | peroxiredoxin 5 | 25824 | ENSG00000126432 |
| 7 | P22570 | NA | FDXR | ferredoxin reductase | 2232 | ENSG00000161513 |
| 8 | P42765 | NA | ACAA2 | acetyl-CoA acyltransferase 2 | 10449 | ENSG00000167315 |
| 9 | P48735 | NA | IDH2 | isocitrate dehydrogenase 2 (NADP+), mitochondrial | 3418 | ENSG00000182054 |
| 10 | P49748 | NA | ACADVL | acyl-CoA dehydrogenase, very long chain | 37 | ENSG00000072778 |
| 11 | Q9P0J0 | NA | NDUFA13 | NADH dehydrogenase (ubiquinone) 1 alpha subcomplex, 13 | 51079 | ENSG00000186010 |
| 12 | Q13011 | NA | ECH1 | enoyl CoA hydratase 1, peroxisomal | 1891 | ENSG00000104823 |

  
  

| **Database:biological process      &nbspName:monocarboxylic acid metabolic process      &nbspID:GO:0032787** | | | | | | |
| --- | --- | --- | --- | --- | --- | --- |
| C=405; O=10; E=1.10; R=9.13; rawP=8.38e-08; adjP=4.06e-06 | | | | | | |
| Index | UserID | Value | Gene Symbol | Gene Name | EntrezGene | Ensembl |
| 1 | Q9UBQ7 | NA | GRHPR | glyoxylate reductase/hydroxypyruvate reductase | 9380 | ENSG00000137106 |
| 2 | P51649 | NA | ALDH5A1 | aldehyde dehydrogenase 5 family, member A1 | 7915 | ENSG00000112294 |
| 3 | Q16851 | NA | UGP2 | UDP-glucose pyrophosphorylase 2 | 7360 | ENSG00000169764 |
| 4 | P00338 | NA | LDHA | lactate dehydrogenase A | 3939 | ENSG00000134333 |
| 5 | P42765 | NA | ACAA2 | acetyl-CoA acyltransferase 2 | 10449 | ENSG00000167315 |
| 6 | P13674 | NA | P4HA1 | prolyl 4-hydroxylase, alpha polypeptide I | 5033 | ENSG00000122884 |
| 7 | P48735 | NA | IDH2 | isocitrate dehydrogenase 2 (NADP+), mitochondrial | 3418 | ENSG00000182054 |
| 8 | P38571 | NA | LIPA | lipase A, lysosomal acid, cholesterol esterase | 3988 | ENSG00000107798 |
| 9 | P49748 | NA | ACADVL | acyl-CoA dehydrogenase, very long chain | 37 | ENSG00000072778 |
| 10 | Q13011 | NA | ECH1 | enoyl CoA hydratase 1, peroxisomal | 1891 | ENSG00000104823 |

  
  

| **Database:biological process      &nbspName:generation of precursor metabolites and energy      &nbspID:GO:0006091** | | | | | | |
| --- | --- | --- | --- | --- | --- | --- |
| C=447; O=10; E=1.21; R=8.27; rawP=2.10e-07; adjP=8.73e-06 | | | | | | |
| Index | UserID | Value | Gene Symbol | Gene Name | EntrezGene | Ensembl |
| 1 | P04040 | NA | CAT | catalase | 847 | ENSG00000121691 |
| 2 | P10253 | NA | GAA | glucosidase, alpha; acid | 2548 | ENSG00000171298 |
| 3 | P51649 | NA | ALDH5A1 | aldehyde dehydrogenase 5 family, member A1 | 7915 | ENSG00000112294 |
| 4 | Q16851 | NA | UGP2 | UDP-glucose pyrophosphorylase 2 | 7360 | ENSG00000169764 |
| 5 | P22570 | NA | FDXR | ferredoxin reductase | 2232 | ENSG00000161513 |
| 6 | P00338 | NA | LDHA | lactate dehydrogenase A | 3939 | ENSG00000134333 |
| 7 | P48735 | NA | IDH2 | isocitrate dehydrogenase 2 (NADP+), mitochondrial | 3418 | ENSG00000182054 |
| 8 | P06744 | NA | GPI | glucose-6-phosphate isomerase | 2821 | ENSG00000105220 |
| 9 | P49748 | NA | ACADVL | acyl-CoA dehydrogenase, very long chain | 37 | ENSG00000072778 |
| 10 | Q9P0J0 | NA | NDUFA13 | NADH dehydrogenase (ubiquinone) 1 alpha subcomplex, 13 | 51079 | ENSG00000186010 |

  
  

| **Database:biological process      &nbspName:organic acid catabolic process      &nbspID:GO:0016054** | | | | | | |
| --- | --- | --- | --- | --- | --- | --- |
| C=198; O=7; E=0.54; R=13.07; rawP=8.77e-07; adjP=2.84e-05 | | | | | | |
| Index | UserID | Value | Gene Symbol | Gene Name | EntrezGene | Ensembl |
| 1 | P24752 | NA | ACAT1 | acetyl-CoA acetyltransferase 1 | 38 | ENSG00000075239 |
| 2 | P49419 | NA | ALDH7A1 | aldehyde dehydrogenase 7 family, member A1 | 501 | ENSG00000164904 |
| 3 | P00367 | NA | GLUD1 | glutamate dehydrogenase 1 | 2746 | ENSG00000148672 |
| 4 | P51649 | NA | ALDH5A1 | aldehyde dehydrogenase 5 family, member A1 | 7915 | ENSG00000112294 |
| 5 | P49748 | NA | ACADVL | acyl-CoA dehydrogenase, very long chain | 37 | ENSG00000072778 |
| 6 | Q13011 | NA | ECH1 | enoyl CoA hydratase 1, peroxisomal | 1891 | ENSG00000104823 |
| 7 | P42765 | NA | ACAA2 | acetyl-CoA acyltransferase 2 | 10449 | ENSG00000167315 |

  
  

| **Database:biological process      &nbspName:carboxylic acid catabolic process      &nbspID:GO:0046395** | | | | | | |
| --- | --- | --- | --- | --- | --- | --- |
| C=198; O=7; E=0.54; R=13.07; rawP=8.77e-07; adjP=2.84e-05 | | | | | | |
| Index | UserID | Value | Gene Symbol | Gene Name | EntrezGene | Ensembl |
| 1 | P24752 | NA | ACAT1 | acetyl-CoA acetyltransferase 1 | 38 | ENSG00000075239 |
| 2 | P49419 | NA | ALDH7A1 | aldehyde dehydrogenase 7 family, member A1 | 501 | ENSG00000164904 |
| 3 | P00367 | NA | GLUD1 | glutamate dehydrogenase 1 | 2746 | ENSG00000148672 |
| 4 | P51649 | NA | ALDH5A1 | aldehyde dehydrogenase 5 family, member A1 | 7915 | ENSG00000112294 |
| 5 | P49748 | NA | ACADVL | acyl-CoA dehydrogenase, very long chain | 37 | ENSG00000072778 |
| 6 | Q13011 | NA | ECH1 | enoyl CoA hydratase 1, peroxisomal | 1891 | ENSG00000104823 |
| 7 | P42765 | NA | ACAA2 | acetyl-CoA acyltransferase 2 | 10449 | ENSG00000167315 |

  
  

| **Database:biological process      &nbspName:catabolic process      &nbspID:GO:0009056** | | | | | | |
| --- | --- | --- | --- | --- | --- | --- |
| C=1991; O=18; E=5.38; R=3.34; rawP=1.06e-06; adjP=3.08e-05 | | | | | | |
| Index | UserID | Value | Gene Symbol | Gene Name | EntrezGene | Ensembl |
| 1 | P10253 | NA | GAA | glucosidase, alpha; acid | 2548 | ENSG00000171298 |
| 2 | P00338 | NA | LDHA | lactate dehydrogenase A | 3939 | ENSG00000134333 |
| 3 | P00367 | NA | GLUD1 | glutamate dehydrogenase 1 | 2746 | ENSG00000148672 |
| 4 | Q06323 | NA | PSME1 | proteasome (prosome, macropain) activator subunit 1 (PA28 alpha) | 5720 | ENSG00000092010 |
| 5 | P48735 | NA | IDH2 | isocitrate dehydrogenase 2 (NADP+), mitochondrial | 3418 | ENSG00000182054 |
| 6 | P38571 | NA | LIPA | lipase A, lysosomal acid, cholesterol esterase | 3988 | ENSG00000107798 |
| 7 | Q13011 | NA | ECH1 | enoyl CoA hydratase 1, peroxisomal | 1891 | ENSG00000104823 |
| 8 | O14773 | NA | TPP1 | tripeptidyl peptidase I | 1200 | ENSG00000166340 |
| 9 | P04040 | NA | CAT | catalase | 847 | ENSG00000121691 |
| 10 | P51649 | NA | ALDH5A1 | aldehyde dehydrogenase 5 family, member A1 | 7915 | ENSG00000112294 |
| 11 | P30044 | NA | PRDX5 | peroxiredoxin 5 | 25824 | ENSG00000126432 |
| 12 | P42765 | NA | ACAA2 | acetyl-CoA acyltransferase 2 | 10449 | ENSG00000167315 |
| 13 | P24752 | NA | ACAT1 | acetyl-CoA acetyltransferase 1 | 38 | ENSG00000075239 |
| 14 | P49419 | NA | ALDH7A1 | aldehyde dehydrogenase 7 family, member A1 | 501 | ENSG00000164904 |
| 15 | P16278 | NA | GLB1 | galactosidase, beta 1 | 2720 | ENSG00000170266 |
| 16 | P06744 | NA | GPI | glucose-6-phosphate isomerase | 2821 | ENSG00000105220 |
| 17 | P49748 | NA | ACADVL | acyl-CoA dehydrogenase, very long chain | 37 | ENSG00000072778 |
| 18 | P07384 | NA | CAPN1 | calpain 1, (mu/I) large subunit | 823 | ENSG00000014216 |

  
  

| **Database:biological process      &nbspName:organic substance catabolic process      &nbspID:GO:1901575** | | | | | | |
| --- | --- | --- | --- | --- | --- | --- |
| C=1851; O=17; E=5.01; R=3.40; rawP=2.01e-06; adjP=5.32e-05 | | | | | | |
| Index | UserID | Value | Gene Symbol | Gene Name | EntrezGene | Ensembl |
| 1 | P10253 | NA | GAA | glucosidase, alpha; acid | 2548 | ENSG00000171298 |
| 2 | P00338 | NA | LDHA | lactate dehydrogenase A | 3939 | ENSG00000134333 |
| 3 | P00367 | NA | GLUD1 | glutamate dehydrogenase 1 | 2746 | ENSG00000148672 |
| 4 | Q06323 | NA | PSME1 | proteasome (prosome, macropain) activator subunit 1 (PA28 alpha) | 5720 | ENSG00000092010 |
| 5 | P48735 | NA | IDH2 | isocitrate dehydrogenase 2 (NADP+), mitochondrial | 3418 | ENSG00000182054 |
| 6 | P38571 | NA | LIPA | lipase A, lysosomal acid, cholesterol esterase | 3988 | ENSG00000107798 |
| 7 | Q13011 | NA | ECH1 | enoyl CoA hydratase 1, peroxisomal | 1891 | ENSG00000104823 |
| 8 | O14773 | NA | TPP1 | tripeptidyl peptidase I | 1200 | ENSG00000166340 |
| 9 | P04040 | NA | CAT | catalase | 847 | ENSG00000121691 |
| 10 | P51649 | NA | ALDH5A1 | aldehyde dehydrogenase 5 family, member A1 | 7915 | ENSG00000112294 |
| 11 | P42765 | NA | ACAA2 | acetyl-CoA acyltransferase 2 | 10449 | ENSG00000167315 |
| 12 | P24752 | NA | ACAT1 | acetyl-CoA acetyltransferase 1 | 38 | ENSG00000075239 |
| 13 | P49419 | NA | ALDH7A1 | aldehyde dehydrogenase 7 family, member A1 | 501 | ENSG00000164904 |
| 14 | P16278 | NA | GLB1 | galactosidase, beta 1 | 2720 | ENSG00000170266 |
| 15 | P06744 | NA | GPI | glucose-6-phosphate isomerase | 2821 | ENSG00000105220 |
| 16 | P49748 | NA | ACADVL | acyl-CoA dehydrogenase, very long chain | 37 | ENSG00000072778 |
| 17 | P07384 | NA | CAPN1 | calpain 1, (mu/I) large subunit | 823 | ENSG00000014216 |

  
  

| **Database:biological process      &nbspName:hexose metabolic process      &nbspID:GO:0019318** | | | | | | |
| --- | --- | --- | --- | --- | --- | --- |
| C=249; O=7; E=0.67; R=10.40; rawP=4.03e-06; adjP=9.02e-05 | | | | | | |
| Index | UserID | Value | Gene Symbol | Gene Name | EntrezGene | Ensembl |
| 1 | P16278 | NA | GLB1 | galactosidase, beta 1 | 2720 | ENSG00000170266 |
| 2 | P10253 | NA | GAA | glucosidase, alpha; acid | 2548 | ENSG00000171298 |
| 3 | P06744 | NA | GPI | glucose-6-phosphate isomerase | 2821 | ENSG00000105220 |
| 4 | P51649 | NA | ALDH5A1 | aldehyde dehydrogenase 5 family, member A1 | 7915 | ENSG00000112294 |
| 5 | Q16851 | NA | UGP2 | UDP-glucose pyrophosphorylase 2 | 7360 | ENSG00000169764 |
| 6 | P00338 | NA | LDHA | lactate dehydrogenase A | 3939 | ENSG00000134333 |
| 7 | P53007 | NA | SLC25A1 | solute carrier family 25 (mitochondrial carrier; citrate transporter), member 1 | 6576 | ENSG00000100075 |

  
  

| **Database:biological process      &nbspName:cellular aldehyde metabolic process      &nbspID:GO:0006081** | | | | | | |
| --- | --- | --- | --- | --- | --- | --- |
| C=40; O=4; E=0.11; R=36.98; rawP=3.89e-06; adjP=9.02e-05 | | | | | | |
| Index | UserID | Value | Gene Symbol | Gene Name | EntrezGene | Ensembl |
| 1 | Q9UBQ7 | NA | GRHPR | glyoxylate reductase/hydroxypyruvate reductase | 9380 | ENSG00000137106 |
| 2 | P49419 | NA | ALDH7A1 | aldehyde dehydrogenase 7 family, member A1 | 501 | ENSG00000164904 |
| 3 | P48735 | NA | IDH2 | isocitrate dehydrogenase 2 (NADP+), mitochondrial | 3418 | ENSG00000182054 |
| 4 | P06744 | NA | GPI | glucose-6-phosphate isomerase | 2821 | ENSG00000105220 |

  
  

| **Database:biological process      &nbspName:small molecule catabolic process      &nbspID:GO:0044282** | | | | | | |
| --- | --- | --- | --- | --- | --- | --- |
| C=258; O=7; E=0.70; R=10.03; rawP=5.10e-06; adjP=0.0001 | | | | | | |
| Index | UserID | Value | Gene Symbol | Gene Name | EntrezGene | Ensembl |
| 1 | P24752 | NA | ACAT1 | acetyl-CoA acetyltransferase 1 | 38 | ENSG00000075239 |
| 2 | P49419 | NA | ALDH7A1 | aldehyde dehydrogenase 7 family, member A1 | 501 | ENSG00000164904 |
| 3 | P00367 | NA | GLUD1 | glutamate dehydrogenase 1 | 2746 | ENSG00000148672 |
| 4 | P51649 | NA | ALDH5A1 | aldehyde dehydrogenase 5 family, member A1 | 7915 | ENSG00000112294 |
| 5 | P49748 | NA | ACADVL | acyl-CoA dehydrogenase, very long chain | 37 | ENSG00000072778 |
| 6 | Q13011 | NA | ECH1 | enoyl CoA hydratase 1, peroxisomal | 1891 | ENSG00000104823 |
| 7 | P42765 | NA | ACAA2 | acetyl-CoA acyltransferase 2 | 10449 | ENSG00000167315 |

  
  

| **Database:biological process      &nbspName:single-organism catabolic process      &nbspID:GO:0044712** | | | | | | |
| --- | --- | --- | --- | --- | --- | --- |
| C=259; O=7; E=0.70; R=9.99; rawP=5.23e-06; adjP=0.0001 | | | | | | |
| Index | UserID | Value | Gene Symbol | Gene Name | EntrezGene | Ensembl |
| 1 | P24752 | NA | ACAT1 | acetyl-CoA acetyltransferase 1 | 38 | ENSG00000075239 |
| 2 | P49419 | NA | ALDH7A1 | aldehyde dehydrogenase 7 family, member A1 | 501 | ENSG00000164904 |
| 3 | P00367 | NA | GLUD1 | glutamate dehydrogenase 1 | 2746 | ENSG00000148672 |
| 4 | P51649 | NA | ALDH5A1 | aldehyde dehydrogenase 5 family, member A1 | 7915 | ENSG00000112294 |
| 5 | P49748 | NA | ACADVL | acyl-CoA dehydrogenase, very long chain | 37 | ENSG00000072778 |
| 6 | Q13011 | NA | ECH1 | enoyl CoA hydratase 1, peroxisomal | 1891 | ENSG00000104823 |
| 7 | P42765 | NA | ACAA2 | acetyl-CoA acyltransferase 2 | 10449 | ENSG00000167315 |

  
  

| **Database:biological process      &nbspName:monosaccharide metabolic process      &nbspID:GO:0005996** | | | | | | |
| --- | --- | --- | --- | --- | --- | --- |
| C=279; O=7; E=0.75; R=9.28; rawP=8.51e-06; adjP=0.0002 | | | | | | |
| Index | UserID | Value | Gene Symbol | Gene Name | EntrezGene | Ensembl |
| 1 | P16278 | NA | GLB1 | galactosidase, beta 1 | 2720 | ENSG00000170266 |
| 2 | P10253 | NA | GAA | glucosidase, alpha; acid | 2548 | ENSG00000171298 |
| 3 | P06744 | NA | GPI | glucose-6-phosphate isomerase | 2821 | ENSG00000105220 |
| 4 | P51649 | NA | ALDH5A1 | aldehyde dehydrogenase 5 family, member A1 | 7915 | ENSG00000112294 |
| 5 | Q16851 | NA | UGP2 | UDP-glucose pyrophosphorylase 2 | 7360 | ENSG00000169764 |
| 6 | P00338 | NA | LDHA | lactate dehydrogenase A | 3939 | ENSG00000134333 |
| 7 | P53007 | NA | SLC25A1 | solute carrier family 25 (mitochondrial carrier; citrate transporter), member 1 | 6576 | ENSG00000100075 |

  
  

| **Database:biological process      &nbspName:cellular catabolic process      &nbspID:GO:0044248** | | | | | | |
| --- | --- | --- | --- | --- | --- | --- |
| C=1665; O=15; E=4.50; R=3.33; rawP=1.36e-05; adjP=0.0002 | | | | | | |
| Index | UserID | Value | Gene Symbol | Gene Name | EntrezGene | Ensembl |
| 1 | P04040 | NA | CAT | catalase | 847 | ENSG00000121691 |
| 2 | O14773 | NA | TPP1 | tripeptidyl peptidase I | 1200 | ENSG00000166340 |
| 3 | P51649 | NA | ALDH5A1 | aldehyde dehydrogenase 5 family, member A1 | 7915 | ENSG00000112294 |
| 4 | P30044 | NA | PRDX5 | peroxiredoxin 5 | 25824 | ENSG00000126432 |
| 5 | P42765 | NA | ACAA2 | acetyl-CoA acyltransferase 2 | 10449 | ENSG00000167315 |
| 6 | P24752 | NA | ACAT1 | acetyl-CoA acetyltransferase 1 | 38 | ENSG00000075239 |
| 7 | P16278 | NA | GLB1 | galactosidase, beta 1 | 2720 | ENSG00000170266 |
| 8 | P00367 | NA | GLUD1 | glutamate dehydrogenase 1 | 2746 | ENSG00000148672 |
| 9 | Q06323 | NA | PSME1 | proteasome (prosome, macropain) activator subunit 1 (PA28 alpha) | 5720 | ENSG00000092010 |
| 10 | P48735 | NA | IDH2 | isocitrate dehydrogenase 2 (NADP+), mitochondrial | 3418 | ENSG00000182054 |
| 11 | P49419 | NA | ALDH7A1 | aldehyde dehydrogenase 7 family, member A1 | 501 | ENSG00000164904 |
| 12 | P06744 | NA | GPI | glucose-6-phosphate isomerase | 2821 | ENSG00000105220 |
| 13 | P49748 | NA | ACADVL | acyl-CoA dehydrogenase, very long chain | 37 | ENSG00000072778 |
| 14 | P07384 | NA | CAPN1 | calpain 1, (mu/I) large subunit | 823 | ENSG00000014216 |
| 15 | Q13011 | NA | ECH1 | enoyl CoA hydratase 1, peroxisomal | 1891 | ENSG00000104823 |

  
  

| **Database:biological process      &nbspName:sterol metabolic process      &nbspID:GO:0016125** | | | | | | |
| --- | --- | --- | --- | --- | --- | --- |
| C=122; O=5; E=0.33; R=15.16; rawP=1.82e-05; adjP=0.0003 | | | | | | |
| Index | UserID | Value | Gene Symbol | Gene Name | EntrezGene | Ensembl |
| 1 | P04040 | NA | CAT | catalase | 847 | ENSG00000121691 |
| 2 | P49748 | NA | ACADVL | acyl-CoA dehydrogenase, very long chain | 37 | ENSG00000072778 |
| 3 | P38571 | NA | LIPA | lipase A, lysosomal acid, cholesterol esterase | 3988 | ENSG00000107798 |
| 4 | P22570 | NA | FDXR | ferredoxin reductase | 2232 | ENSG00000161513 |
| 5 | P42765 | NA | ACAA2 | acetyl-CoA acyltransferase 2 | 10449 | ENSG00000167315 |

  
  

| **Database:biological process      &nbspName:energy derivation by oxidation of organic compounds      &nbspID:GO:0015980** | | | | | | |
| --- | --- | --- | --- | --- | --- | --- |
| C=326; O=7; E=0.88; R=7.94; rawP=2.33e-05; adjP=0.0003 | | | | | | |
| Index | UserID | Value | Gene Symbol | Gene Name | EntrezGene | Ensembl |
| 1 | P04040 | NA | CAT | catalase | 847 | ENSG00000121691 |
| 2 | P48735 | NA | IDH2 | isocitrate dehydrogenase 2 (NADP+), mitochondrial | 3418 | ENSG00000182054 |
| 3 | P10253 | NA | GAA | glucosidase, alpha; acid | 2548 | ENSG00000171298 |
| 4 | P51649 | NA | ALDH5A1 | aldehyde dehydrogenase 5 family, member A1 | 7915 | ENSG00000112294 |
| 5 | Q16851 | NA | UGP2 | UDP-glucose pyrophosphorylase 2 | 7360 | ENSG00000169764 |
| 6 | P49748 | NA | ACADVL | acyl-CoA dehydrogenase, very long chain | 37 | ENSG00000072778 |
| 7 | Q9P0J0 | NA | NDUFA13 | NADH dehydrogenase (ubiquinone) 1 alpha subcomplex, 13 | 51079 | ENSG00000186010 |

  
  

| **Database:biological process      &nbspName:glucose metabolic process      &nbspID:GO:0006006** | | | | | | |
| --- | --- | --- | --- | --- | --- | --- |
| C=213; O=6; E=0.58; R=10.42; rawP=2.10e-05; adjP=0.0003 | | | | | | |
| Index | UserID | Value | Gene Symbol | Gene Name | EntrezGene | Ensembl |
| 1 | P10253 | NA | GAA | glucosidase, alpha; acid | 2548 | ENSG00000171298 |
| 2 | P06744 | NA | GPI | glucose-6-phosphate isomerase | 2821 | ENSG00000105220 |
| 3 | P51649 | NA | ALDH5A1 | aldehyde dehydrogenase 5 family, member A1 | 7915 | ENSG00000112294 |
| 4 | Q16851 | NA | UGP2 | UDP-glucose pyrophosphorylase 2 | 7360 | ENSG00000169764 |
| 5 | P00338 | NA | LDHA | lactate dehydrogenase A | 3939 | ENSG00000134333 |
| 6 | P53007 | NA | SLC25A1 | solute carrier family 25 (mitochondrial carrier; citrate transporter), member 1 | 6576 | ENSG00000100075 |

  
  

| **Database:biological process      &nbspName:lipid metabolic process      &nbspID:GO:0006629** | | | | | | |
| --- | --- | --- | --- | --- | --- | --- |
| C=1137; O=12; E=3.07; R=3.90; rawP=2.87e-05; adjP=0.0004 | | | | | | |
| Index | UserID | Value | Gene Symbol | Gene Name | EntrezGene | Ensembl |
| 1 | P04040 | NA | CAT | catalase | 847 | ENSG00000121691 |
| 2 | O14773 | NA | TPP1 | tripeptidyl peptidase I | 1200 | ENSG00000166340 |
| 3 | P51649 | NA | ALDH5A1 | aldehyde dehydrogenase 5 family, member A1 | 7915 | ENSG00000112294 |
| 4 | P22570 | NA | FDXR | ferredoxin reductase | 2232 | ENSG00000161513 |
| 5 | P42765 | NA | ACAA2 | acetyl-CoA acyltransferase 2 | 10449 | ENSG00000167315 |
| 6 | P24752 | NA | ACAT1 | acetyl-CoA acetyltransferase 1 | 38 | ENSG00000075239 |
| 7 | P16278 | NA | GLB1 | galactosidase, beta 1 | 2720 | ENSG00000170266 |
| 8 | P07602 | NA | PSAP | prosaposin | 5660 | ENSG00000197746 |
| 9 | P49748 | NA | ACADVL | acyl-CoA dehydrogenase, very long chain | 37 | ENSG00000072778 |
| 10 | P38571 | NA | LIPA | lipase A, lysosomal acid, cholesterol esterase | 3988 | ENSG00000107798 |
| 11 | Q13011 | NA | ECH1 | enoyl CoA hydratase 1, peroxisomal | 1891 | ENSG00000104823 |
| 12 | P53007 | NA | SLC25A1 | solute carrier family 25 (mitochondrial carrier; citrate transporter), member 1 | 6576 | ENSG00000100075 |

  
  

| **Database:biological process      &nbspName:cellular lipid metabolic process      &nbspID:GO:0044255** | | | | | | |
| --- | --- | --- | --- | --- | --- | --- |
| C=809; O=10; E=2.19; R=4.57; rawP=4.19e-05; adjP=0.0006 | | | | | | |
| Index | UserID | Value | Gene Symbol | Gene Name | EntrezGene | Ensembl |
| 1 | P04040 | NA | CAT | catalase | 847 | ENSG00000121691 |
| 2 | P51649 | NA | ALDH5A1 | aldehyde dehydrogenase 5 family, member A1 | 7915 | ENSG00000112294 |
| 3 | P42765 | NA | ACAA2 | acetyl-CoA acyltransferase 2 | 10449 | ENSG00000167315 |
| 4 | P24752 | NA | ACAT1 | acetyl-CoA acetyltransferase 1 | 38 | ENSG00000075239 |
| 5 | P16278 | NA | GLB1 | galactosidase, beta 1 | 2720 | ENSG00000170266 |
| 6 | P07602 | NA | PSAP | prosaposin | 5660 | ENSG00000197746 |
| 7 | P49748 | NA | ACADVL | acyl-CoA dehydrogenase, very long chain | 37 | ENSG00000072778 |
| 8 | P38571 | NA | LIPA | lipase A, lysosomal acid, cholesterol esterase | 3988 | ENSG00000107798 |
| 9 | Q13011 | NA | ECH1 | enoyl CoA hydratase 1, peroxisomal | 1891 | ENSG00000104823 |
| 10 | P53007 | NA | SLC25A1 | solute carrier family 25 (mitochondrial carrier; citrate transporter), member 1 | 6576 | ENSG00000100075 |

  
  

| **Database:biological process      &nbspName:fatty acid catabolic process      &nbspID:GO:0009062** | | | | | | |
| --- | --- | --- | --- | --- | --- | --- |
| C=75; O=4; E=0.20; R=19.72; rawP=4.83e-05; adjP=0.0006 | | | | | | |
| Index | UserID | Value | Gene Symbol | Gene Name | EntrezGene | Ensembl |
| 1 | P51649 | NA | ALDH5A1 | aldehyde dehydrogenase 5 family, member A1 | 7915 | ENSG00000112294 |
| 2 | P49748 | NA | ACADVL | acyl-CoA dehydrogenase, very long chain | 37 | ENSG00000072778 |
| 3 | Q13011 | NA | ECH1 | enoyl CoA hydratase 1, peroxisomal | 1891 | ENSG00000104823 |
| 4 | P42765 | NA | ACAA2 | acetyl-CoA acyltransferase 2 | 10449 | ENSG00000167315 |

  
  

| **Database:biological process      &nbspName:organonitrogen compound metabolic process      &nbspID:GO:1901564** | | | | | | |
| --- | --- | --- | --- | --- | --- | --- |
| C=1673; O=14; E=4.52; R=3.09; rawP=6.83e-05; adjP=0.0008 | | | | | | |
| Index | UserID | Value | Gene Symbol | Gene Name | EntrezGene | Ensembl |
| 1 | P04040 | NA | CAT | catalase | 847 | ENSG00000121691 |
| 2 | O14773 | NA | TPP1 | tripeptidyl peptidase I | 1200 | ENSG00000166340 |
| 3 | P51649 | NA | ALDH5A1 | aldehyde dehydrogenase 5 family, member A1 | 7915 | ENSG00000112294 |
| 4 | P30044 | NA | PRDX5 | peroxiredoxin 5 | 25824 | ENSG00000126432 |
| 5 | P30086 | NA | PEBP1 | phosphatidylethanolamine binding protein 1 | 5037 | ENSG00000089220 |
| 6 | P13674 | NA | P4HA1 | prolyl 4-hydroxylase, alpha polypeptide I | 5033 | ENSG00000122884 |
| 7 | P24752 | NA | ACAT1 | acetyl-CoA acetyltransferase 1 | 38 | ENSG00000075239 |
| 8 | P16278 | NA | GLB1 | galactosidase, beta 1 | 2720 | ENSG00000170266 |
| 9 | P00367 | NA | GLUD1 | glutamate dehydrogenase 1 | 2746 | ENSG00000148672 |
| 10 | Q06323 | NA | PSME1 | proteasome (prosome, macropain) activator subunit 1 (PA28 alpha) | 5720 | ENSG00000092010 |
| 11 | P49419 | NA | ALDH7A1 | aldehyde dehydrogenase 7 family, member A1 | 501 | ENSG00000164904 |
| 12 | P07602 | NA | PSAP | prosaposin | 5660 | ENSG00000197746 |
| 13 | P09211 | NA | GSTP1 | glutathione S-transferase pi 1 | 2950 | ENSG00000084207 |
| 14 | P23381 | NA | WARS | tryptophanyl-tRNA synthetase | 7453 | ENSG00000140105 |

  
  

| **Database:molecular function      &nbspName:oxidoreductase activity      &nbspID:GO:0016491** | | | | | | |
| --- | --- | --- | --- | --- | --- | --- |
| C=697; O=13; E=1.77; R=7.35; rawP=7.92e-09; adjP=3.96e-07 | | | | | | |
| Index | UserID | Value | Gene Symbol | Gene Name | EntrezGene | Ensembl |
| 1 | P04040 | NA | CAT | catalase | 847 | ENSG00000121691 |
| 2 | Q9UBQ7 | NA | GRHPR | glyoxylate reductase/hydroxypyruvate reductase | 9380 | ENSG00000137106 |
| 3 | P02792 | NA | FTL | ferritin, light polypeptide | 2512 | ENSG00000087086 |
| 4 | P51649 | NA | ALDH5A1 | aldehyde dehydrogenase 5 family, member A1 | 7915 | ENSG00000112294 |
| 5 | P30044 | NA | PRDX5 | peroxiredoxin 5 | 25824 | ENSG00000126432 |
| 6 | P22570 | NA | FDXR | ferredoxin reductase | 2232 | ENSG00000161513 |
| 7 | P00338 | NA | LDHA | lactate dehydrogenase A | 3939 | ENSG00000134333 |
| 8 | P13674 | NA | P4HA1 | prolyl 4-hydroxylase, alpha polypeptide I | 5033 | ENSG00000122884 |
| 9 | P00367 | NA | GLUD1 | glutamate dehydrogenase 1 | 2746 | ENSG00000148672 |
| 10 | P48735 | NA | IDH2 | isocitrate dehydrogenase 2 (NADP+), mitochondrial | 3418 | ENSG00000182054 |
| 11 | P49419 | NA | ALDH7A1 | aldehyde dehydrogenase 7 family, member A1 | 501 | ENSG00000164904 |
| 12 | P49748 | NA | ACADVL | acyl-CoA dehydrogenase, very long chain | 37 | ENSG00000072778 |
| 13 | Q9P0J0 | NA | NDUFA13 | NADH dehydrogenase (ubiquinone) 1 alpha subcomplex, 13 | 51079 | ENSG00000186010 |

  
  

| **Database:molecular function      &nbspName:catalytic activity      &nbspID:GO:0003824** | | | | | | |
| --- | --- | --- | --- | --- | --- | --- |
| C=5299; O=29; E=13.44; R=2.16; rawP=3.02e-07; adjP=7.55e-06 | | | | | | |
| Index | UserID | Value | Gene Symbol | Gene Name | EntrezGene | Ensembl |
| 1 | P02792 | NA | FTL | ferritin, light polypeptide | 2512 | ENSG00000087086 |
| 2 | P10253 | NA | GAA | glucosidase, alpha; acid | 2548 | ENSG00000171298 |
| 3 | Q71U36 | NA | TUBA1A | tubulin, alpha 1a | 7846 | ENSG00000167552 |
| 4 | Q16851 | NA | UGP2 | UDP-glucose pyrophosphorylase 2 | 7360 | ENSG00000169764 |
| 5 | P00338 | NA | LDHA | lactate dehydrogenase A | 3939 | ENSG00000134333 |
| 6 | P00367 | NA | GLUD1 | glutamate dehydrogenase 1 | 2746 | ENSG00000148672 |
| 7 | P48735 | NA | IDH2 | isocitrate dehydrogenase 2 (NADP+), mitochondrial | 3418 | ENSG00000182054 |
| 8 | P38571 | NA | LIPA | lipase A, lysosomal acid, cholesterol esterase | 3988 | ENSG00000107798 |
| 9 | P09211 | NA | GSTP1 | glutathione S-transferase pi 1 | 2950 | ENSG00000084207 |
| 10 | Q9P0J0 | NA | NDUFA13 | NADH dehydrogenase (ubiquinone) 1 alpha subcomplex, 13 | 51079 | ENSG00000186010 |
| 11 | Q13011 | NA | ECH1 | enoyl CoA hydratase 1, peroxisomal | 1891 | ENSG00000104823 |
| 12 | P23381 | NA | WARS | tryptophanyl-tRNA synthetase | 7453 | ENSG00000140105 |
| 13 | P04040 | NA | CAT | catalase | 847 | ENSG00000121691 |
| 14 | O14773 | NA | TPP1 | tripeptidyl peptidase I | 1200 | ENSG00000166340 |
| 15 | Q9UBQ7 | NA | GRHPR | glyoxylate reductase/hydroxypyruvate reductase | 9380 | ENSG00000137106 |
| 16 | P51649 | NA | ALDH5A1 | aldehyde dehydrogenase 5 family, member A1 | 7915 | ENSG00000112294 |
| 17 | P30044 | NA | PRDX5 | peroxiredoxin 5 | 25824 | ENSG00000126432 |
| 18 | Q05639 | NA | EEF1A2 | eukaryotic translation elongation factor 1 alpha 2 | 1917 | ENSG00000101210 |
| 19 | P22570 | NA | FDXR | ferredoxin reductase | 2232 | ENSG00000161513 |
| 20 | P42765 | NA | ACAA2 | acetyl-CoA acyltransferase 2 | 10449 | ENSG00000167315 |
| 21 | P13674 | NA | P4HA1 | prolyl 4-hydroxylase, alpha polypeptide I | 5033 | ENSG00000122884 |
| 22 | P24752 | NA | ACAT1 | acetyl-CoA acetyltransferase 1 | 38 | ENSG00000075239 |
| 23 | P16278 | NA | GLB1 | galactosidase, beta 1 | 2720 | ENSG00000170266 |
| 24 | P49419 | NA | ALDH7A1 | aldehyde dehydrogenase 7 family, member A1 | 501 | ENSG00000164904 |
| 25 | P06744 | NA | GPI | glucose-6-phosphate isomerase | 2821 | ENSG00000105220 |
| 26 | P07339 | NA | CTSD | cathepsin D | 1509 | ENSG00000117984 |
| 27 | P49748 | NA | ACADVL | acyl-CoA dehydrogenase, very long chain | 37 | ENSG00000072778 |
| 28 | Q14697 | NA | GANAB | glucosidase, alpha; neutral AB | 23193 | ENSG00000089597 |
| 29 | P07384 | NA | CAPN1 | calpain 1, (mu/I) large subunit | 823 | ENSG00000014216 |

  
  

| **Database:molecular function      &nbspName:coenzyme binding      &nbspID:GO:0050662** | | | | | | |
| --- | --- | --- | --- | --- | --- | --- |
| C=184; O=6; E=0.47; R=12.85; rawP=6.31e-06; adjP=0.0001 | | | | | | |
| Index | UserID | Value | Gene Symbol | Gene Name | EntrezGene | Ensembl |
| 1 | P24752 | NA | ACAT1 | acetyl-CoA acetyltransferase 1 | 38 | ENSG00000075239 |
| 2 | P04040 | NA | CAT | catalase | 847 | ENSG00000121691 |
| 3 | Q9UBQ7 | NA | GRHPR | glyoxylate reductase/hydroxypyruvate reductase | 9380 | ENSG00000137106 |
| 4 | P48735 | NA | IDH2 | isocitrate dehydrogenase 2 (NADP+), mitochondrial | 3418 | ENSG00000182054 |
| 5 | P00367 | NA | GLUD1 | glutamate dehydrogenase 1 | 2746 | ENSG00000148672 |
| 6 | P49748 | NA | ACADVL | acyl-CoA dehydrogenase, very long chain | 37 | ENSG00000072778 |

  
  

| **Database:molecular function      &nbspName:carbohydrate binding      &nbspID:GO:0030246** | | | | | | |
| --- | --- | --- | --- | --- | --- | --- |
| C=267; O=6; E=0.68; R=8.86; rawP=5.19e-05; adjP=0.0005 | | | | | | |
| Index | UserID | Value | Gene Symbol | Gene Name | EntrezGene | Ensembl |
| 1 | P13674 | NA | P4HA1 | prolyl 4-hydroxylase, alpha polypeptide I | 5033 | ENSG00000122884 |
| 2 | P10253 | NA | GAA | glucosidase, alpha; acid | 2548 | ENSG00000171298 |
| 3 | P06744 | NA | GPI | glucose-6-phosphate isomerase | 2821 | ENSG00000105220 |
| 4 | P17931 | NA | LGALS3 | lectin, galactoside-binding, soluble, 3 | 3958 | ENSG00000131981 |
| 5 | Q16851 | NA | UGP2 | UDP-glucose pyrophosphorylase 2 | 7360 | ENSG00000169764 |
| 6 | Q14697 | NA | GANAB | glucosidase, alpha; neutral AB | 23193 | ENSG00000089597 |

  
  

| **Database:molecular function      &nbspName:cofactor binding      &nbspID:GO:0048037** | | | | | | |
| --- | --- | --- | --- | --- | --- | --- |
| C=259; O=6; E=0.66; R=9.13; rawP=4.38e-05; adjP=0.0005 | | | | | | |
| Index | UserID | Value | Gene Symbol | Gene Name | EntrezGene | Ensembl |
| 1 | P24752 | NA | ACAT1 | acetyl-CoA acetyltransferase 1 | 38 | ENSG00000075239 |
| 2 | P04040 | NA | CAT | catalase | 847 | ENSG00000121691 |
| 3 | Q9UBQ7 | NA | GRHPR | glyoxylate reductase/hydroxypyruvate reductase | 9380 | ENSG00000137106 |
| 4 | P48735 | NA | IDH2 | isocitrate dehydrogenase 2 (NADP+), mitochondrial | 3418 | ENSG00000182054 |
| 5 | P00367 | NA | GLUD1 | glutamate dehydrogenase 1 | 2746 | ENSG00000148672 |
| 6 | P49748 | NA | ACADVL | acyl-CoA dehydrogenase, very long chain | 37 | ENSG00000072778 |

  
  

| **Database:cellular component      &nbspName:cytoplasmic part      &nbspID:GO:0044444** | | | | | | |
| --- | --- | --- | --- | --- | --- | --- |
| C=6728; O=38; E=16.13; R=2.36; rawP=2.80e-13; adjP=1.99e-11 | | | | | | |
| Index | UserID | Value | Gene Symbol | Gene Name | EntrezGene | Ensembl |
| 1 | P10253 | NA | GAA | glucosidase, alpha; acid | 2548 | ENSG00000171298 |
| 2 | P17931 | NA | LGALS3 | lectin, galactoside-binding, soluble, 3 | 3958 | ENSG00000131981 |
| 3 | P08133 | NA | ANXA6 | annexin A6 | 309 | ENSG00000197043 |
| 4 | Q16851 | NA | UGP2 | UDP-glucose pyrophosphorylase 2 | 7360 | ENSG00000169764 |
| 5 | P30086 | NA | PEBP1 | phosphatidylethanolamine binding protein 1 | 5037 | ENSG00000089220 |
| 6 | P00367 | NA | GLUD1 | glutamate dehydrogenase 1 | 2746 | ENSG00000148672 |
| 7 | Q06323 | NA | PSME1 | proteasome (prosome, macropain) activator subunit 1 (PA28 alpha) | 5720 | ENSG00000092010 |
| 8 | P48735 | NA | IDH2 | isocitrate dehydrogenase 2 (NADP+), mitochondrial | 3418 | ENSG00000182054 |
| 9 | Q15019 | NA | SEPT2 | septin 2 | 4735 | ENSG00000168385 |
| 10 | Q9P0J0 | NA | NDUFA13 | NADH dehydrogenase (ubiquinone) 1 alpha subcomplex, 13 | 51079 | ENSG00000186010 |
| 11 | P09211 | NA | GSTP1 | glutathione S-transferase pi 1 | 2950 | ENSG00000084207 |
| 12 | O14773 | NA | TPP1 | tripeptidyl peptidase I | 1200 | ENSG00000166340 |
| 13 | Q05639 | NA | EEF1A2 | eukaryotic translation elongation factor 1 alpha 2 | 1917 | ENSG00000101210 |
| 14 | P22570 | NA | FDXR | ferredoxin reductase | 2232 | ENSG00000161513 |
| 15 | P40121 | NA | CAPG | capping protein (actin filament), gelsolin-like | 822 | ENSG00000042493 |
| 16 | P42765 | NA | ACAA2 | acetyl-CoA acyltransferase 2 | 10449 | ENSG00000167315 |
| 17 | P49419 | NA | ALDH7A1 | aldehyde dehydrogenase 7 family, member A1 | 501 | ENSG00000164904 |
| 18 | P07339 | NA | CTSD | cathepsin D | 1509 | ENSG00000117984 |
| 19 | Q14697 | NA | GANAB | glucosidase, alpha; neutral AB | 23193 | ENSG00000089597 |
| 20 | P53007 | NA | SLC25A1 | solute carrier family 25 (mitochondrial carrier; citrate transporter), member 1 | 6576 | ENSG00000100075 |
| 21 | Q9BPW8 | NA | NIPSNAP1 | nipsnap homolog 1 (C. elegans) | 8508 | ENSG00000184117 |
| 22 | P02792 | NA | FTL | ferritin, light polypeptide | 2512 | ENSG00000087086 |
| 23 | Q71U36 | NA | TUBA1A | tubulin, alpha 1a | 7846 | ENSG00000167552 |
| 24 | P00338 | NA | LDHA | lactate dehydrogenase A | 3939 | ENSG00000134333 |
| 25 | P38571 | NA | LIPA | lipase A, lysosomal acid, cholesterol esterase | 3988 | ENSG00000107798 |
| 26 | Q13011 | NA | ECH1 | enoyl CoA hydratase 1, peroxisomal | 1891 | ENSG00000104823 |
| 27 | P23381 | NA | WARS | tryptophanyl-tRNA synthetase | 7453 | ENSG00000140105 |
| 28 | Q14980 | NA | NUMA1 | nuclear mitotic apparatus protein 1 | 4926 | ENSG00000137497 |
| 29 | P04040 | NA | CAT | catalase | 847 | ENSG00000121691 |
| 30 | Q9UBQ7 | NA | GRHPR | glyoxylate reductase/hydroxypyruvate reductase | 9380 | ENSG00000137106 |
| 31 | P51649 | NA | ALDH5A1 | aldehyde dehydrogenase 5 family, member A1 | 7915 | ENSG00000112294 |
| 32 | P30044 | NA | PRDX5 | peroxiredoxin 5 | 25824 | ENSG00000126432 |
| 33 | P13674 | NA | P4HA1 | prolyl 4-hydroxylase, alpha polypeptide I | 5033 | ENSG00000122884 |
| 34 | P24752 | NA | ACAT1 | acetyl-CoA acetyltransferase 1 | 38 | ENSG00000075239 |
| 35 | P16278 | NA | GLB1 | galactosidase, beta 1 | 2720 | ENSG00000170266 |
| 36 | P06744 | NA | GPI | glucose-6-phosphate isomerase | 2821 | ENSG00000105220 |
| 37 | P07602 | NA | PSAP | prosaposin | 5660 | ENSG00000197746 |
| 38 | P49748 | NA | ACADVL | acyl-CoA dehydrogenase, very long chain | 37 | ENSG00000072778 |

  
  

| **Database:cellular component      &nbspName:mitochondrion      &nbspID:GO:0005739** | | | | | | |
| --- | --- | --- | --- | --- | --- | --- |
| C=1515; O=21; E=3.63; R=5.78; rawP=2.80e-12; adjP=9.94e-11 | | | | | | |
| Index | UserID | Value | Gene Symbol | Gene Name | EntrezGene | Ensembl |
| 1 | P17931 | NA | LGALS3 | lectin, galactoside-binding, soluble, 3 | 3958 | ENSG00000131981 |
| 2 | P30086 | NA | PEBP1 | phosphatidylethanolamine binding protein 1 | 5037 | ENSG00000089220 |
| 3 | P00338 | NA | LDHA | lactate dehydrogenase A | 3939 | ENSG00000134333 |
| 4 | P00367 | NA | GLUD1 | glutamate dehydrogenase 1 | 2746 | ENSG00000148672 |
| 5 | P48735 | NA | IDH2 | isocitrate dehydrogenase 2 (NADP+), mitochondrial | 3418 | ENSG00000182054 |
| 6 | P09211 | NA | GSTP1 | glutathione S-transferase pi 1 | 2950 | ENSG00000084207 |
| 7 | Q9P0J0 | NA | NDUFA13 | NADH dehydrogenase (ubiquinone) 1 alpha subcomplex, 13 | 51079 | ENSG00000186010 |
| 8 | Q13011 | NA | ECH1 | enoyl CoA hydratase 1, peroxisomal | 1891 | ENSG00000104823 |
| 9 | O14773 | NA | TPP1 | tripeptidyl peptidase I | 1200 | ENSG00000166340 |
| 10 | P04040 | NA | CAT | catalase | 847 | ENSG00000121691 |
| 11 | P51649 | NA | ALDH5A1 | aldehyde dehydrogenase 5 family, member A1 | 7915 | ENSG00000112294 |
| 12 | P22570 | NA | FDXR | ferredoxin reductase | 2232 | ENSG00000161513 |
| 13 | P30044 | NA | PRDX5 | peroxiredoxin 5 | 25824 | ENSG00000126432 |
| 14 | P42765 | NA | ACAA2 | acetyl-CoA acyltransferase 2 | 10449 | ENSG00000167315 |
| 15 | P13674 | NA | P4HA1 | prolyl 4-hydroxylase, alpha polypeptide I | 5033 | ENSG00000122884 |
| 16 | P24752 | NA | ACAT1 | acetyl-CoA acetyltransferase 1 | 38 | ENSG00000075239 |
| 17 | P49419 | NA | ALDH7A1 | aldehyde dehydrogenase 7 family, member A1 | 501 | ENSG00000164904 |
| 18 | P07602 | NA | PSAP | prosaposin | 5660 | ENSG00000197746 |
| 19 | P49748 | NA | ACADVL | acyl-CoA dehydrogenase, very long chain | 37 | ENSG00000072778 |
| 20 | Q9BPW8 | NA | NIPSNAP1 | nipsnap homolog 1 (C. elegans) | 8508 | ENSG00000184117 |
| 21 | P53007 | NA | SLC25A1 | solute carrier family 25 (mitochondrial carrier; citrate transporter), member 1 | 6576 | ENSG00000100075 |

  
  

| **Database:cellular component      &nbspName:cytoplasm      &nbspID:GO:0005737** | | | | | | |
| --- | --- | --- | --- | --- | --- | --- |
| C=9051; O=40; E=21.70; R=1.84; rawP=2.29e-11; adjP=5.42e-10 | | | | | | |
| Index | UserID | Value | Gene Symbol | Gene Name | EntrezGene | Ensembl |
| 1 | P10253 | NA | GAA | glucosidase, alpha; acid | 2548 | ENSG00000171298 |
| 2 | P17931 | NA | LGALS3 | lectin, galactoside-binding, soluble, 3 | 3958 | ENSG00000131981 |
| 3 | P08133 | NA | ANXA6 | annexin A6 | 309 | ENSG00000197043 |
| 4 | Q16851 | NA | UGP2 | UDP-glucose pyrophosphorylase 2 | 7360 | ENSG00000169764 |
| 5 | P30086 | NA | PEBP1 | phosphatidylethanolamine binding protein 1 | 5037 | ENSG00000089220 |
| 6 | P00367 | NA | GLUD1 | glutamate dehydrogenase 1 | 2746 | ENSG00000148672 |
| 7 | Q06323 | NA | PSME1 | proteasome (prosome, macropain) activator subunit 1 (PA28 alpha) | 5720 | ENSG00000092010 |
| 8 | P48735 | NA | IDH2 | isocitrate dehydrogenase 2 (NADP+), mitochondrial | 3418 | ENSG00000182054 |
| 9 | Q15019 | NA | SEPT2 | septin 2 | 4735 | ENSG00000168385 |
| 10 | Q9P0J0 | NA | NDUFA13 | NADH dehydrogenase (ubiquinone) 1 alpha subcomplex, 13 | 51079 | ENSG00000186010 |
| 11 | P09211 | NA | GSTP1 | glutathione S-transferase pi 1 | 2950 | ENSG00000084207 |
| 12 | O14773 | NA | TPP1 | tripeptidyl peptidase I | 1200 | ENSG00000166340 |
| 13 | Q05639 | NA | EEF1A2 | eukaryotic translation elongation factor 1 alpha 2 | 1917 | ENSG00000101210 |
| 14 | P22570 | NA | FDXR | ferredoxin reductase | 2232 | ENSG00000161513 |
| 15 | P40121 | NA | CAPG | capping protein (actin filament), gelsolin-like | 822 | ENSG00000042493 |
| 16 | P42765 | NA | ACAA2 | acetyl-CoA acyltransferase 2 | 10449 | ENSG00000167315 |
| 17 | P49419 | NA | ALDH7A1 | aldehyde dehydrogenase 7 family, member A1 | 501 | ENSG00000164904 |
| 18 | P07339 | NA | CTSD | cathepsin D | 1509 | ENSG00000117984 |
| 19 | Q14764 | NA | MVP | major vault protein | 9961 | ENSG00000013364 |
| 20 | Q14697 | NA | GANAB | glucosidase, alpha; neutral AB | 23193 | ENSG00000089597 |
| 21 | P53007 | NA | SLC25A1 | solute carrier family 25 (mitochondrial carrier; citrate transporter), member 1 | 6576 | ENSG00000100075 |
| 22 | Q9BPW8 | NA | NIPSNAP1 | nipsnap homolog 1 (C. elegans) | 8508 | ENSG00000184117 |
| 23 | P02792 | NA | FTL | ferritin, light polypeptide | 2512 | ENSG00000087086 |
| 24 | Q71U36 | NA | TUBA1A | tubulin, alpha 1a | 7846 | ENSG00000167552 |
| 25 | P00338 | NA | LDHA | lactate dehydrogenase A | 3939 | ENSG00000134333 |
| 26 | P38571 | NA | LIPA | lipase A, lysosomal acid, cholesterol esterase | 3988 | ENSG00000107798 |
| 27 | Q13011 | NA | ECH1 | enoyl CoA hydratase 1, peroxisomal | 1891 | ENSG00000104823 |
| 28 | P23381 | NA | WARS | tryptophanyl-tRNA synthetase | 7453 | ENSG00000140105 |
| 29 | Q14980 | NA | NUMA1 | nuclear mitotic apparatus protein 1 | 4926 | ENSG00000137497 |
| 30 | P04040 | NA | CAT | catalase | 847 | ENSG00000121691 |
| 31 | Q9UBQ7 | NA | GRHPR | glyoxylate reductase/hydroxypyruvate reductase | 9380 | ENSG00000137106 |
| 32 | P51649 | NA | ALDH5A1 | aldehyde dehydrogenase 5 family, member A1 | 7915 | ENSG00000112294 |
| 33 | P30044 | NA | PRDX5 | peroxiredoxin 5 | 25824 | ENSG00000126432 |
| 34 | P13674 | NA | P4HA1 | prolyl 4-hydroxylase, alpha polypeptide I | 5033 | ENSG00000122884 |
| 35 | P24752 | NA | ACAT1 | acetyl-CoA acetyltransferase 1 | 38 | ENSG00000075239 |
| 36 | P16278 | NA | GLB1 | galactosidase, beta 1 | 2720 | ENSG00000170266 |
| 37 | P06744 | NA | GPI | glucose-6-phosphate isomerase | 2821 | ENSG00000105220 |
| 38 | P07602 | NA | PSAP | prosaposin | 5660 | ENSG00000197746 |
| 39 | P49748 | NA | ACADVL | acyl-CoA dehydrogenase, very long chain | 37 | ENSG00000072778 |
| 40 | P07384 | NA | CAPN1 | calpain 1, (mu/I) large subunit | 823 | ENSG00000014216 |

  
  

| **Database:cellular component      &nbspName:intracellular organelle lumen      &nbspID:GO:0070013** | | | | | | |
| --- | --- | --- | --- | --- | --- | --- |
| C=3285; O=22; E=7.88; R=2.79; rawP=7.75e-07; adjP=6.88e-06 | | | | | | |
| Index | UserID | Value | Gene Symbol | Gene Name | EntrezGene | Ensembl |
| 1 | P00367 | NA | GLUD1 | glutamate dehydrogenase 1 | 2746 | ENSG00000148672 |
| 2 | Q06323 | NA | PSME1 | proteasome (prosome, macropain) activator subunit 1 (PA28 alpha) | 5720 | ENSG00000092010 |
| 3 | P48735 | NA | IDH2 | isocitrate dehydrogenase 2 (NADP+), mitochondrial | 3418 | ENSG00000182054 |
| 4 | Q15019 | NA | SEPT2 | septin 2 | 4735 | ENSG00000168385 |
| 5 | Q9P0J0 | NA | NDUFA13 | NADH dehydrogenase (ubiquinone) 1 alpha subcomplex, 13 | 51079 | ENSG00000186010 |
| 6 | O14773 | NA | TPP1 | tripeptidyl peptidase I | 1200 | ENSG00000166340 |
| 7 | Q14980 | NA | NUMA1 | nuclear mitotic apparatus protein 1 | 4926 | ENSG00000137497 |
| 8 | P04040 | NA | CAT | catalase | 847 | ENSG00000121691 |
| 9 | Q9UBQ7 | NA | GRHPR | glyoxylate reductase/hydroxypyruvate reductase | 9380 | ENSG00000137106 |
| 10 | P51649 | NA | ALDH5A1 | aldehyde dehydrogenase 5 family, member A1 | 7915 | ENSG00000112294 |
| 11 | P22570 | NA | FDXR | ferredoxin reductase | 2232 | ENSG00000161513 |
| 12 | P30044 | NA | PRDX5 | peroxiredoxin 5 | 25824 | ENSG00000126432 |
| 13 | P40121 | NA | CAPG | capping protein (actin filament), gelsolin-like | 822 | ENSG00000042493 |
| 14 | P13674 | NA | P4HA1 | prolyl 4-hydroxylase, alpha polypeptide I | 5033 | ENSG00000122884 |
| 15 | P24752 | NA | ACAT1 | acetyl-CoA acetyltransferase 1 | 38 | ENSG00000075239 |
| 16 | P16278 | NA | GLB1 | galactosidase, beta 1 | 2720 | ENSG00000170266 |
| 17 | P49419 | NA | ALDH7A1 | aldehyde dehydrogenase 7 family, member A1 | 501 | ENSG00000164904 |
| 18 | P06744 | NA | GPI | glucose-6-phosphate isomerase | 2821 | ENSG00000105220 |
| 19 | P07602 | NA | PSAP | prosaposin | 5660 | ENSG00000197746 |
| 20 | P07339 | NA | CTSD | cathepsin D | 1509 | ENSG00000117984 |
| 21 | P49748 | NA | ACADVL | acyl-CoA dehydrogenase, very long chain | 37 | ENSG00000072778 |
| 22 | Q14697 | NA | GANAB | glucosidase, alpha; neutral AB | 23193 | ENSG00000089597 |

  
  

| **Database:cellular component      &nbspName:organelle lumen      &nbspID:GO:0043233** | | | | | | |
| --- | --- | --- | --- | --- | --- | --- |
| C=3331; O=22; E=7.99; R=2.75; rawP=9.94e-07; adjP=7.84e-06 | | | | | | |
| Index | UserID | Value | Gene Symbol | Gene Name | EntrezGene | Ensembl |
| 1 | P00367 | NA | GLUD1 | glutamate dehydrogenase 1 | 2746 | ENSG00000148672 |
| 2 | Q06323 | NA | PSME1 | proteasome (prosome, macropain) activator subunit 1 (PA28 alpha) | 5720 | ENSG00000092010 |
| 3 | P48735 | NA | IDH2 | isocitrate dehydrogenase 2 (NADP+), mitochondrial | 3418 | ENSG00000182054 |
| 4 | Q15019 | NA | SEPT2 | septin 2 | 4735 | ENSG00000168385 |
| 5 | Q9P0J0 | NA | NDUFA13 | NADH dehydrogenase (ubiquinone) 1 alpha subcomplex, 13 | 51079 | ENSG00000186010 |
| 6 | O14773 | NA | TPP1 | tripeptidyl peptidase I | 1200 | ENSG00000166340 |
| 7 | Q14980 | NA | NUMA1 | nuclear mitotic apparatus protein 1 | 4926 | ENSG00000137497 |
| 8 | P04040 | NA | CAT | catalase | 847 | ENSG00000121691 |
| 9 | Q9UBQ7 | NA | GRHPR | glyoxylate reductase/hydroxypyruvate reductase | 9380 | ENSG00000137106 |
| 10 | P51649 | NA | ALDH5A1 | aldehyde dehydrogenase 5 family, member A1 | 7915 | ENSG00000112294 |
| 11 | P22570 | NA | FDXR | ferredoxin reductase | 2232 | ENSG00000161513 |
| 12 | P30044 | NA | PRDX5 | peroxiredoxin 5 | 25824 | ENSG00000126432 |
| 13 | P40121 | NA | CAPG | capping protein (actin filament), gelsolin-like | 822 | ENSG00000042493 |
| 14 | P13674 | NA | P4HA1 | prolyl 4-hydroxylase, alpha polypeptide I | 5033 | ENSG00000122884 |
| 15 | P24752 | NA | ACAT1 | acetyl-CoA acetyltransferase 1 | 38 | ENSG00000075239 |
| 16 | P16278 | NA | GLB1 | galactosidase, beta 1 | 2720 | ENSG00000170266 |
| 17 | P49419 | NA | ALDH7A1 | aldehyde dehydrogenase 7 family, member A1 | 501 | ENSG00000164904 |
| 18 | P06744 | NA | GPI | glucose-6-phosphate isomerase | 2821 | ENSG00000105220 |
| 19 | P07602 | NA | PSAP | prosaposin | 5660 | ENSG00000197746 |
| 20 | P07339 | NA | CTSD | cathepsin D | 1509 | ENSG00000117984 |
| 21 | P49748 | NA | ACADVL | acyl-CoA dehydrogenase, very long chain | 37 | ENSG00000072778 |
| 22 | Q14697 | NA | GANAB | glucosidase, alpha; neutral AB | 23193 | ENSG00000089597 |

  
  

| **Database:cellular component      &nbspName:membrane-enclosed lumen      &nbspID:GO:0031974** | | | | | | |
| --- | --- | --- | --- | --- | --- | --- |
| C=3375; O=22; E=8.09; R=2.72; rawP=1.26e-06; adjP=8.95e-06 | | | | | | |
| Index | UserID | Value | Gene Symbol | Gene Name | EntrezGene | Ensembl |
| 1 | P00367 | NA | GLUD1 | glutamate dehydrogenase 1 | 2746 | ENSG00000148672 |
| 2 | Q06323 | NA | PSME1 | proteasome (prosome, macropain) activator subunit 1 (PA28 alpha) | 5720 | ENSG00000092010 |
| 3 | P48735 | NA | IDH2 | isocitrate dehydrogenase 2 (NADP+), mitochondrial | 3418 | ENSG00000182054 |
| 4 | Q15019 | NA | SEPT2 | septin 2 | 4735 | ENSG00000168385 |
| 5 | Q9P0J0 | NA | NDUFA13 | NADH dehydrogenase (ubiquinone) 1 alpha subcomplex, 13 | 51079 | ENSG00000186010 |
| 6 | O14773 | NA | TPP1 | tripeptidyl peptidase I | 1200 | ENSG00000166340 |
| 7 | Q14980 | NA | NUMA1 | nuclear mitotic apparatus protein 1 | 4926 | ENSG00000137497 |
| 8 | P04040 | NA | CAT | catalase | 847 | ENSG00000121691 |
| 9 | Q9UBQ7 | NA | GRHPR | glyoxylate reductase/hydroxypyruvate reductase | 9380 | ENSG00000137106 |
| 10 | P51649 | NA | ALDH5A1 | aldehyde dehydrogenase 5 family, member A1 | 7915 | ENSG00000112294 |
| 11 | P22570 | NA | FDXR | ferredoxin reductase | 2232 | ENSG00000161513 |
| 12 | P30044 | NA | PRDX5 | peroxiredoxin 5 | 25824 | ENSG00000126432 |
| 13 | P40121 | NA | CAPG | capping protein (actin filament), gelsolin-like | 822 | ENSG00000042493 |
| 14 | P13674 | NA | P4HA1 | prolyl 4-hydroxylase, alpha polypeptide I | 5033 | ENSG00000122884 |
| 15 | P24752 | NA | ACAT1 | acetyl-CoA acetyltransferase 1 | 38 | ENSG00000075239 |
| 16 | P16278 | NA | GLB1 | galactosidase, beta 1 | 2720 | ENSG00000170266 |
| 17 | P49419 | NA | ALDH7A1 | aldehyde dehydrogenase 7 family, member A1 | 501 | ENSG00000164904 |
| 18 | P06744 | NA | GPI | glucose-6-phosphate isomerase | 2821 | ENSG00000105220 |
| 19 | P07602 | NA | PSAP | prosaposin | 5660 | ENSG00000197746 |
| 20 | P07339 | NA | CTSD | cathepsin D | 1509 | ENSG00000117984 |
| 21 | P49748 | NA | ACADVL | acyl-CoA dehydrogenase, very long chain | 37 | ENSG00000072778 |
| 22 | Q14697 | NA | GANAB | glucosidase, alpha; neutral AB | 23193 | ENSG00000089597 |

  
  

| **Database:cellular component      &nbspName:mitochondrial inner membrane      &nbspID:GO:0005743** | | | | | | |
| --- | --- | --- | --- | --- | --- | --- |
| C=351; O=8; E=0.84; R=9.51; rawP=1.52e-06; adjP=9.81e-06 | | | | | | |
| Index | UserID | Value | Gene Symbol | Gene Name | EntrezGene | Ensembl |
| 1 | P17931 | NA | LGALS3 | lectin, galactoside-binding, soluble, 3 | 3958 | ENSG00000131981 |
| 2 | P42765 | NA | ACAA2 | acetyl-CoA acyltransferase 2 | 10449 | ENSG00000167315 |
| 3 | P24752 | NA | ACAT1 | acetyl-CoA acetyltransferase 1 | 38 | ENSG00000075239 |
| 4 | P48735 | NA | IDH2 | isocitrate dehydrogenase 2 (NADP+), mitochondrial | 3418 | ENSG00000182054 |
| 5 | Q9P0J0 | NA | NDUFA13 | NADH dehydrogenase (ubiquinone) 1 alpha subcomplex, 13 | 51079 | ENSG00000186010 |
| 6 | P49748 | NA | ACADVL | acyl-CoA dehydrogenase, very long chain | 37 | ENSG00000072778 |
| 7 | Q9BPW8 | NA | NIPSNAP1 | nipsnap homolog 1 (C. elegans) | 8508 | ENSG00000184117 |
| 8 | P53007 | NA | SLC25A1 | solute carrier family 25 (mitochondrial carrier; citrate transporter), member 1 | 6576 | ENSG00000100075 |

  
  

| **Database:cellular component      &nbspName:mitochondrial membrane      &nbspID:GO:0031966** | | | | | | |
| --- | --- | --- | --- | --- | --- | --- |
| C=505; O=9; E=1.21; R=7.43; rawP=2.35e-06; adjP=1.24e-05 | | | | | | |
| Index | UserID | Value | Gene Symbol | Gene Name | EntrezGene | Ensembl |
| 1 | P17931 | NA | LGALS3 | lectin, galactoside-binding, soluble, 3 | 3958 | ENSG00000131981 |
| 2 | P30086 | NA | PEBP1 | phosphatidylethanolamine binding protein 1 | 5037 | ENSG00000089220 |
| 3 | P42765 | NA | ACAA2 | acetyl-CoA acyltransferase 2 | 10449 | ENSG00000167315 |
| 4 | P24752 | NA | ACAT1 | acetyl-CoA acetyltransferase 1 | 38 | ENSG00000075239 |
| 5 | P48735 | NA | IDH2 | isocitrate dehydrogenase 2 (NADP+), mitochondrial | 3418 | ENSG00000182054 |
| 6 | Q9P0J0 | NA | NDUFA13 | NADH dehydrogenase (ubiquinone) 1 alpha subcomplex, 13 | 51079 | ENSG00000186010 |
| 7 | P49748 | NA | ACADVL | acyl-CoA dehydrogenase, very long chain | 37 | ENSG00000072778 |
| 8 | Q9BPW8 | NA | NIPSNAP1 | nipsnap homolog 1 (C. elegans) | 8508 | ENSG00000184117 |
| 9 | P53007 | NA | SLC25A1 | solute carrier family 25 (mitochondrial carrier; citrate transporter), member 1 | 6576 | ENSG00000100075 |

  
  

| **Database:cellular component      &nbspName:pigment granule      &nbspID:GO:0048770** | | | | | | |
| --- | --- | --- | --- | --- | --- | --- |
| C=93; O=5; E=0.22; R=22.42; rawP=2.72e-06; adjP=1.24e-05 | | | | | | |
| Index | UserID | Value | Gene Symbol | Gene Name | EntrezGene | Ensembl |
| 1 | O14773 | NA | TPP1 | tripeptidyl peptidase I | 1200 | ENSG00000166340 |
| 2 | P07339 | NA | CTSD | cathepsin D | 1509 | ENSG00000117984 |
| 3 | P08133 | NA | ANXA6 | annexin A6 | 309 | ENSG00000197043 |
| 4 | Q14697 | NA | GANAB | glucosidase, alpha; neutral AB | 23193 | ENSG00000089597 |
| 5 | P40121 | NA | CAPG | capping protein (actin filament), gelsolin-like | 822 | ENSG00000042493 |

  
  

| **Database:cellular component      &nbspName:organelle inner membrane      &nbspID:GO:0019866** | | | | | | |
| --- | --- | --- | --- | --- | --- | --- |
| C=381; O=8; E=0.91; R=8.76; rawP=2.79e-06; adjP=1.24e-05 | | | | | | |
| Index | UserID | Value | Gene Symbol | Gene Name | EntrezGene | Ensembl |
| 1 | P17931 | NA | LGALS3 | lectin, galactoside-binding, soluble, 3 | 3958 | ENSG00000131981 |
| 2 | P42765 | NA | ACAA2 | acetyl-CoA acyltransferase 2 | 10449 | ENSG00000167315 |
| 3 | P24752 | NA | ACAT1 | acetyl-CoA acetyltransferase 1 | 38 | ENSG00000075239 |
| 4 | P48735 | NA | IDH2 | isocitrate dehydrogenase 2 (NADP+), mitochondrial | 3418 | ENSG00000182054 |
| 5 | Q9P0J0 | NA | NDUFA13 | NADH dehydrogenase (ubiquinone) 1 alpha subcomplex, 13 | 51079 | ENSG00000186010 |
| 6 | P49748 | NA | ACADVL | acyl-CoA dehydrogenase, very long chain | 37 | ENSG00000072778 |
| 7 | Q9BPW8 | NA | NIPSNAP1 | nipsnap homolog 1 (C. elegans) | 8508 | ENSG00000184117 |
| 8 | P53007 | NA | SLC25A1 | solute carrier family 25 (mitochondrial carrier; citrate transporter), member 1 | 6576 | ENSG00000100075 |

  
  

| **Database:cellular component      &nbspName:melanosome      &nbspID:GO:0042470** | | | | | | |
| --- | --- | --- | --- | --- | --- | --- |
| C=93; O=5; E=0.22; R=22.42; rawP=2.72e-06; adjP=1.24e-05 | | | | | | |
| Index | UserID | Value | Gene Symbol | Gene Name | EntrezGene | Ensembl |
| 1 | O14773 | NA | TPP1 | tripeptidyl peptidase I | 1200 | ENSG00000166340 |
| 2 | P07339 | NA | CTSD | cathepsin D | 1509 | ENSG00000117984 |
| 3 | P08133 | NA | ANXA6 | annexin A6 | 309 | ENSG00000197043 |
| 4 | Q14697 | NA | GANAB | glucosidase, alpha; neutral AB | 23193 | ENSG00000089597 |
| 5 | P40121 | NA | CAPG | capping protein (actin filament), gelsolin-like | 822 | ENSG00000042493 |

  
  

| **Database:cellular component      &nbspName:intracellular part      &nbspID:GO:0044424** | | | | | | |
| --- | --- | --- | --- | --- | --- | --- |
| C=12096; O=40; E=29.00; R=1.38; rawP=2.56e-06; adjP=1.24e-05 | | | | | | |
| Index | UserID | Value | Gene Symbol | Gene Name | EntrezGene | Ensembl |
| 1 | P10253 | NA | GAA | glucosidase, alpha; acid | 2548 | ENSG00000171298 |
| 2 | P17931 | NA | LGALS3 | lectin, galactoside-binding, soluble, 3 | 3958 | ENSG00000131981 |
| 3 | P08133 | NA | ANXA6 | annexin A6 | 309 | ENSG00000197043 |
| 4 | Q16851 | NA | UGP2 | UDP-glucose pyrophosphorylase 2 | 7360 | ENSG00000169764 |
| 5 | P30086 | NA | PEBP1 | phosphatidylethanolamine binding protein 1 | 5037 | ENSG00000089220 |
| 6 | P00367 | NA | GLUD1 | glutamate dehydrogenase 1 | 2746 | ENSG00000148672 |
| 7 | Q06323 | NA | PSME1 | proteasome (prosome, macropain) activator subunit 1 (PA28 alpha) | 5720 | ENSG00000092010 |
| 8 | P48735 | NA | IDH2 | isocitrate dehydrogenase 2 (NADP+), mitochondrial | 3418 | ENSG00000182054 |
| 9 | Q15019 | NA | SEPT2 | septin 2 | 4735 | ENSG00000168385 |
| 10 | Q9P0J0 | NA | NDUFA13 | NADH dehydrogenase (ubiquinone) 1 alpha subcomplex, 13 | 51079 | ENSG00000186010 |
| 11 | P09211 | NA | GSTP1 | glutathione S-transferase pi 1 | 2950 | ENSG00000084207 |
| 12 | O14773 | NA | TPP1 | tripeptidyl peptidase I | 1200 | ENSG00000166340 |
| 13 | Q05639 | NA | EEF1A2 | eukaryotic translation elongation factor 1 alpha 2 | 1917 | ENSG00000101210 |
| 14 | P22570 | NA | FDXR | ferredoxin reductase | 2232 | ENSG00000161513 |
| 15 | P40121 | NA | CAPG | capping protein (actin filament), gelsolin-like | 822 | ENSG00000042493 |
| 16 | P42765 | NA | ACAA2 | acetyl-CoA acyltransferase 2 | 10449 | ENSG00000167315 |
| 17 | P49419 | NA | ALDH7A1 | aldehyde dehydrogenase 7 family, member A1 | 501 | ENSG00000164904 |
| 18 | P07339 | NA | CTSD | cathepsin D | 1509 | ENSG00000117984 |
| 19 | Q14764 | NA | MVP | major vault protein | 9961 | ENSG00000013364 |
| 20 | Q14697 | NA | GANAB | glucosidase, alpha; neutral AB | 23193 | ENSG00000089597 |
| 21 | P53007 | NA | SLC25A1 | solute carrier family 25 (mitochondrial carrier; citrate transporter), member 1 | 6576 | ENSG00000100075 |
| 22 | Q9BPW8 | NA | NIPSNAP1 | nipsnap homolog 1 (C. elegans) | 8508 | ENSG00000184117 |
| 23 | P02792 | NA | FTL | ferritin, light polypeptide | 2512 | ENSG00000087086 |
| 24 | Q71U36 | NA | TUBA1A | tubulin, alpha 1a | 7846 | ENSG00000167552 |
| 25 | P00338 | NA | LDHA | lactate dehydrogenase A | 3939 | ENSG00000134333 |
| 26 | P38571 | NA | LIPA | lipase A, lysosomal acid, cholesterol esterase | 3988 | ENSG00000107798 |
| 27 | Q13011 | NA | ECH1 | enoyl CoA hydratase 1, peroxisomal | 1891 | ENSG00000104823 |
| 28 | P23381 | NA | WARS | tryptophanyl-tRNA synthetase | 7453 | ENSG00000140105 |
| 29 | Q14980 | NA | NUMA1 | nuclear mitotic apparatus protein 1 | 4926 | ENSG00000137497 |
| 30 | P04040 | NA | CAT | catalase | 847 | ENSG00000121691 |
| 31 | Q9UBQ7 | NA | GRHPR | glyoxylate reductase/hydroxypyruvate reductase | 9380 | ENSG00000137106 |
| 32 | P51649 | NA | ALDH5A1 | aldehyde dehydrogenase 5 family, member A1 | 7915 | ENSG00000112294 |
| 33 | P30044 | NA | PRDX5 | peroxiredoxin 5 | 25824 | ENSG00000126432 |
| 34 | P13674 | NA | P4HA1 | prolyl 4-hydroxylase, alpha polypeptide I | 5033 | ENSG00000122884 |
| 35 | P24752 | NA | ACAT1 | acetyl-CoA acetyltransferase 1 | 38 | ENSG00000075239 |
| 36 | P16278 | NA | GLB1 | galactosidase, beta 1 | 2720 | ENSG00000170266 |
| 37 | P06744 | NA | GPI | glucose-6-phosphate isomerase | 2821 | ENSG00000105220 |
| 38 | P07602 | NA | PSAP | prosaposin | 5660 | ENSG00000197746 |
| 39 | P49748 | NA | ACADVL | acyl-CoA dehydrogenase, very long chain | 37 | ENSG00000072778 |
| 40 | P07384 | NA | CAPN1 | calpain 1, (mu/I) large subunit | 823 | ENSG00000014216 |

  
  

| **Database:cellular component      &nbspName:mitochondrial matrix      &nbspID:GO:0005759** | | | | | | |
| --- | --- | --- | --- | --- | --- | --- |
| C=278; O=7; E=0.67; R=10.50; rawP=3.85e-06; adjP=1.61e-05 | | | | | | |
| Index | UserID | Value | Gene Symbol | Gene Name | EntrezGene | Ensembl |
| 1 | P24752 | NA | ACAT1 | acetyl-CoA acetyltransferase 1 | 38 | ENSG00000075239 |
| 2 | P49419 | NA | ALDH7A1 | aldehyde dehydrogenase 7 family, member A1 | 501 | ENSG00000164904 |
| 3 | P48735 | NA | IDH2 | isocitrate dehydrogenase 2 (NADP+), mitochondrial | 3418 | ENSG00000182054 |
| 4 | P00367 | NA | GLUD1 | glutamate dehydrogenase 1 | 2746 | ENSG00000148672 |
| 5 | P51649 | NA | ALDH5A1 | aldehyde dehydrogenase 5 family, member A1 | 7915 | ENSG00000112294 |
| 6 | P49748 | NA | ACADVL | acyl-CoA dehydrogenase, very long chain | 37 | ENSG00000072778 |
| 7 | P22570 | NA | FDXR | ferredoxin reductase | 2232 | ENSG00000161513 |

  
  

| **Database:cellular component      &nbspName:intracellular      &nbspID:GO:0005622** | | | | | | |
| --- | --- | --- | --- | --- | --- | --- |
| C=12412; O=40; E=29.76; R=1.34; rawP=7.19e-06; adjP=2.84e-05 | | | | | | |
| Index | UserID | Value | Gene Symbol | Gene Name | EntrezGene | Ensembl |
| 1 | P10253 | NA | GAA | glucosidase, alpha; acid | 2548 | ENSG00000171298 |
| 2 | P17931 | NA | LGALS3 | lectin, galactoside-binding, soluble, 3 | 3958 | ENSG00000131981 |
| 3 | P08133 | NA | ANXA6 | annexin A6 | 309 | ENSG00000197043 |
| 4 | Q16851 | NA | UGP2 | UDP-glucose pyrophosphorylase 2 | 7360 | ENSG00000169764 |
| 5 | P30086 | NA | PEBP1 | phosphatidylethanolamine binding protein 1 | 5037 | ENSG00000089220 |
| 6 | P00367 | NA | GLUD1 | glutamate dehydrogenase 1 | 2746 | ENSG00000148672 |
| 7 | Q06323 | NA | PSME1 | proteasome (prosome, macropain) activator subunit 1 (PA28 alpha) | 5720 | ENSG00000092010 |
| 8 | P48735 | NA | IDH2 | isocitrate dehydrogenase 2 (NADP+), mitochondrial | 3418 | ENSG00000182054 |
| 9 | Q15019 | NA | SEPT2 | septin 2 | 4735 | ENSG00000168385 |
| 10 | Q9P0J0 | NA | NDUFA13 | NADH dehydrogenase (ubiquinone) 1 alpha subcomplex, 13 | 51079 | ENSG00000186010 |
| 11 | P09211 | NA | GSTP1 | glutathione S-transferase pi 1 | 2950 | ENSG00000084207 |
| 12 | O14773 | NA | TPP1 | tripeptidyl peptidase I | 1200 | ENSG00000166340 |
| 13 | Q05639 | NA | EEF1A2 | eukaryotic translation elongation factor 1 alpha 2 | 1917 | ENSG00000101210 |
| 14 | P22570 | NA | FDXR | ferredoxin reductase | 2232 | ENSG00000161513 |
| 15 | P40121 | NA | CAPG | capping protein (actin filament), gelsolin-like | 822 | ENSG00000042493 |
| 16 | P42765 | NA | ACAA2 | acetyl-CoA acyltransferase 2 | 10449 | ENSG00000167315 |
| 17 | P49419 | NA | ALDH7A1 | aldehyde dehydrogenase 7 family, member A1 | 501 | ENSG00000164904 |
| 18 | P07339 | NA | CTSD | cathepsin D | 1509 | ENSG00000117984 |
| 19 | Q14764 | NA | MVP | major vault protein | 9961 | ENSG00000013364 |
| 20 | Q14697 | NA | GANAB | glucosidase, alpha; neutral AB | 23193 | ENSG00000089597 |
| 21 | P53007 | NA | SLC25A1 | solute carrier family 25 (mitochondrial carrier; citrate transporter), member 1 | 6576 | ENSG00000100075 |
| 22 | Q9BPW8 | NA | NIPSNAP1 | nipsnap homolog 1 (C. elegans) | 8508 | ENSG00000184117 |
| 23 | P02792 | NA | FTL | ferritin, light polypeptide | 2512 | ENSG00000087086 |
| 24 | Q71U36 | NA | TUBA1A | tubulin, alpha 1a | 7846 | ENSG00000167552 |
| 25 | P00338 | NA | LDHA | lactate dehydrogenase A | 3939 | ENSG00000134333 |
| 26 | P38571 | NA | LIPA | lipase A, lysosomal acid, cholesterol esterase | 3988 | ENSG00000107798 |
| 27 | Q13011 | NA | ECH1 | enoyl CoA hydratase 1, peroxisomal | 1891 | ENSG00000104823 |
| 28 | P23381 | NA | WARS | tryptophanyl-tRNA synthetase | 7453 | ENSG00000140105 |
| 29 | Q14980 | NA | NUMA1 | nuclear mitotic apparatus protein 1 | 4926 | ENSG00000137497 |
| 30 | P04040 | NA | CAT | catalase | 847 | ENSG00000121691 |
| 31 | Q9UBQ7 | NA | GRHPR | glyoxylate reductase/hydroxypyruvate reductase | 9380 | ENSG00000137106 |
| 32 | P51649 | NA | ALDH5A1 | aldehyde dehydrogenase 5 family, member A1 | 7915 | ENSG00000112294 |
| 33 | P30044 | NA | PRDX5 | peroxiredoxin 5 | 25824 | ENSG00000126432 |
| 34 | P13674 | NA | P4HA1 | prolyl 4-hydroxylase, alpha polypeptide I | 5033 | ENSG00000122884 |
| 35 | P24752 | NA | ACAT1 | acetyl-CoA acetyltransferase 1 | 38 | ENSG00000075239 |
| 36 | P16278 | NA | GLB1 | galactosidase, beta 1 | 2720 | ENSG00000170266 |
| 37 | P06744 | NA | GPI | glucose-6-phosphate isomerase | 2821 | ENSG00000105220 |
| 38 | P07602 | NA | PSAP | prosaposin | 5660 | ENSG00000197746 |
| 39 | P49748 | NA | ACADVL | acyl-CoA dehydrogenase, very long chain | 37 | ENSG00000072778 |
| 40 | P07384 | NA | CAPN1 | calpain 1, (mu/I) large subunit | 823 | ENSG00000014216 |

  
  

| **Database:cellular component      &nbspName:intracellular organelle part      &nbspID:GO:0044446** | | | | | | |
| --- | --- | --- | --- | --- | --- | --- |
| C=6690; O=30; E=16.04; R=1.87; rawP=7.75e-06; adjP=2.90e-05 | | | | | | |
| Index | UserID | Value | Gene Symbol | Gene Name | EntrezGene | Ensembl |
| 1 | P10253 | NA | GAA | glucosidase, alpha; acid | 2548 | ENSG00000171298 |
| 2 | Q71U36 | NA | TUBA1A | tubulin, alpha 1a | 7846 | ENSG00000167552 |
| 3 | P17931 | NA | LGALS3 | lectin, galactoside-binding, soluble, 3 | 3958 | ENSG00000131981 |
| 4 | P30086 | NA | PEBP1 | phosphatidylethanolamine binding protein 1 | 5037 | ENSG00000089220 |
| 5 | P00367 | NA | GLUD1 | glutamate dehydrogenase 1 | 2746 | ENSG00000148672 |
| 6 | Q06323 | NA | PSME1 | proteasome (prosome, macropain) activator subunit 1 (PA28 alpha) | 5720 | ENSG00000092010 |
| 7 | P48735 | NA | IDH2 | isocitrate dehydrogenase 2 (NADP+), mitochondrial | 3418 | ENSG00000182054 |
| 8 | Q15019 | NA | SEPT2 | septin 2 | 4735 | ENSG00000168385 |
| 9 | Q9P0J0 | NA | NDUFA13 | NADH dehydrogenase (ubiquinone) 1 alpha subcomplex, 13 | 51079 | ENSG00000186010 |
| 10 | P04040 | NA | CAT | catalase | 847 | ENSG00000121691 |
| 11 | Q14980 | NA | NUMA1 | nuclear mitotic apparatus protein 1 | 4926 | ENSG00000137497 |
| 12 | O14773 | NA | TPP1 | tripeptidyl peptidase I | 1200 | ENSG00000166340 |
| 13 | Q9UBQ7 | NA | GRHPR | glyoxylate reductase/hydroxypyruvate reductase | 9380 | ENSG00000137106 |
| 14 | P51649 | NA | ALDH5A1 | aldehyde dehydrogenase 5 family, member A1 | 7915 | ENSG00000112294 |
| 15 | P30044 | NA | PRDX5 | peroxiredoxin 5 | 25824 | ENSG00000126432 |
| 16 | P22570 | NA | FDXR | ferredoxin reductase | 2232 | ENSG00000161513 |
| 17 | P40121 | NA | CAPG | capping protein (actin filament), gelsolin-like | 822 | ENSG00000042493 |
| 18 | P42765 | NA | ACAA2 | acetyl-CoA acyltransferase 2 | 10449 | ENSG00000167315 |
| 19 | P13674 | NA | P4HA1 | prolyl 4-hydroxylase, alpha polypeptide I | 5033 | ENSG00000122884 |
| 20 | P24752 | NA | ACAT1 | acetyl-CoA acetyltransferase 1 | 38 | ENSG00000075239 |
| 21 | P16278 | NA | GLB1 | galactosidase, beta 1 | 2720 | ENSG00000170266 |
| 22 | P49419 | NA | ALDH7A1 | aldehyde dehydrogenase 7 family, member A1 | 501 | ENSG00000164904 |
| 23 | P06744 | NA | GPI | glucose-6-phosphate isomerase | 2821 | ENSG00000105220 |
| 24 | P07602 | NA | PSAP | prosaposin | 5660 | ENSG00000197746 |
| 25 | Q14764 | NA | MVP | major vault protein | 9961 | ENSG00000013364 |
| 26 | P07339 | NA | CTSD | cathepsin D | 1509 | ENSG00000117984 |
| 27 | P49748 | NA | ACADVL | acyl-CoA dehydrogenase, very long chain | 37 | ENSG00000072778 |
| 28 | Q14697 | NA | GANAB | glucosidase, alpha; neutral AB | 23193 | ENSG00000089597 |
| 29 | P53007 | NA | SLC25A1 | solute carrier family 25 (mitochondrial carrier; citrate transporter), member 1 | 6576 | ENSG00000100075 |
| 30 | Q9BPW8 | NA | NIPSNAP1 | nipsnap homolog 1 (C. elegans) | 8508 | ENSG00000184117 |

  
  

| **Database:cellular component      &nbspName:organelle part      &nbspID:GO:0044422** | | | | | | |
| --- | --- | --- | --- | --- | --- | --- |
| C=6777; O=30; E=16.25; R=1.85; rawP=1.05e-05; adjP=3.73e-05 | | | | | | |
| Index | UserID | Value | Gene Symbol | Gene Name | EntrezGene | Ensembl |
| 1 | P10253 | NA | GAA | glucosidase, alpha; acid | 2548 | ENSG00000171298 |
| 2 | Q71U36 | NA | TUBA1A | tubulin, alpha 1a | 7846 | ENSG00000167552 |
| 3 | P17931 | NA | LGALS3 | lectin, galactoside-binding, soluble, 3 | 3958 | ENSG00000131981 |
| 4 | P30086 | NA | PEBP1 | phosphatidylethanolamine binding protein 1 | 5037 | ENSG00000089220 |
| 5 | P00367 | NA | GLUD1 | glutamate dehydrogenase 1 | 2746 | ENSG00000148672 |
| 6 | Q06323 | NA | PSME1 | proteasome (prosome, macropain) activator subunit 1 (PA28 alpha) | 5720 | ENSG00000092010 |
| 7 | P48735 | NA | IDH2 | isocitrate dehydrogenase 2 (NADP+), mitochondrial | 3418 | ENSG00000182054 |
| 8 | Q15019 | NA | SEPT2 | septin 2 | 4735 | ENSG00000168385 |
| 9 | Q9P0J0 | NA | NDUFA13 | NADH dehydrogenase (ubiquinone) 1 alpha subcomplex, 13 | 51079 | ENSG00000186010 |
| 10 | P04040 | NA | CAT | catalase | 847 | ENSG00000121691 |
| 11 | Q14980 | NA | NUMA1 | nuclear mitotic apparatus protein 1 | 4926 | ENSG00000137497 |
| 12 | O14773 | NA | TPP1 | tripeptidyl peptidase I | 1200 | ENSG00000166340 |
| 13 | Q9UBQ7 | NA | GRHPR | glyoxylate reductase/hydroxypyruvate reductase | 9380 | ENSG00000137106 |
| 14 | P51649 | NA | ALDH5A1 | aldehyde dehydrogenase 5 family, member A1 | 7915 | ENSG00000112294 |
| 15 | P30044 | NA | PRDX5 | peroxiredoxin 5 | 25824 | ENSG00000126432 |
| 16 | P22570 | NA | FDXR | ferredoxin reductase | 2232 | ENSG00000161513 |
| 17 | P40121 | NA | CAPG | capping protein (actin filament), gelsolin-like | 822 | ENSG00000042493 |
| 18 | P42765 | NA | ACAA2 | acetyl-CoA acyltransferase 2 | 10449 | ENSG00000167315 |
| 19 | P13674 | NA | P4HA1 | prolyl 4-hydroxylase, alpha polypeptide I | 5033 | ENSG00000122884 |
| 20 | P24752 | NA | ACAT1 | acetyl-CoA acetyltransferase 1 | 38 | ENSG00000075239 |
| 21 | P16278 | NA | GLB1 | galactosidase, beta 1 | 2720 | ENSG00000170266 |
| 22 | P49419 | NA | ALDH7A1 | aldehyde dehydrogenase 7 family, member A1 | 501 | ENSG00000164904 |
| 23 | P06744 | NA | GPI | glucose-6-phosphate isomerase | 2821 | ENSG00000105220 |
| 24 | P07602 | NA | PSAP | prosaposin | 5660 | ENSG00000197746 |
| 25 | Q14764 | NA | MVP | major vault protein | 9961 | ENSG00000013364 |
| 26 | P07339 | NA | CTSD | cathepsin D | 1509 | ENSG00000117984 |
| 27 | P49748 | NA | ACADVL | acyl-CoA dehydrogenase, very long chain | 37 | ENSG00000072778 |
| 28 | Q14697 | NA | GANAB | glucosidase, alpha; neutral AB | 23193 | ENSG00000089597 |
| 29 | P53007 | NA | SLC25A1 | solute carrier family 25 (mitochondrial carrier; citrate transporter), member 1 | 6576 | ENSG00000100075 |
| 30 | Q9BPW8 | NA | NIPSNAP1 | nipsnap homolog 1 (C. elegans) | 8508 | ENSG00000184117 |

  
  

| **Database:cellular component      &nbspName:lytic vacuole      &nbspID:GO:0000323** | | | | | | |
| --- | --- | --- | --- | --- | --- | --- |
| C=345; O=7; E=0.83; R=8.46; rawP=1.58e-05; adjP=5.10e-05 | | | | | | |
| Index | UserID | Value | Gene Symbol | Gene Name | EntrezGene | Ensembl |
| 1 | P04040 | NA | CAT | catalase | 847 | ENSG00000121691 |
| 2 | O14773 | NA | TPP1 | tripeptidyl peptidase I | 1200 | ENSG00000166340 |
| 3 | P16278 | NA | GLB1 | galactosidase, beta 1 | 2720 | ENSG00000170266 |
| 4 | P10253 | NA | GAA | glucosidase, alpha; acid | 2548 | ENSG00000171298 |
| 5 | P07602 | NA | PSAP | prosaposin | 5660 | ENSG00000197746 |
| 6 | P07339 | NA | CTSD | cathepsin D | 1509 | ENSG00000117984 |
| 7 | P38571 | NA | LIPA | lipase A, lysosomal acid, cholesterol esterase | 3988 | ENSG00000107798 |

  
  

| **Database:cellular component      &nbspName:lysosome      &nbspID:GO:0005764** | | | | | | |
| --- | --- | --- | --- | --- | --- | --- |
| C=345; O=7; E=0.83; R=8.46; rawP=1.58e-05; adjP=5.10e-05 | | | | | | |
| Index | UserID | Value | Gene Symbol | Gene Name | EntrezGene | Ensembl |
| 1 | P04040 | NA | CAT | catalase | 847 | ENSG00000121691 |
| 2 | O14773 | NA | TPP1 | tripeptidyl peptidase I | 1200 | ENSG00000166340 |
| 3 | P16278 | NA | GLB1 | galactosidase, beta 1 | 2720 | ENSG00000170266 |
| 4 | P10253 | NA | GAA | glucosidase, alpha; acid | 2548 | ENSG00000171298 |
| 5 | P07602 | NA | PSAP | prosaposin | 5660 | ENSG00000197746 |
| 6 | P07339 | NA | CTSD | cathepsin D | 1509 | ENSG00000117984 |
| 7 | P38571 | NA | LIPA | lipase A, lysosomal acid, cholesterol esterase | 3988 | ENSG00000107798 |

  
  

| **Database:cellular component      &nbspName:lysosomal lumen      &nbspID:GO:0043202** | | | | | | |
| --- | --- | --- | --- | --- | --- | --- |
| C=71; O=4; E=0.17; R=23.50; rawP=2.45e-05; adjP=7.56e-05 | | | | | | |
| Index | UserID | Value | Gene Symbol | Gene Name | EntrezGene | Ensembl |
| 1 | O14773 | NA | TPP1 | tripeptidyl peptidase I | 1200 | ENSG00000166340 |
| 2 | P16278 | NA | GLB1 | galactosidase, beta 1 | 2720 | ENSG00000170266 |
| 3 | P07602 | NA | PSAP | prosaposin | 5660 | ENSG00000197746 |
| 4 | P07339 | NA | CTSD | cathepsin D | 1509 | ENSG00000117984 |

  
  

| **Database:cellular component      &nbspName:vacuolar lumen      &nbspID:GO:0005775** | | | | | | |
| --- | --- | --- | --- | --- | --- | --- |
| C=75; O=4; E=0.18; R=22.24; rawP=3.05e-05; adjP=8.63e-05 | | | | | | |
| Index | UserID | Value | Gene Symbol | Gene Name | EntrezGene | Ensembl |
| 1 | O14773 | NA | TPP1 | tripeptidyl peptidase I | 1200 | ENSG00000166340 |
| 2 | P16278 | NA | GLB1 | galactosidase, beta 1 | 2720 | ENSG00000170266 |
| 3 | P07602 | NA | PSAP | prosaposin | 5660 | ENSG00000197746 |
| 4 | P07339 | NA | CTSD | cathepsin D | 1509 | ENSG00000117984 |

  
  

| **Database:cellular component      &nbspName:intracellular membrane-bounded organelle      &nbspID:GO:0043231** | | | | | | |
| --- | --- | --- | --- | --- | --- | --- |
| C=9484; O=35; E=22.74; R=1.54; rawP=3.05e-05; adjP=8.63e-05 | | | | | | |
| Index | UserID | Value | Gene Symbol | Gene Name | EntrezGene | Ensembl |
| 1 | P10253 | NA | GAA | glucosidase, alpha; acid | 2548 | ENSG00000171298 |
| 2 | P17931 | NA | LGALS3 | lectin, galactoside-binding, soluble, 3 | 3958 | ENSG00000131981 |
| 3 | P08133 | NA | ANXA6 | annexin A6 | 309 | ENSG00000197043 |
| 4 | P30086 | NA | PEBP1 | phosphatidylethanolamine binding protein 1 | 5037 | ENSG00000089220 |
| 5 | P00367 | NA | GLUD1 | glutamate dehydrogenase 1 | 2746 | ENSG00000148672 |
| 6 | Q06323 | NA | PSME1 | proteasome (prosome, macropain) activator subunit 1 (PA28 alpha) | 5720 | ENSG00000092010 |
| 7 | P48735 | NA | IDH2 | isocitrate dehydrogenase 2 (NADP+), mitochondrial | 3418 | ENSG00000182054 |
| 8 | Q15019 | NA | SEPT2 | septin 2 | 4735 | ENSG00000168385 |
| 9 | Q9P0J0 | NA | NDUFA13 | NADH dehydrogenase (ubiquinone) 1 alpha subcomplex, 13 | 51079 | ENSG00000186010 |
| 10 | P09211 | NA | GSTP1 | glutathione S-transferase pi 1 | 2950 | ENSG00000084207 |
| 11 | O14773 | NA | TPP1 | tripeptidyl peptidase I | 1200 | ENSG00000166340 |
| 12 | Q05639 | NA | EEF1A2 | eukaryotic translation elongation factor 1 alpha 2 | 1917 | ENSG00000101210 |
| 13 | P22570 | NA | FDXR | ferredoxin reductase | 2232 | ENSG00000161513 |
| 14 | P40121 | NA | CAPG | capping protein (actin filament), gelsolin-like | 822 | ENSG00000042493 |
| 15 | P42765 | NA | ACAA2 | acetyl-CoA acyltransferase 2 | 10449 | ENSG00000167315 |
| 16 | P49419 | NA | ALDH7A1 | aldehyde dehydrogenase 7 family, member A1 | 501 | ENSG00000164904 |
| 17 | P07339 | NA | CTSD | cathepsin D | 1509 | ENSG00000117984 |
| 18 | Q14764 | NA | MVP | major vault protein | 9961 | ENSG00000013364 |
| 19 | Q14697 | NA | GANAB | glucosidase, alpha; neutral AB | 23193 | ENSG00000089597 |
| 20 | P53007 | NA | SLC25A1 | solute carrier family 25 (mitochondrial carrier; citrate transporter), member 1 | 6576 | ENSG00000100075 |
| 21 | Q9BPW8 | NA | NIPSNAP1 | nipsnap homolog 1 (C. elegans) | 8508 | ENSG00000184117 |
| 22 | P00338 | NA | LDHA | lactate dehydrogenase A | 3939 | ENSG00000134333 |
| 23 | P38571 | NA | LIPA | lipase A, lysosomal acid, cholesterol esterase | 3988 | ENSG00000107798 |
| 24 | Q13011 | NA | ECH1 | enoyl CoA hydratase 1, peroxisomal | 1891 | ENSG00000104823 |
| 25 | Q14980 | NA | NUMA1 | nuclear mitotic apparatus protein 1 | 4926 | ENSG00000137497 |
| 26 | P04040 | NA | CAT | catalase | 847 | ENSG00000121691 |
| 27 | Q9UBQ7 | NA | GRHPR | glyoxylate reductase/hydroxypyruvate reductase | 9380 | ENSG00000137106 |
| 28 | P51649 | NA | ALDH5A1 | aldehyde dehydrogenase 5 family, member A1 | 7915 | ENSG00000112294 |
| 29 | P30044 | NA | PRDX5 | peroxiredoxin 5 | 25824 | ENSG00000126432 |
| 30 | P24752 | NA | ACAT1 | acetyl-CoA acetyltransferase 1 | 38 | ENSG00000075239 |
| 31 | P13674 | NA | P4HA1 | prolyl 4-hydroxylase, alpha polypeptide I | 5033 | ENSG00000122884 |
| 32 | P16278 | NA | GLB1 | galactosidase, beta 1 | 2720 | ENSG00000170266 |
| 33 | P06744 | NA | GPI | glucose-6-phosphate isomerase | 2821 | ENSG00000105220 |
| 34 | P07602 | NA | PSAP | prosaposin | 5660 | ENSG00000197746 |
| 35 | P49748 | NA | ACADVL | acyl-CoA dehydrogenase, very long chain | 37 | ENSG00000072778 |

  
  

| **Database:cellular component      &nbspName:membrane-bounded organelle      &nbspID:GO:0043227** | | | | | | |
| --- | --- | --- | --- | --- | --- | --- |
| C=9495; O=35; E=22.77; R=1.54; rawP=3.16e-05; adjP=8.63e-05 | | | | | | |
| Index | UserID | Value | Gene Symbol | Gene Name | EntrezGene | Ensembl |
| 1 | P10253 | NA | GAA | glucosidase, alpha; acid | 2548 | ENSG00000171298 |
| 2 | P17931 | NA | LGALS3 | lectin, galactoside-binding, soluble, 3 | 3958 | ENSG00000131981 |
| 3 | P08133 | NA | ANXA6 | annexin A6 | 309 | ENSG00000197043 |
| 4 | P30086 | NA | PEBP1 | phosphatidylethanolamine binding protein 1 | 5037 | ENSG00000089220 |
| 5 | P00367 | NA | GLUD1 | glutamate dehydrogenase 1 | 2746 | ENSG00000148672 |
| 6 | Q06323 | NA | PSME1 | proteasome (prosome, macropain) activator subunit 1 (PA28 alpha) | 5720 | ENSG00000092010 |
| 7 | P48735 | NA | IDH2 | isocitrate dehydrogenase 2 (NADP+), mitochondrial | 3418 | ENSG00000182054 |
| 8 | Q15019 | NA | SEPT2 | septin 2 | 4735 | ENSG00000168385 |
| 9 | Q9P0J0 | NA | NDUFA13 | NADH dehydrogenase (ubiquinone) 1 alpha subcomplex, 13 | 51079 | ENSG00000186010 |
| 10 | P09211 | NA | GSTP1 | glutathione S-transferase pi 1 | 2950 | ENSG00000084207 |
| 11 | O14773 | NA | TPP1 | tripeptidyl peptidase I | 1200 | ENSG00000166340 |
| 12 | Q05639 | NA | EEF1A2 | eukaryotic translation elongation factor 1 alpha 2 | 1917 | ENSG00000101210 |
| 13 | P22570 | NA | FDXR | ferredoxin reductase | 2232 | ENSG00000161513 |
| 14 | P40121 | NA | CAPG | capping protein (actin filament), gelsolin-like | 822 | ENSG00000042493 |
| 15 | P42765 | NA | ACAA2 | acetyl-CoA acyltransferase 2 | 10449 | ENSG00000167315 |
| 16 | P49419 | NA | ALDH7A1 | aldehyde dehydrogenase 7 family, member A1 | 501 | ENSG00000164904 |
| 17 | P07339 | NA | CTSD | cathepsin D | 1509 | ENSG00000117984 |
| 18 | Q14764 | NA | MVP | major vault protein | 9961 | ENSG00000013364 |
| 19 | Q14697 | NA | GANAB | glucosidase, alpha; neutral AB | 23193 | ENSG00000089597 |
| 20 | P53007 | NA | SLC25A1 | solute carrier family 25 (mitochondrial carrier; citrate transporter), member 1 | 6576 | ENSG00000100075 |
| 21 | Q9BPW8 | NA | NIPSNAP1 | nipsnap homolog 1 (C. elegans) | 8508 | ENSG00000184117 |
| 22 | P00338 | NA | LDHA | lactate dehydrogenase A | 3939 | ENSG00000134333 |
| 23 | P38571 | NA | LIPA | lipase A, lysosomal acid, cholesterol esterase | 3988 | ENSG00000107798 |
| 24 | Q13011 | NA | ECH1 | enoyl CoA hydratase 1, peroxisomal | 1891 | ENSG00000104823 |
| 25 | Q14980 | NA | NUMA1 | nuclear mitotic apparatus protein 1 | 4926 | ENSG00000137497 |
| 26 | P04040 | NA | CAT | catalase | 847 | ENSG00000121691 |
| 27 | Q9UBQ7 | NA | GRHPR | glyoxylate reductase/hydroxypyruvate reductase | 9380 | ENSG00000137106 |
| 28 | P51649 | NA | ALDH5A1 | aldehyde dehydrogenase 5 family, member A1 | 7915 | ENSG00000112294 |
| 29 | P30044 | NA | PRDX5 | peroxiredoxin 5 | 25824 | ENSG00000126432 |
| 30 | P24752 | NA | ACAT1 | acetyl-CoA acetyltransferase 1 | 38 | ENSG00000075239 |
| 31 | P13674 | NA | P4HA1 | prolyl 4-hydroxylase, alpha polypeptide I | 5033 | ENSG00000122884 |
| 32 | P16278 | NA | GLB1 | galactosidase, beta 1 | 2720 | ENSG00000170266 |
| 33 | P06744 | NA | GPI | glucose-6-phosphate isomerase | 2821 | ENSG00000105220 |
| 34 | P07602 | NA | PSAP | prosaposin | 5660 | ENSG00000197746 |
| 35 | P49748 | NA | ACADVL | acyl-CoA dehydrogenase, very long chain | 37 | ENSG00000072778 |

  
  

| **Database:cellular component      &nbspName:vacuole      &nbspID:GO:0005773** | | | | | | |
| --- | --- | --- | --- | --- | --- | --- |
| C=408; O=7; E=0.98; R=7.16; rawP=4.62e-05; adjP=0.0001 | | | | | | |
| Index | UserID | Value | Gene Symbol | Gene Name | EntrezGene | Ensembl |
| 1 | P04040 | NA | CAT | catalase | 847 | ENSG00000121691 |
| 2 | O14773 | NA | TPP1 | tripeptidyl peptidase I | 1200 | ENSG00000166340 |
| 3 | P16278 | NA | GLB1 | galactosidase, beta 1 | 2720 | ENSG00000170266 |
| 4 | P10253 | NA | GAA | glucosidase, alpha; acid | 2548 | ENSG00000171298 |
| 5 | P07602 | NA | PSAP | prosaposin | 5660 | ENSG00000197746 |
| 6 | P07339 | NA | CTSD | cathepsin D | 1509 | ENSG00000117984 |
| 7 | P38571 | NA | LIPA | lipase A, lysosomal acid, cholesterol esterase | 3988 | ENSG00000107798 |

  
  

| **Database:cellular component      &nbspName:microbody lumen      &nbspID:GO:0031907** | | | | | | |
| --- | --- | --- | --- | --- | --- | --- |
| C=33; O=3; E=0.08; R=37.91; rawP=6.63e-05; adjP=0.0002 | | | | | | |
| Index | UserID | Value | Gene Symbol | Gene Name | EntrezGene | Ensembl |
| 1 | P04040 | NA | CAT | catalase | 847 | ENSG00000121691 |
| 2 | Q9UBQ7 | NA | GRHPR | glyoxylate reductase/hydroxypyruvate reductase | 9380 | ENSG00000137106 |
| 3 | P30044 | NA | PRDX5 | peroxiredoxin 5 | 25824 | ENSG00000126432 |

  
  

| **Database:cellular component      &nbspName:organelle      &nbspID:GO:0043226** | | | | | | |
| --- | --- | --- | --- | --- | --- | --- |
| C=10536; O=36; E=25.26; R=1.43; rawP=0.0001; adjP=0.0002 | | | | | | |
| Index | UserID | Value | Gene Symbol | Gene Name | EntrezGene | Ensembl |
| 1 | P10253 | NA | GAA | glucosidase, alpha; acid | 2548 | ENSG00000171298 |
| 2 | P17931 | NA | LGALS3 | lectin, galactoside-binding, soluble, 3 | 3958 | ENSG00000131981 |
| 3 | P08133 | NA | ANXA6 | annexin A6 | 309 | ENSG00000197043 |
| 4 | P30086 | NA | PEBP1 | phosphatidylethanolamine binding protein 1 | 5037 | ENSG00000089220 |
| 5 | P00367 | NA | GLUD1 | glutamate dehydrogenase 1 | 2746 | ENSG00000148672 |
| 6 | Q06323 | NA | PSME1 | proteasome (prosome, macropain) activator subunit 1 (PA28 alpha) | 5720 | ENSG00000092010 |
| 7 | P48735 | NA | IDH2 | isocitrate dehydrogenase 2 (NADP+), mitochondrial | 3418 | ENSG00000182054 |
| 8 | Q15019 | NA | SEPT2 | septin 2 | 4735 | ENSG00000168385 |
| 9 | Q9P0J0 | NA | NDUFA13 | NADH dehydrogenase (ubiquinone) 1 alpha subcomplex, 13 | 51079 | ENSG00000186010 |
| 10 | P09211 | NA | GSTP1 | glutathione S-transferase pi 1 | 2950 | ENSG00000084207 |
| 11 | O14773 | NA | TPP1 | tripeptidyl peptidase I | 1200 | ENSG00000166340 |
| 12 | Q05639 | NA | EEF1A2 | eukaryotic translation elongation factor 1 alpha 2 | 1917 | ENSG00000101210 |
| 13 | P22570 | NA | FDXR | ferredoxin reductase | 2232 | ENSG00000161513 |
| 14 | P40121 | NA | CAPG | capping protein (actin filament), gelsolin-like | 822 | ENSG00000042493 |
| 15 | P42765 | NA | ACAA2 | acetyl-CoA acyltransferase 2 | 10449 | ENSG00000167315 |
| 16 | P49419 | NA | ALDH7A1 | aldehyde dehydrogenase 7 family, member A1 | 501 | ENSG00000164904 |
| 17 | P07339 | NA | CTSD | cathepsin D | 1509 | ENSG00000117984 |
| 18 | Q14764 | NA | MVP | major vault protein | 9961 | ENSG00000013364 |
| 19 | Q14697 | NA | GANAB | glucosidase, alpha; neutral AB | 23193 | ENSG00000089597 |
| 20 | P53007 | NA | SLC25A1 | solute carrier family 25 (mitochondrial carrier; citrate transporter), member 1 | 6576 | ENSG00000100075 |
| 21 | Q9BPW8 | NA | NIPSNAP1 | nipsnap homolog 1 (C. elegans) | 8508 | ENSG00000184117 |
| 22 | Q71U36 | NA | TUBA1A | tubulin, alpha 1a | 7846 | ENSG00000167552 |
| 23 | P00338 | NA | LDHA | lactate dehydrogenase A | 3939 | ENSG00000134333 |
| 24 | P38571 | NA | LIPA | lipase A, lysosomal acid, cholesterol esterase | 3988 | ENSG00000107798 |
| 25 | Q13011 | NA | ECH1 | enoyl CoA hydratase 1, peroxisomal | 1891 | ENSG00000104823 |
| 26 | Q14980 | NA | NUMA1 | nuclear mitotic apparatus protein 1 | 4926 | ENSG00000137497 |
| 27 | P04040 | NA | CAT | catalase | 847 | ENSG00000121691 |
| 28 | Q9UBQ7 | NA | GRHPR | glyoxylate reductase/hydroxypyruvate reductase | 9380 | ENSG00000137106 |
| 29 | P51649 | NA | ALDH5A1 | aldehyde dehydrogenase 5 family, member A1 | 7915 | ENSG00000112294 |
| 30 | P30044 | NA | PRDX5 | peroxiredoxin 5 | 25824 | ENSG00000126432 |
| 31 | P24752 | NA | ACAT1 | acetyl-CoA acetyltransferase 1 | 38 | ENSG00000075239 |
| 32 | P13674 | NA | P4HA1 | prolyl 4-hydroxylase, alpha polypeptide I | 5033 | ENSG00000122884 |
| 33 | P16278 | NA | GLB1 | galactosidase, beta 1 | 2720 | ENSG00000170266 |
| 34 | P06744 | NA | GPI | glucose-6-phosphate isomerase | 2821 | ENSG00000105220 |
| 35 | P07602 | NA | PSAP | prosaposin | 5660 | ENSG00000197746 |
| 36 | P49748 | NA | ACADVL | acyl-CoA dehydrogenase, very long chain | 37 | ENSG00000072778 |

  
  

| **Database:cellular component      &nbspName:intracellular organelle      &nbspID:GO:0043229** | | | | | | |
| --- | --- | --- | --- | --- | --- | --- |
| C=10521; O=36; E=25.23; R=1.43; rawP=0.0001; adjP=0.0002 | | | | | | |
| Index | UserID | Value | Gene Symbol | Gene Name | EntrezGene | Ensembl |
| 1 | P10253 | NA | GAA | glucosidase, alpha; acid | 2548 | ENSG00000171298 |
| 2 | P17931 | NA | LGALS3 | lectin, galactoside-binding, soluble, 3 | 3958 | ENSG00000131981 |
| 3 | P08133 | NA | ANXA6 | annexin A6 | 309 | ENSG00000197043 |
| 4 | P30086 | NA | PEBP1 | phosphatidylethanolamine binding protein 1 | 5037 | ENSG00000089220 |
| 5 | P00367 | NA | GLUD1 | glutamate dehydrogenase 1 | 2746 | ENSG00000148672 |
| 6 | Q06323 | NA | PSME1 | proteasome (prosome, macropain) activator subunit 1 (PA28 alpha) | 5720 | ENSG00000092010 |
| 7 | P48735 | NA | IDH2 | isocitrate dehydrogenase 2 (NADP+), mitochondrial | 3418 | ENSG00000182054 |
| 8 | Q15019 | NA | SEPT2 | septin 2 | 4735 | ENSG00000168385 |
| 9 | Q9P0J0 | NA | NDUFA13 | NADH dehydrogenase (ubiquinone) 1 alpha subcomplex, 13 | 51079 | ENSG00000186010 |
| 10 | P09211 | NA | GSTP1 | glutathione S-transferase pi 1 | 2950 | ENSG00000084207 |
| 11 | O14773 | NA | TPP1 | tripeptidyl peptidase I | 1200 | ENSG00000166340 |
| 12 | Q05639 | NA | EEF1A2 | eukaryotic translation elongation factor 1 alpha 2 | 1917 | ENSG00000101210 |
| 13 | P22570 | NA | FDXR | ferredoxin reductase | 2232 | ENSG00000161513 |
| 14 | P40121 | NA | CAPG | capping protein (actin filament), gelsolin-like | 822 | ENSG00000042493 |
| 15 | P42765 | NA | ACAA2 | acetyl-CoA acyltransferase 2 | 10449 | ENSG00000167315 |
| 16 | P49419 | NA | ALDH7A1 | aldehyde dehydrogenase 7 family, member A1 | 501 | ENSG00000164904 |
| 17 | P07339 | NA | CTSD | cathepsin D | 1509 | ENSG00000117984 |
| 18 | Q14764 | NA | MVP | major vault protein | 9961 | ENSG00000013364 |
| 19 | Q14697 | NA | GANAB | glucosidase, alpha; neutral AB | 23193 | ENSG00000089597 |
| 20 | P53007 | NA | SLC25A1 | solute carrier family 25 (mitochondrial carrier; citrate transporter), member 1 | 6576 | ENSG00000100075 |
| 21 | Q9BPW8 | NA | NIPSNAP1 | nipsnap homolog 1 (C. elegans) | 8508 | ENSG00000184117 |
| 22 | Q71U36 | NA | TUBA1A | tubulin, alpha 1a | 7846 | ENSG00000167552 |
| 23 | P00338 | NA | LDHA | lactate dehydrogenase A | 3939 | ENSG00000134333 |
| 24 | P38571 | NA | LIPA | lipase A, lysosomal acid, cholesterol esterase | 3988 | ENSG00000107798 |
| 25 | Q13011 | NA | ECH1 | enoyl CoA hydratase 1, peroxisomal | 1891 | ENSG00000104823 |
| 26 | Q14980 | NA | NUMA1 | nuclear mitotic apparatus protein 1 | 4926 | ENSG00000137497 |
| 27 | P04040 | NA | CAT | catalase | 847 | ENSG00000121691 |
| 28 | Q9UBQ7 | NA | GRHPR | glyoxylate reductase/hydroxypyruvate reductase | 9380 | ENSG00000137106 |
| 29 | P51649 | NA | ALDH5A1 | aldehyde dehydrogenase 5 family, member A1 | 7915 | ENSG00000112294 |
| 30 | P30044 | NA | PRDX5 | peroxiredoxin 5 | 25824 | ENSG00000126432 |
| 31 | P24752 | NA | ACAT1 | acetyl-CoA acetyltransferase 1 | 38 | ENSG00000075239 |
| 32 | P13674 | NA | P4HA1 | prolyl 4-hydroxylase, alpha polypeptide I | 5033 | ENSG00000122884 |
| 33 | P16278 | NA | GLB1 | galactosidase, beta 1 | 2720 | ENSG00000170266 |
| 34 | P06744 | NA | GPI | glucose-6-phosphate isomerase | 2821 | ENSG00000105220 |
| 35 | P07602 | NA | PSAP | prosaposin | 5660 | ENSG00000197746 |
| 36 | P49748 | NA | ACADVL | acyl-CoA dehydrogenase, very long chain | 37 | ENSG00000072778 |

  
  

| **Database:cellular component      &nbspName:peroxisomal matrix      &nbspID:GO:0005782** | | | | | | |
| --- | --- | --- | --- | --- | --- | --- |
| C=33; O=3; E=0.08; R=37.91; rawP=6.63e-05; adjP=0.0002 | | | | | | |
| Index | UserID | Value | Gene Symbol | Gene Name | EntrezGene | Ensembl |
| 1 | P04040 | NA | CAT | catalase | 847 | ENSG00000121691 |
| 2 | Q9UBQ7 | NA | GRHPR | glyoxylate reductase/hydroxypyruvate reductase | 9380 | ENSG00000137106 |
| 3 | P30044 | NA | PRDX5 | peroxiredoxin 5 | 25824 | ENSG00000126432 |

  
  

| **Database:cellular component      &nbspName:microbody      &nbspID:GO:0042579** | | | | | | |
| --- | --- | --- | --- | --- | --- | --- |
| C=122; O=4; E=0.29; R=13.67; rawP=0.0002; adjP=0.0004 | | | | | | |
| Index | UserID | Value | Gene Symbol | Gene Name | EntrezGene | Ensembl |
| 1 | P04040 | NA | CAT | catalase | 847 | ENSG00000121691 |
| 2 | Q9UBQ7 | NA | GRHPR | glyoxylate reductase/hydroxypyruvate reductase | 9380 | ENSG00000137106 |
| 3 | P30044 | NA | PRDX5 | peroxiredoxin 5 | 25824 | ENSG00000126432 |
| 4 | Q13011 | NA | ECH1 | enoyl CoA hydratase 1, peroxisomal | 1891 | ENSG00000104823 |

  
  

| **Database:cellular component      &nbspName:peroxisome      &nbspID:GO:0005777** | | | | | | |
| --- | --- | --- | --- | --- | --- | --- |
| C=122; O=4; E=0.29; R=13.67; rawP=0.0002; adjP=0.0004 | | | | | | |
| Index | UserID | Value | Gene Symbol | Gene Name | EntrezGene | Ensembl |
| 1 | P04040 | NA | CAT | catalase | 847 | ENSG00000121691 |
| 2 | Q9UBQ7 | NA | GRHPR | glyoxylate reductase/hydroxypyruvate reductase | 9380 | ENSG00000137106 |
| 3 | P30044 | NA | PRDX5 | peroxiredoxin 5 | 25824 | ENSG00000126432 |
| 4 | Q13011 | NA | ECH1 | enoyl CoA hydratase 1, peroxisomal | 1891 | ENSG00000104823 |

  
  

| **Database:cellular component      &nbspName:vacuolar part      &nbspID:GO:0044437** | | | | | | |
| --- | --- | --- | --- | --- | --- | --- |
| C=253; O=5; E=0.61; R=8.24; rawP=0.0003; adjP=0.0006 | | | | | | |
| Index | UserID | Value | Gene Symbol | Gene Name | EntrezGene | Ensembl |
| 1 | O14773 | NA | TPP1 | tripeptidyl peptidase I | 1200 | ENSG00000166340 |
| 2 | P16278 | NA | GLB1 | galactosidase, beta 1 | 2720 | ENSG00000170266 |
| 3 | P10253 | NA | GAA | glucosidase, alpha; acid | 2548 | ENSG00000171298 |
| 4 | P07602 | NA | PSAP | prosaposin | 5660 | ENSG00000197746 |
| 5 | P07339 | NA | CTSD | cathepsin D | 1509 | ENSG00000117984 |

  
  
  
  

---

WebGestalt is currently developed and maintained by Jing Wang and Bing Zhang at the  Zhang Lab. Other people who have made significant contribution to the project include Dexter Duncan, Stefan Kirov, Zhiao Shi, and Jay Snoddy.  
  
**Funding credits:** NIH/NIAAA (U01 AA016662, U01 AA013512); NIH/NIDA (P01 DA015027); NIH/NIMH (P50 MH078028, P50 MH096972); NIH/NCI (U24 CA159988); NIH/NIGMS (R01 GM088822).
