## Supplementary material for "PGRMC1 phosphorylation and cell plasticity 1: glycolysis, mitochondria, tumor growth": File S7: final_sig_kegg_file_1438746600.html

Anchored HTML File of EIDs

|  |  |
| --- | --- |
|  | WEB-based GEne SeT AnaLysis Toolkit |
|  | ***Translating gene lists into biological insights...*** |

---

  

| **Database:KEGG pathway      &nbspName:Metabolic pathways      &nbspID:01100** | | | | | | |
| --- | --- | --- | --- | --- | --- | --- |
| C=1130; O=16; E=2.23; R=7.18; rawP=1.37e-10; adjP=1.64e-09 | | | | | | |
| Index | UserID | Value | Gene Symbol | Gene Name | EntrezGene | Ensembl |
| 1 | P10253 | NA | GAA | glucosidase, alpha; acid | 2548 | ENSG00000171298 |
| 2 | Q16851 | NA | UGP2 | UDP-glucose pyrophosphorylase 2 | 7360 | ENSG00000169764 |
| 3 | P00338 | NA | LDHA | lactate dehydrogenase A | 3939 | ENSG00000134333 |
| 4 | P00367 | NA | GLUD1 | glutamate dehydrogenase 1 | 2746 | ENSG00000148672 |
| 5 | P48735 | NA | IDH2 | isocitrate dehydrogenase 2 (NADP+), mitochondrial | 3418 | ENSG00000182054 |
| 6 | P04040 | NA | CAT | catalase | 847 | ENSG00000121691 |
| 7 | Q9UBQ7 | NA | GRHPR | glyoxylate reductase/hydroxypyruvate reductase | 9380 | ENSG00000137106 |
| 8 | P51649 | NA | ALDH5A1 | aldehyde dehydrogenase 5 family, member A1 | 7915 | ENSG00000112294 |
| 9 | P42765 | NA | ACAA2 | acetyl-CoA acyltransferase 2 | 10449 | ENSG00000167315 |
| 10 | P13674 | NA | P4HA1 | prolyl 4-hydroxylase, alpha polypeptide I | 5033 | ENSG00000122884 |
| 11 | P24752 | NA | ACAT1 | acetyl-CoA acetyltransferase 1 | 38 | ENSG00000075239 |
| 12 | P49419 | NA | ALDH7A1 | aldehyde dehydrogenase 7 family, member A1 | 501 | ENSG00000164904 |
| 13 | P16278 | NA | GLB1 | galactosidase, beta 1 | 2720 | ENSG00000170266 |
| 14 | P06744 | NA | GPI | glucose-6-phosphate isomerase | 2821 | ENSG00000105220 |
| 15 | P49748 | NA | ACADVL | acyl-CoA dehydrogenase, very long chain | 37 | ENSG00000072778 |
| 16 | Q14697 | NA | GANAB | glucosidase, alpha; neutral AB | 23193 | ENSG00000089597 |

  
  

| **Database:KEGG pathway      &nbspName:Peroxisome      &nbspID:04146** | | | | | | |
| --- | --- | --- | --- | --- | --- | --- |
| C=79; O=4; E=0.16; R=25.67; rawP=1.75e-05; adjP=3.45e-05 | | | | | | |
| Index | UserID | Value | Gene Symbol | Gene Name | EntrezGene | Ensembl |
| 1 | P04040 | NA | CAT | catalase | 847 | ENSG00000121691 |
| 2 | P48735 | NA | IDH2 | isocitrate dehydrogenase 2 (NADP+), mitochondrial | 3418 | ENSG00000182054 |
| 3 | P30044 | NA | PRDX5 | peroxiredoxin 5 | 25824 | ENSG00000126432 |
| 4 | Q13011 | NA | ECH1 | enoyl CoA hydratase 1, peroxisomal | 1891 | ENSG00000104823 |

  
  

| **Database:KEGG pathway      &nbspName:Propanoate metabolism      &nbspID:00640** | | | | | | |
| --- | --- | --- | --- | --- | --- | --- |
| C=32; O=3; E=0.06; R=47.54; rawP=3.39e-05; adjP=5.09e-05 | | | | | | |
| Index | UserID | Value | Gene Symbol | Gene Name | EntrezGene | Ensembl |
| 1 | P24752 | NA | ACAT1 | acetyl-CoA acetyltransferase 1 | 38 | ENSG00000075239 |
| 2 | P49419 | NA | ALDH7A1 | aldehyde dehydrogenase 7 family, member A1 | 501 | ENSG00000164904 |
| 3 | P00338 | NA | LDHA | lactate dehydrogenase A | 3939 | ENSG00000134333 |

  
  

| **Database:KEGG pathway      &nbspName:Valine, leucine and isoleucine degradation      &nbspID:00280** | | | | | | |
| --- | --- | --- | --- | --- | --- | --- |
| C=44; O=3; E=0.09; R=34.57; rawP=8.90e-05; adjP=0.0001 | | | | | | |
| Index | UserID | Value | Gene Symbol | Gene Name | EntrezGene | Ensembl |
| 1 | P24752 | NA | ACAT1 | acetyl-CoA acetyltransferase 1 | 38 | ENSG00000075239 |
| 2 | P49419 | NA | ALDH7A1 | aldehyde dehydrogenase 7 family, member A1 | 501 | ENSG00000164904 |
| 3 | P42765 | NA | ACAA2 | acetyl-CoA acyltransferase 2 | 10449 | ENSG00000167315 |

  
  

| **Database:KEGG pathway      &nbspName:Starch and sucrose metabolism      &nbspID:00500** | | | | | | |
| --- | --- | --- | --- | --- | --- | --- |
| C=54; O=3; E=0.11; R=28.17; rawP=0.0002; adjP=0.0002 | | | | | | |
| Index | UserID | Value | Gene Symbol | Gene Name | EntrezGene | Ensembl |
| 1 | P10253 | NA | GAA | glucosidase, alpha; acid | 2548 | ENSG00000171298 |
| 2 | P06744 | NA | GPI | glucose-6-phosphate isomerase | 2821 | ENSG00000105220 |
| 3 | Q16851 | NA | UGP2 | UDP-glucose pyrophosphorylase 2 | 7360 | ENSG00000169764 |

  
  

| **Database:KEGG pathway      &nbspName:Arginine and proline metabolism      &nbspID:00330** | | | | | | |
| --- | --- | --- | --- | --- | --- | --- |
| C=54; O=3; E=0.11; R=28.17; rawP=0.0002; adjP=0.0002 | | | | | | |
| Index | UserID | Value | Gene Symbol | Gene Name | EntrezGene | Ensembl |
| 1 | P13674 | NA | P4HA1 | prolyl 4-hydroxylase, alpha polypeptide I | 5033 | ENSG00000122884 |
| 2 | P49419 | NA | ALDH7A1 | aldehyde dehydrogenase 7 family, member A1 | 501 | ENSG00000164904 |
| 3 | P00367 | NA | GLUD1 | glutamate dehydrogenase 1 | 2746 | ENSG00000148672 |

  
  

| **Database:KEGG pathway      &nbspName:Glycolysis / Gluconeogenesis      &nbspID:00010** | | | | | | |
| --- | --- | --- | --- | --- | --- | --- |
| C=65; O=3; E=0.13; R=23.40; rawP=0.0003; adjP=0.0003 | | | | | | |
| Index | UserID | Value | Gene Symbol | Gene Name | EntrezGene | Ensembl |
| 1 | P49419 | NA | ALDH7A1 | aldehyde dehydrogenase 7 family, member A1 | 501 | ENSG00000164904 |
| 2 | P06744 | NA | GPI | glucose-6-phosphate isomerase | 2821 | ENSG00000105220 |
| 3 | P00338 | NA | LDHA | lactate dehydrogenase A | 3939 | ENSG00000134333 |

  
  

  
  

---

WebGestalt is currently developed and maintained by Jing Wang and Bing Zhang at the  Zhang Lab. Other people who have made significant contribution to the project include Dexter Duncan, Stefan Kirov, Zhiao Shi, and Jay Snoddy.  
  
**Funding credits:** NIH/NIAAA (U01 AA016662, U01 AA013512); NIH/NIDA (P01 DA015027); NIH/NIMH (P50 MH078028, P50 MH096972); NIH/NCI (U24 CA159988); NIH/NIGMS (R01 GM088822).
