## Supplementary material for "PGRMC1 phosphorylation and cell plasticity 1: glycolysis, mitochondria, tumor growth": File S7: final_sig_file_1438748351.html

Anchored HTML File of EIDs

|  |  |
| --- | --- |
|  | WEB-based GEne SeT AnaLysis Toolkit |
|  | ***Translating gene lists into biological insights...*** |

---

  

| **Database:molecular function      &nbspName:translation factor activity, nucleic acid binding      &nbspID:GO:0008135** | | | | | | |
| --- | --- | --- | --- | --- | --- | --- |
| C=79; O=5; E=0.23; R=21.54; rawP=3.32e-06; adjP=0.0001 | | | | | | |
| Index | UserID | Value | Gene Symbol | Gene Name | EntrezGene | Ensembl |
| 1 | Q92616 | NA | GCN1L1 | GCN1 general control of amino-acid synthesis 1-like 1 (yeast) | 10985 | ENSG00000089154 |
| 2 | P20042 | NA | EIF2S2 | eukaryotic translation initiation factor 2, subunit 2 beta, 38kDa | 8894 | ENSG00000125977 |
| 3 | P41091 | NA | EIF2S3 | eukaryotic translation initiation factor 2, subunit 3 gamma, 52kDa | 1968 | ENSG00000130741 |
| 4 | Q15056 | NA | EIF4H | eukaryotic translation initiation factor 4H | 7458 | ENSG00000106682 |
| 5 | P68104 | NA | EEF1A1 | eukaryotic translation elongation factor 1 alpha 1 | 1915 | ENSG00000156508 |

  
  

| **Database:molecular function      &nbspName:RNA binding      &nbspID:GO:0003723** | | | | | | |
| --- | --- | --- | --- | --- | --- | --- |
| C=843; O=12; E=2.48; R=4.85; rawP=3.70e-06; adjP=0.0001 | | | | | | |
| Index | UserID | Value | Gene Symbol | Gene Name | EntrezGene | Ensembl |
| 1 | Q9NR30 | NA | DDX21 | DEAD (Asp-Glu-Ala-Asp) box helicase 21 | 9188 | ENSG00000165732 |
| 2 | P49591 | NA | SARS | seryl-tRNA synthetase | 6301 | ENSG00000031698 |
| 3 | P20042 | NA | EIF2S2 | eukaryotic translation initiation factor 2, subunit 2 beta, 38kDa | 8894 | ENSG00000125977 |
| 4 | P41091 | NA | EIF2S3 | eukaryotic translation initiation factor 2, subunit 3 gamma, 52kDa | 1968 | ENSG00000130741 |
| 5 | Q9HAV4 | NA | XPO5 | exportin 5 | 57510 | ENSG00000124571 |
| 6 | Q92616 | NA | GCN1L1 | GCN1 general control of amino-acid synthesis 1-like 1 (yeast) | 10985 | ENSG00000089154 |
| 7 | P56192 | NA | MARS | methionyl-tRNA synthetase | 4141 | ENSG00000166986 |
| 8 | Q15056 | NA | EIF4H | eukaryotic translation initiation factor 4H | 7458 | ENSG00000106682 |
| 9 | P49588 | NA | AARS | alanyl-tRNA synthetase | 16 | ENSG00000090861 |
| 10 | P68104 | NA | EEF1A1 | eukaryotic translation elongation factor 1 alpha 1 | 1915 | ENSG00000156508 |
| 11 | P04083 | NA | ANXA1 | annexin A1 | 301 | ENSG00000135046 |
| 12 | O00425 | NA | IGF2BP3 | insulin-like growth factor 2 mRNA binding protein 3 | 10643 | ENSG00000136231 |

  
  

| **Database:molecular function      &nbspName:small molecule binding      &nbspID:GO:0036094** | | | | | | |
| --- | --- | --- | --- | --- | --- | --- |
| C=2595; O=20; E=7.62; R=2.62; rawP=1.37e-05; adjP=0.0003 | | | | | | |
| Index | UserID | Value | Gene Symbol | Gene Name | EntrezGene | Ensembl |
| 1 | Q16881 | NA | TXNRD1 | thioredoxin reductase 1 | 7296 | ENSG00000198431 |
| 2 | Q9NR30 | NA | DDX21 | DEAD (Asp-Glu-Ala-Asp) box helicase 21 | 9188 | ENSG00000165732 |
| 3 | P49591 | NA | SARS | seryl-tRNA synthetase | 6301 | ENSG00000031698 |
| 4 | Q92598 | NA | HSPH1 | heat shock 105kDa/110kDa protein 1 | 10808 | ENSG00000120694 |
| 5 | P00491 | NA | PNP | purine nucleoside phosphorylase | 4860 | ENSG00000198805 |
| 6 | P61225 | NA | RAP2B | RAP2B, member of RAS oncogene family | 5912 | ENSG00000181467 |
| 7 | P56192 | NA | MARS | methionyl-tRNA synthetase | 4141 | ENSG00000166986 |
| 8 | P49736 | NA | MCM2 | minichromosome maintenance complex component 2 | 4171 | ENSG00000073111 |
| 9 | Q15056 | NA | EIF4H | eukaryotic translation initiation factor 4H | 7458 | ENSG00000106682 |
| 10 | P33993 | NA | MCM7 | minichromosome maintenance complex component 7 | 4176 | ENSG00000166508 |
| 11 | P49588 | NA | AARS | alanyl-tRNA synthetase | 16 | ENSG00000090861 |
| 12 | P68104 | NA | EEF1A1 | eukaryotic translation elongation factor 1 alpha 1 | 1915 | ENSG00000156508 |
| 13 | O00425 | NA | IGF2BP3 | insulin-like growth factor 2 mRNA binding protein 3 | 10643 | ENSG00000136231 |
| 14 | P22102 | NA | GART | phosphoribosylglycinamide formyltransferase, phosphoribosylglycinamide synthetase, phosphoribosylaminoimidazole synthetase | 2618 | ENSG00000159131 |
| 15 | P16615 | NA | ATP2A2 | ATPase, Ca++ transporting, cardiac muscle, slow twitch 2 | 488 | ENSG00000174437 |
| 16 | P41091 | NA | EIF2S3 | eukaryotic translation initiation factor 2, subunit 3 gamma, 52kDa | 1968 | ENSG00000130741 |
| 17 | P02768 | NA | ALB | albumin | 213 | ENSG00000163631 |
| 18 | P17174 | NA | GOT1 | glutamic-oxaloacetic transaminase 1, soluble (aspartate aminotransferase 1) | 2805 | ENSG00000120053 |
| 19 | P21589 | NA | NT5E | 5'-nucleotidase, ecto (CD73) | 4907 | ENSG00000135318 |
| 20 | P11387 | NA | TOP1 | topoisomerase (DNA) I | 7150 | ENSG00000198900 |

  
  

| **Database:molecular function      &nbspName:antioxidant activity      &nbspID:GO:0016209** | | | | | | |
| --- | --- | --- | --- | --- | --- | --- |
| C=61; O=4; E=0.18; R=22.32; rawP=2.99e-05; adjP=0.0005 | | | | | | |
| Index | UserID | Value | Gene Symbol | Gene Name | EntrezGene | Ensembl |
| 1 | Q16881 | NA | TXNRD1 | thioredoxin reductase 1 | 7296 | ENSG00000198431 |
| 2 | P78417 | NA | GSTO1 | glutathione S-transferase omega 1 | 9446 | ENSG00000148834 |
| 3 | P02768 | NA | ALB | albumin | 213 | ENSG00000163631 |
| 4 | Q06830 | NA | PRDX1 | peroxiredoxin 1 | 5052 | ENSG00000117450 |

  
  

| **Database:cellular component      &nbspName:cytosol      &nbspID:GO:0005829** | | | | | | |
| --- | --- | --- | --- | --- | --- | --- |
| C=2367; O=21; E=6.53; R=3.22; rawP=2.74e-07; adjP=9.58e-06 | | | | | | |
| Index | UserID | Value | Gene Symbol | Gene Name | EntrezGene | Ensembl |
| 1 | Q16881 | NA | TXNRD1 | thioredoxin reductase 1 | 7296 | ENSG00000198431 |
| 2 | Q04446 | NA | GBE1 | glucan (1,4-alpha-), branching enzyme 1 | 2632 | ENSG00000114480 |
| 3 | P49591 | NA | SARS | seryl-tRNA synthetase | 6301 | ENSG00000031698 |
| 4 | P00491 | NA | PNP | purine nucleoside phosphorylase | 4860 | ENSG00000198805 |
| 5 | Q9HAV4 | NA | XPO5 | exportin 5 | 57510 | ENSG00000124571 |
| 6 | P61225 | NA | RAP2B | RAP2B, member of RAS oncogene family | 5912 | ENSG00000181467 |
| 7 | P56192 | NA | MARS | methionyl-tRNA synthetase | 4141 | ENSG00000166986 |
| 8 | P78417 | NA | GSTO1 | glutathione S-transferase omega 1 | 9446 | ENSG00000148834 |
| 9 | Q15056 | NA | EIF4H | eukaryotic translation initiation factor 4H | 7458 | ENSG00000106682 |
| 10 | P16949 | NA | STMN1 | stathmin 1 | 3925 | ENSG00000117632 |
| 11 | P49588 | NA | AARS | alanyl-tRNA synthetase | 16 | ENSG00000090861 |
| 12 | P68104 | NA | EEF1A1 | eukaryotic translation elongation factor 1 alpha 1 | 1915 | ENSG00000156508 |
| 13 | O75369 | NA | FLNB | filamin B, beta | 2317 | ENSG00000136068 |
| 14 | P43490 | NA | NAMPT | nicotinamide phosphoribosyltransferase | 10135 | ENSG00000105835 |
| 15 | O00425 | NA | IGF2BP3 | insulin-like growth factor 2 mRNA binding protein 3 | 10643 | ENSG00000136231 |
| 16 | P22102 | NA | GART | phosphoribosylglycinamide formyltransferase, phosphoribosylglycinamide synthetase, phosphoribosylaminoimidazole synthetase | 2618 | ENSG00000159131 |
| 17 | Q9P258 | NA | RCC2 | regulator of chromosome condensation 2 | 55920 | ENSG00000179051 |
| 18 | P20042 | NA | EIF2S2 | eukaryotic translation initiation factor 2, subunit 2 beta, 38kDa | 8894 | ENSG00000125977 |
| 19 | P41091 | NA | EIF2S3 | eukaryotic translation initiation factor 2, subunit 3 gamma, 52kDa | 1968 | ENSG00000130741 |
| 20 | P17174 | NA | GOT1 | glutamic-oxaloacetic transaminase 1, soluble (aspartate aminotransferase 1) | 2805 | ENSG00000120053 |
| 21 | P18206 | NA | VCL | vinculin | 7414 | ENSG00000035403 |

  
  

| **Database:cellular component      &nbspName:intracellular part      &nbspID:GO:0044424** | | | | | | |
| --- | --- | --- | --- | --- | --- | --- |
| C=12096; O=46; E=33.35; R=1.38; rawP=3.70e-07; adjP=9.58e-06 | | | | | | |
| Index | UserID | Value | Gene Symbol | Gene Name | EntrezGene | Ensembl |
| 1 | Q04446 | NA | GBE1 | glucan (1,4-alpha-), branching enzyme 1 | 2632 | ENSG00000114480 |
| 2 | P49591 | NA | SARS | seryl-tRNA synthetase | 6301 | ENSG00000031698 |
| 3 | Q92598 | NA | HSPH1 | heat shock 105kDa/110kDa protein 1 | 10808 | ENSG00000120694 |
| 4 | P61225 | NA | RAP2B | RAP2B, member of RAS oncogene family | 5912 | ENSG00000181467 |
| 5 | P55060 | NA | CSE1L | CSE1 chromosome segregation 1-like (yeast) | 1434 | ENSG00000124207 |
| 6 | Q9NQC3 | NA | RTN4 | reticulon 4 | 57142 | ENSG00000115310 |
| 7 | P16949 | NA | STMN1 | stathmin 1 | 3925 | ENSG00000117632 |
| 8 | P33993 | NA | MCM7 | minichromosome maintenance complex component 7 | 4176 | ENSG00000166508 |
| 9 | P68104 | NA | EEF1A1 | eukaryotic translation elongation factor 1 alpha 1 | 1915 | ENSG00000156508 |
| 10 | Q86UP2 | NA | KTN1 | kinectin 1 (kinesin receptor) | 3895 | ENSG00000126777 |
| 11 | P49588 | NA | AARS | alanyl-tRNA synthetase | 16 | ENSG00000090861 |
| 12 | P02786 | NA | TFRC | transferrin receptor (p90, CD71) | 7037 | ENSG00000072274 |
| 13 | O75369 | NA | FLNB | filamin B, beta | 2317 | ENSG00000136068 |
| 14 | P43490 | NA | NAMPT | nicotinamide phosphoribosyltransferase | 10135 | ENSG00000105835 |
| 15 | O00425 | NA | IGF2BP3 | insulin-like growth factor 2 mRNA binding protein 3 | 10643 | ENSG00000136231 |
| 16 | P17655 | NA | CAPN2 | calpain 2, (m/II) large subunit | 824 | ENSG00000162909 |
| 17 | P20042 | NA | EIF2S2 | eukaryotic translation initiation factor 2, subunit 2 beta, 38kDa | 8894 | ENSG00000125977 |
| 18 | Q9P258 | NA | RCC2 | regulator of chromosome condensation 2 | 55920 | ENSG00000179051 |
| 19 | P22102 | NA | GART | phosphoribosylglycinamide formyltransferase, phosphoribosylglycinamide synthetase, phosphoribosylaminoimidazole synthetase | 2618 | ENSG00000159131 |
| 20 | P16615 | NA | ATP2A2 | ATPase, Ca++ transporting, cardiac muscle, slow twitch 2 | 488 | ENSG00000174437 |
| 21 | Q92616 | NA | GCN1L1 | GCN1 general control of amino-acid synthesis 1-like 1 (yeast) | 10985 | ENSG00000089154 |
| 22 | P17174 | NA | GOT1 | glutamic-oxaloacetic transaminase 1, soluble (aspartate aminotransferase 1) | 2805 | ENSG00000120053 |
| 23 | P37802 | NA | TAGLN2 | transgelin 2 | 8407 | ENSG00000158710 |
| 24 | P08727 | NA | KRT19 | keratin 19 | 3880 | ENSG00000171345 |
| 25 | Q6PIU2 | NA | NCEH1 | neutral cholesterol ester hydrolase 1 | 57552 | ENSG00000144959 |
| 26 | P18206 | NA | VCL | vinculin | 7414 | ENSG00000035403 |
| 27 | P11387 | NA | TOP1 | topoisomerase (DNA) I | 7150 | ENSG00000198900 |
| 28 | Q9UK76 | NA | HN1 | hematological and neurological expressed 1 | 51155 | ENSG00000189159 |
| 29 | P21589 | NA | NT5E | 5'-nucleotidase, ecto (CD73) | 4907 | ENSG00000135318 |
| 30 | Q16881 | NA | TXNRD1 | thioredoxin reductase 1 | 7296 | ENSG00000198431 |
| 31 | Q9NR30 | NA | DDX21 | DEAD (Asp-Glu-Ala-Asp) box helicase 21 | 9188 | ENSG00000165732 |
| 32 | P08195 | NA | SLC3A2 | solute carrier family 3 (activators of dibasic and neutral amino acid transport), member 2 | 6520 | ENSG00000168003 |
| 33 | P00491 | NA | PNP | purine nucleoside phosphorylase | 4860 | ENSG00000198805 |
| 34 | P49321 | NA | NASP | nuclear autoantigenic sperm protein (histone-binding) | 4678 | ENSG00000132780 |
| 35 | Q9HAV4 | NA | XPO5 | exportin 5 | 57510 | ENSG00000124571 |
| 36 | P56192 | NA | MARS | methionyl-tRNA synthetase | 4141 | ENSG00000166986 |
| 37 | P78417 | NA | GSTO1 | glutathione S-transferase omega 1 | 9446 | ENSG00000148834 |
| 38 | P49736 | NA | MCM2 | minichromosome maintenance complex component 2 | 4171 | ENSG00000073111 |
| 39 | Q15056 | NA | EIF4H | eukaryotic translation initiation factor 4H | 7458 | ENSG00000106682 |
| 40 | O60610 | NA | DIAPH1 | diaphanous homolog 1 (Drosophila) | 1729 | ENSG00000131504 |
| 41 | Q06830 | NA | PRDX1 | peroxiredoxin 1 | 5052 | ENSG00000117450 |
| 42 | P04083 | NA | ANXA1 | annexin A1 | 301 | ENSG00000135046 |
| 43 | P41091 | NA | EIF2S3 | eukaryotic translation initiation factor 2, subunit 3 gamma, 52kDa | 1968 | ENSG00000130741 |
| 44 | P02768 | NA | ALB | albumin | 213 | ENSG00000163631 |
| 45 | P05556 | NA | ITGB1 | integrin, beta 1 (fibronectin receptor, beta polypeptide, antigen CD29 includes MDF2, MSK12) | 3688 | ENSG00000150093 |
| 46 | Q15758 | NA | SLC1A5 | solute carrier family 1 (neutral amino acid transporter), member 5 | 6510 | ENSG00000105281 |

  
  

| **Database:cellular component      &nbspName:cytoplasm      &nbspID:GO:0005737** | | | | | | |
| --- | --- | --- | --- | --- | --- | --- |
| C=9051; O=41; E=24.96; R=1.64; rawP=3.99e-07; adjP=9.58e-06 | | | | | | |
| Index | UserID | Value | Gene Symbol | Gene Name | EntrezGene | Ensembl |
| 1 | Q04446 | NA | GBE1 | glucan (1,4-alpha-), branching enzyme 1 | 2632 | ENSG00000114480 |
| 2 | P49591 | NA | SARS | seryl-tRNA synthetase | 6301 | ENSG00000031698 |
| 3 | Q92598 | NA | HSPH1 | heat shock 105kDa/110kDa protein 1 | 10808 | ENSG00000120694 |
| 4 | P61225 | NA | RAP2B | RAP2B, member of RAS oncogene family | 5912 | ENSG00000181467 |
| 5 | P55060 | NA | CSE1L | CSE1 chromosome segregation 1-like (yeast) | 1434 | ENSG00000124207 |
| 6 | Q9NQC3 | NA | RTN4 | reticulon 4 | 57142 | ENSG00000115310 |
| 7 | P16949 | NA | STMN1 | stathmin 1 | 3925 | ENSG00000117632 |
| 8 | P68104 | NA | EEF1A1 | eukaryotic translation elongation factor 1 alpha 1 | 1915 | ENSG00000156508 |
| 9 | P49588 | NA | AARS | alanyl-tRNA synthetase | 16 | ENSG00000090861 |
| 10 | Q86UP2 | NA | KTN1 | kinectin 1 (kinesin receptor) | 3895 | ENSG00000126777 |
| 11 | P02786 | NA | TFRC | transferrin receptor (p90, CD71) | 7037 | ENSG00000072274 |
| 12 | O75369 | NA | FLNB | filamin B, beta | 2317 | ENSG00000136068 |
| 13 | P43490 | NA | NAMPT | nicotinamide phosphoribosyltransferase | 10135 | ENSG00000105835 |
| 14 | O00425 | NA | IGF2BP3 | insulin-like growth factor 2 mRNA binding protein 3 | 10643 | ENSG00000136231 |
| 15 | P17655 | NA | CAPN2 | calpain 2, (m/II) large subunit | 824 | ENSG00000162909 |
| 16 | P20042 | NA | EIF2S2 | eukaryotic translation initiation factor 2, subunit 2 beta, 38kDa | 8894 | ENSG00000125977 |
| 17 | P22102 | NA | GART | phosphoribosylglycinamide formyltransferase, phosphoribosylglycinamide synthetase, phosphoribosylaminoimidazole synthetase | 2618 | ENSG00000159131 |
| 18 | Q9P258 | NA | RCC2 | regulator of chromosome condensation 2 | 55920 | ENSG00000179051 |
| 19 | P16615 | NA | ATP2A2 | ATPase, Ca++ transporting, cardiac muscle, slow twitch 2 | 488 | ENSG00000174437 |
| 20 | Q92616 | NA | GCN1L1 | GCN1 general control of amino-acid synthesis 1-like 1 (yeast) | 10985 | ENSG00000089154 |
| 21 | P17174 | NA | GOT1 | glutamic-oxaloacetic transaminase 1, soluble (aspartate aminotransferase 1) | 2805 | ENSG00000120053 |
| 22 | P08727 | NA | KRT19 | keratin 19 | 3880 | ENSG00000171345 |
| 23 | Q6PIU2 | NA | NCEH1 | neutral cholesterol ester hydrolase 1 | 57552 | ENSG00000144959 |
| 24 | P18206 | NA | VCL | vinculin | 7414 | ENSG00000035403 |
| 25 | P11387 | NA | TOP1 | topoisomerase (DNA) I | 7150 | ENSG00000198900 |
| 26 | P21589 | NA | NT5E | 5'-nucleotidase, ecto (CD73) | 4907 | ENSG00000135318 |
| 27 | Q16881 | NA | TXNRD1 | thioredoxin reductase 1 | 7296 | ENSG00000198431 |
| 28 | P08195 | NA | SLC3A2 | solute carrier family 3 (activators of dibasic and neutral amino acid transport), member 2 | 6520 | ENSG00000168003 |
| 29 | P00491 | NA | PNP | purine nucleoside phosphorylase | 4860 | ENSG00000198805 |
| 30 | P49321 | NA | NASP | nuclear autoantigenic sperm protein (histone-binding) | 4678 | ENSG00000132780 |
| 31 | Q9HAV4 | NA | XPO5 | exportin 5 | 57510 | ENSG00000124571 |
| 32 | P56192 | NA | MARS | methionyl-tRNA synthetase | 4141 | ENSG00000166986 |
| 33 | P78417 | NA | GSTO1 | glutathione S-transferase omega 1 | 9446 | ENSG00000148834 |
| 34 | Q15056 | NA | EIF4H | eukaryotic translation initiation factor 4H | 7458 | ENSG00000106682 |
| 35 | O60610 | NA | DIAPH1 | diaphanous homolog 1 (Drosophila) | 1729 | ENSG00000131504 |
| 36 | Q06830 | NA | PRDX1 | peroxiredoxin 1 | 5052 | ENSG00000117450 |
| 37 | P04083 | NA | ANXA1 | annexin A1 | 301 | ENSG00000135046 |
| 38 | P41091 | NA | EIF2S3 | eukaryotic translation initiation factor 2, subunit 3 gamma, 52kDa | 1968 | ENSG00000130741 |
| 39 | P02768 | NA | ALB | albumin | 213 | ENSG00000163631 |
| 40 | P05556 | NA | ITGB1 | integrin, beta 1 (fibronectin receptor, beta polypeptide, antigen CD29 includes MDF2, MSK12) | 3688 | ENSG00000150093 |
| 41 | Q15758 | NA | SLC1A5 | solute carrier family 1 (neutral amino acid transporter), member 5 | 6510 | ENSG00000105281 |

  
  

| **Database:cellular component      &nbspName:cytoplasmic part      &nbspID:GO:0044444** | | | | | | |
| --- | --- | --- | --- | --- | --- | --- |
| C=6728; O=35; E=18.55; R=1.89; rawP=8.68e-07; adjP=1.56e-05 | | | | | | |
| Index | UserID | Value | Gene Symbol | Gene Name | EntrezGene | Ensembl |
| 1 | Q04446 | NA | GBE1 | glucan (1,4-alpha-), branching enzyme 1 | 2632 | ENSG00000114480 |
| 2 | P49591 | NA | SARS | seryl-tRNA synthetase | 6301 | ENSG00000031698 |
| 3 | P61225 | NA | RAP2B | RAP2B, member of RAS oncogene family | 5912 | ENSG00000181467 |
| 4 | Q9NQC3 | NA | RTN4 | reticulon 4 | 57142 | ENSG00000115310 |
| 5 | P16949 | NA | STMN1 | stathmin 1 | 3925 | ENSG00000117632 |
| 6 | P49588 | NA | AARS | alanyl-tRNA synthetase | 16 | ENSG00000090861 |
| 7 | Q86UP2 | NA | KTN1 | kinectin 1 (kinesin receptor) | 3895 | ENSG00000126777 |
| 8 | P68104 | NA | EEF1A1 | eukaryotic translation elongation factor 1 alpha 1 | 1915 | ENSG00000156508 |
| 9 | P02786 | NA | TFRC | transferrin receptor (p90, CD71) | 7037 | ENSG00000072274 |
| 10 | O75369 | NA | FLNB | filamin B, beta | 2317 | ENSG00000136068 |
| 11 | P43490 | NA | NAMPT | nicotinamide phosphoribosyltransferase | 10135 | ENSG00000105835 |
| 12 | O00425 | NA | IGF2BP3 | insulin-like growth factor 2 mRNA binding protein 3 | 10643 | ENSG00000136231 |
| 13 | P22102 | NA | GART | phosphoribosylglycinamide formyltransferase, phosphoribosylglycinamide synthetase, phosphoribosylaminoimidazole synthetase | 2618 | ENSG00000159131 |
| 14 | Q9P258 | NA | RCC2 | regulator of chromosome condensation 2 | 55920 | ENSG00000179051 |
| 15 | P20042 | NA | EIF2S2 | eukaryotic translation initiation factor 2, subunit 2 beta, 38kDa | 8894 | ENSG00000125977 |
| 16 | P16615 | NA | ATP2A2 | ATPase, Ca++ transporting, cardiac muscle, slow twitch 2 | 488 | ENSG00000174437 |
| 17 | Q92616 | NA | GCN1L1 | GCN1 general control of amino-acid synthesis 1-like 1 (yeast) | 10985 | ENSG00000089154 |
| 18 | P17174 | NA | GOT1 | glutamic-oxaloacetic transaminase 1, soluble (aspartate aminotransferase 1) | 2805 | ENSG00000120053 |
| 19 | P08727 | NA | KRT19 | keratin 19 | 3880 | ENSG00000171345 |
| 20 | Q6PIU2 | NA | NCEH1 | neutral cholesterol ester hydrolase 1 | 57552 | ENSG00000144959 |
| 21 | P18206 | NA | VCL | vinculin | 7414 | ENSG00000035403 |
| 22 | P11387 | NA | TOP1 | topoisomerase (DNA) I | 7150 | ENSG00000198900 |
| 23 | Q16881 | NA | TXNRD1 | thioredoxin reductase 1 | 7296 | ENSG00000198431 |
| 24 | P08195 | NA | SLC3A2 | solute carrier family 3 (activators of dibasic and neutral amino acid transport), member 2 | 6520 | ENSG00000168003 |
| 25 | P00491 | NA | PNP | purine nucleoside phosphorylase | 4860 | ENSG00000198805 |
| 26 | Q9HAV4 | NA | XPO5 | exportin 5 | 57510 | ENSG00000124571 |
| 27 | P78417 | NA | GSTO1 | glutathione S-transferase omega 1 | 9446 | ENSG00000148834 |
| 28 | P56192 | NA | MARS | methionyl-tRNA synthetase | 4141 | ENSG00000166986 |
| 29 | Q15056 | NA | EIF4H | eukaryotic translation initiation factor 4H | 7458 | ENSG00000106682 |
| 30 | Q06830 | NA | PRDX1 | peroxiredoxin 1 | 5052 | ENSG00000117450 |
| 31 | P04083 | NA | ANXA1 | annexin A1 | 301 | ENSG00000135046 |
| 32 | P02768 | NA | ALB | albumin | 213 | ENSG00000163631 |
| 33 | P41091 | NA | EIF2S3 | eukaryotic translation initiation factor 2, subunit 3 gamma, 52kDa | 1968 | ENSG00000130741 |
| 34 | P05556 | NA | ITGB1 | integrin, beta 1 (fibronectin receptor, beta polypeptide, antigen CD29 includes MDF2, MSK12) | 3688 | ENSG00000150093 |
| 35 | Q15758 | NA | SLC1A5 | solute carrier family 1 (neutral amino acid transporter), member 5 | 6510 | ENSG00000105281 |

  
  

| **Database:cellular component      &nbspName:intracellular      &nbspID:GO:0005622** | | | | | | |
| --- | --- | --- | --- | --- | --- | --- |
| C=12412; O=46; E=34.23; R=1.34; rawP=1.21e-06; adjP=1.74e-05 | | | | | | |
| Index | UserID | Value | Gene Symbol | Gene Name | EntrezGene | Ensembl |
| 1 | Q04446 | NA | GBE1 | glucan (1,4-alpha-), branching enzyme 1 | 2632 | ENSG00000114480 |
| 2 | P49591 | NA | SARS | seryl-tRNA synthetase | 6301 | ENSG00000031698 |
| 3 | Q92598 | NA | HSPH1 | heat shock 105kDa/110kDa protein 1 | 10808 | ENSG00000120694 |
| 4 | P61225 | NA | RAP2B | RAP2B, member of RAS oncogene family | 5912 | ENSG00000181467 |
| 5 | P55060 | NA | CSE1L | CSE1 chromosome segregation 1-like (yeast) | 1434 | ENSG00000124207 |
| 6 | Q9NQC3 | NA | RTN4 | reticulon 4 | 57142 | ENSG00000115310 |
| 7 | P16949 | NA | STMN1 | stathmin 1 | 3925 | ENSG00000117632 |
| 8 | P33993 | NA | MCM7 | minichromosome maintenance complex component 7 | 4176 | ENSG00000166508 |
| 9 | P68104 | NA | EEF1A1 | eukaryotic translation elongation factor 1 alpha 1 | 1915 | ENSG00000156508 |
| 10 | Q86UP2 | NA | KTN1 | kinectin 1 (kinesin receptor) | 3895 | ENSG00000126777 |
| 11 | P49588 | NA | AARS | alanyl-tRNA synthetase | 16 | ENSG00000090861 |
| 12 | P02786 | NA | TFRC | transferrin receptor (p90, CD71) | 7037 | ENSG00000072274 |
| 13 | O75369 | NA | FLNB | filamin B, beta | 2317 | ENSG00000136068 |
| 14 | P43490 | NA | NAMPT | nicotinamide phosphoribosyltransferase | 10135 | ENSG00000105835 |
| 15 | O00425 | NA | IGF2BP3 | insulin-like growth factor 2 mRNA binding protein 3 | 10643 | ENSG00000136231 |
| 16 | P17655 | NA | CAPN2 | calpain 2, (m/II) large subunit | 824 | ENSG00000162909 |
| 17 | P20042 | NA | EIF2S2 | eukaryotic translation initiation factor 2, subunit 2 beta, 38kDa | 8894 | ENSG00000125977 |
| 18 | Q9P258 | NA | RCC2 | regulator of chromosome condensation 2 | 55920 | ENSG00000179051 |
| 19 | P22102 | NA | GART | phosphoribosylglycinamide formyltransferase, phosphoribosylglycinamide synthetase, phosphoribosylaminoimidazole synthetase | 2618 | ENSG00000159131 |
| 20 | P16615 | NA | ATP2A2 | ATPase, Ca++ transporting, cardiac muscle, slow twitch 2 | 488 | ENSG00000174437 |
| 21 | Q92616 | NA | GCN1L1 | GCN1 general control of amino-acid synthesis 1-like 1 (yeast) | 10985 | ENSG00000089154 |
| 22 | P17174 | NA | GOT1 | glutamic-oxaloacetic transaminase 1, soluble (aspartate aminotransferase 1) | 2805 | ENSG00000120053 |
| 23 | P37802 | NA | TAGLN2 | transgelin 2 | 8407 | ENSG00000158710 |
| 24 | P08727 | NA | KRT19 | keratin 19 | 3880 | ENSG00000171345 |
| 25 | Q6PIU2 | NA | NCEH1 | neutral cholesterol ester hydrolase 1 | 57552 | ENSG00000144959 |
| 26 | P18206 | NA | VCL | vinculin | 7414 | ENSG00000035403 |
| 27 | P11387 | NA | TOP1 | topoisomerase (DNA) I | 7150 | ENSG00000198900 |
| 28 | Q9UK76 | NA | HN1 | hematological and neurological expressed 1 | 51155 | ENSG00000189159 |
| 29 | P21589 | NA | NT5E | 5'-nucleotidase, ecto (CD73) | 4907 | ENSG00000135318 |
| 30 | Q16881 | NA | TXNRD1 | thioredoxin reductase 1 | 7296 | ENSG00000198431 |
| 31 | Q9NR30 | NA | DDX21 | DEAD (Asp-Glu-Ala-Asp) box helicase 21 | 9188 | ENSG00000165732 |
| 32 | P08195 | NA | SLC3A2 | solute carrier family 3 (activators of dibasic and neutral amino acid transport), member 2 | 6520 | ENSG00000168003 |
| 33 | P00491 | NA | PNP | purine nucleoside phosphorylase | 4860 | ENSG00000198805 |
| 34 | P49321 | NA | NASP | nuclear autoantigenic sperm protein (histone-binding) | 4678 | ENSG00000132780 |
| 35 | Q9HAV4 | NA | XPO5 | exportin 5 | 57510 | ENSG00000124571 |
| 36 | P56192 | NA | MARS | methionyl-tRNA synthetase | 4141 | ENSG00000166986 |
| 37 | P78417 | NA | GSTO1 | glutathione S-transferase omega 1 | 9446 | ENSG00000148834 |
| 38 | P49736 | NA | MCM2 | minichromosome maintenance complex component 2 | 4171 | ENSG00000073111 |
| 39 | Q15056 | NA | EIF4H | eukaryotic translation initiation factor 4H | 7458 | ENSG00000106682 |
| 40 | O60610 | NA | DIAPH1 | diaphanous homolog 1 (Drosophila) | 1729 | ENSG00000131504 |
| 41 | Q06830 | NA | PRDX1 | peroxiredoxin 1 | 5052 | ENSG00000117450 |
| 42 | P04083 | NA | ANXA1 | annexin A1 | 301 | ENSG00000135046 |
| 43 | P41091 | NA | EIF2S3 | eukaryotic translation initiation factor 2, subunit 3 gamma, 52kDa | 1968 | ENSG00000130741 |
| 44 | P02768 | NA | ALB | albumin | 213 | ENSG00000163631 |
| 45 | P05556 | NA | ITGB1 | integrin, beta 1 (fibronectin receptor, beta polypeptide, antigen CD29 includes MDF2, MSK12) | 3688 | ENSG00000150093 |
| 46 | Q15758 | NA | SLC1A5 | solute carrier family 1 (neutral amino acid transporter), member 5 | 6510 | ENSG00000105281 |

  
  

| **Database:cellular component      &nbspName:melanosome      &nbspID:GO:0042470** | | | | | | |
| --- | --- | --- | --- | --- | --- | --- |
| C=93; O=5; E=0.26; R=19.50; rawP=5.53e-06; adjP=5.69e-05 | | | | | | |
| Index | UserID | Value | Gene Symbol | Gene Name | EntrezGene | Ensembl |
| 1 | P08195 | NA | SLC3A2 | solute carrier family 3 (activators of dibasic and neutral amino acid transport), member 2 | 6520 | ENSG00000168003 |
| 2 | Q15758 | NA | SLC1A5 | solute carrier family 1 (neutral amino acid transporter), member 5 | 6510 | ENSG00000105281 |
| 3 | Q06830 | NA | PRDX1 | peroxiredoxin 1 | 5052 | ENSG00000117450 |
| 4 | P02786 | NA | TFRC | transferrin receptor (p90, CD71) | 7037 | ENSG00000072274 |
| 5 | P05556 | NA | ITGB1 | integrin, beta 1 (fibronectin receptor, beta polypeptide, antigen CD29 includes MDF2, MSK12) | 3688 | ENSG00000150093 |

  
  

| **Database:cellular component      &nbspName:pigment granule      &nbspID:GO:0048770** | | | | | | |
| --- | --- | --- | --- | --- | --- | --- |
| C=93; O=5; E=0.26; R=19.50; rawP=5.53e-06; adjP=5.69e-05 | | | | | | |
| Index | UserID | Value | Gene Symbol | Gene Name | EntrezGene | Ensembl |
| 1 | P08195 | NA | SLC3A2 | solute carrier family 3 (activators of dibasic and neutral amino acid transport), member 2 | 6520 | ENSG00000168003 |
| 2 | Q15758 | NA | SLC1A5 | solute carrier family 1 (neutral amino acid transporter), member 5 | 6510 | ENSG00000105281 |
| 3 | Q06830 | NA | PRDX1 | peroxiredoxin 1 | 5052 | ENSG00000117450 |
| 4 | P02786 | NA | TFRC | transferrin receptor (p90, CD71) | 7037 | ENSG00000072274 |
| 5 | P05556 | NA | ITGB1 | integrin, beta 1 (fibronectin receptor, beta polypeptide, antigen CD29 includes MDF2, MSK12) | 3688 | ENSG00000150093 |

  
  
  
  

---

WebGestalt is currently developed and maintained by Jing Wang and Bing Zhang at the  Zhang Lab. Other people who have made significant contribution to the project include Dexter Duncan, Stefan Kirov, Zhiao Shi, and Jay Snoddy.  
  
**Funding credits:** NIH/NIAAA (U01 AA016662, U01 AA013512); NIH/NIDA (P01 DA015027); NIH/NIMH (P50 MH078028, P50 MH096972); NIH/NCI (U24 CA159988); NIH/NIGMS (R01 GM088822).
