## Supplementary material for "PGRMC1 phosphorylation and cell plasticity 1: glycolysis, mitochondria, tumor growth": File S7: final_pc_geneset_file_1438748351.html

  
  

| **Database:Pathway Commons pathway      &nbspName:CDC42 signaling events      &nbspID:DB\_ID:1488** | | | | | | |
| --- | --- | --- | --- | --- | --- | --- |
| C=757; O=9; E=1.72; R=5.24; rawP=4.27e-05; adjP=0.0002 | | | | | | |
| Index | UserID | Value | Gene Symbol | Gene Name | EntrezGene | Ensembl |
| 1 | P08195 | NA | SLC3A2 | solute carrier family 3 (activators of dibasic and neutral amino acid transport), member 2 | 6520 | ENSG00000168003 |
| 2 | P05556 | NA | ITGB1 | integrin, beta 1 (fibronectin receptor, beta polypeptide, antigen CD29 includes MDF2, MSK12) | 3688 | ENSG00000150093 |
| 3 | P08727 | NA | KRT19 | keratin 19 | 3880 | ENSG00000171345 |
| 4 | P16949 | NA | STMN1 | stathmin 1 | 3925 | ENSG00000117632 |
| 5 | P18206 | NA | VCL | vinculin | 7414 | ENSG00000035403 |
| 6 | O60610 | NA | DIAPH1 | diaphanous homolog 1 (Drosophila) | 1729 | ENSG00000131504 |
| 7 | P02786 | NA | TFRC | transferrin receptor (p90, CD71) | 7037 | ENSG00000072274 |
| 8 | Q06830 | NA | PRDX1 | peroxiredoxin 1 | 5052 | ENSG00000117450 |
| 9 | P21589 | NA | NT5E | 5'-nucleotidase, ecto (CD73) | 4907 | ENSG00000135318 |

  
  

| **Database:Pathway Commons pathway      &nbspName:Regulation of CDC42 activity      &nbspID:DB\_ID:1456** | | | | | | |
| --- | --- | --- | --- | --- | --- | --- |
| C=770; O=9; E=1.75; R=5.15; rawP=4.87e-05; adjP=0.0002 | | | | | | |
| Index | UserID | Value | Gene Symbol | Gene Name | EntrezGene | Ensembl |
| 1 | P08195 | NA | SLC3A2 | solute carrier family 3 (activators of dibasic and neutral amino acid transport), member 2 | 6520 | ENSG00000168003 |
| 2 | P05556 | NA | ITGB1 | integrin, beta 1 (fibronectin receptor, beta polypeptide, antigen CD29 includes MDF2, MSK12) | 3688 | ENSG00000150093 |
| 3 | P08727 | NA | KRT19 | keratin 19 | 3880 | ENSG00000171345 |
| 4 | P16949 | NA | STMN1 | stathmin 1 | 3925 | ENSG00000117632 |
| 5 | P18206 | NA | VCL | vinculin | 7414 | ENSG00000035403 |
| 6 | O60610 | NA | DIAPH1 | diaphanous homolog 1 (Drosophila) | 1729 | ENSG00000131504 |
| 7 | P02786 | NA | TFRC | transferrin receptor (p90, CD71) | 7037 | ENSG00000072274 |
| 8 | Q06830 | NA | PRDX1 | peroxiredoxin 1 | 5052 | ENSG00000117450 |
| 9 | P21589 | NA | NT5E | 5'-nucleotidase, ecto (CD73) | 4907 | ENSG00000135318 |

  
  

| **Database:Pathway Commons pathway      &nbspName:Purine metabolism      &nbspID:DB\_ID:638** | | | | | | |
| --- | --- | --- | --- | --- | --- | --- |
| C=24; O=3; E=0.05; R=55.12; rawP=2.14e-05; adjP=0.0002 | | | | | | |
| Index | UserID | Value | Gene Symbol | Gene Name | EntrezGene | Ensembl |
| 1 | P22102 | NA | GART | phosphoribosylglycinamide formyltransferase, phosphoribosylglycinamide synthetase, phosphoribosylaminoimidazole synthetase | 2618 | ENSG00000159131 |
| 2 | P00491 | NA | PNP | purine nucleoside phosphorylase | 4860 | ENSG00000198805 |
| 3 | P21589 | NA | NT5E | 5'-nucleotidase, ecto (CD73) | 4907 | ENSG00000135318 |

  
  

| **Database:Pathway Commons pathway      &nbspName:Activation of the mRNA upon binding of the cap-binding complex and eIFs, and subsequent binding to 43S      &nbspID:DB\_ID:720** | | | | | | |
| --- | --- | --- | --- | --- | --- | --- |
| C=56; O=3; E=0.13; R=23.62; rawP=0.0003; adjP=0.0005 | | | | | | |
| Index | UserID | Value | Gene Symbol | Gene Name | EntrezGene | Ensembl |
| 1 | P20042 | NA | EIF2S2 | eukaryotic translation initiation factor 2, subunit 2 beta, 38kDa | 8894 | ENSG00000125977 |
| 2 | P41091 | NA | EIF2S3 | eukaryotic translation initiation factor 2, subunit 3 gamma, 52kDa | 1968 | ENSG00000130741 |
| 3 | Q15056 | NA | EIF4H | eukaryotic translation initiation factor 4H | 7458 | ENSG00000106682 |

  
  

| **Database:Pathway Commons pathway      &nbspName:Ribosomal scanning and start codon recognition      &nbspID:DB\_ID:717** | | | | | | |
| --- | --- | --- | --- | --- | --- | --- |
| C=55; O=3; E=0.12; R=24.05; rawP=0.0003; adjP=0.0005 | | | | | | |
| Index | UserID | Value | Gene Symbol | Gene Name | EntrezGene | Ensembl |
| 1 | P20042 | NA | EIF2S2 | eukaryotic translation initiation factor 2, subunit 2 beta, 38kDa | 8894 | ENSG00000125977 |
| 2 | P41091 | NA | EIF2S3 | eukaryotic translation initiation factor 2, subunit 3 gamma, 52kDa | 1968 | ENSG00000130741 |
| 3 | Q15056 | NA | EIF4H | eukaryotic translation initiation factor 4H | 7458 | ENSG00000106682 |

  
  

| **Database:Pathway Commons pathway      &nbspName:Translation initiation complex formation      &nbspID:DB\_ID:719** | | | | | | |
| --- | --- | --- | --- | --- | --- | --- |
| C=55; O=3; E=0.12; R=24.05; rawP=0.0003; adjP=0.0005 | | | | | | |
| Index | UserID | Value | Gene Symbol | Gene Name | EntrezGene | Ensembl |
| 1 | P20042 | NA | EIF2S2 | eukaryotic translation initiation factor 2, subunit 2 beta, 38kDa | 8894 | ENSG00000125977 |
| 2 | P41091 | NA | EIF2S3 | eukaryotic translation initiation factor 2, subunit 3 gamma, 52kDa | 1968 | ENSG00000130741 |
| 3 | Q15056 | NA | EIF4H | eukaryotic translation initiation factor 4H | 7458 | ENSG00000106682 |

  
  
  
  

---

WebGestalt is currently developed and maintained by Jing Wang and Bing Zhang at the  Zhang Lab. Other people who have made significant contribution to the project include Dexter Duncan, Stefan Kirov, Zhiao Shi, and Jay Snoddy.  
  
**Funding credits:** NIH/NIAAA (U01 AA016662, U01 AA013512); NIH/NIDA (P01 DA015027); NIH/NIMH (P50 MH078028, P50 MH096972); NIH/NCI (U24 CA159988); NIH/NIGMS (R01 GM088822).
