## Supplementary material for "PGRMC1 phosphorylation and cell plasticity 1: glycolysis, mitochondria, tumor growth": File S7: final_tf_geneset_file_1438748351.html

Anchored HTML File of EIDs

|  |  |
| --- | --- |
|  | WEB-based GEne SeT AnaLysis Toolkit |
|  | ***Translating gene lists into biological insights...*** |

---

  

| **Database:Transcrription Target      &nbspName:hsa\_GGGCGGR\_V$SP1\_Q6      &nbspID:DB\_ID:2452** | | | | | | |
| --- | --- | --- | --- | --- | --- | --- |
| C=2863; O=20; E=6.49; R=3.08; rawP=1.28e-06; adjP=0.0001 | | | | | | |
| Index | UserID | Value | Gene Symbol | Gene Name | EntrezGene | Ensembl |
| 1 | Q16881 | NA | TXNRD1 | thioredoxin reductase 1 | 7296 | ENSG00000198431 |
| 2 | Q92598 | NA | HSPH1 | heat shock 105kDa/110kDa protein 1 | 10808 | ENSG00000120694 |
| 3 | P49321 | NA | NASP | nuclear autoantigenic sperm protein (histone-binding) | 4678 | ENSG00000132780 |
| 4 | Q9HAV4 | NA | XPO5 | exportin 5 | 57510 | ENSG00000124571 |
| 5 | P55060 | NA | CSE1L | CSE1 chromosome segregation 1-like (yeast) | 1434 | ENSG00000124207 |
| 6 | Q9NQC3 | NA | RTN4 | reticulon 4 | 57142 | ENSG00000115310 |
| 7 | P16949 | NA | STMN1 | stathmin 1 | 3925 | ENSG00000117632 |
| 8 | P33993 | NA | MCM7 | minichromosome maintenance complex component 7 | 4176 | ENSG00000166508 |
| 9 | O60610 | NA | DIAPH1 | diaphanous homolog 1 (Drosophila) | 1729 | ENSG00000131504 |
| 10 | Q86UP2 | NA | KTN1 | kinectin 1 (kinesin receptor) | 3895 | ENSG00000126777 |
| 11 | P68104 | NA | EEF1A1 | eukaryotic translation elongation factor 1 alpha 1 | 1915 | ENSG00000156508 |
| 12 | O00425 | NA | IGF2BP3 | insulin-like growth factor 2 mRNA binding protein 3 | 10643 | ENSG00000136231 |
| 13 | P20042 | NA | EIF2S2 | eukaryotic translation initiation factor 2, subunit 2 beta, 38kDa | 8894 | ENSG00000125977 |
| 14 | P41091 | NA | EIF2S3 | eukaryotic translation initiation factor 2, subunit 3 gamma, 52kDa | 1968 | ENSG00000130741 |
| 15 | P05556 | NA | ITGB1 | integrin, beta 1 (fibronectin receptor, beta polypeptide, antigen CD29 includes MDF2, MSK12) | 3688 | ENSG00000150093 |
| 16 | P08727 | NA | KRT19 | keratin 19 | 3880 | ENSG00000171345 |
| 17 | P37802 | NA | TAGLN2 | transgelin 2 | 8407 | ENSG00000158710 |
| 18 | P18206 | NA | VCL | vinculin | 7414 | ENSG00000035403 |
| 19 | Q9UK76 | NA | HN1 | hematological and neurological expressed 1 | 51155 | ENSG00000189159 |
| 20 | P11387 | NA | TOP1 | topoisomerase (DNA) I | 7150 | ENSG00000198900 |

  
  
  
  

---

WebGestalt is currently developed and maintained by Jing Wang and Bing Zhang at the  Zhang Lab. Other people who have made significant contribution to the project include Dexter Duncan, Stefan Kirov, Zhiao Shi, and Jay Snoddy.  
  
**Funding credits:** NIH/NIAAA (U01 AA016662, U01 AA013512); NIH/NIDA (P01 DA015027); NIH/NIMH (P50 MH078028, P50 MH096972); NIH/NCI (U24 CA159988); NIH/NIGMS (R01 GM088822).
