## Supplementary figures and images for "PGRMC1 phosphorylation and cell plasticity 1: glycolysis, mitochondria, tumor growth"

### final_DAG_file_1438480126.gif

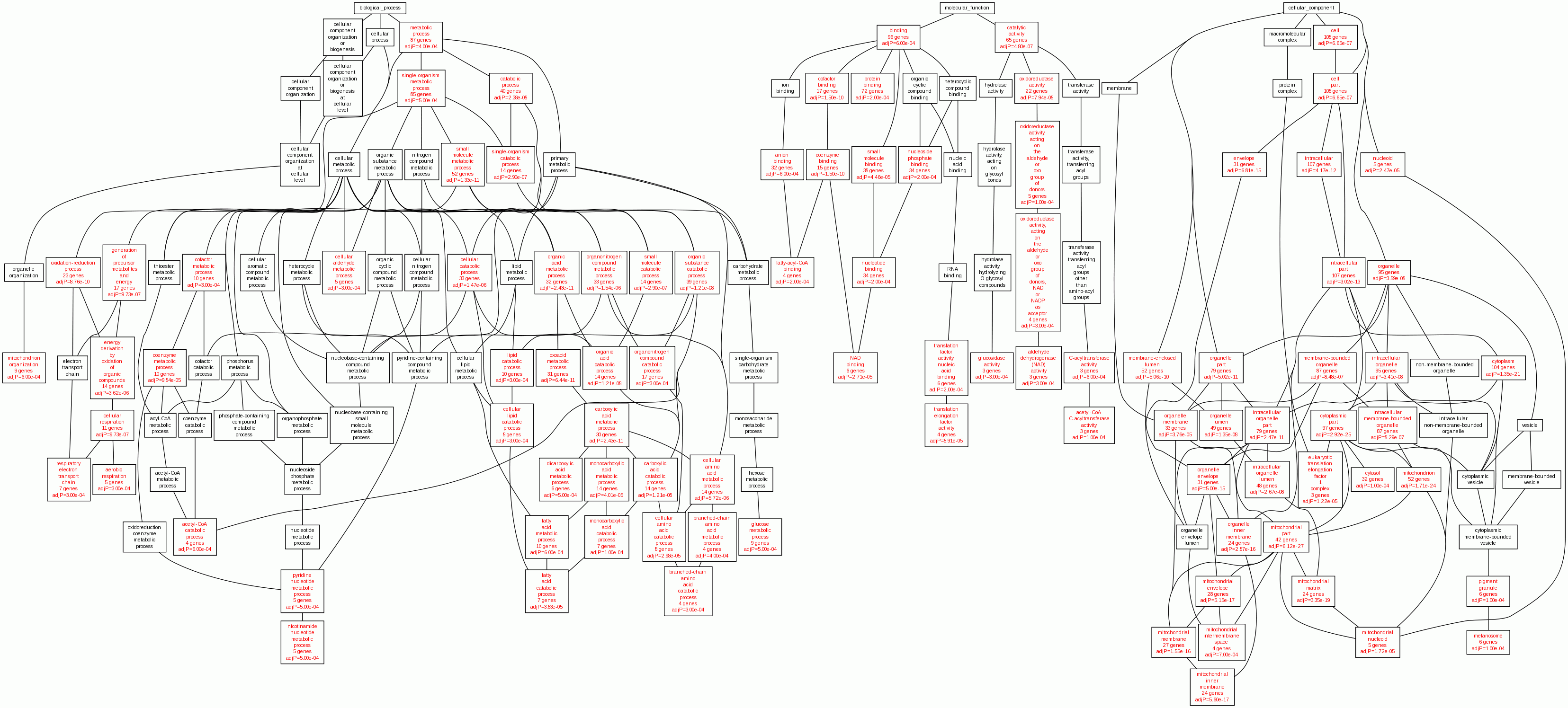

### final_DAG_file_1438486678.gif

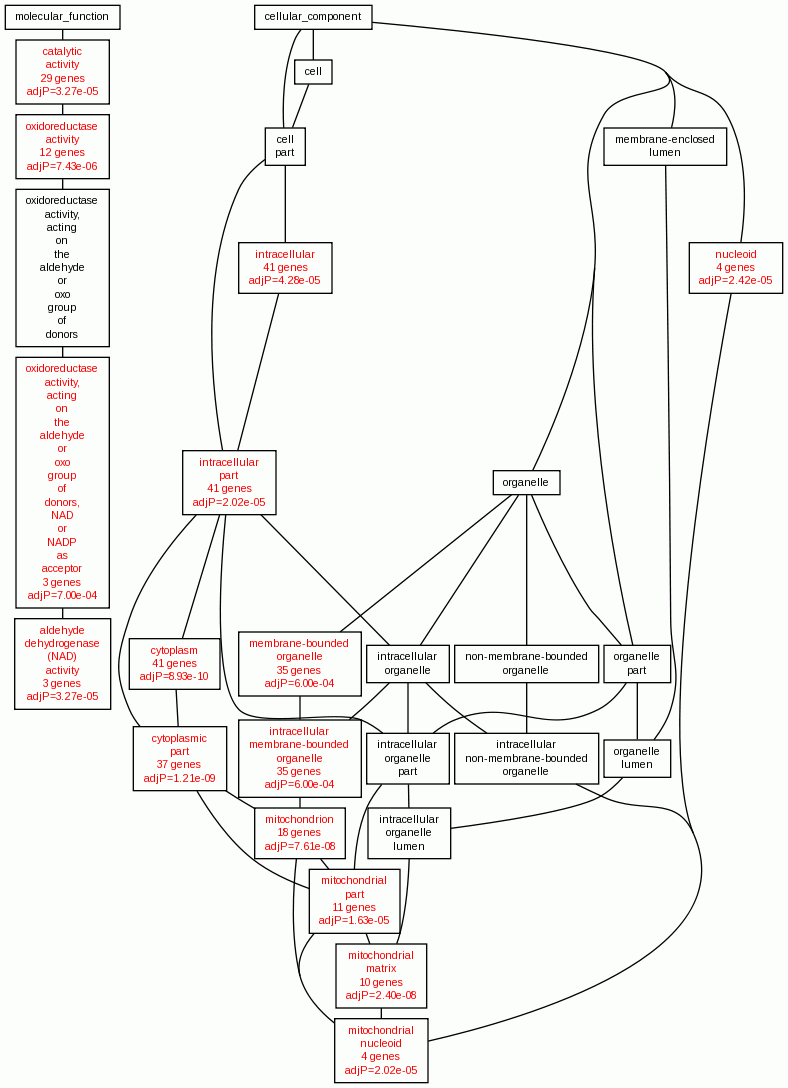

### final_DAG_file_1438487532.gif

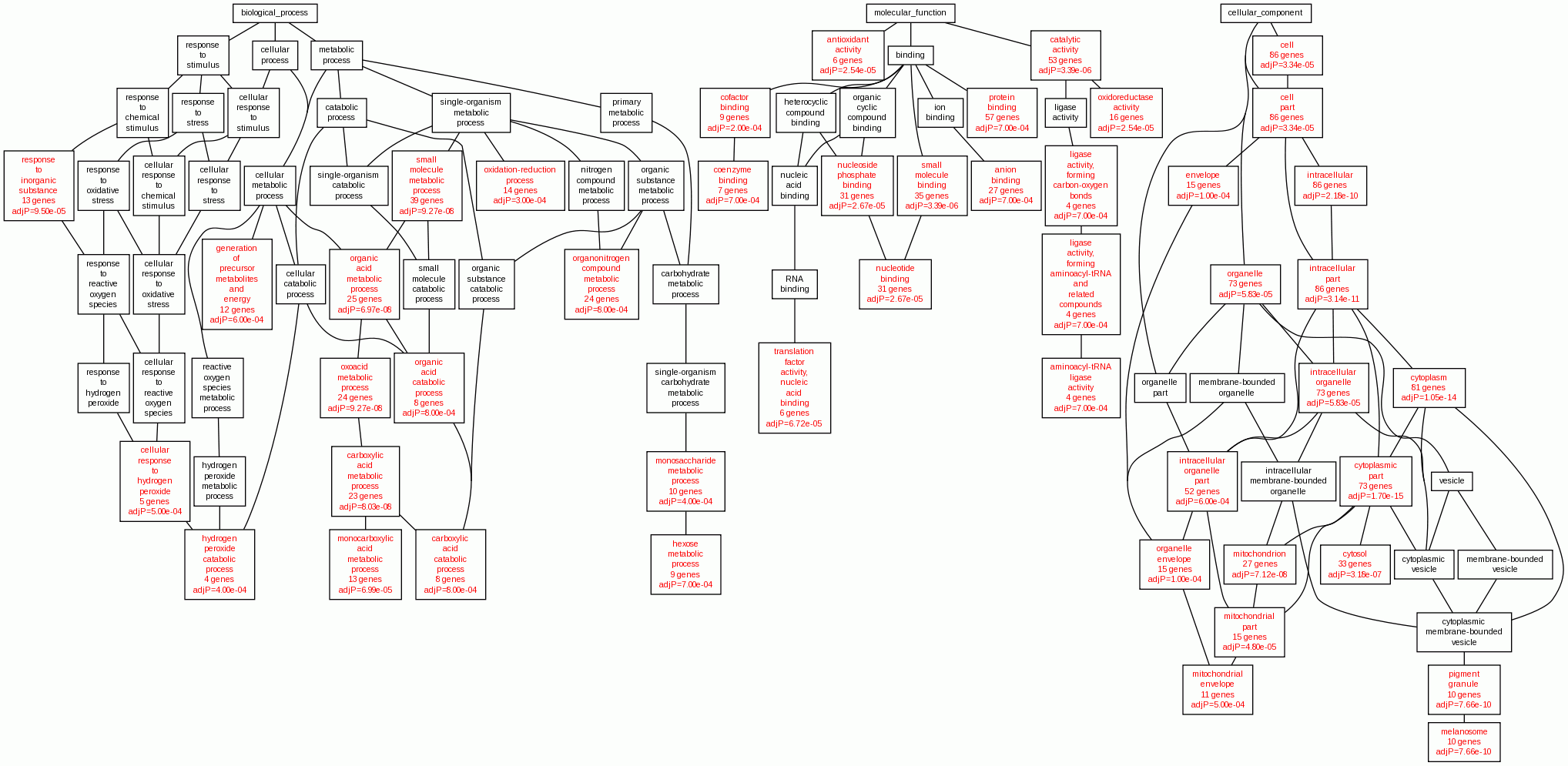

### final_DAG_file_1438746600.gif

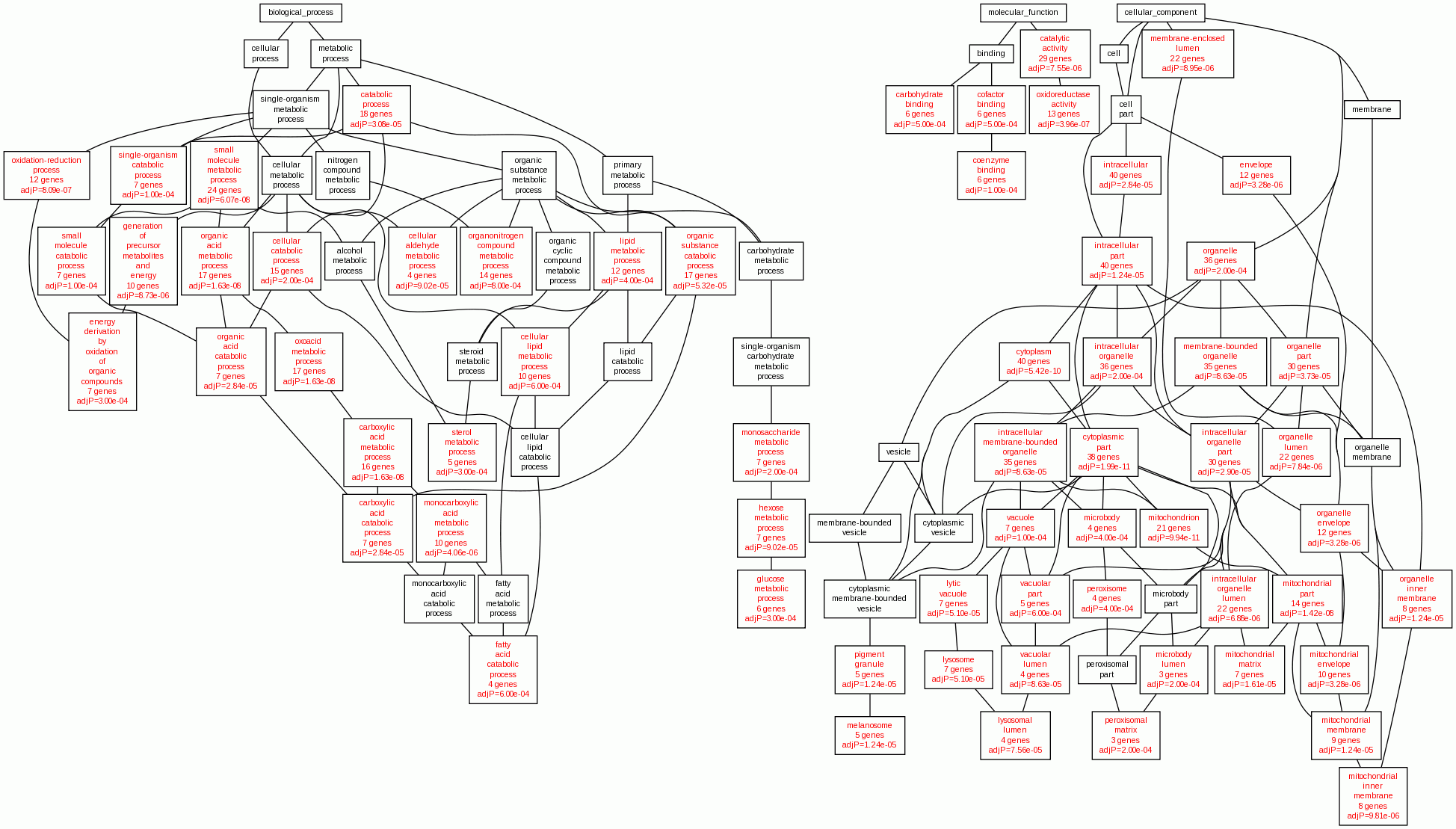

### final_DAG_file_1438748351.gif

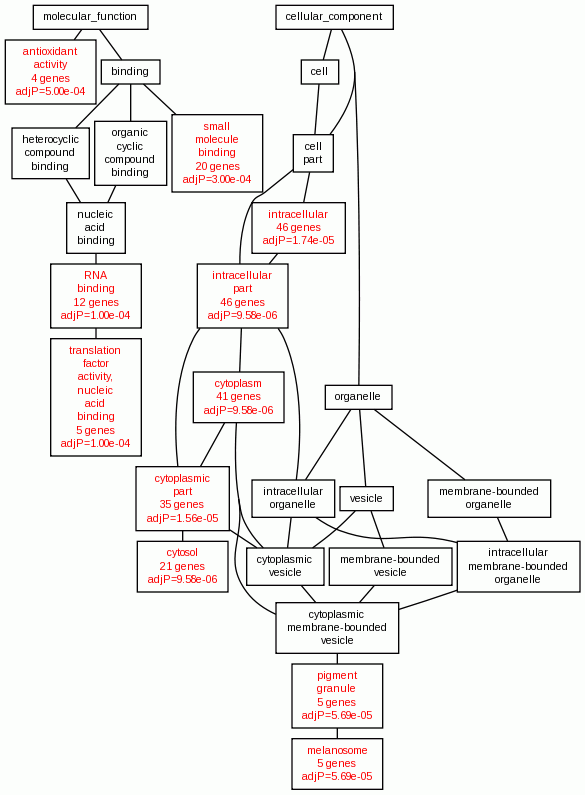

### final_DAG_file_1438753413.gif

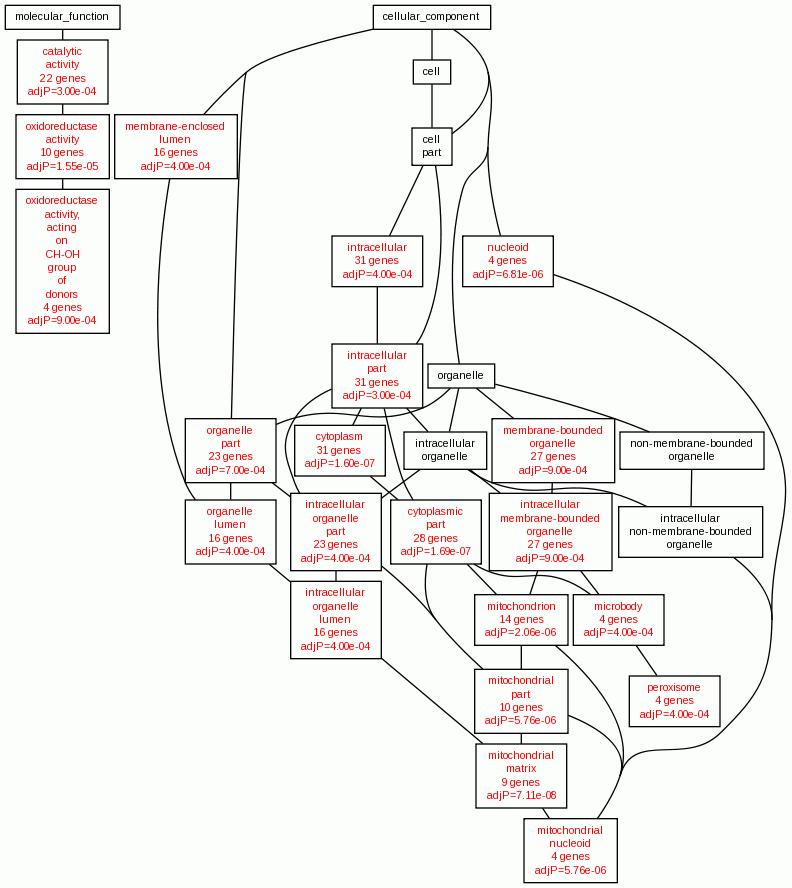

### final_DAG_file_1439165549.gif

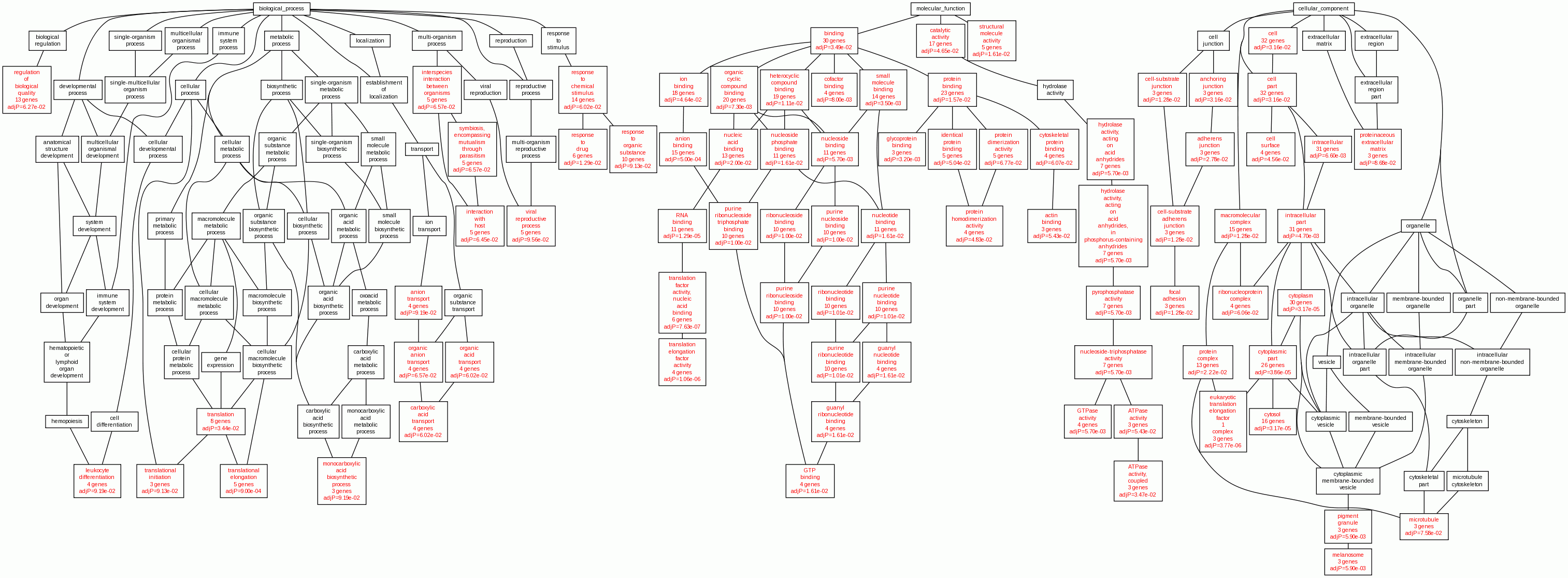

### final_DAG_file_1439165895.gif

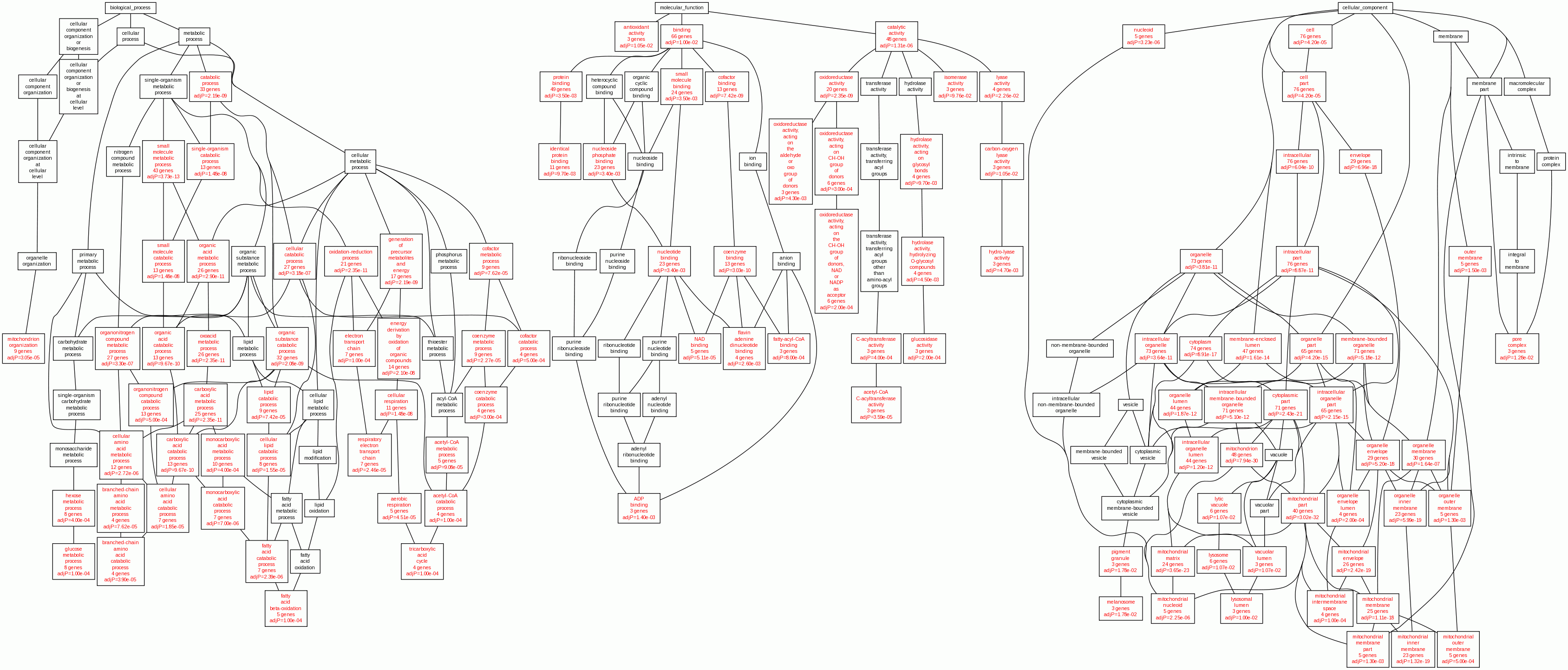

### final_DAG_file_1439166314.gif

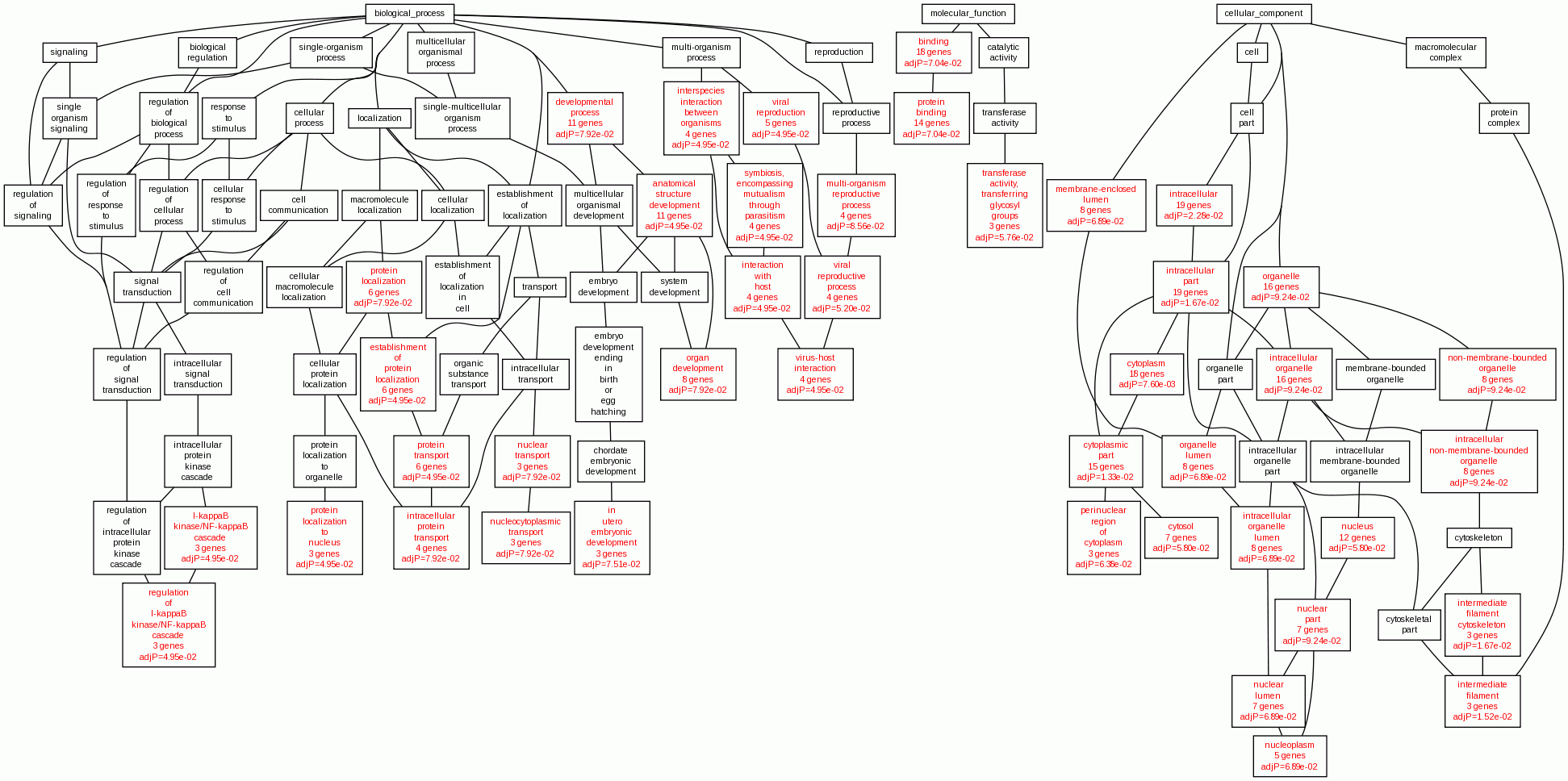

### final_DAG_file_1439166598.gif

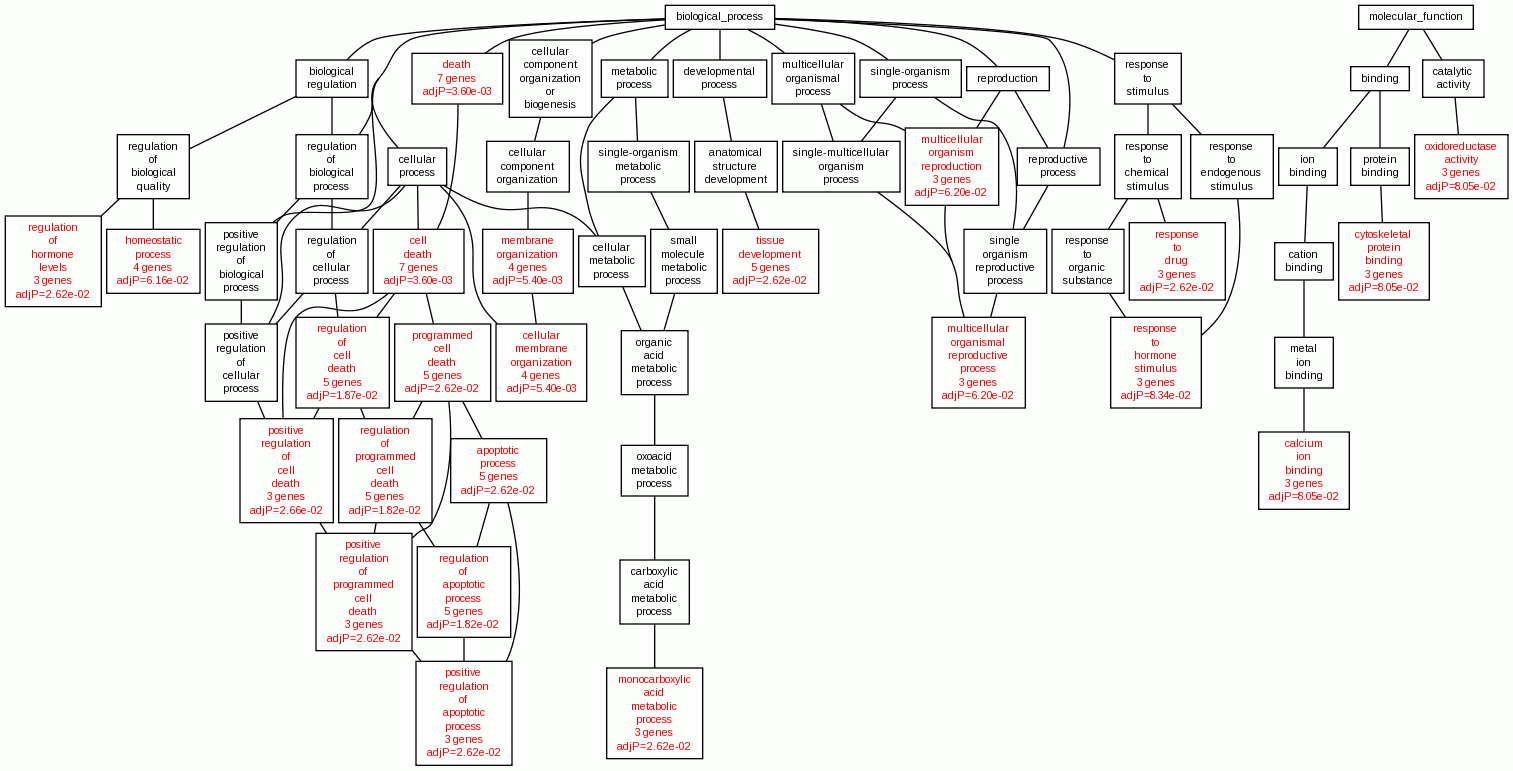

### final_DAG_file_1439166893.gif

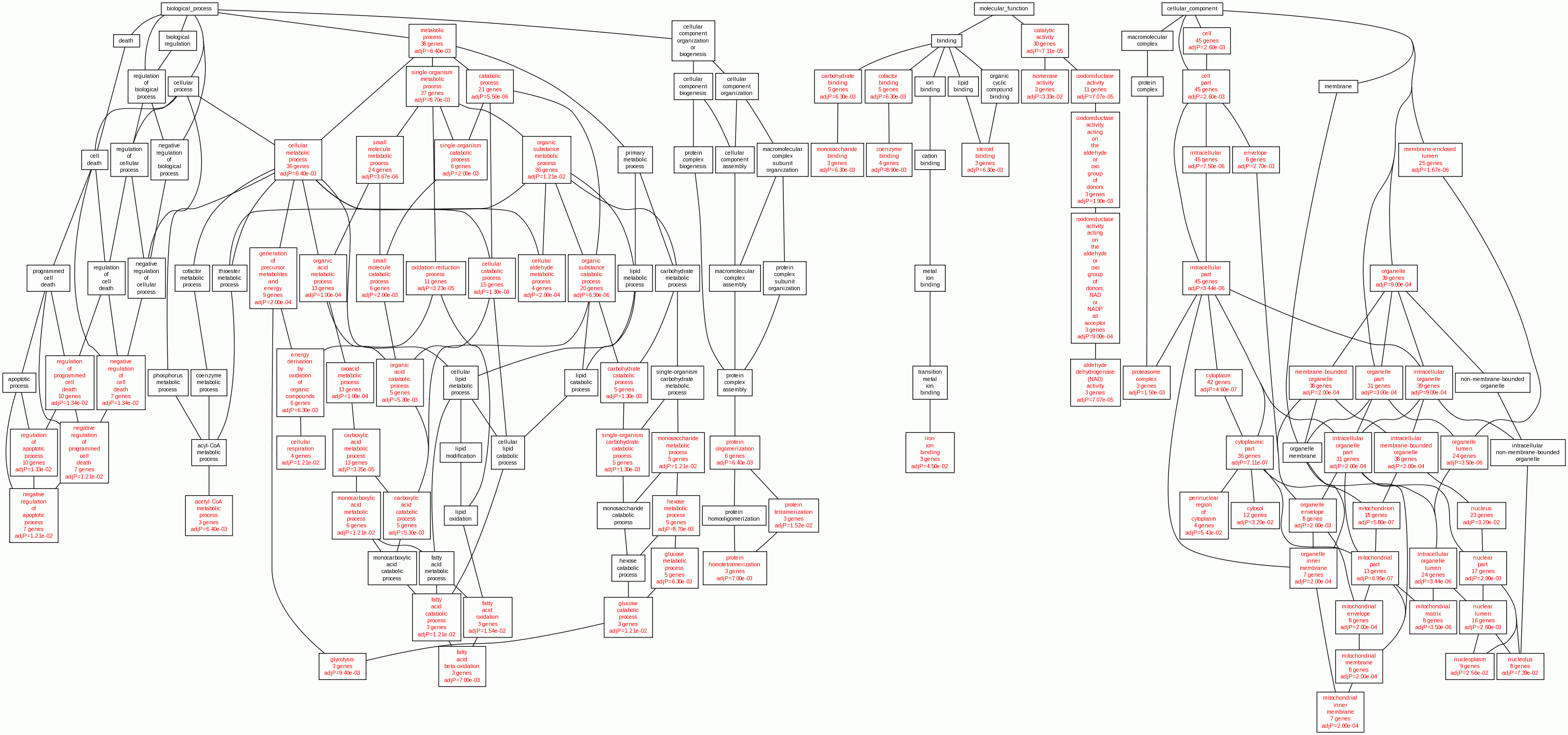

### final_DAG_file_1439167077.gif

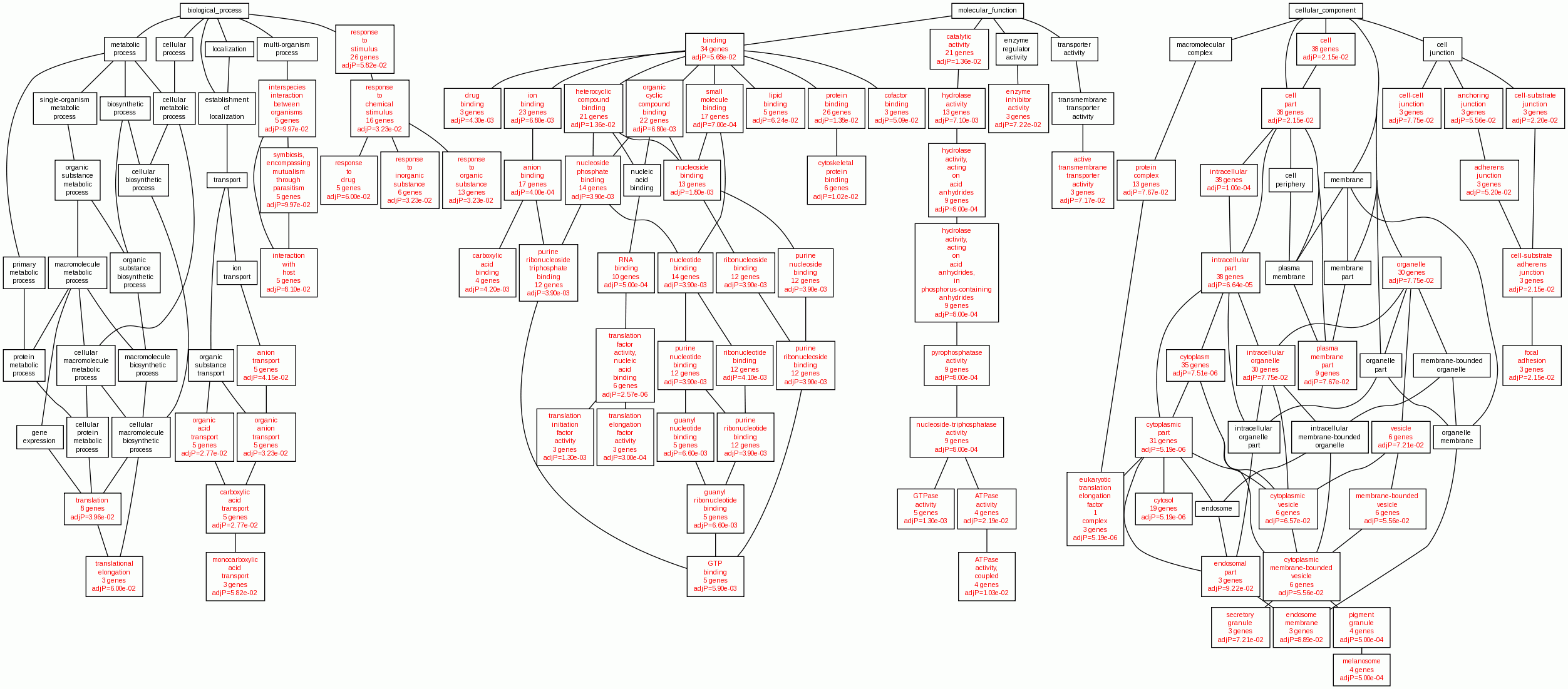

### final_DAG_file_1439167306.gif

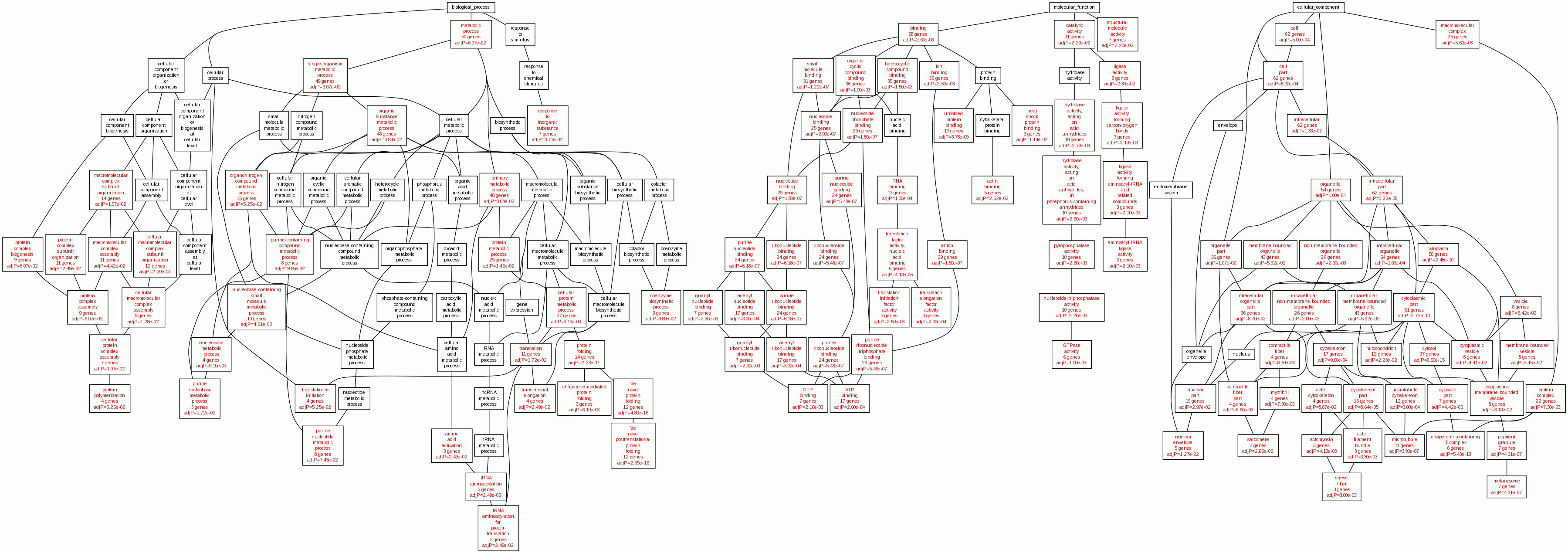

### final_ppi_DAG_file_1438487532.gif

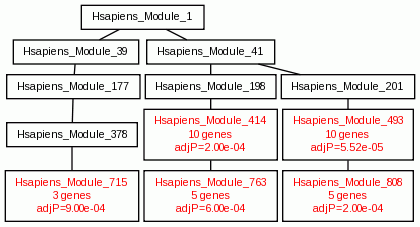

### final_ppi_DAG_file_1438746600.gif

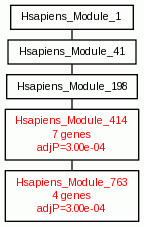
