## Supplementary material for "PGRMC1 phosphorylation and cell plasticity 1: glycolysis, mitochondria, tumor growth": File S8: final_sig_file_1439165895.html

Anchored HTML File of EIDs

|  |  |
| --- | --- |
|  | WEB-based GEne SeT AnaLysis Toolkit |
|  | ***Translating gene lists into biological insights...*** |

---

  

| **Database:biological process      &nbspName:small molecule metabolic process      &nbspID:GO:0044281** | | | | | | |
| --- | --- | --- | --- | --- | --- | --- |
| C=2500; O=43; E=12.48; R=3.45; rawP=7.99e-16; adjP=3.73e-13 | | | | | | |
| Index | UserID | Value | Gene Symbol | Gene Name | EntrezGene | Ensembl |
| 1 | P54819 | NA | AK2 | adenylate kinase 2 | 204 | ENSG00000004455 |
| 2 | Q9UL46 | NA | PSME2 | proteasome (prosome, macropain) activator subunit 2 (PA28 beta) | 5721 | ENSG00000100911 |
| 3 | P40926 | NA | MDH2 | malate dehydrogenase 2, NAD (mitochondrial) | 4191 | ENSG00000146701 |
| 4 | Q99714 | NA | HSD17B10 | hydroxysteroid (17-beta) dehydrogenase 10 | 3028 | ENSG00000072506 |
| 5 | P04179 | NA | SOD2 | superoxide dismutase 2, mitochondrial | 6648 | ENSG00000112096 |
| 6 | P07741 | NA | APRT | adenine phosphoribosyltransferase | 353 | ENSG00000198931 |
| 7 | P00367 | NA | GLUD1 | glutamate dehydrogenase 1 | 2746 | ENSG00000148672 |
| 8 | Q06323 | NA | PSME1 | proteasome (prosome, macropain) activator subunit 1 (PA28 alpha) | 5720 | ENSG00000092010 |
| 9 | P48735 | NA | IDH2 | isocitrate dehydrogenase 2 (NADP+), mitochondrial | 3418 | ENSG00000182054 |
| 10 | O60313 | NA | OPA1 | optic atrophy 1 (autosomal dominant) | 4976 | ENSG00000198836 |
| 11 | P35270 | NA | SPR | sepiapterin reductase (7,8-dihydrobiopterin:NADP+ oxidoreductase) | 6697 | ENSG00000116096 |
| 12 | P31040 | NA | SDHA | succinate dehydrogenase complex, subunit A, flavoprotein (Fp) | 6389 | ENSG00000073578 |
| 13 | P38117 | NA | ETFB | electron-transfer-flavoprotein, beta polypeptide | 2109 | ENSG00000105379 |
| 14 | P43490 | NA | NAMPT | nicotinamide phosphoribosyltransferase | 10135 | ENSG00000105835 |
| 15 | P13804 | NA | ETFA | electron-transfer-flavoprotein, alpha polypeptide | 2108 | ENSG00000140374 |
| 16 | P42765 | NA | ACAA2 | acetyl-CoA acyltransferase 2 | 10449 | ENSG00000167315 |
| 17 | O75521 | NA | ECI2 | enoyl-CoA delta isomerase 2 | 10455 | ENSG00000198721 |
| 18 | P49419 | NA | ALDH7A1 | aldehyde dehydrogenase 7 family, member A1 | 501 | ENSG00000164904 |
| 19 | P21953 | NA | BCKDHB | branched chain keto acid dehydrogenase E1, beta polypeptide | 594 | ENSG00000083123 |
| 20 | P40939 | NA | HADHA | hydroxyacyl-CoA dehydrogenase/3-ketoacyl-CoA thiolase/enoyl-CoA hydratase (trifunctional protein), alpha subunit | 3030 | ENSG00000084754 |
| 21 | P30084 | NA | ECHS1 | enoyl CoA hydratase, short chain, 1, mitochondrial | 1892 | ENSG00000127884 |
| 22 | P38571 | NA | LIPA | lipase A, lysosomal acid, cholesterol esterase | 3988 | ENSG00000107798 |
| 23 | Q13011 | NA | ECH1 | enoyl CoA hydratase 1, peroxisomal | 1891 | ENSG00000104823 |
| 24 | P32322 | NA | PYCR1 | pyrroline-5-carboxylate reductase 1 | 5831 | ENSG00000183010 |
| 25 | P04040 | NA | CAT | catalase | 847 | ENSG00000121691 |
| 26 | Q99798 | NA | ACO2 | aconitase 2, mitochondrial | 50 | ENSG00000100412 |
| 27 | P51688 | NA | SGSH | N-sulfoglucosamine sulfohydrolase | 6448 | ENSG00000181523 |
| 28 | P51649 | NA | ALDH5A1 | aldehyde dehydrogenase 5 family, member A1 | 7915 | ENSG00000112294 |
| 29 | P12235 | NA | SLC25A4 | solute carrier family 25 (mitochondrial carrier; adenine nucleotide translocator), member 4 | 291 | ENSG00000151729 |
| 30 | P00558 | NA | PGK1 | phosphoglycerate kinase 1 | 5230 | ENSG00000102144 |
| 31 | Q9NSE4 | NA | IARS2 | isoleucyl-tRNA synthetase 2, mitochondrial | 55699 | ENSG00000067704 |
| 32 | Q6PI48 | NA | DARS2 | aspartyl-tRNA synthetase 2, mitochondrial | 55157 | ENSG00000117593 |
| 33 | P24752 | NA | ACAT1 | acetyl-CoA acetyltransferase 1 | 38 | ENSG00000075239 |
| 34 | P13674 | NA | P4HA1 | prolyl 4-hydroxylase, alpha polypeptide I | 5033 | ENSG00000122884 |
| 35 | Q02978 | NA | SLC25A11 | solute carrier family 25 (mitochondrial carrier; oxoglutarate carrier), member 11 | 8402 | ENSG00000108528 |
| 36 | P19367 | NA | HK1 | hexokinase 1 | 3098 | ENSG00000156515 |
| 37 | P25705 | NA | ATP5A1 | ATP synthase, H+ transporting, mitochondrial F1 complex, alpha subunit 1, cardiac muscle | 498 | ENSG00000152234 |
| 38 | Q9BQ69 | NA | MACROD1 | MACRO domain containing 1 | 28992 | ENSG00000133315 |
| 39 | P06865 | NA | HEXA | hexosaminidase A (alpha polypeptide) | 3073 | ENSG00000213614 |
| 40 | P49748 | NA | ACADVL | acyl-CoA dehydrogenase, very long chain | 37 | ENSG00000072778 |
| 41 | P48047 | NA | ATP5O | ATP synthase, H+ transporting, mitochondrial F1 complex, O subunit | 539 | ENSG00000241837 |
| 42 | P60174 | NA | TPI1 | triosephosphate isomerase 1 | 7167 | ENSG00000111669 |
| 43 | P31937 | NA | HIBADH | 3-hydroxyisobutyrate dehydrogenase | 11112 | ENSG00000106049 |

  
  

| **Database:biological process      &nbspName:oxoacid metabolic process      &nbspID:GO:0043436** | | | | | | |
| --- | --- | --- | --- | --- | --- | --- |
| C=949; O=26; E=4.74; R=5.49; rawP=2.01e-13; adjP=2.35e-11 | | | | | | |
| Index | UserID | Value | Gene Symbol | Gene Name | EntrezGene | Ensembl |
| 1 | P21953 | NA | BCKDHB | branched chain keto acid dehydrogenase E1, beta polypeptide | 594 | ENSG00000083123 |
| 2 | P40939 | NA | HADHA | hydroxyacyl-CoA dehydrogenase/3-ketoacyl-CoA thiolase/enoyl-CoA hydratase (trifunctional protein), alpha subunit | 3030 | ENSG00000084754 |
| 3 | Q9UL46 | NA | PSME2 | proteasome (prosome, macropain) activator subunit 2 (PA28 beta) | 5721 | ENSG00000100911 |
| 4 | P40926 | NA | MDH2 | malate dehydrogenase 2, NAD (mitochondrial) | 4191 | ENSG00000146701 |
| 5 | P30084 | NA | ECHS1 | enoyl CoA hydratase, short chain, 1, mitochondrial | 1892 | ENSG00000127884 |
| 6 | Q99714 | NA | HSD17B10 | hydroxysteroid (17-beta) dehydrogenase 10 | 3028 | ENSG00000072506 |
| 7 | P04179 | NA | SOD2 | superoxide dismutase 2, mitochondrial | 6648 | ENSG00000112096 |
| 8 | P00367 | NA | GLUD1 | glutamate dehydrogenase 1 | 2746 | ENSG00000148672 |
| 9 | Q06323 | NA | PSME1 | proteasome (prosome, macropain) activator subunit 1 (PA28 alpha) | 5720 | ENSG00000092010 |
| 10 | P48735 | NA | IDH2 | isocitrate dehydrogenase 2 (NADP+), mitochondrial | 3418 | ENSG00000182054 |
| 11 | P38571 | NA | LIPA | lipase A, lysosomal acid, cholesterol esterase | 3988 | ENSG00000107798 |
| 12 | P31040 | NA | SDHA | succinate dehydrogenase complex, subunit A, flavoprotein (Fp) | 6389 | ENSG00000073578 |
| 13 | Q13011 | NA | ECH1 | enoyl CoA hydratase 1, peroxisomal | 1891 | ENSG00000104823 |
| 14 | P32322 | NA | PYCR1 | pyrroline-5-carboxylate reductase 1 | 5831 | ENSG00000183010 |
| 15 | Q99798 | NA | ACO2 | aconitase 2, mitochondrial | 50 | ENSG00000100412 |
| 16 | P51649 | NA | ALDH5A1 | aldehyde dehydrogenase 5 family, member A1 | 7915 | ENSG00000112294 |
| 17 | P42765 | NA | ACAA2 | acetyl-CoA acyltransferase 2 | 10449 | ENSG00000167315 |
| 18 | Q9NSE4 | NA | IARS2 | isoleucyl-tRNA synthetase 2, mitochondrial | 55699 | ENSG00000067704 |
| 19 | Q6PI48 | NA | DARS2 | aspartyl-tRNA synthetase 2, mitochondrial | 55157 | ENSG00000117593 |
| 20 | P24752 | NA | ACAT1 | acetyl-CoA acetyltransferase 1 | 38 | ENSG00000075239 |
| 21 | O75521 | NA | ECI2 | enoyl-CoA delta isomerase 2 | 10455 | ENSG00000198721 |
| 22 | P13674 | NA | P4HA1 | prolyl 4-hydroxylase, alpha polypeptide I | 5033 | ENSG00000122884 |
| 23 | P49419 | NA | ALDH7A1 | aldehyde dehydrogenase 7 family, member A1 | 501 | ENSG00000164904 |
| 24 | P06865 | NA | HEXA | hexosaminidase A (alpha polypeptide) | 3073 | ENSG00000213614 |
| 25 | P49748 | NA | ACADVL | acyl-CoA dehydrogenase, very long chain | 37 | ENSG00000072778 |
| 26 | P31937 | NA | HIBADH | 3-hydroxyisobutyrate dehydrogenase | 11112 | ENSG00000106049 |

  
  

| **Database:biological process      &nbspName:oxidation-reduction process      &nbspID:GO:0055114** | | | | | | |
| --- | --- | --- | --- | --- | --- | --- |
| C=554; O=21; E=2.77; R=7.59; rawP=1.60e-13; adjP=2.35e-11 | | | | | | |
| Index | UserID | Value | Gene Symbol | Gene Name | EntrezGene | Ensembl |
| 1 | P10253 | NA | GAA | glucosidase, alpha; acid | 2548 | ENSG00000171298 |
| 2 | P40939 | NA | HADHA | hydroxyacyl-CoA dehydrogenase/3-ketoacyl-CoA thiolase/enoyl-CoA hydratase (trifunctional protein), alpha subunit | 3030 | ENSG00000084754 |
| 3 | P40926 | NA | MDH2 | malate dehydrogenase 2, NAD (mitochondrial) | 4191 | ENSG00000146701 |
| 4 | P30084 | NA | ECHS1 | enoyl CoA hydratase, short chain, 1, mitochondrial | 1892 | ENSG00000127884 |
| 5 | P04179 | NA | SOD2 | superoxide dismutase 2, mitochondrial | 6648 | ENSG00000112096 |
| 6 | P48735 | NA | IDH2 | isocitrate dehydrogenase 2 (NADP+), mitochondrial | 3418 | ENSG00000182054 |
| 7 | P35270 | NA | SPR | sepiapterin reductase (7,8-dihydrobiopterin:NADP+ oxidoreductase) | 6697 | ENSG00000116096 |
| 8 | P31040 | NA | SDHA | succinate dehydrogenase complex, subunit A, flavoprotein (Fp) | 6389 | ENSG00000073578 |
| 9 | Q13011 | NA | ECH1 | enoyl CoA hydratase 1, peroxisomal | 1891 | ENSG00000104823 |
| 10 | P38117 | NA | ETFB | electron-transfer-flavoprotein, beta polypeptide | 2109 | ENSG00000105379 |
| 11 | P04040 | NA | CAT | catalase | 847 | ENSG00000121691 |
| 12 | Q99798 | NA | ACO2 | aconitase 2, mitochondrial | 50 | ENSG00000100412 |
| 13 | P51649 | NA | ALDH5A1 | aldehyde dehydrogenase 5 family, member A1 | 7915 | ENSG00000112294 |
| 14 | P12235 | NA | SLC25A4 | solute carrier family 25 (mitochondrial carrier; adenine nucleotide translocator), member 4 | 291 | ENSG00000151729 |
| 15 | P13804 | NA | ETFA | electron-transfer-flavoprotein, alpha polypeptide | 2108 | ENSG00000140374 |
| 16 | P42765 | NA | ACAA2 | acetyl-CoA acyltransferase 2 | 10449 | ENSG00000167315 |
| 17 | P25705 | NA | ATP5A1 | ATP synthase, H+ transporting, mitochondrial F1 complex, alpha subunit 1, cardiac muscle | 498 | ENSG00000152234 |
| 18 | P49748 | NA | ACADVL | acyl-CoA dehydrogenase, very long chain | 37 | ENSG00000072778 |
| 19 | P48047 | NA | ATP5O | ATP synthase, H+ transporting, mitochondrial F1 complex, O subunit | 539 | ENSG00000241837 |
| 20 | P60174 | NA | TPI1 | triosephosphate isomerase 1 | 7167 | ENSG00000111669 |
| 21 | P31937 | NA | HIBADH | 3-hydroxyisobutyrate dehydrogenase | 11112 | ENSG00000106049 |

  
  

| **Database:biological process      &nbspName:organic acid metabolic process      &nbspID:GO:0006082** | | | | | | |
| --- | --- | --- | --- | --- | --- | --- |
| C=967; O=26; E=4.83; R=5.39; rawP=3.10e-13; adjP=2.90e-11 | | | | | | |
| Index | UserID | Value | Gene Symbol | Gene Name | EntrezGene | Ensembl |
| 1 | P21953 | NA | BCKDHB | branched chain keto acid dehydrogenase E1, beta polypeptide | 594 | ENSG00000083123 |
| 2 | P40939 | NA | HADHA | hydroxyacyl-CoA dehydrogenase/3-ketoacyl-CoA thiolase/enoyl-CoA hydratase (trifunctional protein), alpha subunit | 3030 | ENSG00000084754 |
| 3 | Q9UL46 | NA | PSME2 | proteasome (prosome, macropain) activator subunit 2 (PA28 beta) | 5721 | ENSG00000100911 |
| 4 | P40926 | NA | MDH2 | malate dehydrogenase 2, NAD (mitochondrial) | 4191 | ENSG00000146701 |
| 5 | P30084 | NA | ECHS1 | enoyl CoA hydratase, short chain, 1, mitochondrial | 1892 | ENSG00000127884 |
| 6 | Q99714 | NA | HSD17B10 | hydroxysteroid (17-beta) dehydrogenase 10 | 3028 | ENSG00000072506 |
| 7 | P04179 | NA | SOD2 | superoxide dismutase 2, mitochondrial | 6648 | ENSG00000112096 |
| 8 | P00367 | NA | GLUD1 | glutamate dehydrogenase 1 | 2746 | ENSG00000148672 |
| 9 | Q06323 | NA | PSME1 | proteasome (prosome, macropain) activator subunit 1 (PA28 alpha) | 5720 | ENSG00000092010 |
| 10 | P48735 | NA | IDH2 | isocitrate dehydrogenase 2 (NADP+), mitochondrial | 3418 | ENSG00000182054 |
| 11 | P38571 | NA | LIPA | lipase A, lysosomal acid, cholesterol esterase | 3988 | ENSG00000107798 |
| 12 | P31040 | NA | SDHA | succinate dehydrogenase complex, subunit A, flavoprotein (Fp) | 6389 | ENSG00000073578 |
| 13 | Q13011 | NA | ECH1 | enoyl CoA hydratase 1, peroxisomal | 1891 | ENSG00000104823 |
| 14 | P32322 | NA | PYCR1 | pyrroline-5-carboxylate reductase 1 | 5831 | ENSG00000183010 |
| 15 | Q99798 | NA | ACO2 | aconitase 2, mitochondrial | 50 | ENSG00000100412 |
| 16 | P51649 | NA | ALDH5A1 | aldehyde dehydrogenase 5 family, member A1 | 7915 | ENSG00000112294 |
| 17 | P42765 | NA | ACAA2 | acetyl-CoA acyltransferase 2 | 10449 | ENSG00000167315 |
| 18 | Q9NSE4 | NA | IARS2 | isoleucyl-tRNA synthetase 2, mitochondrial | 55699 | ENSG00000067704 |
| 19 | Q6PI48 | NA | DARS2 | aspartyl-tRNA synthetase 2, mitochondrial | 55157 | ENSG00000117593 |
| 20 | P24752 | NA | ACAT1 | acetyl-CoA acetyltransferase 1 | 38 | ENSG00000075239 |
| 21 | O75521 | NA | ECI2 | enoyl-CoA delta isomerase 2 | 10455 | ENSG00000198721 |
| 22 | P13674 | NA | P4HA1 | prolyl 4-hydroxylase, alpha polypeptide I | 5033 | ENSG00000122884 |
| 23 | P49419 | NA | ALDH7A1 | aldehyde dehydrogenase 7 family, member A1 | 501 | ENSG00000164904 |
| 24 | P06865 | NA | HEXA | hexosaminidase A (alpha polypeptide) | 3073 | ENSG00000213614 |
| 25 | P49748 | NA | ACADVL | acyl-CoA dehydrogenase, very long chain | 37 | ENSG00000072778 |
| 26 | P31937 | NA | HIBADH | 3-hydroxyisobutyrate dehydrogenase | 11112 | ENSG00000106049 |

  
  

| **Database:biological process      &nbspName:organic substance catabolic process      &nbspID:GO:1901575** | | | | | | |
| --- | --- | --- | --- | --- | --- | --- |
| C=1851; O=32; E=9.24; R=3.46; rawP=3.57e-11; adjP=2.08e-09 | | | | | | |
| Index | UserID | Value | Gene Symbol | Gene Name | EntrezGene | Ensembl |
| 1 | P10253 | NA | GAA | glucosidase, alpha; acid | 2548 | ENSG00000171298 |
| 2 | P21953 | NA | BCKDHB | branched chain keto acid dehydrogenase E1, beta polypeptide | 594 | ENSG00000083123 |
| 3 | P40939 | NA | HADHA | hydroxyacyl-CoA dehydrogenase/3-ketoacyl-CoA thiolase/enoyl-CoA hydratase (trifunctional protein), alpha subunit | 3030 | ENSG00000084754 |
| 4 | Q9UL46 | NA | PSME2 | proteasome (prosome, macropain) activator subunit 2 (PA28 beta) | 5721 | ENSG00000100911 |
| 5 | P40926 | NA | MDH2 | malate dehydrogenase 2, NAD (mitochondrial) | 4191 | ENSG00000146701 |
| 6 | P30084 | NA | ECHS1 | enoyl CoA hydratase, short chain, 1, mitochondrial | 1892 | ENSG00000127884 |
| 7 | O95831 | NA | AIFM1 | apoptosis-inducing factor, mitochondrion-associated, 1 | 9131 | ENSG00000156709 |
| 8 | Q99714 | NA | HSD17B10 | hydroxysteroid (17-beta) dehydrogenase 10 | 3028 | ENSG00000072506 |
| 9 | P00367 | NA | GLUD1 | glutamate dehydrogenase 1 | 2746 | ENSG00000148672 |
| 10 | Q06323 | NA | PSME1 | proteasome (prosome, macropain) activator subunit 1 (PA28 alpha) | 5720 | ENSG00000092010 |
| 11 | P48735 | NA | IDH2 | isocitrate dehydrogenase 2 (NADP+), mitochondrial | 3418 | ENSG00000182054 |
| 12 | O60313 | NA | OPA1 | optic atrophy 1 (autosomal dominant) | 4976 | ENSG00000198836 |
| 13 | P38571 | NA | LIPA | lipase A, lysosomal acid, cholesterol esterase | 3988 | ENSG00000107798 |
| 14 | P31040 | NA | SDHA | succinate dehydrogenase complex, subunit A, flavoprotein (Fp) | 6389 | ENSG00000073578 |
| 15 | Q13011 | NA | ECH1 | enoyl CoA hydratase 1, peroxisomal | 1891 | ENSG00000104823 |
| 16 | P04040 | NA | CAT | catalase | 847 | ENSG00000121691 |
| 17 | O14773 | NA | TPP1 | tripeptidyl peptidase I | 1200 | ENSG00000166340 |
| 18 | Q99798 | NA | ACO2 | aconitase 2, mitochondrial | 50 | ENSG00000100412 |
| 19 | P51688 | NA | SGSH | N-sulfoglucosamine sulfohydrolase | 6448 | ENSG00000181523 |
| 20 | P51649 | NA | ALDH5A1 | aldehyde dehydrogenase 5 family, member A1 | 7915 | ENSG00000112294 |
| 21 | P00558 | NA | PGK1 | phosphoglycerate kinase 1 | 5230 | ENSG00000102144 |
| 22 | P42765 | NA | ACAA2 | acetyl-CoA acyltransferase 2 | 10449 | ENSG00000167315 |
| 23 | P24752 | NA | ACAT1 | acetyl-CoA acetyltransferase 1 | 38 | ENSG00000075239 |
| 24 | O75521 | NA | ECI2 | enoyl-CoA delta isomerase 2 | 10455 | ENSG00000198721 |
| 25 | P49419 | NA | ALDH7A1 | aldehyde dehydrogenase 7 family, member A1 | 501 | ENSG00000164904 |
| 26 | P19367 | NA | HK1 | hexokinase 1 | 3098 | ENSG00000156515 |
| 27 | P06865 | NA | HEXA | hexosaminidase A (alpha polypeptide) | 3073 | ENSG00000213614 |
| 28 | P49748 | NA | ACADVL | acyl-CoA dehydrogenase, very long chain | 37 | ENSG00000072778 |
| 29 | P48047 | NA | ATP5O | ATP synthase, H+ transporting, mitochondrial F1 complex, O subunit | 539 | ENSG00000241837 |
| 30 | Q16531 | NA | DDB1 | damage-specific DNA binding protein 1, 127kDa | 1642 | ENSG00000167986 |
| 31 | P60174 | NA | TPI1 | triosephosphate isomerase 1 | 7167 | ENSG00000111669 |
| 32 | P31937 | NA | HIBADH | 3-hydroxyisobutyrate dehydrogenase | 11112 | ENSG00000106049 |

  
  

| **Database:biological process      &nbspName:catabolic process      &nbspID:GO:0009056** | | | | | | |
| --- | --- | --- | --- | --- | --- | --- |
| C=1991; O=33; E=9.94; R=3.32; rawP=4.62e-11; adjP=2.19e-09 | | | | | | |
| Index | UserID | Value | Gene Symbol | Gene Name | EntrezGene | Ensembl |
| 1 | P10253 | NA | GAA | glucosidase, alpha; acid | 2548 | ENSG00000171298 |
| 2 | Q9UL46 | NA | PSME2 | proteasome (prosome, macropain) activator subunit 2 (PA28 beta) | 5721 | ENSG00000100911 |
| 3 | P40926 | NA | MDH2 | malate dehydrogenase 2, NAD (mitochondrial) | 4191 | ENSG00000146701 |
| 4 | O95831 | NA | AIFM1 | apoptosis-inducing factor, mitochondrion-associated, 1 | 9131 | ENSG00000156709 |
| 5 | Q99714 | NA | HSD17B10 | hydroxysteroid (17-beta) dehydrogenase 10 | 3028 | ENSG00000072506 |
| 6 | P00367 | NA | GLUD1 | glutamate dehydrogenase 1 | 2746 | ENSG00000148672 |
| 7 | Q06323 | NA | PSME1 | proteasome (prosome, macropain) activator subunit 1 (PA28 alpha) | 5720 | ENSG00000092010 |
| 8 | P48735 | NA | IDH2 | isocitrate dehydrogenase 2 (NADP+), mitochondrial | 3418 | ENSG00000182054 |
| 9 | O60313 | NA | OPA1 | optic atrophy 1 (autosomal dominant) | 4976 | ENSG00000198836 |
| 10 | P31040 | NA | SDHA | succinate dehydrogenase complex, subunit A, flavoprotein (Fp) | 6389 | ENSG00000073578 |
| 11 | O14773 | NA | TPP1 | tripeptidyl peptidase I | 1200 | ENSG00000166340 |
| 12 | P42765 | NA | ACAA2 | acetyl-CoA acyltransferase 2 | 10449 | ENSG00000167315 |
| 13 | O75521 | NA | ECI2 | enoyl-CoA delta isomerase 2 | 10455 | ENSG00000198721 |
| 14 | P49419 | NA | ALDH7A1 | aldehyde dehydrogenase 7 family, member A1 | 501 | ENSG00000164904 |
| 15 | P21953 | NA | BCKDHB | branched chain keto acid dehydrogenase E1, beta polypeptide | 594 | ENSG00000083123 |
| 16 | P40939 | NA | HADHA | hydroxyacyl-CoA dehydrogenase/3-ketoacyl-CoA thiolase/enoyl-CoA hydratase (trifunctional protein), alpha subunit | 3030 | ENSG00000084754 |
| 17 | P30084 | NA | ECHS1 | enoyl CoA hydratase, short chain, 1, mitochondrial | 1892 | ENSG00000127884 |
| 18 | P38571 | NA | LIPA | lipase A, lysosomal acid, cholesterol esterase | 3988 | ENSG00000107798 |
| 19 | Q13011 | NA | ECH1 | enoyl CoA hydratase 1, peroxisomal | 1891 | ENSG00000104823 |
| 20 | P04040 | NA | CAT | catalase | 847 | ENSG00000121691 |
| 21 | Q99798 | NA | ACO2 | aconitase 2, mitochondrial | 50 | ENSG00000100412 |
| 22 | P51649 | NA | ALDH5A1 | aldehyde dehydrogenase 5 family, member A1 | 7915 | ENSG00000112294 |
| 23 | P51688 | NA | SGSH | N-sulfoglucosamine sulfohydrolase | 6448 | ENSG00000181523 |
| 24 | P30048 | NA | PRDX3 | peroxiredoxin 3 | 10935 | ENSG00000165672 |
| 25 | P00558 | NA | PGK1 | phosphoglycerate kinase 1 | 5230 | ENSG00000102144 |
| 26 | P24752 | NA | ACAT1 | acetyl-CoA acetyltransferase 1 | 38 | ENSG00000075239 |
| 27 | P19367 | NA | HK1 | hexokinase 1 | 3098 | ENSG00000156515 |
| 28 | P49748 | NA | ACADVL | acyl-CoA dehydrogenase, very long chain | 37 | ENSG00000072778 |
| 29 | P06865 | NA | HEXA | hexosaminidase A (alpha polypeptide) | 3073 | ENSG00000213614 |
| 30 | P48047 | NA | ATP5O | ATP synthase, H+ transporting, mitochondrial F1 complex, O subunit | 539 | ENSG00000241837 |
| 31 | P60174 | NA | TPI1 | triosephosphate isomerase 1 | 7167 | ENSG00000111669 |
| 32 | Q16531 | NA | DDB1 | damage-specific DNA binding protein 1, 127kDa | 1642 | ENSG00000167986 |
| 33 | P31937 | NA | HIBADH | 3-hydroxyisobutyrate dehydrogenase | 11112 | ENSG00000106049 |

  
  

| **Database:biological process      &nbspName:cellular catabolic process      &nbspID:GO:0044248** | | | | | | |
| --- | --- | --- | --- | --- | --- | --- |
| C=1665; O=27; E=8.31; R=3.25; rawP=1.02e-08; adjP=3.18e-07 | | | | | | |
| Index | UserID | Value | Gene Symbol | Gene Name | EntrezGene | Ensembl |
| 1 | P21953 | NA | BCKDHB | branched chain keto acid dehydrogenase E1, beta polypeptide | 594 | ENSG00000083123 |
| 2 | P40939 | NA | HADHA | hydroxyacyl-CoA dehydrogenase/3-ketoacyl-CoA thiolase/enoyl-CoA hydratase (trifunctional protein), alpha subunit | 3030 | ENSG00000084754 |
| 3 | Q9UL46 | NA | PSME2 | proteasome (prosome, macropain) activator subunit 2 (PA28 beta) | 5721 | ENSG00000100911 |
| 4 | P40926 | NA | MDH2 | malate dehydrogenase 2, NAD (mitochondrial) | 4191 | ENSG00000146701 |
| 5 | P30084 | NA | ECHS1 | enoyl CoA hydratase, short chain, 1, mitochondrial | 1892 | ENSG00000127884 |
| 6 | O95831 | NA | AIFM1 | apoptosis-inducing factor, mitochondrion-associated, 1 | 9131 | ENSG00000156709 |
| 7 | Q99714 | NA | HSD17B10 | hydroxysteroid (17-beta) dehydrogenase 10 | 3028 | ENSG00000072506 |
| 8 | P00367 | NA | GLUD1 | glutamate dehydrogenase 1 | 2746 | ENSG00000148672 |
| 9 | Q06323 | NA | PSME1 | proteasome (prosome, macropain) activator subunit 1 (PA28 alpha) | 5720 | ENSG00000092010 |
| 10 | P48735 | NA | IDH2 | isocitrate dehydrogenase 2 (NADP+), mitochondrial | 3418 | ENSG00000182054 |
| 11 | O60313 | NA | OPA1 | optic atrophy 1 (autosomal dominant) | 4976 | ENSG00000198836 |
| 12 | P31040 | NA | SDHA | succinate dehydrogenase complex, subunit A, flavoprotein (Fp) | 6389 | ENSG00000073578 |
| 13 | Q13011 | NA | ECH1 | enoyl CoA hydratase 1, peroxisomal | 1891 | ENSG00000104823 |
| 14 | P04040 | NA | CAT | catalase | 847 | ENSG00000121691 |
| 15 | O14773 | NA | TPP1 | tripeptidyl peptidase I | 1200 | ENSG00000166340 |
| 16 | Q99798 | NA | ACO2 | aconitase 2, mitochondrial | 50 | ENSG00000100412 |
| 17 | P51649 | NA | ALDH5A1 | aldehyde dehydrogenase 5 family, member A1 | 7915 | ENSG00000112294 |
| 18 | P30048 | NA | PRDX3 | peroxiredoxin 3 | 10935 | ENSG00000165672 |
| 19 | P42765 | NA | ACAA2 | acetyl-CoA acyltransferase 2 | 10449 | ENSG00000167315 |
| 20 | P24752 | NA | ACAT1 | acetyl-CoA acetyltransferase 1 | 38 | ENSG00000075239 |
| 21 | O75521 | NA | ECI2 | enoyl-CoA delta isomerase 2 | 10455 | ENSG00000198721 |
| 22 | P49419 | NA | ALDH7A1 | aldehyde dehydrogenase 7 family, member A1 | 501 | ENSG00000164904 |
| 23 | P06865 | NA | HEXA | hexosaminidase A (alpha polypeptide) | 3073 | ENSG00000213614 |
| 24 | P49748 | NA | ACADVL | acyl-CoA dehydrogenase, very long chain | 37 | ENSG00000072778 |
| 25 | P48047 | NA | ATP5O | ATP synthase, H+ transporting, mitochondrial F1 complex, O subunit | 539 | ENSG00000241837 |
| 26 | Q16531 | NA | DDB1 | damage-specific DNA binding protein 1, 127kDa | 1642 | ENSG00000167986 |
| 27 | P31937 | NA | HIBADH | 3-hydroxyisobutyrate dehydrogenase | 11112 | ENSG00000106049 |

  
  

| **Database:biological process      &nbspName:fatty acid catabolic process      &nbspID:GO:0009062** | | | | | | |
| --- | --- | --- | --- | --- | --- | --- |
| C=75; O=7; E=0.37; R=18.70; rawP=8.69e-08; adjP=2.39e-06 | | | | | | |
| Index | UserID | Value | Gene Symbol | Gene Name | EntrezGene | Ensembl |
| 1 | O75521 | NA | ECI2 | enoyl-CoA delta isomerase 2 | 10455 | ENSG00000198721 |
| 2 | P51649 | NA | ALDH5A1 | aldehyde dehydrogenase 5 family, member A1 | 7915 | ENSG00000112294 |
| 3 | P49748 | NA | ACADVL | acyl-CoA dehydrogenase, very long chain | 37 | ENSG00000072778 |
| 4 | P40939 | NA | HADHA | hydroxyacyl-CoA dehydrogenase/3-ketoacyl-CoA thiolase/enoyl-CoA hydratase (trifunctional protein), alpha subunit | 3030 | ENSG00000084754 |
| 5 | Q13011 | NA | ECH1 | enoyl CoA hydratase 1, peroxisomal | 1891 | ENSG00000104823 |
| 6 | P30084 | NA | ECHS1 | enoyl CoA hydratase, short chain, 1, mitochondrial | 1892 | ENSG00000127884 |
| 7 | P42765 | NA | ACAA2 | acetyl-CoA acyltransferase 2 | 10449 | ENSG00000167315 |

  
  

| **Database:biological process      &nbspName:cellular amino acid metabolic process      &nbspID:GO:0006520** | | | | | | |
| --- | --- | --- | --- | --- | --- | --- |
| C=340; O=12; E=1.70; R=7.07; rawP=1.05e-07; adjP=2.72e-06 | | | | | | |
| Index | UserID | Value | Gene Symbol | Gene Name | EntrezGene | Ensembl |
| 1 | P51649 | NA | ALDH5A1 | aldehyde dehydrogenase 5 family, member A1 | 7915 | ENSG00000112294 |
| 2 | P21953 | NA | BCKDHB | branched chain keto acid dehydrogenase E1, beta polypeptide | 594 | ENSG00000083123 |
| 3 | Q9UL46 | NA | PSME2 | proteasome (prosome, macropain) activator subunit 2 (PA28 beta) | 5721 | ENSG00000100911 |
| 4 | Q99714 | NA | HSD17B10 | hydroxysteroid (17-beta) dehydrogenase 10 | 3028 | ENSG00000072506 |
| 5 | Q6PI48 | NA | DARS2 | aspartyl-tRNA synthetase 2, mitochondrial | 55157 | ENSG00000117593 |
| 6 | Q9NSE4 | NA | IARS2 | isoleucyl-tRNA synthetase 2, mitochondrial | 55699 | ENSG00000067704 |
| 7 | P24752 | NA | ACAT1 | acetyl-CoA acetyltransferase 1 | 38 | ENSG00000075239 |
| 8 | P49419 | NA | ALDH7A1 | aldehyde dehydrogenase 7 family, member A1 | 501 | ENSG00000164904 |
| 9 | P00367 | NA | GLUD1 | glutamate dehydrogenase 1 | 2746 | ENSG00000148672 |
| 10 | Q06323 | NA | PSME1 | proteasome (prosome, macropain) activator subunit 1 (PA28 alpha) | 5720 | ENSG00000092010 |
| 11 | P31937 | NA | HIBADH | 3-hydroxyisobutyrate dehydrogenase | 11112 | ENSG00000106049 |
| 12 | P32322 | NA | PYCR1 | pyrroline-5-carboxylate reductase 1 | 5831 | ENSG00000183010 |

  
  

| **Database:biological process      &nbspName:monocarboxylic acid catabolic process      &nbspID:GO:0072329** | | | | | | |
| --- | --- | --- | --- | --- | --- | --- |
| C=89; O=7; E=0.44; R=15.75; rawP=2.85e-07; adjP=7.00e-06 | | | | | | |
| Index | UserID | Value | Gene Symbol | Gene Name | EntrezGene | Ensembl |
| 1 | O75521 | NA | ECI2 | enoyl-CoA delta isomerase 2 | 10455 | ENSG00000198721 |
| 2 | P51649 | NA | ALDH5A1 | aldehyde dehydrogenase 5 family, member A1 | 7915 | ENSG00000112294 |
| 3 | P49748 | NA | ACADVL | acyl-CoA dehydrogenase, very long chain | 37 | ENSG00000072778 |
| 4 | P40939 | NA | HADHA | hydroxyacyl-CoA dehydrogenase/3-ketoacyl-CoA thiolase/enoyl-CoA hydratase (trifunctional protein), alpha subunit | 3030 | ENSG00000084754 |
| 5 | Q13011 | NA | ECH1 | enoyl CoA hydratase 1, peroxisomal | 1891 | ENSG00000104823 |
| 6 | P30084 | NA | ECHS1 | enoyl CoA hydratase, short chain, 1, mitochondrial | 1892 | ENSG00000127884 |
| 7 | P42765 | NA | ACAA2 | acetyl-CoA acyltransferase 2 | 10449 | ENSG00000167315 |

  
  

| **Database:biological process      &nbspName:cellular lipid catabolic process      &nbspID:GO:0044242** | | | | | | |
| --- | --- | --- | --- | --- | --- | --- |
| C=147; O=8; E=0.73; R=10.90; rawP=6.64e-07; adjP=1.55e-05 | | | | | | |
| Index | UserID | Value | Gene Symbol | Gene Name | EntrezGene | Ensembl |
| 1 | P51649 | NA | ALDH5A1 | aldehyde dehydrogenase 5 family, member A1 | 7915 | ENSG00000112294 |
| 2 | P40939 | NA | HADHA | hydroxyacyl-CoA dehydrogenase/3-ketoacyl-CoA thiolase/enoyl-CoA hydratase (trifunctional protein), alpha subunit | 3030 | ENSG00000084754 |
| 3 | P30084 | NA | ECHS1 | enoyl CoA hydratase, short chain, 1, mitochondrial | 1892 | ENSG00000127884 |
| 4 | P42765 | NA | ACAA2 | acetyl-CoA acyltransferase 2 | 10449 | ENSG00000167315 |
| 5 | O75521 | NA | ECI2 | enoyl-CoA delta isomerase 2 | 10455 | ENSG00000198721 |
| 6 | P49748 | NA | ACADVL | acyl-CoA dehydrogenase, very long chain | 37 | ENSG00000072778 |
| 7 | P06865 | NA | HEXA | hexosaminidase A (alpha polypeptide) | 3073 | ENSG00000213614 |
| 8 | Q13011 | NA | ECH1 | enoyl CoA hydratase 1, peroxisomal | 1891 | ENSG00000104823 |

  
  

| **Database:biological process      &nbspName:cellular amino acid catabolic process      &nbspID:GO:0009063** | | | | | | |
| --- | --- | --- | --- | --- | --- | --- |
| C=104; O=7; E=0.52; R=13.48; rawP=8.30e-07; adjP=1.85e-05 | | | | | | |
| Index | UserID | Value | Gene Symbol | Gene Name | EntrezGene | Ensembl |
| 1 | P24752 | NA | ACAT1 | acetyl-CoA acetyltransferase 1 | 38 | ENSG00000075239 |
| 2 | P49419 | NA | ALDH7A1 | aldehyde dehydrogenase 7 family, member A1 | 501 | ENSG00000164904 |
| 3 | P00367 | NA | GLUD1 | glutamate dehydrogenase 1 | 2746 | ENSG00000148672 |
| 4 | P51649 | NA | ALDH5A1 | aldehyde dehydrogenase 5 family, member A1 | 7915 | ENSG00000112294 |
| 5 | P21953 | NA | BCKDHB | branched chain keto acid dehydrogenase E1, beta polypeptide | 594 | ENSG00000083123 |
| 6 | P31937 | NA | HIBADH | 3-hydroxyisobutyrate dehydrogenase | 11112 | ENSG00000106049 |
| 7 | Q99714 | NA | HSD17B10 | hydroxysteroid (17-beta) dehydrogenase 10 | 3028 | ENSG00000072506 |

  
  

| **Database:biological process      &nbspName:coenzyme metabolic process      &nbspID:GO:0006732** | | | | | | |
| --- | --- | --- | --- | --- | --- | --- |
| C=213; O=9; E=1.06; R=8.46; rawP=1.07e-06; adjP=2.27e-05 | | | | | | |
| Index | UserID | Value | Gene Symbol | Gene Name | EntrezGene | Ensembl |
| 1 | Q99798 | NA | ACO2 | aconitase 2, mitochondrial | 50 | ENSG00000100412 |
| 2 | P40926 | NA | MDH2 | malate dehydrogenase 2, NAD (mitochondrial) | 4191 | ENSG00000146701 |
| 3 | P42765 | NA | ACAA2 | acetyl-CoA acyltransferase 2 | 10449 | ENSG00000167315 |
| 4 | P48735 | NA | IDH2 | isocitrate dehydrogenase 2 (NADP+), mitochondrial | 3418 | ENSG00000182054 |
| 5 | P35270 | NA | SPR | sepiapterin reductase (7,8-dihydrobiopterin:NADP+ oxidoreductase) | 6697 | ENSG00000116096 |
| 6 | P31040 | NA | SDHA | succinate dehydrogenase complex, subunit A, flavoprotein (Fp) | 6389 | ENSG00000073578 |
| 7 | P60174 | NA | TPI1 | triosephosphate isomerase 1 | 7167 | ENSG00000111669 |
| 8 | P31937 | NA | HIBADH | 3-hydroxyisobutyrate dehydrogenase | 11112 | ENSG00000106049 |
| 9 | P43490 | NA | NAMPT | nicotinamide phosphoribosyltransferase | 10135 | ENSG00000105835 |

  
  

| **Database:biological process      &nbspName:cofactor metabolic process      &nbspID:GO:0051186** | | | | | | |
| --- | --- | --- | --- | --- | --- | --- |
| C=255; O=9; E=1.27; R=7.07; rawP=4.73e-06; adjP=7.62e-05 | | | | | | |
| Index | UserID | Value | Gene Symbol | Gene Name | EntrezGene | Ensembl |
| 1 | Q99798 | NA | ACO2 | aconitase 2, mitochondrial | 50 | ENSG00000100412 |
| 2 | P40926 | NA | MDH2 | malate dehydrogenase 2, NAD (mitochondrial) | 4191 | ENSG00000146701 |
| 3 | P42765 | NA | ACAA2 | acetyl-CoA acyltransferase 2 | 10449 | ENSG00000167315 |
| 4 | P48735 | NA | IDH2 | isocitrate dehydrogenase 2 (NADP+), mitochondrial | 3418 | ENSG00000182054 |
| 5 | P35270 | NA | SPR | sepiapterin reductase (7,8-dihydrobiopterin:NADP+ oxidoreductase) | 6697 | ENSG00000116096 |
| 6 | P31040 | NA | SDHA | succinate dehydrogenase complex, subunit A, flavoprotein (Fp) | 6389 | ENSG00000073578 |
| 7 | P60174 | NA | TPI1 | triosephosphate isomerase 1 | 7167 | ENSG00000111669 |
| 8 | P31937 | NA | HIBADH | 3-hydroxyisobutyrate dehydrogenase | 11112 | ENSG00000106049 |
| 9 | P43490 | NA | NAMPT | nicotinamide phosphoribosyltransferase | 10135 | ENSG00000105835 |

  
  

| **Database:biological process      &nbspName:branched-chain amino acid metabolic process      &nbspID:GO:0009081** | | | | | | |
| --- | --- | --- | --- | --- | --- | --- |
| C=23; O=4; E=0.11; R=34.84; rawP=4.71e-06; adjP=7.62e-05 | | | | | | |
| Index | UserID | Value | Gene Symbol | Gene Name | EntrezGene | Ensembl |
| 1 | P24752 | NA | ACAT1 | acetyl-CoA acetyltransferase 1 | 38 | ENSG00000075239 |
| 2 | P21953 | NA | BCKDHB | branched chain keto acid dehydrogenase E1, beta polypeptide | 594 | ENSG00000083123 |
| 3 | P31937 | NA | HIBADH | 3-hydroxyisobutyrate dehydrogenase | 11112 | ENSG00000106049 |
| 4 | Q99714 | NA | HSD17B10 | hydroxysteroid (17-beta) dehydrogenase 10 | 3028 | ENSG00000072506 |

  
  

| **Database:biological process      &nbspName:acetyl-CoA metabolic process      &nbspID:GO:0006084** | | | | | | |
| --- | --- | --- | --- | --- | --- | --- |
| C=52; O=5; E=0.26; R=19.26; rawP=5.83e-06; adjP=9.08e-05 | | | | | | |
| Index | UserID | Value | Gene Symbol | Gene Name | EntrezGene | Ensembl |
| 1 | Q99798 | NA | ACO2 | aconitase 2, mitochondrial | 50 | ENSG00000100412 |
| 2 | P48735 | NA | IDH2 | isocitrate dehydrogenase 2 (NADP+), mitochondrial | 3418 | ENSG00000182054 |
| 3 | P31040 | NA | SDHA | succinate dehydrogenase complex, subunit A, flavoprotein (Fp) | 6389 | ENSG00000073578 |
| 4 | P40926 | NA | MDH2 | malate dehydrogenase 2, NAD (mitochondrial) | 4191 | ENSG00000146701 |
| 5 | P42765 | NA | ACAA2 | acetyl-CoA acyltransferase 2 | 10449 | ENSG00000167315 |

  
  

| **Database:biological process      &nbspName:acetyl-CoA catabolic process      &nbspID:GO:0046356** | | | | | | |
| --- | --- | --- | --- | --- | --- | --- |
| C=27; O=4; E=0.13; R=29.67; rawP=9.19e-06; adjP=0.0001 | | | | | | |
| Index | UserID | Value | Gene Symbol | Gene Name | EntrezGene | Ensembl |
| 1 | Q99798 | NA | ACO2 | aconitase 2, mitochondrial | 50 | ENSG00000100412 |
| 2 | P48735 | NA | IDH2 | isocitrate dehydrogenase 2 (NADP+), mitochondrial | 3418 | ENSG00000182054 |
| 3 | P31040 | NA | SDHA | succinate dehydrogenase complex, subunit A, flavoprotein (Fp) | 6389 | ENSG00000073578 |
| 4 | P40926 | NA | MDH2 | malate dehydrogenase 2, NAD (mitochondrial) | 4191 | ENSG00000146701 |

  
  

| **Database:biological process      &nbspName:fatty acid beta-oxidation      &nbspID:GO:0006635** | | | | | | |
| --- | --- | --- | --- | --- | --- | --- |
| C=57; O=5; E=0.28; R=17.57; rawP=9.22e-06; adjP=0.0001 | | | | | | |
| Index | UserID | Value | Gene Symbol | Gene Name | EntrezGene | Ensembl |
| 1 | P49748 | NA | ACADVL | acyl-CoA dehydrogenase, very long chain | 37 | ENSG00000072778 |
| 2 | P40939 | NA | HADHA | hydroxyacyl-CoA dehydrogenase/3-ketoacyl-CoA thiolase/enoyl-CoA hydratase (trifunctional protein), alpha subunit | 3030 | ENSG00000084754 |
| 3 | Q13011 | NA | ECH1 | enoyl CoA hydratase 1, peroxisomal | 1891 | ENSG00000104823 |
| 4 | P30084 | NA | ECHS1 | enoyl CoA hydratase, short chain, 1, mitochondrial | 1892 | ENSG00000127884 |
| 5 | P42765 | NA | ACAA2 | acetyl-CoA acyltransferase 2 | 10449 | ENSG00000167315 |

  
  

| **Database:biological process      &nbspName:tricarboxylic acid cycle      &nbspID:GO:0006099** | | | | | | |
| --- | --- | --- | --- | --- | --- | --- |
| C=26; O=4; E=0.13; R=30.82; rawP=7.86e-06; adjP=0.0001 | | | | | | |
| Index | UserID | Value | Gene Symbol | Gene Name | EntrezGene | Ensembl |
| 1 | Q99798 | NA | ACO2 | aconitase 2, mitochondrial | 50 | ENSG00000100412 |
| 2 | P48735 | NA | IDH2 | isocitrate dehydrogenase 2 (NADP+), mitochondrial | 3418 | ENSG00000182054 |
| 3 | P31040 | NA | SDHA | succinate dehydrogenase complex, subunit A, flavoprotein (Fp) | 6389 | ENSG00000073578 |
| 4 | P40926 | NA | MDH2 | malate dehydrogenase 2, NAD (mitochondrial) | 4191 | ENSG00000146701 |

  
  

| **Database:biological process      &nbspName:glucose metabolic process      &nbspID:GO:0006006** | | | | | | |
| --- | --- | --- | --- | --- | --- | --- |
| C=213; O=8; E=1.06; R=7.52; rawP=1.06e-05; adjP=0.0001 | | | | | | |
| Index | UserID | Value | Gene Symbol | Gene Name | EntrezGene | Ensembl |
| 1 | Q02978 | NA | SLC25A11 | solute carrier family 25 (mitochondrial carrier; oxoglutarate carrier), member 11 | 8402 | ENSG00000108528 |
| 2 | P10253 | NA | GAA | glucosidase, alpha; acid | 2548 | ENSG00000171298 |
| 3 | P19367 | NA | HK1 | hexokinase 1 | 3098 | ENSG00000156515 |
| 4 | P51649 | NA | ALDH5A1 | aldehyde dehydrogenase 5 family, member A1 | 7915 | ENSG00000112294 |
| 5 | P00558 | NA | PGK1 | phosphoglycerate kinase 1 | 5230 | ENSG00000102144 |
| 6 | P40926 | NA | MDH2 | malate dehydrogenase 2, NAD (mitochondrial) | 4191 | ENSG00000146701 |
| 7 | P60174 | NA | TPI1 | triosephosphate isomerase 1 | 7167 | ENSG00000111669 |
| 8 | P31937 | NA | HIBADH | 3-hydroxyisobutyrate dehydrogenase | 11112 | ENSG00000106049 |

  
  

| **Database:biological process      &nbspName:electron transport chain      &nbspID:GO:0022900** | | | | | | |
| --- | --- | --- | --- | --- | --- | --- |
| C=147; O=7; E=0.73; R=9.54; rawP=8.39e-06; adjP=0.0001 | | | | | | |
| Index | UserID | Value | Gene Symbol | Gene Name | EntrezGene | Ensembl |
| 1 | P25705 | NA | ATP5A1 | ATP synthase, H+ transporting, mitochondrial F1 complex, alpha subunit 1, cardiac muscle | 498 | ENSG00000152234 |
| 2 | P51649 | NA | ALDH5A1 | aldehyde dehydrogenase 5 family, member A1 | 7915 | ENSG00000112294 |
| 3 | P48047 | NA | ATP5O | ATP synthase, H+ transporting, mitochondrial F1 complex, O subunit | 539 | ENSG00000241837 |
| 4 | P31040 | NA | SDHA | succinate dehydrogenase complex, subunit A, flavoprotein (Fp) | 6389 | ENSG00000073578 |
| 5 | P38117 | NA | ETFB | electron-transfer-flavoprotein, beta polypeptide | 2109 | ENSG00000105379 |
| 6 | P13804 | NA | ETFA | electron-transfer-flavoprotein, alpha polypeptide | 2108 | ENSG00000140374 |
| 7 | P04179 | NA | SOD2 | superoxide dismutase 2, mitochondrial | 6648 | ENSG00000112096 |

  
  

| **Database:biological process      &nbspName:hexose metabolic process      &nbspID:GO:0019318** | | | | | | |
| --- | --- | --- | --- | --- | --- | --- |
| C=249; O=8; E=1.24; R=6.44; rawP=3.26e-05; adjP=0.0004 | | | | | | |
| Index | UserID | Value | Gene Symbol | Gene Name | EntrezGene | Ensembl |
| 1 | Q02978 | NA | SLC25A11 | solute carrier family 25 (mitochondrial carrier; oxoglutarate carrier), member 11 | 8402 | ENSG00000108528 |
| 2 | P10253 | NA | GAA | glucosidase, alpha; acid | 2548 | ENSG00000171298 |
| 3 | P19367 | NA | HK1 | hexokinase 1 | 3098 | ENSG00000156515 |
| 4 | P51649 | NA | ALDH5A1 | aldehyde dehydrogenase 5 family, member A1 | 7915 | ENSG00000112294 |
| 5 | P00558 | NA | PGK1 | phosphoglycerate kinase 1 | 5230 | ENSG00000102144 |
| 6 | P40926 | NA | MDH2 | malate dehydrogenase 2, NAD (mitochondrial) | 4191 | ENSG00000146701 |
| 7 | P60174 | NA | TPI1 | triosephosphate isomerase 1 | 7167 | ENSG00000111669 |
| 8 | P31937 | NA | HIBADH | 3-hydroxyisobutyrate dehydrogenase | 11112 | ENSG00000106049 |

  
  

| **Database:biological process      &nbspName:cofactor catabolic process      &nbspID:GO:0051187** | | | | | | |
| --- | --- | --- | --- | --- | --- | --- |
| C=39; O=4; E=0.19; R=20.54; rawP=4.12e-05; adjP=0.0005 | | | | | | |
| Index | UserID | Value | Gene Symbol | Gene Name | EntrezGene | Ensembl |
| 1 | Q99798 | NA | ACO2 | aconitase 2, mitochondrial | 50 | ENSG00000100412 |
| 2 | P48735 | NA | IDH2 | isocitrate dehydrogenase 2 (NADP+), mitochondrial | 3418 | ENSG00000182054 |
| 3 | P31040 | NA | SDHA | succinate dehydrogenase complex, subunit A, flavoprotein (Fp) | 6389 | ENSG00000073578 |
| 4 | P40926 | NA | MDH2 | malate dehydrogenase 2, NAD (mitochondrial) | 4191 | ENSG00000146701 |

  
  

| **Database:biological process      &nbspName:organonitrogen compound catabolic process      &nbspID:GO:1901565** | | | | | | |
| --- | --- | --- | --- | --- | --- | --- |
| C=711; O=13; E=3.55; R=3.66; rawP=4.37e-05; adjP=0.0005 | | | | | | |
| Index | UserID | Value | Gene Symbol | Gene Name | EntrezGene | Ensembl |
| 1 | P04040 | NA | CAT | catalase | 847 | ENSG00000121691 |
| 2 | O14773 | NA | TPP1 | tripeptidyl peptidase I | 1200 | ENSG00000166340 |
| 3 | P51688 | NA | SGSH | N-sulfoglucosamine sulfohydrolase | 6448 | ENSG00000181523 |
| 4 | P51649 | NA | ALDH5A1 | aldehyde dehydrogenase 5 family, member A1 | 7915 | ENSG00000112294 |
| 5 | P21953 | NA | BCKDHB | branched chain keto acid dehydrogenase E1, beta polypeptide | 594 | ENSG00000083123 |
| 6 | Q99714 | NA | HSD17B10 | hydroxysteroid (17-beta) dehydrogenase 10 | 3028 | ENSG00000072506 |
| 7 | P24752 | NA | ACAT1 | acetyl-CoA acetyltransferase 1 | 38 | ENSG00000075239 |
| 8 | P00367 | NA | GLUD1 | glutamate dehydrogenase 1 | 2746 | ENSG00000148672 |
| 9 | P49419 | NA | ALDH7A1 | aldehyde dehydrogenase 7 family, member A1 | 501 | ENSG00000164904 |
| 10 | O60313 | NA | OPA1 | optic atrophy 1 (autosomal dominant) | 4976 | ENSG00000198836 |
| 11 | P06865 | NA | HEXA | hexosaminidase A (alpha polypeptide) | 3073 | ENSG00000213614 |
| 12 | P48047 | NA | ATP5O | ATP synthase, H+ transporting, mitochondrial F1 complex, O subunit | 539 | ENSG00000241837 |
| 13 | P31937 | NA | HIBADH | 3-hydroxyisobutyrate dehydrogenase | 11112 | ENSG00000106049 |

  
  

| **Database:molecular function      &nbspName:coenzyme binding      &nbspID:GO:0050662** | | | | | | |
| --- | --- | --- | --- | --- | --- | --- |
| C=184; O=13; E=0.88; R=14.70; rawP=3.57e-12; adjP=3.03e-10 | | | | | | |
| Index | UserID | Value | Gene Symbol | Gene Name | EntrezGene | Ensembl |
| 1 | P04040 | NA | CAT | catalase | 847 | ENSG00000121691 |
| 2 | P40939 | NA | HADHA | hydroxyacyl-CoA dehydrogenase/3-ketoacyl-CoA thiolase/enoyl-CoA hydratase (trifunctional protein), alpha subunit | 3030 | ENSG00000084754 |
| 3 | O95831 | NA | AIFM1 | apoptosis-inducing factor, mitochondrion-associated, 1 | 9131 | ENSG00000156709 |
| 4 | Q99714 | NA | HSD17B10 | hydroxysteroid (17-beta) dehydrogenase 10 | 3028 | ENSG00000072506 |
| 5 | P13804 | NA | ETFA | electron-transfer-flavoprotein, alpha polypeptide | 2108 | ENSG00000140374 |
| 6 | P24752 | NA | ACAT1 | acetyl-CoA acetyltransferase 1 | 38 | ENSG00000075239 |
| 7 | O75521 | NA | ECI2 | enoyl-CoA delta isomerase 2 | 10455 | ENSG00000198721 |
| 8 | P00367 | NA | GLUD1 | glutamate dehydrogenase 1 | 2746 | ENSG00000148672 |
| 9 | P48735 | NA | IDH2 | isocitrate dehydrogenase 2 (NADP+), mitochondrial | 3418 | ENSG00000182054 |
| 10 | P49748 | NA | ACADVL | acyl-CoA dehydrogenase, very long chain | 37 | ENSG00000072778 |
| 11 | P35270 | NA | SPR | sepiapterin reductase (7,8-dihydrobiopterin:NADP+ oxidoreductase) | 6697 | ENSG00000116096 |
| 12 | P31040 | NA | SDHA | succinate dehydrogenase complex, subunit A, flavoprotein (Fp) | 6389 | ENSG00000073578 |
| 13 | P31937 | NA | HIBADH | 3-hydroxyisobutyrate dehydrogenase | 11112 | ENSG00000106049 |

  
  

| **Database:molecular function      &nbspName:cofactor binding      &nbspID:GO:0048037** | | | | | | |
| --- | --- | --- | --- | --- | --- | --- |
| C=259; O=13; E=1.25; R=10.44; rawP=2.62e-10; adjP=7.42e-09 | | | | | | |
| Index | UserID | Value | Gene Symbol | Gene Name | EntrezGene | Ensembl |
| 1 | P04040 | NA | CAT | catalase | 847 | ENSG00000121691 |
| 2 | P40939 | NA | HADHA | hydroxyacyl-CoA dehydrogenase/3-ketoacyl-CoA thiolase/enoyl-CoA hydratase (trifunctional protein), alpha subunit | 3030 | ENSG00000084754 |
| 3 | O95831 | NA | AIFM1 | apoptosis-inducing factor, mitochondrion-associated, 1 | 9131 | ENSG00000156709 |
| 4 | Q99714 | NA | HSD17B10 | hydroxysteroid (17-beta) dehydrogenase 10 | 3028 | ENSG00000072506 |
| 5 | P13804 | NA | ETFA | electron-transfer-flavoprotein, alpha polypeptide | 2108 | ENSG00000140374 |
| 6 | P24752 | NA | ACAT1 | acetyl-CoA acetyltransferase 1 | 38 | ENSG00000075239 |
| 7 | O75521 | NA | ECI2 | enoyl-CoA delta isomerase 2 | 10455 | ENSG00000198721 |
| 8 | P00367 | NA | GLUD1 | glutamate dehydrogenase 1 | 2746 | ENSG00000148672 |
| 9 | P48735 | NA | IDH2 | isocitrate dehydrogenase 2 (NADP+), mitochondrial | 3418 | ENSG00000182054 |
| 10 | P49748 | NA | ACADVL | acyl-CoA dehydrogenase, very long chain | 37 | ENSG00000072778 |
| 11 | P35270 | NA | SPR | sepiapterin reductase (7,8-dihydrobiopterin:NADP+ oxidoreductase) | 6697 | ENSG00000116096 |
| 12 | P31040 | NA | SDHA | succinate dehydrogenase complex, subunit A, flavoprotein (Fp) | 6389 | ENSG00000073578 |
| 13 | P31937 | NA | HIBADH | 3-hydroxyisobutyrate dehydrogenase | 11112 | ENSG00000106049 |

  
  

| **Database:molecular function      &nbspName:catalytic activity      &nbspID:GO:0003824** | | | | | | |
| --- | --- | --- | --- | --- | --- | --- |
| C=5299; O=48; E=25.47; R=1.88; rawP=6.17e-08; adjP=1.31e-06 | | | | | | |
| Index | UserID | Value | Gene Symbol | Gene Name | EntrezGene | Ensembl |
| 1 | P10253 | NA | GAA | glucosidase, alpha; acid | 2548 | ENSG00000171298 |
| 2 | P54819 | NA | AK2 | adenylate kinase 2 | 204 | ENSG00000004455 |
| 3 | P40926 | NA | MDH2 | malate dehydrogenase 2, NAD (mitochondrial) | 4191 | ENSG00000146701 |
| 4 | O95831 | NA | AIFM1 | apoptosis-inducing factor, mitochondrion-associated, 1 | 9131 | ENSG00000156709 |
| 5 | Q99714 | NA | HSD17B10 | hydroxysteroid (17-beta) dehydrogenase 10 | 3028 | ENSG00000072506 |
| 6 | P04179 | NA | SOD2 | superoxide dismutase 2, mitochondrial | 6648 | ENSG00000112096 |
| 7 | Q13724 | NA | MOGS | mannosyl-oligosaccharide glucosidase | 7841 | ENSG00000115275 |
| 8 | P07741 | NA | APRT | adenine phosphoribosyltransferase | 353 | ENSG00000198931 |
| 9 | P00367 | NA | GLUD1 | glutamate dehydrogenase 1 | 2746 | ENSG00000148672 |
| 10 | P48735 | NA | IDH2 | isocitrate dehydrogenase 2 (NADP+), mitochondrial | 3418 | ENSG00000182054 |
| 11 | O60313 | NA | OPA1 | optic atrophy 1 (autosomal dominant) | 4976 | ENSG00000198836 |
| 12 | P35270 | NA | SPR | sepiapterin reductase (7,8-dihydrobiopterin:NADP+ oxidoreductase) | 6697 | ENSG00000116096 |
| 13 | P35580 | NA | MYH10 | myosin, heavy chain 10, non-muscle | 4628 | ENSG00000133026 |
| 14 | P31040 | NA | SDHA | succinate dehydrogenase complex, subunit A, flavoprotein (Fp) | 6389 | ENSG00000073578 |
| 15 | P43490 | NA | NAMPT | nicotinamide phosphoribosyltransferase | 10135 | ENSG00000105835 |
| 16 | O14773 | NA | TPP1 | tripeptidyl peptidase I | 1200 | ENSG00000166340 |
| 17 | P13804 | NA | ETFA | electron-transfer-flavoprotein, alpha polypeptide | 2108 | ENSG00000140374 |
| 18 | P42765 | NA | ACAA2 | acetyl-CoA acyltransferase 2 | 10449 | ENSG00000167315 |
| 19 | O75521 | NA | ECI2 | enoyl-CoA delta isomerase 2 | 10455 | ENSG00000198721 |
| 20 | P49419 | NA | ALDH7A1 | aldehyde dehydrogenase 7 family, member A1 | 501 | ENSG00000164904 |
| 21 | Q14697 | NA | GANAB | glucosidase, alpha; neutral AB | 23193 | ENSG00000089597 |
| 22 | P02792 | NA | FTL | ferritin, light polypeptide | 2512 | ENSG00000087086 |
| 23 | O75439 | NA | PMPCB | peptidase (mitochondrial processing) beta | 9512 | ENSG00000105819 |
| 24 | P21953 | NA | BCKDHB | branched chain keto acid dehydrogenase E1, beta polypeptide | 594 | ENSG00000083123 |
| 25 | P40939 | NA | HADHA | hydroxyacyl-CoA dehydrogenase/3-ketoacyl-CoA thiolase/enoyl-CoA hydratase (trifunctional protein), alpha subunit | 3030 | ENSG00000084754 |
| 26 | P30084 | NA | ECHS1 | enoyl CoA hydratase, short chain, 1, mitochondrial | 1892 | ENSG00000127884 |
| 27 | Q9BXW7 | NA | CECR5 | cat eye syndrome chromosome region, candidate 5 | 27440 | ENSG00000069998 |
| 28 | P38571 | NA | LIPA | lipase A, lysosomal acid, cholesterol esterase | 3988 | ENSG00000107798 |
| 29 | Q13011 | NA | ECH1 | enoyl CoA hydratase 1, peroxisomal | 1891 | ENSG00000104823 |
| 30 | P32322 | NA | PYCR1 | pyrroline-5-carboxylate reductase 1 | 5831 | ENSG00000183010 |
| 31 | P04040 | NA | CAT | catalase | 847 | ENSG00000121691 |
| 32 | Q99798 | NA | ACO2 | aconitase 2, mitochondrial | 50 | ENSG00000100412 |
| 33 | P51688 | NA | SGSH | N-sulfoglucosamine sulfohydrolase | 6448 | ENSG00000181523 |
| 34 | P51649 | NA | ALDH5A1 | aldehyde dehydrogenase 5 family, member A1 | 7915 | ENSG00000112294 |
| 35 | P00558 | NA | PGK1 | phosphoglycerate kinase 1 | 5230 | ENSG00000102144 |
| 36 | P30048 | NA | PRDX3 | peroxiredoxin 3 | 10935 | ENSG00000165672 |
| 37 | Q9NSE4 | NA | IARS2 | isoleucyl-tRNA synthetase 2, mitochondrial | 55699 | ENSG00000067704 |
| 38 | Q6PI48 | NA | DARS2 | aspartyl-tRNA synthetase 2, mitochondrial | 55157 | ENSG00000117593 |
| 39 | P24752 | NA | ACAT1 | acetyl-CoA acetyltransferase 1 | 38 | ENSG00000075239 |
| 40 | P13674 | NA | P4HA1 | prolyl 4-hydroxylase, alpha polypeptide I | 5033 | ENSG00000122884 |
| 41 | P19367 | NA | HK1 | hexokinase 1 | 3098 | ENSG00000156515 |
| 42 | P25705 | NA | ATP5A1 | ATP synthase, H+ transporting, mitochondrial F1 complex, alpha subunit 1, cardiac muscle | 498 | ENSG00000152234 |
| 43 | Q9BQ69 | NA | MACROD1 | MACRO domain containing 1 | 28992 | ENSG00000133315 |
| 44 | P06865 | NA | HEXA | hexosaminidase A (alpha polypeptide) | 3073 | ENSG00000213614 |
| 45 | P49748 | NA | ACADVL | acyl-CoA dehydrogenase, very long chain | 37 | ENSG00000072778 |
| 46 | P48047 | NA | ATP5O | ATP synthase, H+ transporting, mitochondrial F1 complex, O subunit | 539 | ENSG00000241837 |
| 47 | P60174 | NA | TPI1 | triosephosphate isomerase 1 | 7167 | ENSG00000111669 |
| 48 | P31937 | NA | HIBADH | 3-hydroxyisobutyrate dehydrogenase | 11112 | ENSG00000106049 |

  
  

| **Database:molecular function      &nbspName:oxidoreductase activity, acting on the CH-OH group of donors, NAD or NADP as acceptor      &nbspID:GO:0016616** | | | | | | |
| --- | --- | --- | --- | --- | --- | --- |
| C=116; O=6; E=0.56; R=10.76; rawP=1.95e-05; adjP=0.0002 | | | | | | |
| Index | UserID | Value | Gene Symbol | Gene Name | EntrezGene | Ensembl |
| 1 | P48735 | NA | IDH2 | isocitrate dehydrogenase 2 (NADP+), mitochondrial | 3418 | ENSG00000182054 |
| 2 | P35270 | NA | SPR | sepiapterin reductase (7,8-dihydrobiopterin:NADP+ oxidoreductase) | 6697 | ENSG00000116096 |
| 3 | P40939 | NA | HADHA | hydroxyacyl-CoA dehydrogenase/3-ketoacyl-CoA thiolase/enoyl-CoA hydratase (trifunctional protein), alpha subunit | 3030 | ENSG00000084754 |
| 4 | P40926 | NA | MDH2 | malate dehydrogenase 2, NAD (mitochondrial) | 4191 | ENSG00000146701 |
| 5 | P31937 | NA | HIBADH | 3-hydroxyisobutyrate dehydrogenase | 11112 | ENSG00000106049 |
| 6 | Q99714 | NA | HSD17B10 | hydroxysteroid (17-beta) dehydrogenase 10 | 3028 | ENSG00000072506 |

  
  

| **Database:molecular function      &nbspName:ADP binding      &nbspID:GO:0043531** | | | | | | |
| --- | --- | --- | --- | --- | --- | --- |
| C=25; O=3; E=0.12; R=24.96; rawP=0.0002; adjP=0.0014 | | | | | | |
| Index | UserID | Value | Gene Symbol | Gene Name | EntrezGene | Ensembl |
| 1 | P00367 | NA | GLUD1 | glutamate dehydrogenase 1 | 2746 | ENSG00000148672 |
| 2 | P25705 | NA | ATP5A1 | ATP synthase, H+ transporting, mitochondrial F1 complex, alpha subunit 1, cardiac muscle | 498 | ENSG00000152234 |
| 3 | P35580 | NA | MYH10 | myosin, heavy chain 10, non-muscle | 4628 | ENSG00000133026 |

  
  

| **Database:molecular function      &nbspName:flavin adenine dinucleotide binding      &nbspID:GO:0050660** | | | | | | |
| --- | --- | --- | --- | --- | --- | --- |
| C=72; O=4; E=0.35; R=11.56; rawP=0.0004; adjP=0.0026 | | | | | | |
| Index | UserID | Value | Gene Symbol | Gene Name | EntrezGene | Ensembl |
| 1 | P49748 | NA | ACADVL | acyl-CoA dehydrogenase, very long chain | 37 | ENSG00000072778 |
| 2 | P31040 | NA | SDHA | succinate dehydrogenase complex, subunit A, flavoprotein (Fp) | 6389 | ENSG00000073578 |
| 3 | O95831 | NA | AIFM1 | apoptosis-inducing factor, mitochondrion-associated, 1 | 9131 | ENSG00000156709 |
| 4 | P13804 | NA | ETFA | electron-transfer-flavoprotein, alpha polypeptide | 2108 | ENSG00000140374 |

  
  

| **Database:molecular function      &nbspName:protein binding      &nbspID:GO:0005515** | | | | | | |
| --- | --- | --- | --- | --- | --- | --- |
| C=7301; O=49; E=35.10; R=1.40; rawP=0.0007; adjP=0.0035 | | | | | | |
| Index | UserID | Value | Gene Symbol | Gene Name | EntrezGene | Ensembl |
| 1 | P17931 | NA | LGALS3 | lectin, galactoside-binding, soluble, 3 | 3958 | ENSG00000131981 |
| 2 | Q99623 | NA | PHB2 | prohibitin 2 | 11331 | ENSG00000215021 |
| 3 | P05787 | NA | KRT8 | keratin 8 | 3856 | ENSG00000170421 |
| 4 | P40926 | NA | MDH2 | malate dehydrogenase 2, NAD (mitochondrial) | 4191 | ENSG00000146701 |
| 5 | P35232 | NA | PHB | prohibitin | 5245 | ENSG00000167085 |
| 6 | P50453 | NA | SERPINB9 | serpin peptidase inhibitor, clade B (ovalbumin), member 9 | 5272 | ENSG00000170542 |
| 7 | O95831 | NA | AIFM1 | apoptosis-inducing factor, mitochondrion-associated, 1 | 9131 | ENSG00000156709 |
| 8 | Q99714 | NA | HSD17B10 | hydroxysteroid (17-beta) dehydrogenase 10 | 3028 | ENSG00000072506 |
| 9 | P04179 | NA | SOD2 | superoxide dismutase 2, mitochondrial | 6648 | ENSG00000112096 |
| 10 | P00367 | NA | GLUD1 | glutamate dehydrogenase 1 | 2746 | ENSG00000148672 |
| 11 | O60313 | NA | OPA1 | optic atrophy 1 (autosomal dominant) | 4976 | ENSG00000198836 |
| 12 | P35580 | NA | MYH10 | myosin, heavy chain 10, non-muscle | 4628 | ENSG00000133026 |
| 13 | P31040 | NA | SDHA | succinate dehydrogenase complex, subunit A, flavoprotein (Fp) | 6389 | ENSG00000073578 |
| 14 | P43490 | NA | NAMPT | nicotinamide phosphoribosyltransferase | 10135 | ENSG00000105835 |
| 15 | O14773 | NA | TPP1 | tripeptidyl peptidase I | 1200 | ENSG00000166340 |
| 16 | P40121 | NA | CAPG | capping protein (actin filament), gelsolin-like | 822 | ENSG00000042493 |
| 17 | P42765 | NA | ACAA2 | acetyl-CoA acyltransferase 2 | 10449 | ENSG00000167315 |
| 18 | O75521 | NA | ECI2 | enoyl-CoA delta isomerase 2 | 10455 | ENSG00000198721 |
| 19 | O75367 | NA | H2AFY | H2A histone family, member Y | 9555 | ENSG00000113648 |
| 20 | P49419 | NA | ALDH7A1 | aldehyde dehydrogenase 7 family, member A1 | 501 | ENSG00000164904 |
| 21 | Q04837 | NA | SSBP1 | single-stranded DNA binding protein 1, mitochondrial | 6742 | ENSG00000106028 |
| 22 | P08727 | NA | KRT19 | keratin 19 | 3880 | ENSG00000171345 |
| 23 | Q14764 | NA | MVP | major vault protein | 9961 | ENSG00000013364 |
| 24 | P50454 | NA | SERPINH1 | serpin peptidase inhibitor, clade H (heat shock protein 47), member 1, (collagen binding protein 1) | 871 | ENSG00000149257 |
| 25 | Q13813 | NA | SPTAN1 | spectrin, alpha, non-erythrocytic 1 | 6709 | ENSG00000197694 |
| 26 | P02792 | NA | FTL | ferritin, light polypeptide | 2512 | ENSG00000087086 |
| 27 | P21953 | NA | BCKDHB | branched chain keto acid dehydrogenase E1, beta polypeptide | 594 | ENSG00000083123 |
| 28 | P40939 | NA | HADHA | hydroxyacyl-CoA dehydrogenase/3-ketoacyl-CoA thiolase/enoyl-CoA hydratase (trifunctional protein), alpha subunit | 3030 | ENSG00000084754 |
| 29 | P30084 | NA | ECHS1 | enoyl CoA hydratase, short chain, 1, mitochondrial | 1892 | ENSG00000127884 |
| 30 | P52815 | NA | MRPL12 | mitochondrial ribosomal protein L12 | 6182 | ENSG00000262814 |
| 31 | P21796 | NA | VDAC1 | voltage-dependent anion channel 1 | 7416 | ENSG00000213585 |
| 32 | Q13011 | NA | ECH1 | enoyl CoA hydratase 1, peroxisomal | 1891 | ENSG00000104823 |
| 33 | P32322 | NA | PYCR1 | pyrroline-5-carboxylate reductase 1 | 5831 | ENSG00000183010 |
| 34 | Q9UJZ1 | NA | STOML2 | stomatin (EPB72)-like 2 | 30968 | ENSG00000165283 |
| 35 | P04040 | NA | CAT | catalase | 847 | ENSG00000121691 |
| 36 | Q14980 | NA | NUMA1 | nuclear mitotic apparatus protein 1 | 4926 | ENSG00000137497 |
| 37 | O75340 | NA | PDCD6 | programmed cell death 6 | 10016 | ENSG00000249915 |
| 38 | P51649 | NA | ALDH5A1 | aldehyde dehydrogenase 5 family, member A1 | 7915 | ENSG00000112294 |
| 39 | P12235 | NA | SLC25A4 | solute carrier family 25 (mitochondrial carrier; adenine nucleotide translocator), member 4 | 291 | ENSG00000151729 |
| 40 | P30048 | NA | PRDX3 | peroxiredoxin 3 | 10935 | ENSG00000165672 |
| 41 | Q6PI48 | NA | DARS2 | aspartyl-tRNA synthetase 2, mitochondrial | 55157 | ENSG00000117593 |
| 42 | P24752 | NA | ACAT1 | acetyl-CoA acetyltransferase 1 | 38 | ENSG00000075239 |
| 43 | P19367 | NA | HK1 | hexokinase 1 | 3098 | ENSG00000156515 |
| 44 | P25705 | NA | ATP5A1 | ATP synthase, H+ transporting, mitochondrial F1 complex, alpha subunit 1, cardiac muscle | 498 | ENSG00000152234 |
| 45 | Q9BQ69 | NA | MACROD1 | MACRO domain containing 1 | 28992 | ENSG00000133315 |
| 46 | P06865 | NA | HEXA | hexosaminidase A (alpha polypeptide) | 3073 | ENSG00000213614 |
| 47 | P05783 | NA | KRT18 | keratin 18 | 3875 | ENSG00000111057 |
| 48 | P48047 | NA | ATP5O | ATP synthase, H+ transporting, mitochondrial F1 complex, O subunit | 539 | ENSG00000241837 |
| 49 | Q16531 | NA | DDB1 | damage-specific DNA binding protein 1, 127kDa | 1642 | ENSG00000167986 |

  
  

| **Database:molecular function      &nbspName:hydrolase activity, hydrolyzing O-glycosyl compounds      &nbspID:GO:0004553** | | | | | | |
| --- | --- | --- | --- | --- | --- | --- |
| C=93; O=4; E=0.45; R=8.95; rawP=0.0010; adjP=0.0045 | | | | | | |
| Index | UserID | Value | Gene Symbol | Gene Name | EntrezGene | Ensembl |
| 1 | P10253 | NA | GAA | glucosidase, alpha; acid | 2548 | ENSG00000171298 |
| 2 | P06865 | NA | HEXA | hexosaminidase A (alpha polypeptide) | 3073 | ENSG00000213614 |
| 3 | Q14697 | NA | GANAB | glucosidase, alpha; neutral AB | 23193 | ENSG00000089597 |
| 4 | Q13724 | NA | MOGS | mannosyl-oligosaccharide glucosidase | 7841 | ENSG00000115275 |

  
  

| **Database:molecular function      &nbspName:hydro-lyase activity      &nbspID:GO:0016836** | | | | | | |
| --- | --- | --- | --- | --- | --- | --- |
| C=42; O=3; E=0.20; R=14.86; rawP=0.0011; adjP=0.0047 | | | | | | |
| Index | UserID | Value | Gene Symbol | Gene Name | EntrezGene | Ensembl |
| 1 | Q99798 | NA | ACO2 | aconitase 2, mitochondrial | 50 | ENSG00000100412 |
| 2 | P40939 | NA | HADHA | hydroxyacyl-CoA dehydrogenase/3-ketoacyl-CoA thiolase/enoyl-CoA hydratase (trifunctional protein), alpha subunit | 3030 | ENSG00000084754 |
| 3 | P30084 | NA | ECHS1 | enoyl CoA hydratase, short chain, 1, mitochondrial | 1892 | ENSG00000127884 |

  
  

| **Database:molecular function      &nbspName:identical protein binding      &nbspID:GO:0042802** | | | | | | |
| --- | --- | --- | --- | --- | --- | --- |
| C=860; O=11; E=4.13; R=2.66; rawP=0.0025; adjP=0.0097 | | | | | | |
| Index | UserID | Value | Gene Symbol | Gene Name | EntrezGene | Ensembl |
| 1 | P04040 | NA | CAT | catalase | 847 | ENSG00000121691 |
| 2 | P02792 | NA | FTL | ferritin, light polypeptide | 2512 | ENSG00000087086 |
| 3 | O75340 | NA | PDCD6 | programmed cell death 6 | 10016 | ENSG00000249915 |
| 4 | P51649 | NA | ALDH5A1 | aldehyde dehydrogenase 5 family, member A1 | 7915 | ENSG00000112294 |
| 5 | P30048 | NA | PRDX3 | peroxiredoxin 3 | 10935 | ENSG00000165672 |
| 6 | Q99714 | NA | HSD17B10 | hydroxysteroid (17-beta) dehydrogenase 10 | 3028 | ENSG00000072506 |
| 7 | Q6PI48 | NA | DARS2 | aspartyl-tRNA synthetase 2, mitochondrial | 55157 | ENSG00000117593 |
| 8 | P04179 | NA | SOD2 | superoxide dismutase 2, mitochondrial | 6648 | ENSG00000112096 |
| 9 | P24752 | NA | ACAT1 | acetyl-CoA acetyltransferase 1 | 38 | ENSG00000075239 |
| 10 | P00367 | NA | GLUD1 | glutamate dehydrogenase 1 | 2746 | ENSG00000148672 |
| 11 | P32322 | NA | PYCR1 | pyrroline-5-carboxylate reductase 1 | 5831 | ENSG00000183010 |

  
  

| **Database:molecular function      &nbspName:hydrolase activity, acting on glycosyl bonds      &nbspID:GO:0016798** | | | | | | |
| --- | --- | --- | --- | --- | --- | --- |
| C=117; O=4; E=0.56; R=7.11; rawP=0.0024; adjP=0.0097 | | | | | | |
| Index | UserID | Value | Gene Symbol | Gene Name | EntrezGene | Ensembl |
| 1 | P10253 | NA | GAA | glucosidase, alpha; acid | 2548 | ENSG00000171298 |
| 2 | P06865 | NA | HEXA | hexosaminidase A (alpha polypeptide) | 3073 | ENSG00000213614 |
| 3 | Q14697 | NA | GANAB | glucosidase, alpha; neutral AB | 23193 | ENSG00000089597 |
| 4 | Q13724 | NA | MOGS | mannosyl-oligosaccharide glucosidase | 7841 | ENSG00000115275 |

  
  

| **Database:molecular function      &nbspName:binding      &nbspID:GO:0005488** | | | | | | |
| --- | --- | --- | --- | --- | --- | --- |
| C=11779; O=66; E=56.63; R=1.17; rawP=0.0027; adjP=0.0100 | | | | | | |
| Index | UserID | Value | Gene Symbol | Gene Name | EntrezGene | Ensembl |
| 1 | P10253 | NA | GAA | glucosidase, alpha; acid | 2548 | ENSG00000171298 |
| 2 | P54819 | NA | AK2 | adenylate kinase 2 | 204 | ENSG00000004455 |
| 3 | P17931 | NA | LGALS3 | lectin, galactoside-binding, soluble, 3 | 3958 | ENSG00000131981 |
| 4 | Q99623 | NA | PHB2 | prohibitin 2 | 11331 | ENSG00000215021 |
| 5 | P05787 | NA | KRT8 | keratin 8 | 3856 | ENSG00000170421 |
| 6 | P40926 | NA | MDH2 | malate dehydrogenase 2, NAD (mitochondrial) | 4191 | ENSG00000146701 |
| 7 | P35232 | NA | PHB | prohibitin | 5245 | ENSG00000167085 |
| 8 | P50453 | NA | SERPINB9 | serpin peptidase inhibitor, clade B (ovalbumin), member 9 | 5272 | ENSG00000170542 |
| 9 | O95831 | NA | AIFM1 | apoptosis-inducing factor, mitochondrion-associated, 1 | 9131 | ENSG00000156709 |
| 10 | Q99714 | NA | HSD17B10 | hydroxysteroid (17-beta) dehydrogenase 10 | 3028 | ENSG00000072506 |
| 11 | P04179 | NA | SOD2 | superoxide dismutase 2, mitochondrial | 6648 | ENSG00000112096 |
| 12 | P07741 | NA | APRT | adenine phosphoribosyltransferase | 353 | ENSG00000198931 |
| 13 | P00367 | NA | GLUD1 | glutamate dehydrogenase 1 | 2746 | ENSG00000148672 |
| 14 | P48735 | NA | IDH2 | isocitrate dehydrogenase 2 (NADP+), mitochondrial | 3418 | ENSG00000182054 |
| 15 | O60313 | NA | OPA1 | optic atrophy 1 (autosomal dominant) | 4976 | ENSG00000198836 |
| 16 | P35270 | NA | SPR | sepiapterin reductase (7,8-dihydrobiopterin:NADP+ oxidoreductase) | 6697 | ENSG00000116096 |
| 17 | P35580 | NA | MYH10 | myosin, heavy chain 10, non-muscle | 4628 | ENSG00000133026 |
| 18 | P31040 | NA | SDHA | succinate dehydrogenase complex, subunit A, flavoprotein (Fp) | 6389 | ENSG00000073578 |
| 19 | P43490 | NA | NAMPT | nicotinamide phosphoribosyltransferase | 10135 | ENSG00000105835 |
| 20 | O14773 | NA | TPP1 | tripeptidyl peptidase I | 1200 | ENSG00000166340 |
| 21 | P13804 | NA | ETFA | electron-transfer-flavoprotein, alpha polypeptide | 2108 | ENSG00000140374 |
| 22 | P40121 | NA | CAPG | capping protein (actin filament), gelsolin-like | 822 | ENSG00000042493 |
| 23 | P42765 | NA | ACAA2 | acetyl-CoA acyltransferase 2 | 10449 | ENSG00000167315 |
| 24 | O75521 | NA | ECI2 | enoyl-CoA delta isomerase 2 | 10455 | ENSG00000198721 |
| 25 | O75367 | NA | H2AFY | H2A histone family, member Y | 9555 | ENSG00000113648 |
| 26 | Q9Y277 | NA | VDAC3 | voltage-dependent anion channel 3 | 7419 | ENSG00000078668 |
| 27 | P49419 | NA | ALDH7A1 | aldehyde dehydrogenase 7 family, member A1 | 501 | ENSG00000164904 |
| 28 | Q04837 | NA | SSBP1 | single-stranded DNA binding protein 1, mitochondrial | 6742 | ENSG00000106028 |
| 29 | P08727 | NA | KRT19 | keratin 19 | 3880 | ENSG00000171345 |
| 30 | Q14764 | NA | MVP | major vault protein | 9961 | ENSG00000013364 |
| 31 | P50454 | NA | SERPINH1 | serpin peptidase inhibitor, clade H (heat shock protein 47), member 1, (collagen binding protein 1) | 871 | ENSG00000149257 |
| 32 | Q14697 | NA | GANAB | glucosidase, alpha; neutral AB | 23193 | ENSG00000089597 |
| 33 | Q13813 | NA | SPTAN1 | spectrin, alpha, non-erythrocytic 1 | 6709 | ENSG00000197694 |
| 34 | Q5SSJ5 | NA | HP1BP3 | heterochromatin protein 1, binding protein 3 | 50809 | ENSG00000127483 |
| 35 | P02792 | NA | FTL | ferritin, light polypeptide | 2512 | ENSG00000087086 |
| 36 | O75439 | NA | PMPCB | peptidase (mitochondrial processing) beta | 9512 | ENSG00000105819 |
| 37 | P21953 | NA | BCKDHB | branched chain keto acid dehydrogenase E1, beta polypeptide | 594 | ENSG00000083123 |
| 38 | P40939 | NA | HADHA | hydroxyacyl-CoA dehydrogenase/3-ketoacyl-CoA thiolase/enoyl-CoA hydratase (trifunctional protein), alpha subunit | 3030 | ENSG00000084754 |
| 39 | P30084 | NA | ECHS1 | enoyl CoA hydratase, short chain, 1, mitochondrial | 1892 | ENSG00000127884 |
| 40 | P52815 | NA | MRPL12 | mitochondrial ribosomal protein L12 | 6182 | ENSG00000262814 |
| 41 | P21796 | NA | VDAC1 | voltage-dependent anion channel 1 | 7416 | ENSG00000213585 |
| 42 | Q13011 | NA | ECH1 | enoyl CoA hydratase 1, peroxisomal | 1891 | ENSG00000104823 |
| 43 | P32322 | NA | PYCR1 | pyrroline-5-carboxylate reductase 1 | 5831 | ENSG00000183010 |
| 44 | Q9UJZ1 | NA | STOML2 | stomatin (EPB72)-like 2 | 30968 | ENSG00000165283 |
| 45 | P04040 | NA | CAT | catalase | 847 | ENSG00000121691 |
| 46 | Q14980 | NA | NUMA1 | nuclear mitotic apparatus protein 1 | 4926 | ENSG00000137497 |
| 47 | Q99798 | NA | ACO2 | aconitase 2, mitochondrial | 50 | ENSG00000100412 |
| 48 | O75340 | NA | PDCD6 | programmed cell death 6 | 10016 | ENSG00000249915 |
| 49 | P51688 | NA | SGSH | N-sulfoglucosamine sulfohydrolase | 6448 | ENSG00000181523 |
| 50 | P51649 | NA | ALDH5A1 | aldehyde dehydrogenase 5 family, member A1 | 7915 | ENSG00000112294 |
| 51 | P12235 | NA | SLC25A4 | solute carrier family 25 (mitochondrial carrier; adenine nucleotide translocator), member 4 | 291 | ENSG00000151729 |
| 52 | P00558 | NA | PGK1 | phosphoglycerate kinase 1 | 5230 | ENSG00000102144 |
| 53 | P30048 | NA | PRDX3 | peroxiredoxin 3 | 10935 | ENSG00000165672 |
| 54 | Q9NSE4 | NA | IARS2 | isoleucyl-tRNA synthetase 2, mitochondrial | 55699 | ENSG00000067704 |
| 55 | Q6PI48 | NA | DARS2 | aspartyl-tRNA synthetase 2, mitochondrial | 55157 | ENSG00000117593 |
| 56 | P24752 | NA | ACAT1 | acetyl-CoA acetyltransferase 1 | 38 | ENSG00000075239 |
| 57 | P13674 | NA | P4HA1 | prolyl 4-hydroxylase, alpha polypeptide I | 5033 | ENSG00000122884 |
| 58 | P19367 | NA | HK1 | hexokinase 1 | 3098 | ENSG00000156515 |
| 59 | P25705 | NA | ATP5A1 | ATP synthase, H+ transporting, mitochondrial F1 complex, alpha subunit 1, cardiac muscle | 498 | ENSG00000152234 |
| 60 | Q9BQ69 | NA | MACROD1 | MACRO domain containing 1 | 28992 | ENSG00000133315 |
| 61 | P06865 | NA | HEXA | hexosaminidase A (alpha polypeptide) | 3073 | ENSG00000213614 |
| 62 | P49748 | NA | ACADVL | acyl-CoA dehydrogenase, very long chain | 37 | ENSG00000072778 |
| 63 | P05783 | NA | KRT18 | keratin 18 | 3875 | ENSG00000111057 |
| 64 | P48047 | NA | ATP5O | ATP synthase, H+ transporting, mitochondrial F1 complex, O subunit | 539 | ENSG00000241837 |
| 65 | Q16531 | NA | DDB1 | damage-specific DNA binding protein 1, 127kDa | 1642 | ENSG00000167986 |
| 66 | P31937 | NA | HIBADH | 3-hydroxyisobutyrate dehydrogenase | 11112 | ENSG00000106049 |

  
  

| **Database:molecular function      &nbspName:carbon-oxygen lyase activity      &nbspID:GO:0016835** | | | | | | |
| --- | --- | --- | --- | --- | --- | --- |
| C=61; O=3; E=0.29; R=10.23; rawP=0.0031; adjP=0.0105 | | | | | | |
| Index | UserID | Value | Gene Symbol | Gene Name | EntrezGene | Ensembl |
| 1 | Q99798 | NA | ACO2 | aconitase 2, mitochondrial | 50 | ENSG00000100412 |
| 2 | P40939 | NA | HADHA | hydroxyacyl-CoA dehydrogenase/3-ketoacyl-CoA thiolase/enoyl-CoA hydratase (trifunctional protein), alpha subunit | 3030 | ENSG00000084754 |
| 3 | P30084 | NA | ECHS1 | enoyl CoA hydratase, short chain, 1, mitochondrial | 1892 | ENSG00000127884 |

  
  

| **Database:molecular function      &nbspName:antioxidant activity      &nbspID:GO:0016209** | | | | | | |
| --- | --- | --- | --- | --- | --- | --- |
| C=61; O=3; E=0.29; R=10.23; rawP=0.0031; adjP=0.0105 | | | | | | |
| Index | UserID | Value | Gene Symbol | Gene Name | EntrezGene | Ensembl |
| 1 | P04040 | NA | CAT | catalase | 847 | ENSG00000121691 |
| 2 | P30048 | NA | PRDX3 | peroxiredoxin 3 | 10935 | ENSG00000165672 |
| 3 | P04179 | NA | SOD2 | superoxide dismutase 2, mitochondrial | 6648 | ENSG00000112096 |

  
  

| **Database:molecular function      &nbspName:lyase activity      &nbspID:GO:0016829** | | | | | | |
| --- | --- | --- | --- | --- | --- | --- |
| C=157; O=4; E=0.75; R=5.30; rawP=0.0069; adjP=0.0226 | | | | | | |
| Index | UserID | Value | Gene Symbol | Gene Name | EntrezGene | Ensembl |
| 1 | Q99798 | NA | ACO2 | aconitase 2, mitochondrial | 50 | ENSG00000100412 |
| 2 | P21953 | NA | BCKDHB | branched chain keto acid dehydrogenase E1, beta polypeptide | 594 | ENSG00000083123 |
| 3 | P40939 | NA | HADHA | hydroxyacyl-CoA dehydrogenase/3-ketoacyl-CoA thiolase/enoyl-CoA hydratase (trifunctional protein), alpha subunit | 3030 | ENSG00000084754 |
| 4 | P30084 | NA | ECHS1 | enoyl CoA hydratase, short chain, 1, mitochondrial | 1892 | ENSG00000127884 |

  
  

| **Database:molecular function      &nbspName:isomerase activity      &nbspID:GO:0016853** | | | | | | |
| --- | --- | --- | --- | --- | --- | --- |
| C=142; O=3; E=0.68; R=4.39; rawP=0.0310; adjP=0.0976 | | | | | | |
| Index | UserID | Value | Gene Symbol | Gene Name | EntrezGene | Ensembl |
| 1 | O75521 | NA | ECI2 | enoyl-CoA delta isomerase 2 | 10455 | ENSG00000198721 |
| 2 | Q13011 | NA | ECH1 | enoyl CoA hydratase 1, peroxisomal | 1891 | ENSG00000104823 |
| 3 | P60174 | NA | TPI1 | triosephosphate isomerase 1 | 7167 | ENSG00000111669 |

  
  

| **Database:cellular component      &nbspName:mitochondrial part      &nbspID:GO:0044429** | | | | | | |
| --- | --- | --- | --- | --- | --- | --- |
| C=740; O=40; E=3.37; R=11.86; rawP=3.73e-34; adjP=3.02e-32 | | | | | | |
| Index | UserID | Value | Gene Symbol | Gene Name | EntrezGene | Ensembl |
| 1 | P54819 | NA | AK2 | adenylate kinase 2 | 204 | ENSG00000004455 |
| 2 | P17931 | NA | LGALS3 | lectin, galactoside-binding, soluble, 3 | 3958 | ENSG00000131981 |
| 3 | Q99623 | NA | PHB2 | prohibitin 2 | 11331 | ENSG00000215021 |
| 4 | P40926 | NA | MDH2 | malate dehydrogenase 2, NAD (mitochondrial) | 4191 | ENSG00000146701 |
| 5 | P35232 | NA | PHB | prohibitin | 5245 | ENSG00000167085 |
| 6 | Q99714 | NA | HSD17B10 | hydroxysteroid (17-beta) dehydrogenase 10 | 3028 | ENSG00000072506 |
| 7 | O95831 | NA | AIFM1 | apoptosis-inducing factor, mitochondrion-associated, 1 | 9131 | ENSG00000156709 |
| 8 | P04179 | NA | SOD2 | superoxide dismutase 2, mitochondrial | 6648 | ENSG00000112096 |
| 9 | P00367 | NA | GLUD1 | glutamate dehydrogenase 1 | 2746 | ENSG00000148672 |
| 10 | P48735 | NA | IDH2 | isocitrate dehydrogenase 2 (NADP+), mitochondrial | 3418 | ENSG00000182054 |
| 11 | O60313 | NA | OPA1 | optic atrophy 1 (autosomal dominant) | 4976 | ENSG00000198836 |
| 12 | P31040 | NA | SDHA | succinate dehydrogenase complex, subunit A, flavoprotein (Fp) | 6389 | ENSG00000073578 |
| 13 | P38117 | NA | ETFB | electron-transfer-flavoprotein, beta polypeptide | 2109 | ENSG00000105379 |
| 14 | P13804 | NA | ETFA | electron-transfer-flavoprotein, alpha polypeptide | 2108 | ENSG00000140374 |
| 15 | P42765 | NA | ACAA2 | acetyl-CoA acyltransferase 2 | 10449 | ENSG00000167315 |
| 16 | Q9Y277 | NA | VDAC3 | voltage-dependent anion channel 3 | 7419 | ENSG00000078668 |
| 17 | P49419 | NA | ALDH7A1 | aldehyde dehydrogenase 7 family, member A1 | 501 | ENSG00000164904 |
| 18 | Q04837 | NA | SSBP1 | single-stranded DNA binding protein 1, mitochondrial | 6742 | ENSG00000106028 |
| 19 | Q9BPW8 | NA | NIPSNAP1 | nipsnap homolog 1 (C. elegans) | 8508 | ENSG00000184117 |
| 20 | O75439 | NA | PMPCB | peptidase (mitochondrial processing) beta | 9512 | ENSG00000105819 |
| 21 | P21953 | NA | BCKDHB | branched chain keto acid dehydrogenase E1, beta polypeptide | 594 | ENSG00000083123 |
| 22 | P40939 | NA | HADHA | hydroxyacyl-CoA dehydrogenase/3-ketoacyl-CoA thiolase/enoyl-CoA hydratase (trifunctional protein), alpha subunit | 3030 | ENSG00000084754 |
| 23 | P30084 | NA | ECHS1 | enoyl CoA hydratase, short chain, 1, mitochondrial | 1892 | ENSG00000127884 |
| 24 | P52815 | NA | MRPL12 | mitochondrial ribosomal protein L12 | 6182 | ENSG00000262814 |
| 25 | P21796 | NA | VDAC1 | voltage-dependent anion channel 1 | 7416 | ENSG00000213585 |
| 26 | P32322 | NA | PYCR1 | pyrroline-5-carboxylate reductase 1 | 5831 | ENSG00000183010 |
| 27 | Q9UJZ1 | NA | STOML2 | stomatin (EPB72)-like 2 | 30968 | ENSG00000165283 |
| 28 | P04040 | NA | CAT | catalase | 847 | ENSG00000121691 |
| 29 | Q99798 | NA | ACO2 | aconitase 2, mitochondrial | 50 | ENSG00000100412 |
| 30 | P51649 | NA | ALDH5A1 | aldehyde dehydrogenase 5 family, member A1 | 7915 | ENSG00000112294 |
| 31 | P12235 | NA | SLC25A4 | solute carrier family 25 (mitochondrial carrier; adenine nucleotide translocator), member 4 | 291 | ENSG00000151729 |
| 32 | Q9NSE4 | NA | IARS2 | isoleucyl-tRNA synthetase 2, mitochondrial | 55699 | ENSG00000067704 |
| 33 | Q6PI48 | NA | DARS2 | aspartyl-tRNA synthetase 2, mitochondrial | 55157 | ENSG00000117593 |
| 34 | P24752 | NA | ACAT1 | acetyl-CoA acetyltransferase 1 | 38 | ENSG00000075239 |
| 35 | Q02978 | NA | SLC25A11 | solute carrier family 25 (mitochondrial carrier; oxoglutarate carrier), member 11 | 8402 | ENSG00000108528 |
| 36 | P19367 | NA | HK1 | hexokinase 1 | 3098 | ENSG00000156515 |
| 37 | P25705 | NA | ATP5A1 | ATP synthase, H+ transporting, mitochondrial F1 complex, alpha subunit 1, cardiac muscle | 498 | ENSG00000152234 |
| 38 | P49748 | NA | ACADVL | acyl-CoA dehydrogenase, very long chain | 37 | ENSG00000072778 |
| 39 | P48047 | NA | ATP5O | ATP synthase, H+ transporting, mitochondrial F1 complex, O subunit | 539 | ENSG00000241837 |
| 40 | P31937 | NA | HIBADH | 3-hydroxyisobutyrate dehydrogenase | 11112 | ENSG00000106049 |

  
  

| **Database:cellular component      &nbspName:mitochondrion      &nbspID:GO:0005739** | | | | | | |
| --- | --- | --- | --- | --- | --- | --- |
| C=1515; O=48; E=6.90; R=6.95; rawP=1.96e-31; adjP=7.94e-30 | | | | | | |
| Index | UserID | Value | Gene Symbol | Gene Name | EntrezGene | Ensembl |
| 1 | P54819 | NA | AK2 | adenylate kinase 2 | 204 | ENSG00000004455 |
| 2 | P17931 | NA | LGALS3 | lectin, galactoside-binding, soluble, 3 | 3958 | ENSG00000131981 |
| 3 | Q99623 | NA | PHB2 | prohibitin 2 | 11331 | ENSG00000215021 |
| 4 | P40926 | NA | MDH2 | malate dehydrogenase 2, NAD (mitochondrial) | 4191 | ENSG00000146701 |
| 5 | P35232 | NA | PHB | prohibitin | 5245 | ENSG00000167085 |
| 6 | Q99714 | NA | HSD17B10 | hydroxysteroid (17-beta) dehydrogenase 10 | 3028 | ENSG00000072506 |
| 7 | O95831 | NA | AIFM1 | apoptosis-inducing factor, mitochondrion-associated, 1 | 9131 | ENSG00000156709 |
| 8 | P04179 | NA | SOD2 | superoxide dismutase 2, mitochondrial | 6648 | ENSG00000112096 |
| 9 | P00367 | NA | GLUD1 | glutamate dehydrogenase 1 | 2746 | ENSG00000148672 |
| 10 | P48735 | NA | IDH2 | isocitrate dehydrogenase 2 (NADP+), mitochondrial | 3418 | ENSG00000182054 |
| 11 | O60313 | NA | OPA1 | optic atrophy 1 (autosomal dominant) | 4976 | ENSG00000198836 |
| 12 | P31040 | NA | SDHA | succinate dehydrogenase complex, subunit A, flavoprotein (Fp) | 6389 | ENSG00000073578 |
| 13 | P38117 | NA | ETFB | electron-transfer-flavoprotein, beta polypeptide | 2109 | ENSG00000105379 |
| 14 | O14773 | NA | TPP1 | tripeptidyl peptidase I | 1200 | ENSG00000166340 |
| 15 | P13804 | NA | ETFA | electron-transfer-flavoprotein, alpha polypeptide | 2108 | ENSG00000140374 |
| 16 | P42765 | NA | ACAA2 | acetyl-CoA acyltransferase 2 | 10449 | ENSG00000167315 |
| 17 | O75521 | NA | ECI2 | enoyl-CoA delta isomerase 2 | 10455 | ENSG00000198721 |
| 18 | Q9Y277 | NA | VDAC3 | voltage-dependent anion channel 3 | 7419 | ENSG00000078668 |
| 19 | P49419 | NA | ALDH7A1 | aldehyde dehydrogenase 7 family, member A1 | 501 | ENSG00000164904 |
| 20 | Q04837 | NA | SSBP1 | single-stranded DNA binding protein 1, mitochondrial | 6742 | ENSG00000106028 |
| 21 | Q9BPW8 | NA | NIPSNAP1 | nipsnap homolog 1 (C. elegans) | 8508 | ENSG00000184117 |
| 22 | O75439 | NA | PMPCB | peptidase (mitochondrial processing) beta | 9512 | ENSG00000105819 |
| 23 | P21953 | NA | BCKDHB | branched chain keto acid dehydrogenase E1, beta polypeptide | 594 | ENSG00000083123 |
| 24 | P40939 | NA | HADHA | hydroxyacyl-CoA dehydrogenase/3-ketoacyl-CoA thiolase/enoyl-CoA hydratase (trifunctional protein), alpha subunit | 3030 | ENSG00000084754 |
| 25 | P30084 | NA | ECHS1 | enoyl CoA hydratase, short chain, 1, mitochondrial | 1892 | ENSG00000127884 |
| 26 | P52815 | NA | MRPL12 | mitochondrial ribosomal protein L12 | 6182 | ENSG00000262814 |
| 27 | Q9BXW7 | NA | CECR5 | cat eye syndrome chromosome region, candidate 5 | 27440 | ENSG00000069998 |
| 28 | P21796 | NA | VDAC1 | voltage-dependent anion channel 1 | 7416 | ENSG00000213585 |
| 29 | Q13011 | NA | ECH1 | enoyl CoA hydratase 1, peroxisomal | 1891 | ENSG00000104823 |
| 30 | P32322 | NA | PYCR1 | pyrroline-5-carboxylate reductase 1 | 5831 | ENSG00000183010 |
| 31 | Q9UJZ1 | NA | STOML2 | stomatin (EPB72)-like 2 | 30968 | ENSG00000165283 |
| 32 | P04040 | NA | CAT | catalase | 847 | ENSG00000121691 |
| 33 | Q99798 | NA | ACO2 | aconitase 2, mitochondrial | 50 | ENSG00000100412 |
| 34 | P51649 | NA | ALDH5A1 | aldehyde dehydrogenase 5 family, member A1 | 7915 | ENSG00000112294 |
| 35 | P12235 | NA | SLC25A4 | solute carrier family 25 (mitochondrial carrier; adenine nucleotide translocator), member 4 | 291 | ENSG00000151729 |
| 36 | P30048 | NA | PRDX3 | peroxiredoxin 3 | 10935 | ENSG00000165672 |
| 37 | Q9NSE4 | NA | IARS2 | isoleucyl-tRNA synthetase 2, mitochondrial | 55699 | ENSG00000067704 |
| 38 | Q6PI48 | NA | DARS2 | aspartyl-tRNA synthetase 2, mitochondrial | 55157 | ENSG00000117593 |
| 39 | P24752 | NA | ACAT1 | acetyl-CoA acetyltransferase 1 | 38 | ENSG00000075239 |
| 40 | P13674 | NA | P4HA1 | prolyl 4-hydroxylase, alpha polypeptide I | 5033 | ENSG00000122884 |
| 41 | Q02978 | NA | SLC25A11 | solute carrier family 25 (mitochondrial carrier; oxoglutarate carrier), member 11 | 8402 | ENSG00000108528 |
| 42 | P19367 | NA | HK1 | hexokinase 1 | 3098 | ENSG00000156515 |
| 43 | P25705 | NA | ATP5A1 | ATP synthase, H+ transporting, mitochondrial F1 complex, alpha subunit 1, cardiac muscle | 498 | ENSG00000152234 |
| 44 | Q9BQ69 | NA | MACROD1 | MACRO domain containing 1 | 28992 | ENSG00000133315 |
| 45 | P49748 | NA | ACADVL | acyl-CoA dehydrogenase, very long chain | 37 | ENSG00000072778 |
| 46 | P48047 | NA | ATP5O | ATP synthase, H+ transporting, mitochondrial F1 complex, O subunit | 539 | ENSG00000241837 |
| 47 | P31937 | NA | HIBADH | 3-hydroxyisobutyrate dehydrogenase | 11112 | ENSG00000106049 |
| 48 | P30042 | NA | C21orf33 | chromosome 21 open reading frame 33 | 8209 | ENSG00000160221 |

  
  

| **Database:cellular component      &nbspName:cytoplasmic part      &nbspID:GO:0044444** | | | | | | |
| --- | --- | --- | --- | --- | --- | --- |
| C=6728; O=71; E=30.65; R=2.32; rawP=1.20e-22; adjP=2.43e-21 | | | | | | |
| Index | UserID | Value | Gene Symbol | Gene Name | EntrezGene | Ensembl |
| 1 | P10253 | NA | GAA | glucosidase, alpha; acid | 2548 | ENSG00000171298 |
| 2 | Q9UL46 | NA | PSME2 | proteasome (prosome, macropain) activator subunit 2 (PA28 beta) | 5721 | ENSG00000100911 |
| 3 | P40926 | NA | MDH2 | malate dehydrogenase 2, NAD (mitochondrial) | 4191 | ENSG00000146701 |
| 4 | P04179 | NA | SOD2 | superoxide dismutase 2, mitochondrial | 6648 | ENSG00000112096 |
| 5 | Q13724 | NA | MOGS | mannosyl-oligosaccharide glucosidase | 7841 | ENSG00000115275 |
| 6 | P00367 | NA | GLUD1 | glutamate dehydrogenase 1 | 2746 | ENSG00000148672 |
| 7 | Q06323 | NA | PSME1 | proteasome (prosome, macropain) activator subunit 1 (PA28 alpha) | 5720 | ENSG00000092010 |
| 8 | P48735 | NA | IDH2 | isocitrate dehydrogenase 2 (NADP+), mitochondrial | 3418 | ENSG00000182054 |
| 9 | P35580 | NA | MYH10 | myosin, heavy chain 10, non-muscle | 4628 | ENSG00000133026 |
| 10 | P31040 | NA | SDHA | succinate dehydrogenase complex, subunit A, flavoprotein (Fp) | 6389 | ENSG00000073578 |
| 11 | O14773 | NA | TPP1 | tripeptidyl peptidase I | 1200 | ENSG00000166340 |
| 12 | P13804 | NA | ETFA | electron-transfer-flavoprotein, alpha polypeptide | 2108 | ENSG00000140374 |
| 13 | P40121 | NA | CAPG | capping protein (actin filament), gelsolin-like | 822 | ENSG00000042493 |
| 14 | P42765 | NA | ACAA2 | acetyl-CoA acyltransferase 2 | 10449 | ENSG00000167315 |
| 15 | Q9Y277 | NA | VDAC3 | voltage-dependent anion channel 3 | 7419 | ENSG00000078668 |
| 16 | Q04837 | NA | SSBP1 | single-stranded DNA binding protein 1, mitochondrial | 6742 | ENSG00000106028 |
| 17 | P08727 | NA | KRT19 | keratin 19 | 3880 | ENSG00000171345 |
| 18 | P50454 | NA | SERPINH1 | serpin peptidase inhibitor, clade H (heat shock protein 47), member 1, (collagen binding protein 1) | 871 | ENSG00000149257 |
| 19 | Q14697 | NA | GANAB | glucosidase, alpha; neutral AB | 23193 | ENSG00000089597 |
| 20 | P40939 | NA | HADHA | hydroxyacyl-CoA dehydrogenase/3-ketoacyl-CoA thiolase/enoyl-CoA hydratase (trifunctional protein), alpha subunit | 3030 | ENSG00000084754 |
| 21 | P52815 | NA | MRPL12 | mitochondrial ribosomal protein L12 | 6182 | ENSG00000262814 |
| 22 | Q9BXW7 | NA | CECR5 | cat eye syndrome chromosome region, candidate 5 | 27440 | ENSG00000069998 |
| 23 | P21796 | NA | VDAC1 | voltage-dependent anion channel 1 | 7416 | ENSG00000213585 |
| 24 | P38571 | NA | LIPA | lipase A, lysosomal acid, cholesterol esterase | 3988 | ENSG00000107798 |
| 25 | P32322 | NA | PYCR1 | pyrroline-5-carboxylate reductase 1 | 5831 | ENSG00000183010 |
| 26 | Q9UJZ1 | NA | STOML2 | stomatin (EPB72)-like 2 | 30968 | ENSG00000165283 |
| 27 | Q14980 | NA | NUMA1 | nuclear mitotic apparatus protein 1 | 4926 | ENSG00000137497 |
| 28 | Q99798 | NA | ACO2 | aconitase 2, mitochondrial | 50 | ENSG00000100412 |
| 29 | P51649 | NA | ALDH5A1 | aldehyde dehydrogenase 5 family, member A1 | 7915 | ENSG00000112294 |
| 30 | P12235 | NA | SLC25A4 | solute carrier family 25 (mitochondrial carrier; adenine nucleotide translocator), member 4 | 291 | ENSG00000151729 |
| 31 | Q9NSE4 | NA | IARS2 | isoleucyl-tRNA synthetase 2, mitochondrial | 55699 | ENSG00000067704 |
| 32 | Q6PI48 | NA | DARS2 | aspartyl-tRNA synthetase 2, mitochondrial | 55157 | ENSG00000117593 |
| 33 | P13674 | NA | P4HA1 | prolyl 4-hydroxylase, alpha polypeptide I | 5033 | ENSG00000122884 |
| 34 | Q02978 | NA | SLC25A11 | solute carrier family 25 (mitochondrial carrier; oxoglutarate carrier), member 11 | 8402 | ENSG00000108528 |
| 35 | P60174 | NA | TPI1 | triosephosphate isomerase 1 | 7167 | ENSG00000111669 |
| 36 | P30042 | NA | C21orf33 | chromosome 21 open reading frame 33 | 8209 | ENSG00000160221 |
| 37 | P54819 | NA | AK2 | adenylate kinase 2 | 204 | ENSG00000004455 |
| 38 | P17931 | NA | LGALS3 | lectin, galactoside-binding, soluble, 3 | 3958 | ENSG00000131981 |
| 39 | Q99623 | NA | PHB2 | prohibitin 2 | 11331 | ENSG00000215021 |
| 40 | P50453 | NA | SERPINB9 | serpin peptidase inhibitor, clade B (ovalbumin), member 9 | 5272 | ENSG00000170542 |
| 41 | P35232 | NA | PHB | prohibitin | 5245 | ENSG00000167085 |
| 42 | O95831 | NA | AIFM1 | apoptosis-inducing factor, mitochondrion-associated, 1 | 9131 | ENSG00000156709 |
| 43 | Q99714 | NA | HSD17B10 | hydroxysteroid (17-beta) dehydrogenase 10 | 3028 | ENSG00000072506 |
| 44 | P07741 | NA | APRT | adenine phosphoribosyltransferase | 353 | ENSG00000198931 |
| 45 | O60313 | NA | OPA1 | optic atrophy 1 (autosomal dominant) | 4976 | ENSG00000198836 |
| 46 | P35270 | NA | SPR | sepiapterin reductase (7,8-dihydrobiopterin:NADP+ oxidoreductase) | 6697 | ENSG00000116096 |
| 47 | P38117 | NA | ETFB | electron-transfer-flavoprotein, beta polypeptide | 2109 | ENSG00000105379 |
| 48 | P43490 | NA | NAMPT | nicotinamide phosphoribosyltransferase | 10135 | ENSG00000105835 |
| 49 | O75521 | NA | ECI2 | enoyl-CoA delta isomerase 2 | 10455 | ENSG00000198721 |
| 50 | P49419 | NA | ALDH7A1 | aldehyde dehydrogenase 7 family, member A1 | 501 | ENSG00000164904 |
| 51 | Q9BPW8 | NA | NIPSNAP1 | nipsnap homolog 1 (C. elegans) | 8508 | ENSG00000184117 |
| 52 | Q13813 | NA | SPTAN1 | spectrin, alpha, non-erythrocytic 1 | 6709 | ENSG00000197694 |
| 53 | O75439 | NA | PMPCB | peptidase (mitochondrial processing) beta | 9512 | ENSG00000105819 |
| 54 | P02792 | NA | FTL | ferritin, light polypeptide | 2512 | ENSG00000087086 |
| 55 | P21953 | NA | BCKDHB | branched chain keto acid dehydrogenase E1, beta polypeptide | 594 | ENSG00000083123 |
| 56 | P30084 | NA | ECHS1 | enoyl CoA hydratase, short chain, 1, mitochondrial | 1892 | ENSG00000127884 |
| 57 | Q13011 | NA | ECH1 | enoyl CoA hydratase 1, peroxisomal | 1891 | ENSG00000104823 |
| 58 | P04040 | NA | CAT | catalase | 847 | ENSG00000121691 |
| 59 | P51688 | NA | SGSH | N-sulfoglucosamine sulfohydrolase | 6448 | ENSG00000181523 |
| 60 | O75340 | NA | PDCD6 | programmed cell death 6 | 10016 | ENSG00000249915 |
| 61 | P30048 | NA | PRDX3 | peroxiredoxin 3 | 10935 | ENSG00000165672 |
| 62 | P00558 | NA | PGK1 | phosphoglycerate kinase 1 | 5230 | ENSG00000102144 |
| 63 | P24752 | NA | ACAT1 | acetyl-CoA acetyltransferase 1 | 38 | ENSG00000075239 |
| 64 | P25705 | NA | ATP5A1 | ATP synthase, H+ transporting, mitochondrial F1 complex, alpha subunit 1, cardiac muscle | 498 | ENSG00000152234 |
| 65 | P19367 | NA | HK1 | hexokinase 1 | 3098 | ENSG00000156515 |
| 66 | Q9BQ69 | NA | MACROD1 | MACRO domain containing 1 | 28992 | ENSG00000133315 |
| 67 | P05783 | NA | KRT18 | keratin 18 | 3875 | ENSG00000111057 |
| 68 | P06865 | NA | HEXA | hexosaminidase A (alpha polypeptide) | 3073 | ENSG00000213614 |
| 69 | P49748 | NA | ACADVL | acyl-CoA dehydrogenase, very long chain | 37 | ENSG00000072778 |
| 70 | P48047 | NA | ATP5O | ATP synthase, H+ transporting, mitochondrial F1 complex, O subunit | 539 | ENSG00000241837 |
| 71 | P31937 | NA | HIBADH | 3-hydroxyisobutyrate dehydrogenase | 11112 | ENSG00000106049 |

  
  

| **Database:cellular component      &nbspName:cytoplasm      &nbspID:GO:0005737** | | | | | | |
| --- | --- | --- | --- | --- | --- | --- |
| C=9051; O=74; E=41.23; R=1.79; rawP=1.21e-17; adjP=8.91e-17 | | | | | | |
| Index | UserID | Value | Gene Symbol | Gene Name | EntrezGene | Ensembl |
| 1 | P10253 | NA | GAA | glucosidase, alpha; acid | 2548 | ENSG00000171298 |
| 2 | Q9UL46 | NA | PSME2 | proteasome (prosome, macropain) activator subunit 2 (PA28 beta) | 5721 | ENSG00000100911 |
| 3 | P40926 | NA | MDH2 | malate dehydrogenase 2, NAD (mitochondrial) | 4191 | ENSG00000146701 |
| 4 | P04179 | NA | SOD2 | superoxide dismutase 2, mitochondrial | 6648 | ENSG00000112096 |
| 5 | Q13724 | NA | MOGS | mannosyl-oligosaccharide glucosidase | 7841 | ENSG00000115275 |
| 6 | P00367 | NA | GLUD1 | glutamate dehydrogenase 1 | 2746 | ENSG00000148672 |
| 7 | Q06323 | NA | PSME1 | proteasome (prosome, macropain) activator subunit 1 (PA28 alpha) | 5720 | ENSG00000092010 |
| 8 | P48735 | NA | IDH2 | isocitrate dehydrogenase 2 (NADP+), mitochondrial | 3418 | ENSG00000182054 |
| 9 | P35580 | NA | MYH10 | myosin, heavy chain 10, non-muscle | 4628 | ENSG00000133026 |
| 10 | P31040 | NA | SDHA | succinate dehydrogenase complex, subunit A, flavoprotein (Fp) | 6389 | ENSG00000073578 |
| 11 | O14773 | NA | TPP1 | tripeptidyl peptidase I | 1200 | ENSG00000166340 |
| 12 | P13804 | NA | ETFA | electron-transfer-flavoprotein, alpha polypeptide | 2108 | ENSG00000140374 |
| 13 | P40121 | NA | CAPG | capping protein (actin filament), gelsolin-like | 822 | ENSG00000042493 |
| 14 | P42765 | NA | ACAA2 | acetyl-CoA acyltransferase 2 | 10449 | ENSG00000167315 |
| 15 | Q9Y277 | NA | VDAC3 | voltage-dependent anion channel 3 | 7419 | ENSG00000078668 |
| 16 | Q04837 | NA | SSBP1 | single-stranded DNA binding protein 1, mitochondrial | 6742 | ENSG00000106028 |
| 17 | P08727 | NA | KRT19 | keratin 19 | 3880 | ENSG00000171345 |
| 18 | P50454 | NA | SERPINH1 | serpin peptidase inhibitor, clade H (heat shock protein 47), member 1, (collagen binding protein 1) | 871 | ENSG00000149257 |
| 19 | Q14697 | NA | GANAB | glucosidase, alpha; neutral AB | 23193 | ENSG00000089597 |
| 20 | P40939 | NA | HADHA | hydroxyacyl-CoA dehydrogenase/3-ketoacyl-CoA thiolase/enoyl-CoA hydratase (trifunctional protein), alpha subunit | 3030 | ENSG00000084754 |
| 21 | P52815 | NA | MRPL12 | mitochondrial ribosomal protein L12 | 6182 | ENSG00000262814 |
| 22 | Q9BXW7 | NA | CECR5 | cat eye syndrome chromosome region, candidate 5 | 27440 | ENSG00000069998 |
| 23 | P21796 | NA | VDAC1 | voltage-dependent anion channel 1 | 7416 | ENSG00000213585 |
| 24 | P38571 | NA | LIPA | lipase A, lysosomal acid, cholesterol esterase | 3988 | ENSG00000107798 |
| 25 | P32322 | NA | PYCR1 | pyrroline-5-carboxylate reductase 1 | 5831 | ENSG00000183010 |
| 26 | Q9UJZ1 | NA | STOML2 | stomatin (EPB72)-like 2 | 30968 | ENSG00000165283 |
| 27 | Q14980 | NA | NUMA1 | nuclear mitotic apparatus protein 1 | 4926 | ENSG00000137497 |
| 28 | Q99798 | NA | ACO2 | aconitase 2, mitochondrial | 50 | ENSG00000100412 |
| 29 | P51649 | NA | ALDH5A1 | aldehyde dehydrogenase 5 family, member A1 | 7915 | ENSG00000112294 |
| 30 | P12235 | NA | SLC25A4 | solute carrier family 25 (mitochondrial carrier; adenine nucleotide translocator), member 4 | 291 | ENSG00000151729 |
| 31 | Q9NSE4 | NA | IARS2 | isoleucyl-tRNA synthetase 2, mitochondrial | 55699 | ENSG00000067704 |
| 32 | Q6PI48 | NA | DARS2 | aspartyl-tRNA synthetase 2, mitochondrial | 55157 | ENSG00000117593 |
| 33 | P13674 | NA | P4HA1 | prolyl 4-hydroxylase, alpha polypeptide I | 5033 | ENSG00000122884 |
| 34 | Q02978 | NA | SLC25A11 | solute carrier family 25 (mitochondrial carrier; oxoglutarate carrier), member 11 | 8402 | ENSG00000108528 |
| 35 | P60174 | NA | TPI1 | triosephosphate isomerase 1 | 7167 | ENSG00000111669 |
| 36 | P30042 | NA | C21orf33 | chromosome 21 open reading frame 33 | 8209 | ENSG00000160221 |
| 37 | P54819 | NA | AK2 | adenylate kinase 2 | 204 | ENSG00000004455 |
| 38 | P17931 | NA | LGALS3 | lectin, galactoside-binding, soluble, 3 | 3958 | ENSG00000131981 |
| 39 | P05787 | NA | KRT8 | keratin 8 | 3856 | ENSG00000170421 |
| 40 | Q99623 | NA | PHB2 | prohibitin 2 | 11331 | ENSG00000215021 |
| 41 | P50453 | NA | SERPINB9 | serpin peptidase inhibitor, clade B (ovalbumin), member 9 | 5272 | ENSG00000170542 |
| 42 | P35232 | NA | PHB | prohibitin | 5245 | ENSG00000167085 |
| 43 | O95831 | NA | AIFM1 | apoptosis-inducing factor, mitochondrion-associated, 1 | 9131 | ENSG00000156709 |
| 44 | Q99714 | NA | HSD17B10 | hydroxysteroid (17-beta) dehydrogenase 10 | 3028 | ENSG00000072506 |
| 45 | P07741 | NA | APRT | adenine phosphoribosyltransferase | 353 | ENSG00000198931 |
| 46 | O60313 | NA | OPA1 | optic atrophy 1 (autosomal dominant) | 4976 | ENSG00000198836 |
| 47 | P35270 | NA | SPR | sepiapterin reductase (7,8-dihydrobiopterin:NADP+ oxidoreductase) | 6697 | ENSG00000116096 |
| 48 | P38117 | NA | ETFB | electron-transfer-flavoprotein, beta polypeptide | 2109 | ENSG00000105379 |
| 49 | P43490 | NA | NAMPT | nicotinamide phosphoribosyltransferase | 10135 | ENSG00000105835 |
| 50 | O75521 | NA | ECI2 | enoyl-CoA delta isomerase 2 | 10455 | ENSG00000198721 |
| 51 | P49419 | NA | ALDH7A1 | aldehyde dehydrogenase 7 family, member A1 | 501 | ENSG00000164904 |
| 52 | Q14764 | NA | MVP | major vault protein | 9961 | ENSG00000013364 |
| 53 | Q9BPW8 | NA | NIPSNAP1 | nipsnap homolog 1 (C. elegans) | 8508 | ENSG00000184117 |
| 54 | Q13813 | NA | SPTAN1 | spectrin, alpha, non-erythrocytic 1 | 6709 | ENSG00000197694 |
| 55 | O75439 | NA | PMPCB | peptidase (mitochondrial processing) beta | 9512 | ENSG00000105819 |
| 56 | P02792 | NA | FTL | ferritin, light polypeptide | 2512 | ENSG00000087086 |
| 57 | P21953 | NA | BCKDHB | branched chain keto acid dehydrogenase E1, beta polypeptide | 594 | ENSG00000083123 |
| 58 | P30084 | NA | ECHS1 | enoyl CoA hydratase, short chain, 1, mitochondrial | 1892 | ENSG00000127884 |
| 59 | Q13011 | NA | ECH1 | enoyl CoA hydratase 1, peroxisomal | 1891 | ENSG00000104823 |
| 60 | P04040 | NA | CAT | catalase | 847 | ENSG00000121691 |
| 61 | P51688 | NA | SGSH | N-sulfoglucosamine sulfohydrolase | 6448 | ENSG00000181523 |
| 62 | O75340 | NA | PDCD6 | programmed cell death 6 | 10016 | ENSG00000249915 |
| 63 | P30048 | NA | PRDX3 | peroxiredoxin 3 | 10935 | ENSG00000165672 |
| 64 | P00558 | NA | PGK1 | phosphoglycerate kinase 1 | 5230 | ENSG00000102144 |
| 65 | P24752 | NA | ACAT1 | acetyl-CoA acetyltransferase 1 | 38 | ENSG00000075239 |
| 66 | P19367 | NA | HK1 | hexokinase 1 | 3098 | ENSG00000156515 |
| 67 | P25705 | NA | ATP5A1 | ATP synthase, H+ transporting, mitochondrial F1 complex, alpha subunit 1, cardiac muscle | 498 | ENSG00000152234 |
| 68 | Q9BQ69 | NA | MACROD1 | MACRO domain containing 1 | 28992 | ENSG00000133315 |
| 69 | P05783 | NA | KRT18 | keratin 18 | 3875 | ENSG00000111057 |
| 70 | P06865 | NA | HEXA | hexosaminidase A (alpha polypeptide) | 3073 | ENSG00000213614 |
| 71 | P49748 | NA | ACADVL | acyl-CoA dehydrogenase, very long chain | 37 | ENSG00000072778 |
| 72 | P48047 | NA | ATP5O | ATP synthase, H+ transporting, mitochondrial F1 complex, O subunit | 539 | ENSG00000241837 |
| 73 | Q16531 | NA | DDB1 | damage-specific DNA binding protein 1, 127kDa | 1642 | ENSG00000167986 |
| 74 | P31937 | NA | HIBADH | 3-hydroxyisobutyrate dehydrogenase | 11112 | ENSG00000106049 |

  
  

| **Database:cellular component      &nbspName:intracellular organelle part      &nbspID:GO:0044446** | | | | | | |
| --- | --- | --- | --- | --- | --- | --- |
| C=6690; O=65; E=30.48; R=2.13; rawP=3.18e-16; adjP=2.15e-15 | | | | | | |
| Index | UserID | Value | Gene Symbol | Gene Name | EntrezGene | Ensembl |
| 1 | P10253 | NA | GAA | glucosidase, alpha; acid | 2548 | ENSG00000171298 |
| 2 | P54819 | NA | AK2 | adenylate kinase 2 | 204 | ENSG00000004455 |
| 3 | P17931 | NA | LGALS3 | lectin, galactoside-binding, soluble, 3 | 3958 | ENSG00000131981 |
| 4 | Q99623 | NA | PHB2 | prohibitin 2 | 11331 | ENSG00000215021 |
| 5 | P05787 | NA | KRT8 | keratin 8 | 3856 | ENSG00000170421 |
| 6 | Q9UL46 | NA | PSME2 | proteasome (prosome, macropain) activator subunit 2 (PA28 beta) | 5721 | ENSG00000100911 |
| 7 | P40926 | NA | MDH2 | malate dehydrogenase 2, NAD (mitochondrial) | 4191 | ENSG00000146701 |
| 8 | P35232 | NA | PHB | prohibitin | 5245 | ENSG00000167085 |
| 9 | Q99714 | NA | HSD17B10 | hydroxysteroid (17-beta) dehydrogenase 10 | 3028 | ENSG00000072506 |
| 10 | O95831 | NA | AIFM1 | apoptosis-inducing factor, mitochondrion-associated, 1 | 9131 | ENSG00000156709 |
| 11 | P04179 | NA | SOD2 | superoxide dismutase 2, mitochondrial | 6648 | ENSG00000112096 |
| 12 | Q13724 | NA | MOGS | mannosyl-oligosaccharide glucosidase | 7841 | ENSG00000115275 |
| 13 | P07741 | NA | APRT | adenine phosphoribosyltransferase | 353 | ENSG00000198931 |
| 14 | P00367 | NA | GLUD1 | glutamate dehydrogenase 1 | 2746 | ENSG00000148672 |
| 15 | Q06323 | NA | PSME1 | proteasome (prosome, macropain) activator subunit 1 (PA28 alpha) | 5720 | ENSG00000092010 |
| 16 | P48735 | NA | IDH2 | isocitrate dehydrogenase 2 (NADP+), mitochondrial | 3418 | ENSG00000182054 |
| 17 | O60313 | NA | OPA1 | optic atrophy 1 (autosomal dominant) | 4976 | ENSG00000198836 |
| 18 | P35270 | NA | SPR | sepiapterin reductase (7,8-dihydrobiopterin:NADP+ oxidoreductase) | 6697 | ENSG00000116096 |
| 19 | P35580 | NA | MYH10 | myosin, heavy chain 10, non-muscle | 4628 | ENSG00000133026 |
| 20 | P31040 | NA | SDHA | succinate dehydrogenase complex, subunit A, flavoprotein (Fp) | 6389 | ENSG00000073578 |
| 21 | P38117 | NA | ETFB | electron-transfer-flavoprotein, beta polypeptide | 2109 | ENSG00000105379 |
| 22 | O14773 | NA | TPP1 | tripeptidyl peptidase I | 1200 | ENSG00000166340 |
| 23 | P13804 | NA | ETFA | electron-transfer-flavoprotein, alpha polypeptide | 2108 | ENSG00000140374 |
| 24 | P40121 | NA | CAPG | capping protein (actin filament), gelsolin-like | 822 | ENSG00000042493 |
| 25 | P42765 | NA | ACAA2 | acetyl-CoA acyltransferase 2 | 10449 | ENSG00000167315 |
| 26 | O75521 | NA | ECI2 | enoyl-CoA delta isomerase 2 | 10455 | ENSG00000198721 |
| 27 | O75367 | NA | H2AFY | H2A histone family, member Y | 9555 | ENSG00000113648 |
| 28 | Q9Y277 | NA | VDAC3 | voltage-dependent anion channel 3 | 7419 | ENSG00000078668 |
| 29 | P49419 | NA | ALDH7A1 | aldehyde dehydrogenase 7 family, member A1 | 501 | ENSG00000164904 |
| 30 | Q04837 | NA | SSBP1 | single-stranded DNA binding protein 1, mitochondrial | 6742 | ENSG00000106028 |
| 31 | P08727 | NA | KRT19 | keratin 19 | 3880 | ENSG00000171345 |
| 32 | Q14764 | NA | MVP | major vault protein | 9961 | ENSG00000013364 |
| 33 | P50454 | NA | SERPINH1 | serpin peptidase inhibitor, clade H (heat shock protein 47), member 1, (collagen binding protein 1) | 871 | ENSG00000149257 |
| 34 | Q14697 | NA | GANAB | glucosidase, alpha; neutral AB | 23193 | ENSG00000089597 |
| 35 | Q13813 | NA | SPTAN1 | spectrin, alpha, non-erythrocytic 1 | 6709 | ENSG00000197694 |
| 36 | Q5SSJ5 | NA | HP1BP3 | heterochromatin protein 1, binding protein 3 | 50809 | ENSG00000127483 |
| 37 | Q9BPW8 | NA | NIPSNAP1 | nipsnap homolog 1 (C. elegans) | 8508 | ENSG00000184117 |
| 38 | O75439 | NA | PMPCB | peptidase (mitochondrial processing) beta | 9512 | ENSG00000105819 |
| 39 | P21953 | NA | BCKDHB | branched chain keto acid dehydrogenase E1, beta polypeptide | 594 | ENSG00000083123 |
| 40 | P40939 | NA | HADHA | hydroxyacyl-CoA dehydrogenase/3-ketoacyl-CoA thiolase/enoyl-CoA hydratase (trifunctional protein), alpha subunit | 3030 | ENSG00000084754 |
| 41 | P30084 | NA | ECHS1 | enoyl CoA hydratase, short chain, 1, mitochondrial | 1892 | ENSG00000127884 |
| 42 | P52815 | NA | MRPL12 | mitochondrial ribosomal protein L12 | 6182 | ENSG00000262814 |
| 43 | P21796 | NA | VDAC1 | voltage-dependent anion channel 1 | 7416 | ENSG00000213585 |
| 44 | P32322 | NA | PYCR1 | pyrroline-5-carboxylate reductase 1 | 5831 | ENSG00000183010 |
| 45 | Q9UJZ1 | NA | STOML2 | stomatin (EPB72)-like 2 | 30968 | ENSG00000165283 |
| 46 | P04040 | NA | CAT | catalase | 847 | ENSG00000121691 |
| 47 | Q14980 | NA | NUMA1 | nuclear mitotic apparatus protein 1 | 4926 | ENSG00000137497 |
| 48 | Q99798 | NA | ACO2 | aconitase 2, mitochondrial | 50 | ENSG00000100412 |
| 49 | O75340 | NA | PDCD6 | programmed cell death 6 | 10016 | ENSG00000249915 |
| 50 | P51688 | NA | SGSH | N-sulfoglucosamine sulfohydrolase | 6448 | ENSG00000181523 |
| 51 | P51649 | NA | ALDH5A1 | aldehyde dehydrogenase 5 family, member A1 | 7915 | ENSG00000112294 |
| 52 | P12235 | NA | SLC25A4 | solute carrier family 25 (mitochondrial carrier; adenine nucleotide translocator), member 4 | 291 | ENSG00000151729 |
| 53 | Q9NSE4 | NA | IARS2 | isoleucyl-tRNA synthetase 2, mitochondrial | 55699 | ENSG00000067704 |
| 54 | Q6PI48 | NA | DARS2 | aspartyl-tRNA synthetase 2, mitochondrial | 55157 | ENSG00000117593 |
| 55 | P24752 | NA | ACAT1 | acetyl-CoA acetyltransferase 1 | 38 | ENSG00000075239 |
| 56 | P13674 | NA | P4HA1 | prolyl 4-hydroxylase, alpha polypeptide I | 5033 | ENSG00000122884 |
| 57 | Q02978 | NA | SLC25A11 | solute carrier family 25 (mitochondrial carrier; oxoglutarate carrier), member 11 | 8402 | ENSG00000108528 |
| 58 | P19367 | NA | HK1 | hexokinase 1 | 3098 | ENSG00000156515 |
| 59 | P25705 | NA | ATP5A1 | ATP synthase, H+ transporting, mitochondrial F1 complex, alpha subunit 1, cardiac muscle | 498 | ENSG00000152234 |
| 60 | P05783 | NA | KRT18 | keratin 18 | 3875 | ENSG00000111057 |
| 61 | P06865 | NA | HEXA | hexosaminidase A (alpha polypeptide) | 3073 | ENSG00000213614 |
| 62 | P49748 | NA | ACADVL | acyl-CoA dehydrogenase, very long chain | 37 | ENSG00000072778 |
| 63 | P48047 | NA | ATP5O | ATP synthase, H+ transporting, mitochondrial F1 complex, O subunit | 539 | ENSG00000241837 |
| 64 | Q16531 | NA | DDB1 | damage-specific DNA binding protein 1, 127kDa | 1642 | ENSG00000167986 |
| 65 | P31937 | NA | HIBADH | 3-hydroxyisobutyrate dehydrogenase | 11112 | ENSG00000106049 |

  
  

| **Database:cellular component      &nbspName:organelle part      &nbspID:GO:0044422** | | | | | | |
| --- | --- | --- | --- | --- | --- | --- |
| C=6777; O=65; E=30.87; R=2.11; rawP=6.74e-16; adjP=4.20e-15 | | | | | | |
| Index | UserID | Value | Gene Symbol | Gene Name | EntrezGene | Ensembl |
| 1 | P10253 | NA | GAA | glucosidase, alpha; acid | 2548 | ENSG00000171298 |
| 2 | P54819 | NA | AK2 | adenylate kinase 2 | 204 | ENSG00000004455 |
| 3 | P17931 | NA | LGALS3 | lectin, galactoside-binding, soluble, 3 | 3958 | ENSG00000131981 |
| 4 | Q99623 | NA | PHB2 | prohibitin 2 | 11331 | ENSG00000215021 |
| 5 | P05787 | NA | KRT8 | keratin 8 | 3856 | ENSG00000170421 |
| 6 | Q9UL46 | NA | PSME2 | proteasome (prosome, macropain) activator subunit 2 (PA28 beta) | 5721 | ENSG00000100911 |
| 7 | P40926 | NA | MDH2 | malate dehydrogenase 2, NAD (mitochondrial) | 4191 | ENSG00000146701 |
| 8 | P35232 | NA | PHB | prohibitin | 5245 | ENSG00000167085 |
| 9 | Q99714 | NA | HSD17B10 | hydroxysteroid (17-beta) dehydrogenase 10 | 3028 | ENSG00000072506 |
| 10 | O95831 | NA | AIFM1 | apoptosis-inducing factor, mitochondrion-associated, 1 | 9131 | ENSG00000156709 |
| 11 | P04179 | NA | SOD2 | superoxide dismutase 2, mitochondrial | 6648 | ENSG00000112096 |
| 12 | Q13724 | NA | MOGS | mannosyl-oligosaccharide glucosidase | 7841 | ENSG00000115275 |
| 13 | P07741 | NA | APRT | adenine phosphoribosyltransferase | 353 | ENSG00000198931 |
| 14 | P00367 | NA | GLUD1 | glutamate dehydrogenase 1 | 2746 | ENSG00000148672 |
| 15 | Q06323 | NA | PSME1 | proteasome (prosome, macropain) activator subunit 1 (PA28 alpha) | 5720 | ENSG00000092010 |
| 16 | P48735 | NA | IDH2 | isocitrate dehydrogenase 2 (NADP+), mitochondrial | 3418 | ENSG00000182054 |
| 17 | O60313 | NA | OPA1 | optic atrophy 1 (autosomal dominant) | 4976 | ENSG00000198836 |
| 18 | P35270 | NA | SPR | sepiapterin reductase (7,8-dihydrobiopterin:NADP+ oxidoreductase) | 6697 | ENSG00000116096 |
| 19 | P35580 | NA | MYH10 | myosin, heavy chain 10, non-muscle | 4628 | ENSG00000133026 |
| 20 | P31040 | NA | SDHA | succinate dehydrogenase complex, subunit A, flavoprotein (Fp) | 6389 | ENSG00000073578 |
| 21 | P38117 | NA | ETFB | electron-transfer-flavoprotein, beta polypeptide | 2109 | ENSG00000105379 |
| 22 | O14773 | NA | TPP1 | tripeptidyl peptidase I | 1200 | ENSG00000166340 |
| 23 | P13804 | NA | ETFA | electron-transfer-flavoprotein, alpha polypeptide | 2108 | ENSG00000140374 |
| 24 | P40121 | NA | CAPG | capping protein (actin filament), gelsolin-like | 822 | ENSG00000042493 |
| 25 | P42765 | NA | ACAA2 | acetyl-CoA acyltransferase 2 | 10449 | ENSG00000167315 |
| 26 | O75521 | NA | ECI2 | enoyl-CoA delta isomerase 2 | 10455 | ENSG00000198721 |
| 27 | O75367 | NA | H2AFY | H2A histone family, member Y | 9555 | ENSG00000113648 |
| 28 | Q9Y277 | NA | VDAC3 | voltage-dependent anion channel 3 | 7419 | ENSG00000078668 |
| 29 | P49419 | NA | ALDH7A1 | aldehyde dehydrogenase 7 family, member A1 | 501 | ENSG00000164904 |
| 30 | Q04837 | NA | SSBP1 | single-stranded DNA binding protein 1, mitochondrial | 6742 | ENSG00000106028 |
| 31 | P08727 | NA | KRT19 | keratin 19 | 3880 | ENSG00000171345 |
| 32 | Q14764 | NA | MVP | major vault protein | 9961 | ENSG00000013364 |
| 33 | P50454 | NA | SERPINH1 | serpin peptidase inhibitor, clade H (heat shock protein 47), member 1, (collagen binding protein 1) | 871 | ENSG00000149257 |
| 34 | Q14697 | NA | GANAB | glucosidase, alpha; neutral AB | 23193 | ENSG00000089597 |
| 35 | Q13813 | NA | SPTAN1 | spectrin, alpha, non-erythrocytic 1 | 6709 | ENSG00000197694 |
| 36 | Q5SSJ5 | NA | HP1BP3 | heterochromatin protein 1, binding protein 3 | 50809 | ENSG00000127483 |
| 37 | Q9BPW8 | NA | NIPSNAP1 | nipsnap homolog 1 (C. elegans) | 8508 | ENSG00000184117 |
| 38 | O75439 | NA | PMPCB | peptidase (mitochondrial processing) beta | 9512 | ENSG00000105819 |
| 39 | P21953 | NA | BCKDHB | branched chain keto acid dehydrogenase E1, beta polypeptide | 594 | ENSG00000083123 |
| 40 | P40939 | NA | HADHA | hydroxyacyl-CoA dehydrogenase/3-ketoacyl-CoA thiolase/enoyl-CoA hydratase (trifunctional protein), alpha subunit | 3030 | ENSG00000084754 |
| 41 | P30084 | NA | ECHS1 | enoyl CoA hydratase, short chain, 1, mitochondrial | 1892 | ENSG00000127884 |
| 42 | P52815 | NA | MRPL12 | mitochondrial ribosomal protein L12 | 6182 | ENSG00000262814 |
| 43 | P21796 | NA | VDAC1 | voltage-dependent anion channel 1 | 7416 | ENSG00000213585 |
| 44 | P32322 | NA | PYCR1 | pyrroline-5-carboxylate reductase 1 | 5831 | ENSG00000183010 |
| 45 | Q9UJZ1 | NA | STOML2 | stomatin (EPB72)-like 2 | 30968 | ENSG00000165283 |
| 46 | P04040 | NA | CAT | catalase | 847 | ENSG00000121691 |
| 47 | Q14980 | NA | NUMA1 | nuclear mitotic apparatus protein 1 | 4926 | ENSG00000137497 |
| 48 | Q99798 | NA | ACO2 | aconitase 2, mitochondrial | 50 | ENSG00000100412 |
| 49 | O75340 | NA | PDCD6 | programmed cell death 6 | 10016 | ENSG00000249915 |
| 50 | P51688 | NA | SGSH | N-sulfoglucosamine sulfohydrolase | 6448 | ENSG00000181523 |
| 51 | P51649 | NA | ALDH5A1 | aldehyde dehydrogenase 5 family, member A1 | 7915 | ENSG00000112294 |
| 52 | P12235 | NA | SLC25A4 | solute carrier family 25 (mitochondrial carrier; adenine nucleotide translocator), member 4 | 291 | ENSG00000151729 |
| 53 | Q9NSE4 | NA | IARS2 | isoleucyl-tRNA synthetase 2, mitochondrial | 55699 | ENSG00000067704 |
| 54 | Q6PI48 | NA | DARS2 | aspartyl-tRNA synthetase 2, mitochondrial | 55157 | ENSG00000117593 |
| 55 | P24752 | NA | ACAT1 | acetyl-CoA acetyltransferase 1 | 38 | ENSG00000075239 |
| 56 | P13674 | NA | P4HA1 | prolyl 4-hydroxylase, alpha polypeptide I | 5033 | ENSG00000122884 |
| 57 | Q02978 | NA | SLC25A11 | solute carrier family 25 (mitochondrial carrier; oxoglutarate carrier), member 11 | 8402 | ENSG00000108528 |
| 58 | P19367 | NA | HK1 | hexokinase 1 | 3098 | ENSG00000156515 |
| 59 | P25705 | NA | ATP5A1 | ATP synthase, H+ transporting, mitochondrial F1 complex, alpha subunit 1, cardiac muscle | 498 | ENSG00000152234 |
| 60 | P05783 | NA | KRT18 | keratin 18 | 3875 | ENSG00000111057 |
| 61 | P06865 | NA | HEXA | hexosaminidase A (alpha polypeptide) | 3073 | ENSG00000213614 |
| 62 | P49748 | NA | ACADVL | acyl-CoA dehydrogenase, very long chain | 37 | ENSG00000072778 |
| 63 | P48047 | NA | ATP5O | ATP synthase, H+ transporting, mitochondrial F1 complex, O subunit | 539 | ENSG00000241837 |
| 64 | Q16531 | NA | DDB1 | damage-specific DNA binding protein 1, 127kDa | 1642 | ENSG00000167986 |
| 65 | P31937 | NA | HIBADH | 3-hydroxyisobutyrate dehydrogenase | 11112 | ENSG00000106049 |

  
  

| **Database:cellular component      &nbspName:intracellular membrane-bounded organelle      &nbspID:GO:0043231** | | | | | | |
| --- | --- | --- | --- | --- | --- | --- |
| C=9484; O=71; E=43.21; R=1.64; rawP=1.07e-12; adjP=5.10e-12 | | | | | | |
| Index | UserID | Value | Gene Symbol | Gene Name | EntrezGene | Ensembl |
| 1 | P10253 | NA | GAA | glucosidase, alpha; acid | 2548 | ENSG00000171298 |
| 2 | Q9UL46 | NA | PSME2 | proteasome (prosome, macropain) activator subunit 2 (PA28 beta) | 5721 | ENSG00000100911 |
| 3 | P40926 | NA | MDH2 | malate dehydrogenase 2, NAD (mitochondrial) | 4191 | ENSG00000146701 |
| 4 | P04179 | NA | SOD2 | superoxide dismutase 2, mitochondrial | 6648 | ENSG00000112096 |
| 5 | Q13724 | NA | MOGS | mannosyl-oligosaccharide glucosidase | 7841 | ENSG00000115275 |
| 6 | P00367 | NA | GLUD1 | glutamate dehydrogenase 1 | 2746 | ENSG00000148672 |
| 7 | Q06323 | NA | PSME1 | proteasome (prosome, macropain) activator subunit 1 (PA28 alpha) | 5720 | ENSG00000092010 |
| 8 | P48735 | NA | IDH2 | isocitrate dehydrogenase 2 (NADP+), mitochondrial | 3418 | ENSG00000182054 |
| 9 | P31040 | NA | SDHA | succinate dehydrogenase complex, subunit A, flavoprotein (Fp) | 6389 | ENSG00000073578 |
| 10 | O14773 | NA | TPP1 | tripeptidyl peptidase I | 1200 | ENSG00000166340 |
| 11 | P13804 | NA | ETFA | electron-transfer-flavoprotein, alpha polypeptide | 2108 | ENSG00000140374 |
| 12 | P40121 | NA | CAPG | capping protein (actin filament), gelsolin-like | 822 | ENSG00000042493 |
| 13 | P42765 | NA | ACAA2 | acetyl-CoA acyltransferase 2 | 10449 | ENSG00000167315 |
| 14 | Q9Y277 | NA | VDAC3 | voltage-dependent anion channel 3 | 7419 | ENSG00000078668 |
| 15 | Q04837 | NA | SSBP1 | single-stranded DNA binding protein 1, mitochondrial | 6742 | ENSG00000106028 |
| 16 | P50454 | NA | SERPINH1 | serpin peptidase inhibitor, clade H (heat shock protein 47), member 1, (collagen binding protein 1) | 871 | ENSG00000149257 |
| 17 | Q14697 | NA | GANAB | glucosidase, alpha; neutral AB | 23193 | ENSG00000089597 |
| 18 | P40939 | NA | HADHA | hydroxyacyl-CoA dehydrogenase/3-ketoacyl-CoA thiolase/enoyl-CoA hydratase (trifunctional protein), alpha subunit | 3030 | ENSG00000084754 |
| 19 | P52815 | NA | MRPL12 | mitochondrial ribosomal protein L12 | 6182 | ENSG00000262814 |
| 20 | Q9BXW7 | NA | CECR5 | cat eye syndrome chromosome region, candidate 5 | 27440 | ENSG00000069998 |
| 21 | P21796 | NA | VDAC1 | voltage-dependent anion channel 1 | 7416 | ENSG00000213585 |
| 22 | P38571 | NA | LIPA | lipase A, lysosomal acid, cholesterol esterase | 3988 | ENSG00000107798 |
| 23 | P32322 | NA | PYCR1 | pyrroline-5-carboxylate reductase 1 | 5831 | ENSG00000183010 |
| 24 | Q9UJZ1 | NA | STOML2 | stomatin (EPB72)-like 2 | 30968 | ENSG00000165283 |
| 25 | Q14980 | NA | NUMA1 | nuclear mitotic apparatus protein 1 | 4926 | ENSG00000137497 |
| 26 | Q99798 | NA | ACO2 | aconitase 2, mitochondrial | 50 | ENSG00000100412 |
| 27 | P51649 | NA | ALDH5A1 | aldehyde dehydrogenase 5 family, member A1 | 7915 | ENSG00000112294 |
| 28 | P12235 | NA | SLC25A4 | solute carrier family 25 (mitochondrial carrier; adenine nucleotide translocator), member 4 | 291 | ENSG00000151729 |
| 29 | Q9NSE4 | NA | IARS2 | isoleucyl-tRNA synthetase 2, mitochondrial | 55699 | ENSG00000067704 |
| 30 | Q6PI48 | NA | DARS2 | aspartyl-tRNA synthetase 2, mitochondrial | 55157 | ENSG00000117593 |
| 31 | P13674 | NA | P4HA1 | prolyl 4-hydroxylase, alpha polypeptide I | 5033 | ENSG00000122884 |
| 32 | Q02978 | NA | SLC25A11 | solute carrier family 25 (mitochondrial carrier; oxoglutarate carrier), member 11 | 8402 | ENSG00000108528 |
| 33 | P60174 | NA | TPI1 | triosephosphate isomerase 1 | 7167 | ENSG00000111669 |
| 34 | P30042 | NA | C21orf33 | chromosome 21 open reading frame 33 | 8209 | ENSG00000160221 |
| 35 | P54819 | NA | AK2 | adenylate kinase 2 | 204 | ENSG00000004455 |
| 36 | P17931 | NA | LGALS3 | lectin, galactoside-binding, soluble, 3 | 3958 | ENSG00000131981 |
| 37 | P05787 | NA | KRT8 | keratin 8 | 3856 | ENSG00000170421 |
| 38 | Q99623 | NA | PHB2 | prohibitin 2 | 11331 | ENSG00000215021 |
| 39 | P50453 | NA | SERPINB9 | serpin peptidase inhibitor, clade B (ovalbumin), member 9 | 5272 | ENSG00000170542 |
| 40 | P35232 | NA | PHB | prohibitin | 5245 | ENSG00000167085 |
| 41 | O95831 | NA | AIFM1 | apoptosis-inducing factor, mitochondrion-associated, 1 | 9131 | ENSG00000156709 |
| 42 | Q99714 | NA | HSD17B10 | hydroxysteroid (17-beta) dehydrogenase 10 | 3028 | ENSG00000072506 |
| 43 | P07741 | NA | APRT | adenine phosphoribosyltransferase | 353 | ENSG00000198931 |
| 44 | O60313 | NA | OPA1 | optic atrophy 1 (autosomal dominant) | 4976 | ENSG00000198836 |
| 45 | P35270 | NA | SPR | sepiapterin reductase (7,8-dihydrobiopterin:NADP+ oxidoreductase) | 6697 | ENSG00000116096 |
| 46 | P38117 | NA | ETFB | electron-transfer-flavoprotein, beta polypeptide | 2109 | ENSG00000105379 |
| 47 | O75367 | NA | H2AFY | H2A histone family, member Y | 9555 | ENSG00000113648 |
| 48 | O75521 | NA | ECI2 | enoyl-CoA delta isomerase 2 | 10455 | ENSG00000198721 |
| 49 | P49419 | NA | ALDH7A1 | aldehyde dehydrogenase 7 family, member A1 | 501 | ENSG00000164904 |
| 50 | Q14764 | NA | MVP | major vault protein | 9961 | ENSG00000013364 |
| 51 | Q9BPW8 | NA | NIPSNAP1 | nipsnap homolog 1 (C. elegans) | 8508 | ENSG00000184117 |
| 52 | Q5SSJ5 | NA | HP1BP3 | heterochromatin protein 1, binding protein 3 | 50809 | ENSG00000127483 |
| 53 | Q13813 | NA | SPTAN1 | spectrin, alpha, non-erythrocytic 1 | 6709 | ENSG00000197694 |
| 54 | O75439 | NA | PMPCB | peptidase (mitochondrial processing) beta | 9512 | ENSG00000105819 |
| 55 | P21953 | NA | BCKDHB | branched chain keto acid dehydrogenase E1, beta polypeptide | 594 | ENSG00000083123 |
| 56 | P30084 | NA | ECHS1 | enoyl CoA hydratase, short chain, 1, mitochondrial | 1892 | ENSG00000127884 |
| 57 | Q13011 | NA | ECH1 | enoyl CoA hydratase 1, peroxisomal | 1891 | ENSG00000104823 |
| 58 | P04040 | NA | CAT | catalase | 847 | ENSG00000121691 |
| 59 | P51688 | NA | SGSH | N-sulfoglucosamine sulfohydrolase | 6448 | ENSG00000181523 |
| 60 | O75340 | NA | PDCD6 | programmed cell death 6 | 10016 | ENSG00000249915 |
| 61 | P30048 | NA | PRDX3 | peroxiredoxin 3 | 10935 | ENSG00000165672 |
| 62 | P24752 | NA | ACAT1 | acetyl-CoA acetyltransferase 1 | 38 | ENSG00000075239 |
| 63 | P25705 | NA | ATP5A1 | ATP synthase, H+ transporting, mitochondrial F1 complex, alpha subunit 1, cardiac muscle | 498 | ENSG00000152234 |
| 64 | P19367 | NA | HK1 | hexokinase 1 | 3098 | ENSG00000156515 |
| 65 | Q9BQ69 | NA | MACROD1 | MACRO domain containing 1 | 28992 | ENSG00000133315 |
| 66 | P05783 | NA | KRT18 | keratin 18 | 3875 | ENSG00000111057 |
| 67 | P06865 | NA | HEXA | hexosaminidase A (alpha polypeptide) | 3073 | ENSG00000213614 |
| 68 | P49748 | NA | ACADVL | acyl-CoA dehydrogenase, very long chain | 37 | ENSG00000072778 |
| 69 | P48047 | NA | ATP5O | ATP synthase, H+ transporting, mitochondrial F1 complex, O subunit | 539 | ENSG00000241837 |
| 70 | Q16531 | NA | DDB1 | damage-specific DNA binding protein 1, 127kDa | 1642 | ENSG00000167986 |
| 71 | P31937 | NA | HIBADH | 3-hydroxyisobutyrate dehydrogenase | 11112 | ENSG00000106049 |

  
  

| **Database:cellular component      &nbspName:membrane-bounded organelle      &nbspID:GO:0043227** | | | | | | |
| --- | --- | --- | --- | --- | --- | --- |
| C=9495; O=71; E=43.26; R=1.64; rawP=1.15e-12; adjP=5.18e-12 | | | | | | |
| Index | UserID | Value | Gene Symbol | Gene Name | EntrezGene | Ensembl |
| 1 | P10253 | NA | GAA | glucosidase, alpha; acid | 2548 | ENSG00000171298 |
| 2 | Q9UL46 | NA | PSME2 | proteasome (prosome, macropain) activator subunit 2 (PA28 beta) | 5721 | ENSG00000100911 |
| 3 | P40926 | NA | MDH2 | malate dehydrogenase 2, NAD (mitochondrial) | 4191 | ENSG00000146701 |
| 4 | P04179 | NA | SOD2 | superoxide dismutase 2, mitochondrial | 6648 | ENSG00000112096 |
| 5 | Q13724 | NA | MOGS | mannosyl-oligosaccharide glucosidase | 7841 | ENSG00000115275 |
| 6 | P00367 | NA | GLUD1 | glutamate dehydrogenase 1 | 2746 | ENSG00000148672 |
| 7 | Q06323 | NA | PSME1 | proteasome (prosome, macropain) activator subunit 1 (PA28 alpha) | 5720 | ENSG00000092010 |
| 8 | P48735 | NA | IDH2 | isocitrate dehydrogenase 2 (NADP+), mitochondrial | 3418 | ENSG00000182054 |
| 9 | P31040 | NA | SDHA | succinate dehydrogenase complex, subunit A, flavoprotein (Fp) | 6389 | ENSG00000073578 |
| 10 | O14773 | NA | TPP1 | tripeptidyl peptidase I | 1200 | ENSG00000166340 |
| 11 | P13804 | NA | ETFA | electron-transfer-flavoprotein, alpha polypeptide | 2108 | ENSG00000140374 |
| 12 | P40121 | NA | CAPG | capping protein (actin filament), gelsolin-like | 822 | ENSG00000042493 |
| 13 | P42765 | NA | ACAA2 | acetyl-CoA acyltransferase 2 | 10449 | ENSG00000167315 |
| 14 | Q9Y277 | NA | VDAC3 | voltage-dependent anion channel 3 | 7419 | ENSG00000078668 |
| 15 | Q04837 | NA | SSBP1 | single-stranded DNA binding protein 1, mitochondrial | 6742 | ENSG00000106028 |
| 16 | P50454 | NA | SERPINH1 | serpin peptidase inhibitor, clade H (heat shock protein 47), member 1, (collagen binding protein 1) | 871 | ENSG00000149257 |
| 17 | Q14697 | NA | GANAB | glucosidase, alpha; neutral AB | 23193 | ENSG00000089597 |
| 18 | P40939 | NA | HADHA | hydroxyacyl-CoA dehydrogenase/3-ketoacyl-CoA thiolase/enoyl-CoA hydratase (trifunctional protein), alpha subunit | 3030 | ENSG00000084754 |
| 19 | P52815 | NA | MRPL12 | mitochondrial ribosomal protein L12 | 6182 | ENSG00000262814 |
| 20 | Q9BXW7 | NA | CECR5 | cat eye syndrome chromosome region, candidate 5 | 27440 | ENSG00000069998 |
| 21 | P21796 | NA | VDAC1 | voltage-dependent anion channel 1 | 7416 | ENSG00000213585 |
| 22 | P38571 | NA | LIPA | lipase A, lysosomal acid, cholesterol esterase | 3988 | ENSG00000107798 |
| 23 | P32322 | NA | PYCR1 | pyrroline-5-carboxylate reductase 1 | 5831 | ENSG00000183010 |
| 24 | Q9UJZ1 | NA | STOML2 | stomatin (EPB72)-like 2 | 30968 | ENSG00000165283 |
| 25 | Q14980 | NA | NUMA1 | nuclear mitotic apparatus protein 1 | 4926 | ENSG00000137497 |
| 26 | Q99798 | NA | ACO2 | aconitase 2, mitochondrial | 50 | ENSG00000100412 |
| 27 | P51649 | NA | ALDH5A1 | aldehyde dehydrogenase 5 family, member A1 | 7915 | ENSG00000112294 |
| 28 | P12235 | NA | SLC25A4 | solute carrier family 25 (mitochondrial carrier; adenine nucleotide translocator), member 4 | 291 | ENSG00000151729 |
| 29 | Q9NSE4 | NA | IARS2 | isoleucyl-tRNA synthetase 2, mitochondrial | 55699 | ENSG00000067704 |
| 30 | Q6PI48 | NA | DARS2 | aspartyl-tRNA synthetase 2, mitochondrial | 55157 | ENSG00000117593 |
| 31 | P13674 | NA | P4HA1 | prolyl 4-hydroxylase, alpha polypeptide I | 5033 | ENSG00000122884 |
| 32 | Q02978 | NA | SLC25A11 | solute carrier family 25 (mitochondrial carrier; oxoglutarate carrier), member 11 | 8402 | ENSG00000108528 |
| 33 | P60174 | NA | TPI1 | triosephosphate isomerase 1 | 7167 | ENSG00000111669 |
| 34 | P30042 | NA | C21orf33 | chromosome 21 open reading frame 33 | 8209 | ENSG00000160221 |
| 35 | P54819 | NA | AK2 | adenylate kinase 2 | 204 | ENSG00000004455 |
| 36 | P17931 | NA | LGALS3 | lectin, galactoside-binding, soluble, 3 | 3958 | ENSG00000131981 |
| 37 | P05787 | NA | KRT8 | keratin 8 | 3856 | ENSG00000170421 |
| 38 | Q99623 | NA | PHB2 | prohibitin 2 | 11331 | ENSG00000215021 |
| 39 | P50453 | NA | SERPINB9 | serpin peptidase inhibitor, clade B (ovalbumin), member 9 | 5272 | ENSG00000170542 |
| 40 | P35232 | NA | PHB | prohibitin | 5245 | ENSG00000167085 |
| 41 | O95831 | NA | AIFM1 | apoptosis-inducing factor, mitochondrion-associated, 1 | 9131 | ENSG00000156709 |
| 42 | Q99714 | NA | HSD17B10 | hydroxysteroid (17-beta) dehydrogenase 10 | 3028 | ENSG00000072506 |
| 43 | P07741 | NA | APRT | adenine phosphoribosyltransferase | 353 | ENSG00000198931 |
| 44 | O60313 | NA | OPA1 | optic atrophy 1 (autosomal dominant) | 4976 | ENSG00000198836 |
| 45 | P35270 | NA | SPR | sepiapterin reductase (7,8-dihydrobiopterin:NADP+ oxidoreductase) | 6697 | ENSG00000116096 |
| 46 | P38117 | NA | ETFB | electron-transfer-flavoprotein, beta polypeptide | 2109 | ENSG00000105379 |
| 47 | O75367 | NA | H2AFY | H2A histone family, member Y | 9555 | ENSG00000113648 |
| 48 | O75521 | NA | ECI2 | enoyl-CoA delta isomerase 2 | 10455 | ENSG00000198721 |
| 49 | P49419 | NA | ALDH7A1 | aldehyde dehydrogenase 7 family, member A1 | 501 | ENSG00000164904 |
| 50 | Q14764 | NA | MVP | major vault protein | 9961 | ENSG00000013364 |
| 51 | Q9BPW8 | NA | NIPSNAP1 | nipsnap homolog 1 (C. elegans) | 8508 | ENSG00000184117 |
| 52 | Q5SSJ5 | NA | HP1BP3 | heterochromatin protein 1, binding protein 3 | 50809 | ENSG00000127483 |
| 53 | Q13813 | NA | SPTAN1 | spectrin, alpha, non-erythrocytic 1 | 6709 | ENSG00000197694 |
| 54 | O75439 | NA | PMPCB | peptidase (mitochondrial processing) beta | 9512 | ENSG00000105819 |
| 55 | P21953 | NA | BCKDHB | branched chain keto acid dehydrogenase E1, beta polypeptide | 594 | ENSG00000083123 |
| 56 | P30084 | NA | ECHS1 | enoyl CoA hydratase, short chain, 1, mitochondrial | 1892 | ENSG00000127884 |
| 57 | Q13011 | NA | ECH1 | enoyl CoA hydratase 1, peroxisomal | 1891 | ENSG00000104823 |
| 58 | P04040 | NA | CAT | catalase | 847 | ENSG00000121691 |
| 59 | P51688 | NA | SGSH | N-sulfoglucosamine sulfohydrolase | 6448 | ENSG00000181523 |
| 60 | O75340 | NA | PDCD6 | programmed cell death 6 | 10016 | ENSG00000249915 |
| 61 | P30048 | NA | PRDX3 | peroxiredoxin 3 | 10935 | ENSG00000165672 |
| 62 | P24752 | NA | ACAT1 | acetyl-CoA acetyltransferase 1 | 38 | ENSG00000075239 |
| 63 | P25705 | NA | ATP5A1 | ATP synthase, H+ transporting, mitochondrial F1 complex, alpha subunit 1, cardiac muscle | 498 | ENSG00000152234 |
| 64 | P19367 | NA | HK1 | hexokinase 1 | 3098 | ENSG00000156515 |
| 65 | Q9BQ69 | NA | MACROD1 | MACRO domain containing 1 | 28992 | ENSG00000133315 |
| 66 | P05783 | NA | KRT18 | keratin 18 | 3875 | ENSG00000111057 |
| 67 | P06865 | NA | HEXA | hexosaminidase A (alpha polypeptide) | 3073 | ENSG00000213614 |
| 68 | P49748 | NA | ACADVL | acyl-CoA dehydrogenase, very long chain | 37 | ENSG00000072778 |
| 69 | P48047 | NA | ATP5O | ATP synthase, H+ transporting, mitochondrial F1 complex, O subunit | 539 | ENSG00000241837 |
| 70 | Q16531 | NA | DDB1 | damage-specific DNA binding protein 1, 127kDa | 1642 | ENSG00000167986 |
| 71 | P31937 | NA | HIBADH | 3-hydroxyisobutyrate dehydrogenase | 11112 | ENSG00000106049 |

  
  

| **Database:cellular component      &nbspName:intracellular organelle      &nbspID:GO:0043229** | | | | | | |
| --- | --- | --- | --- | --- | --- | --- |
| C=10521; O=73; E=47.93; R=1.52; rawP=8.53e-12; adjP=3.64e-11 | | | | | | |
| Index | UserID | Value | Gene Symbol | Gene Name | EntrezGene | Ensembl |
| 1 | P10253 | NA | GAA | glucosidase, alpha; acid | 2548 | ENSG00000171298 |
| 2 | Q9UL46 | NA | PSME2 | proteasome (prosome, macropain) activator subunit 2 (PA28 beta) | 5721 | ENSG00000100911 |
| 3 | P40926 | NA | MDH2 | malate dehydrogenase 2, NAD (mitochondrial) | 4191 | ENSG00000146701 |
| 4 | P04179 | NA | SOD2 | superoxide dismutase 2, mitochondrial | 6648 | ENSG00000112096 |
| 5 | Q13724 | NA | MOGS | mannosyl-oligosaccharide glucosidase | 7841 | ENSG00000115275 |
| 6 | P00367 | NA | GLUD1 | glutamate dehydrogenase 1 | 2746 | ENSG00000148672 |
| 7 | Q06323 | NA | PSME1 | proteasome (prosome, macropain) activator subunit 1 (PA28 alpha) | 5720 | ENSG00000092010 |
| 8 | P48735 | NA | IDH2 | isocitrate dehydrogenase 2 (NADP+), mitochondrial | 3418 | ENSG00000182054 |
| 9 | P35580 | NA | MYH10 | myosin, heavy chain 10, non-muscle | 4628 | ENSG00000133026 |
| 10 | P31040 | NA | SDHA | succinate dehydrogenase complex, subunit A, flavoprotein (Fp) | 6389 | ENSG00000073578 |
| 11 | O14773 | NA | TPP1 | tripeptidyl peptidase I | 1200 | ENSG00000166340 |
| 12 | P13804 | NA | ETFA | electron-transfer-flavoprotein, alpha polypeptide | 2108 | ENSG00000140374 |
| 13 | P40121 | NA | CAPG | capping protein (actin filament), gelsolin-like | 822 | ENSG00000042493 |
| 14 | P42765 | NA | ACAA2 | acetyl-CoA acyltransferase 2 | 10449 | ENSG00000167315 |
| 15 | Q9Y277 | NA | VDAC3 | voltage-dependent anion channel 3 | 7419 | ENSG00000078668 |
| 16 | Q04837 | NA | SSBP1 | single-stranded DNA binding protein 1, mitochondrial | 6742 | ENSG00000106028 |
| 17 | P08727 | NA | KRT19 | keratin 19 | 3880 | ENSG00000171345 |
| 18 | P50454 | NA | SERPINH1 | serpin peptidase inhibitor, clade H (heat shock protein 47), member 1, (collagen binding protein 1) | 871 | ENSG00000149257 |
| 19 | Q14697 | NA | GANAB | glucosidase, alpha; neutral AB | 23193 | ENSG00000089597 |
| 20 | P40939 | NA | HADHA | hydroxyacyl-CoA dehydrogenase/3-ketoacyl-CoA thiolase/enoyl-CoA hydratase (trifunctional protein), alpha subunit | 3030 | ENSG00000084754 |
| 21 | P52815 | NA | MRPL12 | mitochondrial ribosomal protein L12 | 6182 | ENSG00000262814 |
| 22 | Q9BXW7 | NA | CECR5 | cat eye syndrome chromosome region, candidate 5 | 27440 | ENSG00000069998 |
| 23 | P21796 | NA | VDAC1 | voltage-dependent anion channel 1 | 7416 | ENSG00000213585 |
| 24 | P38571 | NA | LIPA | lipase A, lysosomal acid, cholesterol esterase | 3988 | ENSG00000107798 |
| 25 | P32322 | NA | PYCR1 | pyrroline-5-carboxylate reductase 1 | 5831 | ENSG00000183010 |
| 26 | Q9UJZ1 | NA | STOML2 | stomatin (EPB72)-like 2 | 30968 | ENSG00000165283 |
| 27 | Q14980 | NA | NUMA1 | nuclear mitotic apparatus protein 1 | 4926 | ENSG00000137497 |
| 28 | Q99798 | NA | ACO2 | aconitase 2, mitochondrial | 50 | ENSG00000100412 |
| 29 | P51649 | NA | ALDH5A1 | aldehyde dehydrogenase 5 family, member A1 | 7915 | ENSG00000112294 |
| 30 | P12235 | NA | SLC25A4 | solute carrier family 25 (mitochondrial carrier; adenine nucleotide translocator), member 4 | 291 | ENSG00000151729 |
| 31 | Q9NSE4 | NA | IARS2 | isoleucyl-tRNA synthetase 2, mitochondrial | 55699 | ENSG00000067704 |
| 32 | Q6PI48 | NA | DARS2 | aspartyl-tRNA synthetase 2, mitochondrial | 55157 | ENSG00000117593 |
| 33 | P13674 | NA | P4HA1 | prolyl 4-hydroxylase, alpha polypeptide I | 5033 | ENSG00000122884 |
| 34 | Q02978 | NA | SLC25A11 | solute carrier family 25 (mitochondrial carrier; oxoglutarate carrier), member 11 | 8402 | ENSG00000108528 |
| 35 | P60174 | NA | TPI1 | triosephosphate isomerase 1 | 7167 | ENSG00000111669 |
| 36 | P30042 | NA | C21orf33 | chromosome 21 open reading frame 33 | 8209 | ENSG00000160221 |
| 37 | P54819 | NA | AK2 | adenylate kinase 2 | 204 | ENSG00000004455 |
| 38 | P17931 | NA | LGALS3 | lectin, galactoside-binding, soluble, 3 | 3958 | ENSG00000131981 |
| 39 | P05787 | NA | KRT8 | keratin 8 | 3856 | ENSG00000170421 |
| 40 | Q99623 | NA | PHB2 | prohibitin 2 | 11331 | ENSG00000215021 |
| 41 | P50453 | NA | SERPINB9 | serpin peptidase inhibitor, clade B (ovalbumin), member 9 | 5272 | ENSG00000170542 |
| 42 | P35232 | NA | PHB | prohibitin | 5245 | ENSG00000167085 |
| 43 | O95831 | NA | AIFM1 | apoptosis-inducing factor, mitochondrion-associated, 1 | 9131 | ENSG00000156709 |
| 44 | Q99714 | NA | HSD17B10 | hydroxysteroid (17-beta) dehydrogenase 10 | 3028 | ENSG00000072506 |
| 45 | P07741 | NA | APRT | adenine phosphoribosyltransferase | 353 | ENSG00000198931 |
| 46 | O60313 | NA | OPA1 | optic atrophy 1 (autosomal dominant) | 4976 | ENSG00000198836 |
| 47 | P35270 | NA | SPR | sepiapterin reductase (7,8-dihydrobiopterin:NADP+ oxidoreductase) | 6697 | ENSG00000116096 |
| 48 | P38117 | NA | ETFB | electron-transfer-flavoprotein, beta polypeptide | 2109 | ENSG00000105379 |
| 49 | O75367 | NA | H2AFY | H2A histone family, member Y | 9555 | ENSG00000113648 |
| 50 | O75521 | NA | ECI2 | enoyl-CoA delta isomerase 2 | 10455 | ENSG00000198721 |
| 51 | P49419 | NA | ALDH7A1 | aldehyde dehydrogenase 7 family, member A1 | 501 | ENSG00000164904 |
| 52 | Q14764 | NA | MVP | major vault protein | 9961 | ENSG00000013364 |
| 53 | Q9BPW8 | NA | NIPSNAP1 | nipsnap homolog 1 (C. elegans) | 8508 | ENSG00000184117 |
| 54 | Q5SSJ5 | NA | HP1BP3 | heterochromatin protein 1, binding protein 3 | 50809 | ENSG00000127483 |
| 55 | Q13813 | NA | SPTAN1 | spectrin, alpha, non-erythrocytic 1 | 6709 | ENSG00000197694 |
| 56 | O75439 | NA | PMPCB | peptidase (mitochondrial processing) beta | 9512 | ENSG00000105819 |
| 57 | P21953 | NA | BCKDHB | branched chain keto acid dehydrogenase E1, beta polypeptide | 594 | ENSG00000083123 |
| 58 | P30084 | NA | ECHS1 | enoyl CoA hydratase, short chain, 1, mitochondrial | 1892 | ENSG00000127884 |
| 59 | Q13011 | NA | ECH1 | enoyl CoA hydratase 1, peroxisomal | 1891 | ENSG00000104823 |
| 60 | P04040 | NA | CAT | catalase | 847 | ENSG00000121691 |
| 61 | P51688 | NA | SGSH | N-sulfoglucosamine sulfohydrolase | 6448 | ENSG00000181523 |
| 62 | O75340 | NA | PDCD6 | programmed cell death 6 | 10016 | ENSG00000249915 |
| 63 | P30048 | NA | PRDX3 | peroxiredoxin 3 | 10935 | ENSG00000165672 |
| 64 | P24752 | NA | ACAT1 | acetyl-CoA acetyltransferase 1 | 38 | ENSG00000075239 |
| 65 | P19367 | NA | HK1 | hexokinase 1 | 3098 | ENSG00000156515 |
| 66 | P25705 | NA | ATP5A1 | ATP synthase, H+ transporting, mitochondrial F1 complex, alpha subunit 1, cardiac muscle | 498 | ENSG00000152234 |
| 67 | Q9BQ69 | NA | MACROD1 | MACRO domain containing 1 | 28992 | ENSG00000133315 |
| 68 | P05783 | NA | KRT18 | keratin 18 | 3875 | ENSG00000111057 |
| 69 | P06865 | NA | HEXA | hexosaminidase A (alpha polypeptide) | 3073 | ENSG00000213614 |
| 70 | P49748 | NA | ACADVL | acyl-CoA dehydrogenase, very long chain | 37 | ENSG00000072778 |
| 71 | P48047 | NA | ATP5O | ATP synthase, H+ transporting, mitochondrial F1 complex, O subunit | 539 | ENSG00000241837 |
| 72 | Q16531 | NA | DDB1 | damage-specific DNA binding protein 1, 127kDa | 1642 | ENSG00000167986 |
| 73 | P31937 | NA | HIBADH | 3-hydroxyisobutyrate dehydrogenase | 11112 | ENSG00000106049 |

  
  

| **Database:cellular component      &nbspName:organelle      &nbspID:GO:0043226** | | | | | | |
| --- | --- | --- | --- | --- | --- | --- |
| C=10536; O=73; E=48.00; R=1.52; rawP=9.40e-12; adjP=3.81e-11 | | | | | | |
| Index | UserID | Value | Gene Symbol | Gene Name | EntrezGene | Ensembl |
| 1 | P10253 | NA | GAA | glucosidase, alpha; acid | 2548 | ENSG00000171298 |
| 2 | Q9UL46 | NA | PSME2 | proteasome (prosome, macropain) activator subunit 2 (PA28 beta) | 5721 | ENSG00000100911 |
| 3 | P40926 | NA | MDH2 | malate dehydrogenase 2, NAD (mitochondrial) | 4191 | ENSG00000146701 |
| 4 | P04179 | NA | SOD2 | superoxide dismutase 2, mitochondrial | 6648 | ENSG00000112096 |
| 5 | Q13724 | NA | MOGS | mannosyl-oligosaccharide glucosidase | 7841 | ENSG00000115275 |
| 6 | P00367 | NA | GLUD1 | glutamate dehydrogenase 1 | 2746 | ENSG00000148672 |
| 7 | Q06323 | NA | PSME1 | proteasome (prosome, macropain) activator subunit 1 (PA28 alpha) | 5720 | ENSG00000092010 |
| 8 | P48735 | NA | IDH2 | isocitrate dehydrogenase 2 (NADP+), mitochondrial | 3418 | ENSG00000182054 |
| 9 | P35580 | NA | MYH10 | myosin, heavy chain 10, non-muscle | 4628 | ENSG00000133026 |
| 10 | P31040 | NA | SDHA | succinate dehydrogenase complex, subunit A, flavoprotein (Fp) | 6389 | ENSG00000073578 |
| 11 | O14773 | NA | TPP1 | tripeptidyl peptidase I | 1200 | ENSG00000166340 |
| 12 | P13804 | NA | ETFA | electron-transfer-flavoprotein, alpha polypeptide | 2108 | ENSG00000140374 |
| 13 | P40121 | NA | CAPG | capping protein (actin filament), gelsolin-like | 822 | ENSG00000042493 |
| 14 | P42765 | NA | ACAA2 | acetyl-CoA acyltransferase 2 | 10449 | ENSG00000167315 |
| 15 | Q9Y277 | NA | VDAC3 | voltage-dependent anion channel 3 | 7419 | ENSG00000078668 |
| 16 | Q04837 | NA | SSBP1 | single-stranded DNA binding protein 1, mitochondrial | 6742 | ENSG00000106028 |
| 17 | P08727 | NA | KRT19 | keratin 19 | 3880 | ENSG00000171345 |
| 18 | P50454 | NA | SERPINH1 | serpin peptidase inhibitor, clade H (heat shock protein 47), member 1, (collagen binding protein 1) | 871 | ENSG00000149257 |
| 19 | Q14697 | NA | GANAB | glucosidase, alpha; neutral AB | 23193 | ENSG00000089597 |
| 20 | P40939 | NA | HADHA | hydroxyacyl-CoA dehydrogenase/3-ketoacyl-CoA thiolase/enoyl-CoA hydratase (trifunctional protein), alpha subunit | 3030 | ENSG00000084754 |
| 21 | P52815 | NA | MRPL12 | mitochondrial ribosomal protein L12 | 6182 | ENSG00000262814 |
| 22 | Q9BXW7 | NA | CECR5 | cat eye syndrome chromosome region, candidate 5 | 27440 | ENSG00000069998 |
| 23 | P21796 | NA | VDAC1 | voltage-dependent anion channel 1 | 7416 | ENSG00000213585 |
| 24 | P38571 | NA | LIPA | lipase A, lysosomal acid, cholesterol esterase | 3988 | ENSG00000107798 |
| 25 | P32322 | NA | PYCR1 | pyrroline-5-carboxylate reductase 1 | 5831 | ENSG00000183010 |
| 26 | Q9UJZ1 | NA | STOML2 | stomatin (EPB72)-like 2 | 30968 | ENSG00000165283 |
| 27 | Q14980 | NA | NUMA1 | nuclear mitotic apparatus protein 1 | 4926 | ENSG00000137497 |
| 28 | Q99798 | NA | ACO2 | aconitase 2, mitochondrial | 50 | ENSG00000100412 |
| 29 | P51649 | NA | ALDH5A1 | aldehyde dehydrogenase 5 family, member A1 | 7915 | ENSG00000112294 |
| 30 | P12235 | NA | SLC25A4 | solute carrier family 25 (mitochondrial carrier; adenine nucleotide translocator), member 4 | 291 | ENSG00000151729 |
| 31 | Q9NSE4 | NA | IARS2 | isoleucyl-tRNA synthetase 2, mitochondrial | 55699 | ENSG00000067704 |
| 32 | Q6PI48 | NA | DARS2 | aspartyl-tRNA synthetase 2, mitochondrial | 55157 | ENSG00000117593 |
| 33 | P13674 | NA | P4HA1 | prolyl 4-hydroxylase, alpha polypeptide I | 5033 | ENSG00000122884 |
| 34 | Q02978 | NA | SLC25A11 | solute carrier family 25 (mitochondrial carrier; oxoglutarate carrier), member 11 | 8402 | ENSG00000108528 |
| 35 | P60174 | NA | TPI1 | triosephosphate isomerase 1 | 7167 | ENSG00000111669 |
| 36 | P30042 | NA | C21orf33 | chromosome 21 open reading frame 33 | 8209 | ENSG00000160221 |
| 37 | P54819 | NA | AK2 | adenylate kinase 2 | 204 | ENSG00000004455 |
| 38 | P17931 | NA | LGALS3 | lectin, galactoside-binding, soluble, 3 | 3958 | ENSG00000131981 |
| 39 | P05787 | NA | KRT8 | keratin 8 | 3856 | ENSG00000170421 |
| 40 | Q99623 | NA | PHB2 | prohibitin 2 | 11331 | ENSG00000215021 |
| 41 | P50453 | NA | SERPINB9 | serpin peptidase inhibitor, clade B (ovalbumin), member 9 | 5272 | ENSG00000170542 |
| 42 | P35232 | NA | PHB | prohibitin | 5245 | ENSG00000167085 |
| 43 | O95831 | NA | AIFM1 | apoptosis-inducing factor, mitochondrion-associated, 1 | 9131 | ENSG00000156709 |
| 44 | Q99714 | NA | HSD17B10 | hydroxysteroid (17-beta) dehydrogenase 10 | 3028 | ENSG00000072506 |
| 45 | P07741 | NA | APRT | adenine phosphoribosyltransferase | 353 | ENSG00000198931 |
| 46 | O60313 | NA | OPA1 | optic atrophy 1 (autosomal dominant) | 4976 | ENSG00000198836 |
| 47 | P35270 | NA | SPR | sepiapterin reductase (7,8-dihydrobiopterin:NADP+ oxidoreductase) | 6697 | ENSG00000116096 |
| 48 | P38117 | NA | ETFB | electron-transfer-flavoprotein, beta polypeptide | 2109 | ENSG00000105379 |
| 49 | O75367 | NA | H2AFY | H2A histone family, member Y | 9555 | ENSG00000113648 |
| 50 | O75521 | NA | ECI2 | enoyl-CoA delta isomerase 2 | 10455 | ENSG00000198721 |
| 51 | P49419 | NA | ALDH7A1 | aldehyde dehydrogenase 7 family, member A1 | 501 | ENSG00000164904 |
| 52 | Q14764 | NA | MVP | major vault protein | 9961 | ENSG00000013364 |
| 53 | Q9BPW8 | NA | NIPSNAP1 | nipsnap homolog 1 (C. elegans) | 8508 | ENSG00000184117 |
| 54 | Q5SSJ5 | NA | HP1BP3 | heterochromatin protein 1, binding protein 3 | 50809 | ENSG00000127483 |
| 55 | Q13813 | NA | SPTAN1 | spectrin, alpha, non-erythrocytic 1 | 6709 | ENSG00000197694 |
| 56 | O75439 | NA | PMPCB | peptidase (mitochondrial processing) beta | 9512 | ENSG00000105819 |
| 57 | P21953 | NA | BCKDHB | branched chain keto acid dehydrogenase E1, beta polypeptide | 594 | ENSG00000083123 |
| 58 | P30084 | NA | ECHS1 | enoyl CoA hydratase, short chain, 1, mitochondrial | 1892 | ENSG00000127884 |
| 59 | Q13011 | NA | ECH1 | enoyl CoA hydratase 1, peroxisomal | 1891 | ENSG00000104823 |
| 60 | P04040 | NA | CAT | catalase | 847 | ENSG00000121691 |
| 61 | P51688 | NA | SGSH | N-sulfoglucosamine sulfohydrolase | 6448 | ENSG00000181523 |
| 62 | O75340 | NA | PDCD6 | programmed cell death 6 | 10016 | ENSG00000249915 |
| 63 | P30048 | NA | PRDX3 | peroxiredoxin 3 | 10935 | ENSG00000165672 |
| 64 | P24752 | NA | ACAT1 | acetyl-CoA acetyltransferase 1 | 38 | ENSG00000075239 |
| 65 | P19367 | NA | HK1 | hexokinase 1 | 3098 | ENSG00000156515 |
| 66 | P25705 | NA | ATP5A1 | ATP synthase, H+ transporting, mitochondrial F1 complex, alpha subunit 1, cardiac muscle | 498 | ENSG00000152234 |
| 67 | Q9BQ69 | NA | MACROD1 | MACRO domain containing 1 | 28992 | ENSG00000133315 |
| 68 | P05783 | NA | KRT18 | keratin 18 | 3875 | ENSG00000111057 |
| 69 | P06865 | NA | HEXA | hexosaminidase A (alpha polypeptide) | 3073 | ENSG00000213614 |
| 70 | P49748 | NA | ACADVL | acyl-CoA dehydrogenase, very long chain | 37 | ENSG00000072778 |
| 71 | P48047 | NA | ATP5O | ATP synthase, H+ transporting, mitochondrial F1 complex, O subunit | 539 | ENSG00000241837 |
| 72 | Q16531 | NA | DDB1 | damage-specific DNA binding protein 1, 127kDa | 1642 | ENSG00000167986 |
| 73 | P31937 | NA | HIBADH | 3-hydroxyisobutyrate dehydrogenase | 11112 | ENSG00000106049 |

  
  

| **Database:cellular component      &nbspName:intracellular part      &nbspID:GO:0044424** | | | | | | |
| --- | --- | --- | --- | --- | --- | --- |
| C=12096; O=76; E=55.11; R=1.38; rawP=2.30e-11; adjP=8.87e-11 | | | | | | |
| Index | UserID | Value | Gene Symbol | Gene Name | EntrezGene | Ensembl |
| 1 | P10253 | NA | GAA | glucosidase, alpha; acid | 2548 | ENSG00000171298 |
| 2 | Q9UL46 | NA | PSME2 | proteasome (prosome, macropain) activator subunit 2 (PA28 beta) | 5721 | ENSG00000100911 |
| 3 | P40926 | NA | MDH2 | malate dehydrogenase 2, NAD (mitochondrial) | 4191 | ENSG00000146701 |
| 4 | P04179 | NA | SOD2 | superoxide dismutase 2, mitochondrial | 6648 | ENSG00000112096 |
| 5 | Q13724 | NA | MOGS | mannosyl-oligosaccharide glucosidase | 7841 | ENSG00000115275 |
| 6 | P00367 | NA | GLUD1 | glutamate dehydrogenase 1 | 2746 | ENSG00000148672 |
| 7 | Q06323 | NA | PSME1 | proteasome (prosome, macropain) activator subunit 1 (PA28 alpha) | 5720 | ENSG00000092010 |
| 8 | P48735 | NA | IDH2 | isocitrate dehydrogenase 2 (NADP+), mitochondrial | 3418 | ENSG00000182054 |
| 9 | P35580 | NA | MYH10 | myosin, heavy chain 10, non-muscle | 4628 | ENSG00000133026 |
| 10 | P31040 | NA | SDHA | succinate dehydrogenase complex, subunit A, flavoprotein (Fp) | 6389 | ENSG00000073578 |
| 11 | O14773 | NA | TPP1 | tripeptidyl peptidase I | 1200 | ENSG00000166340 |
| 12 | P13804 | NA | ETFA | electron-transfer-flavoprotein, alpha polypeptide | 2108 | ENSG00000140374 |
| 13 | P40121 | NA | CAPG | capping protein (actin filament), gelsolin-like | 822 | ENSG00000042493 |
| 14 | P42765 | NA | ACAA2 | acetyl-CoA acyltransferase 2 | 10449 | ENSG00000167315 |
| 15 | Q9Y277 | NA | VDAC3 | voltage-dependent anion channel 3 | 7419 | ENSG00000078668 |
| 16 | Q04837 | NA | SSBP1 | single-stranded DNA binding protein 1, mitochondrial | 6742 | ENSG00000106028 |
| 17 | P08727 | NA | KRT19 | keratin 19 | 3880 | ENSG00000171345 |
| 18 | P50454 | NA | SERPINH1 | serpin peptidase inhibitor, clade H (heat shock protein 47), member 1, (collagen binding protein 1) | 871 | ENSG00000149257 |
| 19 | Q14697 | NA | GANAB | glucosidase, alpha; neutral AB | 23193 | ENSG00000089597 |
| 20 | P40939 | NA | HADHA | hydroxyacyl-CoA dehydrogenase/3-ketoacyl-CoA thiolase/enoyl-CoA hydratase (trifunctional protein), alpha subunit | 3030 | ENSG00000084754 |
| 21 | P52815 | NA | MRPL12 | mitochondrial ribosomal protein L12 | 6182 | ENSG00000262814 |
| 22 | Q9BXW7 | NA | CECR5 | cat eye syndrome chromosome region, candidate 5 | 27440 | ENSG00000069998 |
| 23 | P21796 | NA | VDAC1 | voltage-dependent anion channel 1 | 7416 | ENSG00000213585 |
| 24 | P38571 | NA | LIPA | lipase A, lysosomal acid, cholesterol esterase | 3988 | ENSG00000107798 |
| 25 | P32322 | NA | PYCR1 | pyrroline-5-carboxylate reductase 1 | 5831 | ENSG00000183010 |
| 26 | Q9UJZ1 | NA | STOML2 | stomatin (EPB72)-like 2 | 30968 | ENSG00000165283 |
| 27 | Q14980 | NA | NUMA1 | nuclear mitotic apparatus protein 1 | 4926 | ENSG00000137497 |
| 28 | Q99798 | NA | ACO2 | aconitase 2, mitochondrial | 50 | ENSG00000100412 |
| 29 | P51649 | NA | ALDH5A1 | aldehyde dehydrogenase 5 family, member A1 | 7915 | ENSG00000112294 |
| 30 | P12235 | NA | SLC25A4 | solute carrier family 25 (mitochondrial carrier; adenine nucleotide translocator), member 4 | 291 | ENSG00000151729 |
| 31 | Q9NSE4 | NA | IARS2 | isoleucyl-tRNA synthetase 2, mitochondrial | 55699 | ENSG00000067704 |
| 32 | Q6PI48 | NA | DARS2 | aspartyl-tRNA synthetase 2, mitochondrial | 55157 | ENSG00000117593 |
| 33 | P13674 | NA | P4HA1 | prolyl 4-hydroxylase, alpha polypeptide I | 5033 | ENSG00000122884 |
| 34 | Q02978 | NA | SLC25A11 | solute carrier family 25 (mitochondrial carrier; oxoglutarate carrier), member 11 | 8402 | ENSG00000108528 |
| 35 | P60174 | NA | TPI1 | triosephosphate isomerase 1 | 7167 | ENSG00000111669 |
| 36 | P30042 | NA | C21orf33 | chromosome 21 open reading frame 33 | 8209 | ENSG00000160221 |
| 37 | P54819 | NA | AK2 | adenylate kinase 2 | 204 | ENSG00000004455 |
| 38 | P17931 | NA | LGALS3 | lectin, galactoside-binding, soluble, 3 | 3958 | ENSG00000131981 |
| 39 | P05787 | NA | KRT8 | keratin 8 | 3856 | ENSG00000170421 |
| 40 | Q99623 | NA | PHB2 | prohibitin 2 | 11331 | ENSG00000215021 |
| 41 | P50453 | NA | SERPINB9 | serpin peptidase inhibitor, clade B (ovalbumin), member 9 | 5272 | ENSG00000170542 |
| 42 | P35232 | NA | PHB | prohibitin | 5245 | ENSG00000167085 |
| 43 | O95831 | NA | AIFM1 | apoptosis-inducing factor, mitochondrion-associated, 1 | 9131 | ENSG00000156709 |
| 44 | Q99714 | NA | HSD17B10 | hydroxysteroid (17-beta) dehydrogenase 10 | 3028 | ENSG00000072506 |
| 45 | P07741 | NA | APRT | adenine phosphoribosyltransferase | 353 | ENSG00000198931 |
| 46 | O60313 | NA | OPA1 | optic atrophy 1 (autosomal dominant) | 4976 | ENSG00000198836 |
| 47 | P35270 | NA | SPR | sepiapterin reductase (7,8-dihydrobiopterin:NADP+ oxidoreductase) | 6697 | ENSG00000116096 |
| 48 | P38117 | NA | ETFB | electron-transfer-flavoprotein, beta polypeptide | 2109 | ENSG00000105379 |
| 49 | P43490 | NA | NAMPT | nicotinamide phosphoribosyltransferase | 10135 | ENSG00000105835 |
| 50 | O75367 | NA | H2AFY | H2A histone family, member Y | 9555 | ENSG00000113648 |
| 51 | O75521 | NA | ECI2 | enoyl-CoA delta isomerase 2 | 10455 | ENSG00000198721 |
| 52 | P49419 | NA | ALDH7A1 | aldehyde dehydrogenase 7 family, member A1 | 501 | ENSG00000164904 |
| 53 | Q14764 | NA | MVP | major vault protein | 9961 | ENSG00000013364 |
| 54 | Q9BPW8 | NA | NIPSNAP1 | nipsnap homolog 1 (C. elegans) | 8508 | ENSG00000184117 |
| 55 | Q5SSJ5 | NA | HP1BP3 | heterochromatin protein 1, binding protein 3 | 50809 | ENSG00000127483 |
| 56 | Q13813 | NA | SPTAN1 | spectrin, alpha, non-erythrocytic 1 | 6709 | ENSG00000197694 |
| 57 | O75439 | NA | PMPCB | peptidase (mitochondrial processing) beta | 9512 | ENSG00000105819 |
| 58 | P02792 | NA | FTL | ferritin, light polypeptide | 2512 | ENSG00000087086 |
| 59 | P21953 | NA | BCKDHB | branched chain keto acid dehydrogenase E1, beta polypeptide | 594 | ENSG00000083123 |
| 60 | P30084 | NA | ECHS1 | enoyl CoA hydratase, short chain, 1, mitochondrial | 1892 | ENSG00000127884 |
| 61 | Q13011 | NA | ECH1 | enoyl CoA hydratase 1, peroxisomal | 1891 | ENSG00000104823 |
| 62 | P04040 | NA | CAT | catalase | 847 | ENSG00000121691 |
| 63 | P51688 | NA | SGSH | N-sulfoglucosamine sulfohydrolase | 6448 | ENSG00000181523 |
| 64 | O75340 | NA | PDCD6 | programmed cell death 6 | 10016 | ENSG00000249915 |
| 65 | P00558 | NA | PGK1 | phosphoglycerate kinase 1 | 5230 | ENSG00000102144 |
| 66 | P30048 | NA | PRDX3 | peroxiredoxin 3 | 10935 | ENSG00000165672 |
| 67 | P24752 | NA | ACAT1 | acetyl-CoA acetyltransferase 1 | 38 | ENSG00000075239 |
| 68 | P19367 | NA | HK1 | hexokinase 1 | 3098 | ENSG00000156515 |
| 69 | P25705 | NA | ATP5A1 | ATP synthase, H+ transporting, mitochondrial F1 complex, alpha subunit 1, cardiac muscle | 498 | ENSG00000152234 |
| 70 | Q9BQ69 | NA | MACROD1 | MACRO domain containing 1 | 28992 | ENSG00000133315 |
| 71 | P05783 | NA | KRT18 | keratin 18 | 3875 | ENSG00000111057 |
| 72 | P06865 | NA | HEXA | hexosaminidase A (alpha polypeptide) | 3073 | ENSG00000213614 |
| 73 | P49748 | NA | ACADVL | acyl-CoA dehydrogenase, very long chain | 37 | ENSG00000072778 |
| 74 | P48047 | NA | ATP5O | ATP synthase, H+ transporting, mitochondrial F1 complex, O subunit | 539 | ENSG00000241837 |
| 75 | Q16531 | NA | DDB1 | damage-specific DNA binding protein 1, 127kDa | 1642 | ENSG00000167986 |
| 76 | P31937 | NA | HIBADH | 3-hydroxyisobutyrate dehydrogenase | 11112 | ENSG00000106049 |

  
  

| **Database:cellular component      &nbspName:intracellular      &nbspID:GO:0005622** | | | | | | |
| --- | --- | --- | --- | --- | --- | --- |
| C=12412; O=76; E=56.55; R=1.34; rawP=1.64e-10; adjP=6.04e-10 | | | | | | |
| Index | UserID | Value | Gene Symbol | Gene Name | EntrezGene | Ensembl |
| 1 | P10253 | NA | GAA | glucosidase, alpha; acid | 2548 | ENSG00000171298 |
| 2 | Q9UL46 | NA | PSME2 | proteasome (prosome, macropain) activator subunit 2 (PA28 beta) | 5721 | ENSG00000100911 |
| 3 | P40926 | NA | MDH2 | malate dehydrogenase 2, NAD (mitochondrial) | 4191 | ENSG00000146701 |
| 4 | P04179 | NA | SOD2 | superoxide dismutase 2, mitochondrial | 6648 | ENSG00000112096 |
| 5 | Q13724 | NA | MOGS | mannosyl-oligosaccharide glucosidase | 7841 | ENSG00000115275 |
| 6 | P00367 | NA | GLUD1 | glutamate dehydrogenase 1 | 2746 | ENSG00000148672 |
| 7 | Q06323 | NA | PSME1 | proteasome (prosome, macropain) activator subunit 1 (PA28 alpha) | 5720 | ENSG00000092010 |
| 8 | P48735 | NA | IDH2 | isocitrate dehydrogenase 2 (NADP+), mitochondrial | 3418 | ENSG00000182054 |
| 9 | P35580 | NA | MYH10 | myosin, heavy chain 10, non-muscle | 4628 | ENSG00000133026 |
| 10 | P31040 | NA | SDHA | succinate dehydrogenase complex, subunit A, flavoprotein (Fp) | 6389 | ENSG00000073578 |
| 11 | O14773 | NA | TPP1 | tripeptidyl peptidase I | 1200 | ENSG00000166340 |
| 12 | P13804 | NA | ETFA | electron-transfer-flavoprotein, alpha polypeptide | 2108 | ENSG00000140374 |
| 13 | P40121 | NA | CAPG | capping protein (actin filament), gelsolin-like | 822 | ENSG00000042493 |
| 14 | P42765 | NA | ACAA2 | acetyl-CoA acyltransferase 2 | 10449 | ENSG00000167315 |
| 15 | Q9Y277 | NA | VDAC3 | voltage-dependent anion channel 3 | 7419 | ENSG00000078668 |
| 16 | Q04837 | NA | SSBP1 | single-stranded DNA binding protein 1, mitochondrial | 6742 | ENSG00000106028 |
| 17 | P08727 | NA | KRT19 | keratin 19 | 3880 | ENSG00000171345 |
| 18 | P50454 | NA | SERPINH1 | serpin peptidase inhibitor, clade H (heat shock protein 47), member 1, (collagen binding protein 1) | 871 | ENSG00000149257 |
| 19 | Q14697 | NA | GANAB | glucosidase, alpha; neutral AB | 23193 | ENSG00000089597 |
| 20 | P40939 | NA | HADHA | hydroxyacyl-CoA dehydrogenase/3-ketoacyl-CoA thiolase/enoyl-CoA hydratase (trifunctional protein), alpha subunit | 3030 | ENSG00000084754 |
| 21 | P52815 | NA | MRPL12 | mitochondrial ribosomal protein L12 | 6182 | ENSG00000262814 |
| 22 | Q9BXW7 | NA | CECR5 | cat eye syndrome chromosome region, candidate 5 | 27440 | ENSG00000069998 |
| 23 | P21796 | NA | VDAC1 | voltage-dependent anion channel 1 | 7416 | ENSG00000213585 |
| 24 | P38571 | NA | LIPA | lipase A, lysosomal acid, cholesterol esterase | 3988 | ENSG00000107798 |
| 25 | P32322 | NA | PYCR1 | pyrroline-5-carboxylate reductase 1 | 5831 | ENSG00000183010 |
| 26 | Q9UJZ1 | NA | STOML2 | stomatin (EPB72)-like 2 | 30968 | ENSG00000165283 |
| 27 | Q14980 | NA | NUMA1 | nuclear mitotic apparatus protein 1 | 4926 | ENSG00000137497 |
| 28 | Q99798 | NA | ACO2 | aconitase 2, mitochondrial | 50 | ENSG00000100412 |
| 29 | P51649 | NA | ALDH5A1 | aldehyde dehydrogenase 5 family, member A1 | 7915 | ENSG00000112294 |
| 30 | P12235 | NA | SLC25A4 | solute carrier family 25 (mitochondrial carrier; adenine nucleotide translocator), member 4 | 291 | ENSG00000151729 |
| 31 | Q9NSE4 | NA | IARS2 | isoleucyl-tRNA synthetase 2, mitochondrial | 55699 | ENSG00000067704 |
| 32 | Q6PI48 | NA | DARS2 | aspartyl-tRNA synthetase 2, mitochondrial | 55157 | ENSG00000117593 |
| 33 | P13674 | NA | P4HA1 | prolyl 4-hydroxylase, alpha polypeptide I | 5033 | ENSG00000122884 |
| 34 | Q02978 | NA | SLC25A11 | solute carrier family 25 (mitochondrial carrier; oxoglutarate carrier), member 11 | 8402 | ENSG00000108528 |
| 35 | P60174 | NA | TPI1 | triosephosphate isomerase 1 | 7167 | ENSG00000111669 |
| 36 | P30042 | NA | C21orf33 | chromosome 21 open reading frame 33 | 8209 | ENSG00000160221 |
| 37 | P54819 | NA | AK2 | adenylate kinase 2 | 204 | ENSG00000004455 |
| 38 | P17931 | NA | LGALS3 | lectin, galactoside-binding, soluble, 3 | 3958 | ENSG00000131981 |
| 39 | P05787 | NA | KRT8 | keratin 8 | 3856 | ENSG00000170421 |
| 40 | Q99623 | NA | PHB2 | prohibitin 2 | 11331 | ENSG00000215021 |
| 41 | P50453 | NA | SERPINB9 | serpin peptidase inhibitor, clade B (ovalbumin), member 9 | 5272 | ENSG00000170542 |
| 42 | P35232 | NA | PHB | prohibitin | 5245 | ENSG00000167085 |
| 43 | O95831 | NA | AIFM1 | apoptosis-inducing factor, mitochondrion-associated, 1 | 9131 | ENSG00000156709 |
| 44 | Q99714 | NA | HSD17B10 | hydroxysteroid (17-beta) dehydrogenase 10 | 3028 | ENSG00000072506 |
| 45 | P07741 | NA | APRT | adenine phosphoribosyltransferase | 353 | ENSG00000198931 |
| 46 | O60313 | NA | OPA1 | optic atrophy 1 (autosomal dominant) | 4976 | ENSG00000198836 |
| 47 | P35270 | NA | SPR | sepiapterin reductase (7,8-dihydrobiopterin:NADP+ oxidoreductase) | 6697 | ENSG00000116096 |
| 48 | P38117 | NA | ETFB | electron-transfer-flavoprotein, beta polypeptide | 2109 | ENSG00000105379 |
| 49 | P43490 | NA | NAMPT | nicotinamide phosphoribosyltransferase | 10135 | ENSG00000105835 |
| 50 | O75367 | NA | H2AFY | H2A histone family, member Y | 9555 | ENSG00000113648 |
| 51 | O75521 | NA | ECI2 | enoyl-CoA delta isomerase 2 | 10455 | ENSG00000198721 |
| 52 | P49419 | NA | ALDH7A1 | aldehyde dehydrogenase 7 family, member A1 | 501 | ENSG00000164904 |
| 53 | Q14764 | NA | MVP | major vault protein | 9961 | ENSG00000013364 |
| 54 | Q9BPW8 | NA | NIPSNAP1 | nipsnap homolog 1 (C. elegans) | 8508 | ENSG00000184117 |
| 55 | Q5SSJ5 | NA | HP1BP3 | heterochromatin protein 1, binding protein 3 | 50809 | ENSG00000127483 |
| 56 | Q13813 | NA | SPTAN1 | spectrin, alpha, non-erythrocytic 1 | 6709 | ENSG00000197694 |
| 57 | O75439 | NA | PMPCB | peptidase (mitochondrial processing) beta | 9512 | ENSG00000105819 |
| 58 | P02792 | NA | FTL | ferritin, light polypeptide | 2512 | ENSG00000087086 |
| 59 | P21953 | NA | BCKDHB | branched chain keto acid dehydrogenase E1, beta polypeptide | 594 | ENSG00000083123 |
| 60 | P30084 | NA | ECHS1 | enoyl CoA hydratase, short chain, 1, mitochondrial | 1892 | ENSG00000127884 |
| 61 | Q13011 | NA | ECH1 | enoyl CoA hydratase 1, peroxisomal | 1891 | ENSG00000104823 |
| 62 | P04040 | NA | CAT | catalase | 847 | ENSG00000121691 |
| 63 | P51688 | NA | SGSH | N-sulfoglucosamine sulfohydrolase | 6448 | ENSG00000181523 |
| 64 | O75340 | NA | PDCD6 | programmed cell death 6 | 10016 | ENSG00000249915 |
| 65 | P00558 | NA | PGK1 | phosphoglycerate kinase 1 | 5230 | ENSG00000102144 |
| 66 | P30048 | NA | PRDX3 | peroxiredoxin 3 | 10935 | ENSG00000165672 |
| 67 | P24752 | NA | ACAT1 | acetyl-CoA acetyltransferase 1 | 38 | ENSG00000075239 |
| 68 | P19367 | NA | HK1 | hexokinase 1 | 3098 | ENSG00000156515 |
| 69 | P25705 | NA | ATP5A1 | ATP synthase, H+ transporting, mitochondrial F1 complex, alpha subunit 1, cardiac muscle | 498 | ENSG00000152234 |
| 70 | Q9BQ69 | NA | MACROD1 | MACRO domain containing 1 | 28992 | ENSG00000133315 |
| 71 | P05783 | NA | KRT18 | keratin 18 | 3875 | ENSG00000111057 |
| 72 | P06865 | NA | HEXA | hexosaminidase A (alpha polypeptide) | 3073 | ENSG00000213614 |
| 73 | P49748 | NA | ACADVL | acyl-CoA dehydrogenase, very long chain | 37 | ENSG00000072778 |
| 74 | P48047 | NA | ATP5O | ATP synthase, H+ transporting, mitochondrial F1 complex, O subunit | 539 | ENSG00000241837 |
| 75 | Q16531 | NA | DDB1 | damage-specific DNA binding protein 1, 127kDa | 1642 | ENSG00000167986 |
| 76 | P31937 | NA | HIBADH | 3-hydroxyisobutyrate dehydrogenase | 11112 | ENSG00000106049 |

  
  

| **Database:cellular component      &nbspName:cell part      &nbspID:GO:0044464** | | | | | | |
| --- | --- | --- | --- | --- | --- | --- |
| C=14405; O=76; E=65.63; R=1.16; rawP=1.39e-05; adjP=4.20e-05 | | | | | | |
| Index | UserID | Value | Gene Symbol | Gene Name | EntrezGene | Ensembl |
| 1 | P10253 | NA | GAA | glucosidase, alpha; acid | 2548 | ENSG00000171298 |
| 2 | Q9UL46 | NA | PSME2 | proteasome (prosome, macropain) activator subunit 2 (PA28 beta) | 5721 | ENSG00000100911 |
| 3 | P40926 | NA | MDH2 | malate dehydrogenase 2, NAD (mitochondrial) | 4191 | ENSG00000146701 |
| 4 | P04179 | NA | SOD2 | superoxide dismutase 2, mitochondrial | 6648 | ENSG00000112096 |
| 5 | Q13724 | NA | MOGS | mannosyl-oligosaccharide glucosidase | 7841 | ENSG00000115275 |
| 6 | P00367 | NA | GLUD1 | glutamate dehydrogenase 1 | 2746 | ENSG00000148672 |
| 7 | Q06323 | NA | PSME1 | proteasome (prosome, macropain) activator subunit 1 (PA28 alpha) | 5720 | ENSG00000092010 |
| 8 | P48735 | NA | IDH2 | isocitrate dehydrogenase 2 (NADP+), mitochondrial | 3418 | ENSG00000182054 |
| 9 | P35580 | NA | MYH10 | myosin, heavy chain 10, non-muscle | 4628 | ENSG00000133026 |
| 10 | P31040 | NA | SDHA | succinate dehydrogenase complex, subunit A, flavoprotein (Fp) | 6389 | ENSG00000073578 |
| 11 | O14773 | NA | TPP1 | tripeptidyl peptidase I | 1200 | ENSG00000166340 |
| 12 | P13804 | NA | ETFA | electron-transfer-flavoprotein, alpha polypeptide | 2108 | ENSG00000140374 |
| 13 | P40121 | NA | CAPG | capping protein (actin filament), gelsolin-like | 822 | ENSG00000042493 |
| 14 | P42765 | NA | ACAA2 | acetyl-CoA acyltransferase 2 | 10449 | ENSG00000167315 |
| 15 | Q9Y277 | NA | VDAC3 | voltage-dependent anion channel 3 | 7419 | ENSG00000078668 |
| 16 | Q04837 | NA | SSBP1 | single-stranded DNA binding protein 1, mitochondrial | 6742 | ENSG00000106028 |
| 17 | P08727 | NA | KRT19 | keratin 19 | 3880 | ENSG00000171345 |
| 18 | P50454 | NA | SERPINH1 | serpin peptidase inhibitor, clade H (heat shock protein 47), member 1, (collagen binding protein 1) | 871 | ENSG00000149257 |
| 19 | Q14697 | NA | GANAB | glucosidase, alpha; neutral AB | 23193 | ENSG00000089597 |
| 20 | P40939 | NA | HADHA | hydroxyacyl-CoA dehydrogenase/3-ketoacyl-CoA thiolase/enoyl-CoA hydratase (trifunctional protein), alpha subunit | 3030 | ENSG00000084754 |
| 21 | P52815 | NA | MRPL12 | mitochondrial ribosomal protein L12 | 6182 | ENSG00000262814 |
| 22 | Q9BXW7 | NA | CECR5 | cat eye syndrome chromosome region, candidate 5 | 27440 | ENSG00000069998 |
| 23 | P21796 | NA | VDAC1 | voltage-dependent anion channel 1 | 7416 | ENSG00000213585 |
| 24 | P38571 | NA | LIPA | lipase A, lysosomal acid, cholesterol esterase | 3988 | ENSG00000107798 |
| 25 | P32322 | NA | PYCR1 | pyrroline-5-carboxylate reductase 1 | 5831 | ENSG00000183010 |
| 26 | Q9UJZ1 | NA | STOML2 | stomatin (EPB72)-like 2 | 30968 | ENSG00000165283 |
| 27 | Q14980 | NA | NUMA1 | nuclear mitotic apparatus protein 1 | 4926 | ENSG00000137497 |
| 28 | Q99798 | NA | ACO2 | aconitase 2, mitochondrial | 50 | ENSG00000100412 |
| 29 | P51649 | NA | ALDH5A1 | aldehyde dehydrogenase 5 family, member A1 | 7915 | ENSG00000112294 |
| 30 | P12235 | NA | SLC25A4 | solute carrier family 25 (mitochondrial carrier; adenine nucleotide translocator), member 4 | 291 | ENSG00000151729 |
| 31 | Q9NSE4 | NA | IARS2 | isoleucyl-tRNA synthetase 2, mitochondrial | 55699 | ENSG00000067704 |
| 32 | Q6PI48 | NA | DARS2 | aspartyl-tRNA synthetase 2, mitochondrial | 55157 | ENSG00000117593 |
| 33 | P13674 | NA | P4HA1 | prolyl 4-hydroxylase, alpha polypeptide I | 5033 | ENSG00000122884 |
| 34 | Q02978 | NA | SLC25A11 | solute carrier family 25 (mitochondrial carrier; oxoglutarate carrier), member 11 | 8402 | ENSG00000108528 |
| 35 | P60174 | NA | TPI1 | triosephosphate isomerase 1 | 7167 | ENSG00000111669 |
| 36 | P30042 | NA | C21orf33 | chromosome 21 open reading frame 33 | 8209 | ENSG00000160221 |
| 37 | P54819 | NA | AK2 | adenylate kinase 2 | 204 | ENSG00000004455 |
| 38 | P17931 | NA | LGALS3 | lectin, galactoside-binding, soluble, 3 | 3958 | ENSG00000131981 |
| 39 | P05787 | NA | KRT8 | keratin 8 | 3856 | ENSG00000170421 |
| 40 | Q99623 | NA | PHB2 | prohibitin 2 | 11331 | ENSG00000215021 |
| 41 | P50453 | NA | SERPINB9 | serpin peptidase inhibitor, clade B (ovalbumin), member 9 | 5272 | ENSG00000170542 |
| 42 | P35232 | NA | PHB | prohibitin | 5245 | ENSG00000167085 |
| 43 | O95831 | NA | AIFM1 | apoptosis-inducing factor, mitochondrion-associated, 1 | 9131 | ENSG00000156709 |
| 44 | Q99714 | NA | HSD17B10 | hydroxysteroid (17-beta) dehydrogenase 10 | 3028 | ENSG00000072506 |
| 45 | P07741 | NA | APRT | adenine phosphoribosyltransferase | 353 | ENSG00000198931 |
| 46 | O60313 | NA | OPA1 | optic atrophy 1 (autosomal dominant) | 4976 | ENSG00000198836 |
| 47 | P35270 | NA | SPR | sepiapterin reductase (7,8-dihydrobiopterin:NADP+ oxidoreductase) | 6697 | ENSG00000116096 |
| 48 | P38117 | NA | ETFB | electron-transfer-flavoprotein, beta polypeptide | 2109 | ENSG00000105379 |
| 49 | P43490 | NA | NAMPT | nicotinamide phosphoribosyltransferase | 10135 | ENSG00000105835 |
| 50 | O75367 | NA | H2AFY | H2A histone family, member Y | 9555 | ENSG00000113648 |
| 51 | O75521 | NA | ECI2 | enoyl-CoA delta isomerase 2 | 10455 | ENSG00000198721 |
| 52 | P49419 | NA | ALDH7A1 | aldehyde dehydrogenase 7 family, member A1 | 501 | ENSG00000164904 |
| 53 | Q14764 | NA | MVP | major vault protein | 9961 | ENSG00000013364 |
| 54 | Q9BPW8 | NA | NIPSNAP1 | nipsnap homolog 1 (C. elegans) | 8508 | ENSG00000184117 |
| 55 | Q5SSJ5 | NA | HP1BP3 | heterochromatin protein 1, binding protein 3 | 50809 | ENSG00000127483 |
| 56 | Q13813 | NA | SPTAN1 | spectrin, alpha, non-erythrocytic 1 | 6709 | ENSG00000197694 |
| 57 | O75439 | NA | PMPCB | peptidase (mitochondrial processing) beta | 9512 | ENSG00000105819 |
| 58 | P02792 | NA | FTL | ferritin, light polypeptide | 2512 | ENSG00000087086 |
| 59 | P21953 | NA | BCKDHB | branched chain keto acid dehydrogenase E1, beta polypeptide | 594 | ENSG00000083123 |
| 60 | P30084 | NA | ECHS1 | enoyl CoA hydratase, short chain, 1, mitochondrial | 1892 | ENSG00000127884 |
| 61 | Q13011 | NA | ECH1 | enoyl CoA hydratase 1, peroxisomal | 1891 | ENSG00000104823 |
| 62 | P04040 | NA | CAT | catalase | 847 | ENSG00000121691 |
| 63 | P51688 | NA | SGSH | N-sulfoglucosamine sulfohydrolase | 6448 | ENSG00000181523 |
| 64 | O75340 | NA | PDCD6 | programmed cell death 6 | 10016 | ENSG00000249915 |
| 65 | P00558 | NA | PGK1 | phosphoglycerate kinase 1 | 5230 | ENSG00000102144 |
| 66 | P30048 | NA | PRDX3 | peroxiredoxin 3 | 10935 | ENSG00000165672 |
| 67 | P24752 | NA | ACAT1 | acetyl-CoA acetyltransferase 1 | 38 | ENSG00000075239 |
| 68 | P19367 | NA | HK1 | hexokinase 1 | 3098 | ENSG00000156515 |
| 69 | P25705 | NA | ATP5A1 | ATP synthase, H+ transporting, mitochondrial F1 complex, alpha subunit 1, cardiac muscle | 498 | ENSG00000152234 |
| 70 | Q9BQ69 | NA | MACROD1 | MACRO domain containing 1 | 28992 | ENSG00000133315 |
| 71 | P05783 | NA | KRT18 | keratin 18 | 3875 | ENSG00000111057 |
| 72 | P06865 | NA | HEXA | hexosaminidase A (alpha polypeptide) | 3073 | ENSG00000213614 |
| 73 | P49748 | NA | ACADVL | acyl-CoA dehydrogenase, very long chain | 37 | ENSG00000072778 |
| 74 | P48047 | NA | ATP5O | ATP synthase, H+ transporting, mitochondrial F1 complex, O subunit | 539 | ENSG00000241837 |
| 75 | Q16531 | NA | DDB1 | damage-specific DNA binding protein 1, 127kDa | 1642 | ENSG00000167986 |
| 76 | P31937 | NA | HIBADH | 3-hydroxyisobutyrate dehydrogenase | 11112 | ENSG00000106049 |

  
  

| **Database:cellular component      &nbspName:cell      &nbspID:GO:0005623** | | | | | | |
| --- | --- | --- | --- | --- | --- | --- |
| C=14406; O=76; E=65.63; R=1.16; rawP=1.40e-05; adjP=4.20e-05 | | | | | | |
| Index | UserID | Value | Gene Symbol | Gene Name | EntrezGene | Ensembl |
| 1 | P10253 | NA | GAA | glucosidase, alpha; acid | 2548 | ENSG00000171298 |
| 2 | Q9UL46 | NA | PSME2 | proteasome (prosome, macropain) activator subunit 2 (PA28 beta) | 5721 | ENSG00000100911 |
| 3 | P40926 | NA | MDH2 | malate dehydrogenase 2, NAD (mitochondrial) | 4191 | ENSG00000146701 |
| 4 | P04179 | NA | SOD2 | superoxide dismutase 2, mitochondrial | 6648 | ENSG00000112096 |
| 5 | Q13724 | NA | MOGS | mannosyl-oligosaccharide glucosidase | 7841 | ENSG00000115275 |
| 6 | P00367 | NA | GLUD1 | glutamate dehydrogenase 1 | 2746 | ENSG00000148672 |
| 7 | Q06323 | NA | PSME1 | proteasome (prosome, macropain) activator subunit 1 (PA28 alpha) | 5720 | ENSG00000092010 |
| 8 | P48735 | NA | IDH2 | isocitrate dehydrogenase 2 (NADP+), mitochondrial | 3418 | ENSG00000182054 |
| 9 | P35580 | NA | MYH10 | myosin, heavy chain 10, non-muscle | 4628 | ENSG00000133026 |
| 10 | P31040 | NA | SDHA | succinate dehydrogenase complex, subunit A, flavoprotein (Fp) | 6389 | ENSG00000073578 |
| 11 | O14773 | NA | TPP1 | tripeptidyl peptidase I | 1200 | ENSG00000166340 |
| 12 | P13804 | NA | ETFA | electron-transfer-flavoprotein, alpha polypeptide | 2108 | ENSG00000140374 |
| 13 | P40121 | NA | CAPG | capping protein (actin filament), gelsolin-like | 822 | ENSG00000042493 |
| 14 | P42765 | NA | ACAA2 | acetyl-CoA acyltransferase 2 | 10449 | ENSG00000167315 |
| 15 | Q9Y277 | NA | VDAC3 | voltage-dependent anion channel 3 | 7419 | ENSG00000078668 |
| 16 | Q04837 | NA | SSBP1 | single-stranded DNA binding protein 1, mitochondrial | 6742 | ENSG00000106028 |
| 17 | P08727 | NA | KRT19 | keratin 19 | 3880 | ENSG00000171345 |
| 18 | P50454 | NA | SERPINH1 | serpin peptidase inhibitor, clade H (heat shock protein 47), member 1, (collagen binding protein 1) | 871 | ENSG00000149257 |
| 19 | Q14697 | NA | GANAB | glucosidase, alpha; neutral AB | 23193 | ENSG00000089597 |
| 20 | P40939 | NA | HADHA | hydroxyacyl-CoA dehydrogenase/3-ketoacyl-CoA thiolase/enoyl-CoA hydratase (trifunctional protein), alpha subunit | 3030 | ENSG00000084754 |
| 21 | P52815 | NA | MRPL12 | mitochondrial ribosomal protein L12 | 6182 | ENSG00000262814 |
| 22 | Q9BXW7 | NA | CECR5 | cat eye syndrome chromosome region, candidate 5 | 27440 | ENSG00000069998 |
| 23 | P21796 | NA | VDAC1 | voltage-dependent anion channel 1 | 7416 | ENSG00000213585 |
| 24 | P38571 | NA | LIPA | lipase A, lysosomal acid, cholesterol esterase | 3988 | ENSG00000107798 |
| 25 | P32322 | NA | PYCR1 | pyrroline-5-carboxylate reductase 1 | 5831 | ENSG00000183010 |
| 26 | Q9UJZ1 | NA | STOML2 | stomatin (EPB72)-like 2 | 30968 | ENSG00000165283 |
| 27 | Q14980 | NA | NUMA1 | nuclear mitotic apparatus protein 1 | 4926 | ENSG00000137497 |
| 28 | Q99798 | NA | ACO2 | aconitase 2, mitochondrial | 50 | ENSG00000100412 |
| 29 | P51649 | NA | ALDH5A1 | aldehyde dehydrogenase 5 family, member A1 | 7915 | ENSG00000112294 |
| 30 | P12235 | NA | SLC25A4 | solute carrier family 25 (mitochondrial carrier; adenine nucleotide translocator), member 4 | 291 | ENSG00000151729 |
| 31 | Q9NSE4 | NA | IARS2 | isoleucyl-tRNA synthetase 2, mitochondrial | 55699 | ENSG00000067704 |
| 32 | Q6PI48 | NA | DARS2 | aspartyl-tRNA synthetase 2, mitochondrial | 55157 | ENSG00000117593 |
| 33 | P13674 | NA | P4HA1 | prolyl 4-hydroxylase, alpha polypeptide I | 5033 | ENSG00000122884 |
| 34 | Q02978 | NA | SLC25A11 | solute carrier family 25 (mitochondrial carrier; oxoglutarate carrier), member 11 | 8402 | ENSG00000108528 |
| 35 | P60174 | NA | TPI1 | triosephosphate isomerase 1 | 7167 | ENSG00000111669 |
| 36 | P30042 | NA | C21orf33 | chromosome 21 open reading frame 33 | 8209 | ENSG00000160221 |
| 37 | P54819 | NA | AK2 | adenylate kinase 2 | 204 | ENSG00000004455 |
| 38 | P17931 | NA | LGALS3 | lectin, galactoside-binding, soluble, 3 | 3958 | ENSG00000131981 |
| 39 | P05787 | NA | KRT8 | keratin 8 | 3856 | ENSG00000170421 |
| 40 | Q99623 | NA | PHB2 | prohibitin 2 | 11331 | ENSG00000215021 |
| 41 | P50453 | NA | SERPINB9 | serpin peptidase inhibitor, clade B (ovalbumin), member 9 | 5272 | ENSG00000170542 |
| 42 | P35232 | NA | PHB | prohibitin | 5245 | ENSG00000167085 |
| 43 | O95831 | NA | AIFM1 | apoptosis-inducing factor, mitochondrion-associated, 1 | 9131 | ENSG00000156709 |
| 44 | Q99714 | NA | HSD17B10 | hydroxysteroid (17-beta) dehydrogenase 10 | 3028 | ENSG00000072506 |
| 45 | P07741 | NA | APRT | adenine phosphoribosyltransferase | 353 | ENSG00000198931 |
| 46 | O60313 | NA | OPA1 | optic atrophy 1 (autosomal dominant) | 4976 | ENSG00000198836 |
| 47 | P35270 | NA | SPR | sepiapterin reductase (7,8-dihydrobiopterin:NADP+ oxidoreductase) | 6697 | ENSG00000116096 |
| 48 | P38117 | NA | ETFB | electron-transfer-flavoprotein, beta polypeptide | 2109 | ENSG00000105379 |
| 49 | P43490 | NA | NAMPT | nicotinamide phosphoribosyltransferase | 10135 | ENSG00000105835 |
| 50 | O75367 | NA | H2AFY | H2A histone family, member Y | 9555 | ENSG00000113648 |
| 51 | O75521 | NA | ECI2 | enoyl-CoA delta isomerase 2 | 10455 | ENSG00000198721 |
| 52 | P49419 | NA | ALDH7A1 | aldehyde dehydrogenase 7 family, member A1 | 501 | ENSG00000164904 |
| 53 | Q14764 | NA | MVP | major vault protein | 9961 | ENSG00000013364 |
| 54 | Q9BPW8 | NA | NIPSNAP1 | nipsnap homolog 1 (C. elegans) | 8508 | ENSG00000184117 |
| 55 | Q5SSJ5 | NA | HP1BP3 | heterochromatin protein 1, binding protein 3 | 50809 | ENSG00000127483 |
| 56 | Q13813 | NA | SPTAN1 | spectrin, alpha, non-erythrocytic 1 | 6709 | ENSG00000197694 |
| 57 | O75439 | NA | PMPCB | peptidase (mitochondrial processing) beta | 9512 | ENSG00000105819 |
| 58 | P02792 | NA | FTL | ferritin, light polypeptide | 2512 | ENSG00000087086 |
| 59 | P21953 | NA | BCKDHB | branched chain keto acid dehydrogenase E1, beta polypeptide | 594 | ENSG00000083123 |
| 60 | P30084 | NA | ECHS1 | enoyl CoA hydratase, short chain, 1, mitochondrial | 1892 | ENSG00000127884 |
| 61 | Q13011 | NA | ECH1 | enoyl CoA hydratase 1, peroxisomal | 1891 | ENSG00000104823 |
| 62 | P04040 | NA | CAT | catalase | 847 | ENSG00000121691 |
| 63 | P51688 | NA | SGSH | N-sulfoglucosamine sulfohydrolase | 6448 | ENSG00000181523 |
| 64 | O75340 | NA | PDCD6 | programmed cell death 6 | 10016 | ENSG00000249915 |
| 65 | P00558 | NA | PGK1 | phosphoglycerate kinase 1 | 5230 | ENSG00000102144 |
| 66 | P30048 | NA | PRDX3 | peroxiredoxin 3 | 10935 | ENSG00000165672 |
| 67 | P24752 | NA | ACAT1 | acetyl-CoA acetyltransferase 1 | 38 | ENSG00000075239 |
| 68 | P19367 | NA | HK1 | hexokinase 1 | 3098 | ENSG00000156515 |
| 69 | P25705 | NA | ATP5A1 | ATP synthase, H+ transporting, mitochondrial F1 complex, alpha subunit 1, cardiac muscle | 498 | ENSG00000152234 |
| 70 | Q9BQ69 | NA | MACROD1 | MACRO domain containing 1 | 28992 | ENSG00000133315 |
| 71 | P05783 | NA | KRT18 | keratin 18 | 3875 | ENSG00000111057 |
| 72 | P06865 | NA | HEXA | hexosaminidase A (alpha polypeptide) | 3073 | ENSG00000213614 |
| 73 | P49748 | NA | ACADVL | acyl-CoA dehydrogenase, very long chain | 37 | ENSG00000072778 |
| 74 | P48047 | NA | ATP5O | ATP synthase, H+ transporting, mitochondrial F1 complex, O subunit | 539 | ENSG00000241837 |
| 75 | Q16531 | NA | DDB1 | damage-specific DNA binding protein 1, 127kDa | 1642 | ENSG00000167986 |
| 76 | P31937 | NA | HIBADH | 3-hydroxyisobutyrate dehydrogenase | 11112 | ENSG00000106049 |

  
  

| **Database:cellular component      &nbspName:mitochondrial intermembrane space      &nbspID:GO:0005758** | | | | | | |
| --- | --- | --- | --- | --- | --- | --- |
| C=44; O=4; E=0.20; R=19.95; rawP=4.70e-05; adjP=0.0001 | | | | | | |
| Index | UserID | Value | Gene Symbol | Gene Name | EntrezGene | Ensembl |
| 1 | P04040 | NA | CAT | catalase | 847 | ENSG00000121691 |
| 2 | P54819 | NA | AK2 | adenylate kinase 2 | 204 | ENSG00000004455 |
| 3 | O60313 | NA | OPA1 | optic atrophy 1 (autosomal dominant) | 4976 | ENSG00000198836 |
| 4 | O95831 | NA | AIFM1 | apoptosis-inducing factor, mitochondrion-associated, 1 | 9131 | ENSG00000156709 |

  
  

| **Database:cellular component      &nbspName:organelle envelope lumen      &nbspID:GO:0031970** | | | | | | |
| --- | --- | --- | --- | --- | --- | --- |
| C=51; O=4; E=0.23; R=17.22; rawP=8.45e-05; adjP=0.0002 | | | | | | |
| Index | UserID | Value | Gene Symbol | Gene Name | EntrezGene | Ensembl |
| 1 | P04040 | NA | CAT | catalase | 847 | ENSG00000121691 |
| 2 | P54819 | NA | AK2 | adenylate kinase 2 | 204 | ENSG00000004455 |
| 3 | O60313 | NA | OPA1 | optic atrophy 1 (autosomal dominant) | 4976 | ENSG00000198836 |
| 4 | O95831 | NA | AIFM1 | apoptosis-inducing factor, mitochondrion-associated, 1 | 9131 | ENSG00000156709 |

  
  

| **Database:cellular component      &nbspName:mitochondrial outer membrane      &nbspID:GO:0005741** | | | | | | |
| --- | --- | --- | --- | --- | --- | --- |
| C=122; O=5; E=0.56; R=9.00; rawP=0.0002; adjP=0.0005 | | | | | | |
| Index | UserID | Value | Gene Symbol | Gene Name | EntrezGene | Ensembl |
| 1 | Q9Y277 | NA | VDAC3 | voltage-dependent anion channel 3 | 7419 | ENSG00000078668 |
| 2 | P19367 | NA | HK1 | hexokinase 1 | 3098 | ENSG00000156515 |
| 3 | O60313 | NA | OPA1 | optic atrophy 1 (autosomal dominant) | 4976 | ENSG00000198836 |
| 4 | P12235 | NA | SLC25A4 | solute carrier family 25 (mitochondrial carrier; adenine nucleotide translocator), member 4 | 291 | ENSG00000151729 |
| 5 | P21796 | NA | VDAC1 | voltage-dependent anion channel 1 | 7416 | ENSG00000213585 |

  
  

| **Database:cellular component      &nbspName:organelle outer membrane      &nbspID:GO:0031968** | | | | | | |
| --- | --- | --- | --- | --- | --- | --- |
| C=143; O=5; E=0.65; R=7.67; rawP=0.0005; adjP=0.0013 | | | | | | |
| Index | UserID | Value | Gene Symbol | Gene Name | EntrezGene | Ensembl |
| 1 | Q9Y277 | NA | VDAC3 | voltage-dependent anion channel 3 | 7419 | ENSG00000078668 |
| 2 | P19367 | NA | HK1 | hexokinase 1 | 3098 | ENSG00000156515 |
| 3 | O60313 | NA | OPA1 | optic atrophy 1 (autosomal dominant) | 4976 | ENSG00000198836 |
| 4 | P12235 | NA | SLC25A4 | solute carrier family 25 (mitochondrial carrier; adenine nucleotide translocator), member 4 | 291 | ENSG00000151729 |
| 5 | P21796 | NA | VDAC1 | voltage-dependent anion channel 1 | 7416 | ENSG00000213585 |

  
  

| **Database:cellular component      &nbspName:mitochondrial membrane part      &nbspID:GO:0044455** | | | | | | |
| --- | --- | --- | --- | --- | --- | --- |
| C=145; O=5; E=0.66; R=7.57; rawP=0.0005; adjP=0.0013 | | | | | | |
| Index | UserID | Value | Gene Symbol | Gene Name | EntrezGene | Ensembl |
| 1 | P25705 | NA | ATP5A1 | ATP synthase, H+ transporting, mitochondrial F1 complex, alpha subunit 1, cardiac muscle | 498 | ENSG00000152234 |
| 2 | O60313 | NA | OPA1 | optic atrophy 1 (autosomal dominant) | 4976 | ENSG00000198836 |
| 3 | P48047 | NA | ATP5O | ATP synthase, H+ transporting, mitochondrial F1 complex, O subunit | 539 | ENSG00000241837 |
| 4 | P31040 | NA | SDHA | succinate dehydrogenase complex, subunit A, flavoprotein (Fp) | 6389 | ENSG00000073578 |
| 5 | O95831 | NA | AIFM1 | apoptosis-inducing factor, mitochondrion-associated, 1 | 9131 | ENSG00000156709 |

  
  

| **Database:cellular component      &nbspName:outer membrane      &nbspID:GO:0019867** | | | | | | |
| --- | --- | --- | --- | --- | --- | --- |
| C=148; O=5; E=0.67; R=7.42; rawP=0.0006; adjP=0.0015 | | | | | | |
| Index | UserID | Value | Gene Symbol | Gene Name | EntrezGene | Ensembl |
| 1 | Q9Y277 | NA | VDAC3 | voltage-dependent anion channel 3 | 7419 | ENSG00000078668 |
| 2 | P19367 | NA | HK1 | hexokinase 1 | 3098 | ENSG00000156515 |
| 3 | O60313 | NA | OPA1 | optic atrophy 1 (autosomal dominant) | 4976 | ENSG00000198836 |
| 4 | P12235 | NA | SLC25A4 | solute carrier family 25 (mitochondrial carrier; adenine nucleotide translocator), member 4 | 291 | ENSG00000151729 |
| 5 | P21796 | NA | VDAC1 | voltage-dependent anion channel 1 | 7416 | ENSG00000213585 |

  
  

| **Database:cellular component      &nbspName:lysosomal lumen      &nbspID:GO:0043202** | | | | | | |
| --- | --- | --- | --- | --- | --- | --- |
| C=71; O=3; E=0.32; R=9.27; rawP=0.0042; adjP=0.0100 | | | | | | |
| Index | UserID | Value | Gene Symbol | Gene Name | EntrezGene | Ensembl |
| 1 | O14773 | NA | TPP1 | tripeptidyl peptidase I | 1200 | ENSG00000166340 |
| 2 | P51688 | NA | SGSH | N-sulfoglucosamine sulfohydrolase | 6448 | ENSG00000181523 |
| 3 | P06865 | NA | HEXA | hexosaminidase A (alpha polypeptide) | 3073 | ENSG00000213614 |

  
  

| **Database:cellular component      &nbspName:lytic vacuole      &nbspID:GO:0000323** | | | | | | |
| --- | --- | --- | --- | --- | --- | --- |
| C=345; O=6; E=1.57; R=3.82; rawP=0.0049; adjP=0.0107 | | | | | | |
| Index | UserID | Value | Gene Symbol | Gene Name | EntrezGene | Ensembl |
| 1 | P04040 | NA | CAT | catalase | 847 | ENSG00000121691 |
| 2 | O14773 | NA | TPP1 | tripeptidyl peptidase I | 1200 | ENSG00000166340 |
| 3 | P10253 | NA | GAA | glucosidase, alpha; acid | 2548 | ENSG00000171298 |
| 4 | P51688 | NA | SGSH | N-sulfoglucosamine sulfohydrolase | 6448 | ENSG00000181523 |
| 5 | P06865 | NA | HEXA | hexosaminidase A (alpha polypeptide) | 3073 | ENSG00000213614 |
| 6 | P38571 | NA | LIPA | lipase A, lysosomal acid, cholesterol esterase | 3988 | ENSG00000107798 |

  
  

| **Database:cellular component      &nbspName:vacuolar lumen      &nbspID:GO:0005775** | | | | | | |
| --- | --- | --- | --- | --- | --- | --- |
| C=75; O=3; E=0.34; R=8.78; rawP=0.0048; adjP=0.0107 | | | | | | |
| Index | UserID | Value | Gene Symbol | Gene Name | EntrezGene | Ensembl |
| 1 | O14773 | NA | TPP1 | tripeptidyl peptidase I | 1200 | ENSG00000166340 |
| 2 | P51688 | NA | SGSH | N-sulfoglucosamine sulfohydrolase | 6448 | ENSG00000181523 |
| 3 | P06865 | NA | HEXA | hexosaminidase A (alpha polypeptide) | 3073 | ENSG00000213614 |

  
  

| **Database:cellular component      &nbspName:lysosome      &nbspID:GO:0005764** | | | | | | |
| --- | --- | --- | --- | --- | --- | --- |
| C=345; O=6; E=1.57; R=3.82; rawP=0.0049; adjP=0.0107 | | | | | | |
| Index | UserID | Value | Gene Symbol | Gene Name | EntrezGene | Ensembl |
| 1 | P04040 | NA | CAT | catalase | 847 | ENSG00000121691 |
| 2 | O14773 | NA | TPP1 | tripeptidyl peptidase I | 1200 | ENSG00000166340 |
| 3 | P10253 | NA | GAA | glucosidase, alpha; acid | 2548 | ENSG00000171298 |
| 4 | P51688 | NA | SGSH | N-sulfoglucosamine sulfohydrolase | 6448 | ENSG00000181523 |
| 5 | P06865 | NA | HEXA | hexosaminidase A (alpha polypeptide) | 3073 | ENSG00000213614 |
| 6 | P38571 | NA | LIPA | lipase A, lysosomal acid, cholesterol esterase | 3988 | ENSG00000107798 |

  
  

| **Database:cellular component      &nbspName:pore complex      &nbspID:GO:0046930** | | | | | | |
| --- | --- | --- | --- | --- | --- | --- |
| C=81; O=3; E=0.37; R=8.13; rawP=0.0060; adjP=0.0128 | | | | | | |
| Index | UserID | Value | Gene Symbol | Gene Name | EntrezGene | Ensembl |
| 1 | Q9Y277 | NA | VDAC3 | voltage-dependent anion channel 3 | 7419 | ENSG00000078668 |
| 2 | Q14764 | NA | MVP | major vault protein | 9961 | ENSG00000013364 |
| 3 | P21796 | NA | VDAC1 | voltage-dependent anion channel 1 | 7416 | ENSG00000213585 |

  
  

| **Database:cellular component      &nbspName:melanosome      &nbspID:GO:0042470** | | | | | | |
| --- | --- | --- | --- | --- | --- | --- |
| C=93; O=3; E=0.42; R=7.08; rawP=0.0088; adjP=0.0178 | | | | | | |
| Index | UserID | Value | Gene Symbol | Gene Name | EntrezGene | Ensembl |
| 1 | O14773 | NA | TPP1 | tripeptidyl peptidase I | 1200 | ENSG00000166340 |
| 2 | Q14697 | NA | GANAB | glucosidase, alpha; neutral AB | 23193 | ENSG00000089597 |
| 3 | P40121 | NA | CAPG | capping protein (actin filament), gelsolin-like | 822 | ENSG00000042493 |

  
  

| **Database:cellular component      &nbspName:pigment granule      &nbspID:GO:0048770** | | | | | | |
| --- | --- | --- | --- | --- | --- | --- |
| C=93; O=3; E=0.42; R=7.08; rawP=0.0088; adjP=0.0178 | | | | | | |
| Index | UserID | Value | Gene Symbol | Gene Name | EntrezGene | Ensembl |
| 1 | O14773 | NA | TPP1 | tripeptidyl peptidase I | 1200 | ENSG00000166340 |
| 2 | Q14697 | NA | GANAB | glucosidase, alpha; neutral AB | 23193 | ENSG00000089597 |
| 3 | P40121 | NA | CAPG | capping protein (actin filament), gelsolin-like | 822 | ENSG00000042493 |

  
  
  
  

---

WebGestalt is currently developed and maintained by Jing Wang and Bing Zhang at the  Zhang Lab. Other people who have made significant contribution to the project include Dexter Duncan, Stefan Kirov, Zhiao Shi, and Jay Snoddy.  
  
**Funding credits:** NIH/NIAAA (U01 AA016662, U01 AA013512); NIH/NIDA (P01 DA015027); NIH/NIMH (P50 MH078028, P50 MH096972); NIH/NCI (U24 CA159988); NIH/NIGMS (R01 GM088822).
