## Supplementary material for "PGRMC1 phosphorylation and cell plasticity 1: glycolysis, mitochondria, tumor growth": File S8: final_sig_kegg_file_1439185596.html

Anchored HTML File of EIDs

|  |  |
| --- | --- |
|  | WEB-based GEne SeT AnaLysis Toolkit |
|  | ***Translating gene lists into biological insights...*** |

---

  

| **Database:KEGG pathway      &nbspName:Metabolic pathways      &nbspID:01100** | | | | | | |
| --- | --- | --- | --- | --- | --- | --- |
| C=1130; O=31; E=4.23; R=7.32; rawP=1.52e-19; adjP=3.19e-18 | | | | | | |
| Index | UserID | Value | Gene Symbol | Gene Name | EntrezGene | Ensembl |
| 1 | P10253 | NA | GAA | glucosidase, alpha; acid | 2548 | ENSG00000171298 |
| 2 | P54819 | NA | AK2 | adenylate kinase 2 | 204 | ENSG00000004455 |
| 3 | P21953 | NA | BCKDHB | branched chain keto acid dehydrogenase E1, beta polypeptide | 594 | ENSG00000083123 |
| 4 | P40939 | NA | HADHA | hydroxyacyl-CoA dehydrogenase/3-ketoacyl-CoA thiolase/enoyl-CoA hydratase (trifunctional protein), alpha subunit | 3030 | ENSG00000084754 |
| 5 | P40926 | NA | MDH2 | malate dehydrogenase 2, NAD (mitochondrial) | 4191 | ENSG00000146701 |
| 6 | P30084 | NA | ECHS1 | enoyl CoA hydratase, short chain, 1, mitochondrial | 1892 | ENSG00000127884 |
| 7 | Q13724 | NA | MOGS | mannosyl-oligosaccharide glucosidase | 7841 | ENSG00000115275 |
| 8 | Q99714 | NA | HSD17B10 | hydroxysteroid (17-beta) dehydrogenase 10 | 3028 | ENSG00000072506 |
| 9 | P00367 | NA | GLUD1 | glutamate dehydrogenase 1 | 2746 | ENSG00000148672 |
| 10 | P07741 | NA | APRT | adenine phosphoribosyltransferase | 353 | ENSG00000198931 |
| 11 | P48735 | NA | IDH2 | isocitrate dehydrogenase 2 (NADP+), mitochondrial | 3418 | ENSG00000182054 |
| 12 | P35270 | NA | SPR | sepiapterin reductase (7,8-dihydrobiopterin:NADP+ oxidoreductase) | 6697 | ENSG00000116096 |
| 13 | P31040 | NA | SDHA | succinate dehydrogenase complex, subunit A, flavoprotein (Fp) | 6389 | ENSG00000073578 |
| 14 | P32322 | NA | PYCR1 | pyrroline-5-carboxylate reductase 1 | 5831 | ENSG00000183010 |
| 15 | P04040 | NA | CAT | catalase | 847 | ENSG00000121691 |
| 16 | Q99798 | NA | ACO2 | aconitase 2, mitochondrial | 50 | ENSG00000100412 |
| 17 | P51649 | NA | ALDH5A1 | aldehyde dehydrogenase 5 family, member A1 | 7915 | ENSG00000112294 |
| 18 | P51688 | NA | SGSH | N-sulfoglucosamine sulfohydrolase | 6448 | ENSG00000181523 |
| 19 | P00558 | NA | PGK1 | phosphoglycerate kinase 1 | 5230 | ENSG00000102144 |
| 20 | P42765 | NA | ACAA2 | acetyl-CoA acyltransferase 2 | 10449 | ENSG00000167315 |
| 21 | P24752 | NA | ACAT1 | acetyl-CoA acetyltransferase 1 | 38 | ENSG00000075239 |
| 22 | P13674 | NA | P4HA1 | prolyl 4-hydroxylase, alpha polypeptide I | 5033 | ENSG00000122884 |
| 23 | P49419 | NA | ALDH7A1 | aldehyde dehydrogenase 7 family, member A1 | 501 | ENSG00000164904 |
| 24 | P19367 | NA | HK1 | hexokinase 1 | 3098 | ENSG00000156515 |
| 25 | P25705 | NA | ATP5A1 | ATP synthase, H+ transporting, mitochondrial F1 complex, alpha subunit 1, cardiac muscle | 498 | ENSG00000152234 |
| 26 | P49748 | NA | ACADVL | acyl-CoA dehydrogenase, very long chain | 37 | ENSG00000072778 |
| 27 | P06865 | NA | HEXA | hexosaminidase A (alpha polypeptide) | 3073 | ENSG00000213614 |
| 28 | Q14697 | NA | GANAB | glucosidase, alpha; neutral AB | 23193 | ENSG00000089597 |
| 29 | P48047 | NA | ATP5O | ATP synthase, H+ transporting, mitochondrial F1 complex, O subunit | 539 | ENSG00000241837 |
| 30 | P60174 | NA | TPI1 | triosephosphate isomerase 1 | 7167 | ENSG00000111669 |
| 31 | P31937 | NA | HIBADH | 3-hydroxyisobutyrate dehydrogenase | 11112 | ENSG00000106049 |

  
  

| **Database:KEGG pathway      &nbspName:Alzheimer's disease      &nbspID:05010** | | | | | | |
| --- | --- | --- | --- | --- | --- | --- |
| C=167; O=4; E=0.63; R=6.39; rawP=0.0036; adjP=0.0040 | | | | | | |
| Index | UserID | Value | Gene Symbol | Gene Name | EntrezGene | Ensembl |
| 1 | P25705 | NA | ATP5A1 | ATP synthase, H+ transporting, mitochondrial F1 complex, alpha subunit 1, cardiac muscle | 498 | ENSG00000152234 |
| 2 | P48047 | NA | ATP5O | ATP synthase, H+ transporting, mitochondrial F1 complex, O subunit | 539 | ENSG00000241837 |
| 3 | P31040 | NA | SDHA | succinate dehydrogenase complex, subunit A, flavoprotein (Fp) | 6389 | ENSG00000073578 |
| 4 | Q99714 | NA | HSD17B10 | hydroxysteroid (17-beta) dehydrogenase 10 | 3028 | ENSG00000072506 |

  
  

| **Database:KEGG pathway      &nbspName:Oxidative phosphorylation      &nbspID:00190** | | | | | | |
| --- | --- | --- | --- | --- | --- | --- |
| C=132; O=3; E=0.49; R=6.07; rawP=0.0134; adjP=0.0141 | | | | | | |
| Index | UserID | Value | Gene Symbol | Gene Name | EntrezGene | Ensembl |
| 1 | P25705 | NA | ATP5A1 | ATP synthase, H+ transporting, mitochondrial F1 complex, alpha subunit 1, cardiac muscle | 498 | ENSG00000152234 |
| 2 | P48047 | NA | ATP5O | ATP synthase, H+ transporting, mitochondrial F1 complex, O subunit | 539 | ENSG00000241837 |
| 3 | P31040 | NA | SDHA | succinate dehydrogenase complex, subunit A, flavoprotein (Fp) | 6389 | ENSG00000073578 |

  
  

| **Database:KEGG pathway      &nbspName:Calcium signaling pathway      &nbspID:04020** | | | | | | |
| --- | --- | --- | --- | --- | --- | --- |
| C=177; O=3; E=0.66; R=4.52; rawP=0.0289; adjP=0.0289 | | | | | | |
| Index | UserID | Value | Gene Symbol | Gene Name | EntrezGene | Ensembl |
| 1 | Q9Y277 | NA | VDAC3 | voltage-dependent anion channel 3 | 7419 | ENSG00000078668 |
| 2 | P12235 | NA | SLC25A4 | solute carrier family 25 (mitochondrial carrier; adenine nucleotide translocator), member 4 | 291 | ENSG00000151729 |
| 3 | P21796 | NA | VDAC1 | voltage-dependent anion channel 1 | 7416 | ENSG00000213585 |

  
  

  
  

---

WebGestalt is currently developed and maintained by Jing Wang and Bing Zhang at the  Zhang Lab. Other people who have made significant contribution to the project include Dexter Duncan, Stefan Kirov, Zhiao Shi, and Jay Snoddy.  
  
**Funding credits:** NIH/NIAAA (U01 AA016662, U01 AA013512); NIH/NIDA (P01 DA015027); NIH/NIMH (P50 MH078028, P50 MH096972); NIH/NCI (U24 CA159988); NIH/NIGMS (R01 GM088822).
