## Supplementary material for "PGRMC1 phosphorylation and cell plasticity 1: glycolysis, mitochondria, tumor growth": File S8: final_pc_geneset_file_1439165895.html

  
  

| **Database:Pathway Commons pathway      &nbspName:Gluconeogenesis      &nbspID:DB\_ID:929** | | | | | | |
| --- | --- | --- | --- | --- | --- | --- |
| C=20; O=4; E=0.07; R=53.38; rawP=8.42e-07; adjP=7.58e-06 | | | | | | |
| Index | UserID | Value | Gene Symbol | Gene Name | EntrezGene | Ensembl |
| 1 | Q02978 | NA | SLC25A11 | solute carrier family 25 (mitochondrial carrier; oxoglutarate carrier), member 11 | 8402 | ENSG00000108528 |
| 2 | P00558 | NA | PGK1 | phosphoglycerate kinase 1 | 5230 | ENSG00000102144 |
| 3 | P40926 | NA | MDH2 | malate dehydrogenase 2, NAD (mitochondrial) | 4191 | ENSG00000146701 |
| 4 | P60174 | NA | TPI1 | triosephosphate isomerase 1 | 7167 | ENSG00000111669 |

  
  

| **Database:Pathway Commons pathway      &nbspName:mitochondrial fatty acid beta-oxidation of saturated fatty acids      &nbspID:DB\_ID:911** | | | | | | |
| --- | --- | --- | --- | --- | --- | --- |
| C=8; O=3; E=0.03; R=100.08; rawP=2.79e-06; adjP=2.23e-05 | | | | | | |
| Index | UserID | Value | Gene Symbol | Gene Name | EntrezGene | Ensembl |
| 1 | P49748 | NA | ACADVL | acyl-CoA dehydrogenase, very long chain | 37 | ENSG00000072778 |
| 2 | P40939 | NA | HADHA | hydroxyacyl-CoA dehydrogenase/3-ketoacyl-CoA thiolase/enoyl-CoA hydratase (trifunctional protein), alpha subunit | 3030 | ENSG00000084754 |
| 3 | P30084 | NA | ECHS1 | enoyl CoA hydratase, short chain, 1, mitochondrial | 1892 | ENSG00000127884 |

  
  

| **Database:Pathway Commons pathway      &nbspName:Pyruvate metabolism and Citric Acid (TCA) cycle      &nbspID:DB\_ID:546** | | | | | | |
| --- | --- | --- | --- | --- | --- | --- |
| C=31; O=4; E=0.12; R=34.44; rawP=5.30e-06; adjP=3.82e-05 | | | | | | |
| Index | UserID | Value | Gene Symbol | Gene Name | EntrezGene | Ensembl |
| 1 | Q99798 | NA | ACO2 | aconitase 2, mitochondrial | 50 | ENSG00000100412 |
| 2 | P48735 | NA | IDH2 | isocitrate dehydrogenase 2 (NADP+), mitochondrial | 3418 | ENSG00000182054 |
| 3 | P31040 | NA | SDHA | succinate dehydrogenase complex, subunit A, flavoprotein (Fp) | 6389 | ENSG00000073578 |
| 4 | P40926 | NA | MDH2 | malate dehydrogenase 2, NAD (mitochondrial) | 4191 | ENSG00000146701 |

  
  

| **Database:Pathway Commons pathway      &nbspName:Glucose metabolism      &nbspID:DB\_ID:898** | | | | | | |
| --- | --- | --- | --- | --- | --- | --- |
| C=36; O=4; E=0.13; R=29.65; rawP=9.79e-06; adjP=6.41e-05 | | | | | | |
| Index | UserID | Value | Gene Symbol | Gene Name | EntrezGene | Ensembl |
| 1 | Q02978 | NA | SLC25A11 | solute carrier family 25 (mitochondrial carrier; oxoglutarate carrier), member 11 | 8402 | ENSG00000108528 |
| 2 | P00558 | NA | PGK1 | phosphoglycerate kinase 1 | 5230 | ENSG00000102144 |
| 3 | P40926 | NA | MDH2 | malate dehydrogenase 2, NAD (mitochondrial) | 4191 | ENSG00000146701 |
| 4 | P60174 | NA | TPI1 | triosephosphate isomerase 1 | 7167 | ENSG00000111669 |

  
  

| **Database:Pathway Commons pathway      &nbspName:TNF receptor signaling pathway      &nbspID:DB\_ID:1600** | | | | | | |
| --- | --- | --- | --- | --- | --- | --- |
| C=299; O=6; E=1.12; R=5.36; rawP=0.0009; adjP=0.0038 | | | | | | |
| Index | UserID | Value | Gene Symbol | Gene Name | EntrezGene | Ensembl |
| 1 | Q14980 | NA | NUMA1 | nuclear mitotic apparatus protein 1 | 4926 | ENSG00000137497 |
| 2 | P08727 | NA | KRT19 | keratin 19 | 3880 | ENSG00000171345 |
| 3 | P05783 | NA | KRT18 | keratin 18 | 3875 | ENSG00000111057 |
| 4 | P05787 | NA | KRT8 | keratin 8 | 3856 | ENSG00000170421 |
| 5 | Q13813 | NA | SPTAN1 | spectrin, alpha, non-erythrocytic 1 | 6709 | ENSG00000197694 |
| 6 | O95831 | NA | AIFM1 | apoptosis-inducing factor, mitochondrion-associated, 1 | 9131 | ENSG00000156709 |

  
  

| **Database:Pathway Commons pathway      &nbspName:Caspase cascade in apoptosis      &nbspID:DB\_ID:1501** | | | | | | |
| --- | --- | --- | --- | --- | --- | --- |
| C=52; O=3; E=0.19; R=15.40; rawP=0.0010; adjP=0.0040 | | | | | | |
| Index | UserID | Value | Gene Symbol | Gene Name | EntrezGene | Ensembl |
| 1 | Q14980 | NA | NUMA1 | nuclear mitotic apparatus protein 1 | 4926 | ENSG00000137497 |
| 2 | P05783 | NA | KRT18 | keratin 18 | 3875 | ENSG00000111057 |
| 3 | Q13813 | NA | SPTAN1 | spectrin, alpha, non-erythrocytic 1 | 6709 | ENSG00000197694 |

  
  

| **Database:Pathway Commons pathway      &nbspName:Metabolism of nucleotides      &nbspID:DB\_ID:639** | | | | | | |
| --- | --- | --- | --- | --- | --- | --- |
| C=58; O=3; E=0.22; R=13.80; rawP=0.0013; adjP=0.0049 | | | | | | |
| Index | UserID | Value | Gene Symbol | Gene Name | EntrezGene | Ensembl |
| 1 | P04040 | NA | CAT | catalase | 847 | ENSG00000121691 |
| 2 | P07741 | NA | APRT | adenine phosphoribosyltransferase | 353 | ENSG00000198931 |
| 3 | P54819 | NA | AK2 | adenylate kinase 2 | 204 | ENSG00000004455 |

  
  

| **Database:Pathway Commons pathway      &nbspName:FAS (CD95) signaling pathway      &nbspID:DB\_ID:1559** | | | | | | |
| --- | --- | --- | --- | --- | --- | --- |
| C=130; O=4; E=0.49; R=8.21; rawP=0.0014; adjP=0.0050 | | | | | | |
| Index | UserID | Value | Gene Symbol | Gene Name | EntrezGene | Ensembl |
| 1 | Q14980 | NA | NUMA1 | nuclear mitotic apparatus protein 1 | 4926 | ENSG00000137497 |
| 2 | P05783 | NA | KRT18 | keratin 18 | 3875 | ENSG00000111057 |
| 3 | Q13813 | NA | SPTAN1 | spectrin, alpha, non-erythrocytic 1 | 6709 | ENSG00000197694 |
| 4 | O95831 | NA | AIFM1 | apoptosis-inducing factor, mitochondrion-associated, 1 | 9131 | ENSG00000156709 |

  
  

| **Database:Pathway Commons pathway      &nbspName:Respiratory electron transport      &nbspID:DB\_ID:867** | | | | | | |
| --- | --- | --- | --- | --- | --- | --- |
| C=76; O=3; E=0.28; R=10.53; rawP=0.0029; adjP=0.0099 | | | | | | |
| Index | UserID | Value | Gene Symbol | Gene Name | EntrezGene | Ensembl |
| 1 | P31040 | NA | SDHA | succinate dehydrogenase complex, subunit A, flavoprotein (Fp) | 6389 | ENSG00000073578 |
| 2 | P38117 | NA | ETFB | electron-transfer-flavoprotein, beta polypeptide | 2109 | ENSG00000105379 |
| 3 | P13804 | NA | ETFA | electron-transfer-flavoprotein, alpha polypeptide | 2108 | ENSG00000140374 |

  
  

| **Database:Pathway Commons pathway      &nbspName:TRAIL signaling pathway      &nbspID:DB\_ID:1480** | | | | | | |
| --- | --- | --- | --- | --- | --- | --- |
| C=1328; O=12; E=4.98; R=2.41; rawP=0.0037; adjP=0.0121 | | | | | | |
| Index | UserID | Value | Gene Symbol | Gene Name | EntrezGene | Ensembl |
| 1 | Q14980 | NA | NUMA1 | nuclear mitotic apparatus protein 1 | 4926 | ENSG00000137497 |
| 2 | P04040 | NA | CAT | catalase | 847 | ENSG00000121691 |
| 3 | P00558 | NA | PGK1 | phosphoglycerate kinase 1 | 5230 | ENSG00000102144 |
| 4 | P30048 | NA | PRDX3 | peroxiredoxin 3 | 10935 | ENSG00000165672 |
| 5 | P05787 | NA | KRT8 | keratin 8 | 3856 | ENSG00000170421 |
| 6 | P04179 | NA | SOD2 | superoxide dismutase 2, mitochondrial | 6648 | ENSG00000112096 |
| 7 | O95831 | NA | AIFM1 | apoptosis-inducing factor, mitochondrion-associated, 1 | 9131 | ENSG00000156709 |
| 8 | O75367 | NA | H2AFY | H2A histone family, member Y | 9555 | ENSG00000113648 |
| 9 | P19367 | NA | HK1 | hexokinase 1 | 3098 | ENSG00000156515 |
| 10 | P08727 | NA | KRT19 | keratin 19 | 3880 | ENSG00000171345 |
| 11 | P05783 | NA | KRT18 | keratin 18 | 3875 | ENSG00000111057 |
| 12 | Q13813 | NA | SPTAN1 | spectrin, alpha, non-erythrocytic 1 | 6709 | ENSG00000197694 |

  
  

| **Database:Pathway Commons pathway      &nbspName:Host Interactions of HIV factors      &nbspID:DB\_ID:201** | | | | | | |
| --- | --- | --- | --- | --- | --- | --- |
| C=142; O=3; E=0.53; R=5.64; rawP=0.0163; adjP=0.0489 | | | | | | |
| Index | UserID | Value | Gene Symbol | Gene Name | EntrezGene | Ensembl |
| 1 | Q06323 | NA | PSME1 | proteasome (prosome, macropain) activator subunit 1 (PA28 alpha) | 5720 | ENSG00000092010 |
| 2 | P12235 | NA | SLC25A4 | solute carrier family 25 (mitochondrial carrier; adenine nucleotide translocator), member 4 | 291 | ENSG00000151729 |
| 3 | Q9UL46 | NA | PSME2 | proteasome (prosome, macropain) activator subunit 2 (PA28 beta) | 5721 | ENSG00000100911 |

  
  

| **Database:Pathway Commons pathway      &nbspName:Metabolism of lipids and lipoproteins      &nbspID:DB\_ID:984** | | | | | | |
| --- | --- | --- | --- | --- | --- | --- |
| C=258; O=4; E=0.97; R=4.14; rawP=0.0161; adjP=0.0489 | | | | | | |
| Index | UserID | Value | Gene Symbol | Gene Name | EntrezGene | Ensembl |
| 1 | P24752 | NA | ACAT1 | acetyl-CoA acetyltransferase 1 | 38 | ENSG00000075239 |
| 2 | P49748 | NA | ACADVL | acyl-CoA dehydrogenase, very long chain | 37 | ENSG00000072778 |
| 3 | P40939 | NA | HADHA | hydroxyacyl-CoA dehydrogenase/3-ketoacyl-CoA thiolase/enoyl-CoA hydratase (trifunctional protein), alpha subunit | 3030 | ENSG00000084754 |
| 4 | P30084 | NA | ECHS1 | enoyl CoA hydratase, short chain, 1, mitochondrial | 1892 | ENSG00000127884 |

  
  

| **Database:Pathway Commons pathway      &nbspName:Apoptosis      &nbspID:DB\_ID:1068** | | | | | | |
| --- | --- | --- | --- | --- | --- | --- |
| C=158; O=3; E=0.59; R=5.07; rawP=0.0215; adjP=0.0619 | | | | | | |
| Index | UserID | Value | Gene Symbol | Gene Name | EntrezGene | Ensembl |
| 1 | Q06323 | NA | PSME1 | proteasome (prosome, macropain) activator subunit 1 (PA28 alpha) | 5720 | ENSG00000092010 |
| 2 | Q9UL46 | NA | PSME2 | proteasome (prosome, macropain) activator subunit 2 (PA28 beta) | 5721 | ENSG00000100911 |
| 3 | Q13813 | NA | SPTAN1 | spectrin, alpha, non-erythrocytic 1 | 6709 | ENSG00000197694 |

  
  
  
  

---

WebGestalt is currently developed and maintained by Jing Wang and Bing Zhang at the  Zhang Lab. Other people who have made significant contribution to the project include Dexter Duncan, Stefan Kirov, Zhiao Shi, and Jay Snoddy.  
  
**Funding credits:** NIH/NIAAA (U01 AA016662, U01 AA013512); NIH/NIDA (P01 DA015027); NIH/NIMH (P50 MH078028, P50 MH096972); NIH/NCI (U24 CA159988); NIH/NIGMS (R01 GM088822).
