## Supplementary material for "PGRMC1 phosphorylation and cell plasticity 1: glycolysis, mitochondria, tumor growth": File S8: final_tf_geneset_file_1439185596.html

Anchored HTML File of EIDs

|  |  |
| --- | --- |
|  | WEB-based GEne SeT AnaLysis Toolkit |
|  | ***Translating gene lists into biological insights...*** |

---

  

| **Database:Transcrription Target      &nbspName:hsa\_TGACCTY\_V$ERR1\_Q2      &nbspID:DB\_ID:2414** | | | | | | |
| --- | --- | --- | --- | --- | --- | --- |
| C=1005; O=15; E=3.77; R=3.98; rawP=3.98e-06; adjP=0.0002 | | | | | | |
| Index | UserID | Value | Gene Symbol | Gene Name | EntrezGene | Ensembl |
| 1 | Q99798 | NA | ACO2 | aconitase 2, mitochondrial | 50 | ENSG00000100412 |
| 2 | P54819 | NA | AK2 | adenylate kinase 2 | 204 | ENSG00000004455 |
| 3 | P12235 | NA | SLC25A4 | solute carrier family 25 (mitochondrial carrier; adenine nucleotide translocator), member 4 | 291 | ENSG00000151729 |
| 4 | P00558 | NA | PGK1 | phosphoglycerate kinase 1 | 5230 | ENSG00000102144 |
| 5 | P35232 | NA | PHB | prohibitin | 5245 | ENSG00000167085 |
| 6 | O95831 | NA | AIFM1 | apoptosis-inducing factor, mitochondrion-associated, 1 | 9131 | ENSG00000156709 |
| 7 | P52815 | NA | MRPL12 | mitochondrial ribosomal protein L12 | 6182 | ENSG00000262814 |
| 8 | P24752 | NA | ACAT1 | acetyl-CoA acetyltransferase 1 | 38 | ENSG00000075239 |
| 9 | P00367 | NA | GLUD1 | glutamate dehydrogenase 1 | 2746 | ENSG00000148672 |
| 10 | P25705 | NA | ATP5A1 | ATP synthase, H+ transporting, mitochondrial F1 complex, alpha subunit 1, cardiac muscle | 498 | ENSG00000152234 |
| 11 | Q16531 | NA | DDB1 | damage-specific DNA binding protein 1, 127kDa | 1642 | ENSG00000167986 |
| 12 | P60174 | NA | TPI1 | triosephosphate isomerase 1 | 7167 | ENSG00000111669 |
| 13 | P31937 | NA | HIBADH | 3-hydroxyisobutyrate dehydrogenase | 11112 | ENSG00000106049 |
| 14 | Q13813 | NA | SPTAN1 | spectrin, alpha, non-erythrocytic 1 | 6709 | ENSG00000197694 |
| 15 | Q9BPW8 | NA | NIPSNAP1 | nipsnap homolog 1 (C. elegans) | 8508 | ENSG00000184117 |

  
  

| **Database:Transcrription Target      &nbspName:hsa\_V$ERR1\_Q2      &nbspID:DB\_ID:2154** | | | | | | |
| --- | --- | --- | --- | --- | --- | --- |
| C=254; O=6; E=0.95; R=6.30; rawP=0.0004; adjP=0.0076 | | | | | | |
| Index | UserID | Value | Gene Symbol | Gene Name | EntrezGene | Ensembl |
| 1 | Q99798 | NA | ACO2 | aconitase 2, mitochondrial | 50 | ENSG00000100412 |
| 2 | P25705 | NA | ATP5A1 | ATP synthase, H+ transporting, mitochondrial F1 complex, alpha subunit 1, cardiac muscle | 498 | ENSG00000152234 |
| 3 | P21796 | NA | VDAC1 | voltage-dependent anion channel 1 | 7416 | ENSG00000213585 |
| 4 | P21953 | NA | BCKDHB | branched chain keto acid dehydrogenase E1, beta polypeptide | 594 | ENSG00000083123 |
| 5 | P35232 | NA | PHB | prohibitin | 5245 | ENSG00000167085 |
| 6 | O95831 | NA | AIFM1 | apoptosis-inducing factor, mitochondrion-associated, 1 | 9131 | ENSG00000156709 |

  
  

| **Database:Transcrription Target      &nbspName:hsa\_V$PPAR\_DR1\_Q2      &nbspID:DB\_ID:2248** | | | | | | |
| --- | --- | --- | --- | --- | --- | --- |
| C=252; O=6; E=0.94; R=6.35; rawP=0.0004; adjP=0.0076 | | | | | | |
| Index | UserID | Value | Gene Symbol | Gene Name | EntrezGene | Ensembl |
| 1 | P49748 | NA | ACADVL | acyl-CoA dehydrogenase, very long chain | 37 | ENSG00000072778 |
| 2 | P21953 | NA | BCKDHB | branched chain keto acid dehydrogenase E1, beta polypeptide | 594 | ENSG00000083123 |
| 3 | Q99623 | NA | PHB2 | prohibitin 2 | 11331 | ENSG00000215021 |
| 4 | P35580 | NA | MYH10 | myosin, heavy chain 10, non-muscle | 4628 | ENSG00000133026 |
| 5 | P31937 | NA | HIBADH | 3-hydroxyisobutyrate dehydrogenase | 11112 | ENSG00000106049 |
| 6 | Q9UJZ1 | NA | STOML2 | stomatin (EPB72)-like 2 | 30968 | ENSG00000165283 |

  
  

| **Database:Transcrription Target      &nbspName:hsa\_TMTCGCGANR\_UNKNOWN      &nbspID:DB\_ID:2370** | | | | | | |
| --- | --- | --- | --- | --- | --- | --- |
| C=156; O=4; E=0.58; R=6.84; rawP=0.0028; adjP=0.0266 | | | | | | |
| Index | UserID | Value | Gene Symbol | Gene Name | EntrezGene | Ensembl |
| 1 | Q9Y277 | NA | VDAC3 | voltage-dependent anion channel 3 | 7419 | ENSG00000078668 |
| 2 | Q02978 | NA | SLC25A11 | solute carrier family 25 (mitochondrial carrier; oxoglutarate carrier), member 11 | 8402 | ENSG00000108528 |
| 3 | P12235 | NA | SLC25A4 | solute carrier family 25 (mitochondrial carrier; adenine nucleotide translocator), member 4 | 291 | ENSG00000151729 |
| 4 | P40926 | NA | MDH2 | malate dehydrogenase 2, NAD (mitochondrial) | 4191 | ENSG00000146701 |

  
  

| **Database:Transcrription Target      &nbspName:hsa\_V$COUP\_DR1\_Q6      &nbspID:DB\_ID:2250** | | | | | | |
| --- | --- | --- | --- | --- | --- | --- |
| C=240; O=5; E=0.90; R=5.56; rawP=0.0021; adjP=0.0266 | | | | | | |
| Index | UserID | Value | Gene Symbol | Gene Name | EntrezGene | Ensembl |
| 1 | Q99623 | NA | PHB2 | prohibitin 2 | 11331 | ENSG00000215021 |
| 2 | P35580 | NA | MYH10 | myosin, heavy chain 10, non-muscle | 4628 | ENSG00000133026 |
| 3 | P48047 | NA | ATP5O | ATP synthase, H+ transporting, mitochondrial F1 complex, O subunit | 539 | ENSG00000241837 |
| 4 | P31937 | NA | HIBADH | 3-hydroxyisobutyrate dehydrogenase | 11112 | ENSG00000106049 |
| 5 | Q9UJZ1 | NA | STOML2 | stomatin (EPB72)-like 2 | 30968 | ENSG00000165283 |

  
  

| **Database:Transcrription Target      &nbspName:hsa\_V$DR1\_Q3      &nbspID:DB\_ID:2246** | | | | | | |
| --- | --- | --- | --- | --- | --- | --- |
| C=247; O=5; E=0.93; R=5.40; rawP=0.0024; adjP=0.0266 | | | | | | |
| Index | UserID | Value | Gene Symbol | Gene Name | EntrezGene | Ensembl |
| 1 | Q99623 | NA | PHB2 | prohibitin 2 | 11331 | ENSG00000215021 |
| 2 | P35580 | NA | MYH10 | myosin, heavy chain 10, non-muscle | 4628 | ENSG00000133026 |
| 3 | P48047 | NA | ATP5O | ATP synthase, H+ transporting, mitochondrial F1 complex, O subunit | 539 | ENSG00000241837 |
| 4 | P31937 | NA | HIBADH | 3-hydroxyisobutyrate dehydrogenase | 11112 | ENSG00000106049 |
| 5 | Q9UJZ1 | NA | STOML2 | stomatin (EPB72)-like 2 | 30968 | ENSG00000165283 |

  
  

| **Database:Transcrription Target      &nbspName:hsa\_WCAANNNYCAG\_UNKNOWN      &nbspID:DB\_ID:2376** | | | | | | |
| --- | --- | --- | --- | --- | --- | --- |
| C=244; O=4; E=0.91; R=4.38; rawP=0.0133; adjP=0.0886 | | | | | | |
| Index | UserID | Value | Gene Symbol | Gene Name | EntrezGene | Ensembl |
| 1 | Q14980 | NA | NUMA1 | nuclear mitotic apparatus protein 1 | 4926 | ENSG00000137497 |
| 2 | P25705 | NA | ATP5A1 | ATP synthase, H+ transporting, mitochondrial F1 complex, alpha subunit 1, cardiac muscle | 498 | ENSG00000152234 |
| 3 | Q9BQ69 | NA | MACROD1 | MACRO domain containing 1 | 28992 | ENSG00000133315 |
| 4 | P48047 | NA | ATP5O | ATP synthase, H+ transporting, mitochondrial F1 complex, O subunit | 539 | ENSG00000241837 |

  
  

| **Database:Transcrription Target      &nbspName:hsa\_V$HNF4\_01      &nbspID:DB\_ID:1938** | | | | | | |
| --- | --- | --- | --- | --- | --- | --- |
| C=263; O=4; E=0.99; R=4.06; rawP=0.0171; adjP=0.0886 | | | | | | |
| Index | UserID | Value | Gene Symbol | Gene Name | EntrezGene | Ensembl |
| 1 | P12235 | NA | SLC25A4 | solute carrier family 25 (mitochondrial carrier; adenine nucleotide translocator), member 4 | 291 | ENSG00000151729 |
| 2 | P35580 | NA | MYH10 | myosin, heavy chain 10, non-muscle | 4628 | ENSG00000133026 |
| 3 | P60174 | NA | TPI1 | triosephosphate isomerase 1 | 7167 | ENSG00000111669 |
| 4 | Q9BPW8 | NA | NIPSNAP1 | nipsnap homolog 1 (C. elegans) | 8508 | ENSG00000184117 |

  
  

| **Database:Transcrription Target      &nbspName:hsa\_V$BACH1\_01      &nbspID:DB\_ID:2146** | | | | | | |
| --- | --- | --- | --- | --- | --- | --- |
| C=256; O=4; E=0.96; R=4.17; rawP=0.0157; adjP=0.0886 | | | | | | |
| Index | UserID | Value | Gene Symbol | Gene Name | EntrezGene | Ensembl |
| 1 | O14773 | NA | TPP1 | tripeptidyl peptidase I | 1200 | ENSG00000166340 |
| 2 | P08727 | NA | KRT19 | keratin 19 | 3880 | ENSG00000171345 |
| 3 | P49748 | NA | ACADVL | acyl-CoA dehydrogenase, very long chain | 37 | ENSG00000072778 |
| 4 | P05787 | NA | KRT8 | keratin 8 | 3856 | ENSG00000170421 |

  
  

| **Database:Transcrription Target      &nbspName:hsa\_V$ER\_Q6\_02      &nbspID:DB\_ID:2322** | | | | | | |
| --- | --- | --- | --- | --- | --- | --- |
| C=250; O=4; E=0.94; R=4.27; rawP=0.0145; adjP=0.0886 | | | | | | |
| Index | UserID | Value | Gene Symbol | Gene Name | EntrezGene | Ensembl |
| 1 | P00367 | NA | GLUD1 | glutamate dehydrogenase 1 | 2746 | ENSG00000148672 |
| 2 | P48047 | NA | ATP5O | ATP synthase, H+ transporting, mitochondrial F1 complex, O subunit | 539 | ENSG00000241837 |
| 3 | P31937 | NA | HIBADH | 3-hydroxyisobutyrate dehydrogenase | 11112 | ENSG00000106049 |
| 4 | Q9UJZ1 | NA | STOML2 | stomatin (EPB72)-like 2 | 30968 | ENSG00000165283 |

  
  

| **Database:Transcrription Target      &nbspName:hsa\_V$E2F1\_Q6\_01      &nbspID:DB\_ID:2314** | | | | | | |
| --- | --- | --- | --- | --- | --- | --- |
| C=232; O=4; E=0.87; R=4.60; rawP=0.0113; adjP=0.0886 | | | | | | |
| Index | UserID | Value | Gene Symbol | Gene Name | EntrezGene | Ensembl |
| 1 | Q14980 | NA | NUMA1 | nuclear mitotic apparatus protein 1 | 4926 | ENSG00000137497 |
| 2 | Q99798 | NA | ACO2 | aconitase 2, mitochondrial | 50 | ENSG00000100412 |
| 3 | P54819 | NA | AK2 | adenylate kinase 2 | 204 | ENSG00000004455 |
| 4 | P35580 | NA | MYH10 | myosin, heavy chain 10, non-muscle | 4628 | ENSG00000133026 |

  
  

| **Database:Transcrription Target      &nbspName:hsa\_TGANTCA\_V$AP1\_C      &nbspID:DB\_ID:2402** | | | | | | |
| --- | --- | --- | --- | --- | --- | --- |
| C=1092; O=9; E=4.09; R=2.20; rawP=0.0206; adjP=0.0979 | | | | | | |
| Index | UserID | Value | Gene Symbol | Gene Name | EntrezGene | Ensembl |
| 1 | O14773 | NA | TPP1 | tripeptidyl peptidase I | 1200 | ENSG00000166340 |
| 2 | P17931 | NA | LGALS3 | lectin, galactoside-binding, soluble, 3 | 3958 | ENSG00000131981 |
| 3 | P05787 | NA | KRT8 | keratin 8 | 3856 | ENSG00000170421 |
| 4 | P08727 | NA | KRT19 | keratin 19 | 3880 | ENSG00000171345 |
| 5 | P25705 | NA | ATP5A1 | ATP synthase, H+ transporting, mitochondrial F1 complex, alpha subunit 1, cardiac muscle | 498 | ENSG00000152234 |
| 6 | P49748 | NA | ACADVL | acyl-CoA dehydrogenase, very long chain | 37 | ENSG00000072778 |
| 7 | P35270 | NA | SPR | sepiapterin reductase (7,8-dihydrobiopterin:NADP+ oxidoreductase) | 6697 | ENSG00000116096 |
| 8 | P38117 | NA | ETFB | electron-transfer-flavoprotein, beta polypeptide | 2109 | ENSG00000105379 |
| 9 | Q9UJZ1 | NA | STOML2 | stomatin (EPB72)-like 2 | 30968 | ENSG00000165283 |

  
  
  
  

---

WebGestalt is currently developed and maintained by Jing Wang and Bing Zhang at the  Zhang Lab. Other people who have made significant contribution to the project include Dexter Duncan, Stefan Kirov, Zhiao Shi, and Jay Snoddy.  
  
**Funding credits:** NIH/NIAAA (U01 AA016662, U01 AA013512); NIH/NIDA (P01 DA015027); NIH/NIMH (P50 MH078028, P50 MH096972); NIH/NCI (U24 CA159988); NIH/NIGMS (R01 GM088822).
