## Supplementary material for "PGRMC1 phosphorylation and cell plasticity 1: glycolysis, mitochondria, tumor growth": File S8: final_wiki_geneset_file_1439185596.html

Anchored HTML File of EIDs

|  |  |
| --- | --- |
|  | WEB-based GEne SeT AnaLysis Toolkit |
|  | ***Translating gene lists into biological insights...*** |

---

  

| **Database:Wikipathways pathway      &nbspName:TCA Cycle      &nbspID:WP78** | | | | | | |
| --- | --- | --- | --- | --- | --- | --- |
| C=20; O=4; E=0.07; R=53.38; rawP=8.42e-07; adjP=7.58e-06 | | | | | | |
| Index | UserID | Value | Gene Symbol | Gene Name | EntrezGene | Ensembl |
| 1 | Q99798 | NA | ACO2 | aconitase 2, mitochondrial | 50 | ENSG00000100412 |
| 2 | P48735 | NA | IDH2 | isocitrate dehydrogenase 2 (NADP+), mitochondrial | 3418 | ENSG00000182054 |
| 3 | P31040 | NA | SDHA | succinate dehydrogenase complex, subunit A, flavoprotein (Fp) | 6389 | ENSG00000073578 |
| 4 | P40926 | NA | MDH2 | malate dehydrogenase 2, NAD (mitochondrial) | 4191 | ENSG00000146701 |

  
  

| **Database:Wikipathways pathway      &nbspName:Fatty Acid Beta Oxidation      &nbspID:WP143** | | | | | | |
| --- | --- | --- | --- | --- | --- | --- |
| C=73; O=5; E=0.27; R=18.28; rawP=7.96e-06; adjP=3.58e-05 | | | | | | |
| Index | UserID | Value | Gene Symbol | Gene Name | EntrezGene | Ensembl |
| 1 | P24752 | NA | ACAT1 | acetyl-CoA acetyltransferase 1 | 38 | ENSG00000075239 |
| 2 | P49748 | NA | ACADVL | acyl-CoA dehydrogenase, very long chain | 37 | ENSG00000072778 |
| 3 | P40939 | NA | HADHA | hydroxyacyl-CoA dehydrogenase/3-ketoacyl-CoA thiolase/enoyl-CoA hydratase (trifunctional protein), alpha subunit | 3030 | ENSG00000084754 |
| 4 | P60174 | NA | TPI1 | triosephosphate isomerase 1 | 7167 | ENSG00000111669 |
| 5 | P30084 | NA | ECHS1 | enoyl CoA hydratase, short chain, 1, mitochondrial | 1892 | ENSG00000127884 |

  
  

| **Database:Wikipathways pathway      &nbspName:Glycolysis and Gluconeogenesis      &nbspID:WP534** | | | | | | |
| --- | --- | --- | --- | --- | --- | --- |
| C=50; O=4; E=0.19; R=21.35; rawP=3.68e-05; adjP=0.0001 | | | | | | |
| Index | UserID | Value | Gene Symbol | Gene Name | EntrezGene | Ensembl |
| 1 | P19367 | NA | HK1 | hexokinase 1 | 3098 | ENSG00000156515 |
| 2 | P00558 | NA | PGK1 | phosphoglycerate kinase 1 | 5230 | ENSG00000102144 |
| 3 | P40926 | NA | MDH2 | malate dehydrogenase 2, NAD (mitochondrial) | 4191 | ENSG00000146701 |
| 4 | P60174 | NA | TPI1 | triosephosphate isomerase 1 | 7167 | ENSG00000111669 |

  
  

| **Database:Wikipathways pathway      &nbspName:Tryptophan metabolism      &nbspID:WP465** | | | | | | |
| --- | --- | --- | --- | --- | --- | --- |
| C=69; O=4; E=0.26; R=15.47; rawP=0.0001; adjP=0.0002 | | | | | | |
| Index | UserID | Value | Gene Symbol | Gene Name | EntrezGene | Ensembl |
| 1 | P24752 | NA | ACAT1 | acetyl-CoA acetyltransferase 1 | 38 | ENSG00000075239 |
| 2 | P04040 | NA | CAT | catalase | 847 | ENSG00000121691 |
| 3 | P30084 | NA | ECHS1 | enoyl CoA hydratase, short chain, 1, mitochondrial | 1892 | ENSG00000127884 |
| 4 | Q99714 | NA | HSD17B10 | hydroxysteroid (17-beta) dehydrogenase 10 | 3028 | ENSG00000072506 |

  
  

| **Database:Wikipathways pathway      &nbspName:Fatty Acid Biosynthesis      &nbspID:WP357** | | | | | | |
| --- | --- | --- | --- | --- | --- | --- |
| C=29; O=3; E=0.11; R=27.61; rawP=0.0002; adjP=0.0004 | | | | | | |
| Index | UserID | Value | Gene Symbol | Gene Name | EntrezGene | Ensembl |
| 1 | Q13011 | NA | ECH1 | enoyl CoA hydratase 1, peroxisomal | 1891 | ENSG00000104823 |
| 2 | P30084 | NA | ECHS1 | enoyl CoA hydratase, short chain, 1, mitochondrial | 1892 | ENSG00000127884 |
| 3 | P42765 | NA | ACAA2 | acetyl-CoA acyltransferase 2 | 10449 | ENSG00000167315 |

  
  

| **Database:Wikipathways pathway      &nbspName:Electron Transport Chain      &nbspID:WP111** | | | | | | |
| --- | --- | --- | --- | --- | --- | --- |
| C=103; O=4; E=0.39; R=10.36; rawP=0.0006; adjP=0.0009 | | | | | | |
| Index | UserID | Value | Gene Symbol | Gene Name | EntrezGene | Ensembl |
| 1 | P25705 | NA | ATP5A1 | ATP synthase, H+ transporting, mitochondrial F1 complex, alpha subunit 1, cardiac muscle | 498 | ENSG00000152234 |
| 2 | P12235 | NA | SLC25A4 | solute carrier family 25 (mitochondrial carrier; adenine nucleotide translocator), member 4 | 291 | ENSG00000151729 |
| 3 | P48047 | NA | ATP5O | ATP synthase, H+ transporting, mitochondrial F1 complex, O subunit | 539 | ENSG00000241837 |
| 4 | P31040 | NA | SDHA | succinate dehydrogenase complex, subunit A, flavoprotein (Fp) | 6389 | ENSG00000073578 |

  
  

| **Database:Wikipathways pathway      &nbspName:Selenium Pathway      &nbspID:WP15** | | | | | | |
| --- | --- | --- | --- | --- | --- | --- |
| C=107; O=3; E=0.40; R=7.48; rawP=0.0076; adjP=0.0098 | | | | | | |
| Index | UserID | Value | Gene Symbol | Gene Name | EntrezGene | Ensembl |
| 1 | P04040 | NA | CAT | catalase | 847 | ENSG00000121691 |
| 2 | P30048 | NA | PRDX3 | peroxiredoxin 3 | 10935 | ENSG00000165672 |
| 3 | P04179 | NA | SOD2 | superoxide dismutase 2, mitochondrial | 6648 | ENSG00000112096 |

  
  
  
  

---

WebGestalt is currently developed and maintained by Jing Wang and Bing Zhang at the  Zhang Lab. Other people who have made significant contribution to the project include Dexter Duncan, Stefan Kirov, Zhiao Shi, and Jay Snoddy.  
  
**Funding credits:** NIH/NIAAA (U01 AA016662, U01 AA013512); NIH/NIDA (P01 DA015027); NIH/NIMH (P50 MH078028, P50 MH096972); NIH/NCI (U24 CA159988); NIH/NIGMS (R01 GM088822).
