## Supplementary material for "PGRMC1 phosphorylation and cell plasticity 1: glycolysis, mitochondria, tumor growth": File S8: final_sig_file_1439165549.html

Anchored HTML File of EIDs

|  |  |
| --- | --- |
|  | WEB-based GEne SeT AnaLysis Toolkit |
|  | ***Translating gene lists into biological insights...*** |

---

  

| **Database:biological process      &nbspName:translational elongation      &nbspID:GO:0006414** | | | | | | |
| --- | --- | --- | --- | --- | --- | --- |
| C=108; O=5; E=0.24; R=20.87; rawP=3.68e-06; adjP=0.0009 | | | | | | |
| Index | UserID | Value | Gene Symbol | Gene Name | EntrezGene | Ensembl |
| 1 | P61313 | NA | RPL15 | ribosomal protein L15 | 6138 | ENSG00000174748 |
| 2 | P13639 | NA | EEF2 | eukaryotic translation elongation factor 2 | 1938 | ENSG00000167658 |
| 3 | P68104 | NA | EEF1A1 | eukaryotic translation elongation factor 1 alpha 1 | 1915 | ENSG00000156508 |
| 4 | P29692 | NA | EEF1D | eukaryotic translation elongation factor 1 delta (guanine nucleotide exchange protein) | 1936 | ENSG00000104529 |
| 5 | P24534 | NA | EEF1B2 | eukaryotic translation elongation factor 1 beta 2 | 1933 | ENSG00000114942 |

  
  

| **Database:biological process      &nbspName:response to drug      &nbspID:GO:0042493** | | | | | | |
| --- | --- | --- | --- | --- | --- | --- |
| C=371; O=6; E=0.82; R=7.29; rawP=0.0001; adjP=0.0129 | | | | | | |
| Index | UserID | Value | Gene Symbol | Gene Name | EntrezGene | Ensembl |
| 1 | P00491 | NA | PNP | purine nucleoside phosphorylase | 4860 | ENSG00000198805 |
| 2 | P47895 | NA | ALDH1A3 | aldehyde dehydrogenase 1 family, member A3 | 220 | ENSG00000184254 |
| 3 | P05556 | NA | ITGB1 | integrin, beta 1 (fibronectin receptor, beta polypeptide, antigen CD29 includes MDF2, MSK12) | 3688 | ENSG00000150093 |
| 4 | P04083 | NA | ANXA1 | annexin A1 | 301 | ENSG00000135046 |
| 5 | P09382 | NA | LGALS1 | lectin, galactoside-binding, soluble, 1 | 3956 | ENSG00000100097 |
| 6 | P11387 | NA | TOP1 | topoisomerase (DNA) I | 7150 | ENSG00000198900 |

  
  

| **Database:biological process      &nbspName:translation      &nbspID:GO:0006412** | | | | | | |
| --- | --- | --- | --- | --- | --- | --- |
| C=832; O=8; E=1.85; R=4.33; rawP=0.0004; adjP=0.0344 | | | | | | |
| Index | UserID | Value | Gene Symbol | Gene Name | EntrezGene | Ensembl |
| 1 | P61313 | NA | RPL15 | ribosomal protein L15 | 6138 | ENSG00000174748 |
| 2 | P49589 | NA | CARS | cysteinyl-tRNA synthetase | 833 | ENSG00000110619 |
| 3 | P60842 | NA | EIF4A1 | eukaryotic translation initiation factor 4A1 | 1973 | ENSG00000161960 |
| 4 | P13639 | NA | EEF2 | eukaryotic translation elongation factor 2 | 1938 | ENSG00000167658 |
| 5 | P41091 | NA | EIF2S3 | eukaryotic translation initiation factor 2, subunit 3 gamma, 52kDa | 1968 | ENSG00000130741 |
| 6 | P68104 | NA | EEF1A1 | eukaryotic translation elongation factor 1 alpha 1 | 1915 | ENSG00000156508 |
| 7 | P29692 | NA | EEF1D | eukaryotic translation elongation factor 1 delta (guanine nucleotide exchange protein) | 1936 | ENSG00000104529 |
| 8 | P24534 | NA | EEF1B2 | eukaryotic translation elongation factor 1 beta 2 | 1933 | ENSG00000114942 |

  
  

| **Database:biological process      &nbspName:response to chemical stimulus      &nbspID:GO:0042221** | | | | | | |
| --- | --- | --- | --- | --- | --- | --- |
| C=2741; O=14; E=6.08; R=2.30; rawP=0.0012; adjP=0.0602 | | | | | | |
| Index | UserID | Value | Gene Symbol | Gene Name | EntrezGene | Ensembl |
| 1 | Q9NR30 | NA | DDX21 | DEAD (Asp-Glu-Ala-Asp) box helicase 21 | 9188 | ENSG00000165732 |
| 2 | P22102 | NA | GART | phosphoribosylglycinamide formyltransferase, phosphoribosylglycinamide synthetase, phosphoribosylaminoimidazole synthetase | 2618 | ENSG00000159131 |
| 3 | P00491 | NA | PNP | purine nucleoside phosphorylase | 4860 | ENSG00000198805 |
| 4 | P02768 | NA | ALB | albumin | 213 | ENSG00000163631 |
| 5 | P47895 | NA | ALDH1A3 | aldehyde dehydrogenase 1 family, member A3 | 220 | ENSG00000184254 |
| 6 | P05556 | NA | ITGB1 | integrin, beta 1 (fibronectin receptor, beta polypeptide, antigen CD29 includes MDF2, MSK12) | 3688 | ENSG00000150093 |
| 7 | P09382 | NA | LGALS1 | lectin, galactoside-binding, soluble, 1 | 3956 | ENSG00000100097 |
| 8 | P60842 | NA | EIF4A1 | eukaryotic translation initiation factor 4A1 | 1973 | ENSG00000161960 |
| 9 | P17174 | NA | GOT1 | glutamic-oxaloacetic transaminase 1, soluble (aspartate aminotransferase 1) | 2805 | ENSG00000120053 |
| 10 | P00352 | NA | ALDH1A1 | aldehyde dehydrogenase 1 family, member A1 | 216 | ENSG00000165092 |
| 11 | P02786 | NA | TFRC | transferrin receptor (p90, CD71) | 7037 | ENSG00000072274 |
| 12 | O75369 | NA | FLNB | filamin B, beta | 2317 | ENSG00000136068 |
| 13 | P04083 | NA | ANXA1 | annexin A1 | 301 | ENSG00000135046 |
| 14 | P11387 | NA | TOP1 | topoisomerase (DNA) I | 7150 | ENSG00000198900 |

  
  

| **Database:biological process      &nbspName:organic acid transport      &nbspID:GO:0015849** | | | | | | |
| --- | --- | --- | --- | --- | --- | --- |
| C=221; O=4; E=0.49; R=8.16; rawP=0.0014; adjP=0.0602 | | | | | | |
| Index | UserID | Value | Gene Symbol | Gene Name | EntrezGene | Ensembl |
| 1 | P08195 | NA | SLC3A2 | solute carrier family 3 (activators of dibasic and neutral amino acid transport), member 2 | 6520 | ENSG00000168003 |
| 2 | P02768 | NA | ALB | albumin | 213 | ENSG00000163631 |
| 3 | P04083 | NA | ANXA1 | annexin A1 | 301 | ENSG00000135046 |
| 4 | Q96QD8 | NA | SLC38A2 | solute carrier family 38, member 2 | 54407 | ENSG00000134294 |

  
  

| **Database:biological process      &nbspName:carboxylic acid transport      &nbspID:GO:0046942** | | | | | | |
| --- | --- | --- | --- | --- | --- | --- |
| C=219; O=4; E=0.49; R=8.23; rawP=0.0013; adjP=0.0602 | | | | | | |
| Index | UserID | Value | Gene Symbol | Gene Name | EntrezGene | Ensembl |
| 1 | P08195 | NA | SLC3A2 | solute carrier family 3 (activators of dibasic and neutral amino acid transport), member 2 | 6520 | ENSG00000168003 |
| 2 | P02768 | NA | ALB | albumin | 213 | ENSG00000163631 |
| 3 | P04083 | NA | ANXA1 | annexin A1 | 301 | ENSG00000135046 |
| 4 | Q96QD8 | NA | SLC38A2 | solute carrier family 38, member 2 | 54407 | ENSG00000134294 |

  
  

| **Database:biological process      &nbspName:regulation of biological quality      &nbspID:GO:0065008** | | | | | | |
| --- | --- | --- | --- | --- | --- | --- |
| C=2507; O=13; E=5.56; R=2.34; rawP=0.0017; adjP=0.0627 | | | | | | |
| Index | UserID | Value | Gene Symbol | Gene Name | EntrezGene | Ensembl |
| 1 | P08195 | NA | SLC3A2 | solute carrier family 3 (activators of dibasic and neutral amino acid transport), member 2 | 6520 | ENSG00000168003 |
| 2 | P02768 | NA | ALB | albumin | 213 | ENSG00000163631 |
| 3 | P47895 | NA | ALDH1A3 | aldehyde dehydrogenase 1 family, member A3 | 220 | ENSG00000184254 |
| 4 | P05556 | NA | ITGB1 | integrin, beta 1 (fibronectin receptor, beta polypeptide, antigen CD29 includes MDF2, MSK12) | 3688 | ENSG00000150093 |
| 5 | P09382 | NA | LGALS1 | lectin, galactoside-binding, soluble, 1 | 3956 | ENSG00000100097 |
| 6 | Q96QD8 | NA | SLC38A2 | solute carrier family 38, member 2 | 54407 | ENSG00000134294 |
| 7 | P67809 | NA | YBX1 | Y box binding protein 1 | 4904 | ENSG00000065978 |
| 8 | P17174 | NA | GOT1 | glutamic-oxaloacetic transaminase 1, soluble (aspartate aminotransferase 1) | 2805 | ENSG00000120053 |
| 9 | P00352 | NA | ALDH1A1 | aldehyde dehydrogenase 1 family, member A1 | 216 | ENSG00000165092 |
| 10 | P18206 | NA | VCL | vinculin | 7414 | ENSG00000035403 |
| 11 | P02786 | NA | TFRC | transferrin receptor (p90, CD71) | 7037 | ENSG00000072274 |
| 12 | O75369 | NA | FLNB | filamin B, beta | 2317 | ENSG00000136068 |
| 13 | P04083 | NA | ANXA1 | annexin A1 | 301 | ENSG00000135046 |

  
  

| **Database:biological process      &nbspName:interaction with host      &nbspID:GO:0051701** | | | | | | |
| --- | --- | --- | --- | --- | --- | --- |
| C=411; O=5; E=0.91; R=5.48; rawP=0.0020; adjP=0.0645 | | | | | | |
| Index | UserID | Value | Gene Symbol | Gene Name | EntrezGene | Ensembl |
| 1 | P60842 | NA | EIF4A1 | eukaryotic translation initiation factor 4A1 | 1973 | ENSG00000161960 |
| 2 | P02768 | NA | ALB | albumin | 213 | ENSG00000163631 |
| 3 | P02786 | NA | TFRC | transferrin receptor (p90, CD71) | 7037 | ENSG00000072274 |
| 4 | P05556 | NA | ITGB1 | integrin, beta 1 (fibronectin receptor, beta polypeptide, antigen CD29 includes MDF2, MSK12) | 3688 | ENSG00000150093 |
| 5 | P11387 | NA | TOP1 | topoisomerase (DNA) I | 7150 | ENSG00000198900 |

  
  

| **Database:biological process      &nbspName:organic anion transport      &nbspID:GO:0015711** | | | | | | |
| --- | --- | --- | --- | --- | --- | --- |
| C=264; O=4; E=0.59; R=6.83; rawP=0.0026; adjP=0.0657 | | | | | | |
| Index | UserID | Value | Gene Symbol | Gene Name | EntrezGene | Ensembl |
| 1 | P08195 | NA | SLC3A2 | solute carrier family 3 (activators of dibasic and neutral amino acid transport), member 2 | 6520 | ENSG00000168003 |
| 2 | P02768 | NA | ALB | albumin | 213 | ENSG00000163631 |
| 3 | P04083 | NA | ANXA1 | annexin A1 | 301 | ENSG00000135046 |
| 4 | Q96QD8 | NA | SLC38A2 | solute carrier family 38, member 2 | 54407 | ENSG00000134294 |

  
  

| **Database:biological process      &nbspName:symbiosis, encompassing mutualism through parasitism      &nbspID:GO:0044403** | | | | | | |
| --- | --- | --- | --- | --- | --- | --- |
| C=447; O=5; E=0.99; R=5.04; rawP=0.0028; adjP=0.0657 | | | | | | |
| Index | UserID | Value | Gene Symbol | Gene Name | EntrezGene | Ensembl |
| 1 | P60842 | NA | EIF4A1 | eukaryotic translation initiation factor 4A1 | 1973 | ENSG00000161960 |
| 2 | P02768 | NA | ALB | albumin | 213 | ENSG00000163631 |
| 3 | P02786 | NA | TFRC | transferrin receptor (p90, CD71) | 7037 | ENSG00000072274 |
| 4 | P05556 | NA | ITGB1 | integrin, beta 1 (fibronectin receptor, beta polypeptide, antigen CD29 includes MDF2, MSK12) | 3688 | ENSG00000150093 |
| 5 | P11387 | NA | TOP1 | topoisomerase (DNA) I | 7150 | ENSG00000198900 |

  
  

| **Database:biological process      &nbspName:interspecies interaction between organisms      &nbspID:GO:0044419** | | | | | | |
| --- | --- | --- | --- | --- | --- | --- |
| C=447; O=5; E=0.99; R=5.04; rawP=0.0028; adjP=0.0657 | | | | | | |
| Index | UserID | Value | Gene Symbol | Gene Name | EntrezGene | Ensembl |
| 1 | P60842 | NA | EIF4A1 | eukaryotic translation initiation factor 4A1 | 1973 | ENSG00000161960 |
| 2 | P02768 | NA | ALB | albumin | 213 | ENSG00000163631 |
| 3 | P02786 | NA | TFRC | transferrin receptor (p90, CD71) | 7037 | ENSG00000072274 |
| 4 | P05556 | NA | ITGB1 | integrin, beta 1 (fibronectin receptor, beta polypeptide, antigen CD29 includes MDF2, MSK12) | 3688 | ENSG00000150093 |
| 5 | P11387 | NA | TOP1 | topoisomerase (DNA) I | 7150 | ENSG00000198900 |

  
  

| **Database:biological process      &nbspName:response to organic substance      &nbspID:GO:0010033** | | | | | | |
| --- | --- | --- | --- | --- | --- | --- |
| C=1814; O=10; E=4.02; R=2.48; rawP=0.0045; adjP=0.0913 | | | | | | |
| Index | UserID | Value | Gene Symbol | Gene Name | EntrezGene | Ensembl |
| 1 | Q9NR30 | NA | DDX21 | DEAD (Asp-Glu-Ala-Asp) box helicase 21 | 9188 | ENSG00000165732 |
| 2 | P22102 | NA | GART | phosphoribosylglycinamide formyltransferase, phosphoribosylglycinamide synthetase, phosphoribosylaminoimidazole synthetase | 2618 | ENSG00000159131 |
| 3 | P02768 | NA | ALB | albumin | 213 | ENSG00000163631 |
| 4 | P05556 | NA | ITGB1 | integrin, beta 1 (fibronectin receptor, beta polypeptide, antigen CD29 includes MDF2, MSK12) | 3688 | ENSG00000150093 |
| 5 | P09382 | NA | LGALS1 | lectin, galactoside-binding, soluble, 1 | 3956 | ENSG00000100097 |
| 6 | P17174 | NA | GOT1 | glutamic-oxaloacetic transaminase 1, soluble (aspartate aminotransferase 1) | 2805 | ENSG00000120053 |
| 7 | P60842 | NA | EIF4A1 | eukaryotic translation initiation factor 4A1 | 1973 | ENSG00000161960 |
| 8 | P02786 | NA | TFRC | transferrin receptor (p90, CD71) | 7037 | ENSG00000072274 |
| 9 | O75369 | NA | FLNB | filamin B, beta | 2317 | ENSG00000136068 |
| 10 | P04083 | NA | ANXA1 | annexin A1 | 301 | ENSG00000135046 |

  
  

| **Database:biological process      &nbspName:translational initiation      &nbspID:GO:0006413** | | | | | | |
| --- | --- | --- | --- | --- | --- | --- |
| C=152; O=3; E=0.34; R=8.90; rawP=0.0046; adjP=0.0913 | | | | | | |
| Index | UserID | Value | Gene Symbol | Gene Name | EntrezGene | Ensembl |
| 1 | P61313 | NA | RPL15 | ribosomal protein L15 | 6138 | ENSG00000174748 |
| 2 | P60842 | NA | EIF4A1 | eukaryotic translation initiation factor 4A1 | 1973 | ENSG00000161960 |
| 3 | P41091 | NA | EIF2S3 | eukaryotic translation initiation factor 2, subunit 3 gamma, 52kDa | 1968 | ENSG00000130741 |

  
  

| **Database:biological process      &nbspName:monocarboxylic acid biosynthetic process      &nbspID:GO:0072330** | | | | | | |
| --- | --- | --- | --- | --- | --- | --- |
| C=165; O=3; E=0.37; R=8.19; rawP=0.0057; adjP=0.0919 | | | | | | |
| Index | UserID | Value | Gene Symbol | Gene Name | EntrezGene | Ensembl |
| 1 | P17174 | NA | GOT1 | glutamic-oxaloacetic transaminase 1, soluble (aspartate aminotransferase 1) | 2805 | ENSG00000120053 |
| 2 | P47895 | NA | ALDH1A3 | aldehyde dehydrogenase 1 family, member A3 | 220 | ENSG00000184254 |
| 3 | P04083 | NA | ANXA1 | annexin A1 | 301 | ENSG00000135046 |

  
  

| **Database:biological process      &nbspName:leukocyte differentiation      &nbspID:GO:0002521** | | | | | | |
| --- | --- | --- | --- | --- | --- | --- |
| C=324; O=4; E=0.72; R=5.56; rawP=0.0055; adjP=0.0919 | | | | | | |
| Index | UserID | Value | Gene Symbol | Gene Name | EntrezGene | Ensembl |
| 1 | P00491 | NA | PNP | purine nucleoside phosphorylase | 4860 | ENSG00000198805 |
| 2 | P02786 | NA | TFRC | transferrin receptor (p90, CD71) | 7037 | ENSG00000072274 |
| 3 | P05556 | NA | ITGB1 | integrin, beta 1 (fibronectin receptor, beta polypeptide, antigen CD29 includes MDF2, MSK12) | 3688 | ENSG00000150093 |
| 4 | P04083 | NA | ANXA1 | annexin A1 | 301 | ENSG00000135046 |

  
  

| **Database:biological process      &nbspName:anion transport      &nbspID:GO:0006820** | | | | | | |
| --- | --- | --- | --- | --- | --- | --- |
| C=315; O=4; E=0.70; R=5.72; rawP=0.0050; adjP=0.0919 | | | | | | |
| Index | UserID | Value | Gene Symbol | Gene Name | EntrezGene | Ensembl |
| 1 | P08195 | NA | SLC3A2 | solute carrier family 3 (activators of dibasic and neutral amino acid transport), member 2 | 6520 | ENSG00000168003 |
| 2 | P02768 | NA | ALB | albumin | 213 | ENSG00000163631 |
| 3 | P04083 | NA | ANXA1 | annexin A1 | 301 | ENSG00000135046 |
| 4 | Q96QD8 | NA | SLC38A2 | solute carrier family 38, member 2 | 54407 | ENSG00000134294 |

  
  

| **Database:biological process      &nbspName:viral reproductive process      &nbspID:GO:0022415** | | | | | | |
| --- | --- | --- | --- | --- | --- | --- |
| C=540; O=5; E=1.20; R=4.17; rawP=0.0063; adjP=0.0956 | | | | | | |
| Index | UserID | Value | Gene Symbol | Gene Name | EntrezGene | Ensembl |
| 1 | P61313 | NA | RPL15 | ribosomal protein L15 | 6138 | ENSG00000174748 |
| 2 | P60842 | NA | EIF4A1 | eukaryotic translation initiation factor 4A1 | 1973 | ENSG00000161960 |
| 3 | P02786 | NA | TFRC | transferrin receptor (p90, CD71) | 7037 | ENSG00000072274 |
| 4 | P05556 | NA | ITGB1 | integrin, beta 1 (fibronectin receptor, beta polypeptide, antigen CD29 includes MDF2, MSK12) | 3688 | ENSG00000150093 |
| 5 | P11387 | NA | TOP1 | topoisomerase (DNA) I | 7150 | ENSG00000198900 |

  
  

| **Database:molecular function      &nbspName:translation factor activity, nucleic acid binding      &nbspID:GO:0008135** | | | | | | |
| --- | --- | --- | --- | --- | --- | --- |
| C=79; O=6; E=0.17; R=35.55; rawP=1.44e-08; adjP=7.63e-07 | | | | | | |
| Index | UserID | Value | Gene Symbol | Gene Name | EntrezGene | Ensembl |
| 1 | P60842 | NA | EIF4A1 | eukaryotic translation initiation factor 4A1 | 1973 | ENSG00000161960 |
| 2 | P13639 | NA | EEF2 | eukaryotic translation elongation factor 2 | 1938 | ENSG00000167658 |
| 3 | P41091 | NA | EIF2S3 | eukaryotic translation initiation factor 2, subunit 3 gamma, 52kDa | 1968 | ENSG00000130741 |
| 4 | P68104 | NA | EEF1A1 | eukaryotic translation elongation factor 1 alpha 1 | 1915 | ENSG00000156508 |
| 5 | P29692 | NA | EEF1D | eukaryotic translation elongation factor 1 delta (guanine nucleotide exchange protein) | 1936 | ENSG00000104529 |
| 6 | P24534 | NA | EEF1B2 | eukaryotic translation elongation factor 1 beta 2 | 1933 | ENSG00000114942 |

  
  

| **Database:molecular function      &nbspName:translation elongation factor activity      &nbspID:GO:0003746** | | | | | | |
| --- | --- | --- | --- | --- | --- | --- |
| C=17; O=4; E=0.04; R=110.12; rawP=4.01e-08; adjP=1.06e-06 | | | | | | |
| Index | UserID | Value | Gene Symbol | Gene Name | EntrezGene | Ensembl |
| 1 | P13639 | NA | EEF2 | eukaryotic translation elongation factor 2 | 1938 | ENSG00000167658 |
| 2 | P68104 | NA | EEF1A1 | eukaryotic translation elongation factor 1 alpha 1 | 1915 | ENSG00000156508 |
| 3 | P29692 | NA | EEF1D | eukaryotic translation elongation factor 1 delta (guanine nucleotide exchange protein) | 1936 | ENSG00000104529 |
| 4 | P24534 | NA | EEF1B2 | eukaryotic translation elongation factor 1 beta 2 | 1933 | ENSG00000114942 |

  
  

| **Database:molecular function      &nbspName:RNA binding      &nbspID:GO:0003723** | | | | | | |
| --- | --- | --- | --- | --- | --- | --- |
| C=843; O=11; E=1.80; R=6.11; rawP=7.29e-07; adjP=1.29e-05 | | | | | | |
| Index | UserID | Value | Gene Symbol | Gene Name | EntrezGene | Ensembl |
| 1 | P49589 | NA | CARS | cysteinyl-tRNA synthetase | 833 | ENSG00000110619 |
| 2 | Q9NR30 | NA | DDX21 | DEAD (Asp-Glu-Ala-Asp) box helicase 21 | 9188 | ENSG00000165732 |
| 3 | P13639 | NA | EEF2 | eukaryotic translation elongation factor 2 | 1938 | ENSG00000167658 |
| 4 | P41091 | NA | EIF2S3 | eukaryotic translation initiation factor 2, subunit 3 gamma, 52kDa | 1968 | ENSG00000130741 |
| 5 | P61313 | NA | RPL15 | ribosomal protein L15 | 6138 | ENSG00000174748 |
| 6 | P60842 | NA | EIF4A1 | eukaryotic translation initiation factor 4A1 | 1973 | ENSG00000161960 |
| 7 | P67809 | NA | YBX1 | Y box binding protein 1 | 4904 | ENSG00000065978 |
| 8 | P68104 | NA | EEF1A1 | eukaryotic translation elongation factor 1 alpha 1 | 1915 | ENSG00000156508 |
| 9 | P29692 | NA | EEF1D | eukaryotic translation elongation factor 1 delta (guanine nucleotide exchange protein) | 1936 | ENSG00000104529 |
| 10 | P04083 | NA | ANXA1 | annexin A1 | 301 | ENSG00000135046 |
| 11 | P24534 | NA | EEF1B2 | eukaryotic translation elongation factor 1 beta 2 | 1933 | ENSG00000114942 |

  
  

| **Database:molecular function      &nbspName:anion binding      &nbspID:GO:0043168** | | | | | | |
| --- | --- | --- | --- | --- | --- | --- |
| C=2371; O=15; E=5.07; R=2.96; rawP=3.60e-05; adjP=0.0005 | | | | | | |
| Index | UserID | Value | Gene Symbol | Gene Name | EntrezGene | Ensembl |
| 1 | P49589 | NA | CARS | cysteinyl-tRNA synthetase | 833 | ENSG00000110619 |
| 2 | Q9NR30 | NA | DDX21 | DEAD (Asp-Glu-Ala-Asp) box helicase 21 | 9188 | ENSG00000165732 |
| 3 | P22102 | NA | GART | phosphoribosylglycinamide formyltransferase, phosphoribosylglycinamide synthetase, phosphoribosylaminoimidazole synthetase | 2618 | ENSG00000159131 |
| 4 | P00491 | NA | PNP | purine nucleoside phosphorylase | 4860 | ENSG00000198805 |
| 5 | P13639 | NA | EEF2 | eukaryotic translation elongation factor 2 | 1938 | ENSG00000167658 |
| 6 | P02768 | NA | ALB | albumin | 213 | ENSG00000163631 |
| 7 | P41091 | NA | EIF2S3 | eukaryotic translation initiation factor 2, subunit 3 gamma, 52kDa | 1968 | ENSG00000130741 |
| 8 | P68371 | NA | TUBB4B | tubulin, beta 4B class IVb | 10383 | ENSG00000188229 |
| 9 | P07108 | NA | DBI | diazepam binding inhibitor (GABA receptor modulator, acyl-CoA binding protein) | 1622 | ENSG00000155368 |
| 10 | P47895 | NA | ALDH1A3 | aldehyde dehydrogenase 1 family, member A3 | 220 | ENSG00000184254 |
| 11 | P40227 | NA | CCT6A | chaperonin containing TCP1, subunit 6A (zeta 1) | 908 | ENSG00000146731 |
| 12 | P60842 | NA | EIF4A1 | eukaryotic translation initiation factor 4A1 | 1973 | ENSG00000161960 |
| 13 | P17174 | NA | GOT1 | glutamic-oxaloacetic transaminase 1, soluble (aspartate aminotransferase 1) | 2805 | ENSG00000120053 |
| 14 | P68104 | NA | EEF1A1 | eukaryotic translation elongation factor 1 alpha 1 | 1915 | ENSG00000156508 |
| 15 | P11387 | NA | TOP1 | topoisomerase (DNA) I | 7150 | ENSG00000198900 |

  
  

| **Database:molecular function      &nbspName:glycoprotein binding      &nbspID:GO:0001948** | | | | | | |
| --- | --- | --- | --- | --- | --- | --- |
| C=61; O=3; E=0.13; R=23.02; rawP=0.0003; adjP=0.0032 | | | | | | |
| Index | UserID | Value | Gene Symbol | Gene Name | EntrezGene | Ensembl |
| 1 | P18206 | NA | VCL | vinculin | 7414 | ENSG00000035403 |
| 2 | P05556 | NA | ITGB1 | integrin, beta 1 (fibronectin receptor, beta polypeptide, antigen CD29 includes MDF2, MSK12) | 3688 | ENSG00000150093 |
| 3 | P09382 | NA | LGALS1 | lectin, galactoside-binding, soluble, 1 | 3956 | ENSG00000100097 |

  
  

| **Database:molecular function      &nbspName:small molecule binding      &nbspID:GO:0036094** | | | | | | |
| --- | --- | --- | --- | --- | --- | --- |
| C=2595; O=14; E=5.54; R=2.53; rawP=0.0004; adjP=0.0035 | | | | | | |
| Index | UserID | Value | Gene Symbol | Gene Name | EntrezGene | Ensembl |
| 1 | P49589 | NA | CARS | cysteinyl-tRNA synthetase | 833 | ENSG00000110619 |
| 2 | Q9NR30 | NA | DDX21 | DEAD (Asp-Glu-Ala-Asp) box helicase 21 | 9188 | ENSG00000165732 |
| 3 | P22102 | NA | GART | phosphoribosylglycinamide formyltransferase, phosphoribosylglycinamide synthetase, phosphoribosylaminoimidazole synthetase | 2618 | ENSG00000159131 |
| 4 | P00491 | NA | PNP | purine nucleoside phosphorylase | 4860 | ENSG00000198805 |
| 5 | P13639 | NA | EEF2 | eukaryotic translation elongation factor 2 | 1938 | ENSG00000167658 |
| 6 | P02768 | NA | ALB | albumin | 213 | ENSG00000163631 |
| 7 | P41091 | NA | EIF2S3 | eukaryotic translation initiation factor 2, subunit 3 gamma, 52kDa | 1968 | ENSG00000130741 |
| 8 | P68371 | NA | TUBB4B | tubulin, beta 4B class IVb | 10383 | ENSG00000188229 |
| 9 | P47895 | NA | ALDH1A3 | aldehyde dehydrogenase 1 family, member A3 | 220 | ENSG00000184254 |
| 10 | P40227 | NA | CCT6A | chaperonin containing TCP1, subunit 6A (zeta 1) | 908 | ENSG00000146731 |
| 11 | P60842 | NA | EIF4A1 | eukaryotic translation initiation factor 4A1 | 1973 | ENSG00000161960 |
| 12 | P17174 | NA | GOT1 | glutamic-oxaloacetic transaminase 1, soluble (aspartate aminotransferase 1) | 2805 | ENSG00000120053 |
| 13 | P68104 | NA | EEF1A1 | eukaryotic translation elongation factor 1 alpha 1 | 1915 | ENSG00000156508 |
| 14 | P11387 | NA | TOP1 | topoisomerase (DNA) I | 7150 | ENSG00000198900 |

  
  

| **Database:molecular function      &nbspName:nucleoside-triphosphatase activity      &nbspID:GO:0017111** | | | | | | |
| --- | --- | --- | --- | --- | --- | --- |
| C=749; O=7; E=1.60; R=4.37; rawP=0.0009; adjP=0.0057 | | | | | | |
| Index | UserID | Value | Gene Symbol | Gene Name | EntrezGene | Ensembl |
| 1 | P60842 | NA | EIF4A1 | eukaryotic translation initiation factor 4A1 | 1973 | ENSG00000161960 |
| 2 | Q9NR30 | NA | DDX21 | DEAD (Asp-Glu-Ala-Asp) box helicase 21 | 9188 | ENSG00000165732 |
| 3 | P13639 | NA | EEF2 | eukaryotic translation elongation factor 2 | 1938 | ENSG00000167658 |
| 4 | P41091 | NA | EIF2S3 | eukaryotic translation initiation factor 2, subunit 3 gamma, 52kDa | 1968 | ENSG00000130741 |
| 5 | P68371 | NA | TUBB4B | tubulin, beta 4B class IVb | 10383 | ENSG00000188229 |
| 6 | P68104 | NA | EEF1A1 | eukaryotic translation elongation factor 1 alpha 1 | 1915 | ENSG00000156508 |
| 7 | P11387 | NA | TOP1 | topoisomerase (DNA) I | 7150 | ENSG00000198900 |

  
  

| **Database:molecular function      &nbspName:hydrolase activity, acting on acid anhydrides, in phosphorus-containing anhydrides      &nbspID:GO:0016818** | | | | | | |
| --- | --- | --- | --- | --- | --- | --- |
| C=786; O=7; E=1.68; R=4.17; rawP=0.0011; adjP=0.0057 | | | | | | |
| Index | UserID | Value | Gene Symbol | Gene Name | EntrezGene | Ensembl |
| 1 | P60842 | NA | EIF4A1 | eukaryotic translation initiation factor 4A1 | 1973 | ENSG00000161960 |
| 2 | Q9NR30 | NA | DDX21 | DEAD (Asp-Glu-Ala-Asp) box helicase 21 | 9188 | ENSG00000165732 |
| 3 | P13639 | NA | EEF2 | eukaryotic translation elongation factor 2 | 1938 | ENSG00000167658 |
| 4 | P41091 | NA | EIF2S3 | eukaryotic translation initiation factor 2, subunit 3 gamma, 52kDa | 1968 | ENSG00000130741 |
| 5 | P68371 | NA | TUBB4B | tubulin, beta 4B class IVb | 10383 | ENSG00000188229 |
| 6 | P68104 | NA | EEF1A1 | eukaryotic translation elongation factor 1 alpha 1 | 1915 | ENSG00000156508 |
| 7 | P11387 | NA | TOP1 | topoisomerase (DNA) I | 7150 | ENSG00000198900 |

  
  

| **Database:molecular function      &nbspName:nucleoside binding      &nbspID:GO:0001882** | | | | | | |
| --- | --- | --- | --- | --- | --- | --- |
| C=1823; O=11; E=3.90; R=2.82; rawP=0.0009; adjP=0.0057 | | | | | | |
| Index | UserID | Value | Gene Symbol | Gene Name | EntrezGene | Ensembl |
| 1 | P49589 | NA | CARS | cysteinyl-tRNA synthetase | 833 | ENSG00000110619 |
| 2 | Q9NR30 | NA | DDX21 | DEAD (Asp-Glu-Ala-Asp) box helicase 21 | 9188 | ENSG00000165732 |
| 3 | P22102 | NA | GART | phosphoribosylglycinamide formyltransferase, phosphoribosylglycinamide synthetase, phosphoribosylaminoimidazole synthetase | 2618 | ENSG00000159131 |
| 4 | P00491 | NA | PNP | purine nucleoside phosphorylase | 4860 | ENSG00000198805 |
| 5 | P13639 | NA | EEF2 | eukaryotic translation elongation factor 2 | 1938 | ENSG00000167658 |
| 6 | P41091 | NA | EIF2S3 | eukaryotic translation initiation factor 2, subunit 3 gamma, 52kDa | 1968 | ENSG00000130741 |
| 7 | P68371 | NA | TUBB4B | tubulin, beta 4B class IVb | 10383 | ENSG00000188229 |
| 8 | P40227 | NA | CCT6A | chaperonin containing TCP1, subunit 6A (zeta 1) | 908 | ENSG00000146731 |
| 9 | P60842 | NA | EIF4A1 | eukaryotic translation initiation factor 4A1 | 1973 | ENSG00000161960 |
| 10 | P68104 | NA | EEF1A1 | eukaryotic translation elongation factor 1 alpha 1 | 1915 | ENSG00000156508 |
| 11 | P11387 | NA | TOP1 | topoisomerase (DNA) I | 7150 | ENSG00000198900 |

  
  

| **Database:molecular function      &nbspName:GTPase activity      &nbspID:GO:0003924** | | | | | | |
| --- | --- | --- | --- | --- | --- | --- |
| C=225; O=4; E=0.48; R=8.32; rawP=0.0013; adjP=0.0057 | | | | | | |
| Index | UserID | Value | Gene Symbol | Gene Name | EntrezGene | Ensembl |
| 1 | P13639 | NA | EEF2 | eukaryotic translation elongation factor 2 | 1938 | ENSG00000167658 |
| 2 | P41091 | NA | EIF2S3 | eukaryotic translation initiation factor 2, subunit 3 gamma, 52kDa | 1968 | ENSG00000130741 |
| 3 | P68371 | NA | TUBB4B | tubulin, beta 4B class IVb | 10383 | ENSG00000188229 |
| 4 | P68104 | NA | EEF1A1 | eukaryotic translation elongation factor 1 alpha 1 | 1915 | ENSG00000156508 |

  
  

| **Database:molecular function      &nbspName:hydrolase activity, acting on acid anhydrides      &nbspID:GO:0016817** | | | | | | |
| --- | --- | --- | --- | --- | --- | --- |
| C=791; O=7; E=1.69; R=4.14; rawP=0.0012; adjP=0.0057 | | | | | | |
| Index | UserID | Value | Gene Symbol | Gene Name | EntrezGene | Ensembl |
| 1 | P60842 | NA | EIF4A1 | eukaryotic translation initiation factor 4A1 | 1973 | ENSG00000161960 |
| 2 | Q9NR30 | NA | DDX21 | DEAD (Asp-Glu-Ala-Asp) box helicase 21 | 9188 | ENSG00000165732 |
| 3 | P13639 | NA | EEF2 | eukaryotic translation elongation factor 2 | 1938 | ENSG00000167658 |
| 4 | P41091 | NA | EIF2S3 | eukaryotic translation initiation factor 2, subunit 3 gamma, 52kDa | 1968 | ENSG00000130741 |
| 5 | P68371 | NA | TUBB4B | tubulin, beta 4B class IVb | 10383 | ENSG00000188229 |
| 6 | P68104 | NA | EEF1A1 | eukaryotic translation elongation factor 1 alpha 1 | 1915 | ENSG00000156508 |
| 7 | P11387 | NA | TOP1 | topoisomerase (DNA) I | 7150 | ENSG00000198900 |

  
  

| **Database:molecular function      &nbspName:pyrophosphatase activity      &nbspID:GO:0016462** | | | | | | |
| --- | --- | --- | --- | --- | --- | --- |
| C=783; O=7; E=1.67; R=4.18; rawP=0.0011; adjP=0.0057 | | | | | | |
| Index | UserID | Value | Gene Symbol | Gene Name | EntrezGene | Ensembl |
| 1 | P60842 | NA | EIF4A1 | eukaryotic translation initiation factor 4A1 | 1973 | ENSG00000161960 |
| 2 | Q9NR30 | NA | DDX21 | DEAD (Asp-Glu-Ala-Asp) box helicase 21 | 9188 | ENSG00000165732 |
| 3 | P13639 | NA | EEF2 | eukaryotic translation elongation factor 2 | 1938 | ENSG00000167658 |
| 4 | P41091 | NA | EIF2S3 | eukaryotic translation initiation factor 2, subunit 3 gamma, 52kDa | 1968 | ENSG00000130741 |
| 5 | P68371 | NA | TUBB4B | tubulin, beta 4B class IVb | 10383 | ENSG00000188229 |
| 6 | P68104 | NA | EEF1A1 | eukaryotic translation elongation factor 1 alpha 1 | 1915 | ENSG00000156508 |
| 7 | P11387 | NA | TOP1 | topoisomerase (DNA) I | 7150 | ENSG00000198900 |

  
  

| **Database:molecular function      &nbspName:organic cyclic compound binding      &nbspID:GO:0097159** | | | | | | |
| --- | --- | --- | --- | --- | --- | --- |
| C=5342; O=20; E=11.41; R=1.75; rawP=0.0018; adjP=0.0073 | | | | | | |
| Index | UserID | Value | Gene Symbol | Gene Name | EntrezGene | Ensembl |
| 1 | P49589 | NA | CARS | cysteinyl-tRNA synthetase | 833 | ENSG00000110619 |
| 2 | Q9NR30 | NA | DDX21 | DEAD (Asp-Glu-Ala-Asp) box helicase 21 | 9188 | ENSG00000165732 |
| 3 | P00491 | NA | PNP | purine nucleoside phosphorylase | 4860 | ENSG00000198805 |
| 4 | P68371 | NA | TUBB4B | tubulin, beta 4B class IVb | 10383 | ENSG00000188229 |
| 5 | P61313 | NA | RPL15 | ribosomal protein L15 | 6138 | ENSG00000174748 |
| 6 | P60842 | NA | EIF4A1 | eukaryotic translation initiation factor 4A1 | 1973 | ENSG00000161960 |
| 7 | P68104 | NA | EEF1A1 | eukaryotic translation elongation factor 1 alpha 1 | 1915 | ENSG00000156508 |
| 8 | P04083 | NA | ANXA1 | annexin A1 | 301 | ENSG00000135046 |
| 9 | P24534 | NA | EEF1B2 | eukaryotic translation elongation factor 1 beta 2 | 1933 | ENSG00000114942 |
| 10 | P22102 | NA | GART | phosphoribosylglycinamide formyltransferase, phosphoribosylglycinamide synthetase, phosphoribosylaminoimidazole synthetase | 2618 | ENSG00000159131 |
| 11 | P13639 | NA | EEF2 | eukaryotic translation elongation factor 2 | 1938 | ENSG00000167658 |
| 12 | P02768 | NA | ALB | albumin | 213 | ENSG00000163631 |
| 13 | P41091 | NA | EIF2S3 | eukaryotic translation initiation factor 2, subunit 3 gamma, 52kDa | 1968 | ENSG00000130741 |
| 14 | P47895 | NA | ALDH1A3 | aldehyde dehydrogenase 1 family, member A3 | 220 | ENSG00000184254 |
| 15 | P40227 | NA | CCT6A | chaperonin containing TCP1, subunit 6A (zeta 1) | 908 | ENSG00000146731 |
| 16 | P17174 | NA | GOT1 | glutamic-oxaloacetic transaminase 1, soluble (aspartate aminotransferase 1) | 2805 | ENSG00000120053 |
| 17 | P67809 | NA | YBX1 | Y box binding protein 1 | 4904 | ENSG00000065978 |
| 18 | P00352 | NA | ALDH1A1 | aldehyde dehydrogenase 1 family, member A1 | 216 | ENSG00000165092 |
| 19 | P29692 | NA | EEF1D | eukaryotic translation elongation factor 1 delta (guanine nucleotide exchange protein) | 1936 | ENSG00000104529 |
| 20 | P11387 | NA | TOP1 | topoisomerase (DNA) I | 7150 | ENSG00000198900 |

  
  

| **Database:molecular function      &nbspName:cofactor binding      &nbspID:GO:0048037** | | | | | | |
| --- | --- | --- | --- | --- | --- | --- |
| C=259; O=4; E=0.55; R=7.23; rawP=0.0021; adjP=0.0080 | | | | | | |
| Index | UserID | Value | Gene Symbol | Gene Name | EntrezGene | Ensembl |
| 1 | P17174 | NA | GOT1 | glutamic-oxaloacetic transaminase 1, soluble (aspartate aminotransferase 1) | 2805 | ENSG00000120053 |
| 2 | P02768 | NA | ALB | albumin | 213 | ENSG00000163631 |
| 3 | P07108 | NA | DBI | diazepam binding inhibitor (GABA receptor modulator, acyl-CoA binding protein) | 1622 | ENSG00000155368 |
| 4 | P47895 | NA | ALDH1A3 | aldehyde dehydrogenase 1 family, member A3 | 220 | ENSG00000184254 |

  
  

| **Database:molecular function      &nbspName:purine nucleoside binding      &nbspID:GO:0001883** | | | | | | |
| --- | --- | --- | --- | --- | --- | --- |
| C=1812; O=10; E=3.87; R=2.58; rawP=0.0034; adjP=0.0100 | | | | | | |
| Index | UserID | Value | Gene Symbol | Gene Name | EntrezGene | Ensembl |
| 1 | P49589 | NA | CARS | cysteinyl-tRNA synthetase | 833 | ENSG00000110619 |
| 2 | Q9NR30 | NA | DDX21 | DEAD (Asp-Glu-Ala-Asp) box helicase 21 | 9188 | ENSG00000165732 |
| 3 | P22102 | NA | GART | phosphoribosylglycinamide formyltransferase, phosphoribosylglycinamide synthetase, phosphoribosylaminoimidazole synthetase | 2618 | ENSG00000159131 |
| 4 | P13639 | NA | EEF2 | eukaryotic translation elongation factor 2 | 1938 | ENSG00000167658 |
| 5 | P41091 | NA | EIF2S3 | eukaryotic translation initiation factor 2, subunit 3 gamma, 52kDa | 1968 | ENSG00000130741 |
| 6 | P68371 | NA | TUBB4B | tubulin, beta 4B class IVb | 10383 | ENSG00000188229 |
| 7 | P40227 | NA | CCT6A | chaperonin containing TCP1, subunit 6A (zeta 1) | 908 | ENSG00000146731 |
| 8 | P60842 | NA | EIF4A1 | eukaryotic translation initiation factor 4A1 | 1973 | ENSG00000161960 |
| 9 | P68104 | NA | EEF1A1 | eukaryotic translation elongation factor 1 alpha 1 | 1915 | ENSG00000156508 |
| 10 | P11387 | NA | TOP1 | topoisomerase (DNA) I | 7150 | ENSG00000198900 |

  
  

| **Database:molecular function      &nbspName:purine ribonucleoside binding      &nbspID:GO:0032550** | | | | | | |
| --- | --- | --- | --- | --- | --- | --- |
| C=1809; O=10; E=3.87; R=2.59; rawP=0.0034; adjP=0.0100 | | | | | | |
| Index | UserID | Value | Gene Symbol | Gene Name | EntrezGene | Ensembl |
| 1 | P49589 | NA | CARS | cysteinyl-tRNA synthetase | 833 | ENSG00000110619 |
| 2 | Q9NR30 | NA | DDX21 | DEAD (Asp-Glu-Ala-Asp) box helicase 21 | 9188 | ENSG00000165732 |
| 3 | P22102 | NA | GART | phosphoribosylglycinamide formyltransferase, phosphoribosylglycinamide synthetase, phosphoribosylaminoimidazole synthetase | 2618 | ENSG00000159131 |
| 4 | P13639 | NA | EEF2 | eukaryotic translation elongation factor 2 | 1938 | ENSG00000167658 |
| 5 | P41091 | NA | EIF2S3 | eukaryotic translation initiation factor 2, subunit 3 gamma, 52kDa | 1968 | ENSG00000130741 |
| 6 | P68371 | NA | TUBB4B | tubulin, beta 4B class IVb | 10383 | ENSG00000188229 |
| 7 | P40227 | NA | CCT6A | chaperonin containing TCP1, subunit 6A (zeta 1) | 908 | ENSG00000146731 |
| 8 | P60842 | NA | EIF4A1 | eukaryotic translation initiation factor 4A1 | 1973 | ENSG00000161960 |
| 9 | P68104 | NA | EEF1A1 | eukaryotic translation elongation factor 1 alpha 1 | 1915 | ENSG00000156508 |
| 10 | P11387 | NA | TOP1 | topoisomerase (DNA) I | 7150 | ENSG00000198900 |

  
  

| **Database:molecular function      &nbspName:purine ribonucleoside triphosphate binding      &nbspID:GO:0035639** | | | | | | |
| --- | --- | --- | --- | --- | --- | --- |
| C=1800; O=10; E=3.85; R=2.60; rawP=0.0033; adjP=0.0100 | | | | | | |
| Index | UserID | Value | Gene Symbol | Gene Name | EntrezGene | Ensembl |
| 1 | P49589 | NA | CARS | cysteinyl-tRNA synthetase | 833 | ENSG00000110619 |
| 2 | Q9NR30 | NA | DDX21 | DEAD (Asp-Glu-Ala-Asp) box helicase 21 | 9188 | ENSG00000165732 |
| 3 | P22102 | NA | GART | phosphoribosylglycinamide formyltransferase, phosphoribosylglycinamide synthetase, phosphoribosylaminoimidazole synthetase | 2618 | ENSG00000159131 |
| 4 | P13639 | NA | EEF2 | eukaryotic translation elongation factor 2 | 1938 | ENSG00000167658 |
| 5 | P41091 | NA | EIF2S3 | eukaryotic translation initiation factor 2, subunit 3 gamma, 52kDa | 1968 | ENSG00000130741 |
| 6 | P68371 | NA | TUBB4B | tubulin, beta 4B class IVb | 10383 | ENSG00000188229 |
| 7 | P40227 | NA | CCT6A | chaperonin containing TCP1, subunit 6A (zeta 1) | 908 | ENSG00000146731 |
| 8 | P60842 | NA | EIF4A1 | eukaryotic translation initiation factor 4A1 | 1973 | ENSG00000161960 |
| 9 | P68104 | NA | EEF1A1 | eukaryotic translation elongation factor 1 alpha 1 | 1915 | ENSG00000156508 |
| 10 | P11387 | NA | TOP1 | topoisomerase (DNA) I | 7150 | ENSG00000198900 |

  
  

| **Database:molecular function      &nbspName:ribonucleoside binding      &nbspID:GO:0032549** | | | | | | |
| --- | --- | --- | --- | --- | --- | --- |
| C=1813; O=10; E=3.87; R=2.58; rawP=0.0034; adjP=0.0100 | | | | | | |
| Index | UserID | Value | Gene Symbol | Gene Name | EntrezGene | Ensembl |
| 1 | P49589 | NA | CARS | cysteinyl-tRNA synthetase | 833 | ENSG00000110619 |
| 2 | Q9NR30 | NA | DDX21 | DEAD (Asp-Glu-Ala-Asp) box helicase 21 | 9188 | ENSG00000165732 |
| 3 | P22102 | NA | GART | phosphoribosylglycinamide formyltransferase, phosphoribosylglycinamide synthetase, phosphoribosylaminoimidazole synthetase | 2618 | ENSG00000159131 |
| 4 | P13639 | NA | EEF2 | eukaryotic translation elongation factor 2 | 1938 | ENSG00000167658 |
| 5 | P41091 | NA | EIF2S3 | eukaryotic translation initiation factor 2, subunit 3 gamma, 52kDa | 1968 | ENSG00000130741 |
| 6 | P68371 | NA | TUBB4B | tubulin, beta 4B class IVb | 10383 | ENSG00000188229 |
| 7 | P40227 | NA | CCT6A | chaperonin containing TCP1, subunit 6A (zeta 1) | 908 | ENSG00000146731 |
| 8 | P60842 | NA | EIF4A1 | eukaryotic translation initiation factor 4A1 | 1973 | ENSG00000161960 |
| 9 | P68104 | NA | EEF1A1 | eukaryotic translation elongation factor 1 alpha 1 | 1915 | ENSG00000156508 |
| 10 | P11387 | NA | TOP1 | topoisomerase (DNA) I | 7150 | ENSG00000198900 |

  
  

| **Database:molecular function      &nbspName:purine ribonucleotide binding      &nbspID:GO:0032555** | | | | | | |
| --- | --- | --- | --- | --- | --- | --- |
| C=1835; O=10; E=3.92; R=2.55; rawP=0.0037; adjP=0.0101 | | | | | | |
| Index | UserID | Value | Gene Symbol | Gene Name | EntrezGene | Ensembl |
| 1 | P49589 | NA | CARS | cysteinyl-tRNA synthetase | 833 | ENSG00000110619 |
| 2 | Q9NR30 | NA | DDX21 | DEAD (Asp-Glu-Ala-Asp) box helicase 21 | 9188 | ENSG00000165732 |
| 3 | P22102 | NA | GART | phosphoribosylglycinamide formyltransferase, phosphoribosylglycinamide synthetase, phosphoribosylaminoimidazole synthetase | 2618 | ENSG00000159131 |
| 4 | P13639 | NA | EEF2 | eukaryotic translation elongation factor 2 | 1938 | ENSG00000167658 |
| 5 | P41091 | NA | EIF2S3 | eukaryotic translation initiation factor 2, subunit 3 gamma, 52kDa | 1968 | ENSG00000130741 |
| 6 | P68371 | NA | TUBB4B | tubulin, beta 4B class IVb | 10383 | ENSG00000188229 |
| 7 | P40227 | NA | CCT6A | chaperonin containing TCP1, subunit 6A (zeta 1) | 908 | ENSG00000146731 |
| 8 | P60842 | NA | EIF4A1 | eukaryotic translation initiation factor 4A1 | 1973 | ENSG00000161960 |
| 9 | P68104 | NA | EEF1A1 | eukaryotic translation elongation factor 1 alpha 1 | 1915 | ENSG00000156508 |
| 10 | P11387 | NA | TOP1 | topoisomerase (DNA) I | 7150 | ENSG00000198900 |

  
  

| **Database:molecular function      &nbspName:purine nucleotide binding      &nbspID:GO:0017076** | | | | | | |
| --- | --- | --- | --- | --- | --- | --- |
| C=1842; O=10; E=3.94; R=2.54; rawP=0.0039; adjP=0.0101 | | | | | | |
| Index | UserID | Value | Gene Symbol | Gene Name | EntrezGene | Ensembl |
| 1 | P49589 | NA | CARS | cysteinyl-tRNA synthetase | 833 | ENSG00000110619 |
| 2 | Q9NR30 | NA | DDX21 | DEAD (Asp-Glu-Ala-Asp) box helicase 21 | 9188 | ENSG00000165732 |
| 3 | P22102 | NA | GART | phosphoribosylglycinamide formyltransferase, phosphoribosylglycinamide synthetase, phosphoribosylaminoimidazole synthetase | 2618 | ENSG00000159131 |
| 4 | P13639 | NA | EEF2 | eukaryotic translation elongation factor 2 | 1938 | ENSG00000167658 |
| 5 | P41091 | NA | EIF2S3 | eukaryotic translation initiation factor 2, subunit 3 gamma, 52kDa | 1968 | ENSG00000130741 |
| 6 | P68371 | NA | TUBB4B | tubulin, beta 4B class IVb | 10383 | ENSG00000188229 |
| 7 | P40227 | NA | CCT6A | chaperonin containing TCP1, subunit 6A (zeta 1) | 908 | ENSG00000146731 |
| 8 | P60842 | NA | EIF4A1 | eukaryotic translation initiation factor 4A1 | 1973 | ENSG00000161960 |
| 9 | P68104 | NA | EEF1A1 | eukaryotic translation elongation factor 1 alpha 1 | 1915 | ENSG00000156508 |
| 10 | P11387 | NA | TOP1 | topoisomerase (DNA) I | 7150 | ENSG00000198900 |

  
  

| **Database:molecular function      &nbspName:ribonucleotide binding      &nbspID:GO:0032553** | | | | | | |
| --- | --- | --- | --- | --- | --- | --- |
| C=1850; O=10; E=3.95; R=2.53; rawP=0.0040; adjP=0.0101 | | | | | | |
| Index | UserID | Value | Gene Symbol | Gene Name | EntrezGene | Ensembl |
| 1 | P49589 | NA | CARS | cysteinyl-tRNA synthetase | 833 | ENSG00000110619 |
| 2 | Q9NR30 | NA | DDX21 | DEAD (Asp-Glu-Ala-Asp) box helicase 21 | 9188 | ENSG00000165732 |
| 3 | P22102 | NA | GART | phosphoribosylglycinamide formyltransferase, phosphoribosylglycinamide synthetase, phosphoribosylaminoimidazole synthetase | 2618 | ENSG00000159131 |
| 4 | P13639 | NA | EEF2 | eukaryotic translation elongation factor 2 | 1938 | ENSG00000167658 |
| 5 | P41091 | NA | EIF2S3 | eukaryotic translation initiation factor 2, subunit 3 gamma, 52kDa | 1968 | ENSG00000130741 |
| 6 | P68371 | NA | TUBB4B | tubulin, beta 4B class IVb | 10383 | ENSG00000188229 |
| 7 | P40227 | NA | CCT6A | chaperonin containing TCP1, subunit 6A (zeta 1) | 908 | ENSG00000146731 |
| 8 | P60842 | NA | EIF4A1 | eukaryotic translation initiation factor 4A1 | 1973 | ENSG00000161960 |
| 9 | P68104 | NA | EEF1A1 | eukaryotic translation elongation factor 1 alpha 1 | 1915 | ENSG00000156508 |
| 10 | P11387 | NA | TOP1 | topoisomerase (DNA) I | 7150 | ENSG00000198900 |

  
  

| **Database:molecular function      &nbspName:heterocyclic compound binding      &nbspID:GO:1901363** | | | | | | |
| --- | --- | --- | --- | --- | --- | --- |
| C=5290; O=19; E=11.30; R=1.68; rawP=0.0046; adjP=0.0111 | | | | | | |
| Index | UserID | Value | Gene Symbol | Gene Name | EntrezGene | Ensembl |
| 1 | P49589 | NA | CARS | cysteinyl-tRNA synthetase | 833 | ENSG00000110619 |
| 2 | Q9NR30 | NA | DDX21 | DEAD (Asp-Glu-Ala-Asp) box helicase 21 | 9188 | ENSG00000165732 |
| 3 | P00491 | NA | PNP | purine nucleoside phosphorylase | 4860 | ENSG00000198805 |
| 4 | P68371 | NA | TUBB4B | tubulin, beta 4B class IVb | 10383 | ENSG00000188229 |
| 5 | P61313 | NA | RPL15 | ribosomal protein L15 | 6138 | ENSG00000174748 |
| 6 | P60842 | NA | EIF4A1 | eukaryotic translation initiation factor 4A1 | 1973 | ENSG00000161960 |
| 7 | P68104 | NA | EEF1A1 | eukaryotic translation elongation factor 1 alpha 1 | 1915 | ENSG00000156508 |
| 8 | P04083 | NA | ANXA1 | annexin A1 | 301 | ENSG00000135046 |
| 9 | P24534 | NA | EEF1B2 | eukaryotic translation elongation factor 1 beta 2 | 1933 | ENSG00000114942 |
| 10 | P22102 | NA | GART | phosphoribosylglycinamide formyltransferase, phosphoribosylglycinamide synthetase, phosphoribosylaminoimidazole synthetase | 2618 | ENSG00000159131 |
| 11 | P13639 | NA | EEF2 | eukaryotic translation elongation factor 2 | 1938 | ENSG00000167658 |
| 12 | P02768 | NA | ALB | albumin | 213 | ENSG00000163631 |
| 13 | P41091 | NA | EIF2S3 | eukaryotic translation initiation factor 2, subunit 3 gamma, 52kDa | 1968 | ENSG00000130741 |
| 14 | P47895 | NA | ALDH1A3 | aldehyde dehydrogenase 1 family, member A3 | 220 | ENSG00000184254 |
| 15 | P40227 | NA | CCT6A | chaperonin containing TCP1, subunit 6A (zeta 1) | 908 | ENSG00000146731 |
| 16 | P17174 | NA | GOT1 | glutamic-oxaloacetic transaminase 1, soluble (aspartate aminotransferase 1) | 2805 | ENSG00000120053 |
| 17 | P67809 | NA | YBX1 | Y box binding protein 1 | 4904 | ENSG00000065978 |
| 18 | P29692 | NA | EEF1D | eukaryotic translation elongation factor 1 delta (guanine nucleotide exchange protein) | 1936 | ENSG00000104529 |
| 19 | P11387 | NA | TOP1 | topoisomerase (DNA) I | 7150 | ENSG00000198900 |

  
  

| **Database:molecular function      &nbspName:protein binding      &nbspID:GO:0005515** | | | | | | |
| --- | --- | --- | --- | --- | --- | --- |
| C=7301; O=23; E=15.60; R=1.47; rawP=0.0068; adjP=0.0157 | | | | | | |
| Index | UserID | Value | Gene Symbol | Gene Name | EntrezGene | Ensembl |
| 1 | P49589 | NA | CARS | cysteinyl-tRNA synthetase | 833 | ENSG00000110619 |
| 2 | P08195 | NA | SLC3A2 | solute carrier family 3 (activators of dibasic and neutral amino acid transport), member 2 | 6520 | ENSG00000168003 |
| 3 | P68371 | NA | TUBB4B | tubulin, beta 4B class IVb | 10383 | ENSG00000188229 |
| 4 | P09382 | NA | LGALS1 | lectin, galactoside-binding, soluble, 1 | 3956 | ENSG00000100097 |
| 5 | P61313 | NA | RPL15 | ribosomal protein L15 | 6138 | ENSG00000174748 |
| 6 | P60842 | NA | EIF4A1 | eukaryotic translation initiation factor 4A1 | 1973 | ENSG00000161960 |
| 7 | P04080 | NA | CSTB | cystatin B (stefin B) | 1476 | ENSG00000160213 |
| 8 | P68104 | NA | EEF1A1 | eukaryotic translation elongation factor 1 alpha 1 | 1915 | ENSG00000156508 |
| 9 | P02786 | NA | TFRC | transferrin receptor (p90, CD71) | 7037 | ENSG00000072274 |
| 10 | O75369 | NA | FLNB | filamin B, beta | 2317 | ENSG00000136068 |
| 11 | P04083 | NA | ANXA1 | annexin A1 | 301 | ENSG00000135046 |
| 12 | P24534 | NA | EEF1B2 | eukaryotic translation elongation factor 1 beta 2 | 1933 | ENSG00000114942 |
| 13 | P02768 | NA | ALB | albumin | 213 | ENSG00000163631 |
| 14 | P41091 | NA | EIF2S3 | eukaryotic translation initiation factor 2, subunit 3 gamma, 52kDa | 1968 | ENSG00000130741 |
| 15 | P07108 | NA | DBI | diazepam binding inhibitor (GABA receptor modulator, acyl-CoA binding protein) | 1622 | ENSG00000155368 |
| 16 | P27816 | NA | MAP4 | microtubule-associated protein 4 | 4134 | ENSG00000047849 |
| 17 | P47895 | NA | ALDH1A3 | aldehyde dehydrogenase 1 family, member A3 | 220 | ENSG00000184254 |
| 18 | P05556 | NA | ITGB1 | integrin, beta 1 (fibronectin receptor, beta polypeptide, antigen CD29 includes MDF2, MSK12) | 3688 | ENSG00000150093 |
| 19 | P40227 | NA | CCT6A | chaperonin containing TCP1, subunit 6A (zeta 1) | 908 | ENSG00000146731 |
| 20 | P67809 | NA | YBX1 | Y box binding protein 1 | 4904 | ENSG00000065978 |
| 21 | P18206 | NA | VCL | vinculin | 7414 | ENSG00000035403 |
| 22 | P29692 | NA | EEF1D | eukaryotic translation elongation factor 1 delta (guanine nucleotide exchange protein) | 1936 | ENSG00000104529 |
| 23 | P11387 | NA | TOP1 | topoisomerase (DNA) I | 7150 | ENSG00000198900 |

  
  

| **Database:molecular function      &nbspName:structural molecule activity      &nbspID:GO:0005198** | | | | | | |
| --- | --- | --- | --- | --- | --- | --- |
| C=608; O=5; E=1.30; R=3.85; rawP=0.0088; adjP=0.0161 | | | | | | |
| Index | UserID | Value | Gene Symbol | Gene Name | EntrezGene | Ensembl |
| 1 | P61313 | NA | RPL15 | ribosomal protein L15 | 6138 | ENSG00000174748 |
| 2 | P18206 | NA | VCL | vinculin | 7414 | ENSG00000035403 |
| 3 | P68371 | NA | TUBB4B | tubulin, beta 4B class IVb | 10383 | ENSG00000188229 |
| 4 | P27816 | NA | MAP4 | microtubule-associated protein 4 | 4134 | ENSG00000047849 |
| 5 | P04083 | NA | ANXA1 | annexin A1 | 301 | ENSG00000135046 |

  
  

| **Database:molecular function      &nbspName:GTP binding      &nbspID:GO:0005525** | | | | | | |
| --- | --- | --- | --- | --- | --- | --- |
| C=370; O=4; E=0.79; R=5.06; rawP=0.0076; adjP=0.0161 | | | | | | |
| Index | UserID | Value | Gene Symbol | Gene Name | EntrezGene | Ensembl |
| 1 | P13639 | NA | EEF2 | eukaryotic translation elongation factor 2 | 1938 | ENSG00000167658 |
| 2 | P41091 | NA | EIF2S3 | eukaryotic translation initiation factor 2, subunit 3 gamma, 52kDa | 1968 | ENSG00000130741 |
| 3 | P68371 | NA | TUBB4B | tubulin, beta 4B class IVb | 10383 | ENSG00000188229 |
| 4 | P68104 | NA | EEF1A1 | eukaryotic translation elongation factor 1 alpha 1 | 1915 | ENSG00000156508 |

  
  

| **Database:molecular function      &nbspName:guanyl nucleotide binding      &nbspID:GO:0019001** | | | | | | |
| --- | --- | --- | --- | --- | --- | --- |
| C=383; O=4; E=0.82; R=4.89; rawP=0.0086; adjP=0.0161 | | | | | | |
| Index | UserID | Value | Gene Symbol | Gene Name | EntrezGene | Ensembl |
| 1 | P13639 | NA | EEF2 | eukaryotic translation elongation factor 2 | 1938 | ENSG00000167658 |
| 2 | P41091 | NA | EIF2S3 | eukaryotic translation initiation factor 2, subunit 3 gamma, 52kDa | 1968 | ENSG00000130741 |
| 3 | P68371 | NA | TUBB4B | tubulin, beta 4B class IVb | 10383 | ENSG00000188229 |
| 4 | P68104 | NA | EEF1A1 | eukaryotic translation elongation factor 1 alpha 1 | 1915 | ENSG00000156508 |

  
  

| **Database:molecular function      &nbspName:nucleotide binding      &nbspID:GO:0000166** | | | | | | |
| --- | --- | --- | --- | --- | --- | --- |
| C=2403; O=11; E=5.13; R=2.14; rawP=0.0086; adjP=0.0161 | | | | | | |
| Index | UserID | Value | Gene Symbol | Gene Name | EntrezGene | Ensembl |
| 1 | P49589 | NA | CARS | cysteinyl-tRNA synthetase | 833 | ENSG00000110619 |
| 2 | Q9NR30 | NA | DDX21 | DEAD (Asp-Glu-Ala-Asp) box helicase 21 | 9188 | ENSG00000165732 |
| 3 | P22102 | NA | GART | phosphoribosylglycinamide formyltransferase, phosphoribosylglycinamide synthetase, phosphoribosylaminoimidazole synthetase | 2618 | ENSG00000159131 |
| 4 | P13639 | NA | EEF2 | eukaryotic translation elongation factor 2 | 1938 | ENSG00000167658 |
| 5 | P41091 | NA | EIF2S3 | eukaryotic translation initiation factor 2, subunit 3 gamma, 52kDa | 1968 | ENSG00000130741 |
| 6 | P68371 | NA | TUBB4B | tubulin, beta 4B class IVb | 10383 | ENSG00000188229 |
| 7 | P47895 | NA | ALDH1A3 | aldehyde dehydrogenase 1 family, member A3 | 220 | ENSG00000184254 |
| 8 | P40227 | NA | CCT6A | chaperonin containing TCP1, subunit 6A (zeta 1) | 908 | ENSG00000146731 |
| 9 | P60842 | NA | EIF4A1 | eukaryotic translation initiation factor 4A1 | 1973 | ENSG00000161960 |
| 10 | P68104 | NA | EEF1A1 | eukaryotic translation elongation factor 1 alpha 1 | 1915 | ENSG00000156508 |
| 11 | P11387 | NA | TOP1 | topoisomerase (DNA) I | 7150 | ENSG00000198900 |

  
  

| **Database:molecular function      &nbspName:nucleoside phosphate binding      &nbspID:GO:1901265** | | | | | | |
| --- | --- | --- | --- | --- | --- | --- |
| C=2404; O=11; E=5.14; R=2.14; rawP=0.0087; adjP=0.0161 | | | | | | |
| Index | UserID | Value | Gene Symbol | Gene Name | EntrezGene | Ensembl |
| 1 | P49589 | NA | CARS | cysteinyl-tRNA synthetase | 833 | ENSG00000110619 |
| 2 | Q9NR30 | NA | DDX21 | DEAD (Asp-Glu-Ala-Asp) box helicase 21 | 9188 | ENSG00000165732 |
| 3 | P22102 | NA | GART | phosphoribosylglycinamide formyltransferase, phosphoribosylglycinamide synthetase, phosphoribosylaminoimidazole synthetase | 2618 | ENSG00000159131 |
| 4 | P13639 | NA | EEF2 | eukaryotic translation elongation factor 2 | 1938 | ENSG00000167658 |
| 5 | P41091 | NA | EIF2S3 | eukaryotic translation initiation factor 2, subunit 3 gamma, 52kDa | 1968 | ENSG00000130741 |
| 6 | P68371 | NA | TUBB4B | tubulin, beta 4B class IVb | 10383 | ENSG00000188229 |
| 7 | P47895 | NA | ALDH1A3 | aldehyde dehydrogenase 1 family, member A3 | 220 | ENSG00000184254 |
| 8 | P40227 | NA | CCT6A | chaperonin containing TCP1, subunit 6A (zeta 1) | 908 | ENSG00000146731 |
| 9 | P60842 | NA | EIF4A1 | eukaryotic translation initiation factor 4A1 | 1973 | ENSG00000161960 |
| 10 | P68104 | NA | EEF1A1 | eukaryotic translation elongation factor 1 alpha 1 | 1915 | ENSG00000156508 |
| 11 | P11387 | NA | TOP1 | topoisomerase (DNA) I | 7150 | ENSG00000198900 |

  
  

| **Database:molecular function      &nbspName:guanyl ribonucleotide binding      &nbspID:GO:0032561** | | | | | | |
| --- | --- | --- | --- | --- | --- | --- |
| C=383; O=4; E=0.82; R=4.89; rawP=0.0086; adjP=0.0161 | | | | | | |
| Index | UserID | Value | Gene Symbol | Gene Name | EntrezGene | Ensembl |
| 1 | P13639 | NA | EEF2 | eukaryotic translation elongation factor 2 | 1938 | ENSG00000167658 |
| 2 | P41091 | NA | EIF2S3 | eukaryotic translation initiation factor 2, subunit 3 gamma, 52kDa | 1968 | ENSG00000130741 |
| 3 | P68371 | NA | TUBB4B | tubulin, beta 4B class IVb | 10383 | ENSG00000188229 |
| 4 | P68104 | NA | EEF1A1 | eukaryotic translation elongation factor 1 alpha 1 | 1915 | ENSG00000156508 |

  
  

| **Database:molecular function      &nbspName:nucleic acid binding      &nbspID:GO:0003676** | | | | | | |
| --- | --- | --- | --- | --- | --- | --- |
| C=3222; O=13; E=6.88; R=1.89; rawP=0.0113; adjP=0.0200 | | | | | | |
| Index | UserID | Value | Gene Symbol | Gene Name | EntrezGene | Ensembl |
| 1 | P49589 | NA | CARS | cysteinyl-tRNA synthetase | 833 | ENSG00000110619 |
| 2 | Q9NR30 | NA | DDX21 | DEAD (Asp-Glu-Ala-Asp) box helicase 21 | 9188 | ENSG00000165732 |
| 3 | P13639 | NA | EEF2 | eukaryotic translation elongation factor 2 | 1938 | ENSG00000167658 |
| 4 | P02768 | NA | ALB | albumin | 213 | ENSG00000163631 |
| 5 | P41091 | NA | EIF2S3 | eukaryotic translation initiation factor 2, subunit 3 gamma, 52kDa | 1968 | ENSG00000130741 |
| 6 | P61313 | NA | RPL15 | ribosomal protein L15 | 6138 | ENSG00000174748 |
| 7 | P60842 | NA | EIF4A1 | eukaryotic translation initiation factor 4A1 | 1973 | ENSG00000161960 |
| 8 | P67809 | NA | YBX1 | Y box binding protein 1 | 4904 | ENSG00000065978 |
| 9 | P68104 | NA | EEF1A1 | eukaryotic translation elongation factor 1 alpha 1 | 1915 | ENSG00000156508 |
| 10 | P29692 | NA | EEF1D | eukaryotic translation elongation factor 1 delta (guanine nucleotide exchange protein) | 1936 | ENSG00000104529 |
| 11 | P04083 | NA | ANXA1 | annexin A1 | 301 | ENSG00000135046 |
| 12 | P24534 | NA | EEF1B2 | eukaryotic translation elongation factor 1 beta 2 | 1933 | ENSG00000114942 |
| 13 | P11387 | NA | TOP1 | topoisomerase (DNA) I | 7150 | ENSG00000198900 |

  
  

| **Database:molecular function      &nbspName:ATPase activity, coupled      &nbspID:GO:0042623** | | | | | | |
| --- | --- | --- | --- | --- | --- | --- |
| C=274; O=3; E=0.59; R=5.12; rawP=0.0203; adjP=0.0347 | | | | | | |
| Index | UserID | Value | Gene Symbol | Gene Name | EntrezGene | Ensembl |
| 1 | P60842 | NA | EIF4A1 | eukaryotic translation initiation factor 4A1 | 1973 | ENSG00000161960 |
| 2 | Q9NR30 | NA | DDX21 | DEAD (Asp-Glu-Ala-Asp) box helicase 21 | 9188 | ENSG00000165732 |
| 3 | P11387 | NA | TOP1 | topoisomerase (DNA) I | 7150 | ENSG00000198900 |

  
  

| **Database:molecular function      &nbspName:binding      &nbspID:GO:0005488** | | | | | | |
| --- | --- | --- | --- | --- | --- | --- |
| C=11779; O=30; E=25.17; R=1.19; rawP=0.0211; adjP=0.0349 | | | | | | |
| Index | UserID | Value | Gene Symbol | Gene Name | EntrezGene | Ensembl |
| 1 | P49589 | NA | CARS | cysteinyl-tRNA synthetase | 833 | ENSG00000110619 |
| 2 | Q9NR30 | NA | DDX21 | DEAD (Asp-Glu-Ala-Asp) box helicase 21 | 9188 | ENSG00000165732 |
| 3 | P08195 | NA | SLC3A2 | solute carrier family 3 (activators of dibasic and neutral amino acid transport), member 2 | 6520 | ENSG00000168003 |
| 4 | P00491 | NA | PNP | purine nucleoside phosphorylase | 4860 | ENSG00000198805 |
| 5 | P68371 | NA | TUBB4B | tubulin, beta 4B class IVb | 10383 | ENSG00000188229 |
| 6 | P09382 | NA | LGALS1 | lectin, galactoside-binding, soluble, 1 | 3956 | ENSG00000100097 |
| 7 | P61313 | NA | RPL15 | ribosomal protein L15 | 6138 | ENSG00000174748 |
| 8 | P60842 | NA | EIF4A1 | eukaryotic translation initiation factor 4A1 | 1973 | ENSG00000161960 |
| 9 | P53701 | NA | HCCS | holocytochrome c synthase | 3052 | ENSG00000004961 |
| 10 | P04080 | NA | CSTB | cystatin B (stefin B) | 1476 | ENSG00000160213 |
| 11 | P68104 | NA | EEF1A1 | eukaryotic translation elongation factor 1 alpha 1 | 1915 | ENSG00000156508 |
| 12 | P02786 | NA | TFRC | transferrin receptor (p90, CD71) | 7037 | ENSG00000072274 |
| 13 | O75369 | NA | FLNB | filamin B, beta | 2317 | ENSG00000136068 |
| 14 | P04083 | NA | ANXA1 | annexin A1 | 301 | ENSG00000135046 |
| 15 | P24534 | NA | EEF1B2 | eukaryotic translation elongation factor 1 beta 2 | 1933 | ENSG00000114942 |
| 16 | P22102 | NA | GART | phosphoribosylglycinamide formyltransferase, phosphoribosylglycinamide synthetase, phosphoribosylaminoimidazole synthetase | 2618 | ENSG00000159131 |
| 17 | P13639 | NA | EEF2 | eukaryotic translation elongation factor 2 | 1938 | ENSG00000167658 |
| 18 | P02768 | NA | ALB | albumin | 213 | ENSG00000163631 |
| 19 | P41091 | NA | EIF2S3 | eukaryotic translation initiation factor 2, subunit 3 gamma, 52kDa | 1968 | ENSG00000130741 |
| 20 | P07108 | NA | DBI | diazepam binding inhibitor (GABA receptor modulator, acyl-CoA binding protein) | 1622 | ENSG00000155368 |
| 21 | P27816 | NA | MAP4 | microtubule-associated protein 4 | 4134 | ENSG00000047849 |
| 22 | P47895 | NA | ALDH1A3 | aldehyde dehydrogenase 1 family, member A3 | 220 | ENSG00000184254 |
| 23 | P05556 | NA | ITGB1 | integrin, beta 1 (fibronectin receptor, beta polypeptide, antigen CD29 includes MDF2, MSK12) | 3688 | ENSG00000150093 |
| 24 | P40227 | NA | CCT6A | chaperonin containing TCP1, subunit 6A (zeta 1) | 908 | ENSG00000146731 |
| 25 | P67809 | NA | YBX1 | Y box binding protein 1 | 4904 | ENSG00000065978 |
| 26 | P17174 | NA | GOT1 | glutamic-oxaloacetic transaminase 1, soluble (aspartate aminotransferase 1) | 2805 | ENSG00000120053 |
| 27 | P00352 | NA | ALDH1A1 | aldehyde dehydrogenase 1 family, member A1 | 216 | ENSG00000165092 |
| 28 | P18206 | NA | VCL | vinculin | 7414 | ENSG00000035403 |
| 29 | P29692 | NA | EEF1D | eukaryotic translation elongation factor 1 delta (guanine nucleotide exchange protein) | 1936 | ENSG00000104529 |
| 30 | P11387 | NA | TOP1 | topoisomerase (DNA) I | 7150 | ENSG00000198900 |

  
  

| **Database:molecular function      &nbspName:ion binding      &nbspID:GO:0043167** | | | | | | |
| --- | --- | --- | --- | --- | --- | --- |
| C=5721; O=18; E=12.22; R=1.47; rawP=0.0289; adjP=0.0464 | | | | | | |
| Index | UserID | Value | Gene Symbol | Gene Name | EntrezGene | Ensembl |
| 1 | P49589 | NA | CARS | cysteinyl-tRNA synthetase | 833 | ENSG00000110619 |
| 2 | Q9NR30 | NA | DDX21 | DEAD (Asp-Glu-Ala-Asp) box helicase 21 | 9188 | ENSG00000165732 |
| 3 | P08195 | NA | SLC3A2 | solute carrier family 3 (activators of dibasic and neutral amino acid transport), member 2 | 6520 | ENSG00000168003 |
| 4 | P00491 | NA | PNP | purine nucleoside phosphorylase | 4860 | ENSG00000198805 |
| 5 | P68371 | NA | TUBB4B | tubulin, beta 4B class IVb | 10383 | ENSG00000188229 |
| 6 | P60842 | NA | EIF4A1 | eukaryotic translation initiation factor 4A1 | 1973 | ENSG00000161960 |
| 7 | P53701 | NA | HCCS | holocytochrome c synthase | 3052 | ENSG00000004961 |
| 8 | P68104 | NA | EEF1A1 | eukaryotic translation elongation factor 1 alpha 1 | 1915 | ENSG00000156508 |
| 9 | P04083 | NA | ANXA1 | annexin A1 | 301 | ENSG00000135046 |
| 10 | P22102 | NA | GART | phosphoribosylglycinamide formyltransferase, phosphoribosylglycinamide synthetase, phosphoribosylaminoimidazole synthetase | 2618 | ENSG00000159131 |
| 11 | P13639 | NA | EEF2 | eukaryotic translation elongation factor 2 | 1938 | ENSG00000167658 |
| 12 | P41091 | NA | EIF2S3 | eukaryotic translation initiation factor 2, subunit 3 gamma, 52kDa | 1968 | ENSG00000130741 |
| 13 | P02768 | NA | ALB | albumin | 213 | ENSG00000163631 |
| 14 | P07108 | NA | DBI | diazepam binding inhibitor (GABA receptor modulator, acyl-CoA binding protein) | 1622 | ENSG00000155368 |
| 15 | P47895 | NA | ALDH1A3 | aldehyde dehydrogenase 1 family, member A3 | 220 | ENSG00000184254 |
| 16 | P40227 | NA | CCT6A | chaperonin containing TCP1, subunit 6A (zeta 1) | 908 | ENSG00000146731 |
| 17 | P17174 | NA | GOT1 | glutamic-oxaloacetic transaminase 1, soluble (aspartate aminotransferase 1) | 2805 | ENSG00000120053 |
| 18 | P11387 | NA | TOP1 | topoisomerase (DNA) I | 7150 | ENSG00000198900 |

  
  

| **Database:molecular function      &nbspName:catalytic activity      &nbspID:GO:0003824** | | | | | | |
| --- | --- | --- | --- | --- | --- | --- |
| C=5299; O=17; E=11.32; R=1.50; rawP=0.0298; adjP=0.0465 | | | | | | |
| Index | UserID | Value | Gene Symbol | Gene Name | EntrezGene | Ensembl |
| 1 | P49589 | NA | CARS | cysteinyl-tRNA synthetase | 833 | ENSG00000110619 |
| 2 | Q9NR30 | NA | DDX21 | DEAD (Asp-Glu-Ala-Asp) box helicase 21 | 9188 | ENSG00000165732 |
| 3 | P08195 | NA | SLC3A2 | solute carrier family 3 (activators of dibasic and neutral amino acid transport), member 2 | 6520 | ENSG00000168003 |
| 4 | P00491 | NA | PNP | purine nucleoside phosphorylase | 4860 | ENSG00000198805 |
| 5 | P68371 | NA | TUBB4B | tubulin, beta 4B class IVb | 10383 | ENSG00000188229 |
| 6 | P60842 | NA | EIF4A1 | eukaryotic translation initiation factor 4A1 | 1973 | ENSG00000161960 |
| 7 | P53701 | NA | HCCS | holocytochrome c synthase | 3052 | ENSG00000004961 |
| 8 | P68104 | NA | EEF1A1 | eukaryotic translation elongation factor 1 alpha 1 | 1915 | ENSG00000156508 |
| 9 | P02786 | NA | TFRC | transferrin receptor (p90, CD71) | 7037 | ENSG00000072274 |
| 10 | P22102 | NA | GART | phosphoribosylglycinamide formyltransferase, phosphoribosylglycinamide synthetase, phosphoribosylaminoimidazole synthetase | 2618 | ENSG00000159131 |
| 11 | P13639 | NA | EEF2 | eukaryotic translation elongation factor 2 | 1938 | ENSG00000167658 |
| 12 | P41091 | NA | EIF2S3 | eukaryotic translation initiation factor 2, subunit 3 gamma, 52kDa | 1968 | ENSG00000130741 |
| 13 | P47895 | NA | ALDH1A3 | aldehyde dehydrogenase 1 family, member A3 | 220 | ENSG00000184254 |
| 14 | P17174 | NA | GOT1 | glutamic-oxaloacetic transaminase 1, soluble (aspartate aminotransferase 1) | 2805 | ENSG00000120053 |
| 15 | Q6PIU2 | NA | NCEH1 | neutral cholesterol ester hydrolase 1 | 57552 | ENSG00000144959 |
| 16 | P00352 | NA | ALDH1A1 | aldehyde dehydrogenase 1 family, member A1 | 216 | ENSG00000165092 |
| 17 | P11387 | NA | TOP1 | topoisomerase (DNA) I | 7150 | ENSG00000198900 |

  
  

| **Database:molecular function      &nbspName:protein homodimerization activity      &nbspID:GO:0042803** | | | | | | |
| --- | --- | --- | --- | --- | --- | --- |
| C=569; O=4; E=1.22; R=3.29; rawP=0.0319; adjP=0.0483 | | | | | | |
| Index | UserID | Value | Gene Symbol | Gene Name | EntrezGene | Ensembl |
| 1 | P49589 | NA | CARS | cysteinyl-tRNA synthetase | 833 | ENSG00000110619 |
| 2 | P47895 | NA | ALDH1A3 | aldehyde dehydrogenase 1 family, member A3 | 220 | ENSG00000184254 |
| 3 | P04083 | NA | ANXA1 | annexin A1 | 301 | ENSG00000135046 |
| 4 | P09382 | NA | LGALS1 | lectin, galactoside-binding, soluble, 1 | 3956 | ENSG00000100097 |

  
  

| **Database:molecular function      &nbspName:identical protein binding      &nbspID:GO:0042802** | | | | | | |
| --- | --- | --- | --- | --- | --- | --- |
| C=860; O=5; E=1.84; R=2.72; rawP=0.0342; adjP=0.0504 | | | | | | |
| Index | UserID | Value | Gene Symbol | Gene Name | EntrezGene | Ensembl |
| 1 | P49589 | NA | CARS | cysteinyl-tRNA synthetase | 833 | ENSG00000110619 |
| 2 | P47895 | NA | ALDH1A3 | aldehyde dehydrogenase 1 family, member A3 | 220 | ENSG00000184254 |
| 3 | O75369 | NA | FLNB | filamin B, beta | 2317 | ENSG00000136068 |
| 4 | P04083 | NA | ANXA1 | annexin A1 | 301 | ENSG00000135046 |
| 5 | P09382 | NA | LGALS1 | lectin, galactoside-binding, soluble, 1 | 3956 | ENSG00000100097 |

  
  

| **Database:molecular function      &nbspName:ATPase activity      &nbspID:GO:0016887** | | | | | | |
| --- | --- | --- | --- | --- | --- | --- |
| C=352; O=3; E=0.75; R=3.99; rawP=0.0386; adjP=0.0543 | | | | | | |
| Index | UserID | Value | Gene Symbol | Gene Name | EntrezGene | Ensembl |
| 1 | P60842 | NA | EIF4A1 | eukaryotic translation initiation factor 4A1 | 1973 | ENSG00000161960 |
| 2 | Q9NR30 | NA | DDX21 | DEAD (Asp-Glu-Ala-Asp) box helicase 21 | 9188 | ENSG00000165732 |
| 3 | P11387 | NA | TOP1 | topoisomerase (DNA) I | 7150 | ENSG00000198900 |

  
  

| **Database:molecular function      &nbspName:actin binding      &nbspID:GO:0003779** | | | | | | |
| --- | --- | --- | --- | --- | --- | --- |
| C=353; O=3; E=0.75; R=3.98; rawP=0.0389; adjP=0.0543 | | | | | | |
| Index | UserID | Value | Gene Symbol | Gene Name | EntrezGene | Ensembl |
| 1 | P18206 | NA | VCL | vinculin | 7414 | ENSG00000035403 |
| 2 | O75369 | NA | FLNB | filamin B, beta | 2317 | ENSG00000136068 |
| 3 | P05556 | NA | ITGB1 | integrin, beta 1 (fibronectin receptor, beta polypeptide, antigen CD29 includes MDF2, MSK12) | 3688 | ENSG00000150093 |

  
  

| **Database:molecular function      &nbspName:cytoskeletal protein binding      &nbspID:GO:0008092** | | | | | | |
| --- | --- | --- | --- | --- | --- | --- |
| C=634; O=4; E=1.35; R=2.95; rawP=0.0447; adjP=0.0607 | | | | | | |
| Index | UserID | Value | Gene Symbol | Gene Name | EntrezGene | Ensembl |
| 1 | P18206 | NA | VCL | vinculin | 7414 | ENSG00000035403 |
| 2 | P27816 | NA | MAP4 | microtubule-associated protein 4 | 4134 | ENSG00000047849 |
| 3 | O75369 | NA | FLNB | filamin B, beta | 2317 | ENSG00000136068 |
| 4 | P05556 | NA | ITGB1 | integrin, beta 1 (fibronectin receptor, beta polypeptide, antigen CD29 includes MDF2, MSK12) | 3688 | ENSG00000150093 |

  
  

| **Database:molecular function      &nbspName:protein dimerization activity      &nbspID:GO:0046983** | | | | | | |
| --- | --- | --- | --- | --- | --- | --- |
| C=967; O=5; E=2.07; R=2.42; rawP=0.0524; adjP=0.0677 | | | | | | |
| Index | UserID | Value | Gene Symbol | Gene Name | EntrezGene | Ensembl |
| 1 | P49589 | NA | CARS | cysteinyl-tRNA synthetase | 833 | ENSG00000110619 |
| 2 | P47895 | NA | ALDH1A3 | aldehyde dehydrogenase 1 family, member A3 | 220 | ENSG00000184254 |
| 3 | P05556 | NA | ITGB1 | integrin, beta 1 (fibronectin receptor, beta polypeptide, antigen CD29 includes MDF2, MSK12) | 3688 | ENSG00000150093 |
| 4 | P04083 | NA | ANXA1 | annexin A1 | 301 | ENSG00000135046 |
| 5 | P09382 | NA | LGALS1 | lectin, galactoside-binding, soluble, 1 | 3956 | ENSG00000100097 |

  
  

| **Database:cellular component      &nbspName:eukaryotic translation elongation factor 1 complex      &nbspID:GO:0005853** | | | | | | |
| --- | --- | --- | --- | --- | --- | --- |
| C=5; O=3; E=0.01; R=312.79; rawP=6.39e-08; adjP=3.77e-06 | | | | | | |
| Index | UserID | Value | Gene Symbol | Gene Name | EntrezGene | Ensembl |
| 1 | P68104 | NA | EEF1A1 | eukaryotic translation elongation factor 1 alpha 1 | 1915 | ENSG00000156508 |
| 2 | P29692 | NA | EEF1D | eukaryotic translation elongation factor 1 delta (guanine nucleotide exchange protein) | 1936 | ENSG00000104529 |
| 3 | P24534 | NA | EEF1B2 | eukaryotic translation elongation factor 1 beta 2 | 1933 | ENSG00000114942 |

  
  

| **Database:cellular component      &nbspName:cytosol      &nbspID:GO:0005829** | | | | | | |
| --- | --- | --- | --- | --- | --- | --- |
| C=2367; O=16; E=4.54; R=3.52; rawP=1.61e-06; adjP=3.17e-05 | | | | | | |
| Index | UserID | Value | Gene Symbol | Gene Name | EntrezGene | Ensembl |
| 1 | P49589 | NA | CARS | cysteinyl-tRNA synthetase | 833 | ENSG00000110619 |
| 2 | P00491 | NA | PNP | purine nucleoside phosphorylase | 4860 | ENSG00000198805 |
| 3 | P68371 | NA | TUBB4B | tubulin, beta 4B class IVb | 10383 | ENSG00000188229 |
| 4 | P61313 | NA | RPL15 | ribosomal protein L15 | 6138 | ENSG00000174748 |
| 5 | P60842 | NA | EIF4A1 | eukaryotic translation initiation factor 4A1 | 1973 | ENSG00000161960 |
| 6 | P68104 | NA | EEF1A1 | eukaryotic translation elongation factor 1 alpha 1 | 1915 | ENSG00000156508 |
| 7 | O75369 | NA | FLNB | filamin B, beta | 2317 | ENSG00000136068 |
| 8 | P24534 | NA | EEF1B2 | eukaryotic translation elongation factor 1 beta 2 | 1933 | ENSG00000114942 |
| 9 | P22102 | NA | GART | phosphoribosylglycinamide formyltransferase, phosphoribosylglycinamide synthetase, phosphoribosylaminoimidazole synthetase | 2618 | ENSG00000159131 |
| 10 | P13639 | NA | EEF2 | eukaryotic translation elongation factor 2 | 1938 | ENSG00000167658 |
| 11 | P41091 | NA | EIF2S3 | eukaryotic translation initiation factor 2, subunit 3 gamma, 52kDa | 1968 | ENSG00000130741 |
| 12 | P40227 | NA | CCT6A | chaperonin containing TCP1, subunit 6A (zeta 1) | 908 | ENSG00000146731 |
| 13 | P17174 | NA | GOT1 | glutamic-oxaloacetic transaminase 1, soluble (aspartate aminotransferase 1) | 2805 | ENSG00000120053 |
| 14 | P18206 | NA | VCL | vinculin | 7414 | ENSG00000035403 |
| 15 | P00352 | NA | ALDH1A1 | aldehyde dehydrogenase 1 family, member A1 | 216 | ENSG00000165092 |
| 16 | P29692 | NA | EEF1D | eukaryotic translation elongation factor 1 delta (guanine nucleotide exchange protein) | 1936 | ENSG00000104529 |

  
  

| **Database:cellular component      &nbspName:cytoplasm      &nbspID:GO:0005737** | | | | | | |
| --- | --- | --- | --- | --- | --- | --- |
| C=9051; O=30; E=17.36; R=1.73; rawP=1.19e-06; adjP=3.17e-05 | | | | | | |
| Index | UserID | Value | Gene Symbol | Gene Name | EntrezGene | Ensembl |
| 1 | P49589 | NA | CARS | cysteinyl-tRNA synthetase | 833 | ENSG00000110619 |
| 2 | P08195 | NA | SLC3A2 | solute carrier family 3 (activators of dibasic and neutral amino acid transport), member 2 | 6520 | ENSG00000168003 |
| 3 | P00491 | NA | PNP | purine nucleoside phosphorylase | 4860 | ENSG00000198805 |
| 4 | P68371 | NA | TUBB4B | tubulin, beta 4B class IVb | 10383 | ENSG00000188229 |
| 5 | P09382 | NA | LGALS1 | lectin, galactoside-binding, soluble, 1 | 3956 | ENSG00000100097 |
| 6 | P61313 | NA | RPL15 | ribosomal protein L15 | 6138 | ENSG00000174748 |
| 7 | P60842 | NA | EIF4A1 | eukaryotic translation initiation factor 4A1 | 1973 | ENSG00000161960 |
| 8 | P53701 | NA | HCCS | holocytochrome c synthase | 3052 | ENSG00000004961 |
| 9 | P04080 | NA | CSTB | cystatin B (stefin B) | 1476 | ENSG00000160213 |
| 10 | P68104 | NA | EEF1A1 | eukaryotic translation elongation factor 1 alpha 1 | 1915 | ENSG00000156508 |
| 11 | P02786 | NA | TFRC | transferrin receptor (p90, CD71) | 7037 | ENSG00000072274 |
| 12 | O75369 | NA | FLNB | filamin B, beta | 2317 | ENSG00000136068 |
| 13 | P04083 | NA | ANXA1 | annexin A1 | 301 | ENSG00000135046 |
| 14 | P24534 | NA | EEF1B2 | eukaryotic translation elongation factor 1 beta 2 | 1933 | ENSG00000114942 |
| 15 | P22102 | NA | GART | phosphoribosylglycinamide formyltransferase, phosphoribosylglycinamide synthetase, phosphoribosylaminoimidazole synthetase | 2618 | ENSG00000159131 |
| 16 | P13639 | NA | EEF2 | eukaryotic translation elongation factor 2 | 1938 | ENSG00000167658 |
| 17 | P02768 | NA | ALB | albumin | 213 | ENSG00000163631 |
| 18 | P41091 | NA | EIF2S3 | eukaryotic translation initiation factor 2, subunit 3 gamma, 52kDa | 1968 | ENSG00000130741 |
| 19 | P07108 | NA | DBI | diazepam binding inhibitor (GABA receptor modulator, acyl-CoA binding protein) | 1622 | ENSG00000155368 |
| 20 | P27816 | NA | MAP4 | microtubule-associated protein 4 | 4134 | ENSG00000047849 |
| 21 | P47895 | NA | ALDH1A3 | aldehyde dehydrogenase 1 family, member A3 | 220 | ENSG00000184254 |
| 22 | P05556 | NA | ITGB1 | integrin, beta 1 (fibronectin receptor, beta polypeptide, antigen CD29 includes MDF2, MSK12) | 3688 | ENSG00000150093 |
| 23 | P40227 | NA | CCT6A | chaperonin containing TCP1, subunit 6A (zeta 1) | 908 | ENSG00000146731 |
| 24 | P67809 | NA | YBX1 | Y box binding protein 1 | 4904 | ENSG00000065978 |
| 25 | P17174 | NA | GOT1 | glutamic-oxaloacetic transaminase 1, soluble (aspartate aminotransferase 1) | 2805 | ENSG00000120053 |
| 26 | Q6PIU2 | NA | NCEH1 | neutral cholesterol ester hydrolase 1 | 57552 | ENSG00000144959 |
| 27 | P00352 | NA | ALDH1A1 | aldehyde dehydrogenase 1 family, member A1 | 216 | ENSG00000165092 |
| 28 | P18206 | NA | VCL | vinculin | 7414 | ENSG00000035403 |
| 29 | P29692 | NA | EEF1D | eukaryotic translation elongation factor 1 delta (guanine nucleotide exchange protein) | 1936 | ENSG00000104529 |
| 30 | P11387 | NA | TOP1 | topoisomerase (DNA) I | 7150 | ENSG00000198900 |

  
  

| **Database:cellular component      &nbspName:cytoplasmic part      &nbspID:GO:0044444** | | | | | | |
| --- | --- | --- | --- | --- | --- | --- |
| C=6728; O=26; E=12.91; R=2.01; rawP=2.62e-06; adjP=3.86e-05 | | | | | | |
| Index | UserID | Value | Gene Symbol | Gene Name | EntrezGene | Ensembl |
| 1 | P49589 | NA | CARS | cysteinyl-tRNA synthetase | 833 | ENSG00000110619 |
| 2 | P08195 | NA | SLC3A2 | solute carrier family 3 (activators of dibasic and neutral amino acid transport), member 2 | 6520 | ENSG00000168003 |
| 3 | P00491 | NA | PNP | purine nucleoside phosphorylase | 4860 | ENSG00000198805 |
| 4 | P68371 | NA | TUBB4B | tubulin, beta 4B class IVb | 10383 | ENSG00000188229 |
| 5 | P61313 | NA | RPL15 | ribosomal protein L15 | 6138 | ENSG00000174748 |
| 6 | P60842 | NA | EIF4A1 | eukaryotic translation initiation factor 4A1 | 1973 | ENSG00000161960 |
| 7 | P53701 | NA | HCCS | holocytochrome c synthase | 3052 | ENSG00000004961 |
| 8 | P68104 | NA | EEF1A1 | eukaryotic translation elongation factor 1 alpha 1 | 1915 | ENSG00000156508 |
| 9 | P02786 | NA | TFRC | transferrin receptor (p90, CD71) | 7037 | ENSG00000072274 |
| 10 | O75369 | NA | FLNB | filamin B, beta | 2317 | ENSG00000136068 |
| 11 | P04083 | NA | ANXA1 | annexin A1 | 301 | ENSG00000135046 |
| 12 | P24534 | NA | EEF1B2 | eukaryotic translation elongation factor 1 beta 2 | 1933 | ENSG00000114942 |
| 13 | P22102 | NA | GART | phosphoribosylglycinamide formyltransferase, phosphoribosylglycinamide synthetase, phosphoribosylaminoimidazole synthetase | 2618 | ENSG00000159131 |
| 14 | P13639 | NA | EEF2 | eukaryotic translation elongation factor 2 | 1938 | ENSG00000167658 |
| 15 | P02768 | NA | ALB | albumin | 213 | ENSG00000163631 |
| 16 | P41091 | NA | EIF2S3 | eukaryotic translation initiation factor 2, subunit 3 gamma, 52kDa | 1968 | ENSG00000130741 |
| 17 | P07108 | NA | DBI | diazepam binding inhibitor (GABA receptor modulator, acyl-CoA binding protein) | 1622 | ENSG00000155368 |
| 18 | P05556 | NA | ITGB1 | integrin, beta 1 (fibronectin receptor, beta polypeptide, antigen CD29 includes MDF2, MSK12) | 3688 | ENSG00000150093 |
| 19 | P40227 | NA | CCT6A | chaperonin containing TCP1, subunit 6A (zeta 1) | 908 | ENSG00000146731 |
| 20 | P67809 | NA | YBX1 | Y box binding protein 1 | 4904 | ENSG00000065978 |
| 21 | P17174 | NA | GOT1 | glutamic-oxaloacetic transaminase 1, soluble (aspartate aminotransferase 1) | 2805 | ENSG00000120053 |
| 22 | Q6PIU2 | NA | NCEH1 | neutral cholesterol ester hydrolase 1 | 57552 | ENSG00000144959 |
| 23 | P00352 | NA | ALDH1A1 | aldehyde dehydrogenase 1 family, member A1 | 216 | ENSG00000165092 |
| 24 | P18206 | NA | VCL | vinculin | 7414 | ENSG00000035403 |
| 25 | P29692 | NA | EEF1D | eukaryotic translation elongation factor 1 delta (guanine nucleotide exchange protein) | 1936 | ENSG00000104529 |
| 26 | P11387 | NA | TOP1 | topoisomerase (DNA) I | 7150 | ENSG00000198900 |

  
  

| **Database:cellular component      &nbspName:intracellular part      &nbspID:GO:0044424** | | | | | | |
| --- | --- | --- | --- | --- | --- | --- |
| C=12096; O=31; E=23.20; R=1.34; rawP=0.0004; adjP=0.0047 | | | | | | |
| Index | UserID | Value | Gene Symbol | Gene Name | EntrezGene | Ensembl |
| 1 | P49589 | NA | CARS | cysteinyl-tRNA synthetase | 833 | ENSG00000110619 |
| 2 | Q9NR30 | NA | DDX21 | DEAD (Asp-Glu-Ala-Asp) box helicase 21 | 9188 | ENSG00000165732 |
| 3 | P08195 | NA | SLC3A2 | solute carrier family 3 (activators of dibasic and neutral amino acid transport), member 2 | 6520 | ENSG00000168003 |
| 4 | P00491 | NA | PNP | purine nucleoside phosphorylase | 4860 | ENSG00000198805 |
| 5 | P68371 | NA | TUBB4B | tubulin, beta 4B class IVb | 10383 | ENSG00000188229 |
| 6 | P09382 | NA | LGALS1 | lectin, galactoside-binding, soluble, 1 | 3956 | ENSG00000100097 |
| 7 | P61313 | NA | RPL15 | ribosomal protein L15 | 6138 | ENSG00000174748 |
| 8 | P60842 | NA | EIF4A1 | eukaryotic translation initiation factor 4A1 | 1973 | ENSG00000161960 |
| 9 | P53701 | NA | HCCS | holocytochrome c synthase | 3052 | ENSG00000004961 |
| 10 | P04080 | NA | CSTB | cystatin B (stefin B) | 1476 | ENSG00000160213 |
| 11 | P68104 | NA | EEF1A1 | eukaryotic translation elongation factor 1 alpha 1 | 1915 | ENSG00000156508 |
| 12 | P02786 | NA | TFRC | transferrin receptor (p90, CD71) | 7037 | ENSG00000072274 |
| 13 | O75369 | NA | FLNB | filamin B, beta | 2317 | ENSG00000136068 |
| 14 | P04083 | NA | ANXA1 | annexin A1 | 301 | ENSG00000135046 |
| 15 | P24534 | NA | EEF1B2 | eukaryotic translation elongation factor 1 beta 2 | 1933 | ENSG00000114942 |
| 16 | P22102 | NA | GART | phosphoribosylglycinamide formyltransferase, phosphoribosylglycinamide synthetase, phosphoribosylaminoimidazole synthetase | 2618 | ENSG00000159131 |
| 17 | P13639 | NA | EEF2 | eukaryotic translation elongation factor 2 | 1938 | ENSG00000167658 |
| 18 | P02768 | NA | ALB | albumin | 213 | ENSG00000163631 |
| 19 | P41091 | NA | EIF2S3 | eukaryotic translation initiation factor 2, subunit 3 gamma, 52kDa | 1968 | ENSG00000130741 |
| 20 | P07108 | NA | DBI | diazepam binding inhibitor (GABA receptor modulator, acyl-CoA binding protein) | 1622 | ENSG00000155368 |
| 21 | P27816 | NA | MAP4 | microtubule-associated protein 4 | 4134 | ENSG00000047849 |
| 22 | P47895 | NA | ALDH1A3 | aldehyde dehydrogenase 1 family, member A3 | 220 | ENSG00000184254 |
| 23 | P05556 | NA | ITGB1 | integrin, beta 1 (fibronectin receptor, beta polypeptide, antigen CD29 includes MDF2, MSK12) | 3688 | ENSG00000150093 |
| 24 | P40227 | NA | CCT6A | chaperonin containing TCP1, subunit 6A (zeta 1) | 908 | ENSG00000146731 |
| 25 | P67809 | NA | YBX1 | Y box binding protein 1 | 4904 | ENSG00000065978 |
| 26 | P17174 | NA | GOT1 | glutamic-oxaloacetic transaminase 1, soluble (aspartate aminotransferase 1) | 2805 | ENSG00000120053 |
| 27 | Q6PIU2 | NA | NCEH1 | neutral cholesterol ester hydrolase 1 | 57552 | ENSG00000144959 |
| 28 | P00352 | NA | ALDH1A1 | aldehyde dehydrogenase 1 family, member A1 | 216 | ENSG00000165092 |
| 29 | P18206 | NA | VCL | vinculin | 7414 | ENSG00000035403 |
| 30 | P29692 | NA | EEF1D | eukaryotic translation elongation factor 1 delta (guanine nucleotide exchange protein) | 1936 | ENSG00000104529 |
| 31 | P11387 | NA | TOP1 | topoisomerase (DNA) I | 7150 | ENSG00000198900 |

  
  

| **Database:cellular component      &nbspName:melanosome      &nbspID:GO:0042470** | | | | | | |
| --- | --- | --- | --- | --- | --- | --- |
| C=93; O=3; E=0.18; R=16.82; rawP=0.0007; adjP=0.0059 | | | | | | |
| Index | UserID | Value | Gene Symbol | Gene Name | EntrezGene | Ensembl |
| 1 | P08195 | NA | SLC3A2 | solute carrier family 3 (activators of dibasic and neutral amino acid transport), member 2 | 6520 | ENSG00000168003 |
| 2 | P02786 | NA | TFRC | transferrin receptor (p90, CD71) | 7037 | ENSG00000072274 |
| 3 | P05556 | NA | ITGB1 | integrin, beta 1 (fibronectin receptor, beta polypeptide, antigen CD29 includes MDF2, MSK12) | 3688 | ENSG00000150093 |

  
  

| **Database:cellular component      &nbspName:pigment granule      &nbspID:GO:0048770** | | | | | | |
| --- | --- | --- | --- | --- | --- | --- |
| C=93; O=3; E=0.18; R=16.82; rawP=0.0007; adjP=0.0059 | | | | | | |
| Index | UserID | Value | Gene Symbol | Gene Name | EntrezGene | Ensembl |
| 1 | P08195 | NA | SLC3A2 | solute carrier family 3 (activators of dibasic and neutral amino acid transport), member 2 | 6520 | ENSG00000168003 |
| 2 | P02786 | NA | TFRC | transferrin receptor (p90, CD71) | 7037 | ENSG00000072274 |
| 3 | P05556 | NA | ITGB1 | integrin, beta 1 (fibronectin receptor, beta polypeptide, antigen CD29 includes MDF2, MSK12) | 3688 | ENSG00000150093 |

  
  

| **Database:cellular component      &nbspName:intracellular      &nbspID:GO:0005622** | | | | | | |
| --- | --- | --- | --- | --- | --- | --- |
| C=12412; O=31; E=23.81; R=1.30; rawP=0.0009; adjP=0.0066 | | | | | | |
| Index | UserID | Value | Gene Symbol | Gene Name | EntrezGene | Ensembl |
| 1 | P49589 | NA | CARS | cysteinyl-tRNA synthetase | 833 | ENSG00000110619 |
| 2 | Q9NR30 | NA | DDX21 | DEAD (Asp-Glu-Ala-Asp) box helicase 21 | 9188 | ENSG00000165732 |
| 3 | P08195 | NA | SLC3A2 | solute carrier family 3 (activators of dibasic and neutral amino acid transport), member 2 | 6520 | ENSG00000168003 |
| 4 | P00491 | NA | PNP | purine nucleoside phosphorylase | 4860 | ENSG00000198805 |
| 5 | P68371 | NA | TUBB4B | tubulin, beta 4B class IVb | 10383 | ENSG00000188229 |
| 6 | P09382 | NA | LGALS1 | lectin, galactoside-binding, soluble, 1 | 3956 | ENSG00000100097 |
| 7 | P61313 | NA | RPL15 | ribosomal protein L15 | 6138 | ENSG00000174748 |
| 8 | P60842 | NA | EIF4A1 | eukaryotic translation initiation factor 4A1 | 1973 | ENSG00000161960 |
| 9 | P53701 | NA | HCCS | holocytochrome c synthase | 3052 | ENSG00000004961 |
| 10 | P04080 | NA | CSTB | cystatin B (stefin B) | 1476 | ENSG00000160213 |
| 11 | P68104 | NA | EEF1A1 | eukaryotic translation elongation factor 1 alpha 1 | 1915 | ENSG00000156508 |
| 12 | P02786 | NA | TFRC | transferrin receptor (p90, CD71) | 7037 | ENSG00000072274 |
| 13 | O75369 | NA | FLNB | filamin B, beta | 2317 | ENSG00000136068 |
| 14 | P04083 | NA | ANXA1 | annexin A1 | 301 | ENSG00000135046 |
| 15 | P24534 | NA | EEF1B2 | eukaryotic translation elongation factor 1 beta 2 | 1933 | ENSG00000114942 |
| 16 | P22102 | NA | GART | phosphoribosylglycinamide formyltransferase, phosphoribosylglycinamide synthetase, phosphoribosylaminoimidazole synthetase | 2618 | ENSG00000159131 |
| 17 | P13639 | NA | EEF2 | eukaryotic translation elongation factor 2 | 1938 | ENSG00000167658 |
| 18 | P02768 | NA | ALB | albumin | 213 | ENSG00000163631 |
| 19 | P41091 | NA | EIF2S3 | eukaryotic translation initiation factor 2, subunit 3 gamma, 52kDa | 1968 | ENSG00000130741 |
| 20 | P07108 | NA | DBI | diazepam binding inhibitor (GABA receptor modulator, acyl-CoA binding protein) | 1622 | ENSG00000155368 |
| 21 | P27816 | NA | MAP4 | microtubule-associated protein 4 | 4134 | ENSG00000047849 |
| 22 | P47895 | NA | ALDH1A3 | aldehyde dehydrogenase 1 family, member A3 | 220 | ENSG00000184254 |
| 23 | P05556 | NA | ITGB1 | integrin, beta 1 (fibronectin receptor, beta polypeptide, antigen CD29 includes MDF2, MSK12) | 3688 | ENSG00000150093 |
| 24 | P40227 | NA | CCT6A | chaperonin containing TCP1, subunit 6A (zeta 1) | 908 | ENSG00000146731 |
| 25 | P67809 | NA | YBX1 | Y box binding protein 1 | 4904 | ENSG00000065978 |
| 26 | P17174 | NA | GOT1 | glutamic-oxaloacetic transaminase 1, soluble (aspartate aminotransferase 1) | 2805 | ENSG00000120053 |
| 27 | Q6PIU2 | NA | NCEH1 | neutral cholesterol ester hydrolase 1 | 57552 | ENSG00000144959 |
| 28 | P00352 | NA | ALDH1A1 | aldehyde dehydrogenase 1 family, member A1 | 216 | ENSG00000165092 |
| 29 | P18206 | NA | VCL | vinculin | 7414 | ENSG00000035403 |
| 30 | P29692 | NA | EEF1D | eukaryotic translation elongation factor 1 delta (guanine nucleotide exchange protein) | 1936 | ENSG00000104529 |
| 31 | P11387 | NA | TOP1 | topoisomerase (DNA) I | 7150 | ENSG00000198900 |

  
  

| **Database:cellular component      &nbspName:macromolecular complex      &nbspID:GO:0032991** | | | | | | |
| --- | --- | --- | --- | --- | --- | --- |
| C=3836; O=15; E=7.36; R=2.04; rawP=0.0025; adjP=0.0128 | | | | | | |
| Index | UserID | Value | Gene Symbol | Gene Name | EntrezGene | Ensembl |
| 1 | P13639 | NA | EEF2 | eukaryotic translation elongation factor 2 | 1938 | ENSG00000167658 |
| 2 | P02768 | NA | ALB | albumin | 213 | ENSG00000163631 |
| 3 | P68371 | NA | TUBB4B | tubulin, beta 4B class IVb | 10383 | ENSG00000188229 |
| 4 | P27816 | NA | MAP4 | microtubule-associated protein 4 | 4134 | ENSG00000047849 |
| 5 | P05556 | NA | ITGB1 | integrin, beta 1 (fibronectin receptor, beta polypeptide, antigen CD29 includes MDF2, MSK12) | 3688 | ENSG00000150093 |
| 6 | P40227 | NA | CCT6A | chaperonin containing TCP1, subunit 6A (zeta 1) | 908 | ENSG00000146731 |
| 7 | P61313 | NA | RPL15 | ribosomal protein L15 | 6138 | ENSG00000174748 |
| 8 | P60842 | NA | EIF4A1 | eukaryotic translation initiation factor 4A1 | 1973 | ENSG00000161960 |
| 9 | P67809 | NA | YBX1 | Y box binding protein 1 | 4904 | ENSG00000065978 |
| 10 | P18206 | NA | VCL | vinculin | 7414 | ENSG00000035403 |
| 11 | P68104 | NA | EEF1A1 | eukaryotic translation elongation factor 1 alpha 1 | 1915 | ENSG00000156508 |
| 12 | P29692 | NA | EEF1D | eukaryotic translation elongation factor 1 delta (guanine nucleotide exchange protein) | 1936 | ENSG00000104529 |
| 13 | P04083 | NA | ANXA1 | annexin A1 | 301 | ENSG00000135046 |
| 14 | P24534 | NA | EEF1B2 | eukaryotic translation elongation factor 1 beta 2 | 1933 | ENSG00000114942 |
| 15 | P11387 | NA | TOP1 | topoisomerase (DNA) I | 7150 | ENSG00000198900 |

  
  

| **Database:cellular component      &nbspName:focal adhesion      &nbspID:GO:0005925** | | | | | | |
| --- | --- | --- | --- | --- | --- | --- |
| C=131; O=3; E=0.25; R=11.94; rawP=0.0020; adjP=0.0128 | | | | | | |
| Index | UserID | Value | Gene Symbol | Gene Name | EntrezGene | Ensembl |
| 1 | P18206 | NA | VCL | vinculin | 7414 | ENSG00000035403 |
| 2 | O75369 | NA | FLNB | filamin B, beta | 2317 | ENSG00000136068 |
| 3 | P05556 | NA | ITGB1 | integrin, beta 1 (fibronectin receptor, beta polypeptide, antigen CD29 includes MDF2, MSK12) | 3688 | ENSG00000150093 |

  
  

| **Database:cellular component      &nbspName:cell-substrate adherens junction      &nbspID:GO:0005924** | | | | | | |
| --- | --- | --- | --- | --- | --- | --- |
| C=136; O=3; E=0.26; R=11.50; rawP=0.0022; adjP=0.0128 | | | | | | |
| Index | UserID | Value | Gene Symbol | Gene Name | EntrezGene | Ensembl |
| 1 | P18206 | NA | VCL | vinculin | 7414 | ENSG00000035403 |
| 2 | O75369 | NA | FLNB | filamin B, beta | 2317 | ENSG00000136068 |
| 3 | P05556 | NA | ITGB1 | integrin, beta 1 (fibronectin receptor, beta polypeptide, antigen CD29 includes MDF2, MSK12) | 3688 | ENSG00000150093 |

  
  

| **Database:cellular component      &nbspName:cell-substrate junction      &nbspID:GO:0030055** | | | | | | |
| --- | --- | --- | --- | --- | --- | --- |
| C=143; O=3; E=0.27; R=10.94; rawP=0.0026; adjP=0.0128 | | | | | | |
| Index | UserID | Value | Gene Symbol | Gene Name | EntrezGene | Ensembl |
| 1 | P18206 | NA | VCL | vinculin | 7414 | ENSG00000035403 |
| 2 | O75369 | NA | FLNB | filamin B, beta | 2317 | ENSG00000136068 |
| 3 | P05556 | NA | ITGB1 | integrin, beta 1 (fibronectin receptor, beta polypeptide, antigen CD29 includes MDF2, MSK12) | 3688 | ENSG00000150093 |

  
  

| **Database:cellular component      &nbspName:protein complex      &nbspID:GO:0043234** | | | | | | |
| --- | --- | --- | --- | --- | --- | --- |
| C=3262; O=13; E=6.26; R=2.08; rawP=0.0049; adjP=0.0222 | | | | | | |
| Index | UserID | Value | Gene Symbol | Gene Name | EntrezGene | Ensembl |
| 1 | P02768 | NA | ALB | albumin | 213 | ENSG00000163631 |
| 2 | P68371 | NA | TUBB4B | tubulin, beta 4B class IVb | 10383 | ENSG00000188229 |
| 3 | P27816 | NA | MAP4 | microtubule-associated protein 4 | 4134 | ENSG00000047849 |
| 4 | P05556 | NA | ITGB1 | integrin, beta 1 (fibronectin receptor, beta polypeptide, antigen CD29 includes MDF2, MSK12) | 3688 | ENSG00000150093 |
| 5 | P40227 | NA | CCT6A | chaperonin containing TCP1, subunit 6A (zeta 1) | 908 | ENSG00000146731 |
| 6 | P67809 | NA | YBX1 | Y box binding protein 1 | 4904 | ENSG00000065978 |
| 7 | P60842 | NA | EIF4A1 | eukaryotic translation initiation factor 4A1 | 1973 | ENSG00000161960 |
| 8 | P18206 | NA | VCL | vinculin | 7414 | ENSG00000035403 |
| 9 | P68104 | NA | EEF1A1 | eukaryotic translation elongation factor 1 alpha 1 | 1915 | ENSG00000156508 |
| 10 | P29692 | NA | EEF1D | eukaryotic translation elongation factor 1 delta (guanine nucleotide exchange protein) | 1936 | ENSG00000104529 |
| 11 | P04083 | NA | ANXA1 | annexin A1 | 301 | ENSG00000135046 |
| 12 | P24534 | NA | EEF1B2 | eukaryotic translation elongation factor 1 beta 2 | 1933 | ENSG00000114942 |
| 13 | P11387 | NA | TOP1 | topoisomerase (DNA) I | 7150 | ENSG00000198900 |

  
  

| **Database:cellular component      &nbspName:adherens junction      &nbspID:GO:0005912** | | | | | | |
| --- | --- | --- | --- | --- | --- | --- |
| C=201; O=3; E=0.39; R=7.78; rawP=0.0066; adjP=0.0278 | | | | | | |
| Index | UserID | Value | Gene Symbol | Gene Name | EntrezGene | Ensembl |
| 1 | P18206 | NA | VCL | vinculin | 7414 | ENSG00000035403 |
| 2 | O75369 | NA | FLNB | filamin B, beta | 2317 | ENSG00000136068 |
| 3 | P05556 | NA | ITGB1 | integrin, beta 1 (fibronectin receptor, beta polypeptide, antigen CD29 includes MDF2, MSK12) | 3688 | ENSG00000150093 |

  
  

| **Database:cellular component      &nbspName:anchoring junction      &nbspID:GO:0070161** | | | | | | |
| --- | --- | --- | --- | --- | --- | --- |
| C=218; O=3; E=0.42; R=7.17; rawP=0.0083; adjP=0.0316 | | | | | | |
| Index | UserID | Value | Gene Symbol | Gene Name | EntrezGene | Ensembl |
| 1 | P18206 | NA | VCL | vinculin | 7414 | ENSG00000035403 |
| 2 | O75369 | NA | FLNB | filamin B, beta | 2317 | ENSG00000136068 |
| 3 | P05556 | NA | ITGB1 | integrin, beta 1 (fibronectin receptor, beta polypeptide, antigen CD29 includes MDF2, MSK12) | 3688 | ENSG00000150093 |

  
  

| **Database:cellular component      &nbspName:cell      &nbspID:GO:0005623** | | | | | | |
| --- | --- | --- | --- | --- | --- | --- |
| C=14406; O=32; E=27.63; R=1.16; rawP=0.0091; adjP=0.0316 | | | | | | |
| Index | UserID | Value | Gene Symbol | Gene Name | EntrezGene | Ensembl |
| 1 | P49589 | NA | CARS | cysteinyl-tRNA synthetase | 833 | ENSG00000110619 |
| 2 | P68371 | NA | TUBB4B | tubulin, beta 4B class IVb | 10383 | ENSG00000188229 |
| 3 | P61313 | NA | RPL15 | ribosomal protein L15 | 6138 | ENSG00000174748 |
| 4 | P53701 | NA | HCCS | holocytochrome c synthase | 3052 | ENSG00000004961 |
| 5 | P68104 | NA | EEF1A1 | eukaryotic translation elongation factor 1 alpha 1 | 1915 | ENSG00000156508 |
| 6 | P02786 | NA | TFRC | transferrin receptor (p90, CD71) | 7037 | ENSG00000072274 |
| 7 | O75369 | NA | FLNB | filamin B, beta | 2317 | ENSG00000136068 |
| 8 | P24534 | NA | EEF1B2 | eukaryotic translation elongation factor 1 beta 2 | 1933 | ENSG00000114942 |
| 9 | P22102 | NA | GART | phosphoribosylglycinamide formyltransferase, phosphoribosylglycinamide synthetase, phosphoribosylaminoimidazole synthetase | 2618 | ENSG00000159131 |
| 10 | P13639 | NA | EEF2 | eukaryotic translation elongation factor 2 | 1938 | ENSG00000167658 |
| 11 | P47895 | NA | ALDH1A3 | aldehyde dehydrogenase 1 family, member A3 | 220 | ENSG00000184254 |
| 12 | Q96QD8 | NA | SLC38A2 | solute carrier family 38, member 2 | 54407 | ENSG00000134294 |
| 13 | P67809 | NA | YBX1 | Y box binding protein 1 | 4904 | ENSG00000065978 |
| 14 | P17174 | NA | GOT1 | glutamic-oxaloacetic transaminase 1, soluble (aspartate aminotransferase 1) | 2805 | ENSG00000120053 |
| 15 | Q6PIU2 | NA | NCEH1 | neutral cholesterol ester hydrolase 1 | 57552 | ENSG00000144959 |
| 16 | P18206 | NA | VCL | vinculin | 7414 | ENSG00000035403 |
| 17 | P29692 | NA | EEF1D | eukaryotic translation elongation factor 1 delta (guanine nucleotide exchange protein) | 1936 | ENSG00000104529 |
| 18 | P11387 | NA | TOP1 | topoisomerase (DNA) I | 7150 | ENSG00000198900 |
| 19 | Q9NR30 | NA | DDX21 | DEAD (Asp-Glu-Ala-Asp) box helicase 21 | 9188 | ENSG00000165732 |
| 20 | P08195 | NA | SLC3A2 | solute carrier family 3 (activators of dibasic and neutral amino acid transport), member 2 | 6520 | ENSG00000168003 |
| 21 | P00491 | NA | PNP | purine nucleoside phosphorylase | 4860 | ENSG00000198805 |
| 22 | P09382 | NA | LGALS1 | lectin, galactoside-binding, soluble, 1 | 3956 | ENSG00000100097 |
| 23 | P60842 | NA | EIF4A1 | eukaryotic translation initiation factor 4A1 | 1973 | ENSG00000161960 |
| 24 | P04080 | NA | CSTB | cystatin B (stefin B) | 1476 | ENSG00000160213 |
| 25 | P04083 | NA | ANXA1 | annexin A1 | 301 | ENSG00000135046 |
| 26 | P41091 | NA | EIF2S3 | eukaryotic translation initiation factor 2, subunit 3 gamma, 52kDa | 1968 | ENSG00000130741 |
| 27 | P02768 | NA | ALB | albumin | 213 | ENSG00000163631 |
| 28 | P27816 | NA | MAP4 | microtubule-associated protein 4 | 4134 | ENSG00000047849 |
| 29 | P07108 | NA | DBI | diazepam binding inhibitor (GABA receptor modulator, acyl-CoA binding protein) | 1622 | ENSG00000155368 |
| 30 | P05556 | NA | ITGB1 | integrin, beta 1 (fibronectin receptor, beta polypeptide, antigen CD29 includes MDF2, MSK12) | 3688 | ENSG00000150093 |
| 31 | P40227 | NA | CCT6A | chaperonin containing TCP1, subunit 6A (zeta 1) | 908 | ENSG00000146731 |
| 32 | P00352 | NA | ALDH1A1 | aldehyde dehydrogenase 1 family, member A1 | 216 | ENSG00000165092 |

  
  

| **Database:cellular component      &nbspName:cell part      &nbspID:GO:0044464** | | | | | | |
| --- | --- | --- | --- | --- | --- | --- |
| C=14405; O=32; E=27.63; R=1.16; rawP=0.0091; adjP=0.0316 | | | | | | |
| Index | UserID | Value | Gene Symbol | Gene Name | EntrezGene | Ensembl |
| 1 | P49589 | NA | CARS | cysteinyl-tRNA synthetase | 833 | ENSG00000110619 |
| 2 | P68371 | NA | TUBB4B | tubulin, beta 4B class IVb | 10383 | ENSG00000188229 |
| 3 | P61313 | NA | RPL15 | ribosomal protein L15 | 6138 | ENSG00000174748 |
| 4 | P53701 | NA | HCCS | holocytochrome c synthase | 3052 | ENSG00000004961 |
| 5 | P68104 | NA | EEF1A1 | eukaryotic translation elongation factor 1 alpha 1 | 1915 | ENSG00000156508 |
| 6 | P02786 | NA | TFRC | transferrin receptor (p90, CD71) | 7037 | ENSG00000072274 |
| 7 | O75369 | NA | FLNB | filamin B, beta | 2317 | ENSG00000136068 |
| 8 | P24534 | NA | EEF1B2 | eukaryotic translation elongation factor 1 beta 2 | 1933 | ENSG00000114942 |
| 9 | P22102 | NA | GART | phosphoribosylglycinamide formyltransferase, phosphoribosylglycinamide synthetase, phosphoribosylaminoimidazole synthetase | 2618 | ENSG00000159131 |
| 10 | P13639 | NA | EEF2 | eukaryotic translation elongation factor 2 | 1938 | ENSG00000167658 |
| 11 | P47895 | NA | ALDH1A3 | aldehyde dehydrogenase 1 family, member A3 | 220 | ENSG00000184254 |
| 12 | Q96QD8 | NA | SLC38A2 | solute carrier family 38, member 2 | 54407 | ENSG00000134294 |
| 13 | P67809 | NA | YBX1 | Y box binding protein 1 | 4904 | ENSG00000065978 |
| 14 | P17174 | NA | GOT1 | glutamic-oxaloacetic transaminase 1, soluble (aspartate aminotransferase 1) | 2805 | ENSG00000120053 |
| 15 | Q6PIU2 | NA | NCEH1 | neutral cholesterol ester hydrolase 1 | 57552 | ENSG00000144959 |
| 16 | P18206 | NA | VCL | vinculin | 7414 | ENSG00000035403 |
| 17 | P29692 | NA | EEF1D | eukaryotic translation elongation factor 1 delta (guanine nucleotide exchange protein) | 1936 | ENSG00000104529 |
| 18 | P11387 | NA | TOP1 | topoisomerase (DNA) I | 7150 | ENSG00000198900 |
| 19 | Q9NR30 | NA | DDX21 | DEAD (Asp-Glu-Ala-Asp) box helicase 21 | 9188 | ENSG00000165732 |
| 20 | P08195 | NA | SLC3A2 | solute carrier family 3 (activators of dibasic and neutral amino acid transport), member 2 | 6520 | ENSG00000168003 |
| 21 | P00491 | NA | PNP | purine nucleoside phosphorylase | 4860 | ENSG00000198805 |
| 22 | P09382 | NA | LGALS1 | lectin, galactoside-binding, soluble, 1 | 3956 | ENSG00000100097 |
| 23 | P60842 | NA | EIF4A1 | eukaryotic translation initiation factor 4A1 | 1973 | ENSG00000161960 |
| 24 | P04080 | NA | CSTB | cystatin B (stefin B) | 1476 | ENSG00000160213 |
| 25 | P04083 | NA | ANXA1 | annexin A1 | 301 | ENSG00000135046 |
| 26 | P41091 | NA | EIF2S3 | eukaryotic translation initiation factor 2, subunit 3 gamma, 52kDa | 1968 | ENSG00000130741 |
| 27 | P02768 | NA | ALB | albumin | 213 | ENSG00000163631 |
| 28 | P27816 | NA | MAP4 | microtubule-associated protein 4 | 4134 | ENSG00000047849 |
| 29 | P07108 | NA | DBI | diazepam binding inhibitor (GABA receptor modulator, acyl-CoA binding protein) | 1622 | ENSG00000155368 |
| 30 | P05556 | NA | ITGB1 | integrin, beta 1 (fibronectin receptor, beta polypeptide, antigen CD29 includes MDF2, MSK12) | 3688 | ENSG00000150093 |
| 31 | P40227 | NA | CCT6A | chaperonin containing TCP1, subunit 6A (zeta 1) | 908 | ENSG00000146731 |
| 32 | P00352 | NA | ALDH1A1 | aldehyde dehydrogenase 1 family, member A1 | 216 | ENSG00000165092 |

  
  

| **Database:cellular component      &nbspName:cell surface      &nbspID:GO:0009986** | | | | | | |
| --- | --- | --- | --- | --- | --- | --- |
| C=491; O=4; E=0.94; R=4.25; rawP=0.0139; adjP=0.0456 | | | | | | |
| Index | UserID | Value | Gene Symbol | Gene Name | EntrezGene | Ensembl |
| 1 | P08195 | NA | SLC3A2 | solute carrier family 3 (activators of dibasic and neutral amino acid transport), member 2 | 6520 | ENSG00000168003 |
| 2 | P02786 | NA | TFRC | transferrin receptor (p90, CD71) | 7037 | ENSG00000072274 |
| 3 | P05556 | NA | ITGB1 | integrin, beta 1 (fibronectin receptor, beta polypeptide, antigen CD29 includes MDF2, MSK12) | 3688 | ENSG00000150093 |
| 4 | P09382 | NA | LGALS1 | lectin, galactoside-binding, soluble, 1 | 3956 | ENSG00000100097 |

  
  

| **Database:cellular component      &nbspName:ribonucleoprotein complex      &nbspID:GO:0030529** | | | | | | |
| --- | --- | --- | --- | --- | --- | --- |
| C=544; O=4; E=1.04; R=3.83; rawP=0.0195; adjP=0.0606 | | | | | | |
| Index | UserID | Value | Gene Symbol | Gene Name | EntrezGene | Ensembl |
| 1 | P61313 | NA | RPL15 | ribosomal protein L15 | 6138 | ENSG00000174748 |
| 2 | P67809 | NA | YBX1 | Y box binding protein 1 | 4904 | ENSG00000065978 |
| 3 | P13639 | NA | EEF2 | eukaryotic translation elongation factor 2 | 1938 | ENSG00000167658 |
| 4 | P11387 | NA | TOP1 | topoisomerase (DNA) I | 7150 | ENSG00000198900 |

  
  

| **Database:cellular component      &nbspName:microtubule      &nbspID:GO:0005874** | | | | | | |
| --- | --- | --- | --- | --- | --- | --- |
| C=334; O=3; E=0.64; R=4.68; rawP=0.0257; adjP=0.0758 | | | | | | |
| Index | UserID | Value | Gene Symbol | Gene Name | EntrezGene | Ensembl |
| 1 | P40227 | NA | CCT6A | chaperonin containing TCP1, subunit 6A (zeta 1) | 908 | ENSG00000146731 |
| 2 | P68371 | NA | TUBB4B | tubulin, beta 4B class IVb | 10383 | ENSG00000188229 |
| 3 | P27816 | NA | MAP4 | microtubule-associated protein 4 | 4134 | ENSG00000047849 |

  
  

| **Database:cellular component      &nbspName:proteinaceous extracellular matrix      &nbspID:GO:0005578** | | | | | | |
| --- | --- | --- | --- | --- | --- | --- |
| C=359; O=3; E=0.69; R=4.36; rawP=0.0309; adjP=0.0868 | | | | | | |
| Index | UserID | Value | Gene Symbol | Gene Name | EntrezGene | Ensembl |
| 1 | P02768 | NA | ALB | albumin | 213 | ENSG00000163631 |
| 2 | P05556 | NA | ITGB1 | integrin, beta 1 (fibronectin receptor, beta polypeptide, antigen CD29 includes MDF2, MSK12) | 3688 | ENSG00000150093 |
| 3 | P09382 | NA | LGALS1 | lectin, galactoside-binding, soluble, 1 | 3956 | ENSG00000100097 |

  
  
  
  

---

WebGestalt is currently developed and maintained by Jing Wang and Bing Zhang at the  Zhang Lab. Other people who have made significant contribution to the project include Dexter Duncan, Stefan Kirov, Zhiao Shi, and Jay Snoddy.  
  
**Funding credits:** NIH/NIAAA (U01 AA016662, U01 AA013512); NIH/NIDA (P01 DA015027); NIH/NIMH (P50 MH078028, P50 MH096972); NIH/NCI (U24 CA159988); NIH/NIGMS (R01 GM088822).
