## Supplementary material for "PGRMC1 phosphorylation and cell plasticity 1: glycolysis, mitochondria, tumor growth": File S8: final_pc_geneset_file_1439165549.html

  
  

| **Database:Pathway Commons pathway      &nbspName:Gene Expression      &nbspID:DB\_ID:531** | | | | | | |
| --- | --- | --- | --- | --- | --- | --- |
| C=379; O=9; E=0.60; R=15.05; rawP=4.85e-09; adjP=9.05e-08 | | | | | | |
| Index | UserID | Value | Gene Symbol | Gene Name | EntrezGene | Ensembl |
| 1 | P49589 | NA | CARS | cysteinyl-tRNA synthetase | 833 | ENSG00000110619 |
| 2 | P13639 | NA | EEF2 | eukaryotic translation elongation factor 2 | 1938 | ENSG00000167658 |
| 3 | P41091 | NA | EIF2S3 | eukaryotic translation initiation factor 2, subunit 3 gamma, 52kDa | 1968 | ENSG00000130741 |
| 4 | P61313 | NA | RPL15 | ribosomal protein L15 | 6138 | ENSG00000174748 |
| 5 | P60842 | NA | EIF4A1 | eukaryotic translation initiation factor 4A1 | 1973 | ENSG00000161960 |
| 6 | P67809 | NA | YBX1 | Y box binding protein 1 | 4904 | ENSG00000065978 |
| 7 | P68104 | NA | EEF1A1 | eukaryotic translation elongation factor 1 alpha 1 | 1915 | ENSG00000156508 |
| 8 | P29692 | NA | EEF1D | eukaryotic translation elongation factor 1 delta (guanine nucleotide exchange protein) | 1936 | ENSG00000104529 |
| 9 | P24534 | NA | EEF1B2 | eukaryotic translation elongation factor 1 beta 2 | 1933 | ENSG00000114942 |

  
  

| **Database:Pathway Commons pathway      &nbspName:Eukaryotic Translation Elongation      &nbspID:DB\_ID:713** | | | | | | |
| --- | --- | --- | --- | --- | --- | --- |
| C=86; O=5; E=0.14; R=36.85; rawP=2.24e-07; adjP=3.14e-06 | | | | | | |
| Index | UserID | Value | Gene Symbol | Gene Name | EntrezGene | Ensembl |
| 1 | P61313 | NA | RPL15 | ribosomal protein L15 | 6138 | ENSG00000174748 |
| 2 | P13639 | NA | EEF2 | eukaryotic translation elongation factor 2 | 1938 | ENSG00000167658 |
| 3 | P68104 | NA | EEF1A1 | eukaryotic translation elongation factor 1 alpha 1 | 1915 | ENSG00000156508 |
| 4 | P29692 | NA | EEF1D | eukaryotic translation elongation factor 1 delta (guanine nucleotide exchange protein) | 1936 | ENSG00000104529 |
| 5 | P24534 | NA | EEF1B2 | eukaryotic translation elongation factor 1 beta 2 | 1933 | ENSG00000114942 |

  
  

| **Database:Pathway Commons pathway      &nbspName:C-MYC pathway      &nbspID:DB\_ID:1467** | | | | | | |
| --- | --- | --- | --- | --- | --- | --- |
| C=149; O=4; E=0.24; R=17.02; rawP=8.57e-05; adjP=0.0010 | | | | | | |
| Index | UserID | Value | Gene Symbol | Gene Name | EntrezGene | Ensembl |
| 1 | P60842 | NA | EIF4A1 | eukaryotic translation initiation factor 4A1 | 1973 | ENSG00000161960 |
| 2 | P02786 | NA | TFRC | transferrin receptor (p90, CD71) | 7037 | ENSG00000072274 |
| 3 | P05556 | NA | ITGB1 | integrin, beta 1 (fibronectin receptor, beta polypeptide, antigen CD29 includes MDF2, MSK12) | 3688 | ENSG00000150093 |
| 4 | P09382 | NA | LGALS1 | lectin, galactoside-binding, soluble, 1 | 3956 | ENSG00000100097 |

  
  

| **Database:Pathway Commons pathway      &nbspName:Peptide chain elongation      &nbspID:DB\_ID:714** | | | | | | |
| --- | --- | --- | --- | --- | --- | --- |
| C=83; O=3; E=0.13; R=22.91; rawP=0.0003; adjP=0.0028 | | | | | | |
| Index | UserID | Value | Gene Symbol | Gene Name | EntrezGene | Ensembl |
| 1 | P61313 | NA | RPL15 | ribosomal protein L15 | 6138 | ENSG00000174748 |
| 2 | P13639 | NA | EEF2 | eukaryotic translation elongation factor 2 | 1938 | ENSG00000167658 |
| 3 | P68104 | NA | EEF1A1 | eukaryotic translation elongation factor 1 alpha 1 | 1915 | ENSG00000156508 |

  
  

| **Database:Pathway Commons pathway      &nbspName:Cap-dependent Translation Initiation      &nbspID:DB\_ID:724** | | | | | | |
| --- | --- | --- | --- | --- | --- | --- |
| C=111; O=3; E=0.18; R=17.13; rawP=0.0007; adjP=0.0036 | | | | | | |
| Index | UserID | Value | Gene Symbol | Gene Name | EntrezGene | Ensembl |
| 1 | P61313 | NA | RPL15 | ribosomal protein L15 | 6138 | ENSG00000174748 |
| 2 | P60842 | NA | EIF4A1 | eukaryotic translation initiation factor 4A1 | 1973 | ENSG00000161960 |
| 3 | P41091 | NA | EIF2S3 | eukaryotic translation initiation factor 2, subunit 3 gamma, 52kDa | 1968 | ENSG00000130741 |

  
  

| **Database:Pathway Commons pathway      &nbspName:L13a-mediated translational silencing of Ceruloplasmin expression      &nbspID:DB\_ID:635** | | | | | | |
| --- | --- | --- | --- | --- | --- | --- |
| C=103; O=3; E=0.16; R=18.46; rawP=0.0006; adjP=0.0036 | | | | | | |
| Index | UserID | Value | Gene Symbol | Gene Name | EntrezGene | Ensembl |
| 1 | P61313 | NA | RPL15 | ribosomal protein L15 | 6138 | ENSG00000174748 |
| 2 | P60842 | NA | EIF4A1 | eukaryotic translation initiation factor 4A1 | 1973 | ENSG00000161960 |
| 3 | P41091 | NA | EIF2S3 | eukaryotic translation initiation factor 2, subunit 3 gamma, 52kDa | 1968 | ENSG00000130741 |

  
  

| **Database:Pathway Commons pathway      &nbspName:Eukaryotic Translation Initiation      &nbspID:DB\_ID:723** | | | | | | |
| --- | --- | --- | --- | --- | --- | --- |
| C=111; O=3; E=0.18; R=17.13; rawP=0.0007; adjP=0.0036 | | | | | | |
| Index | UserID | Value | Gene Symbol | Gene Name | EntrezGene | Ensembl |
| 1 | P61313 | NA | RPL15 | ribosomal protein L15 | 6138 | ENSG00000174748 |
| 2 | P60842 | NA | EIF4A1 | eukaryotic translation initiation factor 4A1 | 1973 | ENSG00000161960 |
| 3 | P41091 | NA | EIF2S3 | eukaryotic translation initiation factor 2, subunit 3 gamma, 52kDa | 1968 | ENSG00000130741 |

  
  

| **Database:Pathway Commons pathway      &nbspName:GTP hydrolysis and joining of the 60S ribosomal subunit      &nbspID:DB\_ID:718** | | | | | | |
| --- | --- | --- | --- | --- | --- | --- |
| C=104; O=3; E=0.16; R=18.28; rawP=0.0006; adjP=0.0036 | | | | | | |
| Index | UserID | Value | Gene Symbol | Gene Name | EntrezGene | Ensembl |
| 1 | P61313 | NA | RPL15 | ribosomal protein L15 | 6138 | ENSG00000174748 |
| 2 | P60842 | NA | EIF4A1 | eukaryotic translation initiation factor 4A1 | 1973 | ENSG00000161960 |
| 3 | P41091 | NA | EIF2S3 | eukaryotic translation initiation factor 2, subunit 3 gamma, 52kDa | 1968 | ENSG00000130741 |

  
  

| **Database:Pathway Commons pathway      &nbspName:3' -UTR-mediated translational regulation      &nbspID:DB\_ID:644** | | | | | | |
| --- | --- | --- | --- | --- | --- | --- |
| C=103; O=3; E=0.16; R=18.46; rawP=0.0006; adjP=0.0036 | | | | | | |
| Index | UserID | Value | Gene Symbol | Gene Name | EntrezGene | Ensembl |
| 1 | P61313 | NA | RPL15 | ribosomal protein L15 | 6138 | ENSG00000174748 |
| 2 | P60842 | NA | EIF4A1 | eukaryotic translation initiation factor 4A1 | 1973 | ENSG00000161960 |
| 3 | P41091 | NA | EIF2S3 | eukaryotic translation initiation factor 2, subunit 3 gamma, 52kDa | 1968 | ENSG00000130741 |

  
  

| **Database:Pathway Commons pathway      &nbspName:TRAIL signaling pathway      &nbspID:DB\_ID:1480** | | | | | | |
| --- | --- | --- | --- | --- | --- | --- |
| C=1328; O=8; E=2.10; R=3.82; rawP=0.0008; adjP=0.0037 | | | | | | |
| Index | UserID | Value | Gene Symbol | Gene Name | EntrezGene | Ensembl |
| 1 | P08195 | NA | SLC3A2 | solute carrier family 3 (activators of dibasic and neutral amino acid transport), member 2 | 6520 | ENSG00000168003 |
| 2 | P13639 | NA | EEF2 | eukaryotic translation elongation factor 2 | 1938 | ENSG00000167658 |
| 3 | P05556 | NA | ITGB1 | integrin, beta 1 (fibronectin receptor, beta polypeptide, antigen CD29 includes MDF2, MSK12) | 3688 | ENSG00000150093 |
| 4 | P09382 | NA | LGALS1 | lectin, galactoside-binding, soluble, 1 | 3956 | ENSG00000100097 |
| 5 | P60842 | NA | EIF4A1 | eukaryotic translation initiation factor 4A1 | 1973 | ENSG00000161960 |
| 6 | P18206 | NA | VCL | vinculin | 7414 | ENSG00000035403 |
| 7 | P02786 | NA | TFRC | transferrin receptor (p90, CD71) | 7037 | ENSG00000072274 |
| 8 | P11387 | NA | TOP1 | topoisomerase (DNA) I | 7150 | ENSG00000198900 |

  
  

| **Database:Pathway Commons pathway      &nbspName:EGFR-dependent Endothelin signaling events      &nbspID:DB\_ID:1603** | | | | | | |
| --- | --- | --- | --- | --- | --- | --- |
| C=1289; O=7; E=2.03; R=3.44; rawP=0.0034; adjP=0.0043 | | | | | | |
| Index | UserID | Value | Gene Symbol | Gene Name | EntrezGene | Ensembl |
| 1 | P60842 | NA | EIF4A1 | eukaryotic translation initiation factor 4A1 | 1973 | ENSG00000161960 |
| 2 | P08195 | NA | SLC3A2 | solute carrier family 3 (activators of dibasic and neutral amino acid transport), member 2 | 6520 | ENSG00000168003 |
| 3 | P13639 | NA | EEF2 | eukaryotic translation elongation factor 2 | 1938 | ENSG00000167658 |
| 4 | P18206 | NA | VCL | vinculin | 7414 | ENSG00000035403 |
| 5 | P02786 | NA | TFRC | transferrin receptor (p90, CD71) | 7037 | ENSG00000072274 |
| 6 | P05556 | NA | ITGB1 | integrin, beta 1 (fibronectin receptor, beta polypeptide, antigen CD29 includes MDF2, MSK12) | 3688 | ENSG00000150093 |
| 7 | P09382 | NA | LGALS1 | lectin, galactoside-binding, soluble, 1 | 3956 | ENSG00000100097 |

  
  

| **Database:Pathway Commons pathway      &nbspName:PDGF receptor signaling network      &nbspID:DB\_ID:1497** | | | | | | |
| --- | --- | --- | --- | --- | --- | --- |
| C=1293; O=7; E=2.04; R=3.43; rawP=0.0035; adjP=0.0043 | | | | | | |
| Index | UserID | Value | Gene Symbol | Gene Name | EntrezGene | Ensembl |
| 1 | P60842 | NA | EIF4A1 | eukaryotic translation initiation factor 4A1 | 1973 | ENSG00000161960 |
| 2 | P08195 | NA | SLC3A2 | solute carrier family 3 (activators of dibasic and neutral amino acid transport), member 2 | 6520 | ENSG00000168003 |
| 3 | P13639 | NA | EEF2 | eukaryotic translation elongation factor 2 | 1938 | ENSG00000167658 |
| 4 | P18206 | NA | VCL | vinculin | 7414 | ENSG00000035403 |
| 5 | P02786 | NA | TFRC | transferrin receptor (p90, CD71) | 7037 | ENSG00000072274 |
| 6 | P05556 | NA | ITGB1 | integrin, beta 1 (fibronectin receptor, beta polypeptide, antigen CD29 includes MDF2, MSK12) | 3688 | ENSG00000150093 |
| 7 | P09382 | NA | LGALS1 | lectin, galactoside-binding, soluble, 1 | 3956 | ENSG00000100097 |

  
  

| **Database:Pathway Commons pathway      &nbspName:EGF receptor (ErbB1) signaling pathway      &nbspID:DB\_ID:1550** | | | | | | |
| --- | --- | --- | --- | --- | --- | --- |
| C=1288; O=7; E=2.03; R=3.44; rawP=0.0034; adjP=0.0043 | | | | | | |
| Index | UserID | Value | Gene Symbol | Gene Name | EntrezGene | Ensembl |
| 1 | P60842 | NA | EIF4A1 | eukaryotic translation initiation factor 4A1 | 1973 | ENSG00000161960 |
| 2 | P08195 | NA | SLC3A2 | solute carrier family 3 (activators of dibasic and neutral amino acid transport), member 2 | 6520 | ENSG00000168003 |
| 3 | P13639 | NA | EEF2 | eukaryotic translation elongation factor 2 | 1938 | ENSG00000167658 |
| 4 | P18206 | NA | VCL | vinculin | 7414 | ENSG00000035403 |
| 5 | P02786 | NA | TFRC | transferrin receptor (p90, CD71) | 7037 | ENSG00000072274 |
| 6 | P05556 | NA | ITGB1 | integrin, beta 1 (fibronectin receptor, beta polypeptide, antigen CD29 includes MDF2, MSK12) | 3688 | ENSG00000150093 |
| 7 | P09382 | NA | LGALS1 | lectin, galactoside-binding, soluble, 1 | 3956 | ENSG00000100097 |

  
  

| **Database:Pathway Commons pathway      &nbspName:VEGF and VEGFR signaling network      &nbspID:DB\_ID:1575** | | | | | | |
| --- | --- | --- | --- | --- | --- | --- |
| C=1304; O=7; E=2.06; R=3.40; rawP=0.0036; adjP=0.0043 | | | | | | |
| Index | UserID | Value | Gene Symbol | Gene Name | EntrezGene | Ensembl |
| 1 | P60842 | NA | EIF4A1 | eukaryotic translation initiation factor 4A1 | 1973 | ENSG00000161960 |
| 2 | P08195 | NA | SLC3A2 | solute carrier family 3 (activators of dibasic and neutral amino acid transport), member 2 | 6520 | ENSG00000168003 |
| 3 | P13639 | NA | EEF2 | eukaryotic translation elongation factor 2 | 1938 | ENSG00000167658 |
| 4 | P18206 | NA | VCL | vinculin | 7414 | ENSG00000035403 |
| 5 | P02786 | NA | TFRC | transferrin receptor (p90, CD71) | 7037 | ENSG00000072274 |
| 6 | P05556 | NA | ITGB1 | integrin, beta 1 (fibronectin receptor, beta polypeptide, antigen CD29 includes MDF2, MSK12) | 3688 | ENSG00000150093 |
| 7 | P09382 | NA | LGALS1 | lectin, galactoside-binding, soluble, 1 | 3956 | ENSG00000100097 |

  
  

| **Database:Pathway Commons pathway      &nbspName:PAR1-mediated thrombin signaling events      &nbspID:DB\_ID:1531** | | | | | | |
| --- | --- | --- | --- | --- | --- | --- |
| C=1299; O=7; E=2.05; R=3.42; rawP=0.0036; adjP=0.0043 | | | | | | |
| Index | UserID | Value | Gene Symbol | Gene Name | EntrezGene | Ensembl |
| 1 | P60842 | NA | EIF4A1 | eukaryotic translation initiation factor 4A1 | 1973 | ENSG00000161960 |
| 2 | P08195 | NA | SLC3A2 | solute carrier family 3 (activators of dibasic and neutral amino acid transport), member 2 | 6520 | ENSG00000168003 |
| 3 | P13639 | NA | EEF2 | eukaryotic translation elongation factor 2 | 1938 | ENSG00000167658 |
| 4 | P18206 | NA | VCL | vinculin | 7414 | ENSG00000035403 |
| 5 | P02786 | NA | TFRC | transferrin receptor (p90, CD71) | 7037 | ENSG00000072274 |
| 6 | P05556 | NA | ITGB1 | integrin, beta 1 (fibronectin receptor, beta polypeptide, antigen CD29 includes MDF2, MSK12) | 3688 | ENSG00000150093 |
| 7 | P09382 | NA | LGALS1 | lectin, galactoside-binding, soluble, 1 | 3956 | ENSG00000100097 |

  
  

| **Database:Pathway Commons pathway      &nbspName:Endothelins      &nbspID:DB\_ID:1619** | | | | | | |
| --- | --- | --- | --- | --- | --- | --- |
| C=1307; O=7; E=2.06; R=3.39; rawP=0.0037; adjP=0.0043 | | | | | | |
| Index | UserID | Value | Gene Symbol | Gene Name | EntrezGene | Ensembl |
| 1 | P60842 | NA | EIF4A1 | eukaryotic translation initiation factor 4A1 | 1973 | ENSG00000161960 |
| 2 | P08195 | NA | SLC3A2 | solute carrier family 3 (activators of dibasic and neutral amino acid transport), member 2 | 6520 | ENSG00000168003 |
| 3 | P13639 | NA | EEF2 | eukaryotic translation elongation factor 2 | 1938 | ENSG00000167658 |
| 4 | P18206 | NA | VCL | vinculin | 7414 | ENSG00000035403 |
| 5 | P02786 | NA | TFRC | transferrin receptor (p90, CD71) | 7037 | ENSG00000072274 |
| 6 | P05556 | NA | ITGB1 | integrin, beta 1 (fibronectin receptor, beta polypeptide, antigen CD29 includes MDF2, MSK12) | 3688 | ENSG00000150093 |
| 7 | P09382 | NA | LGALS1 | lectin, galactoside-binding, soluble, 1 | 3956 | ENSG00000100097 |

  
  

| **Database:Pathway Commons pathway      &nbspName:LKB1 signaling events      &nbspID:DB\_ID:1649** | | | | | | |
| --- | --- | --- | --- | --- | --- | --- |
| C=1308; O=7; E=2.06; R=3.39; rawP=0.0037; adjP=0.0043 | | | | | | |
| Index | UserID | Value | Gene Symbol | Gene Name | EntrezGene | Ensembl |
| 1 | P60842 | NA | EIF4A1 | eukaryotic translation initiation factor 4A1 | 1973 | ENSG00000161960 |
| 2 | P08195 | NA | SLC3A2 | solute carrier family 3 (activators of dibasic and neutral amino acid transport), member 2 | 6520 | ENSG00000168003 |
| 3 | P13639 | NA | EEF2 | eukaryotic translation elongation factor 2 | 1938 | ENSG00000167658 |
| 4 | P18206 | NA | VCL | vinculin | 7414 | ENSG00000035403 |
| 5 | P02786 | NA | TFRC | transferrin receptor (p90, CD71) | 7037 | ENSG00000072274 |
| 6 | P05556 | NA | ITGB1 | integrin, beta 1 (fibronectin receptor, beta polypeptide, antigen CD29 includes MDF2, MSK12) | 3688 | ENSG00000150093 |
| 7 | P09382 | NA | LGALS1 | lectin, galactoside-binding, soluble, 1 | 3956 | ENSG00000100097 |

  
  

| **Database:Pathway Commons pathway      &nbspName:IL5-mediated signaling events      &nbspID:DB\_ID:1627** | | | | | | |
| --- | --- | --- | --- | --- | --- | --- |
| C=1292; O=7; E=2.04; R=3.43; rawP=0.0034; adjP=0.0043 | | | | | | |
| Index | UserID | Value | Gene Symbol | Gene Name | EntrezGene | Ensembl |
| 1 | P60842 | NA | EIF4A1 | eukaryotic translation initiation factor 4A1 | 1973 | ENSG00000161960 |
| 2 | P08195 | NA | SLC3A2 | solute carrier family 3 (activators of dibasic and neutral amino acid transport), member 2 | 6520 | ENSG00000168003 |
| 3 | P13639 | NA | EEF2 | eukaryotic translation elongation factor 2 | 1938 | ENSG00000167658 |
| 4 | P18206 | NA | VCL | vinculin | 7414 | ENSG00000035403 |
| 5 | P02786 | NA | TFRC | transferrin receptor (p90, CD71) | 7037 | ENSG00000072274 |
| 6 | P05556 | NA | ITGB1 | integrin, beta 1 (fibronectin receptor, beta polypeptide, antigen CD29 includes MDF2, MSK12) | 3688 | ENSG00000150093 |
| 7 | P09382 | NA | LGALS1 | lectin, galactoside-binding, soluble, 1 | 3956 | ENSG00000100097 |

  
  

| **Database:Pathway Commons pathway      &nbspName:Arf6 signaling events      &nbspID:DB\_ID:1554** | | | | | | |
| --- | --- | --- | --- | --- | --- | --- |
| C=1288; O=7; E=2.03; R=3.44; rawP=0.0034; adjP=0.0043 | | | | | | |
| Index | UserID | Value | Gene Symbol | Gene Name | EntrezGene | Ensembl |
| 1 | P60842 | NA | EIF4A1 | eukaryotic translation initiation factor 4A1 | 1973 | ENSG00000161960 |
| 2 | P08195 | NA | SLC3A2 | solute carrier family 3 (activators of dibasic and neutral amino acid transport), member 2 | 6520 | ENSG00000168003 |
| 3 | P13639 | NA | EEF2 | eukaryotic translation elongation factor 2 | 1938 | ENSG00000167658 |
| 4 | P18206 | NA | VCL | vinculin | 7414 | ENSG00000035403 |
| 5 | P02786 | NA | TFRC | transferrin receptor (p90, CD71) | 7037 | ENSG00000072274 |
| 6 | P05556 | NA | ITGB1 | integrin, beta 1 (fibronectin receptor, beta polypeptide, antigen CD29 includes MDF2, MSK12) | 3688 | ENSG00000150093 |
| 7 | P09382 | NA | LGALS1 | lectin, galactoside-binding, soluble, 1 | 3956 | ENSG00000100097 |

  
  

| **Database:Pathway Commons pathway      &nbspName:Signaling events mediated by Hepatocyte Growth Factor Receptor (c-Met)      &nbspID:DB\_ID:1491** | | | | | | |
| --- | --- | --- | --- | --- | --- | --- |
| C=1293; O=7; E=2.04; R=3.43; rawP=0.0035; adjP=0.0043 | | | | | | |
| Index | UserID | Value | Gene Symbol | Gene Name | EntrezGene | Ensembl |
| 1 | P60842 | NA | EIF4A1 | eukaryotic translation initiation factor 4A1 | 1973 | ENSG00000161960 |
| 2 | P08195 | NA | SLC3A2 | solute carrier family 3 (activators of dibasic and neutral amino acid transport), member 2 | 6520 | ENSG00000168003 |
| 3 | P13639 | NA | EEF2 | eukaryotic translation elongation factor 2 | 1938 | ENSG00000167658 |
| 4 | P18206 | NA | VCL | vinculin | 7414 | ENSG00000035403 |
| 5 | P02786 | NA | TFRC | transferrin receptor (p90, CD71) | 7037 | ENSG00000072274 |
| 6 | P05556 | NA | ITGB1 | integrin, beta 1 (fibronectin receptor, beta polypeptide, antigen CD29 includes MDF2, MSK12) | 3688 | ENSG00000150093 |
| 7 | P09382 | NA | LGALS1 | lectin, galactoside-binding, soluble, 1 | 3956 | ENSG00000100097 |

  
  

| **Database:Pathway Commons pathway      &nbspName:Nectin adhesion pathway      &nbspID:DB\_ID:1472** | | | | | | |
| --- | --- | --- | --- | --- | --- | --- |
| C=1295; O=7; E=2.04; R=3.43; rawP=0.0035; adjP=0.0043 | | | | | | |
| Index | UserID | Value | Gene Symbol | Gene Name | EntrezGene | Ensembl |
| 1 | P60842 | NA | EIF4A1 | eukaryotic translation initiation factor 4A1 | 1973 | ENSG00000161960 |
| 2 | P08195 | NA | SLC3A2 | solute carrier family 3 (activators of dibasic and neutral amino acid transport), member 2 | 6520 | ENSG00000168003 |
| 3 | P13639 | NA | EEF2 | eukaryotic translation elongation factor 2 | 1938 | ENSG00000167658 |
| 4 | P18206 | NA | VCL | vinculin | 7414 | ENSG00000035403 |
| 5 | P02786 | NA | TFRC | transferrin receptor (p90, CD71) | 7037 | ENSG00000072274 |
| 6 | P05556 | NA | ITGB1 | integrin, beta 1 (fibronectin receptor, beta polypeptide, antigen CD29 includes MDF2, MSK12) | 3688 | ENSG00000150093 |
| 7 | P09382 | NA | LGALS1 | lectin, galactoside-binding, soluble, 1 | 3956 | ENSG00000100097 |

  
  

| **Database:Pathway Commons pathway      &nbspName:Internalization of ErbB1      &nbspID:DB\_ID:1509** | | | | | | |
| --- | --- | --- | --- | --- | --- | --- |
| C=1288; O=7; E=2.03; R=3.44; rawP=0.0034; adjP=0.0043 | | | | | | |
| Index | UserID | Value | Gene Symbol | Gene Name | EntrezGene | Ensembl |
| 1 | P60842 | NA | EIF4A1 | eukaryotic translation initiation factor 4A1 | 1973 | ENSG00000161960 |
| 2 | P08195 | NA | SLC3A2 | solute carrier family 3 (activators of dibasic and neutral amino acid transport), member 2 | 6520 | ENSG00000168003 |
| 3 | P13639 | NA | EEF2 | eukaryotic translation elongation factor 2 | 1938 | ENSG00000167658 |
| 4 | P18206 | NA | VCL | vinculin | 7414 | ENSG00000035403 |
| 5 | P02786 | NA | TFRC | transferrin receptor (p90, CD71) | 7037 | ENSG00000072274 |
| 6 | P05556 | NA | ITGB1 | integrin, beta 1 (fibronectin receptor, beta polypeptide, antigen CD29 includes MDF2, MSK12) | 3688 | ENSG00000150093 |
| 7 | P09382 | NA | LGALS1 | lectin, galactoside-binding, soluble, 1 | 3956 | ENSG00000100097 |

  
  

| **Database:Pathway Commons pathway      &nbspName:Urokinase-type plasminogen activator (uPA) and uPAR-mediated signaling      &nbspID:DB\_ID:1519** | | | | | | |
| --- | --- | --- | --- | --- | --- | --- |
| C=1288; O=7; E=2.03; R=3.44; rawP=0.0034; adjP=0.0043 | | | | | | |
| Index | UserID | Value | Gene Symbol | Gene Name | EntrezGene | Ensembl |
| 1 | P60842 | NA | EIF4A1 | eukaryotic translation initiation factor 4A1 | 1973 | ENSG00000161960 |
| 2 | P08195 | NA | SLC3A2 | solute carrier family 3 (activators of dibasic and neutral amino acid transport), member 2 | 6520 | ENSG00000168003 |
| 3 | P13639 | NA | EEF2 | eukaryotic translation elongation factor 2 | 1938 | ENSG00000167658 |
| 4 | P18206 | NA | VCL | vinculin | 7414 | ENSG00000035403 |
| 5 | P02786 | NA | TFRC | transferrin receptor (p90, CD71) | 7037 | ENSG00000072274 |
| 6 | P05556 | NA | ITGB1 | integrin, beta 1 (fibronectin receptor, beta polypeptide, antigen CD29 includes MDF2, MSK12) | 3688 | ENSG00000150093 |
| 7 | P09382 | NA | LGALS1 | lectin, galactoside-binding, soluble, 1 | 3956 | ENSG00000100097 |

  
  

| **Database:Pathway Commons pathway      &nbspName:Metabolism      &nbspID:DB\_ID:634** | | | | | | |
| --- | --- | --- | --- | --- | --- | --- |
| C=824; O=6; E=1.30; R=4.62; rawP=0.0016; adjP=0.0043 | | | | | | |
| Index | UserID | Value | Gene Symbol | Gene Name | EntrezGene | Ensembl |
| 1 | P61313 | NA | RPL15 | ribosomal protein L15 | 6138 | ENSG00000174748 |
| 2 | P17174 | NA | GOT1 | glutamic-oxaloacetic transaminase 1, soluble (aspartate aminotransferase 1) | 2805 | ENSG00000120053 |
| 3 | P60842 | NA | EIF4A1 | eukaryotic translation initiation factor 4A1 | 1973 | ENSG00000161960 |
| 4 | P22102 | NA | GART | phosphoribosylglycinamide formyltransferase, phosphoribosylglycinamide synthetase, phosphoribosylaminoimidazole synthetase | 2618 | ENSG00000159131 |
| 5 | P00491 | NA | PNP | purine nucleoside phosphorylase | 4860 | ENSG00000198805 |
| 6 | P00352 | NA | ALDH1A1 | aldehyde dehydrogenase 1 family, member A1 | 216 | ENSG00000165092 |

  
  

| **Database:Pathway Commons pathway      &nbspName:Class I PI3K signaling events      &nbspID:DB\_ID:1553** | | | | | | |
| --- | --- | --- | --- | --- | --- | --- |
| C=1288; O=7; E=2.03; R=3.44; rawP=0.0034; adjP=0.0043 | | | | | | |
| Index | UserID | Value | Gene Symbol | Gene Name | EntrezGene | Ensembl |
| 1 | P60842 | NA | EIF4A1 | eukaryotic translation initiation factor 4A1 | 1973 | ENSG00000161960 |
| 2 | P08195 | NA | SLC3A2 | solute carrier family 3 (activators of dibasic and neutral amino acid transport), member 2 | 6520 | ENSG00000168003 |
| 3 | P13639 | NA | EEF2 | eukaryotic translation elongation factor 2 | 1938 | ENSG00000167658 |
| 4 | P18206 | NA | VCL | vinculin | 7414 | ENSG00000035403 |
| 5 | P02786 | NA | TFRC | transferrin receptor (p90, CD71) | 7037 | ENSG00000072274 |
| 6 | P05556 | NA | ITGB1 | integrin, beta 1 (fibronectin receptor, beta polypeptide, antigen CD29 includes MDF2, MSK12) | 3688 | ENSG00000150093 |
| 7 | P09382 | NA | LGALS1 | lectin, galactoside-binding, soluble, 1 | 3956 | ENSG00000100097 |

  
  

| **Database:Pathway Commons pathway      &nbspName:Alpha9 beta1 integrin signaling events      &nbspID:DB\_ID:1578** | | | | | | |
| --- | --- | --- | --- | --- | --- | --- |
| C=1305; O=7; E=2.06; R=3.40; rawP=0.0036; adjP=0.0043 | | | | | | |
| Index | UserID | Value | Gene Symbol | Gene Name | EntrezGene | Ensembl |
| 1 | P60842 | NA | EIF4A1 | eukaryotic translation initiation factor 4A1 | 1973 | ENSG00000161960 |
| 2 | P08195 | NA | SLC3A2 | solute carrier family 3 (activators of dibasic and neutral amino acid transport), member 2 | 6520 | ENSG00000168003 |
| 3 | P13639 | NA | EEF2 | eukaryotic translation elongation factor 2 | 1938 | ENSG00000167658 |
| 4 | P18206 | NA | VCL | vinculin | 7414 | ENSG00000035403 |
| 5 | P02786 | NA | TFRC | transferrin receptor (p90, CD71) | 7037 | ENSG00000072274 |
| 6 | P05556 | NA | ITGB1 | integrin, beta 1 (fibronectin receptor, beta polypeptide, antigen CD29 includes MDF2, MSK12) | 3688 | ENSG00000150093 |
| 7 | P09382 | NA | LGALS1 | lectin, galactoside-binding, soluble, 1 | 3956 | ENSG00000100097 |

  
  

| **Database:Pathway Commons pathway      &nbspName:IFN-gamma pathway      &nbspID:DB\_ID:1529** | | | | | | |
| --- | --- | --- | --- | --- | --- | --- |
| C=1296; O=7; E=2.04; R=3.42; rawP=0.0035; adjP=0.0043 | | | | | | |
| Index | UserID | Value | Gene Symbol | Gene Name | EntrezGene | Ensembl |
| 1 | P60842 | NA | EIF4A1 | eukaryotic translation initiation factor 4A1 | 1973 | ENSG00000161960 |
| 2 | P08195 | NA | SLC3A2 | solute carrier family 3 (activators of dibasic and neutral amino acid transport), member 2 | 6520 | ENSG00000168003 |
| 3 | P13639 | NA | EEF2 | eukaryotic translation elongation factor 2 | 1938 | ENSG00000167658 |
| 4 | P18206 | NA | VCL | vinculin | 7414 | ENSG00000035403 |
| 5 | P02786 | NA | TFRC | transferrin receptor (p90, CD71) | 7037 | ENSG00000072274 |
| 6 | P05556 | NA | ITGB1 | integrin, beta 1 (fibronectin receptor, beta polypeptide, antigen CD29 includes MDF2, MSK12) | 3688 | ENSG00000150093 |
| 7 | P09382 | NA | LGALS1 | lectin, galactoside-binding, soluble, 1 | 3956 | ENSG00000100097 |

  
  

| **Database:Pathway Commons pathway      &nbspName:Plasma membrane estrogen receptor signaling      &nbspID:DB\_ID:1556** | | | | | | |
| --- | --- | --- | --- | --- | --- | --- |
| C=1301; O=7; E=2.05; R=3.41; rawP=0.0036; adjP=0.0043 | | | | | | |
| Index | UserID | Value | Gene Symbol | Gene Name | EntrezGene | Ensembl |
| 1 | P60842 | NA | EIF4A1 | eukaryotic translation initiation factor 4A1 | 1973 | ENSG00000161960 |
| 2 | P08195 | NA | SLC3A2 | solute carrier family 3 (activators of dibasic and neutral amino acid transport), member 2 | 6520 | ENSG00000168003 |
| 3 | P13639 | NA | EEF2 | eukaryotic translation elongation factor 2 | 1938 | ENSG00000167658 |
| 4 | P18206 | NA | VCL | vinculin | 7414 | ENSG00000035403 |
| 5 | P02786 | NA | TFRC | transferrin receptor (p90, CD71) | 7037 | ENSG00000072274 |
| 6 | P05556 | NA | ITGB1 | integrin, beta 1 (fibronectin receptor, beta polypeptide, antigen CD29 includes MDF2, MSK12) | 3688 | ENSG00000150093 |
| 7 | P09382 | NA | LGALS1 | lectin, galactoside-binding, soluble, 1 | 3956 | ENSG00000100097 |

  
  

| **Database:Pathway Commons pathway      &nbspName:Signaling events mediated by VEGFR1 and VEGFR2      &nbspID:DB\_ID:1516** | | | | | | |
| --- | --- | --- | --- | --- | --- | --- |
| C=1296; O=7; E=2.04; R=3.42; rawP=0.0035; adjP=0.0043 | | | | | | |
| Index | UserID | Value | Gene Symbol | Gene Name | EntrezGene | Ensembl |
| 1 | P60842 | NA | EIF4A1 | eukaryotic translation initiation factor 4A1 | 1973 | ENSG00000161960 |
| 2 | P08195 | NA | SLC3A2 | solute carrier family 3 (activators of dibasic and neutral amino acid transport), member 2 | 6520 | ENSG00000168003 |
| 3 | P13639 | NA | EEF2 | eukaryotic translation elongation factor 2 | 1938 | ENSG00000167658 |
| 4 | P18206 | NA | VCL | vinculin | 7414 | ENSG00000035403 |
| 5 | P02786 | NA | TFRC | transferrin receptor (p90, CD71) | 7037 | ENSG00000072274 |
| 6 | P05556 | NA | ITGB1 | integrin, beta 1 (fibronectin receptor, beta polypeptide, antigen CD29 includes MDF2, MSK12) | 3688 | ENSG00000150093 |
| 7 | P09382 | NA | LGALS1 | lectin, galactoside-binding, soluble, 1 | 3956 | ENSG00000100097 |

  
  

| **Database:Pathway Commons pathway      &nbspName:Arf6 downstream pathway      &nbspID:DB\_ID:1585** | | | | | | |
| --- | --- | --- | --- | --- | --- | --- |
| C=1288; O=7; E=2.03; R=3.44; rawP=0.0034; adjP=0.0043 | | | | | | |
| Index | UserID | Value | Gene Symbol | Gene Name | EntrezGene | Ensembl |
| 1 | P60842 | NA | EIF4A1 | eukaryotic translation initiation factor 4A1 | 1973 | ENSG00000161960 |
| 2 | P08195 | NA | SLC3A2 | solute carrier family 3 (activators of dibasic and neutral amino acid transport), member 2 | 6520 | ENSG00000168003 |
| 3 | P13639 | NA | EEF2 | eukaryotic translation elongation factor 2 | 1938 | ENSG00000167658 |
| 4 | P18206 | NA | VCL | vinculin | 7414 | ENSG00000035403 |
| 5 | P02786 | NA | TFRC | transferrin receptor (p90, CD71) | 7037 | ENSG00000072274 |
| 6 | P05556 | NA | ITGB1 | integrin, beta 1 (fibronectin receptor, beta polypeptide, antigen CD29 includes MDF2, MSK12) | 3688 | ENSG00000150093 |
| 7 | P09382 | NA | LGALS1 | lectin, galactoside-binding, soluble, 1 | 3956 | ENSG00000100097 |

  
  

| **Database:Pathway Commons pathway      &nbspName:Sphingosine 1-phosphate (S1P) pathway      &nbspID:DB\_ID:1635** | | | | | | |
| --- | --- | --- | --- | --- | --- | --- |
| C=1311; O=7; E=2.07; R=3.38; rawP=0.0037; adjP=0.0043 | | | | | | |
| Index | UserID | Value | Gene Symbol | Gene Name | EntrezGene | Ensembl |
| 1 | P60842 | NA | EIF4A1 | eukaryotic translation initiation factor 4A1 | 1973 | ENSG00000161960 |
| 2 | P08195 | NA | SLC3A2 | solute carrier family 3 (activators of dibasic and neutral amino acid transport), member 2 | 6520 | ENSG00000168003 |
| 3 | P13639 | NA | EEF2 | eukaryotic translation elongation factor 2 | 1938 | ENSG00000167658 |
| 4 | P18206 | NA | VCL | vinculin | 7414 | ENSG00000035403 |
| 5 | P02786 | NA | TFRC | transferrin receptor (p90, CD71) | 7037 | ENSG00000072274 |
| 6 | P05556 | NA | ITGB1 | integrin, beta 1 (fibronectin receptor, beta polypeptide, antigen CD29 includes MDF2, MSK12) | 3688 | ENSG00000150093 |
| 7 | P09382 | NA | LGALS1 | lectin, galactoside-binding, soluble, 1 | 3956 | ENSG00000100097 |

  
  

| **Database:Pathway Commons pathway      &nbspName:Signaling events mediated by focal adhesion kinase      &nbspID:DB\_ID:1574** | | | | | | |
| --- | --- | --- | --- | --- | --- | --- |
| C=1288; O=7; E=2.03; R=3.44; rawP=0.0034; adjP=0.0043 | | | | | | |
| Index | UserID | Value | Gene Symbol | Gene Name | EntrezGene | Ensembl |
| 1 | P60842 | NA | EIF4A1 | eukaryotic translation initiation factor 4A1 | 1973 | ENSG00000161960 |
| 2 | P08195 | NA | SLC3A2 | solute carrier family 3 (activators of dibasic and neutral amino acid transport), member 2 | 6520 | ENSG00000168003 |
| 3 | P13639 | NA | EEF2 | eukaryotic translation elongation factor 2 | 1938 | ENSG00000167658 |
| 4 | P18206 | NA | VCL | vinculin | 7414 | ENSG00000035403 |
| 5 | P02786 | NA | TFRC | transferrin receptor (p90, CD71) | 7037 | ENSG00000072274 |
| 6 | P05556 | NA | ITGB1 | integrin, beta 1 (fibronectin receptor, beta polypeptide, antigen CD29 includes MDF2, MSK12) | 3688 | ENSG00000150093 |
| 7 | P09382 | NA | LGALS1 | lectin, galactoside-binding, soluble, 1 | 3956 | ENSG00000100097 |

  
  

| **Database:Pathway Commons pathway      &nbspName:ErbB receptor signaling network      &nbspID:DB\_ID:1573** | | | | | | |
| --- | --- | --- | --- | --- | --- | --- |
| C=1312; O=7; E=2.07; R=3.38; rawP=0.0038; adjP=0.0043 | | | | | | |
| Index | UserID | Value | Gene Symbol | Gene Name | EntrezGene | Ensembl |
| 1 | P60842 | NA | EIF4A1 | eukaryotic translation initiation factor 4A1 | 1973 | ENSG00000161960 |
| 2 | P08195 | NA | SLC3A2 | solute carrier family 3 (activators of dibasic and neutral amino acid transport), member 2 | 6520 | ENSG00000168003 |
| 3 | P13639 | NA | EEF2 | eukaryotic translation elongation factor 2 | 1938 | ENSG00000167658 |
| 4 | P18206 | NA | VCL | vinculin | 7414 | ENSG00000035403 |
| 5 | P02786 | NA | TFRC | transferrin receptor (p90, CD71) | 7037 | ENSG00000072274 |
| 6 | P05556 | NA | ITGB1 | integrin, beta 1 (fibronectin receptor, beta polypeptide, antigen CD29 includes MDF2, MSK12) | 3688 | ENSG00000150093 |
| 7 | P09382 | NA | LGALS1 | lectin, galactoside-binding, soluble, 1 | 3956 | ENSG00000100097 |

  
  

| **Database:Pathway Commons pathway      &nbspName:IL3-mediated signaling events      &nbspID:DB\_ID:1564** | | | | | | |
| --- | --- | --- | --- | --- | --- | --- |
| C=1295; O=7; E=2.04; R=3.43; rawP=0.0035; adjP=0.0043 | | | | | | |
| Index | UserID | Value | Gene Symbol | Gene Name | EntrezGene | Ensembl |
| 1 | P60842 | NA | EIF4A1 | eukaryotic translation initiation factor 4A1 | 1973 | ENSG00000161960 |
| 2 | P08195 | NA | SLC3A2 | solute carrier family 3 (activators of dibasic and neutral amino acid transport), member 2 | 6520 | ENSG00000168003 |
| 3 | P13639 | NA | EEF2 | eukaryotic translation elongation factor 2 | 1938 | ENSG00000167658 |
| 4 | P18206 | NA | VCL | vinculin | 7414 | ENSG00000035403 |
| 5 | P02786 | NA | TFRC | transferrin receptor (p90, CD71) | 7037 | ENSG00000072274 |
| 6 | P05556 | NA | ITGB1 | integrin, beta 1 (fibronectin receptor, beta polypeptide, antigen CD29 includes MDF2, MSK12) | 3688 | ENSG00000150093 |
| 7 | P09382 | NA | LGALS1 | lectin, galactoside-binding, soluble, 1 | 3956 | ENSG00000100097 |

  
  

| **Database:Pathway Commons pathway      &nbspName:Syndecan-1-mediated signaling events      &nbspID:DB\_ID:1454** | | | | | | |
| --- | --- | --- | --- | --- | --- | --- |
| C=1300; O=7; E=2.05; R=3.41; rawP=0.0036; adjP=0.0043 | | | | | | |
| Index | UserID | Value | Gene Symbol | Gene Name | EntrezGene | Ensembl |
| 1 | P60842 | NA | EIF4A1 | eukaryotic translation initiation factor 4A1 | 1973 | ENSG00000161960 |
| 2 | P08195 | NA | SLC3A2 | solute carrier family 3 (activators of dibasic and neutral amino acid transport), member 2 | 6520 | ENSG00000168003 |
| 3 | P13639 | NA | EEF2 | eukaryotic translation elongation factor 2 | 1938 | ENSG00000167658 |
| 4 | P18206 | NA | VCL | vinculin | 7414 | ENSG00000035403 |
| 5 | P02786 | NA | TFRC | transferrin receptor (p90, CD71) | 7037 | ENSG00000072274 |
| 6 | P05556 | NA | ITGB1 | integrin, beta 1 (fibronectin receptor, beta polypeptide, antigen CD29 includes MDF2, MSK12) | 3688 | ENSG00000150093 |
| 7 | P09382 | NA | LGALS1 | lectin, galactoside-binding, soluble, 1 | 3956 | ENSG00000100097 |

  
  

| **Database:Pathway Commons pathway      &nbspName:S1P1 pathway      &nbspID:DB\_ID:1594** | | | | | | |
| --- | --- | --- | --- | --- | --- | --- |
| C=1288; O=7; E=2.03; R=3.44; rawP=0.0034; adjP=0.0043 | | | | | | |
| Index | UserID | Value | Gene Symbol | Gene Name | EntrezGene | Ensembl |
| 1 | P60842 | NA | EIF4A1 | eukaryotic translation initiation factor 4A1 | 1973 | ENSG00000161960 |
| 2 | P08195 | NA | SLC3A2 | solute carrier family 3 (activators of dibasic and neutral amino acid transport), member 2 | 6520 | ENSG00000168003 |
| 3 | P13639 | NA | EEF2 | eukaryotic translation elongation factor 2 | 1938 | ENSG00000167658 |
| 4 | P18206 | NA | VCL | vinculin | 7414 | ENSG00000035403 |
| 5 | P02786 | NA | TFRC | transferrin receptor (p90, CD71) | 7037 | ENSG00000072274 |
| 6 | P05556 | NA | ITGB1 | integrin, beta 1 (fibronectin receptor, beta polypeptide, antigen CD29 includes MDF2, MSK12) | 3688 | ENSG00000150093 |
| 7 | P09382 | NA | LGALS1 | lectin, galactoside-binding, soluble, 1 | 3956 | ENSG00000100097 |

  
  

| **Database:Pathway Commons pathway      &nbspName:PDGFR-beta signaling pathway      &nbspID:DB\_ID:1540** | | | | | | |
| --- | --- | --- | --- | --- | --- | --- |
| C=1288; O=7; E=2.03; R=3.44; rawP=0.0034; adjP=0.0043 | | | | | | |
| Index | UserID | Value | Gene Symbol | Gene Name | EntrezGene | Ensembl |
| 1 | P60842 | NA | EIF4A1 | eukaryotic translation initiation factor 4A1 | 1973 | ENSG00000161960 |
| 2 | P08195 | NA | SLC3A2 | solute carrier family 3 (activators of dibasic and neutral amino acid transport), member 2 | 6520 | ENSG00000168003 |
| 3 | P13639 | NA | EEF2 | eukaryotic translation elongation factor 2 | 1938 | ENSG00000167658 |
| 4 | P18206 | NA | VCL | vinculin | 7414 | ENSG00000035403 |
| 5 | P02786 | NA | TFRC | transferrin receptor (p90, CD71) | 7037 | ENSG00000072274 |
| 6 | P05556 | NA | ITGB1 | integrin, beta 1 (fibronectin receptor, beta polypeptide, antigen CD29 includes MDF2, MSK12) | 3688 | ENSG00000150093 |
| 7 | P09382 | NA | LGALS1 | lectin, galactoside-binding, soluble, 1 | 3956 | ENSG00000100097 |

  
  

| **Database:Pathway Commons pathway      &nbspName:Thrombin/protease-activated receptor (PAR) pathway      &nbspID:DB\_ID:1552** | | | | | | |
| --- | --- | --- | --- | --- | --- | --- |
| C=1300; O=7; E=2.05; R=3.41; rawP=0.0036; adjP=0.0043 | | | | | | |
| Index | UserID | Value | Gene Symbol | Gene Name | EntrezGene | Ensembl |
| 1 | P60842 | NA | EIF4A1 | eukaryotic translation initiation factor 4A1 | 1973 | ENSG00000161960 |
| 2 | P08195 | NA | SLC3A2 | solute carrier family 3 (activators of dibasic and neutral amino acid transport), member 2 | 6520 | ENSG00000168003 |
| 3 | P13639 | NA | EEF2 | eukaryotic translation elongation factor 2 | 1938 | ENSG00000167658 |
| 4 | P18206 | NA | VCL | vinculin | 7414 | ENSG00000035403 |
| 5 | P02786 | NA | TFRC | transferrin receptor (p90, CD71) | 7037 | ENSG00000072274 |
| 6 | P05556 | NA | ITGB1 | integrin, beta 1 (fibronectin receptor, beta polypeptide, antigen CD29 includes MDF2, MSK12) | 3688 | ENSG00000150093 |
| 7 | P09382 | NA | LGALS1 | lectin, galactoside-binding, soluble, 1 | 3956 | ENSG00000100097 |

  
  

| **Database:Pathway Commons pathway      &nbspName:IGF1 pathway      &nbspID:DB\_ID:1482** | | | | | | |
| --- | --- | --- | --- | --- | --- | --- |
| C=1291; O=7; E=2.04; R=3.44; rawP=0.0034; adjP=0.0043 | | | | | | |
| Index | UserID | Value | Gene Symbol | Gene Name | EntrezGene | Ensembl |
| 1 | P60842 | NA | EIF4A1 | eukaryotic translation initiation factor 4A1 | 1973 | ENSG00000161960 |
| 2 | P08195 | NA | SLC3A2 | solute carrier family 3 (activators of dibasic and neutral amino acid transport), member 2 | 6520 | ENSG00000168003 |
| 3 | P13639 | NA | EEF2 | eukaryotic translation elongation factor 2 | 1938 | ENSG00000167658 |
| 4 | P18206 | NA | VCL | vinculin | 7414 | ENSG00000035403 |
| 5 | P02786 | NA | TFRC | transferrin receptor (p90, CD71) | 7037 | ENSG00000072274 |
| 6 | P05556 | NA | ITGB1 | integrin, beta 1 (fibronectin receptor, beta polypeptide, antigen CD29 includes MDF2, MSK12) | 3688 | ENSG00000150093 |
| 7 | P09382 | NA | LGALS1 | lectin, galactoside-binding, soluble, 1 | 3956 | ENSG00000100097 |

  
  

| **Database:Pathway Commons pathway      &nbspName:Insulin Pathway      &nbspID:DB\_ID:1466** | | | | | | |
| --- | --- | --- | --- | --- | --- | --- |
| C=1288; O=7; E=2.03; R=3.44; rawP=0.0034; adjP=0.0043 | | | | | | |
| Index | UserID | Value | Gene Symbol | Gene Name | EntrezGene | Ensembl |
| 1 | P60842 | NA | EIF4A1 | eukaryotic translation initiation factor 4A1 | 1973 | ENSG00000161960 |
| 2 | P08195 | NA | SLC3A2 | solute carrier family 3 (activators of dibasic and neutral amino acid transport), member 2 | 6520 | ENSG00000168003 |
| 3 | P13639 | NA | EEF2 | eukaryotic translation elongation factor 2 | 1938 | ENSG00000167658 |
| 4 | P18206 | NA | VCL | vinculin | 7414 | ENSG00000035403 |
| 5 | P02786 | NA | TFRC | transferrin receptor (p90, CD71) | 7037 | ENSG00000072274 |
| 6 | P05556 | NA | ITGB1 | integrin, beta 1 (fibronectin receptor, beta polypeptide, antigen CD29 includes MDF2, MSK12) | 3688 | ENSG00000150093 |
| 7 | P09382 | NA | LGALS1 | lectin, galactoside-binding, soluble, 1 | 3956 | ENSG00000100097 |

  
  

| **Database:Pathway Commons pathway      &nbspName:ErbB1 downstream signaling      &nbspID:DB\_ID:1602** | | | | | | |
| --- | --- | --- | --- | --- | --- | --- |
| C=1288; O=7; E=2.03; R=3.44; rawP=0.0034; adjP=0.0043 | | | | | | |
| Index | UserID | Value | Gene Symbol | Gene Name | EntrezGene | Ensembl |
| 1 | P60842 | NA | EIF4A1 | eukaryotic translation initiation factor 4A1 | 1973 | ENSG00000161960 |
| 2 | P08195 | NA | SLC3A2 | solute carrier family 3 (activators of dibasic and neutral amino acid transport), member 2 | 6520 | ENSG00000168003 |
| 3 | P13639 | NA | EEF2 | eukaryotic translation elongation factor 2 | 1938 | ENSG00000167658 |
| 4 | P18206 | NA | VCL | vinculin | 7414 | ENSG00000035403 |
| 5 | P02786 | NA | TFRC | transferrin receptor (p90, CD71) | 7037 | ENSG00000072274 |
| 6 | P05556 | NA | ITGB1 | integrin, beta 1 (fibronectin receptor, beta polypeptide, antigen CD29 includes MDF2, MSK12) | 3688 | ENSG00000150093 |
| 7 | P09382 | NA | LGALS1 | lectin, galactoside-binding, soluble, 1 | 3956 | ENSG00000100097 |

  
  

| **Database:Pathway Commons pathway      &nbspName:mTOR signaling pathway      &nbspID:DB\_ID:1571** | | | | | | |
| --- | --- | --- | --- | --- | --- | --- |
| C=1288; O=7; E=2.03; R=3.44; rawP=0.0034; adjP=0.0043 | | | | | | |
| Index | UserID | Value | Gene Symbol | Gene Name | EntrezGene | Ensembl |
| 1 | P60842 | NA | EIF4A1 | eukaryotic translation initiation factor 4A1 | 1973 | ENSG00000161960 |
| 2 | P08195 | NA | SLC3A2 | solute carrier family 3 (activators of dibasic and neutral amino acid transport), member 2 | 6520 | ENSG00000168003 |
| 3 | P13639 | NA | EEF2 | eukaryotic translation elongation factor 2 | 1938 | ENSG00000167658 |
| 4 | P18206 | NA | VCL | vinculin | 7414 | ENSG00000035403 |
| 5 | P02786 | NA | TFRC | transferrin receptor (p90, CD71) | 7037 | ENSG00000072274 |
| 6 | P05556 | NA | ITGB1 | integrin, beta 1 (fibronectin receptor, beta polypeptide, antigen CD29 includes MDF2, MSK12) | 3688 | ENSG00000150093 |
| 7 | P09382 | NA | LGALS1 | lectin, galactoside-binding, soluble, 1 | 3956 | ENSG00000100097 |

  
  

| **Database:Pathway Commons pathway      &nbspName:Class I PI3K signaling events mediated by Akt      &nbspID:DB\_ID:1648** | | | | | | |
| --- | --- | --- | --- | --- | --- | --- |
| C=1288; O=7; E=2.03; R=3.44; rawP=0.0034; adjP=0.0043 | | | | | | |
| Index | UserID | Value | Gene Symbol | Gene Name | EntrezGene | Ensembl |
| 1 | P60842 | NA | EIF4A1 | eukaryotic translation initiation factor 4A1 | 1973 | ENSG00000161960 |
| 2 | P08195 | NA | SLC3A2 | solute carrier family 3 (activators of dibasic and neutral amino acid transport), member 2 | 6520 | ENSG00000168003 |
| 3 | P13639 | NA | EEF2 | eukaryotic translation elongation factor 2 | 1938 | ENSG00000167658 |
| 4 | P18206 | NA | VCL | vinculin | 7414 | ENSG00000035403 |
| 5 | P02786 | NA | TFRC | transferrin receptor (p90, CD71) | 7037 | ENSG00000072274 |
| 6 | P05556 | NA | ITGB1 | integrin, beta 1 (fibronectin receptor, beta polypeptide, antigen CD29 includes MDF2, MSK12) | 3688 | ENSG00000150093 |
| 7 | P09382 | NA | LGALS1 | lectin, galactoside-binding, soluble, 1 | 3956 | ENSG00000100097 |

  
  

| **Database:Pathway Commons pathway      &nbspName:Arf6 trafficking events      &nbspID:DB\_ID:1615** | | | | | | |
| --- | --- | --- | --- | --- | --- | --- |
| C=1288; O=7; E=2.03; R=3.44; rawP=0.0034; adjP=0.0043 | | | | | | |
| Index | UserID | Value | Gene Symbol | Gene Name | EntrezGene | Ensembl |
| 1 | P60842 | NA | EIF4A1 | eukaryotic translation initiation factor 4A1 | 1973 | ENSG00000161960 |
| 2 | P08195 | NA | SLC3A2 | solute carrier family 3 (activators of dibasic and neutral amino acid transport), member 2 | 6520 | ENSG00000168003 |
| 3 | P13639 | NA | EEF2 | eukaryotic translation elongation factor 2 | 1938 | ENSG00000167658 |
| 4 | P18206 | NA | VCL | vinculin | 7414 | ENSG00000035403 |
| 5 | P02786 | NA | TFRC | transferrin receptor (p90, CD71) | 7037 | ENSG00000072274 |
| 6 | P05556 | NA | ITGB1 | integrin, beta 1 (fibronectin receptor, beta polypeptide, antigen CD29 includes MDF2, MSK12) | 3688 | ENSG00000150093 |
| 7 | P09382 | NA | LGALS1 | lectin, galactoside-binding, soluble, 1 | 3956 | ENSG00000100097 |

  
  

| **Database:Pathway Commons pathway      &nbspName:Transmembrane transport of small molecules      &nbspID:DB\_ID:937** | | | | | | |
| --- | --- | --- | --- | --- | --- | --- |
| C=379; O=4; E=0.60; R=6.69; rawP=0.0029; adjP=0.0043 | | | | | | |
| Index | UserID | Value | Gene Symbol | Gene Name | EntrezGene | Ensembl |
| 1 | P08195 | NA | SLC3A2 | solute carrier family 3 (activators of dibasic and neutral amino acid transport), member 2 | 6520 | ENSG00000168003 |
| 2 | P02768 | NA | ALB | albumin | 213 | ENSG00000163631 |
| 3 | P02786 | NA | TFRC | transferrin receptor (p90, CD71) | 7037 | ENSG00000072274 |
| 4 | Q96QD8 | NA | SLC38A2 | solute carrier family 38, member 2 | 54407 | ENSG00000134294 |

  
  

| **Database:Pathway Commons pathway      &nbspName:Glypican 1 network      &nbspID:DB\_ID:1492** | | | | | | |
| --- | --- | --- | --- | --- | --- | --- |
| C=1299; O=7; E=2.05; R=3.42; rawP=0.0036; adjP=0.0043 | | | | | | |
| Index | UserID | Value | Gene Symbol | Gene Name | EntrezGene | Ensembl |
| 1 | P60842 | NA | EIF4A1 | eukaryotic translation initiation factor 4A1 | 1973 | ENSG00000161960 |
| 2 | P08195 | NA | SLC3A2 | solute carrier family 3 (activators of dibasic and neutral amino acid transport), member 2 | 6520 | ENSG00000168003 |
| 3 | P13639 | NA | EEF2 | eukaryotic translation elongation factor 2 | 1938 | ENSG00000167658 |
| 4 | P18206 | NA | VCL | vinculin | 7414 | ENSG00000035403 |
| 5 | P02786 | NA | TFRC | transferrin receptor (p90, CD71) | 7037 | ENSG00000072274 |
| 6 | P05556 | NA | ITGB1 | integrin, beta 1 (fibronectin receptor, beta polypeptide, antigen CD29 includes MDF2, MSK12) | 3688 | ENSG00000150093 |
| 7 | P09382 | NA | LGALS1 | lectin, galactoside-binding, soluble, 1 | 3956 | ENSG00000100097 |

  
  

| **Database:Pathway Commons pathway      &nbspName:GMCSF-mediated signaling events      &nbspID:DB\_ID:1461** | | | | | | |
| --- | --- | --- | --- | --- | --- | --- |
| C=1292; O=7; E=2.04; R=3.43; rawP=0.0034; adjP=0.0043 | | | | | | |
| Index | UserID | Value | Gene Symbol | Gene Name | EntrezGene | Ensembl |
| 1 | P60842 | NA | EIF4A1 | eukaryotic translation initiation factor 4A1 | 1973 | ENSG00000161960 |
| 2 | P08195 | NA | SLC3A2 | solute carrier family 3 (activators of dibasic and neutral amino acid transport), member 2 | 6520 | ENSG00000168003 |
| 3 | P13639 | NA | EEF2 | eukaryotic translation elongation factor 2 | 1938 | ENSG00000167658 |
| 4 | P18206 | NA | VCL | vinculin | 7414 | ENSG00000035403 |
| 5 | P02786 | NA | TFRC | transferrin receptor (p90, CD71) | 7037 | ENSG00000072274 |
| 6 | P05556 | NA | ITGB1 | integrin, beta 1 (fibronectin receptor, beta polypeptide, antigen CD29 includes MDF2, MSK12) | 3688 | ENSG00000150093 |
| 7 | P09382 | NA | LGALS1 | lectin, galactoside-binding, soluble, 1 | 3956 | ENSG00000100097 |

  
  

| **Database:Pathway Commons pathway      &nbspName:Proteoglycan syndecan-mediated signaling events      &nbspID:DB\_ID:1637** | | | | | | |
| --- | --- | --- | --- | --- | --- | --- |
| C=1345; O=7; E=2.12; R=3.30; rawP=0.0043; adjP=0.0047 | | | | | | |
| Index | UserID | Value | Gene Symbol | Gene Name | EntrezGene | Ensembl |
| 1 | P60842 | NA | EIF4A1 | eukaryotic translation initiation factor 4A1 | 1973 | ENSG00000161960 |
| 2 | P08195 | NA | SLC3A2 | solute carrier family 3 (activators of dibasic and neutral amino acid transport), member 2 | 6520 | ENSG00000168003 |
| 3 | P13639 | NA | EEF2 | eukaryotic translation elongation factor 2 | 1938 | ENSG00000167658 |
| 4 | P18206 | NA | VCL | vinculin | 7414 | ENSG00000035403 |
| 5 | P02786 | NA | TFRC | transferrin receptor (p90, CD71) | 7037 | ENSG00000072274 |
| 6 | P05556 | NA | ITGB1 | integrin, beta 1 (fibronectin receptor, beta polypeptide, antigen CD29 includes MDF2, MSK12) | 3688 | ENSG00000150093 |
| 7 | P09382 | NA | LGALS1 | lectin, galactoside-binding, soluble, 1 | 3956 | ENSG00000100097 |

  
  

| **Database:Pathway Commons pathway      &nbspName:Beta1 integrin cell surface interactions      &nbspID:DB\_ID:1517** | | | | | | |
| --- | --- | --- | --- | --- | --- | --- |
| C=1351; O=7; E=2.13; R=3.28; rawP=0.0044; adjP=0.0047 | | | | | | |
| Index | UserID | Value | Gene Symbol | Gene Name | EntrezGene | Ensembl |
| 1 | P60842 | NA | EIF4A1 | eukaryotic translation initiation factor 4A1 | 1973 | ENSG00000161960 |
| 2 | P08195 | NA | SLC3A2 | solute carrier family 3 (activators of dibasic and neutral amino acid transport), member 2 | 6520 | ENSG00000168003 |
| 3 | P13639 | NA | EEF2 | eukaryotic translation elongation factor 2 | 1938 | ENSG00000167658 |
| 4 | P18206 | NA | VCL | vinculin | 7414 | ENSG00000035403 |
| 5 | P02786 | NA | TFRC | transferrin receptor (p90, CD71) | 7037 | ENSG00000072274 |
| 6 | P05556 | NA | ITGB1 | integrin, beta 1 (fibronectin receptor, beta polypeptide, antigen CD29 includes MDF2, MSK12) | 3688 | ENSG00000150093 |
| 7 | P09382 | NA | LGALS1 | lectin, galactoside-binding, soluble, 1 | 3956 | ENSG00000100097 |

  
  

| **Database:Pathway Commons pathway      &nbspName:Glypican pathway      &nbspID:DB\_ID:1459** | | | | | | |
| --- | --- | --- | --- | --- | --- | --- |
| C=1338; O=7; E=2.11; R=3.32; rawP=0.0042; adjP=0.0047 | | | | | | |
| Index | UserID | Value | Gene Symbol | Gene Name | EntrezGene | Ensembl |
| 1 | P60842 | NA | EIF4A1 | eukaryotic translation initiation factor 4A1 | 1973 | ENSG00000161960 |
| 2 | P08195 | NA | SLC3A2 | solute carrier family 3 (activators of dibasic and neutral amino acid transport), member 2 | 6520 | ENSG00000168003 |
| 3 | P13639 | NA | EEF2 | eukaryotic translation elongation factor 2 | 1938 | ENSG00000167658 |
| 4 | P18206 | NA | VCL | vinculin | 7414 | ENSG00000035403 |
| 5 | P02786 | NA | TFRC | transferrin receptor (p90, CD71) | 7037 | ENSG00000072274 |
| 6 | P05556 | NA | ITGB1 | integrin, beta 1 (fibronectin receptor, beta polypeptide, antigen CD29 includes MDF2, MSK12) | 3688 | ENSG00000150093 |
| 7 | P09382 | NA | LGALS1 | lectin, galactoside-binding, soluble, 1 | 3956 | ENSG00000100097 |

  
  

| **Database:Pathway Commons pathway      &nbspName:Integrin family cell surface interactions      &nbspID:DB\_ID:1499** | | | | | | |
| --- | --- | --- | --- | --- | --- | --- |
| C=1378; O=7; E=2.17; R=3.22; rawP=0.0049; adjP=0.0052 | | | | | | |
| Index | UserID | Value | Gene Symbol | Gene Name | EntrezGene | Ensembl |
| 1 | P60842 | NA | EIF4A1 | eukaryotic translation initiation factor 4A1 | 1973 | ENSG00000161960 |
| 2 | P08195 | NA | SLC3A2 | solute carrier family 3 (activators of dibasic and neutral amino acid transport), member 2 | 6520 | ENSG00000168003 |
| 3 | P13639 | NA | EEF2 | eukaryotic translation elongation factor 2 | 1938 | ENSG00000167658 |
| 4 | P18206 | NA | VCL | vinculin | 7414 | ENSG00000035403 |
| 5 | P02786 | NA | TFRC | transferrin receptor (p90, CD71) | 7037 | ENSG00000072274 |
| 6 | P05556 | NA | ITGB1 | integrin, beta 1 (fibronectin receptor, beta polypeptide, antigen CD29 includes MDF2, MSK12) | 3688 | ENSG00000150093 |
| 7 | P09382 | NA | LGALS1 | lectin, galactoside-binding, soluble, 1 | 3956 | ENSG00000100097 |

  
  

| **Database:Pathway Commons pathway      &nbspName:SLC-mediated transmembrane transport      &nbspID:DB\_ID:943** | | | | | | |
| --- | --- | --- | --- | --- | --- | --- |
| C=248; O=3; E=0.39; R=7.67; rawP=0.0069; adjP=0.0072 | | | | | | |
| Index | UserID | Value | Gene Symbol | Gene Name | EntrezGene | Ensembl |
| 1 | P08195 | NA | SLC3A2 | solute carrier family 3 (activators of dibasic and neutral amino acid transport), member 2 | 6520 | ENSG00000168003 |
| 2 | P02768 | NA | ALB | albumin | 213 | ENSG00000163631 |
| 3 | Q96QD8 | NA | SLC38A2 | solute carrier family 38, member 2 | 54407 | ENSG00000134294 |

  
  

| **Database:Pathway Commons pathway      &nbspName:CDC42 signaling events      &nbspID:DB\_ID:1488** | | | | | | |
| --- | --- | --- | --- | --- | --- | --- |
| C=757; O=4; E=1.19; R=3.35; rawP=0.0302; adjP=0.0307 | | | | | | |
| Index | UserID | Value | Gene Symbol | Gene Name | EntrezGene | Ensembl |
| 1 | P08195 | NA | SLC3A2 | solute carrier family 3 (activators of dibasic and neutral amino acid transport), member 2 | 6520 | ENSG00000168003 |
| 2 | P18206 | NA | VCL | vinculin | 7414 | ENSG00000035403 |
| 3 | P02786 | NA | TFRC | transferrin receptor (p90, CD71) | 7037 | ENSG00000072274 |
| 4 | P05556 | NA | ITGB1 | integrin, beta 1 (fibronectin receptor, beta polypeptide, antigen CD29 includes MDF2, MSK12) | 3688 | ENSG00000150093 |

  
  

| **Database:Pathway Commons pathway      &nbspName:Regulation of CDC42 activity      &nbspID:DB\_ID:1456** | | | | | | |
| --- | --- | --- | --- | --- | --- | --- |
| C=770; O=4; E=1.21; R=3.29; rawP=0.0319; adjP=0.0319 | | | | | | |
| Index | UserID | Value | Gene Symbol | Gene Name | EntrezGene | Ensembl |
| 1 | P08195 | NA | SLC3A2 | solute carrier family 3 (activators of dibasic and neutral amino acid transport), member 2 | 6520 | ENSG00000168003 |
| 2 | P18206 | NA | VCL | vinculin | 7414 | ENSG00000035403 |
| 3 | P02786 | NA | TFRC | transferrin receptor (p90, CD71) | 7037 | ENSG00000072274 |
| 4 | P05556 | NA | ITGB1 | integrin, beta 1 (fibronectin receptor, beta polypeptide, antigen CD29 includes MDF2, MSK12) | 3688 | ENSG00000150093 |

  
  
  
  

---

WebGestalt is currently developed and maintained by Jing Wang and Bing Zhang at the  Zhang Lab. Other people who have made significant contribution to the project include Dexter Duncan, Stefan Kirov, Zhiao Shi, and Jay Snoddy.  
  
**Funding credits:** NIH/NIAAA (U01 AA016662, U01 AA013512); NIH/NIDA (P01 DA015027); NIH/NIMH (P50 MH078028, P50 MH096972); NIH/NCI (U24 CA159988); NIH/NIGMS (R01 GM088822).
