## Supplementary material for "PGRMC1 phosphorylation and cell plasticity 1: glycolysis, mitochondria, tumor growth": File S8: final_tf_geneset_file_1439185880.html

Anchored HTML File of EIDs

|  |  |
| --- | --- |
|  | WEB-based GEne SeT AnaLysis Toolkit |
|  | ***Translating gene lists into biological insights...*** |

---

  

| **Database:Transcrription Target      &nbspName:hsa\_V$HLF\_01      &nbspID:DB\_ID:2030** | | | | | | |
| --- | --- | --- | --- | --- | --- | --- |
| C=250; O=6; E=0.39; R=15.21; rawP=2.29e-06; adjP=6.87e-05 | | | | | | |
| Index | UserID | Value | Gene Symbol | Gene Name | EntrezGene | Ensembl |
| 1 | P60842 | NA | EIF4A1 | eukaryotic translation initiation factor 4A1 | 1973 | ENSG00000161960 |
| 2 | P13639 | NA | EEF2 | eukaryotic translation elongation factor 2 | 1938 | ENSG00000167658 |
| 3 | P00352 | NA | ALDH1A1 | aldehyde dehydrogenase 1 family, member A1 | 216 | ENSG00000165092 |
| 4 | P02768 | NA | ALB | albumin | 213 | ENSG00000163631 |
| 5 | P27816 | NA | MAP4 | microtubule-associated protein 4 | 4134 | ENSG00000047849 |
| 6 | P11387 | NA | TOP1 | topoisomerase (DNA) I | 7150 | ENSG00000198900 |

  
  

| **Database:Transcrription Target      &nbspName:hsa\_GGCNNMSMYNTTG\_UNKNOWN      &nbspID:DB\_ID:2313** | | | | | | |
| --- | --- | --- | --- | --- | --- | --- |
| C=74; O=3; E=0.12; R=25.70; rawP=0.0002; adjP=0.0030 | | | | | | |
| Index | UserID | Value | Gene Symbol | Gene Name | EntrezGene | Ensembl |
| 1 | P67809 | NA | YBX1 | Y box binding protein 1 | 4904 | ENSG00000065978 |
| 2 | P60842 | NA | EIF4A1 | eukaryotic translation initiation factor 4A1 | 1973 | ENSG00000161960 |
| 3 | P68104 | NA | EEF1A1 | eukaryotic translation elongation factor 1 alpha 1 | 1915 | ENSG00000156508 |

  
  

| **Database:Transcrription Target      &nbspName:hsa\_V$AP1\_01      &nbspID:DB\_ID:2160** | | | | | | |
| --- | --- | --- | --- | --- | --- | --- |
| C=256; O=4; E=0.40; R=9.90; rawP=0.0007; adjP=0.0042 | | | | | | |
| Index | UserID | Value | Gene Symbol | Gene Name | EntrezGene | Ensembl |
| 1 | P18206 | NA | VCL | vinculin | 7414 | ENSG00000035403 |
| 2 | P68104 | NA | EEF1A1 | eukaryotic translation elongation factor 1 alpha 1 | 1915 | ENSG00000156508 |
| 3 | P27816 | NA | MAP4 | microtubule-associated protein 4 | 4134 | ENSG00000047849 |
| 4 | Q96QD8 | NA | SLC38A2 | solute carrier family 38, member 2 | 54407 | ENSG00000134294 |

  
  

| **Database:Transcrription Target      &nbspName:hsa\_V$MAX\_01      &nbspID:DB\_ID:1924** | | | | | | |
| --- | --- | --- | --- | --- | --- | --- |
| C=256; O=4; E=0.40; R=9.90; rawP=0.0007; adjP=0.0042 | | | | | | |
| Index | UserID | Value | Gene Symbol | Gene Name | EntrezGene | Ensembl |
| 1 | P02786 | NA | TFRC | transferrin receptor (p90, CD71) | 7037 | ENSG00000072274 |
| 2 | P11387 | NA | TOP1 | topoisomerase (DNA) I | 7150 | ENSG00000198900 |
| 3 | Q96QD8 | NA | SLC38A2 | solute carrier family 38, member 2 | 54407 | ENSG00000134294 |
| 4 | P24534 | NA | EEF1B2 | eukaryotic translation elongation factor 1 beta 2 | 1933 | ENSG00000114942 |

  
  

| **Database:Transcrription Target      &nbspName:hsa\_V$CHOP\_01      &nbspID:DB\_ID:2022** | | | | | | |
| --- | --- | --- | --- | --- | --- | --- |
| C=232; O=4; E=0.37; R=10.93; rawP=0.0005; adjP=0.0042 | | | | | | |
| Index | UserID | Value | Gene Symbol | Gene Name | EntrezGene | Ensembl |
| 1 | P49589 | NA | CARS | cysteinyl-tRNA synthetase | 833 | ENSG00000110619 |
| 2 | P60842 | NA | EIF4A1 | eukaryotic translation initiation factor 4A1 | 1973 | ENSG00000161960 |
| 3 | P27816 | NA | MAP4 | microtubule-associated protein 4 | 4134 | ENSG00000047849 |
| 4 | Q96QD8 | NA | SLC38A2 | solute carrier family 38, member 2 | 54407 | ENSG00000134294 |

  
  

| **Database:Transcrription Target      &nbspName:hsa\_V$CEBPB\_02      &nbspID:DB\_ID:1922** | | | | | | |
| --- | --- | --- | --- | --- | --- | --- |
| C=250; O=3; E=0.39; R=7.61; rawP=0.0070; adjP=0.0171 | | | | | | |
| Index | UserID | Value | Gene Symbol | Gene Name | EntrezGene | Ensembl |
| 1 | P17174 | NA | GOT1 | glutamic-oxaloacetic transaminase 1, soluble (aspartate aminotransferase 1) | 2805 | ENSG00000120053 |
| 2 | P60842 | NA | EIF4A1 | eukaryotic translation initiation factor 4A1 | 1973 | ENSG00000161960 |
| 3 | P11387 | NA | TOP1 | topoisomerase (DNA) I | 7150 | ENSG00000198900 |

  
  

| **Database:Transcrription Target      &nbspName:hsa\_V$E4BP4\_01      &nbspID:DB\_ID:1869** | | | | | | |
| --- | --- | --- | --- | --- | --- | --- |
| C=217; O=3; E=0.34; R=8.76; rawP=0.0048; adjP=0.0171 | | | | | | |
| Index | UserID | Value | Gene Symbol | Gene Name | EntrezGene | Ensembl |
| 1 | P13639 | NA | EEF2 | eukaryotic translation elongation factor 2 | 1938 | ENSG00000167658 |
| 2 | P00352 | NA | ALDH1A1 | aldehyde dehydrogenase 1 family, member A1 | 216 | ENSG00000165092 |
| 3 | P27816 | NA | MAP4 | microtubule-associated protein 4 | 4134 | ENSG00000047849 |

  
  

| **Database:Transcrription Target      &nbspName:hsa\_V$MYCMAX\_01      &nbspID:DB\_ID:1923** | | | | | | |
| --- | --- | --- | --- | --- | --- | --- |
| C=250; O=3; E=0.39; R=7.61; rawP=0.0070; adjP=0.0171 | | | | | | |
| Index | UserID | Value | Gene Symbol | Gene Name | EntrezGene | Ensembl |
| 1 | P02786 | NA | TFRC | transferrin receptor (p90, CD71) | 7037 | ENSG00000072274 |
| 2 | P11387 | NA | TOP1 | topoisomerase (DNA) I | 7150 | ENSG00000198900 |
| 3 | P24534 | NA | EEF1B2 | eukaryotic translation elongation factor 1 beta 2 | 1933 | ENSG00000114942 |

  
  

| **Database:Transcrription Target      &nbspName:hsa\_V$USF\_01      &nbspID:DB\_ID:1925** | | | | | | |
| --- | --- | --- | --- | --- | --- | --- |
| C=255; O=3; E=0.40; R=7.46; rawP=0.0074; adjP=0.0171 | | | | | | |
| Index | UserID | Value | Gene Symbol | Gene Name | EntrezGene | Ensembl |
| 1 | P02786 | NA | TFRC | transferrin receptor (p90, CD71) | 7037 | ENSG00000072274 |
| 2 | Q96QD8 | NA | SLC38A2 | solute carrier family 38, member 2 | 54407 | ENSG00000134294 |
| 3 | P24534 | NA | EEF1B2 | eukaryotic translation elongation factor 1 beta 2 | 1933 | ENSG00000114942 |

  
  

| **Database:Transcrription Target      &nbspName:hsa\_CACGTG\_V$MYC\_Q2      &nbspID:DB\_ID:2434** | | | | | | |
| --- | --- | --- | --- | --- | --- | --- |
| C=1008; O=6; E=1.59; R=3.77; rawP=0.0044; adjP=0.0171 | | | | | | |
| Index | UserID | Value | Gene Symbol | Gene Name | EntrezGene | Ensembl |
| 1 | P67809 | NA | YBX1 | Y box binding protein 1 | 4904 | ENSG00000065978 |
| 2 | P60842 | NA | EIF4A1 | eukaryotic translation initiation factor 4A1 | 1973 | ENSG00000161960 |
| 3 | P02786 | NA | TFRC | transferrin receptor (p90, CD71) | 7037 | ENSG00000072274 |
| 4 | P11387 | NA | TOP1 | topoisomerase (DNA) I | 7150 | ENSG00000198900 |
| 5 | Q96QD8 | NA | SLC38A2 | solute carrier family 38, member 2 | 54407 | ENSG00000134294 |
| 6 | P24534 | NA | EEF1B2 | eukaryotic translation elongation factor 1 beta 2 | 1933 | ENSG00000114942 |

  
  

| **Database:Transcrription Target      &nbspName:hsa\_V$ATF1\_Q6      &nbspID:DB\_ID:2205** | | | | | | |
| --- | --- | --- | --- | --- | --- | --- |
| C=228; O=3; E=0.36; R=8.34; rawP=0.0055; adjP=0.0171 | | | | | | |
| Index | UserID | Value | Gene Symbol | Gene Name | EntrezGene | Ensembl |
| 1 | P60842 | NA | EIF4A1 | eukaryotic translation initiation factor 4A1 | 1973 | ENSG00000161960 |
| 2 | P13639 | NA | EEF2 | eukaryotic translation elongation factor 2 | 1938 | ENSG00000167658 |
| 3 | P11387 | NA | TOP1 | topoisomerase (DNA) I | 7150 | ENSG00000198900 |

  
  

| **Database:Transcrription Target      &nbspName:hsa\_V$YY1\_Q6      &nbspID:DB\_ID:2268** | | | | | | |
| --- | --- | --- | --- | --- | --- | --- |
| C=236; O=3; E=0.37; R=8.06; rawP=0.0060; adjP=0.0171 | | | | | | |
| Index | UserID | Value | Gene Symbol | Gene Name | EntrezGene | Ensembl |
| 1 | P67809 | NA | YBX1 | Y box binding protein 1 | 4904 | ENSG00000065978 |
| 2 | P60842 | NA | EIF4A1 | eukaryotic translation initiation factor 4A1 | 1973 | ENSG00000161960 |
| 3 | P68104 | NA | EEF1A1 | eukaryotic translation elongation factor 1 alpha 1 | 1915 | ENSG00000156508 |

  
  

| **Database:Transcrription Target      &nbspName:hsa\_V$CEBP\_Q2      &nbspID:DB\_ID:1979** | | | | | | |
| --- | --- | --- | --- | --- | --- | --- |
| C=227; O=3; E=0.36; R=8.38; rawP=0.0054; adjP=0.0171 | | | | | | |
| Index | UserID | Value | Gene Symbol | Gene Name | EntrezGene | Ensembl |
| 1 | P60842 | NA | EIF4A1 | eukaryotic translation initiation factor 4A1 | 1973 | ENSG00000161960 |
| 2 | P27816 | NA | MAP4 | microtubule-associated protein 4 | 4134 | ENSG00000047849 |
| 3 | Q96QD8 | NA | SLC38A2 | solute carrier family 38, member 2 | 54407 | ENSG00000134294 |

  
  

| **Database:Transcrription Target      &nbspName:hsa\_TTGTTT\_V$FOXO4\_01      &nbspID:DB\_ID:2416** | | | | | | |
| --- | --- | --- | --- | --- | --- | --- |
| C=2012; O=8; E=3.17; R=2.52; rawP=0.0111; adjP=0.0238 | | | | | | |
| Index | UserID | Value | Gene Symbol | Gene Name | EntrezGene | Ensembl |
| 1 | P02768 | NA | ALB | albumin | 213 | ENSG00000163631 |
| 2 | Q96QD8 | NA | SLC38A2 | solute carrier family 38, member 2 | 54407 | ENSG00000134294 |
| 3 | P67809 | NA | YBX1 | Y box binding protein 1 | 4904 | ENSG00000065978 |
| 4 | P53701 | NA | HCCS | holocytochrome c synthase | 3052 | ENSG00000004961 |
| 5 | P00352 | NA | ALDH1A1 | aldehyde dehydrogenase 1 family, member A1 | 216 | ENSG00000165092 |
| 6 | P18206 | NA | VCL | vinculin | 7414 | ENSG00000035403 |
| 7 | P04083 | NA | ANXA1 | annexin A1 | 301 | ENSG00000135046 |
| 8 | P11387 | NA | TOP1 | topoisomerase (DNA) I | 7150 | ENSG00000198900 |

  
  

| **Database:Transcrription Target      &nbspName:hsa\_GGGYGTGNY\_UNKNOWN      &nbspID:DB\_ID:2333** | | | | | | |
| --- | --- | --- | --- | --- | --- | --- |
| C=648; O=4; E=1.02; R=3.91; rawP=0.0183; adjP=0.0366 | | | | | | |
| Index | UserID | Value | Gene Symbol | Gene Name | EntrezGene | Ensembl |
| 1 | P07108 | NA | DBI | diazepam binding inhibitor (GABA receptor modulator, acyl-CoA binding protein) | 1622 | ENSG00000155368 |
| 2 | O75369 | NA | FLNB | filamin B, beta | 2317 | ENSG00000136068 |
| 3 | Q96QD8 | NA | SLC38A2 | solute carrier family 38, member 2 | 54407 | ENSG00000134294 |
| 4 | P09382 | NA | LGALS1 | lectin, galactoside-binding, soluble, 1 | 3956 | ENSG00000100097 |

  
  

| **Database:Transcrription Target      &nbspName:hsa\_GGGAGGRR\_V$MAZ\_Q6      &nbspID:DB\_ID:2430** | | | | | | |
| --- | --- | --- | --- | --- | --- | --- |
| C=2223; O=8; E=3.51; R=2.28; rawP=0.0195; adjP=0.0366 | | | | | | |
| Index | UserID | Value | Gene Symbol | Gene Name | EntrezGene | Ensembl |
| 1 | P61313 | NA | RPL15 | ribosomal protein L15 | 6138 | ENSG00000174748 |
| 2 | P17174 | NA | GOT1 | glutamic-oxaloacetic transaminase 1, soluble (aspartate aminotransferase 1) | 2805 | ENSG00000120053 |
| 3 | P60842 | NA | EIF4A1 | eukaryotic translation initiation factor 4A1 | 1973 | ENSG00000161960 |
| 4 | P22102 | NA | GART | phosphoribosylglycinamide formyltransferase, phosphoribosylglycinamide synthetase, phosphoribosylaminoimidazole synthetase | 2618 | ENSG00000159131 |
| 5 | P18206 | NA | VCL | vinculin | 7414 | ENSG00000035403 |
| 6 | P68104 | NA | EEF1A1 | eukaryotic translation elongation factor 1 alpha 1 | 1915 | ENSG00000156508 |
| 7 | P27816 | NA | MAP4 | microtubule-associated protein 4 | 4134 | ENSG00000047849 |
| 8 | P09382 | NA | LGALS1 | lectin, galactoside-binding, soluble, 1 | 3956 | ENSG00000100097 |

  
  

| **Database:Transcrription Target      &nbspName:hsa\_TGANTCA\_V$AP1\_C      &nbspID:DB\_ID:2402** | | | | | | |
| --- | --- | --- | --- | --- | --- | --- |
| C=1092; O=5; E=1.72; R=2.90; rawP=0.0269; adjP=0.0457 | | | | | | |
| Index | UserID | Value | Gene Symbol | Gene Name | EntrezGene | Ensembl |
| 1 | P18206 | NA | VCL | vinculin | 7414 | ENSG00000035403 |
| 2 | P68104 | NA | EEF1A1 | eukaryotic translation elongation factor 1 alpha 1 | 1915 | ENSG00000156508 |
| 3 | P27816 | NA | MAP4 | microtubule-associated protein 4 | 4134 | ENSG00000047849 |
| 4 | Q96QD8 | NA | SLC38A2 | solute carrier family 38, member 2 | 54407 | ENSG00000134294 |
| 5 | P09382 | NA | LGALS1 | lectin, galactoside-binding, soluble, 1 | 3956 | ENSG00000100097 |

  
  

| **Database:Transcrription Target      &nbspName:hsa\_GCCATNTTG\_V$YY1\_Q6      &nbspID:DB\_ID:2459** | | | | | | |
| --- | --- | --- | --- | --- | --- | --- |
| C=416; O=3; E=0.66; R=4.57; rawP=0.0274; adjP=0.0457 | | | | | | |
| Index | UserID | Value | Gene Symbol | Gene Name | EntrezGene | Ensembl |
| 1 | P67809 | NA | YBX1 | Y box binding protein 1 | 4904 | ENSG00000065978 |
| 2 | P60842 | NA | EIF4A1 | eukaryotic translation initiation factor 4A1 | 1973 | ENSG00000161960 |
| 3 | P24534 | NA | EEF1B2 | eukaryotic translation elongation factor 1 beta 2 | 1933 | ENSG00000114942 |

  
  

| **Database:Transcrription Target      &nbspName:hsa\_GGGCGGR\_V$SP1\_Q6      &nbspID:DB\_ID:2452** | | | | | | |
| --- | --- | --- | --- | --- | --- | --- |
| C=2863; O=9; E=4.52; R=1.99; rawP=0.0291; adjP=0.0459 | | | | | | |
| Index | UserID | Value | Gene Symbol | Gene Name | EntrezGene | Ensembl |
| 1 | P41091 | NA | EIF2S3 | eukaryotic translation initiation factor 2, subunit 3 gamma, 52kDa | 1968 | ENSG00000130741 |
| 2 | P27816 | NA | MAP4 | microtubule-associated protein 4 | 4134 | ENSG00000047849 |
| 3 | P05556 | NA | ITGB1 | integrin, beta 1 (fibronectin receptor, beta polypeptide, antigen CD29 includes MDF2, MSK12) | 3688 | ENSG00000150093 |
| 4 | P09382 | NA | LGALS1 | lectin, galactoside-binding, soluble, 1 | 3956 | ENSG00000100097 |
| 5 | P60842 | NA | EIF4A1 | eukaryotic translation initiation factor 4A1 | 1973 | ENSG00000161960 |
| 6 | P67809 | NA | YBX1 | Y box binding protein 1 | 4904 | ENSG00000065978 |
| 7 | P18206 | NA | VCL | vinculin | 7414 | ENSG00000035403 |
| 8 | P68104 | NA | EEF1A1 | eukaryotic translation elongation factor 1 alpha 1 | 1915 | ENSG00000156508 |
| 9 | P11387 | NA | TOP1 | topoisomerase (DNA) I | 7150 | ENSG00000198900 |

  
  

| **Database:Transcrription Target      &nbspName:hsa\_SCGGAAGY\_V$ELK1\_02      &nbspID:DB\_ID:2413** | | | | | | |
| --- | --- | --- | --- | --- | --- | --- |
| C=1170; O=5; E=1.85; R=2.71; rawP=0.0348; adjP=0.0522 | | | | | | |
| Index | UserID | Value | Gene Symbol | Gene Name | EntrezGene | Ensembl |
| 1 | P49589 | NA | CARS | cysteinyl-tRNA synthetase | 833 | ENSG00000110619 |
| 2 | P60842 | NA | EIF4A1 | eukaryotic translation initiation factor 4A1 | 1973 | ENSG00000161960 |
| 3 | P22102 | NA | GART | phosphoribosylglycinamide formyltransferase, phosphoribosylglycinamide synthetase, phosphoribosylaminoimidazole synthetase | 2618 | ENSG00000159131 |
| 4 | P53701 | NA | HCCS | holocytochrome c synthase | 3052 | ENSG00000004961 |
| 5 | P24534 | NA | EEF1B2 | eukaryotic translation elongation factor 1 beta 2 | 1933 | ENSG00000114942 |

  
  
  
  

---

WebGestalt is currently developed and maintained by Jing Wang and Bing Zhang at the  Zhang Lab. Other people who have made significant contribution to the project include Dexter Duncan, Stefan Kirov, Zhiao Shi, and Jay Snoddy.  
  
**Funding credits:** NIH/NIAAA (U01 AA016662, U01 AA013512); NIH/NIDA (P01 DA015027); NIH/NIMH (P50 MH078028, P50 MH096972); NIH/NCI (U24 CA159988); NIH/NIGMS (R01 GM088822).
