## Supplementary material for "PGRMC1 phosphorylation and cell plasticity 1: glycolysis, mitochondria, tumor growth": File S8: final_sig_file_1439166314.html

Anchored HTML File of EIDs

|  |  |
| --- | --- |
|  | WEB-based GEne SeT AnaLysis Toolkit |
|  | ***Translating gene lists into biological insights...*** |

---

  

| **Database:biological process      &nbspName:protein transport      &nbspID:GO:0015031** | | | | | | |
| --- | --- | --- | --- | --- | --- | --- |
| C=1231; O=6; E=1.54; R=3.91; rawP=0.0029; adjP=0.0495 | | | | | | |
| Index | UserID | Value | Gene Symbol | Gene Name | EntrezGene | Ensembl |
| 1 | O14980 | NA | XPO1 | exportin 1 (CRM1 homolog, yeast) | 7514 | ENSG00000082898 |
| 2 | P49321 | NA | NASP | nuclear autoantigenic sperm protein (histone-binding) | 4678 | ENSG00000132780 |
| 3 | P00491 | NA | PNP | purine nucleoside phosphorylase | 4860 | ENSG00000198805 |
| 4 | P05783 | NA | KRT18 | keratin 18 | 3875 | ENSG00000111057 |
| 5 | Q9Y2X3 | NA | NOP58 | NOP58 ribonucleoprotein homolog (yeast) | 51602 | ENSG00000055044 |
| 6 | Q06830 | NA | PRDX1 | peroxiredoxin 1 | 5052 | ENSG00000117450 |

  
  

| **Database:biological process      &nbspName:establishment of protein localization      &nbspID:GO:0045184** | | | | | | |
| --- | --- | --- | --- | --- | --- | --- |
| C=1269; O=6; E=1.58; R=3.79; rawP=0.0034; adjP=0.0495 | | | | | | |
| Index | UserID | Value | Gene Symbol | Gene Name | EntrezGene | Ensembl |
| 1 | O14980 | NA | XPO1 | exportin 1 (CRM1 homolog, yeast) | 7514 | ENSG00000082898 |
| 2 | P49321 | NA | NASP | nuclear autoantigenic sperm protein (histone-binding) | 4678 | ENSG00000132780 |
| 3 | P00491 | NA | PNP | purine nucleoside phosphorylase | 4860 | ENSG00000198805 |
| 4 | P05783 | NA | KRT18 | keratin 18 | 3875 | ENSG00000111057 |
| 5 | Q9Y2X3 | NA | NOP58 | NOP58 ribonucleoprotein homolog (yeast) | 51602 | ENSG00000055044 |
| 6 | Q06830 | NA | PRDX1 | peroxiredoxin 1 | 5052 | ENSG00000117450 |

  
  

| **Database:biological process      &nbspName:anatomical structure development      &nbspID:GO:0048856** | | | | | | |
| --- | --- | --- | --- | --- | --- | --- |
| C=4024; O=11; E=5.02; R=2.19; rawP=0.0033; adjP=0.0495 | | | | | | |
| Index | UserID | Value | Gene Symbol | Gene Name | EntrezGene | Ensembl |
| 1 | Q7Z2K6 | NA | ERMP1 | endoplasmic reticulum metallopeptidase 1 | 79956 | ENSG00000099219 |
| 2 | P49321 | NA | NASP | nuclear autoantigenic sperm protein (histone-binding) | 4678 | ENSG00000132780 |
| 3 | P00491 | NA | PNP | purine nucleoside phosphorylase | 4860 | ENSG00000198805 |
| 4 | P26447 | NA | S100A4 | S100 calcium binding protein A4 | 6275 | ENSG00000196154 |
| 5 | Q9UL46 | NA | PSME2 | proteasome (prosome, macropain) activator subunit 2 (PA28 beta) | 5721 | ENSG00000100911 |
| 6 | P08727 | NA | KRT19 | keratin 19 | 3880 | ENSG00000171345 |
| 7 | P21980 | NA | TGM2 | transglutaminase 2 (C polypeptide, protein-glutamine-gamma-glutamyltransferase) | 7052 | ENSG00000198959 |
| 8 | P05783 | NA | KRT18 | keratin 18 | 3875 | ENSG00000111057 |
| 9 | Q13263 | NA | TRIM28 | tripartite motif containing 28 | 10155 | ENSG00000130726 |
| 10 | Q06830 | NA | PRDX1 | peroxiredoxin 1 | 5052 | ENSG00000117450 |
| 11 | P43490 | NA | NAMPT | nicotinamide phosphoribosyltransferase | 10135 | ENSG00000105835 |

  
  

| **Database:biological process      &nbspName:viral reproduction      &nbspID:GO:0016032** | | | | | | |
| --- | --- | --- | --- | --- | --- | --- |
| C=688; O=5; E=0.86; R=5.82; rawP=0.0012; adjP=0.0495 | | | | | | |
| Index | UserID | Value | Gene Symbol | Gene Name | EntrezGene | Ensembl |
| 1 | P08727 | NA | KRT19 | keratin 19 | 3880 | ENSG00000171345 |
| 2 | O14980 | NA | XPO1 | exportin 1 (CRM1 homolog, yeast) | 7514 | ENSG00000082898 |
| 3 | P05783 | NA | KRT18 | keratin 18 | 3875 | ENSG00000111057 |
| 4 | P05787 | NA | KRT8 | keratin 8 | 3856 | ENSG00000170421 |
| 5 | Q9UL46 | NA | PSME2 | proteasome (prosome, macropain) activator subunit 2 (PA28 beta) | 5721 | ENSG00000100911 |

  
  

| **Database:biological process      &nbspName:virus-host interaction      &nbspID:GO:0019048** | | | | | | |
| --- | --- | --- | --- | --- | --- | --- |
| C=369; O=4; E=0.46; R=8.69; rawP=0.0010; adjP=0.0495 | | | | | | |
| Index | UserID | Value | Gene Symbol | Gene Name | EntrezGene | Ensembl |
| 1 | P08727 | NA | KRT19 | keratin 19 | 3880 | ENSG00000171345 |
| 2 | O14980 | NA | XPO1 | exportin 1 (CRM1 homolog, yeast) | 7514 | ENSG00000082898 |
| 3 | P05783 | NA | KRT18 | keratin 18 | 3875 | ENSG00000111057 |
| 4 | P05787 | NA | KRT8 | keratin 8 | 3856 | ENSG00000170421 |

  
  

| **Database:biological process      &nbspName:regulation of I-kappaB kinase/NF-kappaB cascade      &nbspID:GO:0043122** | | | | | | |
| --- | --- | --- | --- | --- | --- | --- |
| C=196; O=3; E=0.24; R=12.26; rawP=0.0017; adjP=0.0495 | | | | | | |
| Index | UserID | Value | Gene Symbol | Gene Name | EntrezGene | Ensembl |
| 1 | P21980 | NA | TGM2 | transglutaminase 2 (C polypeptide, protein-glutamine-gamma-glutamyltransferase) | 7052 | ENSG00000198959 |
| 2 | P26447 | NA | S100A4 | S100 calcium binding protein A4 | 6275 | ENSG00000196154 |
| 3 | Q06830 | NA | PRDX1 | peroxiredoxin 1 | 5052 | ENSG00000117450 |

  
  

| **Database:biological process      &nbspName:interaction with host      &nbspID:GO:0051701** | | | | | | |
| --- | --- | --- | --- | --- | --- | --- |
| C=411; O=4; E=0.51; R=7.80; rawP=0.0014; adjP=0.0495 | | | | | | |
| Index | UserID | Value | Gene Symbol | Gene Name | EntrezGene | Ensembl |
| 1 | P08727 | NA | KRT19 | keratin 19 | 3880 | ENSG00000171345 |
| 2 | O14980 | NA | XPO1 | exportin 1 (CRM1 homolog, yeast) | 7514 | ENSG00000082898 |
| 3 | P05783 | NA | KRT18 | keratin 18 | 3875 | ENSG00000111057 |
| 4 | P05787 | NA | KRT8 | keratin 8 | 3856 | ENSG00000170421 |

  
  

| **Database:biological process      &nbspName:interspecies interaction between organisms      &nbspID:GO:0044419** | | | | | | |
| --- | --- | --- | --- | --- | --- | --- |
| C=447; O=4; E=0.56; R=7.17; rawP=0.0020; adjP=0.0495 | | | | | | |
| Index | UserID | Value | Gene Symbol | Gene Name | EntrezGene | Ensembl |
| 1 | P08727 | NA | KRT19 | keratin 19 | 3880 | ENSG00000171345 |
| 2 | O14980 | NA | XPO1 | exportin 1 (CRM1 homolog, yeast) | 7514 | ENSG00000082898 |
| 3 | P05783 | NA | KRT18 | keratin 18 | 3875 | ENSG00000111057 |
| 4 | P05787 | NA | KRT8 | keratin 8 | 3856 | ENSG00000170421 |

  
  

| **Database:biological process      &nbspName:symbiosis, encompassing mutualism through parasitism      &nbspID:GO:0044403** | | | | | | |
| --- | --- | --- | --- | --- | --- | --- |
| C=447; O=4; E=0.56; R=7.17; rawP=0.0020; adjP=0.0495 | | | | | | |
| Index | UserID | Value | Gene Symbol | Gene Name | EntrezGene | Ensembl |
| 1 | P08727 | NA | KRT19 | keratin 19 | 3880 | ENSG00000171345 |
| 2 | O14980 | NA | XPO1 | exportin 1 (CRM1 homolog, yeast) | 7514 | ENSG00000082898 |
| 3 | P05783 | NA | KRT18 | keratin 18 | 3875 | ENSG00000111057 |
| 4 | P05787 | NA | KRT8 | keratin 8 | 3856 | ENSG00000170421 |

  
  

| **Database:biological process      &nbspName:protein localization to nucleus      &nbspID:GO:0034504** | | | | | | |
| --- | --- | --- | --- | --- | --- | --- |
| C=239; O=3; E=0.30; R=10.06; rawP=0.0031; adjP=0.0495 | | | | | | |
| Index | UserID | Value | Gene Symbol | Gene Name | EntrezGene | Ensembl |
| 1 | O14980 | NA | XPO1 | exportin 1 (CRM1 homolog, yeast) | 7514 | ENSG00000082898 |
| 2 | Q9Y2X3 | NA | NOP58 | NOP58 ribonucleoprotein homolog (yeast) | 51602 | ENSG00000055044 |
| 3 | Q06830 | NA | PRDX1 | peroxiredoxin 1 | 5052 | ENSG00000117450 |

  
  

| **Database:biological process      &nbspName:I-kappaB kinase/NF-kappaB cascade      &nbspID:GO:0007249** | | | | | | |
| --- | --- | --- | --- | --- | --- | --- |
| C=220; O=3; E=0.27; R=10.93; rawP=0.0024; adjP=0.0495 | | | | | | |
| Index | UserID | Value | Gene Symbol | Gene Name | EntrezGene | Ensembl |
| 1 | P21980 | NA | TGM2 | transglutaminase 2 (C polypeptide, protein-glutamine-gamma-glutamyltransferase) | 7052 | ENSG00000198959 |
| 2 | P26447 | NA | S100A4 | S100 calcium binding protein A4 | 6275 | ENSG00000196154 |
| 3 | Q06830 | NA | PRDX1 | peroxiredoxin 1 | 5052 | ENSG00000117450 |

  
  

| **Database:biological process      &nbspName:viral reproductive process      &nbspID:GO:0022415** | | | | | | |
| --- | --- | --- | --- | --- | --- | --- |
| C=540; O=4; E=0.67; R=5.93; rawP=0.0039; adjP=0.0520 | | | | | | |
| Index | UserID | Value | Gene Symbol | Gene Name | EntrezGene | Ensembl |
| 1 | P08727 | NA | KRT19 | keratin 19 | 3880 | ENSG00000171345 |
| 2 | O14980 | NA | XPO1 | exportin 1 (CRM1 homolog, yeast) | 7514 | ENSG00000082898 |
| 3 | P05783 | NA | KRT18 | keratin 18 | 3875 | ENSG00000111057 |
| 4 | P05787 | NA | KRT8 | keratin 8 | 3856 | ENSG00000170421 |

  
  

| **Database:biological process      &nbspName:in utero embryonic development      &nbspID:GO:0001701** | | | | | | |
| --- | --- | --- | --- | --- | --- | --- |
| C=306; O=3; E=0.38; R=7.86; rawP=0.0061; adjP=0.0751 | | | | | | |
| Index | UserID | Value | Gene Symbol | Gene Name | EntrezGene | Ensembl |
| 1 | P08727 | NA | KRT19 | keratin 19 | 3880 | ENSG00000171345 |
| 2 | P49321 | NA | NASP | nuclear autoantigenic sperm protein (histone-binding) | 4678 | ENSG00000132780 |
| 3 | Q13263 | NA | TRIM28 | tripartite motif containing 28 | 10155 | ENSG00000130726 |

  
  

| **Database:biological process      &nbspName:developmental process      &nbspID:GO:0032502** | | | | | | |
| --- | --- | --- | --- | --- | --- | --- |
| C=4561; O=11; E=5.69; R=1.93; rawP=0.0094; adjP=0.0792 | | | | | | |
| Index | UserID | Value | Gene Symbol | Gene Name | EntrezGene | Ensembl |
| 1 | Q7Z2K6 | NA | ERMP1 | endoplasmic reticulum metallopeptidase 1 | 79956 | ENSG00000099219 |
| 2 | P49321 | NA | NASP | nuclear autoantigenic sperm protein (histone-binding) | 4678 | ENSG00000132780 |
| 3 | P00491 | NA | PNP | purine nucleoside phosphorylase | 4860 | ENSG00000198805 |
| 4 | P26447 | NA | S100A4 | S100 calcium binding protein A4 | 6275 | ENSG00000196154 |
| 5 | Q9UL46 | NA | PSME2 | proteasome (prosome, macropain) activator subunit 2 (PA28 beta) | 5721 | ENSG00000100911 |
| 6 | P08727 | NA | KRT19 | keratin 19 | 3880 | ENSG00000171345 |
| 7 | P21980 | NA | TGM2 | transglutaminase 2 (C polypeptide, protein-glutamine-gamma-glutamyltransferase) | 7052 | ENSG00000198959 |
| 8 | P05783 | NA | KRT18 | keratin 18 | 3875 | ENSG00000111057 |
| 9 | Q13263 | NA | TRIM28 | tripartite motif containing 28 | 10155 | ENSG00000130726 |
| 10 | Q06830 | NA | PRDX1 | peroxiredoxin 1 | 5052 | ENSG00000117450 |
| 11 | P43490 | NA | NAMPT | nicotinamide phosphoribosyltransferase | 10135 | ENSG00000105835 |

  
  

| **Database:biological process      &nbspName:intracellular protein transport      &nbspID:GO:0006886** | | | | | | |
| --- | --- | --- | --- | --- | --- | --- |
| C=691; O=4; E=0.86; R=4.64; rawP=0.0093; adjP=0.0792 | | | | | | |
| Index | UserID | Value | Gene Symbol | Gene Name | EntrezGene | Ensembl |
| 1 | O14980 | NA | XPO1 | exportin 1 (CRM1 homolog, yeast) | 7514 | ENSG00000082898 |
| 2 | P05783 | NA | KRT18 | keratin 18 | 3875 | ENSG00000111057 |
| 3 | Q9Y2X3 | NA | NOP58 | NOP58 ribonucleoprotein homolog (yeast) | 51602 | ENSG00000055044 |
| 4 | Q06830 | NA | PRDX1 | peroxiredoxin 1 | 5052 | ENSG00000117450 |

  
  

| **Database:biological process      &nbspName:organ development      &nbspID:GO:0048513** | | | | | | |
| --- | --- | --- | --- | --- | --- | --- |
| C=2550; O=8; E=3.18; R=2.51; rawP=0.0077; adjP=0.0792 | | | | | | |
| Index | UserID | Value | Gene Symbol | Gene Name | EntrezGene | Ensembl |
| 1 | Q7Z2K6 | NA | ERMP1 | endoplasmic reticulum metallopeptidase 1 | 79956 | ENSG00000099219 |
| 2 | P00491 | NA | PNP | purine nucleoside phosphorylase | 4860 | ENSG00000198805 |
| 3 | P26447 | NA | S100A4 | S100 calcium binding protein A4 | 6275 | ENSG00000196154 |
| 4 | Q9UL46 | NA | PSME2 | proteasome (prosome, macropain) activator subunit 2 (PA28 beta) | 5721 | ENSG00000100911 |
| 5 | P21980 | NA | TGM2 | transglutaminase 2 (C polypeptide, protein-glutamine-gamma-glutamyltransferase) | 7052 | ENSG00000198959 |
| 6 | P08727 | NA | KRT19 | keratin 19 | 3880 | ENSG00000171345 |
| 7 | Q13263 | NA | TRIM28 | tripartite motif containing 28 | 10155 | ENSG00000130726 |
| 8 | P43490 | NA | NAMPT | nicotinamide phosphoribosyltransferase | 10135 | ENSG00000105835 |

  
  

| **Database:biological process      &nbspName:nucleocytoplasmic transport      &nbspID:GO:0006913** | | | | | | |
| --- | --- | --- | --- | --- | --- | --- |
| C=337; O=3; E=0.42; R=7.13; rawP=0.0080; adjP=0.0792 | | | | | | |
| Index | UserID | Value | Gene Symbol | Gene Name | EntrezGene | Ensembl |
| 1 | O14980 | NA | XPO1 | exportin 1 (CRM1 homolog, yeast) | 7514 | ENSG00000082898 |
| 2 | Q9Y2X3 | NA | NOP58 | NOP58 ribonucleoprotein homolog (yeast) | 51602 | ENSG00000055044 |
| 3 | Q06830 | NA | PRDX1 | peroxiredoxin 1 | 5052 | ENSG00000117450 |

  
  

| **Database:biological process      &nbspName:nuclear transport      &nbspID:GO:0051169** | | | | | | |
| --- | --- | --- | --- | --- | --- | --- |
| C=340; O=3; E=0.42; R=7.07; rawP=0.0081; adjP=0.0792 | | | | | | |
| Index | UserID | Value | Gene Symbol | Gene Name | EntrezGene | Ensembl |
| 1 | O14980 | NA | XPO1 | exportin 1 (CRM1 homolog, yeast) | 7514 | ENSG00000082898 |
| 2 | Q9Y2X3 | NA | NOP58 | NOP58 ribonucleoprotein homolog (yeast) | 51602 | ENSG00000055044 |
| 3 | Q06830 | NA | PRDX1 | peroxiredoxin 1 | 5052 | ENSG00000117450 |

  
  

| **Database:biological process      &nbspName:protein localization      &nbspID:GO:0008104** | | | | | | |
| --- | --- | --- | --- | --- | --- | --- |
| C=1559; O=6; E=1.95; R=3.08; rawP=0.0093; adjP=0.0792 | | | | | | |
| Index | UserID | Value | Gene Symbol | Gene Name | EntrezGene | Ensembl |
| 1 | O14980 | NA | XPO1 | exportin 1 (CRM1 homolog, yeast) | 7514 | ENSG00000082898 |
| 2 | P49321 | NA | NASP | nuclear autoantigenic sperm protein (histone-binding) | 4678 | ENSG00000132780 |
| 3 | P00491 | NA | PNP | purine nucleoside phosphorylase | 4860 | ENSG00000198805 |
| 4 | P05783 | NA | KRT18 | keratin 18 | 3875 | ENSG00000111057 |
| 5 | Q9Y2X3 | NA | NOP58 | NOP58 ribonucleoprotein homolog (yeast) | 51602 | ENSG00000055044 |
| 6 | Q06830 | NA | PRDX1 | peroxiredoxin 1 | 5052 | ENSG00000117450 |

  
  

| **Database:biological process      &nbspName:multi-organism reproductive process      &nbspID:GO:0044703** | | | | | | |
| --- | --- | --- | --- | --- | --- | --- |
| C=718; O=4; E=0.90; R=4.46; rawP=0.0107; adjP=0.0856 | | | | | | |
| Index | UserID | Value | Gene Symbol | Gene Name | EntrezGene | Ensembl |
| 1 | P08727 | NA | KRT19 | keratin 19 | 3880 | ENSG00000171345 |
| 2 | O14980 | NA | XPO1 | exportin 1 (CRM1 homolog, yeast) | 7514 | ENSG00000082898 |
| 3 | P05783 | NA | KRT18 | keratin 18 | 3875 | ENSG00000111057 |
| 4 | P05787 | NA | KRT8 | keratin 8 | 3856 | ENSG00000170421 |

  
  

| **Database:molecular function      &nbspName:transferase activity, transferring glycosyl groups      &nbspID:GO:0016757** | | | | | | |
| --- | --- | --- | --- | --- | --- | --- |
| C=263; O=3; E=0.32; R=9.49; rawP=0.0036; adjP=0.0576 | | | | | | |
| Index | UserID | Value | Gene Symbol | Gene Name | EntrezGene | Ensembl |
| 1 | Q04446 | NA | GBE1 | glucan (1,4-alpha-), branching enzyme 1 | 2632 | ENSG00000114480 |
| 2 | P00491 | NA | PNP | purine nucleoside phosphorylase | 4860 | ENSG00000198805 |
| 3 | P43490 | NA | NAMPT | nicotinamide phosphoribosyltransferase | 10135 | ENSG00000105835 |

  
  

| **Database:molecular function      &nbspName:binding      &nbspID:GO:0005488** | | | | | | |
| --- | --- | --- | --- | --- | --- | --- |
| C=11779; O=18; E=14.16; R=1.27; rawP=0.0132; adjP=0.0704 | | | | | | |
| Index | UserID | Value | Gene Symbol | Gene Name | EntrezGene | Ensembl |
| 1 | Q04446 | NA | GBE1 | glucan (1,4-alpha-), branching enzyme 1 | 2632 | ENSG00000114480 |
| 2 | Q7Z2K6 | NA | ERMP1 | endoplasmic reticulum metallopeptidase 1 | 79956 | ENSG00000099219 |
| 3 | P49321 | NA | NASP | nuclear autoantigenic sperm protein (histone-binding) | 4678 | ENSG00000132780 |
| 4 | P00491 | NA | PNP | purine nucleoside phosphorylase | 4860 | ENSG00000198805 |
| 5 | P05787 | NA | KRT8 | keratin 8 | 3856 | ENSG00000170421 |
| 6 | P26447 | NA | S100A4 | S100 calcium binding protein A4 | 6275 | ENSG00000196154 |
| 7 | Q9Y2X3 | NA | NOP58 | NOP58 ribonucleoprotein homolog (yeast) | 51602 | ENSG00000055044 |
| 8 | Q13263 | NA | TRIM28 | tripartite motif containing 28 | 10155 | ENSG00000130726 |
| 9 | Q06830 | NA | PRDX1 | peroxiredoxin 1 | 5052 | ENSG00000117450 |
| 10 | P43490 | NA | NAMPT | nicotinamide phosphoribosyltransferase | 10135 | ENSG00000105835 |
| 11 | O14980 | NA | XPO1 | exportin 1 (CRM1 homolog, yeast) | 7514 | ENSG00000082898 |
| 12 | O00264 | NA | PGRMC1 | progesterone receptor membrane component 1 | 10857 | ENSG00000101856 |
| 13 | P49419 | NA | ALDH7A1 | aldehyde dehydrogenase 7 family, member A1 | 501 | ENSG00000164904 |
| 14 | P21980 | NA | TGM2 | transglutaminase 2 (C polypeptide, protein-glutamine-gamma-glutamyltransferase) | 7052 | ENSG00000198959 |
| 15 | P08727 | NA | KRT19 | keratin 19 | 3880 | ENSG00000171345 |
| 16 | P05783 | NA | KRT18 | keratin 18 | 3875 | ENSG00000111057 |
| 17 | P00352 | NA | ALDH1A1 | aldehyde dehydrogenase 1 family, member A1 | 216 | ENSG00000165092 |
| 18 | Q8NC51 | NA | SERBP1 | SERPINE1 mRNA binding protein 1 | 26135 | ENSG00000142864 |

  
  

| **Database:molecular function      &nbspName:protein binding      &nbspID:GO:0005515** | | | | | | |
| --- | --- | --- | --- | --- | --- | --- |
| C=7301; O=14; E=8.77; R=1.60; rawP=0.0117; adjP=0.0704 | | | | | | |
| Index | UserID | Value | Gene Symbol | Gene Name | EntrezGene | Ensembl |
| 1 | O14980 | NA | XPO1 | exportin 1 (CRM1 homolog, yeast) | 7514 | ENSG00000082898 |
| 2 | P49321 | NA | NASP | nuclear autoantigenic sperm protein (histone-binding) | 4678 | ENSG00000132780 |
| 3 | P05787 | NA | KRT8 | keratin 8 | 3856 | ENSG00000170421 |
| 4 | P26447 | NA | S100A4 | S100 calcium binding protein A4 | 6275 | ENSG00000196154 |
| 5 | O00264 | NA | PGRMC1 | progesterone receptor membrane component 1 | 10857 | ENSG00000101856 |
| 6 | P49419 | NA | ALDH7A1 | aldehyde dehydrogenase 7 family, member A1 | 501 | ENSG00000164904 |
| 7 | P08727 | NA | KRT19 | keratin 19 | 3880 | ENSG00000171345 |
| 8 | P21980 | NA | TGM2 | transglutaminase 2 (C polypeptide, protein-glutamine-gamma-glutamyltransferase) | 7052 | ENSG00000198959 |
| 9 | P05783 | NA | KRT18 | keratin 18 | 3875 | ENSG00000111057 |
| 10 | Q9Y2X3 | NA | NOP58 | NOP58 ribonucleoprotein homolog (yeast) | 51602 | ENSG00000055044 |
| 11 | Q13263 | NA | TRIM28 | tripartite motif containing 28 | 10155 | ENSG00000130726 |
| 12 | Q8NC51 | NA | SERBP1 | SERPINE1 mRNA binding protein 1 | 26135 | ENSG00000142864 |
| 13 | Q06830 | NA | PRDX1 | peroxiredoxin 1 | 5052 | ENSG00000117450 |
| 14 | P43490 | NA | NAMPT | nicotinamide phosphoribosyltransferase | 10135 | ENSG00000105835 |

  
  

| **Database:cellular component      &nbspName:cytoplasm      &nbspID:GO:0005737** | | | | | | |
| --- | --- | --- | --- | --- | --- | --- |
| C=9051; O=18; E=10.31; R=1.75; rawP=0.0002; adjP=0.0076 | | | | | | |
| Index | UserID | Value | Gene Symbol | Gene Name | EntrezGene | Ensembl |
| 1 | Q04446 | NA | GBE1 | glucan (1,4-alpha-), branching enzyme 1 | 2632 | ENSG00000114480 |
| 2 | Q7Z2K6 | NA | ERMP1 | endoplasmic reticulum metallopeptidase 1 | 79956 | ENSG00000099219 |
| 3 | P49321 | NA | NASP | nuclear autoantigenic sperm protein (histone-binding) | 4678 | ENSG00000132780 |
| 4 | P00491 | NA | PNP | purine nucleoside phosphorylase | 4860 | ENSG00000198805 |
| 5 | P05787 | NA | KRT8 | keratin 8 | 3856 | ENSG00000170421 |
| 6 | P26447 | NA | S100A4 | S100 calcium binding protein A4 | 6275 | ENSG00000196154 |
| 7 | Q9UL46 | NA | PSME2 | proteasome (prosome, macropain) activator subunit 2 (PA28 beta) | 5721 | ENSG00000100911 |
| 8 | Q9Y2X3 | NA | NOP58 | NOP58 ribonucleoprotein homolog (yeast) | 51602 | ENSG00000055044 |
| 9 | Q06830 | NA | PRDX1 | peroxiredoxin 1 | 5052 | ENSG00000117450 |
| 10 | P43490 | NA | NAMPT | nicotinamide phosphoribosyltransferase | 10135 | ENSG00000105835 |
| 11 | O14980 | NA | XPO1 | exportin 1 (CRM1 homolog, yeast) | 7514 | ENSG00000082898 |
| 12 | O00264 | NA | PGRMC1 | progesterone receptor membrane component 1 | 10857 | ENSG00000101856 |
| 13 | P49419 | NA | ALDH7A1 | aldehyde dehydrogenase 7 family, member A1 | 501 | ENSG00000164904 |
| 14 | P21980 | NA | TGM2 | transglutaminase 2 (C polypeptide, protein-glutamine-gamma-glutamyltransferase) | 7052 | ENSG00000198959 |
| 15 | P08727 | NA | KRT19 | keratin 19 | 3880 | ENSG00000171345 |
| 16 | P05783 | NA | KRT18 | keratin 18 | 3875 | ENSG00000111057 |
| 17 | P00352 | NA | ALDH1A1 | aldehyde dehydrogenase 1 family, member A1 | 216 | ENSG00000165092 |
| 18 | Q8NC51 | NA | SERBP1 | SERPINE1 mRNA binding protein 1 | 26135 | ENSG00000142864 |

  
  

| **Database:cellular component      &nbspName:cytoplasmic part      &nbspID:GO:0044444** | | | | | | |
| --- | --- | --- | --- | --- | --- | --- |
| C=6728; O=15; E=7.66; R=1.96; rawP=0.0007; adjP=0.0133 | | | | | | |
| Index | UserID | Value | Gene Symbol | Gene Name | EntrezGene | Ensembl |
| 1 | Q04446 | NA | GBE1 | glucan (1,4-alpha-), branching enzyme 1 | 2632 | ENSG00000114480 |
| 2 | O14980 | NA | XPO1 | exportin 1 (CRM1 homolog, yeast) | 7514 | ENSG00000082898 |
| 3 | Q7Z2K6 | NA | ERMP1 | endoplasmic reticulum metallopeptidase 1 | 79956 | ENSG00000099219 |
| 4 | P00491 | NA | PNP | purine nucleoside phosphorylase | 4860 | ENSG00000198805 |
| 5 | P26447 | NA | S100A4 | S100 calcium binding protein A4 | 6275 | ENSG00000196154 |
| 6 | Q9UL46 | NA | PSME2 | proteasome (prosome, macropain) activator subunit 2 (PA28 beta) | 5721 | ENSG00000100911 |
| 7 | O00264 | NA | PGRMC1 | progesterone receptor membrane component 1 | 10857 | ENSG00000101856 |
| 8 | P49419 | NA | ALDH7A1 | aldehyde dehydrogenase 7 family, member A1 | 501 | ENSG00000164904 |
| 9 | P08727 | NA | KRT19 | keratin 19 | 3880 | ENSG00000171345 |
| 10 | P21980 | NA | TGM2 | transglutaminase 2 (C polypeptide, protein-glutamine-gamma-glutamyltransferase) | 7052 | ENSG00000198959 |
| 11 | P00352 | NA | ALDH1A1 | aldehyde dehydrogenase 1 family, member A1 | 216 | ENSG00000165092 |
| 12 | P05783 | NA | KRT18 | keratin 18 | 3875 | ENSG00000111057 |
| 13 | Q8NC51 | NA | SERBP1 | SERPINE1 mRNA binding protein 1 | 26135 | ENSG00000142864 |
| 14 | Q06830 | NA | PRDX1 | peroxiredoxin 1 | 5052 | ENSG00000117450 |
| 15 | P43490 | NA | NAMPT | nicotinamide phosphoribosyltransferase | 10135 | ENSG00000105835 |

  
  

| **Database:cellular component      &nbspName:intermediate filament      &nbspID:GO:0005882** | | | | | | |
| --- | --- | --- | --- | --- | --- | --- |
| C=189; O=3; E=0.22; R=13.94; rawP=0.0012; adjP=0.0152 | | | | | | |
| Index | UserID | Value | Gene Symbol | Gene Name | EntrezGene | Ensembl |
| 1 | P08727 | NA | KRT19 | keratin 19 | 3880 | ENSG00000171345 |
| 2 | P05783 | NA | KRT18 | keratin 18 | 3875 | ENSG00000111057 |
| 3 | P05787 | NA | KRT8 | keratin 8 | 3856 | ENSG00000170421 |

  
  

| **Database:cellular component      &nbspName:intermediate filament cytoskeleton      &nbspID:GO:0045111** | | | | | | |
| --- | --- | --- | --- | --- | --- | --- |
| C=229; O=3; E=0.26; R=11.50; rawP=0.0021; adjP=0.0167 | | | | | | |
| Index | UserID | Value | Gene Symbol | Gene Name | EntrezGene | Ensembl |
| 1 | P08727 | NA | KRT19 | keratin 19 | 3880 | ENSG00000171345 |
| 2 | P05783 | NA | KRT18 | keratin 18 | 3875 | ENSG00000111057 |
| 3 | P05787 | NA | KRT8 | keratin 8 | 3856 | ENSG00000170421 |

  
  

| **Database:cellular component      &nbspName:intracellular part      &nbspID:GO:0044424** | | | | | | |
| --- | --- | --- | --- | --- | --- | --- |
| C=12096; O=19; E=13.78; R=1.38; rawP=0.0022; adjP=0.0167 | | | | | | |
| Index | UserID | Value | Gene Symbol | Gene Name | EntrezGene | Ensembl |
| 1 | Q04446 | NA | GBE1 | glucan (1,4-alpha-), branching enzyme 1 | 2632 | ENSG00000114480 |
| 2 | Q7Z2K6 | NA | ERMP1 | endoplasmic reticulum metallopeptidase 1 | 79956 | ENSG00000099219 |
| 3 | P49321 | NA | NASP | nuclear autoantigenic sperm protein (histone-binding) | 4678 | ENSG00000132780 |
| 4 | P00491 | NA | PNP | purine nucleoside phosphorylase | 4860 | ENSG00000198805 |
| 5 | P05787 | NA | KRT8 | keratin 8 | 3856 | ENSG00000170421 |
| 6 | P26447 | NA | S100A4 | S100 calcium binding protein A4 | 6275 | ENSG00000196154 |
| 7 | Q9UL46 | NA | PSME2 | proteasome (prosome, macropain) activator subunit 2 (PA28 beta) | 5721 | ENSG00000100911 |
| 8 | Q13263 | NA | TRIM28 | tripartite motif containing 28 | 10155 | ENSG00000130726 |
| 9 | Q9Y2X3 | NA | NOP58 | NOP58 ribonucleoprotein homolog (yeast) | 51602 | ENSG00000055044 |
| 10 | Q06830 | NA | PRDX1 | peroxiredoxin 1 | 5052 | ENSG00000117450 |
| 11 | P43490 | NA | NAMPT | nicotinamide phosphoribosyltransferase | 10135 | ENSG00000105835 |
| 12 | O14980 | NA | XPO1 | exportin 1 (CRM1 homolog, yeast) | 7514 | ENSG00000082898 |
| 13 | O00264 | NA | PGRMC1 | progesterone receptor membrane component 1 | 10857 | ENSG00000101856 |
| 14 | P49419 | NA | ALDH7A1 | aldehyde dehydrogenase 7 family, member A1 | 501 | ENSG00000164904 |
| 15 | P21980 | NA | TGM2 | transglutaminase 2 (C polypeptide, protein-glutamine-gamma-glutamyltransferase) | 7052 | ENSG00000198959 |
| 16 | P08727 | NA | KRT19 | keratin 19 | 3880 | ENSG00000171345 |
| 17 | P00352 | NA | ALDH1A1 | aldehyde dehydrogenase 1 family, member A1 | 216 | ENSG00000165092 |
| 18 | P05783 | NA | KRT18 | keratin 18 | 3875 | ENSG00000111057 |
| 19 | Q8NC51 | NA | SERBP1 | SERPINE1 mRNA binding protein 1 | 26135 | ENSG00000142864 |

  
  

| **Database:cellular component      &nbspName:intracellular      &nbspID:GO:0005622** | | | | | | |
| --- | --- | --- | --- | --- | --- | --- |
| C=12412; O=19; E=14.14; R=1.34; rawP=0.0036; adjP=0.0228 | | | | | | |
| Index | UserID | Value | Gene Symbol | Gene Name | EntrezGene | Ensembl |
| 1 | Q04446 | NA | GBE1 | glucan (1,4-alpha-), branching enzyme 1 | 2632 | ENSG00000114480 |
| 2 | Q7Z2K6 | NA | ERMP1 | endoplasmic reticulum metallopeptidase 1 | 79956 | ENSG00000099219 |
| 3 | P49321 | NA | NASP | nuclear autoantigenic sperm protein (histone-binding) | 4678 | ENSG00000132780 |
| 4 | P00491 | NA | PNP | purine nucleoside phosphorylase | 4860 | ENSG00000198805 |
| 5 | P05787 | NA | KRT8 | keratin 8 | 3856 | ENSG00000170421 |
| 6 | P26447 | NA | S100A4 | S100 calcium binding protein A4 | 6275 | ENSG00000196154 |
| 7 | Q9UL46 | NA | PSME2 | proteasome (prosome, macropain) activator subunit 2 (PA28 beta) | 5721 | ENSG00000100911 |
| 8 | Q13263 | NA | TRIM28 | tripartite motif containing 28 | 10155 | ENSG00000130726 |
| 9 | Q9Y2X3 | NA | NOP58 | NOP58 ribonucleoprotein homolog (yeast) | 51602 | ENSG00000055044 |
| 10 | Q06830 | NA | PRDX1 | peroxiredoxin 1 | 5052 | ENSG00000117450 |
| 11 | P43490 | NA | NAMPT | nicotinamide phosphoribosyltransferase | 10135 | ENSG00000105835 |
| 12 | O14980 | NA | XPO1 | exportin 1 (CRM1 homolog, yeast) | 7514 | ENSG00000082898 |
| 13 | O00264 | NA | PGRMC1 | progesterone receptor membrane component 1 | 10857 | ENSG00000101856 |
| 14 | P49419 | NA | ALDH7A1 | aldehyde dehydrogenase 7 family, member A1 | 501 | ENSG00000164904 |
| 15 | P21980 | NA | TGM2 | transglutaminase 2 (C polypeptide, protein-glutamine-gamma-glutamyltransferase) | 7052 | ENSG00000198959 |
| 16 | P08727 | NA | KRT19 | keratin 19 | 3880 | ENSG00000171345 |
| 17 | P00352 | NA | ALDH1A1 | aldehyde dehydrogenase 1 family, member A1 | 216 | ENSG00000165092 |
| 18 | P05783 | NA | KRT18 | keratin 18 | 3875 | ENSG00000111057 |
| 19 | Q8NC51 | NA | SERBP1 | SERPINE1 mRNA binding protein 1 | 26135 | ENSG00000142864 |

  
  

| **Database:cellular component      &nbspName:cytosol      &nbspID:GO:0005829** | | | | | | |
| --- | --- | --- | --- | --- | --- | --- |
| C=2367; O=7; E=2.70; R=2.60; rawP=0.0121; adjP=0.0580 | | | | | | |
| Index | UserID | Value | Gene Symbol | Gene Name | EntrezGene | Ensembl |
| 1 | Q04446 | NA | GBE1 | glucan (1,4-alpha-), branching enzyme 1 | 2632 | ENSG00000114480 |
| 2 | P49419 | NA | ALDH7A1 | aldehyde dehydrogenase 7 family, member A1 | 501 | ENSG00000164904 |
| 3 | O14980 | NA | XPO1 | exportin 1 (CRM1 homolog, yeast) | 7514 | ENSG00000082898 |
| 4 | P00491 | NA | PNP | purine nucleoside phosphorylase | 4860 | ENSG00000198805 |
| 5 | P00352 | NA | ALDH1A1 | aldehyde dehydrogenase 1 family, member A1 | 216 | ENSG00000165092 |
| 6 | Q9UL46 | NA | PSME2 | proteasome (prosome, macropain) activator subunit 2 (PA28 beta) | 5721 | ENSG00000100911 |
| 7 | P43490 | NA | NAMPT | nicotinamide phosphoribosyltransferase | 10135 | ENSG00000105835 |

  
  

| **Database:cellular component      &nbspName:nucleus      &nbspID:GO:0005634** | | | | | | |
| --- | --- | --- | --- | --- | --- | --- |
| C=5883; O=12; E=6.70; R=1.79; rawP=0.0122; adjP=0.0580 | | | | | | |
| Index | UserID | Value | Gene Symbol | Gene Name | EntrezGene | Ensembl |
| 1 | O14980 | NA | XPO1 | exportin 1 (CRM1 homolog, yeast) | 7514 | ENSG00000082898 |
| 2 | P49321 | NA | NASP | nuclear autoantigenic sperm protein (histone-binding) | 4678 | ENSG00000132780 |
| 3 | P05787 | NA | KRT8 | keratin 8 | 3856 | ENSG00000170421 |
| 4 | P26447 | NA | S100A4 | S100 calcium binding protein A4 | 6275 | ENSG00000196154 |
| 5 | Q9UL46 | NA | PSME2 | proteasome (prosome, macropain) activator subunit 2 (PA28 beta) | 5721 | ENSG00000100911 |
| 6 | O00264 | NA | PGRMC1 | progesterone receptor membrane component 1 | 10857 | ENSG00000101856 |
| 7 | P49419 | NA | ALDH7A1 | aldehyde dehydrogenase 7 family, member A1 | 501 | ENSG00000164904 |
| 8 | P05783 | NA | KRT18 | keratin 18 | 3875 | ENSG00000111057 |
| 9 | Q9Y2X3 | NA | NOP58 | NOP58 ribonucleoprotein homolog (yeast) | 51602 | ENSG00000055044 |
| 10 | Q13263 | NA | TRIM28 | tripartite motif containing 28 | 10155 | ENSG00000130726 |
| 11 | Q8NC51 | NA | SERBP1 | SERPINE1 mRNA binding protein 1 | 26135 | ENSG00000142864 |
| 12 | Q06830 | NA | PRDX1 | peroxiredoxin 1 | 5052 | ENSG00000117450 |

  
  

| **Database:cellular component      &nbspName:perinuclear region of cytoplasm      &nbspID:GO:0048471** | | | | | | |
| --- | --- | --- | --- | --- | --- | --- |
| C=467; O=3; E=0.53; R=5.64; rawP=0.0151; adjP=0.0638 | | | | | | |
| Index | UserID | Value | Gene Symbol | Gene Name | EntrezGene | Ensembl |
| 1 | P05783 | NA | KRT18 | keratin 18 | 3875 | ENSG00000111057 |
| 2 | P26447 | NA | S100A4 | S100 calcium binding protein A4 | 6275 | ENSG00000196154 |
| 3 | Q8NC51 | NA | SERBP1 | SERPINE1 mRNA binding protein 1 | 26135 | ENSG00000142864 |

  
  

| **Database:cellular component      &nbspName:nuclear lumen      &nbspID:GO:0031981** | | | | | | |
| --- | --- | --- | --- | --- | --- | --- |
| C=2728; O=7; E=3.11; R=2.25; rawP=0.0254; adjP=0.0689 | | | | | | |
| Index | UserID | Value | Gene Symbol | Gene Name | EntrezGene | Ensembl |
| 1 | O14980 | NA | XPO1 | exportin 1 (CRM1 homolog, yeast) | 7514 | ENSG00000082898 |
| 2 | P05783 | NA | KRT18 | keratin 18 | 3875 | ENSG00000111057 |
| 3 | P05787 | NA | KRT8 | keratin 8 | 3856 | ENSG00000170421 |
| 4 | Q13263 | NA | TRIM28 | tripartite motif containing 28 | 10155 | ENSG00000130726 |
| 5 | Q9Y2X3 | NA | NOP58 | NOP58 ribonucleoprotein homolog (yeast) | 51602 | ENSG00000055044 |
| 6 | Q9UL46 | NA | PSME2 | proteasome (prosome, macropain) activator subunit 2 (PA28 beta) | 5721 | ENSG00000100911 |
| 7 | O00264 | NA | PGRMC1 | progesterone receptor membrane component 1 | 10857 | ENSG00000101856 |

  
  

| **Database:cellular component      &nbspName:intracellular organelle lumen      &nbspID:GO:0070013** | | | | | | |
| --- | --- | --- | --- | --- | --- | --- |
| C=3285; O=8; E=3.74; R=2.14; rawP=0.0212; adjP=0.0689 | | | | | | |
| Index | UserID | Value | Gene Symbol | Gene Name | EntrezGene | Ensembl |
| 1 | O14980 | NA | XPO1 | exportin 1 (CRM1 homolog, yeast) | 7514 | ENSG00000082898 |
| 2 | P05787 | NA | KRT8 | keratin 8 | 3856 | ENSG00000170421 |
| 3 | Q9UL46 | NA | PSME2 | proteasome (prosome, macropain) activator subunit 2 (PA28 beta) | 5721 | ENSG00000100911 |
| 4 | O00264 | NA | PGRMC1 | progesterone receptor membrane component 1 | 10857 | ENSG00000101856 |
| 5 | P49419 | NA | ALDH7A1 | aldehyde dehydrogenase 7 family, member A1 | 501 | ENSG00000164904 |
| 6 | P05783 | NA | KRT18 | keratin 18 | 3875 | ENSG00000111057 |
| 7 | Q9Y2X3 | NA | NOP58 | NOP58 ribonucleoprotein homolog (yeast) | 51602 | ENSG00000055044 |
| 8 | Q13263 | NA | TRIM28 | tripartite motif containing 28 | 10155 | ENSG00000130726 |

  
  

| **Database:cellular component      &nbspName:nucleoplasm      &nbspID:GO:0005654** | | | | | | |
| --- | --- | --- | --- | --- | --- | --- |
| C=1476; O=5; E=1.68; R=2.97; rawP=0.0219; adjP=0.0689 | | | | | | |
| Index | UserID | Value | Gene Symbol | Gene Name | EntrezGene | Ensembl |
| 1 | O14980 | NA | XPO1 | exportin 1 (CRM1 homolog, yeast) | 7514 | ENSG00000082898 |
| 2 | P05787 | NA | KRT8 | keratin 8 | 3856 | ENSG00000170421 |
| 3 | Q13263 | NA | TRIM28 | tripartite motif containing 28 | 10155 | ENSG00000130726 |
| 4 | Q9Y2X3 | NA | NOP58 | NOP58 ribonucleoprotein homolog (yeast) | 51602 | ENSG00000055044 |
| 5 | Q9UL46 | NA | PSME2 | proteasome (prosome, macropain) activator subunit 2 (PA28 beta) | 5721 | ENSG00000100911 |

  
  

| **Database:cellular component      &nbspName:organelle lumen      &nbspID:GO:0043233** | | | | | | |
| --- | --- | --- | --- | --- | --- | --- |
| C=3331; O=8; E=3.79; R=2.11; rawP=0.0230; adjP=0.0689 | | | | | | |
| Index | UserID | Value | Gene Symbol | Gene Name | EntrezGene | Ensembl |
| 1 | O14980 | NA | XPO1 | exportin 1 (CRM1 homolog, yeast) | 7514 | ENSG00000082898 |
| 2 | P05787 | NA | KRT8 | keratin 8 | 3856 | ENSG00000170421 |
| 3 | Q9UL46 | NA | PSME2 | proteasome (prosome, macropain) activator subunit 2 (PA28 beta) | 5721 | ENSG00000100911 |
| 4 | O00264 | NA | PGRMC1 | progesterone receptor membrane component 1 | 10857 | ENSG00000101856 |
| 5 | P49419 | NA | ALDH7A1 | aldehyde dehydrogenase 7 family, member A1 | 501 | ENSG00000164904 |
| 6 | P05783 | NA | KRT18 | keratin 18 | 3875 | ENSG00000111057 |
| 7 | Q9Y2X3 | NA | NOP58 | NOP58 ribonucleoprotein homolog (yeast) | 51602 | ENSG00000055044 |
| 8 | Q13263 | NA | TRIM28 | tripartite motif containing 28 | 10155 | ENSG00000130726 |

  
  

| **Database:cellular component      &nbspName:membrane-enclosed lumen      &nbspID:GO:0031974** | | | | | | |
| --- | --- | --- | --- | --- | --- | --- |
| C=3375; O=8; E=3.84; R=2.08; rawP=0.0248; adjP=0.0689 | | | | | | |
| Index | UserID | Value | Gene Symbol | Gene Name | EntrezGene | Ensembl |
| 1 | O14980 | NA | XPO1 | exportin 1 (CRM1 homolog, yeast) | 7514 | ENSG00000082898 |
| 2 | P05787 | NA | KRT8 | keratin 8 | 3856 | ENSG00000170421 |
| 3 | Q9UL46 | NA | PSME2 | proteasome (prosome, macropain) activator subunit 2 (PA28 beta) | 5721 | ENSG00000100911 |
| 4 | O00264 | NA | PGRMC1 | progesterone receptor membrane component 1 | 10857 | ENSG00000101856 |
| 5 | P49419 | NA | ALDH7A1 | aldehyde dehydrogenase 7 family, member A1 | 501 | ENSG00000164904 |
| 6 | P05783 | NA | KRT18 | keratin 18 | 3875 | ENSG00000111057 |
| 7 | Q9Y2X3 | NA | NOP58 | NOP58 ribonucleoprotein homolog (yeast) | 51602 | ENSG00000055044 |
| 8 | Q13263 | NA | TRIM28 | tripartite motif containing 28 | 10155 | ENSG00000130726 |

  
  

| **Database:cellular component      &nbspName:organelle      &nbspID:GO:0043226** | | | | | | |
| --- | --- | --- | --- | --- | --- | --- |
| C=10536; O=16; E=12.00; R=1.33; rawP=0.0423; adjP=0.0924 | | | | | | |
| Index | UserID | Value | Gene Symbol | Gene Name | EntrezGene | Ensembl |
| 1 | Q7Z2K6 | NA | ERMP1 | endoplasmic reticulum metallopeptidase 1 | 79956 | ENSG00000099219 |
| 2 | P49321 | NA | NASP | nuclear autoantigenic sperm protein (histone-binding) | 4678 | ENSG00000132780 |
| 3 | P00491 | NA | PNP | purine nucleoside phosphorylase | 4860 | ENSG00000198805 |
| 4 | P05787 | NA | KRT8 | keratin 8 | 3856 | ENSG00000170421 |
| 5 | P26447 | NA | S100A4 | S100 calcium binding protein A4 | 6275 | ENSG00000196154 |
| 6 | Q9UL46 | NA | PSME2 | proteasome (prosome, macropain) activator subunit 2 (PA28 beta) | 5721 | ENSG00000100911 |
| 7 | Q9Y2X3 | NA | NOP58 | NOP58 ribonucleoprotein homolog (yeast) | 51602 | ENSG00000055044 |
| 8 | Q13263 | NA | TRIM28 | tripartite motif containing 28 | 10155 | ENSG00000130726 |
| 9 | Q06830 | NA | PRDX1 | peroxiredoxin 1 | 5052 | ENSG00000117450 |
| 10 | O14980 | NA | XPO1 | exportin 1 (CRM1 homolog, yeast) | 7514 | ENSG00000082898 |
| 11 | O00264 | NA | PGRMC1 | progesterone receptor membrane component 1 | 10857 | ENSG00000101856 |
| 12 | P49419 | NA | ALDH7A1 | aldehyde dehydrogenase 7 family, member A1 | 501 | ENSG00000164904 |
| 13 | P21980 | NA | TGM2 | transglutaminase 2 (C polypeptide, protein-glutamine-gamma-glutamyltransferase) | 7052 | ENSG00000198959 |
| 14 | P08727 | NA | KRT19 | keratin 19 | 3880 | ENSG00000171345 |
| 15 | P05783 | NA | KRT18 | keratin 18 | 3875 | ENSG00000111057 |
| 16 | Q8NC51 | NA | SERBP1 | SERPINE1 mRNA binding protein 1 | 26135 | ENSG00000142864 |

  
  

| **Database:cellular component      &nbspName:nuclear part      &nbspID:GO:0044428** | | | | | | |
| --- | --- | --- | --- | --- | --- | --- |
| C=3057; O=7; E=3.48; R=2.01; rawP=0.0447; adjP=0.0924 | | | | | | |
| Index | UserID | Value | Gene Symbol | Gene Name | EntrezGene | Ensembl |
| 1 | O14980 | NA | XPO1 | exportin 1 (CRM1 homolog, yeast) | 7514 | ENSG00000082898 |
| 2 | P05783 | NA | KRT18 | keratin 18 | 3875 | ENSG00000111057 |
| 3 | P05787 | NA | KRT8 | keratin 8 | 3856 | ENSG00000170421 |
| 4 | Q13263 | NA | TRIM28 | tripartite motif containing 28 | 10155 | ENSG00000130726 |
| 5 | Q9Y2X3 | NA | NOP58 | NOP58 ribonucleoprotein homolog (yeast) | 51602 | ENSG00000055044 |
| 6 | Q9UL46 | NA | PSME2 | proteasome (prosome, macropain) activator subunit 2 (PA28 beta) | 5721 | ENSG00000100911 |
| 7 | O00264 | NA | PGRMC1 | progesterone receptor membrane component 1 | 10857 | ENSG00000101856 |

  
  

| **Database:cellular component      &nbspName:intracellular organelle      &nbspID:GO:0043229** | | | | | | |
| --- | --- | --- | --- | --- | --- | --- |
| C=10521; O=16; E=11.98; R=1.34; rawP=0.0416; adjP=0.0924 | | | | | | |
| Index | UserID | Value | Gene Symbol | Gene Name | EntrezGene | Ensembl |
| 1 | Q7Z2K6 | NA | ERMP1 | endoplasmic reticulum metallopeptidase 1 | 79956 | ENSG00000099219 |
| 2 | P49321 | NA | NASP | nuclear autoantigenic sperm protein (histone-binding) | 4678 | ENSG00000132780 |
| 3 | P00491 | NA | PNP | purine nucleoside phosphorylase | 4860 | ENSG00000198805 |
| 4 | P05787 | NA | KRT8 | keratin 8 | 3856 | ENSG00000170421 |
| 5 | P26447 | NA | S100A4 | S100 calcium binding protein A4 | 6275 | ENSG00000196154 |
| 6 | Q9UL46 | NA | PSME2 | proteasome (prosome, macropain) activator subunit 2 (PA28 beta) | 5721 | ENSG00000100911 |
| 7 | Q9Y2X3 | NA | NOP58 | NOP58 ribonucleoprotein homolog (yeast) | 51602 | ENSG00000055044 |
| 8 | Q13263 | NA | TRIM28 | tripartite motif containing 28 | 10155 | ENSG00000130726 |
| 9 | Q06830 | NA | PRDX1 | peroxiredoxin 1 | 5052 | ENSG00000117450 |
| 10 | O14980 | NA | XPO1 | exportin 1 (CRM1 homolog, yeast) | 7514 | ENSG00000082898 |
| 11 | O00264 | NA | PGRMC1 | progesterone receptor membrane component 1 | 10857 | ENSG00000101856 |
| 12 | P49419 | NA | ALDH7A1 | aldehyde dehydrogenase 7 family, member A1 | 501 | ENSG00000164904 |
| 13 | P21980 | NA | TGM2 | transglutaminase 2 (C polypeptide, protein-glutamine-gamma-glutamyltransferase) | 7052 | ENSG00000198959 |
| 14 | P08727 | NA | KRT19 | keratin 19 | 3880 | ENSG00000171345 |
| 15 | P05783 | NA | KRT18 | keratin 18 | 3875 | ENSG00000111057 |
| 16 | Q8NC51 | NA | SERBP1 | SERPINE1 mRNA binding protein 1 | 26135 | ENSG00000142864 |

  
  

| **Database:cellular component      &nbspName:intracellular non-membrane-bounded organelle      &nbspID:GO:0043232** | | | | | | |
| --- | --- | --- | --- | --- | --- | --- |
| C=3778; O=8; E=4.30; R=1.86; rawP=0.0462; adjP=0.0924 | | | | | | |
| Index | UserID | Value | Gene Symbol | Gene Name | EntrezGene | Ensembl |
| 1 | O14980 | NA | XPO1 | exportin 1 (CRM1 homolog, yeast) | 7514 | ENSG00000082898 |
| 2 | P00491 | NA | PNP | purine nucleoside phosphorylase | 4860 | ENSG00000198805 |
| 3 | P05787 | NA | KRT8 | keratin 8 | 3856 | ENSG00000170421 |
| 4 | O00264 | NA | PGRMC1 | progesterone receptor membrane component 1 | 10857 | ENSG00000101856 |
| 5 | P08727 | NA | KRT19 | keratin 19 | 3880 | ENSG00000171345 |
| 6 | P05783 | NA | KRT18 | keratin 18 | 3875 | ENSG00000111057 |
| 7 | Q9Y2X3 | NA | NOP58 | NOP58 ribonucleoprotein homolog (yeast) | 51602 | ENSG00000055044 |
| 8 | Q13263 | NA | TRIM28 | tripartite motif containing 28 | 10155 | ENSG00000130726 |

  
  

| **Database:cellular component      &nbspName:non-membrane-bounded organelle      &nbspID:GO:0043228** | | | | | | |
| --- | --- | --- | --- | --- | --- | --- |
| C=3778; O=8; E=4.30; R=1.86; rawP=0.0462; adjP=0.0924 | | | | | | |
| Index | UserID | Value | Gene Symbol | Gene Name | EntrezGene | Ensembl |
| 1 | O14980 | NA | XPO1 | exportin 1 (CRM1 homolog, yeast) | 7514 | ENSG00000082898 |
| 2 | P00491 | NA | PNP | purine nucleoside phosphorylase | 4860 | ENSG00000198805 |
| 3 | P05787 | NA | KRT8 | keratin 8 | 3856 | ENSG00000170421 |
| 4 | O00264 | NA | PGRMC1 | progesterone receptor membrane component 1 | 10857 | ENSG00000101856 |
| 5 | P08727 | NA | KRT19 | keratin 19 | 3880 | ENSG00000171345 |
| 6 | P05783 | NA | KRT18 | keratin 18 | 3875 | ENSG00000111057 |
| 7 | Q9Y2X3 | NA | NOP58 | NOP58 ribonucleoprotein homolog (yeast) | 51602 | ENSG00000055044 |
| 8 | Q13263 | NA | TRIM28 | tripartite motif containing 28 | 10155 | ENSG00000130726 |

  
  
  
  

---

WebGestalt is currently developed and maintained by Jing Wang and Bing Zhang at the  Zhang Lab. Other people who have made significant contribution to the project include Dexter Duncan, Stefan Kirov, Zhiao Shi, and Jay Snoddy.  
  
**Funding credits:** NIH/NIAAA (U01 AA016662, U01 AA013512); NIH/NIDA (P01 DA015027); NIH/NIMH (P50 MH078028, P50 MH096972); NIH/NCI (U24 CA159988); NIH/NIGMS (R01 GM088822).
