## Supplementary material for "PGRMC1 phosphorylation and cell plasticity 1: glycolysis, mitochondria, tumor growth": File S8: final_tf_geneset_file_1439189707.html

Anchored HTML File of EIDs

|  |  |
| --- | --- |
|  | WEB-based GEne SeT AnaLysis Toolkit |
|  | ***Translating gene lists into biological insights...*** |

---

  

| **Database:Transcrription Target      &nbspName:hsa\_V$TATA\_01      &nbspID:DB\_ID:2025** | | | | | | |
| --- | --- | --- | --- | --- | --- | --- |
| C=254; O=4; E=0.24; R=16.81; rawP=8.03e-05; adjP=0.0010 | | | | | | |
| Index | UserID | Value | Gene Symbol | Gene Name | EntrezGene | Ensembl |
| 1 | Q04446 | NA | GBE1 | glucan (1,4-alpha-), branching enzyme 1 | 2632 | ENSG00000114480 |
| 2 | P49321 | NA | NASP | nuclear autoantigenic sperm protein (histone-binding) | 4678 | ENSG00000132780 |
| 3 | P26447 | NA | S100A4 | S100 calcium binding protein A4 | 6275 | ENSG00000196154 |
| 4 | Q06830 | NA | PRDX1 | peroxiredoxin 1 | 5052 | ENSG00000117450 |

  
  

| **Database:Transcrription Target      &nbspName:hsa\_STTTCRNTTT\_V$IRF\_Q6      &nbspID:DB\_ID:2426** | | | | | | |
| --- | --- | --- | --- | --- | --- | --- |
| C=184; O=3; E=0.17; R=17.41; rawP=0.0006; adjP=0.0039 | | | | | | |
| Index | UserID | Value | Gene Symbol | Gene Name | EntrezGene | Ensembl |
| 1 | P00352 | NA | ALDH1A1 | aldehyde dehydrogenase 1 family, member A1 | 216 | ENSG00000165092 |
| 2 | Q9UL46 | NA | PSME2 | proteasome (prosome, macropain) activator subunit 2 (PA28 beta) | 5721 | ENSG00000100911 |
| 3 | P43490 | NA | NAMPT | nicotinamide phosphoribosyltransferase | 10135 | ENSG00000105835 |

  
  

| **Database:Transcrription Target      &nbspName:hsa\_V$ARNT\_01      &nbspID:DB\_ID:2011** | | | | | | |
| --- | --- | --- | --- | --- | --- | --- |
| C=255; O=3; E=0.24; R=12.56; rawP=0.0016; adjP=0.0055 | | | | | | |
| Index | UserID | Value | Gene Symbol | Gene Name | EntrezGene | Ensembl |
| 1 | O14980 | NA | XPO1 | exportin 1 (CRM1 homolog, yeast) | 7514 | ENSG00000082898 |
| 2 | Q9Y2X3 | NA | NOP58 | NOP58 ribonucleoprotein homolog (yeast) | 51602 | ENSG00000055044 |
| 3 | Q8NC51 | NA | SERBP1 | SERPINE1 mRNA binding protein 1 | 26135 | ENSG00000142864 |

  
  

| **Database:Transcrription Target      &nbspName:hsa\_V$MAX\_01      &nbspID:DB\_ID:1924** | | | | | | |
| --- | --- | --- | --- | --- | --- | --- |
| C=256; O=3; E=0.24; R=12.51; rawP=0.0017; adjP=0.0055 | | | | | | |
| Index | UserID | Value | Gene Symbol | Gene Name | EntrezGene | Ensembl |
| 1 | O14980 | NA | XPO1 | exportin 1 (CRM1 homolog, yeast) | 7514 | ENSG00000082898 |
| 2 | Q9Y2X3 | NA | NOP58 | NOP58 ribonucleoprotein homolog (yeast) | 51602 | ENSG00000055044 |
| 3 | Q8NC51 | NA | SERBP1 | SERPINE1 mRNA binding protein 1 | 26135 | ENSG00000142864 |

  
  

| **Database:Transcrription Target      &nbspName:hsa\_GCANCTGNY\_V$MYOD\_Q6      &nbspID:DB\_ID:2435** | | | | | | |
| --- | --- | --- | --- | --- | --- | --- |
| C=898; O=4; E=0.84; R=4.76; rawP=0.0087; adjP=0.0226 | | | | | | |
| Index | UserID | Value | Gene Symbol | Gene Name | EntrezGene | Ensembl |
| 1 | P08727 | NA | KRT19 | keratin 19 | 3880 | ENSG00000171345 |
| 2 | O14980 | NA | XPO1 | exportin 1 (CRM1 homolog, yeast) | 7514 | ENSG00000082898 |
| 3 | P00352 | NA | ALDH1A1 | aldehyde dehydrogenase 1 family, member A1 | 216 | ENSG00000165092 |
| 4 | P05787 | NA | KRT8 | keratin 8 | 3856 | ENSG00000170421 |

  
  

| **Database:Transcrription Target      &nbspName:hsa\_GGGCGGR\_V$SP1\_Q6      &nbspID:DB\_ID:2452** | | | | | | |
| --- | --- | --- | --- | --- | --- | --- |
| C=2863; O=7; E=2.68; R=2.61; rawP=0.0118; adjP=0.0256 | | | | | | |
| Index | UserID | Value | Gene Symbol | Gene Name | EntrezGene | Ensembl |
| 1 | P49419 | NA | ALDH7A1 | aldehyde dehydrogenase 7 family, member A1 | 501 | ENSG00000164904 |
| 2 | P08727 | NA | KRT19 | keratin 19 | 3880 | ENSG00000171345 |
| 3 | O14980 | NA | XPO1 | exportin 1 (CRM1 homolog, yeast) | 7514 | ENSG00000082898 |
| 4 | P49321 | NA | NASP | nuclear autoantigenic sperm protein (histone-binding) | 4678 | ENSG00000132780 |
| 5 | Q13263 | NA | TRIM28 | tripartite motif containing 28 | 10155 | ENSG00000130726 |
| 6 | Q8NC51 | NA | SERBP1 | SERPINE1 mRNA binding protein 1 | 26135 | ENSG00000142864 |
| 7 | O00264 | NA | PGRMC1 | progesterone receptor membrane component 1 | 10857 | ENSG00000101856 |

  
  

| **Database:Transcrription Target      &nbspName:hsa\_TATAAA\_V$TATA\_01      &nbspID:DB\_ID:2456** | | | | | | |
| --- | --- | --- | --- | --- | --- | --- |
| C=1260; O=4; E=1.18; R=3.39; rawP=0.0271; adjP=0.0503 | | | | | | |
| Index | UserID | Value | Gene Symbol | Gene Name | EntrezGene | Ensembl |
| 1 | Q04446 | NA | GBE1 | glucan (1,4-alpha-), branching enzyme 1 | 2632 | ENSG00000114480 |
| 2 | P05787 | NA | KRT8 | keratin 8 | 3856 | ENSG00000170421 |
| 3 | P26447 | NA | S100A4 | S100 calcium binding protein A4 | 6275 | ENSG00000196154 |
| 4 | Q06830 | NA | PRDX1 | peroxiredoxin 1 | 5052 | ENSG00000117450 |

  
  

| **Database:Transcrription Target      &nbspName:hsa\_TTGTTT\_V$FOXO4\_01      &nbspID:DB\_ID:2416** | | | | | | |
| --- | --- | --- | --- | --- | --- | --- |
| C=2012; O=5; E=1.88; R=2.65; rawP=0.0341; adjP=0.0554 | | | | | | |
| Index | UserID | Value | Gene Symbol | Gene Name | EntrezGene | Ensembl |
| 1 | P49321 | NA | NASP | nuclear autoantigenic sperm protein (histone-binding) | 4678 | ENSG00000132780 |
| 2 | P00352 | NA | ALDH1A1 | aldehyde dehydrogenase 1 family, member A1 | 216 | ENSG00000165092 |
| 3 | Q8NC51 | NA | SERBP1 | SERPINE1 mRNA binding protein 1 | 26135 | ENSG00000142864 |
| 4 | Q06830 | NA | PRDX1 | peroxiredoxin 1 | 5052 | ENSG00000117450 |
| 5 | O00264 | NA | PGRMC1 | progesterone receptor membrane component 1 | 10857 | ENSG00000101856 |

  
  

| **Database:Transcrription Target      &nbspName:hsa\_CTTTAAR\_UNKNOWN      &nbspID:DB\_ID:2170** | | | | | | |
| --- | --- | --- | --- | --- | --- | --- |
| C=946; O=3; E=0.89; R=3.39; rawP=0.0561; adjP=0.0810 | | | | | | |
| Index | UserID | Value | Gene Symbol | Gene Name | EntrezGene | Ensembl |
| 1 | P49321 | NA | NASP | nuclear autoantigenic sperm protein (histone-binding) | 4678 | ENSG00000132780 |
| 2 | Q8NC51 | NA | SERBP1 | SERPINE1 mRNA binding protein 1 | 26135 | ENSG00000142864 |
| 3 | P43490 | NA | NAMPT | nicotinamide phosphoribosyltransferase | 10135 | ENSG00000105835 |

  
  

| **Database:Transcrription Target      &nbspName:hsa\_CACGTG\_V$MYC\_Q2      &nbspID:DB\_ID:2434** | | | | | | |
| --- | --- | --- | --- | --- | --- | --- |
| C=1008; O=3; E=0.94; R=3.18; rawP=0.0655; adjP=0.0852 | | | | | | |
| Index | UserID | Value | Gene Symbol | Gene Name | EntrezGene | Ensembl |
| 1 | O14980 | NA | XPO1 | exportin 1 (CRM1 homolog, yeast) | 7514 | ENSG00000082898 |
| 2 | Q9Y2X3 | NA | NOP58 | NOP58 ribonucleoprotein homolog (yeast) | 51602 | ENSG00000055044 |
| 3 | Q8NC51 | NA | SERBP1 | SERPINE1 mRNA binding protein 1 | 26135 | ENSG00000142864 |

  
  

| **Database:Transcrription Target      &nbspName:hsa\_TGANTCA\_V$AP1\_C      &nbspID:DB\_ID:2402** | | | | | | |
| --- | --- | --- | --- | --- | --- | --- |
| C=1092; O=3; E=1.02; R=2.93; rawP=0.0793; adjP=0.0937 | | | | | | |
| Index | UserID | Value | Gene Symbol | Gene Name | EntrezGene | Ensembl |
| 1 | P08727 | NA | KRT19 | keratin 19 | 3880 | ENSG00000171345 |
| 2 | P49321 | NA | NASP | nuclear autoantigenic sperm protein (histone-binding) | 4678 | ENSG00000132780 |
| 3 | P05787 | NA | KRT8 | keratin 8 | 3856 | ENSG00000170421 |

  
  
  
  

---

WebGestalt is currently developed and maintained by Jing Wang and Bing Zhang at the  Zhang Lab. Other people who have made significant contribution to the project include Dexter Duncan, Stefan Kirov, Zhiao Shi, and Jay Snoddy.  
  
**Funding credits:** NIH/NIAAA (U01 AA016662, U01 AA013512); NIH/NIDA (P01 DA015027); NIH/NIMH (P50 MH078028, P50 MH096972); NIH/NCI (U24 CA159988); NIH/NIGMS (R01 GM088822).
