## Supplementary material for "PGRMC1 phosphorylation and cell plasticity 1: glycolysis, mitochondria, tumor growth": File S8: final_sig_file_1439166598.html

Anchored HTML File of EIDs

|  |  |
| --- | --- |
|  | WEB-based GEne SeT AnaLysis Toolkit |
|  | ***Translating gene lists into biological insights...*** |

---

  

| **Database:biological process      &nbspName:cell death      &nbspID:GO:0008219** | | | | | | |
| --- | --- | --- | --- | --- | --- | --- |
| C=1697; O=7; E=1.29; R=5.41; rawP=6.62e-05; adjP=0.0036 | | | | | | |
| Index | UserID | Value | Gene Symbol | Gene Name | EntrezGene | Ensembl |
| 1 | P02792 | NA | FTL | ferritin, light polypeptide | 2512 | ENSG00000087086 |
| 2 | P07339 | NA | CTSD | cathepsin D | 1509 | ENSG00000117984 |
| 3 | P04080 | NA | CSTB | cystatin B (stefin B) | 1476 | ENSG00000160213 |
| 4 | P02545 | NA | LMNA | lamin A/C | 4000 | ENSG00000160789 |
| 5 | P47895 | NA | ALDH1A3 | aldehyde dehydrogenase 1 family, member A3 | 220 | ENSG00000184254 |
| 6 | P04083 | NA | ANXA1 | annexin A1 | 301 | ENSG00000135046 |
| 7 | P05556 | NA | ITGB1 | integrin, beta 1 (fibronectin receptor, beta polypeptide, antigen CD29 includes MDF2, MSK12) | 3688 | ENSG00000150093 |

  
  

| **Database:biological process      &nbspName:death      &nbspID:GO:0016265** | | | | | | |
| --- | --- | --- | --- | --- | --- | --- |
| C=1699; O=7; E=1.30; R=5.40; rawP=6.67e-05; adjP=0.0036 | | | | | | |
| Index | UserID | Value | Gene Symbol | Gene Name | EntrezGene | Ensembl |
| 1 | P02792 | NA | FTL | ferritin, light polypeptide | 2512 | ENSG00000087086 |
| 2 | P07339 | NA | CTSD | cathepsin D | 1509 | ENSG00000117984 |
| 3 | P04080 | NA | CSTB | cystatin B (stefin B) | 1476 | ENSG00000160213 |
| 4 | P02545 | NA | LMNA | lamin A/C | 4000 | ENSG00000160789 |
| 5 | P47895 | NA | ALDH1A3 | aldehyde dehydrogenase 1 family, member A3 | 220 | ENSG00000184254 |
| 6 | P04083 | NA | ANXA1 | annexin A1 | 301 | ENSG00000135046 |
| 7 | P05556 | NA | ITGB1 | integrin, beta 1 (fibronectin receptor, beta polypeptide, antigen CD29 includes MDF2, MSK12) | 3688 | ENSG00000150093 |

  
  

| **Database:biological process      &nbspName:membrane organization      &nbspID:GO:0061024** | | | | | | |
| --- | --- | --- | --- | --- | --- | --- |
| C=413; O=4; E=0.32; R=12.70; rawP=0.0002; adjP=0.0054 | | | | | | |
| Index | UserID | Value | Gene Symbol | Gene Name | EntrezGene | Ensembl |
| 1 | P02792 | NA | FTL | ferritin, light polypeptide | 2512 | ENSG00000087086 |
| 2 | P02545 | NA | LMNA | lamin A/C | 4000 | ENSG00000160789 |
| 3 | P04083 | NA | ANXA1 | annexin A1 | 301 | ENSG00000135046 |
| 4 | P05556 | NA | ITGB1 | integrin, beta 1 (fibronectin receptor, beta polypeptide, antigen CD29 includes MDF2, MSK12) | 3688 | ENSG00000150093 |

  
  

| **Database:biological process      &nbspName:cellular membrane organization      &nbspID:GO:0016044** | | | | | | |
| --- | --- | --- | --- | --- | --- | --- |
| C=409; O=4; E=0.31; R=12.82; rawP=0.0002; adjP=0.0054 | | | | | | |
| Index | UserID | Value | Gene Symbol | Gene Name | EntrezGene | Ensembl |
| 1 | P02792 | NA | FTL | ferritin, light polypeptide | 2512 | ENSG00000087086 |
| 2 | P02545 | NA | LMNA | lamin A/C | 4000 | ENSG00000160789 |
| 3 | P04083 | NA | ANXA1 | annexin A1 | 301 | ENSG00000135046 |
| 4 | P05556 | NA | ITGB1 | integrin, beta 1 (fibronectin receptor, beta polypeptide, antigen CD29 includes MDF2, MSK12) | 3688 | ENSG00000150093 |

  
  

| **Database:biological process      &nbspName:regulation of programmed cell death      &nbspID:GO:0043067** | | | | | | |
| --- | --- | --- | --- | --- | --- | --- |
| C=1164; O=5; E=0.89; R=5.63; rawP=0.0010; adjP=0.0182 | | | | | | |
| Index | UserID | Value | Gene Symbol | Gene Name | EntrezGene | Ensembl |
| 1 | P04080 | NA | CSTB | cystatin B (stefin B) | 1476 | ENSG00000160213 |
| 2 | P02545 | NA | LMNA | lamin A/C | 4000 | ENSG00000160789 |
| 3 | P47895 | NA | ALDH1A3 | aldehyde dehydrogenase 1 family, member A3 | 220 | ENSG00000184254 |
| 4 | P04083 | NA | ANXA1 | annexin A1 | 301 | ENSG00000135046 |
| 5 | P05556 | NA | ITGB1 | integrin, beta 1 (fibronectin receptor, beta polypeptide, antigen CD29 includes MDF2, MSK12) | 3688 | ENSG00000150093 |

  
  

| **Database:biological process      &nbspName:regulation of apoptotic process      &nbspID:GO:0042981** | | | | | | |
| --- | --- | --- | --- | --- | --- | --- |
| C=1150; O=5; E=0.88; R=5.70; rawP=0.0010; adjP=0.0182 | | | | | | |
| Index | UserID | Value | Gene Symbol | Gene Name | EntrezGene | Ensembl |
| 1 | P04080 | NA | CSTB | cystatin B (stefin B) | 1476 | ENSG00000160213 |
| 2 | P02545 | NA | LMNA | lamin A/C | 4000 | ENSG00000160789 |
| 3 | P47895 | NA | ALDH1A3 | aldehyde dehydrogenase 1 family, member A3 | 220 | ENSG00000184254 |
| 4 | P04083 | NA | ANXA1 | annexin A1 | 301 | ENSG00000135046 |
| 5 | P05556 | NA | ITGB1 | integrin, beta 1 (fibronectin receptor, beta polypeptide, antigen CD29 includes MDF2, MSK12) | 3688 | ENSG00000150093 |

  
  

| **Database:biological process      &nbspName:regulation of cell death      &nbspID:GO:0010941** | | | | | | |
| --- | --- | --- | --- | --- | --- | --- |
| C=1192; O=5; E=0.91; R=5.50; rawP=0.0012; adjP=0.0187 | | | | | | |
| Index | UserID | Value | Gene Symbol | Gene Name | EntrezGene | Ensembl |
| 1 | P04080 | NA | CSTB | cystatin B (stefin B) | 1476 | ENSG00000160213 |
| 2 | P02545 | NA | LMNA | lamin A/C | 4000 | ENSG00000160789 |
| 3 | P47895 | NA | ALDH1A3 | aldehyde dehydrogenase 1 family, member A3 | 220 | ENSG00000184254 |
| 4 | P04083 | NA | ANXA1 | annexin A1 | 301 | ENSG00000135046 |
| 5 | P05556 | NA | ITGB1 | integrin, beta 1 (fibronectin receptor, beta polypeptide, antigen CD29 includes MDF2, MSK12) | 3688 | ENSG00000150093 |

  
  

| **Database:biological process      &nbspName:response to drug      &nbspID:GO:0042493** | | | | | | |
| --- | --- | --- | --- | --- | --- | --- |
| C=371; O=3; E=0.28; R=10.60; rawP=0.0024; adjP=0.0262 | | | | | | |
| Index | UserID | Value | Gene Symbol | Gene Name | EntrezGene | Ensembl |
| 1 | P47895 | NA | ALDH1A3 | aldehyde dehydrogenase 1 family, member A3 | 220 | ENSG00000184254 |
| 2 | P04083 | NA | ANXA1 | annexin A1 | 301 | ENSG00000135046 |
| 3 | P05556 | NA | ITGB1 | integrin, beta 1 (fibronectin receptor, beta polypeptide, antigen CD29 includes MDF2, MSK12) | 3688 | ENSG00000150093 |

  
  

| **Database:biological process      &nbspName:regulation of hormone levels      &nbspID:GO:0010817** | | | | | | |
| --- | --- | --- | --- | --- | --- | --- |
| C=378; O=3; E=0.29; R=10.41; rawP=0.0025; adjP=0.0262 | | | | | | |
| Index | UserID | Value | Gene Symbol | Gene Name | EntrezGene | Ensembl |
| 1 | Q04828 | NA | AKR1C1 | aldo-keto reductase family 1, member C1 (dihydrodiol dehydrogenase 1; 20-alpha (3-alpha)-hydroxysteroid dehydrogenase) | 1645 | ENSG00000187134 |
| 2 | P47895 | NA | ALDH1A3 | aldehyde dehydrogenase 1 family, member A3 | 220 | ENSG00000184254 |
| 3 | P04083 | NA | ANXA1 | annexin A1 | 301 | ENSG00000135046 |

  
  

| **Database:biological process      &nbspName:programmed cell death      &nbspID:GO:0012501** | | | | | | |
| --- | --- | --- | --- | --- | --- | --- |
| C=1538; O=5; E=1.17; R=4.26; rawP=0.0036; adjP=0.0262 | | | | | | |
| Index | UserID | Value | Gene Symbol | Gene Name | EntrezGene | Ensembl |
| 1 | P04080 | NA | CSTB | cystatin B (stefin B) | 1476 | ENSG00000160213 |
| 2 | P02545 | NA | LMNA | lamin A/C | 4000 | ENSG00000160789 |
| 3 | P47895 | NA | ALDH1A3 | aldehyde dehydrogenase 1 family, member A3 | 220 | ENSG00000184254 |
| 4 | P04083 | NA | ANXA1 | annexin A1 | 301 | ENSG00000135046 |
| 5 | P05556 | NA | ITGB1 | integrin, beta 1 (fibronectin receptor, beta polypeptide, antigen CD29 includes MDF2, MSK12) | 3688 | ENSG00000150093 |

  
  

| **Database:biological process      &nbspName:apoptotic process      &nbspID:GO:0006915** | | | | | | |
| --- | --- | --- | --- | --- | --- | --- |
| C=1521; O=5; E=1.16; R=4.31; rawP=0.0035; adjP=0.0262 | | | | | | |
| Index | UserID | Value | Gene Symbol | Gene Name | EntrezGene | Ensembl |
| 1 | P04080 | NA | CSTB | cystatin B (stefin B) | 1476 | ENSG00000160213 |
| 2 | P02545 | NA | LMNA | lamin A/C | 4000 | ENSG00000160789 |
| 3 | P47895 | NA | ALDH1A3 | aldehyde dehydrogenase 1 family, member A3 | 220 | ENSG00000184254 |
| 4 | P04083 | NA | ANXA1 | annexin A1 | 301 | ENSG00000135046 |
| 5 | P05556 | NA | ITGB1 | integrin, beta 1 (fibronectin receptor, beta polypeptide, antigen CD29 includes MDF2, MSK12) | 3688 | ENSG00000150093 |

  
  

| **Database:biological process      &nbspName:positive regulation of apoptotic process      &nbspID:GO:0043065** | | | | | | |
| --- | --- | --- | --- | --- | --- | --- |
| C=421; O=3; E=0.32; R=9.34; rawP=0.0034; adjP=0.0262 | | | | | | |
| Index | UserID | Value | Gene Symbol | Gene Name | EntrezGene | Ensembl |
| 1 | P47895 | NA | ALDH1A3 | aldehyde dehydrogenase 1 family, member A3 | 220 | ENSG00000184254 |
| 2 | P04083 | NA | ANXA1 | annexin A1 | 301 | ENSG00000135046 |
| 3 | P05556 | NA | ITGB1 | integrin, beta 1 (fibronectin receptor, beta polypeptide, antigen CD29 includes MDF2, MSK12) | 3688 | ENSG00000150093 |

  
  

| **Database:biological process      &nbspName:positive regulation of programmed cell death      &nbspID:GO:0043068** | | | | | | |
| --- | --- | --- | --- | --- | --- | --- |
| C=427; O=3; E=0.33; R=9.21; rawP=0.0036; adjP=0.0262 | | | | | | |
| Index | UserID | Value | Gene Symbol | Gene Name | EntrezGene | Ensembl |
| 1 | P47895 | NA | ALDH1A3 | aldehyde dehydrogenase 1 family, member A3 | 220 | ENSG00000184254 |
| 2 | P04083 | NA | ANXA1 | annexin A1 | 301 | ENSG00000135046 |
| 3 | P05556 | NA | ITGB1 | integrin, beta 1 (fibronectin receptor, beta polypeptide, antigen CD29 includes MDF2, MSK12) | 3688 | ENSG00000150093 |

  
  

| **Database:biological process      &nbspName:tissue development      &nbspID:GO:0009888** | | | | | | |
| --- | --- | --- | --- | --- | --- | --- |
| C=1448; O=5; E=1.10; R=4.53; rawP=0.0028; adjP=0.0262 | | | | | | |
| Index | UserID | Value | Gene Symbol | Gene Name | EntrezGene | Ensembl |
| 1 | P07339 | NA | CTSD | cathepsin D | 1509 | ENSG00000117984 |
| 2 | P02545 | NA | LMNA | lamin A/C | 4000 | ENSG00000160789 |
| 3 | P47895 | NA | ALDH1A3 | aldehyde dehydrogenase 1 family, member A3 | 220 | ENSG00000184254 |
| 4 | P04083 | NA | ANXA1 | annexin A1 | 301 | ENSG00000135046 |
| 5 | P05556 | NA | ITGB1 | integrin, beta 1 (fibronectin receptor, beta polypeptide, antigen CD29 includes MDF2, MSK12) | 3688 | ENSG00000150093 |

  
  

| **Database:biological process      &nbspName:monocarboxylic acid metabolic process      &nbspID:GO:0032787** | | | | | | |
| --- | --- | --- | --- | --- | --- | --- |
| C=405; O=3; E=0.31; R=9.71; rawP=0.0031; adjP=0.0262 | | | | | | |
| Index | UserID | Value | Gene Symbol | Gene Name | EntrezGene | Ensembl |
| 1 | Q04828 | NA | AKR1C1 | aldo-keto reductase family 1, member C1 (dihydrodiol dehydrogenase 1; 20-alpha (3-alpha)-hydroxysteroid dehydrogenase) | 1645 | ENSG00000187134 |
| 2 | P47895 | NA | ALDH1A3 | aldehyde dehydrogenase 1 family, member A3 | 220 | ENSG00000184254 |
| 3 | P04083 | NA | ANXA1 | annexin A1 | 301 | ENSG00000135046 |

  
  

| **Database:biological process      &nbspName:positive regulation of cell death      &nbspID:GO:0010942** | | | | | | |
| --- | --- | --- | --- | --- | --- | --- |
| C=442; O=3; E=0.34; R=8.90; rawP=0.0039; adjP=0.0266 | | | | | | |
| Index | UserID | Value | Gene Symbol | Gene Name | EntrezGene | Ensembl |
| 1 | P47895 | NA | ALDH1A3 | aldehyde dehydrogenase 1 family, member A3 | 220 | ENSG00000184254 |
| 2 | P04083 | NA | ANXA1 | annexin A1 | 301 | ENSG00000135046 |
| 3 | P05556 | NA | ITGB1 | integrin, beta 1 (fibronectin receptor, beta polypeptide, antigen CD29 includes MDF2, MSK12) | 3688 | ENSG00000150093 |

  
  

| **Database:biological process      &nbspName:homeostatic process      &nbspID:GO:0042592** | | | | | | |
| --- | --- | --- | --- | --- | --- | --- |
| C=1193; O=4; E=0.91; R=4.40; rawP=0.0096; adjP=0.0616 | | | | | | |
| Index | UserID | Value | Gene Symbol | Gene Name | EntrezGene | Ensembl |
| 1 | Q04828 | NA | AKR1C1 | aldo-keto reductase family 1, member C1 (dihydrodiol dehydrogenase 1; 20-alpha (3-alpha)-hydroxysteroid dehydrogenase) | 1645 | ENSG00000187134 |
| 2 | P02792 | NA | FTL | ferritin, light polypeptide | 2512 | ENSG00000087086 |
| 3 | P04083 | NA | ANXA1 | annexin A1 | 301 | ENSG00000135046 |
| 4 | P05556 | NA | ITGB1 | integrin, beta 1 (fibronectin receptor, beta polypeptide, antigen CD29 includes MDF2, MSK12) | 3688 | ENSG00000150093 |

  
  

| **Database:biological process      &nbspName:multicellular organism reproduction      &nbspID:GO:0032504** | | | | | | |
| --- | --- | --- | --- | --- | --- | --- |
| C=635; O=3; E=0.48; R=6.19; rawP=0.0108; adjP=0.0620 | | | | | | |
| Index | UserID | Value | Gene Symbol | Gene Name | EntrezGene | Ensembl |
| 1 | P02545 | NA | LMNA | lamin A/C | 4000 | ENSG00000160789 |
| 2 | P04083 | NA | ANXA1 | annexin A1 | 301 | ENSG00000135046 |
| 3 | P05556 | NA | ITGB1 | integrin, beta 1 (fibronectin receptor, beta polypeptide, antigen CD29 includes MDF2, MSK12) | 3688 | ENSG00000150093 |

  
  

| **Database:biological process      &nbspName:multicellular organismal reproductive process      &nbspID:GO:0048609** | | | | | | |
| --- | --- | --- | --- | --- | --- | --- |
| C=635; O=3; E=0.48; R=6.19; rawP=0.0108; adjP=0.0620 | | | | | | |
| Index | UserID | Value | Gene Symbol | Gene Name | EntrezGene | Ensembl |
| 1 | P02545 | NA | LMNA | lamin A/C | 4000 | ENSG00000160789 |
| 2 | P04083 | NA | ANXA1 | annexin A1 | 301 | ENSG00000135046 |
| 3 | P05556 | NA | ITGB1 | integrin, beta 1 (fibronectin receptor, beta polypeptide, antigen CD29 includes MDF2, MSK12) | 3688 | ENSG00000150093 |

  
  

| **Database:biological process      &nbspName:response to hormone stimulus      &nbspID:GO:0009725** | | | | | | |
| --- | --- | --- | --- | --- | --- | --- |
| C=723; O=3; E=0.55; R=5.44; rawP=0.0153; adjP=0.0834 | | | | | | |
| Index | UserID | Value | Gene Symbol | Gene Name | EntrezGene | Ensembl |
| 1 | Q04828 | NA | AKR1C1 | aldo-keto reductase family 1, member C1 (dihydrodiol dehydrogenase 1; 20-alpha (3-alpha)-hydroxysteroid dehydrogenase) | 1645 | ENSG00000187134 |
| 2 | P04083 | NA | ANXA1 | annexin A1 | 301 | ENSG00000135046 |
| 3 | P05556 | NA | ITGB1 | integrin, beta 1 (fibronectin receptor, beta polypeptide, antigen CD29 includes MDF2, MSK12) | 3688 | ENSG00000150093 |

  
  

| **Database:molecular function      &nbspName:calcium ion binding      &nbspID:GO:0005509** | | | | | | |
| --- | --- | --- | --- | --- | --- | --- |
| C=665; O=3; E=0.53; R=5.63; rawP=0.0142; adjP=0.0805 | | | | | | |
| Index | UserID | Value | Gene Symbol | Gene Name | EntrezGene | Ensembl |
| 1 | P07384 | NA | CAPN1 | calpain 1, (mu/I) large subunit | 823 | ENSG00000014216 |
| 2 | P04083 | NA | ANXA1 | annexin A1 | 301 | ENSG00000135046 |
| 3 | P13797 | NA | PLS3 | plastin 3 | 5358 | ENSG00000102024 |

  
  

| **Database:molecular function      &nbspName:cytoskeletal protein binding      &nbspID:GO:0008092** | | | | | | |
| --- | --- | --- | --- | --- | --- | --- |
| C=634; O=3; E=0.51; R=5.91; rawP=0.0125; adjP=0.0805 | | | | | | |
| Index | UserID | Value | Gene Symbol | Gene Name | EntrezGene | Ensembl |
| 1 | P07384 | NA | CAPN1 | calpain 1, (mu/I) large subunit | 823 | ENSG00000014216 |
| 2 | P13797 | NA | PLS3 | plastin 3 | 5358 | ENSG00000102024 |
| 3 | P05556 | NA | ITGB1 | integrin, beta 1 (fibronectin receptor, beta polypeptide, antigen CD29 includes MDF2, MSK12) | 3688 | ENSG00000150093 |

  
  

| **Database:molecular function      &nbspName:oxidoreductase activity      &nbspID:GO:0016491** | | | | | | |
| --- | --- | --- | --- | --- | --- | --- |
| C=697; O=3; E=0.56; R=5.37; rawP=0.0161; adjP=0.0805 | | | | | | |
| Index | UserID | Value | Gene Symbol | Gene Name | EntrezGene | Ensembl |
| 1 | Q04828 | NA | AKR1C1 | aldo-keto reductase family 1, member C1 (dihydrodiol dehydrogenase 1; 20-alpha (3-alpha)-hydroxysteroid dehydrogenase) | 1645 | ENSG00000187134 |
| 2 | P02792 | NA | FTL | ferritin, light polypeptide | 2512 | ENSG00000087086 |
| 3 | P47895 | NA | ALDH1A3 | aldehyde dehydrogenase 1 family, member A3 | 220 | ENSG00000184254 |

  
  
  
  

---

WebGestalt is currently developed and maintained by Jing Wang and Bing Zhang at the  Zhang Lab. Other people who have made significant contribution to the project include Dexter Duncan, Stefan Kirov, Zhiao Shi, and Jay Snoddy.  
  
**Funding credits:** NIH/NIAAA (U01 AA016662, U01 AA013512); NIH/NIDA (P01 DA015027); NIH/NIMH (P50 MH078028, P50 MH096972); NIH/NCI (U24 CA159988); NIH/NIGMS (R01 GM088822).
