## Supplementary material for "PGRMC1 phosphorylation and cell plasticity 1: glycolysis, mitochondria, tumor growth": File S8: final_sig_file_1439166893.html

Anchored HTML File of EIDs

|  |  |
| --- | --- |
|  | WEB-based GEne SeT AnaLysis Toolkit |
|  | ***Translating gene lists into biological insights...*** |

---

  

| **Database:biological process      &nbspName:small molecule metabolic process      &nbspID:GO:0044281** | | | | | | |
| --- | --- | --- | --- | --- | --- | --- |
| C=2500; O=24; E=7.45; R=3.22; rawP=1.30e-08; adjP=3.67e-06 | | | | | | |
| Index | UserID | Value | Gene Symbol | Gene Name | EntrezGene | Ensembl |
| 1 | P54819 | NA | AK2 | adenylate kinase 2 | 204 | ENSG00000004455 |
| 2 | Q9UL46 | NA | PSME2 | proteasome (prosome, macropain) activator subunit 2 (PA28 beta) | 5721 | ENSG00000100911 |
| 3 | Q99714 | NA | HSD17B10 | hydroxysteroid (17-beta) dehydrogenase 10 | 3028 | ENSG00000072506 |
| 4 | P04179 | NA | SOD2 | superoxide dismutase 2, mitochondrial | 6648 | ENSG00000112096 |
| 5 | P06737 | NA | PYGL | phosphorylase, glycogen, liver | 5836 | ENSG00000100504 |
| 6 | P07741 | NA | APRT | adenine phosphoribosyltransferase | 353 | ENSG00000198931 |
| 7 | Q06323 | NA | PSME1 | proteasome (prosome, macropain) activator subunit 1 (PA28 alpha) | 5720 | ENSG00000092010 |
| 8 | P30837 | NA | ALDH1B1 | aldehyde dehydrogenase 1 family, member B1 | 219 | ENSG00000137124 |
| 9 | P38571 | NA | LIPA | lipase A, lysosomal acid, cholesterol esterase | 3988 | ENSG00000107798 |
| 10 | P31040 | NA | SDHA | succinate dehydrogenase complex, subunit A, flavoprotein (Fp) | 6389 | ENSG00000073578 |
| 11 | Q13011 | NA | ECH1 | enoyl CoA hydratase 1, peroxisomal | 1891 | ENSG00000104823 |
| 12 | P38117 | NA | ETFB | electron-transfer-flavoprotein, beta polypeptide | 2109 | ENSG00000105379 |
| 13 | Q99798 | NA | ACO2 | aconitase 2, mitochondrial | 50 | ENSG00000100412 |
| 14 | Q16698 | NA | DECR1 | 2,4-dienoyl CoA reductase 1, mitochondrial | 1666 | ENSG00000104325 |
| 15 | Q9UBQ7 | NA | GRHPR | glyoxylate reductase/hydroxypyruvate reductase | 9380 | ENSG00000137106 |
| 16 | P00558 | NA | PGK1 | phosphoglycerate kinase 1 | 5230 | ENSG00000102144 |
| 17 | P42765 | NA | ACAA2 | acetyl-CoA acyltransferase 2 | 10449 | ENSG00000167315 |
| 18 | P13674 | NA | P4HA1 | prolyl 4-hydroxylase, alpha polypeptide I | 5033 | ENSG00000122884 |
| 19 | O43598 | NA | C6orf108 | chromosome 6 open reading frame 108 | 10591 | ENSG00000112667 |
| 20 | P49419 | NA | ALDH7A1 | aldehyde dehydrogenase 7 family, member A1 | 501 | ENSG00000164904 |
| 21 | P06744 | NA | GPI | glucose-6-phosphate isomerase | 2821 | ENSG00000105220 |
| 22 | P19367 | NA | HK1 | hexokinase 1 | 3098 | ENSG00000156515 |
| 23 | Q9BQ69 | NA | MACROD1 | MACRO domain containing 1 | 28992 | ENSG00000133315 |
| 24 | P00352 | NA | ALDH1A1 | aldehyde dehydrogenase 1 family, member A1 | 216 | ENSG00000165092 |

  
  

| **Database:biological process      &nbspName:catabolic process      &nbspID:GO:0009056** | | | | | | |
| --- | --- | --- | --- | --- | --- | --- |
| C=1991; O=21; E=5.94; R=3.54; rawP=3.90e-08; adjP=5.50e-06 | | | | | | |
| Index | UserID | Value | Gene Symbol | Gene Name | EntrezGene | Ensembl |
| 1 | P10253 | NA | GAA | glucosidase, alpha; acid | 2548 | ENSG00000171298 |
| 2 | Q9UL46 | NA | PSME2 | proteasome (prosome, macropain) activator subunit 2 (PA28 beta) | 5721 | ENSG00000100911 |
| 3 | Q99714 | NA | HSD17B10 | hydroxysteroid (17-beta) dehydrogenase 10 | 3028 | ENSG00000072506 |
| 4 | P06737 | NA | PYGL | phosphorylase, glycogen, liver | 5836 | ENSG00000100504 |
| 5 | Q06323 | NA | PSME1 | proteasome (prosome, macropain) activator subunit 1 (PA28 alpha) | 5720 | ENSG00000092010 |
| 6 | P30837 | NA | ALDH1B1 | aldehyde dehydrogenase 1 family, member B1 | 219 | ENSG00000137124 |
| 7 | P38571 | NA | LIPA | lipase A, lysosomal acid, cholesterol esterase | 3988 | ENSG00000107798 |
| 8 | P32119 | NA | PRDX2 | peroxiredoxin 2 | 7001 | ENSG00000167815 |
| 9 | P31040 | NA | SDHA | succinate dehydrogenase complex, subunit A, flavoprotein (Fp) | 6389 | ENSG00000073578 |
| 10 | Q13011 | NA | ECH1 | enoyl CoA hydratase 1, peroxisomal | 1891 | ENSG00000104823 |
| 11 | Q16698 | NA | DECR1 | 2,4-dienoyl CoA reductase 1, mitochondrial | 1666 | ENSG00000104325 |
| 12 | Q99798 | NA | ACO2 | aconitase 2, mitochondrial | 50 | ENSG00000100412 |
| 13 | P00558 | NA | PGK1 | phosphoglycerate kinase 1 | 5230 | ENSG00000102144 |
| 14 | P54727 | NA | RAD23B | RAD23 homolog B (S. cerevisiae) | 5887 | ENSG00000119318 |
| 15 | P42765 | NA | ACAA2 | acetyl-CoA acyltransferase 2 | 10449 | ENSG00000167315 |
| 16 | P49419 | NA | ALDH7A1 | aldehyde dehydrogenase 7 family, member A1 | 501 | ENSG00000164904 |
| 17 | O43598 | NA | C6orf108 | chromosome 6 open reading frame 108 | 10591 | ENSG00000112667 |
| 18 | P06744 | NA | GPI | glucose-6-phosphate isomerase | 2821 | ENSG00000105220 |
| 19 | P19367 | NA | HK1 | hexokinase 1 | 3098 | ENSG00000156515 |
| 20 | P00352 | NA | ALDH1A1 | aldehyde dehydrogenase 1 family, member A1 | 216 | ENSG00000165092 |
| 21 | Q16531 | NA | DDB1 | damage-specific DNA binding protein 1, 127kDa | 1642 | ENSG00000167986 |

  
  

| **Database:biological process      &nbspName:organic substance catabolic process      &nbspID:GO:1901575** | | | | | | |
| --- | --- | --- | --- | --- | --- | --- |
| C=1851; O=20; E=5.52; R=3.62; rawP=6.70e-08; adjP=6.30e-06 | | | | | | |
| Index | UserID | Value | Gene Symbol | Gene Name | EntrezGene | Ensembl |
| 1 | P10253 | NA | GAA | glucosidase, alpha; acid | 2548 | ENSG00000171298 |
| 2 | Q9UL46 | NA | PSME2 | proteasome (prosome, macropain) activator subunit 2 (PA28 beta) | 5721 | ENSG00000100911 |
| 3 | Q99714 | NA | HSD17B10 | hydroxysteroid (17-beta) dehydrogenase 10 | 3028 | ENSG00000072506 |
| 4 | P06737 | NA | PYGL | phosphorylase, glycogen, liver | 5836 | ENSG00000100504 |
| 5 | Q06323 | NA | PSME1 | proteasome (prosome, macropain) activator subunit 1 (PA28 alpha) | 5720 | ENSG00000092010 |
| 6 | P30837 | NA | ALDH1B1 | aldehyde dehydrogenase 1 family, member B1 | 219 | ENSG00000137124 |
| 7 | P38571 | NA | LIPA | lipase A, lysosomal acid, cholesterol esterase | 3988 | ENSG00000107798 |
| 8 | P31040 | NA | SDHA | succinate dehydrogenase complex, subunit A, flavoprotein (Fp) | 6389 | ENSG00000073578 |
| 9 | Q13011 | NA | ECH1 | enoyl CoA hydratase 1, peroxisomal | 1891 | ENSG00000104823 |
| 10 | Q16698 | NA | DECR1 | 2,4-dienoyl CoA reductase 1, mitochondrial | 1666 | ENSG00000104325 |
| 11 | Q99798 | NA | ACO2 | aconitase 2, mitochondrial | 50 | ENSG00000100412 |
| 12 | P00558 | NA | PGK1 | phosphoglycerate kinase 1 | 5230 | ENSG00000102144 |
| 13 | P54727 | NA | RAD23B | RAD23 homolog B (S. cerevisiae) | 5887 | ENSG00000119318 |
| 14 | P42765 | NA | ACAA2 | acetyl-CoA acyltransferase 2 | 10449 | ENSG00000167315 |
| 15 | P49419 | NA | ALDH7A1 | aldehyde dehydrogenase 7 family, member A1 | 501 | ENSG00000164904 |
| 16 | O43598 | NA | C6orf108 | chromosome 6 open reading frame 108 | 10591 | ENSG00000112667 |
| 17 | P06744 | NA | GPI | glucose-6-phosphate isomerase | 2821 | ENSG00000105220 |
| 18 | P19367 | NA | HK1 | hexokinase 1 | 3098 | ENSG00000156515 |
| 19 | P00352 | NA | ALDH1A1 | aldehyde dehydrogenase 1 family, member A1 | 216 | ENSG00000165092 |
| 20 | Q16531 | NA | DDB1 | damage-specific DNA binding protein 1, 127kDa | 1642 | ENSG00000167986 |

  
  

| **Database:biological process      &nbspName:oxidation-reduction process      &nbspID:GO:0055114** | | | | | | |
| --- | --- | --- | --- | --- | --- | --- |
| C=554; O=11; E=1.65; R=6.66; rawP=4.58e-07; adjP=3.23e-05 | | | | | | |
| Index | UserID | Value | Gene Symbol | Gene Name | EntrezGene | Ensembl |
| 1 | Q99798 | NA | ACO2 | aconitase 2, mitochondrial | 50 | ENSG00000100412 |
| 2 | Q9UBQ7 | NA | GRHPR | glyoxylate reductase/hydroxypyruvate reductase | 9380 | ENSG00000137106 |
| 3 | Q16698 | NA | DECR1 | 2,4-dienoyl CoA reductase 1, mitochondrial | 1666 | ENSG00000104325 |
| 4 | P10253 | NA | GAA | glucosidase, alpha; acid | 2548 | ENSG00000171298 |
| 5 | P42765 | NA | ACAA2 | acetyl-CoA acyltransferase 2 | 10449 | ENSG00000167315 |
| 6 | P04179 | NA | SOD2 | superoxide dismutase 2, mitochondrial | 6648 | ENSG00000112096 |
| 7 | P06737 | NA | PYGL | phosphorylase, glycogen, liver | 5836 | ENSG00000100504 |
| 8 | P00352 | NA | ALDH1A1 | aldehyde dehydrogenase 1 family, member A1 | 216 | ENSG00000165092 |
| 9 | P31040 | NA | SDHA | succinate dehydrogenase complex, subunit A, flavoprotein (Fp) | 6389 | ENSG00000073578 |
| 10 | P38117 | NA | ETFB | electron-transfer-flavoprotein, beta polypeptide | 2109 | ENSG00000105379 |
| 11 | Q13011 | NA | ECH1 | enoyl CoA hydratase 1, peroxisomal | 1891 | ENSG00000104823 |

  
  

| **Database:biological process      &nbspName:carboxylic acid metabolic process      &nbspID:GO:0019752** | | | | | | |
| --- | --- | --- | --- | --- | --- | --- |
| C=842; O=13; E=2.51; R=5.18; rawP=5.94e-07; adjP=3.35e-05 | | | | | | |
| Index | UserID | Value | Gene Symbol | Gene Name | EntrezGene | Ensembl |
| 1 | Q99798 | NA | ACO2 | aconitase 2, mitochondrial | 50 | ENSG00000100412 |
| 2 | Q9UBQ7 | NA | GRHPR | glyoxylate reductase/hydroxypyruvate reductase | 9380 | ENSG00000137106 |
| 3 | Q16698 | NA | DECR1 | 2,4-dienoyl CoA reductase 1, mitochondrial | 1666 | ENSG00000104325 |
| 4 | Q9UL46 | NA | PSME2 | proteasome (prosome, macropain) activator subunit 2 (PA28 beta) | 5721 | ENSG00000100911 |
| 5 | P42765 | NA | ACAA2 | acetyl-CoA acyltransferase 2 | 10449 | ENSG00000167315 |
| 6 | Q99714 | NA | HSD17B10 | hydroxysteroid (17-beta) dehydrogenase 10 | 3028 | ENSG00000072506 |
| 7 | P04179 | NA | SOD2 | superoxide dismutase 2, mitochondrial | 6648 | ENSG00000112096 |
| 8 | P13674 | NA | P4HA1 | prolyl 4-hydroxylase, alpha polypeptide I | 5033 | ENSG00000122884 |
| 9 | Q06323 | NA | PSME1 | proteasome (prosome, macropain) activator subunit 1 (PA28 alpha) | 5720 | ENSG00000092010 |
| 10 | P49419 | NA | ALDH7A1 | aldehyde dehydrogenase 7 family, member A1 | 501 | ENSG00000164904 |
| 11 | P38571 | NA | LIPA | lipase A, lysosomal acid, cholesterol esterase | 3988 | ENSG00000107798 |
| 12 | P31040 | NA | SDHA | succinate dehydrogenase complex, subunit A, flavoprotein (Fp) | 6389 | ENSG00000073578 |
| 13 | Q13011 | NA | ECH1 | enoyl CoA hydratase 1, peroxisomal | 1891 | ENSG00000104823 |

  
  

| **Database:biological process      &nbspName:organic acid metabolic process      &nbspID:GO:0006082** | | | | | | |
| --- | --- | --- | --- | --- | --- | --- |
| C=967; O=13; E=2.88; R=4.51; rawP=2.82e-06; adjP=0.0001 | | | | | | |
| Index | UserID | Value | Gene Symbol | Gene Name | EntrezGene | Ensembl |
| 1 | Q99798 | NA | ACO2 | aconitase 2, mitochondrial | 50 | ENSG00000100412 |
| 2 | Q9UBQ7 | NA | GRHPR | glyoxylate reductase/hydroxypyruvate reductase | 9380 | ENSG00000137106 |
| 3 | Q16698 | NA | DECR1 | 2,4-dienoyl CoA reductase 1, mitochondrial | 1666 | ENSG00000104325 |
| 4 | Q9UL46 | NA | PSME2 | proteasome (prosome, macropain) activator subunit 2 (PA28 beta) | 5721 | ENSG00000100911 |
| 5 | P42765 | NA | ACAA2 | acetyl-CoA acyltransferase 2 | 10449 | ENSG00000167315 |
| 6 | Q99714 | NA | HSD17B10 | hydroxysteroid (17-beta) dehydrogenase 10 | 3028 | ENSG00000072506 |
| 7 | P04179 | NA | SOD2 | superoxide dismutase 2, mitochondrial | 6648 | ENSG00000112096 |
| 8 | P13674 | NA | P4HA1 | prolyl 4-hydroxylase, alpha polypeptide I | 5033 | ENSG00000122884 |
| 9 | Q06323 | NA | PSME1 | proteasome (prosome, macropain) activator subunit 1 (PA28 alpha) | 5720 | ENSG00000092010 |
| 10 | P49419 | NA | ALDH7A1 | aldehyde dehydrogenase 7 family, member A1 | 501 | ENSG00000164904 |
| 11 | P38571 | NA | LIPA | lipase A, lysosomal acid, cholesterol esterase | 3988 | ENSG00000107798 |
| 12 | P31040 | NA | SDHA | succinate dehydrogenase complex, subunit A, flavoprotein (Fp) | 6389 | ENSG00000073578 |
| 13 | Q13011 | NA | ECH1 | enoyl CoA hydratase 1, peroxisomal | 1891 | ENSG00000104823 |

  
  

| **Database:biological process      &nbspName:oxoacid metabolic process      &nbspID:GO:0043436** | | | | | | |
| --- | --- | --- | --- | --- | --- | --- |
| C=949; O=13; E=2.83; R=4.59; rawP=2.29e-06; adjP=0.0001 | | | | | | |
| Index | UserID | Value | Gene Symbol | Gene Name | EntrezGene | Ensembl |
| 1 | Q99798 | NA | ACO2 | aconitase 2, mitochondrial | 50 | ENSG00000100412 |
| 2 | Q9UBQ7 | NA | GRHPR | glyoxylate reductase/hydroxypyruvate reductase | 9380 | ENSG00000137106 |
| 3 | Q16698 | NA | DECR1 | 2,4-dienoyl CoA reductase 1, mitochondrial | 1666 | ENSG00000104325 |
| 4 | Q9UL46 | NA | PSME2 | proteasome (prosome, macropain) activator subunit 2 (PA28 beta) | 5721 | ENSG00000100911 |
| 5 | P42765 | NA | ACAA2 | acetyl-CoA acyltransferase 2 | 10449 | ENSG00000167315 |
| 6 | Q99714 | NA | HSD17B10 | hydroxysteroid (17-beta) dehydrogenase 10 | 3028 | ENSG00000072506 |
| 7 | P04179 | NA | SOD2 | superoxide dismutase 2, mitochondrial | 6648 | ENSG00000112096 |
| 8 | P13674 | NA | P4HA1 | prolyl 4-hydroxylase, alpha polypeptide I | 5033 | ENSG00000122884 |
| 9 | Q06323 | NA | PSME1 | proteasome (prosome, macropain) activator subunit 1 (PA28 alpha) | 5720 | ENSG00000092010 |
| 10 | P49419 | NA | ALDH7A1 | aldehyde dehydrogenase 7 family, member A1 | 501 | ENSG00000164904 |
| 11 | P38571 | NA | LIPA | lipase A, lysosomal acid, cholesterol esterase | 3988 | ENSG00000107798 |
| 12 | P31040 | NA | SDHA | succinate dehydrogenase complex, subunit A, flavoprotein (Fp) | 6389 | ENSG00000073578 |
| 13 | Q13011 | NA | ECH1 | enoyl CoA hydratase 1, peroxisomal | 1891 | ENSG00000104823 |

  
  

| **Database:biological process      &nbspName:cellular aldehyde metabolic process      &nbspID:GO:0006081** | | | | | | |
| --- | --- | --- | --- | --- | --- | --- |
| C=40; O=4; E=0.12; R=33.54; rawP=5.79e-06; adjP=0.0002 | | | | | | |
| Index | UserID | Value | Gene Symbol | Gene Name | EntrezGene | Ensembl |
| 1 | Q9UBQ7 | NA | GRHPR | glyoxylate reductase/hydroxypyruvate reductase | 9380 | ENSG00000137106 |
| 2 | P49419 | NA | ALDH7A1 | aldehyde dehydrogenase 7 family, member A1 | 501 | ENSG00000164904 |
| 3 | P06744 | NA | GPI | glucose-6-phosphate isomerase | 2821 | ENSG00000105220 |
| 4 | P00352 | NA | ALDH1A1 | aldehyde dehydrogenase 1 family, member A1 | 216 | ENSG00000165092 |

  
  

| **Database:biological process      &nbspName:generation of precursor metabolites and energy      &nbspID:GO:0006091** | | | | | | |
| --- | --- | --- | --- | --- | --- | --- |
| C=447; O=9; E=1.33; R=6.75; rawP=5.38e-06; adjP=0.0002 | | | | | | |
| Index | UserID | Value | Gene Symbol | Gene Name | EntrezGene | Ensembl |
| 1 | Q99798 | NA | ACO2 | aconitase 2, mitochondrial | 50 | ENSG00000100412 |
| 2 | P10253 | NA | GAA | glucosidase, alpha; acid | 2548 | ENSG00000171298 |
| 3 | P00558 | NA | PGK1 | phosphoglycerate kinase 1 | 5230 | ENSG00000102144 |
| 4 | P04179 | NA | SOD2 | superoxide dismutase 2, mitochondrial | 6648 | ENSG00000112096 |
| 5 | P06737 | NA | PYGL | phosphorylase, glycogen, liver | 5836 | ENSG00000100504 |
| 6 | P19367 | NA | HK1 | hexokinase 1 | 3098 | ENSG00000156515 |
| 7 | P06744 | NA | GPI | glucose-6-phosphate isomerase | 2821 | ENSG00000105220 |
| 8 | P31040 | NA | SDHA | succinate dehydrogenase complex, subunit A, flavoprotein (Fp) | 6389 | ENSG00000073578 |
| 9 | P38117 | NA | ETFB | electron-transfer-flavoprotein, beta polypeptide | 2109 | ENSG00000105379 |

  
  

| **Database:biological process      &nbspName:cellular catabolic process      &nbspID:GO:0044248** | | | | | | |
| --- | --- | --- | --- | --- | --- | --- |
| C=1665; O=15; E=4.96; R=3.02; rawP=5.25e-05; adjP=0.0013 | | | | | | |
| Index | UserID | Value | Gene Symbol | Gene Name | EntrezGene | Ensembl |
| 1 | Q99798 | NA | ACO2 | aconitase 2, mitochondrial | 50 | ENSG00000100412 |
| 2 | Q16698 | NA | DECR1 | 2,4-dienoyl CoA reductase 1, mitochondrial | 1666 | ENSG00000104325 |
| 3 | Q9UL46 | NA | PSME2 | proteasome (prosome, macropain) activator subunit 2 (PA28 beta) | 5721 | ENSG00000100911 |
| 4 | P54727 | NA | RAD23B | RAD23 homolog B (S. cerevisiae) | 5887 | ENSG00000119318 |
| 5 | P42765 | NA | ACAA2 | acetyl-CoA acyltransferase 2 | 10449 | ENSG00000167315 |
| 6 | Q99714 | NA | HSD17B10 | hydroxysteroid (17-beta) dehydrogenase 10 | 3028 | ENSG00000072506 |
| 7 | O43598 | NA | C6orf108 | chromosome 6 open reading frame 108 | 10591 | ENSG00000112667 |
| 8 | Q06323 | NA | PSME1 | proteasome (prosome, macropain) activator subunit 1 (PA28 alpha) | 5720 | ENSG00000092010 |
| 9 | P49419 | NA | ALDH7A1 | aldehyde dehydrogenase 7 family, member A1 | 501 | ENSG00000164904 |
| 10 | P06744 | NA | GPI | glucose-6-phosphate isomerase | 2821 | ENSG00000105220 |
| 11 | P00352 | NA | ALDH1A1 | aldehyde dehydrogenase 1 family, member A1 | 216 | ENSG00000165092 |
| 12 | P32119 | NA | PRDX2 | peroxiredoxin 2 | 7001 | ENSG00000167815 |
| 13 | P31040 | NA | SDHA | succinate dehydrogenase complex, subunit A, flavoprotein (Fp) | 6389 | ENSG00000073578 |
| 14 | Q16531 | NA | DDB1 | damage-specific DNA binding protein 1, 127kDa | 1642 | ENSG00000167986 |
| 15 | Q13011 | NA | ECH1 | enoyl CoA hydratase 1, peroxisomal | 1891 | ENSG00000104823 |

  
  

| **Database:biological process      &nbspName:carbohydrate catabolic process      &nbspID:GO:0016052** | | | | | | |
| --- | --- | --- | --- | --- | --- | --- |
| C=140; O=5; E=0.42; R=11.98; rawP=5.74e-05; adjP=0.0013 | | | | | | |
| Index | UserID | Value | Gene Symbol | Gene Name | EntrezGene | Ensembl |
| 1 | P06737 | NA | PYGL | phosphorylase, glycogen, liver | 5836 | ENSG00000100504 |
| 2 | P10253 | NA | GAA | glucosidase, alpha; acid | 2548 | ENSG00000171298 |
| 3 | P06744 | NA | GPI | glucose-6-phosphate isomerase | 2821 | ENSG00000105220 |
| 4 | P19367 | NA | HK1 | hexokinase 1 | 3098 | ENSG00000156515 |
| 5 | P00558 | NA | PGK1 | phosphoglycerate kinase 1 | 5230 | ENSG00000102144 |

  
  

| **Database:biological process      &nbspName:single-organism carbohydrate catabolic process      &nbspID:GO:0044724** | | | | | | |
| --- | --- | --- | --- | --- | --- | --- |
| C=137; O=5; E=0.41; R=12.24; rawP=5.18e-05; adjP=0.0013 | | | | | | |
| Index | UserID | Value | Gene Symbol | Gene Name | EntrezGene | Ensembl |
| 1 | P06737 | NA | PYGL | phosphorylase, glycogen, liver | 5836 | ENSG00000100504 |
| 2 | P10253 | NA | GAA | glucosidase, alpha; acid | 2548 | ENSG00000171298 |
| 3 | P06744 | NA | GPI | glucose-6-phosphate isomerase | 2821 | ENSG00000105220 |
| 4 | P19367 | NA | HK1 | hexokinase 1 | 3098 | ENSG00000156515 |
| 5 | P00558 | NA | PGK1 | phosphoglycerate kinase 1 | 5230 | ENSG00000102144 |

  
  

| **Database:biological process      &nbspName:small molecule catabolic process      &nbspID:GO:0044282** | | | | | | |
| --- | --- | --- | --- | --- | --- | --- |
| C=258; O=6; E=0.77; R=7.80; rawP=0.0001; adjP=0.0020 | | | | | | |
| Index | UserID | Value | Gene Symbol | Gene Name | EntrezGene | Ensembl |
| 1 | P49419 | NA | ALDH7A1 | aldehyde dehydrogenase 7 family, member A1 | 501 | ENSG00000164904 |
| 2 | Q16698 | NA | DECR1 | 2,4-dienoyl CoA reductase 1, mitochondrial | 1666 | ENSG00000104325 |
| 3 | P30837 | NA | ALDH1B1 | aldehyde dehydrogenase 1 family, member B1 | 219 | ENSG00000137124 |
| 4 | Q13011 | NA | ECH1 | enoyl CoA hydratase 1, peroxisomal | 1891 | ENSG00000104823 |
| 5 | P42765 | NA | ACAA2 | acetyl-CoA acyltransferase 2 | 10449 | ENSG00000167315 |
| 6 | Q99714 | NA | HSD17B10 | hydroxysteroid (17-beta) dehydrogenase 10 | 3028 | ENSG00000072506 |

  
  

| **Database:biological process      &nbspName:single-organism catabolic process      &nbspID:GO:0044712** | | | | | | |
| --- | --- | --- | --- | --- | --- | --- |
| C=259; O=6; E=0.77; R=7.77; rawP=0.0001; adjP=0.0020 | | | | | | |
| Index | UserID | Value | Gene Symbol | Gene Name | EntrezGene | Ensembl |
| 1 | P49419 | NA | ALDH7A1 | aldehyde dehydrogenase 7 family, member A1 | 501 | ENSG00000164904 |
| 2 | Q16698 | NA | DECR1 | 2,4-dienoyl CoA reductase 1, mitochondrial | 1666 | ENSG00000104325 |
| 3 | P30837 | NA | ALDH1B1 | aldehyde dehydrogenase 1 family, member B1 | 219 | ENSG00000137124 |
| 4 | Q13011 | NA | ECH1 | enoyl CoA hydratase 1, peroxisomal | 1891 | ENSG00000104823 |
| 5 | P42765 | NA | ACAA2 | acetyl-CoA acyltransferase 2 | 10449 | ENSG00000167315 |
| 6 | Q99714 | NA | HSD17B10 | hydroxysteroid (17-beta) dehydrogenase 10 | 3028 | ENSG00000072506 |

  
  

| **Database:biological process      &nbspName:organic acid catabolic process      &nbspID:GO:0016054** | | | | | | |
| --- | --- | --- | --- | --- | --- | --- |
| C=198; O=5; E=0.59; R=8.47; rawP=0.0003; adjP=0.0053 | | | | | | |
| Index | UserID | Value | Gene Symbol | Gene Name | EntrezGene | Ensembl |
| 1 | P49419 | NA | ALDH7A1 | aldehyde dehydrogenase 7 family, member A1 | 501 | ENSG00000164904 |
| 2 | Q16698 | NA | DECR1 | 2,4-dienoyl CoA reductase 1, mitochondrial | 1666 | ENSG00000104325 |
| 3 | Q13011 | NA | ECH1 | enoyl CoA hydratase 1, peroxisomal | 1891 | ENSG00000104823 |
| 4 | P42765 | NA | ACAA2 | acetyl-CoA acyltransferase 2 | 10449 | ENSG00000167315 |
| 5 | Q99714 | NA | HSD17B10 | hydroxysteroid (17-beta) dehydrogenase 10 | 3028 | ENSG00000072506 |

  
  

| **Database:biological process      &nbspName:carboxylic acid catabolic process      &nbspID:GO:0046395** | | | | | | |
| --- | --- | --- | --- | --- | --- | --- |
| C=198; O=5; E=0.59; R=8.47; rawP=0.0003; adjP=0.0053 | | | | | | |
| Index | UserID | Value | Gene Symbol | Gene Name | EntrezGene | Ensembl |
| 1 | P49419 | NA | ALDH7A1 | aldehyde dehydrogenase 7 family, member A1 | 501 | ENSG00000164904 |
| 2 | Q16698 | NA | DECR1 | 2,4-dienoyl CoA reductase 1, mitochondrial | 1666 | ENSG00000104325 |
| 3 | Q13011 | NA | ECH1 | enoyl CoA hydratase 1, peroxisomal | 1891 | ENSG00000104823 |
| 4 | P42765 | NA | ACAA2 | acetyl-CoA acyltransferase 2 | 10449 | ENSG00000167315 |
| 5 | Q99714 | NA | HSD17B10 | hydroxysteroid (17-beta) dehydrogenase 10 | 3028 | ENSG00000072506 |

  
  

| **Database:biological process      &nbspName:energy derivation by oxidation of organic compounds      &nbspID:GO:0015980** | | | | | | |
| --- | --- | --- | --- | --- | --- | --- |
| C=326; O=6; E=0.97; R=6.17; rawP=0.0004; adjP=0.0063 | | | | | | |
| Index | UserID | Value | Gene Symbol | Gene Name | EntrezGene | Ensembl |
| 1 | P06737 | NA | PYGL | phosphorylase, glycogen, liver | 5836 | ENSG00000100504 |
| 2 | Q99798 | NA | ACO2 | aconitase 2, mitochondrial | 50 | ENSG00000100412 |
| 3 | P10253 | NA | GAA | glucosidase, alpha; acid | 2548 | ENSG00000171298 |
| 4 | P31040 | NA | SDHA | succinate dehydrogenase complex, subunit A, flavoprotein (Fp) | 6389 | ENSG00000073578 |
| 5 | P38117 | NA | ETFB | electron-transfer-flavoprotein, beta polypeptide | 2109 | ENSG00000105379 |
| 6 | P04179 | NA | SOD2 | superoxide dismutase 2, mitochondrial | 6648 | ENSG00000112096 |

  
  

| **Database:biological process      &nbspName:glucose metabolic process      &nbspID:GO:0006006** | | | | | | |
| --- | --- | --- | --- | --- | --- | --- |
| C=213; O=5; E=0.64; R=7.87; rawP=0.0004; adjP=0.0063 | | | | | | |
| Index | UserID | Value | Gene Symbol | Gene Name | EntrezGene | Ensembl |
| 1 | P06737 | NA | PYGL | phosphorylase, glycogen, liver | 5836 | ENSG00000100504 |
| 2 | P10253 | NA | GAA | glucosidase, alpha; acid | 2548 | ENSG00000171298 |
| 3 | P06744 | NA | GPI | glucose-6-phosphate isomerase | 2821 | ENSG00000105220 |
| 4 | P19367 | NA | HK1 | hexokinase 1 | 3098 | ENSG00000156515 |
| 5 | P00558 | NA | PGK1 | phosphoglycerate kinase 1 | 5230 | ENSG00000102144 |

  
  

| **Database:biological process      &nbspName:cellular metabolic process      &nbspID:GO:0044237** | | | | | | |
| --- | --- | --- | --- | --- | --- | --- |
| C=8514; O=36; E=25.38; R=1.42; rawP=0.0005; adjP=0.0064 | | | | | | |
| Index | UserID | Value | Gene Symbol | Gene Name | EntrezGene | Ensembl |
| 1 | P09661 | NA | SNRPA1 | small nuclear ribonucleoprotein polypeptide A' | 6627 | ENSG00000131876 |
| 2 | P10253 | NA | GAA | glucosidase, alpha; acid | 2548 | ENSG00000171298 |
| 3 | P54819 | NA | AK2 | adenylate kinase 2 | 204 | ENSG00000004455 |
| 4 | P17931 | NA | LGALS3 | lectin, galactoside-binding, soluble, 3 | 3958 | ENSG00000131981 |
| 5 | Q9UL46 | NA | PSME2 | proteasome (prosome, macropain) activator subunit 2 (PA28 beta) | 5721 | ENSG00000100911 |
| 6 | Q99714 | NA | HSD17B10 | hydroxysteroid (17-beta) dehydrogenase 10 | 3028 | ENSG00000072506 |
| 7 | P04179 | NA | SOD2 | superoxide dismutase 2, mitochondrial | 6648 | ENSG00000112096 |
| 8 | Q06323 | NA | PSME1 | proteasome (prosome, macropain) activator subunit 1 (PA28 alpha) | 5720 | ENSG00000092010 |
| 9 | P07741 | NA | APRT | adenine phosphoribosyltransferase | 353 | ENSG00000198931 |
| 10 | P30046 | NA | DDT | D-dopachrome tautomerase | 1652 | ENSG00000099977 |
| 11 | P31040 | NA | SDHA | succinate dehydrogenase complex, subunit A, flavoprotein (Fp) | 6389 | ENSG00000073578 |
| 12 | P38117 | NA | ETFB | electron-transfer-flavoprotein, beta polypeptide | 2109 | ENSG00000105379 |
| 13 | Q16698 | NA | DECR1 | 2,4-dienoyl CoA reductase 1, mitochondrial | 1666 | ENSG00000104325 |
| 14 | P54727 | NA | RAD23B | RAD23 homolog B (S. cerevisiae) | 5887 | ENSG00000119318 |
| 15 | P42765 | NA | ACAA2 | acetyl-CoA acyltransferase 2 | 10449 | ENSG00000167315 |
| 16 | Q10713 | NA | PMPCA | peptidase (mitochondrial processing) alpha | 23203 | ENSG00000165688 |
| 17 | P39687 | NA | ANP32A | acidic (leucine-rich) nuclear phosphoprotein 32 family, member A | 8125 | ENSG00000140350 |
| 18 | P49419 | NA | ALDH7A1 | aldehyde dehydrogenase 7 family, member A1 | 501 | ENSG00000164904 |
| 19 | P21980 | NA | TGM2 | transglutaminase 2 (C polypeptide, protein-glutamine-gamma-glutamyltransferase) | 7052 | ENSG00000198959 |
| 20 | Q5SSJ5 | NA | HP1BP3 | heterochromatin protein 1, binding protein 3 | 50809 | ENSG00000127483 |
| 21 | P06737 | NA | PYGL | phosphorylase, glycogen, liver | 5836 | ENSG00000100504 |
| 22 | P38571 | NA | LIPA | lipase A, lysosomal acid, cholesterol esterase | 3988 | ENSG00000107798 |
| 23 | Q13263 | NA | TRIM28 | tripartite motif containing 28 | 10155 | ENSG00000130726 |
| 24 | P32119 | NA | PRDX2 | peroxiredoxin 2 | 7001 | ENSG00000167815 |
| 25 | Q13011 | NA | ECH1 | enoyl CoA hydratase 1, peroxisomal | 1891 | ENSG00000104823 |
| 26 | Q9UBQ7 | NA | GRHPR | glyoxylate reductase/hydroxypyruvate reductase | 9380 | ENSG00000137106 |
| 27 | Q99798 | NA | ACO2 | aconitase 2, mitochondrial | 50 | ENSG00000100412 |
| 28 | P00558 | NA | PGK1 | phosphoglycerate kinase 1 | 5230 | ENSG00000102144 |
| 29 | Q01105 | NA | SET | SET nuclear oncogene | 6418 | ENSG00000119335 |
| 30 | P13674 | NA | P4HA1 | prolyl 4-hydroxylase, alpha polypeptide I | 5033 | ENSG00000122884 |
| 31 | O43598 | NA | C6orf108 | chromosome 6 open reading frame 108 | 10591 | ENSG00000112667 |
| 32 | P06744 | NA | GPI | glucose-6-phosphate isomerase | 2821 | ENSG00000105220 |
| 33 | P19367 | NA | HK1 | hexokinase 1 | 3098 | ENSG00000156515 |
| 34 | Q9BQ69 | NA | MACROD1 | MACRO domain containing 1 | 28992 | ENSG00000133315 |
| 35 | P00352 | NA | ALDH1A1 | aldehyde dehydrogenase 1 family, member A1 | 216 | ENSG00000165092 |
| 36 | Q16531 | NA | DDB1 | damage-specific DNA binding protein 1, 127kDa | 1642 | ENSG00000167986 |

  
  

| **Database:biological process      &nbspName:protein oligomerization      &nbspID:GO:0051259** | | | | | | |
| --- | --- | --- | --- | --- | --- | --- |
| C=337; O=6; E=1.00; R=5.97; rawP=0.0005; adjP=0.0064 | | | | | | |
| Index | UserID | Value | Gene Symbol | Gene Name | EntrezGene | Ensembl |
| 1 | Q9UBQ7 | NA | GRHPR | glyoxylate reductase/hydroxypyruvate reductase | 9380 | ENSG00000137106 |
| 2 | Q16698 | NA | DECR1 | 2,4-dienoyl CoA reductase 1, mitochondrial | 1666 | ENSG00000104325 |
| 3 | P21980 | NA | TGM2 | transglutaminase 2 (C polypeptide, protein-glutamine-gamma-glutamyltransferase) | 7052 | ENSG00000198959 |
| 4 | Q13263 | NA | TRIM28 | tripartite motif containing 28 | 10155 | ENSG00000130726 |
| 5 | Q99714 | NA | HSD17B10 | hydroxysteroid (17-beta) dehydrogenase 10 | 3028 | ENSG00000072506 |
| 6 | P04179 | NA | SOD2 | superoxide dismutase 2, mitochondrial | 6648 | ENSG00000112096 |

  
  

| **Database:biological process      &nbspName:metabolic process      &nbspID:GO:0008152** | | | | | | |
| --- | --- | --- | --- | --- | --- | --- |
| C=9363; O=38; E=27.92; R=1.36; rawP=0.0005; adjP=0.0064 | | | | | | |
| Index | UserID | Value | Gene Symbol | Gene Name | EntrezGene | Ensembl |
| 1 | P09661 | NA | SNRPA1 | small nuclear ribonucleoprotein polypeptide A' | 6627 | ENSG00000131876 |
| 2 | P10253 | NA | GAA | glucosidase, alpha; acid | 2548 | ENSG00000171298 |
| 3 | P54819 | NA | AK2 | adenylate kinase 2 | 204 | ENSG00000004455 |
| 4 | P17931 | NA | LGALS3 | lectin, galactoside-binding, soluble, 3 | 3958 | ENSG00000131981 |
| 5 | Q9UL46 | NA | PSME2 | proteasome (prosome, macropain) activator subunit 2 (PA28 beta) | 5721 | ENSG00000100911 |
| 6 | P50453 | NA | SERPINB9 | serpin peptidase inhibitor, clade B (ovalbumin), member 9 | 5272 | ENSG00000170542 |
| 7 | Q99714 | NA | HSD17B10 | hydroxysteroid (17-beta) dehydrogenase 10 | 3028 | ENSG00000072506 |
| 8 | P04179 | NA | SOD2 | superoxide dismutase 2, mitochondrial | 6648 | ENSG00000112096 |
| 9 | Q06323 | NA | PSME1 | proteasome (prosome, macropain) activator subunit 1 (PA28 alpha) | 5720 | ENSG00000092010 |
| 10 | P07741 | NA | APRT | adenine phosphoribosyltransferase | 353 | ENSG00000198931 |
| 11 | P30046 | NA | DDT | D-dopachrome tautomerase | 1652 | ENSG00000099977 |
| 12 | P31040 | NA | SDHA | succinate dehydrogenase complex, subunit A, flavoprotein (Fp) | 6389 | ENSG00000073578 |
| 13 | P38117 | NA | ETFB | electron-transfer-flavoprotein, beta polypeptide | 2109 | ENSG00000105379 |
| 14 | Q16698 | NA | DECR1 | 2,4-dienoyl CoA reductase 1, mitochondrial | 1666 | ENSG00000104325 |
| 15 | P54727 | NA | RAD23B | RAD23 homolog B (S. cerevisiae) | 5887 | ENSG00000119318 |
| 16 | Q10713 | NA | PMPCA | peptidase (mitochondrial processing) alpha | 23203 | ENSG00000165688 |
| 17 | P42765 | NA | ACAA2 | acetyl-CoA acyltransferase 2 | 10449 | ENSG00000167315 |
| 18 | P39687 | NA | ANP32A | acidic (leucine-rich) nuclear phosphoprotein 32 family, member A | 8125 | ENSG00000140350 |
| 19 | P49419 | NA | ALDH7A1 | aldehyde dehydrogenase 7 family, member A1 | 501 | ENSG00000164904 |
| 20 | P21980 | NA | TGM2 | transglutaminase 2 (C polypeptide, protein-glutamine-gamma-glutamyltransferase) | 7052 | ENSG00000198959 |
| 21 | Q5SSJ5 | NA | HP1BP3 | heterochromatin protein 1, binding protein 3 | 50809 | ENSG00000127483 |
| 22 | P06737 | NA | PYGL | phosphorylase, glycogen, liver | 5836 | ENSG00000100504 |
| 23 | P30837 | NA | ALDH1B1 | aldehyde dehydrogenase 1 family, member B1 | 219 | ENSG00000137124 |
| 24 | P38571 | NA | LIPA | lipase A, lysosomal acid, cholesterol esterase | 3988 | ENSG00000107798 |
| 25 | Q13263 | NA | TRIM28 | tripartite motif containing 28 | 10155 | ENSG00000130726 |
| 26 | P32119 | NA | PRDX2 | peroxiredoxin 2 | 7001 | ENSG00000167815 |
| 27 | Q13011 | NA | ECH1 | enoyl CoA hydratase 1, peroxisomal | 1891 | ENSG00000104823 |
| 28 | Q9UBQ7 | NA | GRHPR | glyoxylate reductase/hydroxypyruvate reductase | 9380 | ENSG00000137106 |
| 29 | Q99798 | NA | ACO2 | aconitase 2, mitochondrial | 50 | ENSG00000100412 |
| 30 | P00558 | NA | PGK1 | phosphoglycerate kinase 1 | 5230 | ENSG00000102144 |
| 31 | Q01105 | NA | SET | SET nuclear oncogene | 6418 | ENSG00000119335 |
| 32 | P13674 | NA | P4HA1 | prolyl 4-hydroxylase, alpha polypeptide I | 5033 | ENSG00000122884 |
| 33 | O43598 | NA | C6orf108 | chromosome 6 open reading frame 108 | 10591 | ENSG00000112667 |
| 34 | P06744 | NA | GPI | glucose-6-phosphate isomerase | 2821 | ENSG00000105220 |
| 35 | P19367 | NA | HK1 | hexokinase 1 | 3098 | ENSG00000156515 |
| 36 | Q9BQ69 | NA | MACROD1 | MACRO domain containing 1 | 28992 | ENSG00000133315 |
| 37 | P00352 | NA | ALDH1A1 | aldehyde dehydrogenase 1 family, member A1 | 216 | ENSG00000165092 |
| 38 | Q16531 | NA | DDB1 | damage-specific DNA binding protein 1, 127kDa | 1642 | ENSG00000167986 |

  
  

| **Database:biological process      &nbspName:acetyl-CoA metabolic process      &nbspID:GO:0006084** | | | | | | |
| --- | --- | --- | --- | --- | --- | --- |
| C=52; O=3; E=0.16; R=19.35; rawP=0.0005; adjP=0.0064 | | | | | | |
| Index | UserID | Value | Gene Symbol | Gene Name | EntrezGene | Ensembl |
| 1 | Q99798 | NA | ACO2 | aconitase 2, mitochondrial | 50 | ENSG00000100412 |
| 2 | P31040 | NA | SDHA | succinate dehydrogenase complex, subunit A, flavoprotein (Fp) | 6389 | ENSG00000073578 |
| 3 | P42765 | NA | ACAA2 | acetyl-CoA acyltransferase 2 | 10449 | ENSG00000167315 |

  
  

| **Database:biological process      &nbspName:fatty acid beta-oxidation      &nbspID:GO:0006635** | | | | | | |
| --- | --- | --- | --- | --- | --- | --- |
| C=57; O=3; E=0.17; R=17.65; rawP=0.0006; adjP=0.0070 | | | | | | |
| Index | UserID | Value | Gene Symbol | Gene Name | EntrezGene | Ensembl |
| 1 | Q16698 | NA | DECR1 | 2,4-dienoyl CoA reductase 1, mitochondrial | 1666 | ENSG00000104325 |
| 2 | Q13011 | NA | ECH1 | enoyl CoA hydratase 1, peroxisomal | 1891 | ENSG00000104823 |
| 3 | P42765 | NA | ACAA2 | acetyl-CoA acyltransferase 2 | 10449 | ENSG00000167315 |

  
  

| **Database:biological process      &nbspName:protein homotetramerization      &nbspID:GO:0051289** | | | | | | |
| --- | --- | --- | --- | --- | --- | --- |
| C=55; O=3; E=0.16; R=18.29; rawP=0.0006; adjP=0.0070 | | | | | | |
| Index | UserID | Value | Gene Symbol | Gene Name | EntrezGene | Ensembl |
| 1 | Q16698 | NA | DECR1 | 2,4-dienoyl CoA reductase 1, mitochondrial | 1666 | ENSG00000104325 |
| 2 | Q99714 | NA | HSD17B10 | hydroxysteroid (17-beta) dehydrogenase 10 | 3028 | ENSG00000072506 |
| 3 | P04179 | NA | SOD2 | superoxide dismutase 2, mitochondrial | 6648 | ENSG00000112096 |

  
  

| **Database:biological process      &nbspName:hexose metabolic process      &nbspID:GO:0019318** | | | | | | |
| --- | --- | --- | --- | --- | --- | --- |
| C=249; O=5; E=0.74; R=6.73; rawP=0.0008; adjP=0.0087 | | | | | | |
| Index | UserID | Value | Gene Symbol | Gene Name | EntrezGene | Ensembl |
| 1 | P06737 | NA | PYGL | phosphorylase, glycogen, liver | 5836 | ENSG00000100504 |
| 2 | P10253 | NA | GAA | glucosidase, alpha; acid | 2548 | ENSG00000171298 |
| 3 | P06744 | NA | GPI | glucose-6-phosphate isomerase | 2821 | ENSG00000105220 |
| 4 | P19367 | NA | HK1 | hexokinase 1 | 3098 | ENSG00000156515 |
| 5 | P00558 | NA | PGK1 | phosphoglycerate kinase 1 | 5230 | ENSG00000102144 |

  
  

| **Database:biological process      &nbspName:single-organism metabolic process      &nbspID:GO:0044710** | | | | | | |
| --- | --- | --- | --- | --- | --- | --- |
| C=9109; O=37; E=27.16; R=1.36; rawP=0.0008; adjP=0.0087 | | | | | | |
| Index | UserID | Value | Gene Symbol | Gene Name | EntrezGene | Ensembl |
| 1 | P09661 | NA | SNRPA1 | small nuclear ribonucleoprotein polypeptide A' | 6627 | ENSG00000131876 |
| 2 | P10253 | NA | GAA | glucosidase, alpha; acid | 2548 | ENSG00000171298 |
| 3 | P54819 | NA | AK2 | adenylate kinase 2 | 204 | ENSG00000004455 |
| 4 | P17931 | NA | LGALS3 | lectin, galactoside-binding, soluble, 3 | 3958 | ENSG00000131981 |
| 5 | Q9UL46 | NA | PSME2 | proteasome (prosome, macropain) activator subunit 2 (PA28 beta) | 5721 | ENSG00000100911 |
| 6 | P50453 | NA | SERPINB9 | serpin peptidase inhibitor, clade B (ovalbumin), member 9 | 5272 | ENSG00000170542 |
| 7 | Q99714 | NA | HSD17B10 | hydroxysteroid (17-beta) dehydrogenase 10 | 3028 | ENSG00000072506 |
| 8 | P04179 | NA | SOD2 | superoxide dismutase 2, mitochondrial | 6648 | ENSG00000112096 |
| 9 | Q06323 | NA | PSME1 | proteasome (prosome, macropain) activator subunit 1 (PA28 alpha) | 5720 | ENSG00000092010 |
| 10 | P07741 | NA | APRT | adenine phosphoribosyltransferase | 353 | ENSG00000198931 |
| 11 | P30046 | NA | DDT | D-dopachrome tautomerase | 1652 | ENSG00000099977 |
| 12 | P31040 | NA | SDHA | succinate dehydrogenase complex, subunit A, flavoprotein (Fp) | 6389 | ENSG00000073578 |
| 13 | P38117 | NA | ETFB | electron-transfer-flavoprotein, beta polypeptide | 2109 | ENSG00000105379 |
| 14 | Q16698 | NA | DECR1 | 2,4-dienoyl CoA reductase 1, mitochondrial | 1666 | ENSG00000104325 |
| 15 | P54727 | NA | RAD23B | RAD23 homolog B (S. cerevisiae) | 5887 | ENSG00000119318 |
| 16 | Q10713 | NA | PMPCA | peptidase (mitochondrial processing) alpha | 23203 | ENSG00000165688 |
| 17 | P42765 | NA | ACAA2 | acetyl-CoA acyltransferase 2 | 10449 | ENSG00000167315 |
| 18 | P39687 | NA | ANP32A | acidic (leucine-rich) nuclear phosphoprotein 32 family, member A | 8125 | ENSG00000140350 |
| 19 | P49419 | NA | ALDH7A1 | aldehyde dehydrogenase 7 family, member A1 | 501 | ENSG00000164904 |
| 20 | P21980 | NA | TGM2 | transglutaminase 2 (C polypeptide, protein-glutamine-gamma-glutamyltransferase) | 7052 | ENSG00000198959 |
| 21 | Q5SSJ5 | NA | HP1BP3 | heterochromatin protein 1, binding protein 3 | 50809 | ENSG00000127483 |
| 22 | P06737 | NA | PYGL | phosphorylase, glycogen, liver | 5836 | ENSG00000100504 |
| 23 | P30837 | NA | ALDH1B1 | aldehyde dehydrogenase 1 family, member B1 | 219 | ENSG00000137124 |
| 24 | P38571 | NA | LIPA | lipase A, lysosomal acid, cholesterol esterase | 3988 | ENSG00000107798 |
| 25 | Q13263 | NA | TRIM28 | tripartite motif containing 28 | 10155 | ENSG00000130726 |
| 26 | Q13011 | NA | ECH1 | enoyl CoA hydratase 1, peroxisomal | 1891 | ENSG00000104823 |
| 27 | Q9UBQ7 | NA | GRHPR | glyoxylate reductase/hydroxypyruvate reductase | 9380 | ENSG00000137106 |
| 28 | Q99798 | NA | ACO2 | aconitase 2, mitochondrial | 50 | ENSG00000100412 |
| 29 | P00558 | NA | PGK1 | phosphoglycerate kinase 1 | 5230 | ENSG00000102144 |
| 30 | Q01105 | NA | SET | SET nuclear oncogene | 6418 | ENSG00000119335 |
| 31 | P13674 | NA | P4HA1 | prolyl 4-hydroxylase, alpha polypeptide I | 5033 | ENSG00000122884 |
| 32 | O43598 | NA | C6orf108 | chromosome 6 open reading frame 108 | 10591 | ENSG00000112667 |
| 33 | P06744 | NA | GPI | glucose-6-phosphate isomerase | 2821 | ENSG00000105220 |
| 34 | P19367 | NA | HK1 | hexokinase 1 | 3098 | ENSG00000156515 |
| 35 | Q9BQ69 | NA | MACROD1 | MACRO domain containing 1 | 28992 | ENSG00000133315 |
| 36 | P00352 | NA | ALDH1A1 | aldehyde dehydrogenase 1 family, member A1 | 216 | ENSG00000165092 |
| 37 | Q16531 | NA | DDB1 | damage-specific DNA binding protein 1, 127kDa | 1642 | ENSG00000167986 |

  
  

| **Database:biological process      &nbspName:glycolysis      &nbspID:GO:0006096** | | | | | | |
| --- | --- | --- | --- | --- | --- | --- |
| C=63; O=3; E=0.19; R=15.97; rawP=0.0009; adjP=0.0094 | | | | | | |
| Index | UserID | Value | Gene Symbol | Gene Name | EntrezGene | Ensembl |
| 1 | P06744 | NA | GPI | glucose-6-phosphate isomerase | 2821 | ENSG00000105220 |
| 2 | P19367 | NA | HK1 | hexokinase 1 | 3098 | ENSG00000156515 |
| 3 | P00558 | NA | PGK1 | phosphoglycerate kinase 1 | 5230 | ENSG00000102144 |

  
  

| **Database:biological process      &nbspName:fatty acid catabolic process      &nbspID:GO:0009062** | | | | | | |
| --- | --- | --- | --- | --- | --- | --- |
| C=75; O=3; E=0.22; R=13.42; rawP=0.0014; adjP=0.0121 | | | | | | |
| Index | UserID | Value | Gene Symbol | Gene Name | EntrezGene | Ensembl |
| 1 | Q16698 | NA | DECR1 | 2,4-dienoyl CoA reductase 1, mitochondrial | 1666 | ENSG00000104325 |
| 2 | Q13011 | NA | ECH1 | enoyl CoA hydratase 1, peroxisomal | 1891 | ENSG00000104823 |
| 3 | P42765 | NA | ACAA2 | acetyl-CoA acyltransferase 2 | 10449 | ENSG00000167315 |

  
  

| **Database:biological process      &nbspName:glucose catabolic process      &nbspID:GO:0006007** | | | | | | |
| --- | --- | --- | --- | --- | --- | --- |
| C=77; O=3; E=0.23; R=13.07; rawP=0.0015; adjP=0.0121 | | | | | | |
| Index | UserID | Value | Gene Symbol | Gene Name | EntrezGene | Ensembl |
| 1 | P06744 | NA | GPI | glucose-6-phosphate isomerase | 2821 | ENSG00000105220 |
| 2 | P19367 | NA | HK1 | hexokinase 1 | 3098 | ENSG00000156515 |
| 3 | P00558 | NA | PGK1 | phosphoglycerate kinase 1 | 5230 | ENSG00000102144 |

  
  

| **Database:biological process      &nbspName:monocarboxylic acid metabolic process      &nbspID:GO:0032787** | | | | | | |
| --- | --- | --- | --- | --- | --- | --- |
| C=405; O=6; E=1.21; R=4.97; rawP=0.0012; adjP=0.0121 | | | | | | |
| Index | UserID | Value | Gene Symbol | Gene Name | EntrezGene | Ensembl |
| 1 | P13674 | NA | P4HA1 | prolyl 4-hydroxylase, alpha polypeptide I | 5033 | ENSG00000122884 |
| 2 | Q9UBQ7 | NA | GRHPR | glyoxylate reductase/hydroxypyruvate reductase | 9380 | ENSG00000137106 |
| 3 | Q16698 | NA | DECR1 | 2,4-dienoyl CoA reductase 1, mitochondrial | 1666 | ENSG00000104325 |
| 4 | P38571 | NA | LIPA | lipase A, lysosomal acid, cholesterol esterase | 3988 | ENSG00000107798 |
| 5 | Q13011 | NA | ECH1 | enoyl CoA hydratase 1, peroxisomal | 1891 | ENSG00000104823 |
| 6 | P42765 | NA | ACAA2 | acetyl-CoA acyltransferase 2 | 10449 | ENSG00000167315 |

  
  

| **Database:biological process      &nbspName:negative regulation of programmed cell death      &nbspID:GO:0043069** | | | | | | |
| --- | --- | --- | --- | --- | --- | --- |
| C=580; O=7; E=1.73; R=4.05; rawP=0.0015; adjP=0.0121 | | | | | | |
| Index | UserID | Value | Gene Symbol | Gene Name | EntrezGene | Ensembl |
| 1 | P06744 | NA | GPI | glucose-6-phosphate isomerase | 2821 | ENSG00000105220 |
| 2 | P05783 | NA | KRT18 | keratin 18 | 3875 | ENSG00000111057 |
| 3 | P32119 | NA | PRDX2 | peroxiredoxin 2 | 7001 | ENSG00000167815 |
| 4 | Q01105 | NA | SET | SET nuclear oncogene | 6418 | ENSG00000119335 |
| 5 | P50453 | NA | SERPINB9 | serpin peptidase inhibitor, clade B (ovalbumin), member 9 | 5272 | ENSG00000170542 |
| 6 | P42765 | NA | ACAA2 | acetyl-CoA acyltransferase 2 | 10449 | ENSG00000167315 |
| 7 | P04179 | NA | SOD2 | superoxide dismutase 2, mitochondrial | 6648 | ENSG00000112096 |

  
  

| **Database:biological process      &nbspName:organic substance metabolic process      &nbspID:GO:0071704** | | | | | | |
| --- | --- | --- | --- | --- | --- | --- |
| C=8902; O=36; E=26.54; R=1.36; rawP=0.0015; adjP=0.0121 | | | | | | |
| Index | UserID | Value | Gene Symbol | Gene Name | EntrezGene | Ensembl |
| 1 | P09661 | NA | SNRPA1 | small nuclear ribonucleoprotein polypeptide A' | 6627 | ENSG00000131876 |
| 2 | P10253 | NA | GAA | glucosidase, alpha; acid | 2548 | ENSG00000171298 |
| 3 | P54819 | NA | AK2 | adenylate kinase 2 | 204 | ENSG00000004455 |
| 4 | P17931 | NA | LGALS3 | lectin, galactoside-binding, soluble, 3 | 3958 | ENSG00000131981 |
| 5 | Q9UL46 | NA | PSME2 | proteasome (prosome, macropain) activator subunit 2 (PA28 beta) | 5721 | ENSG00000100911 |
| 6 | P50453 | NA | SERPINB9 | serpin peptidase inhibitor, clade B (ovalbumin), member 9 | 5272 | ENSG00000170542 |
| 7 | Q99714 | NA | HSD17B10 | hydroxysteroid (17-beta) dehydrogenase 10 | 3028 | ENSG00000072506 |
| 8 | P04179 | NA | SOD2 | superoxide dismutase 2, mitochondrial | 6648 | ENSG00000112096 |
| 9 | Q06323 | NA | PSME1 | proteasome (prosome, macropain) activator subunit 1 (PA28 alpha) | 5720 | ENSG00000092010 |
| 10 | P07741 | NA | APRT | adenine phosphoribosyltransferase | 353 | ENSG00000198931 |
| 11 | P30046 | NA | DDT | D-dopachrome tautomerase | 1652 | ENSG00000099977 |
| 12 | P31040 | NA | SDHA | succinate dehydrogenase complex, subunit A, flavoprotein (Fp) | 6389 | ENSG00000073578 |
| 13 | Q16698 | NA | DECR1 | 2,4-dienoyl CoA reductase 1, mitochondrial | 1666 | ENSG00000104325 |
| 14 | P54727 | NA | RAD23B | RAD23 homolog B (S. cerevisiae) | 5887 | ENSG00000119318 |
| 15 | P42765 | NA | ACAA2 | acetyl-CoA acyltransferase 2 | 10449 | ENSG00000167315 |
| 16 | Q10713 | NA | PMPCA | peptidase (mitochondrial processing) alpha | 23203 | ENSG00000165688 |
| 17 | P39687 | NA | ANP32A | acidic (leucine-rich) nuclear phosphoprotein 32 family, member A | 8125 | ENSG00000140350 |
| 18 | P49419 | NA | ALDH7A1 | aldehyde dehydrogenase 7 family, member A1 | 501 | ENSG00000164904 |
| 19 | P21980 | NA | TGM2 | transglutaminase 2 (C polypeptide, protein-glutamine-gamma-glutamyltransferase) | 7052 | ENSG00000198959 |
| 20 | Q5SSJ5 | NA | HP1BP3 | heterochromatin protein 1, binding protein 3 | 50809 | ENSG00000127483 |
| 21 | P06737 | NA | PYGL | phosphorylase, glycogen, liver | 5836 | ENSG00000100504 |
| 22 | P30837 | NA | ALDH1B1 | aldehyde dehydrogenase 1 family, member B1 | 219 | ENSG00000137124 |
| 23 | P38571 | NA | LIPA | lipase A, lysosomal acid, cholesterol esterase | 3988 | ENSG00000107798 |
| 24 | Q13263 | NA | TRIM28 | tripartite motif containing 28 | 10155 | ENSG00000130726 |
| 25 | Q13011 | NA | ECH1 | enoyl CoA hydratase 1, peroxisomal | 1891 | ENSG00000104823 |
| 26 | Q9UBQ7 | NA | GRHPR | glyoxylate reductase/hydroxypyruvate reductase | 9380 | ENSG00000137106 |
| 27 | Q99798 | NA | ACO2 | aconitase 2, mitochondrial | 50 | ENSG00000100412 |
| 28 | P00558 | NA | PGK1 | phosphoglycerate kinase 1 | 5230 | ENSG00000102144 |
| 29 | Q01105 | NA | SET | SET nuclear oncogene | 6418 | ENSG00000119335 |
| 30 | P13674 | NA | P4HA1 | prolyl 4-hydroxylase, alpha polypeptide I | 5033 | ENSG00000122884 |
| 31 | O43598 | NA | C6orf108 | chromosome 6 open reading frame 108 | 10591 | ENSG00000112667 |
| 32 | P06744 | NA | GPI | glucose-6-phosphate isomerase | 2821 | ENSG00000105220 |
| 33 | P19367 | NA | HK1 | hexokinase 1 | 3098 | ENSG00000156515 |
| 34 | Q9BQ69 | NA | MACROD1 | MACRO domain containing 1 | 28992 | ENSG00000133315 |
| 35 | P00352 | NA | ALDH1A1 | aldehyde dehydrogenase 1 family, member A1 | 216 | ENSG00000165092 |
| 36 | Q16531 | NA | DDB1 | damage-specific DNA binding protein 1, 127kDa | 1642 | ENSG00000167986 |

  
  

| **Database:biological process      &nbspName:monosaccharide metabolic process      &nbspID:GO:0005996** | | | | | | |
| --- | --- | --- | --- | --- | --- | --- |
| C=279; O=5; E=0.83; R=6.01; rawP=0.0014; adjP=0.0121 | | | | | | |
| Index | UserID | Value | Gene Symbol | Gene Name | EntrezGene | Ensembl |
| 1 | P06737 | NA | PYGL | phosphorylase, glycogen, liver | 5836 | ENSG00000100504 |
| 2 | P10253 | NA | GAA | glucosidase, alpha; acid | 2548 | ENSG00000171298 |
| 3 | P06744 | NA | GPI | glucose-6-phosphate isomerase | 2821 | ENSG00000105220 |
| 4 | P19367 | NA | HK1 | hexokinase 1 | 3098 | ENSG00000156515 |
| 5 | P00558 | NA | PGK1 | phosphoglycerate kinase 1 | 5230 | ENSG00000102144 |

  
  

| **Database:biological process      &nbspName:negative regulation of apoptotic process      &nbspID:GO:0043066** | | | | | | |
| --- | --- | --- | --- | --- | --- | --- |
| C=570; O=7; E=1.70; R=4.12; rawP=0.0014; adjP=0.0121 | | | | | | |
| Index | UserID | Value | Gene Symbol | Gene Name | EntrezGene | Ensembl |
| 1 | P06744 | NA | GPI | glucose-6-phosphate isomerase | 2821 | ENSG00000105220 |
| 2 | P05783 | NA | KRT18 | keratin 18 | 3875 | ENSG00000111057 |
| 3 | P32119 | NA | PRDX2 | peroxiredoxin 2 | 7001 | ENSG00000167815 |
| 4 | Q01105 | NA | SET | SET nuclear oncogene | 6418 | ENSG00000119335 |
| 5 | P50453 | NA | SERPINB9 | serpin peptidase inhibitor, clade B (ovalbumin), member 9 | 5272 | ENSG00000170542 |
| 6 | P42765 | NA | ACAA2 | acetyl-CoA acyltransferase 2 | 10449 | ENSG00000167315 |
| 7 | P04179 | NA | SOD2 | superoxide dismutase 2, mitochondrial | 6648 | ENSG00000112096 |

  
  

| **Database:biological process      &nbspName:cellular respiration      &nbspID:GO:0045333** | | | | | | |
| --- | --- | --- | --- | --- | --- | --- |
| C=159; O=4; E=0.47; R=8.44; rawP=0.0013; adjP=0.0121 | | | | | | |
| Index | UserID | Value | Gene Symbol | Gene Name | EntrezGene | Ensembl |
| 1 | Q99798 | NA | ACO2 | aconitase 2, mitochondrial | 50 | ENSG00000100412 |
| 2 | P31040 | NA | SDHA | succinate dehydrogenase complex, subunit A, flavoprotein (Fp) | 6389 | ENSG00000073578 |
| 3 | P38117 | NA | ETFB | electron-transfer-flavoprotein, beta polypeptide | 2109 | ENSG00000105379 |
| 4 | P04179 | NA | SOD2 | superoxide dismutase 2, mitochondrial | 6648 | ENSG00000112096 |

  
  

| **Database:biological process      &nbspName:regulation of apoptotic process      &nbspID:GO:0042981** | | | | | | |
| --- | --- | --- | --- | --- | --- | --- |
| C=1150; O=10; E=3.43; R=2.92; rawP=0.0017; adjP=0.0133 | | | | | | |
| Index | UserID | Value | Gene Symbol | Gene Name | EntrezGene | Ensembl |
| 1 | Q9UL46 | NA | PSME2 | proteasome (prosome, macropain) activator subunit 2 (PA28 beta) | 5721 | ENSG00000100911 |
| 2 | Q01105 | NA | SET | SET nuclear oncogene | 6418 | ENSG00000119335 |
| 3 | P50453 | NA | SERPINB9 | serpin peptidase inhibitor, clade B (ovalbumin), member 9 | 5272 | ENSG00000170542 |
| 4 | P42765 | NA | ACAA2 | acetyl-CoA acyltransferase 2 | 10449 | ENSG00000167315 |
| 5 | P04179 | NA | SOD2 | superoxide dismutase 2, mitochondrial | 6648 | ENSG00000112096 |
| 6 | Q06323 | NA | PSME1 | proteasome (prosome, macropain) activator subunit 1 (PA28 alpha) | 5720 | ENSG00000092010 |
| 7 | P06744 | NA | GPI | glucose-6-phosphate isomerase | 2821 | ENSG00000105220 |
| 8 | P21980 | NA | TGM2 | transglutaminase 2 (C polypeptide, protein-glutamine-gamma-glutamyltransferase) | 7052 | ENSG00000198959 |
| 9 | P05783 | NA | KRT18 | keratin 18 | 3875 | ENSG00000111057 |
| 10 | P32119 | NA | PRDX2 | peroxiredoxin 2 | 7001 | ENSG00000167815 |

  
  

| **Database:biological process      &nbspName:negative regulation of cell death      &nbspID:GO:0060548** | | | | | | |
| --- | --- | --- | --- | --- | --- | --- |
| C=598; O=7; E=1.78; R=3.93; rawP=0.0018; adjP=0.0134 | | | | | | |
| Index | UserID | Value | Gene Symbol | Gene Name | EntrezGene | Ensembl |
| 1 | P06744 | NA | GPI | glucose-6-phosphate isomerase | 2821 | ENSG00000105220 |
| 2 | P05783 | NA | KRT18 | keratin 18 | 3875 | ENSG00000111057 |
| 3 | P32119 | NA | PRDX2 | peroxiredoxin 2 | 7001 | ENSG00000167815 |
| 4 | Q01105 | NA | SET | SET nuclear oncogene | 6418 | ENSG00000119335 |
| 5 | P50453 | NA | SERPINB9 | serpin peptidase inhibitor, clade B (ovalbumin), member 9 | 5272 | ENSG00000170542 |
| 6 | P42765 | NA | ACAA2 | acetyl-CoA acyltransferase 2 | 10449 | ENSG00000167315 |
| 7 | P04179 | NA | SOD2 | superoxide dismutase 2, mitochondrial | 6648 | ENSG00000112096 |

  
  

| **Database:biological process      &nbspName:regulation of programmed cell death      &nbspID:GO:0043067** | | | | | | |
| --- | --- | --- | --- | --- | --- | --- |
| C=1164; O=10; E=3.47; R=2.88; rawP=0.0018; adjP=0.0134 | | | | | | |
| Index | UserID | Value | Gene Symbol | Gene Name | EntrezGene | Ensembl |
| 1 | Q9UL46 | NA | PSME2 | proteasome (prosome, macropain) activator subunit 2 (PA28 beta) | 5721 | ENSG00000100911 |
| 2 | Q01105 | NA | SET | SET nuclear oncogene | 6418 | ENSG00000119335 |
| 3 | P50453 | NA | SERPINB9 | serpin peptidase inhibitor, clade B (ovalbumin), member 9 | 5272 | ENSG00000170542 |
| 4 | P42765 | NA | ACAA2 | acetyl-CoA acyltransferase 2 | 10449 | ENSG00000167315 |
| 5 | P04179 | NA | SOD2 | superoxide dismutase 2, mitochondrial | 6648 | ENSG00000112096 |
| 6 | Q06323 | NA | PSME1 | proteasome (prosome, macropain) activator subunit 1 (PA28 alpha) | 5720 | ENSG00000092010 |
| 7 | P06744 | NA | GPI | glucose-6-phosphate isomerase | 2821 | ENSG00000105220 |
| 8 | P21980 | NA | TGM2 | transglutaminase 2 (C polypeptide, protein-glutamine-gamma-glutamyltransferase) | 7052 | ENSG00000198959 |
| 9 | P05783 | NA | KRT18 | keratin 18 | 3875 | ENSG00000111057 |
| 10 | P32119 | NA | PRDX2 | peroxiredoxin 2 | 7001 | ENSG00000167815 |

  
  

| **Database:biological process      &nbspName:protein tetramerization      &nbspID:GO:0051262** | | | | | | |
| --- | --- | --- | --- | --- | --- | --- |
| C=85; O=3; E=0.25; R=11.84; rawP=0.0021; adjP=0.0152 | | | | | | |
| Index | UserID | Value | Gene Symbol | Gene Name | EntrezGene | Ensembl |
| 1 | Q16698 | NA | DECR1 | 2,4-dienoyl CoA reductase 1, mitochondrial | 1666 | ENSG00000104325 |
| 2 | Q99714 | NA | HSD17B10 | hydroxysteroid (17-beta) dehydrogenase 10 | 3028 | ENSG00000072506 |
| 3 | P04179 | NA | SOD2 | superoxide dismutase 2, mitochondrial | 6648 | ENSG00000112096 |

  
  

| **Database:biological process      &nbspName:fatty acid oxidation      &nbspID:GO:0019395** | | | | | | |
| --- | --- | --- | --- | --- | --- | --- |
| C=89; O=3; E=0.27; R=11.31; rawP=0.0023; adjP=0.0154 | | | | | | |
| Index | UserID | Value | Gene Symbol | Gene Name | EntrezGene | Ensembl |
| 1 | Q16698 | NA | DECR1 | 2,4-dienoyl CoA reductase 1, mitochondrial | 1666 | ENSG00000104325 |
| 2 | Q13011 | NA | ECH1 | enoyl CoA hydratase 1, peroxisomal | 1891 | ENSG00000104823 |
| 3 | P42765 | NA | ACAA2 | acetyl-CoA acyltransferase 2 | 10449 | ENSG00000167315 |

  
  

| **Database:molecular function      &nbspName:aldehyde dehydrogenase (NAD) activity      &nbspID:GO:0004029** | | | | | | |
| --- | --- | --- | --- | --- | --- | --- |
| C=10; O=3; E=0.03; R=104.49; rawP=2.61e-06; adjP=7.07e-05 | | | | | | |
| Index | UserID | Value | Gene Symbol | Gene Name | EntrezGene | Ensembl |
| 1 | P49419 | NA | ALDH7A1 | aldehyde dehydrogenase 7 family, member A1 | 501 | ENSG00000164904 |
| 2 | P30837 | NA | ALDH1B1 | aldehyde dehydrogenase 1 family, member B1 | 219 | ENSG00000137124 |
| 3 | P00352 | NA | ALDH1A1 | aldehyde dehydrogenase 1 family, member A1 | 216 | ENSG00000165092 |

  
  

| **Database:molecular function      &nbspName:oxidoreductase activity      &nbspID:GO:0016491** | | | | | | |
| --- | --- | --- | --- | --- | --- | --- |
| C=697; O=11; E=2.00; R=5.50; rawP=3.01e-06; adjP=7.07e-05 | | | | | | |
| Index | UserID | Value | Gene Symbol | Gene Name | EntrezGene | Ensembl |
| 1 | Q9UBQ7 | NA | GRHPR | glyoxylate reductase/hydroxypyruvate reductase | 9380 | ENSG00000137106 |
| 2 | P02792 | NA | FTL | ferritin, light polypeptide | 2512 | ENSG00000087086 |
| 3 | Q16698 | NA | DECR1 | 2,4-dienoyl CoA reductase 1, mitochondrial | 1666 | ENSG00000104325 |
| 4 | Q99714 | NA | HSD17B10 | hydroxysteroid (17-beta) dehydrogenase 10 | 3028 | ENSG00000072506 |
| 5 | P04179 | NA | SOD2 | superoxide dismutase 2, mitochondrial | 6648 | ENSG00000112096 |
| 6 | P13674 | NA | P4HA1 | prolyl 4-hydroxylase, alpha polypeptide I | 5033 | ENSG00000122884 |
| 7 | P49419 | NA | ALDH7A1 | aldehyde dehydrogenase 7 family, member A1 | 501 | ENSG00000164904 |
| 8 | P30837 | NA | ALDH1B1 | aldehyde dehydrogenase 1 family, member B1 | 219 | ENSG00000137124 |
| 9 | P00352 | NA | ALDH1A1 | aldehyde dehydrogenase 1 family, member A1 | 216 | ENSG00000165092 |
| 10 | P31040 | NA | SDHA | succinate dehydrogenase complex, subunit A, flavoprotein (Fp) | 6389 | ENSG00000073578 |
| 11 | P32119 | NA | PRDX2 | peroxiredoxin 2 | 7001 | ENSG00000167815 |

  
  

| **Database:molecular function      &nbspName:catalytic activity      &nbspID:GO:0003824** | | | | | | |
| --- | --- | --- | --- | --- | --- | --- |
| C=5299; O=30; E=15.21; R=1.97; rawP=4.54e-06; adjP=7.11e-05 | | | | | | |
| Index | UserID | Value | Gene Symbol | Gene Name | EntrezGene | Ensembl |
| 1 | P02792 | NA | FTL | ferritin, light polypeptide | 2512 | ENSG00000087086 |
| 2 | P10253 | NA | GAA | glucosidase, alpha; acid | 2548 | ENSG00000171298 |
| 3 | P54819 | NA | AK2 | adenylate kinase 2 | 204 | ENSG00000004455 |
| 4 | Q99714 | NA | HSD17B10 | hydroxysteroid (17-beta) dehydrogenase 10 | 3028 | ENSG00000072506 |
| 5 | P04179 | NA | SOD2 | superoxide dismutase 2, mitochondrial | 6648 | ENSG00000112096 |
| 6 | P06737 | NA | PYGL | phosphorylase, glycogen, liver | 5836 | ENSG00000100504 |
| 7 | P30046 | NA | DDT | D-dopachrome tautomerase | 1652 | ENSG00000099977 |
| 8 | P07741 | NA | APRT | adenine phosphoribosyltransferase | 353 | ENSG00000198931 |
| 9 | Q9BXW7 | NA | CECR5 | cat eye syndrome chromosome region, candidate 5 | 27440 | ENSG00000069998 |
| 10 | P30837 | NA | ALDH1B1 | aldehyde dehydrogenase 1 family, member B1 | 219 | ENSG00000137124 |
| 11 | P38571 | NA | LIPA | lipase A, lysosomal acid, cholesterol esterase | 3988 | ENSG00000107798 |
| 12 | P32119 | NA | PRDX2 | peroxiredoxin 2 | 7001 | ENSG00000167815 |
| 13 | Q13263 | NA | TRIM28 | tripartite motif containing 28 | 10155 | ENSG00000130726 |
| 14 | P31040 | NA | SDHA | succinate dehydrogenase complex, subunit A, flavoprotein (Fp) | 6389 | ENSG00000073578 |
| 15 | Q13011 | NA | ECH1 | enoyl CoA hydratase 1, peroxisomal | 1891 | ENSG00000104823 |
| 16 | Q99798 | NA | ACO2 | aconitase 2, mitochondrial | 50 | ENSG00000100412 |
| 17 | Q9UBQ7 | NA | GRHPR | glyoxylate reductase/hydroxypyruvate reductase | 9380 | ENSG00000137106 |
| 18 | Q16698 | NA | DECR1 | 2,4-dienoyl CoA reductase 1, mitochondrial | 1666 | ENSG00000104325 |
| 19 | P00558 | NA | PGK1 | phosphoglycerate kinase 1 | 5230 | ENSG00000102144 |
| 20 | Q05639 | NA | EEF1A2 | eukaryotic translation elongation factor 1 alpha 2 | 1917 | ENSG00000101210 |
| 21 | P42765 | NA | ACAA2 | acetyl-CoA acyltransferase 2 | 10449 | ENSG00000167315 |
| 22 | Q10713 | NA | PMPCA | peptidase (mitochondrial processing) alpha | 23203 | ENSG00000165688 |
| 23 | P13674 | NA | P4HA1 | prolyl 4-hydroxylase, alpha polypeptide I | 5033 | ENSG00000122884 |
| 24 | O43598 | NA | C6orf108 | chromosome 6 open reading frame 108 | 10591 | ENSG00000112667 |
| 25 | P49419 | NA | ALDH7A1 | aldehyde dehydrogenase 7 family, member A1 | 501 | ENSG00000164904 |
| 26 | P06744 | NA | GPI | glucose-6-phosphate isomerase | 2821 | ENSG00000105220 |
| 27 | P19367 | NA | HK1 | hexokinase 1 | 3098 | ENSG00000156515 |
| 28 | P21980 | NA | TGM2 | transglutaminase 2 (C polypeptide, protein-glutamine-gamma-glutamyltransferase) | 7052 | ENSG00000198959 |
| 29 | Q9BQ69 | NA | MACROD1 | MACRO domain containing 1 | 28992 | ENSG00000133315 |
| 30 | P00352 | NA | ALDH1A1 | aldehyde dehydrogenase 1 family, member A1 | 216 | ENSG00000165092 |

  
  

| **Database:molecular function      &nbspName:oxidoreductase activity, acting on the aldehyde or oxo group of donors, NAD or NADP as acceptor      &nbspID:GO:0016620** | | | | | | |
| --- | --- | --- | --- | --- | --- | --- |
| C=29; O=3; E=0.08; R=36.03; rawP=7.65e-05; adjP=0.0009 | | | | | | |
| Index | UserID | Value | Gene Symbol | Gene Name | EntrezGene | Ensembl |
| 1 | P49419 | NA | ALDH7A1 | aldehyde dehydrogenase 7 family, member A1 | 501 | ENSG00000164904 |
| 2 | P30837 | NA | ALDH1B1 | aldehyde dehydrogenase 1 family, member B1 | 219 | ENSG00000137124 |
| 3 | P00352 | NA | ALDH1A1 | aldehyde dehydrogenase 1 family, member A1 | 216 | ENSG00000165092 |

  
  

| **Database:molecular function      &nbspName:oxidoreductase activity, acting on the aldehyde or oxo group of donors      &nbspID:GO:0016903** | | | | | | |
| --- | --- | --- | --- | --- | --- | --- |
| C=40; O=3; E=0.11; R=26.12; rawP=0.0002; adjP=0.0019 | | | | | | |
| Index | UserID | Value | Gene Symbol | Gene Name | EntrezGene | Ensembl |
| 1 | P49419 | NA | ALDH7A1 | aldehyde dehydrogenase 7 family, member A1 | 501 | ENSG00000164904 |
| 2 | P30837 | NA | ALDH1B1 | aldehyde dehydrogenase 1 family, member B1 | 219 | ENSG00000137124 |
| 3 | P00352 | NA | ALDH1A1 | aldehyde dehydrogenase 1 family, member A1 | 216 | ENSG00000165092 |

  
  

| **Database:molecular function      &nbspName:carbohydrate binding      &nbspID:GO:0030246** | | | | | | |
| --- | --- | --- | --- | --- | --- | --- |
| C=267; O=5; E=0.77; R=6.52; rawP=0.0010; adjP=0.0063 | | | | | | |
| Index | UserID | Value | Gene Symbol | Gene Name | EntrezGene | Ensembl |
| 1 | P13674 | NA | P4HA1 | prolyl 4-hydroxylase, alpha polypeptide I | 5033 | ENSG00000122884 |
| 2 | P06737 | NA | PYGL | phosphorylase, glycogen, liver | 5836 | ENSG00000100504 |
| 3 | P10253 | NA | GAA | glucosidase, alpha; acid | 2548 | ENSG00000171298 |
| 4 | P06744 | NA | GPI | glucose-6-phosphate isomerase | 2821 | ENSG00000105220 |
| 5 | P17931 | NA | LGALS3 | lectin, galactoside-binding, soluble, 3 | 3958 | ENSG00000131981 |

  
  

| **Database:molecular function      &nbspName:cofactor binding      &nbspID:GO:0048037** | | | | | | |
| --- | --- | --- | --- | --- | --- | --- |
| C=259; O=5; E=0.74; R=6.72; rawP=0.0008; adjP=0.0063 | | | | | | |
| Index | UserID | Value | Gene Symbol | Gene Name | EntrezGene | Ensembl |
| 1 | P06737 | NA | PYGL | phosphorylase, glycogen, liver | 5836 | ENSG00000100504 |
| 2 | Q9UBQ7 | NA | GRHPR | glyoxylate reductase/hydroxypyruvate reductase | 9380 | ENSG00000137106 |
| 3 | Q16698 | NA | DECR1 | 2,4-dienoyl CoA reductase 1, mitochondrial | 1666 | ENSG00000104325 |
| 4 | P31040 | NA | SDHA | succinate dehydrogenase complex, subunit A, flavoprotein (Fp) | 6389 | ENSG00000073578 |
| 5 | Q99714 | NA | HSD17B10 | hydroxysteroid (17-beta) dehydrogenase 10 | 3028 | ENSG00000072506 |

  
  

| **Database:molecular function      &nbspName:monosaccharide binding      &nbspID:GO:0048029** | | | | | | |
| --- | --- | --- | --- | --- | --- | --- |
| C=72; O=3; E=0.21; R=14.51; rawP=0.0011; adjP=0.0063 | | | | | | |
| Index | UserID | Value | Gene Symbol | Gene Name | EntrezGene | Ensembl |
| 1 | P13674 | NA | P4HA1 | prolyl 4-hydroxylase, alpha polypeptide I | 5033 | ENSG00000122884 |
| 2 | P06737 | NA | PYGL | phosphorylase, glycogen, liver | 5836 | ENSG00000100504 |
| 3 | P06744 | NA | GPI | glucose-6-phosphate isomerase | 2821 | ENSG00000105220 |

  
  

| **Database:molecular function      &nbspName:steroid binding      &nbspID:GO:0005496** | | | | | | |
| --- | --- | --- | --- | --- | --- | --- |
| C=73; O=3; E=0.21; R=14.31; rawP=0.0012; adjP=0.0063 | | | | | | |
| Index | UserID | Value | Gene Symbol | Gene Name | EntrezGene | Ensembl |
| 1 | P06737 | NA | PYGL | phosphorylase, glycogen, liver | 5836 | ENSG00000100504 |
| 2 | P00352 | NA | ALDH1A1 | aldehyde dehydrogenase 1 family, member A1 | 216 | ENSG00000165092 |
| 3 | Q99714 | NA | HSD17B10 | hydroxysteroid (17-beta) dehydrogenase 10 | 3028 | ENSG00000072506 |

  
  

| **Database:molecular function      &nbspName:coenzyme binding      &nbspID:GO:0050662** | | | | | | |
| --- | --- | --- | --- | --- | --- | --- |
| C=184; O=4; E=0.53; R=7.57; rawP=0.0019; adjP=0.0089 | | | | | | |
| Index | UserID | Value | Gene Symbol | Gene Name | EntrezGene | Ensembl |
| 1 | Q9UBQ7 | NA | GRHPR | glyoxylate reductase/hydroxypyruvate reductase | 9380 | ENSG00000137106 |
| 2 | Q16698 | NA | DECR1 | 2,4-dienoyl CoA reductase 1, mitochondrial | 1666 | ENSG00000104325 |
| 3 | P31040 | NA | SDHA | succinate dehydrogenase complex, subunit A, flavoprotein (Fp) | 6389 | ENSG00000073578 |
| 4 | Q99714 | NA | HSD17B10 | hydroxysteroid (17-beta) dehydrogenase 10 | 3028 | ENSG00000072506 |

  
  

| **Database:molecular function      &nbspName:isomerase activity      &nbspID:GO:0016853** | | | | | | |
| --- | --- | --- | --- | --- | --- | --- |
| C=142; O=3; E=0.41; R=7.36; rawP=0.0078; adjP=0.0333 | | | | | | |
| Index | UserID | Value | Gene Symbol | Gene Name | EntrezGene | Ensembl |
| 1 | P30046 | NA | DDT | D-dopachrome tautomerase | 1652 | ENSG00000099977 |
| 2 | P06744 | NA | GPI | glucose-6-phosphate isomerase | 2821 | ENSG00000105220 |
| 3 | Q13011 | NA | ECH1 | enoyl CoA hydratase 1, peroxisomal | 1891 | ENSG00000104823 |

  
  

| **Database:molecular function      &nbspName:iron ion binding      &nbspID:GO:0005506** | | | | | | |
| --- | --- | --- | --- | --- | --- | --- |
| C=164; O=3; E=0.47; R=6.37; rawP=0.0115; adjP=0.0450 | | | | | | |
| Index | UserID | Value | Gene Symbol | Gene Name | EntrezGene | Ensembl |
| 1 | P13674 | NA | P4HA1 | prolyl 4-hydroxylase, alpha polypeptide I | 5033 | ENSG00000122884 |
| 2 | Q99798 | NA | ACO2 | aconitase 2, mitochondrial | 50 | ENSG00000100412 |
| 3 | P02792 | NA | FTL | ferritin, light polypeptide | 2512 | ENSG00000087086 |

  
  

| **Database:cellular component      &nbspName:cytoplasm      &nbspID:GO:0005737** | | | | | | |
| --- | --- | --- | --- | --- | --- | --- |
| C=9051; O=42; E=24.42; R=1.72; rawP=1.00e-08; adjP=4.60e-07 | | | | | | |
| Index | UserID | Value | Gene Symbol | Gene Name | EntrezGene | Ensembl |
| 1 | P10253 | NA | GAA | glucosidase, alpha; acid | 2548 | ENSG00000171298 |
| 2 | P54819 | NA | AK2 | adenylate kinase 2 | 204 | ENSG00000004455 |
| 3 | P17931 | NA | LGALS3 | lectin, galactoside-binding, soluble, 3 | 3958 | ENSG00000131981 |
| 4 | P05787 | NA | KRT8 | keratin 8 | 3856 | ENSG00000170421 |
| 5 | Q9UL46 | NA | PSME2 | proteasome (prosome, macropain) activator subunit 2 (PA28 beta) | 5721 | ENSG00000100911 |
| 6 | P50453 | NA | SERPINB9 | serpin peptidase inhibitor, clade B (ovalbumin), member 9 | 5272 | ENSG00000170542 |
| 7 | Q99714 | NA | HSD17B10 | hydroxysteroid (17-beta) dehydrogenase 10 | 3028 | ENSG00000072506 |
| 8 | P04179 | NA | SOD2 | superoxide dismutase 2, mitochondrial | 6648 | ENSG00000112096 |
| 9 | Q06323 | NA | PSME1 | proteasome (prosome, macropain) activator subunit 1 (PA28 alpha) | 5720 | ENSG00000092010 |
| 10 | P07741 | NA | APRT | adenine phosphoribosyltransferase | 353 | ENSG00000198931 |
| 11 | P30046 | NA | DDT | D-dopachrome tautomerase | 1652 | ENSG00000099977 |
| 12 | P31040 | NA | SDHA | succinate dehydrogenase complex, subunit A, flavoprotein (Fp) | 6389 | ENSG00000073578 |
| 13 | P38117 | NA | ETFB | electron-transfer-flavoprotein, beta polypeptide | 2109 | ENSG00000105379 |
| 14 | Q16698 | NA | DECR1 | 2,4-dienoyl CoA reductase 1, mitochondrial | 1666 | ENSG00000104325 |
| 15 | Q05639 | NA | EEF1A2 | eukaryotic translation elongation factor 1 alpha 2 | 1917 | ENSG00000101210 |
| 16 | P54727 | NA | RAD23B | RAD23 homolog B (S. cerevisiae) | 5887 | ENSG00000119318 |
| 17 | Q10713 | NA | PMPCA | peptidase (mitochondrial processing) alpha | 23203 | ENSG00000165688 |
| 18 | P42765 | NA | ACAA2 | acetyl-CoA acyltransferase 2 | 10449 | ENSG00000167315 |
| 19 | P39687 | NA | ANP32A | acidic (leucine-rich) nuclear phosphoprotein 32 family, member A | 8125 | ENSG00000140350 |
| 20 | P49419 | NA | ALDH7A1 | aldehyde dehydrogenase 7 family, member A1 | 501 | ENSG00000164904 |
| 21 | P21980 | NA | TGM2 | transglutaminase 2 (C polypeptide, protein-glutamine-gamma-glutamyltransferase) | 7052 | ENSG00000198959 |
| 22 | Q9UDT6 | NA | CLIP2 | CAP-GLY domain containing linker protein 2 | 7461 | ENSG00000106665 |
| 23 | P02792 | NA | FTL | ferritin, light polypeptide | 2512 | ENSG00000087086 |
| 24 | P26447 | NA | S100A4 | S100 calcium binding protein A4 | 6275 | ENSG00000196154 |
| 25 | P06737 | NA | PYGL | phosphorylase, glycogen, liver | 5836 | ENSG00000100504 |
| 26 | Q9BXW7 | NA | CECR5 | cat eye syndrome chromosome region, candidate 5 | 27440 | ENSG00000069998 |
| 27 | P30837 | NA | ALDH1B1 | aldehyde dehydrogenase 1 family, member B1 | 219 | ENSG00000137124 |
| 28 | P38571 | NA | LIPA | lipase A, lysosomal acid, cholesterol esterase | 3988 | ENSG00000107798 |
| 29 | P32119 | NA | PRDX2 | peroxiredoxin 2 | 7001 | ENSG00000167815 |
| 30 | Q13011 | NA | ECH1 | enoyl CoA hydratase 1, peroxisomal | 1891 | ENSG00000104823 |
| 31 | Q99798 | NA | ACO2 | aconitase 2, mitochondrial | 50 | ENSG00000100412 |
| 32 | Q9UBQ7 | NA | GRHPR | glyoxylate reductase/hydroxypyruvate reductase | 9380 | ENSG00000137106 |
| 33 | P00558 | NA | PGK1 | phosphoglycerate kinase 1 | 5230 | ENSG00000102144 |
| 34 | Q01105 | NA | SET | SET nuclear oncogene | 6418 | ENSG00000119335 |
| 35 | P13674 | NA | P4HA1 | prolyl 4-hydroxylase, alpha polypeptide I | 5033 | ENSG00000122884 |
| 36 | O43598 | NA | C6orf108 | chromosome 6 open reading frame 108 | 10591 | ENSG00000112667 |
| 37 | P06744 | NA | GPI | glucose-6-phosphate isomerase | 2821 | ENSG00000105220 |
| 38 | P19367 | NA | HK1 | hexokinase 1 | 3098 | ENSG00000156515 |
| 39 | Q9BQ69 | NA | MACROD1 | MACRO domain containing 1 | 28992 | ENSG00000133315 |
| 40 | P05783 | NA | KRT18 | keratin 18 | 3875 | ENSG00000111057 |
| 41 | P00352 | NA | ALDH1A1 | aldehyde dehydrogenase 1 family, member A1 | 216 | ENSG00000165092 |
| 42 | Q16531 | NA | DDB1 | damage-specific DNA binding protein 1, 127kDa | 1642 | ENSG00000167986 |

  
  

| **Database:cellular component      &nbspName:mitochondrion      &nbspID:GO:0005739** | | | | | | |
| --- | --- | --- | --- | --- | --- | --- |
| C=1515; O=18; E=4.09; R=4.40; rawP=2.52e-08; adjP=5.80e-07 | | | | | | |
| Index | UserID | Value | Gene Symbol | Gene Name | EntrezGene | Ensembl |
| 1 | P54819 | NA | AK2 | adenylate kinase 2 | 204 | ENSG00000004455 |
| 2 | P17931 | NA | LGALS3 | lectin, galactoside-binding, soluble, 3 | 3958 | ENSG00000131981 |
| 3 | Q99714 | NA | HSD17B10 | hydroxysteroid (17-beta) dehydrogenase 10 | 3028 | ENSG00000072506 |
| 4 | P04179 | NA | SOD2 | superoxide dismutase 2, mitochondrial | 6648 | ENSG00000112096 |
| 5 | Q9BXW7 | NA | CECR5 | cat eye syndrome chromosome region, candidate 5 | 27440 | ENSG00000069998 |
| 6 | P30837 | NA | ALDH1B1 | aldehyde dehydrogenase 1 family, member B1 | 219 | ENSG00000137124 |
| 7 | P31040 | NA | SDHA | succinate dehydrogenase complex, subunit A, flavoprotein (Fp) | 6389 | ENSG00000073578 |
| 8 | Q13011 | NA | ECH1 | enoyl CoA hydratase 1, peroxisomal | 1891 | ENSG00000104823 |
| 9 | P38117 | NA | ETFB | electron-transfer-flavoprotein, beta polypeptide | 2109 | ENSG00000105379 |
| 10 | Q16698 | NA | DECR1 | 2,4-dienoyl CoA reductase 1, mitochondrial | 1666 | ENSG00000104325 |
| 11 | Q99798 | NA | ACO2 | aconitase 2, mitochondrial | 50 | ENSG00000100412 |
| 12 | P42765 | NA | ACAA2 | acetyl-CoA acyltransferase 2 | 10449 | ENSG00000167315 |
| 13 | Q10713 | NA | PMPCA | peptidase (mitochondrial processing) alpha | 23203 | ENSG00000165688 |
| 14 | P13674 | NA | P4HA1 | prolyl 4-hydroxylase, alpha polypeptide I | 5033 | ENSG00000122884 |
| 15 | P49419 | NA | ALDH7A1 | aldehyde dehydrogenase 7 family, member A1 | 501 | ENSG00000164904 |
| 16 | P21980 | NA | TGM2 | transglutaminase 2 (C polypeptide, protein-glutamine-gamma-glutamyltransferase) | 7052 | ENSG00000198959 |
| 17 | P19367 | NA | HK1 | hexokinase 1 | 3098 | ENSG00000156515 |
| 18 | Q9BQ69 | NA | MACROD1 | MACRO domain containing 1 | 28992 | ENSG00000133315 |

  
  

| **Database:cellular component      &nbspName:mitochondrial part      &nbspID:GO:0044429** | | | | | | |
| --- | --- | --- | --- | --- | --- | --- |
| C=740; O=13; E=2.00; R=6.51; rawP=4.54e-08; adjP=6.96e-07 | | | | | | |
| Index | UserID | Value | Gene Symbol | Gene Name | EntrezGene | Ensembl |
| 1 | Q99798 | NA | ACO2 | aconitase 2, mitochondrial | 50 | ENSG00000100412 |
| 2 | Q16698 | NA | DECR1 | 2,4-dienoyl CoA reductase 1, mitochondrial | 1666 | ENSG00000104325 |
| 3 | P54819 | NA | AK2 | adenylate kinase 2 | 204 | ENSG00000004455 |
| 4 | P17931 | NA | LGALS3 | lectin, galactoside-binding, soluble, 3 | 3958 | ENSG00000131981 |
| 5 | Q10713 | NA | PMPCA | peptidase (mitochondrial processing) alpha | 23203 | ENSG00000165688 |
| 6 | P42765 | NA | ACAA2 | acetyl-CoA acyltransferase 2 | 10449 | ENSG00000167315 |
| 7 | Q99714 | NA | HSD17B10 | hydroxysteroid (17-beta) dehydrogenase 10 | 3028 | ENSG00000072506 |
| 8 | P04179 | NA | SOD2 | superoxide dismutase 2, mitochondrial | 6648 | ENSG00000112096 |
| 9 | P49419 | NA | ALDH7A1 | aldehyde dehydrogenase 7 family, member A1 | 501 | ENSG00000164904 |
| 10 | P19367 | NA | HK1 | hexokinase 1 | 3098 | ENSG00000156515 |
| 11 | P30837 | NA | ALDH1B1 | aldehyde dehydrogenase 1 family, member B1 | 219 | ENSG00000137124 |
| 12 | P31040 | NA | SDHA | succinate dehydrogenase complex, subunit A, flavoprotein (Fp) | 6389 | ENSG00000073578 |
| 13 | P38117 | NA | ETFB | electron-transfer-flavoprotein, beta polypeptide | 2109 | ENSG00000105379 |

  
  

| **Database:cellular component      &nbspName:cytoplasmic part      &nbspID:GO:0044444** | | | | | | |
| --- | --- | --- | --- | --- | --- | --- |
| C=6728; O=36; E=18.15; R=1.98; rawP=6.18e-08; adjP=7.11e-07 | | | | | | |
| Index | UserID | Value | Gene Symbol | Gene Name | EntrezGene | Ensembl |
| 1 | P10253 | NA | GAA | glucosidase, alpha; acid | 2548 | ENSG00000171298 |
| 2 | P54819 | NA | AK2 | adenylate kinase 2 | 204 | ENSG00000004455 |
| 3 | P17931 | NA | LGALS3 | lectin, galactoside-binding, soluble, 3 | 3958 | ENSG00000131981 |
| 4 | Q9UL46 | NA | PSME2 | proteasome (prosome, macropain) activator subunit 2 (PA28 beta) | 5721 | ENSG00000100911 |
| 5 | P50453 | NA | SERPINB9 | serpin peptidase inhibitor, clade B (ovalbumin), member 9 | 5272 | ENSG00000170542 |
| 6 | Q99714 | NA | HSD17B10 | hydroxysteroid (17-beta) dehydrogenase 10 | 3028 | ENSG00000072506 |
| 7 | P04179 | NA | SOD2 | superoxide dismutase 2, mitochondrial | 6648 | ENSG00000112096 |
| 8 | Q06323 | NA | PSME1 | proteasome (prosome, macropain) activator subunit 1 (PA28 alpha) | 5720 | ENSG00000092010 |
| 9 | P07741 | NA | APRT | adenine phosphoribosyltransferase | 353 | ENSG00000198931 |
| 10 | P31040 | NA | SDHA | succinate dehydrogenase complex, subunit A, flavoprotein (Fp) | 6389 | ENSG00000073578 |
| 11 | P38117 | NA | ETFB | electron-transfer-flavoprotein, beta polypeptide | 2109 | ENSG00000105379 |
| 12 | Q16698 | NA | DECR1 | 2,4-dienoyl CoA reductase 1, mitochondrial | 1666 | ENSG00000104325 |
| 13 | Q05639 | NA | EEF1A2 | eukaryotic translation elongation factor 1 alpha 2 | 1917 | ENSG00000101210 |
| 14 | P42765 | NA | ACAA2 | acetyl-CoA acyltransferase 2 | 10449 | ENSG00000167315 |
| 15 | Q10713 | NA | PMPCA | peptidase (mitochondrial processing) alpha | 23203 | ENSG00000165688 |
| 16 | P39687 | NA | ANP32A | acidic (leucine-rich) nuclear phosphoprotein 32 family, member A | 8125 | ENSG00000140350 |
| 17 | P49419 | NA | ALDH7A1 | aldehyde dehydrogenase 7 family, member A1 | 501 | ENSG00000164904 |
| 18 | P21980 | NA | TGM2 | transglutaminase 2 (C polypeptide, protein-glutamine-gamma-glutamyltransferase) | 7052 | ENSG00000198959 |
| 19 | Q9UDT6 | NA | CLIP2 | CAP-GLY domain containing linker protein 2 | 7461 | ENSG00000106665 |
| 20 | P02792 | NA | FTL | ferritin, light polypeptide | 2512 | ENSG00000087086 |
| 21 | P26447 | NA | S100A4 | S100 calcium binding protein A4 | 6275 | ENSG00000196154 |
| 22 | P06737 | NA | PYGL | phosphorylase, glycogen, liver | 5836 | ENSG00000100504 |
| 23 | Q9BXW7 | NA | CECR5 | cat eye syndrome chromosome region, candidate 5 | 27440 | ENSG00000069998 |
| 24 | P30837 | NA | ALDH1B1 | aldehyde dehydrogenase 1 family, member B1 | 219 | ENSG00000137124 |
| 25 | P38571 | NA | LIPA | lipase A, lysosomal acid, cholesterol esterase | 3988 | ENSG00000107798 |
| 26 | Q13011 | NA | ECH1 | enoyl CoA hydratase 1, peroxisomal | 1891 | ENSG00000104823 |
| 27 | Q99798 | NA | ACO2 | aconitase 2, mitochondrial | 50 | ENSG00000100412 |
| 28 | Q9UBQ7 | NA | GRHPR | glyoxylate reductase/hydroxypyruvate reductase | 9380 | ENSG00000137106 |
| 29 | P00558 | NA | PGK1 | phosphoglycerate kinase 1 | 5230 | ENSG00000102144 |
| 30 | Q01105 | NA | SET | SET nuclear oncogene | 6418 | ENSG00000119335 |
| 31 | P13674 | NA | P4HA1 | prolyl 4-hydroxylase, alpha polypeptide I | 5033 | ENSG00000122884 |
| 32 | P06744 | NA | GPI | glucose-6-phosphate isomerase | 2821 | ENSG00000105220 |
| 33 | P19367 | NA | HK1 | hexokinase 1 | 3098 | ENSG00000156515 |
| 34 | Q9BQ69 | NA | MACROD1 | MACRO domain containing 1 | 28992 | ENSG00000133315 |
| 35 | P05783 | NA | KRT18 | keratin 18 | 3875 | ENSG00000111057 |
| 36 | P00352 | NA | ALDH1A1 | aldehyde dehydrogenase 1 family, member A1 | 216 | ENSG00000165092 |

  
  

| **Database:cellular component      &nbspName:membrane-enclosed lumen      &nbspID:GO:0031974** | | | | | | |
| --- | --- | --- | --- | --- | --- | --- |
| C=3375; O=25; E=9.10; R=2.75; rawP=1.82e-07; adjP=1.67e-06 | | | | | | |
| Index | UserID | Value | Gene Symbol | Gene Name | EntrezGene | Ensembl |
| 1 | P09661 | NA | SNRPA1 | small nuclear ribonucleoprotein polypeptide A' | 6627 | ENSG00000131876 |
| 2 | P54819 | NA | AK2 | adenylate kinase 2 | 204 | ENSG00000004455 |
| 3 | P05787 | NA | KRT8 | keratin 8 | 3856 | ENSG00000170421 |
| 4 | Q9UL46 | NA | PSME2 | proteasome (prosome, macropain) activator subunit 2 (PA28 beta) | 5721 | ENSG00000100911 |
| 5 | Q99714 | NA | HSD17B10 | hydroxysteroid (17-beta) dehydrogenase 10 | 3028 | ENSG00000072506 |
| 6 | P04179 | NA | SOD2 | superoxide dismutase 2, mitochondrial | 6648 | ENSG00000112096 |
| 7 | P07741 | NA | APRT | adenine phosphoribosyltransferase | 353 | ENSG00000198931 |
| 8 | Q06323 | NA | PSME1 | proteasome (prosome, macropain) activator subunit 1 (PA28 alpha) | 5720 | ENSG00000092010 |
| 9 | P30837 | NA | ALDH1B1 | aldehyde dehydrogenase 1 family, member B1 | 219 | ENSG00000137124 |
| 10 | Q13263 | NA | TRIM28 | tripartite motif containing 28 | 10155 | ENSG00000130726 |
| 11 | P38117 | NA | ETFB | electron-transfer-flavoprotein, beta polypeptide | 2109 | ENSG00000105379 |
| 12 | Q9UBQ7 | NA | GRHPR | glyoxylate reductase/hydroxypyruvate reductase | 9380 | ENSG00000137106 |
| 13 | Q16698 | NA | DECR1 | 2,4-dienoyl CoA reductase 1, mitochondrial | 1666 | ENSG00000104325 |
| 14 | Q99798 | NA | ACO2 | aconitase 2, mitochondrial | 50 | ENSG00000100412 |
| 15 | Q01105 | NA | SET | SET nuclear oncogene | 6418 | ENSG00000119335 |
| 16 | P54727 | NA | RAD23B | RAD23 homolog B (S. cerevisiae) | 5887 | ENSG00000119318 |
| 17 | Q10713 | NA | PMPCA | peptidase (mitochondrial processing) alpha | 23203 | ENSG00000165688 |
| 18 | P13674 | NA | P4HA1 | prolyl 4-hydroxylase, alpha polypeptide I | 5033 | ENSG00000122884 |
| 19 | P39687 | NA | ANP32A | acidic (leucine-rich) nuclear phosphoprotein 32 family, member A | 8125 | ENSG00000140350 |
| 20 | O43598 | NA | C6orf108 | chromosome 6 open reading frame 108 | 10591 | ENSG00000112667 |
| 21 | P49419 | NA | ALDH7A1 | aldehyde dehydrogenase 7 family, member A1 | 501 | ENSG00000164904 |
| 22 | P06744 | NA | GPI | glucose-6-phosphate isomerase | 2821 | ENSG00000105220 |
| 23 | P19367 | NA | HK1 | hexokinase 1 | 3098 | ENSG00000156515 |
| 24 | P05783 | NA | KRT18 | keratin 18 | 3875 | ENSG00000111057 |
| 25 | Q16531 | NA | DDB1 | damage-specific DNA binding protein 1, 127kDa | 1642 | ENSG00000167986 |

  
  

| **Database:cellular component      &nbspName:intracellular organelle lumen      &nbspID:GO:0070013** | | | | | | |
| --- | --- | --- | --- | --- | --- | --- |
| C=3285; O=24; E=8.86; R=2.71; rawP=5.24e-07; adjP=3.44e-06 | | | | | | |
| Index | UserID | Value | Gene Symbol | Gene Name | EntrezGene | Ensembl |
| 1 | P09661 | NA | SNRPA1 | small nuclear ribonucleoprotein polypeptide A' | 6627 | ENSG00000131876 |
| 2 | P05787 | NA | KRT8 | keratin 8 | 3856 | ENSG00000170421 |
| 3 | Q9UL46 | NA | PSME2 | proteasome (prosome, macropain) activator subunit 2 (PA28 beta) | 5721 | ENSG00000100911 |
| 4 | Q99714 | NA | HSD17B10 | hydroxysteroid (17-beta) dehydrogenase 10 | 3028 | ENSG00000072506 |
| 5 | P04179 | NA | SOD2 | superoxide dismutase 2, mitochondrial | 6648 | ENSG00000112096 |
| 6 | P07741 | NA | APRT | adenine phosphoribosyltransferase | 353 | ENSG00000198931 |
| 7 | Q06323 | NA | PSME1 | proteasome (prosome, macropain) activator subunit 1 (PA28 alpha) | 5720 | ENSG00000092010 |
| 8 | P30837 | NA | ALDH1B1 | aldehyde dehydrogenase 1 family, member B1 | 219 | ENSG00000137124 |
| 9 | Q13263 | NA | TRIM28 | tripartite motif containing 28 | 10155 | ENSG00000130726 |
| 10 | P38117 | NA | ETFB | electron-transfer-flavoprotein, beta polypeptide | 2109 | ENSG00000105379 |
| 11 | Q9UBQ7 | NA | GRHPR | glyoxylate reductase/hydroxypyruvate reductase | 9380 | ENSG00000137106 |
| 12 | Q16698 | NA | DECR1 | 2,4-dienoyl CoA reductase 1, mitochondrial | 1666 | ENSG00000104325 |
| 13 | Q99798 | NA | ACO2 | aconitase 2, mitochondrial | 50 | ENSG00000100412 |
| 14 | Q01105 | NA | SET | SET nuclear oncogene | 6418 | ENSG00000119335 |
| 15 | P54727 | NA | RAD23B | RAD23 homolog B (S. cerevisiae) | 5887 | ENSG00000119318 |
| 16 | Q10713 | NA | PMPCA | peptidase (mitochondrial processing) alpha | 23203 | ENSG00000165688 |
| 17 | P13674 | NA | P4HA1 | prolyl 4-hydroxylase, alpha polypeptide I | 5033 | ENSG00000122884 |
| 18 | P39687 | NA | ANP32A | acidic (leucine-rich) nuclear phosphoprotein 32 family, member A | 8125 | ENSG00000140350 |
| 19 | O43598 | NA | C6orf108 | chromosome 6 open reading frame 108 | 10591 | ENSG00000112667 |
| 20 | P49419 | NA | ALDH7A1 | aldehyde dehydrogenase 7 family, member A1 | 501 | ENSG00000164904 |
| 21 | P06744 | NA | GPI | glucose-6-phosphate isomerase | 2821 | ENSG00000105220 |
| 22 | P19367 | NA | HK1 | hexokinase 1 | 3098 | ENSG00000156515 |
| 23 | P05783 | NA | KRT18 | keratin 18 | 3875 | ENSG00000111057 |
| 24 | Q16531 | NA | DDB1 | damage-specific DNA binding protein 1, 127kDa | 1642 | ENSG00000167986 |

  
  

| **Database:cellular component      &nbspName:intracellular part      &nbspID:GO:0044424** | | | | | | |
| --- | --- | --- | --- | --- | --- | --- |
| C=12096; O=45; E=32.63; R=1.38; rawP=5.10e-07; adjP=3.44e-06 | | | | | | |
| Index | UserID | Value | Gene Symbol | Gene Name | EntrezGene | Ensembl |
| 1 | P09661 | NA | SNRPA1 | small nuclear ribonucleoprotein polypeptide A' | 6627 | ENSG00000131876 |
| 2 | P10253 | NA | GAA | glucosidase, alpha; acid | 2548 | ENSG00000171298 |
| 3 | P54819 | NA | AK2 | adenylate kinase 2 | 204 | ENSG00000004455 |
| 4 | P17931 | NA | LGALS3 | lectin, galactoside-binding, soluble, 3 | 3958 | ENSG00000131981 |
| 5 | P05787 | NA | KRT8 | keratin 8 | 3856 | ENSG00000170421 |
| 6 | Q9UL46 | NA | PSME2 | proteasome (prosome, macropain) activator subunit 2 (PA28 beta) | 5721 | ENSG00000100911 |
| 7 | P50453 | NA | SERPINB9 | serpin peptidase inhibitor, clade B (ovalbumin), member 9 | 5272 | ENSG00000170542 |
| 8 | Q99714 | NA | HSD17B10 | hydroxysteroid (17-beta) dehydrogenase 10 | 3028 | ENSG00000072506 |
| 9 | P04179 | NA | SOD2 | superoxide dismutase 2, mitochondrial | 6648 | ENSG00000112096 |
| 10 | Q06323 | NA | PSME1 | proteasome (prosome, macropain) activator subunit 1 (PA28 alpha) | 5720 | ENSG00000092010 |
| 11 | P07741 | NA | APRT | adenine phosphoribosyltransferase | 353 | ENSG00000198931 |
| 12 | P30046 | NA | DDT | D-dopachrome tautomerase | 1652 | ENSG00000099977 |
| 13 | P31040 | NA | SDHA | succinate dehydrogenase complex, subunit A, flavoprotein (Fp) | 6389 | ENSG00000073578 |
| 14 | P38117 | NA | ETFB | electron-transfer-flavoprotein, beta polypeptide | 2109 | ENSG00000105379 |
| 15 | Q16698 | NA | DECR1 | 2,4-dienoyl CoA reductase 1, mitochondrial | 1666 | ENSG00000104325 |
| 16 | P54727 | NA | RAD23B | RAD23 homolog B (S. cerevisiae) | 5887 | ENSG00000119318 |
| 17 | Q05639 | NA | EEF1A2 | eukaryotic translation elongation factor 1 alpha 2 | 1917 | ENSG00000101210 |
| 18 | Q10713 | NA | PMPCA | peptidase (mitochondrial processing) alpha | 23203 | ENSG00000165688 |
| 19 | P42765 | NA | ACAA2 | acetyl-CoA acyltransferase 2 | 10449 | ENSG00000167315 |
| 20 | P39687 | NA | ANP32A | acidic (leucine-rich) nuclear phosphoprotein 32 family, member A | 8125 | ENSG00000140350 |
| 21 | P49419 | NA | ALDH7A1 | aldehyde dehydrogenase 7 family, member A1 | 501 | ENSG00000164904 |
| 22 | P21980 | NA | TGM2 | transglutaminase 2 (C polypeptide, protein-glutamine-gamma-glutamyltransferase) | 7052 | ENSG00000198959 |
| 23 | Q9UDT6 | NA | CLIP2 | CAP-GLY domain containing linker protein 2 | 7461 | ENSG00000106665 |
| 24 | Q5SSJ5 | NA | HP1BP3 | heterochromatin protein 1, binding protein 3 | 50809 | ENSG00000127483 |
| 25 | P02792 | NA | FTL | ferritin, light polypeptide | 2512 | ENSG00000087086 |
| 26 | P26447 | NA | S100A4 | S100 calcium binding protein A4 | 6275 | ENSG00000196154 |
| 27 | P06737 | NA | PYGL | phosphorylase, glycogen, liver | 5836 | ENSG00000100504 |
| 28 | Q9BXW7 | NA | CECR5 | cat eye syndrome chromosome region, candidate 5 | 27440 | ENSG00000069998 |
| 29 | P30837 | NA | ALDH1B1 | aldehyde dehydrogenase 1 family, member B1 | 219 | ENSG00000137124 |
| 30 | P38571 | NA | LIPA | lipase A, lysosomal acid, cholesterol esterase | 3988 | ENSG00000107798 |
| 31 | Q13263 | NA | TRIM28 | tripartite motif containing 28 | 10155 | ENSG00000130726 |
| 32 | P32119 | NA | PRDX2 | peroxiredoxin 2 | 7001 | ENSG00000167815 |
| 33 | Q13011 | NA | ECH1 | enoyl CoA hydratase 1, peroxisomal | 1891 | ENSG00000104823 |
| 34 | Q9UBQ7 | NA | GRHPR | glyoxylate reductase/hydroxypyruvate reductase | 9380 | ENSG00000137106 |
| 35 | Q99798 | NA | ACO2 | aconitase 2, mitochondrial | 50 | ENSG00000100412 |
| 36 | P00558 | NA | PGK1 | phosphoglycerate kinase 1 | 5230 | ENSG00000102144 |
| 37 | Q01105 | NA | SET | SET nuclear oncogene | 6418 | ENSG00000119335 |
| 38 | P13674 | NA | P4HA1 | prolyl 4-hydroxylase, alpha polypeptide I | 5033 | ENSG00000122884 |
| 39 | O43598 | NA | C6orf108 | chromosome 6 open reading frame 108 | 10591 | ENSG00000112667 |
| 40 | P06744 | NA | GPI | glucose-6-phosphate isomerase | 2821 | ENSG00000105220 |
| 41 | P19367 | NA | HK1 | hexokinase 1 | 3098 | ENSG00000156515 |
| 42 | Q9BQ69 | NA | MACROD1 | MACRO domain containing 1 | 28992 | ENSG00000133315 |
| 43 | P05783 | NA | KRT18 | keratin 18 | 3875 | ENSG00000111057 |
| 44 | P00352 | NA | ALDH1A1 | aldehyde dehydrogenase 1 family, member A1 | 216 | ENSG00000165092 |
| 45 | Q16531 | NA | DDB1 | damage-specific DNA binding protein 1, 127kDa | 1642 | ENSG00000167986 |

  
  

| **Database:cellular component      &nbspName:organelle lumen      &nbspID:GO:0043233** | | | | | | |
| --- | --- | --- | --- | --- | --- | --- |
| C=3331; O=24; E=8.99; R=2.67; rawP=6.84e-07; adjP=3.50e-06 | | | | | | |
| Index | UserID | Value | Gene Symbol | Gene Name | EntrezGene | Ensembl |
| 1 | P09661 | NA | SNRPA1 | small nuclear ribonucleoprotein polypeptide A' | 6627 | ENSG00000131876 |
| 2 | P05787 | NA | KRT8 | keratin 8 | 3856 | ENSG00000170421 |
| 3 | Q9UL46 | NA | PSME2 | proteasome (prosome, macropain) activator subunit 2 (PA28 beta) | 5721 | ENSG00000100911 |
| 4 | Q99714 | NA | HSD17B10 | hydroxysteroid (17-beta) dehydrogenase 10 | 3028 | ENSG00000072506 |
| 5 | P04179 | NA | SOD2 | superoxide dismutase 2, mitochondrial | 6648 | ENSG00000112096 |
| 6 | P07741 | NA | APRT | adenine phosphoribosyltransferase | 353 | ENSG00000198931 |
| 7 | Q06323 | NA | PSME1 | proteasome (prosome, macropain) activator subunit 1 (PA28 alpha) | 5720 | ENSG00000092010 |
| 8 | P30837 | NA | ALDH1B1 | aldehyde dehydrogenase 1 family, member B1 | 219 | ENSG00000137124 |
| 9 | Q13263 | NA | TRIM28 | tripartite motif containing 28 | 10155 | ENSG00000130726 |
| 10 | P38117 | NA | ETFB | electron-transfer-flavoprotein, beta polypeptide | 2109 | ENSG00000105379 |
| 11 | Q9UBQ7 | NA | GRHPR | glyoxylate reductase/hydroxypyruvate reductase | 9380 | ENSG00000137106 |
| 12 | Q16698 | NA | DECR1 | 2,4-dienoyl CoA reductase 1, mitochondrial | 1666 | ENSG00000104325 |
| 13 | Q99798 | NA | ACO2 | aconitase 2, mitochondrial | 50 | ENSG00000100412 |
| 14 | Q01105 | NA | SET | SET nuclear oncogene | 6418 | ENSG00000119335 |
| 15 | P54727 | NA | RAD23B | RAD23 homolog B (S. cerevisiae) | 5887 | ENSG00000119318 |
| 16 | Q10713 | NA | PMPCA | peptidase (mitochondrial processing) alpha | 23203 | ENSG00000165688 |
| 17 | P13674 | NA | P4HA1 | prolyl 4-hydroxylase, alpha polypeptide I | 5033 | ENSG00000122884 |
| 18 | P39687 | NA | ANP32A | acidic (leucine-rich) nuclear phosphoprotein 32 family, member A | 8125 | ENSG00000140350 |
| 19 | O43598 | NA | C6orf108 | chromosome 6 open reading frame 108 | 10591 | ENSG00000112667 |
| 20 | P49419 | NA | ALDH7A1 | aldehyde dehydrogenase 7 family, member A1 | 501 | ENSG00000164904 |
| 21 | P06744 | NA | GPI | glucose-6-phosphate isomerase | 2821 | ENSG00000105220 |
| 22 | P19367 | NA | HK1 | hexokinase 1 | 3098 | ENSG00000156515 |
| 23 | P05783 | NA | KRT18 | keratin 18 | 3875 | ENSG00000111057 |
| 24 | Q16531 | NA | DDB1 | damage-specific DNA binding protein 1, 127kDa | 1642 | ENSG00000167986 |

  
  

| **Database:cellular component      &nbspName:mitochondrial matrix      &nbspID:GO:0005759** | | | | | | |
| --- | --- | --- | --- | --- | --- | --- |
| C=278; O=8; E=0.75; R=10.67; rawP=6.79e-07; adjP=3.50e-06 | | | | | | |
| Index | UserID | Value | Gene Symbol | Gene Name | EntrezGene | Ensembl |
| 1 | Q99798 | NA | ACO2 | aconitase 2, mitochondrial | 50 | ENSG00000100412 |
| 2 | Q16698 | NA | DECR1 | 2,4-dienoyl CoA reductase 1, mitochondrial | 1666 | ENSG00000104325 |
| 3 | Q10713 | NA | PMPCA | peptidase (mitochondrial processing) alpha | 23203 | ENSG00000165688 |
| 4 | Q99714 | NA | HSD17B10 | hydroxysteroid (17-beta) dehydrogenase 10 | 3028 | ENSG00000072506 |
| 5 | P04179 | NA | SOD2 | superoxide dismutase 2, mitochondrial | 6648 | ENSG00000112096 |
| 6 | P49419 | NA | ALDH7A1 | aldehyde dehydrogenase 7 family, member A1 | 501 | ENSG00000164904 |
| 7 | P30837 | NA | ALDH1B1 | aldehyde dehydrogenase 1 family, member B1 | 219 | ENSG00000137124 |
| 8 | P38117 | NA | ETFB | electron-transfer-flavoprotein, beta polypeptide | 2109 | ENSG00000105379 |

  
  

| **Database:cellular component      &nbspName:intracellular      &nbspID:GO:0005622** | | | | | | |
| --- | --- | --- | --- | --- | --- | --- |
| C=12412; O=45; E=33.48; R=1.34; rawP=1.63e-06; adjP=7.50e-06 | | | | | | |
| Index | UserID | Value | Gene Symbol | Gene Name | EntrezGene | Ensembl |
| 1 | P09661 | NA | SNRPA1 | small nuclear ribonucleoprotein polypeptide A' | 6627 | ENSG00000131876 |
| 2 | P10253 | NA | GAA | glucosidase, alpha; acid | 2548 | ENSG00000171298 |
| 3 | P54819 | NA | AK2 | adenylate kinase 2 | 204 | ENSG00000004455 |
| 4 | P17931 | NA | LGALS3 | lectin, galactoside-binding, soluble, 3 | 3958 | ENSG00000131981 |
| 5 | P05787 | NA | KRT8 | keratin 8 | 3856 | ENSG00000170421 |
| 6 | Q9UL46 | NA | PSME2 | proteasome (prosome, macropain) activator subunit 2 (PA28 beta) | 5721 | ENSG00000100911 |
| 7 | P50453 | NA | SERPINB9 | serpin peptidase inhibitor, clade B (ovalbumin), member 9 | 5272 | ENSG00000170542 |
| 8 | Q99714 | NA | HSD17B10 | hydroxysteroid (17-beta) dehydrogenase 10 | 3028 | ENSG00000072506 |
| 9 | P04179 | NA | SOD2 | superoxide dismutase 2, mitochondrial | 6648 | ENSG00000112096 |
| 10 | Q06323 | NA | PSME1 | proteasome (prosome, macropain) activator subunit 1 (PA28 alpha) | 5720 | ENSG00000092010 |
| 11 | P07741 | NA | APRT | adenine phosphoribosyltransferase | 353 | ENSG00000198931 |
| 12 | P30046 | NA | DDT | D-dopachrome tautomerase | 1652 | ENSG00000099977 |
| 13 | P31040 | NA | SDHA | succinate dehydrogenase complex, subunit A, flavoprotein (Fp) | 6389 | ENSG00000073578 |
| 14 | P38117 | NA | ETFB | electron-transfer-flavoprotein, beta polypeptide | 2109 | ENSG00000105379 |
| 15 | Q16698 | NA | DECR1 | 2,4-dienoyl CoA reductase 1, mitochondrial | 1666 | ENSG00000104325 |
| 16 | P54727 | NA | RAD23B | RAD23 homolog B (S. cerevisiae) | 5887 | ENSG00000119318 |
| 17 | Q05639 | NA | EEF1A2 | eukaryotic translation elongation factor 1 alpha 2 | 1917 | ENSG00000101210 |
| 18 | Q10713 | NA | PMPCA | peptidase (mitochondrial processing) alpha | 23203 | ENSG00000165688 |
| 19 | P42765 | NA | ACAA2 | acetyl-CoA acyltransferase 2 | 10449 | ENSG00000167315 |
| 20 | P39687 | NA | ANP32A | acidic (leucine-rich) nuclear phosphoprotein 32 family, member A | 8125 | ENSG00000140350 |
| 21 | P49419 | NA | ALDH7A1 | aldehyde dehydrogenase 7 family, member A1 | 501 | ENSG00000164904 |
| 22 | P21980 | NA | TGM2 | transglutaminase 2 (C polypeptide, protein-glutamine-gamma-glutamyltransferase) | 7052 | ENSG00000198959 |
| 23 | Q9UDT6 | NA | CLIP2 | CAP-GLY domain containing linker protein 2 | 7461 | ENSG00000106665 |
| 24 | Q5SSJ5 | NA | HP1BP3 | heterochromatin protein 1, binding protein 3 | 50809 | ENSG00000127483 |
| 25 | P02792 | NA | FTL | ferritin, light polypeptide | 2512 | ENSG00000087086 |
| 26 | P26447 | NA | S100A4 | S100 calcium binding protein A4 | 6275 | ENSG00000196154 |
| 27 | P06737 | NA | PYGL | phosphorylase, glycogen, liver | 5836 | ENSG00000100504 |
| 28 | Q9BXW7 | NA | CECR5 | cat eye syndrome chromosome region, candidate 5 | 27440 | ENSG00000069998 |
| 29 | P30837 | NA | ALDH1B1 | aldehyde dehydrogenase 1 family, member B1 | 219 | ENSG00000137124 |
| 30 | P38571 | NA | LIPA | lipase A, lysosomal acid, cholesterol esterase | 3988 | ENSG00000107798 |
| 31 | Q13263 | NA | TRIM28 | tripartite motif containing 28 | 10155 | ENSG00000130726 |
| 32 | P32119 | NA | PRDX2 | peroxiredoxin 2 | 7001 | ENSG00000167815 |
| 33 | Q13011 | NA | ECH1 | enoyl CoA hydratase 1, peroxisomal | 1891 | ENSG00000104823 |
| 34 | Q9UBQ7 | NA | GRHPR | glyoxylate reductase/hydroxypyruvate reductase | 9380 | ENSG00000137106 |
| 35 | Q99798 | NA | ACO2 | aconitase 2, mitochondrial | 50 | ENSG00000100412 |
| 36 | P00558 | NA | PGK1 | phosphoglycerate kinase 1 | 5230 | ENSG00000102144 |
| 37 | Q01105 | NA | SET | SET nuclear oncogene | 6418 | ENSG00000119335 |
| 38 | P13674 | NA | P4HA1 | prolyl 4-hydroxylase, alpha polypeptide I | 5033 | ENSG00000122884 |
| 39 | O43598 | NA | C6orf108 | chromosome 6 open reading frame 108 | 10591 | ENSG00000112667 |
| 40 | P06744 | NA | GPI | glucose-6-phosphate isomerase | 2821 | ENSG00000105220 |
| 41 | P19367 | NA | HK1 | hexokinase 1 | 3098 | ENSG00000156515 |
| 42 | Q9BQ69 | NA | MACROD1 | MACRO domain containing 1 | 28992 | ENSG00000133315 |
| 43 | P05783 | NA | KRT18 | keratin 18 | 3875 | ENSG00000111057 |
| 44 | P00352 | NA | ALDH1A1 | aldehyde dehydrogenase 1 family, member A1 | 216 | ENSG00000165092 |
| 45 | Q16531 | NA | DDB1 | damage-specific DNA binding protein 1, 127kDa | 1642 | ENSG00000167986 |

  
  

| **Database:cellular component      &nbspName:membrane-bounded organelle      &nbspID:GO:0043227** | | | | | | |
| --- | --- | --- | --- | --- | --- | --- |
| C=9495; O=38; E=25.61; R=1.48; rawP=7.94e-05; adjP=0.0002 | | | | | | |
| Index | UserID | Value | Gene Symbol | Gene Name | EntrezGene | Ensembl |
| 1 | P09661 | NA | SNRPA1 | small nuclear ribonucleoprotein polypeptide A' | 6627 | ENSG00000131876 |
| 2 | P10253 | NA | GAA | glucosidase, alpha; acid | 2548 | ENSG00000171298 |
| 3 | P54819 | NA | AK2 | adenylate kinase 2 | 204 | ENSG00000004455 |
| 4 | P17931 | NA | LGALS3 | lectin, galactoside-binding, soluble, 3 | 3958 | ENSG00000131981 |
| 5 | P05787 | NA | KRT8 | keratin 8 | 3856 | ENSG00000170421 |
| 6 | Q9UL46 | NA | PSME2 | proteasome (prosome, macropain) activator subunit 2 (PA28 beta) | 5721 | ENSG00000100911 |
| 7 | P50453 | NA | SERPINB9 | serpin peptidase inhibitor, clade B (ovalbumin), member 9 | 5272 | ENSG00000170542 |
| 8 | Q99714 | NA | HSD17B10 | hydroxysteroid (17-beta) dehydrogenase 10 | 3028 | ENSG00000072506 |
| 9 | P04179 | NA | SOD2 | superoxide dismutase 2, mitochondrial | 6648 | ENSG00000112096 |
| 10 | Q06323 | NA | PSME1 | proteasome (prosome, macropain) activator subunit 1 (PA28 alpha) | 5720 | ENSG00000092010 |
| 11 | P07741 | NA | APRT | adenine phosphoribosyltransferase | 353 | ENSG00000198931 |
| 12 | P31040 | NA | SDHA | succinate dehydrogenase complex, subunit A, flavoprotein (Fp) | 6389 | ENSG00000073578 |
| 13 | P38117 | NA | ETFB | electron-transfer-flavoprotein, beta polypeptide | 2109 | ENSG00000105379 |
| 14 | Q16698 | NA | DECR1 | 2,4-dienoyl CoA reductase 1, mitochondrial | 1666 | ENSG00000104325 |
| 15 | Q05639 | NA | EEF1A2 | eukaryotic translation elongation factor 1 alpha 2 | 1917 | ENSG00000101210 |
| 16 | P54727 | NA | RAD23B | RAD23 homolog B (S. cerevisiae) | 5887 | ENSG00000119318 |
| 17 | Q10713 | NA | PMPCA | peptidase (mitochondrial processing) alpha | 23203 | ENSG00000165688 |
| 18 | P42765 | NA | ACAA2 | acetyl-CoA acyltransferase 2 | 10449 | ENSG00000167315 |
| 19 | P39687 | NA | ANP32A | acidic (leucine-rich) nuclear phosphoprotein 32 family, member A | 8125 | ENSG00000140350 |
| 20 | P49419 | NA | ALDH7A1 | aldehyde dehydrogenase 7 family, member A1 | 501 | ENSG00000164904 |
| 21 | P21980 | NA | TGM2 | transglutaminase 2 (C polypeptide, protein-glutamine-gamma-glutamyltransferase) | 7052 | ENSG00000198959 |
| 22 | Q5SSJ5 | NA | HP1BP3 | heterochromatin protein 1, binding protein 3 | 50809 | ENSG00000127483 |
| 23 | P26447 | NA | S100A4 | S100 calcium binding protein A4 | 6275 | ENSG00000196154 |
| 24 | Q9BXW7 | NA | CECR5 | cat eye syndrome chromosome region, candidate 5 | 27440 | ENSG00000069998 |
| 25 | P30837 | NA | ALDH1B1 | aldehyde dehydrogenase 1 family, member B1 | 219 | ENSG00000137124 |
| 26 | P38571 | NA | LIPA | lipase A, lysosomal acid, cholesterol esterase | 3988 | ENSG00000107798 |
| 27 | Q13263 | NA | TRIM28 | tripartite motif containing 28 | 10155 | ENSG00000130726 |
| 28 | Q13011 | NA | ECH1 | enoyl CoA hydratase 1, peroxisomal | 1891 | ENSG00000104823 |
| 29 | Q99798 | NA | ACO2 | aconitase 2, mitochondrial | 50 | ENSG00000100412 |
| 30 | Q9UBQ7 | NA | GRHPR | glyoxylate reductase/hydroxypyruvate reductase | 9380 | ENSG00000137106 |
| 31 | Q01105 | NA | SET | SET nuclear oncogene | 6418 | ENSG00000119335 |
| 32 | P13674 | NA | P4HA1 | prolyl 4-hydroxylase, alpha polypeptide I | 5033 | ENSG00000122884 |
| 33 | O43598 | NA | C6orf108 | chromosome 6 open reading frame 108 | 10591 | ENSG00000112667 |
| 34 | P06744 | NA | GPI | glucose-6-phosphate isomerase | 2821 | ENSG00000105220 |
| 35 | P19367 | NA | HK1 | hexokinase 1 | 3098 | ENSG00000156515 |
| 36 | Q9BQ69 | NA | MACROD1 | MACRO domain containing 1 | 28992 | ENSG00000133315 |
| 37 | P05783 | NA | KRT18 | keratin 18 | 3875 | ENSG00000111057 |
| 38 | Q16531 | NA | DDB1 | damage-specific DNA binding protein 1, 127kDa | 1642 | ENSG00000167986 |

  
  

| **Database:cellular component      &nbspName:mitochondrial membrane      &nbspID:GO:0031966** | | | | | | |
| --- | --- | --- | --- | --- | --- | --- |
| C=505; O=8; E=1.36; R=5.87; rawP=5.36e-05; adjP=0.0002 | | | | | | |
| Index | UserID | Value | Gene Symbol | Gene Name | EntrezGene | Ensembl |
| 1 | P54819 | NA | AK2 | adenylate kinase 2 | 204 | ENSG00000004455 |
| 2 | P17931 | NA | LGALS3 | lectin, galactoside-binding, soluble, 3 | 3958 | ENSG00000131981 |
| 3 | Q10713 | NA | PMPCA | peptidase (mitochondrial processing) alpha | 23203 | ENSG00000165688 |
| 4 | P42765 | NA | ACAA2 | acetyl-CoA acyltransferase 2 | 10449 | ENSG00000167315 |
| 5 | Q99714 | NA | HSD17B10 | hydroxysteroid (17-beta) dehydrogenase 10 | 3028 | ENSG00000072506 |
| 6 | P04179 | NA | SOD2 | superoxide dismutase 2, mitochondrial | 6648 | ENSG00000112096 |
| 7 | P19367 | NA | HK1 | hexokinase 1 | 3098 | ENSG00000156515 |
| 8 | P31040 | NA | SDHA | succinate dehydrogenase complex, subunit A, flavoprotein (Fp) | 6389 | ENSG00000073578 |

  
  

| **Database:cellular component      &nbspName:mitochondrial inner membrane      &nbspID:GO:0005743** | | | | | | |
| --- | --- | --- | --- | --- | --- | --- |
| C=351; O=7; E=0.95; R=7.39; rawP=3.92e-05; adjP=0.0002 | | | | | | |
| Index | UserID | Value | Gene Symbol | Gene Name | EntrezGene | Ensembl |
| 1 | P54819 | NA | AK2 | adenylate kinase 2 | 204 | ENSG00000004455 |
| 2 | P17931 | NA | LGALS3 | lectin, galactoside-binding, soluble, 3 | 3958 | ENSG00000131981 |
| 3 | P31040 | NA | SDHA | succinate dehydrogenase complex, subunit A, flavoprotein (Fp) | 6389 | ENSG00000073578 |
| 4 | Q10713 | NA | PMPCA | peptidase (mitochondrial processing) alpha | 23203 | ENSG00000165688 |
| 5 | P42765 | NA | ACAA2 | acetyl-CoA acyltransferase 2 | 10449 | ENSG00000167315 |
| 6 | Q99714 | NA | HSD17B10 | hydroxysteroid (17-beta) dehydrogenase 10 | 3028 | ENSG00000072506 |
| 7 | P04179 | NA | SOD2 | superoxide dismutase 2, mitochondrial | 6648 | ENSG00000112096 |

  
  

| **Database:cellular component      &nbspName:intracellular membrane-bounded organelle      &nbspID:GO:0043231** | | | | | | |
| --- | --- | --- | --- | --- | --- | --- |
| C=9484; O=38; E=25.58; R=1.49; rawP=7.67e-05; adjP=0.0002 | | | | | | |
| Index | UserID | Value | Gene Symbol | Gene Name | EntrezGene | Ensembl |
| 1 | P09661 | NA | SNRPA1 | small nuclear ribonucleoprotein polypeptide A' | 6627 | ENSG00000131876 |
| 2 | P10253 | NA | GAA | glucosidase, alpha; acid | 2548 | ENSG00000171298 |
| 3 | P54819 | NA | AK2 | adenylate kinase 2 | 204 | ENSG00000004455 |
| 4 | P17931 | NA | LGALS3 | lectin, galactoside-binding, soluble, 3 | 3958 | ENSG00000131981 |
| 5 | P05787 | NA | KRT8 | keratin 8 | 3856 | ENSG00000170421 |
| 6 | Q9UL46 | NA | PSME2 | proteasome (prosome, macropain) activator subunit 2 (PA28 beta) | 5721 | ENSG00000100911 |
| 7 | P50453 | NA | SERPINB9 | serpin peptidase inhibitor, clade B (ovalbumin), member 9 | 5272 | ENSG00000170542 |
| 8 | Q99714 | NA | HSD17B10 | hydroxysteroid (17-beta) dehydrogenase 10 | 3028 | ENSG00000072506 |
| 9 | P04179 | NA | SOD2 | superoxide dismutase 2, mitochondrial | 6648 | ENSG00000112096 |
| 10 | Q06323 | NA | PSME1 | proteasome (prosome, macropain) activator subunit 1 (PA28 alpha) | 5720 | ENSG00000092010 |
| 11 | P07741 | NA | APRT | adenine phosphoribosyltransferase | 353 | ENSG00000198931 |
| 12 | P31040 | NA | SDHA | succinate dehydrogenase complex, subunit A, flavoprotein (Fp) | 6389 | ENSG00000073578 |
| 13 | P38117 | NA | ETFB | electron-transfer-flavoprotein, beta polypeptide | 2109 | ENSG00000105379 |
| 14 | Q16698 | NA | DECR1 | 2,4-dienoyl CoA reductase 1, mitochondrial | 1666 | ENSG00000104325 |
| 15 | Q05639 | NA | EEF1A2 | eukaryotic translation elongation factor 1 alpha 2 | 1917 | ENSG00000101210 |
| 16 | P54727 | NA | RAD23B | RAD23 homolog B (S. cerevisiae) | 5887 | ENSG00000119318 |
| 17 | Q10713 | NA | PMPCA | peptidase (mitochondrial processing) alpha | 23203 | ENSG00000165688 |
| 18 | P42765 | NA | ACAA2 | acetyl-CoA acyltransferase 2 | 10449 | ENSG00000167315 |
| 19 | P39687 | NA | ANP32A | acidic (leucine-rich) nuclear phosphoprotein 32 family, member A | 8125 | ENSG00000140350 |
| 20 | P49419 | NA | ALDH7A1 | aldehyde dehydrogenase 7 family, member A1 | 501 | ENSG00000164904 |
| 21 | P21980 | NA | TGM2 | transglutaminase 2 (C polypeptide, protein-glutamine-gamma-glutamyltransferase) | 7052 | ENSG00000198959 |
| 22 | Q5SSJ5 | NA | HP1BP3 | heterochromatin protein 1, binding protein 3 | 50809 | ENSG00000127483 |
| 23 | P26447 | NA | S100A4 | S100 calcium binding protein A4 | 6275 | ENSG00000196154 |
| 24 | Q9BXW7 | NA | CECR5 | cat eye syndrome chromosome region, candidate 5 | 27440 | ENSG00000069998 |
| 25 | P30837 | NA | ALDH1B1 | aldehyde dehydrogenase 1 family, member B1 | 219 | ENSG00000137124 |
| 26 | P38571 | NA | LIPA | lipase A, lysosomal acid, cholesterol esterase | 3988 | ENSG00000107798 |
| 27 | Q13263 | NA | TRIM28 | tripartite motif containing 28 | 10155 | ENSG00000130726 |
| 28 | Q13011 | NA | ECH1 | enoyl CoA hydratase 1, peroxisomal | 1891 | ENSG00000104823 |
| 29 | Q99798 | NA | ACO2 | aconitase 2, mitochondrial | 50 | ENSG00000100412 |
| 30 | Q9UBQ7 | NA | GRHPR | glyoxylate reductase/hydroxypyruvate reductase | 9380 | ENSG00000137106 |
| 31 | Q01105 | NA | SET | SET nuclear oncogene | 6418 | ENSG00000119335 |
| 32 | P13674 | NA | P4HA1 | prolyl 4-hydroxylase, alpha polypeptide I | 5033 | ENSG00000122884 |
| 33 | O43598 | NA | C6orf108 | chromosome 6 open reading frame 108 | 10591 | ENSG00000112667 |
| 34 | P06744 | NA | GPI | glucose-6-phosphate isomerase | 2821 | ENSG00000105220 |
| 35 | P19367 | NA | HK1 | hexokinase 1 | 3098 | ENSG00000156515 |
| 36 | Q9BQ69 | NA | MACROD1 | MACRO domain containing 1 | 28992 | ENSG00000133315 |
| 37 | P05783 | NA | KRT18 | keratin 18 | 3875 | ENSG00000111057 |
| 38 | Q16531 | NA | DDB1 | damage-specific DNA binding protein 1, 127kDa | 1642 | ENSG00000167986 |

  
  

| **Database:cellular component      &nbspName:intracellular organelle part      &nbspID:GO:0044446** | | | | | | |
| --- | --- | --- | --- | --- | --- | --- |
| C=6690; O=31; E=18.05; R=1.72; rawP=8.70e-05; adjP=0.0002 | | | | | | |
| Index | UserID | Value | Gene Symbol | Gene Name | EntrezGene | Ensembl |
| 1 | P09661 | NA | SNRPA1 | small nuclear ribonucleoprotein polypeptide A' | 6627 | ENSG00000131876 |
| 2 | P10253 | NA | GAA | glucosidase, alpha; acid | 2548 | ENSG00000171298 |
| 3 | P54819 | NA | AK2 | adenylate kinase 2 | 204 | ENSG00000004455 |
| 4 | P17931 | NA | LGALS3 | lectin, galactoside-binding, soluble, 3 | 3958 | ENSG00000131981 |
| 5 | P05787 | NA | KRT8 | keratin 8 | 3856 | ENSG00000170421 |
| 6 | Q9UL46 | NA | PSME2 | proteasome (prosome, macropain) activator subunit 2 (PA28 beta) | 5721 | ENSG00000100911 |
| 7 | Q99714 | NA | HSD17B10 | hydroxysteroid (17-beta) dehydrogenase 10 | 3028 | ENSG00000072506 |
| 8 | P04179 | NA | SOD2 | superoxide dismutase 2, mitochondrial | 6648 | ENSG00000112096 |
| 9 | Q06323 | NA | PSME1 | proteasome (prosome, macropain) activator subunit 1 (PA28 alpha) | 5720 | ENSG00000092010 |
| 10 | P07741 | NA | APRT | adenine phosphoribosyltransferase | 353 | ENSG00000198931 |
| 11 | P30837 | NA | ALDH1B1 | aldehyde dehydrogenase 1 family, member B1 | 219 | ENSG00000137124 |
| 12 | Q13263 | NA | TRIM28 | tripartite motif containing 28 | 10155 | ENSG00000130726 |
| 13 | P31040 | NA | SDHA | succinate dehydrogenase complex, subunit A, flavoprotein (Fp) | 6389 | ENSG00000073578 |
| 14 | P38117 | NA | ETFB | electron-transfer-flavoprotein, beta polypeptide | 2109 | ENSG00000105379 |
| 15 | Q9UBQ7 | NA | GRHPR | glyoxylate reductase/hydroxypyruvate reductase | 9380 | ENSG00000137106 |
| 16 | Q99798 | NA | ACO2 | aconitase 2, mitochondrial | 50 | ENSG00000100412 |
| 17 | Q16698 | NA | DECR1 | 2,4-dienoyl CoA reductase 1, mitochondrial | 1666 | ENSG00000104325 |
| 18 | Q01105 | NA | SET | SET nuclear oncogene | 6418 | ENSG00000119335 |
| 19 | P54727 | NA | RAD23B | RAD23 homolog B (S. cerevisiae) | 5887 | ENSG00000119318 |
| 20 | P42765 | NA | ACAA2 | acetyl-CoA acyltransferase 2 | 10449 | ENSG00000167315 |
| 21 | Q10713 | NA | PMPCA | peptidase (mitochondrial processing) alpha | 23203 | ENSG00000165688 |
| 22 | P13674 | NA | P4HA1 | prolyl 4-hydroxylase, alpha polypeptide I | 5033 | ENSG00000122884 |
| 23 | P39687 | NA | ANP32A | acidic (leucine-rich) nuclear phosphoprotein 32 family, member A | 8125 | ENSG00000140350 |
| 24 | O43598 | NA | C6orf108 | chromosome 6 open reading frame 108 | 10591 | ENSG00000112667 |
| 25 | P49419 | NA | ALDH7A1 | aldehyde dehydrogenase 7 family, member A1 | 501 | ENSG00000164904 |
| 26 | P06744 | NA | GPI | glucose-6-phosphate isomerase | 2821 | ENSG00000105220 |
| 27 | P19367 | NA | HK1 | hexokinase 1 | 3098 | ENSG00000156515 |
| 28 | P05783 | NA | KRT18 | keratin 18 | 3875 | ENSG00000111057 |
| 29 | Q9UDT6 | NA | CLIP2 | CAP-GLY domain containing linker protein 2 | 7461 | ENSG00000106665 |
| 30 | Q16531 | NA | DDB1 | damage-specific DNA binding protein 1, 127kDa | 1642 | ENSG00000167986 |
| 31 | Q5SSJ5 | NA | HP1BP3 | heterochromatin protein 1, binding protein 3 | 50809 | ENSG00000127483 |

  
  

| **Database:cellular component      &nbspName:organelle inner membrane      &nbspID:GO:0019866** | | | | | | |
| --- | --- | --- | --- | --- | --- | --- |
| C=381; O=7; E=1.03; R=6.81; rawP=6.59e-05; adjP=0.0002 | | | | | | |
| Index | UserID | Value | Gene Symbol | Gene Name | EntrezGene | Ensembl |
| 1 | P54819 | NA | AK2 | adenylate kinase 2 | 204 | ENSG00000004455 |
| 2 | P17931 | NA | LGALS3 | lectin, galactoside-binding, soluble, 3 | 3958 | ENSG00000131981 |
| 3 | P31040 | NA | SDHA | succinate dehydrogenase complex, subunit A, flavoprotein (Fp) | 6389 | ENSG00000073578 |
| 4 | Q10713 | NA | PMPCA | peptidase (mitochondrial processing) alpha | 23203 | ENSG00000165688 |
| 5 | P42765 | NA | ACAA2 | acetyl-CoA acyltransferase 2 | 10449 | ENSG00000167315 |
| 6 | Q99714 | NA | HSD17B10 | hydroxysteroid (17-beta) dehydrogenase 10 | 3028 | ENSG00000072506 |
| 7 | P04179 | NA | SOD2 | superoxide dismutase 2, mitochondrial | 6648 | ENSG00000112096 |

  
  

| **Database:cellular component      &nbspName:mitochondrial envelope      &nbspID:GO:0005740** | | | | | | |
| --- | --- | --- | --- | --- | --- | --- |
| C=526; O=8; E=1.42; R=5.64; rawP=7.14e-05; adjP=0.0002 | | | | | | |
| Index | UserID | Value | Gene Symbol | Gene Name | EntrezGene | Ensembl |
| 1 | P54819 | NA | AK2 | adenylate kinase 2 | 204 | ENSG00000004455 |
| 2 | P17931 | NA | LGALS3 | lectin, galactoside-binding, soluble, 3 | 3958 | ENSG00000131981 |
| 3 | Q10713 | NA | PMPCA | peptidase (mitochondrial processing) alpha | 23203 | ENSG00000165688 |
| 4 | P42765 | NA | ACAA2 | acetyl-CoA acyltransferase 2 | 10449 | ENSG00000167315 |
| 5 | Q99714 | NA | HSD17B10 | hydroxysteroid (17-beta) dehydrogenase 10 | 3028 | ENSG00000072506 |
| 6 | P04179 | NA | SOD2 | superoxide dismutase 2, mitochondrial | 6648 | ENSG00000112096 |
| 7 | P19367 | NA | HK1 | hexokinase 1 | 3098 | ENSG00000156515 |
| 8 | P31040 | NA | SDHA | succinate dehydrogenase complex, subunit A, flavoprotein (Fp) | 6389 | ENSG00000073578 |

  
  

| **Database:cellular component      &nbspName:organelle part      &nbspID:GO:0044422** | | | | | | |
| --- | --- | --- | --- | --- | --- | --- |
| C=6777; O=31; E=18.28; R=1.70; rawP=0.0001; adjP=0.0003 | | | | | | |
| Index | UserID | Value | Gene Symbol | Gene Name | EntrezGene | Ensembl |
| 1 | P09661 | NA | SNRPA1 | small nuclear ribonucleoprotein polypeptide A' | 6627 | ENSG00000131876 |
| 2 | P10253 | NA | GAA | glucosidase, alpha; acid | 2548 | ENSG00000171298 |
| 3 | P54819 | NA | AK2 | adenylate kinase 2 | 204 | ENSG00000004455 |
| 4 | P17931 | NA | LGALS3 | lectin, galactoside-binding, soluble, 3 | 3958 | ENSG00000131981 |
| 5 | P05787 | NA | KRT8 | keratin 8 | 3856 | ENSG00000170421 |
| 6 | Q9UL46 | NA | PSME2 | proteasome (prosome, macropain) activator subunit 2 (PA28 beta) | 5721 | ENSG00000100911 |
| 7 | Q99714 | NA | HSD17B10 | hydroxysteroid (17-beta) dehydrogenase 10 | 3028 | ENSG00000072506 |
| 8 | P04179 | NA | SOD2 | superoxide dismutase 2, mitochondrial | 6648 | ENSG00000112096 |
| 9 | Q06323 | NA | PSME1 | proteasome (prosome, macropain) activator subunit 1 (PA28 alpha) | 5720 | ENSG00000092010 |
| 10 | P07741 | NA | APRT | adenine phosphoribosyltransferase | 353 | ENSG00000198931 |
| 11 | P30837 | NA | ALDH1B1 | aldehyde dehydrogenase 1 family, member B1 | 219 | ENSG00000137124 |
| 12 | Q13263 | NA | TRIM28 | tripartite motif containing 28 | 10155 | ENSG00000130726 |
| 13 | P31040 | NA | SDHA | succinate dehydrogenase complex, subunit A, flavoprotein (Fp) | 6389 | ENSG00000073578 |
| 14 | P38117 | NA | ETFB | electron-transfer-flavoprotein, beta polypeptide | 2109 | ENSG00000105379 |
| 15 | Q9UBQ7 | NA | GRHPR | glyoxylate reductase/hydroxypyruvate reductase | 9380 | ENSG00000137106 |
| 16 | Q99798 | NA | ACO2 | aconitase 2, mitochondrial | 50 | ENSG00000100412 |
| 17 | Q16698 | NA | DECR1 | 2,4-dienoyl CoA reductase 1, mitochondrial | 1666 | ENSG00000104325 |
| 18 | Q01105 | NA | SET | SET nuclear oncogene | 6418 | ENSG00000119335 |
| 19 | P54727 | NA | RAD23B | RAD23 homolog B (S. cerevisiae) | 5887 | ENSG00000119318 |
| 20 | P42765 | NA | ACAA2 | acetyl-CoA acyltransferase 2 | 10449 | ENSG00000167315 |
| 21 | Q10713 | NA | PMPCA | peptidase (mitochondrial processing) alpha | 23203 | ENSG00000165688 |
| 22 | P13674 | NA | P4HA1 | prolyl 4-hydroxylase, alpha polypeptide I | 5033 | ENSG00000122884 |
| 23 | P39687 | NA | ANP32A | acidic (leucine-rich) nuclear phosphoprotein 32 family, member A | 8125 | ENSG00000140350 |
| 24 | O43598 | NA | C6orf108 | chromosome 6 open reading frame 108 | 10591 | ENSG00000112667 |
| 25 | P49419 | NA | ALDH7A1 | aldehyde dehydrogenase 7 family, member A1 | 501 | ENSG00000164904 |
| 26 | P06744 | NA | GPI | glucose-6-phosphate isomerase | 2821 | ENSG00000105220 |
| 27 | P19367 | NA | HK1 | hexokinase 1 | 3098 | ENSG00000156515 |
| 28 | P05783 | NA | KRT18 | keratin 18 | 3875 | ENSG00000111057 |
| 29 | Q9UDT6 | NA | CLIP2 | CAP-GLY domain containing linker protein 2 | 7461 | ENSG00000106665 |
| 30 | Q16531 | NA | DDB1 | damage-specific DNA binding protein 1, 127kDa | 1642 | ENSG00000167986 |
| 31 | Q5SSJ5 | NA | HP1BP3 | heterochromatin protein 1, binding protein 3 | 50809 | ENSG00000127483 |

  
  

| **Database:cellular component      &nbspName:intracellular organelle      &nbspID:GO:0043229** | | | | | | |
| --- | --- | --- | --- | --- | --- | --- |
| C=10521; O=39; E=28.38; R=1.37; rawP=0.0004; adjP=0.0009 | | | | | | |
| Index | UserID | Value | Gene Symbol | Gene Name | EntrezGene | Ensembl |
| 1 | P09661 | NA | SNRPA1 | small nuclear ribonucleoprotein polypeptide A' | 6627 | ENSG00000131876 |
| 2 | P10253 | NA | GAA | glucosidase, alpha; acid | 2548 | ENSG00000171298 |
| 3 | P54819 | NA | AK2 | adenylate kinase 2 | 204 | ENSG00000004455 |
| 4 | P17931 | NA | LGALS3 | lectin, galactoside-binding, soluble, 3 | 3958 | ENSG00000131981 |
| 5 | P05787 | NA | KRT8 | keratin 8 | 3856 | ENSG00000170421 |
| 6 | Q9UL46 | NA | PSME2 | proteasome (prosome, macropain) activator subunit 2 (PA28 beta) | 5721 | ENSG00000100911 |
| 7 | P50453 | NA | SERPINB9 | serpin peptidase inhibitor, clade B (ovalbumin), member 9 | 5272 | ENSG00000170542 |
| 8 | Q99714 | NA | HSD17B10 | hydroxysteroid (17-beta) dehydrogenase 10 | 3028 | ENSG00000072506 |
| 9 | P04179 | NA | SOD2 | superoxide dismutase 2, mitochondrial | 6648 | ENSG00000112096 |
| 10 | Q06323 | NA | PSME1 | proteasome (prosome, macropain) activator subunit 1 (PA28 alpha) | 5720 | ENSG00000092010 |
| 11 | P07741 | NA | APRT | adenine phosphoribosyltransferase | 353 | ENSG00000198931 |
| 12 | P31040 | NA | SDHA | succinate dehydrogenase complex, subunit A, flavoprotein (Fp) | 6389 | ENSG00000073578 |
| 13 | P38117 | NA | ETFB | electron-transfer-flavoprotein, beta polypeptide | 2109 | ENSG00000105379 |
| 14 | Q16698 | NA | DECR1 | 2,4-dienoyl CoA reductase 1, mitochondrial | 1666 | ENSG00000104325 |
| 15 | Q05639 | NA | EEF1A2 | eukaryotic translation elongation factor 1 alpha 2 | 1917 | ENSG00000101210 |
| 16 | P54727 | NA | RAD23B | RAD23 homolog B (S. cerevisiae) | 5887 | ENSG00000119318 |
| 17 | Q10713 | NA | PMPCA | peptidase (mitochondrial processing) alpha | 23203 | ENSG00000165688 |
| 18 | P42765 | NA | ACAA2 | acetyl-CoA acyltransferase 2 | 10449 | ENSG00000167315 |
| 19 | P39687 | NA | ANP32A | acidic (leucine-rich) nuclear phosphoprotein 32 family, member A | 8125 | ENSG00000140350 |
| 20 | P49419 | NA | ALDH7A1 | aldehyde dehydrogenase 7 family, member A1 | 501 | ENSG00000164904 |
| 21 | P21980 | NA | TGM2 | transglutaminase 2 (C polypeptide, protein-glutamine-gamma-glutamyltransferase) | 7052 | ENSG00000198959 |
| 22 | Q9UDT6 | NA | CLIP2 | CAP-GLY domain containing linker protein 2 | 7461 | ENSG00000106665 |
| 23 | Q5SSJ5 | NA | HP1BP3 | heterochromatin protein 1, binding protein 3 | 50809 | ENSG00000127483 |
| 24 | P26447 | NA | S100A4 | S100 calcium binding protein A4 | 6275 | ENSG00000196154 |
| 25 | Q9BXW7 | NA | CECR5 | cat eye syndrome chromosome region, candidate 5 | 27440 | ENSG00000069998 |
| 26 | P30837 | NA | ALDH1B1 | aldehyde dehydrogenase 1 family, member B1 | 219 | ENSG00000137124 |
| 27 | P38571 | NA | LIPA | lipase A, lysosomal acid, cholesterol esterase | 3988 | ENSG00000107798 |
| 28 | Q13263 | NA | TRIM28 | tripartite motif containing 28 | 10155 | ENSG00000130726 |
| 29 | Q13011 | NA | ECH1 | enoyl CoA hydratase 1, peroxisomal | 1891 | ENSG00000104823 |
| 30 | Q99798 | NA | ACO2 | aconitase 2, mitochondrial | 50 | ENSG00000100412 |
| 31 | Q9UBQ7 | NA | GRHPR | glyoxylate reductase/hydroxypyruvate reductase | 9380 | ENSG00000137106 |
| 32 | Q01105 | NA | SET | SET nuclear oncogene | 6418 | ENSG00000119335 |
| 33 | P13674 | NA | P4HA1 | prolyl 4-hydroxylase, alpha polypeptide I | 5033 | ENSG00000122884 |
| 34 | O43598 | NA | C6orf108 | chromosome 6 open reading frame 108 | 10591 | ENSG00000112667 |
| 35 | P06744 | NA | GPI | glucose-6-phosphate isomerase | 2821 | ENSG00000105220 |
| 36 | P19367 | NA | HK1 | hexokinase 1 | 3098 | ENSG00000156515 |
| 37 | Q9BQ69 | NA | MACROD1 | MACRO domain containing 1 | 28992 | ENSG00000133315 |
| 38 | P05783 | NA | KRT18 | keratin 18 | 3875 | ENSG00000111057 |
| 39 | Q16531 | NA | DDB1 | damage-specific DNA binding protein 1, 127kDa | 1642 | ENSG00000167986 |

  
  

| **Database:cellular component      &nbspName:organelle      &nbspID:GO:0043226** | | | | | | |
| --- | --- | --- | --- | --- | --- | --- |
| C=10536; O=39; E=28.42; R=1.37; rawP=0.0004; adjP=0.0009 | | | | | | |
| Index | UserID | Value | Gene Symbol | Gene Name | EntrezGene | Ensembl |
| 1 | P09661 | NA | SNRPA1 | small nuclear ribonucleoprotein polypeptide A' | 6627 | ENSG00000131876 |
| 2 | P10253 | NA | GAA | glucosidase, alpha; acid | 2548 | ENSG00000171298 |
| 3 | P54819 | NA | AK2 | adenylate kinase 2 | 204 | ENSG00000004455 |
| 4 | P17931 | NA | LGALS3 | lectin, galactoside-binding, soluble, 3 | 3958 | ENSG00000131981 |
| 5 | P05787 | NA | KRT8 | keratin 8 | 3856 | ENSG00000170421 |
| 6 | Q9UL46 | NA | PSME2 | proteasome (prosome, macropain) activator subunit 2 (PA28 beta) | 5721 | ENSG00000100911 |
| 7 | P50453 | NA | SERPINB9 | serpin peptidase inhibitor, clade B (ovalbumin), member 9 | 5272 | ENSG00000170542 |
| 8 | Q99714 | NA | HSD17B10 | hydroxysteroid (17-beta) dehydrogenase 10 | 3028 | ENSG00000072506 |
| 9 | P04179 | NA | SOD2 | superoxide dismutase 2, mitochondrial | 6648 | ENSG00000112096 |
| 10 | Q06323 | NA | PSME1 | proteasome (prosome, macropain) activator subunit 1 (PA28 alpha) | 5720 | ENSG00000092010 |
| 11 | P07741 | NA | APRT | adenine phosphoribosyltransferase | 353 | ENSG00000198931 |
| 12 | P31040 | NA | SDHA | succinate dehydrogenase complex, subunit A, flavoprotein (Fp) | 6389 | ENSG00000073578 |
| 13 | P38117 | NA | ETFB | electron-transfer-flavoprotein, beta polypeptide | 2109 | ENSG00000105379 |
| 14 | Q16698 | NA | DECR1 | 2,4-dienoyl CoA reductase 1, mitochondrial | 1666 | ENSG00000104325 |
| 15 | Q05639 | NA | EEF1A2 | eukaryotic translation elongation factor 1 alpha 2 | 1917 | ENSG00000101210 |
| 16 | P54727 | NA | RAD23B | RAD23 homolog B (S. cerevisiae) | 5887 | ENSG00000119318 |
| 17 | Q10713 | NA | PMPCA | peptidase (mitochondrial processing) alpha | 23203 | ENSG00000165688 |
| 18 | P42765 | NA | ACAA2 | acetyl-CoA acyltransferase 2 | 10449 | ENSG00000167315 |
| 19 | P39687 | NA | ANP32A | acidic (leucine-rich) nuclear phosphoprotein 32 family, member A | 8125 | ENSG00000140350 |
| 20 | P49419 | NA | ALDH7A1 | aldehyde dehydrogenase 7 family, member A1 | 501 | ENSG00000164904 |
| 21 | P21980 | NA | TGM2 | transglutaminase 2 (C polypeptide, protein-glutamine-gamma-glutamyltransferase) | 7052 | ENSG00000198959 |
| 22 | Q9UDT6 | NA | CLIP2 | CAP-GLY domain containing linker protein 2 | 7461 | ENSG00000106665 |
| 23 | Q5SSJ5 | NA | HP1BP3 | heterochromatin protein 1, binding protein 3 | 50809 | ENSG00000127483 |
| 24 | P26447 | NA | S100A4 | S100 calcium binding protein A4 | 6275 | ENSG00000196154 |
| 25 | Q9BXW7 | NA | CECR5 | cat eye syndrome chromosome region, candidate 5 | 27440 | ENSG00000069998 |
| 26 | P30837 | NA | ALDH1B1 | aldehyde dehydrogenase 1 family, member B1 | 219 | ENSG00000137124 |
| 27 | P38571 | NA | LIPA | lipase A, lysosomal acid, cholesterol esterase | 3988 | ENSG00000107798 |
| 28 | Q13263 | NA | TRIM28 | tripartite motif containing 28 | 10155 | ENSG00000130726 |
| 29 | Q13011 | NA | ECH1 | enoyl CoA hydratase 1, peroxisomal | 1891 | ENSG00000104823 |
| 30 | Q99798 | NA | ACO2 | aconitase 2, mitochondrial | 50 | ENSG00000100412 |
| 31 | Q9UBQ7 | NA | GRHPR | glyoxylate reductase/hydroxypyruvate reductase | 9380 | ENSG00000137106 |
| 32 | Q01105 | NA | SET | SET nuclear oncogene | 6418 | ENSG00000119335 |
| 33 | P13674 | NA | P4HA1 | prolyl 4-hydroxylase, alpha polypeptide I | 5033 | ENSG00000122884 |
| 34 | O43598 | NA | C6orf108 | chromosome 6 open reading frame 108 | 10591 | ENSG00000112667 |
| 35 | P06744 | NA | GPI | glucose-6-phosphate isomerase | 2821 | ENSG00000105220 |
| 36 | P19367 | NA | HK1 | hexokinase 1 | 3098 | ENSG00000156515 |
| 37 | Q9BQ69 | NA | MACROD1 | MACRO domain containing 1 | 28992 | ENSG00000133315 |
| 38 | P05783 | NA | KRT18 | keratin 18 | 3875 | ENSG00000111057 |
| 39 | Q16531 | NA | DDB1 | damage-specific DNA binding protein 1, 127kDa | 1642 | ENSG00000167986 |

  
  

| **Database:cellular component      &nbspName:proteasome complex      &nbspID:GO:0000502** | | | | | | |
| --- | --- | --- | --- | --- | --- | --- |
| C=65; O=3; E=0.18; R=17.11; rawP=0.0007; adjP=0.0015 | | | | | | |
| Index | UserID | Value | Gene Symbol | Gene Name | EntrezGene | Ensembl |
| 1 | Q06323 | NA | PSME1 | proteasome (prosome, macropain) activator subunit 1 (PA28 alpha) | 5720 | ENSG00000092010 |
| 2 | Q9UL46 | NA | PSME2 | proteasome (prosome, macropain) activator subunit 2 (PA28 beta) | 5721 | ENSG00000100911 |
| 3 | P54727 | NA | RAD23B | RAD23 homolog B (S. cerevisiae) | 5887 | ENSG00000119318 |

  
  

| **Database:cellular component      &nbspName:organelle envelope      &nbspID:GO:0031967** | | | | | | |
| --- | --- | --- | --- | --- | --- | --- |
| C=817; O=8; E=2.20; R=3.63; rawP=0.0014; adjP=0.0026 | | | | | | |
| Index | UserID | Value | Gene Symbol | Gene Name | EntrezGene | Ensembl |
| 1 | P54819 | NA | AK2 | adenylate kinase 2 | 204 | ENSG00000004455 |
| 2 | P17931 | NA | LGALS3 | lectin, galactoside-binding, soluble, 3 | 3958 | ENSG00000131981 |
| 3 | Q10713 | NA | PMPCA | peptidase (mitochondrial processing) alpha | 23203 | ENSG00000165688 |
| 4 | P42765 | NA | ACAA2 | acetyl-CoA acyltransferase 2 | 10449 | ENSG00000167315 |
| 5 | Q99714 | NA | HSD17B10 | hydroxysteroid (17-beta) dehydrogenase 10 | 3028 | ENSG00000072506 |
| 6 | P04179 | NA | SOD2 | superoxide dismutase 2, mitochondrial | 6648 | ENSG00000112096 |
| 7 | P19367 | NA | HK1 | hexokinase 1 | 3098 | ENSG00000156515 |
| 8 | P31040 | NA | SDHA | succinate dehydrogenase complex, subunit A, flavoprotein (Fp) | 6389 | ENSG00000073578 |

  
  

| **Database:cellular component      &nbspName:cell part      &nbspID:GO:0044464** | | | | | | |
| --- | --- | --- | --- | --- | --- | --- |
| C=14405; O=45; E=38.86; R=1.16; rawP=0.0013; adjP=0.0026 | | | | | | |
| Index | UserID | Value | Gene Symbol | Gene Name | EntrezGene | Ensembl |
| 1 | P09661 | NA | SNRPA1 | small nuclear ribonucleoprotein polypeptide A' | 6627 | ENSG00000131876 |
| 2 | P10253 | NA | GAA | glucosidase, alpha; acid | 2548 | ENSG00000171298 |
| 3 | P54819 | NA | AK2 | adenylate kinase 2 | 204 | ENSG00000004455 |
| 4 | P17931 | NA | LGALS3 | lectin, galactoside-binding, soluble, 3 | 3958 | ENSG00000131981 |
| 5 | P05787 | NA | KRT8 | keratin 8 | 3856 | ENSG00000170421 |
| 6 | Q9UL46 | NA | PSME2 | proteasome (prosome, macropain) activator subunit 2 (PA28 beta) | 5721 | ENSG00000100911 |
| 7 | P50453 | NA | SERPINB9 | serpin peptidase inhibitor, clade B (ovalbumin), member 9 | 5272 | ENSG00000170542 |
| 8 | Q99714 | NA | HSD17B10 | hydroxysteroid (17-beta) dehydrogenase 10 | 3028 | ENSG00000072506 |
| 9 | P04179 | NA | SOD2 | superoxide dismutase 2, mitochondrial | 6648 | ENSG00000112096 |
| 10 | Q06323 | NA | PSME1 | proteasome (prosome, macropain) activator subunit 1 (PA28 alpha) | 5720 | ENSG00000092010 |
| 11 | P07741 | NA | APRT | adenine phosphoribosyltransferase | 353 | ENSG00000198931 |
| 12 | P30046 | NA | DDT | D-dopachrome tautomerase | 1652 | ENSG00000099977 |
| 13 | P31040 | NA | SDHA | succinate dehydrogenase complex, subunit A, flavoprotein (Fp) | 6389 | ENSG00000073578 |
| 14 | P38117 | NA | ETFB | electron-transfer-flavoprotein, beta polypeptide | 2109 | ENSG00000105379 |
| 15 | Q16698 | NA | DECR1 | 2,4-dienoyl CoA reductase 1, mitochondrial | 1666 | ENSG00000104325 |
| 16 | P54727 | NA | RAD23B | RAD23 homolog B (S. cerevisiae) | 5887 | ENSG00000119318 |
| 17 | Q05639 | NA | EEF1A2 | eukaryotic translation elongation factor 1 alpha 2 | 1917 | ENSG00000101210 |
| 18 | Q10713 | NA | PMPCA | peptidase (mitochondrial processing) alpha | 23203 | ENSG00000165688 |
| 19 | P42765 | NA | ACAA2 | acetyl-CoA acyltransferase 2 | 10449 | ENSG00000167315 |
| 20 | P39687 | NA | ANP32A | acidic (leucine-rich) nuclear phosphoprotein 32 family, member A | 8125 | ENSG00000140350 |
| 21 | P49419 | NA | ALDH7A1 | aldehyde dehydrogenase 7 family, member A1 | 501 | ENSG00000164904 |
| 22 | P21980 | NA | TGM2 | transglutaminase 2 (C polypeptide, protein-glutamine-gamma-glutamyltransferase) | 7052 | ENSG00000198959 |
| 23 | Q9UDT6 | NA | CLIP2 | CAP-GLY domain containing linker protein 2 | 7461 | ENSG00000106665 |
| 24 | Q5SSJ5 | NA | HP1BP3 | heterochromatin protein 1, binding protein 3 | 50809 | ENSG00000127483 |
| 25 | P02792 | NA | FTL | ferritin, light polypeptide | 2512 | ENSG00000087086 |
| 26 | P26447 | NA | S100A4 | S100 calcium binding protein A4 | 6275 | ENSG00000196154 |
| 27 | P06737 | NA | PYGL | phosphorylase, glycogen, liver | 5836 | ENSG00000100504 |
| 28 | Q9BXW7 | NA | CECR5 | cat eye syndrome chromosome region, candidate 5 | 27440 | ENSG00000069998 |
| 29 | P30837 | NA | ALDH1B1 | aldehyde dehydrogenase 1 family, member B1 | 219 | ENSG00000137124 |
| 30 | P38571 | NA | LIPA | lipase A, lysosomal acid, cholesterol esterase | 3988 | ENSG00000107798 |
| 31 | Q13263 | NA | TRIM28 | tripartite motif containing 28 | 10155 | ENSG00000130726 |
| 32 | P32119 | NA | PRDX2 | peroxiredoxin 2 | 7001 | ENSG00000167815 |
| 33 | Q13011 | NA | ECH1 | enoyl CoA hydratase 1, peroxisomal | 1891 | ENSG00000104823 |
| 34 | Q9UBQ7 | NA | GRHPR | glyoxylate reductase/hydroxypyruvate reductase | 9380 | ENSG00000137106 |
| 35 | Q99798 | NA | ACO2 | aconitase 2, mitochondrial | 50 | ENSG00000100412 |
| 36 | P00558 | NA | PGK1 | phosphoglycerate kinase 1 | 5230 | ENSG00000102144 |
| 37 | Q01105 | NA | SET | SET nuclear oncogene | 6418 | ENSG00000119335 |
| 38 | P13674 | NA | P4HA1 | prolyl 4-hydroxylase, alpha polypeptide I | 5033 | ENSG00000122884 |
| 39 | O43598 | NA | C6orf108 | chromosome 6 open reading frame 108 | 10591 | ENSG00000112667 |
| 40 | P06744 | NA | GPI | glucose-6-phosphate isomerase | 2821 | ENSG00000105220 |
| 41 | P19367 | NA | HK1 | hexokinase 1 | 3098 | ENSG00000156515 |
| 42 | Q9BQ69 | NA | MACROD1 | MACRO domain containing 1 | 28992 | ENSG00000133315 |
| 43 | P05783 | NA | KRT18 | keratin 18 | 3875 | ENSG00000111057 |
| 44 | P00352 | NA | ALDH1A1 | aldehyde dehydrogenase 1 family, member A1 | 216 | ENSG00000165092 |
| 45 | Q16531 | NA | DDB1 | damage-specific DNA binding protein 1, 127kDa | 1642 | ENSG00000167986 |

  
  

| **Database:cellular component      &nbspName:cell      &nbspID:GO:0005623** | | | | | | |
| --- | --- | --- | --- | --- | --- | --- |
| C=14406; O=45; E=38.86; R=1.16; rawP=0.0013; adjP=0.0026 | | | | | | |
| Index | UserID | Value | Gene Symbol | Gene Name | EntrezGene | Ensembl |
| 1 | P09661 | NA | SNRPA1 | small nuclear ribonucleoprotein polypeptide A' | 6627 | ENSG00000131876 |
| 2 | P10253 | NA | GAA | glucosidase, alpha; acid | 2548 | ENSG00000171298 |
| 3 | P54819 | NA | AK2 | adenylate kinase 2 | 204 | ENSG00000004455 |
| 4 | P17931 | NA | LGALS3 | lectin, galactoside-binding, soluble, 3 | 3958 | ENSG00000131981 |
| 5 | P05787 | NA | KRT8 | keratin 8 | 3856 | ENSG00000170421 |
| 6 | Q9UL46 | NA | PSME2 | proteasome (prosome, macropain) activator subunit 2 (PA28 beta) | 5721 | ENSG00000100911 |
| 7 | P50453 | NA | SERPINB9 | serpin peptidase inhibitor, clade B (ovalbumin), member 9 | 5272 | ENSG00000170542 |
| 8 | Q99714 | NA | HSD17B10 | hydroxysteroid (17-beta) dehydrogenase 10 | 3028 | ENSG00000072506 |
| 9 | P04179 | NA | SOD2 | superoxide dismutase 2, mitochondrial | 6648 | ENSG00000112096 |
| 10 | Q06323 | NA | PSME1 | proteasome (prosome, macropain) activator subunit 1 (PA28 alpha) | 5720 | ENSG00000092010 |
| 11 | P07741 | NA | APRT | adenine phosphoribosyltransferase | 353 | ENSG00000198931 |
| 12 | P30046 | NA | DDT | D-dopachrome tautomerase | 1652 | ENSG00000099977 |
| 13 | P31040 | NA | SDHA | succinate dehydrogenase complex, subunit A, flavoprotein (Fp) | 6389 | ENSG00000073578 |
| 14 | P38117 | NA | ETFB | electron-transfer-flavoprotein, beta polypeptide | 2109 | ENSG00000105379 |
| 15 | Q16698 | NA | DECR1 | 2,4-dienoyl CoA reductase 1, mitochondrial | 1666 | ENSG00000104325 |
| 16 | P54727 | NA | RAD23B | RAD23 homolog B (S. cerevisiae) | 5887 | ENSG00000119318 |
| 17 | Q05639 | NA | EEF1A2 | eukaryotic translation elongation factor 1 alpha 2 | 1917 | ENSG00000101210 |
| 18 | Q10713 | NA | PMPCA | peptidase (mitochondrial processing) alpha | 23203 | ENSG00000165688 |
| 19 | P42765 | NA | ACAA2 | acetyl-CoA acyltransferase 2 | 10449 | ENSG00000167315 |
| 20 | P39687 | NA | ANP32A | acidic (leucine-rich) nuclear phosphoprotein 32 family, member A | 8125 | ENSG00000140350 |
| 21 | P49419 | NA | ALDH7A1 | aldehyde dehydrogenase 7 family, member A1 | 501 | ENSG00000164904 |
| 22 | P21980 | NA | TGM2 | transglutaminase 2 (C polypeptide, protein-glutamine-gamma-glutamyltransferase) | 7052 | ENSG00000198959 |
| 23 | Q9UDT6 | NA | CLIP2 | CAP-GLY domain containing linker protein 2 | 7461 | ENSG00000106665 |
| 24 | Q5SSJ5 | NA | HP1BP3 | heterochromatin protein 1, binding protein 3 | 50809 | ENSG00000127483 |
| 25 | P02792 | NA | FTL | ferritin, light polypeptide | 2512 | ENSG00000087086 |
| 26 | P26447 | NA | S100A4 | S100 calcium binding protein A4 | 6275 | ENSG00000196154 |
| 27 | P06737 | NA | PYGL | phosphorylase, glycogen, liver | 5836 | ENSG00000100504 |
| 28 | Q9BXW7 | NA | CECR5 | cat eye syndrome chromosome region, candidate 5 | 27440 | ENSG00000069998 |
| 29 | P30837 | NA | ALDH1B1 | aldehyde dehydrogenase 1 family, member B1 | 219 | ENSG00000137124 |
| 30 | P38571 | NA | LIPA | lipase A, lysosomal acid, cholesterol esterase | 3988 | ENSG00000107798 |
| 31 | Q13263 | NA | TRIM28 | tripartite motif containing 28 | 10155 | ENSG00000130726 |
| 32 | P32119 | NA | PRDX2 | peroxiredoxin 2 | 7001 | ENSG00000167815 |
| 33 | Q13011 | NA | ECH1 | enoyl CoA hydratase 1, peroxisomal | 1891 | ENSG00000104823 |
| 34 | Q9UBQ7 | NA | GRHPR | glyoxylate reductase/hydroxypyruvate reductase | 9380 | ENSG00000137106 |
| 35 | Q99798 | NA | ACO2 | aconitase 2, mitochondrial | 50 | ENSG00000100412 |
| 36 | P00558 | NA | PGK1 | phosphoglycerate kinase 1 | 5230 | ENSG00000102144 |
| 37 | Q01105 | NA | SET | SET nuclear oncogene | 6418 | ENSG00000119335 |
| 38 | P13674 | NA | P4HA1 | prolyl 4-hydroxylase, alpha polypeptide I | 5033 | ENSG00000122884 |
| 39 | O43598 | NA | C6orf108 | chromosome 6 open reading frame 108 | 10591 | ENSG00000112667 |
| 40 | P06744 | NA | GPI | glucose-6-phosphate isomerase | 2821 | ENSG00000105220 |
| 41 | P19367 | NA | HK1 | hexokinase 1 | 3098 | ENSG00000156515 |
| 42 | Q9BQ69 | NA | MACROD1 | MACRO domain containing 1 | 28992 | ENSG00000133315 |
| 43 | P05783 | NA | KRT18 | keratin 18 | 3875 | ENSG00000111057 |
| 44 | P00352 | NA | ALDH1A1 | aldehyde dehydrogenase 1 family, member A1 | 216 | ENSG00000165092 |
| 45 | Q16531 | NA | DDB1 | damage-specific DNA binding protein 1, 127kDa | 1642 | ENSG00000167986 |

  
  

| **Database:cellular component      &nbspName:nuclear lumen      &nbspID:GO:0031981** | | | | | | |
| --- | --- | --- | --- | --- | --- | --- |
| C=2728; O=16; E=7.36; R=2.17; rawP=0.0014; adjP=0.0026 | | | | | | |
| Index | UserID | Value | Gene Symbol | Gene Name | EntrezGene | Ensembl |
| 1 | P09661 | NA | SNRPA1 | small nuclear ribonucleoprotein polypeptide A' | 6627 | ENSG00000131876 |
| 2 | Q16698 | NA | DECR1 | 2,4-dienoyl CoA reductase 1, mitochondrial | 1666 | ENSG00000104325 |
| 3 | P05787 | NA | KRT8 | keratin 8 | 3856 | ENSG00000170421 |
| 4 | Q9UL46 | NA | PSME2 | proteasome (prosome, macropain) activator subunit 2 (PA28 beta) | 5721 | ENSG00000100911 |
| 5 | Q01105 | NA | SET | SET nuclear oncogene | 6418 | ENSG00000119335 |
| 6 | P54727 | NA | RAD23B | RAD23 homolog B (S. cerevisiae) | 5887 | ENSG00000119318 |
| 7 | P39687 | NA | ANP32A | acidic (leucine-rich) nuclear phosphoprotein 32 family, member A | 8125 | ENSG00000140350 |
| 8 | O43598 | NA | C6orf108 | chromosome 6 open reading frame 108 | 10591 | ENSG00000112667 |
| 9 | P07741 | NA | APRT | adenine phosphoribosyltransferase | 353 | ENSG00000198931 |
| 10 | Q06323 | NA | PSME1 | proteasome (prosome, macropain) activator subunit 1 (PA28 alpha) | 5720 | ENSG00000092010 |
| 11 | P06744 | NA | GPI | glucose-6-phosphate isomerase | 2821 | ENSG00000105220 |
| 12 | P19367 | NA | HK1 | hexokinase 1 | 3098 | ENSG00000156515 |
| 13 | P30837 | NA | ALDH1B1 | aldehyde dehydrogenase 1 family, member B1 | 219 | ENSG00000137124 |
| 14 | P05783 | NA | KRT18 | keratin 18 | 3875 | ENSG00000111057 |
| 15 | Q13263 | NA | TRIM28 | tripartite motif containing 28 | 10155 | ENSG00000130726 |
| 16 | Q16531 | NA | DDB1 | damage-specific DNA binding protein 1, 127kDa | 1642 | ENSG00000167986 |

  
  

| **Database:cellular component      &nbspName:envelope      &nbspID:GO:0031975** | | | | | | |
| --- | --- | --- | --- | --- | --- | --- |
| C=829; O=8; E=2.24; R=3.58; rawP=0.0015; adjP=0.0027 | | | | | | |
| Index | UserID | Value | Gene Symbol | Gene Name | EntrezGene | Ensembl |
| 1 | P54819 | NA | AK2 | adenylate kinase 2 | 204 | ENSG00000004455 |
| 2 | P17931 | NA | LGALS3 | lectin, galactoside-binding, soluble, 3 | 3958 | ENSG00000131981 |
| 3 | Q10713 | NA | PMPCA | peptidase (mitochondrial processing) alpha | 23203 | ENSG00000165688 |
| 4 | P42765 | NA | ACAA2 | acetyl-CoA acyltransferase 2 | 10449 | ENSG00000167315 |
| 5 | Q99714 | NA | HSD17B10 | hydroxysteroid (17-beta) dehydrogenase 10 | 3028 | ENSG00000072506 |
| 6 | P04179 | NA | SOD2 | superoxide dismutase 2, mitochondrial | 6648 | ENSG00000112096 |
| 7 | P19367 | NA | HK1 | hexokinase 1 | 3098 | ENSG00000156515 |
| 8 | P31040 | NA | SDHA | succinate dehydrogenase complex, subunit A, flavoprotein (Fp) | 6389 | ENSG00000073578 |

  
  

| **Database:cellular component      &nbspName:nuclear part      &nbspID:GO:0044428** | | | | | | |
| --- | --- | --- | --- | --- | --- | --- |
| C=3057; O=17; E=8.25; R=2.06; rawP=0.0017; adjP=0.0029 | | | | | | |
| Index | UserID | Value | Gene Symbol | Gene Name | EntrezGene | Ensembl |
| 1 | P09661 | NA | SNRPA1 | small nuclear ribonucleoprotein polypeptide A' | 6627 | ENSG00000131876 |
| 2 | Q16698 | NA | DECR1 | 2,4-dienoyl CoA reductase 1, mitochondrial | 1666 | ENSG00000104325 |
| 3 | P17931 | NA | LGALS3 | lectin, galactoside-binding, soluble, 3 | 3958 | ENSG00000131981 |
| 4 | P05787 | NA | KRT8 | keratin 8 | 3856 | ENSG00000170421 |
| 5 | Q9UL46 | NA | PSME2 | proteasome (prosome, macropain) activator subunit 2 (PA28 beta) | 5721 | ENSG00000100911 |
| 6 | Q01105 | NA | SET | SET nuclear oncogene | 6418 | ENSG00000119335 |
| 7 | P54727 | NA | RAD23B | RAD23 homolog B (S. cerevisiae) | 5887 | ENSG00000119318 |
| 8 | P39687 | NA | ANP32A | acidic (leucine-rich) nuclear phosphoprotein 32 family, member A | 8125 | ENSG00000140350 |
| 9 | O43598 | NA | C6orf108 | chromosome 6 open reading frame 108 | 10591 | ENSG00000112667 |
| 10 | P07741 | NA | APRT | adenine phosphoribosyltransferase | 353 | ENSG00000198931 |
| 11 | Q06323 | NA | PSME1 | proteasome (prosome, macropain) activator subunit 1 (PA28 alpha) | 5720 | ENSG00000092010 |
| 12 | P06744 | NA | GPI | glucose-6-phosphate isomerase | 2821 | ENSG00000105220 |
| 13 | P19367 | NA | HK1 | hexokinase 1 | 3098 | ENSG00000156515 |
| 14 | P30837 | NA | ALDH1B1 | aldehyde dehydrogenase 1 family, member B1 | 219 | ENSG00000137124 |
| 15 | P05783 | NA | KRT18 | keratin 18 | 3875 | ENSG00000111057 |
| 16 | Q13263 | NA | TRIM28 | tripartite motif containing 28 | 10155 | ENSG00000130726 |
| 17 | Q16531 | NA | DDB1 | damage-specific DNA binding protein 1, 127kDa | 1642 | ENSG00000167986 |

  
  

| **Database:cellular component      &nbspName:nucleoplasm      &nbspID:GO:0005654** | | | | | | |
| --- | --- | --- | --- | --- | --- | --- |
| C=1476; O=9; E=3.98; R=2.26; rawP=0.0156; adjP=0.0256 | | | | | | |
| Index | UserID | Value | Gene Symbol | Gene Name | EntrezGene | Ensembl |
| 1 | P09661 | NA | SNRPA1 | small nuclear ribonucleoprotein polypeptide A' | 6627 | ENSG00000131876 |
| 2 | P05787 | NA | KRT8 | keratin 8 | 3856 | ENSG00000170421 |
| 3 | Q9UL46 | NA | PSME2 | proteasome (prosome, macropain) activator subunit 2 (PA28 beta) | 5721 | ENSG00000100911 |
| 4 | Q01105 | NA | SET | SET nuclear oncogene | 6418 | ENSG00000119335 |
| 5 | P54727 | NA | RAD23B | RAD23 homolog B (S. cerevisiae) | 5887 | ENSG00000119318 |
| 6 | P39687 | NA | ANP32A | acidic (leucine-rich) nuclear phosphoprotein 32 family, member A | 8125 | ENSG00000140350 |
| 7 | Q06323 | NA | PSME1 | proteasome (prosome, macropain) activator subunit 1 (PA28 alpha) | 5720 | ENSG00000092010 |
| 8 | Q13263 | NA | TRIM28 | tripartite motif containing 28 | 10155 | ENSG00000130726 |
| 9 | Q16531 | NA | DDB1 | damage-specific DNA binding protein 1, 127kDa | 1642 | ENSG00000167986 |

  
  

| **Database:cellular component      &nbspName:nucleus      &nbspID:GO:0005634** | | | | | | |
| --- | --- | --- | --- | --- | --- | --- |
| C=5883; O=23; E=15.87; R=1.45; rawP=0.0209; adjP=0.0320 | | | | | | |
| Index | UserID | Value | Gene Symbol | Gene Name | EntrezGene | Ensembl |
| 1 | P09661 | NA | SNRPA1 | small nuclear ribonucleoprotein polypeptide A' | 6627 | ENSG00000131876 |
| 2 | P17931 | NA | LGALS3 | lectin, galactoside-binding, soluble, 3 | 3958 | ENSG00000131981 |
| 3 | P05787 | NA | KRT8 | keratin 8 | 3856 | ENSG00000170421 |
| 4 | P26447 | NA | S100A4 | S100 calcium binding protein A4 | 6275 | ENSG00000196154 |
| 5 | Q9UL46 | NA | PSME2 | proteasome (prosome, macropain) activator subunit 2 (PA28 beta) | 5721 | ENSG00000100911 |
| 6 | P50453 | NA | SERPINB9 | serpin peptidase inhibitor, clade B (ovalbumin), member 9 | 5272 | ENSG00000170542 |
| 7 | P07741 | NA | APRT | adenine phosphoribosyltransferase | 353 | ENSG00000198931 |
| 8 | Q06323 | NA | PSME1 | proteasome (prosome, macropain) activator subunit 1 (PA28 alpha) | 5720 | ENSG00000092010 |
| 9 | P30837 | NA | ALDH1B1 | aldehyde dehydrogenase 1 family, member B1 | 219 | ENSG00000137124 |
| 10 | Q13263 | NA | TRIM28 | tripartite motif containing 28 | 10155 | ENSG00000130726 |
| 11 | Q16698 | NA | DECR1 | 2,4-dienoyl CoA reductase 1, mitochondrial | 1666 | ENSG00000104325 |
| 12 | Q99798 | NA | ACO2 | aconitase 2, mitochondrial | 50 | ENSG00000100412 |
| 13 | Q01105 | NA | SET | SET nuclear oncogene | 6418 | ENSG00000119335 |
| 14 | Q05639 | NA | EEF1A2 | eukaryotic translation elongation factor 1 alpha 2 | 1917 | ENSG00000101210 |
| 15 | P54727 | NA | RAD23B | RAD23 homolog B (S. cerevisiae) | 5887 | ENSG00000119318 |
| 16 | P39687 | NA | ANP32A | acidic (leucine-rich) nuclear phosphoprotein 32 family, member A | 8125 | ENSG00000140350 |
| 17 | O43598 | NA | C6orf108 | chromosome 6 open reading frame 108 | 10591 | ENSG00000112667 |
| 18 | P49419 | NA | ALDH7A1 | aldehyde dehydrogenase 7 family, member A1 | 501 | ENSG00000164904 |
| 19 | P06744 | NA | GPI | glucose-6-phosphate isomerase | 2821 | ENSG00000105220 |
| 20 | P19367 | NA | HK1 | hexokinase 1 | 3098 | ENSG00000156515 |
| 21 | P05783 | NA | KRT18 | keratin 18 | 3875 | ENSG00000111057 |
| 22 | Q16531 | NA | DDB1 | damage-specific DNA binding protein 1, 127kDa | 1642 | ENSG00000167986 |
| 23 | Q5SSJ5 | NA | HP1BP3 | heterochromatin protein 1, binding protein 3 | 50809 | ENSG00000127483 |

  
  

| **Database:cellular component      &nbspName:cytosol      &nbspID:GO:0005829** | | | | | | |
| --- | --- | --- | --- | --- | --- | --- |
| C=2367; O=12; E=6.39; R=1.88; rawP=0.0202; adjP=0.0320 | | | | | | |
| Index | UserID | Value | Gene Symbol | Gene Name | EntrezGene | Ensembl |
| 1 | P02792 | NA | FTL | ferritin, light polypeptide | 2512 | ENSG00000087086 |
| 2 | P00558 | NA | PGK1 | phosphoglycerate kinase 1 | 5230 | ENSG00000102144 |
| 3 | Q9UL46 | NA | PSME2 | proteasome (prosome, macropain) activator subunit 2 (PA28 beta) | 5721 | ENSG00000100911 |
| 4 | Q01105 | NA | SET | SET nuclear oncogene | 6418 | ENSG00000119335 |
| 5 | P50453 | NA | SERPINB9 | serpin peptidase inhibitor, clade B (ovalbumin), member 9 | 5272 | ENSG00000170542 |
| 6 | P06737 | NA | PYGL | phosphorylase, glycogen, liver | 5836 | ENSG00000100504 |
| 7 | P07741 | NA | APRT | adenine phosphoribosyltransferase | 353 | ENSG00000198931 |
| 8 | Q06323 | NA | PSME1 | proteasome (prosome, macropain) activator subunit 1 (PA28 alpha) | 5720 | ENSG00000092010 |
| 9 | P49419 | NA | ALDH7A1 | aldehyde dehydrogenase 7 family, member A1 | 501 | ENSG00000164904 |
| 10 | P06744 | NA | GPI | glucose-6-phosphate isomerase | 2821 | ENSG00000105220 |
| 11 | P19367 | NA | HK1 | hexokinase 1 | 3098 | ENSG00000156515 |
| 12 | P00352 | NA | ALDH1A1 | aldehyde dehydrogenase 1 family, member A1 | 216 | ENSG00000165092 |

  
  

| **Database:cellular component      &nbspName:perinuclear region of cytoplasm      &nbspID:GO:0048471** | | | | | | |
| --- | --- | --- | --- | --- | --- | --- |
| C=467; O=4; E=1.26; R=3.18; rawP=0.0366; adjP=0.0543 | | | | | | |
| Index | UserID | Value | Gene Symbol | Gene Name | EntrezGene | Ensembl |
| 1 | P39687 | NA | ANP32A | acidic (leucine-rich) nuclear phosphoprotein 32 family, member A | 8125 | ENSG00000140350 |
| 2 | P05783 | NA | KRT18 | keratin 18 | 3875 | ENSG00000111057 |
| 3 | P26447 | NA | S100A4 | S100 calcium binding protein A4 | 6275 | ENSG00000196154 |
| 4 | Q01105 | NA | SET | SET nuclear oncogene | 6418 | ENSG00000119335 |

  
  

| **Database:cellular component      &nbspName:nucleolus      &nbspID:GO:0005730** | | | | | | |
| --- | --- | --- | --- | --- | --- | --- |
| C=1539; O=8; E=4.15; R=1.93; rawP=0.0514; adjP=0.0739 | | | | | | |
| Index | UserID | Value | Gene Symbol | Gene Name | EntrezGene | Ensembl |
| 1 | O43598 | NA | C6orf108 | chromosome 6 open reading frame 108 | 10591 | ENSG00000112667 |
| 2 | Q16698 | NA | DECR1 | 2,4-dienoyl CoA reductase 1, mitochondrial | 1666 | ENSG00000104325 |
| 3 | P07741 | NA | APRT | adenine phosphoribosyltransferase | 353 | ENSG00000198931 |
| 4 | P06744 | NA | GPI | glucose-6-phosphate isomerase | 2821 | ENSG00000105220 |
| 5 | P19367 | NA | HK1 | hexokinase 1 | 3098 | ENSG00000156515 |
| 6 | P30837 | NA | ALDH1B1 | aldehyde dehydrogenase 1 family, member B1 | 219 | ENSG00000137124 |
| 7 | P05783 | NA | KRT18 | keratin 18 | 3875 | ENSG00000111057 |
| 8 | P54727 | NA | RAD23B | RAD23 homolog B (S. cerevisiae) | 5887 | ENSG00000119318 |

  
  
  
  

---

WebGestalt is currently developed and maintained by Jing Wang and Bing Zhang at the  Zhang Lab. Other people who have made significant contribution to the project include Dexter Duncan, Stefan Kirov, Zhiao Shi, and Jay Snoddy.  
  
**Funding credits:** NIH/NIAAA (U01 AA016662, U01 AA013512); NIH/NIDA (P01 DA015027); NIH/NIMH (P50 MH078028, P50 MH096972); NIH/NCI (U24 CA159988); NIH/NIGMS (R01 GM088822).
