## Supplementary material for "PGRMC1 phosphorylation and cell plasticity 1: glycolysis, mitochondria, tumor growth": File S8: final_sig_kegg_file_1439190287.html

Anchored HTML File of EIDs

|  |  |
| --- | --- |
|  | WEB-based GEne SeT AnaLysis Toolkit |
|  | ***Translating gene lists into biological insights...*** |

---

  

| **Database:KEGG pathway      &nbspName:Metabolic pathways      &nbspID:01100** | | | | | | |
| --- | --- | --- | --- | --- | --- | --- |
| C=1130; O=15; E=2.51; R=5.98; rawP=1.00e-08; adjP=8.00e-08 | | | | | | |
| Index | UserID | Value | Gene Symbol | Gene Name | EntrezGene | Ensembl |
| 1 | Q9UBQ7 | NA | GRHPR | glyoxylate reductase/hydroxypyruvate reductase | 9380 | ENSG00000137106 |
| 2 | Q99798 | NA | ACO2 | aconitase 2, mitochondrial | 50 | ENSG00000100412 |
| 3 | P10253 | NA | GAA | glucosidase, alpha; acid | 2548 | ENSG00000171298 |
| 4 | P54819 | NA | AK2 | adenylate kinase 2 | 204 | ENSG00000004455 |
| 5 | P00558 | NA | PGK1 | phosphoglycerate kinase 1 | 5230 | ENSG00000102144 |
| 6 | P42765 | NA | ACAA2 | acetyl-CoA acyltransferase 2 | 10449 | ENSG00000167315 |
| 7 | Q99714 | NA | HSD17B10 | hydroxysteroid (17-beta) dehydrogenase 10 | 3028 | ENSG00000072506 |
| 8 | P13674 | NA | P4HA1 | prolyl 4-hydroxylase, alpha polypeptide I | 5033 | ENSG00000122884 |
| 9 | P49419 | NA | ALDH7A1 | aldehyde dehydrogenase 7 family, member A1 | 501 | ENSG00000164904 |
| 10 | P07741 | NA | APRT | adenine phosphoribosyltransferase | 353 | ENSG00000198931 |
| 11 | P06744 | NA | GPI | glucose-6-phosphate isomerase | 2821 | ENSG00000105220 |
| 12 | P19367 | NA | HK1 | hexokinase 1 | 3098 | ENSG00000156515 |
| 13 | P30837 | NA | ALDH1B1 | aldehyde dehydrogenase 1 family, member B1 | 219 | ENSG00000137124 |
| 14 | P00352 | NA | ALDH1A1 | aldehyde dehydrogenase 1 family, member A1 | 216 | ENSG00000165092 |
| 15 | P31040 | NA | SDHA | succinate dehydrogenase complex, subunit A, flavoprotein (Fp) | 6389 | ENSG00000073578 |

  
  

| **Database:KEGG pathway      &nbspName:Glycolysis / Gluconeogenesis      &nbspID:00010** | | | | | | |
| --- | --- | --- | --- | --- | --- | --- |
| C=65; O=5; E=0.14; R=34.67; rawP=3.20e-07; adjP=1.28e-06 | | | | | | |
| Index | UserID | Value | Gene Symbol | Gene Name | EntrezGene | Ensembl |
| 1 | P49419 | NA | ALDH7A1 | aldehyde dehydrogenase 7 family, member A1 | 501 | ENSG00000164904 |
| 2 | P06744 | NA | GPI | glucose-6-phosphate isomerase | 2821 | ENSG00000105220 |
| 3 | P19367 | NA | HK1 | hexokinase 1 | 3098 | ENSG00000156515 |
| 4 | P30837 | NA | ALDH1B1 | aldehyde dehydrogenase 1 family, member B1 | 219 | ENSG00000137124 |
| 5 | P00558 | NA | PGK1 | phosphoglycerate kinase 1 | 5230 | ENSG00000102144 |

  
  

| **Database:KEGG pathway      &nbspName:Valine, leucine and isoleucine degradation      &nbspID:00280** | | | | | | |
| --- | --- | --- | --- | --- | --- | --- |
| C=44; O=4; E=0.10; R=40.98; rawP=2.69e-06; adjP=7.17e-06 | | | | | | |
| Index | UserID | Value | Gene Symbol | Gene Name | EntrezGene | Ensembl |
| 1 | P49419 | NA | ALDH7A1 | aldehyde dehydrogenase 7 family, member A1 | 501 | ENSG00000164904 |
| 2 | P30837 | NA | ALDH1B1 | aldehyde dehydrogenase 1 family, member B1 | 219 | ENSG00000137124 |
| 3 | P42765 | NA | ACAA2 | acetyl-CoA acyltransferase 2 | 10449 | ENSG00000167315 |
| 4 | Q99714 | NA | HSD17B10 | hydroxysteroid (17-beta) dehydrogenase 10 | 3028 | ENSG00000072506 |

  
  

| **Database:KEGG pathway      &nbspName:Starch and sucrose metabolism      &nbspID:00500** | | | | | | |
| --- | --- | --- | --- | --- | --- | --- |
| C=54; O=4; E=0.12; R=33.39; rawP=6.16e-06; adjP=1.23e-05 | | | | | | |
| Index | UserID | Value | Gene Symbol | Gene Name | EntrezGene | Ensembl |
| 1 | P06737 | NA | PYGL | phosphorylase, glycogen, liver | 5836 | ENSG00000100504 |
| 2 | P10253 | NA | GAA | glucosidase, alpha; acid | 2548 | ENSG00000171298 |
| 3 | P06744 | NA | GPI | glucose-6-phosphate isomerase | 2821 | ENSG00000105220 |
| 4 | P19367 | NA | HK1 | hexokinase 1 | 3098 | ENSG00000156515 |

  
  

| **Database:KEGG pathway      &nbspName:Pyruvate metabolism      &nbspID:00620** | | | | | | |
| --- | --- | --- | --- | --- | --- | --- |
| C=40; O=3; E=0.09; R=33.80; rawP=9.52e-05; adjP=0.0001 | | | | | | |
| Index | UserID | Value | Gene Symbol | Gene Name | EntrezGene | Ensembl |
| 1 | P49419 | NA | ALDH7A1 | aldehyde dehydrogenase 7 family, member A1 | 501 | ENSG00000164904 |
| 2 | Q9UBQ7 | NA | GRHPR | glyoxylate reductase/hydroxypyruvate reductase | 9380 | ENSG00000137106 |
| 3 | P30837 | NA | ALDH1B1 | aldehyde dehydrogenase 1 family, member B1 | 219 | ENSG00000137124 |

  
  

| **Database:KEGG pathway      &nbspName:Fatty acid metabolism      &nbspID:00071** | | | | | | |
| --- | --- | --- | --- | --- | --- | --- |
| C=43; O=3; E=0.10; R=31.45; rawP=0.0001; adjP=0.0001 | | | | | | |
| Index | UserID | Value | Gene Symbol | Gene Name | EntrezGene | Ensembl |
| 1 | P49419 | NA | ALDH7A1 | aldehyde dehydrogenase 7 family, member A1 | 501 | ENSG00000164904 |
| 2 | P30837 | NA | ALDH1B1 | aldehyde dehydrogenase 1 family, member B1 | 219 | ENSG00000137124 |
| 3 | P42765 | NA | ACAA2 | acetyl-CoA acyltransferase 2 | 10449 | ENSG00000167315 |

  
  

| **Database:KEGG pathway      &nbspName:Arginine and proline metabolism      &nbspID:00330** | | | | | | |
| --- | --- | --- | --- | --- | --- | --- |
| C=54; O=3; E=0.12; R=25.04; rawP=0.0002; adjP=0.0002 | | | | | | |
| Index | UserID | Value | Gene Symbol | Gene Name | EntrezGene | Ensembl |
| 1 | P13674 | NA | P4HA1 | prolyl 4-hydroxylase, alpha polypeptide I | 5033 | ENSG00000122884 |
| 2 | P49419 | NA | ALDH7A1 | aldehyde dehydrogenase 7 family, member A1 | 501 | ENSG00000164904 |
| 3 | P30837 | NA | ALDH1B1 | aldehyde dehydrogenase 1 family, member B1 | 219 | ENSG00000137124 |

  
  

| **Database:KEGG pathway      &nbspName:Huntington's disease      &nbspID:05016** | | | | | | |
| --- | --- | --- | --- | --- | --- | --- |
| C=183; O=3; E=0.41; R=7.39; rawP=0.0078; adjP=0.0078 | | | | | | |
| Index | UserID | Value | Gene Symbol | Gene Name | EntrezGene | Ensembl |
| 1 | P21980 | NA | TGM2 | transglutaminase 2 (C polypeptide, protein-glutamine-gamma-glutamyltransferase) | 7052 | ENSG00000198959 |
| 2 | P31040 | NA | SDHA | succinate dehydrogenase complex, subunit A, flavoprotein (Fp) | 6389 | ENSG00000073578 |
| 3 | P04179 | NA | SOD2 | superoxide dismutase 2, mitochondrial | 6648 | ENSG00000112096 |

  
  

  
  

---

WebGestalt is currently developed and maintained by Jing Wang and Bing Zhang at the  Zhang Lab. Other people who have made significant contribution to the project include Dexter Duncan, Stefan Kirov, Zhiao Shi, and Jay Snoddy.  
  
**Funding credits:** NIH/NIAAA (U01 AA016662, U01 AA013512); NIH/NIDA (P01 DA015027); NIH/NIMH (P50 MH078028, P50 MH096972); NIH/NCI (U24 CA159988); NIH/NIGMS (R01 GM088822).
