## Supplementary material for "PGRMC1 phosphorylation and cell plasticity 1: glycolysis, mitochondria, tumor growth": File S8: final_pc_geneset_file_1439166893.html

Anchored HTML File of EIDs

|  |  |
| --- | --- |
|  | WEB-based GEne SeT AnaLysis Toolkit |
|  | ***Translating gene lists into biological insights...*** |

---

  

| **Database:Pathway Commons pathway      &nbspName:Metabolism      &nbspID:DB\_ID:634** | | | | | | |
| --- | --- | --- | --- | --- | --- | --- |
| C=824; O=13; E=1.83; R=7.11; rawP=1.63e-08; adjP=8.64e-07 | | | | | | |
| Index | UserID | Value | Gene Symbol | Gene Name | EntrezGene | Ensembl |
| 1 | Q9UBQ7 | NA | GRHPR | glyoxylate reductase/hydroxypyruvate reductase | 9380 | ENSG00000137106 |
| 2 | Q99798 | NA | ACO2 | aconitase 2, mitochondrial | 50 | ENSG00000100412 |
| 3 | P54819 | NA | AK2 | adenylate kinase 2 | 204 | ENSG00000004455 |
| 4 | Q9UL46 | NA | PSME2 | proteasome (prosome, macropain) activator subunit 2 (PA28 beta) | 5721 | ENSG00000100911 |
| 5 | Q01105 | NA | SET | SET nuclear oncogene | 6418 | ENSG00000119335 |
| 6 | Q99714 | NA | HSD17B10 | hydroxysteroid (17-beta) dehydrogenase 10 | 3028 | ENSG00000072506 |
| 7 | P39687 | NA | ANP32A | acidic (leucine-rich) nuclear phosphoprotein 32 family, member A | 8125 | ENSG00000140350 |
| 8 | P49419 | NA | ALDH7A1 | aldehyde dehydrogenase 7 family, member A1 | 501 | ENSG00000164904 |
| 9 | P07741 | NA | APRT | adenine phosphoribosyltransferase | 353 | ENSG00000198931 |
| 10 | Q06323 | NA | PSME1 | proteasome (prosome, macropain) activator subunit 1 (PA28 alpha) | 5720 | ENSG00000092010 |
| 11 | P00352 | NA | ALDH1A1 | aldehyde dehydrogenase 1 family, member A1 | 216 | ENSG00000165092 |
| 12 | P31040 | NA | SDHA | succinate dehydrogenase complex, subunit A, flavoprotein (Fp) | 6389 | ENSG00000073578 |
| 13 | P38117 | NA | ETFB | electron-transfer-flavoprotein, beta polypeptide | 2109 | ENSG00000105379 |

  
  

| **Database:Pathway Commons pathway      &nbspName:Regulation of mRNA Stability by Proteins that Bind AU-rich Elements      &nbspID:DB\_ID:858** | | | | | | |
| --- | --- | --- | --- | --- | --- | --- |
| C=104; O=4; E=0.23; R=17.34; rawP=8.27e-05; adjP=0.0009 | | | | | | |
| Index | UserID | Value | Gene Symbol | Gene Name | EntrezGene | Ensembl |
| 1 | P39687 | NA | ANP32A | acidic (leucine-rich) nuclear phosphoprotein 32 family, member A | 8125 | ENSG00000140350 |
| 2 | Q06323 | NA | PSME1 | proteasome (prosome, macropain) activator subunit 1 (PA28 alpha) | 5720 | ENSG00000092010 |
| 3 | Q9UL46 | NA | PSME2 | proteasome (prosome, macropain) activator subunit 2 (PA28 beta) | 5721 | ENSG00000100911 |
| 4 | Q01105 | NA | SET | SET nuclear oncogene | 6418 | ENSG00000119335 |

  
  

| **Database:Pathway Commons pathway      &nbspName:Metabolism of carbohydrates      &nbspID:DB\_ID:905** | | | | | | |
| --- | --- | --- | --- | --- | --- | --- |
| C=92; O=4; E=0.20; R=19.60; rawP=5.12e-05; adjP=0.0009 | | | | | | |
| Index | UserID | Value | Gene Symbol | Gene Name | EntrezGene | Ensembl |
| 1 | P06737 | NA | PYGL | phosphorylase, glycogen, liver | 5836 | ENSG00000100504 |
| 2 | P06744 | NA | GPI | glucose-6-phosphate isomerase | 2821 | ENSG00000105220 |
| 3 | P19367 | NA | HK1 | hexokinase 1 | 3098 | ENSG00000156515 |
| 4 | P00558 | NA | PGK1 | phosphoglycerate kinase 1 | 5230 | ENSG00000102144 |

  
  

| **Database:Pathway Commons pathway      &nbspName:Glucose metabolism      &nbspID:DB\_ID:898** | | | | | | |
| --- | --- | --- | --- | --- | --- | --- |
| C=36; O=3; E=0.08; R=37.56; rawP=6.92e-05; adjP=0.0009 | | | | | | |
| Index | UserID | Value | Gene Symbol | Gene Name | EntrezGene | Ensembl |
| 1 | P06737 | NA | PYGL | phosphorylase, glycogen, liver | 5836 | ENSG00000100504 |
| 2 | P06744 | NA | GPI | glucose-6-phosphate isomerase | 2821 | ENSG00000105220 |
| 3 | P00558 | NA | PGK1 | phosphoglycerate kinase 1 | 5230 | ENSG00000102144 |

  
  

| **Database:Pathway Commons pathway      &nbspName:Metabolism of amino acids and derivatives      &nbspID:DB\_ID:875** | | | | | | |
| --- | --- | --- | --- | --- | --- | --- |
| C=188; O=5; E=0.42; R=11.99; rawP=5.87e-05; adjP=0.0009 | | | | | | |
| Index | UserID | Value | Gene Symbol | Gene Name | EntrezGene | Ensembl |
| 1 | P49419 | NA | ALDH7A1 | aldehyde dehydrogenase 7 family, member A1 | 501 | ENSG00000164904 |
| 2 | Q9UBQ7 | NA | GRHPR | glyoxylate reductase/hydroxypyruvate reductase | 9380 | ENSG00000137106 |
| 3 | Q06323 | NA | PSME1 | proteasome (prosome, macropain) activator subunit 1 (PA28 alpha) | 5720 | ENSG00000092010 |
| 4 | Q9UL46 | NA | PSME2 | proteasome (prosome, macropain) activator subunit 2 (PA28 beta) | 5721 | ENSG00000100911 |
| 5 | Q99714 | NA | HSD17B10 | hydroxysteroid (17-beta) dehydrogenase 10 | 3028 | ENSG00000072506 |

  
  

| **Database:Pathway Commons pathway      &nbspName:Metabolism of mRNA      &nbspID:DB\_ID:864** | | | | | | |
| --- | --- | --- | --- | --- | --- | --- |
| C=235; O=4; E=0.52; R=7.67; rawP=0.0018; adjP=0.0159 | | | | | | |
| Index | UserID | Value | Gene Symbol | Gene Name | EntrezGene | Ensembl |
| 1 | P39687 | NA | ANP32A | acidic (leucine-rich) nuclear phosphoprotein 32 family, member A | 8125 | ENSG00000140350 |
| 2 | Q06323 | NA | PSME1 | proteasome (prosome, macropain) activator subunit 1 (PA28 alpha) | 5720 | ENSG00000092010 |
| 3 | Q9UL46 | NA | PSME2 | proteasome (prosome, macropain) activator subunit 2 (PA28 beta) | 5721 | ENSG00000100911 |
| 4 | Q01105 | NA | SET | SET nuclear oncogene | 6418 | ENSG00000119335 |

  
  

| **Database:Pathway Commons pathway      &nbspName:The citric acid (TCA) cycle and respiratory electron transport      &nbspID:DB\_ID:547** | | | | | | |
| --- | --- | --- | --- | --- | --- | --- |
| C=118; O=3; E=0.26; R=11.46; rawP=0.0023; adjP=0.0174 | | | | | | |
| Index | UserID | Value | Gene Symbol | Gene Name | EntrezGene | Ensembl |
| 1 | Q99798 | NA | ACO2 | aconitase 2, mitochondrial | 50 | ENSG00000100412 |
| 2 | P31040 | NA | SDHA | succinate dehydrogenase complex, subunit A, flavoprotein (Fp) | 6389 | ENSG00000073578 |
| 3 | P38117 | NA | ETFB | electron-transfer-flavoprotein, beta polypeptide | 2109 | ENSG00000105379 |

  
  

| **Database:Pathway Commons pathway      &nbspName:Metabolism of RNA      &nbspID:DB\_ID:865** | | | | | | |
| --- | --- | --- | --- | --- | --- | --- |
| C=282; O=4; E=0.63; R=6.39; rawP=0.0035; adjP=0.0232 | | | | | | |
| Index | UserID | Value | Gene Symbol | Gene Name | EntrezGene | Ensembl |
| 1 | P39687 | NA | ANP32A | acidic (leucine-rich) nuclear phosphoprotein 32 family, member A | 8125 | ENSG00000140350 |
| 2 | Q06323 | NA | PSME1 | proteasome (prosome, macropain) activator subunit 1 (PA28 alpha) | 5720 | ENSG00000092010 |
| 3 | Q9UL46 | NA | PSME2 | proteasome (prosome, macropain) activator subunit 2 (PA28 beta) | 5721 | ENSG00000100911 |
| 4 | Q01105 | NA | SET | SET nuclear oncogene | 6418 | ENSG00000119335 |

  
  
  
  

---

WebGestalt is currently developed and maintained by Jing Wang and Bing Zhang at the  Zhang Lab. Other people who have made significant contribution to the project include Dexter Duncan, Stefan Kirov, Zhiao Shi, and Jay Snoddy.  
  
**Funding credits:** NIH/NIAAA (U01 AA016662, U01 AA013512); NIH/NIDA (P01 DA015027); NIH/NIMH (P50 MH078028, P50 MH096972); NIH/NCI (U24 CA159988); NIH/NIGMS (R01 GM088822).
