## Supplementary material for "PGRMC1 phosphorylation and cell plasticity 1: glycolysis, mitochondria, tumor growth": File S8: final_tf_geneset_file_1439190287.html

Anchored HTML File of EIDs

|  |  |
| --- | --- |
|  | WEB-based GEne SeT AnaLysis Toolkit |
|  | ***Translating gene lists into biological insights...*** |

---

  

| **Database:Transcrription Target      &nbspName:hsa\_V$SP1\_Q4\_01      &nbspID:DB\_ID:2308** | | | | | | |
| --- | --- | --- | --- | --- | --- | --- |
| C=248; O=3; E=0.55; R=5.45; rawP=0.0176; adjP=0.0478 | | | | | | |
| Index | UserID | Value | Gene Symbol | Gene Name | EntrezGene | Ensembl |
| 1 | P39687 | NA | ANP32A | acidic (leucine-rich) nuclear phosphoprotein 32 family, member A | 8125 | ENSG00000140350 |
| 2 | Q13263 | NA | TRIM28 | tripartite motif containing 28 | 10155 | ENSG00000130726 |
| 3 | Q16531 | NA | DDB1 | damage-specific DNA binding protein 1, 127kDa | 1642 | ENSG00000167986 |

  
  

| **Database:Transcrription Target      &nbspName:hsa\_STTTCRNTTT\_V$IRF\_Q6      &nbspID:DB\_ID:2426** | | | | | | |
| --- | --- | --- | --- | --- | --- | --- |
| C=184; O=3; E=0.41; R=7.35; rawP=0.0079; adjP=0.0478 | | | | | | |
| Index | UserID | Value | Gene Symbol | Gene Name | EntrezGene | Ensembl |
| 1 | Q06323 | NA | PSME1 | proteasome (prosome, macropain) activator subunit 1 (PA28 alpha) | 5720 | ENSG00000092010 |
| 2 | P00352 | NA | ALDH1A1 | aldehyde dehydrogenase 1 family, member A1 | 216 | ENSG00000165092 |
| 3 | Q9UL46 | NA | PSME2 | proteasome (prosome, macropain) activator subunit 2 (PA28 beta) | 5721 | ENSG00000100911 |

  
  

| **Database:Transcrription Target      &nbspName:hsa\_V$SP1\_Q6\_01      &nbspID:DB\_ID:2307** | | | | | | |
| --- | --- | --- | --- | --- | --- | --- |
| C=238; O=3; E=0.53; R=5.68; rawP=0.0158; adjP=0.0478 | | | | | | |
| Index | UserID | Value | Gene Symbol | Gene Name | EntrezGene | Ensembl |
| 1 | P39687 | NA | ANP32A | acidic (leucine-rich) nuclear phosphoprotein 32 family, member A | 8125 | ENSG00000140350 |
| 2 | Q13263 | NA | TRIM28 | tripartite motif containing 28 | 10155 | ENSG00000130726 |
| 3 | Q16531 | NA | DDB1 | damage-specific DNA binding protein 1, 127kDa | 1642 | ENSG00000167986 |

  
  

| **Database:Transcrription Target      &nbspName:hsa\_GGGCGGR\_V$SP1\_Q6      &nbspID:DB\_ID:2452** | | | | | | |
| --- | --- | --- | --- | --- | --- | --- |
| C=2863; O=13; E=6.35; R=2.05; rawP=0.0076; adjP=0.0478 | | | | | | |
| Index | UserID | Value | Gene Symbol | Gene Name | EntrezGene | Ensembl |
| 1 | Q99798 | NA | ACO2 | aconitase 2, mitochondrial | 50 | ENSG00000100412 |
| 2 | P54819 | NA | AK2 | adenylate kinase 2 | 204 | ENSG00000004455 |
| 3 | Q01105 | NA | SET | SET nuclear oncogene | 6418 | ENSG00000119335 |
| 4 | P54727 | NA | RAD23B | RAD23 homolog B (S. cerevisiae) | 5887 | ENSG00000119318 |
| 5 | P13674 | NA | P4HA1 | prolyl 4-hydroxylase, alpha polypeptide I | 5033 | ENSG00000122884 |
| 6 | P39687 | NA | ANP32A | acidic (leucine-rich) nuclear phosphoprotein 32 family, member A | 8125 | ENSG00000140350 |
| 7 | P07741 | NA | APRT | adenine phosphoribosyltransferase | 353 | ENSG00000198931 |
| 8 | P49419 | NA | ALDH7A1 | aldehyde dehydrogenase 7 family, member A1 | 501 | ENSG00000164904 |
| 9 | Q9BQ69 | NA | MACROD1 | MACRO domain containing 1 | 28992 | ENSG00000133315 |
| 10 | Q13263 | NA | TRIM28 | tripartite motif containing 28 | 10155 | ENSG00000130726 |
| 11 | Q16531 | NA | DDB1 | damage-specific DNA binding protein 1, 127kDa | 1642 | ENSG00000167986 |
| 12 | P38117 | NA | ETFB | electron-transfer-flavoprotein, beta polypeptide | 2109 | ENSG00000105379 |
| 13 | Q13011 | NA | ECH1 | enoyl CoA hydratase 1, peroxisomal | 1891 | ENSG00000104823 |

  
  

| **Database:Transcrription Target      &nbspName:hsa\_V$SP1\_Q6      &nbspID:DB\_ID:1986** | | | | | | |
| --- | --- | --- | --- | --- | --- | --- |
| C=248; O=3; E=0.55; R=5.45; rawP=0.0176; adjP=0.0478 | | | | | | |
| Index | UserID | Value | Gene Symbol | Gene Name | EntrezGene | Ensembl |
| 1 | P39687 | NA | ANP32A | acidic (leucine-rich) nuclear phosphoprotein 32 family, member A | 8125 | ENSG00000140350 |
| 2 | Q13263 | NA | TRIM28 | tripartite motif containing 28 | 10155 | ENSG00000130726 |
| 3 | Q16531 | NA | DDB1 | damage-specific DNA binding protein 1, 127kDa | 1642 | ENSG00000167986 |

  
  

| **Database:Transcrription Target      &nbspName:hsa\_RCGCANGCGY\_V$NRF1\_Q6      &nbspID:DB\_ID:2441** | | | | | | |
| --- | --- | --- | --- | --- | --- | --- |
| C=891; O=6; E=1.98; R=3.04; rawP=0.0133; adjP=0.0478 | | | | | | |
| Index | UserID | Value | Gene Symbol | Gene Name | EntrezGene | Ensembl |
| 1 | P13674 | NA | P4HA1 | prolyl 4-hydroxylase, alpha polypeptide I | 5033 | ENSG00000122884 |
| 2 | P39687 | NA | ANP32A | acidic (leucine-rich) nuclear phosphoprotein 32 family, member A | 8125 | ENSG00000140350 |
| 3 | Q13263 | NA | TRIM28 | tripartite motif containing 28 | 10155 | ENSG00000130726 |
| 4 | P31040 | NA | SDHA | succinate dehydrogenase complex, subunit A, flavoprotein (Fp) | 6389 | ENSG00000073578 |
| 5 | Q01105 | NA | SET | SET nuclear oncogene | 6418 | ENSG00000119335 |
| 6 | P54727 | NA | RAD23B | RAD23 homolog B (S. cerevisiae) | 5887 | ENSG00000119318 |

  
  

| **Database:Transcrription Target      &nbspName:hsa\_TGACCTTG\_V$SF1\_Q6      &nbspID:DB\_ID:2450** | | | | | | |
| --- | --- | --- | --- | --- | --- | --- |
| C=237; O=3; E=0.53; R=5.71; rawP=0.0156; adjP=0.0478 | | | | | | |
| Index | UserID | Value | Gene Symbol | Gene Name | EntrezGene | Ensembl |
| 1 | Q99798 | NA | ACO2 | aconitase 2, mitochondrial | 50 | ENSG00000100412 |
| 2 | P54819 | NA | AK2 | adenylate kinase 2 | 204 | ENSG00000004455 |
| 3 | Q05639 | NA | EEF1A2 | eukaryotic translation elongation factor 1 alpha 2 | 1917 | ENSG00000101210 |

  
  
  
  

---

WebGestalt is currently developed and maintained by Jing Wang and Bing Zhang at the  Zhang Lab. Other people who have made significant contribution to the project include Dexter Duncan, Stefan Kirov, Zhiao Shi, and Jay Snoddy.  
  
**Funding credits:** NIH/NIAAA (U01 AA016662, U01 AA013512); NIH/NIDA (P01 DA015027); NIH/NIMH (P50 MH078028, P50 MH096972); NIH/NCI (U24 CA159988); NIH/NIGMS (R01 GM088822).
