## Supplementary material for "PGRMC1 phosphorylation and cell plasticity 1: glycolysis, mitochondria, tumor growth": File S8: final_sig_file_1439167077.html

Anchored HTML File of EIDs

|  |  |
| --- | --- |
|  | WEB-based GEne SeT AnaLysis Toolkit |
|  | ***Translating gene lists into biological insights...*** |

---

  

| **Database:biological process      &nbspName:organic acid transport      &nbspID:GO:0015849** | | | | | | |
| --- | --- | --- | --- | --- | --- | --- |
| C=221; O=5; E=0.57; R=8.82; rawP=0.0002; adjP=0.0277 | | | | | | |
| Index | UserID | Value | Gene Symbol | Gene Name | EntrezGene | Ensembl |
| 1 | Q04828 | NA | AKR1C1 | aldo-keto reductase family 1, member C1 (dihydrodiol dehydrogenase 1; 20-alpha (3-alpha)-hydroxysteroid dehydrogenase) | 1645 | ENSG00000187134 |
| 2 | P08195 | NA | SLC3A2 | solute carrier family 3 (activators of dibasic and neutral amino acid transport), member 2 | 6520 | ENSG00000168003 |
| 3 | Q15758 | NA | SLC1A5 | solute carrier family 1 (neutral amino acid transporter), member 5 | 6510 | ENSG00000105281 |
| 4 | P02768 | NA | ALB | albumin | 213 | ENSG00000163631 |
| 5 | P04083 | NA | ANXA1 | annexin A1 | 301 | ENSG00000135046 |

  
  

| **Database:biological process      &nbspName:carboxylic acid transport      &nbspID:GO:0046942** | | | | | | |
| --- | --- | --- | --- | --- | --- | --- |
| C=219; O=5; E=0.56; R=8.90; rawP=0.0002; adjP=0.0277 | | | | | | |
| Index | UserID | Value | Gene Symbol | Gene Name | EntrezGene | Ensembl |
| 1 | Q04828 | NA | AKR1C1 | aldo-keto reductase family 1, member C1 (dihydrodiol dehydrogenase 1; 20-alpha (3-alpha)-hydroxysteroid dehydrogenase) | 1645 | ENSG00000187134 |
| 2 | P08195 | NA | SLC3A2 | solute carrier family 3 (activators of dibasic and neutral amino acid transport), member 2 | 6520 | ENSG00000168003 |
| 3 | Q15758 | NA | SLC1A5 | solute carrier family 1 (neutral amino acid transporter), member 5 | 6510 | ENSG00000105281 |
| 4 | P02768 | NA | ALB | albumin | 213 | ENSG00000163631 |
| 5 | P04083 | NA | ANXA1 | annexin A1 | 301 | ENSG00000135046 |

  
  

| **Database:biological process      &nbspName:response to inorganic substance      &nbspID:GO:0010035** | | | | | | |
| --- | --- | --- | --- | --- | --- | --- |
| C=423; O=6; E=1.09; R=5.53; rawP=0.0007; adjP=0.0323 | | | | | | |
| Index | UserID | Value | Gene Symbol | Gene Name | EntrezGene | Ensembl |
| 1 | Q16881 | NA | TXNRD1 | thioredoxin reductase 1 | 7296 | ENSG00000198431 |
| 2 | P02795 | NA | MT2A | metallothionein 2A | 4502 | ENSG00000125148 |
| 3 | P22102 | NA | GART | phosphoribosylglycinamide formyltransferase, phosphoribosylglycinamide synthetase, phosphoribosylaminoimidazole synthetase | 2618 | ENSG00000159131 |
| 4 | P02768 | NA | ALB | albumin | 213 | ENSG00000163631 |
| 5 | P02786 | NA | TFRC | transferrin receptor (p90, CD71) | 7037 | ENSG00000072274 |
| 6 | P04083 | NA | ANXA1 | annexin A1 | 301 | ENSG00000135046 |

  
  

| **Database:biological process      &nbspName:organic anion transport      &nbspID:GO:0015711** | | | | | | |
| --- | --- | --- | --- | --- | --- | --- |
| C=264; O=5; E=0.68; R=7.38; rawP=0.0005; adjP=0.0323 | | | | | | |
| Index | UserID | Value | Gene Symbol | Gene Name | EntrezGene | Ensembl |
| 1 | Q04828 | NA | AKR1C1 | aldo-keto reductase family 1, member C1 (dihydrodiol dehydrogenase 1; 20-alpha (3-alpha)-hydroxysteroid dehydrogenase) | 1645 | ENSG00000187134 |
| 2 | P08195 | NA | SLC3A2 | solute carrier family 3 (activators of dibasic and neutral amino acid transport), member 2 | 6520 | ENSG00000168003 |
| 3 | Q15758 | NA | SLC1A5 | solute carrier family 1 (neutral amino acid transporter), member 5 | 6510 | ENSG00000105281 |
| 4 | P02768 | NA | ALB | albumin | 213 | ENSG00000163631 |
| 5 | P04083 | NA | ANXA1 | annexin A1 | 301 | ENSG00000135046 |

  
  

| **Database:biological process      &nbspName:response to chemical stimulus      &nbspID:GO:0042221** | | | | | | |
| --- | --- | --- | --- | --- | --- | --- |
| C=2741; O=16; E=7.03; R=2.28; rawP=0.0006; adjP=0.0323 | | | | | | |
| Index | UserID | Value | Gene Symbol | Gene Name | EntrezGene | Ensembl |
| 1 | Q16881 | NA | TXNRD1 | thioredoxin reductase 1 | 7296 | ENSG00000198431 |
| 2 | Q9NR30 | NA | DDX21 | DEAD (Asp-Glu-Ala-Asp) box helicase 21 | 9188 | ENSG00000165732 |
| 3 | P02795 | NA | MT2A | metallothionein 2A | 4502 | ENSG00000125148 |
| 4 | P22102 | NA | GART | phosphoribosylglycinamide formyltransferase, phosphoribosylglycinamide synthetase, phosphoribosylaminoimidazole synthetase | 2618 | ENSG00000159131 |
| 5 | P00491 | NA | PNP | purine nucleoside phosphorylase | 4860 | ENSG00000198805 |
| 6 | P16615 | NA | ATP2A2 | ATPase, Ca++ transporting, cardiac muscle, slow twitch 2 | 488 | ENSG00000174437 |
| 7 | P02768 | NA | ALB | albumin | 213 | ENSG00000163631 |
| 8 | P05556 | NA | ITGB1 | integrin, beta 1 (fibronectin receptor, beta polypeptide, antigen CD29 includes MDF2, MSK12) | 3688 | ENSG00000150093 |
| 9 | Q04828 | NA | AKR1C1 | aldo-keto reductase family 1, member C1 (dihydrodiol dehydrogenase 1; 20-alpha (3-alpha)-hydroxysteroid dehydrogenase) | 1645 | ENSG00000187134 |
| 10 | P60842 | NA | EIF4A1 | eukaryotic translation initiation factor 4A1 | 1973 | ENSG00000161960 |
| 11 | P17174 | NA | GOT1 | glutamic-oxaloacetic transaminase 1, soluble (aspartate aminotransferase 1) | 2805 | ENSG00000120053 |
| 12 | P49588 | NA | AARS | alanyl-tRNA synthetase | 16 | ENSG00000090861 |
| 13 | O75369 | NA | FLNB | filamin B, beta | 2317 | ENSG00000136068 |
| 14 | P02786 | NA | TFRC | transferrin receptor (p90, CD71) | 7037 | ENSG00000072274 |
| 15 | P04083 | NA | ANXA1 | annexin A1 | 301 | ENSG00000135046 |
| 16 | P11387 | NA | TOP1 | topoisomerase (DNA) I | 7150 | ENSG00000198900 |

  
  

| **Database:biological process      &nbspName:response to organic substance      &nbspID:GO:0010033** | | | | | | |
| --- | --- | --- | --- | --- | --- | --- |
| C=1814; O=13; E=4.65; R=2.79; rawP=0.0004; adjP=0.0323 | | | | | | |
| Index | UserID | Value | Gene Symbol | Gene Name | EntrezGene | Ensembl |
| 1 | Q9NR30 | NA | DDX21 | DEAD (Asp-Glu-Ala-Asp) box helicase 21 | 9188 | ENSG00000165732 |
| 2 | P02795 | NA | MT2A | metallothionein 2A | 4502 | ENSG00000125148 |
| 3 | P22102 | NA | GART | phosphoribosylglycinamide formyltransferase, phosphoribosylglycinamide synthetase, phosphoribosylaminoimidazole synthetase | 2618 | ENSG00000159131 |
| 4 | P16615 | NA | ATP2A2 | ATPase, Ca++ transporting, cardiac muscle, slow twitch 2 | 488 | ENSG00000174437 |
| 5 | P02768 | NA | ALB | albumin | 213 | ENSG00000163631 |
| 6 | P05556 | NA | ITGB1 | integrin, beta 1 (fibronectin receptor, beta polypeptide, antigen CD29 includes MDF2, MSK12) | 3688 | ENSG00000150093 |
| 7 | Q04828 | NA | AKR1C1 | aldo-keto reductase family 1, member C1 (dihydrodiol dehydrogenase 1; 20-alpha (3-alpha)-hydroxysteroid dehydrogenase) | 1645 | ENSG00000187134 |
| 8 | P60842 | NA | EIF4A1 | eukaryotic translation initiation factor 4A1 | 1973 | ENSG00000161960 |
| 9 | P17174 | NA | GOT1 | glutamic-oxaloacetic transaminase 1, soluble (aspartate aminotransferase 1) | 2805 | ENSG00000120053 |
| 10 | P49588 | NA | AARS | alanyl-tRNA synthetase | 16 | ENSG00000090861 |
| 11 | O75369 | NA | FLNB | filamin B, beta | 2317 | ENSG00000136068 |
| 12 | P02786 | NA | TFRC | transferrin receptor (p90, CD71) | 7037 | ENSG00000072274 |
| 13 | P04083 | NA | ANXA1 | annexin A1 | 301 | ENSG00000135046 |

  
  

| **Database:biological process      &nbspName:translation      &nbspID:GO:0006412** | | | | | | |
| --- | --- | --- | --- | --- | --- | --- |
| C=832; O=8; E=2.13; R=3.75; rawP=0.0010; adjP=0.0396 | | | | | | |
| Index | UserID | Value | Gene Symbol | Gene Name | EntrezGene | Ensembl |
| 1 | P60842 | NA | EIF4A1 | eukaryotic translation initiation factor 4A1 | 1973 | ENSG00000161960 |
| 2 | P41091 | NA | EIF2S3 | eukaryotic translation initiation factor 2, subunit 3 gamma, 52kDa | 1968 | ENSG00000130741 |
| 3 | P68104 | NA | EEF1A1 | eukaryotic translation elongation factor 1 alpha 1 | 1915 | ENSG00000156508 |
| 4 | P49588 | NA | AARS | alanyl-tRNA synthetase | 16 | ENSG00000090861 |
| 5 | P26639 | NA | TARS | threonyl-tRNA synthetase | 6897 | ENSG00000113407 |
| 6 | P29692 | NA | EEF1D | eukaryotic translation elongation factor 1 delta (guanine nucleotide exchange protein) | 1936 | ENSG00000104529 |
| 7 | P24534 | NA | EEF1B2 | eukaryotic translation elongation factor 1 beta 2 | 1933 | ENSG00000114942 |
| 8 | O00425 | NA | IGF2BP3 | insulin-like growth factor 2 mRNA binding protein 3 | 10643 | ENSG00000136231 |

  
  

| **Database:biological process      &nbspName:anion transport      &nbspID:GO:0006820** | | | | | | |
| --- | --- | --- | --- | --- | --- | --- |
| C=315; O=5; E=0.81; R=6.19; rawP=0.0012; adjP=0.0415 | | | | | | |
| Index | UserID | Value | Gene Symbol | Gene Name | EntrezGene | Ensembl |
| 1 | Q04828 | NA | AKR1C1 | aldo-keto reductase family 1, member C1 (dihydrodiol dehydrogenase 1; 20-alpha (3-alpha)-hydroxysteroid dehydrogenase) | 1645 | ENSG00000187134 |
| 2 | P08195 | NA | SLC3A2 | solute carrier family 3 (activators of dibasic and neutral amino acid transport), member 2 | 6520 | ENSG00000168003 |
| 3 | Q15758 | NA | SLC1A5 | solute carrier family 1 (neutral amino acid transporter), member 5 | 6510 | ENSG00000105281 |
| 4 | P02768 | NA | ALB | albumin | 213 | ENSG00000163631 |
| 5 | P04083 | NA | ANXA1 | annexin A1 | 301 | ENSG00000135046 |

  
  

| **Database:biological process      &nbspName:response to stimulus      &nbspID:GO:0050896** | | | | | | |
| --- | --- | --- | --- | --- | --- | --- |
| C=6561; O=26; E=16.83; R=1.54; rawP=0.0020; adjP=0.0582 | | | | | | |
| Index | UserID | Value | Gene Symbol | Gene Name | EntrezGene | Ensembl |
| 1 | Q16881 | NA | TXNRD1 | thioredoxin reductase 1 | 7296 | ENSG00000198431 |
| 2 | Q9NR30 | NA | DDX21 | DEAD (Asp-Glu-Ala-Asp) box helicase 21 | 9188 | ENSG00000165732 |
| 3 | P02795 | NA | MT2A | metallothionein 2A | 4502 | ENSG00000125148 |
| 4 | P08195 | NA | SLC3A2 | solute carrier family 3 (activators of dibasic and neutral amino acid transport), member 2 | 6520 | ENSG00000168003 |
| 5 | P00491 | NA | PNP | purine nucleoside phosphorylase | 4860 | ENSG00000198805 |
| 6 | P68371 | NA | TUBB4B | tubulin, beta 4B class IVb | 10383 | ENSG00000188229 |
| 7 | P09525 | NA | ANXA4 | annexin A4 | 307 | ENSG00000196975 |
| 8 | P61225 | NA | RAP2B | RAP2B, member of RAS oncogene family | 5912 | ENSG00000181467 |
| 9 | Q04828 | NA | AKR1C1 | aldo-keto reductase family 1, member C1 (dihydrodiol dehydrogenase 1; 20-alpha (3-alpha)-hydroxysteroid dehydrogenase) | 1645 | ENSG00000187134 |
| 10 | P60842 | NA | EIF4A1 | eukaryotic translation initiation factor 4A1 | 1973 | ENSG00000161960 |
| 11 | P04080 | NA | CSTB | cystatin B (stefin B) | 1476 | ENSG00000160213 |
| 12 | P49588 | NA | AARS | alanyl-tRNA synthetase | 16 | ENSG00000090861 |
| 13 | O75369 | NA | FLNB | filamin B, beta | 2317 | ENSG00000136068 |
| 14 | P02786 | NA | TFRC | transferrin receptor (p90, CD71) | 7037 | ENSG00000072274 |
| 15 | P04083 | NA | ANXA1 | annexin A1 | 301 | ENSG00000135046 |
| 16 | P17655 | NA | CAPN2 | calpain 2, (m/II) large subunit | 824 | ENSG00000162909 |
| 17 | P22102 | NA | GART | phosphoribosylglycinamide formyltransferase, phosphoribosylglycinamide synthetase, phosphoribosylaminoimidazole synthetase | 2618 | ENSG00000159131 |
| 18 | P16615 | NA | ATP2A2 | ATPase, Ca++ transporting, cardiac muscle, slow twitch 2 | 488 | ENSG00000174437 |
| 19 | P02768 | NA | ALB | albumin | 213 | ENSG00000163631 |
| 20 | P05556 | NA | ITGB1 | integrin, beta 1 (fibronectin receptor, beta polypeptide, antigen CD29 includes MDF2, MSK12) | 3688 | ENSG00000150093 |
| 21 | Q92974 | NA | ARHGEF2 | Rho/Rac guanine nucleotide exchange factor (GEF) 2 | 9181 | ENSG00000116584 |
| 22 | P17174 | NA | GOT1 | glutamic-oxaloacetic transaminase 1, soluble (aspartate aminotransferase 1) | 2805 | ENSG00000120053 |
| 23 | O00194 | NA | RAB27B | RAB27B, member RAS oncogene family | 5874 | ENSG00000041353 |
| 24 | P18206 | NA | VCL | vinculin | 7414 | ENSG00000035403 |
| 25 | P29692 | NA | EEF1D | eukaryotic translation elongation factor 1 delta (guanine nucleotide exchange protein) | 1936 | ENSG00000104529 |
| 26 | P11387 | NA | TOP1 | topoisomerase (DNA) I | 7150 | ENSG00000198900 |

  
  

| **Database:biological process      &nbspName:monocarboxylic acid transport      &nbspID:GO:0015718** | | | | | | |
| --- | --- | --- | --- | --- | --- | --- |
| C=99; O=3; E=0.25; R=11.81; rawP=0.0021; adjP=0.0582 | | | | | | |
| Index | UserID | Value | Gene Symbol | Gene Name | EntrezGene | Ensembl |
| 1 | Q04828 | NA | AKR1C1 | aldo-keto reductase family 1, member C1 (dihydrodiol dehydrogenase 1; 20-alpha (3-alpha)-hydroxysteroid dehydrogenase) | 1645 | ENSG00000187134 |
| 2 | P02768 | NA | ALB | albumin | 213 | ENSG00000163631 |
| 3 | P04083 | NA | ANXA1 | annexin A1 | 301 | ENSG00000135046 |

  
  

| **Database:biological process      &nbspName:translational elongation      &nbspID:GO:0006414** | | | | | | |
| --- | --- | --- | --- | --- | --- | --- |
| C=108; O=3; E=0.28; R=10.83; rawP=0.0026; adjP=0.0600 | | | | | | |
| Index | UserID | Value | Gene Symbol | Gene Name | EntrezGene | Ensembl |
| 1 | P68104 | NA | EEF1A1 | eukaryotic translation elongation factor 1 alpha 1 | 1915 | ENSG00000156508 |
| 2 | P29692 | NA | EEF1D | eukaryotic translation elongation factor 1 delta (guanine nucleotide exchange protein) | 1936 | ENSG00000104529 |
| 3 | P24534 | NA | EEF1B2 | eukaryotic translation elongation factor 1 beta 2 | 1933 | ENSG00000114942 |

  
  

| **Database:biological process      &nbspName:response to drug      &nbspID:GO:0042493** | | | | | | |
| --- | --- | --- | --- | --- | --- | --- |
| C=371; O=5; E=0.95; R=5.25; rawP=0.0024; adjP=0.0600 | | | | | | |
| Index | UserID | Value | Gene Symbol | Gene Name | EntrezGene | Ensembl |
| 1 | P02795 | NA | MT2A | metallothionein 2A | 4502 | ENSG00000125148 |
| 2 | P00491 | NA | PNP | purine nucleoside phosphorylase | 4860 | ENSG00000198805 |
| 3 | P05556 | NA | ITGB1 | integrin, beta 1 (fibronectin receptor, beta polypeptide, antigen CD29 includes MDF2, MSK12) | 3688 | ENSG00000150093 |
| 4 | P04083 | NA | ANXA1 | annexin A1 | 301 | ENSG00000135046 |
| 5 | P11387 | NA | TOP1 | topoisomerase (DNA) I | 7150 | ENSG00000198900 |

  
  

| **Database:molecular function      &nbspName:translation factor activity, nucleic acid binding      &nbspID:GO:0008135** | | | | | | |
| --- | --- | --- | --- | --- | --- | --- |
| C=79; O=6; E=0.20; R=30.74; rawP=3.62e-08; adjP=2.57e-06 | | | | | | |
| Index | UserID | Value | Gene Symbol | Gene Name | EntrezGene | Ensembl |
| 1 | P60842 | NA | EIF4A1 | eukaryotic translation initiation factor 4A1 | 1973 | ENSG00000161960 |
| 2 | P56537 | NA | EIF6 | eukaryotic translation initiation factor 6 | 3692 | ENSG00000242372 |
| 3 | P41091 | NA | EIF2S3 | eukaryotic translation initiation factor 2, subunit 3 gamma, 52kDa | 1968 | ENSG00000130741 |
| 4 | P68104 | NA | EEF1A1 | eukaryotic translation elongation factor 1 alpha 1 | 1915 | ENSG00000156508 |
| 5 | P29692 | NA | EEF1D | eukaryotic translation elongation factor 1 delta (guanine nucleotide exchange protein) | 1936 | ENSG00000104529 |
| 6 | P24534 | NA | EEF1B2 | eukaryotic translation elongation factor 1 beta 2 | 1933 | ENSG00000114942 |

  
  

| **Database:molecular function      &nbspName:translation elongation factor activity      &nbspID:GO:0003746** | | | | | | |
| --- | --- | --- | --- | --- | --- | --- |
| C=17; O=3; E=0.04; R=71.43; rawP=9.22e-06; adjP=0.0003 | | | | | | |
| Index | UserID | Value | Gene Symbol | Gene Name | EntrezGene | Ensembl |
| 1 | P68104 | NA | EEF1A1 | eukaryotic translation elongation factor 1 alpha 1 | 1915 | ENSG00000156508 |
| 2 | P29692 | NA | EEF1D | eukaryotic translation elongation factor 1 delta (guanine nucleotide exchange protein) | 1936 | ENSG00000104529 |
| 3 | P24534 | NA | EEF1B2 | eukaryotic translation elongation factor 1 beta 2 | 1933 | ENSG00000114942 |

  
  

| **Database:molecular function      &nbspName:anion binding      &nbspID:GO:0043168** | | | | | | |
| --- | --- | --- | --- | --- | --- | --- |
| C=2371; O=17; E=5.86; R=2.90; rawP=1.52e-05; adjP=0.0004 | | | | | | |
| Index | UserID | Value | Gene Symbol | Gene Name | EntrezGene | Ensembl |
| 1 | Q16881 | NA | TXNRD1 | thioredoxin reductase 1 | 7296 | ENSG00000198431 |
| 2 | Q9NR30 | NA | DDX21 | DEAD (Asp-Glu-Ala-Asp) box helicase 21 | 9188 | ENSG00000165732 |
| 3 | P00491 | NA | PNP | purine nucleoside phosphorylase | 4860 | ENSG00000198805 |
| 4 | P68371 | NA | TUBB4B | tubulin, beta 4B class IVb | 10383 | ENSG00000188229 |
| 5 | P61225 | NA | RAP2B | RAP2B, member of RAS oncogene family | 5912 | ENSG00000181467 |
| 6 | Q04828 | NA | AKR1C1 | aldo-keto reductase family 1, member C1 (dihydrodiol dehydrogenase 1; 20-alpha (3-alpha)-hydroxysteroid dehydrogenase) | 1645 | ENSG00000187134 |
| 7 | P60842 | NA | EIF4A1 | eukaryotic translation initiation factor 4A1 | 1973 | ENSG00000161960 |
| 8 | P49588 | NA | AARS | alanyl-tRNA synthetase | 16 | ENSG00000090861 |
| 9 | P68104 | NA | EEF1A1 | eukaryotic translation elongation factor 1 alpha 1 | 1915 | ENSG00000156508 |
| 10 | P26639 | NA | TARS | threonyl-tRNA synthetase | 6897 | ENSG00000113407 |
| 11 | P22102 | NA | GART | phosphoribosylglycinamide formyltransferase, phosphoribosylglycinamide synthetase, phosphoribosylaminoimidazole synthetase | 2618 | ENSG00000159131 |
| 12 | P16615 | NA | ATP2A2 | ATPase, Ca++ transporting, cardiac muscle, slow twitch 2 | 488 | ENSG00000174437 |
| 13 | P41091 | NA | EIF2S3 | eukaryotic translation initiation factor 2, subunit 3 gamma, 52kDa | 1968 | ENSG00000130741 |
| 14 | P02768 | NA | ALB | albumin | 213 | ENSG00000163631 |
| 15 | P17174 | NA | GOT1 | glutamic-oxaloacetic transaminase 1, soluble (aspartate aminotransferase 1) | 2805 | ENSG00000120053 |
| 16 | O00194 | NA | RAB27B | RAB27B, member RAS oncogene family | 5874 | ENSG00000041353 |
| 17 | P11387 | NA | TOP1 | topoisomerase (DNA) I | 7150 | ENSG00000198900 |

  
  

| **Database:molecular function      &nbspName:RNA binding      &nbspID:GO:0003723** | | | | | | |
| --- | --- | --- | --- | --- | --- | --- |
| C=843; O=10; E=2.08; R=4.80; rawP=2.62e-05; adjP=0.0005 | | | | | | |
| Index | UserID | Value | Gene Symbol | Gene Name | EntrezGene | Ensembl |
| 1 | Q9NR30 | NA | DDX21 | DEAD (Asp-Glu-Ala-Asp) box helicase 21 | 9188 | ENSG00000165732 |
| 2 | P56537 | NA | EIF6 | eukaryotic translation initiation factor 6 | 3692 | ENSG00000242372 |
| 3 | P41091 | NA | EIF2S3 | eukaryotic translation initiation factor 2, subunit 3 gamma, 52kDa | 1968 | ENSG00000130741 |
| 4 | P60842 | NA | EIF4A1 | eukaryotic translation initiation factor 4A1 | 1973 | ENSG00000161960 |
| 5 | P49588 | NA | AARS | alanyl-tRNA synthetase | 16 | ENSG00000090861 |
| 6 | P68104 | NA | EEF1A1 | eukaryotic translation elongation factor 1 alpha 1 | 1915 | ENSG00000156508 |
| 7 | P29692 | NA | EEF1D | eukaryotic translation elongation factor 1 delta (guanine nucleotide exchange protein) | 1936 | ENSG00000104529 |
| 8 | P04083 | NA | ANXA1 | annexin A1 | 301 | ENSG00000135046 |
| 9 | O00425 | NA | IGF2BP3 | insulin-like growth factor 2 mRNA binding protein 3 | 10643 | ENSG00000136231 |
| 10 | P24534 | NA | EEF1B2 | eukaryotic translation elongation factor 1 beta 2 | 1933 | ENSG00000114942 |

  
  

| **Database:molecular function      &nbspName:small molecule binding      &nbspID:GO:0036094** | | | | | | |
| --- | --- | --- | --- | --- | --- | --- |
| C=2595; O=17; E=6.41; R=2.65; rawP=5.10e-05; adjP=0.0007 | | | | | | |
| Index | UserID | Value | Gene Symbol | Gene Name | EntrezGene | Ensembl |
| 1 | Q16881 | NA | TXNRD1 | thioredoxin reductase 1 | 7296 | ENSG00000198431 |
| 2 | Q9NR30 | NA | DDX21 | DEAD (Asp-Glu-Ala-Asp) box helicase 21 | 9188 | ENSG00000165732 |
| 3 | P00491 | NA | PNP | purine nucleoside phosphorylase | 4860 | ENSG00000198805 |
| 4 | P68371 | NA | TUBB4B | tubulin, beta 4B class IVb | 10383 | ENSG00000188229 |
| 5 | P61225 | NA | RAP2B | RAP2B, member of RAS oncogene family | 5912 | ENSG00000181467 |
| 6 | P60842 | NA | EIF4A1 | eukaryotic translation initiation factor 4A1 | 1973 | ENSG00000161960 |
| 7 | P49588 | NA | AARS | alanyl-tRNA synthetase | 16 | ENSG00000090861 |
| 8 | P68104 | NA | EEF1A1 | eukaryotic translation elongation factor 1 alpha 1 | 1915 | ENSG00000156508 |
| 9 | P26639 | NA | TARS | threonyl-tRNA synthetase | 6897 | ENSG00000113407 |
| 10 | O00425 | NA | IGF2BP3 | insulin-like growth factor 2 mRNA binding protein 3 | 10643 | ENSG00000136231 |
| 11 | P22102 | NA | GART | phosphoribosylglycinamide formyltransferase, phosphoribosylglycinamide synthetase, phosphoribosylaminoimidazole synthetase | 2618 | ENSG00000159131 |
| 12 | P16615 | NA | ATP2A2 | ATPase, Ca++ transporting, cardiac muscle, slow twitch 2 | 488 | ENSG00000174437 |
| 13 | P41091 | NA | EIF2S3 | eukaryotic translation initiation factor 2, subunit 3 gamma, 52kDa | 1968 | ENSG00000130741 |
| 14 | P02768 | NA | ALB | albumin | 213 | ENSG00000163631 |
| 15 | P17174 | NA | GOT1 | glutamic-oxaloacetic transaminase 1, soluble (aspartate aminotransferase 1) | 2805 | ENSG00000120053 |
| 16 | O00194 | NA | RAB27B | RAB27B, member RAS oncogene family | 5874 | ENSG00000041353 |
| 17 | P11387 | NA | TOP1 | topoisomerase (DNA) I | 7150 | ENSG00000198900 |

  
  

| **Database:molecular function      &nbspName:nucleoside-triphosphatase activity      &nbspID:GO:0017111** | | | | | | |
| --- | --- | --- | --- | --- | --- | --- |
| C=749; O=9; E=1.85; R=4.86; rawP=6.55e-05; adjP=0.0008 | | | | | | |
| Index | UserID | Value | Gene Symbol | Gene Name | EntrezGene | Ensembl |
| 1 | Q9NR30 | NA | DDX21 | DEAD (Asp-Glu-Ala-Asp) box helicase 21 | 9188 | ENSG00000165732 |
| 2 | P16615 | NA | ATP2A2 | ATPase, Ca++ transporting, cardiac muscle, slow twitch 2 | 488 | ENSG00000174437 |
| 3 | P41091 | NA | EIF2S3 | eukaryotic translation initiation factor 2, subunit 3 gamma, 52kDa | 1968 | ENSG00000130741 |
| 4 | P68371 | NA | TUBB4B | tubulin, beta 4B class IVb | 10383 | ENSG00000188229 |
| 5 | P61225 | NA | RAP2B | RAP2B, member of RAS oncogene family | 5912 | ENSG00000181467 |
| 6 | P60842 | NA | EIF4A1 | eukaryotic translation initiation factor 4A1 | 1973 | ENSG00000161960 |
| 7 | O00194 | NA | RAB27B | RAB27B, member RAS oncogene family | 5874 | ENSG00000041353 |
| 8 | P68104 | NA | EEF1A1 | eukaryotic translation elongation factor 1 alpha 1 | 1915 | ENSG00000156508 |
| 9 | P11387 | NA | TOP1 | topoisomerase (DNA) I | 7150 | ENSG00000198900 |

  
  

| **Database:molecular function      &nbspName:hydrolase activity, acting on acid anhydrides      &nbspID:GO:0016817** | | | | | | |
| --- | --- | --- | --- | --- | --- | --- |
| C=791; O=9; E=1.95; R=4.61; rawP=9.98e-05; adjP=0.0008 | | | | | | |
| Index | UserID | Value | Gene Symbol | Gene Name | EntrezGene | Ensembl |
| 1 | Q9NR30 | NA | DDX21 | DEAD (Asp-Glu-Ala-Asp) box helicase 21 | 9188 | ENSG00000165732 |
| 2 | P16615 | NA | ATP2A2 | ATPase, Ca++ transporting, cardiac muscle, slow twitch 2 | 488 | ENSG00000174437 |
| 3 | P41091 | NA | EIF2S3 | eukaryotic translation initiation factor 2, subunit 3 gamma, 52kDa | 1968 | ENSG00000130741 |
| 4 | P68371 | NA | TUBB4B | tubulin, beta 4B class IVb | 10383 | ENSG00000188229 |
| 5 | P61225 | NA | RAP2B | RAP2B, member of RAS oncogene family | 5912 | ENSG00000181467 |
| 6 | P60842 | NA | EIF4A1 | eukaryotic translation initiation factor 4A1 | 1973 | ENSG00000161960 |
| 7 | O00194 | NA | RAB27B | RAB27B, member RAS oncogene family | 5874 | ENSG00000041353 |
| 8 | P68104 | NA | EEF1A1 | eukaryotic translation elongation factor 1 alpha 1 | 1915 | ENSG00000156508 |
| 9 | P11387 | NA | TOP1 | topoisomerase (DNA) I | 7150 | ENSG00000198900 |

  
  

| **Database:molecular function      &nbspName:pyrophosphatase activity      &nbspID:GO:0016462** | | | | | | |
| --- | --- | --- | --- | --- | --- | --- |
| C=783; O=9; E=1.93; R=4.65; rawP=9.23e-05; adjP=0.0008 | | | | | | |
| Index | UserID | Value | Gene Symbol | Gene Name | EntrezGene | Ensembl |
| 1 | Q9NR30 | NA | DDX21 | DEAD (Asp-Glu-Ala-Asp) box helicase 21 | 9188 | ENSG00000165732 |
| 2 | P16615 | NA | ATP2A2 | ATPase, Ca++ transporting, cardiac muscle, slow twitch 2 | 488 | ENSG00000174437 |
| 3 | P41091 | NA | EIF2S3 | eukaryotic translation initiation factor 2, subunit 3 gamma, 52kDa | 1968 | ENSG00000130741 |
| 4 | P68371 | NA | TUBB4B | tubulin, beta 4B class IVb | 10383 | ENSG00000188229 |
| 5 | P61225 | NA | RAP2B | RAP2B, member of RAS oncogene family | 5912 | ENSG00000181467 |
| 6 | P60842 | NA | EIF4A1 | eukaryotic translation initiation factor 4A1 | 1973 | ENSG00000161960 |
| 7 | O00194 | NA | RAB27B | RAB27B, member RAS oncogene family | 5874 | ENSG00000041353 |
| 8 | P68104 | NA | EEF1A1 | eukaryotic translation elongation factor 1 alpha 1 | 1915 | ENSG00000156508 |
| 9 | P11387 | NA | TOP1 | topoisomerase (DNA) I | 7150 | ENSG00000198900 |

  
  

| **Database:molecular function      &nbspName:hydrolase activity, acting on acid anhydrides, in phosphorus-containing anhydrides      &nbspID:GO:0016818** | | | | | | |
| --- | --- | --- | --- | --- | --- | --- |
| C=786; O=9; E=1.94; R=4.63; rawP=9.50e-05; adjP=0.0008 | | | | | | |
| Index | UserID | Value | Gene Symbol | Gene Name | EntrezGene | Ensembl |
| 1 | Q9NR30 | NA | DDX21 | DEAD (Asp-Glu-Ala-Asp) box helicase 21 | 9188 | ENSG00000165732 |
| 2 | P16615 | NA | ATP2A2 | ATPase, Ca++ transporting, cardiac muscle, slow twitch 2 | 488 | ENSG00000174437 |
| 3 | P41091 | NA | EIF2S3 | eukaryotic translation initiation factor 2, subunit 3 gamma, 52kDa | 1968 | ENSG00000130741 |
| 4 | P68371 | NA | TUBB4B | tubulin, beta 4B class IVb | 10383 | ENSG00000188229 |
| 5 | P61225 | NA | RAP2B | RAP2B, member of RAS oncogene family | 5912 | ENSG00000181467 |
| 6 | P60842 | NA | EIF4A1 | eukaryotic translation initiation factor 4A1 | 1973 | ENSG00000161960 |
| 7 | O00194 | NA | RAB27B | RAB27B, member RAS oncogene family | 5874 | ENSG00000041353 |
| 8 | P68104 | NA | EEF1A1 | eukaryotic translation elongation factor 1 alpha 1 | 1915 | ENSG00000156508 |
| 9 | P11387 | NA | TOP1 | topoisomerase (DNA) I | 7150 | ENSG00000198900 |

  
  

| **Database:molecular function      &nbspName:GTPase activity      &nbspID:GO:0003924** | | | | | | |
| --- | --- | --- | --- | --- | --- | --- |
| C=225; O=5; E=0.56; R=9.00; rawP=0.0002; adjP=0.0013 | | | | | | |
| Index | UserID | Value | Gene Symbol | Gene Name | EntrezGene | Ensembl |
| 1 | O00194 | NA | RAB27B | RAB27B, member RAS oncogene family | 5874 | ENSG00000041353 |
| 2 | P41091 | NA | EIF2S3 | eukaryotic translation initiation factor 2, subunit 3 gamma, 52kDa | 1968 | ENSG00000130741 |
| 3 | P68371 | NA | TUBB4B | tubulin, beta 4B class IVb | 10383 | ENSG00000188229 |
| 4 | P68104 | NA | EEF1A1 | eukaryotic translation elongation factor 1 alpha 1 | 1915 | ENSG00000156508 |
| 5 | P61225 | NA | RAP2B | RAP2B, member of RAS oncogene family | 5912 | ENSG00000181467 |

  
  

| **Database:molecular function      &nbspName:translation initiation factor activity      &nbspID:GO:0003743** | | | | | | |
| --- | --- | --- | --- | --- | --- | --- |
| C=49; O=3; E=0.12; R=24.78; rawP=0.0002; adjP=0.0013 | | | | | | |
| Index | UserID | Value | Gene Symbol | Gene Name | EntrezGene | Ensembl |
| 1 | P60842 | NA | EIF4A1 | eukaryotic translation initiation factor 4A1 | 1973 | ENSG00000161960 |
| 2 | P56537 | NA | EIF6 | eukaryotic translation initiation factor 6 | 3692 | ENSG00000242372 |
| 3 | P41091 | NA | EIF2S3 | eukaryotic translation initiation factor 2, subunit 3 gamma, 52kDa | 1968 | ENSG00000130741 |

  
  

| **Database:molecular function      &nbspName:nucleoside binding      &nbspID:GO:0001882** | | | | | | |
| --- | --- | --- | --- | --- | --- | --- |
| C=1823; O=13; E=4.50; R=2.89; rawP=0.0003; adjP=0.0018 | | | | | | |
| Index | UserID | Value | Gene Symbol | Gene Name | EntrezGene | Ensembl |
| 1 | Q9NR30 | NA | DDX21 | DEAD (Asp-Glu-Ala-Asp) box helicase 21 | 9188 | ENSG00000165732 |
| 2 | P22102 | NA | GART | phosphoribosylglycinamide formyltransferase, phosphoribosylglycinamide synthetase, phosphoribosylaminoimidazole synthetase | 2618 | ENSG00000159131 |
| 3 | P00491 | NA | PNP | purine nucleoside phosphorylase | 4860 | ENSG00000198805 |
| 4 | P16615 | NA | ATP2A2 | ATPase, Ca++ transporting, cardiac muscle, slow twitch 2 | 488 | ENSG00000174437 |
| 5 | P41091 | NA | EIF2S3 | eukaryotic translation initiation factor 2, subunit 3 gamma, 52kDa | 1968 | ENSG00000130741 |
| 6 | P68371 | NA | TUBB4B | tubulin, beta 4B class IVb | 10383 | ENSG00000188229 |
| 7 | P61225 | NA | RAP2B | RAP2B, member of RAS oncogene family | 5912 | ENSG00000181467 |
| 8 | P60842 | NA | EIF4A1 | eukaryotic translation initiation factor 4A1 | 1973 | ENSG00000161960 |
| 9 | O00194 | NA | RAB27B | RAB27B, member RAS oncogene family | 5874 | ENSG00000041353 |
| 10 | P49588 | NA | AARS | alanyl-tRNA synthetase | 16 | ENSG00000090861 |
| 11 | P68104 | NA | EEF1A1 | eukaryotic translation elongation factor 1 alpha 1 | 1915 | ENSG00000156508 |
| 12 | P26639 | NA | TARS | threonyl-tRNA synthetase | 6897 | ENSG00000113407 |
| 13 | P11387 | NA | TOP1 | topoisomerase (DNA) I | 7150 | ENSG00000198900 |

  
  

| **Database:molecular function      &nbspName:nucleotide binding      &nbspID:GO:0000166** | | | | | | |
| --- | --- | --- | --- | --- | --- | --- |
| C=2403; O=14; E=5.94; R=2.36; rawP=0.0011; adjP=0.0039 | | | | | | |
| Index | UserID | Value | Gene Symbol | Gene Name | EntrezGene | Ensembl |
| 1 | Q16881 | NA | TXNRD1 | thioredoxin reductase 1 | 7296 | ENSG00000198431 |
| 2 | Q9NR30 | NA | DDX21 | DEAD (Asp-Glu-Ala-Asp) box helicase 21 | 9188 | ENSG00000165732 |
| 3 | P22102 | NA | GART | phosphoribosylglycinamide formyltransferase, phosphoribosylglycinamide synthetase, phosphoribosylaminoimidazole synthetase | 2618 | ENSG00000159131 |
| 4 | P16615 | NA | ATP2A2 | ATPase, Ca++ transporting, cardiac muscle, slow twitch 2 | 488 | ENSG00000174437 |
| 5 | P41091 | NA | EIF2S3 | eukaryotic translation initiation factor 2, subunit 3 gamma, 52kDa | 1968 | ENSG00000130741 |
| 6 | P68371 | NA | TUBB4B | tubulin, beta 4B class IVb | 10383 | ENSG00000188229 |
| 7 | P61225 | NA | RAP2B | RAP2B, member of RAS oncogene family | 5912 | ENSG00000181467 |
| 8 | P60842 | NA | EIF4A1 | eukaryotic translation initiation factor 4A1 | 1973 | ENSG00000161960 |
| 9 | O00194 | NA | RAB27B | RAB27B, member RAS oncogene family | 5874 | ENSG00000041353 |
| 10 | P49588 | NA | AARS | alanyl-tRNA synthetase | 16 | ENSG00000090861 |
| 11 | P68104 | NA | EEF1A1 | eukaryotic translation elongation factor 1 alpha 1 | 1915 | ENSG00000156508 |
| 12 | P26639 | NA | TARS | threonyl-tRNA synthetase | 6897 | ENSG00000113407 |
| 13 | P11387 | NA | TOP1 | topoisomerase (DNA) I | 7150 | ENSG00000198900 |
| 14 | O00425 | NA | IGF2BP3 | insulin-like growth factor 2 mRNA binding protein 3 | 10643 | ENSG00000136231 |

  
  

| **Database:molecular function      &nbspName:purine ribonucleoside triphosphate binding      &nbspID:GO:0035639** | | | | | | |
| --- | --- | --- | --- | --- | --- | --- |
| C=1800; O=12; E=4.45; R=2.70; rawP=0.0009; adjP=0.0039 | | | | | | |
| Index | UserID | Value | Gene Symbol | Gene Name | EntrezGene | Ensembl |
| 1 | Q9NR30 | NA | DDX21 | DEAD (Asp-Glu-Ala-Asp) box helicase 21 | 9188 | ENSG00000165732 |
| 2 | P22102 | NA | GART | phosphoribosylglycinamide formyltransferase, phosphoribosylglycinamide synthetase, phosphoribosylaminoimidazole synthetase | 2618 | ENSG00000159131 |
| 3 | P16615 | NA | ATP2A2 | ATPase, Ca++ transporting, cardiac muscle, slow twitch 2 | 488 | ENSG00000174437 |
| 4 | P41091 | NA | EIF2S3 | eukaryotic translation initiation factor 2, subunit 3 gamma, 52kDa | 1968 | ENSG00000130741 |
| 5 | P68371 | NA | TUBB4B | tubulin, beta 4B class IVb | 10383 | ENSG00000188229 |
| 6 | P61225 | NA | RAP2B | RAP2B, member of RAS oncogene family | 5912 | ENSG00000181467 |
| 7 | P60842 | NA | EIF4A1 | eukaryotic translation initiation factor 4A1 | 1973 | ENSG00000161960 |
| 8 | O00194 | NA | RAB27B | RAB27B, member RAS oncogene family | 5874 | ENSG00000041353 |
| 9 | P49588 | NA | AARS | alanyl-tRNA synthetase | 16 | ENSG00000090861 |
| 10 | P68104 | NA | EEF1A1 | eukaryotic translation elongation factor 1 alpha 1 | 1915 | ENSG00000156508 |
| 11 | P26639 | NA | TARS | threonyl-tRNA synthetase | 6897 | ENSG00000113407 |
| 12 | P11387 | NA | TOP1 | topoisomerase (DNA) I | 7150 | ENSG00000198900 |

  
  

| **Database:molecular function      &nbspName:purine ribonucleotide binding      &nbspID:GO:0032555** | | | | | | |
| --- | --- | --- | --- | --- | --- | --- |
| C=1835; O=12; E=4.53; R=2.65; rawP=0.0011; adjP=0.0039 | | | | | | |
| Index | UserID | Value | Gene Symbol | Gene Name | EntrezGene | Ensembl |
| 1 | Q9NR30 | NA | DDX21 | DEAD (Asp-Glu-Ala-Asp) box helicase 21 | 9188 | ENSG00000165732 |
| 2 | P22102 | NA | GART | phosphoribosylglycinamide formyltransferase, phosphoribosylglycinamide synthetase, phosphoribosylaminoimidazole synthetase | 2618 | ENSG00000159131 |
| 3 | P16615 | NA | ATP2A2 | ATPase, Ca++ transporting, cardiac muscle, slow twitch 2 | 488 | ENSG00000174437 |
| 4 | P41091 | NA | EIF2S3 | eukaryotic translation initiation factor 2, subunit 3 gamma, 52kDa | 1968 | ENSG00000130741 |
| 5 | P68371 | NA | TUBB4B | tubulin, beta 4B class IVb | 10383 | ENSG00000188229 |
| 6 | P61225 | NA | RAP2B | RAP2B, member of RAS oncogene family | 5912 | ENSG00000181467 |
| 7 | P60842 | NA | EIF4A1 | eukaryotic translation initiation factor 4A1 | 1973 | ENSG00000161960 |
| 8 | O00194 | NA | RAB27B | RAB27B, member RAS oncogene family | 5874 | ENSG00000041353 |
| 9 | P49588 | NA | AARS | alanyl-tRNA synthetase | 16 | ENSG00000090861 |
| 10 | P68104 | NA | EEF1A1 | eukaryotic translation elongation factor 1 alpha 1 | 1915 | ENSG00000156508 |
| 11 | P26639 | NA | TARS | threonyl-tRNA synthetase | 6897 | ENSG00000113407 |
| 12 | P11387 | NA | TOP1 | topoisomerase (DNA) I | 7150 | ENSG00000198900 |

  
  

| **Database:molecular function      &nbspName:purine nucleotide binding      &nbspID:GO:0017076** | | | | | | |
| --- | --- | --- | --- | --- | --- | --- |
| C=1842; O=12; E=4.55; R=2.64; rawP=0.0011; adjP=0.0039 | | | | | | |
| Index | UserID | Value | Gene Symbol | Gene Name | EntrezGene | Ensembl |
| 1 | Q9NR30 | NA | DDX21 | DEAD (Asp-Glu-Ala-Asp) box helicase 21 | 9188 | ENSG00000165732 |
| 2 | P22102 | NA | GART | phosphoribosylglycinamide formyltransferase, phosphoribosylglycinamide synthetase, phosphoribosylaminoimidazole synthetase | 2618 | ENSG00000159131 |
| 3 | P16615 | NA | ATP2A2 | ATPase, Ca++ transporting, cardiac muscle, slow twitch 2 | 488 | ENSG00000174437 |
| 4 | P41091 | NA | EIF2S3 | eukaryotic translation initiation factor 2, subunit 3 gamma, 52kDa | 1968 | ENSG00000130741 |
| 5 | P68371 | NA | TUBB4B | tubulin, beta 4B class IVb | 10383 | ENSG00000188229 |
| 6 | P61225 | NA | RAP2B | RAP2B, member of RAS oncogene family | 5912 | ENSG00000181467 |
| 7 | P60842 | NA | EIF4A1 | eukaryotic translation initiation factor 4A1 | 1973 | ENSG00000161960 |
| 8 | O00194 | NA | RAB27B | RAB27B, member RAS oncogene family | 5874 | ENSG00000041353 |
| 9 | P49588 | NA | AARS | alanyl-tRNA synthetase | 16 | ENSG00000090861 |
| 10 | P68104 | NA | EEF1A1 | eukaryotic translation elongation factor 1 alpha 1 | 1915 | ENSG00000156508 |
| 11 | P26639 | NA | TARS | threonyl-tRNA synthetase | 6897 | ENSG00000113407 |
| 12 | P11387 | NA | TOP1 | topoisomerase (DNA) I | 7150 | ENSG00000198900 |

  
  

| **Database:molecular function      &nbspName:purine nucleoside binding      &nbspID:GO:0001883** | | | | | | |
| --- | --- | --- | --- | --- | --- | --- |
| C=1812; O=12; E=4.48; R=2.68; rawP=0.0010; adjP=0.0039 | | | | | | |
| Index | UserID | Value | Gene Symbol | Gene Name | EntrezGene | Ensembl |
| 1 | Q9NR30 | NA | DDX21 | DEAD (Asp-Glu-Ala-Asp) box helicase 21 | 9188 | ENSG00000165732 |
| 2 | P22102 | NA | GART | phosphoribosylglycinamide formyltransferase, phosphoribosylglycinamide synthetase, phosphoribosylaminoimidazole synthetase | 2618 | ENSG00000159131 |
| 3 | P16615 | NA | ATP2A2 | ATPase, Ca++ transporting, cardiac muscle, slow twitch 2 | 488 | ENSG00000174437 |
| 4 | P41091 | NA | EIF2S3 | eukaryotic translation initiation factor 2, subunit 3 gamma, 52kDa | 1968 | ENSG00000130741 |
| 5 | P68371 | NA | TUBB4B | tubulin, beta 4B class IVb | 10383 | ENSG00000188229 |
| 6 | P61225 | NA | RAP2B | RAP2B, member of RAS oncogene family | 5912 | ENSG00000181467 |
| 7 | P60842 | NA | EIF4A1 | eukaryotic translation initiation factor 4A1 | 1973 | ENSG00000161960 |
| 8 | O00194 | NA | RAB27B | RAB27B, member RAS oncogene family | 5874 | ENSG00000041353 |
| 9 | P49588 | NA | AARS | alanyl-tRNA synthetase | 16 | ENSG00000090861 |
| 10 | P68104 | NA | EEF1A1 | eukaryotic translation elongation factor 1 alpha 1 | 1915 | ENSG00000156508 |
| 11 | P26639 | NA | TARS | threonyl-tRNA synthetase | 6897 | ENSG00000113407 |
| 12 | P11387 | NA | TOP1 | topoisomerase (DNA) I | 7150 | ENSG00000198900 |

  
  

| **Database:molecular function      &nbspName:purine ribonucleoside binding      &nbspID:GO:0032550** | | | | | | |
| --- | --- | --- | --- | --- | --- | --- |
| C=1809; O=12; E=4.47; R=2.69; rawP=0.0009; adjP=0.0039 | | | | | | |
| Index | UserID | Value | Gene Symbol | Gene Name | EntrezGene | Ensembl |
| 1 | Q9NR30 | NA | DDX21 | DEAD (Asp-Glu-Ala-Asp) box helicase 21 | 9188 | ENSG00000165732 |
| 2 | P22102 | NA | GART | phosphoribosylglycinamide formyltransferase, phosphoribosylglycinamide synthetase, phosphoribosylaminoimidazole synthetase | 2618 | ENSG00000159131 |
| 3 | P16615 | NA | ATP2A2 | ATPase, Ca++ transporting, cardiac muscle, slow twitch 2 | 488 | ENSG00000174437 |
| 4 | P41091 | NA | EIF2S3 | eukaryotic translation initiation factor 2, subunit 3 gamma, 52kDa | 1968 | ENSG00000130741 |
| 5 | P68371 | NA | TUBB4B | tubulin, beta 4B class IVb | 10383 | ENSG00000188229 |
| 6 | P61225 | NA | RAP2B | RAP2B, member of RAS oncogene family | 5912 | ENSG00000181467 |
| 7 | P60842 | NA | EIF4A1 | eukaryotic translation initiation factor 4A1 | 1973 | ENSG00000161960 |
| 8 | O00194 | NA | RAB27B | RAB27B, member RAS oncogene family | 5874 | ENSG00000041353 |
| 9 | P49588 | NA | AARS | alanyl-tRNA synthetase | 16 | ENSG00000090861 |
| 10 | P68104 | NA | EEF1A1 | eukaryotic translation elongation factor 1 alpha 1 | 1915 | ENSG00000156508 |
| 11 | P26639 | NA | TARS | threonyl-tRNA synthetase | 6897 | ENSG00000113407 |
| 12 | P11387 | NA | TOP1 | topoisomerase (DNA) I | 7150 | ENSG00000198900 |

  
  

| **Database:molecular function      &nbspName:ribonucleoside binding      &nbspID:GO:0032549** | | | | | | |
| --- | --- | --- | --- | --- | --- | --- |
| C=1813; O=12; E=4.48; R=2.68; rawP=0.0010; adjP=0.0039 | | | | | | |
| Index | UserID | Value | Gene Symbol | Gene Name | EntrezGene | Ensembl |
| 1 | Q9NR30 | NA | DDX21 | DEAD (Asp-Glu-Ala-Asp) box helicase 21 | 9188 | ENSG00000165732 |
| 2 | P22102 | NA | GART | phosphoribosylglycinamide formyltransferase, phosphoribosylglycinamide synthetase, phosphoribosylaminoimidazole synthetase | 2618 | ENSG00000159131 |
| 3 | P16615 | NA | ATP2A2 | ATPase, Ca++ transporting, cardiac muscle, slow twitch 2 | 488 | ENSG00000174437 |
| 4 | P41091 | NA | EIF2S3 | eukaryotic translation initiation factor 2, subunit 3 gamma, 52kDa | 1968 | ENSG00000130741 |
| 5 | P68371 | NA | TUBB4B | tubulin, beta 4B class IVb | 10383 | ENSG00000188229 |
| 6 | P61225 | NA | RAP2B | RAP2B, member of RAS oncogene family | 5912 | ENSG00000181467 |
| 7 | P60842 | NA | EIF4A1 | eukaryotic translation initiation factor 4A1 | 1973 | ENSG00000161960 |
| 8 | O00194 | NA | RAB27B | RAB27B, member RAS oncogene family | 5874 | ENSG00000041353 |
| 9 | P49588 | NA | AARS | alanyl-tRNA synthetase | 16 | ENSG00000090861 |
| 10 | P68104 | NA | EEF1A1 | eukaryotic translation elongation factor 1 alpha 1 | 1915 | ENSG00000156508 |
| 11 | P26639 | NA | TARS | threonyl-tRNA synthetase | 6897 | ENSG00000113407 |
| 12 | P11387 | NA | TOP1 | topoisomerase (DNA) I | 7150 | ENSG00000198900 |

  
  

| **Database:molecular function      &nbspName:nucleoside phosphate binding      &nbspID:GO:1901265** | | | | | | |
| --- | --- | --- | --- | --- | --- | --- |
| C=2404; O=14; E=5.94; R=2.36; rawP=0.0011; adjP=0.0039 | | | | | | |
| Index | UserID | Value | Gene Symbol | Gene Name | EntrezGene | Ensembl |
| 1 | Q16881 | NA | TXNRD1 | thioredoxin reductase 1 | 7296 | ENSG00000198431 |
| 2 | Q9NR30 | NA | DDX21 | DEAD (Asp-Glu-Ala-Asp) box helicase 21 | 9188 | ENSG00000165732 |
| 3 | P22102 | NA | GART | phosphoribosylglycinamide formyltransferase, phosphoribosylglycinamide synthetase, phosphoribosylaminoimidazole synthetase | 2618 | ENSG00000159131 |
| 4 | P16615 | NA | ATP2A2 | ATPase, Ca++ transporting, cardiac muscle, slow twitch 2 | 488 | ENSG00000174437 |
| 5 | P41091 | NA | EIF2S3 | eukaryotic translation initiation factor 2, subunit 3 gamma, 52kDa | 1968 | ENSG00000130741 |
| 6 | P68371 | NA | TUBB4B | tubulin, beta 4B class IVb | 10383 | ENSG00000188229 |
| 7 | P61225 | NA | RAP2B | RAP2B, member of RAS oncogene family | 5912 | ENSG00000181467 |
| 8 | P60842 | NA | EIF4A1 | eukaryotic translation initiation factor 4A1 | 1973 | ENSG00000161960 |
| 9 | O00194 | NA | RAB27B | RAB27B, member RAS oncogene family | 5874 | ENSG00000041353 |
| 10 | P49588 | NA | AARS | alanyl-tRNA synthetase | 16 | ENSG00000090861 |
| 11 | P68104 | NA | EEF1A1 | eukaryotic translation elongation factor 1 alpha 1 | 1915 | ENSG00000156508 |
| 12 | P26639 | NA | TARS | threonyl-tRNA synthetase | 6897 | ENSG00000113407 |
| 13 | P11387 | NA | TOP1 | topoisomerase (DNA) I | 7150 | ENSG00000198900 |
| 14 | O00425 | NA | IGF2BP3 | insulin-like growth factor 2 mRNA binding protein 3 | 10643 | ENSG00000136231 |

  
  

| **Database:molecular function      &nbspName:ribonucleotide binding      &nbspID:GO:0032553** | | | | | | |
| --- | --- | --- | --- | --- | --- | --- |
| C=1850; O=12; E=4.57; R=2.63; rawP=0.0012; adjP=0.0041 | | | | | | |
| Index | UserID | Value | Gene Symbol | Gene Name | EntrezGene | Ensembl |
| 1 | Q9NR30 | NA | DDX21 | DEAD (Asp-Glu-Ala-Asp) box helicase 21 | 9188 | ENSG00000165732 |
| 2 | P22102 | NA | GART | phosphoribosylglycinamide formyltransferase, phosphoribosylglycinamide synthetase, phosphoribosylaminoimidazole synthetase | 2618 | ENSG00000159131 |
| 3 | P16615 | NA | ATP2A2 | ATPase, Ca++ transporting, cardiac muscle, slow twitch 2 | 488 | ENSG00000174437 |
| 4 | P41091 | NA | EIF2S3 | eukaryotic translation initiation factor 2, subunit 3 gamma, 52kDa | 1968 | ENSG00000130741 |
| 5 | P68371 | NA | TUBB4B | tubulin, beta 4B class IVb | 10383 | ENSG00000188229 |
| 6 | P61225 | NA | RAP2B | RAP2B, member of RAS oncogene family | 5912 | ENSG00000181467 |
| 7 | P60842 | NA | EIF4A1 | eukaryotic translation initiation factor 4A1 | 1973 | ENSG00000161960 |
| 8 | O00194 | NA | RAB27B | RAB27B, member RAS oncogene family | 5874 | ENSG00000041353 |
| 9 | P49588 | NA | AARS | alanyl-tRNA synthetase | 16 | ENSG00000090861 |
| 10 | P68104 | NA | EEF1A1 | eukaryotic translation elongation factor 1 alpha 1 | 1915 | ENSG00000156508 |
| 11 | P26639 | NA | TARS | threonyl-tRNA synthetase | 6897 | ENSG00000113407 |
| 12 | P11387 | NA | TOP1 | topoisomerase (DNA) I | 7150 | ENSG00000198900 |

  
  

| **Database:molecular function      &nbspName:carboxylic acid binding      &nbspID:GO:0031406** | | | | | | |
| --- | --- | --- | --- | --- | --- | --- |
| C=194; O=4; E=0.48; R=8.35; rawP=0.0013; adjP=0.0042 | | | | | | |
| Index | UserID | Value | Gene Symbol | Gene Name | EntrezGene | Ensembl |
| 1 | Q04828 | NA | AKR1C1 | aldo-keto reductase family 1, member C1 (dihydrodiol dehydrogenase 1; 20-alpha (3-alpha)-hydroxysteroid dehydrogenase) | 1645 | ENSG00000187134 |
| 2 | P17174 | NA | GOT1 | glutamic-oxaloacetic transaminase 1, soluble (aspartate aminotransferase 1) | 2805 | ENSG00000120053 |
| 3 | P02768 | NA | ALB | albumin | 213 | ENSG00000163631 |
| 4 | P49588 | NA | AARS | alanyl-tRNA synthetase | 16 | ENSG00000090861 |

  
  

| **Database:molecular function      &nbspName:drug binding      &nbspID:GO:0008144** | | | | | | |
| --- | --- | --- | --- | --- | --- | --- |
| C=89; O=3; E=0.22; R=13.64; rawP=0.0014; adjP=0.0043 | | | | | | |
| Index | UserID | Value | Gene Symbol | Gene Name | EntrezGene | Ensembl |
| 1 | P02795 | NA | MT2A | metallothionein 2A | 4502 | ENSG00000125148 |
| 2 | P00491 | NA | PNP | purine nucleoside phosphorylase | 4860 | ENSG00000198805 |
| 3 | P02768 | NA | ALB | albumin | 213 | ENSG00000163631 |

  
  

| **Database:molecular function      &nbspName:GTP binding      &nbspID:GO:0005525** | | | | | | |
| --- | --- | --- | --- | --- | --- | --- |
| C=370; O=5; E=0.91; R=5.47; rawP=0.0020; adjP=0.0059 | | | | | | |
| Index | UserID | Value | Gene Symbol | Gene Name | EntrezGene | Ensembl |
| 1 | O00194 | NA | RAB27B | RAB27B, member RAS oncogene family | 5874 | ENSG00000041353 |
| 2 | P41091 | NA | EIF2S3 | eukaryotic translation initiation factor 2, subunit 3 gamma, 52kDa | 1968 | ENSG00000130741 |
| 3 | P68371 | NA | TUBB4B | tubulin, beta 4B class IVb | 10383 | ENSG00000188229 |
| 4 | P68104 | NA | EEF1A1 | eukaryotic translation elongation factor 1 alpha 1 | 1915 | ENSG00000156508 |
| 5 | P61225 | NA | RAP2B | RAP2B, member of RAS oncogene family | 5912 | ENSG00000181467 |

  
  

| **Database:molecular function      &nbspName:guanyl nucleotide binding      &nbspID:GO:0019001** | | | | | | |
| --- | --- | --- | --- | --- | --- | --- |
| C=383; O=5; E=0.95; R=5.28; rawP=0.0024; adjP=0.0066 | | | | | | |
| Index | UserID | Value | Gene Symbol | Gene Name | EntrezGene | Ensembl |
| 1 | O00194 | NA | RAB27B | RAB27B, member RAS oncogene family | 5874 | ENSG00000041353 |
| 2 | P41091 | NA | EIF2S3 | eukaryotic translation initiation factor 2, subunit 3 gamma, 52kDa | 1968 | ENSG00000130741 |
| 3 | P68371 | NA | TUBB4B | tubulin, beta 4B class IVb | 10383 | ENSG00000188229 |
| 4 | P68104 | NA | EEF1A1 | eukaryotic translation elongation factor 1 alpha 1 | 1915 | ENSG00000156508 |
| 5 | P61225 | NA | RAP2B | RAP2B, member of RAS oncogene family | 5912 | ENSG00000181467 |

  
  

| **Database:molecular function      &nbspName:guanyl ribonucleotide binding      &nbspID:GO:0032561** | | | | | | |
| --- | --- | --- | --- | --- | --- | --- |
| C=383; O=5; E=0.95; R=5.28; rawP=0.0024; adjP=0.0066 | | | | | | |
| Index | UserID | Value | Gene Symbol | Gene Name | EntrezGene | Ensembl |
| 1 | O00194 | NA | RAB27B | RAB27B, member RAS oncogene family | 5874 | ENSG00000041353 |
| 2 | P41091 | NA | EIF2S3 | eukaryotic translation initiation factor 2, subunit 3 gamma, 52kDa | 1968 | ENSG00000130741 |
| 3 | P68371 | NA | TUBB4B | tubulin, beta 4B class IVb | 10383 | ENSG00000188229 |
| 4 | P68104 | NA | EEF1A1 | eukaryotic translation elongation factor 1 alpha 1 | 1915 | ENSG00000156508 |
| 5 | P61225 | NA | RAP2B | RAP2B, member of RAS oncogene family | 5912 | ENSG00000181467 |

  
  

| **Database:molecular function      &nbspName:organic cyclic compound binding      &nbspID:GO:0097159** | | | | | | |
| --- | --- | --- | --- | --- | --- | --- |
| C=5342; O=22; E=13.20; R=1.67; rawP=0.0027; adjP=0.0068 | | | | | | |
| Index | UserID | Value | Gene Symbol | Gene Name | EntrezGene | Ensembl |
| 1 | Q16881 | NA | TXNRD1 | thioredoxin reductase 1 | 7296 | ENSG00000198431 |
| 2 | Q9NR30 | NA | DDX21 | DEAD (Asp-Glu-Ala-Asp) box helicase 21 | 9188 | ENSG00000165732 |
| 3 | P00491 | NA | PNP | purine nucleoside phosphorylase | 4860 | ENSG00000198805 |
| 4 | P68371 | NA | TUBB4B | tubulin, beta 4B class IVb | 10383 | ENSG00000188229 |
| 5 | P61225 | NA | RAP2B | RAP2B, member of RAS oncogene family | 5912 | ENSG00000181467 |
| 6 | Q04828 | NA | AKR1C1 | aldo-keto reductase family 1, member C1 (dihydrodiol dehydrogenase 1; 20-alpha (3-alpha)-hydroxysteroid dehydrogenase) | 1645 | ENSG00000187134 |
| 7 | P60842 | NA | EIF4A1 | eukaryotic translation initiation factor 4A1 | 1973 | ENSG00000161960 |
| 8 | P49588 | NA | AARS | alanyl-tRNA synthetase | 16 | ENSG00000090861 |
| 9 | P68104 | NA | EEF1A1 | eukaryotic translation elongation factor 1 alpha 1 | 1915 | ENSG00000156508 |
| 10 | P26639 | NA | TARS | threonyl-tRNA synthetase | 6897 | ENSG00000113407 |
| 11 | P04083 | NA | ANXA1 | annexin A1 | 301 | ENSG00000135046 |
| 12 | O00425 | NA | IGF2BP3 | insulin-like growth factor 2 mRNA binding protein 3 | 10643 | ENSG00000136231 |
| 13 | P24534 | NA | EEF1B2 | eukaryotic translation elongation factor 1 beta 2 | 1933 | ENSG00000114942 |
| 14 | P56537 | NA | EIF6 | eukaryotic translation initiation factor 6 | 3692 | ENSG00000242372 |
| 15 | P22102 | NA | GART | phosphoribosylglycinamide formyltransferase, phosphoribosylglycinamide synthetase, phosphoribosylaminoimidazole synthetase | 2618 | ENSG00000159131 |
| 16 | P16615 | NA | ATP2A2 | ATPase, Ca++ transporting, cardiac muscle, slow twitch 2 | 488 | ENSG00000174437 |
| 17 | P41091 | NA | EIF2S3 | eukaryotic translation initiation factor 2, subunit 3 gamma, 52kDa | 1968 | ENSG00000130741 |
| 18 | P02768 | NA | ALB | albumin | 213 | ENSG00000163631 |
| 19 | P17174 | NA | GOT1 | glutamic-oxaloacetic transaminase 1, soluble (aspartate aminotransferase 1) | 2805 | ENSG00000120053 |
| 20 | O00194 | NA | RAB27B | RAB27B, member RAS oncogene family | 5874 | ENSG00000041353 |
| 21 | P29692 | NA | EEF1D | eukaryotic translation elongation factor 1 delta (guanine nucleotide exchange protein) | 1936 | ENSG00000104529 |
| 22 | P11387 | NA | TOP1 | topoisomerase (DNA) I | 7150 | ENSG00000198900 |

  
  

| **Database:molecular function      &nbspName:ion binding      &nbspID:GO:0043167** | | | | | | |
| --- | --- | --- | --- | --- | --- | --- |
| C=5721; O=23; E=14.13; R=1.63; rawP=0.0027; adjP=0.0068 | | | | | | |
| Index | UserID | Value | Gene Symbol | Gene Name | EntrezGene | Ensembl |
| 1 | Q16881 | NA | TXNRD1 | thioredoxin reductase 1 | 7296 | ENSG00000198431 |
| 2 | Q9NR30 | NA | DDX21 | DEAD (Asp-Glu-Ala-Asp) box helicase 21 | 9188 | ENSG00000165732 |
| 3 | P02795 | NA | MT2A | metallothionein 2A | 4502 | ENSG00000125148 |
| 4 | P08195 | NA | SLC3A2 | solute carrier family 3 (activators of dibasic and neutral amino acid transport), member 2 | 6520 | ENSG00000168003 |
| 5 | P00491 | NA | PNP | purine nucleoside phosphorylase | 4860 | ENSG00000198805 |
| 6 | P68371 | NA | TUBB4B | tubulin, beta 4B class IVb | 10383 | ENSG00000188229 |
| 7 | P09525 | NA | ANXA4 | annexin A4 | 307 | ENSG00000196975 |
| 8 | P61225 | NA | RAP2B | RAP2B, member of RAS oncogene family | 5912 | ENSG00000181467 |
| 9 | Q04828 | NA | AKR1C1 | aldo-keto reductase family 1, member C1 (dihydrodiol dehydrogenase 1; 20-alpha (3-alpha)-hydroxysteroid dehydrogenase) | 1645 | ENSG00000187134 |
| 10 | P60842 | NA | EIF4A1 | eukaryotic translation initiation factor 4A1 | 1973 | ENSG00000161960 |
| 11 | P49588 | NA | AARS | alanyl-tRNA synthetase | 16 | ENSG00000090861 |
| 12 | P68104 | NA | EEF1A1 | eukaryotic translation elongation factor 1 alpha 1 | 1915 | ENSG00000156508 |
| 13 | P26639 | NA | TARS | threonyl-tRNA synthetase | 6897 | ENSG00000113407 |
| 14 | P04083 | NA | ANXA1 | annexin A1 | 301 | ENSG00000135046 |
| 15 | P17655 | NA | CAPN2 | calpain 2, (m/II) large subunit | 824 | ENSG00000162909 |
| 16 | P22102 | NA | GART | phosphoribosylglycinamide formyltransferase, phosphoribosylglycinamide synthetase, phosphoribosylaminoimidazole synthetase | 2618 | ENSG00000159131 |
| 17 | P16615 | NA | ATP2A2 | ATPase, Ca++ transporting, cardiac muscle, slow twitch 2 | 488 | ENSG00000174437 |
| 18 | P41091 | NA | EIF2S3 | eukaryotic translation initiation factor 2, subunit 3 gamma, 52kDa | 1968 | ENSG00000130741 |
| 19 | P02768 | NA | ALB | albumin | 213 | ENSG00000163631 |
| 20 | Q92974 | NA | ARHGEF2 | Rho/Rac guanine nucleotide exchange factor (GEF) 2 | 9181 | ENSG00000116584 |
| 21 | P17174 | NA | GOT1 | glutamic-oxaloacetic transaminase 1, soluble (aspartate aminotransferase 1) | 2805 | ENSG00000120053 |
| 22 | O00194 | NA | RAB27B | RAB27B, member RAS oncogene family | 5874 | ENSG00000041353 |
| 23 | P11387 | NA | TOP1 | topoisomerase (DNA) I | 7150 | ENSG00000198900 |

  
  

| **Database:molecular function      &nbspName:hydrolase activity      &nbspID:GO:0016787** | | | | | | |
| --- | --- | --- | --- | --- | --- | --- |
| C=2343; O=13; E=5.79; R=2.25; rawP=0.0029; adjP=0.0071 | | | | | | |
| Index | UserID | Value | Gene Symbol | Gene Name | EntrezGene | Ensembl |
| 1 | P17655 | NA | CAPN2 | calpain 2, (m/II) large subunit | 824 | ENSG00000162909 |
| 2 | Q9NR30 | NA | DDX21 | DEAD (Asp-Glu-Ala-Asp) box helicase 21 | 9188 | ENSG00000165732 |
| 3 | P16615 | NA | ATP2A2 | ATPase, Ca++ transporting, cardiac muscle, slow twitch 2 | 488 | ENSG00000174437 |
| 4 | P41091 | NA | EIF2S3 | eukaryotic translation initiation factor 2, subunit 3 gamma, 52kDa | 1968 | ENSG00000130741 |
| 5 | P68371 | NA | TUBB4B | tubulin, beta 4B class IVb | 10383 | ENSG00000188229 |
| 6 | P61225 | NA | RAP2B | RAP2B, member of RAS oncogene family | 5912 | ENSG00000181467 |
| 7 | P60842 | NA | EIF4A1 | eukaryotic translation initiation factor 4A1 | 1973 | ENSG00000161960 |
| 8 | Q6PIU2 | NA | NCEH1 | neutral cholesterol ester hydrolase 1 | 57552 | ENSG00000144959 |
| 9 | O00194 | NA | RAB27B | RAB27B, member RAS oncogene family | 5874 | ENSG00000041353 |
| 10 | P49588 | NA | AARS | alanyl-tRNA synthetase | 16 | ENSG00000090861 |
| 11 | P68104 | NA | EEF1A1 | eukaryotic translation elongation factor 1 alpha 1 | 1915 | ENSG00000156508 |
| 12 | P02786 | NA | TFRC | transferrin receptor (p90, CD71) | 7037 | ENSG00000072274 |
| 13 | P11387 | NA | TOP1 | topoisomerase (DNA) I | 7150 | ENSG00000198900 |

  
  

| **Database:molecular function      &nbspName:cytoskeletal protein binding      &nbspID:GO:0008092** | | | | | | |
| --- | --- | --- | --- | --- | --- | --- |
| C=634; O=6; E=1.57; R=3.83; rawP=0.0043; adjP=0.0102 | | | | | | |
| Index | UserID | Value | Gene Symbol | Gene Name | EntrezGene | Ensembl |
| 1 | Q92974 | NA | ARHGEF2 | Rho/Rac guanine nucleotide exchange factor (GEF) 2 | 9181 | ENSG00000116584 |
| 2 | P17655 | NA | CAPN2 | calpain 2, (m/II) large subunit | 824 | ENSG00000162909 |
| 3 | P18206 | NA | VCL | vinculin | 7414 | ENSG00000035403 |
| 4 | Q86UP2 | NA | KTN1 | kinectin 1 (kinesin receptor) | 3895 | ENSG00000126777 |
| 5 | O75369 | NA | FLNB | filamin B, beta | 2317 | ENSG00000136068 |
| 6 | P05556 | NA | ITGB1 | integrin, beta 1 (fibronectin receptor, beta polypeptide, antigen CD29 includes MDF2, MSK12) | 3688 | ENSG00000150093 |

  
  

| **Database:molecular function      &nbspName:ATPase activity, coupled      &nbspID:GO:0042623** | | | | | | |
| --- | --- | --- | --- | --- | --- | --- |
| C=274; O=4; E=0.68; R=5.91; rawP=0.0045; adjP=0.0103 | | | | | | |
| Index | UserID | Value | Gene Symbol | Gene Name | EntrezGene | Ensembl |
| 1 | P60842 | NA | EIF4A1 | eukaryotic translation initiation factor 4A1 | 1973 | ENSG00000161960 |
| 2 | Q9NR30 | NA | DDX21 | DEAD (Asp-Glu-Ala-Asp) box helicase 21 | 9188 | ENSG00000165732 |
| 3 | P16615 | NA | ATP2A2 | ATPase, Ca++ transporting, cardiac muscle, slow twitch 2 | 488 | ENSG00000174437 |
| 4 | P11387 | NA | TOP1 | topoisomerase (DNA) I | 7150 | ENSG00000198900 |

  
  

| **Database:molecular function      &nbspName:heterocyclic compound binding      &nbspID:GO:1901363** | | | | | | |
| --- | --- | --- | --- | --- | --- | --- |
| C=5290; O=21; E=13.07; R=1.61; rawP=0.0062; adjP=0.0136 | | | | | | |
| Index | UserID | Value | Gene Symbol | Gene Name | EntrezGene | Ensembl |
| 1 | Q16881 | NA | TXNRD1 | thioredoxin reductase 1 | 7296 | ENSG00000198431 |
| 2 | Q9NR30 | NA | DDX21 | DEAD (Asp-Glu-Ala-Asp) box helicase 21 | 9188 | ENSG00000165732 |
| 3 | P00491 | NA | PNP | purine nucleoside phosphorylase | 4860 | ENSG00000198805 |
| 4 | P68371 | NA | TUBB4B | tubulin, beta 4B class IVb | 10383 | ENSG00000188229 |
| 5 | P61225 | NA | RAP2B | RAP2B, member of RAS oncogene family | 5912 | ENSG00000181467 |
| 6 | P60842 | NA | EIF4A1 | eukaryotic translation initiation factor 4A1 | 1973 | ENSG00000161960 |
| 7 | P49588 | NA | AARS | alanyl-tRNA synthetase | 16 | ENSG00000090861 |
| 8 | P68104 | NA | EEF1A1 | eukaryotic translation elongation factor 1 alpha 1 | 1915 | ENSG00000156508 |
| 9 | P26639 | NA | TARS | threonyl-tRNA synthetase | 6897 | ENSG00000113407 |
| 10 | P04083 | NA | ANXA1 | annexin A1 | 301 | ENSG00000135046 |
| 11 | O00425 | NA | IGF2BP3 | insulin-like growth factor 2 mRNA binding protein 3 | 10643 | ENSG00000136231 |
| 12 | P24534 | NA | EEF1B2 | eukaryotic translation elongation factor 1 beta 2 | 1933 | ENSG00000114942 |
| 13 | P56537 | NA | EIF6 | eukaryotic translation initiation factor 6 | 3692 | ENSG00000242372 |
| 14 | P22102 | NA | GART | phosphoribosylglycinamide formyltransferase, phosphoribosylglycinamide synthetase, phosphoribosylaminoimidazole synthetase | 2618 | ENSG00000159131 |
| 15 | P16615 | NA | ATP2A2 | ATPase, Ca++ transporting, cardiac muscle, slow twitch 2 | 488 | ENSG00000174437 |
| 16 | P41091 | NA | EIF2S3 | eukaryotic translation initiation factor 2, subunit 3 gamma, 52kDa | 1968 | ENSG00000130741 |
| 17 | P02768 | NA | ALB | albumin | 213 | ENSG00000163631 |
| 18 | P17174 | NA | GOT1 | glutamic-oxaloacetic transaminase 1, soluble (aspartate aminotransferase 1) | 2805 | ENSG00000120053 |
| 19 | O00194 | NA | RAB27B | RAB27B, member RAS oncogene family | 5874 | ENSG00000041353 |
| 20 | P29692 | NA | EEF1D | eukaryotic translation elongation factor 1 delta (guanine nucleotide exchange protein) | 1936 | ENSG00000104529 |
| 21 | P11387 | NA | TOP1 | topoisomerase (DNA) I | 7150 | ENSG00000198900 |

  
  

| **Database:molecular function      &nbspName:catalytic activity      &nbspID:GO:0003824** | | | | | | |
| --- | --- | --- | --- | --- | --- | --- |
| C=5299; O=21; E=13.09; R=1.60; rawP=0.0063; adjP=0.0136 | | | | | | |
| Index | UserID | Value | Gene Symbol | Gene Name | EntrezGene | Ensembl |
| 1 | Q16881 | NA | TXNRD1 | thioredoxin reductase 1 | 7296 | ENSG00000198431 |
| 2 | Q9NR30 | NA | DDX21 | DEAD (Asp-Glu-Ala-Asp) box helicase 21 | 9188 | ENSG00000165732 |
| 3 | P08195 | NA | SLC3A2 | solute carrier family 3 (activators of dibasic and neutral amino acid transport), member 2 | 6520 | ENSG00000168003 |
| 4 | P00491 | NA | PNP | purine nucleoside phosphorylase | 4860 | ENSG00000198805 |
| 5 | P68371 | NA | TUBB4B | tubulin, beta 4B class IVb | 10383 | ENSG00000188229 |
| 6 | P61225 | NA | RAP2B | RAP2B, member of RAS oncogene family | 5912 | ENSG00000181467 |
| 7 | Q04828 | NA | AKR1C1 | aldo-keto reductase family 1, member C1 (dihydrodiol dehydrogenase 1; 20-alpha (3-alpha)-hydroxysteroid dehydrogenase) | 1645 | ENSG00000187134 |
| 8 | P60842 | NA | EIF4A1 | eukaryotic translation initiation factor 4A1 | 1973 | ENSG00000161960 |
| 9 | P49588 | NA | AARS | alanyl-tRNA synthetase | 16 | ENSG00000090861 |
| 10 | P68104 | NA | EEF1A1 | eukaryotic translation elongation factor 1 alpha 1 | 1915 | ENSG00000156508 |
| 11 | P26639 | NA | TARS | threonyl-tRNA synthetase | 6897 | ENSG00000113407 |
| 12 | P02786 | NA | TFRC | transferrin receptor (p90, CD71) | 7037 | ENSG00000072274 |
| 13 | P17655 | NA | CAPN2 | calpain 2, (m/II) large subunit | 824 | ENSG00000162909 |
| 14 | P22102 | NA | GART | phosphoribosylglycinamide formyltransferase, phosphoribosylglycinamide synthetase, phosphoribosylaminoimidazole synthetase | 2618 | ENSG00000159131 |
| 15 | P16615 | NA | ATP2A2 | ATPase, Ca++ transporting, cardiac muscle, slow twitch 2 | 488 | ENSG00000174437 |
| 16 | P41091 | NA | EIF2S3 | eukaryotic translation initiation factor 2, subunit 3 gamma, 52kDa | 1968 | ENSG00000130741 |
| 17 | P17174 | NA | GOT1 | glutamic-oxaloacetic transaminase 1, soluble (aspartate aminotransferase 1) | 2805 | ENSG00000120053 |
| 18 | O00194 | NA | RAB27B | RAB27B, member RAS oncogene family | 5874 | ENSG00000041353 |
| 19 | Q6PIU2 | NA | NCEH1 | neutral cholesterol ester hydrolase 1 | 57552 | ENSG00000144959 |
| 20 | P51648 | NA | ALDH3A2 | aldehyde dehydrogenase 3 family, member A2 | 224 | ENSG00000072210 |
| 21 | P11387 | NA | TOP1 | topoisomerase (DNA) I | 7150 | ENSG00000198900 |

  
  

| **Database:molecular function      &nbspName:protein binding      &nbspID:GO:0005515** | | | | | | |
| --- | --- | --- | --- | --- | --- | --- |
| C=7301; O=26; E=18.04; R=1.44; rawP=0.0066; adjP=0.0138 | | | | | | |
| Index | UserID | Value | Gene Symbol | Gene Name | EntrezGene | Ensembl |
| 1 | P02795 | NA | MT2A | metallothionein 2A | 4502 | ENSG00000125148 |
| 2 | P08195 | NA | SLC3A2 | solute carrier family 3 (activators of dibasic and neutral amino acid transport), member 2 | 6520 | ENSG00000168003 |
| 3 | P68371 | NA | TUBB4B | tubulin, beta 4B class IVb | 10383 | ENSG00000188229 |
| 4 | P61225 | NA | RAP2B | RAP2B, member of RAS oncogene family | 5912 | ENSG00000181467 |
| 5 | P60842 | NA | EIF4A1 | eukaryotic translation initiation factor 4A1 | 1973 | ENSG00000161960 |
| 6 | P04080 | NA | CSTB | cystatin B (stefin B) | 1476 | ENSG00000160213 |
| 7 | Q86UP2 | NA | KTN1 | kinectin 1 (kinesin receptor) | 3895 | ENSG00000126777 |
| 8 | P68104 | NA | EEF1A1 | eukaryotic translation elongation factor 1 alpha 1 | 1915 | ENSG00000156508 |
| 9 | P26639 | NA | TARS | threonyl-tRNA synthetase | 6897 | ENSG00000113407 |
| 10 | O75369 | NA | FLNB | filamin B, beta | 2317 | ENSG00000136068 |
| 11 | P02786 | NA | TFRC | transferrin receptor (p90, CD71) | 7037 | ENSG00000072274 |
| 12 | P04083 | NA | ANXA1 | annexin A1 | 301 | ENSG00000135046 |
| 13 | O00425 | NA | IGF2BP3 | insulin-like growth factor 2 mRNA binding protein 3 | 10643 | ENSG00000136231 |
| 14 | P24534 | NA | EEF1B2 | eukaryotic translation elongation factor 1 beta 2 | 1933 | ENSG00000114942 |
| 15 | P17655 | NA | CAPN2 | calpain 2, (m/II) large subunit | 824 | ENSG00000162909 |
| 16 | P56537 | NA | EIF6 | eukaryotic translation initiation factor 6 | 3692 | ENSG00000242372 |
| 17 | Q09666 | NA | AHNAK | AHNAK nucleoprotein | 79026 | ENSG00000124942 |
| 18 | P16615 | NA | ATP2A2 | ATPase, Ca++ transporting, cardiac muscle, slow twitch 2 | 488 | ENSG00000174437 |
| 19 | P41091 | NA | EIF2S3 | eukaryotic translation initiation factor 2, subunit 3 gamma, 52kDa | 1968 | ENSG00000130741 |
| 20 | P02768 | NA | ALB | albumin | 213 | ENSG00000163631 |
| 21 | P05556 | NA | ITGB1 | integrin, beta 1 (fibronectin receptor, beta polypeptide, antigen CD29 includes MDF2, MSK12) | 3688 | ENSG00000150093 |
| 22 | Q92974 | NA | ARHGEF2 | Rho/Rac guanine nucleotide exchange factor (GEF) 2 | 9181 | ENSG00000116584 |
| 23 | O00194 | NA | RAB27B | RAB27B, member RAS oncogene family | 5874 | ENSG00000041353 |
| 24 | P18206 | NA | VCL | vinculin | 7414 | ENSG00000035403 |
| 25 | P29692 | NA | EEF1D | eukaryotic translation elongation factor 1 delta (guanine nucleotide exchange protein) | 1936 | ENSG00000104529 |
| 26 | P11387 | NA | TOP1 | topoisomerase (DNA) I | 7150 | ENSG00000198900 |

  
  

| **Database:molecular function      &nbspName:ATPase activity      &nbspID:GO:0016887** | | | | | | |
| --- | --- | --- | --- | --- | --- | --- |
| C=352; O=4; E=0.87; R=4.60; rawP=0.0108; adjP=0.0219 | | | | | | |
| Index | UserID | Value | Gene Symbol | Gene Name | EntrezGene | Ensembl |
| 1 | P60842 | NA | EIF4A1 | eukaryotic translation initiation factor 4A1 | 1973 | ENSG00000161960 |
| 2 | Q9NR30 | NA | DDX21 | DEAD (Asp-Glu-Ala-Asp) box helicase 21 | 9188 | ENSG00000165732 |
| 3 | P16615 | NA | ATP2A2 | ATPase, Ca++ transporting, cardiac muscle, slow twitch 2 | 488 | ENSG00000174437 |
| 4 | P11387 | NA | TOP1 | topoisomerase (DNA) I | 7150 | ENSG00000198900 |

  
  

| **Database:molecular function      &nbspName:cofactor binding      &nbspID:GO:0048037** | | | | | | |
| --- | --- | --- | --- | --- | --- | --- |
| C=259; O=3; E=0.64; R=4.69; rawP=0.0258; adjP=0.0509 | | | | | | |
| Index | UserID | Value | Gene Symbol | Gene Name | EntrezGene | Ensembl |
| 1 | Q16881 | NA | TXNRD1 | thioredoxin reductase 1 | 7296 | ENSG00000198431 |
| 2 | P17174 | NA | GOT1 | glutamic-oxaloacetic transaminase 1, soluble (aspartate aminotransferase 1) | 2805 | ENSG00000120053 |
| 3 | P02768 | NA | ALB | albumin | 213 | ENSG00000163631 |

  
  

| **Database:molecular function      &nbspName:binding      &nbspID:GO:0005488** | | | | | | |
| --- | --- | --- | --- | --- | --- | --- |
| C=11779; O=34; E=29.10; R=1.17; rawP=0.0296; adjP=0.0568 | | | | | | |
| Index | UserID | Value | Gene Symbol | Gene Name | EntrezGene | Ensembl |
| 1 | P68371 | NA | TUBB4B | tubulin, beta 4B class IVb | 10383 | ENSG00000188229 |
| 2 | P61225 | NA | RAP2B | RAP2B, member of RAS oncogene family | 5912 | ENSG00000181467 |
| 3 | Q04828 | NA | AKR1C1 | aldo-keto reductase family 1, member C1 (dihydrodiol dehydrogenase 1; 20-alpha (3-alpha)-hydroxysteroid dehydrogenase) | 1645 | ENSG00000187134 |
| 4 | P49588 | NA | AARS | alanyl-tRNA synthetase | 16 | ENSG00000090861 |
| 5 | Q86UP2 | NA | KTN1 | kinectin 1 (kinesin receptor) | 3895 | ENSG00000126777 |
| 6 | P68104 | NA | EEF1A1 | eukaryotic translation elongation factor 1 alpha 1 | 1915 | ENSG00000156508 |
| 7 | O75369 | NA | FLNB | filamin B, beta | 2317 | ENSG00000136068 |
| 8 | P02786 | NA | TFRC | transferrin receptor (p90, CD71) | 7037 | ENSG00000072274 |
| 9 | P24534 | NA | EEF1B2 | eukaryotic translation elongation factor 1 beta 2 | 1933 | ENSG00000114942 |
| 10 | O00425 | NA | IGF2BP3 | insulin-like growth factor 2 mRNA binding protein 3 | 10643 | ENSG00000136231 |
| 11 | P17655 | NA | CAPN2 | calpain 2, (m/II) large subunit | 824 | ENSG00000162909 |
| 12 | P56537 | NA | EIF6 | eukaryotic translation initiation factor 6 | 3692 | ENSG00000242372 |
| 13 | P22102 | NA | GART | phosphoribosylglycinamide formyltransferase, phosphoribosylglycinamide synthetase, phosphoribosylaminoimidazole synthetase | 2618 | ENSG00000159131 |
| 14 | P16615 | NA | ATP2A2 | ATPase, Ca++ transporting, cardiac muscle, slow twitch 2 | 488 | ENSG00000174437 |
| 15 | Q92974 | NA | ARHGEF2 | Rho/Rac guanine nucleotide exchange factor (GEF) 2 | 9181 | ENSG00000116584 |
| 16 | P17174 | NA | GOT1 | glutamic-oxaloacetic transaminase 1, soluble (aspartate aminotransferase 1) | 2805 | ENSG00000120053 |
| 17 | O00194 | NA | RAB27B | RAB27B, member RAS oncogene family | 5874 | ENSG00000041353 |
| 18 | P18206 | NA | VCL | vinculin | 7414 | ENSG00000035403 |
| 19 | P29692 | NA | EEF1D | eukaryotic translation elongation factor 1 delta (guanine nucleotide exchange protein) | 1936 | ENSG00000104529 |
| 20 | P11387 | NA | TOP1 | topoisomerase (DNA) I | 7150 | ENSG00000198900 |
| 21 | Q16881 | NA | TXNRD1 | thioredoxin reductase 1 | 7296 | ENSG00000198431 |
| 22 | Q9NR30 | NA | DDX21 | DEAD (Asp-Glu-Ala-Asp) box helicase 21 | 9188 | ENSG00000165732 |
| 23 | P08195 | NA | SLC3A2 | solute carrier family 3 (activators of dibasic and neutral amino acid transport), member 2 | 6520 | ENSG00000168003 |
| 24 | P02795 | NA | MT2A | metallothionein 2A | 4502 | ENSG00000125148 |
| 25 | P00491 | NA | PNP | purine nucleoside phosphorylase | 4860 | ENSG00000198805 |
| 26 | P09525 | NA | ANXA4 | annexin A4 | 307 | ENSG00000196975 |
| 27 | P60842 | NA | EIF4A1 | eukaryotic translation initiation factor 4A1 | 1973 | ENSG00000161960 |
| 28 | P04080 | NA | CSTB | cystatin B (stefin B) | 1476 | ENSG00000160213 |
| 29 | P26639 | NA | TARS | threonyl-tRNA synthetase | 6897 | ENSG00000113407 |
| 30 | P04083 | NA | ANXA1 | annexin A1 | 301 | ENSG00000135046 |
| 31 | Q09666 | NA | AHNAK | AHNAK nucleoprotein | 79026 | ENSG00000124942 |
| 32 | P02768 | NA | ALB | albumin | 213 | ENSG00000163631 |
| 33 | P41091 | NA | EIF2S3 | eukaryotic translation initiation factor 2, subunit 3 gamma, 52kDa | 1968 | ENSG00000130741 |
| 34 | P05556 | NA | ITGB1 | integrin, beta 1 (fibronectin receptor, beta polypeptide, antigen CD29 includes MDF2, MSK12) | 3688 | ENSG00000150093 |

  
  

| **Database:molecular function      &nbspName:lipid binding      &nbspID:GO:0008289** | | | | | | |
| --- | --- | --- | --- | --- | --- | --- |
| C=735; O=5; E=1.82; R=2.75; rawP=0.0334; adjP=0.0624 | | | | | | |
| Index | UserID | Value | Gene Symbol | Gene Name | EntrezGene | Ensembl |
| 1 | Q92974 | NA | ARHGEF2 | Rho/Rac guanine nucleotide exchange factor (GEF) 2 | 9181 | ENSG00000116584 |
| 2 | Q04828 | NA | AKR1C1 | aldo-keto reductase family 1, member C1 (dihydrodiol dehydrogenase 1; 20-alpha (3-alpha)-hydroxysteroid dehydrogenase) | 1645 | ENSG00000187134 |
| 3 | P02768 | NA | ALB | albumin | 213 | ENSG00000163631 |
| 4 | P09525 | NA | ANXA4 | annexin A4 | 307 | ENSG00000196975 |
| 5 | P04083 | NA | ANXA1 | annexin A1 | 301 | ENSG00000135046 |

  
  

| **Database:molecular function      &nbspName:active transmembrane transporter activity      &nbspID:GO:0022804** | | | | | | |
| --- | --- | --- | --- | --- | --- | --- |
| C=306; O=3; E=0.76; R=3.97; rawP=0.0394; adjP=0.0717 | | | | | | |
| Index | UserID | Value | Gene Symbol | Gene Name | EntrezGene | Ensembl |
| 1 | P08195 | NA | SLC3A2 | solute carrier family 3 (activators of dibasic and neutral amino acid transport), member 2 | 6520 | ENSG00000168003 |
| 2 | Q15758 | NA | SLC1A5 | solute carrier family 1 (neutral amino acid transporter), member 5 | 6510 | ENSG00000105281 |
| 3 | P16615 | NA | ATP2A2 | ATPase, Ca++ transporting, cardiac muscle, slow twitch 2 | 488 | ENSG00000174437 |

  
  

| **Database:molecular function      &nbspName:enzyme inhibitor activity      &nbspID:GO:0004857** | | | | | | |
| --- | --- | --- | --- | --- | --- | --- |
| C=310; O=3; E=0.77; R=3.92; rawP=0.0407; adjP=0.0722 | | | | | | |
| Index | UserID | Value | Gene Symbol | Gene Name | EntrezGene | Ensembl |
| 1 | P04080 | NA | CSTB | cystatin B (stefin B) | 1476 | ENSG00000160213 |
| 2 | P09525 | NA | ANXA4 | annexin A4 | 307 | ENSG00000196975 |
| 3 | P04083 | NA | ANXA1 | annexin A1 | 301 | ENSG00000135046 |

  
  

| **Database:cellular component      &nbspName:eukaryotic translation elongation factor 1 complex      &nbspID:GO:0005853** | | | | | | |
| --- | --- | --- | --- | --- | --- | --- |
| C=5; O=3; E=0.01; R=263.40; rawP=1.09e-07; adjP=5.19e-06 | | | | | | |
| Index | UserID | Value | Gene Symbol | Gene Name | EntrezGene | Ensembl |
| 1 | P68104 | NA | EEF1A1 | eukaryotic translation elongation factor 1 alpha 1 | 1915 | ENSG00000156508 |
| 2 | P29692 | NA | EEF1D | eukaryotic translation elongation factor 1 delta (guanine nucleotide exchange protein) | 1936 | ENSG00000104529 |
| 3 | P24534 | NA | EEF1B2 | eukaryotic translation elongation factor 1 beta 2 | 1933 | ENSG00000114942 |

  
  

| **Database:cellular component      &nbspName:cytoplasmic part      &nbspID:GO:0044444** | | | | | | |
| --- | --- | --- | --- | --- | --- | --- |
| C=6728; O=31; E=15.33; R=2.02; rawP=2.29e-07; adjP=5.19e-06 | | | | | | |
| Index | UserID | Value | Gene Symbol | Gene Name | EntrezGene | Ensembl |
| 1 | Q16881 | NA | TXNRD1 | thioredoxin reductase 1 | 7296 | ENSG00000198431 |
| 2 | P02795 | NA | MT2A | metallothionein 2A | 4502 | ENSG00000125148 |
| 3 | P08195 | NA | SLC3A2 | solute carrier family 3 (activators of dibasic and neutral amino acid transport), member 2 | 6520 | ENSG00000168003 |
| 4 | P00491 | NA | PNP | purine nucleoside phosphorylase | 4860 | ENSG00000198805 |
| 5 | P68371 | NA | TUBB4B | tubulin, beta 4B class IVb | 10383 | ENSG00000188229 |
| 6 | P61225 | NA | RAP2B | RAP2B, member of RAS oncogene family | 5912 | ENSG00000181467 |
| 7 | Q04828 | NA | AKR1C1 | aldo-keto reductase family 1, member C1 (dihydrodiol dehydrogenase 1; 20-alpha (3-alpha)-hydroxysteroid dehydrogenase) | 1645 | ENSG00000187134 |
| 8 | P60842 | NA | EIF4A1 | eukaryotic translation initiation factor 4A1 | 1973 | ENSG00000161960 |
| 9 | P49588 | NA | AARS | alanyl-tRNA synthetase | 16 | ENSG00000090861 |
| 10 | Q86UP2 | NA | KTN1 | kinectin 1 (kinesin receptor) | 3895 | ENSG00000126777 |
| 11 | P68104 | NA | EEF1A1 | eukaryotic translation elongation factor 1 alpha 1 | 1915 | ENSG00000156508 |
| 12 | P26639 | NA | TARS | threonyl-tRNA synthetase | 6897 | ENSG00000113407 |
| 13 | O75369 | NA | FLNB | filamin B, beta | 2317 | ENSG00000136068 |
| 14 | P02786 | NA | TFRC | transferrin receptor (p90, CD71) | 7037 | ENSG00000072274 |
| 15 | P04083 | NA | ANXA1 | annexin A1 | 301 | ENSG00000135046 |
| 16 | P24534 | NA | EEF1B2 | eukaryotic translation elongation factor 1 beta 2 | 1933 | ENSG00000114942 |
| 17 | O00425 | NA | IGF2BP3 | insulin-like growth factor 2 mRNA binding protein 3 | 10643 | ENSG00000136231 |
| 18 | P22102 | NA | GART | phosphoribosylglycinamide formyltransferase, phosphoribosylglycinamide synthetase, phosphoribosylaminoimidazole synthetase | 2618 | ENSG00000159131 |
| 19 | P16615 | NA | ATP2A2 | ATPase, Ca++ transporting, cardiac muscle, slow twitch 2 | 488 | ENSG00000174437 |
| 20 | P41091 | NA | EIF2S3 | eukaryotic translation initiation factor 2, subunit 3 gamma, 52kDa | 1968 | ENSG00000130741 |
| 21 | P02768 | NA | ALB | albumin | 213 | ENSG00000163631 |
| 22 | P05556 | NA | ITGB1 | integrin, beta 1 (fibronectin receptor, beta polypeptide, antigen CD29 includes MDF2, MSK12) | 3688 | ENSG00000150093 |
| 23 | Q92974 | NA | ARHGEF2 | Rho/Rac guanine nucleotide exchange factor (GEF) 2 | 9181 | ENSG00000116584 |
| 24 | P17174 | NA | GOT1 | glutamic-oxaloacetic transaminase 1, soluble (aspartate aminotransferase 1) | 2805 | ENSG00000120053 |
| 25 | Q15758 | NA | SLC1A5 | solute carrier family 1 (neutral amino acid transporter), member 5 | 6510 | ENSG00000105281 |
| 26 | O00194 | NA | RAB27B | RAB27B, member RAS oncogene family | 5874 | ENSG00000041353 |
| 27 | Q6PIU2 | NA | NCEH1 | neutral cholesterol ester hydrolase 1 | 57552 | ENSG00000144959 |
| 28 | P18206 | NA | VCL | vinculin | 7414 | ENSG00000035403 |
| 29 | P51648 | NA | ALDH3A2 | aldehyde dehydrogenase 3 family, member A2 | 224 | ENSG00000072210 |
| 30 | P29692 | NA | EEF1D | eukaryotic translation elongation factor 1 delta (guanine nucleotide exchange protein) | 1936 | ENSG00000104529 |
| 31 | P11387 | NA | TOP1 | topoisomerase (DNA) I | 7150 | ENSG00000198900 |

  
  

| **Database:cellular component      &nbspName:cytosol      &nbspID:GO:0005829** | | | | | | |
| --- | --- | --- | --- | --- | --- | --- |
| C=2367; O=19; E=5.39; R=3.52; rawP=1.69e-07; adjP=5.19e-06 | | | | | | |
| Index | UserID | Value | Gene Symbol | Gene Name | EntrezGene | Ensembl |
| 1 | Q16881 | NA | TXNRD1 | thioredoxin reductase 1 | 7296 | ENSG00000198431 |
| 2 | P02795 | NA | MT2A | metallothionein 2A | 4502 | ENSG00000125148 |
| 3 | P00491 | NA | PNP | purine nucleoside phosphorylase | 4860 | ENSG00000198805 |
| 4 | P68371 | NA | TUBB4B | tubulin, beta 4B class IVb | 10383 | ENSG00000188229 |
| 5 | P61225 | NA | RAP2B | RAP2B, member of RAS oncogene family | 5912 | ENSG00000181467 |
| 6 | Q04828 | NA | AKR1C1 | aldo-keto reductase family 1, member C1 (dihydrodiol dehydrogenase 1; 20-alpha (3-alpha)-hydroxysteroid dehydrogenase) | 1645 | ENSG00000187134 |
| 7 | P60842 | NA | EIF4A1 | eukaryotic translation initiation factor 4A1 | 1973 | ENSG00000161960 |
| 8 | P49588 | NA | AARS | alanyl-tRNA synthetase | 16 | ENSG00000090861 |
| 9 | P68104 | NA | EEF1A1 | eukaryotic translation elongation factor 1 alpha 1 | 1915 | ENSG00000156508 |
| 10 | P26639 | NA | TARS | threonyl-tRNA synthetase | 6897 | ENSG00000113407 |
| 11 | O75369 | NA | FLNB | filamin B, beta | 2317 | ENSG00000136068 |
| 12 | O00425 | NA | IGF2BP3 | insulin-like growth factor 2 mRNA binding protein 3 | 10643 | ENSG00000136231 |
| 13 | P24534 | NA | EEF1B2 | eukaryotic translation elongation factor 1 beta 2 | 1933 | ENSG00000114942 |
| 14 | P22102 | NA | GART | phosphoribosylglycinamide formyltransferase, phosphoribosylglycinamide synthetase, phosphoribosylaminoimidazole synthetase | 2618 | ENSG00000159131 |
| 15 | P41091 | NA | EIF2S3 | eukaryotic translation initiation factor 2, subunit 3 gamma, 52kDa | 1968 | ENSG00000130741 |
| 16 | Q92974 | NA | ARHGEF2 | Rho/Rac guanine nucleotide exchange factor (GEF) 2 | 9181 | ENSG00000116584 |
| 17 | P17174 | NA | GOT1 | glutamic-oxaloacetic transaminase 1, soluble (aspartate aminotransferase 1) | 2805 | ENSG00000120053 |
| 18 | P18206 | NA | VCL | vinculin | 7414 | ENSG00000035403 |
| 19 | P29692 | NA | EEF1D | eukaryotic translation elongation factor 1 delta (guanine nucleotide exchange protein) | 1936 | ENSG00000104529 |

  
  

| **Database:cellular component      &nbspName:cytoplasm      &nbspID:GO:0005737** | | | | | | |
| --- | --- | --- | --- | --- | --- | --- |
| C=9051; O=35; E=20.62; R=1.70; rawP=4.42e-07; adjP=7.51e-06 | | | | | | |
| Index | UserID | Value | Gene Symbol | Gene Name | EntrezGene | Ensembl |
| 1 | P68371 | NA | TUBB4B | tubulin, beta 4B class IVb | 10383 | ENSG00000188229 |
| 2 | P61225 | NA | RAP2B | RAP2B, member of RAS oncogene family | 5912 | ENSG00000181467 |
| 3 | Q04828 | NA | AKR1C1 | aldo-keto reductase family 1, member C1 (dihydrodiol dehydrogenase 1; 20-alpha (3-alpha)-hydroxysteroid dehydrogenase) | 1645 | ENSG00000187134 |
| 4 | P49588 | NA | AARS | alanyl-tRNA synthetase | 16 | ENSG00000090861 |
| 5 | Q86UP2 | NA | KTN1 | kinectin 1 (kinesin receptor) | 3895 | ENSG00000126777 |
| 6 | P68104 | NA | EEF1A1 | eukaryotic translation elongation factor 1 alpha 1 | 1915 | ENSG00000156508 |
| 7 | O75369 | NA | FLNB | filamin B, beta | 2317 | ENSG00000136068 |
| 8 | P02786 | NA | TFRC | transferrin receptor (p90, CD71) | 7037 | ENSG00000072274 |
| 9 | P24534 | NA | EEF1B2 | eukaryotic translation elongation factor 1 beta 2 | 1933 | ENSG00000114942 |
| 10 | O00425 | NA | IGF2BP3 | insulin-like growth factor 2 mRNA binding protein 3 | 10643 | ENSG00000136231 |
| 11 | P17655 | NA | CAPN2 | calpain 2, (m/II) large subunit | 824 | ENSG00000162909 |
| 12 | P56537 | NA | EIF6 | eukaryotic translation initiation factor 6 | 3692 | ENSG00000242372 |
| 13 | P22102 | NA | GART | phosphoribosylglycinamide formyltransferase, phosphoribosylglycinamide synthetase, phosphoribosylaminoimidazole synthetase | 2618 | ENSG00000159131 |
| 14 | P16615 | NA | ATP2A2 | ATPase, Ca++ transporting, cardiac muscle, slow twitch 2 | 488 | ENSG00000174437 |
| 15 | Q92974 | NA | ARHGEF2 | Rho/Rac guanine nucleotide exchange factor (GEF) 2 | 9181 | ENSG00000116584 |
| 16 | P17174 | NA | GOT1 | glutamic-oxaloacetic transaminase 1, soluble (aspartate aminotransferase 1) | 2805 | ENSG00000120053 |
| 17 | O00194 | NA | RAB27B | RAB27B, member RAS oncogene family | 5874 | ENSG00000041353 |
| 18 | Q6PIU2 | NA | NCEH1 | neutral cholesterol ester hydrolase 1 | 57552 | ENSG00000144959 |
| 19 | P18206 | NA | VCL | vinculin | 7414 | ENSG00000035403 |
| 20 | P51648 | NA | ALDH3A2 | aldehyde dehydrogenase 3 family, member A2 | 224 | ENSG00000072210 |
| 21 | P29692 | NA | EEF1D | eukaryotic translation elongation factor 1 delta (guanine nucleotide exchange protein) | 1936 | ENSG00000104529 |
| 22 | P11387 | NA | TOP1 | topoisomerase (DNA) I | 7150 | ENSG00000198900 |
| 23 | Q16881 | NA | TXNRD1 | thioredoxin reductase 1 | 7296 | ENSG00000198431 |
| 24 | P08195 | NA | SLC3A2 | solute carrier family 3 (activators of dibasic and neutral amino acid transport), member 2 | 6520 | ENSG00000168003 |
| 25 | P02795 | NA | MT2A | metallothionein 2A | 4502 | ENSG00000125148 |
| 26 | P00491 | NA | PNP | purine nucleoside phosphorylase | 4860 | ENSG00000198805 |
| 27 | P09525 | NA | ANXA4 | annexin A4 | 307 | ENSG00000196975 |
| 28 | P60842 | NA | EIF4A1 | eukaryotic translation initiation factor 4A1 | 1973 | ENSG00000161960 |
| 29 | P04080 | NA | CSTB | cystatin B (stefin B) | 1476 | ENSG00000160213 |
| 30 | P26639 | NA | TARS | threonyl-tRNA synthetase | 6897 | ENSG00000113407 |
| 31 | P04083 | NA | ANXA1 | annexin A1 | 301 | ENSG00000135046 |
| 32 | P02768 | NA | ALB | albumin | 213 | ENSG00000163631 |
| 33 | P41091 | NA | EIF2S3 | eukaryotic translation initiation factor 2, subunit 3 gamma, 52kDa | 1968 | ENSG00000130741 |
| 34 | P05556 | NA | ITGB1 | integrin, beta 1 (fibronectin receptor, beta polypeptide, antigen CD29 includes MDF2, MSK12) | 3688 | ENSG00000150093 |
| 35 | Q15758 | NA | SLC1A5 | solute carrier family 1 (neutral amino acid transporter), member 5 | 6510 | ENSG00000105281 |

  
  

| **Database:cellular component      &nbspName:intracellular part      &nbspID:GO:0044424** | | | | | | |
| --- | --- | --- | --- | --- | --- | --- |
| C=12096; O=38; E=27.55; R=1.38; rawP=4.88e-06; adjP=6.64e-05 | | | | | | |
| Index | UserID | Value | Gene Symbol | Gene Name | EntrezGene | Ensembl |
| 1 | P68371 | NA | TUBB4B | tubulin, beta 4B class IVb | 10383 | ENSG00000188229 |
| 2 | P61225 | NA | RAP2B | RAP2B, member of RAS oncogene family | 5912 | ENSG00000181467 |
| 3 | Q04828 | NA | AKR1C1 | aldo-keto reductase family 1, member C1 (dihydrodiol dehydrogenase 1; 20-alpha (3-alpha)-hydroxysteroid dehydrogenase) | 1645 | ENSG00000187134 |
| 4 | P49588 | NA | AARS | alanyl-tRNA synthetase | 16 | ENSG00000090861 |
| 5 | Q86UP2 | NA | KTN1 | kinectin 1 (kinesin receptor) | 3895 | ENSG00000126777 |
| 6 | P68104 | NA | EEF1A1 | eukaryotic translation elongation factor 1 alpha 1 | 1915 | ENSG00000156508 |
| 7 | O75369 | NA | FLNB | filamin B, beta | 2317 | ENSG00000136068 |
| 8 | P02786 | NA | TFRC | transferrin receptor (p90, CD71) | 7037 | ENSG00000072274 |
| 9 | P24534 | NA | EEF1B2 | eukaryotic translation elongation factor 1 beta 2 | 1933 | ENSG00000114942 |
| 10 | O00425 | NA | IGF2BP3 | insulin-like growth factor 2 mRNA binding protein 3 | 10643 | ENSG00000136231 |
| 11 | P17655 | NA | CAPN2 | calpain 2, (m/II) large subunit | 824 | ENSG00000162909 |
| 12 | P56537 | NA | EIF6 | eukaryotic translation initiation factor 6 | 3692 | ENSG00000242372 |
| 13 | P22102 | NA | GART | phosphoribosylglycinamide formyltransferase, phosphoribosylglycinamide synthetase, phosphoribosylaminoimidazole synthetase | 2618 | ENSG00000159131 |
| 14 | P16615 | NA | ATP2A2 | ATPase, Ca++ transporting, cardiac muscle, slow twitch 2 | 488 | ENSG00000174437 |
| 15 | Q92974 | NA | ARHGEF2 | Rho/Rac guanine nucleotide exchange factor (GEF) 2 | 9181 | ENSG00000116584 |
| 16 | P17174 | NA | GOT1 | glutamic-oxaloacetic transaminase 1, soluble (aspartate aminotransferase 1) | 2805 | ENSG00000120053 |
| 17 | O00194 | NA | RAB27B | RAB27B, member RAS oncogene family | 5874 | ENSG00000041353 |
| 18 | Q6PIU2 | NA | NCEH1 | neutral cholesterol ester hydrolase 1 | 57552 | ENSG00000144959 |
| 19 | P18206 | NA | VCL | vinculin | 7414 | ENSG00000035403 |
| 20 | P51648 | NA | ALDH3A2 | aldehyde dehydrogenase 3 family, member A2 | 224 | ENSG00000072210 |
| 21 | P29692 | NA | EEF1D | eukaryotic translation elongation factor 1 delta (guanine nucleotide exchange protein) | 1936 | ENSG00000104529 |
| 22 | P11387 | NA | TOP1 | topoisomerase (DNA) I | 7150 | ENSG00000198900 |
| 23 | Q9UK76 | NA | HN1 | hematological and neurological expressed 1 | 51155 | ENSG00000189159 |
| 24 | Q16881 | NA | TXNRD1 | thioredoxin reductase 1 | 7296 | ENSG00000198431 |
| 25 | Q9NR30 | NA | DDX21 | DEAD (Asp-Glu-Ala-Asp) box helicase 21 | 9188 | ENSG00000165732 |
| 26 | P08195 | NA | SLC3A2 | solute carrier family 3 (activators of dibasic and neutral amino acid transport), member 2 | 6520 | ENSG00000168003 |
| 27 | P02795 | NA | MT2A | metallothionein 2A | 4502 | ENSG00000125148 |
| 28 | P00491 | NA | PNP | purine nucleoside phosphorylase | 4860 | ENSG00000198805 |
| 29 | P09525 | NA | ANXA4 | annexin A4 | 307 | ENSG00000196975 |
| 30 | P60842 | NA | EIF4A1 | eukaryotic translation initiation factor 4A1 | 1973 | ENSG00000161960 |
| 31 | P04080 | NA | CSTB | cystatin B (stefin B) | 1476 | ENSG00000160213 |
| 32 | P26639 | NA | TARS | threonyl-tRNA synthetase | 6897 | ENSG00000113407 |
| 33 | P04083 | NA | ANXA1 | annexin A1 | 301 | ENSG00000135046 |
| 34 | Q09666 | NA | AHNAK | AHNAK nucleoprotein | 79026 | ENSG00000124942 |
| 35 | P41091 | NA | EIF2S3 | eukaryotic translation initiation factor 2, subunit 3 gamma, 52kDa | 1968 | ENSG00000130741 |
| 36 | P02768 | NA | ALB | albumin | 213 | ENSG00000163631 |
| 37 | P05556 | NA | ITGB1 | integrin, beta 1 (fibronectin receptor, beta polypeptide, antigen CD29 includes MDF2, MSK12) | 3688 | ENSG00000150093 |
| 38 | Q15758 | NA | SLC1A5 | solute carrier family 1 (neutral amino acid transporter), member 5 | 6510 | ENSG00000105281 |

  
  

| **Database:cellular component      &nbspName:intracellular      &nbspID:GO:0005622** | | | | | | |
| --- | --- | --- | --- | --- | --- | --- |
| C=12412; O=38; E=28.27; R=1.34; rawP=1.30e-05; adjP=0.0001 | | | | | | |
| Index | UserID | Value | Gene Symbol | Gene Name | EntrezGene | Ensembl |
| 1 | P68371 | NA | TUBB4B | tubulin, beta 4B class IVb | 10383 | ENSG00000188229 |
| 2 | P61225 | NA | RAP2B | RAP2B, member of RAS oncogene family | 5912 | ENSG00000181467 |
| 3 | Q04828 | NA | AKR1C1 | aldo-keto reductase family 1, member C1 (dihydrodiol dehydrogenase 1; 20-alpha (3-alpha)-hydroxysteroid dehydrogenase) | 1645 | ENSG00000187134 |
| 4 | P49588 | NA | AARS | alanyl-tRNA synthetase | 16 | ENSG00000090861 |
| 5 | Q86UP2 | NA | KTN1 | kinectin 1 (kinesin receptor) | 3895 | ENSG00000126777 |
| 6 | P68104 | NA | EEF1A1 | eukaryotic translation elongation factor 1 alpha 1 | 1915 | ENSG00000156508 |
| 7 | O75369 | NA | FLNB | filamin B, beta | 2317 | ENSG00000136068 |
| 8 | P02786 | NA | TFRC | transferrin receptor (p90, CD71) | 7037 | ENSG00000072274 |
| 9 | P24534 | NA | EEF1B2 | eukaryotic translation elongation factor 1 beta 2 | 1933 | ENSG00000114942 |
| 10 | O00425 | NA | IGF2BP3 | insulin-like growth factor 2 mRNA binding protein 3 | 10643 | ENSG00000136231 |
| 11 | P17655 | NA | CAPN2 | calpain 2, (m/II) large subunit | 824 | ENSG00000162909 |
| 12 | P56537 | NA | EIF6 | eukaryotic translation initiation factor 6 | 3692 | ENSG00000242372 |
| 13 | P22102 | NA | GART | phosphoribosylglycinamide formyltransferase, phosphoribosylglycinamide synthetase, phosphoribosylaminoimidazole synthetase | 2618 | ENSG00000159131 |
| 14 | P16615 | NA | ATP2A2 | ATPase, Ca++ transporting, cardiac muscle, slow twitch 2 | 488 | ENSG00000174437 |
| 15 | Q92974 | NA | ARHGEF2 | Rho/Rac guanine nucleotide exchange factor (GEF) 2 | 9181 | ENSG00000116584 |
| 16 | P17174 | NA | GOT1 | glutamic-oxaloacetic transaminase 1, soluble (aspartate aminotransferase 1) | 2805 | ENSG00000120053 |
| 17 | O00194 | NA | RAB27B | RAB27B, member RAS oncogene family | 5874 | ENSG00000041353 |
| 18 | Q6PIU2 | NA | NCEH1 | neutral cholesterol ester hydrolase 1 | 57552 | ENSG00000144959 |
| 19 | P18206 | NA | VCL | vinculin | 7414 | ENSG00000035403 |
| 20 | P51648 | NA | ALDH3A2 | aldehyde dehydrogenase 3 family, member A2 | 224 | ENSG00000072210 |
| 21 | P29692 | NA | EEF1D | eukaryotic translation elongation factor 1 delta (guanine nucleotide exchange protein) | 1936 | ENSG00000104529 |
| 22 | P11387 | NA | TOP1 | topoisomerase (DNA) I | 7150 | ENSG00000198900 |
| 23 | Q9UK76 | NA | HN1 | hematological and neurological expressed 1 | 51155 | ENSG00000189159 |
| 24 | Q16881 | NA | TXNRD1 | thioredoxin reductase 1 | 7296 | ENSG00000198431 |
| 25 | Q9NR30 | NA | DDX21 | DEAD (Asp-Glu-Ala-Asp) box helicase 21 | 9188 | ENSG00000165732 |
| 26 | P08195 | NA | SLC3A2 | solute carrier family 3 (activators of dibasic and neutral amino acid transport), member 2 | 6520 | ENSG00000168003 |
| 27 | P02795 | NA | MT2A | metallothionein 2A | 4502 | ENSG00000125148 |
| 28 | P00491 | NA | PNP | purine nucleoside phosphorylase | 4860 | ENSG00000198805 |
| 29 | P09525 | NA | ANXA4 | annexin A4 | 307 | ENSG00000196975 |
| 30 | P60842 | NA | EIF4A1 | eukaryotic translation initiation factor 4A1 | 1973 | ENSG00000161960 |
| 31 | P04080 | NA | CSTB | cystatin B (stefin B) | 1476 | ENSG00000160213 |
| 32 | P26639 | NA | TARS | threonyl-tRNA synthetase | 6897 | ENSG00000113407 |
| 33 | P04083 | NA | ANXA1 | annexin A1 | 301 | ENSG00000135046 |
| 34 | Q09666 | NA | AHNAK | AHNAK nucleoprotein | 79026 | ENSG00000124942 |
| 35 | P41091 | NA | EIF2S3 | eukaryotic translation initiation factor 2, subunit 3 gamma, 52kDa | 1968 | ENSG00000130741 |
| 36 | P02768 | NA | ALB | albumin | 213 | ENSG00000163631 |
| 37 | P05556 | NA | ITGB1 | integrin, beta 1 (fibronectin receptor, beta polypeptide, antigen CD29 includes MDF2, MSK12) | 3688 | ENSG00000150093 |
| 38 | Q15758 | NA | SLC1A5 | solute carrier family 1 (neutral amino acid transporter), member 5 | 6510 | ENSG00000105281 |

  
  

| **Database:cellular component      &nbspName:pigment granule      &nbspID:GO:0048770** | | | | | | |
| --- | --- | --- | --- | --- | --- | --- |
| C=93; O=4; E=0.21; R=18.88; rawP=5.78e-05; adjP=0.0005 | | | | | | |
| Index | UserID | Value | Gene Symbol | Gene Name | EntrezGene | Ensembl |
| 1 | P08195 | NA | SLC3A2 | solute carrier family 3 (activators of dibasic and neutral amino acid transport), member 2 | 6520 | ENSG00000168003 |
| 2 | Q15758 | NA | SLC1A5 | solute carrier family 1 (neutral amino acid transporter), member 5 | 6510 | ENSG00000105281 |
| 3 | P02786 | NA | TFRC | transferrin receptor (p90, CD71) | 7037 | ENSG00000072274 |
| 4 | P05556 | NA | ITGB1 | integrin, beta 1 (fibronectin receptor, beta polypeptide, antigen CD29 includes MDF2, MSK12) | 3688 | ENSG00000150093 |

  
  

| **Database:cellular component      &nbspName:melanosome      &nbspID:GO:0042470** | | | | | | |
| --- | --- | --- | --- | --- | --- | --- |
| C=93; O=4; E=0.21; R=18.88; rawP=5.78e-05; adjP=0.0005 | | | | | | |
| Index | UserID | Value | Gene Symbol | Gene Name | EntrezGene | Ensembl |
| 1 | P08195 | NA | SLC3A2 | solute carrier family 3 (activators of dibasic and neutral amino acid transport), member 2 | 6520 | ENSG00000168003 |
| 2 | Q15758 | NA | SLC1A5 | solute carrier family 1 (neutral amino acid transporter), member 5 | 6510 | ENSG00000105281 |
| 3 | P02786 | NA | TFRC | transferrin receptor (p90, CD71) | 7037 | ENSG00000072274 |
| 4 | P05556 | NA | ITGB1 | integrin, beta 1 (fibronectin receptor, beta polypeptide, antigen CD29 includes MDF2, MSK12) | 3688 | ENSG00000150093 |

  
  

| **Database:cellular component      &nbspName:cell part      &nbspID:GO:0044464** | | | | | | |
| --- | --- | --- | --- | --- | --- | --- |
| C=14405; O=38; E=32.81; R=1.16; rawP=0.0038; adjP=0.0215 | | | | | | |
| Index | UserID | Value | Gene Symbol | Gene Name | EntrezGene | Ensembl |
| 1 | P68371 | NA | TUBB4B | tubulin, beta 4B class IVb | 10383 | ENSG00000188229 |
| 2 | P61225 | NA | RAP2B | RAP2B, member of RAS oncogene family | 5912 | ENSG00000181467 |
| 3 | Q04828 | NA | AKR1C1 | aldo-keto reductase family 1, member C1 (dihydrodiol dehydrogenase 1; 20-alpha (3-alpha)-hydroxysteroid dehydrogenase) | 1645 | ENSG00000187134 |
| 4 | P49588 | NA | AARS | alanyl-tRNA synthetase | 16 | ENSG00000090861 |
| 5 | Q86UP2 | NA | KTN1 | kinectin 1 (kinesin receptor) | 3895 | ENSG00000126777 |
| 6 | P68104 | NA | EEF1A1 | eukaryotic translation elongation factor 1 alpha 1 | 1915 | ENSG00000156508 |
| 7 | O75369 | NA | FLNB | filamin B, beta | 2317 | ENSG00000136068 |
| 8 | P02786 | NA | TFRC | transferrin receptor (p90, CD71) | 7037 | ENSG00000072274 |
| 9 | P24534 | NA | EEF1B2 | eukaryotic translation elongation factor 1 beta 2 | 1933 | ENSG00000114942 |
| 10 | O00425 | NA | IGF2BP3 | insulin-like growth factor 2 mRNA binding protein 3 | 10643 | ENSG00000136231 |
| 11 | P17655 | NA | CAPN2 | calpain 2, (m/II) large subunit | 824 | ENSG00000162909 |
| 12 | P56537 | NA | EIF6 | eukaryotic translation initiation factor 6 | 3692 | ENSG00000242372 |
| 13 | P22102 | NA | GART | phosphoribosylglycinamide formyltransferase, phosphoribosylglycinamide synthetase, phosphoribosylaminoimidazole synthetase | 2618 | ENSG00000159131 |
| 14 | P16615 | NA | ATP2A2 | ATPase, Ca++ transporting, cardiac muscle, slow twitch 2 | 488 | ENSG00000174437 |
| 15 | Q92974 | NA | ARHGEF2 | Rho/Rac guanine nucleotide exchange factor (GEF) 2 | 9181 | ENSG00000116584 |
| 16 | P17174 | NA | GOT1 | glutamic-oxaloacetic transaminase 1, soluble (aspartate aminotransferase 1) | 2805 | ENSG00000120053 |
| 17 | O00194 | NA | RAB27B | RAB27B, member RAS oncogene family | 5874 | ENSG00000041353 |
| 18 | Q6PIU2 | NA | NCEH1 | neutral cholesterol ester hydrolase 1 | 57552 | ENSG00000144959 |
| 19 | P18206 | NA | VCL | vinculin | 7414 | ENSG00000035403 |
| 20 | P51648 | NA | ALDH3A2 | aldehyde dehydrogenase 3 family, member A2 | 224 | ENSG00000072210 |
| 21 | P29692 | NA | EEF1D | eukaryotic translation elongation factor 1 delta (guanine nucleotide exchange protein) | 1936 | ENSG00000104529 |
| 22 | P11387 | NA | TOP1 | topoisomerase (DNA) I | 7150 | ENSG00000198900 |
| 23 | Q9UK76 | NA | HN1 | hematological and neurological expressed 1 | 51155 | ENSG00000189159 |
| 24 | Q16881 | NA | TXNRD1 | thioredoxin reductase 1 | 7296 | ENSG00000198431 |
| 25 | Q9NR30 | NA | DDX21 | DEAD (Asp-Glu-Ala-Asp) box helicase 21 | 9188 | ENSG00000165732 |
| 26 | P08195 | NA | SLC3A2 | solute carrier family 3 (activators of dibasic and neutral amino acid transport), member 2 | 6520 | ENSG00000168003 |
| 27 | P02795 | NA | MT2A | metallothionein 2A | 4502 | ENSG00000125148 |
| 28 | P00491 | NA | PNP | purine nucleoside phosphorylase | 4860 | ENSG00000198805 |
| 29 | P09525 | NA | ANXA4 | annexin A4 | 307 | ENSG00000196975 |
| 30 | P60842 | NA | EIF4A1 | eukaryotic translation initiation factor 4A1 | 1973 | ENSG00000161960 |
| 31 | P04080 | NA | CSTB | cystatin B (stefin B) | 1476 | ENSG00000160213 |
| 32 | P26639 | NA | TARS | threonyl-tRNA synthetase | 6897 | ENSG00000113407 |
| 33 | P04083 | NA | ANXA1 | annexin A1 | 301 | ENSG00000135046 |
| 34 | Q09666 | NA | AHNAK | AHNAK nucleoprotein | 79026 | ENSG00000124942 |
| 35 | P41091 | NA | EIF2S3 | eukaryotic translation initiation factor 2, subunit 3 gamma, 52kDa | 1968 | ENSG00000130741 |
| 36 | P02768 | NA | ALB | albumin | 213 | ENSG00000163631 |
| 37 | P05556 | NA | ITGB1 | integrin, beta 1 (fibronectin receptor, beta polypeptide, antigen CD29 includes MDF2, MSK12) | 3688 | ENSG00000150093 |
| 38 | Q15758 | NA | SLC1A5 | solute carrier family 1 (neutral amino acid transporter), member 5 | 6510 | ENSG00000105281 |

  
  

| **Database:cellular component      &nbspName:cell      &nbspID:GO:0005623** | | | | | | |
| --- | --- | --- | --- | --- | --- | --- |
| C=14406; O=38; E=32.82; R=1.16; rawP=0.0038; adjP=0.0215 | | | | | | |
| Index | UserID | Value | Gene Symbol | Gene Name | EntrezGene | Ensembl |
| 1 | P68371 | NA | TUBB4B | tubulin, beta 4B class IVb | 10383 | ENSG00000188229 |
| 2 | P61225 | NA | RAP2B | RAP2B, member of RAS oncogene family | 5912 | ENSG00000181467 |
| 3 | Q04828 | NA | AKR1C1 | aldo-keto reductase family 1, member C1 (dihydrodiol dehydrogenase 1; 20-alpha (3-alpha)-hydroxysteroid dehydrogenase) | 1645 | ENSG00000187134 |
| 4 | P49588 | NA | AARS | alanyl-tRNA synthetase | 16 | ENSG00000090861 |
| 5 | Q86UP2 | NA | KTN1 | kinectin 1 (kinesin receptor) | 3895 | ENSG00000126777 |
| 6 | P68104 | NA | EEF1A1 | eukaryotic translation elongation factor 1 alpha 1 | 1915 | ENSG00000156508 |
| 7 | O75369 | NA | FLNB | filamin B, beta | 2317 | ENSG00000136068 |
| 8 | P02786 | NA | TFRC | transferrin receptor (p90, CD71) | 7037 | ENSG00000072274 |
| 9 | P24534 | NA | EEF1B2 | eukaryotic translation elongation factor 1 beta 2 | 1933 | ENSG00000114942 |
| 10 | O00425 | NA | IGF2BP3 | insulin-like growth factor 2 mRNA binding protein 3 | 10643 | ENSG00000136231 |
| 11 | P17655 | NA | CAPN2 | calpain 2, (m/II) large subunit | 824 | ENSG00000162909 |
| 12 | P56537 | NA | EIF6 | eukaryotic translation initiation factor 6 | 3692 | ENSG00000242372 |
| 13 | P22102 | NA | GART | phosphoribosylglycinamide formyltransferase, phosphoribosylglycinamide synthetase, phosphoribosylaminoimidazole synthetase | 2618 | ENSG00000159131 |
| 14 | P16615 | NA | ATP2A2 | ATPase, Ca++ transporting, cardiac muscle, slow twitch 2 | 488 | ENSG00000174437 |
| 15 | Q92974 | NA | ARHGEF2 | Rho/Rac guanine nucleotide exchange factor (GEF) 2 | 9181 | ENSG00000116584 |
| 16 | P17174 | NA | GOT1 | glutamic-oxaloacetic transaminase 1, soluble (aspartate aminotransferase 1) | 2805 | ENSG00000120053 |
| 17 | O00194 | NA | RAB27B | RAB27B, member RAS oncogene family | 5874 | ENSG00000041353 |
| 18 | Q6PIU2 | NA | NCEH1 | neutral cholesterol ester hydrolase 1 | 57552 | ENSG00000144959 |
| 19 | P18206 | NA | VCL | vinculin | 7414 | ENSG00000035403 |
| 20 | P51648 | NA | ALDH3A2 | aldehyde dehydrogenase 3 family, member A2 | 224 | ENSG00000072210 |
| 21 | P29692 | NA | EEF1D | eukaryotic translation elongation factor 1 delta (guanine nucleotide exchange protein) | 1936 | ENSG00000104529 |
| 22 | P11387 | NA | TOP1 | topoisomerase (DNA) I | 7150 | ENSG00000198900 |
| 23 | Q9UK76 | NA | HN1 | hematological and neurological expressed 1 | 51155 | ENSG00000189159 |
| 24 | Q16881 | NA | TXNRD1 | thioredoxin reductase 1 | 7296 | ENSG00000198431 |
| 25 | Q9NR30 | NA | DDX21 | DEAD (Asp-Glu-Ala-Asp) box helicase 21 | 9188 | ENSG00000165732 |
| 26 | P08195 | NA | SLC3A2 | solute carrier family 3 (activators of dibasic and neutral amino acid transport), member 2 | 6520 | ENSG00000168003 |
| 27 | P02795 | NA | MT2A | metallothionein 2A | 4502 | ENSG00000125148 |
| 28 | P00491 | NA | PNP | purine nucleoside phosphorylase | 4860 | ENSG00000198805 |
| 29 | P09525 | NA | ANXA4 | annexin A4 | 307 | ENSG00000196975 |
| 30 | P60842 | NA | EIF4A1 | eukaryotic translation initiation factor 4A1 | 1973 | ENSG00000161960 |
| 31 | P04080 | NA | CSTB | cystatin B (stefin B) | 1476 | ENSG00000160213 |
| 32 | P26639 | NA | TARS | threonyl-tRNA synthetase | 6897 | ENSG00000113407 |
| 33 | P04083 | NA | ANXA1 | annexin A1 | 301 | ENSG00000135046 |
| 34 | Q09666 | NA | AHNAK | AHNAK nucleoprotein | 79026 | ENSG00000124942 |
| 35 | P41091 | NA | EIF2S3 | eukaryotic translation initiation factor 2, subunit 3 gamma, 52kDa | 1968 | ENSG00000130741 |
| 36 | P02768 | NA | ALB | albumin | 213 | ENSG00000163631 |
| 37 | P05556 | NA | ITGB1 | integrin, beta 1 (fibronectin receptor, beta polypeptide, antigen CD29 includes MDF2, MSK12) | 3688 | ENSG00000150093 |
| 38 | Q15758 | NA | SLC1A5 | solute carrier family 1 (neutral amino acid transporter), member 5 | 6510 | ENSG00000105281 |

  
  

| **Database:cellular component      &nbspName:membrane-bounded vesicle      &nbspID:GO:0031988** | | | | | | |
| --- | --- | --- | --- | --- | --- | --- |
| C=882; O=6; E=2.01; R=2.99; rawP=0.0139; adjP=0.0556 | | | | | | |
| Index | UserID | Value | Gene Symbol | Gene Name | EntrezGene | Ensembl |
| 1 | P08195 | NA | SLC3A2 | solute carrier family 3 (activators of dibasic and neutral amino acid transport), member 2 | 6520 | ENSG00000168003 |
| 2 | Q15758 | NA | SLC1A5 | solute carrier family 1 (neutral amino acid transporter), member 5 | 6510 | ENSG00000105281 |
| 3 | O00194 | NA | RAB27B | RAB27B, member RAS oncogene family | 5874 | ENSG00000041353 |
| 4 | P02768 | NA | ALB | albumin | 213 | ENSG00000163631 |
| 5 | P02786 | NA | TFRC | transferrin receptor (p90, CD71) | 7037 | ENSG00000072274 |
| 6 | P05556 | NA | ITGB1 | integrin, beta 1 (fibronectin receptor, beta polypeptide, antigen CD29 includes MDF2, MSK12) | 3688 | ENSG00000150093 |

  
  

| **Database:cellular component      &nbspName:cytoplasmic membrane-bounded vesicle      &nbspID:GO:0016023** | | | | | | |
| --- | --- | --- | --- | --- | --- | --- |
| C=861; O=6; E=1.96; R=3.06; rawP=0.0125; adjP=0.0556 | | | | | | |
| Index | UserID | Value | Gene Symbol | Gene Name | EntrezGene | Ensembl |
| 1 | P08195 | NA | SLC3A2 | solute carrier family 3 (activators of dibasic and neutral amino acid transport), member 2 | 6520 | ENSG00000168003 |
| 2 | Q15758 | NA | SLC1A5 | solute carrier family 1 (neutral amino acid transporter), member 5 | 6510 | ENSG00000105281 |
| 3 | O00194 | NA | RAB27B | RAB27B, member RAS oncogene family | 5874 | ENSG00000041353 |
| 4 | P02768 | NA | ALB | albumin | 213 | ENSG00000163631 |
| 5 | P02786 | NA | TFRC | transferrin receptor (p90, CD71) | 7037 | ENSG00000072274 |
| 6 | P05556 | NA | ITGB1 | integrin, beta 1 (fibronectin receptor, beta polypeptide, antigen CD29 includes MDF2, MSK12) | 3688 | ENSG00000150093 |

  
  

| **Database:cellular component      &nbspName:anchoring junction      &nbspID:GO:0070161** | | | | | | |
| --- | --- | --- | --- | --- | --- | --- |
| C=218; O=3; E=0.50; R=6.04; rawP=0.0133; adjP=0.0556 | | | | | | |
| Index | UserID | Value | Gene Symbol | Gene Name | EntrezGene | Ensembl |
| 1 | P18206 | NA | VCL | vinculin | 7414 | ENSG00000035403 |
| 2 | O75369 | NA | FLNB | filamin B, beta | 2317 | ENSG00000136068 |
| 3 | P05556 | NA | ITGB1 | integrin, beta 1 (fibronectin receptor, beta polypeptide, antigen CD29 includes MDF2, MSK12) | 3688 | ENSG00000150093 |

  
  

| **Database:cellular component      &nbspName:cytoplasmic vesicle      &nbspID:GO:0031410** | | | | | | |
| --- | --- | --- | --- | --- | --- | --- |
| C=927; O=6; E=2.11; R=2.84; rawP=0.0174; adjP=0.0657 | | | | | | |
| Index | UserID | Value | Gene Symbol | Gene Name | EntrezGene | Ensembl |
| 1 | P08195 | NA | SLC3A2 | solute carrier family 3 (activators of dibasic and neutral amino acid transport), member 2 | 6520 | ENSG00000168003 |
| 2 | Q15758 | NA | SLC1A5 | solute carrier family 1 (neutral amino acid transporter), member 5 | 6510 | ENSG00000105281 |
| 3 | O00194 | NA | RAB27B | RAB27B, member RAS oncogene family | 5874 | ENSG00000041353 |
| 4 | P02768 | NA | ALB | albumin | 213 | ENSG00000163631 |
| 5 | P02786 | NA | TFRC | transferrin receptor (p90, CD71) | 7037 | ENSG00000072274 |
| 6 | P05556 | NA | ITGB1 | integrin, beta 1 (fibronectin receptor, beta polypeptide, antigen CD29 includes MDF2, MSK12) | 3688 | ENSG00000150093 |

  
  

| **Database:cellular component      &nbspName:secretory granule      &nbspID:GO:0030141** | | | | | | |
| --- | --- | --- | --- | --- | --- | --- |
| C=257; O=3; E=0.59; R=5.12; rawP=0.0205; adjP=0.0721 | | | | | | |
| Index | UserID | Value | Gene Symbol | Gene Name | EntrezGene | Ensembl |
| 1 | O00194 | NA | RAB27B | RAB27B, member RAS oncogene family | 5874 | ENSG00000041353 |
| 2 | P02768 | NA | ALB | albumin | 213 | ENSG00000163631 |
| 3 | P05556 | NA | ITGB1 | integrin, beta 1 (fibronectin receptor, beta polypeptide, antigen CD29 includes MDF2, MSK12) | 3688 | ENSG00000150093 |

  
  

| **Database:cellular component      &nbspName:vesicle      &nbspID:GO:0031982** | | | | | | |
| --- | --- | --- | --- | --- | --- | --- |
| C=969; O=6; E=2.21; R=2.72; rawP=0.0212; adjP=0.0721 | | | | | | |
| Index | UserID | Value | Gene Symbol | Gene Name | EntrezGene | Ensembl |
| 1 | P08195 | NA | SLC3A2 | solute carrier family 3 (activators of dibasic and neutral amino acid transport), member 2 | 6520 | ENSG00000168003 |
| 2 | Q15758 | NA | SLC1A5 | solute carrier family 1 (neutral amino acid transporter), member 5 | 6510 | ENSG00000105281 |
| 3 | O00194 | NA | RAB27B | RAB27B, member RAS oncogene family | 5874 | ENSG00000041353 |
| 4 | P02768 | NA | ALB | albumin | 213 | ENSG00000163631 |
| 5 | P02786 | NA | TFRC | transferrin receptor (p90, CD71) | 7037 | ENSG00000072274 |
| 6 | P05556 | NA | ITGB1 | integrin, beta 1 (fibronectin receptor, beta polypeptide, antigen CD29 includes MDF2, MSK12) | 3688 | ENSG00000150093 |

  
  

| **Database:cellular component      &nbspName:plasma membrane part      &nbspID:GO:0044459** | | | | | | |
| --- | --- | --- | --- | --- | --- | --- |
| C=1908; O=9; E=4.35; R=2.07; rawP=0.0248; adjP=0.0767 | | | | | | |
| Index | UserID | Value | Gene Symbol | Gene Name | EntrezGene | Ensembl |
| 1 | P08195 | NA | SLC3A2 | solute carrier family 3 (activators of dibasic and neutral amino acid transport), member 2 | 6520 | ENSG00000168003 |
| 2 | P16615 | NA | ATP2A2 | ATPase, Ca++ transporting, cardiac muscle, slow twitch 2 | 488 | ENSG00000174437 |
| 3 | P05556 | NA | ITGB1 | integrin, beta 1 (fibronectin receptor, beta polypeptide, antigen CD29 includes MDF2, MSK12) | 3688 | ENSG00000150093 |
| 4 | Q92974 | NA | ARHGEF2 | Rho/Rac guanine nucleotide exchange factor (GEF) 2 | 9181 | ENSG00000116584 |
| 5 | Q15758 | NA | SLC1A5 | solute carrier family 1 (neutral amino acid transporter), member 5 | 6510 | ENSG00000105281 |
| 6 | O00194 | NA | RAB27B | RAB27B, member RAS oncogene family | 5874 | ENSG00000041353 |
| 7 | Q86UP2 | NA | KTN1 | kinectin 1 (kinesin receptor) | 3895 | ENSG00000126777 |
| 8 | P02786 | NA | TFRC | transferrin receptor (p90, CD71) | 7037 | ENSG00000072274 |
| 9 | P04083 | NA | ANXA1 | annexin A1 | 301 | ENSG00000135046 |

  
  

| **Database:cellular component      &nbspName:protein complex      &nbspID:GO:0043234** | | | | | | |
| --- | --- | --- | --- | --- | --- | --- |
| C=3262; O=13; E=7.43; R=1.75; rawP=0.0241; adjP=0.0767 | | | | | | |
| Index | UserID | Value | Gene Symbol | Gene Name | EntrezGene | Ensembl |
| 1 | P56537 | NA | EIF6 | eukaryotic translation initiation factor 6 | 3692 | ENSG00000242372 |
| 2 | P16615 | NA | ATP2A2 | ATPase, Ca++ transporting, cardiac muscle, slow twitch 2 | 488 | ENSG00000174437 |
| 3 | P02768 | NA | ALB | albumin | 213 | ENSG00000163631 |
| 4 | P68371 | NA | TUBB4B | tubulin, beta 4B class IVb | 10383 | ENSG00000188229 |
| 5 | P05556 | NA | ITGB1 | integrin, beta 1 (fibronectin receptor, beta polypeptide, antigen CD29 includes MDF2, MSK12) | 3688 | ENSG00000150093 |
| 6 | Q92974 | NA | ARHGEF2 | Rho/Rac guanine nucleotide exchange factor (GEF) 2 | 9181 | ENSG00000116584 |
| 7 | P60842 | NA | EIF4A1 | eukaryotic translation initiation factor 4A1 | 1973 | ENSG00000161960 |
| 8 | P18206 | NA | VCL | vinculin | 7414 | ENSG00000035403 |
| 9 | P68104 | NA | EEF1A1 | eukaryotic translation elongation factor 1 alpha 1 | 1915 | ENSG00000156508 |
| 10 | P29692 | NA | EEF1D | eukaryotic translation elongation factor 1 delta (guanine nucleotide exchange protein) | 1936 | ENSG00000104529 |
| 11 | P04083 | NA | ANXA1 | annexin A1 | 301 | ENSG00000135046 |
| 12 | P24534 | NA | EEF1B2 | eukaryotic translation elongation factor 1 beta 2 | 1933 | ENSG00000114942 |
| 13 | P11387 | NA | TOP1 | topoisomerase (DNA) I | 7150 | ENSG00000198900 |

  
  

| **Database:cellular component      &nbspName:intracellular organelle      &nbspID:GO:0043229** | | | | | | |
| --- | --- | --- | --- | --- | --- | --- |
| C=10521; O=30; E=23.97; R=1.25; rawP=0.0278; adjP=0.0775 | | | | | | |
| Index | UserID | Value | Gene Symbol | Gene Name | EntrezGene | Ensembl |
| 1 | Q16881 | NA | TXNRD1 | thioredoxin reductase 1 | 7296 | ENSG00000198431 |
| 2 | Q9NR30 | NA | DDX21 | DEAD (Asp-Glu-Ala-Asp) box helicase 21 | 9188 | ENSG00000165732 |
| 3 | P02795 | NA | MT2A | metallothionein 2A | 4502 | ENSG00000125148 |
| 4 | P08195 | NA | SLC3A2 | solute carrier family 3 (activators of dibasic and neutral amino acid transport), member 2 | 6520 | ENSG00000168003 |
| 5 | P00491 | NA | PNP | purine nucleoside phosphorylase | 4860 | ENSG00000198805 |
| 6 | P68371 | NA | TUBB4B | tubulin, beta 4B class IVb | 10383 | ENSG00000188229 |
| 7 | P61225 | NA | RAP2B | RAP2B, member of RAS oncogene family | 5912 | ENSG00000181467 |
| 8 | P04080 | NA | CSTB | cystatin B (stefin B) | 1476 | ENSG00000160213 |
| 9 | Q86UP2 | NA | KTN1 | kinectin 1 (kinesin receptor) | 3895 | ENSG00000126777 |
| 10 | P68104 | NA | EEF1A1 | eukaryotic translation elongation factor 1 alpha 1 | 1915 | ENSG00000156508 |
| 11 | P26639 | NA | TARS | threonyl-tRNA synthetase | 6897 | ENSG00000113407 |
| 12 | O75369 | NA | FLNB | filamin B, beta | 2317 | ENSG00000136068 |
| 13 | P02786 | NA | TFRC | transferrin receptor (p90, CD71) | 7037 | ENSG00000072274 |
| 14 | P04083 | NA | ANXA1 | annexin A1 | 301 | ENSG00000135046 |
| 15 | O00425 | NA | IGF2BP3 | insulin-like growth factor 2 mRNA binding protein 3 | 10643 | ENSG00000136231 |
| 16 | P17655 | NA | CAPN2 | calpain 2, (m/II) large subunit | 824 | ENSG00000162909 |
| 17 | P56537 | NA | EIF6 | eukaryotic translation initiation factor 6 | 3692 | ENSG00000242372 |
| 18 | Q09666 | NA | AHNAK | AHNAK nucleoprotein | 79026 | ENSG00000124942 |
| 19 | P16615 | NA | ATP2A2 | ATPase, Ca++ transporting, cardiac muscle, slow twitch 2 | 488 | ENSG00000174437 |
| 20 | P02768 | NA | ALB | albumin | 213 | ENSG00000163631 |
| 21 | P05556 | NA | ITGB1 | integrin, beta 1 (fibronectin receptor, beta polypeptide, antigen CD29 includes MDF2, MSK12) | 3688 | ENSG00000150093 |
| 22 | Q92974 | NA | ARHGEF2 | Rho/Rac guanine nucleotide exchange factor (GEF) 2 | 9181 | ENSG00000116584 |
| 23 | P17174 | NA | GOT1 | glutamic-oxaloacetic transaminase 1, soluble (aspartate aminotransferase 1) | 2805 | ENSG00000120053 |
| 24 | Q15758 | NA | SLC1A5 | solute carrier family 1 (neutral amino acid transporter), member 5 | 6510 | ENSG00000105281 |
| 25 | O00194 | NA | RAB27B | RAB27B, member RAS oncogene family | 5874 | ENSG00000041353 |
| 26 | Q6PIU2 | NA | NCEH1 | neutral cholesterol ester hydrolase 1 | 57552 | ENSG00000144959 |
| 27 | P18206 | NA | VCL | vinculin | 7414 | ENSG00000035403 |
| 28 | P51648 | NA | ALDH3A2 | aldehyde dehydrogenase 3 family, member A2 | 224 | ENSG00000072210 |
| 29 | P11387 | NA | TOP1 | topoisomerase (DNA) I | 7150 | ENSG00000198900 |
| 30 | Q9UK76 | NA | HN1 | hematological and neurological expressed 1 | 51155 | ENSG00000189159 |

  
  

| **Database:cellular component      &nbspName:cell-cell junction      &nbspID:GO:0005911** | | | | | | |
| --- | --- | --- | --- | --- | --- | --- |
| C=288; O=3; E=0.66; R=4.57; rawP=0.0275; adjP=0.0775 | | | | | | |
| Index | UserID | Value | Gene Symbol | Gene Name | EntrezGene | Ensembl |
| 1 | Q92974 | NA | ARHGEF2 | Rho/Rac guanine nucleotide exchange factor (GEF) 2 | 9181 | ENSG00000116584 |
| 2 | P18206 | NA | VCL | vinculin | 7414 | ENSG00000035403 |
| 3 | P05556 | NA | ITGB1 | integrin, beta 1 (fibronectin receptor, beta polypeptide, antigen CD29 includes MDF2, MSK12) | 3688 | ENSG00000150093 |

  
  

| **Database:cellular component      &nbspName:organelle      &nbspID:GO:0043226** | | | | | | |
| --- | --- | --- | --- | --- | --- | --- |
| C=10536; O=30; E=24.00; R=1.25; rawP=0.0285; adjP=0.0775 | | | | | | |
| Index | UserID | Value | Gene Symbol | Gene Name | EntrezGene | Ensembl |
| 1 | Q16881 | NA | TXNRD1 | thioredoxin reductase 1 | 7296 | ENSG00000198431 |
| 2 | Q9NR30 | NA | DDX21 | DEAD (Asp-Glu-Ala-Asp) box helicase 21 | 9188 | ENSG00000165732 |
| 3 | P02795 | NA | MT2A | metallothionein 2A | 4502 | ENSG00000125148 |
| 4 | P08195 | NA | SLC3A2 | solute carrier family 3 (activators of dibasic and neutral amino acid transport), member 2 | 6520 | ENSG00000168003 |
| 5 | P00491 | NA | PNP | purine nucleoside phosphorylase | 4860 | ENSG00000198805 |
| 6 | P68371 | NA | TUBB4B | tubulin, beta 4B class IVb | 10383 | ENSG00000188229 |
| 7 | P61225 | NA | RAP2B | RAP2B, member of RAS oncogene family | 5912 | ENSG00000181467 |
| 8 | P04080 | NA | CSTB | cystatin B (stefin B) | 1476 | ENSG00000160213 |
| 9 | Q86UP2 | NA | KTN1 | kinectin 1 (kinesin receptor) | 3895 | ENSG00000126777 |
| 10 | P68104 | NA | EEF1A1 | eukaryotic translation elongation factor 1 alpha 1 | 1915 | ENSG00000156508 |
| 11 | P26639 | NA | TARS | threonyl-tRNA synthetase | 6897 | ENSG00000113407 |
| 12 | O75369 | NA | FLNB | filamin B, beta | 2317 | ENSG00000136068 |
| 13 | P02786 | NA | TFRC | transferrin receptor (p90, CD71) | 7037 | ENSG00000072274 |
| 14 | P04083 | NA | ANXA1 | annexin A1 | 301 | ENSG00000135046 |
| 15 | O00425 | NA | IGF2BP3 | insulin-like growth factor 2 mRNA binding protein 3 | 10643 | ENSG00000136231 |
| 16 | P17655 | NA | CAPN2 | calpain 2, (m/II) large subunit | 824 | ENSG00000162909 |
| 17 | P56537 | NA | EIF6 | eukaryotic translation initiation factor 6 | 3692 | ENSG00000242372 |
| 18 | Q09666 | NA | AHNAK | AHNAK nucleoprotein | 79026 | ENSG00000124942 |
| 19 | P16615 | NA | ATP2A2 | ATPase, Ca++ transporting, cardiac muscle, slow twitch 2 | 488 | ENSG00000174437 |
| 20 | P02768 | NA | ALB | albumin | 213 | ENSG00000163631 |
| 21 | P05556 | NA | ITGB1 | integrin, beta 1 (fibronectin receptor, beta polypeptide, antigen CD29 includes MDF2, MSK12) | 3688 | ENSG00000150093 |
| 22 | Q92974 | NA | ARHGEF2 | Rho/Rac guanine nucleotide exchange factor (GEF) 2 | 9181 | ENSG00000116584 |
| 23 | P17174 | NA | GOT1 | glutamic-oxaloacetic transaminase 1, soluble (aspartate aminotransferase 1) | 2805 | ENSG00000120053 |
| 24 | Q15758 | NA | SLC1A5 | solute carrier family 1 (neutral amino acid transporter), member 5 | 6510 | ENSG00000105281 |
| 25 | O00194 | NA | RAB27B | RAB27B, member RAS oncogene family | 5874 | ENSG00000041353 |
| 26 | Q6PIU2 | NA | NCEH1 | neutral cholesterol ester hydrolase 1 | 57552 | ENSG00000144959 |
| 27 | P18206 | NA | VCL | vinculin | 7414 | ENSG00000035403 |
| 28 | P51648 | NA | ALDH3A2 | aldehyde dehydrogenase 3 family, member A2 | 224 | ENSG00000072210 |
| 29 | P11387 | NA | TOP1 | topoisomerase (DNA) I | 7150 | ENSG00000198900 |
| 30 | Q9UK76 | NA | HN1 | hematological and neurological expressed 1 | 51155 | ENSG00000189159 |

  
  

| **Database:cellular component      &nbspName:endosome membrane      &nbspID:GO:0010008** | | | | | | |
| --- | --- | --- | --- | --- | --- | --- |
| C=313; O=3; E=0.71; R=4.21; rawP=0.0340; adjP=0.0889 | | | | | | |
| Index | UserID | Value | Gene Symbol | Gene Name | EntrezGene | Ensembl |
| 1 | O00194 | NA | RAB27B | RAB27B, member RAS oncogene family | 5874 | ENSG00000041353 |
| 2 | P02786 | NA | TFRC | transferrin receptor (p90, CD71) | 7037 | ENSG00000072274 |
| 3 | P61225 | NA | RAP2B | RAP2B, member of RAS oncogene family | 5912 | ENSG00000181467 |

  
  

| **Database:cellular component      &nbspName:endosomal part      &nbspID:GO:0044440** | | | | | | |
| --- | --- | --- | --- | --- | --- | --- |
| C=322; O=3; E=0.73; R=4.09; rawP=0.0366; adjP=0.0922 | | | | | | |
| Index | UserID | Value | Gene Symbol | Gene Name | EntrezGene | Ensembl |
| 1 | O00194 | NA | RAB27B | RAB27B, member RAS oncogene family | 5874 | ENSG00000041353 |
| 2 | P02786 | NA | TFRC | transferrin receptor (p90, CD71) | 7037 | ENSG00000072274 |
| 3 | P61225 | NA | RAP2B | RAP2B, member of RAS oncogene family | 5912 | ENSG00000181467 |

  
  
  
  

---

WebGestalt is currently developed and maintained by Jing Wang and Bing Zhang at the  Zhang Lab. Other people who have made significant contribution to the project include Dexter Duncan, Stefan Kirov, Zhiao Shi, and Jay Snoddy.  
  
**Funding credits:** NIH/NIAAA (U01 AA016662, U01 AA013512); NIH/NIDA (P01 DA015027); NIH/NIMH (P50 MH078028, P50 MH096972); NIH/NCI (U24 CA159988); NIH/NIGMS (R01 GM088822).
