## Supplementary material for "PGRMC1 phosphorylation and cell plasticity 1: glycolysis, mitochondria, tumor growth": File S8: final_tf_geneset_file_1439190558.html

Anchored HTML File of EIDs

|  |  |
| --- | --- |
|  | WEB-based GEne SeT AnaLysis Toolkit |
|  | ***Translating gene lists into biological insights...*** |

---

  

| **Database:Transcrription Target      &nbspName:hsa\_V$HLF\_01      &nbspID:DB\_ID:2030** | | | | | | |
| --- | --- | --- | --- | --- | --- | --- |
| C=250; O=3; E=0.47; R=6.41; rawP=0.0114; adjP=0.0352 | | | | | | |
| Index | UserID | Value | Gene Symbol | Gene Name | EntrezGene | Ensembl |
| 1 | P60842 | NA | EIF4A1 | eukaryotic translation initiation factor 4A1 | 1973 | ENSG00000161960 |
| 2 | P02768 | NA | ALB | albumin | 213 | ENSG00000163631 |
| 3 | P11387 | NA | TOP1 | topoisomerase (DNA) I | 7150 | ENSG00000198900 |

  
  

| **Database:Transcrription Target      &nbspName:hsa\_V$HIF1\_Q3      &nbspID:DB\_ID:2273** | | | | | | |
| --- | --- | --- | --- | --- | --- | --- |
| C=225; O=3; E=0.42; R=7.12; rawP=0.0085; adjP=0.0352 | | | | | | |
| Index | UserID | Value | Gene Symbol | Gene Name | EntrezGene | Ensembl |
| 1 | Q86UP2 | NA | KTN1 | kinectin 1 (kinesin receptor) | 3895 | ENSG00000126777 |
| 2 | P02786 | NA | TFRC | transferrin receptor (p90, CD71) | 7037 | ENSG00000072274 |
| 3 | O00425 | NA | IGF2BP3 | insulin-like growth factor 2 mRNA binding protein 3 | 10643 | ENSG00000136231 |

  
  

| **Database:Transcrription Target      &nbspName:hsa\_V$MYCMAX\_01      &nbspID:DB\_ID:1923** | | | | | | |
| --- | --- | --- | --- | --- | --- | --- |
| C=250; O=3; E=0.47; R=6.41; rawP=0.0114; adjP=0.0352 | | | | | | |
| Index | UserID | Value | Gene Symbol | Gene Name | EntrezGene | Ensembl |
| 1 | P02786 | NA | TFRC | transferrin receptor (p90, CD71) | 7037 | ENSG00000072274 |
| 2 | P11387 | NA | TOP1 | topoisomerase (DNA) I | 7150 | ENSG00000198900 |
| 3 | P24534 | NA | EEF1B2 | eukaryotic translation elongation factor 1 beta 2 | 1933 | ENSG00000114942 |

  
  

| **Database:Transcrription Target      &nbspName:hsa\_V$E2A\_Q2      &nbspID:DB\_ID:2279** | | | | | | |
| --- | --- | --- | --- | --- | --- | --- |
| C=236; O=3; E=0.44; R=6.79; rawP=0.0097; adjP=0.0352 | | | | | | |
| Index | UserID | Value | Gene Symbol | Gene Name | EntrezGene | Ensembl |
| 1 | Q92974 | NA | ARHGEF2 | Rho/Rac guanine nucleotide exchange factor (GEF) 2 | 9181 | ENSG00000116584 |
| 2 | P16615 | NA | ATP2A2 | ATPase, Ca++ transporting, cardiac muscle, slow twitch 2 | 488 | ENSG00000174437 |
| 3 | Q9UK76 | NA | HN1 | hematological and neurological expressed 1 | 51155 | ENSG00000189159 |

  
  

| **Database:Transcrription Target      &nbspName:hsa\_V$NERF\_Q2      &nbspID:DB\_ID:2163** | | | | | | |
| --- | --- | --- | --- | --- | --- | --- |
| C=243; O=3; E=0.46; R=6.59; rawP=0.0105; adjP=0.0352 | | | | | | |
| Index | UserID | Value | Gene Symbol | Gene Name | EntrezGene | Ensembl |
| 1 | P18206 | NA | VCL | vinculin | 7414 | ENSG00000035403 |
| 2 | P02786 | NA | TFRC | transferrin receptor (p90, CD71) | 7037 | ENSG00000072274 |
| 3 | Q9UK76 | NA | HN1 | hematological and neurological expressed 1 | 51155 | ENSG00000189159 |

  
  

| **Database:Transcrription Target      &nbspName:hsa\_V$CEBPB\_02      &nbspID:DB\_ID:1922** | | | | | | |
| --- | --- | --- | --- | --- | --- | --- |
| C=250; O=3; E=0.47; R=6.41; rawP=0.0114; adjP=0.0352 | | | | | | |
| Index | UserID | Value | Gene Symbol | Gene Name | EntrezGene | Ensembl |
| 1 | P17174 | NA | GOT1 | glutamic-oxaloacetic transaminase 1, soluble (aspartate aminotransferase 1) | 2805 | ENSG00000120053 |
| 2 | P60842 | NA | EIF4A1 | eukaryotic translation initiation factor 4A1 | 1973 | ENSG00000161960 |
| 3 | P11387 | NA | TOP1 | topoisomerase (DNA) I | 7150 | ENSG00000198900 |

  
  

| **Database:Transcrription Target      &nbspName:hsa\_TTAYRTAA\_V$E4BP4\_01      &nbspID:DB\_ID:2411** | | | | | | |
| --- | --- | --- | --- | --- | --- | --- |
| C=251; O=3; E=0.47; R=6.38; rawP=0.0115; adjP=0.0352 | | | | | | |
| Index | UserID | Value | Gene Symbol | Gene Name | EntrezGene | Ensembl |
| 1 | Q09666 | NA | AHNAK | AHNAK nucleoprotein | 79026 | ENSG00000124942 |
| 2 | P02768 | NA | ALB | albumin | 213 | ENSG00000163631 |
| 3 | P61225 | NA | RAP2B | RAP2B, member of RAS oncogene family | 5912 | ENSG00000181467 |

  
  

| **Database:Transcrription Target      &nbspName:hsa\_TGGNNNNNNKCCAR\_UNKNOWN      &nbspID:DB\_ID:2368** | | | | | | |
| --- | --- | --- | --- | --- | --- | --- |
| C=406; O=4; E=0.76; R=5.26; rawP=0.0068; adjP=0.0352 | | | | | | |
| Index | UserID | Value | Gene Symbol | Gene Name | EntrezGene | Ensembl |
| 1 | Q92974 | NA | ARHGEF2 | Rho/Rac guanine nucleotide exchange factor (GEF) 2 | 9181 | ENSG00000116584 |
| 2 | P17174 | NA | GOT1 | glutamic-oxaloacetic transaminase 1, soluble (aspartate aminotransferase 1) | 2805 | ENSG00000120053 |
| 3 | P16615 | NA | ATP2A2 | ATPase, Ca++ transporting, cardiac muscle, slow twitch 2 | 488 | ENSG00000174437 |
| 4 | P18206 | NA | VCL | vinculin | 7414 | ENSG00000035403 |

  
  

| **Database:Transcrription Target      &nbspName:hsa\_V$MAX\_01      &nbspID:DB\_ID:1924** | | | | | | |
| --- | --- | --- | --- | --- | --- | --- |
| C=256; O=3; E=0.48; R=6.26; rawP=0.0121; adjP=0.0352 | | | | | | |
| Index | UserID | Value | Gene Symbol | Gene Name | EntrezGene | Ensembl |
| 1 | P02786 | NA | TFRC | transferrin receptor (p90, CD71) | 7037 | ENSG00000072274 |
| 2 | P11387 | NA | TOP1 | topoisomerase (DNA) I | 7150 | ENSG00000198900 |
| 3 | P24534 | NA | EEF1B2 | eukaryotic translation elongation factor 1 beta 2 | 1933 | ENSG00000114942 |

  
  

| **Database:Transcrription Target      &nbspName:hsa\_GGGCGGR\_V$SP1\_Q6      &nbspID:DB\_ID:2452** | | | | | | |
| --- | --- | --- | --- | --- | --- | --- |
| C=2863; O=13; E=5.36; R=2.42; rawP=0.0015; adjP=0.0352 | | | | | | |
| Index | UserID | Value | Gene Symbol | Gene Name | EntrezGene | Ensembl |
| 1 | Q16881 | NA | TXNRD1 | thioredoxin reductase 1 | 7296 | ENSG00000198431 |
| 2 | Q09666 | NA | AHNAK | AHNAK nucleoprotein | 79026 | ENSG00000124942 |
| 3 | P02795 | NA | MT2A | metallothionein 2A | 4502 | ENSG00000125148 |
| 4 | P41091 | NA | EIF2S3 | eukaryotic translation initiation factor 2, subunit 3 gamma, 52kDa | 1968 | ENSG00000130741 |
| 5 | P09525 | NA | ANXA4 | annexin A4 | 307 | ENSG00000196975 |
| 6 | P05556 | NA | ITGB1 | integrin, beta 1 (fibronectin receptor, beta polypeptide, antigen CD29 includes MDF2, MSK12) | 3688 | ENSG00000150093 |
| 7 | P60842 | NA | EIF4A1 | eukaryotic translation initiation factor 4A1 | 1973 | ENSG00000161960 |
| 8 | P18206 | NA | VCL | vinculin | 7414 | ENSG00000035403 |
| 9 | Q86UP2 | NA | KTN1 | kinectin 1 (kinesin receptor) | 3895 | ENSG00000126777 |
| 10 | P68104 | NA | EEF1A1 | eukaryotic translation elongation factor 1 alpha 1 | 1915 | ENSG00000156508 |
| 11 | P11387 | NA | TOP1 | topoisomerase (DNA) I | 7150 | ENSG00000198900 |
| 12 | Q9UK76 | NA | HN1 | hematological and neurological expressed 1 | 51155 | ENSG00000189159 |
| 13 | O00425 | NA | IGF2BP3 | insulin-like growth factor 2 mRNA binding protein 3 | 10643 | ENSG00000136231 |

  
  

| **Database:Transcrription Target      &nbspName:hsa\_V$SMAD4\_Q6      &nbspID:DB\_ID:2229** | | | | | | |
| --- | --- | --- | --- | --- | --- | --- |
| C=235; O=3; E=0.44; R=6.81; rawP=0.0096; adjP=0.0352 | | | | | | |
| Index | UserID | Value | Gene Symbol | Gene Name | EntrezGene | Ensembl |
| 1 | P60842 | NA | EIF4A1 | eukaryotic translation initiation factor 4A1 | 1973 | ENSG00000161960 |
| 2 | P16615 | NA | ATP2A2 | ATPase, Ca++ transporting, cardiac muscle, slow twitch 2 | 488 | ENSG00000174437 |
| 3 | P68104 | NA | EEF1A1 | eukaryotic translation elongation factor 1 alpha 1 | 1915 | ENSG00000156508 |

  
  

| **Database:Transcrription Target      &nbspName:hsa\_TGANTCA\_V$AP1\_C      &nbspID:DB\_ID:2402** | | | | | | |
| --- | --- | --- | --- | --- | --- | --- |
| C=1092; O=6; E=2.05; R=2.93; rawP=0.0151; adjP=0.0403 | | | | | | |
| Index | UserID | Value | Gene Symbol | Gene Name | EntrezGene | Ensembl |
| 1 | Q92974 | NA | ARHGEF2 | Rho/Rac guanine nucleotide exchange factor (GEF) 2 | 9181 | ENSG00000116584 |
| 2 | O00194 | NA | RAB27B | RAB27B, member RAS oncogene family | 5874 | ENSG00000041353 |
| 3 | Q15758 | NA | SLC1A5 | solute carrier family 1 (neutral amino acid transporter), member 5 | 6510 | ENSG00000105281 |
| 4 | P16615 | NA | ATP2A2 | ATPase, Ca++ transporting, cardiac muscle, slow twitch 2 | 488 | ENSG00000174437 |
| 5 | P18206 | NA | VCL | vinculin | 7414 | ENSG00000035403 |
| 6 | P68104 | NA | EEF1A1 | eukaryotic translation elongation factor 1 alpha 1 | 1915 | ENSG00000156508 |

  
  

| **Database:Transcrription Target      &nbspName:hsa\_GCTNWTTGK\_UNKNOWN      &nbspID:DB\_ID:2247** | | | | | | |
| --- | --- | --- | --- | --- | --- | --- |
| C=294; O=3; E=0.55; R=5.45; rawP=0.0175; adjP=0.0431 | | | | | | |
| Index | UserID | Value | Gene Symbol | Gene Name | EntrezGene | Ensembl |
| 1 | P60842 | NA | EIF4A1 | eukaryotic translation initiation factor 4A1 | 1973 | ENSG00000161960 |
| 2 | P16615 | NA | ATP2A2 | ATPase, Ca++ transporting, cardiac muscle, slow twitch 2 | 488 | ENSG00000174437 |
| 3 | Q9UK76 | NA | HN1 | hematological and neurological expressed 1 | 51155 | ENSG00000189159 |

  
  

| **Database:Transcrription Target      &nbspName:hsa\_CTAWWWATA\_V$RSRFC4\_Q2      &nbspID:DB\_ID:2448** | | | | | | |
| --- | --- | --- | --- | --- | --- | --- |
| C=352; O=3; E=0.66; R=4.55; rawP=0.0279; adjP=0.0595 | | | | | | |
| Index | UserID | Value | Gene Symbol | Gene Name | EntrezGene | Ensembl |
| 1 | Q09666 | NA | AHNAK | AHNAK nucleoprotein | 79026 | ENSG00000124942 |
| 2 | Q86UP2 | NA | KTN1 | kinectin 1 (kinesin receptor) | 3895 | ENSG00000126777 |
| 3 | P61225 | NA | RAP2B | RAP2B, member of RAS oncogene family | 5912 | ENSG00000181467 |

  
  

| **Database:Transcrription Target      &nbspName:hsa\_CATTGTYY\_V$SOX9\_B1      &nbspID:DB\_ID:2451** | | | | | | |
| --- | --- | --- | --- | --- | --- | --- |
| C=350; O=3; E=0.66; R=4.58; rawP=0.0275; adjP=0.0595 | | | | | | |
| Index | UserID | Value | Gene Symbol | Gene Name | EntrezGene | Ensembl |
| 1 | P18206 | NA | VCL | vinculin | 7414 | ENSG00000035403 |
| 2 | P11387 | NA | TOP1 | topoisomerase (DNA) I | 7150 | ENSG00000198900 |
| 3 | Q9UK76 | NA | HN1 | hematological and neurological expressed 1 | 51155 | ENSG00000189159 |

  
  

| **Database:Transcrription Target      &nbspName:hsa\_CACGTG\_V$MYC\_Q2      &nbspID:DB\_ID:2434** | | | | | | |
| --- | --- | --- | --- | --- | --- | --- |
| C=1008; O=5; E=1.89; R=2.65; rawP=0.0387; adjP=0.0774 | | | | | | |
| Index | UserID | Value | Gene Symbol | Gene Name | EntrezGene | Ensembl |
| 1 | P60842 | NA | EIF4A1 | eukaryotic translation initiation factor 4A1 | 1973 | ENSG00000161960 |
| 2 | P02786 | NA | TFRC | transferrin receptor (p90, CD71) | 7037 | ENSG00000072274 |
| 3 | P11387 | NA | TOP1 | topoisomerase (DNA) I | 7150 | ENSG00000198900 |
| 4 | P24534 | NA | EEF1B2 | eukaryotic translation elongation factor 1 beta 2 | 1933 | ENSG00000114942 |
| 5 | O00425 | NA | IGF2BP3 | insulin-like growth factor 2 mRNA binding protein 3 | 10643 | ENSG00000136231 |

  
  

| **Database:Transcrription Target      &nbspName:hsa\_GCCATNTTG\_V$YY1\_Q6      &nbspID:DB\_ID:2459** | | | | | | |
| --- | --- | --- | --- | --- | --- | --- |
| C=416; O=3; E=0.78; R=3.85; rawP=0.0426; adjP=0.0802 | | | | | | |
| Index | UserID | Value | Gene Symbol | Gene Name | EntrezGene | Ensembl |
| 1 | P60842 | NA | EIF4A1 | eukaryotic translation initiation factor 4A1 | 1973 | ENSG00000161960 |
| 2 | Q9UK76 | NA | HN1 | hematological and neurological expressed 1 | 51155 | ENSG00000189159 |
| 3 | P24534 | NA | EEF1B2 | eukaryotic translation elongation factor 1 beta 2 | 1933 | ENSG00000114942 |

  
  

| **Database:Transcrription Target      &nbspName:hsa\_RYTTCCTG\_V$ETS2\_B      &nbspID:DB\_ID:2415** | | | | | | |
| --- | --- | --- | --- | --- | --- | --- |
| C=1060; O=5; E=1.99; R=2.52; rawP=0.0464; adjP=0.0825 | | | | | | |
| Index | UserID | Value | Gene Symbol | Gene Name | EntrezGene | Ensembl |
| 1 | Q92974 | NA | ARHGEF2 | Rho/Rac guanine nucleotide exchange factor (GEF) 2 | 9181 | ENSG00000116584 |
| 2 | Q15758 | NA | SLC1A5 | solute carrier family 1 (neutral amino acid transporter), member 5 | 6510 | ENSG00000105281 |
| 3 | P18206 | NA | VCL | vinculin | 7414 | ENSG00000035403 |
| 4 | O75369 | NA | FLNB | filamin B, beta | 2317 | ENSG00000136068 |
| 5 | P05556 | NA | ITGB1 | integrin, beta 1 (fibronectin receptor, beta polypeptide, antigen CD29 includes MDF2, MSK12) | 3688 | ENSG00000150093 |

  
  

| **Database:Transcrription Target      &nbspName:hsa\_GGGAGGRR\_V$MAZ\_Q6      &nbspID:DB\_ID:2430** | | | | | | |
| --- | --- | --- | --- | --- | --- | --- |
| C=2223; O=8; E=4.16; R=1.92; rawP=0.0505; adjP=0.0851 | | | | | | |
| Index | UserID | Value | Gene Symbol | Gene Name | EntrezGene | Ensembl |
| 1 | P17174 | NA | GOT1 | glutamic-oxaloacetic transaminase 1, soluble (aspartate aminotransferase 1) | 2805 | ENSG00000120053 |
| 2 | P60842 | NA | EIF4A1 | eukaryotic translation initiation factor 4A1 | 1973 | ENSG00000161960 |
| 3 | P22102 | NA | GART | phosphoribosylglycinamide formyltransferase, phosphoribosylglycinamide synthetase, phosphoribosylaminoimidazole synthetase | 2618 | ENSG00000159131 |
| 4 | Q09666 | NA | AHNAK | AHNAK nucleoprotein | 79026 | ENSG00000124942 |
| 5 | P16615 | NA | ATP2A2 | ATPase, Ca++ transporting, cardiac muscle, slow twitch 2 | 488 | ENSG00000174437 |
| 6 | P18206 | NA | VCL | vinculin | 7414 | ENSG00000035403 |
| 7 | P68104 | NA | EEF1A1 | eukaryotic translation elongation factor 1 alpha 1 | 1915 | ENSG00000156508 |
| 8 | O00425 | NA | IGF2BP3 | insulin-like growth factor 2 mRNA binding protein 3 | 10643 | ENSG00000136231 |

  
  
  
  

---

WebGestalt is currently developed and maintained by Jing Wang and Bing Zhang at the  Zhang Lab. Other people who have made significant contribution to the project include Dexter Duncan, Stefan Kirov, Zhiao Shi, and Jay Snoddy.  
  
**Funding credits:** NIH/NIAAA (U01 AA016662, U01 AA013512); NIH/NIDA (P01 DA015027); NIH/NIMH (P50 MH078028, P50 MH096972); NIH/NCI (U24 CA159988); NIH/NIGMS (R01 GM088822).
