## Supplementary material for "PGRMC1 phosphorylation and cell plasticity 1: glycolysis, mitochondria, tumor growth": File S8: final_wiki_geneset_file_1439190558.html

Anchored HTML File of EIDs

|  |  |
| --- | --- |
|  | WEB-based GEne SeT AnaLysis Toolkit |
|  | ***Translating gene lists into biological insights...*** |

---

  

| **Database:Wikipathways pathway      &nbspName:Translation Factors      &nbspID:WP107** | | | | | | |
| --- | --- | --- | --- | --- | --- | --- |
| C=51; O=6; E=0.10; R=62.80; rawP=4.84e-10; adjP=3.39e-09 | | | | | | |
| Index | UserID | Value | Gene Symbol | Gene Name | EntrezGene | Ensembl |
| 1 | P56537 | NA | EIF6 | eukaryotic translation initiation factor 6 | 3692 | ENSG00000242372 |
| 2 | P60842 | NA | EIF4A1 | eukaryotic translation initiation factor 4A1 | 1973 | ENSG00000161960 |
| 3 | P41091 | NA | EIF2S3 | eukaryotic translation initiation factor 2, subunit 3 gamma, 52kDa | 1968 | ENSG00000130741 |
| 4 | P68104 | NA | EEF1A1 | eukaryotic translation elongation factor 1 alpha 1 | 1915 | ENSG00000156508 |
| 5 | P29692 | NA | EEF1D | eukaryotic translation elongation factor 1 delta (guanine nucleotide exchange protein) | 1936 | ENSG00000104529 |
| 6 | P24534 | NA | EEF1B2 | eukaryotic translation elongation factor 1 beta 2 | 1933 | ENSG00000114942 |

  
  

| **Database:Wikipathways pathway      &nbspName:Pathogenic Escherichia coli infection      &nbspID:WP2272** | | | | | | |
| --- | --- | --- | --- | --- | --- | --- |
| C=64; O=3; E=0.12; R=25.02; rawP=0.0002; adjP=0.0007 | | | | | | |
| Index | UserID | Value | Gene Symbol | Gene Name | EntrezGene | Ensembl |
| 1 | Q92974 | NA | ARHGEF2 | Rho/Rac guanine nucleotide exchange factor (GEF) 2 | 9181 | ENSG00000116584 |
| 2 | P68371 | NA | TUBB4B | tubulin, beta 4B class IVb | 10383 | ENSG00000188229 |
| 3 | P05556 | NA | ITGB1 | integrin, beta 1 (fibronectin receptor, beta polypeptide, antigen CD29 includes MDF2, MSK12) | 3688 | ENSG00000150093 |

  
  

| **Database:Wikipathways pathway      &nbspName:Integrin-mediated cell adhesion      &nbspID:WP185** | | | | | | |
| --- | --- | --- | --- | --- | --- | --- |
| C=101; O=3; E=0.19; R=15.85; rawP=0.0009; adjP=0.0021 | | | | | | |
| Index | UserID | Value | Gene Symbol | Gene Name | EntrezGene | Ensembl |
| 1 | P17655 | NA | CAPN2 | calpain 2, (m/II) large subunit | 824 | ENSG00000162909 |
| 2 | P18206 | NA | VCL | vinculin | 7414 | ENSG00000035403 |
| 3 | P05556 | NA | ITGB1 | integrin, beta 1 (fibronectin receptor, beta polypeptide, antigen CD29 includes MDF2, MSK12) | 3688 | ENSG00000150093 |

  
  

| **Database:Wikipathways pathway      &nbspName:Lymphocyte TarBase      &nbspID:WP2004** | | | | | | |
| --- | --- | --- | --- | --- | --- | --- |
| C=533; O=5; E=1.00; R=5.01; rawP=0.0030; adjP=0.0039 | | | | | | |
| Index | UserID | Value | Gene Symbol | Gene Name | EntrezGene | Ensembl |
| 1 | Q16881 | NA | TXNRD1 | thioredoxin reductase 1 | 7296 | ENSG00000198431 |
| 2 | Q6PIU2 | NA | NCEH1 | neutral cholesterol ester hydrolase 1 | 57552 | ENSG00000144959 |
| 3 | O00194 | NA | RAB27B | RAB27B, member RAS oncogene family | 5874 | ENSG00000041353 |
| 4 | P16615 | NA | ATP2A2 | ATPase, Ca++ transporting, cardiac muscle, slow twitch 2 | 488 | ENSG00000174437 |
| 5 | P00491 | NA | PNP | purine nucleoside phosphorylase | 4860 | ENSG00000198805 |

  
  

| **Database:Wikipathways pathway      &nbspName:Leukocyte TarBase      &nbspID:WP2003** | | | | | | |
| --- | --- | --- | --- | --- | --- | --- |
| C=160; O=3; E=0.30; R=10.01; rawP=0.0033; adjP=0.0039 | | | | | | |
| Index | UserID | Value | Gene Symbol | Gene Name | EntrezGene | Ensembl |
| 1 | Q16881 | NA | TXNRD1 | thioredoxin reductase 1 | 7296 | ENSG00000198431 |
| 2 | P16615 | NA | ATP2A2 | ATPase, Ca++ transporting, cardiac muscle, slow twitch 2 | 488 | ENSG00000174437 |
| 3 | P00491 | NA | PNP | purine nucleoside phosphorylase | 4860 | ENSG00000198805 |

  
  

| **Database:Wikipathways pathway      &nbspName:Squamous cell TarBase      &nbspID:WP2006** | | | | | | |
| --- | --- | --- | --- | --- | --- | --- |
| C=150; O=3; E=0.28; R=10.68; rawP=0.0028; adjP=0.0039 | | | | | | |
| Index | UserID | Value | Gene Symbol | Gene Name | EntrezGene | Ensembl |
| 1 | Q16881 | NA | TXNRD1 | thioredoxin reductase 1 | 7296 | ENSG00000198431 |
| 2 | P16615 | NA | ATP2A2 | ATPase, Ca++ transporting, cardiac muscle, slow twitch 2 | 488 | ENSG00000174437 |
| 3 | P00491 | NA | PNP | purine nucleoside phosphorylase | 4860 | ENSG00000198805 |

  
  

| **Database:Wikipathways pathway      &nbspName:Muscle cell TarBase      &nbspID:WP2005** | | | | | | |
| --- | --- | --- | --- | --- | --- | --- |
| C=424; O=4; E=0.79; R=5.04; rawP=0.0079; adjP=0.0079 | | | | | | |
| Index | UserID | Value | Gene Symbol | Gene Name | EntrezGene | Ensembl |
| 1 | Q16881 | NA | TXNRD1 | thioredoxin reductase 1 | 7296 | ENSG00000198431 |
| 2 | Q6PIU2 | NA | NCEH1 | neutral cholesterol ester hydrolase 1 | 57552 | ENSG00000144959 |
| 3 | O00194 | NA | RAB27B | RAB27B, member RAS oncogene family | 5874 | ENSG00000041353 |
| 4 | P16615 | NA | ATP2A2 | ATPase, Ca++ transporting, cardiac muscle, slow twitch 2 | 488 | ENSG00000174437 |

  
  
  
  

---

WebGestalt is currently developed and maintained by Jing Wang and Bing Zhang at the  Zhang Lab. Other people who have made significant contribution to the project include Dexter Duncan, Stefan Kirov, Zhiao Shi, and Jay Snoddy.  
  
**Funding credits:** NIH/NIAAA (U01 AA016662, U01 AA013512); NIH/NIDA (P01 DA015027); NIH/NIMH (P50 MH078028, P50 MH096972); NIH/NCI (U24 CA159988); NIH/NIGMS (R01 GM088822).
