## Supplementary material for "PGRMC1 phosphorylation and cell plasticity 1: glycolysis, mitochondria, tumor growth": File S8: final_sig_file_1439167306.html

Anchored HTML File of EIDs

|  |  |
| --- | --- |
|  | WEB-based GEne SeT AnaLysis Toolkit |
|  | ***Translating gene lists into biological insights...*** |

---

  

| **Database:biological process      &nbspName:'de novo' posttranslational protein folding      &nbspID:GO:0051084** | | | | | | |
| --- | --- | --- | --- | --- | --- | --- |
| C=48; O=12; E=0.21; R=58.15; rawP=7.97e-19; adjP=2.55e-16 | | | | | | |
| Index | UserID | Value | Gene Symbol | Gene Name | EntrezGene | Ensembl |
| 1 | P78371 | NA | CCT2 | chaperonin containing TCP1, subunit 2 (beta) | 10576 | ENSG00000166226 |
| 2 | Q92598 | NA | HSPH1 | heat shock 105kDa/110kDa protein 1 | 10808 | ENSG00000120694 |
| 3 | P17987 | NA | TCP1 | t-complex 1 | 6950 | ENSG00000120438 |
| 4 | Q13885 | NA | TUBB2A | tubulin, beta 2A class IIa | 7280 | ENSG00000137267 |
| 5 | P68371 | NA | TUBB4B | tubulin, beta 4B class IVb | 10383 | ENSG00000188229 |
| 6 | P40227 | NA | CCT6A | chaperonin containing TCP1, subunit 6A (zeta 1) | 908 | ENSG00000146731 |
| 7 | P49368 | NA | CCT3 | chaperonin containing TCP1, subunit 3 (gamma) | 7203 | ENSG00000163468 |
| 8 | Q9BQE3 | NA | TUBA1C | tubulin, alpha 1c | 84790 | ENSG00000167553 |
| 9 | P25685 | NA | DNAJB1 | DnaJ (Hsp40) homolog, subfamily B, member 1 | 3337 | ENSG00000132002 |
| 10 | Q99832 | NA | CCT7 | chaperonin containing TCP1, subunit 7 (eta) | 10574 | ENSG00000135624 |
| 11 | P50990 | NA | CCT8 | chaperonin containing TCP1, subunit 8 (theta) | 10694 | ENSG00000156261 |
| 12 | P50991 | NA | CCT4 | chaperonin containing TCP1, subunit 4 (delta) | 10575 | ENSG00000115484 |

  
  

| **Database:biological process      &nbspName:'de novo' protein folding      &nbspID:GO:0006458** | | | | | | |
| --- | --- | --- | --- | --- | --- | --- |
| C=53; O=12; E=0.23; R=52.67; rawP=3.00e-18; adjP=4.80e-16 | | | | | | |
| Index | UserID | Value | Gene Symbol | Gene Name | EntrezGene | Ensembl |
| 1 | P78371 | NA | CCT2 | chaperonin containing TCP1, subunit 2 (beta) | 10576 | ENSG00000166226 |
| 2 | Q92598 | NA | HSPH1 | heat shock 105kDa/110kDa protein 1 | 10808 | ENSG00000120694 |
| 3 | P17987 | NA | TCP1 | t-complex 1 | 6950 | ENSG00000120438 |
| 4 | Q13885 | NA | TUBB2A | tubulin, beta 2A class IIa | 7280 | ENSG00000137267 |
| 5 | P68371 | NA | TUBB4B | tubulin, beta 4B class IVb | 10383 | ENSG00000188229 |
| 6 | P40227 | NA | CCT6A | chaperonin containing TCP1, subunit 6A (zeta 1) | 908 | ENSG00000146731 |
| 7 | P49368 | NA | CCT3 | chaperonin containing TCP1, subunit 3 (gamma) | 7203 | ENSG00000163468 |
| 8 | Q9BQE3 | NA | TUBA1C | tubulin, alpha 1c | 84790 | ENSG00000167553 |
| 9 | P25685 | NA | DNAJB1 | DnaJ (Hsp40) homolog, subfamily B, member 1 | 3337 | ENSG00000132002 |
| 10 | Q99832 | NA | CCT7 | chaperonin containing TCP1, subunit 7 (eta) | 10574 | ENSG00000135624 |
| 11 | P50990 | NA | CCT8 | chaperonin containing TCP1, subunit 8 (theta) | 10694 | ENSG00000156261 |
| 12 | P50991 | NA | CCT4 | chaperonin containing TCP1, subunit 4 (delta) | 10575 | ENSG00000115484 |

  
  

| **Database:biological process      &nbspName:protein folding      &nbspID:GO:0006457** | | | | | | |
| --- | --- | --- | --- | --- | --- | --- |
| C=211; O=14; E=0.91; R=15.43; rawP=2.09e-13; adjP=2.23e-11 | | | | | | |
| Index | UserID | Value | Gene Symbol | Gene Name | EntrezGene | Ensembl |
| 1 | P78371 | NA | CCT2 | chaperonin containing TCP1, subunit 2 (beta) | 10576 | ENSG00000166226 |
| 2 | Q92598 | NA | HSPH1 | heat shock 105kDa/110kDa protein 1 | 10808 | ENSG00000120694 |
| 3 | P17987 | NA | TCP1 | t-complex 1 | 6950 | ENSG00000120438 |
| 4 | Q13885 | NA | TUBB2A | tubulin, beta 2A class IIa | 7280 | ENSG00000137267 |
| 5 | P68371 | NA | TUBB4B | tubulin, beta 4B class IVb | 10383 | ENSG00000188229 |
| 6 | O95433 | NA | AHSA1 | AHA1, activator of heat shock 90kDa protein ATPase homolog 1 (yeast) | 10598 | ENSG00000100591 |
| 7 | P40227 | NA | CCT6A | chaperonin containing TCP1, subunit 6A (zeta 1) | 908 | ENSG00000146731 |
| 8 | P49368 | NA | CCT3 | chaperonin containing TCP1, subunit 3 (gamma) | 7203 | ENSG00000163468 |
| 9 | Q9BQE3 | NA | TUBA1C | tubulin, alpha 1c | 84790 | ENSG00000167553 |
| 10 | P25685 | NA | DNAJB1 | DnaJ (Hsp40) homolog, subfamily B, member 1 | 3337 | ENSG00000132002 |
| 11 | Q15185 | NA | PTGES3 | prostaglandin E synthase 3 (cytosolic) | 10728 | ENSG00000110958 |
| 12 | Q99832 | NA | CCT7 | chaperonin containing TCP1, subunit 7 (eta) | 10574 | ENSG00000135624 |
| 13 | P50990 | NA | CCT8 | chaperonin containing TCP1, subunit 8 (theta) | 10694 | ENSG00000156261 |
| 14 | P50991 | NA | CCT4 | chaperonin containing TCP1, subunit 4 (delta) | 10575 | ENSG00000115484 |

  
  

| **Database:biological process      &nbspName:cellular macromolecular complex subunit organization      &nbspID:GO:0034621** | | | | | | |
| --- | --- | --- | --- | --- | --- | --- |
| C=681; O=12; E=2.93; R=4.10; rawP=2.74e-05; adjP=0.0022 | | | | | | |
| Index | UserID | Value | Gene Symbol | Gene Name | EntrezGene | Ensembl |
| 1 | P63241 | NA | EIF5A | eukaryotic translation initiation factor 5A | 1984 | ENSG00000132507 |
| 2 | P78371 | NA | CCT2 | chaperonin containing TCP1, subunit 2 (beta) | 10576 | ENSG00000166226 |
| 3 | P17987 | NA | TCP1 | t-complex 1 | 6950 | ENSG00000120438 |
| 4 | P55209 | NA | NAP1L1 | nucleosome assembly protein 1-like 1 | 4673 | ENSG00000187109 |
| 5 | P34932 | NA | HSPA4 | heat shock 70kDa protein 4 | 3308 | ENSG00000170606 |
| 6 | Q13885 | NA | TUBB2A | tubulin, beta 2A class IIa | 7280 | ENSG00000137267 |
| 7 | P49321 | NA | NASP | nuclear autoantigenic sperm protein (histone-binding) | 4678 | ENSG00000132780 |
| 8 | P68371 | NA | TUBB4B | tubulin, beta 4B class IVb | 10383 | ENSG00000188229 |
| 9 | Q9BQE3 | NA | TUBA1C | tubulin, alpha 1c | 84790 | ENSG00000167553 |
| 10 | P49736 | NA | MCM2 | minichromosome maintenance complex component 2 | 4171 | ENSG00000073111 |
| 11 | P62906 | NA | RPL10A | ribosomal protein L10a | 4736 | ENSG00000198755 |
| 12 | O60610 | NA | DIAPH1 | diaphanous homolog 1 (Drosophila) | 1729 | ENSG00000131504 |

  
  

| **Database:biological process      &nbspName:chaperone-mediated protein folding      &nbspID:GO:0061077** | | | | | | |
| --- | --- | --- | --- | --- | --- | --- |
| C=21; O=3; E=0.09; R=33.23; rawP=9.52e-05; adjP=0.0061 | | | | | | |
| Index | UserID | Value | Gene Symbol | Gene Name | EntrezGene | Ensembl |
| 1 | Q92598 | NA | HSPH1 | heat shock 105kDa/110kDa protein 1 | 10808 | ENSG00000120694 |
| 2 | Q15185 | NA | PTGES3 | prostaglandin E synthase 3 (cytosolic) | 10728 | ENSG00000110958 |
| 3 | P25685 | NA | DNAJB1 | DnaJ (Hsp40) homolog, subfamily B, member 1 | 3337 | ENSG00000132002 |

  
  

| **Database:biological process      &nbspName:nucleobase metabolic process      &nbspID:GO:0009112** | | | | | | |
| --- | --- | --- | --- | --- | --- | --- |
| C=66; O=4; E=0.28; R=14.10; rawP=0.0002; adjP=0.0091 | | | | | | |
| Index | UserID | Value | Gene Symbol | Gene Name | EntrezGene | Ensembl |
| 1 | P12268 | NA | IMPDH2 | IMP (inosine 5'-monophosphate) dehydrogenase 2 | 3615 | ENSG00000178035 |
| 2 | P22102 | NA | GART | phosphoribosylglycinamide formyltransferase, phosphoribosylglycinamide synthetase, phosphoribosylaminoimidazole synthetase | 2618 | ENSG00000159131 |
| 3 | P00491 | NA | PNP | purine nucleoside phosphorylase | 4860 | ENSG00000198805 |
| 4 | Q16555 | NA | DPYSL2 | dihydropyrimidinase-like 2 | 1808 | ENSG00000092964 |

  
  

| **Database:biological process      &nbspName:cellular protein metabolic process      &nbspID:GO:0044267** | | | | | | |
| --- | --- | --- | --- | --- | --- | --- |
| C=3239; O=27; E=13.92; R=1.94; rawP=0.0002; adjP=0.0091 | | | | | | |
| Index | UserID | Value | Gene Symbol | Gene Name | EntrezGene | Ensembl |
| 1 | P63241 | NA | EIF5A | eukaryotic translation initiation factor 5A | 1984 | ENSG00000132507 |
| 2 | P78371 | NA | CCT2 | chaperonin containing TCP1, subunit 2 (beta) | 10576 | ENSG00000166226 |
| 3 | Q92598 | NA | HSPH1 | heat shock 105kDa/110kDa protein 1 | 10808 | ENSG00000120694 |
| 4 | Q13885 | NA | TUBB2A | tubulin, beta 2A class IIa | 7280 | ENSG00000137267 |
| 5 | P12081 | NA | HARS | histidyl-tRNA synthetase | 3035 | ENSG00000170445 |
| 6 | P68371 | NA | TUBB4B | tubulin, beta 4B class IVb | 10383 | ENSG00000188229 |
| 7 | P49368 | NA | CCT3 | chaperonin containing TCP1, subunit 3 (gamma) | 7203 | ENSG00000163468 |
| 8 | P56192 | NA | MARS | methionyl-tRNA synthetase | 4141 | ENSG00000166986 |
| 9 | Q9BQE3 | NA | TUBA1C | tubulin, alpha 1c | 84790 | ENSG00000167553 |
| 10 | P25685 | NA | DNAJB1 | DnaJ (Hsp40) homolog, subfamily B, member 1 | 3337 | ENSG00000132002 |
| 11 | P62906 | NA | RPL10A | ribosomal protein L10a | 4736 | ENSG00000198755 |
| 12 | Q15056 | NA | EIF4H | eukaryotic translation initiation factor 4H | 7458 | ENSG00000106682 |
| 13 | P50990 | NA | CCT8 | chaperonin containing TCP1, subunit 8 (theta) | 10694 | ENSG00000156261 |
| 14 | P68104 | NA | EEF1A1 | eukaryotic translation elongation factor 1 alpha 1 | 1915 | ENSG00000156508 |
| 15 | P50991 | NA | CCT4 | chaperonin containing TCP1, subunit 4 (delta) | 10575 | ENSG00000115484 |
| 16 | O00425 | NA | IGF2BP3 | insulin-like growth factor 2 mRNA binding protein 3 | 10643 | ENSG00000136231 |
| 17 | P07814 | NA | EPRS | glutamyl-prolyl-tRNA synthetase | 2058 | ENSG00000136628 |
| 18 | P17987 | NA | TCP1 | t-complex 1 | 6950 | ENSG00000120438 |
| 19 | P13639 | NA | EEF2 | eukaryotic translation elongation factor 2 | 1938 | ENSG00000167658 |
| 20 | P41091 | NA | EIF2S3 | eukaryotic translation initiation factor 2, subunit 3 gamma, 52kDa | 1968 | ENSG00000130741 |
| 21 | O95433 | NA | AHSA1 | AHA1, activator of heat shock 90kDa protein ATPase homolog 1 (yeast) | 10598 | ENSG00000100591 |
| 22 | P41567 | NA | EIF1 | eukaryotic translation initiation factor 1 | 10209 | ENSG00000173812 |
| 23 | P40227 | NA | CCT6A | chaperonin containing TCP1, subunit 6A (zeta 1) | 908 | ENSG00000146731 |
| 24 | Q15185 | NA | PTGES3 | prostaglandin E synthase 3 (cytosolic) | 10728 | ENSG00000110958 |
| 25 | Q99832 | NA | CCT7 | chaperonin containing TCP1, subunit 7 (eta) | 10574 | ENSG00000135624 |
| 26 | P21980 | NA | TGM2 | transglutaminase 2 (C polypeptide, protein-glutamine-gamma-glutamyltransferase) | 7052 | ENSG00000198959 |
| 27 | Q14166 | NA | TTLL12 | tubulin tyrosine ligase-like family, member 12 | 23170 | ENSG00000100304 |

  
  

| **Database:biological process      &nbspName:macromolecular complex subunit organization      &nbspID:GO:0043933** | | | | | | |
| --- | --- | --- | --- | --- | --- | --- |
| C=1152; O=14; E=4.95; R=2.83; rawP=0.0003; adjP=0.0107 | | | | | | |
| Index | UserID | Value | Gene Symbol | Gene Name | EntrezGene | Ensembl |
| 1 | P63241 | NA | EIF5A | eukaryotic translation initiation factor 5A | 1984 | ENSG00000132507 |
| 2 | P07814 | NA | EPRS | glutamyl-prolyl-tRNA synthetase | 2058 | ENSG00000136628 |
| 3 | P78371 | NA | CCT2 | chaperonin containing TCP1, subunit 2 (beta) | 10576 | ENSG00000166226 |
| 4 | P17987 | NA | TCP1 | t-complex 1 | 6950 | ENSG00000120438 |
| 5 | P55209 | NA | NAP1L1 | nucleosome assembly protein 1-like 1 | 4673 | ENSG00000187109 |
| 6 | P34932 | NA | HSPA4 | heat shock 70kDa protein 4 | 3308 | ENSG00000170606 |
| 7 | Q13885 | NA | TUBB2A | tubulin, beta 2A class IIa | 7280 | ENSG00000137267 |
| 8 | P49321 | NA | NASP | nuclear autoantigenic sperm protein (histone-binding) | 4678 | ENSG00000132780 |
| 9 | P68371 | NA | TUBB4B | tubulin, beta 4B class IVb | 10383 | ENSG00000188229 |
| 10 | Q9BQE3 | NA | TUBA1C | tubulin, alpha 1c | 84790 | ENSG00000167553 |
| 11 | P21980 | NA | TGM2 | transglutaminase 2 (C polypeptide, protein-glutamine-gamma-glutamyltransferase) | 7052 | ENSG00000198959 |
| 12 | P62906 | NA | RPL10A | ribosomal protein L10a | 4736 | ENSG00000198755 |
| 13 | P49736 | NA | MCM2 | minichromosome maintenance complex component 2 | 4171 | ENSG00000073111 |
| 14 | O60610 | NA | DIAPH1 | diaphanous homolog 1 (Drosophila) | 1729 | ENSG00000131504 |

  
  

| **Database:biological process      &nbspName:cellular protein complex assembly      &nbspID:GO:0043623** | | | | | | |
| --- | --- | --- | --- | --- | --- | --- |
| C=307; O=7; E=1.32; R=5.30; rawP=0.0003; adjP=0.0107 | | | | | | |
| Index | UserID | Value | Gene Symbol | Gene Name | EntrezGene | Ensembl |
| 1 | Q9BQE3 | NA | TUBA1C | tubulin, alpha 1c | 84790 | ENSG00000167553 |
| 2 | P78371 | NA | CCT2 | chaperonin containing TCP1, subunit 2 (beta) | 10576 | ENSG00000166226 |
| 3 | P17987 | NA | TCP1 | t-complex 1 | 6950 | ENSG00000120438 |
| 4 | P34932 | NA | HSPA4 | heat shock 70kDa protein 4 | 3308 | ENSG00000170606 |
| 5 | Q13885 | NA | TUBB2A | tubulin, beta 2A class IIa | 7280 | ENSG00000137267 |
| 6 | P68371 | NA | TUBB4B | tubulin, beta 4B class IVb | 10383 | ENSG00000188229 |
| 7 | O60610 | NA | DIAPH1 | diaphanous homolog 1 (Drosophila) | 1729 | ENSG00000131504 |

  
  

| **Database:biological process      &nbspName:cellular macromolecular complex assembly      &nbspID:GO:0034622** | | | | | | |
| --- | --- | --- | --- | --- | --- | --- |
| C=526; O=9; E=2.26; R=3.98; rawP=0.0004; adjP=0.0128 | | | | | | |
| Index | UserID | Value | Gene Symbol | Gene Name | EntrezGene | Ensembl |
| 1 | P78371 | NA | CCT2 | chaperonin containing TCP1, subunit 2 (beta) | 10576 | ENSG00000166226 |
| 2 | P17987 | NA | TCP1 | t-complex 1 | 6950 | ENSG00000120438 |
| 3 | P55209 | NA | NAP1L1 | nucleosome assembly protein 1-like 1 | 4673 | ENSG00000187109 |
| 4 | P34932 | NA | HSPA4 | heat shock 70kDa protein 4 | 3308 | ENSG00000170606 |
| 5 | Q13885 | NA | TUBB2A | tubulin, beta 2A class IIa | 7280 | ENSG00000137267 |
| 6 | P68371 | NA | TUBB4B | tubulin, beta 4B class IVb | 10383 | ENSG00000188229 |
| 7 | Q9BQE3 | NA | TUBA1C | tubulin, alpha 1c | 84790 | ENSG00000167553 |
| 8 | P49736 | NA | MCM2 | minichromosome maintenance complex component 2 | 4171 | ENSG00000073111 |
| 9 | O60610 | NA | DIAPH1 | diaphanous homolog 1 (Drosophila) | 1729 | ENSG00000131504 |

  
  

| **Database:biological process      &nbspName:protein metabolic process      &nbspID:GO:0019538** | | | | | | |
| --- | --- | --- | --- | --- | --- | --- |
| C=3844; O=29; E=16.53; R=1.75; rawP=0.0005; adjP=0.0145 | | | | | | |
| Index | UserID | Value | Gene Symbol | Gene Name | EntrezGene | Ensembl |
| 1 | P63241 | NA | EIF5A | eukaryotic translation initiation factor 5A | 1984 | ENSG00000132507 |
| 2 | P78371 | NA | CCT2 | chaperonin containing TCP1, subunit 2 (beta) | 10576 | ENSG00000166226 |
| 3 | Q92598 | NA | HSPH1 | heat shock 105kDa/110kDa protein 1 | 10808 | ENSG00000120694 |
| 4 | Q7Z2K6 | NA | ERMP1 | endoplasmic reticulum metallopeptidase 1 | 79956 | ENSG00000099219 |
| 5 | Q13885 | NA | TUBB2A | tubulin, beta 2A class IIa | 7280 | ENSG00000137267 |
| 6 | P12081 | NA | HARS | histidyl-tRNA synthetase | 3035 | ENSG00000170445 |
| 7 | P68371 | NA | TUBB4B | tubulin, beta 4B class IVb | 10383 | ENSG00000188229 |
| 8 | P49368 | NA | CCT3 | chaperonin containing TCP1, subunit 3 (gamma) | 7203 | ENSG00000163468 |
| 9 | P56192 | NA | MARS | methionyl-tRNA synthetase | 4141 | ENSG00000166986 |
| 10 | Q9BQE3 | NA | TUBA1C | tubulin, alpha 1c | 84790 | ENSG00000167553 |
| 11 | P25685 | NA | DNAJB1 | DnaJ (Hsp40) homolog, subfamily B, member 1 | 3337 | ENSG00000132002 |
| 12 | P62906 | NA | RPL10A | ribosomal protein L10a | 4736 | ENSG00000198755 |
| 13 | Q15056 | NA | EIF4H | eukaryotic translation initiation factor 4H | 7458 | ENSG00000106682 |
| 14 | P50990 | NA | CCT8 | chaperonin containing TCP1, subunit 8 (theta) | 10694 | ENSG00000156261 |
| 15 | P68104 | NA | EEF1A1 | eukaryotic translation elongation factor 1 alpha 1 | 1915 | ENSG00000156508 |
| 16 | P02786 | NA | TFRC | transferrin receptor (p90, CD71) | 7037 | ENSG00000072274 |
| 17 | P50991 | NA | CCT4 | chaperonin containing TCP1, subunit 4 (delta) | 10575 | ENSG00000115484 |
| 18 | O00425 | NA | IGF2BP3 | insulin-like growth factor 2 mRNA binding protein 3 | 10643 | ENSG00000136231 |
| 19 | P07814 | NA | EPRS | glutamyl-prolyl-tRNA synthetase | 2058 | ENSG00000136628 |
| 20 | P17987 | NA | TCP1 | t-complex 1 | 6950 | ENSG00000120438 |
| 21 | P13639 | NA | EEF2 | eukaryotic translation elongation factor 2 | 1938 | ENSG00000167658 |
| 22 | P41091 | NA | EIF2S3 | eukaryotic translation initiation factor 2, subunit 3 gamma, 52kDa | 1968 | ENSG00000130741 |
| 23 | O95433 | NA | AHSA1 | AHA1, activator of heat shock 90kDa protein ATPase homolog 1 (yeast) | 10598 | ENSG00000100591 |
| 24 | P41567 | NA | EIF1 | eukaryotic translation initiation factor 1 | 10209 | ENSG00000173812 |
| 25 | P40227 | NA | CCT6A | chaperonin containing TCP1, subunit 6A (zeta 1) | 908 | ENSG00000146731 |
| 26 | Q15185 | NA | PTGES3 | prostaglandin E synthase 3 (cytosolic) | 10728 | ENSG00000110958 |
| 27 | Q99832 | NA | CCT7 | chaperonin containing TCP1, subunit 7 (eta) | 10574 | ENSG00000135624 |
| 28 | P21980 | NA | TGM2 | transglutaminase 2 (C polypeptide, protein-glutamine-gamma-glutamyltransferase) | 7052 | ENSG00000198959 |
| 29 | Q14166 | NA | TTLL12 | tubulin tyrosine ligase-like family, member 12 | 23170 | ENSG00000100304 |

  
  

| **Database:biological process      &nbspName:translation      &nbspID:GO:0006412** | | | | | | |
| --- | --- | --- | --- | --- | --- | --- |
| C=832; O=11; E=3.58; R=3.08; rawP=0.0007; adjP=0.0172 | | | | | | |
| Index | UserID | Value | Gene Symbol | Gene Name | EntrezGene | Ensembl |
| 1 | P63241 | NA | EIF5A | eukaryotic translation initiation factor 5A | 1984 | ENSG00000132507 |
| 2 | P07814 | NA | EPRS | glutamyl-prolyl-tRNA synthetase | 2058 | ENSG00000136628 |
| 3 | P13639 | NA | EEF2 | eukaryotic translation elongation factor 2 | 1938 | ENSG00000167658 |
| 4 | P41091 | NA | EIF2S3 | eukaryotic translation initiation factor 2, subunit 3 gamma, 52kDa | 1968 | ENSG00000130741 |
| 5 | P12081 | NA | HARS | histidyl-tRNA synthetase | 3035 | ENSG00000170445 |
| 6 | P41567 | NA | EIF1 | eukaryotic translation initiation factor 1 | 10209 | ENSG00000173812 |
| 7 | P56192 | NA | MARS | methionyl-tRNA synthetase | 4141 | ENSG00000166986 |
| 8 | P62906 | NA | RPL10A | ribosomal protein L10a | 4736 | ENSG00000198755 |
| 9 | Q15056 | NA | EIF4H | eukaryotic translation initiation factor 4H | 7458 | ENSG00000106682 |
| 10 | P68104 | NA | EEF1A1 | eukaryotic translation elongation factor 1 alpha 1 | 1915 | ENSG00000156508 |
| 11 | O00425 | NA | IGF2BP3 | insulin-like growth factor 2 mRNA binding protein 3 | 10643 | ENSG00000136231 |

  
  

| **Database:biological process      &nbspName:purine nucleobase metabolic process      &nbspID:GO:0006144** | | | | | | |
| --- | --- | --- | --- | --- | --- | --- |
| C=40; O=3; E=0.17; R=17.45; rawP=0.0007; adjP=0.0172 | | | | | | |
| Index | UserID | Value | Gene Symbol | Gene Name | EntrezGene | Ensembl |
| 1 | P12268 | NA | IMPDH2 | IMP (inosine 5'-monophosphate) dehydrogenase 2 | 3615 | ENSG00000178035 |
| 2 | P22102 | NA | GART | phosphoribosylglycinamide formyltransferase, phosphoribosylglycinamide synthetase, phosphoribosylaminoimidazole synthetase | 2618 | ENSG00000159131 |
| 3 | P00491 | NA | PNP | purine nucleoside phosphorylase | 4860 | ENSG00000198805 |

  
  

| **Database:biological process      &nbspName:tRNA aminoacylation      &nbspID:GO:0043039** | | | | | | |
| --- | --- | --- | --- | --- | --- | --- |
| C=51; O=3; E=0.22; R=13.68; rawP=0.0014; adjP=0.0249 | | | | | | |
| Index | UserID | Value | Gene Symbol | Gene Name | EntrezGene | Ensembl |
| 1 | P07814 | NA | EPRS | glutamyl-prolyl-tRNA synthetase | 2058 | ENSG00000136628 |
| 2 | P56192 | NA | MARS | methionyl-tRNA synthetase | 4141 | ENSG00000166986 |
| 3 | P12081 | NA | HARS | histidyl-tRNA synthetase | 3035 | ENSG00000170445 |

  
  

| **Database:biological process      &nbspName:tRNA aminoacylation for protein translation      &nbspID:GO:0006418** | | | | | | |
| --- | --- | --- | --- | --- | --- | --- |
| C=48; O=3; E=0.21; R=14.54; rawP=0.0011; adjP=0.0249 | | | | | | |
| Index | UserID | Value | Gene Symbol | Gene Name | EntrezGene | Ensembl |
| 1 | P07814 | NA | EPRS | glutamyl-prolyl-tRNA synthetase | 2058 | ENSG00000136628 |
| 2 | P56192 | NA | MARS | methionyl-tRNA synthetase | 4141 | ENSG00000166986 |
| 3 | P12081 | NA | HARS | histidyl-tRNA synthetase | 3035 | ENSG00000170445 |

  
  

| **Database:biological process      &nbspName:protein complex subunit organization      &nbspID:GO:0071822** | | | | | | |
| --- | --- | --- | --- | --- | --- | --- |
| C=890; O=11; E=3.83; R=2.87; rawP=0.0013; adjP=0.0249 | | | | | | |
| Index | UserID | Value | Gene Symbol | Gene Name | EntrezGene | Ensembl |
| 1 | P63241 | NA | EIF5A | eukaryotic translation initiation factor 5A | 1984 | ENSG00000132507 |
| 2 | P07814 | NA | EPRS | glutamyl-prolyl-tRNA synthetase | 2058 | ENSG00000136628 |
| 3 | P78371 | NA | CCT2 | chaperonin containing TCP1, subunit 2 (beta) | 10576 | ENSG00000166226 |
| 4 | P17987 | NA | TCP1 | t-complex 1 | 6950 | ENSG00000120438 |
| 5 | P34932 | NA | HSPA4 | heat shock 70kDa protein 4 | 3308 | ENSG00000170606 |
| 6 | Q13885 | NA | TUBB2A | tubulin, beta 2A class IIa | 7280 | ENSG00000137267 |
| 7 | P68371 | NA | TUBB4B | tubulin, beta 4B class IVb | 10383 | ENSG00000188229 |
| 8 | Q9BQE3 | NA | TUBA1C | tubulin, alpha 1c | 84790 | ENSG00000167553 |
| 9 | P21980 | NA | TGM2 | transglutaminase 2 (C polypeptide, protein-glutamine-gamma-glutamyltransferase) | 7052 | ENSG00000198959 |
| 10 | P62906 | NA | RPL10A | ribosomal protein L10a | 4736 | ENSG00000198755 |
| 11 | O60610 | NA | DIAPH1 | diaphanous homolog 1 (Drosophila) | 1729 | ENSG00000131504 |

  
  

| **Database:biological process      &nbspName:translational elongation      &nbspID:GO:0006414** | | | | | | |
| --- | --- | --- | --- | --- | --- | --- |
| C=108; O=4; E=0.46; R=8.62; rawP=0.0012; adjP=0.0249 | | | | | | |
| Index | UserID | Value | Gene Symbol | Gene Name | EntrezGene | Ensembl |
| 1 | P63241 | NA | EIF5A | eukaryotic translation initiation factor 5A | 1984 | ENSG00000132507 |
| 2 | P62906 | NA | RPL10A | ribosomal protein L10a | 4736 | ENSG00000198755 |
| 3 | P13639 | NA | EEF2 | eukaryotic translation elongation factor 2 | 1938 | ENSG00000167658 |
| 4 | P68104 | NA | EEF1A1 | eukaryotic translation elongation factor 1 alpha 1 | 1915 | ENSG00000156508 |

  
  

| **Database:biological process      &nbspName:amino acid activation      &nbspID:GO:0043038** | | | | | | |
| --- | --- | --- | --- | --- | --- | --- |
| C=51; O=3; E=0.22; R=13.68; rawP=0.0014; adjP=0.0249 | | | | | | |
| Index | UserID | Value | Gene Symbol | Gene Name | EntrezGene | Ensembl |
| 1 | P07814 | NA | EPRS | glutamyl-prolyl-tRNA synthetase | 2058 | ENSG00000136628 |
| 2 | P56192 | NA | MARS | methionyl-tRNA synthetase | 4141 | ENSG00000166986 |
| 3 | P12081 | NA | HARS | histidyl-tRNA synthetase | 3035 | ENSG00000170445 |

  
  

| **Database:biological process      &nbspName:response to inorganic substance      &nbspID:GO:0010035** | | | | | | |
| --- | --- | --- | --- | --- | --- | --- |
| C=423; O=7; E=1.82; R=3.85; rawP=0.0022; adjP=0.0371 | | | | | | |
| Index | UserID | Value | Gene Symbol | Gene Name | EntrezGene | Ensembl |
| 1 | Q16881 | NA | TXNRD1 | thioredoxin reductase 1 | 7296 | ENSG00000198431 |
| 2 | P02795 | NA | MT2A | metallothionein 2A | 4502 | ENSG00000125148 |
| 3 | P22102 | NA | GART | phosphoribosylglycinamide formyltransferase, phosphoribosylglycinamide synthetase, phosphoribosylaminoimidazole synthetase | 2618 | ENSG00000159131 |
| 4 | O60610 | NA | DIAPH1 | diaphanous homolog 1 (Drosophila) | 1729 | ENSG00000131504 |
| 5 | Q16555 | NA | DPYSL2 | dihydropyrimidinase-like 2 | 1808 | ENSG00000092964 |
| 6 | Q06830 | NA | PRDX1 | peroxiredoxin 1 | 5052 | ENSG00000117450 |
| 7 | P02786 | NA | TFRC | transferrin receptor (p90, CD71) | 7037 | ENSG00000072274 |

  
  

| **Database:biological process      &nbspName:primary metabolic process      &nbspID:GO:0044238** | | | | | | |
| --- | --- | --- | --- | --- | --- | --- |
| C=8601; O=48; E=36.98; R=1.30; rawP=0.0024; adjP=0.0384 | | | | | | |
| Index | UserID | Value | Gene Symbol | Gene Name | EntrezGene | Ensembl |
| 1 | Q92598 | NA | HSPH1 | heat shock 105kDa/110kDa protein 1 | 10808 | ENSG00000120694 |
| 2 | Q13885 | NA | TUBB2A | tubulin, beta 2A class IIa | 7280 | ENSG00000137267 |
| 3 | P68371 | NA | TUBB4B | tubulin, beta 4B class IVb | 10383 | ENSG00000188229 |
| 4 | P25685 | NA | DNAJB1 | DnaJ (Hsp40) homolog, subfamily B, member 1 | 3337 | ENSG00000132002 |
| 5 | P68104 | NA | EEF1A1 | eukaryotic translation elongation factor 1 alpha 1 | 1915 | ENSG00000156508 |
| 6 | P02786 | NA | TFRC | transferrin receptor (p90, CD71) | 7037 | ENSG00000072274 |
| 7 | P50991 | NA | CCT4 | chaperonin containing TCP1, subunit 4 (delta) | 10575 | ENSG00000115484 |
| 8 | O00425 | NA | IGF2BP3 | insulin-like growth factor 2 mRNA binding protein 3 | 10643 | ENSG00000136231 |
| 9 | P22102 | NA | GART | phosphoribosylglycinamide formyltransferase, phosphoribosylglycinamide synthetase, phosphoribosylaminoimidazole synthetase | 2618 | ENSG00000159131 |
| 10 | P13639 | NA | EEF2 | eukaryotic translation elongation factor 2 | 1938 | ENSG00000167658 |
| 11 | Q16555 | NA | DPYSL2 | dihydropyrimidinase-like 2 | 1808 | ENSG00000092964 |
| 12 | O95433 | NA | AHSA1 | AHA1, activator of heat shock 90kDa protein ATPase homolog 1 (yeast) | 10598 | ENSG00000100591 |
| 13 | P41567 | NA | EIF1 | eukaryotic translation initiation factor 1 | 10209 | ENSG00000173812 |
| 14 | P17174 | NA | GOT1 | glutamic-oxaloacetic transaminase 1, soluble (aspartate aminotransferase 1) | 2805 | ENSG00000120053 |
| 15 | P67809 | NA | YBX1 | Y box binding protein 1 | 4904 | ENSG00000065978 |
| 16 | P21980 | NA | TGM2 | transglutaminase 2 (C polypeptide, protein-glutamine-gamma-glutamyltransferase) | 7052 | ENSG00000198959 |
| 17 | Q99832 | NA | CCT7 | chaperonin containing TCP1, subunit 7 (eta) | 10574 | ENSG00000135624 |
| 18 | Q15185 | NA | PTGES3 | prostaglandin E synthase 3 (cytosolic) | 10728 | ENSG00000110958 |
| 19 | Q6PIU2 | NA | NCEH1 | neutral cholesterol ester hydrolase 1 | 57552 | ENSG00000144959 |
| 20 | Q14166 | NA | TTLL12 | tubulin tyrosine ligase-like family, member 12 | 23170 | ENSG00000100304 |
| 21 | P11387 | NA | TOP1 | topoisomerase (DNA) I | 7150 | ENSG00000198900 |
| 22 | P11586 | NA | MTHFD1 | methylenetetrahydrofolate dehydrogenase (NADP+ dependent) 1, methenyltetrahydrofolate cyclohydrolase, formyltetrahydrofolate synthetase | 4522 | ENSG00000100714 |
| 23 | Q16881 | NA | TXNRD1 | thioredoxin reductase 1 | 7296 | ENSG00000198431 |
| 24 | P63241 | NA | EIF5A | eukaryotic translation initiation factor 5A | 1984 | ENSG00000132507 |
| 25 | P78371 | NA | CCT2 | chaperonin containing TCP1, subunit 2 (beta) | 10576 | ENSG00000166226 |
| 26 | P08195 | NA | SLC3A2 | solute carrier family 3 (activators of dibasic and neutral amino acid transport), member 2 | 6520 | ENSG00000168003 |
| 27 | Q7Z2K6 | NA | ERMP1 | endoplasmic reticulum metallopeptidase 1 | 79956 | ENSG00000099219 |
| 28 | P07951 | NA | TPM2 | tropomyosin 2 (beta) | 7169 | ENSG00000198467 |
| 29 | P00491 | NA | PNP | purine nucleoside phosphorylase | 4860 | ENSG00000198805 |
| 30 | P49321 | NA | NASP | nuclear autoantigenic sperm protein (histone-binding) | 4678 | ENSG00000132780 |
| 31 | P12081 | NA | HARS | histidyl-tRNA synthetase | 3035 | ENSG00000170445 |
| 32 | P49327 | NA | FASN | fatty acid synthase | 2194 | ENSG00000169710 |
| 33 | P49368 | NA | CCT3 | chaperonin containing TCP1, subunit 3 (gamma) | 7203 | ENSG00000163468 |
| 34 | Q9BQE3 | NA | TUBA1C | tubulin, alpha 1c | 84790 | ENSG00000167553 |
| 35 | P56192 | NA | MARS | methionyl-tRNA synthetase | 4141 | ENSG00000166986 |
| 36 | P12268 | NA | IMPDH2 | IMP (inosine 5'-monophosphate) dehydrogenase 2 | 3615 | ENSG00000178035 |
| 37 | P62906 | NA | RPL10A | ribosomal protein L10a | 4736 | ENSG00000198755 |
| 38 | P49736 | NA | MCM2 | minichromosome maintenance complex component 2 | 4171 | ENSG00000073111 |
| 39 | Q15056 | NA | EIF4H | eukaryotic translation initiation factor 4H | 7458 | ENSG00000106682 |
| 40 | P50990 | NA | CCT8 | chaperonin containing TCP1, subunit 8 (theta) | 10694 | ENSG00000156261 |
| 41 | P07814 | NA | EPRS | glutamyl-prolyl-tRNA synthetase | 2058 | ENSG00000136628 |
| 42 | P17987 | NA | TCP1 | t-complex 1 | 6950 | ENSG00000120438 |
| 43 | P55209 | NA | NAP1L1 | nucleosome assembly protein 1-like 1 | 4673 | ENSG00000187109 |
| 44 | P41091 | NA | EIF2S3 | eukaryotic translation initiation factor 2, subunit 3 gamma, 52kDa | 1968 | ENSG00000130741 |
| 45 | P07108 | NA | DBI | diazepam binding inhibitor (GABA receptor modulator, acyl-CoA binding protein) | 1622 | ENSG00000155368 |
| 46 | P40227 | NA | CCT6A | chaperonin containing TCP1, subunit 6A (zeta 1) | 908 | ENSG00000146731 |
| 47 | P52292 | NA | KPNA2 | karyopherin alpha 2 (RAG cohort 1, importin alpha 1) | 3838 | ENSG00000182481 |
| 48 | P00352 | NA | ALDH1A1 | aldehyde dehydrogenase 1 family, member A1 | 216 | ENSG00000165092 |

  
  

| **Database:biological process      &nbspName:macromolecular complex assembly      &nbspID:GO:0065003** | | | | | | |
| --- | --- | --- | --- | --- | --- | --- |
| C=986; O=11; E=4.24; R=2.60; rawP=0.0029; adjP=0.0442 | | | | | | |
| Index | UserID | Value | Gene Symbol | Gene Name | EntrezGene | Ensembl |
| 1 | P07814 | NA | EPRS | glutamyl-prolyl-tRNA synthetase | 2058 | ENSG00000136628 |
| 2 | P78371 | NA | CCT2 | chaperonin containing TCP1, subunit 2 (beta) | 10576 | ENSG00000166226 |
| 3 | P17987 | NA | TCP1 | t-complex 1 | 6950 | ENSG00000120438 |
| 4 | P55209 | NA | NAP1L1 | nucleosome assembly protein 1-like 1 | 4673 | ENSG00000187109 |
| 5 | P34932 | NA | HSPA4 | heat shock 70kDa protein 4 | 3308 | ENSG00000170606 |
| 6 | Q13885 | NA | TUBB2A | tubulin, beta 2A class IIa | 7280 | ENSG00000137267 |
| 7 | P68371 | NA | TUBB4B | tubulin, beta 4B class IVb | 10383 | ENSG00000188229 |
| 8 | Q9BQE3 | NA | TUBA1C | tubulin, alpha 1c | 84790 | ENSG00000167553 |
| 9 | P21980 | NA | TGM2 | transglutaminase 2 (C polypeptide, protein-glutamine-gamma-glutamyltransferase) | 7052 | ENSG00000198959 |
| 10 | P49736 | NA | MCM2 | minichromosome maintenance complex component 2 | 4171 | ENSG00000073111 |
| 11 | O60610 | NA | DIAPH1 | diaphanous homolog 1 (Drosophila) | 1729 | ENSG00000131504 |

  
  

| **Database:biological process      &nbspName:nucleobase-containing small molecule metabolic process      &nbspID:GO:0055086** | | | | | | |
| --- | --- | --- | --- | --- | --- | --- |
| C=848; O=10; E=3.65; R=2.74; rawP=0.0031; adjP=0.0451 | | | | | | |
| Index | UserID | Value | Gene Symbol | Gene Name | EntrezGene | Ensembl |
| 1 | Q16881 | NA | TXNRD1 | thioredoxin reductase 1 | 7296 | ENSG00000198431 |
| 2 | P22102 | NA | GART | phosphoribosylglycinamide formyltransferase, phosphoribosylglycinamide synthetase, phosphoribosylaminoimidazole synthetase | 2618 | ENSG00000159131 |
| 3 | P00491 | NA | PNP | purine nucleoside phosphorylase | 4860 | ENSG00000198805 |
| 4 | P07951 | NA | TPM2 | tropomyosin 2 (beta) | 7169 | ENSG00000198467 |
| 5 | P41091 | NA | EIF2S3 | eukaryotic translation initiation factor 2, subunit 3 gamma, 52kDa | 1968 | ENSG00000130741 |
| 6 | O95433 | NA | AHSA1 | AHA1, activator of heat shock 90kDa protein ATPase homolog 1 (yeast) | 10598 | ENSG00000100591 |
| 7 | Q16555 | NA | DPYSL2 | dihydropyrimidinase-like 2 | 1808 | ENSG00000092964 |
| 8 | P12268 | NA | IMPDH2 | IMP (inosine 5'-monophosphate) dehydrogenase 2 | 3615 | ENSG00000178035 |
| 9 | P00352 | NA | ALDH1A1 | aldehyde dehydrogenase 1 family, member A1 | 216 | ENSG00000165092 |
| 10 | P11586 | NA | MTHFD1 | methylenetetrahydrofolate dehydrogenase (NADP+ dependent) 1, methenyltetrahydrofolate cyclohydrolase, formyltetrahydrofolate synthetase | 4522 | ENSG00000100714 |

  
  

| **Database:biological process      &nbspName:organonitrogen compound metabolic process      &nbspID:GO:1901564** | | | | | | |
| --- | --- | --- | --- | --- | --- | --- |
| C=1673; O=15; E=7.19; R=2.09; rawP=0.0041; adjP=0.0525 | | | | | | |
| Index | UserID | Value | Gene Symbol | Gene Name | EntrezGene | Ensembl |
| 1 | P63241 | NA | EIF5A | eukaryotic translation initiation factor 5A | 1984 | ENSG00000132507 |
| 2 | P07814 | NA | EPRS | glutamyl-prolyl-tRNA synthetase | 2058 | ENSG00000136628 |
| 3 | P22102 | NA | GART | phosphoribosylglycinamide formyltransferase, phosphoribosylglycinamide synthetase, phosphoribosylaminoimidazole synthetase | 2618 | ENSG00000159131 |
| 4 | P00491 | NA | PNP | purine nucleoside phosphorylase | 4860 | ENSG00000198805 |
| 5 | P07951 | NA | TPM2 | tropomyosin 2 (beta) | 7169 | ENSG00000198467 |
| 6 | P41091 | NA | EIF2S3 | eukaryotic translation initiation factor 2, subunit 3 gamma, 52kDa | 1968 | ENSG00000130741 |
| 7 | P12081 | NA | HARS | histidyl-tRNA synthetase | 3035 | ENSG00000170445 |
| 8 | O95433 | NA | AHSA1 | AHA1, activator of heat shock 90kDa protein ATPase homolog 1 (yeast) | 10598 | ENSG00000100591 |
| 9 | Q16555 | NA | DPYSL2 | dihydropyrimidinase-like 2 | 1808 | ENSG00000092964 |
| 10 | P49327 | NA | FASN | fatty acid synthase | 2194 | ENSG00000169710 |
| 11 | P56192 | NA | MARS | methionyl-tRNA synthetase | 4141 | ENSG00000166986 |
| 12 | P17174 | NA | GOT1 | glutamic-oxaloacetic transaminase 1, soluble (aspartate aminotransferase 1) | 2805 | ENSG00000120053 |
| 13 | P12268 | NA | IMPDH2 | IMP (inosine 5'-monophosphate) dehydrogenase 2 | 3615 | ENSG00000178035 |
| 14 | P00352 | NA | ALDH1A1 | aldehyde dehydrogenase 1 family, member A1 | 216 | ENSG00000165092 |
| 15 | P11586 | NA | MTHFD1 | methylenetetrahydrofolate dehydrogenase (NADP+ dependent) 1, methenyltetrahydrofolate cyclohydrolase, formyltetrahydrofolate synthetase | 4522 | ENSG00000100714 |

  
  

| **Database:biological process      &nbspName:translational initiation      &nbspID:GO:0006413** | | | | | | |
| --- | --- | --- | --- | --- | --- | --- |
| C=152; O=4; E=0.65; R=6.12; rawP=0.0041; adjP=0.0525 | | | | | | |
| Index | UserID | Value | Gene Symbol | Gene Name | EntrezGene | Ensembl |
| 1 | P62906 | NA | RPL10A | ribosomal protein L10a | 4736 | ENSG00000198755 |
| 2 | P41091 | NA | EIF2S3 | eukaryotic translation initiation factor 2, subunit 3 gamma, 52kDa | 1968 | ENSG00000130741 |
| 3 | Q15056 | NA | EIF4H | eukaryotic translation initiation factor 4H | 7458 | ENSG00000106682 |
| 4 | P41567 | NA | EIF1 | eukaryotic translation initiation factor 1 | 10209 | ENSG00000173812 |

  
  

| **Database:biological process      &nbspName:protein polymerization      &nbspID:GO:0051258** | | | | | | |
| --- | --- | --- | --- | --- | --- | --- |
| C=151; O=4; E=0.65; R=6.16; rawP=0.0040; adjP=0.0525 | | | | | | |
| Index | UserID | Value | Gene Symbol | Gene Name | EntrezGene | Ensembl |
| 1 | Q9BQE3 | NA | TUBA1C | tubulin, alpha 1c | 84790 | ENSG00000167553 |
| 2 | Q13885 | NA | TUBB2A | tubulin, beta 2A class IIa | 7280 | ENSG00000137267 |
| 3 | P68371 | NA | TUBB4B | tubulin, beta 4B class IVb | 10383 | ENSG00000188229 |
| 4 | O60610 | NA | DIAPH1 | diaphanous homolog 1 (Drosophila) | 1729 | ENSG00000131504 |

  
  

| **Database:biological process      &nbspName:single-organism metabolic process      &nbspID:GO:0044710** | | | | | | |
| --- | --- | --- | --- | --- | --- | --- |
| C=9109; O=49; E=39.16; R=1.25; rawP=0.0054; adjP=0.0607 | | | | | | |
| Index | UserID | Value | Gene Symbol | Gene Name | EntrezGene | Ensembl |
| 1 | Q92598 | NA | HSPH1 | heat shock 105kDa/110kDa protein 1 | 10808 | ENSG00000120694 |
| 2 | Q13885 | NA | TUBB2A | tubulin, beta 2A class IIa | 7280 | ENSG00000137267 |
| 3 | P68371 | NA | TUBB4B | tubulin, beta 4B class IVb | 10383 | ENSG00000188229 |
| 4 | P25685 | NA | DNAJB1 | DnaJ (Hsp40) homolog, subfamily B, member 1 | 3337 | ENSG00000132002 |
| 5 | P53701 | NA | HCCS | holocytochrome c synthase | 3052 | ENSG00000004961 |
| 6 | P68104 | NA | EEF1A1 | eukaryotic translation elongation factor 1 alpha 1 | 1915 | ENSG00000156508 |
| 7 | P02786 | NA | TFRC | transferrin receptor (p90, CD71) | 7037 | ENSG00000072274 |
| 8 | P50991 | NA | CCT4 | chaperonin containing TCP1, subunit 4 (delta) | 10575 | ENSG00000115484 |
| 9 | O00425 | NA | IGF2BP3 | insulin-like growth factor 2 mRNA binding protein 3 | 10643 | ENSG00000136231 |
| 10 | P22102 | NA | GART | phosphoribosylglycinamide formyltransferase, phosphoribosylglycinamide synthetase, phosphoribosylaminoimidazole synthetase | 2618 | ENSG00000159131 |
| 11 | P13639 | NA | EEF2 | eukaryotic translation elongation factor 2 | 1938 | ENSG00000167658 |
| 12 | Q16555 | NA | DPYSL2 | dihydropyrimidinase-like 2 | 1808 | ENSG00000092964 |
| 13 | O95433 | NA | AHSA1 | AHA1, activator of heat shock 90kDa protein ATPase homolog 1 (yeast) | 10598 | ENSG00000100591 |
| 14 | P41567 | NA | EIF1 | eukaryotic translation initiation factor 1 | 10209 | ENSG00000173812 |
| 15 | P17174 | NA | GOT1 | glutamic-oxaloacetic transaminase 1, soluble (aspartate aminotransferase 1) | 2805 | ENSG00000120053 |
| 16 | P67809 | NA | YBX1 | Y box binding protein 1 | 4904 | ENSG00000065978 |
| 17 | P21980 | NA | TGM2 | transglutaminase 2 (C polypeptide, protein-glutamine-gamma-glutamyltransferase) | 7052 | ENSG00000198959 |
| 18 | Q99832 | NA | CCT7 | chaperonin containing TCP1, subunit 7 (eta) | 10574 | ENSG00000135624 |
| 19 | Q15185 | NA | PTGES3 | prostaglandin E synthase 3 (cytosolic) | 10728 | ENSG00000110958 |
| 20 | Q6PIU2 | NA | NCEH1 | neutral cholesterol ester hydrolase 1 | 57552 | ENSG00000144959 |
| 21 | Q14166 | NA | TTLL12 | tubulin tyrosine ligase-like family, member 12 | 23170 | ENSG00000100304 |
| 22 | P11387 | NA | TOP1 | topoisomerase (DNA) I | 7150 | ENSG00000198900 |
| 23 | P11586 | NA | MTHFD1 | methylenetetrahydrofolate dehydrogenase (NADP+ dependent) 1, methenyltetrahydrofolate cyclohydrolase, formyltetrahydrofolate synthetase | 4522 | ENSG00000100714 |
| 24 | Q16881 | NA | TXNRD1 | thioredoxin reductase 1 | 7296 | ENSG00000198431 |
| 25 | P63241 | NA | EIF5A | eukaryotic translation initiation factor 5A | 1984 | ENSG00000132507 |
| 26 | P78371 | NA | CCT2 | chaperonin containing TCP1, subunit 2 (beta) | 10576 | ENSG00000166226 |
| 27 | P08195 | NA | SLC3A2 | solute carrier family 3 (activators of dibasic and neutral amino acid transport), member 2 | 6520 | ENSG00000168003 |
| 28 | Q7Z2K6 | NA | ERMP1 | endoplasmic reticulum metallopeptidase 1 | 79956 | ENSG00000099219 |
| 29 | P07951 | NA | TPM2 | tropomyosin 2 (beta) | 7169 | ENSG00000198467 |
| 30 | P00491 | NA | PNP | purine nucleoside phosphorylase | 4860 | ENSG00000198805 |
| 31 | P49321 | NA | NASP | nuclear autoantigenic sperm protein (histone-binding) | 4678 | ENSG00000132780 |
| 32 | P12081 | NA | HARS | histidyl-tRNA synthetase | 3035 | ENSG00000170445 |
| 33 | P49327 | NA | FASN | fatty acid synthase | 2194 | ENSG00000169710 |
| 34 | P49368 | NA | CCT3 | chaperonin containing TCP1, subunit 3 (gamma) | 7203 | ENSG00000163468 |
| 35 | P56192 | NA | MARS | methionyl-tRNA synthetase | 4141 | ENSG00000166986 |
| 36 | Q9BQE3 | NA | TUBA1C | tubulin, alpha 1c | 84790 | ENSG00000167553 |
| 37 | P12268 | NA | IMPDH2 | IMP (inosine 5'-monophosphate) dehydrogenase 2 | 3615 | ENSG00000178035 |
| 38 | P62906 | NA | RPL10A | ribosomal protein L10a | 4736 | ENSG00000198755 |
| 39 | P49736 | NA | MCM2 | minichromosome maintenance complex component 2 | 4171 | ENSG00000073111 |
| 40 | Q15056 | NA | EIF4H | eukaryotic translation initiation factor 4H | 7458 | ENSG00000106682 |
| 41 | P50990 | NA | CCT8 | chaperonin containing TCP1, subunit 8 (theta) | 10694 | ENSG00000156261 |
| 42 | P07814 | NA | EPRS | glutamyl-prolyl-tRNA synthetase | 2058 | ENSG00000136628 |
| 43 | P17987 | NA | TCP1 | t-complex 1 | 6950 | ENSG00000120438 |
| 44 | P55209 | NA | NAP1L1 | nucleosome assembly protein 1-like 1 | 4673 | ENSG00000187109 |
| 45 | P41091 | NA | EIF2S3 | eukaryotic translation initiation factor 2, subunit 3 gamma, 52kDa | 1968 | ENSG00000130741 |
| 46 | P07108 | NA | DBI | diazepam binding inhibitor (GABA receptor modulator, acyl-CoA binding protein) | 1622 | ENSG00000155368 |
| 47 | P40227 | NA | CCT6A | chaperonin containing TCP1, subunit 6A (zeta 1) | 908 | ENSG00000146731 |
| 48 | P52292 | NA | KPNA2 | karyopherin alpha 2 (RAG cohort 1, importin alpha 1) | 3838 | ENSG00000182481 |
| 49 | P00352 | NA | ALDH1A1 | aldehyde dehydrogenase 1 family, member A1 | 216 | ENSG00000165092 |

  
  

| **Database:biological process      &nbspName:metabolic process      &nbspID:GO:0008152** | | | | | | |
| --- | --- | --- | --- | --- | --- | --- |
| C=9363; O=50; E=40.25; R=1.24; rawP=0.0052; adjP=0.0607 | | | | | | |
| Index | UserID | Value | Gene Symbol | Gene Name | EntrezGene | Ensembl |
| 1 | Q92598 | NA | HSPH1 | heat shock 105kDa/110kDa protein 1 | 10808 | ENSG00000120694 |
| 2 | Q13885 | NA | TUBB2A | tubulin, beta 2A class IIa | 7280 | ENSG00000137267 |
| 3 | P68371 | NA | TUBB4B | tubulin, beta 4B class IVb | 10383 | ENSG00000188229 |
| 4 | P25685 | NA | DNAJB1 | DnaJ (Hsp40) homolog, subfamily B, member 1 | 3337 | ENSG00000132002 |
| 5 | P53701 | NA | HCCS | holocytochrome c synthase | 3052 | ENSG00000004961 |
| 6 | P68104 | NA | EEF1A1 | eukaryotic translation elongation factor 1 alpha 1 | 1915 | ENSG00000156508 |
| 7 | P02786 | NA | TFRC | transferrin receptor (p90, CD71) | 7037 | ENSG00000072274 |
| 8 | P50991 | NA | CCT4 | chaperonin containing TCP1, subunit 4 (delta) | 10575 | ENSG00000115484 |
| 9 | O00425 | NA | IGF2BP3 | insulin-like growth factor 2 mRNA binding protein 3 | 10643 | ENSG00000136231 |
| 10 | P22102 | NA | GART | phosphoribosylglycinamide formyltransferase, phosphoribosylglycinamide synthetase, phosphoribosylaminoimidazole synthetase | 2618 | ENSG00000159131 |
| 11 | P13639 | NA | EEF2 | eukaryotic translation elongation factor 2 | 1938 | ENSG00000167658 |
| 12 | Q16555 | NA | DPYSL2 | dihydropyrimidinase-like 2 | 1808 | ENSG00000092964 |
| 13 | O95433 | NA | AHSA1 | AHA1, activator of heat shock 90kDa protein ATPase homolog 1 (yeast) | 10598 | ENSG00000100591 |
| 14 | P41567 | NA | EIF1 | eukaryotic translation initiation factor 1 | 10209 | ENSG00000173812 |
| 15 | P17174 | NA | GOT1 | glutamic-oxaloacetic transaminase 1, soluble (aspartate aminotransferase 1) | 2805 | ENSG00000120053 |
| 16 | P67809 | NA | YBX1 | Y box binding protein 1 | 4904 | ENSG00000065978 |
| 17 | P21980 | NA | TGM2 | transglutaminase 2 (C polypeptide, protein-glutamine-gamma-glutamyltransferase) | 7052 | ENSG00000198959 |
| 18 | Q99832 | NA | CCT7 | chaperonin containing TCP1, subunit 7 (eta) | 10574 | ENSG00000135624 |
| 19 | Q15185 | NA | PTGES3 | prostaglandin E synthase 3 (cytosolic) | 10728 | ENSG00000110958 |
| 20 | Q6PIU2 | NA | NCEH1 | neutral cholesterol ester hydrolase 1 | 57552 | ENSG00000144959 |
| 21 | Q14166 | NA | TTLL12 | tubulin tyrosine ligase-like family, member 12 | 23170 | ENSG00000100304 |
| 22 | P11387 | NA | TOP1 | topoisomerase (DNA) I | 7150 | ENSG00000198900 |
| 23 | P11586 | NA | MTHFD1 | methylenetetrahydrofolate dehydrogenase (NADP+ dependent) 1, methenyltetrahydrofolate cyclohydrolase, formyltetrahydrofolate synthetase | 4522 | ENSG00000100714 |
| 24 | Q16881 | NA | TXNRD1 | thioredoxin reductase 1 | 7296 | ENSG00000198431 |
| 25 | P63241 | NA | EIF5A | eukaryotic translation initiation factor 5A | 1984 | ENSG00000132507 |
| 26 | P78371 | NA | CCT2 | chaperonin containing TCP1, subunit 2 (beta) | 10576 | ENSG00000166226 |
| 27 | P08195 | NA | SLC3A2 | solute carrier family 3 (activators of dibasic and neutral amino acid transport), member 2 | 6520 | ENSG00000168003 |
| 28 | Q7Z2K6 | NA | ERMP1 | endoplasmic reticulum metallopeptidase 1 | 79956 | ENSG00000099219 |
| 29 | P07951 | NA | TPM2 | tropomyosin 2 (beta) | 7169 | ENSG00000198467 |
| 30 | P00491 | NA | PNP | purine nucleoside phosphorylase | 4860 | ENSG00000198805 |
| 31 | P49321 | NA | NASP | nuclear autoantigenic sperm protein (histone-binding) | 4678 | ENSG00000132780 |
| 32 | P12081 | NA | HARS | histidyl-tRNA synthetase | 3035 | ENSG00000170445 |
| 33 | P49327 | NA | FASN | fatty acid synthase | 2194 | ENSG00000169710 |
| 34 | P49368 | NA | CCT3 | chaperonin containing TCP1, subunit 3 (gamma) | 7203 | ENSG00000163468 |
| 35 | P56192 | NA | MARS | methionyl-tRNA synthetase | 4141 | ENSG00000166986 |
| 36 | Q9BQE3 | NA | TUBA1C | tubulin, alpha 1c | 84790 | ENSG00000167553 |
| 37 | P12268 | NA | IMPDH2 | IMP (inosine 5'-monophosphate) dehydrogenase 2 | 3615 | ENSG00000178035 |
| 38 | P62906 | NA | RPL10A | ribosomal protein L10a | 4736 | ENSG00000198755 |
| 39 | P49736 | NA | MCM2 | minichromosome maintenance complex component 2 | 4171 | ENSG00000073111 |
| 40 | Q15056 | NA | EIF4H | eukaryotic translation initiation factor 4H | 7458 | ENSG00000106682 |
| 41 | P50990 | NA | CCT8 | chaperonin containing TCP1, subunit 8 (theta) | 10694 | ENSG00000156261 |
| 42 | Q06830 | NA | PRDX1 | peroxiredoxin 1 | 5052 | ENSG00000117450 |
| 43 | P07814 | NA | EPRS | glutamyl-prolyl-tRNA synthetase | 2058 | ENSG00000136628 |
| 44 | P17987 | NA | TCP1 | t-complex 1 | 6950 | ENSG00000120438 |
| 45 | P55209 | NA | NAP1L1 | nucleosome assembly protein 1-like 1 | 4673 | ENSG00000187109 |
| 46 | P41091 | NA | EIF2S3 | eukaryotic translation initiation factor 2, subunit 3 gamma, 52kDa | 1968 | ENSG00000130741 |
| 47 | P07108 | NA | DBI | diazepam binding inhibitor (GABA receptor modulator, acyl-CoA binding protein) | 1622 | ENSG00000155368 |
| 48 | P40227 | NA | CCT6A | chaperonin containing TCP1, subunit 6A (zeta 1) | 908 | ENSG00000146731 |
| 49 | P52292 | NA | KPNA2 | karyopherin alpha 2 (RAG cohort 1, importin alpha 1) | 3838 | ENSG00000182481 |
| 50 | P00352 | NA | ALDH1A1 | aldehyde dehydrogenase 1 family, member A1 | 216 | ENSG00000165092 |

  
  

| **Database:biological process      &nbspName:protein complex biogenesis      &nbspID:GO:0070271** | | | | | | |
| --- | --- | --- | --- | --- | --- | --- |
| C=772; O=9; E=3.32; R=2.71; rawP=0.0055; adjP=0.0607 | | | | | | |
| Index | UserID | Value | Gene Symbol | Gene Name | EntrezGene | Ensembl |
| 1 | P07814 | NA | EPRS | glutamyl-prolyl-tRNA synthetase | 2058 | ENSG00000136628 |
| 2 | P78371 | NA | CCT2 | chaperonin containing TCP1, subunit 2 (beta) | 10576 | ENSG00000166226 |
| 3 | P17987 | NA | TCP1 | t-complex 1 | 6950 | ENSG00000120438 |
| 4 | P34932 | NA | HSPA4 | heat shock 70kDa protein 4 | 3308 | ENSG00000170606 |
| 5 | Q13885 | NA | TUBB2A | tubulin, beta 2A class IIa | 7280 | ENSG00000137267 |
| 6 | P68371 | NA | TUBB4B | tubulin, beta 4B class IVb | 10383 | ENSG00000188229 |
| 7 | Q9BQE3 | NA | TUBA1C | tubulin, alpha 1c | 84790 | ENSG00000167553 |
| 8 | P21980 | NA | TGM2 | transglutaminase 2 (C polypeptide, protein-glutamine-gamma-glutamyltransferase) | 7052 | ENSG00000198959 |
| 9 | O60610 | NA | DIAPH1 | diaphanous homolog 1 (Drosophila) | 1729 | ENSG00000131504 |

  
  

| **Database:biological process      &nbspName:protein complex assembly      &nbspID:GO:0006461** | | | | | | |
| --- | --- | --- | --- | --- | --- | --- |
| C=769; O=9; E=3.31; R=2.72; rawP=0.0053; adjP=0.0607 | | | | | | |
| Index | UserID | Value | Gene Symbol | Gene Name | EntrezGene | Ensembl |
| 1 | P07814 | NA | EPRS | glutamyl-prolyl-tRNA synthetase | 2058 | ENSG00000136628 |
| 2 | P78371 | NA | CCT2 | chaperonin containing TCP1, subunit 2 (beta) | 10576 | ENSG00000166226 |
| 3 | P17987 | NA | TCP1 | t-complex 1 | 6950 | ENSG00000120438 |
| 4 | P34932 | NA | HSPA4 | heat shock 70kDa protein 4 | 3308 | ENSG00000170606 |
| 5 | Q13885 | NA | TUBB2A | tubulin, beta 2A class IIa | 7280 | ENSG00000137267 |
| 6 | P68371 | NA | TUBB4B | tubulin, beta 4B class IVb | 10383 | ENSG00000188229 |
| 7 | Q9BQE3 | NA | TUBA1C | tubulin, alpha 1c | 84790 | ENSG00000167553 |
| 8 | P21980 | NA | TGM2 | transglutaminase 2 (C polypeptide, protein-glutamine-gamma-glutamyltransferase) | 7052 | ENSG00000198959 |
| 9 | O60610 | NA | DIAPH1 | diaphanous homolog 1 (Drosophila) | 1729 | ENSG00000131504 |

  
  

| **Database:biological process      &nbspName:organic substance metabolic process      &nbspID:GO:0071704** | | | | | | |
| --- | --- | --- | --- | --- | --- | --- |
| C=8902; O=48; E=38.27; R=1.25; rawP=0.0064; adjP=0.0683 | | | | | | |
| Index | UserID | Value | Gene Symbol | Gene Name | EntrezGene | Ensembl |
| 1 | Q92598 | NA | HSPH1 | heat shock 105kDa/110kDa protein 1 | 10808 | ENSG00000120694 |
| 2 | Q13885 | NA | TUBB2A | tubulin, beta 2A class IIa | 7280 | ENSG00000137267 |
| 3 | P68371 | NA | TUBB4B | tubulin, beta 4B class IVb | 10383 | ENSG00000188229 |
| 4 | P25685 | NA | DNAJB1 | DnaJ (Hsp40) homolog, subfamily B, member 1 | 3337 | ENSG00000132002 |
| 5 | P68104 | NA | EEF1A1 | eukaryotic translation elongation factor 1 alpha 1 | 1915 | ENSG00000156508 |
| 6 | P02786 | NA | TFRC | transferrin receptor (p90, CD71) | 7037 | ENSG00000072274 |
| 7 | P50991 | NA | CCT4 | chaperonin containing TCP1, subunit 4 (delta) | 10575 | ENSG00000115484 |
| 8 | O00425 | NA | IGF2BP3 | insulin-like growth factor 2 mRNA binding protein 3 | 10643 | ENSG00000136231 |
| 9 | P22102 | NA | GART | phosphoribosylglycinamide formyltransferase, phosphoribosylglycinamide synthetase, phosphoribosylaminoimidazole synthetase | 2618 | ENSG00000159131 |
| 10 | P13639 | NA | EEF2 | eukaryotic translation elongation factor 2 | 1938 | ENSG00000167658 |
| 11 | Q16555 | NA | DPYSL2 | dihydropyrimidinase-like 2 | 1808 | ENSG00000092964 |
| 12 | O95433 | NA | AHSA1 | AHA1, activator of heat shock 90kDa protein ATPase homolog 1 (yeast) | 10598 | ENSG00000100591 |
| 13 | P41567 | NA | EIF1 | eukaryotic translation initiation factor 1 | 10209 | ENSG00000173812 |
| 14 | P17174 | NA | GOT1 | glutamic-oxaloacetic transaminase 1, soluble (aspartate aminotransferase 1) | 2805 | ENSG00000120053 |
| 15 | P67809 | NA | YBX1 | Y box binding protein 1 | 4904 | ENSG00000065978 |
| 16 | P21980 | NA | TGM2 | transglutaminase 2 (C polypeptide, protein-glutamine-gamma-glutamyltransferase) | 7052 | ENSG00000198959 |
| 17 | Q99832 | NA | CCT7 | chaperonin containing TCP1, subunit 7 (eta) | 10574 | ENSG00000135624 |
| 18 | Q15185 | NA | PTGES3 | prostaglandin E synthase 3 (cytosolic) | 10728 | ENSG00000110958 |
| 19 | Q6PIU2 | NA | NCEH1 | neutral cholesterol ester hydrolase 1 | 57552 | ENSG00000144959 |
| 20 | Q14166 | NA | TTLL12 | tubulin tyrosine ligase-like family, member 12 | 23170 | ENSG00000100304 |
| 21 | P11387 | NA | TOP1 | topoisomerase (DNA) I | 7150 | ENSG00000198900 |
| 22 | P11586 | NA | MTHFD1 | methylenetetrahydrofolate dehydrogenase (NADP+ dependent) 1, methenyltetrahydrofolate cyclohydrolase, formyltetrahydrofolate synthetase | 4522 | ENSG00000100714 |
| 23 | Q16881 | NA | TXNRD1 | thioredoxin reductase 1 | 7296 | ENSG00000198431 |
| 24 | P63241 | NA | EIF5A | eukaryotic translation initiation factor 5A | 1984 | ENSG00000132507 |
| 25 | P78371 | NA | CCT2 | chaperonin containing TCP1, subunit 2 (beta) | 10576 | ENSG00000166226 |
| 26 | P08195 | NA | SLC3A2 | solute carrier family 3 (activators of dibasic and neutral amino acid transport), member 2 | 6520 | ENSG00000168003 |
| 27 | Q7Z2K6 | NA | ERMP1 | endoplasmic reticulum metallopeptidase 1 | 79956 | ENSG00000099219 |
| 28 | P07951 | NA | TPM2 | tropomyosin 2 (beta) | 7169 | ENSG00000198467 |
| 29 | P00491 | NA | PNP | purine nucleoside phosphorylase | 4860 | ENSG00000198805 |
| 30 | P49321 | NA | NASP | nuclear autoantigenic sperm protein (histone-binding) | 4678 | ENSG00000132780 |
| 31 | P12081 | NA | HARS | histidyl-tRNA synthetase | 3035 | ENSG00000170445 |
| 32 | P49327 | NA | FASN | fatty acid synthase | 2194 | ENSG00000169710 |
| 33 | P49368 | NA | CCT3 | chaperonin containing TCP1, subunit 3 (gamma) | 7203 | ENSG00000163468 |
| 34 | Q9BQE3 | NA | TUBA1C | tubulin, alpha 1c | 84790 | ENSG00000167553 |
| 35 | P56192 | NA | MARS | methionyl-tRNA synthetase | 4141 | ENSG00000166986 |
| 36 | P12268 | NA | IMPDH2 | IMP (inosine 5'-monophosphate) dehydrogenase 2 | 3615 | ENSG00000178035 |
| 37 | P62906 | NA | RPL10A | ribosomal protein L10a | 4736 | ENSG00000198755 |
| 38 | P49736 | NA | MCM2 | minichromosome maintenance complex component 2 | 4171 | ENSG00000073111 |
| 39 | Q15056 | NA | EIF4H | eukaryotic translation initiation factor 4H | 7458 | ENSG00000106682 |
| 40 | P50990 | NA | CCT8 | chaperonin containing TCP1, subunit 8 (theta) | 10694 | ENSG00000156261 |
| 41 | P07814 | NA | EPRS | glutamyl-prolyl-tRNA synthetase | 2058 | ENSG00000136628 |
| 42 | P17987 | NA | TCP1 | t-complex 1 | 6950 | ENSG00000120438 |
| 43 | P55209 | NA | NAP1L1 | nucleosome assembly protein 1-like 1 | 4673 | ENSG00000187109 |
| 44 | P41091 | NA | EIF2S3 | eukaryotic translation initiation factor 2, subunit 3 gamma, 52kDa | 1968 | ENSG00000130741 |
| 45 | P07108 | NA | DBI | diazepam binding inhibitor (GABA receptor modulator, acyl-CoA binding protein) | 1622 | ENSG00000155368 |
| 46 | P40227 | NA | CCT6A | chaperonin containing TCP1, subunit 6A (zeta 1) | 908 | ENSG00000146731 |
| 47 | P52292 | NA | KPNA2 | karyopherin alpha 2 (RAG cohort 1, importin alpha 1) | 3838 | ENSG00000182481 |
| 48 | P00352 | NA | ALDH1A1 | aldehyde dehydrogenase 1 family, member A1 | 216 | ENSG00000165092 |

  
  

| **Database:biological process      &nbspName:purine nucleotide metabolic process      &nbspID:GO:0006163** | | | | | | |
| --- | --- | --- | --- | --- | --- | --- |
| C=662; O=8; E=2.85; R=2.81; rawP=0.0072; adjP=0.0743 | | | | | | |
| Index | UserID | Value | Gene Symbol | Gene Name | EntrezGene | Ensembl |
| 1 | P22102 | NA | GART | phosphoribosylglycinamide formyltransferase, phosphoribosylglycinamide synthetase, phosphoribosylaminoimidazole synthetase | 2618 | ENSG00000159131 |
| 2 | P00491 | NA | PNP | purine nucleoside phosphorylase | 4860 | ENSG00000198805 |
| 3 | P07951 | NA | TPM2 | tropomyosin 2 (beta) | 7169 | ENSG00000198467 |
| 4 | P41091 | NA | EIF2S3 | eukaryotic translation initiation factor 2, subunit 3 gamma, 52kDa | 1968 | ENSG00000130741 |
| 5 | O95433 | NA | AHSA1 | AHA1, activator of heat shock 90kDa protein ATPase homolog 1 (yeast) | 10598 | ENSG00000100591 |
| 6 | P12268 | NA | IMPDH2 | IMP (inosine 5'-monophosphate) dehydrogenase 2 | 3615 | ENSG00000178035 |
| 7 | P00352 | NA | ALDH1A1 | aldehyde dehydrogenase 1 family, member A1 | 216 | ENSG00000165092 |
| 8 | P11586 | NA | MTHFD1 | methylenetetrahydrofolate dehydrogenase (NADP+ dependent) 1, methenyltetrahydrofolate cyclohydrolase, formyltetrahydrofolate synthetase | 4522 | ENSG00000100714 |

  
  

| **Database:biological process      &nbspName:purine-containing compound metabolic process      &nbspID:GO:0072521** | | | | | | |
| --- | --- | --- | --- | --- | --- | --- |
| C=704; O=8; E=3.03; R=2.64; rawP=0.0102; adjP=0.0989 | | | | | | |
| Index | UserID | Value | Gene Symbol | Gene Name | EntrezGene | Ensembl |
| 1 | P22102 | NA | GART | phosphoribosylglycinamide formyltransferase, phosphoribosylglycinamide synthetase, phosphoribosylaminoimidazole synthetase | 2618 | ENSG00000159131 |
| 2 | P00491 | NA | PNP | purine nucleoside phosphorylase | 4860 | ENSG00000198805 |
| 3 | P07951 | NA | TPM2 | tropomyosin 2 (beta) | 7169 | ENSG00000198467 |
| 4 | P41091 | NA | EIF2S3 | eukaryotic translation initiation factor 2, subunit 3 gamma, 52kDa | 1968 | ENSG00000130741 |
| 5 | O95433 | NA | AHSA1 | AHA1, activator of heat shock 90kDa protein ATPase homolog 1 (yeast) | 10598 | ENSG00000100591 |
| 6 | P12268 | NA | IMPDH2 | IMP (inosine 5'-monophosphate) dehydrogenase 2 | 3615 | ENSG00000178035 |
| 7 | P00352 | NA | ALDH1A1 | aldehyde dehydrogenase 1 family, member A1 | 216 | ENSG00000165092 |
| 8 | P11586 | NA | MTHFD1 | methylenetetrahydrofolate dehydrogenase (NADP+ dependent) 1, methenyltetrahydrofolate cyclohydrolase, formyltetrahydrofolate synthetase | 4522 | ENSG00000100714 |

  
  

| **Database:biological process      &nbspName:coenzyme biosynthetic process      &nbspID:GO:0009108** | | | | | | |
| --- | --- | --- | --- | --- | --- | --- |
| C=104; O=3; E=0.45; R=6.71; rawP=0.0101; adjP=0.0989 | | | | | | |
| Index | UserID | Value | Gene Symbol | Gene Name | EntrezGene | Ensembl |
| 1 | P00491 | NA | PNP | purine nucleoside phosphorylase | 4860 | ENSG00000198805 |
| 2 | P49327 | NA | FASN | fatty acid synthase | 2194 | ENSG00000169710 |
| 3 | P11586 | NA | MTHFD1 | methylenetetrahydrofolate dehydrogenase (NADP+ dependent) 1, methenyltetrahydrofolate cyclohydrolase, formyltetrahydrofolate synthetase | 4522 | ENSG00000100714 |

  
  

| **Database:molecular function      &nbspName:unfolded protein binding      &nbspID:GO:0051082** | | | | | | |
| --- | --- | --- | --- | --- | --- | --- |
| C=120; O=10; E=0.50; R=20.13; rawP=5.64e-11; adjP=3.78e-09 | | | | | | |
| Index | UserID | Value | Gene Symbol | Gene Name | EntrezGene | Ensembl |
| 1 | P78371 | NA | CCT2 | chaperonin containing TCP1, subunit 2 (beta) | 10576 | ENSG00000166226 |
| 2 | P17987 | NA | TCP1 | t-complex 1 | 6950 | ENSG00000120438 |
| 3 | P68371 | NA | TUBB4B | tubulin, beta 4B class IVb | 10383 | ENSG00000188229 |
| 4 | P40227 | NA | CCT6A | chaperonin containing TCP1, subunit 6A (zeta 1) | 908 | ENSG00000146731 |
| 5 | P49368 | NA | CCT3 | chaperonin containing TCP1, subunit 3 (gamma) | 7203 | ENSG00000163468 |
| 6 | P25685 | NA | DNAJB1 | DnaJ (Hsp40) homolog, subfamily B, member 1 | 3337 | ENSG00000132002 |
| 7 | Q15185 | NA | PTGES3 | prostaglandin E synthase 3 (cytosolic) | 10728 | ENSG00000110958 |
| 8 | Q99832 | NA | CCT7 | chaperonin containing TCP1, subunit 7 (eta) | 10574 | ENSG00000135624 |
| 9 | P50990 | NA | CCT8 | chaperonin containing TCP1, subunit 8 (theta) | 10694 | ENSG00000156261 |
| 10 | P50991 | NA | CCT4 | chaperonin containing TCP1, subunit 4 (delta) | 10575 | ENSG00000115484 |

  
  

| **Database:molecular function      &nbspName:small molecule binding      &nbspID:GO:0036094** | | | | | | |
| --- | --- | --- | --- | --- | --- | --- |
| C=2595; O=31; E=10.74; R=2.89; rawP=3.65e-09; adjP=1.22e-07 | | | | | | |
| Index | UserID | Value | Gene Symbol | Gene Name | EntrezGene | Ensembl |
| 1 | Q16881 | NA | TXNRD1 | thioredoxin reductase 1 | 7296 | ENSG00000198431 |
| 2 | Q9NR30 | NA | DDX21 | DEAD (Asp-Glu-Ala-Asp) box helicase 21 | 9188 | ENSG00000165732 |
| 3 | P78371 | NA | CCT2 | chaperonin containing TCP1, subunit 2 (beta) | 10576 | ENSG00000166226 |
| 4 | Q92598 | NA | HSPH1 | heat shock 105kDa/110kDa protein 1 | 10808 | ENSG00000120694 |
| 5 | Q13885 | NA | TUBB2A | tubulin, beta 2A class IIa | 7280 | ENSG00000137267 |
| 6 | P00491 | NA | PNP | purine nucleoside phosphorylase | 4860 | ENSG00000198805 |
| 7 | P12081 | NA | HARS | histidyl-tRNA synthetase | 3035 | ENSG00000170445 |
| 8 | P68371 | NA | TUBB4B | tubulin, beta 4B class IVb | 10383 | ENSG00000188229 |
| 9 | P49327 | NA | FASN | fatty acid synthase | 2194 | ENSG00000169710 |
| 10 | P49368 | NA | CCT3 | chaperonin containing TCP1, subunit 3 (gamma) | 7203 | ENSG00000163468 |
| 11 | P56192 | NA | MARS | methionyl-tRNA synthetase | 4141 | ENSG00000166986 |
| 12 | Q9BQE3 | NA | TUBA1C | tubulin, alpha 1c | 84790 | ENSG00000167553 |
| 13 | P12268 | NA | IMPDH2 | IMP (inosine 5'-monophosphate) dehydrogenase 2 | 3615 | ENSG00000178035 |
| 14 | P49736 | NA | MCM2 | minichromosome maintenance complex component 2 | 4171 | ENSG00000073111 |
| 15 | Q15056 | NA | EIF4H | eukaryotic translation initiation factor 4H | 7458 | ENSG00000106682 |
| 16 | P50990 | NA | CCT8 | chaperonin containing TCP1, subunit 8 (theta) | 10694 | ENSG00000156261 |
| 17 | P68104 | NA | EEF1A1 | eukaryotic translation elongation factor 1 alpha 1 | 1915 | ENSG00000156508 |
| 18 | P50991 | NA | CCT4 | chaperonin containing TCP1, subunit 4 (delta) | 10575 | ENSG00000115484 |
| 19 | O00425 | NA | IGF2BP3 | insulin-like growth factor 2 mRNA binding protein 3 | 10643 | ENSG00000136231 |
| 20 | P07814 | NA | EPRS | glutamyl-prolyl-tRNA synthetase | 2058 | ENSG00000136628 |
| 21 | P17987 | NA | TCP1 | t-complex 1 | 6950 | ENSG00000120438 |
| 22 | P34932 | NA | HSPA4 | heat shock 70kDa protein 4 | 3308 | ENSG00000170606 |
| 23 | P22102 | NA | GART | phosphoribosylglycinamide formyltransferase, phosphoribosylglycinamide synthetase, phosphoribosylaminoimidazole synthetase | 2618 | ENSG00000159131 |
| 24 | P13639 | NA | EEF2 | eukaryotic translation elongation factor 2 | 1938 | ENSG00000167658 |
| 25 | P41091 | NA | EIF2S3 | eukaryotic translation initiation factor 2, subunit 3 gamma, 52kDa | 1968 | ENSG00000130741 |
| 26 | P40227 | NA | CCT6A | chaperonin containing TCP1, subunit 6A (zeta 1) | 908 | ENSG00000146731 |
| 27 | P17174 | NA | GOT1 | glutamic-oxaloacetic transaminase 1, soluble (aspartate aminotransferase 1) | 2805 | ENSG00000120053 |
| 28 | Q99832 | NA | CCT7 | chaperonin containing TCP1, subunit 7 (eta) | 10574 | ENSG00000135624 |
| 29 | P21980 | NA | TGM2 | transglutaminase 2 (C polypeptide, protein-glutamine-gamma-glutamyltransferase) | 7052 | ENSG00000198959 |
| 30 | P11586 | NA | MTHFD1 | methylenetetrahydrofolate dehydrogenase (NADP+ dependent) 1, methenyltetrahydrofolate cyclohydrolase, formyltetrahydrofolate synthetase | 4522 | ENSG00000100714 |
| 31 | P11387 | NA | TOP1 | topoisomerase (DNA) I | 7150 | ENSG00000198900 |

  
  

| **Database:molecular function      &nbspName:nucleoside phosphate binding      &nbspID:GO:1901265** | | | | | | |
| --- | --- | --- | --- | --- | --- | --- |
| C=2404; O=29; E=9.95; R=2.91; rawP=1.34e-08; adjP=1.80e-07 | | | | | | |
| Index | UserID | Value | Gene Symbol | Gene Name | EntrezGene | Ensembl |
| 1 | Q16881 | NA | TXNRD1 | thioredoxin reductase 1 | 7296 | ENSG00000198431 |
| 2 | Q9NR30 | NA | DDX21 | DEAD (Asp-Glu-Ala-Asp) box helicase 21 | 9188 | ENSG00000165732 |
| 3 | P78371 | NA | CCT2 | chaperonin containing TCP1, subunit 2 (beta) | 10576 | ENSG00000166226 |
| 4 | Q92598 | NA | HSPH1 | heat shock 105kDa/110kDa protein 1 | 10808 | ENSG00000120694 |
| 5 | Q13885 | NA | TUBB2A | tubulin, beta 2A class IIa | 7280 | ENSG00000137267 |
| 6 | P12081 | NA | HARS | histidyl-tRNA synthetase | 3035 | ENSG00000170445 |
| 7 | P68371 | NA | TUBB4B | tubulin, beta 4B class IVb | 10383 | ENSG00000188229 |
| 8 | P49327 | NA | FASN | fatty acid synthase | 2194 | ENSG00000169710 |
| 9 | P49368 | NA | CCT3 | chaperonin containing TCP1, subunit 3 (gamma) | 7203 | ENSG00000163468 |
| 10 | P56192 | NA | MARS | methionyl-tRNA synthetase | 4141 | ENSG00000166986 |
| 11 | Q9BQE3 | NA | TUBA1C | tubulin, alpha 1c | 84790 | ENSG00000167553 |
| 12 | P12268 | NA | IMPDH2 | IMP (inosine 5'-monophosphate) dehydrogenase 2 | 3615 | ENSG00000178035 |
| 13 | P49736 | NA | MCM2 | minichromosome maintenance complex component 2 | 4171 | ENSG00000073111 |
| 14 | Q15056 | NA | EIF4H | eukaryotic translation initiation factor 4H | 7458 | ENSG00000106682 |
| 15 | P50990 | NA | CCT8 | chaperonin containing TCP1, subunit 8 (theta) | 10694 | ENSG00000156261 |
| 16 | P68104 | NA | EEF1A1 | eukaryotic translation elongation factor 1 alpha 1 | 1915 | ENSG00000156508 |
| 17 | P50991 | NA | CCT4 | chaperonin containing TCP1, subunit 4 (delta) | 10575 | ENSG00000115484 |
| 18 | O00425 | NA | IGF2BP3 | insulin-like growth factor 2 mRNA binding protein 3 | 10643 | ENSG00000136231 |
| 19 | P07814 | NA | EPRS | glutamyl-prolyl-tRNA synthetase | 2058 | ENSG00000136628 |
| 20 | P17987 | NA | TCP1 | t-complex 1 | 6950 | ENSG00000120438 |
| 21 | P34932 | NA | HSPA4 | heat shock 70kDa protein 4 | 3308 | ENSG00000170606 |
| 22 | P22102 | NA | GART | phosphoribosylglycinamide formyltransferase, phosphoribosylglycinamide synthetase, phosphoribosylaminoimidazole synthetase | 2618 | ENSG00000159131 |
| 23 | P13639 | NA | EEF2 | eukaryotic translation elongation factor 2 | 1938 | ENSG00000167658 |
| 24 | P41091 | NA | EIF2S3 | eukaryotic translation initiation factor 2, subunit 3 gamma, 52kDa | 1968 | ENSG00000130741 |
| 25 | P40227 | NA | CCT6A | chaperonin containing TCP1, subunit 6A (zeta 1) | 908 | ENSG00000146731 |
| 26 | Q99832 | NA | CCT7 | chaperonin containing TCP1, subunit 7 (eta) | 10574 | ENSG00000135624 |
| 27 | P21980 | NA | TGM2 | transglutaminase 2 (C polypeptide, protein-glutamine-gamma-glutamyltransferase) | 7052 | ENSG00000198959 |
| 28 | P11586 | NA | MTHFD1 | methylenetetrahydrofolate dehydrogenase (NADP+ dependent) 1, methenyltetrahydrofolate cyclohydrolase, formyltetrahydrofolate synthetase | 4522 | ENSG00000100714 |
| 29 | P11387 | NA | TOP1 | topoisomerase (DNA) I | 7150 | ENSG00000198900 |

  
  

| **Database:molecular function      &nbspName:anion binding      &nbspID:GO:0043168** | | | | | | |
| --- | --- | --- | --- | --- | --- | --- |
| C=2371; O=29; E=9.82; R=2.95; rawP=9.73e-09; adjP=1.80e-07 | | | | | | |
| Index | UserID | Value | Gene Symbol | Gene Name | EntrezGene | Ensembl |
| 1 | Q16881 | NA | TXNRD1 | thioredoxin reductase 1 | 7296 | ENSG00000198431 |
| 2 | Q9NR30 | NA | DDX21 | DEAD (Asp-Glu-Ala-Asp) box helicase 21 | 9188 | ENSG00000165732 |
| 3 | P78371 | NA | CCT2 | chaperonin containing TCP1, subunit 2 (beta) | 10576 | ENSG00000166226 |
| 4 | Q92598 | NA | HSPH1 | heat shock 105kDa/110kDa protein 1 | 10808 | ENSG00000120694 |
| 5 | Q13885 | NA | TUBB2A | tubulin, beta 2A class IIa | 7280 | ENSG00000137267 |
| 6 | P00491 | NA | PNP | purine nucleoside phosphorylase | 4860 | ENSG00000198805 |
| 7 | P12081 | NA | HARS | histidyl-tRNA synthetase | 3035 | ENSG00000170445 |
| 8 | P68371 | NA | TUBB4B | tubulin, beta 4B class IVb | 10383 | ENSG00000188229 |
| 9 | P49327 | NA | FASN | fatty acid synthase | 2194 | ENSG00000169710 |
| 10 | P49368 | NA | CCT3 | chaperonin containing TCP1, subunit 3 (gamma) | 7203 | ENSG00000163468 |
| 11 | P56192 | NA | MARS | methionyl-tRNA synthetase | 4141 | ENSG00000166986 |
| 12 | Q9BQE3 | NA | TUBA1C | tubulin, alpha 1c | 84790 | ENSG00000167553 |
| 13 | P49736 | NA | MCM2 | minichromosome maintenance complex component 2 | 4171 | ENSG00000073111 |
| 14 | P50990 | NA | CCT8 | chaperonin containing TCP1, subunit 8 (theta) | 10694 | ENSG00000156261 |
| 15 | P68104 | NA | EEF1A1 | eukaryotic translation elongation factor 1 alpha 1 | 1915 | ENSG00000156508 |
| 16 | P50991 | NA | CCT4 | chaperonin containing TCP1, subunit 4 (delta) | 10575 | ENSG00000115484 |
| 17 | P07814 | NA | EPRS | glutamyl-prolyl-tRNA synthetase | 2058 | ENSG00000136628 |
| 18 | P17987 | NA | TCP1 | t-complex 1 | 6950 | ENSG00000120438 |
| 19 | P34932 | NA | HSPA4 | heat shock 70kDa protein 4 | 3308 | ENSG00000170606 |
| 20 | P22102 | NA | GART | phosphoribosylglycinamide formyltransferase, phosphoribosylglycinamide synthetase, phosphoribosylaminoimidazole synthetase | 2618 | ENSG00000159131 |
| 21 | P13639 | NA | EEF2 | eukaryotic translation elongation factor 2 | 1938 | ENSG00000167658 |
| 22 | P41091 | NA | EIF2S3 | eukaryotic translation initiation factor 2, subunit 3 gamma, 52kDa | 1968 | ENSG00000130741 |
| 23 | P07108 | NA | DBI | diazepam binding inhibitor (GABA receptor modulator, acyl-CoA binding protein) | 1622 | ENSG00000155368 |
| 24 | P40227 | NA | CCT6A | chaperonin containing TCP1, subunit 6A (zeta 1) | 908 | ENSG00000146731 |
| 25 | P17174 | NA | GOT1 | glutamic-oxaloacetic transaminase 1, soluble (aspartate aminotransferase 1) | 2805 | ENSG00000120053 |
| 26 | Q99832 | NA | CCT7 | chaperonin containing TCP1, subunit 7 (eta) | 10574 | ENSG00000135624 |
| 27 | P21980 | NA | TGM2 | transglutaminase 2 (C polypeptide, protein-glutamine-gamma-glutamyltransferase) | 7052 | ENSG00000198959 |
| 28 | P11586 | NA | MTHFD1 | methylenetetrahydrofolate dehydrogenase (NADP+ dependent) 1, methenyltetrahydrofolate cyclohydrolase, formyltetrahydrofolate synthetase | 4522 | ENSG00000100714 |
| 29 | P11387 | NA | TOP1 | topoisomerase (DNA) I | 7150 | ENSG00000198900 |

  
  

| **Database:molecular function      &nbspName:nucleotide binding      &nbspID:GO:0000166** | | | | | | |
| --- | --- | --- | --- | --- | --- | --- |
| C=2403; O=29; E=9.95; R=2.92; rawP=1.33e-08; adjP=1.80e-07 | | | | | | |
| Index | UserID | Value | Gene Symbol | Gene Name | EntrezGene | Ensembl |
| 1 | Q16881 | NA | TXNRD1 | thioredoxin reductase 1 | 7296 | ENSG00000198431 |
| 2 | Q9NR30 | NA | DDX21 | DEAD (Asp-Glu-Ala-Asp) box helicase 21 | 9188 | ENSG00000165732 |
| 3 | P78371 | NA | CCT2 | chaperonin containing TCP1, subunit 2 (beta) | 10576 | ENSG00000166226 |
| 4 | Q92598 | NA | HSPH1 | heat shock 105kDa/110kDa protein 1 | 10808 | ENSG00000120694 |
| 5 | Q13885 | NA | TUBB2A | tubulin, beta 2A class IIa | 7280 | ENSG00000137267 |
| 6 | P12081 | NA | HARS | histidyl-tRNA synthetase | 3035 | ENSG00000170445 |
| 7 | P68371 | NA | TUBB4B | tubulin, beta 4B class IVb | 10383 | ENSG00000188229 |
| 8 | P49327 | NA | FASN | fatty acid synthase | 2194 | ENSG00000169710 |
| 9 | P49368 | NA | CCT3 | chaperonin containing TCP1, subunit 3 (gamma) | 7203 | ENSG00000163468 |
| 10 | P56192 | NA | MARS | methionyl-tRNA synthetase | 4141 | ENSG00000166986 |
| 11 | Q9BQE3 | NA | TUBA1C | tubulin, alpha 1c | 84790 | ENSG00000167553 |
| 12 | P12268 | NA | IMPDH2 | IMP (inosine 5'-monophosphate) dehydrogenase 2 | 3615 | ENSG00000178035 |
| 13 | P49736 | NA | MCM2 | minichromosome maintenance complex component 2 | 4171 | ENSG00000073111 |
| 14 | Q15056 | NA | EIF4H | eukaryotic translation initiation factor 4H | 7458 | ENSG00000106682 |
| 15 | P50990 | NA | CCT8 | chaperonin containing TCP1, subunit 8 (theta) | 10694 | ENSG00000156261 |
| 16 | P68104 | NA | EEF1A1 | eukaryotic translation elongation factor 1 alpha 1 | 1915 | ENSG00000156508 |
| 17 | P50991 | NA | CCT4 | chaperonin containing TCP1, subunit 4 (delta) | 10575 | ENSG00000115484 |
| 18 | O00425 | NA | IGF2BP3 | insulin-like growth factor 2 mRNA binding protein 3 | 10643 | ENSG00000136231 |
| 19 | P07814 | NA | EPRS | glutamyl-prolyl-tRNA synthetase | 2058 | ENSG00000136628 |
| 20 | P17987 | NA | TCP1 | t-complex 1 | 6950 | ENSG00000120438 |
| 21 | P34932 | NA | HSPA4 | heat shock 70kDa protein 4 | 3308 | ENSG00000170606 |
| 22 | P22102 | NA | GART | phosphoribosylglycinamide formyltransferase, phosphoribosylglycinamide synthetase, phosphoribosylaminoimidazole synthetase | 2618 | ENSG00000159131 |
| 23 | P13639 | NA | EEF2 | eukaryotic translation elongation factor 2 | 1938 | ENSG00000167658 |
| 24 | P41091 | NA | EIF2S3 | eukaryotic translation initiation factor 2, subunit 3 gamma, 52kDa | 1968 | ENSG00000130741 |
| 25 | P40227 | NA | CCT6A | chaperonin containing TCP1, subunit 6A (zeta 1) | 908 | ENSG00000146731 |
| 26 | Q99832 | NA | CCT7 | chaperonin containing TCP1, subunit 7 (eta) | 10574 | ENSG00000135624 |
| 27 | P21980 | NA | TGM2 | transglutaminase 2 (C polypeptide, protein-glutamine-gamma-glutamyltransferase) | 7052 | ENSG00000198959 |
| 28 | P11586 | NA | MTHFD1 | methylenetetrahydrofolate dehydrogenase (NADP+ dependent) 1, methenyltetrahydrofolate cyclohydrolase, formyltetrahydrofolate synthetase | 4522 | ENSG00000100714 |
| 29 | P11387 | NA | TOP1 | topoisomerase (DNA) I | 7150 | ENSG00000198900 |

  
  

| **Database:molecular function      &nbspName:nucleoside binding      &nbspID:GO:0001882** | | | | | | |
| --- | --- | --- | --- | --- | --- | --- |
| C=1823; O=25; E=7.55; R=3.31; rawP=1.87e-08; adjP=2.09e-07 | | | | | | |
| Index | UserID | Value | Gene Symbol | Gene Name | EntrezGene | Ensembl |
| 1 | Q9NR30 | NA | DDX21 | DEAD (Asp-Glu-Ala-Asp) box helicase 21 | 9188 | ENSG00000165732 |
| 2 | P78371 | NA | CCT2 | chaperonin containing TCP1, subunit 2 (beta) | 10576 | ENSG00000166226 |
| 3 | Q92598 | NA | HSPH1 | heat shock 105kDa/110kDa protein 1 | 10808 | ENSG00000120694 |
| 4 | Q13885 | NA | TUBB2A | tubulin, beta 2A class IIa | 7280 | ENSG00000137267 |
| 5 | P00491 | NA | PNP | purine nucleoside phosphorylase | 4860 | ENSG00000198805 |
| 6 | P12081 | NA | HARS | histidyl-tRNA synthetase | 3035 | ENSG00000170445 |
| 7 | P68371 | NA | TUBB4B | tubulin, beta 4B class IVb | 10383 | ENSG00000188229 |
| 8 | P49368 | NA | CCT3 | chaperonin containing TCP1, subunit 3 (gamma) | 7203 | ENSG00000163468 |
| 9 | P56192 | NA | MARS | methionyl-tRNA synthetase | 4141 | ENSG00000166986 |
| 10 | Q9BQE3 | NA | TUBA1C | tubulin, alpha 1c | 84790 | ENSG00000167553 |
| 11 | P49736 | NA | MCM2 | minichromosome maintenance complex component 2 | 4171 | ENSG00000073111 |
| 12 | P50990 | NA | CCT8 | chaperonin containing TCP1, subunit 8 (theta) | 10694 | ENSG00000156261 |
| 13 | P68104 | NA | EEF1A1 | eukaryotic translation elongation factor 1 alpha 1 | 1915 | ENSG00000156508 |
| 14 | P50991 | NA | CCT4 | chaperonin containing TCP1, subunit 4 (delta) | 10575 | ENSG00000115484 |
| 15 | P07814 | NA | EPRS | glutamyl-prolyl-tRNA synthetase | 2058 | ENSG00000136628 |
| 16 | P17987 | NA | TCP1 | t-complex 1 | 6950 | ENSG00000120438 |
| 17 | P34932 | NA | HSPA4 | heat shock 70kDa protein 4 | 3308 | ENSG00000170606 |
| 18 | P22102 | NA | GART | phosphoribosylglycinamide formyltransferase, phosphoribosylglycinamide synthetase, phosphoribosylaminoimidazole synthetase | 2618 | ENSG00000159131 |
| 19 | P13639 | NA | EEF2 | eukaryotic translation elongation factor 2 | 1938 | ENSG00000167658 |
| 20 | P41091 | NA | EIF2S3 | eukaryotic translation initiation factor 2, subunit 3 gamma, 52kDa | 1968 | ENSG00000130741 |
| 21 | P40227 | NA | CCT6A | chaperonin containing TCP1, subunit 6A (zeta 1) | 908 | ENSG00000146731 |
| 22 | Q99832 | NA | CCT7 | chaperonin containing TCP1, subunit 7 (eta) | 10574 | ENSG00000135624 |
| 23 | P21980 | NA | TGM2 | transglutaminase 2 (C polypeptide, protein-glutamine-gamma-glutamyltransferase) | 7052 | ENSG00000198959 |
| 24 | P11586 | NA | MTHFD1 | methylenetetrahydrofolate dehydrogenase (NADP+ dependent) 1, methenyltetrahydrofolate cyclohydrolase, formyltetrahydrofolate synthetase | 4522 | ENSG00000100714 |
| 25 | P11387 | NA | TOP1 | topoisomerase (DNA) I | 7150 | ENSG00000198900 |

  
  

| **Database:molecular function      &nbspName:ribonucleoside binding      &nbspID:GO:0032549** | | | | | | |
| --- | --- | --- | --- | --- | --- | --- |
| C=1813; O=24; E=7.51; R=3.20; rawP=8.18e-08; adjP=5.48e-07 | | | | | | |
| Index | UserID | Value | Gene Symbol | Gene Name | EntrezGene | Ensembl |
| 1 | Q9NR30 | NA | DDX21 | DEAD (Asp-Glu-Ala-Asp) box helicase 21 | 9188 | ENSG00000165732 |
| 2 | P78371 | NA | CCT2 | chaperonin containing TCP1, subunit 2 (beta) | 10576 | ENSG00000166226 |
| 3 | Q92598 | NA | HSPH1 | heat shock 105kDa/110kDa protein 1 | 10808 | ENSG00000120694 |
| 4 | Q13885 | NA | TUBB2A | tubulin, beta 2A class IIa | 7280 | ENSG00000137267 |
| 5 | P12081 | NA | HARS | histidyl-tRNA synthetase | 3035 | ENSG00000170445 |
| 6 | P68371 | NA | TUBB4B | tubulin, beta 4B class IVb | 10383 | ENSG00000188229 |
| 7 | P49368 | NA | CCT3 | chaperonin containing TCP1, subunit 3 (gamma) | 7203 | ENSG00000163468 |
| 8 | P56192 | NA | MARS | methionyl-tRNA synthetase | 4141 | ENSG00000166986 |
| 9 | Q9BQE3 | NA | TUBA1C | tubulin, alpha 1c | 84790 | ENSG00000167553 |
| 10 | P49736 | NA | MCM2 | minichromosome maintenance complex component 2 | 4171 | ENSG00000073111 |
| 11 | P50990 | NA | CCT8 | chaperonin containing TCP1, subunit 8 (theta) | 10694 | ENSG00000156261 |
| 12 | P68104 | NA | EEF1A1 | eukaryotic translation elongation factor 1 alpha 1 | 1915 | ENSG00000156508 |
| 13 | P50991 | NA | CCT4 | chaperonin containing TCP1, subunit 4 (delta) | 10575 | ENSG00000115484 |
| 14 | P07814 | NA | EPRS | glutamyl-prolyl-tRNA synthetase | 2058 | ENSG00000136628 |
| 15 | P17987 | NA | TCP1 | t-complex 1 | 6950 | ENSG00000120438 |
| 16 | P34932 | NA | HSPA4 | heat shock 70kDa protein 4 | 3308 | ENSG00000170606 |
| 17 | P22102 | NA | GART | phosphoribosylglycinamide formyltransferase, phosphoribosylglycinamide synthetase, phosphoribosylaminoimidazole synthetase | 2618 | ENSG00000159131 |
| 18 | P13639 | NA | EEF2 | eukaryotic translation elongation factor 2 | 1938 | ENSG00000167658 |
| 19 | P41091 | NA | EIF2S3 | eukaryotic translation initiation factor 2, subunit 3 gamma, 52kDa | 1968 | ENSG00000130741 |
| 20 | P40227 | NA | CCT6A | chaperonin containing TCP1, subunit 6A (zeta 1) | 908 | ENSG00000146731 |
| 21 | Q99832 | NA | CCT7 | chaperonin containing TCP1, subunit 7 (eta) | 10574 | ENSG00000135624 |
| 22 | P21980 | NA | TGM2 | transglutaminase 2 (C polypeptide, protein-glutamine-gamma-glutamyltransferase) | 7052 | ENSG00000198959 |
| 23 | P11586 | NA | MTHFD1 | methylenetetrahydrofolate dehydrogenase (NADP+ dependent) 1, methenyltetrahydrofolate cyclohydrolase, formyltetrahydrofolate synthetase | 4522 | ENSG00000100714 |
| 24 | P11387 | NA | TOP1 | topoisomerase (DNA) I | 7150 | ENSG00000198900 |

  
  

| **Database:molecular function      &nbspName:purine nucleoside binding      &nbspID:GO:0001883** | | | | | | |
| --- | --- | --- | --- | --- | --- | --- |
| C=1812; O=24; E=7.50; R=3.20; rawP=8.10e-08; adjP=5.48e-07 | | | | | | |
| Index | UserID | Value | Gene Symbol | Gene Name | EntrezGene | Ensembl |
| 1 | Q9NR30 | NA | DDX21 | DEAD (Asp-Glu-Ala-Asp) box helicase 21 | 9188 | ENSG00000165732 |
| 2 | P78371 | NA | CCT2 | chaperonin containing TCP1, subunit 2 (beta) | 10576 | ENSG00000166226 |
| 3 | Q92598 | NA | HSPH1 | heat shock 105kDa/110kDa protein 1 | 10808 | ENSG00000120694 |
| 4 | Q13885 | NA | TUBB2A | tubulin, beta 2A class IIa | 7280 | ENSG00000137267 |
| 5 | P12081 | NA | HARS | histidyl-tRNA synthetase | 3035 | ENSG00000170445 |
| 6 | P68371 | NA | TUBB4B | tubulin, beta 4B class IVb | 10383 | ENSG00000188229 |
| 7 | P49368 | NA | CCT3 | chaperonin containing TCP1, subunit 3 (gamma) | 7203 | ENSG00000163468 |
| 8 | P56192 | NA | MARS | methionyl-tRNA synthetase | 4141 | ENSG00000166986 |
| 9 | Q9BQE3 | NA | TUBA1C | tubulin, alpha 1c | 84790 | ENSG00000167553 |
| 10 | P49736 | NA | MCM2 | minichromosome maintenance complex component 2 | 4171 | ENSG00000073111 |
| 11 | P50990 | NA | CCT8 | chaperonin containing TCP1, subunit 8 (theta) | 10694 | ENSG00000156261 |
| 12 | P68104 | NA | EEF1A1 | eukaryotic translation elongation factor 1 alpha 1 | 1915 | ENSG00000156508 |
| 13 | P50991 | NA | CCT4 | chaperonin containing TCP1, subunit 4 (delta) | 10575 | ENSG00000115484 |
| 14 | P07814 | NA | EPRS | glutamyl-prolyl-tRNA synthetase | 2058 | ENSG00000136628 |
| 15 | P17987 | NA | TCP1 | t-complex 1 | 6950 | ENSG00000120438 |
| 16 | P34932 | NA | HSPA4 | heat shock 70kDa protein 4 | 3308 | ENSG00000170606 |
| 17 | P22102 | NA | GART | phosphoribosylglycinamide formyltransferase, phosphoribosylglycinamide synthetase, phosphoribosylaminoimidazole synthetase | 2618 | ENSG00000159131 |
| 18 | P13639 | NA | EEF2 | eukaryotic translation elongation factor 2 | 1938 | ENSG00000167658 |
| 19 | P41091 | NA | EIF2S3 | eukaryotic translation initiation factor 2, subunit 3 gamma, 52kDa | 1968 | ENSG00000130741 |
| 20 | P40227 | NA | CCT6A | chaperonin containing TCP1, subunit 6A (zeta 1) | 908 | ENSG00000146731 |
| 21 | Q99832 | NA | CCT7 | chaperonin containing TCP1, subunit 7 (eta) | 10574 | ENSG00000135624 |
| 22 | P21980 | NA | TGM2 | transglutaminase 2 (C polypeptide, protein-glutamine-gamma-glutamyltransferase) | 7052 | ENSG00000198959 |
| 23 | P11586 | NA | MTHFD1 | methylenetetrahydrofolate dehydrogenase (NADP+ dependent) 1, methenyltetrahydrofolate cyclohydrolase, formyltetrahydrofolate synthetase | 4522 | ENSG00000100714 |
| 24 | P11387 | NA | TOP1 | topoisomerase (DNA) I | 7150 | ENSG00000198900 |

  
  

| **Database:molecular function      &nbspName:purine ribonucleoside binding      &nbspID:GO:0032550** | | | | | | |
| --- | --- | --- | --- | --- | --- | --- |
| C=1809; O=24; E=7.49; R=3.20; rawP=7.84e-08; adjP=5.48e-07 | | | | | | |
| Index | UserID | Value | Gene Symbol | Gene Name | EntrezGene | Ensembl |
| 1 | Q9NR30 | NA | DDX21 | DEAD (Asp-Glu-Ala-Asp) box helicase 21 | 9188 | ENSG00000165732 |
| 2 | P78371 | NA | CCT2 | chaperonin containing TCP1, subunit 2 (beta) | 10576 | ENSG00000166226 |
| 3 | Q92598 | NA | HSPH1 | heat shock 105kDa/110kDa protein 1 | 10808 | ENSG00000120694 |
| 4 | Q13885 | NA | TUBB2A | tubulin, beta 2A class IIa | 7280 | ENSG00000137267 |
| 5 | P12081 | NA | HARS | histidyl-tRNA synthetase | 3035 | ENSG00000170445 |
| 6 | P68371 | NA | TUBB4B | tubulin, beta 4B class IVb | 10383 | ENSG00000188229 |
| 7 | P49368 | NA | CCT3 | chaperonin containing TCP1, subunit 3 (gamma) | 7203 | ENSG00000163468 |
| 8 | P56192 | NA | MARS | methionyl-tRNA synthetase | 4141 | ENSG00000166986 |
| 9 | Q9BQE3 | NA | TUBA1C | tubulin, alpha 1c | 84790 | ENSG00000167553 |
| 10 | P49736 | NA | MCM2 | minichromosome maintenance complex component 2 | 4171 | ENSG00000073111 |
| 11 | P50990 | NA | CCT8 | chaperonin containing TCP1, subunit 8 (theta) | 10694 | ENSG00000156261 |
| 12 | P68104 | NA | EEF1A1 | eukaryotic translation elongation factor 1 alpha 1 | 1915 | ENSG00000156508 |
| 13 | P50991 | NA | CCT4 | chaperonin containing TCP1, subunit 4 (delta) | 10575 | ENSG00000115484 |
| 14 | P07814 | NA | EPRS | glutamyl-prolyl-tRNA synthetase | 2058 | ENSG00000136628 |
| 15 | P17987 | NA | TCP1 | t-complex 1 | 6950 | ENSG00000120438 |
| 16 | P34932 | NA | HSPA4 | heat shock 70kDa protein 4 | 3308 | ENSG00000170606 |
| 17 | P22102 | NA | GART | phosphoribosylglycinamide formyltransferase, phosphoribosylglycinamide synthetase, phosphoribosylaminoimidazole synthetase | 2618 | ENSG00000159131 |
| 18 | P13639 | NA | EEF2 | eukaryotic translation elongation factor 2 | 1938 | ENSG00000167658 |
| 19 | P41091 | NA | EIF2S3 | eukaryotic translation initiation factor 2, subunit 3 gamma, 52kDa | 1968 | ENSG00000130741 |
| 20 | P40227 | NA | CCT6A | chaperonin containing TCP1, subunit 6A (zeta 1) | 908 | ENSG00000146731 |
| 21 | Q99832 | NA | CCT7 | chaperonin containing TCP1, subunit 7 (eta) | 10574 | ENSG00000135624 |
| 22 | P21980 | NA | TGM2 | transglutaminase 2 (C polypeptide, protein-glutamine-gamma-glutamyltransferase) | 7052 | ENSG00000198959 |
| 23 | P11586 | NA | MTHFD1 | methylenetetrahydrofolate dehydrogenase (NADP+ dependent) 1, methenyltetrahydrofolate cyclohydrolase, formyltetrahydrofolate synthetase | 4522 | ENSG00000100714 |
| 24 | P11387 | NA | TOP1 | topoisomerase (DNA) I | 7150 | ENSG00000198900 |

  
  

| **Database:molecular function      &nbspName:purine ribonucleoside triphosphate binding      &nbspID:GO:0035639** | | | | | | |
| --- | --- | --- | --- | --- | --- | --- |
| C=1800; O=24; E=7.45; R=3.22; rawP=7.13e-08; adjP=5.48e-07 | | | | | | |
| Index | UserID | Value | Gene Symbol | Gene Name | EntrezGene | Ensembl |
| 1 | Q9NR30 | NA | DDX21 | DEAD (Asp-Glu-Ala-Asp) box helicase 21 | 9188 | ENSG00000165732 |
| 2 | P78371 | NA | CCT2 | chaperonin containing TCP1, subunit 2 (beta) | 10576 | ENSG00000166226 |
| 3 | Q92598 | NA | HSPH1 | heat shock 105kDa/110kDa protein 1 | 10808 | ENSG00000120694 |
| 4 | Q13885 | NA | TUBB2A | tubulin, beta 2A class IIa | 7280 | ENSG00000137267 |
| 5 | P12081 | NA | HARS | histidyl-tRNA synthetase | 3035 | ENSG00000170445 |
| 6 | P68371 | NA | TUBB4B | tubulin, beta 4B class IVb | 10383 | ENSG00000188229 |
| 7 | P49368 | NA | CCT3 | chaperonin containing TCP1, subunit 3 (gamma) | 7203 | ENSG00000163468 |
| 8 | P56192 | NA | MARS | methionyl-tRNA synthetase | 4141 | ENSG00000166986 |
| 9 | Q9BQE3 | NA | TUBA1C | tubulin, alpha 1c | 84790 | ENSG00000167553 |
| 10 | P49736 | NA | MCM2 | minichromosome maintenance complex component 2 | 4171 | ENSG00000073111 |
| 11 | P50990 | NA | CCT8 | chaperonin containing TCP1, subunit 8 (theta) | 10694 | ENSG00000156261 |
| 12 | P68104 | NA | EEF1A1 | eukaryotic translation elongation factor 1 alpha 1 | 1915 | ENSG00000156508 |
| 13 | P50991 | NA | CCT4 | chaperonin containing TCP1, subunit 4 (delta) | 10575 | ENSG00000115484 |
| 14 | P07814 | NA | EPRS | glutamyl-prolyl-tRNA synthetase | 2058 | ENSG00000136628 |
| 15 | P17987 | NA | TCP1 | t-complex 1 | 6950 | ENSG00000120438 |
| 16 | P34932 | NA | HSPA4 | heat shock 70kDa protein 4 | 3308 | ENSG00000170606 |
| 17 | P22102 | NA | GART | phosphoribosylglycinamide formyltransferase, phosphoribosylglycinamide synthetase, phosphoribosylaminoimidazole synthetase | 2618 | ENSG00000159131 |
| 18 | P13639 | NA | EEF2 | eukaryotic translation elongation factor 2 | 1938 | ENSG00000167658 |
| 19 | P41091 | NA | EIF2S3 | eukaryotic translation initiation factor 2, subunit 3 gamma, 52kDa | 1968 | ENSG00000130741 |
| 20 | P40227 | NA | CCT6A | chaperonin containing TCP1, subunit 6A (zeta 1) | 908 | ENSG00000146731 |
| 21 | Q99832 | NA | CCT7 | chaperonin containing TCP1, subunit 7 (eta) | 10574 | ENSG00000135624 |
| 22 | P21980 | NA | TGM2 | transglutaminase 2 (C polypeptide, protein-glutamine-gamma-glutamyltransferase) | 7052 | ENSG00000198959 |
| 23 | P11586 | NA | MTHFD1 | methylenetetrahydrofolate dehydrogenase (NADP+ dependent) 1, methenyltetrahydrofolate cyclohydrolase, formyltetrahydrofolate synthetase | 4522 | ENSG00000100714 |
| 24 | P11387 | NA | TOP1 | topoisomerase (DNA) I | 7150 | ENSG00000198900 |

  
  

| **Database:molecular function      &nbspName:purine nucleotide binding      &nbspID:GO:0017076** | | | | | | |
| --- | --- | --- | --- | --- | --- | --- |
| C=1842; O=24; E=7.63; R=3.15; rawP=1.11e-07; adjP=6.18e-07 | | | | | | |
| Index | UserID | Value | Gene Symbol | Gene Name | EntrezGene | Ensembl |
| 1 | Q9NR30 | NA | DDX21 | DEAD (Asp-Glu-Ala-Asp) box helicase 21 | 9188 | ENSG00000165732 |
| 2 | P78371 | NA | CCT2 | chaperonin containing TCP1, subunit 2 (beta) | 10576 | ENSG00000166226 |
| 3 | Q92598 | NA | HSPH1 | heat shock 105kDa/110kDa protein 1 | 10808 | ENSG00000120694 |
| 4 | Q13885 | NA | TUBB2A | tubulin, beta 2A class IIa | 7280 | ENSG00000137267 |
| 5 | P12081 | NA | HARS | histidyl-tRNA synthetase | 3035 | ENSG00000170445 |
| 6 | P68371 | NA | TUBB4B | tubulin, beta 4B class IVb | 10383 | ENSG00000188229 |
| 7 | P49368 | NA | CCT3 | chaperonin containing TCP1, subunit 3 (gamma) | 7203 | ENSG00000163468 |
| 8 | P56192 | NA | MARS | methionyl-tRNA synthetase | 4141 | ENSG00000166986 |
| 9 | Q9BQE3 | NA | TUBA1C | tubulin, alpha 1c | 84790 | ENSG00000167553 |
| 10 | P49736 | NA | MCM2 | minichromosome maintenance complex component 2 | 4171 | ENSG00000073111 |
| 11 | P50990 | NA | CCT8 | chaperonin containing TCP1, subunit 8 (theta) | 10694 | ENSG00000156261 |
| 12 | P68104 | NA | EEF1A1 | eukaryotic translation elongation factor 1 alpha 1 | 1915 | ENSG00000156508 |
| 13 | P50991 | NA | CCT4 | chaperonin containing TCP1, subunit 4 (delta) | 10575 | ENSG00000115484 |
| 14 | P07814 | NA | EPRS | glutamyl-prolyl-tRNA synthetase | 2058 | ENSG00000136628 |
| 15 | P17987 | NA | TCP1 | t-complex 1 | 6950 | ENSG00000120438 |
| 16 | P34932 | NA | HSPA4 | heat shock 70kDa protein 4 | 3308 | ENSG00000170606 |
| 17 | P22102 | NA | GART | phosphoribosylglycinamide formyltransferase, phosphoribosylglycinamide synthetase, phosphoribosylaminoimidazole synthetase | 2618 | ENSG00000159131 |
| 18 | P13639 | NA | EEF2 | eukaryotic translation elongation factor 2 | 1938 | ENSG00000167658 |
| 19 | P41091 | NA | EIF2S3 | eukaryotic translation initiation factor 2, subunit 3 gamma, 52kDa | 1968 | ENSG00000130741 |
| 20 | P40227 | NA | CCT6A | chaperonin containing TCP1, subunit 6A (zeta 1) | 908 | ENSG00000146731 |
| 21 | Q99832 | NA | CCT7 | chaperonin containing TCP1, subunit 7 (eta) | 10574 | ENSG00000135624 |
| 22 | P21980 | NA | TGM2 | transglutaminase 2 (C polypeptide, protein-glutamine-gamma-glutamyltransferase) | 7052 | ENSG00000198959 |
| 23 | P11586 | NA | MTHFD1 | methylenetetrahydrofolate dehydrogenase (NADP+ dependent) 1, methenyltetrahydrofolate cyclohydrolase, formyltetrahydrofolate synthetase | 4522 | ENSG00000100714 |
| 24 | P11387 | NA | TOP1 | topoisomerase (DNA) I | 7150 | ENSG00000198900 |

  
  

| **Database:molecular function      &nbspName:ribonucleotide binding      &nbspID:GO:0032553** | | | | | | |
| --- | --- | --- | --- | --- | --- | --- |
| C=1850; O=24; E=7.66; R=3.13; rawP=1.20e-07; adjP=6.18e-07 | | | | | | |
| Index | UserID | Value | Gene Symbol | Gene Name | EntrezGene | Ensembl |
| 1 | Q9NR30 | NA | DDX21 | DEAD (Asp-Glu-Ala-Asp) box helicase 21 | 9188 | ENSG00000165732 |
| 2 | P78371 | NA | CCT2 | chaperonin containing TCP1, subunit 2 (beta) | 10576 | ENSG00000166226 |
| 3 | Q92598 | NA | HSPH1 | heat shock 105kDa/110kDa protein 1 | 10808 | ENSG00000120694 |
| 4 | Q13885 | NA | TUBB2A | tubulin, beta 2A class IIa | 7280 | ENSG00000137267 |
| 5 | P12081 | NA | HARS | histidyl-tRNA synthetase | 3035 | ENSG00000170445 |
| 6 | P68371 | NA | TUBB4B | tubulin, beta 4B class IVb | 10383 | ENSG00000188229 |
| 7 | P49368 | NA | CCT3 | chaperonin containing TCP1, subunit 3 (gamma) | 7203 | ENSG00000163468 |
| 8 | P56192 | NA | MARS | methionyl-tRNA synthetase | 4141 | ENSG00000166986 |
| 9 | Q9BQE3 | NA | TUBA1C | tubulin, alpha 1c | 84790 | ENSG00000167553 |
| 10 | P49736 | NA | MCM2 | minichromosome maintenance complex component 2 | 4171 | ENSG00000073111 |
| 11 | P50990 | NA | CCT8 | chaperonin containing TCP1, subunit 8 (theta) | 10694 | ENSG00000156261 |
| 12 | P68104 | NA | EEF1A1 | eukaryotic translation elongation factor 1 alpha 1 | 1915 | ENSG00000156508 |
| 13 | P50991 | NA | CCT4 | chaperonin containing TCP1, subunit 4 (delta) | 10575 | ENSG00000115484 |
| 14 | P07814 | NA | EPRS | glutamyl-prolyl-tRNA synthetase | 2058 | ENSG00000136628 |
| 15 | P17987 | NA | TCP1 | t-complex 1 | 6950 | ENSG00000120438 |
| 16 | P34932 | NA | HSPA4 | heat shock 70kDa protein 4 | 3308 | ENSG00000170606 |
| 17 | P22102 | NA | GART | phosphoribosylglycinamide formyltransferase, phosphoribosylglycinamide synthetase, phosphoribosylaminoimidazole synthetase | 2618 | ENSG00000159131 |
| 18 | P13639 | NA | EEF2 | eukaryotic translation elongation factor 2 | 1938 | ENSG00000167658 |
| 19 | P41091 | NA | EIF2S3 | eukaryotic translation initiation factor 2, subunit 3 gamma, 52kDa | 1968 | ENSG00000130741 |
| 20 | P40227 | NA | CCT6A | chaperonin containing TCP1, subunit 6A (zeta 1) | 908 | ENSG00000146731 |
| 21 | Q99832 | NA | CCT7 | chaperonin containing TCP1, subunit 7 (eta) | 10574 | ENSG00000135624 |
| 22 | P21980 | NA | TGM2 | transglutaminase 2 (C polypeptide, protein-glutamine-gamma-glutamyltransferase) | 7052 | ENSG00000198959 |
| 23 | P11586 | NA | MTHFD1 | methylenetetrahydrofolate dehydrogenase (NADP+ dependent) 1, methenyltetrahydrofolate cyclohydrolase, formyltetrahydrofolate synthetase | 4522 | ENSG00000100714 |
| 24 | P11387 | NA | TOP1 | topoisomerase (DNA) I | 7150 | ENSG00000198900 |

  
  

| **Database:molecular function      &nbspName:purine ribonucleotide binding      &nbspID:GO:0032555** | | | | | | |
| --- | --- | --- | --- | --- | --- | --- |
| C=1835; O=24; E=7.60; R=3.16; rawP=1.03e-07; adjP=6.18e-07 | | | | | | |
| Index | UserID | Value | Gene Symbol | Gene Name | EntrezGene | Ensembl |
| 1 | Q9NR30 | NA | DDX21 | DEAD (Asp-Glu-Ala-Asp) box helicase 21 | 9188 | ENSG00000165732 |
| 2 | P78371 | NA | CCT2 | chaperonin containing TCP1, subunit 2 (beta) | 10576 | ENSG00000166226 |
| 3 | Q92598 | NA | HSPH1 | heat shock 105kDa/110kDa protein 1 | 10808 | ENSG00000120694 |
| 4 | Q13885 | NA | TUBB2A | tubulin, beta 2A class IIa | 7280 | ENSG00000137267 |
| 5 | P12081 | NA | HARS | histidyl-tRNA synthetase | 3035 | ENSG00000170445 |
| 6 | P68371 | NA | TUBB4B | tubulin, beta 4B class IVb | 10383 | ENSG00000188229 |
| 7 | P49368 | NA | CCT3 | chaperonin containing TCP1, subunit 3 (gamma) | 7203 | ENSG00000163468 |
| 8 | P56192 | NA | MARS | methionyl-tRNA synthetase | 4141 | ENSG00000166986 |
| 9 | Q9BQE3 | NA | TUBA1C | tubulin, alpha 1c | 84790 | ENSG00000167553 |
| 10 | P49736 | NA | MCM2 | minichromosome maintenance complex component 2 | 4171 | ENSG00000073111 |
| 11 | P50990 | NA | CCT8 | chaperonin containing TCP1, subunit 8 (theta) | 10694 | ENSG00000156261 |
| 12 | P68104 | NA | EEF1A1 | eukaryotic translation elongation factor 1 alpha 1 | 1915 | ENSG00000156508 |
| 13 | P50991 | NA | CCT4 | chaperonin containing TCP1, subunit 4 (delta) | 10575 | ENSG00000115484 |
| 14 | P07814 | NA | EPRS | glutamyl-prolyl-tRNA synthetase | 2058 | ENSG00000136628 |
| 15 | P17987 | NA | TCP1 | t-complex 1 | 6950 | ENSG00000120438 |
| 16 | P34932 | NA | HSPA4 | heat shock 70kDa protein 4 | 3308 | ENSG00000170606 |
| 17 | P22102 | NA | GART | phosphoribosylglycinamide formyltransferase, phosphoribosylglycinamide synthetase, phosphoribosylaminoimidazole synthetase | 2618 | ENSG00000159131 |
| 18 | P13639 | NA | EEF2 | eukaryotic translation elongation factor 2 | 1938 | ENSG00000167658 |
| 19 | P41091 | NA | EIF2S3 | eukaryotic translation initiation factor 2, subunit 3 gamma, 52kDa | 1968 | ENSG00000130741 |
| 20 | P40227 | NA | CCT6A | chaperonin containing TCP1, subunit 6A (zeta 1) | 908 | ENSG00000146731 |
| 21 | Q99832 | NA | CCT7 | chaperonin containing TCP1, subunit 7 (eta) | 10574 | ENSG00000135624 |
| 22 | P21980 | NA | TGM2 | transglutaminase 2 (C polypeptide, protein-glutamine-gamma-glutamyltransferase) | 7052 | ENSG00000198959 |
| 23 | P11586 | NA | MTHFD1 | methylenetetrahydrofolate dehydrogenase (NADP+ dependent) 1, methenyltetrahydrofolate cyclohydrolase, formyltetrahydrofolate synthetase | 4522 | ENSG00000100714 |
| 24 | P11387 | NA | TOP1 | topoisomerase (DNA) I | 7150 | ENSG00000198900 |

  
  

| **Database:molecular function      &nbspName:translation factor activity, nucleic acid binding      &nbspID:GO:0008135** | | | | | | |
| --- | --- | --- | --- | --- | --- | --- |
| C=79; O=6; E=0.33; R=18.35; rawP=8.63e-07; adjP=4.13e-06 | | | | | | |
| Index | UserID | Value | Gene Symbol | Gene Name | EntrezGene | Ensembl |
| 1 | P63241 | NA | EIF5A | eukaryotic translation initiation factor 5A | 1984 | ENSG00000132507 |
| 2 | P13639 | NA | EEF2 | eukaryotic translation elongation factor 2 | 1938 | ENSG00000167658 |
| 3 | P41091 | NA | EIF2S3 | eukaryotic translation initiation factor 2, subunit 3 gamma, 52kDa | 1968 | ENSG00000130741 |
| 4 | Q15056 | NA | EIF4H | eukaryotic translation initiation factor 4H | 7458 | ENSG00000106682 |
| 5 | P68104 | NA | EEF1A1 | eukaryotic translation elongation factor 1 alpha 1 | 1915 | ENSG00000156508 |
| 6 | P41567 | NA | EIF1 | eukaryotic translation initiation factor 1 | 10209 | ENSG00000173812 |

  
  

| **Database:molecular function      &nbspName:RNA binding      &nbspID:GO:0003723** | | | | | | |
| --- | --- | --- | --- | --- | --- | --- |
| C=843; O=13; E=3.49; R=3.73; rawP=3.30e-05; adjP=0.0001 | | | | | | |
| Index | UserID | Value | Gene Symbol | Gene Name | EntrezGene | Ensembl |
| 1 | P63241 | NA | EIF5A | eukaryotic translation initiation factor 5A | 1984 | ENSG00000132507 |
| 2 | P07814 | NA | EPRS | glutamyl-prolyl-tRNA synthetase | 2058 | ENSG00000136628 |
| 3 | Q9NR30 | NA | DDX21 | DEAD (Asp-Glu-Ala-Asp) box helicase 21 | 9188 | ENSG00000165732 |
| 4 | P13639 | NA | EEF2 | eukaryotic translation elongation factor 2 | 1938 | ENSG00000167658 |
| 5 | P41091 | NA | EIF2S3 | eukaryotic translation initiation factor 2, subunit 3 gamma, 52kDa | 1968 | ENSG00000130741 |
| 6 | P41567 | NA | EIF1 | eukaryotic translation initiation factor 1 | 10209 | ENSG00000173812 |
| 7 | P67809 | NA | YBX1 | Y box binding protein 1 | 4904 | ENSG00000065978 |
| 8 | P56192 | NA | MARS | methionyl-tRNA synthetase | 4141 | ENSG00000166986 |
| 9 | P12268 | NA | IMPDH2 | IMP (inosine 5'-monophosphate) dehydrogenase 2 | 3615 | ENSG00000178035 |
| 10 | P62906 | NA | RPL10A | ribosomal protein L10a | 4736 | ENSG00000198755 |
| 11 | Q15056 | NA | EIF4H | eukaryotic translation initiation factor 4H | 7458 | ENSG00000106682 |
| 12 | P68104 | NA | EEF1A1 | eukaryotic translation elongation factor 1 alpha 1 | 1915 | ENSG00000156508 |
| 13 | O00425 | NA | IGF2BP3 | insulin-like growth factor 2 mRNA binding protein 3 | 10643 | ENSG00000136231 |

  
  

| **Database:molecular function      &nbspName:ATP binding      &nbspID:GO:0005524** | | | | | | |
| --- | --- | --- | --- | --- | --- | --- |
| C=1462; O=17; E=6.05; R=2.81; rawP=6.28e-05; adjP=0.0002 | | | | | | |
| Index | UserID | Value | Gene Symbol | Gene Name | EntrezGene | Ensembl |
| 1 | Q9NR30 | NA | DDX21 | DEAD (Asp-Glu-Ala-Asp) box helicase 21 | 9188 | ENSG00000165732 |
| 2 | P78371 | NA | CCT2 | chaperonin containing TCP1, subunit 2 (beta) | 10576 | ENSG00000166226 |
| 3 | Q92598 | NA | HSPH1 | heat shock 105kDa/110kDa protein 1 | 10808 | ENSG00000120694 |
| 4 | P12081 | NA | HARS | histidyl-tRNA synthetase | 3035 | ENSG00000170445 |
| 5 | P49368 | NA | CCT3 | chaperonin containing TCP1, subunit 3 (gamma) | 7203 | ENSG00000163468 |
| 6 | P56192 | NA | MARS | methionyl-tRNA synthetase | 4141 | ENSG00000166986 |
| 7 | P49736 | NA | MCM2 | minichromosome maintenance complex component 2 | 4171 | ENSG00000073111 |
| 8 | P50990 | NA | CCT8 | chaperonin containing TCP1, subunit 8 (theta) | 10694 | ENSG00000156261 |
| 9 | P50991 | NA | CCT4 | chaperonin containing TCP1, subunit 4 (delta) | 10575 | ENSG00000115484 |
| 10 | P07814 | NA | EPRS | glutamyl-prolyl-tRNA synthetase | 2058 | ENSG00000136628 |
| 11 | P22102 | NA | GART | phosphoribosylglycinamide formyltransferase, phosphoribosylglycinamide synthetase, phosphoribosylaminoimidazole synthetase | 2618 | ENSG00000159131 |
| 12 | P34932 | NA | HSPA4 | heat shock 70kDa protein 4 | 3308 | ENSG00000170606 |
| 13 | P17987 | NA | TCP1 | t-complex 1 | 6950 | ENSG00000120438 |
| 14 | P40227 | NA | CCT6A | chaperonin containing TCP1, subunit 6A (zeta 1) | 908 | ENSG00000146731 |
| 15 | Q99832 | NA | CCT7 | chaperonin containing TCP1, subunit 7 (eta) | 10574 | ENSG00000135624 |
| 16 | P11387 | NA | TOP1 | topoisomerase (DNA) I | 7150 | ENSG00000198900 |
| 17 | P11586 | NA | MTHFD1 | methylenetetrahydrofolate dehydrogenase (NADP+ dependent) 1, methenyltetrahydrofolate cyclohydrolase, formyltetrahydrofolate synthetase | 4522 | ENSG00000100714 |

  
  

| **Database:molecular function      &nbspName:translation elongation factor activity      &nbspID:GO:0003746** | | | | | | |
| --- | --- | --- | --- | --- | --- | --- |
| C=17; O=3; E=0.07; R=42.63; rawP=4.41e-05; adjP=0.0002 | | | | | | |
| Index | UserID | Value | Gene Symbol | Gene Name | EntrezGene | Ensembl |
| 1 | P63241 | NA | EIF5A | eukaryotic translation initiation factor 5A | 1984 | ENSG00000132507 |
| 2 | P13639 | NA | EEF2 | eukaryotic translation elongation factor 2 | 1938 | ENSG00000167658 |
| 3 | P68104 | NA | EEF1A1 | eukaryotic translation elongation factor 1 alpha 1 | 1915 | ENSG00000156508 |

  
  

| **Database:molecular function      &nbspName:adenyl nucleotide binding      &nbspID:GO:0030554** | | | | | | |
| --- | --- | --- | --- | --- | --- | --- |
| C=1496; O=17; E=6.19; R=2.75; rawP=8.37e-05; adjP=0.0003 | | | | | | |
| Index | UserID | Value | Gene Symbol | Gene Name | EntrezGene | Ensembl |
| 1 | Q9NR30 | NA | DDX21 | DEAD (Asp-Glu-Ala-Asp) box helicase 21 | 9188 | ENSG00000165732 |
| 2 | P78371 | NA | CCT2 | chaperonin containing TCP1, subunit 2 (beta) | 10576 | ENSG00000166226 |
| 3 | Q92598 | NA | HSPH1 | heat shock 105kDa/110kDa protein 1 | 10808 | ENSG00000120694 |
| 4 | P12081 | NA | HARS | histidyl-tRNA synthetase | 3035 | ENSG00000170445 |
| 5 | P49368 | NA | CCT3 | chaperonin containing TCP1, subunit 3 (gamma) | 7203 | ENSG00000163468 |
| 6 | P56192 | NA | MARS | methionyl-tRNA synthetase | 4141 | ENSG00000166986 |
| 7 | P49736 | NA | MCM2 | minichromosome maintenance complex component 2 | 4171 | ENSG00000073111 |
| 8 | P50990 | NA | CCT8 | chaperonin containing TCP1, subunit 8 (theta) | 10694 | ENSG00000156261 |
| 9 | P50991 | NA | CCT4 | chaperonin containing TCP1, subunit 4 (delta) | 10575 | ENSG00000115484 |
| 10 | P07814 | NA | EPRS | glutamyl-prolyl-tRNA synthetase | 2058 | ENSG00000136628 |
| 11 | P22102 | NA | GART | phosphoribosylglycinamide formyltransferase, phosphoribosylglycinamide synthetase, phosphoribosylaminoimidazole synthetase | 2618 | ENSG00000159131 |
| 12 | P34932 | NA | HSPA4 | heat shock 70kDa protein 4 | 3308 | ENSG00000170606 |
| 13 | P17987 | NA | TCP1 | t-complex 1 | 6950 | ENSG00000120438 |
| 14 | P40227 | NA | CCT6A | chaperonin containing TCP1, subunit 6A (zeta 1) | 908 | ENSG00000146731 |
| 15 | Q99832 | NA | CCT7 | chaperonin containing TCP1, subunit 7 (eta) | 10574 | ENSG00000135624 |
| 16 | P11387 | NA | TOP1 | topoisomerase (DNA) I | 7150 | ENSG00000198900 |
| 17 | P11586 | NA | MTHFD1 | methylenetetrahydrofolate dehydrogenase (NADP+ dependent) 1, methenyltetrahydrofolate cyclohydrolase, formyltetrahydrofolate synthetase | 4522 | ENSG00000100714 |

  
  

| **Database:molecular function      &nbspName:adenyl ribonucleotide binding      &nbspID:GO:0032559** | | | | | | |
| --- | --- | --- | --- | --- | --- | --- |
| C=1492; O=17; E=6.18; R=2.75; rawP=8.10e-05; adjP=0.0003 | | | | | | |
| Index | UserID | Value | Gene Symbol | Gene Name | EntrezGene | Ensembl |
| 1 | Q9NR30 | NA | DDX21 | DEAD (Asp-Glu-Ala-Asp) box helicase 21 | 9188 | ENSG00000165732 |
| 2 | P78371 | NA | CCT2 | chaperonin containing TCP1, subunit 2 (beta) | 10576 | ENSG00000166226 |
| 3 | Q92598 | NA | HSPH1 | heat shock 105kDa/110kDa protein 1 | 10808 | ENSG00000120694 |
| 4 | P12081 | NA | HARS | histidyl-tRNA synthetase | 3035 | ENSG00000170445 |
| 5 | P49368 | NA | CCT3 | chaperonin containing TCP1, subunit 3 (gamma) | 7203 | ENSG00000163468 |
| 6 | P56192 | NA | MARS | methionyl-tRNA synthetase | 4141 | ENSG00000166986 |
| 7 | P49736 | NA | MCM2 | minichromosome maintenance complex component 2 | 4171 | ENSG00000073111 |
| 8 | P50990 | NA | CCT8 | chaperonin containing TCP1, subunit 8 (theta) | 10694 | ENSG00000156261 |
| 9 | P50991 | NA | CCT4 | chaperonin containing TCP1, subunit 4 (delta) | 10575 | ENSG00000115484 |
| 10 | P07814 | NA | EPRS | glutamyl-prolyl-tRNA synthetase | 2058 | ENSG00000136628 |
| 11 | P22102 | NA | GART | phosphoribosylglycinamide formyltransferase, phosphoribosylglycinamide synthetase, phosphoribosylaminoimidazole synthetase | 2618 | ENSG00000159131 |
| 12 | P34932 | NA | HSPA4 | heat shock 70kDa protein 4 | 3308 | ENSG00000170606 |
| 13 | P17987 | NA | TCP1 | t-complex 1 | 6950 | ENSG00000120438 |
| 14 | P40227 | NA | CCT6A | chaperonin containing TCP1, subunit 6A (zeta 1) | 908 | ENSG00000146731 |
| 15 | Q99832 | NA | CCT7 | chaperonin containing TCP1, subunit 7 (eta) | 10574 | ENSG00000135624 |
| 16 | P11387 | NA | TOP1 | topoisomerase (DNA) I | 7150 | ENSG00000198900 |
| 17 | P11586 | NA | MTHFD1 | methylenetetrahydrofolate dehydrogenase (NADP+ dependent) 1, methenyltetrahydrofolate cyclohydrolase, formyltetrahydrofolate synthetase | 4522 | ENSG00000100714 |

  
  

| **Database:molecular function      &nbspName:organic cyclic compound binding      &nbspID:GO:0097159** | | | | | | |
| --- | --- | --- | --- | --- | --- | --- |
| C=5342; O=36; E=22.11; R=1.63; rawP=0.0003; adjP=0.0010 | | | | | | |
| Index | UserID | Value | Gene Symbol | Gene Name | EntrezGene | Ensembl |
| 1 | Q92598 | NA | HSPH1 | heat shock 105kDa/110kDa protein 1 | 10808 | ENSG00000120694 |
| 2 | Q13885 | NA | TUBB2A | tubulin, beta 2A class IIa | 7280 | ENSG00000137267 |
| 3 | P68371 | NA | TUBB4B | tubulin, beta 4B class IVb | 10383 | ENSG00000188229 |
| 4 | P68104 | NA | EEF1A1 | eukaryotic translation elongation factor 1 alpha 1 | 1915 | ENSG00000156508 |
| 5 | P50991 | NA | CCT4 | chaperonin containing TCP1, subunit 4 (delta) | 10575 | ENSG00000115484 |
| 6 | O00425 | NA | IGF2BP3 | insulin-like growth factor 2 mRNA binding protein 3 | 10643 | ENSG00000136231 |
| 7 | P22102 | NA | GART | phosphoribosylglycinamide formyltransferase, phosphoribosylglycinamide synthetase, phosphoribosylaminoimidazole synthetase | 2618 | ENSG00000159131 |
| 8 | P13639 | NA | EEF2 | eukaryotic translation elongation factor 2 | 1938 | ENSG00000167658 |
| 9 | P41567 | NA | EIF1 | eukaryotic translation initiation factor 1 | 10209 | ENSG00000173812 |
| 10 | P17174 | NA | GOT1 | glutamic-oxaloacetic transaminase 1, soluble (aspartate aminotransferase 1) | 2805 | ENSG00000120053 |
| 11 | P67809 | NA | YBX1 | Y box binding protein 1 | 4904 | ENSG00000065978 |
| 12 | Q99832 | NA | CCT7 | chaperonin containing TCP1, subunit 7 (eta) | 10574 | ENSG00000135624 |
| 13 | P21980 | NA | TGM2 | transglutaminase 2 (C polypeptide, protein-glutamine-gamma-glutamyltransferase) | 7052 | ENSG00000198959 |
| 14 | P11586 | NA | MTHFD1 | methylenetetrahydrofolate dehydrogenase (NADP+ dependent) 1, methenyltetrahydrofolate cyclohydrolase, formyltetrahydrofolate synthetase | 4522 | ENSG00000100714 |
| 15 | P11387 | NA | TOP1 | topoisomerase (DNA) I | 7150 | ENSG00000198900 |
| 16 | Q16881 | NA | TXNRD1 | thioredoxin reductase 1 | 7296 | ENSG00000198431 |
| 17 | P63241 | NA | EIF5A | eukaryotic translation initiation factor 5A | 1984 | ENSG00000132507 |
| 18 | P78371 | NA | CCT2 | chaperonin containing TCP1, subunit 2 (beta) | 10576 | ENSG00000166226 |
| 19 | Q9NR30 | NA | DDX21 | DEAD (Asp-Glu-Ala-Asp) box helicase 21 | 9188 | ENSG00000165732 |
| 20 | P00491 | NA | PNP | purine nucleoside phosphorylase | 4860 | ENSG00000198805 |
| 21 | P12081 | NA | HARS | histidyl-tRNA synthetase | 3035 | ENSG00000170445 |
| 22 | P49327 | NA | FASN | fatty acid synthase | 2194 | ENSG00000169710 |
| 23 | P49368 | NA | CCT3 | chaperonin containing TCP1, subunit 3 (gamma) | 7203 | ENSG00000163468 |
| 24 | Q9BQE3 | NA | TUBA1C | tubulin, alpha 1c | 84790 | ENSG00000167553 |
| 25 | P56192 | NA | MARS | methionyl-tRNA synthetase | 4141 | ENSG00000166986 |
| 26 | P12268 | NA | IMPDH2 | IMP (inosine 5'-monophosphate) dehydrogenase 2 | 3615 | ENSG00000178035 |
| 27 | P49736 | NA | MCM2 | minichromosome maintenance complex component 2 | 4171 | ENSG00000073111 |
| 28 | P62906 | NA | RPL10A | ribosomal protein L10a | 4736 | ENSG00000198755 |
| 29 | Q15056 | NA | EIF4H | eukaryotic translation initiation factor 4H | 7458 | ENSG00000106682 |
| 30 | P50990 | NA | CCT8 | chaperonin containing TCP1, subunit 8 (theta) | 10694 | ENSG00000156261 |
| 31 | P07814 | NA | EPRS | glutamyl-prolyl-tRNA synthetase | 2058 | ENSG00000136628 |
| 32 | P17987 | NA | TCP1 | t-complex 1 | 6950 | ENSG00000120438 |
| 33 | P34932 | NA | HSPA4 | heat shock 70kDa protein 4 | 3308 | ENSG00000170606 |
| 34 | P41091 | NA | EIF2S3 | eukaryotic translation initiation factor 2, subunit 3 gamma, 52kDa | 1968 | ENSG00000130741 |
| 35 | P40227 | NA | CCT6A | chaperonin containing TCP1, subunit 6A (zeta 1) | 908 | ENSG00000146731 |
| 36 | P00352 | NA | ALDH1A1 | aldehyde dehydrogenase 1 family, member A1 | 216 | ENSG00000165092 |

  
  

| **Database:molecular function      &nbspName:GTPase activity      &nbspID:GO:0003924** | | | | | | |
| --- | --- | --- | --- | --- | --- | --- |
| C=225; O=6; E=0.93; R=6.44; rawP=0.0003; adjP=0.0010 | | | | | | |
| Index | UserID | Value | Gene Symbol | Gene Name | EntrezGene | Ensembl |
| 1 | Q9BQE3 | NA | TUBA1C | tubulin, alpha 1c | 84790 | ENSG00000167553 |
| 2 | Q13885 | NA | TUBB2A | tubulin, beta 2A class IIa | 7280 | ENSG00000137267 |
| 3 | P13639 | NA | EEF2 | eukaryotic translation elongation factor 2 | 1938 | ENSG00000167658 |
| 4 | P41091 | NA | EIF2S3 | eukaryotic translation initiation factor 2, subunit 3 gamma, 52kDa | 1968 | ENSG00000130741 |
| 5 | P68371 | NA | TUBB4B | tubulin, beta 4B class IVb | 10383 | ENSG00000188229 |
| 6 | P68104 | NA | EEF1A1 | eukaryotic translation elongation factor 1 alpha 1 | 1915 | ENSG00000156508 |

  
  

| **Database:molecular function      &nbspName:heterocyclic compound binding      &nbspID:GO:1901363** | | | | | | |
| --- | --- | --- | --- | --- | --- | --- |
| C=5290; O=35; E=21.90; R=1.60; rawP=0.0005; adjP=0.0015 | | | | | | |
| Index | UserID | Value | Gene Symbol | Gene Name | EntrezGene | Ensembl |
| 1 | Q92598 | NA | HSPH1 | heat shock 105kDa/110kDa protein 1 | 10808 | ENSG00000120694 |
| 2 | Q13885 | NA | TUBB2A | tubulin, beta 2A class IIa | 7280 | ENSG00000137267 |
| 3 | P68371 | NA | TUBB4B | tubulin, beta 4B class IVb | 10383 | ENSG00000188229 |
| 4 | P68104 | NA | EEF1A1 | eukaryotic translation elongation factor 1 alpha 1 | 1915 | ENSG00000156508 |
| 5 | P50991 | NA | CCT4 | chaperonin containing TCP1, subunit 4 (delta) | 10575 | ENSG00000115484 |
| 6 | O00425 | NA | IGF2BP3 | insulin-like growth factor 2 mRNA binding protein 3 | 10643 | ENSG00000136231 |
| 7 | P22102 | NA | GART | phosphoribosylglycinamide formyltransferase, phosphoribosylglycinamide synthetase, phosphoribosylaminoimidazole synthetase | 2618 | ENSG00000159131 |
| 8 | P13639 | NA | EEF2 | eukaryotic translation elongation factor 2 | 1938 | ENSG00000167658 |
| 9 | P41567 | NA | EIF1 | eukaryotic translation initiation factor 1 | 10209 | ENSG00000173812 |
| 10 | P17174 | NA | GOT1 | glutamic-oxaloacetic transaminase 1, soluble (aspartate aminotransferase 1) | 2805 | ENSG00000120053 |
| 11 | P67809 | NA | YBX1 | Y box binding protein 1 | 4904 | ENSG00000065978 |
| 12 | Q99832 | NA | CCT7 | chaperonin containing TCP1, subunit 7 (eta) | 10574 | ENSG00000135624 |
| 13 | P21980 | NA | TGM2 | transglutaminase 2 (C polypeptide, protein-glutamine-gamma-glutamyltransferase) | 7052 | ENSG00000198959 |
| 14 | P11586 | NA | MTHFD1 | methylenetetrahydrofolate dehydrogenase (NADP+ dependent) 1, methenyltetrahydrofolate cyclohydrolase, formyltetrahydrofolate synthetase | 4522 | ENSG00000100714 |
| 15 | P11387 | NA | TOP1 | topoisomerase (DNA) I | 7150 | ENSG00000198900 |
| 16 | Q16881 | NA | TXNRD1 | thioredoxin reductase 1 | 7296 | ENSG00000198431 |
| 17 | P63241 | NA | EIF5A | eukaryotic translation initiation factor 5A | 1984 | ENSG00000132507 |
| 18 | P78371 | NA | CCT2 | chaperonin containing TCP1, subunit 2 (beta) | 10576 | ENSG00000166226 |
| 19 | Q9NR30 | NA | DDX21 | DEAD (Asp-Glu-Ala-Asp) box helicase 21 | 9188 | ENSG00000165732 |
| 20 | P00491 | NA | PNP | purine nucleoside phosphorylase | 4860 | ENSG00000198805 |
| 21 | P12081 | NA | HARS | histidyl-tRNA synthetase | 3035 | ENSG00000170445 |
| 22 | P49327 | NA | FASN | fatty acid synthase | 2194 | ENSG00000169710 |
| 23 | P49368 | NA | CCT3 | chaperonin containing TCP1, subunit 3 (gamma) | 7203 | ENSG00000163468 |
| 24 | Q9BQE3 | NA | TUBA1C | tubulin, alpha 1c | 84790 | ENSG00000167553 |
| 25 | P56192 | NA | MARS | methionyl-tRNA synthetase | 4141 | ENSG00000166986 |
| 26 | P12268 | NA | IMPDH2 | IMP (inosine 5'-monophosphate) dehydrogenase 2 | 3615 | ENSG00000178035 |
| 27 | P49736 | NA | MCM2 | minichromosome maintenance complex component 2 | 4171 | ENSG00000073111 |
| 28 | P62906 | NA | RPL10A | ribosomal protein L10a | 4736 | ENSG00000198755 |
| 29 | Q15056 | NA | EIF4H | eukaryotic translation initiation factor 4H | 7458 | ENSG00000106682 |
| 30 | P50990 | NA | CCT8 | chaperonin containing TCP1, subunit 8 (theta) | 10694 | ENSG00000156261 |
| 31 | P07814 | NA | EPRS | glutamyl-prolyl-tRNA synthetase | 2058 | ENSG00000136628 |
| 32 | P17987 | NA | TCP1 | t-complex 1 | 6950 | ENSG00000120438 |
| 33 | P34932 | NA | HSPA4 | heat shock 70kDa protein 4 | 3308 | ENSG00000170606 |
| 34 | P41091 | NA | EIF2S3 | eukaryotic translation initiation factor 2, subunit 3 gamma, 52kDa | 1968 | ENSG00000130741 |
| 35 | P40227 | NA | CCT6A | chaperonin containing TCP1, subunit 6A (zeta 1) | 908 | ENSG00000146731 |

  
  

| **Database:molecular function      &nbspName:ligase activity, forming aminoacyl-tRNA and related compounds      &nbspID:GO:0016876** | | | | | | |
| --- | --- | --- | --- | --- | --- | --- |
| C=44; O=3; E=0.18; R=16.47; rawP=0.0008; adjP=0.0021 | | | | | | |
| Index | UserID | Value | Gene Symbol | Gene Name | EntrezGene | Ensembl |
| 1 | P07814 | NA | EPRS | glutamyl-prolyl-tRNA synthetase | 2058 | ENSG00000136628 |
| 2 | P56192 | NA | MARS | methionyl-tRNA synthetase | 4141 | ENSG00000166986 |
| 3 | P12081 | NA | HARS | histidyl-tRNA synthetase | 3035 | ENSG00000170445 |

  
  

| **Database:molecular function      &nbspName:GTP binding      &nbspID:GO:0005525** | | | | | | |
| --- | --- | --- | --- | --- | --- | --- |
| C=370; O=7; E=1.53; R=4.57; rawP=0.0008; adjP=0.0021 | | | | | | |
| Index | UserID | Value | Gene Symbol | Gene Name | EntrezGene | Ensembl |
| 1 | Q9BQE3 | NA | TUBA1C | tubulin, alpha 1c | 84790 | ENSG00000167553 |
| 2 | P21980 | NA | TGM2 | transglutaminase 2 (C polypeptide, protein-glutamine-gamma-glutamyltransferase) | 7052 | ENSG00000198959 |
| 3 | Q13885 | NA | TUBB2A | tubulin, beta 2A class IIa | 7280 | ENSG00000137267 |
| 4 | P13639 | NA | EEF2 | eukaryotic translation elongation factor 2 | 1938 | ENSG00000167658 |
| 5 | P41091 | NA | EIF2S3 | eukaryotic translation initiation factor 2, subunit 3 gamma, 52kDa | 1968 | ENSG00000130741 |
| 6 | P68371 | NA | TUBB4B | tubulin, beta 4B class IVb | 10383 | ENSG00000188229 |
| 7 | P68104 | NA | EEF1A1 | eukaryotic translation elongation factor 1 alpha 1 | 1915 | ENSG00000156508 |

  
  

| **Database:molecular function      &nbspName:ligase activity, forming carbon-oxygen bonds      &nbspID:GO:0016875** | | | | | | |
| --- | --- | --- | --- | --- | --- | --- |
| C=44; O=3; E=0.18; R=16.47; rawP=0.0008; adjP=0.0021 | | | | | | |
| Index | UserID | Value | Gene Symbol | Gene Name | EntrezGene | Ensembl |
| 1 | P07814 | NA | EPRS | glutamyl-prolyl-tRNA synthetase | 2058 | ENSG00000136628 |
| 2 | P56192 | NA | MARS | methionyl-tRNA synthetase | 4141 | ENSG00000166986 |
| 3 | P12081 | NA | HARS | histidyl-tRNA synthetase | 3035 | ENSG00000170445 |

  
  

| **Database:molecular function      &nbspName:aminoacyl-tRNA ligase activity      &nbspID:GO:0004812** | | | | | | |
| --- | --- | --- | --- | --- | --- | --- |
| C=44; O=3; E=0.18; R=16.47; rawP=0.0008; adjP=0.0021 | | | | | | |
| Index | UserID | Value | Gene Symbol | Gene Name | EntrezGene | Ensembl |
| 1 | P07814 | NA | EPRS | glutamyl-prolyl-tRNA synthetase | 2058 | ENSG00000136628 |
| 2 | P56192 | NA | MARS | methionyl-tRNA synthetase | 4141 | ENSG00000166986 |
| 3 | P12081 | NA | HARS | histidyl-tRNA synthetase | 3035 | ENSG00000170445 |

  
  

| **Database:molecular function      &nbspName:nucleoside-triphosphatase activity      &nbspID:GO:0017111** | | | | | | |
| --- | --- | --- | --- | --- | --- | --- |
| C=749; O=10; E=3.10; R=3.23; rawP=0.0009; adjP=0.0022 | | | | | | |
| Index | UserID | Value | Gene Symbol | Gene Name | EntrezGene | Ensembl |
| 1 | Q9NR30 | NA | DDX21 | DEAD (Asp-Glu-Ala-Asp) box helicase 21 | 9188 | ENSG00000165732 |
| 2 | Q13885 | NA | TUBB2A | tubulin, beta 2A class IIa | 7280 | ENSG00000137267 |
| 3 | P13639 | NA | EEF2 | eukaryotic translation elongation factor 2 | 1938 | ENSG00000167658 |
| 4 | P41091 | NA | EIF2S3 | eukaryotic translation initiation factor 2, subunit 3 gamma, 52kDa | 1968 | ENSG00000130741 |
| 5 | P68371 | NA | TUBB4B | tubulin, beta 4B class IVb | 10383 | ENSG00000188229 |
| 6 | Q9BQE3 | NA | TUBA1C | tubulin, alpha 1c | 84790 | ENSG00000167553 |
| 7 | P49736 | NA | MCM2 | minichromosome maintenance complex component 2 | 4171 | ENSG00000073111 |
| 8 | P50990 | NA | CCT8 | chaperonin containing TCP1, subunit 8 (theta) | 10694 | ENSG00000156261 |
| 9 | P68104 | NA | EEF1A1 | eukaryotic translation elongation factor 1 alpha 1 | 1915 | ENSG00000156508 |
| 10 | P11387 | NA | TOP1 | topoisomerase (DNA) I | 7150 | ENSG00000198900 |

  
  

| **Database:molecular function      &nbspName:guanyl nucleotide binding      &nbspID:GO:0019001** | | | | | | |
| --- | --- | --- | --- | --- | --- | --- |
| C=383; O=7; E=1.59; R=4.42; rawP=0.0010; adjP=0.0023 | | | | | | |
| Index | UserID | Value | Gene Symbol | Gene Name | EntrezGene | Ensembl |
| 1 | Q9BQE3 | NA | TUBA1C | tubulin, alpha 1c | 84790 | ENSG00000167553 |
| 2 | P21980 | NA | TGM2 | transglutaminase 2 (C polypeptide, protein-glutamine-gamma-glutamyltransferase) | 7052 | ENSG00000198959 |
| 3 | Q13885 | NA | TUBB2A | tubulin, beta 2A class IIa | 7280 | ENSG00000137267 |
| 4 | P13639 | NA | EEF2 | eukaryotic translation elongation factor 2 | 1938 | ENSG00000167658 |
| 5 | P41091 | NA | EIF2S3 | eukaryotic translation initiation factor 2, subunit 3 gamma, 52kDa | 1968 | ENSG00000130741 |
| 6 | P68371 | NA | TUBB4B | tubulin, beta 4B class IVb | 10383 | ENSG00000188229 |
| 7 | P68104 | NA | EEF1A1 | eukaryotic translation elongation factor 1 alpha 1 | 1915 | ENSG00000156508 |

  
  

| **Database:molecular function      &nbspName:guanyl ribonucleotide binding      &nbspID:GO:0032561** | | | | | | |
| --- | --- | --- | --- | --- | --- | --- |
| C=383; O=7; E=1.59; R=4.42; rawP=0.0010; adjP=0.0023 | | | | | | |
| Index | UserID | Value | Gene Symbol | Gene Name | EntrezGene | Ensembl |
| 1 | Q9BQE3 | NA | TUBA1C | tubulin, alpha 1c | 84790 | ENSG00000167553 |
| 2 | P21980 | NA | TGM2 | transglutaminase 2 (C polypeptide, protein-glutamine-gamma-glutamyltransferase) | 7052 | ENSG00000198959 |
| 3 | Q13885 | NA | TUBB2A | tubulin, beta 2A class IIa | 7280 | ENSG00000137267 |
| 4 | P13639 | NA | EEF2 | eukaryotic translation elongation factor 2 | 1938 | ENSG00000167658 |
| 5 | P41091 | NA | EIF2S3 | eukaryotic translation initiation factor 2, subunit 3 gamma, 52kDa | 1968 | ENSG00000130741 |
| 6 | P68371 | NA | TUBB4B | tubulin, beta 4B class IVb | 10383 | ENSG00000188229 |
| 7 | P68104 | NA | EEF1A1 | eukaryotic translation elongation factor 1 alpha 1 | 1915 | ENSG00000156508 |

  
  

| **Database:molecular function      &nbspName:translation initiation factor activity      &nbspID:GO:0003743** | | | | | | |
| --- | --- | --- | --- | --- | --- | --- |
| C=49; O=3; E=0.20; R=14.79; rawP=0.0011; adjP=0.0025 | | | | | | |
| Index | UserID | Value | Gene Symbol | Gene Name | EntrezGene | Ensembl |
| 1 | P41091 | NA | EIF2S3 | eukaryotic translation initiation factor 2, subunit 3 gamma, 52kDa | 1968 | ENSG00000130741 |
| 2 | Q15056 | NA | EIF4H | eukaryotic translation initiation factor 4H | 7458 | ENSG00000106682 |
| 3 | P41567 | NA | EIF1 | eukaryotic translation initiation factor 1 | 10209 | ENSG00000173812 |

  
  

| **Database:molecular function      &nbspName:pyrophosphatase activity      &nbspID:GO:0016462** | | | | | | |
| --- | --- | --- | --- | --- | --- | --- |
| C=783; O=10; E=3.24; R=3.09; rawP=0.0013; adjP=0.0026 | | | | | | |
| Index | UserID | Value | Gene Symbol | Gene Name | EntrezGene | Ensembl |
| 1 | Q9NR30 | NA | DDX21 | DEAD (Asp-Glu-Ala-Asp) box helicase 21 | 9188 | ENSG00000165732 |
| 2 | Q13885 | NA | TUBB2A | tubulin, beta 2A class IIa | 7280 | ENSG00000137267 |
| 3 | P13639 | NA | EEF2 | eukaryotic translation elongation factor 2 | 1938 | ENSG00000167658 |
| 4 | P41091 | NA | EIF2S3 | eukaryotic translation initiation factor 2, subunit 3 gamma, 52kDa | 1968 | ENSG00000130741 |
| 5 | P68371 | NA | TUBB4B | tubulin, beta 4B class IVb | 10383 | ENSG00000188229 |
| 6 | Q9BQE3 | NA | TUBA1C | tubulin, alpha 1c | 84790 | ENSG00000167553 |
| 7 | P49736 | NA | MCM2 | minichromosome maintenance complex component 2 | 4171 | ENSG00000073111 |
| 8 | P50990 | NA | CCT8 | chaperonin containing TCP1, subunit 8 (theta) | 10694 | ENSG00000156261 |
| 9 | P68104 | NA | EEF1A1 | eukaryotic translation elongation factor 1 alpha 1 | 1915 | ENSG00000156508 |
| 10 | P11387 | NA | TOP1 | topoisomerase (DNA) I | 7150 | ENSG00000198900 |

  
  

| **Database:molecular function      &nbspName:hydrolase activity, acting on acid anhydrides, in phosphorus-containing anhydrides      &nbspID:GO:0016818** | | | | | | |
| --- | --- | --- | --- | --- | --- | --- |
| C=786; O=10; E=3.25; R=3.07; rawP=0.0013; adjP=0.0026 | | | | | | |
| Index | UserID | Value | Gene Symbol | Gene Name | EntrezGene | Ensembl |
| 1 | Q9NR30 | NA | DDX21 | DEAD (Asp-Glu-Ala-Asp) box helicase 21 | 9188 | ENSG00000165732 |
| 2 | Q13885 | NA | TUBB2A | tubulin, beta 2A class IIa | 7280 | ENSG00000137267 |
| 3 | P13639 | NA | EEF2 | eukaryotic translation elongation factor 2 | 1938 | ENSG00000167658 |
| 4 | P41091 | NA | EIF2S3 | eukaryotic translation initiation factor 2, subunit 3 gamma, 52kDa | 1968 | ENSG00000130741 |
| 5 | P68371 | NA | TUBB4B | tubulin, beta 4B class IVb | 10383 | ENSG00000188229 |
| 6 | Q9BQE3 | NA | TUBA1C | tubulin, alpha 1c | 84790 | ENSG00000167553 |
| 7 | P49736 | NA | MCM2 | minichromosome maintenance complex component 2 | 4171 | ENSG00000073111 |
| 8 | P50990 | NA | CCT8 | chaperonin containing TCP1, subunit 8 (theta) | 10694 | ENSG00000156261 |
| 9 | P68104 | NA | EEF1A1 | eukaryotic translation elongation factor 1 alpha 1 | 1915 | ENSG00000156508 |
| 10 | P11387 | NA | TOP1 | topoisomerase (DNA) I | 7150 | ENSG00000198900 |

  
  

| **Database:molecular function      &nbspName:ion binding      &nbspID:GO:0043167** | | | | | | |
| --- | --- | --- | --- | --- | --- | --- |
| C=5721; O=36; E=23.68; R=1.52; rawP=0.0012; adjP=0.0026 | | | | | | |
| Index | UserID | Value | Gene Symbol | Gene Name | EntrezGene | Ensembl |
| 1 | Q92598 | NA | HSPH1 | heat shock 105kDa/110kDa protein 1 | 10808 | ENSG00000120694 |
| 2 | Q13885 | NA | TUBB2A | tubulin, beta 2A class IIa | 7280 | ENSG00000137267 |
| 3 | P68371 | NA | TUBB4B | tubulin, beta 4B class IVb | 10383 | ENSG00000188229 |
| 4 | P53701 | NA | HCCS | holocytochrome c synthase | 3052 | ENSG00000004961 |
| 5 | P68104 | NA | EEF1A1 | eukaryotic translation elongation factor 1 alpha 1 | 1915 | ENSG00000156508 |
| 6 | P50991 | NA | CCT4 | chaperonin containing TCP1, subunit 4 (delta) | 10575 | ENSG00000115484 |
| 7 | P22102 | NA | GART | phosphoribosylglycinamide formyltransferase, phosphoribosylglycinamide synthetase, phosphoribosylaminoimidazole synthetase | 2618 | ENSG00000159131 |
| 8 | P13639 | NA | EEF2 | eukaryotic translation elongation factor 2 | 1938 | ENSG00000167658 |
| 9 | P17174 | NA | GOT1 | glutamic-oxaloacetic transaminase 1, soluble (aspartate aminotransferase 1) | 2805 | ENSG00000120053 |
| 10 | Q99832 | NA | CCT7 | chaperonin containing TCP1, subunit 7 (eta) | 10574 | ENSG00000135624 |
| 11 | P21980 | NA | TGM2 | transglutaminase 2 (C polypeptide, protein-glutamine-gamma-glutamyltransferase) | 7052 | ENSG00000198959 |
| 12 | P06703 | NA | S100A6 | S100 calcium binding protein A6 | 6277 | ENSG00000197956 |
| 13 | P11586 | NA | MTHFD1 | methylenetetrahydrofolate dehydrogenase (NADP+ dependent) 1, methenyltetrahydrofolate cyclohydrolase, formyltetrahydrofolate synthetase | 4522 | ENSG00000100714 |
| 14 | P11387 | NA | TOP1 | topoisomerase (DNA) I | 7150 | ENSG00000198900 |
| 15 | Q16881 | NA | TXNRD1 | thioredoxin reductase 1 | 7296 | ENSG00000198431 |
| 16 | P78371 | NA | CCT2 | chaperonin containing TCP1, subunit 2 (beta) | 10576 | ENSG00000166226 |
| 17 | Q9NR30 | NA | DDX21 | DEAD (Asp-Glu-Ala-Asp) box helicase 21 | 9188 | ENSG00000165732 |
| 18 | P08195 | NA | SLC3A2 | solute carrier family 3 (activators of dibasic and neutral amino acid transport), member 2 | 6520 | ENSG00000168003 |
| 19 | Q7Z2K6 | NA | ERMP1 | endoplasmic reticulum metallopeptidase 1 | 79956 | ENSG00000099219 |
| 20 | P02795 | NA | MT2A | metallothionein 2A | 4502 | ENSG00000125148 |
| 21 | P00491 | NA | PNP | purine nucleoside phosphorylase | 4860 | ENSG00000198805 |
| 22 | P12081 | NA | HARS | histidyl-tRNA synthetase | 3035 | ENSG00000170445 |
| 23 | P49327 | NA | FASN | fatty acid synthase | 2194 | ENSG00000169710 |
| 24 | P67936 | NA | TPM4 | tropomyosin 4 | 7171 | ENSG00000167460 |
| 25 | P49368 | NA | CCT3 | chaperonin containing TCP1, subunit 3 (gamma) | 7203 | ENSG00000163468 |
| 26 | Q9BQE3 | NA | TUBA1C | tubulin, alpha 1c | 84790 | ENSG00000167553 |
| 27 | P56192 | NA | MARS | methionyl-tRNA synthetase | 4141 | ENSG00000166986 |
| 28 | P12268 | NA | IMPDH2 | IMP (inosine 5'-monophosphate) dehydrogenase 2 | 3615 | ENSG00000178035 |
| 29 | P49736 | NA | MCM2 | minichromosome maintenance complex component 2 | 4171 | ENSG00000073111 |
| 30 | P50990 | NA | CCT8 | chaperonin containing TCP1, subunit 8 (theta) | 10694 | ENSG00000156261 |
| 31 | P07814 | NA | EPRS | glutamyl-prolyl-tRNA synthetase | 2058 | ENSG00000136628 |
| 32 | P17987 | NA | TCP1 | t-complex 1 | 6950 | ENSG00000120438 |
| 33 | P34932 | NA | HSPA4 | heat shock 70kDa protein 4 | 3308 | ENSG00000170606 |
| 34 | P41091 | NA | EIF2S3 | eukaryotic translation initiation factor 2, subunit 3 gamma, 52kDa | 1968 | ENSG00000130741 |
| 35 | P07108 | NA | DBI | diazepam binding inhibitor (GABA receptor modulator, acyl-CoA binding protein) | 1622 | ENSG00000155368 |
| 36 | P40227 | NA | CCT6A | chaperonin containing TCP1, subunit 6A (zeta 1) | 908 | ENSG00000146731 |

  
  

| **Database:molecular function      &nbspName:binding      &nbspID:GO:0005488** | | | | | | |
| --- | --- | --- | --- | --- | --- | --- |
| C=11779; O=58; E=48.76; R=1.19; rawP=0.0013; adjP=0.0026 | | | | | | |
| Index | UserID | Value | Gene Symbol | Gene Name | EntrezGene | Ensembl |
| 1 | Q92598 | NA | HSPH1 | heat shock 105kDa/110kDa protein 1 | 10808 | ENSG00000120694 |
| 2 | Q13885 | NA | TUBB2A | tubulin, beta 2A class IIa | 7280 | ENSG00000137267 |
| 3 | P68371 | NA | TUBB4B | tubulin, beta 4B class IVb | 10383 | ENSG00000188229 |
| 4 | P25685 | NA | DNAJB1 | DnaJ (Hsp40) homolog, subfamily B, member 1 | 3337 | ENSG00000132002 |
| 5 | P53701 | NA | HCCS | holocytochrome c synthase | 3052 | ENSG00000004961 |
| 6 | P68104 | NA | EEF1A1 | eukaryotic translation elongation factor 1 alpha 1 | 1915 | ENSG00000156508 |
| 7 | O75369 | NA | FLNB | filamin B, beta | 2317 | ENSG00000136068 |
| 8 | P02786 | NA | TFRC | transferrin receptor (p90, CD71) | 7037 | ENSG00000072274 |
| 9 | O00425 | NA | IGF2BP3 | insulin-like growth factor 2 mRNA binding protein 3 | 10643 | ENSG00000136231 |
| 10 | P50991 | NA | CCT4 | chaperonin containing TCP1, subunit 4 (delta) | 10575 | ENSG00000115484 |
| 11 | P22102 | NA | GART | phosphoribosylglycinamide formyltransferase, phosphoribosylglycinamide synthetase, phosphoribosylaminoimidazole synthetase | 2618 | ENSG00000159131 |
| 12 | P13639 | NA | EEF2 | eukaryotic translation elongation factor 2 | 1938 | ENSG00000167658 |
| 13 | Q16555 | NA | DPYSL2 | dihydropyrimidinase-like 2 | 1808 | ENSG00000092964 |
| 14 | O95433 | NA | AHSA1 | AHA1, activator of heat shock 90kDa protein ATPase homolog 1 (yeast) | 10598 | ENSG00000100591 |
| 15 | P41567 | NA | EIF1 | eukaryotic translation initiation factor 1 | 10209 | ENSG00000173812 |
| 16 | P17174 | NA | GOT1 | glutamic-oxaloacetic transaminase 1, soluble (aspartate aminotransferase 1) | 2805 | ENSG00000120053 |
| 17 | P67809 | NA | YBX1 | Y box binding protein 1 | 4904 | ENSG00000065978 |
| 18 | P21980 | NA | TGM2 | transglutaminase 2 (C polypeptide, protein-glutamine-gamma-glutamyltransferase) | 7052 | ENSG00000198959 |
| 19 | P37802 | NA | TAGLN2 | transgelin 2 | 8407 | ENSG00000158710 |
| 20 | Q99832 | NA | CCT7 | chaperonin containing TCP1, subunit 7 (eta) | 10574 | ENSG00000135624 |
| 21 | Q15185 | NA | PTGES3 | prostaglandin E synthase 3 (cytosolic) | 10728 | ENSG00000110958 |
| 22 | P18206 | NA | VCL | vinculin | 7414 | ENSG00000035403 |
| 23 | P06703 | NA | S100A6 | S100 calcium binding protein A6 | 6277 | ENSG00000197956 |
| 24 | P11387 | NA | TOP1 | topoisomerase (DNA) I | 7150 | ENSG00000198900 |
| 25 | P11586 | NA | MTHFD1 | methylenetetrahydrofolate dehydrogenase (NADP+ dependent) 1, methenyltetrahydrofolate cyclohydrolase, formyltetrahydrofolate synthetase | 4522 | ENSG00000100714 |
| 26 | Q16881 | NA | TXNRD1 | thioredoxin reductase 1 | 7296 | ENSG00000198431 |
| 27 | P63241 | NA | EIF5A | eukaryotic translation initiation factor 5A | 1984 | ENSG00000132507 |
| 28 | P78371 | NA | CCT2 | chaperonin containing TCP1, subunit 2 (beta) | 10576 | ENSG00000166226 |
| 29 | Q9NR30 | NA | DDX21 | DEAD (Asp-Glu-Ala-Asp) box helicase 21 | 9188 | ENSG00000165732 |
| 30 | P08195 | NA | SLC3A2 | solute carrier family 3 (activators of dibasic and neutral amino acid transport), member 2 | 6520 | ENSG00000168003 |
| 31 | Q7Z2K6 | NA | ERMP1 | endoplasmic reticulum metallopeptidase 1 | 79956 | ENSG00000099219 |
| 32 | P02795 | NA | MT2A | metallothionein 2A | 4502 | ENSG00000125148 |
| 33 | P49321 | NA | NASP | nuclear autoantigenic sperm protein (histone-binding) | 4678 | ENSG00000132780 |
| 34 | P00491 | NA | PNP | purine nucleoside phosphorylase | 4860 | ENSG00000198805 |
| 35 | Q9NZM1 | NA | MYOF | myoferlin | 26509 | ENSG00000138119 |
| 36 | P07951 | NA | TPM2 | tropomyosin 2 (beta) | 7169 | ENSG00000198467 |
| 37 | P12081 | NA | HARS | histidyl-tRNA synthetase | 3035 | ENSG00000170445 |
| 38 | P67936 | NA | TPM4 | tropomyosin 4 | 7171 | ENSG00000167460 |
| 39 | P49327 | NA | FASN | fatty acid synthase | 2194 | ENSG00000169710 |
| 40 | P49368 | NA | CCT3 | chaperonin containing TCP1, subunit 3 (gamma) | 7203 | ENSG00000163468 |
| 41 | P56192 | NA | MARS | methionyl-tRNA synthetase | 4141 | ENSG00000166986 |
| 42 | Q9BQE3 | NA | TUBA1C | tubulin, alpha 1c | 84790 | ENSG00000167553 |
| 43 | P12268 | NA | IMPDH2 | IMP (inosine 5'-monophosphate) dehydrogenase 2 | 3615 | ENSG00000178035 |
| 44 | P62906 | NA | RPL10A | ribosomal protein L10a | 4736 | ENSG00000198755 |
| 45 | P49736 | NA | MCM2 | minichromosome maintenance complex component 2 | 4171 | ENSG00000073111 |
| 46 | Q15056 | NA | EIF4H | eukaryotic translation initiation factor 4H | 7458 | ENSG00000106682 |
| 47 | O60610 | NA | DIAPH1 | diaphanous homolog 1 (Drosophila) | 1729 | ENSG00000131504 |
| 48 | P50990 | NA | CCT8 | chaperonin containing TCP1, subunit 8 (theta) | 10694 | ENSG00000156261 |
| 49 | Q06830 | NA | PRDX1 | peroxiredoxin 1 | 5052 | ENSG00000117450 |
| 50 | P07814 | NA | EPRS | glutamyl-prolyl-tRNA synthetase | 2058 | ENSG00000136628 |
| 51 | P17987 | NA | TCP1 | t-complex 1 | 6950 | ENSG00000120438 |
| 52 | P55209 | NA | NAP1L1 | nucleosome assembly protein 1-like 1 | 4673 | ENSG00000187109 |
| 53 | P34932 | NA | HSPA4 | heat shock 70kDa protein 4 | 3308 | ENSG00000170606 |
| 54 | P41091 | NA | EIF2S3 | eukaryotic translation initiation factor 2, subunit 3 gamma, 52kDa | 1968 | ENSG00000130741 |
| 55 | P07108 | NA | DBI | diazepam binding inhibitor (GABA receptor modulator, acyl-CoA binding protein) | 1622 | ENSG00000155368 |
| 56 | P40227 | NA | CCT6A | chaperonin containing TCP1, subunit 6A (zeta 1) | 908 | ENSG00000146731 |
| 57 | P52292 | NA | KPNA2 | karyopherin alpha 2 (RAG cohort 1, importin alpha 1) | 3838 | ENSG00000182481 |
| 58 | P00352 | NA | ALDH1A1 | aldehyde dehydrogenase 1 family, member A1 | 216 | ENSG00000165092 |

  
  

| **Database:molecular function      &nbspName:hydrolase activity, acting on acid anhydrides      &nbspID:GO:0016817** | | | | | | |
| --- | --- | --- | --- | --- | --- | --- |
| C=791; O=10; E=3.27; R=3.05; rawP=0.0014; adjP=0.0027 | | | | | | |
| Index | UserID | Value | Gene Symbol | Gene Name | EntrezGene | Ensembl |
| 1 | Q9NR30 | NA | DDX21 | DEAD (Asp-Glu-Ala-Asp) box helicase 21 | 9188 | ENSG00000165732 |
| 2 | Q13885 | NA | TUBB2A | tubulin, beta 2A class IIa | 7280 | ENSG00000137267 |
| 3 | P13639 | NA | EEF2 | eukaryotic translation elongation factor 2 | 1938 | ENSG00000167658 |
| 4 | P41091 | NA | EIF2S3 | eukaryotic translation initiation factor 2, subunit 3 gamma, 52kDa | 1968 | ENSG00000130741 |
| 5 | P68371 | NA | TUBB4B | tubulin, beta 4B class IVb | 10383 | ENSG00000188229 |
| 6 | Q9BQE3 | NA | TUBA1C | tubulin, alpha 1c | 84790 | ENSG00000167553 |
| 7 | P49736 | NA | MCM2 | minichromosome maintenance complex component 2 | 4171 | ENSG00000073111 |
| 8 | P50990 | NA | CCT8 | chaperonin containing TCP1, subunit 8 (theta) | 10694 | ENSG00000156261 |
| 9 | P68104 | NA | EEF1A1 | eukaryotic translation elongation factor 1 alpha 1 | 1915 | ENSG00000156508 |
| 10 | P11387 | NA | TOP1 | topoisomerase (DNA) I | 7150 | ENSG00000198900 |

  
  

| **Database:molecular function      &nbspName:heat shock protein binding      &nbspID:GO:0031072** | | | | | | |
| --- | --- | --- | --- | --- | --- | --- |
| C=90; O=3; E=0.37; R=8.05; rawP=0.0061; adjP=0.0114 | | | | | | |
| Index | UserID | Value | Gene Symbol | Gene Name | EntrezGene | Ensembl |
| 1 | P25685 | NA | DNAJB1 | DnaJ (Hsp40) homolog, subfamily B, member 1 | 3337 | ENSG00000132002 |
| 2 | P49321 | NA | NASP | nuclear autoantigenic sperm protein (histone-binding) | 4678 | ENSG00000132780 |
| 3 | P02786 | NA | TFRC | transferrin receptor (p90, CD71) | 7037 | ENSG00000072274 |

  
  

| **Database:molecular function      &nbspName:structural molecule activity      &nbspID:GO:0005198** | | | | | | |
| --- | --- | --- | --- | --- | --- | --- |
| C=608; O=7; E=2.52; R=2.78; rawP=0.0125; adjP=0.0220 | | | | | | |
| Index | UserID | Value | Gene Symbol | Gene Name | EntrezGene | Ensembl |
| 1 | Q9BQE3 | NA | TUBA1C | tubulin, alpha 1c | 84790 | ENSG00000167553 |
| 2 | Q13885 | NA | TUBB2A | tubulin, beta 2A class IIa | 7280 | ENSG00000137267 |
| 3 | P62906 | NA | RPL10A | ribosomal protein L10a | 4736 | ENSG00000198755 |
| 4 | P07951 | NA | TPM2 | tropomyosin 2 (beta) | 7169 | ENSG00000198467 |
| 5 | P18206 | NA | VCL | vinculin | 7414 | ENSG00000035403 |
| 6 | P68371 | NA | TUBB4B | tubulin, beta 4B class IVb | 10383 | ENSG00000188229 |
| 7 | P67936 | NA | TPM4 | tropomyosin 4 | 7171 | ENSG00000167460 |

  
  

| **Database:molecular function      &nbspName:catalytic activity      &nbspID:GO:0003824** | | | | | | |
| --- | --- | --- | --- | --- | --- | --- |
| C=5299; O=31; E=21.94; R=1.41; rawP=0.0125; adjP=0.0220 | | | | | | |
| Index | UserID | Value | Gene Symbol | Gene Name | EntrezGene | Ensembl |
| 1 | Q16881 | NA | TXNRD1 | thioredoxin reductase 1 | 7296 | ENSG00000198431 |
| 2 | Q9NR30 | NA | DDX21 | DEAD (Asp-Glu-Ala-Asp) box helicase 21 | 9188 | ENSG00000165732 |
| 3 | Q7Z2K6 | NA | ERMP1 | endoplasmic reticulum metallopeptidase 1 | 79956 | ENSG00000099219 |
| 4 | P08195 | NA | SLC3A2 | solute carrier family 3 (activators of dibasic and neutral amino acid transport), member 2 | 6520 | ENSG00000168003 |
| 5 | Q13885 | NA | TUBB2A | tubulin, beta 2A class IIa | 7280 | ENSG00000137267 |
| 6 | P00491 | NA | PNP | purine nucleoside phosphorylase | 4860 | ENSG00000198805 |
| 7 | P12081 | NA | HARS | histidyl-tRNA synthetase | 3035 | ENSG00000170445 |
| 8 | P68371 | NA | TUBB4B | tubulin, beta 4B class IVb | 10383 | ENSG00000188229 |
| 9 | P49327 | NA | FASN | fatty acid synthase | 2194 | ENSG00000169710 |
| 10 | P56192 | NA | MARS | methionyl-tRNA synthetase | 4141 | ENSG00000166986 |
| 11 | Q9BQE3 | NA | TUBA1C | tubulin, alpha 1c | 84790 | ENSG00000167553 |
| 12 | P12268 | NA | IMPDH2 | IMP (inosine 5'-monophosphate) dehydrogenase 2 | 3615 | ENSG00000178035 |
| 13 | P49736 | NA | MCM2 | minichromosome maintenance complex component 2 | 4171 | ENSG00000073111 |
| 14 | P53701 | NA | HCCS | holocytochrome c synthase | 3052 | ENSG00000004961 |
| 15 | P50990 | NA | CCT8 | chaperonin containing TCP1, subunit 8 (theta) | 10694 | ENSG00000156261 |
| 16 | P68104 | NA | EEF1A1 | eukaryotic translation elongation factor 1 alpha 1 | 1915 | ENSG00000156508 |
| 17 | Q06830 | NA | PRDX1 | peroxiredoxin 1 | 5052 | ENSG00000117450 |
| 18 | P02786 | NA | TFRC | transferrin receptor (p90, CD71) | 7037 | ENSG00000072274 |
| 19 | P07814 | NA | EPRS | glutamyl-prolyl-tRNA synthetase | 2058 | ENSG00000136628 |
| 20 | P22102 | NA | GART | phosphoribosylglycinamide formyltransferase, phosphoribosylglycinamide synthetase, phosphoribosylaminoimidazole synthetase | 2618 | ENSG00000159131 |
| 21 | P13639 | NA | EEF2 | eukaryotic translation elongation factor 2 | 1938 | ENSG00000167658 |
| 22 | P41091 | NA | EIF2S3 | eukaryotic translation initiation factor 2, subunit 3 gamma, 52kDa | 1968 | ENSG00000130741 |
| 23 | Q16555 | NA | DPYSL2 | dihydropyrimidinase-like 2 | 1808 | ENSG00000092964 |
| 24 | P17174 | NA | GOT1 | glutamic-oxaloacetic transaminase 1, soluble (aspartate aminotransferase 1) | 2805 | ENSG00000120053 |
| 25 | Q15185 | NA | PTGES3 | prostaglandin E synthase 3 (cytosolic) | 10728 | ENSG00000110958 |
| 26 | P21980 | NA | TGM2 | transglutaminase 2 (C polypeptide, protein-glutamine-gamma-glutamyltransferase) | 7052 | ENSG00000198959 |
| 27 | Q14166 | NA | TTLL12 | tubulin tyrosine ligase-like family, member 12 | 23170 | ENSG00000100304 |
| 28 | Q6PIU2 | NA | NCEH1 | neutral cholesterol ester hydrolase 1 | 57552 | ENSG00000144959 |
| 29 | P00352 | NA | ALDH1A1 | aldehyde dehydrogenase 1 family, member A1 | 216 | ENSG00000165092 |
| 30 | P11586 | NA | MTHFD1 | methylenetetrahydrofolate dehydrogenase (NADP+ dependent) 1, methenyltetrahydrofolate cyclohydrolase, formyltetrahydrofolate synthetase | 4522 | ENSG00000100714 |
| 31 | P11387 | NA | TOP1 | topoisomerase (DNA) I | 7150 | ENSG00000198900 |

  
  

| **Database:molecular function      &nbspName:ligase activity      &nbspID:GO:0016874** | | | | | | |
| --- | --- | --- | --- | --- | --- | --- |
| C=478; O=6; E=1.98; R=3.03; rawP=0.0139; adjP=0.0239 | | | | | | |
| Index | UserID | Value | Gene Symbol | Gene Name | EntrezGene | Ensembl |
| 1 | P07814 | NA | EPRS | glutamyl-prolyl-tRNA synthetase | 2058 | ENSG00000136628 |
| 2 | P56192 | NA | MARS | methionyl-tRNA synthetase | 4141 | ENSG00000166986 |
| 3 | P22102 | NA | GART | phosphoribosylglycinamide formyltransferase, phosphoribosylglycinamide synthetase, phosphoribosylaminoimidazole synthetase | 2618 | ENSG00000159131 |
| 4 | Q14166 | NA | TTLL12 | tubulin tyrosine ligase-like family, member 12 | 23170 | ENSG00000100304 |
| 5 | P12081 | NA | HARS | histidyl-tRNA synthetase | 3035 | ENSG00000170445 |
| 6 | P11586 | NA | MTHFD1 | methylenetetrahydrofolate dehydrogenase (NADP+ dependent) 1, methenyltetrahydrofolate cyclohydrolase, formyltetrahydrofolate synthetase | 4522 | ENSG00000100714 |

  
  

| **Database:molecular function      &nbspName:actin binding      &nbspID:GO:0003779** | | | | | | |
| --- | --- | --- | --- | --- | --- | --- |
| C=353; O=5; E=1.46; R=3.42; rawP=0.0153; adjP=0.0252 | | | | | | |
| Index | UserID | Value | Gene Symbol | Gene Name | EntrezGene | Ensembl |
| 1 | P07951 | NA | TPM2 | tropomyosin 2 (beta) | 7169 | ENSG00000198467 |
| 2 | P18206 | NA | VCL | vinculin | 7414 | ENSG00000035403 |
| 3 | O60610 | NA | DIAPH1 | diaphanous homolog 1 (Drosophila) | 1729 | ENSG00000131504 |
| 4 | O75369 | NA | FLNB | filamin B, beta | 2317 | ENSG00000136068 |
| 5 | P67936 | NA | TPM4 | tropomyosin 4 | 7171 | ENSG00000167460 |

  
  

| **Database:cellular component      &nbspName:cytosol      &nbspID:GO:0005829** | | | | | | |
| --- | --- | --- | --- | --- | --- | --- |
| C=2367; O=37; E=8.80; R=4.21; rawP=1.25e-16; adjP=9.50e-15 | | | | | | |
| Index | UserID | Value | Gene Symbol | Gene Name | EntrezGene | Ensembl |
| 1 | P68371 | NA | TUBB4B | tubulin, beta 4B class IVb | 10383 | ENSG00000188229 |
| 2 | P68104 | NA | EEF1A1 | eukaryotic translation elongation factor 1 alpha 1 | 1915 | ENSG00000156508 |
| 3 | O75369 | NA | FLNB | filamin B, beta | 2317 | ENSG00000136068 |
| 4 | P50991 | NA | CCT4 | chaperonin containing TCP1, subunit 4 (delta) | 10575 | ENSG00000115484 |
| 5 | O00425 | NA | IGF2BP3 | insulin-like growth factor 2 mRNA binding protein 3 | 10643 | ENSG00000136231 |
| 6 | P22102 | NA | GART | phosphoribosylglycinamide formyltransferase, phosphoribosylglycinamide synthetase, phosphoribosylaminoimidazole synthetase | 2618 | ENSG00000159131 |
| 7 | P13639 | NA | EEF2 | eukaryotic translation elongation factor 2 | 1938 | ENSG00000167658 |
| 8 | Q16555 | NA | DPYSL2 | dihydropyrimidinase-like 2 | 1808 | ENSG00000092964 |
| 9 | O95433 | NA | AHSA1 | AHA1, activator of heat shock 90kDa protein ATPase homolog 1 (yeast) | 10598 | ENSG00000100591 |
| 10 | P17174 | NA | GOT1 | glutamic-oxaloacetic transaminase 1, soluble (aspartate aminotransferase 1) | 2805 | ENSG00000120053 |
| 11 | Q15185 | NA | PTGES3 | prostaglandin E synthase 3 (cytosolic) | 10728 | ENSG00000110958 |
| 12 | Q99832 | NA | CCT7 | chaperonin containing TCP1, subunit 7 (eta) | 10574 | ENSG00000135624 |
| 13 | P18206 | NA | VCL | vinculin | 7414 | ENSG00000035403 |
| 14 | P06703 | NA | S100A6 | S100 calcium binding protein A6 | 6277 | ENSG00000197956 |
| 15 | P11586 | NA | MTHFD1 | methylenetetrahydrofolate dehydrogenase (NADP+ dependent) 1, methenyltetrahydrofolate cyclohydrolase, formyltetrahydrofolate synthetase | 4522 | ENSG00000100714 |
| 16 | Q16881 | NA | TXNRD1 | thioredoxin reductase 1 | 7296 | ENSG00000198431 |
| 17 | P63241 | NA | EIF5A | eukaryotic translation initiation factor 5A | 1984 | ENSG00000132507 |
| 18 | P78371 | NA | CCT2 | chaperonin containing TCP1, subunit 2 (beta) | 10576 | ENSG00000166226 |
| 19 | P02795 | NA | MT2A | metallothionein 2A | 4502 | ENSG00000125148 |
| 20 | P07951 | NA | TPM2 | tropomyosin 2 (beta) | 7169 | ENSG00000198467 |
| 21 | P00491 | NA | PNP | purine nucleoside phosphorylase | 4860 | ENSG00000198805 |
| 22 | P12081 | NA | HARS | histidyl-tRNA synthetase | 3035 | ENSG00000170445 |
| 23 | P49327 | NA | FASN | fatty acid synthase | 2194 | ENSG00000169710 |
| 24 | P67936 | NA | TPM4 | tropomyosin 4 | 7171 | ENSG00000167460 |
| 25 | P49368 | NA | CCT3 | chaperonin containing TCP1, subunit 3 (gamma) | 7203 | ENSG00000163468 |
| 26 | P56192 | NA | MARS | methionyl-tRNA synthetase | 4141 | ENSG00000166986 |
| 27 | P12268 | NA | IMPDH2 | IMP (inosine 5'-monophosphate) dehydrogenase 2 | 3615 | ENSG00000178035 |
| 28 | P62906 | NA | RPL10A | ribosomal protein L10a | 4736 | ENSG00000198755 |
| 29 | Q15056 | NA | EIF4H | eukaryotic translation initiation factor 4H | 7458 | ENSG00000106682 |
| 30 | P50990 | NA | CCT8 | chaperonin containing TCP1, subunit 8 (theta) | 10694 | ENSG00000156261 |
| 31 | P07814 | NA | EPRS | glutamyl-prolyl-tRNA synthetase | 2058 | ENSG00000136628 |
| 32 | P17987 | NA | TCP1 | t-complex 1 | 6950 | ENSG00000120438 |
| 33 | P34932 | NA | HSPA4 | heat shock 70kDa protein 4 | 3308 | ENSG00000170606 |
| 34 | P41091 | NA | EIF2S3 | eukaryotic translation initiation factor 2, subunit 3 gamma, 52kDa | 1968 | ENSG00000130741 |
| 35 | P40227 | NA | CCT6A | chaperonin containing TCP1, subunit 6A (zeta 1) | 908 | ENSG00000146731 |
| 36 | P52292 | NA | KPNA2 | karyopherin alpha 2 (RAG cohort 1, importin alpha 1) | 3838 | ENSG00000182481 |
| 37 | P00352 | NA | ALDH1A1 | aldehyde dehydrogenase 1 family, member A1 | 216 | ENSG00000165092 |

  
  

| **Database:cellular component      &nbspName:chaperonin-containing T-complex      &nbspID:GO:0005832** | | | | | | |
| --- | --- | --- | --- | --- | --- | --- |
| C=7; O=6; E=0.03; R=230.63; rawP=1.43e-14; adjP=5.43e-13 | | | | | | |
| Index | UserID | Value | Gene Symbol | Gene Name | EntrezGene | Ensembl |
| 1 | P40227 | NA | CCT6A | chaperonin containing TCP1, subunit 6A (zeta 1) | 908 | ENSG00000146731 |
| 2 | P49368 | NA | CCT3 | chaperonin containing TCP1, subunit 3 (gamma) | 7203 | ENSG00000163468 |
| 3 | P78371 | NA | CCT2 | chaperonin containing TCP1, subunit 2 (beta) | 10576 | ENSG00000166226 |
| 4 | P17987 | NA | TCP1 | t-complex 1 | 6950 | ENSG00000120438 |
| 5 | Q99832 | NA | CCT7 | chaperonin containing TCP1, subunit 7 (eta) | 10574 | ENSG00000135624 |
| 6 | P50991 | NA | CCT4 | chaperonin containing TCP1, subunit 4 (delta) | 10575 | ENSG00000115484 |

  
  

| **Database:cellular component      &nbspName:cytoplasm      &nbspID:GO:0005737** | | | | | | |
| --- | --- | --- | --- | --- | --- | --- |
| C=9051; O=58; E=33.64; R=1.72; rawP=9.79e-12; adjP=2.48e-10 | | | | | | |
| Index | UserID | Value | Gene Symbol | Gene Name | EntrezGene | Ensembl |
| 1 | Q92598 | NA | HSPH1 | heat shock 105kDa/110kDa protein 1 | 10808 | ENSG00000120694 |
| 2 | Q13885 | NA | TUBB2A | tubulin, beta 2A class IIa | 7280 | ENSG00000137267 |
| 3 | P68371 | NA | TUBB4B | tubulin, beta 4B class IVb | 10383 | ENSG00000188229 |
| 4 | P25685 | NA | DNAJB1 | DnaJ (Hsp40) homolog, subfamily B, member 1 | 3337 | ENSG00000132002 |
| 5 | P53701 | NA | HCCS | holocytochrome c synthase | 3052 | ENSG00000004961 |
| 6 | P68104 | NA | EEF1A1 | eukaryotic translation elongation factor 1 alpha 1 | 1915 | ENSG00000156508 |
| 7 | O75369 | NA | FLNB | filamin B, beta | 2317 | ENSG00000136068 |
| 8 | P02786 | NA | TFRC | transferrin receptor (p90, CD71) | 7037 | ENSG00000072274 |
| 9 | O00425 | NA | IGF2BP3 | insulin-like growth factor 2 mRNA binding protein 3 | 10643 | ENSG00000136231 |
| 10 | P50991 | NA | CCT4 | chaperonin containing TCP1, subunit 4 (delta) | 10575 | ENSG00000115484 |
| 11 | P22102 | NA | GART | phosphoribosylglycinamide formyltransferase, phosphoribosylglycinamide synthetase, phosphoribosylaminoimidazole synthetase | 2618 | ENSG00000159131 |
| 12 | P13639 | NA | EEF2 | eukaryotic translation elongation factor 2 | 1938 | ENSG00000167658 |
| 13 | Q16555 | NA | DPYSL2 | dihydropyrimidinase-like 2 | 1808 | ENSG00000092964 |
| 14 | O95433 | NA | AHSA1 | AHA1, activator of heat shock 90kDa protein ATPase homolog 1 (yeast) | 10598 | ENSG00000100591 |
| 15 | P41567 | NA | EIF1 | eukaryotic translation initiation factor 1 | 10209 | ENSG00000173812 |
| 16 | P17174 | NA | GOT1 | glutamic-oxaloacetic transaminase 1, soluble (aspartate aminotransferase 1) | 2805 | ENSG00000120053 |
| 17 | P67809 | NA | YBX1 | Y box binding protein 1 | 4904 | ENSG00000065978 |
| 18 | P21980 | NA | TGM2 | transglutaminase 2 (C polypeptide, protein-glutamine-gamma-glutamyltransferase) | 7052 | ENSG00000198959 |
| 19 | Q99832 | NA | CCT7 | chaperonin containing TCP1, subunit 7 (eta) | 10574 | ENSG00000135624 |
| 20 | Q15185 | NA | PTGES3 | prostaglandin E synthase 3 (cytosolic) | 10728 | ENSG00000110958 |
| 21 | Q6PIU2 | NA | NCEH1 | neutral cholesterol ester hydrolase 1 | 57552 | ENSG00000144959 |
| 22 | P18206 | NA | VCL | vinculin | 7414 | ENSG00000035403 |
| 23 | P06703 | NA | S100A6 | S100 calcium binding protein A6 | 6277 | ENSG00000197956 |
| 24 | P11387 | NA | TOP1 | topoisomerase (DNA) I | 7150 | ENSG00000198900 |
| 25 | P11586 | NA | MTHFD1 | methylenetetrahydrofolate dehydrogenase (NADP+ dependent) 1, methenyltetrahydrofolate cyclohydrolase, formyltetrahydrofolate synthetase | 4522 | ENSG00000100714 |
| 26 | Q16881 | NA | TXNRD1 | thioredoxin reductase 1 | 7296 | ENSG00000198431 |
| 27 | P63241 | NA | EIF5A | eukaryotic translation initiation factor 5A | 1984 | ENSG00000132507 |
| 28 | P78371 | NA | CCT2 | chaperonin containing TCP1, subunit 2 (beta) | 10576 | ENSG00000166226 |
| 29 | P08195 | NA | SLC3A2 | solute carrier family 3 (activators of dibasic and neutral amino acid transport), member 2 | 6520 | ENSG00000168003 |
| 30 | Q7Z2K6 | NA | ERMP1 | endoplasmic reticulum metallopeptidase 1 | 79956 | ENSG00000099219 |
| 31 | P02795 | NA | MT2A | metallothionein 2A | 4502 | ENSG00000125148 |
| 32 | P49321 | NA | NASP | nuclear autoantigenic sperm protein (histone-binding) | 4678 | ENSG00000132780 |
| 33 | P00491 | NA | PNP | purine nucleoside phosphorylase | 4860 | ENSG00000198805 |
| 34 | P07951 | NA | TPM2 | tropomyosin 2 (beta) | 7169 | ENSG00000198467 |
| 35 | Q9NZM1 | NA | MYOF | myoferlin | 26509 | ENSG00000138119 |
| 36 | P12081 | NA | HARS | histidyl-tRNA synthetase | 3035 | ENSG00000170445 |
| 37 | P67936 | NA | TPM4 | tropomyosin 4 | 7171 | ENSG00000167460 |
| 38 | P49327 | NA | FASN | fatty acid synthase | 2194 | ENSG00000169710 |
| 39 | P49368 | NA | CCT3 | chaperonin containing TCP1, subunit 3 (gamma) | 7203 | ENSG00000163468 |
| 40 | P56192 | NA | MARS | methionyl-tRNA synthetase | 4141 | ENSG00000166986 |
| 41 | Q9BQE3 | NA | TUBA1C | tubulin, alpha 1c | 84790 | ENSG00000167553 |
| 42 | P12268 | NA | IMPDH2 | IMP (inosine 5'-monophosphate) dehydrogenase 2 | 3615 | ENSG00000178035 |
| 43 | P62906 | NA | RPL10A | ribosomal protein L10a | 4736 | ENSG00000198755 |
| 44 | Q15056 | NA | EIF4H | eukaryotic translation initiation factor 4H | 7458 | ENSG00000106682 |
| 45 | O60610 | NA | DIAPH1 | diaphanous homolog 1 (Drosophila) | 1729 | ENSG00000131504 |
| 46 | P50990 | NA | CCT8 | chaperonin containing TCP1, subunit 8 (theta) | 10694 | ENSG00000156261 |
| 47 | Q06830 | NA | PRDX1 | peroxiredoxin 1 | 5052 | ENSG00000117450 |
| 48 | P07814 | NA | EPRS | glutamyl-prolyl-tRNA synthetase | 2058 | ENSG00000136628 |
| 49 | P17987 | NA | TCP1 | t-complex 1 | 6950 | ENSG00000120438 |
| 50 | P55209 | NA | NAP1L1 | nucleosome assembly protein 1-like 1 | 4673 | ENSG00000187109 |
| 51 | P34932 | NA | HSPA4 | heat shock 70kDa protein 4 | 3308 | ENSG00000170606 |
| 52 | P41091 | NA | EIF2S3 | eukaryotic translation initiation factor 2, subunit 3 gamma, 52kDa | 1968 | ENSG00000130741 |
| 53 | P07108 | NA | DBI | diazepam binding inhibitor (GABA receptor modulator, acyl-CoA binding protein) | 1622 | ENSG00000155368 |
| 54 | P40227 | NA | CCT6A | chaperonin containing TCP1, subunit 6A (zeta 1) | 908 | ENSG00000146731 |
| 55 | P52292 | NA | KPNA2 | karyopherin alpha 2 (RAG cohort 1, importin alpha 1) | 3838 | ENSG00000182481 |
| 56 | Q15758 | NA | SLC1A5 | solute carrier family 1 (neutral amino acid transporter), member 5 | 6510 | ENSG00000105281 |
| 57 | P00352 | NA | ALDH1A1 | aldehyde dehydrogenase 1 family, member A1 | 216 | ENSG00000165092 |
| 58 | Q8TEX9 | NA | IPO4 | importin 4 | 79711 | ENSG00000196497 |

  
  

| **Database:cellular component      &nbspName:cytoplasmic part      &nbspID:GO:0044444** | | | | | | |
| --- | --- | --- | --- | --- | --- | --- |
| C=6728; O=51; E=25.01; R=2.04; rawP=1.43e-11; adjP=2.72e-10 | | | | | | |
| Index | UserID | Value | Gene Symbol | Gene Name | EntrezGene | Ensembl |
| 1 | P68371 | NA | TUBB4B | tubulin, beta 4B class IVb | 10383 | ENSG00000188229 |
| 2 | P53701 | NA | HCCS | holocytochrome c synthase | 3052 | ENSG00000004961 |
| 3 | P68104 | NA | EEF1A1 | eukaryotic translation elongation factor 1 alpha 1 | 1915 | ENSG00000156508 |
| 4 | O75369 | NA | FLNB | filamin B, beta | 2317 | ENSG00000136068 |
| 5 | P02786 | NA | TFRC | transferrin receptor (p90, CD71) | 7037 | ENSG00000072274 |
| 6 | P50991 | NA | CCT4 | chaperonin containing TCP1, subunit 4 (delta) | 10575 | ENSG00000115484 |
| 7 | O00425 | NA | IGF2BP3 | insulin-like growth factor 2 mRNA binding protein 3 | 10643 | ENSG00000136231 |
| 8 | P22102 | NA | GART | phosphoribosylglycinamide formyltransferase, phosphoribosylglycinamide synthetase, phosphoribosylaminoimidazole synthetase | 2618 | ENSG00000159131 |
| 9 | P13639 | NA | EEF2 | eukaryotic translation elongation factor 2 | 1938 | ENSG00000167658 |
| 10 | Q16555 | NA | DPYSL2 | dihydropyrimidinase-like 2 | 1808 | ENSG00000092964 |
| 11 | O95433 | NA | AHSA1 | AHA1, activator of heat shock 90kDa protein ATPase homolog 1 (yeast) | 10598 | ENSG00000100591 |
| 12 | P17174 | NA | GOT1 | glutamic-oxaloacetic transaminase 1, soluble (aspartate aminotransferase 1) | 2805 | ENSG00000120053 |
| 13 | P67809 | NA | YBX1 | Y box binding protein 1 | 4904 | ENSG00000065978 |
| 14 | Q15185 | NA | PTGES3 | prostaglandin E synthase 3 (cytosolic) | 10728 | ENSG00000110958 |
| 15 | Q99832 | NA | CCT7 | chaperonin containing TCP1, subunit 7 (eta) | 10574 | ENSG00000135624 |
| 16 | P21980 | NA | TGM2 | transglutaminase 2 (C polypeptide, protein-glutamine-gamma-glutamyltransferase) | 7052 | ENSG00000198959 |
| 17 | Q6PIU2 | NA | NCEH1 | neutral cholesterol ester hydrolase 1 | 57552 | ENSG00000144959 |
| 18 | P18206 | NA | VCL | vinculin | 7414 | ENSG00000035403 |
| 19 | P06703 | NA | S100A6 | S100 calcium binding protein A6 | 6277 | ENSG00000197956 |
| 20 | P11387 | NA | TOP1 | topoisomerase (DNA) I | 7150 | ENSG00000198900 |
| 21 | P11586 | NA | MTHFD1 | methylenetetrahydrofolate dehydrogenase (NADP+ dependent) 1, methenyltetrahydrofolate cyclohydrolase, formyltetrahydrofolate synthetase | 4522 | ENSG00000100714 |
| 22 | Q16881 | NA | TXNRD1 | thioredoxin reductase 1 | 7296 | ENSG00000198431 |
| 23 | P63241 | NA | EIF5A | eukaryotic translation initiation factor 5A | 1984 | ENSG00000132507 |
| 24 | P78371 | NA | CCT2 | chaperonin containing TCP1, subunit 2 (beta) | 10576 | ENSG00000166226 |
| 25 | P08195 | NA | SLC3A2 | solute carrier family 3 (activators of dibasic and neutral amino acid transport), member 2 | 6520 | ENSG00000168003 |
| 26 | Q7Z2K6 | NA | ERMP1 | endoplasmic reticulum metallopeptidase 1 | 79956 | ENSG00000099219 |
| 27 | P02795 | NA | MT2A | metallothionein 2A | 4502 | ENSG00000125148 |
| 28 | P07951 | NA | TPM2 | tropomyosin 2 (beta) | 7169 | ENSG00000198467 |
| 29 | Q9NZM1 | NA | MYOF | myoferlin | 26509 | ENSG00000138119 |
| 30 | P00491 | NA | PNP | purine nucleoside phosphorylase | 4860 | ENSG00000198805 |
| 31 | P12081 | NA | HARS | histidyl-tRNA synthetase | 3035 | ENSG00000170445 |
| 32 | P49327 | NA | FASN | fatty acid synthase | 2194 | ENSG00000169710 |
| 33 | P67936 | NA | TPM4 | tropomyosin 4 | 7171 | ENSG00000167460 |
| 34 | P49368 | NA | CCT3 | chaperonin containing TCP1, subunit 3 (gamma) | 7203 | ENSG00000163468 |
| 35 | P56192 | NA | MARS | methionyl-tRNA synthetase | 4141 | ENSG00000166986 |
| 36 | Q9BQE3 | NA | TUBA1C | tubulin, alpha 1c | 84790 | ENSG00000167553 |
| 37 | P12268 | NA | IMPDH2 | IMP (inosine 5'-monophosphate) dehydrogenase 2 | 3615 | ENSG00000178035 |
| 38 | P62906 | NA | RPL10A | ribosomal protein L10a | 4736 | ENSG00000198755 |
| 39 | Q15056 | NA | EIF4H | eukaryotic translation initiation factor 4H | 7458 | ENSG00000106682 |
| 40 | P50990 | NA | CCT8 | chaperonin containing TCP1, subunit 8 (theta) | 10694 | ENSG00000156261 |
| 41 | Q06830 | NA | PRDX1 | peroxiredoxin 1 | 5052 | ENSG00000117450 |
| 42 | P07814 | NA | EPRS | glutamyl-prolyl-tRNA synthetase | 2058 | ENSG00000136628 |
| 43 | P17987 | NA | TCP1 | t-complex 1 | 6950 | ENSG00000120438 |
| 44 | P55209 | NA | NAP1L1 | nucleosome assembly protein 1-like 1 | 4673 | ENSG00000187109 |
| 45 | P34932 | NA | HSPA4 | heat shock 70kDa protein 4 | 3308 | ENSG00000170606 |
| 46 | P41091 | NA | EIF2S3 | eukaryotic translation initiation factor 2, subunit 3 gamma, 52kDa | 1968 | ENSG00000130741 |
| 47 | P07108 | NA | DBI | diazepam binding inhibitor (GABA receptor modulator, acyl-CoA binding protein) | 1622 | ENSG00000155368 |
| 48 | P40227 | NA | CCT6A | chaperonin containing TCP1, subunit 6A (zeta 1) | 908 | ENSG00000146731 |
| 49 | P52292 | NA | KPNA2 | karyopherin alpha 2 (RAG cohort 1, importin alpha 1) | 3838 | ENSG00000182481 |
| 50 | Q15758 | NA | SLC1A5 | solute carrier family 1 (neutral amino acid transporter), member 5 | 6510 | ENSG00000105281 |
| 51 | P00352 | NA | ALDH1A1 | aldehyde dehydrogenase 1 family, member A1 | 216 | ENSG00000165092 |

  
  

| **Database:cellular component      &nbspName:intracellular part      &nbspID:GO:0044424** | | | | | | |
| --- | --- | --- | --- | --- | --- | --- |
| C=12096; O=62; E=44.96; R=1.38; rawP=2.12e-09; adjP=3.22e-08 | | | | | | |
| Index | UserID | Value | Gene Symbol | Gene Name | EntrezGene | Ensembl |
| 1 | Q92598 | NA | HSPH1 | heat shock 105kDa/110kDa protein 1 | 10808 | ENSG00000120694 |
| 2 | Q13885 | NA | TUBB2A | tubulin, beta 2A class IIa | 7280 | ENSG00000137267 |
| 3 | P68371 | NA | TUBB4B | tubulin, beta 4B class IVb | 10383 | ENSG00000188229 |
| 4 | P25685 | NA | DNAJB1 | DnaJ (Hsp40) homolog, subfamily B, member 1 | 3337 | ENSG00000132002 |
| 5 | P53701 | NA | HCCS | holocytochrome c synthase | 3052 | ENSG00000004961 |
| 6 | P68104 | NA | EEF1A1 | eukaryotic translation elongation factor 1 alpha 1 | 1915 | ENSG00000156508 |
| 7 | O75369 | NA | FLNB | filamin B, beta | 2317 | ENSG00000136068 |
| 8 | P02786 | NA | TFRC | transferrin receptor (p90, CD71) | 7037 | ENSG00000072274 |
| 9 | O00425 | NA | IGF2BP3 | insulin-like growth factor 2 mRNA binding protein 3 | 10643 | ENSG00000136231 |
| 10 | P50991 | NA | CCT4 | chaperonin containing TCP1, subunit 4 (delta) | 10575 | ENSG00000115484 |
| 11 | P22102 | NA | GART | phosphoribosylglycinamide formyltransferase, phosphoribosylglycinamide synthetase, phosphoribosylaminoimidazole synthetase | 2618 | ENSG00000159131 |
| 12 | P13639 | NA | EEF2 | eukaryotic translation elongation factor 2 | 1938 | ENSG00000167658 |
| 13 | Q16555 | NA | DPYSL2 | dihydropyrimidinase-like 2 | 1808 | ENSG00000092964 |
| 14 | O95433 | NA | AHSA1 | AHA1, activator of heat shock 90kDa protein ATPase homolog 1 (yeast) | 10598 | ENSG00000100591 |
| 15 | P41567 | NA | EIF1 | eukaryotic translation initiation factor 1 | 10209 | ENSG00000173812 |
| 16 | P17174 | NA | GOT1 | glutamic-oxaloacetic transaminase 1, soluble (aspartate aminotransferase 1) | 2805 | ENSG00000120053 |
| 17 | P67809 | NA | YBX1 | Y box binding protein 1 | 4904 | ENSG00000065978 |
| 18 | P21980 | NA | TGM2 | transglutaminase 2 (C polypeptide, protein-glutamine-gamma-glutamyltransferase) | 7052 | ENSG00000198959 |
| 19 | P37802 | NA | TAGLN2 | transgelin 2 | 8407 | ENSG00000158710 |
| 20 | Q99832 | NA | CCT7 | chaperonin containing TCP1, subunit 7 (eta) | 10574 | ENSG00000135624 |
| 21 | Q15185 | NA | PTGES3 | prostaglandin E synthase 3 (cytosolic) | 10728 | ENSG00000110958 |
| 22 | Q6PIU2 | NA | NCEH1 | neutral cholesterol ester hydrolase 1 | 57552 | ENSG00000144959 |
| 23 | P18206 | NA | VCL | vinculin | 7414 | ENSG00000035403 |
| 24 | P06703 | NA | S100A6 | S100 calcium binding protein A6 | 6277 | ENSG00000197956 |
| 25 | Q9UK76 | NA | HN1 | hematological and neurological expressed 1 | 51155 | ENSG00000189159 |
| 26 | P11387 | NA | TOP1 | topoisomerase (DNA) I | 7150 | ENSG00000198900 |
| 27 | P11586 | NA | MTHFD1 | methylenetetrahydrofolate dehydrogenase (NADP+ dependent) 1, methenyltetrahydrofolate cyclohydrolase, formyltetrahydrofolate synthetase | 4522 | ENSG00000100714 |
| 28 | Q16881 | NA | TXNRD1 | thioredoxin reductase 1 | 7296 | ENSG00000198431 |
| 29 | P63241 | NA | EIF5A | eukaryotic translation initiation factor 5A | 1984 | ENSG00000132507 |
| 30 | P78371 | NA | CCT2 | chaperonin containing TCP1, subunit 2 (beta) | 10576 | ENSG00000166226 |
| 31 | Q9NR30 | NA | DDX21 | DEAD (Asp-Glu-Ala-Asp) box helicase 21 | 9188 | ENSG00000165732 |
| 32 | P02795 | NA | MT2A | metallothionein 2A | 4502 | ENSG00000125148 |
| 33 | Q7Z2K6 | NA | ERMP1 | endoplasmic reticulum metallopeptidase 1 | 79956 | ENSG00000099219 |
| 34 | P08195 | NA | SLC3A2 | solute carrier family 3 (activators of dibasic and neutral amino acid transport), member 2 | 6520 | ENSG00000168003 |
| 35 | P49321 | NA | NASP | nuclear autoantigenic sperm protein (histone-binding) | 4678 | ENSG00000132780 |
| 36 | P00491 | NA | PNP | purine nucleoside phosphorylase | 4860 | ENSG00000198805 |
| 37 | Q9NZM1 | NA | MYOF | myoferlin | 26509 | ENSG00000138119 |
| 38 | P07951 | NA | TPM2 | tropomyosin 2 (beta) | 7169 | ENSG00000198467 |
| 39 | P12081 | NA | HARS | histidyl-tRNA synthetase | 3035 | ENSG00000170445 |
| 40 | P67936 | NA | TPM4 | tropomyosin 4 | 7171 | ENSG00000167460 |
| 41 | P49327 | NA | FASN | fatty acid synthase | 2194 | ENSG00000169710 |
| 42 | P49368 | NA | CCT3 | chaperonin containing TCP1, subunit 3 (gamma) | 7203 | ENSG00000163468 |
| 43 | P56192 | NA | MARS | methionyl-tRNA synthetase | 4141 | ENSG00000166986 |
| 44 | Q9BQE3 | NA | TUBA1C | tubulin, alpha 1c | 84790 | ENSG00000167553 |
| 45 | P12268 | NA | IMPDH2 | IMP (inosine 5'-monophosphate) dehydrogenase 2 | 3615 | ENSG00000178035 |
| 46 | P62906 | NA | RPL10A | ribosomal protein L10a | 4736 | ENSG00000198755 |
| 47 | P49736 | NA | MCM2 | minichromosome maintenance complex component 2 | 4171 | ENSG00000073111 |
| 48 | Q15056 | NA | EIF4H | eukaryotic translation initiation factor 4H | 7458 | ENSG00000106682 |
| 49 | O60610 | NA | DIAPH1 | diaphanous homolog 1 (Drosophila) | 1729 | ENSG00000131504 |
| 50 | P50990 | NA | CCT8 | chaperonin containing TCP1, subunit 8 (theta) | 10694 | ENSG00000156261 |
| 51 | Q06830 | NA | PRDX1 | peroxiredoxin 1 | 5052 | ENSG00000117450 |
| 52 | P07814 | NA | EPRS | glutamyl-prolyl-tRNA synthetase | 2058 | ENSG00000136628 |
| 53 | P17987 | NA | TCP1 | t-complex 1 | 6950 | ENSG00000120438 |
| 54 | P55209 | NA | NAP1L1 | nucleosome assembly protein 1-like 1 | 4673 | ENSG00000187109 |
| 55 | P34932 | NA | HSPA4 | heat shock 70kDa protein 4 | 3308 | ENSG00000170606 |
| 56 | P41091 | NA | EIF2S3 | eukaryotic translation initiation factor 2, subunit 3 gamma, 52kDa | 1968 | ENSG00000130741 |
| 57 | P07108 | NA | DBI | diazepam binding inhibitor (GABA receptor modulator, acyl-CoA binding protein) | 1622 | ENSG00000155368 |
| 58 | P40227 | NA | CCT6A | chaperonin containing TCP1, subunit 6A (zeta 1) | 908 | ENSG00000146731 |
| 59 | P52292 | NA | KPNA2 | karyopherin alpha 2 (RAG cohort 1, importin alpha 1) | 3838 | ENSG00000182481 |
| 60 | Q15758 | NA | SLC1A5 | solute carrier family 1 (neutral amino acid transporter), member 5 | 6510 | ENSG00000105281 |
| 61 | P00352 | NA | ALDH1A1 | aldehyde dehydrogenase 1 family, member A1 | 216 | ENSG00000165092 |
| 62 | Q8TEX9 | NA | IPO4 | importin 4 | 79711 | ENSG00000196497 |

  
  

| **Database:cellular component      &nbspName:intracellular      &nbspID:GO:0005622** | | | | | | |
| --- | --- | --- | --- | --- | --- | --- |
| C=12412; O=62; E=46.13; R=1.34; rawP=1.05e-08; adjP=1.33e-07 | | | | | | |
| Index | UserID | Value | Gene Symbol | Gene Name | EntrezGene | Ensembl |
| 1 | Q92598 | NA | HSPH1 | heat shock 105kDa/110kDa protein 1 | 10808 | ENSG00000120694 |
| 2 | Q13885 | NA | TUBB2A | tubulin, beta 2A class IIa | 7280 | ENSG00000137267 |
| 3 | P68371 | NA | TUBB4B | tubulin, beta 4B class IVb | 10383 | ENSG00000188229 |
| 4 | P25685 | NA | DNAJB1 | DnaJ (Hsp40) homolog, subfamily B, member 1 | 3337 | ENSG00000132002 |
| 5 | P53701 | NA | HCCS | holocytochrome c synthase | 3052 | ENSG00000004961 |
| 6 | P68104 | NA | EEF1A1 | eukaryotic translation elongation factor 1 alpha 1 | 1915 | ENSG00000156508 |
| 7 | O75369 | NA | FLNB | filamin B, beta | 2317 | ENSG00000136068 |
| 8 | P02786 | NA | TFRC | transferrin receptor (p90, CD71) | 7037 | ENSG00000072274 |
| 9 | O00425 | NA | IGF2BP3 | insulin-like growth factor 2 mRNA binding protein 3 | 10643 | ENSG00000136231 |
| 10 | P50991 | NA | CCT4 | chaperonin containing TCP1, subunit 4 (delta) | 10575 | ENSG00000115484 |
| 11 | P22102 | NA | GART | phosphoribosylglycinamide formyltransferase, phosphoribosylglycinamide synthetase, phosphoribosylaminoimidazole synthetase | 2618 | ENSG00000159131 |
| 12 | P13639 | NA | EEF2 | eukaryotic translation elongation factor 2 | 1938 | ENSG00000167658 |
| 13 | Q16555 | NA | DPYSL2 | dihydropyrimidinase-like 2 | 1808 | ENSG00000092964 |
| 14 | O95433 | NA | AHSA1 | AHA1, activator of heat shock 90kDa protein ATPase homolog 1 (yeast) | 10598 | ENSG00000100591 |
| 15 | P41567 | NA | EIF1 | eukaryotic translation initiation factor 1 | 10209 | ENSG00000173812 |
| 16 | P17174 | NA | GOT1 | glutamic-oxaloacetic transaminase 1, soluble (aspartate aminotransferase 1) | 2805 | ENSG00000120053 |
| 17 | P67809 | NA | YBX1 | Y box binding protein 1 | 4904 | ENSG00000065978 |
| 18 | P21980 | NA | TGM2 | transglutaminase 2 (C polypeptide, protein-glutamine-gamma-glutamyltransferase) | 7052 | ENSG00000198959 |
| 19 | P37802 | NA | TAGLN2 | transgelin 2 | 8407 | ENSG00000158710 |
| 20 | Q99832 | NA | CCT7 | chaperonin containing TCP1, subunit 7 (eta) | 10574 | ENSG00000135624 |
| 21 | Q15185 | NA | PTGES3 | prostaglandin E synthase 3 (cytosolic) | 10728 | ENSG00000110958 |
| 22 | Q6PIU2 | NA | NCEH1 | neutral cholesterol ester hydrolase 1 | 57552 | ENSG00000144959 |
| 23 | P18206 | NA | VCL | vinculin | 7414 | ENSG00000035403 |
| 24 | P06703 | NA | S100A6 | S100 calcium binding protein A6 | 6277 | ENSG00000197956 |
| 25 | Q9UK76 | NA | HN1 | hematological and neurological expressed 1 | 51155 | ENSG00000189159 |
| 26 | P11387 | NA | TOP1 | topoisomerase (DNA) I | 7150 | ENSG00000198900 |
| 27 | P11586 | NA | MTHFD1 | methylenetetrahydrofolate dehydrogenase (NADP+ dependent) 1, methenyltetrahydrofolate cyclohydrolase, formyltetrahydrofolate synthetase | 4522 | ENSG00000100714 |
| 28 | Q16881 | NA | TXNRD1 | thioredoxin reductase 1 | 7296 | ENSG00000198431 |
| 29 | P63241 | NA | EIF5A | eukaryotic translation initiation factor 5A | 1984 | ENSG00000132507 |
| 30 | P78371 | NA | CCT2 | chaperonin containing TCP1, subunit 2 (beta) | 10576 | ENSG00000166226 |
| 31 | Q9NR30 | NA | DDX21 | DEAD (Asp-Glu-Ala-Asp) box helicase 21 | 9188 | ENSG00000165732 |
| 32 | P02795 | NA | MT2A | metallothionein 2A | 4502 | ENSG00000125148 |
| 33 | Q7Z2K6 | NA | ERMP1 | endoplasmic reticulum metallopeptidase 1 | 79956 | ENSG00000099219 |
| 34 | P08195 | NA | SLC3A2 | solute carrier family 3 (activators of dibasic and neutral amino acid transport), member 2 | 6520 | ENSG00000168003 |
| 35 | P49321 | NA | NASP | nuclear autoantigenic sperm protein (histone-binding) | 4678 | ENSG00000132780 |
| 36 | P00491 | NA | PNP | purine nucleoside phosphorylase | 4860 | ENSG00000198805 |
| 37 | Q9NZM1 | NA | MYOF | myoferlin | 26509 | ENSG00000138119 |
| 38 | P07951 | NA | TPM2 | tropomyosin 2 (beta) | 7169 | ENSG00000198467 |
| 39 | P12081 | NA | HARS | histidyl-tRNA synthetase | 3035 | ENSG00000170445 |
| 40 | P67936 | NA | TPM4 | tropomyosin 4 | 7171 | ENSG00000167460 |
| 41 | P49327 | NA | FASN | fatty acid synthase | 2194 | ENSG00000169710 |
| 42 | P49368 | NA | CCT3 | chaperonin containing TCP1, subunit 3 (gamma) | 7203 | ENSG00000163468 |
| 43 | P56192 | NA | MARS | methionyl-tRNA synthetase | 4141 | ENSG00000166986 |
| 44 | Q9BQE3 | NA | TUBA1C | tubulin, alpha 1c | 84790 | ENSG00000167553 |
| 45 | P12268 | NA | IMPDH2 | IMP (inosine 5'-monophosphate) dehydrogenase 2 | 3615 | ENSG00000178035 |
| 46 | P62906 | NA | RPL10A | ribosomal protein L10a | 4736 | ENSG00000198755 |
| 47 | P49736 | NA | MCM2 | minichromosome maintenance complex component 2 | 4171 | ENSG00000073111 |
| 48 | Q15056 | NA | EIF4H | eukaryotic translation initiation factor 4H | 7458 | ENSG00000106682 |
| 49 | O60610 | NA | DIAPH1 | diaphanous homolog 1 (Drosophila) | 1729 | ENSG00000131504 |
| 50 | P50990 | NA | CCT8 | chaperonin containing TCP1, subunit 8 (theta) | 10694 | ENSG00000156261 |
| 51 | Q06830 | NA | PRDX1 | peroxiredoxin 1 | 5052 | ENSG00000117450 |
| 52 | P07814 | NA | EPRS | glutamyl-prolyl-tRNA synthetase | 2058 | ENSG00000136628 |
| 53 | P17987 | NA | TCP1 | t-complex 1 | 6950 | ENSG00000120438 |
| 54 | P55209 | NA | NAP1L1 | nucleosome assembly protein 1-like 1 | 4673 | ENSG00000187109 |
| 55 | P34932 | NA | HSPA4 | heat shock 70kDa protein 4 | 3308 | ENSG00000170606 |
| 56 | P41091 | NA | EIF2S3 | eukaryotic translation initiation factor 2, subunit 3 gamma, 52kDa | 1968 | ENSG00000130741 |
| 57 | P07108 | NA | DBI | diazepam binding inhibitor (GABA receptor modulator, acyl-CoA binding protein) | 1622 | ENSG00000155368 |
| 58 | P40227 | NA | CCT6A | chaperonin containing TCP1, subunit 6A (zeta 1) | 908 | ENSG00000146731 |
| 59 | P52292 | NA | KPNA2 | karyopherin alpha 2 (RAG cohort 1, importin alpha 1) | 3838 | ENSG00000182481 |
| 60 | Q15758 | NA | SLC1A5 | solute carrier family 1 (neutral amino acid transporter), member 5 | 6510 | ENSG00000105281 |
| 61 | P00352 | NA | ALDH1A1 | aldehyde dehydrogenase 1 family, member A1 | 216 | ENSG00000165092 |
| 62 | Q8TEX9 | NA | IPO4 | importin 4 | 79711 | ENSG00000196497 |

  
  

| **Database:cellular component      &nbspName:microtubule      &nbspID:GO:0005874** | | | | | | |
| --- | --- | --- | --- | --- | --- | --- |
| C=334; O=11; E=1.24; R=8.86; rawP=3.59e-08; adjP=3.90e-07 | | | | | | |
| Index | UserID | Value | Gene Symbol | Gene Name | EntrezGene | Ensembl |
| 1 | P78371 | NA | CCT2 | chaperonin containing TCP1, subunit 2 (beta) | 10576 | ENSG00000166226 |
| 2 | Q92598 | NA | HSPH1 | heat shock 105kDa/110kDa protein 1 | 10808 | ENSG00000120694 |
| 3 | P17987 | NA | TCP1 | t-complex 1 | 6950 | ENSG00000120438 |
| 4 | Q13885 | NA | TUBB2A | tubulin, beta 2A class IIa | 7280 | ENSG00000137267 |
| 5 | P68371 | NA | TUBB4B | tubulin, beta 4B class IVb | 10383 | ENSG00000188229 |
| 6 | P40227 | NA | CCT6A | chaperonin containing TCP1, subunit 6A (zeta 1) | 908 | ENSG00000146731 |
| 7 | P49368 | NA | CCT3 | chaperonin containing TCP1, subunit 3 (gamma) | 7203 | ENSG00000163468 |
| 8 | Q9BQE3 | NA | TUBA1C | tubulin, alpha 1c | 84790 | ENSG00000167553 |
| 9 | Q99832 | NA | CCT7 | chaperonin containing TCP1, subunit 7 (eta) | 10574 | ENSG00000135624 |
| 10 | P50990 | NA | CCT8 | chaperonin containing TCP1, subunit 8 (theta) | 10694 | ENSG00000156261 |
| 11 | P50991 | NA | CCT4 | chaperonin containing TCP1, subunit 4 (delta) | 10575 | ENSG00000115484 |

  
  

| **Database:cellular component      &nbspName:melanosome      &nbspID:GO:0042470** | | | | | | |
| --- | --- | --- | --- | --- | --- | --- |
| C=93; O=7; E=0.35; R=20.25; rawP=5.10e-08; adjP=4.31e-07 | | | | | | |
| Index | UserID | Value | Gene Symbol | Gene Name | EntrezGene | Ensembl |
| 1 | P55209 | NA | NAP1L1 | nucleosome assembly protein 1-like 1 | 4673 | ENSG00000187109 |
| 2 | P08195 | NA | SLC3A2 | solute carrier family 3 (activators of dibasic and neutral amino acid transport), member 2 | 6520 | ENSG00000168003 |
| 3 | Q15758 | NA | SLC1A5 | solute carrier family 1 (neutral amino acid transporter), member 5 | 6510 | ENSG00000105281 |
| 4 | Q06830 | NA | PRDX1 | peroxiredoxin 1 | 5052 | ENSG00000117450 |
| 5 | P02786 | NA | TFRC | transferrin receptor (p90, CD71) | 7037 | ENSG00000072274 |
| 6 | P49327 | NA | FASN | fatty acid synthase | 2194 | ENSG00000169710 |
| 7 | P50991 | NA | CCT4 | chaperonin containing TCP1, subunit 4 (delta) | 10575 | ENSG00000115484 |

  
  

| **Database:cellular component      &nbspName:pigment granule      &nbspID:GO:0048770** | | | | | | |
| --- | --- | --- | --- | --- | --- | --- |
| C=93; O=7; E=0.35; R=20.25; rawP=5.10e-08; adjP=4.31e-07 | | | | | | |
| Index | UserID | Value | Gene Symbol | Gene Name | EntrezGene | Ensembl |
| 1 | P55209 | NA | NAP1L1 | nucleosome assembly protein 1-like 1 | 4673 | ENSG00000187109 |
| 2 | P08195 | NA | SLC3A2 | solute carrier family 3 (activators of dibasic and neutral amino acid transport), member 2 | 6520 | ENSG00000168003 |
| 3 | Q15758 | NA | SLC1A5 | solute carrier family 1 (neutral amino acid transporter), member 5 | 6510 | ENSG00000105281 |
| 4 | Q06830 | NA | PRDX1 | peroxiredoxin 1 | 5052 | ENSG00000117450 |
| 5 | P02786 | NA | TFRC | transferrin receptor (p90, CD71) | 7037 | ENSG00000072274 |
| 6 | P49327 | NA | FASN | fatty acid synthase | 2194 | ENSG00000169710 |
| 7 | P50991 | NA | CCT4 | chaperonin containing TCP1, subunit 4 (delta) | 10575 | ENSG00000115484 |

  
  

| **Database:cellular component      &nbspName:cytosolic part      &nbspID:GO:0044445** | | | | | | |
| --- | --- | --- | --- | --- | --- | --- |
| C=187; O=7; E=0.70; R=10.07; rawP=5.81e-06; adjP=4.42e-05 | | | | | | |
| Index | UserID | Value | Gene Symbol | Gene Name | EntrezGene | Ensembl |
| 1 | P40227 | NA | CCT6A | chaperonin containing TCP1, subunit 6A (zeta 1) | 908 | ENSG00000146731 |
| 2 | P49368 | NA | CCT3 | chaperonin containing TCP1, subunit 3 (gamma) | 7203 | ENSG00000163468 |
| 3 | P78371 | NA | CCT2 | chaperonin containing TCP1, subunit 2 (beta) | 10576 | ENSG00000166226 |
| 4 | P17987 | NA | TCP1 | t-complex 1 | 6950 | ENSG00000120438 |
| 5 | Q99832 | NA | CCT7 | chaperonin containing TCP1, subunit 7 (eta) | 10574 | ENSG00000135624 |
| 6 | P62906 | NA | RPL10A | ribosomal protein L10a | 4736 | ENSG00000198755 |
| 7 | P50991 | NA | CCT4 | chaperonin containing TCP1, subunit 4 (delta) | 10575 | ENSG00000115484 |

  
  

| **Database:cellular component      &nbspName:cytoskeletal part      &nbspID:GO:0044430** | | | | | | |
| --- | --- | --- | --- | --- | --- | --- |
| C=1282; O=16; E=4.76; R=3.36; rawP=1.25e-05; adjP=8.64e-05 | | | | | | |
| Index | UserID | Value | Gene Symbol | Gene Name | EntrezGene | Ensembl |
| 1 | P78371 | NA | CCT2 | chaperonin containing TCP1, subunit 2 (beta) | 10576 | ENSG00000166226 |
| 2 | Q92598 | NA | HSPH1 | heat shock 105kDa/110kDa protein 1 | 10808 | ENSG00000120694 |
| 3 | Q13885 | NA | TUBB2A | tubulin, beta 2A class IIa | 7280 | ENSG00000137267 |
| 4 | P07951 | NA | TPM2 | tropomyosin 2 (beta) | 7169 | ENSG00000198467 |
| 5 | P68371 | NA | TUBB4B | tubulin, beta 4B class IVb | 10383 | ENSG00000188229 |
| 6 | P67936 | NA | TPM4 | tropomyosin 4 | 7171 | ENSG00000167460 |
| 7 | P49368 | NA | CCT3 | chaperonin containing TCP1, subunit 3 (gamma) | 7203 | ENSG00000163468 |
| 8 | Q9BQE3 | NA | TUBA1C | tubulin, alpha 1c | 84790 | ENSG00000167553 |
| 9 | O60610 | NA | DIAPH1 | diaphanous homolog 1 (Drosophila) | 1729 | ENSG00000131504 |
| 10 | P50990 | NA | CCT8 | chaperonin containing TCP1, subunit 8 (theta) | 10694 | ENSG00000156261 |
| 11 | O75369 | NA | FLNB | filamin B, beta | 2317 | ENSG00000136068 |
| 12 | P50991 | NA | CCT4 | chaperonin containing TCP1, subunit 4 (delta) | 10575 | ENSG00000115484 |
| 13 | P17987 | NA | TCP1 | t-complex 1 | 6950 | ENSG00000120438 |
| 14 | P40227 | NA | CCT6A | chaperonin containing TCP1, subunit 6A (zeta 1) | 908 | ENSG00000146731 |
| 15 | Q99832 | NA | CCT7 | chaperonin containing TCP1, subunit 7 (eta) | 10574 | ENSG00000135624 |
| 16 | P18206 | NA | VCL | vinculin | 7414 | ENSG00000035403 |

  
  

| **Database:cellular component      &nbspName:intracellular organelle      &nbspID:GO:0043229** | | | | | | |
| --- | --- | --- | --- | --- | --- | --- |
| C=10521; O=54; E=39.10; R=1.38; rawP=2.31e-05; adjP=0.0001 | | | | | | |
| Index | UserID | Value | Gene Symbol | Gene Name | EntrezGene | Ensembl |
| 1 | Q92598 | NA | HSPH1 | heat shock 105kDa/110kDa protein 1 | 10808 | ENSG00000120694 |
| 2 | Q13885 | NA | TUBB2A | tubulin, beta 2A class IIa | 7280 | ENSG00000137267 |
| 3 | P68371 | NA | TUBB4B | tubulin, beta 4B class IVb | 10383 | ENSG00000188229 |
| 4 | P25685 | NA | DNAJB1 | DnaJ (Hsp40) homolog, subfamily B, member 1 | 3337 | ENSG00000132002 |
| 5 | P53701 | NA | HCCS | holocytochrome c synthase | 3052 | ENSG00000004961 |
| 6 | P68104 | NA | EEF1A1 | eukaryotic translation elongation factor 1 alpha 1 | 1915 | ENSG00000156508 |
| 7 | O75369 | NA | FLNB | filamin B, beta | 2317 | ENSG00000136068 |
| 8 | P02786 | NA | TFRC | transferrin receptor (p90, CD71) | 7037 | ENSG00000072274 |
| 9 | O00425 | NA | IGF2BP3 | insulin-like growth factor 2 mRNA binding protein 3 | 10643 | ENSG00000136231 |
| 10 | P50991 | NA | CCT4 | chaperonin containing TCP1, subunit 4 (delta) | 10575 | ENSG00000115484 |
| 11 | Q16555 | NA | DPYSL2 | dihydropyrimidinase-like 2 | 1808 | ENSG00000092964 |
| 12 | O95433 | NA | AHSA1 | AHA1, activator of heat shock 90kDa protein ATPase homolog 1 (yeast) | 10598 | ENSG00000100591 |
| 13 | P17174 | NA | GOT1 | glutamic-oxaloacetic transaminase 1, soluble (aspartate aminotransferase 1) | 2805 | ENSG00000120053 |
| 14 | P67809 | NA | YBX1 | Y box binding protein 1 | 4904 | ENSG00000065978 |
| 15 | P21980 | NA | TGM2 | transglutaminase 2 (C polypeptide, protein-glutamine-gamma-glutamyltransferase) | 7052 | ENSG00000198959 |
| 16 | P37802 | NA | TAGLN2 | transgelin 2 | 8407 | ENSG00000158710 |
| 17 | Q15185 | NA | PTGES3 | prostaglandin E synthase 3 (cytosolic) | 10728 | ENSG00000110958 |
| 18 | Q99832 | NA | CCT7 | chaperonin containing TCP1, subunit 7 (eta) | 10574 | ENSG00000135624 |
| 19 | Q6PIU2 | NA | NCEH1 | neutral cholesterol ester hydrolase 1 | 57552 | ENSG00000144959 |
| 20 | P18206 | NA | VCL | vinculin | 7414 | ENSG00000035403 |
| 21 | P06703 | NA | S100A6 | S100 calcium binding protein A6 | 6277 | ENSG00000197956 |
| 22 | Q9UK76 | NA | HN1 | hematological and neurological expressed 1 | 51155 | ENSG00000189159 |
| 23 | P11387 | NA | TOP1 | topoisomerase (DNA) I | 7150 | ENSG00000198900 |
| 24 | P11586 | NA | MTHFD1 | methylenetetrahydrofolate dehydrogenase (NADP+ dependent) 1, methenyltetrahydrofolate cyclohydrolase, formyltetrahydrofolate synthetase | 4522 | ENSG00000100714 |
| 25 | Q16881 | NA | TXNRD1 | thioredoxin reductase 1 | 7296 | ENSG00000198431 |
| 26 | P63241 | NA | EIF5A | eukaryotic translation initiation factor 5A | 1984 | ENSG00000132507 |
| 27 | P78371 | NA | CCT2 | chaperonin containing TCP1, subunit 2 (beta) | 10576 | ENSG00000166226 |
| 28 | Q9NR30 | NA | DDX21 | DEAD (Asp-Glu-Ala-Asp) box helicase 21 | 9188 | ENSG00000165732 |
| 29 | P08195 | NA | SLC3A2 | solute carrier family 3 (activators of dibasic and neutral amino acid transport), member 2 | 6520 | ENSG00000168003 |
| 30 | Q7Z2K6 | NA | ERMP1 | endoplasmic reticulum metallopeptidase 1 | 79956 | ENSG00000099219 |
| 31 | P02795 | NA | MT2A | metallothionein 2A | 4502 | ENSG00000125148 |
| 32 | P49321 | NA | NASP | nuclear autoantigenic sperm protein (histone-binding) | 4678 | ENSG00000132780 |
| 33 | P00491 | NA | PNP | purine nucleoside phosphorylase | 4860 | ENSG00000198805 |
| 34 | P07951 | NA | TPM2 | tropomyosin 2 (beta) | 7169 | ENSG00000198467 |
| 35 | Q9NZM1 | NA | MYOF | myoferlin | 26509 | ENSG00000138119 |
| 36 | P67936 | NA | TPM4 | tropomyosin 4 | 7171 | ENSG00000167460 |
| 37 | P49327 | NA | FASN | fatty acid synthase | 2194 | ENSG00000169710 |
| 38 | P49368 | NA | CCT3 | chaperonin containing TCP1, subunit 3 (gamma) | 7203 | ENSG00000163468 |
| 39 | P56192 | NA | MARS | methionyl-tRNA synthetase | 4141 | ENSG00000166986 |
| 40 | Q9BQE3 | NA | TUBA1C | tubulin, alpha 1c | 84790 | ENSG00000167553 |
| 41 | P12268 | NA | IMPDH2 | IMP (inosine 5'-monophosphate) dehydrogenase 2 | 3615 | ENSG00000178035 |
| 42 | P62906 | NA | RPL10A | ribosomal protein L10a | 4736 | ENSG00000198755 |
| 43 | P49736 | NA | MCM2 | minichromosome maintenance complex component 2 | 4171 | ENSG00000073111 |
| 44 | O60610 | NA | DIAPH1 | diaphanous homolog 1 (Drosophila) | 1729 | ENSG00000131504 |
| 45 | P50990 | NA | CCT8 | chaperonin containing TCP1, subunit 8 (theta) | 10694 | ENSG00000156261 |
| 46 | Q06830 | NA | PRDX1 | peroxiredoxin 1 | 5052 | ENSG00000117450 |
| 47 | P17987 | NA | TCP1 | t-complex 1 | 6950 | ENSG00000120438 |
| 48 | P55209 | NA | NAP1L1 | nucleosome assembly protein 1-like 1 | 4673 | ENSG00000187109 |
| 49 | P34932 | NA | HSPA4 | heat shock 70kDa protein 4 | 3308 | ENSG00000170606 |
| 50 | P07108 | NA | DBI | diazepam binding inhibitor (GABA receptor modulator, acyl-CoA binding protein) | 1622 | ENSG00000155368 |
| 51 | P40227 | NA | CCT6A | chaperonin containing TCP1, subunit 6A (zeta 1) | 908 | ENSG00000146731 |
| 52 | P52292 | NA | KPNA2 | karyopherin alpha 2 (RAG cohort 1, importin alpha 1) | 3838 | ENSG00000182481 |
| 53 | Q15758 | NA | SLC1A5 | solute carrier family 1 (neutral amino acid transporter), member 5 | 6510 | ENSG00000105281 |
| 54 | Q8TEX9 | NA | IPO4 | importin 4 | 79711 | ENSG00000196497 |

  
  

| **Database:cellular component      &nbspName:organelle      &nbspID:GO:0043226** | | | | | | |
| --- | --- | --- | --- | --- | --- | --- |
| C=10536; O=54; E=39.16; R=1.38; rawP=2.45e-05; adjP=0.0001 | | | | | | |
| Index | UserID | Value | Gene Symbol | Gene Name | EntrezGene | Ensembl |
| 1 | Q92598 | NA | HSPH1 | heat shock 105kDa/110kDa protein 1 | 10808 | ENSG00000120694 |
| 2 | Q13885 | NA | TUBB2A | tubulin, beta 2A class IIa | 7280 | ENSG00000137267 |
| 3 | P68371 | NA | TUBB4B | tubulin, beta 4B class IVb | 10383 | ENSG00000188229 |
| 4 | P25685 | NA | DNAJB1 | DnaJ (Hsp40) homolog, subfamily B, member 1 | 3337 | ENSG00000132002 |
| 5 | P53701 | NA | HCCS | holocytochrome c synthase | 3052 | ENSG00000004961 |
| 6 | P68104 | NA | EEF1A1 | eukaryotic translation elongation factor 1 alpha 1 | 1915 | ENSG00000156508 |
| 7 | O75369 | NA | FLNB | filamin B, beta | 2317 | ENSG00000136068 |
| 8 | P02786 | NA | TFRC | transferrin receptor (p90, CD71) | 7037 | ENSG00000072274 |
| 9 | O00425 | NA | IGF2BP3 | insulin-like growth factor 2 mRNA binding protein 3 | 10643 | ENSG00000136231 |
| 10 | P50991 | NA | CCT4 | chaperonin containing TCP1, subunit 4 (delta) | 10575 | ENSG00000115484 |
| 11 | Q16555 | NA | DPYSL2 | dihydropyrimidinase-like 2 | 1808 | ENSG00000092964 |
| 12 | O95433 | NA | AHSA1 | AHA1, activator of heat shock 90kDa protein ATPase homolog 1 (yeast) | 10598 | ENSG00000100591 |
| 13 | P17174 | NA | GOT1 | glutamic-oxaloacetic transaminase 1, soluble (aspartate aminotransferase 1) | 2805 | ENSG00000120053 |
| 14 | P67809 | NA | YBX1 | Y box binding protein 1 | 4904 | ENSG00000065978 |
| 15 | P21980 | NA | TGM2 | transglutaminase 2 (C polypeptide, protein-glutamine-gamma-glutamyltransferase) | 7052 | ENSG00000198959 |
| 16 | P37802 | NA | TAGLN2 | transgelin 2 | 8407 | ENSG00000158710 |
| 17 | Q15185 | NA | PTGES3 | prostaglandin E synthase 3 (cytosolic) | 10728 | ENSG00000110958 |
| 18 | Q99832 | NA | CCT7 | chaperonin containing TCP1, subunit 7 (eta) | 10574 | ENSG00000135624 |
| 19 | Q6PIU2 | NA | NCEH1 | neutral cholesterol ester hydrolase 1 | 57552 | ENSG00000144959 |
| 20 | P18206 | NA | VCL | vinculin | 7414 | ENSG00000035403 |
| 21 | P06703 | NA | S100A6 | S100 calcium binding protein A6 | 6277 | ENSG00000197956 |
| 22 | Q9UK76 | NA | HN1 | hematological and neurological expressed 1 | 51155 | ENSG00000189159 |
| 23 | P11387 | NA | TOP1 | topoisomerase (DNA) I | 7150 | ENSG00000198900 |
| 24 | P11586 | NA | MTHFD1 | methylenetetrahydrofolate dehydrogenase (NADP+ dependent) 1, methenyltetrahydrofolate cyclohydrolase, formyltetrahydrofolate synthetase | 4522 | ENSG00000100714 |
| 25 | Q16881 | NA | TXNRD1 | thioredoxin reductase 1 | 7296 | ENSG00000198431 |
| 26 | P63241 | NA | EIF5A | eukaryotic translation initiation factor 5A | 1984 | ENSG00000132507 |
| 27 | P78371 | NA | CCT2 | chaperonin containing TCP1, subunit 2 (beta) | 10576 | ENSG00000166226 |
| 28 | Q9NR30 | NA | DDX21 | DEAD (Asp-Glu-Ala-Asp) box helicase 21 | 9188 | ENSG00000165732 |
| 29 | P08195 | NA | SLC3A2 | solute carrier family 3 (activators of dibasic and neutral amino acid transport), member 2 | 6520 | ENSG00000168003 |
| 30 | Q7Z2K6 | NA | ERMP1 | endoplasmic reticulum metallopeptidase 1 | 79956 | ENSG00000099219 |
| 31 | P02795 | NA | MT2A | metallothionein 2A | 4502 | ENSG00000125148 |
| 32 | P49321 | NA | NASP | nuclear autoantigenic sperm protein (histone-binding) | 4678 | ENSG00000132780 |
| 33 | P00491 | NA | PNP | purine nucleoside phosphorylase | 4860 | ENSG00000198805 |
| 34 | P07951 | NA | TPM2 | tropomyosin 2 (beta) | 7169 | ENSG00000198467 |
| 35 | Q9NZM1 | NA | MYOF | myoferlin | 26509 | ENSG00000138119 |
| 36 | P67936 | NA | TPM4 | tropomyosin 4 | 7171 | ENSG00000167460 |
| 37 | P49327 | NA | FASN | fatty acid synthase | 2194 | ENSG00000169710 |
| 38 | P49368 | NA | CCT3 | chaperonin containing TCP1, subunit 3 (gamma) | 7203 | ENSG00000163468 |
| 39 | P56192 | NA | MARS | methionyl-tRNA synthetase | 4141 | ENSG00000166986 |
| 40 | Q9BQE3 | NA | TUBA1C | tubulin, alpha 1c | 84790 | ENSG00000167553 |
| 41 | P12268 | NA | IMPDH2 | IMP (inosine 5'-monophosphate) dehydrogenase 2 | 3615 | ENSG00000178035 |
| 42 | P62906 | NA | RPL10A | ribosomal protein L10a | 4736 | ENSG00000198755 |
| 43 | P49736 | NA | MCM2 | minichromosome maintenance complex component 2 | 4171 | ENSG00000073111 |
| 44 | O60610 | NA | DIAPH1 | diaphanous homolog 1 (Drosophila) | 1729 | ENSG00000131504 |
| 45 | P50990 | NA | CCT8 | chaperonin containing TCP1, subunit 8 (theta) | 10694 | ENSG00000156261 |
| 46 | Q06830 | NA | PRDX1 | peroxiredoxin 1 | 5052 | ENSG00000117450 |
| 47 | P17987 | NA | TCP1 | t-complex 1 | 6950 | ENSG00000120438 |
| 48 | P55209 | NA | NAP1L1 | nucleosome assembly protein 1-like 1 | 4673 | ENSG00000187109 |
| 49 | P34932 | NA | HSPA4 | heat shock 70kDa protein 4 | 3308 | ENSG00000170606 |
| 50 | P07108 | NA | DBI | diazepam binding inhibitor (GABA receptor modulator, acyl-CoA binding protein) | 1622 | ENSG00000155368 |
| 51 | P40227 | NA | CCT6A | chaperonin containing TCP1, subunit 6A (zeta 1) | 908 | ENSG00000146731 |
| 52 | P52292 | NA | KPNA2 | karyopherin alpha 2 (RAG cohort 1, importin alpha 1) | 3838 | ENSG00000182481 |
| 53 | Q15758 | NA | SLC1A5 | solute carrier family 1 (neutral amino acid transporter), member 5 | 6510 | ENSG00000105281 |
| 54 | Q8TEX9 | NA | IPO4 | importin 4 | 79711 | ENSG00000196497 |

  
  

| **Database:cellular component      &nbspName:microtubule cytoskeleton      &nbspID:GO:0015630** | | | | | | |
| --- | --- | --- | --- | --- | --- | --- |
| C=859; O=12; E=3.19; R=3.76; rawP=6.42e-05; adjP=0.0003 | | | | | | |
| Index | UserID | Value | Gene Symbol | Gene Name | EntrezGene | Ensembl |
| 1 | P78371 | NA | CCT2 | chaperonin containing TCP1, subunit 2 (beta) | 10576 | ENSG00000166226 |
| 2 | Q92598 | NA | HSPH1 | heat shock 105kDa/110kDa protein 1 | 10808 | ENSG00000120694 |
| 3 | P17987 | NA | TCP1 | t-complex 1 | 6950 | ENSG00000120438 |
| 4 | Q13885 | NA | TUBB2A | tubulin, beta 2A class IIa | 7280 | ENSG00000137267 |
| 5 | P68371 | NA | TUBB4B | tubulin, beta 4B class IVb | 10383 | ENSG00000188229 |
| 6 | P40227 | NA | CCT6A | chaperonin containing TCP1, subunit 6A (zeta 1) | 908 | ENSG00000146731 |
| 7 | P49368 | NA | CCT3 | chaperonin containing TCP1, subunit 3 (gamma) | 7203 | ENSG00000163468 |
| 8 | Q9BQE3 | NA | TUBA1C | tubulin, alpha 1c | 84790 | ENSG00000167553 |
| 9 | Q99832 | NA | CCT7 | chaperonin containing TCP1, subunit 7 (eta) | 10574 | ENSG00000135624 |
| 10 | O60610 | NA | DIAPH1 | diaphanous homolog 1 (Drosophila) | 1729 | ENSG00000131504 |
| 11 | P50990 | NA | CCT8 | chaperonin containing TCP1, subunit 8 (theta) | 10694 | ENSG00000156261 |
| 12 | P50991 | NA | CCT4 | chaperonin containing TCP1, subunit 4 (delta) | 10575 | ENSG00000115484 |

  
  

| **Database:cellular component      &nbspName:cell      &nbspID:GO:0005623** | | | | | | |
| --- | --- | --- | --- | --- | --- | --- |
| C=14406; O=62; E=53.54; R=1.16; rawP=0.0001; adjP=0.0005 | | | | | | |
| Index | UserID | Value | Gene Symbol | Gene Name | EntrezGene | Ensembl |
| 1 | Q92598 | NA | HSPH1 | heat shock 105kDa/110kDa protein 1 | 10808 | ENSG00000120694 |
| 2 | Q13885 | NA | TUBB2A | tubulin, beta 2A class IIa | 7280 | ENSG00000137267 |
| 3 | P68371 | NA | TUBB4B | tubulin, beta 4B class IVb | 10383 | ENSG00000188229 |
| 4 | P25685 | NA | DNAJB1 | DnaJ (Hsp40) homolog, subfamily B, member 1 | 3337 | ENSG00000132002 |
| 5 | P53701 | NA | HCCS | holocytochrome c synthase | 3052 | ENSG00000004961 |
| 6 | P68104 | NA | EEF1A1 | eukaryotic translation elongation factor 1 alpha 1 | 1915 | ENSG00000156508 |
| 7 | O75369 | NA | FLNB | filamin B, beta | 2317 | ENSG00000136068 |
| 8 | P02786 | NA | TFRC | transferrin receptor (p90, CD71) | 7037 | ENSG00000072274 |
| 9 | O00425 | NA | IGF2BP3 | insulin-like growth factor 2 mRNA binding protein 3 | 10643 | ENSG00000136231 |
| 10 | P50991 | NA | CCT4 | chaperonin containing TCP1, subunit 4 (delta) | 10575 | ENSG00000115484 |
| 11 | P22102 | NA | GART | phosphoribosylglycinamide formyltransferase, phosphoribosylglycinamide synthetase, phosphoribosylaminoimidazole synthetase | 2618 | ENSG00000159131 |
| 12 | P13639 | NA | EEF2 | eukaryotic translation elongation factor 2 | 1938 | ENSG00000167658 |
| 13 | Q16555 | NA | DPYSL2 | dihydropyrimidinase-like 2 | 1808 | ENSG00000092964 |
| 14 | O95433 | NA | AHSA1 | AHA1, activator of heat shock 90kDa protein ATPase homolog 1 (yeast) | 10598 | ENSG00000100591 |
| 15 | P41567 | NA | EIF1 | eukaryotic translation initiation factor 1 | 10209 | ENSG00000173812 |
| 16 | P17174 | NA | GOT1 | glutamic-oxaloacetic transaminase 1, soluble (aspartate aminotransferase 1) | 2805 | ENSG00000120053 |
| 17 | P67809 | NA | YBX1 | Y box binding protein 1 | 4904 | ENSG00000065978 |
| 18 | P21980 | NA | TGM2 | transglutaminase 2 (C polypeptide, protein-glutamine-gamma-glutamyltransferase) | 7052 | ENSG00000198959 |
| 19 | P37802 | NA | TAGLN2 | transgelin 2 | 8407 | ENSG00000158710 |
| 20 | Q99832 | NA | CCT7 | chaperonin containing TCP1, subunit 7 (eta) | 10574 | ENSG00000135624 |
| 21 | Q15185 | NA | PTGES3 | prostaglandin E synthase 3 (cytosolic) | 10728 | ENSG00000110958 |
| 22 | Q6PIU2 | NA | NCEH1 | neutral cholesterol ester hydrolase 1 | 57552 | ENSG00000144959 |
| 23 | P18206 | NA | VCL | vinculin | 7414 | ENSG00000035403 |
| 24 | P06703 | NA | S100A6 | S100 calcium binding protein A6 | 6277 | ENSG00000197956 |
| 25 | Q9UK76 | NA | HN1 | hematological and neurological expressed 1 | 51155 | ENSG00000189159 |
| 26 | P11387 | NA | TOP1 | topoisomerase (DNA) I | 7150 | ENSG00000198900 |
| 27 | P11586 | NA | MTHFD1 | methylenetetrahydrofolate dehydrogenase (NADP+ dependent) 1, methenyltetrahydrofolate cyclohydrolase, formyltetrahydrofolate synthetase | 4522 | ENSG00000100714 |
| 28 | Q16881 | NA | TXNRD1 | thioredoxin reductase 1 | 7296 | ENSG00000198431 |
| 29 | P63241 | NA | EIF5A | eukaryotic translation initiation factor 5A | 1984 | ENSG00000132507 |
| 30 | P78371 | NA | CCT2 | chaperonin containing TCP1, subunit 2 (beta) | 10576 | ENSG00000166226 |
| 31 | Q9NR30 | NA | DDX21 | DEAD (Asp-Glu-Ala-Asp) box helicase 21 | 9188 | ENSG00000165732 |
| 32 | P02795 | NA | MT2A | metallothionein 2A | 4502 | ENSG00000125148 |
| 33 | Q7Z2K6 | NA | ERMP1 | endoplasmic reticulum metallopeptidase 1 | 79956 | ENSG00000099219 |
| 34 | P08195 | NA | SLC3A2 | solute carrier family 3 (activators of dibasic and neutral amino acid transport), member 2 | 6520 | ENSG00000168003 |
| 35 | P49321 | NA | NASP | nuclear autoantigenic sperm protein (histone-binding) | 4678 | ENSG00000132780 |
| 36 | P00491 | NA | PNP | purine nucleoside phosphorylase | 4860 | ENSG00000198805 |
| 37 | Q9NZM1 | NA | MYOF | myoferlin | 26509 | ENSG00000138119 |
| 38 | P07951 | NA | TPM2 | tropomyosin 2 (beta) | 7169 | ENSG00000198467 |
| 39 | P12081 | NA | HARS | histidyl-tRNA synthetase | 3035 | ENSG00000170445 |
| 40 | P67936 | NA | TPM4 | tropomyosin 4 | 7171 | ENSG00000167460 |
| 41 | P49327 | NA | FASN | fatty acid synthase | 2194 | ENSG00000169710 |
| 42 | P49368 | NA | CCT3 | chaperonin containing TCP1, subunit 3 (gamma) | 7203 | ENSG00000163468 |
| 43 | P56192 | NA | MARS | methionyl-tRNA synthetase | 4141 | ENSG00000166986 |
| 44 | Q9BQE3 | NA | TUBA1C | tubulin, alpha 1c | 84790 | ENSG00000167553 |
| 45 | P12268 | NA | IMPDH2 | IMP (inosine 5'-monophosphate) dehydrogenase 2 | 3615 | ENSG00000178035 |
| 46 | P62906 | NA | RPL10A | ribosomal protein L10a | 4736 | ENSG00000198755 |
| 47 | P49736 | NA | MCM2 | minichromosome maintenance complex component 2 | 4171 | ENSG00000073111 |
| 48 | Q15056 | NA | EIF4H | eukaryotic translation initiation factor 4H | 7458 | ENSG00000106682 |
| 49 | O60610 | NA | DIAPH1 | diaphanous homolog 1 (Drosophila) | 1729 | ENSG00000131504 |
| 50 | P50990 | NA | CCT8 | chaperonin containing TCP1, subunit 8 (theta) | 10694 | ENSG00000156261 |
| 51 | Q06830 | NA | PRDX1 | peroxiredoxin 1 | 5052 | ENSG00000117450 |
| 52 | P07814 | NA | EPRS | glutamyl-prolyl-tRNA synthetase | 2058 | ENSG00000136628 |
| 53 | P17987 | NA | TCP1 | t-complex 1 | 6950 | ENSG00000120438 |
| 54 | P55209 | NA | NAP1L1 | nucleosome assembly protein 1-like 1 | 4673 | ENSG00000187109 |
| 55 | P34932 | NA | HSPA4 | heat shock 70kDa protein 4 | 3308 | ENSG00000170606 |
| 56 | P41091 | NA | EIF2S3 | eukaryotic translation initiation factor 2, subunit 3 gamma, 52kDa | 1968 | ENSG00000130741 |
| 57 | P07108 | NA | DBI | diazepam binding inhibitor (GABA receptor modulator, acyl-CoA binding protein) | 1622 | ENSG00000155368 |
| 58 | P40227 | NA | CCT6A | chaperonin containing TCP1, subunit 6A (zeta 1) | 908 | ENSG00000146731 |
| 59 | P52292 | NA | KPNA2 | karyopherin alpha 2 (RAG cohort 1, importin alpha 1) | 3838 | ENSG00000182481 |
| 60 | Q15758 | NA | SLC1A5 | solute carrier family 1 (neutral amino acid transporter), member 5 | 6510 | ENSG00000105281 |
| 61 | P00352 | NA | ALDH1A1 | aldehyde dehydrogenase 1 family, member A1 | 216 | ENSG00000165092 |
| 62 | Q8TEX9 | NA | IPO4 | importin 4 | 79711 | ENSG00000196497 |

  
  

| **Database:cellular component      &nbspName:cell part      &nbspID:GO:0044464** | | | | | | |
| --- | --- | --- | --- | --- | --- | --- |
| C=14405; O=62; E=53.54; R=1.16; rawP=0.0001; adjP=0.0005 | | | | | | |
| Index | UserID | Value | Gene Symbol | Gene Name | EntrezGene | Ensembl |
| 1 | Q92598 | NA | HSPH1 | heat shock 105kDa/110kDa protein 1 | 10808 | ENSG00000120694 |
| 2 | Q13885 | NA | TUBB2A | tubulin, beta 2A class IIa | 7280 | ENSG00000137267 |
| 3 | P68371 | NA | TUBB4B | tubulin, beta 4B class IVb | 10383 | ENSG00000188229 |
| 4 | P25685 | NA | DNAJB1 | DnaJ (Hsp40) homolog, subfamily B, member 1 | 3337 | ENSG00000132002 |
| 5 | P53701 | NA | HCCS | holocytochrome c synthase | 3052 | ENSG00000004961 |
| 6 | P68104 | NA | EEF1A1 | eukaryotic translation elongation factor 1 alpha 1 | 1915 | ENSG00000156508 |
| 7 | O75369 | NA | FLNB | filamin B, beta | 2317 | ENSG00000136068 |
| 8 | P02786 | NA | TFRC | transferrin receptor (p90, CD71) | 7037 | ENSG00000072274 |
| 9 | O00425 | NA | IGF2BP3 | insulin-like growth factor 2 mRNA binding protein 3 | 10643 | ENSG00000136231 |
| 10 | P50991 | NA | CCT4 | chaperonin containing TCP1, subunit 4 (delta) | 10575 | ENSG00000115484 |
| 11 | P22102 | NA | GART | phosphoribosylglycinamide formyltransferase, phosphoribosylglycinamide synthetase, phosphoribosylaminoimidazole synthetase | 2618 | ENSG00000159131 |
| 12 | P13639 | NA | EEF2 | eukaryotic translation elongation factor 2 | 1938 | ENSG00000167658 |
| 13 | Q16555 | NA | DPYSL2 | dihydropyrimidinase-like 2 | 1808 | ENSG00000092964 |
| 14 | O95433 | NA | AHSA1 | AHA1, activator of heat shock 90kDa protein ATPase homolog 1 (yeast) | 10598 | ENSG00000100591 |
| 15 | P41567 | NA | EIF1 | eukaryotic translation initiation factor 1 | 10209 | ENSG00000173812 |
| 16 | P17174 | NA | GOT1 | glutamic-oxaloacetic transaminase 1, soluble (aspartate aminotransferase 1) | 2805 | ENSG00000120053 |
| 17 | P67809 | NA | YBX1 | Y box binding protein 1 | 4904 | ENSG00000065978 |
| 18 | P21980 | NA | TGM2 | transglutaminase 2 (C polypeptide, protein-glutamine-gamma-glutamyltransferase) | 7052 | ENSG00000198959 |
| 19 | P37802 | NA | TAGLN2 | transgelin 2 | 8407 | ENSG00000158710 |
| 20 | Q99832 | NA | CCT7 | chaperonin containing TCP1, subunit 7 (eta) | 10574 | ENSG00000135624 |
| 21 | Q15185 | NA | PTGES3 | prostaglandin E synthase 3 (cytosolic) | 10728 | ENSG00000110958 |
| 22 | Q6PIU2 | NA | NCEH1 | neutral cholesterol ester hydrolase 1 | 57552 | ENSG00000144959 |
| 23 | P18206 | NA | VCL | vinculin | 7414 | ENSG00000035403 |
| 24 | P06703 | NA | S100A6 | S100 calcium binding protein A6 | 6277 | ENSG00000197956 |
| 25 | Q9UK76 | NA | HN1 | hematological and neurological expressed 1 | 51155 | ENSG00000189159 |
| 26 | P11387 | NA | TOP1 | topoisomerase (DNA) I | 7150 | ENSG00000198900 |
| 27 | P11586 | NA | MTHFD1 | methylenetetrahydrofolate dehydrogenase (NADP+ dependent) 1, methenyltetrahydrofolate cyclohydrolase, formyltetrahydrofolate synthetase | 4522 | ENSG00000100714 |
| 28 | Q16881 | NA | TXNRD1 | thioredoxin reductase 1 | 7296 | ENSG00000198431 |
| 29 | P63241 | NA | EIF5A | eukaryotic translation initiation factor 5A | 1984 | ENSG00000132507 |
| 30 | P78371 | NA | CCT2 | chaperonin containing TCP1, subunit 2 (beta) | 10576 | ENSG00000166226 |
| 31 | Q9NR30 | NA | DDX21 | DEAD (Asp-Glu-Ala-Asp) box helicase 21 | 9188 | ENSG00000165732 |
| 32 | P02795 | NA | MT2A | metallothionein 2A | 4502 | ENSG00000125148 |
| 33 | Q7Z2K6 | NA | ERMP1 | endoplasmic reticulum metallopeptidase 1 | 79956 | ENSG00000099219 |
| 34 | P08195 | NA | SLC3A2 | solute carrier family 3 (activators of dibasic and neutral amino acid transport), member 2 | 6520 | ENSG00000168003 |
| 35 | P49321 | NA | NASP | nuclear autoantigenic sperm protein (histone-binding) | 4678 | ENSG00000132780 |
| 36 | P00491 | NA | PNP | purine nucleoside phosphorylase | 4860 | ENSG00000198805 |
| 37 | Q9NZM1 | NA | MYOF | myoferlin | 26509 | ENSG00000138119 |
| 38 | P07951 | NA | TPM2 | tropomyosin 2 (beta) | 7169 | ENSG00000198467 |
| 39 | P12081 | NA | HARS | histidyl-tRNA synthetase | 3035 | ENSG00000170445 |
| 40 | P67936 | NA | TPM4 | tropomyosin 4 | 7171 | ENSG00000167460 |
| 41 | P49327 | NA | FASN | fatty acid synthase | 2194 | ENSG00000169710 |
| 42 | P49368 | NA | CCT3 | chaperonin containing TCP1, subunit 3 (gamma) | 7203 | ENSG00000163468 |
| 43 | P56192 | NA | MARS | methionyl-tRNA synthetase | 4141 | ENSG00000166986 |
| 44 | Q9BQE3 | NA | TUBA1C | tubulin, alpha 1c | 84790 | ENSG00000167553 |
| 45 | P12268 | NA | IMPDH2 | IMP (inosine 5'-monophosphate) dehydrogenase 2 | 3615 | ENSG00000178035 |
| 46 | P62906 | NA | RPL10A | ribosomal protein L10a | 4736 | ENSG00000198755 |
| 47 | P49736 | NA | MCM2 | minichromosome maintenance complex component 2 | 4171 | ENSG00000073111 |
| 48 | Q15056 | NA | EIF4H | eukaryotic translation initiation factor 4H | 7458 | ENSG00000106682 |
| 49 | O60610 | NA | DIAPH1 | diaphanous homolog 1 (Drosophila) | 1729 | ENSG00000131504 |
| 50 | P50990 | NA | CCT8 | chaperonin containing TCP1, subunit 8 (theta) | 10694 | ENSG00000156261 |
| 51 | Q06830 | NA | PRDX1 | peroxiredoxin 1 | 5052 | ENSG00000117450 |
| 52 | P07814 | NA | EPRS | glutamyl-prolyl-tRNA synthetase | 2058 | ENSG00000136628 |
| 53 | P17987 | NA | TCP1 | t-complex 1 | 6950 | ENSG00000120438 |
| 54 | P55209 | NA | NAP1L1 | nucleosome assembly protein 1-like 1 | 4673 | ENSG00000187109 |
| 55 | P34932 | NA | HSPA4 | heat shock 70kDa protein 4 | 3308 | ENSG00000170606 |
| 56 | P41091 | NA | EIF2S3 | eukaryotic translation initiation factor 2, subunit 3 gamma, 52kDa | 1968 | ENSG00000130741 |
| 57 | P07108 | NA | DBI | diazepam binding inhibitor (GABA receptor modulator, acyl-CoA binding protein) | 1622 | ENSG00000155368 |
| 58 | P40227 | NA | CCT6A | chaperonin containing TCP1, subunit 6A (zeta 1) | 908 | ENSG00000146731 |
| 59 | P52292 | NA | KPNA2 | karyopherin alpha 2 (RAG cohort 1, importin alpha 1) | 3838 | ENSG00000182481 |
| 60 | Q15758 | NA | SLC1A5 | solute carrier family 1 (neutral amino acid transporter), member 5 | 6510 | ENSG00000105281 |
| 61 | P00352 | NA | ALDH1A1 | aldehyde dehydrogenase 1 family, member A1 | 216 | ENSG00000165092 |
| 62 | Q8TEX9 | NA | IPO4 | importin 4 | 79711 | ENSG00000196497 |

  
  

| **Database:cellular component      &nbspName:cytoskeleton      &nbspID:GO:0005856** | | | | | | |
| --- | --- | --- | --- | --- | --- | --- |
| C=1783; O=17; E=6.63; R=2.57; rawP=0.0002; adjP=0.0009 | | | | | | |
| Index | UserID | Value | Gene Symbol | Gene Name | EntrezGene | Ensembl |
| 1 | P78371 | NA | CCT2 | chaperonin containing TCP1, subunit 2 (beta) | 10576 | ENSG00000166226 |
| 2 | Q92598 | NA | HSPH1 | heat shock 105kDa/110kDa protein 1 | 10808 | ENSG00000120694 |
| 3 | Q13885 | NA | TUBB2A | tubulin, beta 2A class IIa | 7280 | ENSG00000137267 |
| 4 | P00491 | NA | PNP | purine nucleoside phosphorylase | 4860 | ENSG00000198805 |
| 5 | P07951 | NA | TPM2 | tropomyosin 2 (beta) | 7169 | ENSG00000198467 |
| 6 | P68371 | NA | TUBB4B | tubulin, beta 4B class IVb | 10383 | ENSG00000188229 |
| 7 | P67936 | NA | TPM4 | tropomyosin 4 | 7171 | ENSG00000167460 |
| 8 | P49368 | NA | CCT3 | chaperonin containing TCP1, subunit 3 (gamma) | 7203 | ENSG00000163468 |
| 9 | Q9BQE3 | NA | TUBA1C | tubulin, alpha 1c | 84790 | ENSG00000167553 |
| 10 | O60610 | NA | DIAPH1 | diaphanous homolog 1 (Drosophila) | 1729 | ENSG00000131504 |
| 11 | P50990 | NA | CCT8 | chaperonin containing TCP1, subunit 8 (theta) | 10694 | ENSG00000156261 |
| 12 | O75369 | NA | FLNB | filamin B, beta | 2317 | ENSG00000136068 |
| 13 | P50991 | NA | CCT4 | chaperonin containing TCP1, subunit 4 (delta) | 10575 | ENSG00000115484 |
| 14 | P17987 | NA | TCP1 | t-complex 1 | 6950 | ENSG00000120438 |
| 15 | P40227 | NA | CCT6A | chaperonin containing TCP1, subunit 6A (zeta 1) | 908 | ENSG00000146731 |
| 16 | Q99832 | NA | CCT7 | chaperonin containing TCP1, subunit 7 (eta) | 10574 | ENSG00000135624 |
| 17 | P18206 | NA | VCL | vinculin | 7414 | ENSG00000035403 |

  
  

| **Database:cellular component      &nbspName:non-membrane-bounded organelle      &nbspID:GO:0043228** | | | | | | |
| --- | --- | --- | --- | --- | --- | --- |
| C=3778; O=26; E=14.04; R=1.85; rawP=0.0005; adjP=0.0020 | | | | | | |
| Index | UserID | Value | Gene Symbol | Gene Name | EntrezGene | Ensembl |
| 1 | Q16881 | NA | TXNRD1 | thioredoxin reductase 1 | 7296 | ENSG00000198431 |
| 2 | Q9NR30 | NA | DDX21 | DEAD (Asp-Glu-Ala-Asp) box helicase 21 | 9188 | ENSG00000165732 |
| 3 | P78371 | NA | CCT2 | chaperonin containing TCP1, subunit 2 (beta) | 10576 | ENSG00000166226 |
| 4 | Q92598 | NA | HSPH1 | heat shock 105kDa/110kDa protein 1 | 10808 | ENSG00000120694 |
| 5 | Q13885 | NA | TUBB2A | tubulin, beta 2A class IIa | 7280 | ENSG00000137267 |
| 6 | P00491 | NA | PNP | purine nucleoside phosphorylase | 4860 | ENSG00000198805 |
| 7 | P07951 | NA | TPM2 | tropomyosin 2 (beta) | 7169 | ENSG00000198467 |
| 8 | P68371 | NA | TUBB4B | tubulin, beta 4B class IVb | 10383 | ENSG00000188229 |
| 9 | P67936 | NA | TPM4 | tropomyosin 4 | 7171 | ENSG00000167460 |
| 10 | P49368 | NA | CCT3 | chaperonin containing TCP1, subunit 3 (gamma) | 7203 | ENSG00000163468 |
| 11 | Q9BQE3 | NA | TUBA1C | tubulin, alpha 1c | 84790 | ENSG00000167553 |
| 12 | P25685 | NA | DNAJB1 | DnaJ (Hsp40) homolog, subfamily B, member 1 | 3337 | ENSG00000132002 |
| 13 | P62906 | NA | RPL10A | ribosomal protein L10a | 4736 | ENSG00000198755 |
| 14 | P49736 | NA | MCM2 | minichromosome maintenance complex component 2 | 4171 | ENSG00000073111 |
| 15 | O60610 | NA | DIAPH1 | diaphanous homolog 1 (Drosophila) | 1729 | ENSG00000131504 |
| 16 | P50990 | NA | CCT8 | chaperonin containing TCP1, subunit 8 (theta) | 10694 | ENSG00000156261 |
| 17 | O75369 | NA | FLNB | filamin B, beta | 2317 | ENSG00000136068 |
| 18 | P50991 | NA | CCT4 | chaperonin containing TCP1, subunit 4 (delta) | 10575 | ENSG00000115484 |
| 19 | P17987 | NA | TCP1 | t-complex 1 | 6950 | ENSG00000120438 |
| 20 | P34932 | NA | HSPA4 | heat shock 70kDa protein 4 | 3308 | ENSG00000170606 |
| 21 | P40227 | NA | CCT6A | chaperonin containing TCP1, subunit 6A (zeta 1) | 908 | ENSG00000146731 |
| 22 | P67809 | NA | YBX1 | Y box binding protein 1 | 4904 | ENSG00000065978 |
| 23 | Q15185 | NA | PTGES3 | prostaglandin E synthase 3 (cytosolic) | 10728 | ENSG00000110958 |
| 24 | Q99832 | NA | CCT7 | chaperonin containing TCP1, subunit 7 (eta) | 10574 | ENSG00000135624 |
| 25 | P18206 | NA | VCL | vinculin | 7414 | ENSG00000035403 |
| 26 | P11387 | NA | TOP1 | topoisomerase (DNA) I | 7150 | ENSG00000198900 |

  
  

| **Database:cellular component      &nbspName:intracellular non-membrane-bounded organelle      &nbspID:GO:0043232** | | | | | | |
| --- | --- | --- | --- | --- | --- | --- |
| C=3778; O=26; E=14.04; R=1.85; rawP=0.0005; adjP=0.0020 | | | | | | |
| Index | UserID | Value | Gene Symbol | Gene Name | EntrezGene | Ensembl |
| 1 | Q16881 | NA | TXNRD1 | thioredoxin reductase 1 | 7296 | ENSG00000198431 |
| 2 | Q9NR30 | NA | DDX21 | DEAD (Asp-Glu-Ala-Asp) box helicase 21 | 9188 | ENSG00000165732 |
| 3 | P78371 | NA | CCT2 | chaperonin containing TCP1, subunit 2 (beta) | 10576 | ENSG00000166226 |
| 4 | Q92598 | NA | HSPH1 | heat shock 105kDa/110kDa protein 1 | 10808 | ENSG00000120694 |
| 5 | Q13885 | NA | TUBB2A | tubulin, beta 2A class IIa | 7280 | ENSG00000137267 |
| 6 | P00491 | NA | PNP | purine nucleoside phosphorylase | 4860 | ENSG00000198805 |
| 7 | P07951 | NA | TPM2 | tropomyosin 2 (beta) | 7169 | ENSG00000198467 |
| 8 | P68371 | NA | TUBB4B | tubulin, beta 4B class IVb | 10383 | ENSG00000188229 |
| 9 | P67936 | NA | TPM4 | tropomyosin 4 | 7171 | ENSG00000167460 |
| 10 | P49368 | NA | CCT3 | chaperonin containing TCP1, subunit 3 (gamma) | 7203 | ENSG00000163468 |
| 11 | Q9BQE3 | NA | TUBA1C | tubulin, alpha 1c | 84790 | ENSG00000167553 |
| 12 | P25685 | NA | DNAJB1 | DnaJ (Hsp40) homolog, subfamily B, member 1 | 3337 | ENSG00000132002 |
| 13 | P62906 | NA | RPL10A | ribosomal protein L10a | 4736 | ENSG00000198755 |
| 14 | P49736 | NA | MCM2 | minichromosome maintenance complex component 2 | 4171 | ENSG00000073111 |
| 15 | O60610 | NA | DIAPH1 | diaphanous homolog 1 (Drosophila) | 1729 | ENSG00000131504 |
| 16 | P50990 | NA | CCT8 | chaperonin containing TCP1, subunit 8 (theta) | 10694 | ENSG00000156261 |
| 17 | O75369 | NA | FLNB | filamin B, beta | 2317 | ENSG00000136068 |
| 18 | P50991 | NA | CCT4 | chaperonin containing TCP1, subunit 4 (delta) | 10575 | ENSG00000115484 |
| 19 | P17987 | NA | TCP1 | t-complex 1 | 6950 | ENSG00000120438 |
| 20 | P34932 | NA | HSPA4 | heat shock 70kDa protein 4 | 3308 | ENSG00000170606 |
| 21 | P40227 | NA | CCT6A | chaperonin containing TCP1, subunit 6A (zeta 1) | 908 | ENSG00000146731 |
| 22 | P67809 | NA | YBX1 | Y box binding protein 1 | 4904 | ENSG00000065978 |
| 23 | Q15185 | NA | PTGES3 | prostaglandin E synthase 3 (cytosolic) | 10728 | ENSG00000110958 |
| 24 | Q99832 | NA | CCT7 | chaperonin containing TCP1, subunit 7 (eta) | 10574 | ENSG00000135624 |
| 25 | P18206 | NA | VCL | vinculin | 7414 | ENSG00000035403 |
| 26 | P11387 | NA | TOP1 | topoisomerase (DNA) I | 7150 | ENSG00000198900 |

  
  

| **Database:cellular component      &nbspName:stress fiber      &nbspID:GO:0001725** | | | | | | |
| --- | --- | --- | --- | --- | --- | --- |
| C=48; O=3; E=0.18; R=16.82; rawP=0.0008; adjP=0.0030 | | | | | | |
| Index | UserID | Value | Gene Symbol | Gene Name | EntrezGene | Ensembl |
| 1 | P18206 | NA | VCL | vinculin | 7414 | ENSG00000035403 |
| 2 | O75369 | NA | FLNB | filamin B, beta | 2317 | ENSG00000136068 |
| 3 | P67936 | NA | TPM4 | tropomyosin 4 | 7171 | ENSG00000167460 |

  
  

| **Database:cellular component      &nbspName:actin filament bundle      &nbspID:GO:0032432** | | | | | | |
| --- | --- | --- | --- | --- | --- | --- |
| C=51; O=3; E=0.19; R=15.83; rawP=0.0009; adjP=0.0033 | | | | | | |
| Index | UserID | Value | Gene Symbol | Gene Name | EntrezGene | Ensembl |
| 1 | P18206 | NA | VCL | vinculin | 7414 | ENSG00000035403 |
| 2 | O75369 | NA | FLNB | filamin B, beta | 2317 | ENSG00000136068 |
| 3 | P67936 | NA | TPM4 | tropomyosin 4 | 7171 | ENSG00000167460 |

  
  

| **Database:cellular component      &nbspName:actomyosin      &nbspID:GO:0042641** | | | | | | |
| --- | --- | --- | --- | --- | --- | --- |
| C=57; O=3; E=0.21; R=14.16; rawP=0.0012; adjP=0.0041 | | | | | | |
| Index | UserID | Value | Gene Symbol | Gene Name | EntrezGene | Ensembl |
| 1 | P18206 | NA | VCL | vinculin | 7414 | ENSG00000035403 |
| 2 | O75369 | NA | FLNB | filamin B, beta | 2317 | ENSG00000136068 |
| 3 | P67936 | NA | TPM4 | tropomyosin 4 | 7171 | ENSG00000167460 |

  
  

| **Database:cellular component      &nbspName:macromolecular complex      &nbspID:GO:0032991** | | | | | | |
| --- | --- | --- | --- | --- | --- | --- |
| C=3836; O=25; E=14.26; R=1.75; rawP=0.0017; adjP=0.0056 | | | | | | |
| Index | UserID | Value | Gene Symbol | Gene Name | EntrezGene | Ensembl |
| 1 | P63241 | NA | EIF5A | eukaryotic translation initiation factor 5A | 1984 | ENSG00000132507 |
| 2 | P78371 | NA | CCT2 | chaperonin containing TCP1, subunit 2 (beta) | 10576 | ENSG00000166226 |
| 3 | Q92598 | NA | HSPH1 | heat shock 105kDa/110kDa protein 1 | 10808 | ENSG00000120694 |
| 4 | Q13885 | NA | TUBB2A | tubulin, beta 2A class IIa | 7280 | ENSG00000137267 |
| 5 | P68371 | NA | TUBB4B | tubulin, beta 4B class IVb | 10383 | ENSG00000188229 |
| 6 | P67936 | NA | TPM4 | tropomyosin 4 | 7171 | ENSG00000167460 |
| 7 | P49368 | NA | CCT3 | chaperonin containing TCP1, subunit 3 (gamma) | 7203 | ENSG00000163468 |
| 8 | Q9BQE3 | NA | TUBA1C | tubulin, alpha 1c | 84790 | ENSG00000167553 |
| 9 | P62906 | NA | RPL10A | ribosomal protein L10a | 4736 | ENSG00000198755 |
| 10 | P49736 | NA | MCM2 | minichromosome maintenance complex component 2 | 4171 | ENSG00000073111 |
| 11 | Q15056 | NA | EIF4H | eukaryotic translation initiation factor 4H | 7458 | ENSG00000106682 |
| 12 | P50990 | NA | CCT8 | chaperonin containing TCP1, subunit 8 (theta) | 10694 | ENSG00000156261 |
| 13 | P68104 | NA | EEF1A1 | eukaryotic translation elongation factor 1 alpha 1 | 1915 | ENSG00000156508 |
| 14 | P50991 | NA | CCT4 | chaperonin containing TCP1, subunit 4 (delta) | 10575 | ENSG00000115484 |
| 15 | P17987 | NA | TCP1 | t-complex 1 | 6950 | ENSG00000120438 |
| 16 | P55209 | NA | NAP1L1 | nucleosome assembly protein 1-like 1 | 4673 | ENSG00000187109 |
| 17 | P13639 | NA | EEF2 | eukaryotic translation elongation factor 2 | 1938 | ENSG00000167658 |
| 18 | Q16555 | NA | DPYSL2 | dihydropyrimidinase-like 2 | 1808 | ENSG00000092964 |
| 19 | P40227 | NA | CCT6A | chaperonin containing TCP1, subunit 6A (zeta 1) | 908 | ENSG00000146731 |
| 20 | P67809 | NA | YBX1 | Y box binding protein 1 | 4904 | ENSG00000065978 |
| 21 | Q15185 | NA | PTGES3 | prostaglandin E synthase 3 (cytosolic) | 10728 | ENSG00000110958 |
| 22 | Q99832 | NA | CCT7 | chaperonin containing TCP1, subunit 7 (eta) | 10574 | ENSG00000135624 |
| 23 | P18206 | NA | VCL | vinculin | 7414 | ENSG00000035403 |
| 24 | Q8TEX9 | NA | IPO4 | importin 4 | 79711 | ENSG00000196497 |
| 25 | P11387 | NA | TOP1 | topoisomerase (DNA) I | 7150 | ENSG00000198900 |

  
  

| **Database:cellular component      &nbspName:contractile fiber part      &nbspID:GO:0044449** | | | | | | |
| --- | --- | --- | --- | --- | --- | --- |
| C=145; O=4; E=0.54; R=7.42; rawP=0.0021; adjP=0.0066 | | | | | | |
| Index | UserID | Value | Gene Symbol | Gene Name | EntrezGene | Ensembl |
| 1 | P07951 | NA | TPM2 | tropomyosin 2 (beta) | 7169 | ENSG00000198467 |
| 2 | P18206 | NA | VCL | vinculin | 7414 | ENSG00000035403 |
| 3 | O75369 | NA | FLNB | filamin B, beta | 2317 | ENSG00000136068 |
| 4 | P67936 | NA | TPM4 | tropomyosin 4 | 7171 | ENSG00000167460 |

  
  

| **Database:cellular component      &nbspName:protein complex      &nbspID:GO:0043234** | | | | | | |
| --- | --- | --- | --- | --- | --- | --- |
| C=3262; O=22; E=12.12; R=1.81; rawP=0.0024; adjP=0.0073 | | | | | | |
| Index | UserID | Value | Gene Symbol | Gene Name | EntrezGene | Ensembl |
| 1 | P63241 | NA | EIF5A | eukaryotic translation initiation factor 5A | 1984 | ENSG00000132507 |
| 2 | P78371 | NA | CCT2 | chaperonin containing TCP1, subunit 2 (beta) | 10576 | ENSG00000166226 |
| 3 | Q92598 | NA | HSPH1 | heat shock 105kDa/110kDa protein 1 | 10808 | ENSG00000120694 |
| 4 | Q13885 | NA | TUBB2A | tubulin, beta 2A class IIa | 7280 | ENSG00000137267 |
| 5 | P68371 | NA | TUBB4B | tubulin, beta 4B class IVb | 10383 | ENSG00000188229 |
| 6 | P67936 | NA | TPM4 | tropomyosin 4 | 7171 | ENSG00000167460 |
| 7 | P49368 | NA | CCT3 | chaperonin containing TCP1, subunit 3 (gamma) | 7203 | ENSG00000163468 |
| 8 | Q9BQE3 | NA | TUBA1C | tubulin, alpha 1c | 84790 | ENSG00000167553 |
| 9 | P49736 | NA | MCM2 | minichromosome maintenance complex component 2 | 4171 | ENSG00000073111 |
| 10 | Q15056 | NA | EIF4H | eukaryotic translation initiation factor 4H | 7458 | ENSG00000106682 |
| 11 | P50990 | NA | CCT8 | chaperonin containing TCP1, subunit 8 (theta) | 10694 | ENSG00000156261 |
| 12 | P68104 | NA | EEF1A1 | eukaryotic translation elongation factor 1 alpha 1 | 1915 | ENSG00000156508 |
| 13 | P50991 | NA | CCT4 | chaperonin containing TCP1, subunit 4 (delta) | 10575 | ENSG00000115484 |
| 14 | P17987 | NA | TCP1 | t-complex 1 | 6950 | ENSG00000120438 |
| 15 | P55209 | NA | NAP1L1 | nucleosome assembly protein 1-like 1 | 4673 | ENSG00000187109 |
| 16 | Q16555 | NA | DPYSL2 | dihydropyrimidinase-like 2 | 1808 | ENSG00000092964 |
| 17 | P40227 | NA | CCT6A | chaperonin containing TCP1, subunit 6A (zeta 1) | 908 | ENSG00000146731 |
| 18 | P67809 | NA | YBX1 | Y box binding protein 1 | 4904 | ENSG00000065978 |
| 19 | Q99832 | NA | CCT7 | chaperonin containing TCP1, subunit 7 (eta) | 10574 | ENSG00000135624 |
| 20 | P18206 | NA | VCL | vinculin | 7414 | ENSG00000035403 |
| 21 | Q8TEX9 | NA | IPO4 | importin 4 | 79711 | ENSG00000196497 |
| 22 | P11387 | NA | TOP1 | topoisomerase (DNA) I | 7150 | ENSG00000198900 |

  
  

| **Database:cellular component      &nbspName:myofibril      &nbspID:GO:0030016** | | | | | | |
| --- | --- | --- | --- | --- | --- | --- |
| C=152; O=4; E=0.56; R=7.08; rawP=0.0025; adjP=0.0073 | | | | | | |
| Index | UserID | Value | Gene Symbol | Gene Name | EntrezGene | Ensembl |
| 1 | P07951 | NA | TPM2 | tropomyosin 2 (beta) | 7169 | ENSG00000198467 |
| 2 | P18206 | NA | VCL | vinculin | 7414 | ENSG00000035403 |
| 3 | O75369 | NA | FLNB | filamin B, beta | 2317 | ENSG00000136068 |
| 4 | P67936 | NA | TPM4 | tropomyosin 4 | 7171 | ENSG00000167460 |

  
  

| **Database:cellular component      &nbspName:contractile fiber      &nbspID:GO:0043292** | | | | | | |
| --- | --- | --- | --- | --- | --- | --- |
| C=162; O=4; E=0.60; R=6.64; rawP=0.0031; adjP=0.0087 | | | | | | |
| Index | UserID | Value | Gene Symbol | Gene Name | EntrezGene | Ensembl |
| 1 | P07951 | NA | TPM2 | tropomyosin 2 (beta) | 7169 | ENSG00000198467 |
| 2 | P18206 | NA | VCL | vinculin | 7414 | ENSG00000035403 |
| 3 | O75369 | NA | FLNB | filamin B, beta | 2317 | ENSG00000136068 |
| 4 | P67936 | NA | TPM4 | tropomyosin 4 | 7171 | ENSG00000167460 |

  
  

| **Database:cellular component      &nbspName:intracellular organelle part      &nbspID:GO:0044446** | | | | | | |
| --- | --- | --- | --- | --- | --- | --- |
| C=6690; O=36; E=24.86; R=1.45; rawP=0.0032; adjP=0.0087 | | | | | | |
| Index | UserID | Value | Gene Symbol | Gene Name | EntrezGene | Ensembl |
| 1 | Q92598 | NA | HSPH1 | heat shock 105kDa/110kDa protein 1 | 10808 | ENSG00000120694 |
| 2 | Q13885 | NA | TUBB2A | tubulin, beta 2A class IIa | 7280 | ENSG00000137267 |
| 3 | P68371 | NA | TUBB4B | tubulin, beta 4B class IVb | 10383 | ENSG00000188229 |
| 4 | P25685 | NA | DNAJB1 | DnaJ (Hsp40) homolog, subfamily B, member 1 | 3337 | ENSG00000132002 |
| 5 | P53701 | NA | HCCS | holocytochrome c synthase | 3052 | ENSG00000004961 |
| 6 | O75369 | NA | FLNB | filamin B, beta | 2317 | ENSG00000136068 |
| 7 | P02786 | NA | TFRC | transferrin receptor (p90, CD71) | 7037 | ENSG00000072274 |
| 8 | P50991 | NA | CCT4 | chaperonin containing TCP1, subunit 4 (delta) | 10575 | ENSG00000115484 |
| 9 | P67809 | NA | YBX1 | Y box binding protein 1 | 4904 | ENSG00000065978 |
| 10 | Q15185 | NA | PTGES3 | prostaglandin E synthase 3 (cytosolic) | 10728 | ENSG00000110958 |
| 11 | Q99832 | NA | CCT7 | chaperonin containing TCP1, subunit 7 (eta) | 10574 | ENSG00000135624 |
| 12 | P37802 | NA | TAGLN2 | transgelin 2 | 8407 | ENSG00000158710 |
| 13 | P18206 | NA | VCL | vinculin | 7414 | ENSG00000035403 |
| 14 | P06703 | NA | S100A6 | S100 calcium binding protein A6 | 6277 | ENSG00000197956 |
| 15 | P11387 | NA | TOP1 | topoisomerase (DNA) I | 7150 | ENSG00000198900 |
| 16 | Q16881 | NA | TXNRD1 | thioredoxin reductase 1 | 7296 | ENSG00000198431 |
| 17 | P63241 | NA | EIF5A | eukaryotic translation initiation factor 5A | 1984 | ENSG00000132507 |
| 18 | P78371 | NA | CCT2 | chaperonin containing TCP1, subunit 2 (beta) | 10576 | ENSG00000166226 |
| 19 | Q9NR30 | NA | DDX21 | DEAD (Asp-Glu-Ala-Asp) box helicase 21 | 9188 | ENSG00000165732 |
| 20 | Q7Z2K6 | NA | ERMP1 | endoplasmic reticulum metallopeptidase 1 | 79956 | ENSG00000099219 |
| 21 | P07951 | NA | TPM2 | tropomyosin 2 (beta) | 7169 | ENSG00000198467 |
| 22 | Q9NZM1 | NA | MYOF | myoferlin | 26509 | ENSG00000138119 |
| 23 | P67936 | NA | TPM4 | tropomyosin 4 | 7171 | ENSG00000167460 |
| 24 | P49368 | NA | CCT3 | chaperonin containing TCP1, subunit 3 (gamma) | 7203 | ENSG00000163468 |
| 25 | Q9BQE3 | NA | TUBA1C | tubulin, alpha 1c | 84790 | ENSG00000167553 |
| 26 | P12268 | NA | IMPDH2 | IMP (inosine 5'-monophosphate) dehydrogenase 2 | 3615 | ENSG00000178035 |
| 27 | P49736 | NA | MCM2 | minichromosome maintenance complex component 2 | 4171 | ENSG00000073111 |
| 28 | P62906 | NA | RPL10A | ribosomal protein L10a | 4736 | ENSG00000198755 |
| 29 | P50990 | NA | CCT8 | chaperonin containing TCP1, subunit 8 (theta) | 10694 | ENSG00000156261 |
| 30 | O60610 | NA | DIAPH1 | diaphanous homolog 1 (Drosophila) | 1729 | ENSG00000131504 |
| 31 | P17987 | NA | TCP1 | t-complex 1 | 6950 | ENSG00000120438 |
| 32 | P34932 | NA | HSPA4 | heat shock 70kDa protein 4 | 3308 | ENSG00000170606 |
| 33 | P55209 | NA | NAP1L1 | nucleosome assembly protein 1-like 1 | 4673 | ENSG00000187109 |
| 34 | P40227 | NA | CCT6A | chaperonin containing TCP1, subunit 6A (zeta 1) | 908 | ENSG00000146731 |
| 35 | P52292 | NA | KPNA2 | karyopherin alpha 2 (RAG cohort 1, importin alpha 1) | 3838 | ENSG00000182481 |
| 36 | Q8TEX9 | NA | IPO4 | importin 4 | 79711 | ENSG00000196497 |

  
  

| **Database:cellular component      &nbspName:organelle part      &nbspID:GO:0044422** | | | | | | |
| --- | --- | --- | --- | --- | --- | --- |
| C=6777; O=36; E=25.19; R=1.43; rawP=0.0041; adjP=0.0107 | | | | | | |
| Index | UserID | Value | Gene Symbol | Gene Name | EntrezGene | Ensembl |
| 1 | Q92598 | NA | HSPH1 | heat shock 105kDa/110kDa protein 1 | 10808 | ENSG00000120694 |
| 2 | Q13885 | NA | TUBB2A | tubulin, beta 2A class IIa | 7280 | ENSG00000137267 |
| 3 | P68371 | NA | TUBB4B | tubulin, beta 4B class IVb | 10383 | ENSG00000188229 |
| 4 | P25685 | NA | DNAJB1 | DnaJ (Hsp40) homolog, subfamily B, member 1 | 3337 | ENSG00000132002 |
| 5 | P53701 | NA | HCCS | holocytochrome c synthase | 3052 | ENSG00000004961 |
| 6 | O75369 | NA | FLNB | filamin B, beta | 2317 | ENSG00000136068 |
| 7 | P02786 | NA | TFRC | transferrin receptor (p90, CD71) | 7037 | ENSG00000072274 |
| 8 | P50991 | NA | CCT4 | chaperonin containing TCP1, subunit 4 (delta) | 10575 | ENSG00000115484 |
| 9 | P67809 | NA | YBX1 | Y box binding protein 1 | 4904 | ENSG00000065978 |
| 10 | Q15185 | NA | PTGES3 | prostaglandin E synthase 3 (cytosolic) | 10728 | ENSG00000110958 |
| 11 | Q99832 | NA | CCT7 | chaperonin containing TCP1, subunit 7 (eta) | 10574 | ENSG00000135624 |
| 12 | P37802 | NA | TAGLN2 | transgelin 2 | 8407 | ENSG00000158710 |
| 13 | P18206 | NA | VCL | vinculin | 7414 | ENSG00000035403 |
| 14 | P06703 | NA | S100A6 | S100 calcium binding protein A6 | 6277 | ENSG00000197956 |
| 15 | P11387 | NA | TOP1 | topoisomerase (DNA) I | 7150 | ENSG00000198900 |
| 16 | Q16881 | NA | TXNRD1 | thioredoxin reductase 1 | 7296 | ENSG00000198431 |
| 17 | P63241 | NA | EIF5A | eukaryotic translation initiation factor 5A | 1984 | ENSG00000132507 |
| 18 | P78371 | NA | CCT2 | chaperonin containing TCP1, subunit 2 (beta) | 10576 | ENSG00000166226 |
| 19 | Q9NR30 | NA | DDX21 | DEAD (Asp-Glu-Ala-Asp) box helicase 21 | 9188 | ENSG00000165732 |
| 20 | Q7Z2K6 | NA | ERMP1 | endoplasmic reticulum metallopeptidase 1 | 79956 | ENSG00000099219 |
| 21 | P07951 | NA | TPM2 | tropomyosin 2 (beta) | 7169 | ENSG00000198467 |
| 22 | Q9NZM1 | NA | MYOF | myoferlin | 26509 | ENSG00000138119 |
| 23 | P67936 | NA | TPM4 | tropomyosin 4 | 7171 | ENSG00000167460 |
| 24 | P49368 | NA | CCT3 | chaperonin containing TCP1, subunit 3 (gamma) | 7203 | ENSG00000163468 |
| 25 | Q9BQE3 | NA | TUBA1C | tubulin, alpha 1c | 84790 | ENSG00000167553 |
| 26 | P12268 | NA | IMPDH2 | IMP (inosine 5'-monophosphate) dehydrogenase 2 | 3615 | ENSG00000178035 |
| 27 | P49736 | NA | MCM2 | minichromosome maintenance complex component 2 | 4171 | ENSG00000073111 |
| 28 | P62906 | NA | RPL10A | ribosomal protein L10a | 4736 | ENSG00000198755 |
| 29 | P50990 | NA | CCT8 | chaperonin containing TCP1, subunit 8 (theta) | 10694 | ENSG00000156261 |
| 30 | O60610 | NA | DIAPH1 | diaphanous homolog 1 (Drosophila) | 1729 | ENSG00000131504 |
| 31 | P17987 | NA | TCP1 | t-complex 1 | 6950 | ENSG00000120438 |
| 32 | P34932 | NA | HSPA4 | heat shock 70kDa protein 4 | 3308 | ENSG00000170606 |
| 33 | P55209 | NA | NAP1L1 | nucleosome assembly protein 1-like 1 | 4673 | ENSG00000187109 |
| 34 | P40227 | NA | CCT6A | chaperonin containing TCP1, subunit 6A (zeta 1) | 908 | ENSG00000146731 |
| 35 | P52292 | NA | KPNA2 | karyopherin alpha 2 (RAG cohort 1, importin alpha 1) | 3838 | ENSG00000182481 |
| 36 | Q8TEX9 | NA | IPO4 | importin 4 | 79711 | ENSG00000196497 |

  
  

| **Database:cellular component      &nbspName:nuclear envelope      &nbspID:GO:0005635** | | | | | | |
| --- | --- | --- | --- | --- | --- | --- |
| C=299; O=5; E=1.11; R=4.50; rawP=0.0050; adjP=0.0127 | | | | | | |
| Index | UserID | Value | Gene Symbol | Gene Name | EntrezGene | Ensembl |
| 1 | P63241 | NA | EIF5A | eukaryotic translation initiation factor 5A | 1984 | ENSG00000132507 |
| 2 | P37802 | NA | TAGLN2 | transgelin 2 | 8407 | ENSG00000158710 |
| 3 | Q9NZM1 | NA | MYOF | myoferlin | 26509 | ENSG00000138119 |
| 4 | Q8TEX9 | NA | IPO4 | importin 4 | 79711 | ENSG00000196497 |
| 5 | P06703 | NA | S100A6 | S100 calcium binding protein A6 | 6277 | ENSG00000197956 |

  
  

| **Database:cellular component      &nbspName:mitochondrion      &nbspID:GO:0005739** | | | | | | |
| --- | --- | --- | --- | --- | --- | --- |
| C=1515; O=12; E=5.63; R=2.13; rawP=0.0091; adjP=0.0223 | | | | | | |
| Index | UserID | Value | Gene Symbol | Gene Name | EntrezGene | Ensembl |
| 1 | Q16881 | NA | TXNRD1 | thioredoxin reductase 1 | 7296 | ENSG00000198431 |
| 2 | Q16555 | NA | DPYSL2 | dihydropyrimidinase-like 2 | 1808 | ENSG00000092964 |
| 3 | P07108 | NA | DBI | diazepam binding inhibitor (GABA receptor modulator, acyl-CoA binding protein) | 1622 | ENSG00000155368 |
| 4 | P49327 | NA | FASN | fatty acid synthase | 2194 | ENSG00000169710 |
| 5 | P56192 | NA | MARS | methionyl-tRNA synthetase | 4141 | ENSG00000166986 |
| 6 | Q99832 | NA | CCT7 | chaperonin containing TCP1, subunit 7 (eta) | 10574 | ENSG00000135624 |
| 7 | P21980 | NA | TGM2 | transglutaminase 2 (C polypeptide, protein-glutamine-gamma-glutamyltransferase) | 7052 | ENSG00000198959 |
| 8 | P62906 | NA | RPL10A | ribosomal protein L10a | 4736 | ENSG00000198755 |
| 9 | P53701 | NA | HCCS | holocytochrome c synthase | 3052 | ENSG00000004961 |
| 10 | Q06830 | NA | PRDX1 | peroxiredoxin 1 | 5052 | ENSG00000117450 |
| 11 | P02786 | NA | TFRC | transferrin receptor (p90, CD71) | 7037 | ENSG00000072274 |
| 12 | P11586 | NA | MTHFD1 | methylenetetrahydrofolate dehydrogenase (NADP+ dependent) 1, methenyltetrahydrofolate cyclohydrolase, formyltetrahydrofolate synthetase | 4522 | ENSG00000100714 |

  
  

| **Database:cellular component      &nbspName:sarcomere      &nbspID:GO:0030017** | | | | | | |
| --- | --- | --- | --- | --- | --- | --- |
| C=128; O=3; E=0.48; R=6.31; rawP=0.0120; adjP=0.0285 | | | | | | |
| Index | UserID | Value | Gene Symbol | Gene Name | EntrezGene | Ensembl |
| 1 | P07951 | NA | TPM2 | tropomyosin 2 (beta) | 7169 | ENSG00000198467 |
| 2 | O75369 | NA | FLNB | filamin B, beta | 2317 | ENSG00000136068 |
| 3 | P67936 | NA | TPM4 | tropomyosin 4 | 7171 | ENSG00000167460 |

  
  

| **Database:cellular component      &nbspName:nuclear part      &nbspID:GO:0044428** | | | | | | |
| --- | --- | --- | --- | --- | --- | --- |
| C=3057; O=19; E=11.36; R=1.67; rawP=0.0129; adjP=0.0297 | | | | | | |
| Index | UserID | Value | Gene Symbol | Gene Name | EntrezGene | Ensembl |
| 1 | P63241 | NA | EIF5A | eukaryotic translation initiation factor 5A | 1984 | ENSG00000132507 |
| 2 | Q16881 | NA | TXNRD1 | thioredoxin reductase 1 | 7296 | ENSG00000198431 |
| 3 | Q9NR30 | NA | DDX21 | DEAD (Asp-Glu-Ala-Asp) box helicase 21 | 9188 | ENSG00000165732 |
| 4 | P78371 | NA | CCT2 | chaperonin containing TCP1, subunit 2 (beta) | 10576 | ENSG00000166226 |
| 5 | Q92598 | NA | HSPH1 | heat shock 105kDa/110kDa protein 1 | 10808 | ENSG00000120694 |
| 6 | Q9NZM1 | NA | MYOF | myoferlin | 26509 | ENSG00000138119 |
| 7 | P25685 | NA | DNAJB1 | DnaJ (Hsp40) homolog, subfamily B, member 1 | 3337 | ENSG00000132002 |
| 8 | P49736 | NA | MCM2 | minichromosome maintenance complex component 2 | 4171 | ENSG00000073111 |
| 9 | P50991 | NA | CCT4 | chaperonin containing TCP1, subunit 4 (delta) | 10575 | ENSG00000115484 |
| 10 | P17987 | NA | TCP1 | t-complex 1 | 6950 | ENSG00000120438 |
| 11 | P34932 | NA | HSPA4 | heat shock 70kDa protein 4 | 3308 | ENSG00000170606 |
| 12 | P55209 | NA | NAP1L1 | nucleosome assembly protein 1-like 1 | 4673 | ENSG00000187109 |
| 13 | P52292 | NA | KPNA2 | karyopherin alpha 2 (RAG cohort 1, importin alpha 1) | 3838 | ENSG00000182481 |
| 14 | P67809 | NA | YBX1 | Y box binding protein 1 | 4904 | ENSG00000065978 |
| 15 | P37802 | NA | TAGLN2 | transgelin 2 | 8407 | ENSG00000158710 |
| 16 | Q15185 | NA | PTGES3 | prostaglandin E synthase 3 (cytosolic) | 10728 | ENSG00000110958 |
| 17 | Q8TEX9 | NA | IPO4 | importin 4 | 79711 | ENSG00000196497 |
| 18 | P06703 | NA | S100A6 | S100 calcium binding protein A6 | 6277 | ENSG00000197956 |
| 19 | P11387 | NA | TOP1 | topoisomerase (DNA) I | 7150 | ENSG00000198900 |

  
  

| **Database:cellular component      &nbspName:cytoplasmic membrane-bounded vesicle      &nbspID:GO:0016023** | | | | | | |
| --- | --- | --- | --- | --- | --- | --- |
| C=861; O=8; E=3.20; R=2.50; rawP=0.0140; adjP=0.0313 | | | | | | |
| Index | UserID | Value | Gene Symbol | Gene Name | EntrezGene | Ensembl |
| 1 | P55209 | NA | NAP1L1 | nucleosome assembly protein 1-like 1 | 4673 | ENSG00000187109 |
| 2 | P08195 | NA | SLC3A2 | solute carrier family 3 (activators of dibasic and neutral amino acid transport), member 2 | 6520 | ENSG00000168003 |
| 3 | Q9NZM1 | NA | MYOF | myoferlin | 26509 | ENSG00000138119 |
| 4 | P49327 | NA | FASN | fatty acid synthase | 2194 | ENSG00000169710 |
| 5 | Q15758 | NA | SLC1A5 | solute carrier family 1 (neutral amino acid transporter), member 5 | 6510 | ENSG00000105281 |
| 6 | P02786 | NA | TFRC | transferrin receptor (p90, CD71) | 7037 | ENSG00000072274 |
| 7 | Q06830 | NA | PRDX1 | peroxiredoxin 1 | 5052 | ENSG00000117450 |
| 8 | P50991 | NA | CCT4 | chaperonin containing TCP1, subunit 4 (delta) | 10575 | ENSG00000115484 |

  
  

| **Database:cellular component      &nbspName:membrane-bounded vesicle      &nbspID:GO:0031988** | | | | | | |
| --- | --- | --- | --- | --- | --- | --- |
| C=882; O=8; E=3.28; R=2.44; rawP=0.0159; adjP=0.0345 | | | | | | |
| Index | UserID | Value | Gene Symbol | Gene Name | EntrezGene | Ensembl |
| 1 | P55209 | NA | NAP1L1 | nucleosome assembly protein 1-like 1 | 4673 | ENSG00000187109 |
| 2 | P08195 | NA | SLC3A2 | solute carrier family 3 (activators of dibasic and neutral amino acid transport), member 2 | 6520 | ENSG00000168003 |
| 3 | Q9NZM1 | NA | MYOF | myoferlin | 26509 | ENSG00000138119 |
| 4 | P49327 | NA | FASN | fatty acid synthase | 2194 | ENSG00000169710 |
| 5 | Q15758 | NA | SLC1A5 | solute carrier family 1 (neutral amino acid transporter), member 5 | 6510 | ENSG00000105281 |
| 6 | P02786 | NA | TFRC | transferrin receptor (p90, CD71) | 7037 | ENSG00000072274 |
| 7 | Q06830 | NA | PRDX1 | peroxiredoxin 1 | 5052 | ENSG00000117450 |
| 8 | P50991 | NA | CCT4 | chaperonin containing TCP1, subunit 4 (delta) | 10575 | ENSG00000115484 |

  
  

| **Database:cellular component      &nbspName:cytoplasmic vesicle      &nbspID:GO:0031410** | | | | | | |
| --- | --- | --- | --- | --- | --- | --- |
| C=927; O=8; E=3.45; R=2.32; rawP=0.0209; adjP=0.0441 | | | | | | |
| Index | UserID | Value | Gene Symbol | Gene Name | EntrezGene | Ensembl |
| 1 | P55209 | NA | NAP1L1 | nucleosome assembly protein 1-like 1 | 4673 | ENSG00000187109 |
| 2 | P08195 | NA | SLC3A2 | solute carrier family 3 (activators of dibasic and neutral amino acid transport), member 2 | 6520 | ENSG00000168003 |
| 3 | Q9NZM1 | NA | MYOF | myoferlin | 26509 | ENSG00000138119 |
| 4 | P49327 | NA | FASN | fatty acid synthase | 2194 | ENSG00000169710 |
| 5 | Q15758 | NA | SLC1A5 | solute carrier family 1 (neutral amino acid transporter), member 5 | 6510 | ENSG00000105281 |
| 6 | P02786 | NA | TFRC | transferrin receptor (p90, CD71) | 7037 | ENSG00000072274 |
| 7 | Q06830 | NA | PRDX1 | peroxiredoxin 1 | 5052 | ENSG00000117450 |
| 8 | P50991 | NA | CCT4 | chaperonin containing TCP1, subunit 4 (delta) | 10575 | ENSG00000115484 |

  
  

| **Database:cellular component      &nbspName:vesicle      &nbspID:GO:0031982** | | | | | | |
| --- | --- | --- | --- | --- | --- | --- |
| C=969; O=8; E=3.60; R=2.22; rawP=0.0264; adjP=0.0542 | | | | | | |
| Index | UserID | Value | Gene Symbol | Gene Name | EntrezGene | Ensembl |
| 1 | P55209 | NA | NAP1L1 | nucleosome assembly protein 1-like 1 | 4673 | ENSG00000187109 |
| 2 | P08195 | NA | SLC3A2 | solute carrier family 3 (activators of dibasic and neutral amino acid transport), member 2 | 6520 | ENSG00000168003 |
| 3 | Q9NZM1 | NA | MYOF | myoferlin | 26509 | ENSG00000138119 |
| 4 | P49327 | NA | FASN | fatty acid synthase | 2194 | ENSG00000169710 |
| 5 | Q15758 | NA | SLC1A5 | solute carrier family 1 (neutral amino acid transporter), member 5 | 6510 | ENSG00000105281 |
| 6 | P02786 | NA | TFRC | transferrin receptor (p90, CD71) | 7037 | ENSG00000072274 |
| 7 | Q06830 | NA | PRDX1 | peroxiredoxin 1 | 5052 | ENSG00000117450 |
| 8 | P50991 | NA | CCT4 | chaperonin containing TCP1, subunit 4 (delta) | 10575 | ENSG00000115484 |

  
  

| **Database:cellular component      &nbspName:membrane-bounded organelle      &nbspID:GO:0043227** | | | | | | |
| --- | --- | --- | --- | --- | --- | --- |
| C=9495; O=43; E=35.29; R=1.22; rawP=0.0304; adjP=0.0592 | | | | | | |
| Index | UserID | Value | Gene Symbol | Gene Name | EntrezGene | Ensembl |
| 1 | Q92598 | NA | HSPH1 | heat shock 105kDa/110kDa protein 1 | 10808 | ENSG00000120694 |
| 2 | P68371 | NA | TUBB4B | tubulin, beta 4B class IVb | 10383 | ENSG00000188229 |
| 3 | P25685 | NA | DNAJB1 | DnaJ (Hsp40) homolog, subfamily B, member 1 | 3337 | ENSG00000132002 |
| 4 | P53701 | NA | HCCS | holocytochrome c synthase | 3052 | ENSG00000004961 |
| 5 | P68104 | NA | EEF1A1 | eukaryotic translation elongation factor 1 alpha 1 | 1915 | ENSG00000156508 |
| 6 | P02786 | NA | TFRC | transferrin receptor (p90, CD71) | 7037 | ENSG00000072274 |
| 7 | P50991 | NA | CCT4 | chaperonin containing TCP1, subunit 4 (delta) | 10575 | ENSG00000115484 |
| 8 | O00425 | NA | IGF2BP3 | insulin-like growth factor 2 mRNA binding protein 3 | 10643 | ENSG00000136231 |
| 9 | Q16555 | NA | DPYSL2 | dihydropyrimidinase-like 2 | 1808 | ENSG00000092964 |
| 10 | O95433 | NA | AHSA1 | AHA1, activator of heat shock 90kDa protein ATPase homolog 1 (yeast) | 10598 | ENSG00000100591 |
| 11 | P17174 | NA | GOT1 | glutamic-oxaloacetic transaminase 1, soluble (aspartate aminotransferase 1) | 2805 | ENSG00000120053 |
| 12 | P67809 | NA | YBX1 | Y box binding protein 1 | 4904 | ENSG00000065978 |
| 13 | Q15185 | NA | PTGES3 | prostaglandin E synthase 3 (cytosolic) | 10728 | ENSG00000110958 |
| 14 | Q99832 | NA | CCT7 | chaperonin containing TCP1, subunit 7 (eta) | 10574 | ENSG00000135624 |
| 15 | P37802 | NA | TAGLN2 | transgelin 2 | 8407 | ENSG00000158710 |
| 16 | P21980 | NA | TGM2 | transglutaminase 2 (C polypeptide, protein-glutamine-gamma-glutamyltransferase) | 7052 | ENSG00000198959 |
| 17 | Q6PIU2 | NA | NCEH1 | neutral cholesterol ester hydrolase 1 | 57552 | ENSG00000144959 |
| 18 | P06703 | NA | S100A6 | S100 calcium binding protein A6 | 6277 | ENSG00000197956 |
| 19 | Q9UK76 | NA | HN1 | hematological and neurological expressed 1 | 51155 | ENSG00000189159 |
| 20 | P11387 | NA | TOP1 | topoisomerase (DNA) I | 7150 | ENSG00000198900 |
| 21 | P11586 | NA | MTHFD1 | methylenetetrahydrofolate dehydrogenase (NADP+ dependent) 1, methenyltetrahydrofolate cyclohydrolase, formyltetrahydrofolate synthetase | 4522 | ENSG00000100714 |
| 22 | Q16881 | NA | TXNRD1 | thioredoxin reductase 1 | 7296 | ENSG00000198431 |
| 23 | P63241 | NA | EIF5A | eukaryotic translation initiation factor 5A | 1984 | ENSG00000132507 |
| 24 | P78371 | NA | CCT2 | chaperonin containing TCP1, subunit 2 (beta) | 10576 | ENSG00000166226 |
| 25 | Q9NR30 | NA | DDX21 | DEAD (Asp-Glu-Ala-Asp) box helicase 21 | 9188 | ENSG00000165732 |
| 26 | P08195 | NA | SLC3A2 | solute carrier family 3 (activators of dibasic and neutral amino acid transport), member 2 | 6520 | ENSG00000168003 |
| 27 | Q7Z2K6 | NA | ERMP1 | endoplasmic reticulum metallopeptidase 1 | 79956 | ENSG00000099219 |
| 28 | P02795 | NA | MT2A | metallothionein 2A | 4502 | ENSG00000125148 |
| 29 | Q9NZM1 | NA | MYOF | myoferlin | 26509 | ENSG00000138119 |
| 30 | P49321 | NA | NASP | nuclear autoantigenic sperm protein (histone-binding) | 4678 | ENSG00000132780 |
| 31 | P49327 | NA | FASN | fatty acid synthase | 2194 | ENSG00000169710 |
| 32 | P56192 | NA | MARS | methionyl-tRNA synthetase | 4141 | ENSG00000166986 |
| 33 | P12268 | NA | IMPDH2 | IMP (inosine 5'-monophosphate) dehydrogenase 2 | 3615 | ENSG00000178035 |
| 34 | P62906 | NA | RPL10A | ribosomal protein L10a | 4736 | ENSG00000198755 |
| 35 | P49736 | NA | MCM2 | minichromosome maintenance complex component 2 | 4171 | ENSG00000073111 |
| 36 | Q06830 | NA | PRDX1 | peroxiredoxin 1 | 5052 | ENSG00000117450 |
| 37 | P17987 | NA | TCP1 | t-complex 1 | 6950 | ENSG00000120438 |
| 38 | P55209 | NA | NAP1L1 | nucleosome assembly protein 1-like 1 | 4673 | ENSG00000187109 |
| 39 | P34932 | NA | HSPA4 | heat shock 70kDa protein 4 | 3308 | ENSG00000170606 |
| 40 | P07108 | NA | DBI | diazepam binding inhibitor (GABA receptor modulator, acyl-CoA binding protein) | 1622 | ENSG00000155368 |
| 41 | P52292 | NA | KPNA2 | karyopherin alpha 2 (RAG cohort 1, importin alpha 1) | 3838 | ENSG00000182481 |
| 42 | Q15758 | NA | SLC1A5 | solute carrier family 1 (neutral amino acid transporter), member 5 | 6510 | ENSG00000105281 |
| 43 | Q8TEX9 | NA | IPO4 | importin 4 | 79711 | ENSG00000196497 |

  
  

| **Database:cellular component      &nbspName:intracellular membrane-bounded organelle      &nbspID:GO:0043231** | | | | | | |
| --- | --- | --- | --- | --- | --- | --- |
| C=9484; O=43; E=35.25; R=1.22; rawP=0.0297; adjP=0.0592 | | | | | | |
| Index | UserID | Value | Gene Symbol | Gene Name | EntrezGene | Ensembl |
| 1 | Q92598 | NA | HSPH1 | heat shock 105kDa/110kDa protein 1 | 10808 | ENSG00000120694 |
| 2 | P68371 | NA | TUBB4B | tubulin, beta 4B class IVb | 10383 | ENSG00000188229 |
| 3 | P25685 | NA | DNAJB1 | DnaJ (Hsp40) homolog, subfamily B, member 1 | 3337 | ENSG00000132002 |
| 4 | P53701 | NA | HCCS | holocytochrome c synthase | 3052 | ENSG00000004961 |
| 5 | P68104 | NA | EEF1A1 | eukaryotic translation elongation factor 1 alpha 1 | 1915 | ENSG00000156508 |
| 6 | P02786 | NA | TFRC | transferrin receptor (p90, CD71) | 7037 | ENSG00000072274 |
| 7 | P50991 | NA | CCT4 | chaperonin containing TCP1, subunit 4 (delta) | 10575 | ENSG00000115484 |
| 8 | O00425 | NA | IGF2BP3 | insulin-like growth factor 2 mRNA binding protein 3 | 10643 | ENSG00000136231 |
| 9 | Q16555 | NA | DPYSL2 | dihydropyrimidinase-like 2 | 1808 | ENSG00000092964 |
| 10 | O95433 | NA | AHSA1 | AHA1, activator of heat shock 90kDa protein ATPase homolog 1 (yeast) | 10598 | ENSG00000100591 |
| 11 | P17174 | NA | GOT1 | glutamic-oxaloacetic transaminase 1, soluble (aspartate aminotransferase 1) | 2805 | ENSG00000120053 |
| 12 | P67809 | NA | YBX1 | Y box binding protein 1 | 4904 | ENSG00000065978 |
| 13 | Q15185 | NA | PTGES3 | prostaglandin E synthase 3 (cytosolic) | 10728 | ENSG00000110958 |
| 14 | Q99832 | NA | CCT7 | chaperonin containing TCP1, subunit 7 (eta) | 10574 | ENSG00000135624 |
| 15 | P37802 | NA | TAGLN2 | transgelin 2 | 8407 | ENSG00000158710 |
| 16 | P21980 | NA | TGM2 | transglutaminase 2 (C polypeptide, protein-glutamine-gamma-glutamyltransferase) | 7052 | ENSG00000198959 |
| 17 | Q6PIU2 | NA | NCEH1 | neutral cholesterol ester hydrolase 1 | 57552 | ENSG00000144959 |
| 18 | P06703 | NA | S100A6 | S100 calcium binding protein A6 | 6277 | ENSG00000197956 |
| 19 | Q9UK76 | NA | HN1 | hematological and neurological expressed 1 | 51155 | ENSG00000189159 |
| 20 | P11387 | NA | TOP1 | topoisomerase (DNA) I | 7150 | ENSG00000198900 |
| 21 | P11586 | NA | MTHFD1 | methylenetetrahydrofolate dehydrogenase (NADP+ dependent) 1, methenyltetrahydrofolate cyclohydrolase, formyltetrahydrofolate synthetase | 4522 | ENSG00000100714 |
| 22 | Q16881 | NA | TXNRD1 | thioredoxin reductase 1 | 7296 | ENSG00000198431 |
| 23 | P63241 | NA | EIF5A | eukaryotic translation initiation factor 5A | 1984 | ENSG00000132507 |
| 24 | P78371 | NA | CCT2 | chaperonin containing TCP1, subunit 2 (beta) | 10576 | ENSG00000166226 |
| 25 | Q9NR30 | NA | DDX21 | DEAD (Asp-Glu-Ala-Asp) box helicase 21 | 9188 | ENSG00000165732 |
| 26 | P08195 | NA | SLC3A2 | solute carrier family 3 (activators of dibasic and neutral amino acid transport), member 2 | 6520 | ENSG00000168003 |
| 27 | Q7Z2K6 | NA | ERMP1 | endoplasmic reticulum metallopeptidase 1 | 79956 | ENSG00000099219 |
| 28 | P02795 | NA | MT2A | metallothionein 2A | 4502 | ENSG00000125148 |
| 29 | Q9NZM1 | NA | MYOF | myoferlin | 26509 | ENSG00000138119 |
| 30 | P49321 | NA | NASP | nuclear autoantigenic sperm protein (histone-binding) | 4678 | ENSG00000132780 |
| 31 | P49327 | NA | FASN | fatty acid synthase | 2194 | ENSG00000169710 |
| 32 | P56192 | NA | MARS | methionyl-tRNA synthetase | 4141 | ENSG00000166986 |
| 33 | P12268 | NA | IMPDH2 | IMP (inosine 5'-monophosphate) dehydrogenase 2 | 3615 | ENSG00000178035 |
| 34 | P62906 | NA | RPL10A | ribosomal protein L10a | 4736 | ENSG00000198755 |
| 35 | P49736 | NA | MCM2 | minichromosome maintenance complex component 2 | 4171 | ENSG00000073111 |
| 36 | Q06830 | NA | PRDX1 | peroxiredoxin 1 | 5052 | ENSG00000117450 |
| 37 | P17987 | NA | TCP1 | t-complex 1 | 6950 | ENSG00000120438 |
| 38 | P55209 | NA | NAP1L1 | nucleosome assembly protein 1-like 1 | 4673 | ENSG00000187109 |
| 39 | P34932 | NA | HSPA4 | heat shock 70kDa protein 4 | 3308 | ENSG00000170606 |
| 40 | P07108 | NA | DBI | diazepam binding inhibitor (GABA receptor modulator, acyl-CoA binding protein) | 1622 | ENSG00000155368 |
| 41 | P52292 | NA | KPNA2 | karyopherin alpha 2 (RAG cohort 1, importin alpha 1) | 3838 | ENSG00000182481 |
| 42 | Q15758 | NA | SLC1A5 | solute carrier family 1 (neutral amino acid transporter), member 5 | 6510 | ENSG00000105281 |
| 43 | Q8TEX9 | NA | IPO4 | importin 4 | 79711 | ENSG00000196497 |

  
  

| **Database:cellular component      &nbspName:actin cytoskeleton      &nbspID:GO:0015629** | | | | | | |
| --- | --- | --- | --- | --- | --- | --- |
| C=365; O=4; E=1.36; R=2.95; rawP=0.0467; adjP=0.0887 | | | | | | |
| Index | UserID | Value | Gene Symbol | Gene Name | EntrezGene | Ensembl |
| 1 | P07951 | NA | TPM2 | tropomyosin 2 (beta) | 7169 | ENSG00000198467 |
| 2 | P18206 | NA | VCL | vinculin | 7414 | ENSG00000035403 |
| 3 | O75369 | NA | FLNB | filamin B, beta | 2317 | ENSG00000136068 |
| 4 | P67936 | NA | TPM4 | tropomyosin 4 | 7171 | ENSG00000167460 |

  
  
  
  

---

WebGestalt is currently developed and maintained by Jing Wang and Bing Zhang at the  Zhang Lab. Other people who have made significant contribution to the project include Dexter Duncan, Stefan Kirov, Zhiao Shi, and Jay Snoddy.  
  
**Funding credits:** NIH/NIAAA (U01 AA016662, U01 AA013512); NIH/NIDA (P01 DA015027); NIH/NIMH (P50 MH078028, P50 MH096972); NIH/NCI (U24 CA159988); NIH/NIGMS (R01 GM088822).
