## Supplementary material for "PGRMC1 phosphorylation and cell plasticity 1: glycolysis, mitochondria, tumor growth": File S8: final_pc_geneset_file_1439167306.html

Anchored HTML File of EIDs

|  |  |
| --- | --- |
|  | WEB-based GEne SeT AnaLysis Toolkit |
|  | ***Translating gene lists into biological insights...*** |

---

  

| **Database:Pathway Commons pathway      &nbspName:Formation of tubulin folding intermediates by CCT/TriC      &nbspID:DB\_ID:704** | | | | | | |
| --- | --- | --- | --- | --- | --- | --- |
| C=8; O=7; E=0.02; R=281.71; rawP=1.58e-17; adjP=1.15e-15 | | | | | | |
| Index | UserID | Value | Gene Symbol | Gene Name | EntrezGene | Ensembl |
| 1 | P40227 | NA | CCT6A | chaperonin containing TCP1, subunit 6A (zeta 1) | 908 | ENSG00000146731 |
| 2 | P49368 | NA | CCT3 | chaperonin containing TCP1, subunit 3 (gamma) | 7203 | ENSG00000163468 |
| 3 | P78371 | NA | CCT2 | chaperonin containing TCP1, subunit 2 (beta) | 10576 | ENSG00000166226 |
| 4 | P17987 | NA | TCP1 | t-complex 1 | 6950 | ENSG00000120438 |
| 5 | Q99832 | NA | CCT7 | chaperonin containing TCP1, subunit 7 (eta) | 10574 | ENSG00000135624 |
| 6 | P50990 | NA | CCT8 | chaperonin containing TCP1, subunit 8 (theta) | 10694 | ENSG00000156261 |
| 7 | P50991 | NA | CCT4 | chaperonin containing TCP1, subunit 4 (delta) | 10575 | ENSG00000115484 |

  
  

| **Database:Pathway Commons pathway      &nbspName:Folding of actin by CCT/TriC      &nbspID:DB\_ID:699** | | | | | | |
| --- | --- | --- | --- | --- | --- | --- |
| C=9; O=7; E=0.03; R=250.41; rawP=7.08e-17; adjP=1.72e-15 | | | | | | |
| Index | UserID | Value | Gene Symbol | Gene Name | EntrezGene | Ensembl |
| 1 | P40227 | NA | CCT6A | chaperonin containing TCP1, subunit 6A (zeta 1) | 908 | ENSG00000146731 |
| 2 | P49368 | NA | CCT3 | chaperonin containing TCP1, subunit 3 (gamma) | 7203 | ENSG00000163468 |
| 3 | P78371 | NA | CCT2 | chaperonin containing TCP1, subunit 2 (beta) | 10576 | ENSG00000166226 |
| 4 | P17987 | NA | TCP1 | t-complex 1 | 6950 | ENSG00000120438 |
| 5 | Q99832 | NA | CCT7 | chaperonin containing TCP1, subunit 7 (eta) | 10574 | ENSG00000135624 |
| 6 | P50990 | NA | CCT8 | chaperonin containing TCP1, subunit 8 (theta) | 10694 | ENSG00000156261 |
| 7 | P50991 | NA | CCT4 | chaperonin containing TCP1, subunit 4 (delta) | 10575 | ENSG00000115484 |

  
  

| **Database:Pathway Commons pathway      &nbspName:Association of TriC/CCT with target proteins during biosynthesis      &nbspID:DB\_ID:700** | | | | | | |
| --- | --- | --- | --- | --- | --- | --- |
| C=9; O=7; E=0.03; R=250.41; rawP=7.08e-17; adjP=1.72e-15 | | | | | | |
| Index | UserID | Value | Gene Symbol | Gene Name | EntrezGene | Ensembl |
| 1 | P40227 | NA | CCT6A | chaperonin containing TCP1, subunit 6A (zeta 1) | 908 | ENSG00000146731 |
| 2 | P49368 | NA | CCT3 | chaperonin containing TCP1, subunit 3 (gamma) | 7203 | ENSG00000163468 |
| 3 | P78371 | NA | CCT2 | chaperonin containing TCP1, subunit 2 (beta) | 10576 | ENSG00000166226 |
| 4 | P17987 | NA | TCP1 | t-complex 1 | 6950 | ENSG00000120438 |
| 5 | Q99832 | NA | CCT7 | chaperonin containing TCP1, subunit 7 (eta) | 10574 | ENSG00000135624 |
| 6 | P50990 | NA | CCT8 | chaperonin containing TCP1, subunit 8 (theta) | 10694 | ENSG00000156261 |
| 7 | P50991 | NA | CCT4 | chaperonin containing TCP1, subunit 4 (delta) | 10575 | ENSG00000115484 |

  
  

| **Database:Pathway Commons pathway      &nbspName:Protein folding      &nbspID:DB\_ID:712** | | | | | | |
| --- | --- | --- | --- | --- | --- | --- |
| C=28; O=9; E=0.09; R=103.48; rawP=9.77e-17; adjP=1.78e-15 | | | | | | |
| Index | UserID | Value | Gene Symbol | Gene Name | EntrezGene | Ensembl |
| 1 | P78371 | NA | CCT2 | chaperonin containing TCP1, subunit 2 (beta) | 10576 | ENSG00000166226 |
| 2 | P17987 | NA | TCP1 | t-complex 1 | 6950 | ENSG00000120438 |
| 3 | Q13885 | NA | TUBB2A | tubulin, beta 2A class IIa | 7280 | ENSG00000137267 |
| 4 | P68371 | NA | TUBB4B | tubulin, beta 4B class IVb | 10383 | ENSG00000188229 |
| 5 | P40227 | NA | CCT6A | chaperonin containing TCP1, subunit 6A (zeta 1) | 908 | ENSG00000146731 |
| 6 | P49368 | NA | CCT3 | chaperonin containing TCP1, subunit 3 (gamma) | 7203 | ENSG00000163468 |
| 7 | Q99832 | NA | CCT7 | chaperonin containing TCP1, subunit 7 (eta) | 10574 | ENSG00000135624 |
| 8 | P50990 | NA | CCT8 | chaperonin containing TCP1, subunit 8 (theta) | 10694 | ENSG00000156261 |
| 9 | P50991 | NA | CCT4 | chaperonin containing TCP1, subunit 4 (delta) | 10575 | ENSG00000115484 |

  
  

| **Database:Pathway Commons pathway      &nbspName:Prefoldin mediated transfer of substrate to CCT/TriC      &nbspID:DB\_ID:703** | | | | | | |
| --- | --- | --- | --- | --- | --- | --- |
| C=14; O=7; E=0.04; R=160.98; rawP=6.67e-15; adjP=9.74e-14 | | | | | | |
| Index | UserID | Value | Gene Symbol | Gene Name | EntrezGene | Ensembl |
| 1 | P40227 | NA | CCT6A | chaperonin containing TCP1, subunit 6A (zeta 1) | 908 | ENSG00000146731 |
| 2 | P49368 | NA | CCT3 | chaperonin containing TCP1, subunit 3 (gamma) | 7203 | ENSG00000163468 |
| 3 | P78371 | NA | CCT2 | chaperonin containing TCP1, subunit 2 (beta) | 10576 | ENSG00000166226 |
| 4 | P17987 | NA | TCP1 | t-complex 1 | 6950 | ENSG00000120438 |
| 5 | Q99832 | NA | CCT7 | chaperonin containing TCP1, subunit 7 (eta) | 10574 | ENSG00000135624 |
| 6 | P50990 | NA | CCT8 | chaperonin containing TCP1, subunit 8 (theta) | 10694 | ENSG00000156261 |
| 7 | P50991 | NA | CCT4 | chaperonin containing TCP1, subunit 4 (delta) | 10575 | ENSG00000115484 |

  
  

| **Database:Pathway Commons pathway      &nbspName:Cooperation of Prefoldin and TriC/CCT in actin and tubulin folding      &nbspID:DB\_ID:709** | | | | | | |
| --- | --- | --- | --- | --- | --- | --- |
| C=15; O=7; E=0.05; R=150.24; rawP=1.25e-14; adjP=1.52e-13 | | | | | | |
| Index | UserID | Value | Gene Symbol | Gene Name | EntrezGene | Ensembl |
| 1 | P40227 | NA | CCT6A | chaperonin containing TCP1, subunit 6A (zeta 1) | 908 | ENSG00000146731 |
| 2 | P49368 | NA | CCT3 | chaperonin containing TCP1, subunit 3 (gamma) | 7203 | ENSG00000163468 |
| 3 | P78371 | NA | CCT2 | chaperonin containing TCP1, subunit 2 (beta) | 10576 | ENSG00000166226 |
| 4 | P17987 | NA | TCP1 | t-complex 1 | 6950 | ENSG00000120438 |
| 5 | Q99832 | NA | CCT7 | chaperonin containing TCP1, subunit 7 (eta) | 10574 | ENSG00000135624 |
| 6 | P50990 | NA | CCT8 | chaperonin containing TCP1, subunit 8 (theta) | 10694 | ENSG00000156261 |
| 7 | P50991 | NA | CCT4 | chaperonin containing TCP1, subunit 4 (delta) | 10575 | ENSG00000115484 |

  
  

| **Database:Pathway Commons pathway      &nbspName:Chaperonin-mediated protein folding      &nbspID:DB\_ID:710** | | | | | | |
| --- | --- | --- | --- | --- | --- | --- |
| C=16; O=7; E=0.05; R=140.85; rawP=2.21e-14; adjP=2.30e-13 | | | | | | |
| Index | UserID | Value | Gene Symbol | Gene Name | EntrezGene | Ensembl |
| 1 | P40227 | NA | CCT6A | chaperonin containing TCP1, subunit 6A (zeta 1) | 908 | ENSG00000146731 |
| 2 | P49368 | NA | CCT3 | chaperonin containing TCP1, subunit 3 (gamma) | 7203 | ENSG00000163468 |
| 3 | P78371 | NA | CCT2 | chaperonin containing TCP1, subunit 2 (beta) | 10576 | ENSG00000166226 |
| 4 | P17987 | NA | TCP1 | t-complex 1 | 6950 | ENSG00000120438 |
| 5 | Q99832 | NA | CCT7 | chaperonin containing TCP1, subunit 7 (eta) | 10574 | ENSG00000135624 |
| 6 | P50990 | NA | CCT8 | chaperonin containing TCP1, subunit 8 (theta) | 10694 | ENSG00000156261 |
| 7 | P50991 | NA | CCT4 | chaperonin containing TCP1, subunit 4 (delta) | 10575 | ENSG00000115484 |

  
  

| **Database:Pathway Commons pathway      &nbspName:Metabolism of proteins      &nbspID:DB\_ID:728** | | | | | | |
| --- | --- | --- | --- | --- | --- | --- |
| C=260; O=14; E=0.81; R=17.34; rawP=4.87e-14; adjP=4.44e-13 | | | | | | |
| Index | UserID | Value | Gene Symbol | Gene Name | EntrezGene | Ensembl |
| 1 | P78371 | NA | CCT2 | chaperonin containing TCP1, subunit 2 (beta) | 10576 | ENSG00000166226 |
| 2 | P17987 | NA | TCP1 | t-complex 1 | 6950 | ENSG00000120438 |
| 3 | Q13885 | NA | TUBB2A | tubulin, beta 2A class IIa | 7280 | ENSG00000137267 |
| 4 | P13639 | NA | EEF2 | eukaryotic translation elongation factor 2 | 1938 | ENSG00000167658 |
| 5 | P41091 | NA | EIF2S3 | eukaryotic translation initiation factor 2, subunit 3 gamma, 52kDa | 1968 | ENSG00000130741 |
| 6 | P68371 | NA | TUBB4B | tubulin, beta 4B class IVb | 10383 | ENSG00000188229 |
| 7 | P40227 | NA | CCT6A | chaperonin containing TCP1, subunit 6A (zeta 1) | 908 | ENSG00000146731 |
| 8 | P49368 | NA | CCT3 | chaperonin containing TCP1, subunit 3 (gamma) | 7203 | ENSG00000163468 |
| 9 | Q99832 | NA | CCT7 | chaperonin containing TCP1, subunit 7 (eta) | 10574 | ENSG00000135624 |
| 10 | P62906 | NA | RPL10A | ribosomal protein L10a | 4736 | ENSG00000198755 |
| 11 | Q15056 | NA | EIF4H | eukaryotic translation initiation factor 4H | 7458 | ENSG00000106682 |
| 12 | P50990 | NA | CCT8 | chaperonin containing TCP1, subunit 8 (theta) | 10694 | ENSG00000156261 |
| 13 | P68104 | NA | EEF1A1 | eukaryotic translation elongation factor 1 alpha 1 | 1915 | ENSG00000156508 |
| 14 | P50991 | NA | CCT4 | chaperonin containing TCP1, subunit 4 (delta) | 10575 | ENSG00000115484 |

  
  

| **Database:Pathway Commons pathway      &nbspName:Gene Expression      &nbspID:DB\_ID:531** | | | | | | |
| --- | --- | --- | --- | --- | --- | --- |
| C=379; O=9; E=1.18; R=7.65; rawP=2.45e-06; adjP=1.99e-05 | | | | | | |
| Index | UserID | Value | Gene Symbol | Gene Name | EntrezGene | Ensembl |
| 1 | P07814 | NA | EPRS | glutamyl-prolyl-tRNA synthetase | 2058 | ENSG00000136628 |
| 2 | P13639 | NA | EEF2 | eukaryotic translation elongation factor 2 | 1938 | ENSG00000167658 |
| 3 | P41091 | NA | EIF2S3 | eukaryotic translation initiation factor 2, subunit 3 gamma, 52kDa | 1968 | ENSG00000130741 |
| 4 | P12081 | NA | HARS | histidyl-tRNA synthetase | 3035 | ENSG00000170445 |
| 5 | P67809 | NA | YBX1 | Y box binding protein 1 | 4904 | ENSG00000065978 |
| 6 | P56192 | NA | MARS | methionyl-tRNA synthetase | 4141 | ENSG00000166986 |
| 7 | P62906 | NA | RPL10A | ribosomal protein L10a | 4736 | ENSG00000198755 |
| 8 | Q15056 | NA | EIF4H | eukaryotic translation initiation factor 4H | 7458 | ENSG00000106682 |
| 9 | P68104 | NA | EEF1A1 | eukaryotic translation elongation factor 1 alpha 1 | 1915 | ENSG00000156508 |

  
  

| **Database:Pathway Commons pathway      &nbspName:Translation      &nbspID:DB\_ID:722** | | | | | | |
| --- | --- | --- | --- | --- | --- | --- |
| C=118; O=5; E=0.37; R=13.64; rawP=3.29e-05; adjP=0.0002 | | | | | | |
| Index | UserID | Value | Gene Symbol | Gene Name | EntrezGene | Ensembl |
| 1 | P62906 | NA | RPL10A | ribosomal protein L10a | 4736 | ENSG00000198755 |
| 2 | P13639 | NA | EEF2 | eukaryotic translation elongation factor 2 | 1938 | ENSG00000167658 |
| 3 | P41091 | NA | EIF2S3 | eukaryotic translation initiation factor 2, subunit 3 gamma, 52kDa | 1968 | ENSG00000130741 |
| 4 | Q15056 | NA | EIF4H | eukaryotic translation initiation factor 4H | 7458 | ENSG00000106682 |
| 5 | P68104 | NA | EEF1A1 | eukaryotic translation elongation factor 1 alpha 1 | 1915 | ENSG00000156508 |

  
  

| **Database:Pathway Commons pathway      &nbspName:Smooth Muscle Contraction      &nbspID:DB\_ID:144** | | | | | | |
| --- | --- | --- | --- | --- | --- | --- |
| C=24; O=3; E=0.07; R=40.24; rawP=5.52e-05; adjP=0.0003 | | | | | | |
| Index | UserID | Value | Gene Symbol | Gene Name | EntrezGene | Ensembl |
| 1 | P07951 | NA | TPM2 | tropomyosin 2 (beta) | 7169 | ENSG00000198467 |
| 2 | P18206 | NA | VCL | vinculin | 7414 | ENSG00000035403 |
| 3 | P67936 | NA | TPM4 | tropomyosin 4 | 7171 | ENSG00000167460 |

  
  

| **Database:Pathway Commons pathway      &nbspName:Cytosolic tRNA aminoacylation      &nbspID:DB\_ID:520** | | | | | | |
| --- | --- | --- | --- | --- | --- | --- |
| C=24; O=3; E=0.07; R=40.24; rawP=5.52e-05; adjP=0.0003 | | | | | | |
| Index | UserID | Value | Gene Symbol | Gene Name | EntrezGene | Ensembl |
| 1 | P07814 | NA | EPRS | glutamyl-prolyl-tRNA synthetase | 2058 | ENSG00000136628 |
| 2 | P56192 | NA | MARS | methionyl-tRNA synthetase | 4141 | ENSG00000166986 |
| 3 | P12081 | NA | HARS | histidyl-tRNA synthetase | 3035 | ENSG00000170445 |

  
  

| **Database:Pathway Commons pathway      &nbspName:tRNA Aminoacylation      &nbspID:DB\_ID:519** | | | | | | |
| --- | --- | --- | --- | --- | --- | --- |
| C=42; O=3; E=0.13; R=23.00; rawP=0.0003; adjP=0.0017 | | | | | | |
| Index | UserID | Value | Gene Symbol | Gene Name | EntrezGene | Ensembl |
| 1 | P07814 | NA | EPRS | glutamyl-prolyl-tRNA synthetase | 2058 | ENSG00000136628 |
| 2 | P56192 | NA | MARS | methionyl-tRNA synthetase | 4141 | ENSG00000166986 |
| 3 | P12081 | NA | HARS | histidyl-tRNA synthetase | 3035 | ENSG00000170445 |

  
  

| **Database:Pathway Commons pathway      &nbspName:Muscle contraction      &nbspID:DB\_ID:146** | | | | | | |
| --- | --- | --- | --- | --- | --- | --- |
| C=50; O=3; E=0.16; R=19.32; rawP=0.0005; adjP=0.0026 | | | | | | |
| Index | UserID | Value | Gene Symbol | Gene Name | EntrezGene | Ensembl |
| 1 | P07951 | NA | TPM2 | tropomyosin 2 (beta) | 7169 | ENSG00000198467 |
| 2 | P18206 | NA | VCL | vinculin | 7414 | ENSG00000035403 |
| 3 | P67936 | NA | TPM4 | tropomyosin 4 | 7171 | ENSG00000167460 |

  
  

| **Database:Pathway Commons pathway      &nbspName:Metabolism of nucleotides      &nbspID:DB\_ID:639** | | | | | | |
| --- | --- | --- | --- | --- | --- | --- |
| C=58; O=3; E=0.18; R=16.65; rawP=0.0008; adjP=0.0039 | | | | | | |
| Index | UserID | Value | Gene Symbol | Gene Name | EntrezGene | Ensembl |
| 1 | Q16881 | NA | TXNRD1 | thioredoxin reductase 1 | 7296 | ENSG00000198431 |
| 2 | P22102 | NA | GART | phosphoribosylglycinamide formyltransferase, phosphoribosylglycinamide synthetase, phosphoribosylaminoimidazole synthetase | 2618 | ENSG00000159131 |
| 3 | P00491 | NA | PNP | purine nucleoside phosphorylase | 4860 | ENSG00000198805 |

  
  

| **Database:Pathway Commons pathway      &nbspName:Peptide chain elongation      &nbspID:DB\_ID:714** | | | | | | |
| --- | --- | --- | --- | --- | --- | --- |
| C=83; O=3; E=0.26; R=11.64; rawP=0.0022; adjP=0.0100 | | | | | | |
| Index | UserID | Value | Gene Symbol | Gene Name | EntrezGene | Ensembl |
| 1 | P62906 | NA | RPL10A | ribosomal protein L10a | 4736 | ENSG00000198755 |
| 2 | P13639 | NA | EEF2 | eukaryotic translation elongation factor 2 | 1938 | ENSG00000167658 |
| 3 | P68104 | NA | EEF1A1 | eukaryotic translation elongation factor 1 alpha 1 | 1915 | ENSG00000156508 |

  
  

| **Database:Pathway Commons pathway      &nbspName:Eukaryotic Translation Elongation      &nbspID:DB\_ID:713** | | | | | | |
| --- | --- | --- | --- | --- | --- | --- |
| C=86; O=3; E=0.27; R=11.23; rawP=0.0024; adjP=0.0103 | | | | | | |
| Index | UserID | Value | Gene Symbol | Gene Name | EntrezGene | Ensembl |
| 1 | P62906 | NA | RPL10A | ribosomal protein L10a | 4736 | ENSG00000198755 |
| 2 | P13639 | NA | EEF2 | eukaryotic translation elongation factor 2 | 1938 | ENSG00000167658 |
| 3 | P68104 | NA | EEF1A1 | eukaryotic translation elongation factor 1 alpha 1 | 1915 | ENSG00000156508 |

  
  

| **Database:Pathway Commons pathway      &nbspName:Signaling events mediated by VEGFR1 and VEGFR2      &nbspID:DB\_ID:1516** | | | | | | |
| --- | --- | --- | --- | --- | --- | --- |
| C=1296; O=10; E=4.03; R=2.48; rawP=0.0063; adjP=0.0185 | | | | | | |
| Index | UserID | Value | Gene Symbol | Gene Name | EntrezGene | Ensembl |
| 1 | P34932 | NA | HSPA4 | heat shock 70kDa protein 4 | 3308 | ENSG00000170606 |
| 2 | P02795 | NA | MT2A | metallothionein 2A | 4502 | ENSG00000125148 |
| 3 | P08195 | NA | SLC3A2 | solute carrier family 3 (activators of dibasic and neutral amino acid transport), member 2 | 6520 | ENSG00000168003 |
| 4 | Q9NZM1 | NA | MYOF | myoferlin | 26509 | ENSG00000138119 |
| 5 | P13639 | NA | EEF2 | eukaryotic translation elongation factor 2 | 1938 | ENSG00000167658 |
| 6 | P52292 | NA | KPNA2 | karyopherin alpha 2 (RAG cohort 1, importin alpha 1) | 3838 | ENSG00000182481 |
| 7 | P18206 | NA | VCL | vinculin | 7414 | ENSG00000035403 |
| 8 | O60610 | NA | DIAPH1 | diaphanous homolog 1 (Drosophila) | 1729 | ENSG00000131504 |
| 9 | P02786 | NA | TFRC | transferrin receptor (p90, CD71) | 7037 | ENSG00000072274 |
| 10 | Q06830 | NA | PRDX1 | peroxiredoxin 1 | 5052 | ENSG00000117450 |

  
  

| **Database:Pathway Commons pathway      &nbspName:Alpha9 beta1 integrin signaling events      &nbspID:DB\_ID:1578** | | | | | | |
| --- | --- | --- | --- | --- | --- | --- |
| C=1305; O=10; E=4.05; R=2.47; rawP=0.0066; adjP=0.0185 | | | | | | |
| Index | UserID | Value | Gene Symbol | Gene Name | EntrezGene | Ensembl |
| 1 | P34932 | NA | HSPA4 | heat shock 70kDa protein 4 | 3308 | ENSG00000170606 |
| 2 | P02795 | NA | MT2A | metallothionein 2A | 4502 | ENSG00000125148 |
| 3 | P08195 | NA | SLC3A2 | solute carrier family 3 (activators of dibasic and neutral amino acid transport), member 2 | 6520 | ENSG00000168003 |
| 4 | P13639 | NA | EEF2 | eukaryotic translation elongation factor 2 | 1938 | ENSG00000167658 |
| 5 | P52292 | NA | KPNA2 | karyopherin alpha 2 (RAG cohort 1, importin alpha 1) | 3838 | ENSG00000182481 |
| 6 | P21980 | NA | TGM2 | transglutaminase 2 (C polypeptide, protein-glutamine-gamma-glutamyltransferase) | 7052 | ENSG00000198959 |
| 7 | P18206 | NA | VCL | vinculin | 7414 | ENSG00000035403 |
| 8 | O60610 | NA | DIAPH1 | diaphanous homolog 1 (Drosophila) | 1729 | ENSG00000131504 |
| 9 | P02786 | NA | TFRC | transferrin receptor (p90, CD71) | 7037 | ENSG00000072274 |
| 10 | Q06830 | NA | PRDX1 | peroxiredoxin 1 | 5052 | ENSG00000117450 |

  
  

| **Database:Pathway Commons pathway      &nbspName:VEGF and VEGFR signaling network      &nbspID:DB\_ID:1575** | | | | | | |
| --- | --- | --- | --- | --- | --- | --- |
| C=1304; O=10; E=4.05; R=2.47; rawP=0.0066; adjP=0.0185 | | | | | | |
| Index | UserID | Value | Gene Symbol | Gene Name | EntrezGene | Ensembl |
| 1 | P34932 | NA | HSPA4 | heat shock 70kDa protein 4 | 3308 | ENSG00000170606 |
| 2 | P02795 | NA | MT2A | metallothionein 2A | 4502 | ENSG00000125148 |
| 3 | P08195 | NA | SLC3A2 | solute carrier family 3 (activators of dibasic and neutral amino acid transport), member 2 | 6520 | ENSG00000168003 |
| 4 | Q9NZM1 | NA | MYOF | myoferlin | 26509 | ENSG00000138119 |
| 5 | P13639 | NA | EEF2 | eukaryotic translation elongation factor 2 | 1938 | ENSG00000167658 |
| 6 | P52292 | NA | KPNA2 | karyopherin alpha 2 (RAG cohort 1, importin alpha 1) | 3838 | ENSG00000182481 |
| 7 | P18206 | NA | VCL | vinculin | 7414 | ENSG00000035403 |
| 8 | O60610 | NA | DIAPH1 | diaphanous homolog 1 (Drosophila) | 1729 | ENSG00000131504 |
| 9 | P02786 | NA | TFRC | transferrin receptor (p90, CD71) | 7037 | ENSG00000072274 |
| 10 | Q06830 | NA | PRDX1 | peroxiredoxin 1 | 5052 | ENSG00000117450 |

  
  

| **Database:Pathway Commons pathway      &nbspName:TRAIL signaling pathway      &nbspID:DB\_ID:1480** | | | | | | |
| --- | --- | --- | --- | --- | --- | --- |
| C=1328; O=10; E=4.12; R=2.42; rawP=0.0075; adjP=0.0203 | | | | | | |
| Index | UserID | Value | Gene Symbol | Gene Name | EntrezGene | Ensembl |
| 1 | P34932 | NA | HSPA4 | heat shock 70kDa protein 4 | 3308 | ENSG00000170606 |
| 2 | P02795 | NA | MT2A | metallothionein 2A | 4502 | ENSG00000125148 |
| 3 | P08195 | NA | SLC3A2 | solute carrier family 3 (activators of dibasic and neutral amino acid transport), member 2 | 6520 | ENSG00000168003 |
| 4 | P13639 | NA | EEF2 | eukaryotic translation elongation factor 2 | 1938 | ENSG00000167658 |
| 5 | P52292 | NA | KPNA2 | karyopherin alpha 2 (RAG cohort 1, importin alpha 1) | 3838 | ENSG00000182481 |
| 6 | P18206 | NA | VCL | vinculin | 7414 | ENSG00000035403 |
| 7 | O60610 | NA | DIAPH1 | diaphanous homolog 1 (Drosophila) | 1729 | ENSG00000131504 |
| 8 | P02786 | NA | TFRC | transferrin receptor (p90, CD71) | 7037 | ENSG00000072274 |
| 9 | Q06830 | NA | PRDX1 | peroxiredoxin 1 | 5052 | ENSG00000117450 |
| 10 | P11387 | NA | TOP1 | topoisomerase (DNA) I | 7150 | ENSG00000198900 |

  
  

| **Database:Pathway Commons pathway      &nbspName:Beta1 integrin cell surface interactions      &nbspID:DB\_ID:1517** | | | | | | |
| --- | --- | --- | --- | --- | --- | --- |
| C=1351; O=10; E=4.20; R=2.38; rawP=0.0084; adjP=0.0219 | | | | | | |
| Index | UserID | Value | Gene Symbol | Gene Name | EntrezGene | Ensembl |
| 1 | P34932 | NA | HSPA4 | heat shock 70kDa protein 4 | 3308 | ENSG00000170606 |
| 2 | P02795 | NA | MT2A | metallothionein 2A | 4502 | ENSG00000125148 |
| 3 | P08195 | NA | SLC3A2 | solute carrier family 3 (activators of dibasic and neutral amino acid transport), member 2 | 6520 | ENSG00000168003 |
| 4 | P13639 | NA | EEF2 | eukaryotic translation elongation factor 2 | 1938 | ENSG00000167658 |
| 5 | P52292 | NA | KPNA2 | karyopherin alpha 2 (RAG cohort 1, importin alpha 1) | 3838 | ENSG00000182481 |
| 6 | P21980 | NA | TGM2 | transglutaminase 2 (C polypeptide, protein-glutamine-gamma-glutamyltransferase) | 7052 | ENSG00000198959 |
| 7 | P18206 | NA | VCL | vinculin | 7414 | ENSG00000035403 |
| 8 | O60610 | NA | DIAPH1 | diaphanous homolog 1 (Drosophila) | 1729 | ENSG00000131504 |
| 9 | P02786 | NA | TFRC | transferrin receptor (p90, CD71) | 7037 | ENSG00000072274 |
| 10 | Q06830 | NA | PRDX1 | peroxiredoxin 1 | 5052 | ENSG00000117450 |

  
  

| **Database:Pathway Commons pathway      &nbspName:Regulation of CDC42 activity      &nbspID:DB\_ID:1456** | | | | | | |
| --- | --- | --- | --- | --- | --- | --- |
| C=770; O=7; E=2.39; R=2.93; rawP=0.0097; adjP=0.0222 | | | | | | |
| Index | UserID | Value | Gene Symbol | Gene Name | EntrezGene | Ensembl |
| 1 | P52292 | NA | KPNA2 | karyopherin alpha 2 (RAG cohort 1, importin alpha 1) | 3838 | ENSG00000182481 |
| 2 | P02795 | NA | MT2A | metallothionein 2A | 4502 | ENSG00000125148 |
| 3 | P08195 | NA | SLC3A2 | solute carrier family 3 (activators of dibasic and neutral amino acid transport), member 2 | 6520 | ENSG00000168003 |
| 4 | P18206 | NA | VCL | vinculin | 7414 | ENSG00000035403 |
| 5 | O60610 | NA | DIAPH1 | diaphanous homolog 1 (Drosophila) | 1729 | ENSG00000131504 |
| 6 | Q06830 | NA | PRDX1 | peroxiredoxin 1 | 5052 | ENSG00000117450 |
| 7 | P02786 | NA | TFRC | transferrin receptor (p90, CD71) | 7037 | ENSG00000072274 |

  
  

| **Database:Pathway Commons pathway      &nbspName:Signaling events mediated by Hepatocyte Growth Factor Receptor (c-Met)      &nbspID:DB\_ID:1491** | | | | | | |
| --- | --- | --- | --- | --- | --- | --- |
| C=1293; O=9; E=4.02; R=2.24; rawP=0.0179; adjP=0.0222 | | | | | | |
| Index | UserID | Value | Gene Symbol | Gene Name | EntrezGene | Ensembl |
| 1 | P34932 | NA | HSPA4 | heat shock 70kDa protein 4 | 3308 | ENSG00000170606 |
| 2 | P02795 | NA | MT2A | metallothionein 2A | 4502 | ENSG00000125148 |
| 3 | P08195 | NA | SLC3A2 | solute carrier family 3 (activators of dibasic and neutral amino acid transport), member 2 | 6520 | ENSG00000168003 |
| 4 | P13639 | NA | EEF2 | eukaryotic translation elongation factor 2 | 1938 | ENSG00000167658 |
| 5 | P52292 | NA | KPNA2 | karyopherin alpha 2 (RAG cohort 1, importin alpha 1) | 3838 | ENSG00000182481 |
| 6 | P18206 | NA | VCL | vinculin | 7414 | ENSG00000035403 |
| 7 | O60610 | NA | DIAPH1 | diaphanous homolog 1 (Drosophila) | 1729 | ENSG00000131504 |
| 8 | P02786 | NA | TFRC | transferrin receptor (p90, CD71) | 7037 | ENSG00000072274 |
| 9 | Q06830 | NA | PRDX1 | peroxiredoxin 1 | 5052 | ENSG00000117450 |

  
  

| **Database:Pathway Commons pathway      &nbspName:S1P1 pathway      &nbspID:DB\_ID:1594** | | | | | | |
| --- | --- | --- | --- | --- | --- | --- |
| C=1288; O=9; E=4.00; R=2.25; rawP=0.0175; adjP=0.0222 | | | | | | |
| Index | UserID | Value | Gene Symbol | Gene Name | EntrezGene | Ensembl |
| 1 | P34932 | NA | HSPA4 | heat shock 70kDa protein 4 | 3308 | ENSG00000170606 |
| 2 | P02795 | NA | MT2A | metallothionein 2A | 4502 | ENSG00000125148 |
| 3 | P08195 | NA | SLC3A2 | solute carrier family 3 (activators of dibasic and neutral amino acid transport), member 2 | 6520 | ENSG00000168003 |
| 4 | P13639 | NA | EEF2 | eukaryotic translation elongation factor 2 | 1938 | ENSG00000167658 |
| 5 | P52292 | NA | KPNA2 | karyopherin alpha 2 (RAG cohort 1, importin alpha 1) | 3838 | ENSG00000182481 |
| 6 | P18206 | NA | VCL | vinculin | 7414 | ENSG00000035403 |
| 7 | O60610 | NA | DIAPH1 | diaphanous homolog 1 (Drosophila) | 1729 | ENSG00000131504 |
| 8 | P02786 | NA | TFRC | transferrin receptor (p90, CD71) | 7037 | ENSG00000072274 |
| 9 | Q06830 | NA | PRDX1 | peroxiredoxin 1 | 5052 | ENSG00000117450 |

  
  

| **Database:Pathway Commons pathway      &nbspName:Arf6 downstream pathway      &nbspID:DB\_ID:1585** | | | | | | |
| --- | --- | --- | --- | --- | --- | --- |
| C=1288; O=9; E=4.00; R=2.25; rawP=0.0175; adjP=0.0222 | | | | | | |
| Index | UserID | Value | Gene Symbol | Gene Name | EntrezGene | Ensembl |
| 1 | P34932 | NA | HSPA4 | heat shock 70kDa protein 4 | 3308 | ENSG00000170606 |
| 2 | P02795 | NA | MT2A | metallothionein 2A | 4502 | ENSG00000125148 |
| 3 | P08195 | NA | SLC3A2 | solute carrier family 3 (activators of dibasic and neutral amino acid transport), member 2 | 6520 | ENSG00000168003 |
| 4 | P13639 | NA | EEF2 | eukaryotic translation elongation factor 2 | 1938 | ENSG00000167658 |
| 5 | P52292 | NA | KPNA2 | karyopherin alpha 2 (RAG cohort 1, importin alpha 1) | 3838 | ENSG00000182481 |
| 6 | P18206 | NA | VCL | vinculin | 7414 | ENSG00000035403 |
| 7 | O60610 | NA | DIAPH1 | diaphanous homolog 1 (Drosophila) | 1729 | ENSG00000131504 |
| 8 | P02786 | NA | TFRC | transferrin receptor (p90, CD71) | 7037 | ENSG00000072274 |
| 9 | Q06830 | NA | PRDX1 | peroxiredoxin 1 | 5052 | ENSG00000117450 |

  
  

| **Database:Pathway Commons pathway      &nbspName:Insulin Pathway      &nbspID:DB\_ID:1466** | | | | | | |
| --- | --- | --- | --- | --- | --- | --- |
| C=1288; O=9; E=4.00; R=2.25; rawP=0.0175; adjP=0.0222 | | | | | | |
| Index | UserID | Value | Gene Symbol | Gene Name | EntrezGene | Ensembl |
| 1 | P34932 | NA | HSPA4 | heat shock 70kDa protein 4 | 3308 | ENSG00000170606 |
| 2 | P02795 | NA | MT2A | metallothionein 2A | 4502 | ENSG00000125148 |
| 3 | P08195 | NA | SLC3A2 | solute carrier family 3 (activators of dibasic and neutral amino acid transport), member 2 | 6520 | ENSG00000168003 |
| 4 | P13639 | NA | EEF2 | eukaryotic translation elongation factor 2 | 1938 | ENSG00000167658 |
| 5 | P52292 | NA | KPNA2 | karyopherin alpha 2 (RAG cohort 1, importin alpha 1) | 3838 | ENSG00000182481 |
| 6 | P18206 | NA | VCL | vinculin | 7414 | ENSG00000035403 |
| 7 | O60610 | NA | DIAPH1 | diaphanous homolog 1 (Drosophila) | 1729 | ENSG00000131504 |
| 8 | P02786 | NA | TFRC | transferrin receptor (p90, CD71) | 7037 | ENSG00000072274 |
| 9 | Q06830 | NA | PRDX1 | peroxiredoxin 1 | 5052 | ENSG00000117450 |

  
  

| **Database:Pathway Commons pathway      &nbspName:ErbB1 downstream signaling      &nbspID:DB\_ID:1602** | | | | | | |
| --- | --- | --- | --- | --- | --- | --- |
| C=1288; O=9; E=4.00; R=2.25; rawP=0.0175; adjP=0.0222 | | | | | | |
| Index | UserID | Value | Gene Symbol | Gene Name | EntrezGene | Ensembl |
| 1 | P34932 | NA | HSPA4 | heat shock 70kDa protein 4 | 3308 | ENSG00000170606 |
| 2 | P02795 | NA | MT2A | metallothionein 2A | 4502 | ENSG00000125148 |
| 3 | P08195 | NA | SLC3A2 | solute carrier family 3 (activators of dibasic and neutral amino acid transport), member 2 | 6520 | ENSG00000168003 |
| 4 | P13639 | NA | EEF2 | eukaryotic translation elongation factor 2 | 1938 | ENSG00000167658 |
| 5 | P52292 | NA | KPNA2 | karyopherin alpha 2 (RAG cohort 1, importin alpha 1) | 3838 | ENSG00000182481 |
| 6 | P18206 | NA | VCL | vinculin | 7414 | ENSG00000035403 |
| 7 | O60610 | NA | DIAPH1 | diaphanous homolog 1 (Drosophila) | 1729 | ENSG00000131504 |
| 8 | P02786 | NA | TFRC | transferrin receptor (p90, CD71) | 7037 | ENSG00000072274 |
| 9 | Q06830 | NA | PRDX1 | peroxiredoxin 1 | 5052 | ENSG00000117450 |

  
  

| **Database:Pathway Commons pathway      &nbspName:Urokinase-type plasminogen activator (uPA) and uPAR-mediated signaling      &nbspID:DB\_ID:1519** | | | | | | |
| --- | --- | --- | --- | --- | --- | --- |
| C=1288; O=9; E=4.00; R=2.25; rawP=0.0175; adjP=0.0222 | | | | | | |
| Index | UserID | Value | Gene Symbol | Gene Name | EntrezGene | Ensembl |
| 1 | P34932 | NA | HSPA4 | heat shock 70kDa protein 4 | 3308 | ENSG00000170606 |
| 2 | P02795 | NA | MT2A | metallothionein 2A | 4502 | ENSG00000125148 |
| 3 | P08195 | NA | SLC3A2 | solute carrier family 3 (activators of dibasic and neutral amino acid transport), member 2 | 6520 | ENSG00000168003 |
| 4 | P13639 | NA | EEF2 | eukaryotic translation elongation factor 2 | 1938 | ENSG00000167658 |
| 5 | P52292 | NA | KPNA2 | karyopherin alpha 2 (RAG cohort 1, importin alpha 1) | 3838 | ENSG00000182481 |
| 6 | P18206 | NA | VCL | vinculin | 7414 | ENSG00000035403 |
| 7 | O60610 | NA | DIAPH1 | diaphanous homolog 1 (Drosophila) | 1729 | ENSG00000131504 |
| 8 | P02786 | NA | TFRC | transferrin receptor (p90, CD71) | 7037 | ENSG00000072274 |
| 9 | Q06830 | NA | PRDX1 | peroxiredoxin 1 | 5052 | ENSG00000117450 |

  
  

| **Database:Pathway Commons pathway      &nbspName:EGFR-dependent Endothelin signaling events      &nbspID:DB\_ID:1603** | | | | | | |
| --- | --- | --- | --- | --- | --- | --- |
| C=1289; O=9; E=4.00; R=2.25; rawP=0.0175; adjP=0.0222 | | | | | | |
| Index | UserID | Value | Gene Symbol | Gene Name | EntrezGene | Ensembl |
| 1 | P34932 | NA | HSPA4 | heat shock 70kDa protein 4 | 3308 | ENSG00000170606 |
| 2 | P02795 | NA | MT2A | metallothionein 2A | 4502 | ENSG00000125148 |
| 3 | P08195 | NA | SLC3A2 | solute carrier family 3 (activators of dibasic and neutral amino acid transport), member 2 | 6520 | ENSG00000168003 |
| 4 | P13639 | NA | EEF2 | eukaryotic translation elongation factor 2 | 1938 | ENSG00000167658 |
| 5 | P52292 | NA | KPNA2 | karyopherin alpha 2 (RAG cohort 1, importin alpha 1) | 3838 | ENSG00000182481 |
| 6 | P18206 | NA | VCL | vinculin | 7414 | ENSG00000035403 |
| 7 | O60610 | NA | DIAPH1 | diaphanous homolog 1 (Drosophila) | 1729 | ENSG00000131504 |
| 8 | P02786 | NA | TFRC | transferrin receptor (p90, CD71) | 7037 | ENSG00000072274 |
| 9 | Q06830 | NA | PRDX1 | peroxiredoxin 1 | 5052 | ENSG00000117450 |

  
  

| **Database:Pathway Commons pathway      &nbspName:EGF receptor (ErbB1) signaling pathway      &nbspID:DB\_ID:1550** | | | | | | |
| --- | --- | --- | --- | --- | --- | --- |
| C=1288; O=9; E=4.00; R=2.25; rawP=0.0175; adjP=0.0222 | | | | | | |
| Index | UserID | Value | Gene Symbol | Gene Name | EntrezGene | Ensembl |
| 1 | P34932 | NA | HSPA4 | heat shock 70kDa protein 4 | 3308 | ENSG00000170606 |
| 2 | P02795 | NA | MT2A | metallothionein 2A | 4502 | ENSG00000125148 |
| 3 | P08195 | NA | SLC3A2 | solute carrier family 3 (activators of dibasic and neutral amino acid transport), member 2 | 6520 | ENSG00000168003 |
| 4 | P13639 | NA | EEF2 | eukaryotic translation elongation factor 2 | 1938 | ENSG00000167658 |
| 5 | P52292 | NA | KPNA2 | karyopherin alpha 2 (RAG cohort 1, importin alpha 1) | 3838 | ENSG00000182481 |
| 6 | P18206 | NA | VCL | vinculin | 7414 | ENSG00000035403 |
| 7 | O60610 | NA | DIAPH1 | diaphanous homolog 1 (Drosophila) | 1729 | ENSG00000131504 |
| 8 | P02786 | NA | TFRC | transferrin receptor (p90, CD71) | 7037 | ENSG00000072274 |
| 9 | Q06830 | NA | PRDX1 | peroxiredoxin 1 | 5052 | ENSG00000117450 |

  
  

| **Database:Pathway Commons pathway      &nbspName:Integrin-linked kinase signaling      &nbspID:DB\_ID:1546** | | | | | | |
| --- | --- | --- | --- | --- | --- | --- |
| C=656; O=6; E=2.04; R=2.94; rawP=0.0160; adjP=0.0222 | | | | | | |
| Index | UserID | Value | Gene Symbol | Gene Name | EntrezGene | Ensembl |
| 1 | P52292 | NA | KPNA2 | karyopherin alpha 2 (RAG cohort 1, importin alpha 1) | 3838 | ENSG00000182481 |
| 2 | P02795 | NA | MT2A | metallothionein 2A | 4502 | ENSG00000125148 |
| 3 | P08195 | NA | SLC3A2 | solute carrier family 3 (activators of dibasic and neutral amino acid transport), member 2 | 6520 | ENSG00000168003 |
| 4 | O60610 | NA | DIAPH1 | diaphanous homolog 1 (Drosophila) | 1729 | ENSG00000131504 |
| 5 | Q06830 | NA | PRDX1 | peroxiredoxin 1 | 5052 | ENSG00000117450 |
| 6 | P02786 | NA | TFRC | transferrin receptor (p90, CD71) | 7037 | ENSG00000072274 |

  
  

| **Database:Pathway Commons pathway      &nbspName:GMCSF-mediated signaling events      &nbspID:DB\_ID:1461** | | | | | | |
| --- | --- | --- | --- | --- | --- | --- |
| C=1292; O=9; E=4.01; R=2.24; rawP=0.0178; adjP=0.0222 | | | | | | |
| Index | UserID | Value | Gene Symbol | Gene Name | EntrezGene | Ensembl |
| 1 | P34932 | NA | HSPA4 | heat shock 70kDa protein 4 | 3308 | ENSG00000170606 |
| 2 | P02795 | NA | MT2A | metallothionein 2A | 4502 | ENSG00000125148 |
| 3 | P08195 | NA | SLC3A2 | solute carrier family 3 (activators of dibasic and neutral amino acid transport), member 2 | 6520 | ENSG00000168003 |
| 4 | P13639 | NA | EEF2 | eukaryotic translation elongation factor 2 | 1938 | ENSG00000167658 |
| 5 | P52292 | NA | KPNA2 | karyopherin alpha 2 (RAG cohort 1, importin alpha 1) | 3838 | ENSG00000182481 |
| 6 | P18206 | NA | VCL | vinculin | 7414 | ENSG00000035403 |
| 7 | O60610 | NA | DIAPH1 | diaphanous homolog 1 (Drosophila) | 1729 | ENSG00000131504 |
| 8 | P02786 | NA | TFRC | transferrin receptor (p90, CD71) | 7037 | ENSG00000072274 |
| 9 | Q06830 | NA | PRDX1 | peroxiredoxin 1 | 5052 | ENSG00000117450 |

  
  

| **Database:Pathway Commons pathway      &nbspName:Arf6 trafficking events      &nbspID:DB\_ID:1615** | | | | | | |
| --- | --- | --- | --- | --- | --- | --- |
| C=1288; O=9; E=4.00; R=2.25; rawP=0.0175; adjP=0.0222 | | | | | | |
| Index | UserID | Value | Gene Symbol | Gene Name | EntrezGene | Ensembl |
| 1 | P34932 | NA | HSPA4 | heat shock 70kDa protein 4 | 3308 | ENSG00000170606 |
| 2 | P02795 | NA | MT2A | metallothionein 2A | 4502 | ENSG00000125148 |
| 3 | P08195 | NA | SLC3A2 | solute carrier family 3 (activators of dibasic and neutral amino acid transport), member 2 | 6520 | ENSG00000168003 |
| 4 | P13639 | NA | EEF2 | eukaryotic translation elongation factor 2 | 1938 | ENSG00000167658 |
| 5 | P52292 | NA | KPNA2 | karyopherin alpha 2 (RAG cohort 1, importin alpha 1) | 3838 | ENSG00000182481 |
| 6 | P18206 | NA | VCL | vinculin | 7414 | ENSG00000035403 |
| 7 | O60610 | NA | DIAPH1 | diaphanous homolog 1 (Drosophila) | 1729 | ENSG00000131504 |
| 8 | P02786 | NA | TFRC | transferrin receptor (p90, CD71) | 7037 | ENSG00000072274 |
| 9 | Q06830 | NA | PRDX1 | peroxiredoxin 1 | 5052 | ENSG00000117450 |

  
  

| **Database:Pathway Commons pathway      &nbspName:Sphingosine 1-phosphate (S1P) pathway      &nbspID:DB\_ID:1635** | | | | | | |
| --- | --- | --- | --- | --- | --- | --- |
| C=1311; O=9; E=4.07; R=2.21; rawP=0.0194; adjP=0.0222 | | | | | | |
| Index | UserID | Value | Gene Symbol | Gene Name | EntrezGene | Ensembl |
| 1 | P34932 | NA | HSPA4 | heat shock 70kDa protein 4 | 3308 | ENSG00000170606 |
| 2 | P02795 | NA | MT2A | metallothionein 2A | 4502 | ENSG00000125148 |
| 3 | P08195 | NA | SLC3A2 | solute carrier family 3 (activators of dibasic and neutral amino acid transport), member 2 | 6520 | ENSG00000168003 |
| 4 | P13639 | NA | EEF2 | eukaryotic translation elongation factor 2 | 1938 | ENSG00000167658 |
| 5 | P52292 | NA | KPNA2 | karyopherin alpha 2 (RAG cohort 1, importin alpha 1) | 3838 | ENSG00000182481 |
| 6 | P18206 | NA | VCL | vinculin | 7414 | ENSG00000035403 |
| 7 | O60610 | NA | DIAPH1 | diaphanous homolog 1 (Drosophila) | 1729 | ENSG00000131504 |
| 8 | P02786 | NA | TFRC | transferrin receptor (p90, CD71) | 7037 | ENSG00000072274 |
| 9 | Q06830 | NA | PRDX1 | peroxiredoxin 1 | 5052 | ENSG00000117450 |

  
  

| **Database:Pathway Commons pathway      &nbspName:Arf6 signaling events      &nbspID:DB\_ID:1554** | | | | | | |
| --- | --- | --- | --- | --- | --- | --- |
| C=1288; O=9; E=4.00; R=2.25; rawP=0.0175; adjP=0.0222 | | | | | | |
| Index | UserID | Value | Gene Symbol | Gene Name | EntrezGene | Ensembl |
| 1 | P34932 | NA | HSPA4 | heat shock 70kDa protein 4 | 3308 | ENSG00000170606 |
| 2 | P02795 | NA | MT2A | metallothionein 2A | 4502 | ENSG00000125148 |
| 3 | P08195 | NA | SLC3A2 | solute carrier family 3 (activators of dibasic and neutral amino acid transport), member 2 | 6520 | ENSG00000168003 |
| 4 | P13639 | NA | EEF2 | eukaryotic translation elongation factor 2 | 1938 | ENSG00000167658 |
| 5 | P52292 | NA | KPNA2 | karyopherin alpha 2 (RAG cohort 1, importin alpha 1) | 3838 | ENSG00000182481 |
| 6 | P18206 | NA | VCL | vinculin | 7414 | ENSG00000035403 |
| 7 | O60610 | NA | DIAPH1 | diaphanous homolog 1 (Drosophila) | 1729 | ENSG00000131504 |
| 8 | P02786 | NA | TFRC | transferrin receptor (p90, CD71) | 7037 | ENSG00000072274 |
| 9 | Q06830 | NA | PRDX1 | peroxiredoxin 1 | 5052 | ENSG00000117450 |

  
  

| **Database:Pathway Commons pathway      &nbspName:PDGFR-beta signaling pathway      &nbspID:DB\_ID:1540** | | | | | | |
| --- | --- | --- | --- | --- | --- | --- |
| C=1288; O=9; E=4.00; R=2.25; rawP=0.0175; adjP=0.0222 | | | | | | |
| Index | UserID | Value | Gene Symbol | Gene Name | EntrezGene | Ensembl |
| 1 | P34932 | NA | HSPA4 | heat shock 70kDa protein 4 | 3308 | ENSG00000170606 |
| 2 | P02795 | NA | MT2A | metallothionein 2A | 4502 | ENSG00000125148 |
| 3 | P08195 | NA | SLC3A2 | solute carrier family 3 (activators of dibasic and neutral amino acid transport), member 2 | 6520 | ENSG00000168003 |
| 4 | P13639 | NA | EEF2 | eukaryotic translation elongation factor 2 | 1938 | ENSG00000167658 |
| 5 | P52292 | NA | KPNA2 | karyopherin alpha 2 (RAG cohort 1, importin alpha 1) | 3838 | ENSG00000182481 |
| 6 | P18206 | NA | VCL | vinculin | 7414 | ENSG00000035403 |
| 7 | O60610 | NA | DIAPH1 | diaphanous homolog 1 (Drosophila) | 1729 | ENSG00000131504 |
| 8 | P02786 | NA | TFRC | transferrin receptor (p90, CD71) | 7037 | ENSG00000072274 |
| 9 | Q06830 | NA | PRDX1 | peroxiredoxin 1 | 5052 | ENSG00000117450 |

  
  

| **Database:Pathway Commons pathway      &nbspName:IFN-gamma pathway      &nbspID:DB\_ID:1529** | | | | | | |
| --- | --- | --- | --- | --- | --- | --- |
| C=1296; O=9; E=4.03; R=2.24; rawP=0.0181; adjP=0.0222 | | | | | | |
| Index | UserID | Value | Gene Symbol | Gene Name | EntrezGene | Ensembl |
| 1 | P34932 | NA | HSPA4 | heat shock 70kDa protein 4 | 3308 | ENSG00000170606 |
| 2 | P02795 | NA | MT2A | metallothionein 2A | 4502 | ENSG00000125148 |
| 3 | P08195 | NA | SLC3A2 | solute carrier family 3 (activators of dibasic and neutral amino acid transport), member 2 | 6520 | ENSG00000168003 |
| 4 | P13639 | NA | EEF2 | eukaryotic translation elongation factor 2 | 1938 | ENSG00000167658 |
| 5 | P52292 | NA | KPNA2 | karyopherin alpha 2 (RAG cohort 1, importin alpha 1) | 3838 | ENSG00000182481 |
| 6 | P18206 | NA | VCL | vinculin | 7414 | ENSG00000035403 |
| 7 | O60610 | NA | DIAPH1 | diaphanous homolog 1 (Drosophila) | 1729 | ENSG00000131504 |
| 8 | P02786 | NA | TFRC | transferrin receptor (p90, CD71) | 7037 | ENSG00000072274 |
| 9 | Q06830 | NA | PRDX1 | peroxiredoxin 1 | 5052 | ENSG00000117450 |

  
  

| **Database:Pathway Commons pathway      &nbspName:PAR1-mediated thrombin signaling events      &nbspID:DB\_ID:1531** | | | | | | |
| --- | --- | --- | --- | --- | --- | --- |
| C=1299; O=9; E=4.03; R=2.23; rawP=0.0184; adjP=0.0222 | | | | | | |
| Index | UserID | Value | Gene Symbol | Gene Name | EntrezGene | Ensembl |
| 1 | P34932 | NA | HSPA4 | heat shock 70kDa protein 4 | 3308 | ENSG00000170606 |
| 2 | P02795 | NA | MT2A | metallothionein 2A | 4502 | ENSG00000125148 |
| 3 | P08195 | NA | SLC3A2 | solute carrier family 3 (activators of dibasic and neutral amino acid transport), member 2 | 6520 | ENSG00000168003 |
| 4 | P13639 | NA | EEF2 | eukaryotic translation elongation factor 2 | 1938 | ENSG00000167658 |
| 5 | P52292 | NA | KPNA2 | karyopherin alpha 2 (RAG cohort 1, importin alpha 1) | 3838 | ENSG00000182481 |
| 6 | P18206 | NA | VCL | vinculin | 7414 | ENSG00000035403 |
| 7 | O60610 | NA | DIAPH1 | diaphanous homolog 1 (Drosophila) | 1729 | ENSG00000131504 |
| 8 | P02786 | NA | TFRC | transferrin receptor (p90, CD71) | 7037 | ENSG00000072274 |
| 9 | Q06830 | NA | PRDX1 | peroxiredoxin 1 | 5052 | ENSG00000117450 |

  
  

| **Database:Pathway Commons pathway      &nbspName:Glypican 1 network      &nbspID:DB\_ID:1492** | | | | | | |
| --- | --- | --- | --- | --- | --- | --- |
| C=1299; O=9; E=4.03; R=2.23; rawP=0.0184; adjP=0.0222 | | | | | | |
| Index | UserID | Value | Gene Symbol | Gene Name | EntrezGene | Ensembl |
| 1 | P34932 | NA | HSPA4 | heat shock 70kDa protein 4 | 3308 | ENSG00000170606 |
| 2 | P02795 | NA | MT2A | metallothionein 2A | 4502 | ENSG00000125148 |
| 3 | P08195 | NA | SLC3A2 | solute carrier family 3 (activators of dibasic and neutral amino acid transport), member 2 | 6520 | ENSG00000168003 |
| 4 | P13639 | NA | EEF2 | eukaryotic translation elongation factor 2 | 1938 | ENSG00000167658 |
| 5 | P52292 | NA | KPNA2 | karyopherin alpha 2 (RAG cohort 1, importin alpha 1) | 3838 | ENSG00000182481 |
| 6 | P18206 | NA | VCL | vinculin | 7414 | ENSG00000035403 |
| 7 | O60610 | NA | DIAPH1 | diaphanous homolog 1 (Drosophila) | 1729 | ENSG00000131504 |
| 8 | P02786 | NA | TFRC | transferrin receptor (p90, CD71) | 7037 | ENSG00000072274 |
| 9 | Q06830 | NA | PRDX1 | peroxiredoxin 1 | 5052 | ENSG00000117450 |

  
  

| **Database:Pathway Commons pathway      &nbspName:Nectin adhesion pathway      &nbspID:DB\_ID:1472** | | | | | | |
| --- | --- | --- | --- | --- | --- | --- |
| C=1295; O=9; E=4.02; R=2.24; rawP=0.0180; adjP=0.0222 | | | | | | |
| Index | UserID | Value | Gene Symbol | Gene Name | EntrezGene | Ensembl |
| 1 | P34932 | NA | HSPA4 | heat shock 70kDa protein 4 | 3308 | ENSG00000170606 |
| 2 | P02795 | NA | MT2A | metallothionein 2A | 4502 | ENSG00000125148 |
| 3 | P08195 | NA | SLC3A2 | solute carrier family 3 (activators of dibasic and neutral amino acid transport), member 2 | 6520 | ENSG00000168003 |
| 4 | P13639 | NA | EEF2 | eukaryotic translation elongation factor 2 | 1938 | ENSG00000167658 |
| 5 | P52292 | NA | KPNA2 | karyopherin alpha 2 (RAG cohort 1, importin alpha 1) | 3838 | ENSG00000182481 |
| 6 | P18206 | NA | VCL | vinculin | 7414 | ENSG00000035403 |
| 7 | O60610 | NA | DIAPH1 | diaphanous homolog 1 (Drosophila) | 1729 | ENSG00000131504 |
| 8 | P02786 | NA | TFRC | transferrin receptor (p90, CD71) | 7037 | ENSG00000072274 |
| 9 | Q06830 | NA | PRDX1 | peroxiredoxin 1 | 5052 | ENSG00000117450 |

  
  

| **Database:Pathway Commons pathway      &nbspName:IL5-mediated signaling events      &nbspID:DB\_ID:1627** | | | | | | |
| --- | --- | --- | --- | --- | --- | --- |
| C=1292; O=9; E=4.01; R=2.24; rawP=0.0178; adjP=0.0222 | | | | | | |
| Index | UserID | Value | Gene Symbol | Gene Name | EntrezGene | Ensembl |
| 1 | P34932 | NA | HSPA4 | heat shock 70kDa protein 4 | 3308 | ENSG00000170606 |
| 2 | P02795 | NA | MT2A | metallothionein 2A | 4502 | ENSG00000125148 |
| 3 | P08195 | NA | SLC3A2 | solute carrier family 3 (activators of dibasic and neutral amino acid transport), member 2 | 6520 | ENSG00000168003 |
| 4 | P13639 | NA | EEF2 | eukaryotic translation elongation factor 2 | 1938 | ENSG00000167658 |
| 5 | P52292 | NA | KPNA2 | karyopherin alpha 2 (RAG cohort 1, importin alpha 1) | 3838 | ENSG00000182481 |
| 6 | P18206 | NA | VCL | vinculin | 7414 | ENSG00000035403 |
| 7 | O60610 | NA | DIAPH1 | diaphanous homolog 1 (Drosophila) | 1729 | ENSG00000131504 |
| 8 | P02786 | NA | TFRC | transferrin receptor (p90, CD71) | 7037 | ENSG00000072274 |
| 9 | Q06830 | NA | PRDX1 | peroxiredoxin 1 | 5052 | ENSG00000117450 |

  
  

| **Database:Pathway Commons pathway      &nbspName:PDGF receptor signaling network      &nbspID:DB\_ID:1497** | | | | | | |
| --- | --- | --- | --- | --- | --- | --- |
| C=1293; O=9; E=4.02; R=2.24; rawP=0.0179; adjP=0.0222 | | | | | | |
| Index | UserID | Value | Gene Symbol | Gene Name | EntrezGene | Ensembl |
| 1 | P34932 | NA | HSPA4 | heat shock 70kDa protein 4 | 3308 | ENSG00000170606 |
| 2 | P02795 | NA | MT2A | metallothionein 2A | 4502 | ENSG00000125148 |
| 3 | P08195 | NA | SLC3A2 | solute carrier family 3 (activators of dibasic and neutral amino acid transport), member 2 | 6520 | ENSG00000168003 |
| 4 | P13639 | NA | EEF2 | eukaryotic translation elongation factor 2 | 1938 | ENSG00000167658 |
| 5 | P52292 | NA | KPNA2 | karyopherin alpha 2 (RAG cohort 1, importin alpha 1) | 3838 | ENSG00000182481 |
| 6 | P18206 | NA | VCL | vinculin | 7414 | ENSG00000035403 |
| 7 | O60610 | NA | DIAPH1 | diaphanous homolog 1 (Drosophila) | 1729 | ENSG00000131504 |
| 8 | P02786 | NA | TFRC | transferrin receptor (p90, CD71) | 7037 | ENSG00000072274 |
| 9 | Q06830 | NA | PRDX1 | peroxiredoxin 1 | 5052 | ENSG00000117450 |

  
  

| **Database:Pathway Commons pathway      &nbspName:CDC42 signaling events      &nbspID:DB\_ID:1488** | | | | | | |
| --- | --- | --- | --- | --- | --- | --- |
| C=757; O=7; E=2.35; R=2.98; rawP=0.0089; adjP=0.0222 | | | | | | |
| Index | UserID | Value | Gene Symbol | Gene Name | EntrezGene | Ensembl |
| 1 | P52292 | NA | KPNA2 | karyopherin alpha 2 (RAG cohort 1, importin alpha 1) | 3838 | ENSG00000182481 |
| 2 | P02795 | NA | MT2A | metallothionein 2A | 4502 | ENSG00000125148 |
| 3 | P08195 | NA | SLC3A2 | solute carrier family 3 (activators of dibasic and neutral amino acid transport), member 2 | 6520 | ENSG00000168003 |
| 4 | P18206 | NA | VCL | vinculin | 7414 | ENSG00000035403 |
| 5 | O60610 | NA | DIAPH1 | diaphanous homolog 1 (Drosophila) | 1729 | ENSG00000131504 |
| 6 | Q06830 | NA | PRDX1 | peroxiredoxin 1 | 5052 | ENSG00000117450 |
| 7 | P02786 | NA | TFRC | transferrin receptor (p90, CD71) | 7037 | ENSG00000072274 |

  
  

| **Database:Pathway Commons pathway      &nbspName:ErbB receptor signaling network      &nbspID:DB\_ID:1573** | | | | | | |
| --- | --- | --- | --- | --- | --- | --- |
| C=1312; O=9; E=4.08; R=2.21; rawP=0.0195; adjP=0.0222 | | | | | | |
| Index | UserID | Value | Gene Symbol | Gene Name | EntrezGene | Ensembl |
| 1 | P34932 | NA | HSPA4 | heat shock 70kDa protein 4 | 3308 | ENSG00000170606 |
| 2 | P02795 | NA | MT2A | metallothionein 2A | 4502 | ENSG00000125148 |
| 3 | P08195 | NA | SLC3A2 | solute carrier family 3 (activators of dibasic and neutral amino acid transport), member 2 | 6520 | ENSG00000168003 |
| 4 | P13639 | NA | EEF2 | eukaryotic translation elongation factor 2 | 1938 | ENSG00000167658 |
| 5 | P52292 | NA | KPNA2 | karyopherin alpha 2 (RAG cohort 1, importin alpha 1) | 3838 | ENSG00000182481 |
| 6 | P18206 | NA | VCL | vinculin | 7414 | ENSG00000035403 |
| 7 | O60610 | NA | DIAPH1 | diaphanous homolog 1 (Drosophila) | 1729 | ENSG00000131504 |
| 8 | P02786 | NA | TFRC | transferrin receptor (p90, CD71) | 7037 | ENSG00000072274 |
| 9 | Q06830 | NA | PRDX1 | peroxiredoxin 1 | 5052 | ENSG00000117450 |

  
  

| **Database:Pathway Commons pathway      &nbspName:Thrombin/protease-activated receptor (PAR) pathway      &nbspID:DB\_ID:1552** | | | | | | |
| --- | --- | --- | --- | --- | --- | --- |
| C=1300; O=9; E=4.04; R=2.23; rawP=0.0184; adjP=0.0222 | | | | | | |
| Index | UserID | Value | Gene Symbol | Gene Name | EntrezGene | Ensembl |
| 1 | P34932 | NA | HSPA4 | heat shock 70kDa protein 4 | 3308 | ENSG00000170606 |
| 2 | P02795 | NA | MT2A | metallothionein 2A | 4502 | ENSG00000125148 |
| 3 | P08195 | NA | SLC3A2 | solute carrier family 3 (activators of dibasic and neutral amino acid transport), member 2 | 6520 | ENSG00000168003 |
| 4 | P13639 | NA | EEF2 | eukaryotic translation elongation factor 2 | 1938 | ENSG00000167658 |
| 5 | P52292 | NA | KPNA2 | karyopherin alpha 2 (RAG cohort 1, importin alpha 1) | 3838 | ENSG00000182481 |
| 6 | P18206 | NA | VCL | vinculin | 7414 | ENSG00000035403 |
| 7 | O60610 | NA | DIAPH1 | diaphanous homolog 1 (Drosophila) | 1729 | ENSG00000131504 |
| 8 | P02786 | NA | TFRC | transferrin receptor (p90, CD71) | 7037 | ENSG00000072274 |
| 9 | Q06830 | NA | PRDX1 | peroxiredoxin 1 | 5052 | ENSG00000117450 |

  
  

| **Database:Pathway Commons pathway      &nbspName:Syndecan-1-mediated signaling events      &nbspID:DB\_ID:1454** | | | | | | |
| --- | --- | --- | --- | --- | --- | --- |
| C=1300; O=9; E=4.04; R=2.23; rawP=0.0184; adjP=0.0222 | | | | | | |
| Index | UserID | Value | Gene Symbol | Gene Name | EntrezGene | Ensembl |
| 1 | P34932 | NA | HSPA4 | heat shock 70kDa protein 4 | 3308 | ENSG00000170606 |
| 2 | P02795 | NA | MT2A | metallothionein 2A | 4502 | ENSG00000125148 |
| 3 | P08195 | NA | SLC3A2 | solute carrier family 3 (activators of dibasic and neutral amino acid transport), member 2 | 6520 | ENSG00000168003 |
| 4 | P13639 | NA | EEF2 | eukaryotic translation elongation factor 2 | 1938 | ENSG00000167658 |
| 5 | P52292 | NA | KPNA2 | karyopherin alpha 2 (RAG cohort 1, importin alpha 1) | 3838 | ENSG00000182481 |
| 6 | P18206 | NA | VCL | vinculin | 7414 | ENSG00000035403 |
| 7 | O60610 | NA | DIAPH1 | diaphanous homolog 1 (Drosophila) | 1729 | ENSG00000131504 |
| 8 | P02786 | NA | TFRC | transferrin receptor (p90, CD71) | 7037 | ENSG00000072274 |
| 9 | Q06830 | NA | PRDX1 | peroxiredoxin 1 | 5052 | ENSG00000117450 |

  
  

| **Database:Pathway Commons pathway      &nbspName:Internalization of ErbB1      &nbspID:DB\_ID:1509** | | | | | | |
| --- | --- | --- | --- | --- | --- | --- |
| C=1288; O=9; E=4.00; R=2.25; rawP=0.0175; adjP=0.0222 | | | | | | |
| Index | UserID | Value | Gene Symbol | Gene Name | EntrezGene | Ensembl |
| 1 | P34932 | NA | HSPA4 | heat shock 70kDa protein 4 | 3308 | ENSG00000170606 |
| 2 | P02795 | NA | MT2A | metallothionein 2A | 4502 | ENSG00000125148 |
| 3 | P08195 | NA | SLC3A2 | solute carrier family 3 (activators of dibasic and neutral amino acid transport), member 2 | 6520 | ENSG00000168003 |
| 4 | P13639 | NA | EEF2 | eukaryotic translation elongation factor 2 | 1938 | ENSG00000167658 |
| 5 | P52292 | NA | KPNA2 | karyopherin alpha 2 (RAG cohort 1, importin alpha 1) | 3838 | ENSG00000182481 |
| 6 | P18206 | NA | VCL | vinculin | 7414 | ENSG00000035403 |
| 7 | O60610 | NA | DIAPH1 | diaphanous homolog 1 (Drosophila) | 1729 | ENSG00000131504 |
| 8 | P02786 | NA | TFRC | transferrin receptor (p90, CD71) | 7037 | ENSG00000072274 |
| 9 | Q06830 | NA | PRDX1 | peroxiredoxin 1 | 5052 | ENSG00000117450 |

  
  

| **Database:Pathway Commons pathway      &nbspName:IGF1 pathway      &nbspID:DB\_ID:1482** | | | | | | |
| --- | --- | --- | --- | --- | --- | --- |
| C=1291; O=9; E=4.01; R=2.24; rawP=0.0177; adjP=0.0222 | | | | | | |
| Index | UserID | Value | Gene Symbol | Gene Name | EntrezGene | Ensembl |
| 1 | P34932 | NA | HSPA4 | heat shock 70kDa protein 4 | 3308 | ENSG00000170606 |
| 2 | P02795 | NA | MT2A | metallothionein 2A | 4502 | ENSG00000125148 |
| 3 | P08195 | NA | SLC3A2 | solute carrier family 3 (activators of dibasic and neutral amino acid transport), member 2 | 6520 | ENSG00000168003 |
| 4 | P13639 | NA | EEF2 | eukaryotic translation elongation factor 2 | 1938 | ENSG00000167658 |
| 5 | P52292 | NA | KPNA2 | karyopherin alpha 2 (RAG cohort 1, importin alpha 1) | 3838 | ENSG00000182481 |
| 6 | P18206 | NA | VCL | vinculin | 7414 | ENSG00000035403 |
| 7 | O60610 | NA | DIAPH1 | diaphanous homolog 1 (Drosophila) | 1729 | ENSG00000131504 |
| 8 | P02786 | NA | TFRC | transferrin receptor (p90, CD71) | 7037 | ENSG00000072274 |
| 9 | Q06830 | NA | PRDX1 | peroxiredoxin 1 | 5052 | ENSG00000117450 |

  
  

| **Database:Pathway Commons pathway      &nbspName:IL3-mediated signaling events      &nbspID:DB\_ID:1564** | | | | | | |
| --- | --- | --- | --- | --- | --- | --- |
| C=1295; O=9; E=4.02; R=2.24; rawP=0.0180; adjP=0.0222 | | | | | | |
| Index | UserID | Value | Gene Symbol | Gene Name | EntrezGene | Ensembl |
| 1 | P34932 | NA | HSPA4 | heat shock 70kDa protein 4 | 3308 | ENSG00000170606 |
| 2 | P02795 | NA | MT2A | metallothionein 2A | 4502 | ENSG00000125148 |
| 3 | P08195 | NA | SLC3A2 | solute carrier family 3 (activators of dibasic and neutral amino acid transport), member 2 | 6520 | ENSG00000168003 |
| 4 | P13639 | NA | EEF2 | eukaryotic translation elongation factor 2 | 1938 | ENSG00000167658 |
| 5 | P52292 | NA | KPNA2 | karyopherin alpha 2 (RAG cohort 1, importin alpha 1) | 3838 | ENSG00000182481 |
| 6 | P18206 | NA | VCL | vinculin | 7414 | ENSG00000035403 |
| 7 | O60610 | NA | DIAPH1 | diaphanous homolog 1 (Drosophila) | 1729 | ENSG00000131504 |
| 8 | P02786 | NA | TFRC | transferrin receptor (p90, CD71) | 7037 | ENSG00000072274 |
| 9 | Q06830 | NA | PRDX1 | peroxiredoxin 1 | 5052 | ENSG00000117450 |

  
  

| **Database:Pathway Commons pathway      &nbspName:Plasma membrane estrogen receptor signaling      &nbspID:DB\_ID:1556** | | | | | | |
| --- | --- | --- | --- | --- | --- | --- |
| C=1301; O=9; E=4.04; R=2.23; rawP=0.0185; adjP=0.0222 | | | | | | |
| Index | UserID | Value | Gene Symbol | Gene Name | EntrezGene | Ensembl |
| 1 | P34932 | NA | HSPA4 | heat shock 70kDa protein 4 | 3308 | ENSG00000170606 |
| 2 | P02795 | NA | MT2A | metallothionein 2A | 4502 | ENSG00000125148 |
| 3 | P08195 | NA | SLC3A2 | solute carrier family 3 (activators of dibasic and neutral amino acid transport), member 2 | 6520 | ENSG00000168003 |
| 4 | P13639 | NA | EEF2 | eukaryotic translation elongation factor 2 | 1938 | ENSG00000167658 |
| 5 | P52292 | NA | KPNA2 | karyopherin alpha 2 (RAG cohort 1, importin alpha 1) | 3838 | ENSG00000182481 |
| 6 | P18206 | NA | VCL | vinculin | 7414 | ENSG00000035403 |
| 7 | O60610 | NA | DIAPH1 | diaphanous homolog 1 (Drosophila) | 1729 | ENSG00000131504 |
| 8 | P02786 | NA | TFRC | transferrin receptor (p90, CD71) | 7037 | ENSG00000072274 |
| 9 | Q06830 | NA | PRDX1 | peroxiredoxin 1 | 5052 | ENSG00000117450 |

  
  

| **Database:Pathway Commons pathway      &nbspName:LKB1 signaling events      &nbspID:DB\_ID:1649** | | | | | | |
| --- | --- | --- | --- | --- | --- | --- |
| C=1308; O=9; E=4.06; R=2.22; rawP=0.0191; adjP=0.0222 | | | | | | |
| Index | UserID | Value | Gene Symbol | Gene Name | EntrezGene | Ensembl |
| 1 | P34932 | NA | HSPA4 | heat shock 70kDa protein 4 | 3308 | ENSG00000170606 |
| 2 | P02795 | NA | MT2A | metallothionein 2A | 4502 | ENSG00000125148 |
| 3 | P08195 | NA | SLC3A2 | solute carrier family 3 (activators of dibasic and neutral amino acid transport), member 2 | 6520 | ENSG00000168003 |
| 4 | P13639 | NA | EEF2 | eukaryotic translation elongation factor 2 | 1938 | ENSG00000167658 |
| 5 | P52292 | NA | KPNA2 | karyopherin alpha 2 (RAG cohort 1, importin alpha 1) | 3838 | ENSG00000182481 |
| 6 | P18206 | NA | VCL | vinculin | 7414 | ENSG00000035403 |
| 7 | O60610 | NA | DIAPH1 | diaphanous homolog 1 (Drosophila) | 1729 | ENSG00000131504 |
| 8 | P02786 | NA | TFRC | transferrin receptor (p90, CD71) | 7037 | ENSG00000072274 |
| 9 | Q06830 | NA | PRDX1 | peroxiredoxin 1 | 5052 | ENSG00000117450 |

  
  

| **Database:Pathway Commons pathway      &nbspName:mTOR signaling pathway      &nbspID:DB\_ID:1571** | | | | | | |
| --- | --- | --- | --- | --- | --- | --- |
| C=1288; O=9; E=4.00; R=2.25; rawP=0.0175; adjP=0.0222 | | | | | | |
| Index | UserID | Value | Gene Symbol | Gene Name | EntrezGene | Ensembl |
| 1 | P34932 | NA | HSPA4 | heat shock 70kDa protein 4 | 3308 | ENSG00000170606 |
| 2 | P02795 | NA | MT2A | metallothionein 2A | 4502 | ENSG00000125148 |
| 3 | P08195 | NA | SLC3A2 | solute carrier family 3 (activators of dibasic and neutral amino acid transport), member 2 | 6520 | ENSG00000168003 |
| 4 | P13639 | NA | EEF2 | eukaryotic translation elongation factor 2 | 1938 | ENSG00000167658 |
| 5 | P52292 | NA | KPNA2 | karyopherin alpha 2 (RAG cohort 1, importin alpha 1) | 3838 | ENSG00000182481 |
| 6 | P18206 | NA | VCL | vinculin | 7414 | ENSG00000035403 |
| 7 | O60610 | NA | DIAPH1 | diaphanous homolog 1 (Drosophila) | 1729 | ENSG00000131504 |
| 8 | P02786 | NA | TFRC | transferrin receptor (p90, CD71) | 7037 | ENSG00000072274 |
| 9 | Q06830 | NA | PRDX1 | peroxiredoxin 1 | 5052 | ENSG00000117450 |

  
  

| **Database:Pathway Commons pathway      &nbspName:Endothelins      &nbspID:DB\_ID:1619** | | | | | | |
| --- | --- | --- | --- | --- | --- | --- |
| C=1307; O=9; E=4.06; R=2.22; rawP=0.0190; adjP=0.0222 | | | | | | |
| Index | UserID | Value | Gene Symbol | Gene Name | EntrezGene | Ensembl |
| 1 | P34932 | NA | HSPA4 | heat shock 70kDa protein 4 | 3308 | ENSG00000170606 |
| 2 | P02795 | NA | MT2A | metallothionein 2A | 4502 | ENSG00000125148 |
| 3 | P08195 | NA | SLC3A2 | solute carrier family 3 (activators of dibasic and neutral amino acid transport), member 2 | 6520 | ENSG00000168003 |
| 4 | P13639 | NA | EEF2 | eukaryotic translation elongation factor 2 | 1938 | ENSG00000167658 |
| 5 | P52292 | NA | KPNA2 | karyopherin alpha 2 (RAG cohort 1, importin alpha 1) | 3838 | ENSG00000182481 |
| 6 | P18206 | NA | VCL | vinculin | 7414 | ENSG00000035403 |
| 7 | O60610 | NA | DIAPH1 | diaphanous homolog 1 (Drosophila) | 1729 | ENSG00000131504 |
| 8 | P02786 | NA | TFRC | transferrin receptor (p90, CD71) | 7037 | ENSG00000072274 |
| 9 | Q06830 | NA | PRDX1 | peroxiredoxin 1 | 5052 | ENSG00000117450 |

  
  

| **Database:Pathway Commons pathway      &nbspName:Class I PI3K signaling events mediated by Akt      &nbspID:DB\_ID:1648** | | | | | | |
| --- | --- | --- | --- | --- | --- | --- |
| C=1288; O=9; E=4.00; R=2.25; rawP=0.0175; adjP=0.0222 | | | | | | |
| Index | UserID | Value | Gene Symbol | Gene Name | EntrezGene | Ensembl |
| 1 | P34932 | NA | HSPA4 | heat shock 70kDa protein 4 | 3308 | ENSG00000170606 |
| 2 | P02795 | NA | MT2A | metallothionein 2A | 4502 | ENSG00000125148 |
| 3 | P08195 | NA | SLC3A2 | solute carrier family 3 (activators of dibasic and neutral amino acid transport), member 2 | 6520 | ENSG00000168003 |
| 4 | P13639 | NA | EEF2 | eukaryotic translation elongation factor 2 | 1938 | ENSG00000167658 |
| 5 | P52292 | NA | KPNA2 | karyopherin alpha 2 (RAG cohort 1, importin alpha 1) | 3838 | ENSG00000182481 |
| 6 | P18206 | NA | VCL | vinculin | 7414 | ENSG00000035403 |
| 7 | O60610 | NA | DIAPH1 | diaphanous homolog 1 (Drosophila) | 1729 | ENSG00000131504 |
| 8 | P02786 | NA | TFRC | transferrin receptor (p90, CD71) | 7037 | ENSG00000072274 |
| 9 | Q06830 | NA | PRDX1 | peroxiredoxin 1 | 5052 | ENSG00000117450 |

  
  

| **Database:Pathway Commons pathway      &nbspName:Signaling events mediated by focal adhesion kinase      &nbspID:DB\_ID:1574** | | | | | | |
| --- | --- | --- | --- | --- | --- | --- |
| C=1288; O=9; E=4.00; R=2.25; rawP=0.0175; adjP=0.0222 | | | | | | |
| Index | UserID | Value | Gene Symbol | Gene Name | EntrezGene | Ensembl |
| 1 | P34932 | NA | HSPA4 | heat shock 70kDa protein 4 | 3308 | ENSG00000170606 |
| 2 | P02795 | NA | MT2A | metallothionein 2A | 4502 | ENSG00000125148 |
| 3 | P08195 | NA | SLC3A2 | solute carrier family 3 (activators of dibasic and neutral amino acid transport), member 2 | 6520 | ENSG00000168003 |
| 4 | P13639 | NA | EEF2 | eukaryotic translation elongation factor 2 | 1938 | ENSG00000167658 |
| 5 | P52292 | NA | KPNA2 | karyopherin alpha 2 (RAG cohort 1, importin alpha 1) | 3838 | ENSG00000182481 |
| 6 | P18206 | NA | VCL | vinculin | 7414 | ENSG00000035403 |
| 7 | O60610 | NA | DIAPH1 | diaphanous homolog 1 (Drosophila) | 1729 | ENSG00000131504 |
| 8 | P02786 | NA | TFRC | transferrin receptor (p90, CD71) | 7037 | ENSG00000072274 |
| 9 | Q06830 | NA | PRDX1 | peroxiredoxin 1 | 5052 | ENSG00000117450 |

  
  

| **Database:Pathway Commons pathway      &nbspName:Class I PI3K signaling events      &nbspID:DB\_ID:1553** | | | | | | |
| --- | --- | --- | --- | --- | --- | --- |
| C=1288; O=9; E=4.00; R=2.25; rawP=0.0175; adjP=0.0222 | | | | | | |
| Index | UserID | Value | Gene Symbol | Gene Name | EntrezGene | Ensembl |
| 1 | P34932 | NA | HSPA4 | heat shock 70kDa protein 4 | 3308 | ENSG00000170606 |
| 2 | P02795 | NA | MT2A | metallothionein 2A | 4502 | ENSG00000125148 |
| 3 | P08195 | NA | SLC3A2 | solute carrier family 3 (activators of dibasic and neutral amino acid transport), member 2 | 6520 | ENSG00000168003 |
| 4 | P13639 | NA | EEF2 | eukaryotic translation elongation factor 2 | 1938 | ENSG00000167658 |
| 5 | P52292 | NA | KPNA2 | karyopherin alpha 2 (RAG cohort 1, importin alpha 1) | 3838 | ENSG00000182481 |
| 6 | P18206 | NA | VCL | vinculin | 7414 | ENSG00000035403 |
| 7 | O60610 | NA | DIAPH1 | diaphanous homolog 1 (Drosophila) | 1729 | ENSG00000131504 |
| 8 | P02786 | NA | TFRC | transferrin receptor (p90, CD71) | 7037 | ENSG00000072274 |
| 9 | Q06830 | NA | PRDX1 | peroxiredoxin 1 | 5052 | ENSG00000117450 |

  
  

| **Database:Pathway Commons pathway      &nbspName:Integrin family cell surface interactions      &nbspID:DB\_ID:1499** | | | | | | |
| --- | --- | --- | --- | --- | --- | --- |
| C=1378; O=10; E=4.28; R=2.34; rawP=0.0096; adjP=0.0222 | | | | | | |
| Index | UserID | Value | Gene Symbol | Gene Name | EntrezGene | Ensembl |
| 1 | P34932 | NA | HSPA4 | heat shock 70kDa protein 4 | 3308 | ENSG00000170606 |
| 2 | P02795 | NA | MT2A | metallothionein 2A | 4502 | ENSG00000125148 |
| 3 | P08195 | NA | SLC3A2 | solute carrier family 3 (activators of dibasic and neutral amino acid transport), member 2 | 6520 | ENSG00000168003 |
| 4 | P13639 | NA | EEF2 | eukaryotic translation elongation factor 2 | 1938 | ENSG00000167658 |
| 5 | P52292 | NA | KPNA2 | karyopherin alpha 2 (RAG cohort 1, importin alpha 1) | 3838 | ENSG00000182481 |
| 6 | P21980 | NA | TGM2 | transglutaminase 2 (C polypeptide, protein-glutamine-gamma-glutamyltransferase) | 7052 | ENSG00000198959 |
| 7 | P18206 | NA | VCL | vinculin | 7414 | ENSG00000035403 |
| 8 | O60610 | NA | DIAPH1 | diaphanous homolog 1 (Drosophila) | 1729 | ENSG00000131504 |
| 9 | P02786 | NA | TFRC | transferrin receptor (p90, CD71) | 7037 | ENSG00000072274 |
| 10 | Q06830 | NA | PRDX1 | peroxiredoxin 1 | 5052 | ENSG00000117450 |

  
  

| **Database:Pathway Commons pathway      &nbspName:Glypican pathway      &nbspID:DB\_ID:1459** | | | | | | |
| --- | --- | --- | --- | --- | --- | --- |
| C=1338; O=9; E=4.16; R=2.17; rawP=0.0218; adjP=0.0245 | | | | | | |
| Index | UserID | Value | Gene Symbol | Gene Name | EntrezGene | Ensembl |
| 1 | P34932 | NA | HSPA4 | heat shock 70kDa protein 4 | 3308 | ENSG00000170606 |
| 2 | P02795 | NA | MT2A | metallothionein 2A | 4502 | ENSG00000125148 |
| 3 | P08195 | NA | SLC3A2 | solute carrier family 3 (activators of dibasic and neutral amino acid transport), member 2 | 6520 | ENSG00000168003 |
| 4 | P13639 | NA | EEF2 | eukaryotic translation elongation factor 2 | 1938 | ENSG00000167658 |
| 5 | P52292 | NA | KPNA2 | karyopherin alpha 2 (RAG cohort 1, importin alpha 1) | 3838 | ENSG00000182481 |
| 6 | P18206 | NA | VCL | vinculin | 7414 | ENSG00000035403 |
| 7 | O60610 | NA | DIAPH1 | diaphanous homolog 1 (Drosophila) | 1729 | ENSG00000131504 |
| 8 | P02786 | NA | TFRC | transferrin receptor (p90, CD71) | 7037 | ENSG00000072274 |
| 9 | Q06830 | NA | PRDX1 | peroxiredoxin 1 | 5052 | ENSG00000117450 |

  
  

| **Database:Pathway Commons pathway      &nbspName:Proteoglycan syndecan-mediated signaling events      &nbspID:DB\_ID:1637** | | | | | | |
| --- | --- | --- | --- | --- | --- | --- |
| C=1345; O=9; E=4.18; R=2.15; rawP=0.0225; adjP=0.0249 | | | | | | |
| Index | UserID | Value | Gene Symbol | Gene Name | EntrezGene | Ensembl |
| 1 | P34932 | NA | HSPA4 | heat shock 70kDa protein 4 | 3308 | ENSG00000170606 |
| 2 | P02795 | NA | MT2A | metallothionein 2A | 4502 | ENSG00000125148 |
| 3 | P08195 | NA | SLC3A2 | solute carrier family 3 (activators of dibasic and neutral amino acid transport), member 2 | 6520 | ENSG00000168003 |
| 4 | P13639 | NA | EEF2 | eukaryotic translation elongation factor 2 | 1938 | ENSG00000167658 |
| 5 | P52292 | NA | KPNA2 | karyopherin alpha 2 (RAG cohort 1, importin alpha 1) | 3838 | ENSG00000182481 |
| 6 | P18206 | NA | VCL | vinculin | 7414 | ENSG00000035403 |
| 7 | O60610 | NA | DIAPH1 | diaphanous homolog 1 (Drosophila) | 1729 | ENSG00000131504 |
| 8 | P02786 | NA | TFRC | transferrin receptor (p90, CD71) | 7037 | ENSG00000072274 |
| 9 | Q06830 | NA | PRDX1 | peroxiredoxin 1 | 5052 | ENSG00000117450 |

  
  

| **Database:Pathway Commons pathway      &nbspName:AP-1 transcription factor network      &nbspID:DB\_ID:1565** | | | | | | |
| --- | --- | --- | --- | --- | --- | --- |
| C=623; O=5; E=1.94; R=2.58; rawP=0.0440; adjP=0.0479 | | | | | | |
| Index | UserID | Value | Gene Symbol | Gene Name | EntrezGene | Ensembl |
| 1 | P52292 | NA | KPNA2 | karyopherin alpha 2 (RAG cohort 1, importin alpha 1) | 3838 | ENSG00000182481 |
| 2 | P02795 | NA | MT2A | metallothionein 2A | 4502 | ENSG00000125148 |
| 3 | P08195 | NA | SLC3A2 | solute carrier family 3 (activators of dibasic and neutral amino acid transport), member 2 | 6520 | ENSG00000168003 |
| 4 | Q06830 | NA | PRDX1 | peroxiredoxin 1 | 5052 | ENSG00000117450 |
| 5 | P02786 | NA | TFRC | transferrin receptor (p90, CD71) | 7037 | ENSG00000072274 |

  
  

| **Database:Pathway Commons pathway      &nbspName:Metabolism of lipids and lipoproteins      &nbspID:DB\_ID:984** | | | | | | |
| --- | --- | --- | --- | --- | --- | --- |
| C=258; O=3; E=0.80; R=3.74; rawP=0.0462; adjP=0.0496 | | | | | | |
| Index | UserID | Value | Gene Symbol | Gene Name | EntrezGene | Ensembl |
| 1 | Q16881 | NA | TXNRD1 | thioredoxin reductase 1 | 7296 | ENSG00000198431 |
| 2 | Q15185 | NA | PTGES3 | prostaglandin E synthase 3 (cytosolic) | 10728 | ENSG00000110958 |
| 3 | P49327 | NA | FASN | fatty acid synthase | 2194 | ENSG00000169710 |

  
  

| **Database:Pathway Commons pathway      &nbspName:Stabilization and expansion of the E-cadherin adherens junction      &nbspID:DB\_ID:1469** | | | | | | |
| --- | --- | --- | --- | --- | --- | --- |
| C=275; O=3; E=0.85; R=3.51; rawP=0.0540; adjP=0.0563 | | | | | | |
| Index | UserID | Value | Gene Symbol | Gene Name | EntrezGene | Ensembl |
| 1 | P18206 | NA | VCL | vinculin | 7414 | ENSG00000035403 |
| 2 | O60610 | NA | DIAPH1 | diaphanous homolog 1 (Drosophila) | 1729 | ENSG00000131504 |
| 3 | Q06830 | NA | PRDX1 | peroxiredoxin 1 | 5052 | ENSG00000117450 |

  
  

| **Database:Pathway Commons pathway      &nbspName:E-cadherin signaling in the nascent adherens junction      &nbspID:DB\_ID:1544** | | | | | | |
| --- | --- | --- | --- | --- | --- | --- |
| C=275; O=3; E=0.85; R=3.51; rawP=0.0540; adjP=0.0563 | | | | | | |
| Index | UserID | Value | Gene Symbol | Gene Name | EntrezGene | Ensembl |
| 1 | P18206 | NA | VCL | vinculin | 7414 | ENSG00000035403 |
| 2 | O60610 | NA | DIAPH1 | diaphanous homolog 1 (Drosophila) | 1729 | ENSG00000131504 |
| 3 | Q06830 | NA | PRDX1 | peroxiredoxin 1 | 5052 | ENSG00000117450 |

  
  

| **Database:Pathway Commons pathway      &nbspName:E-cadherin signaling events      &nbspID:DB\_ID:1617** | | | | | | |
| --- | --- | --- | --- | --- | --- | --- |
| C=280; O=3; E=0.87; R=3.45; rawP=0.0564; adjP=0.0580 | | | | | | |
| Index | UserID | Value | Gene Symbol | Gene Name | EntrezGene | Ensembl |
| 1 | P18206 | NA | VCL | vinculin | 7414 | ENSG00000035403 |
| 2 | O60610 | NA | DIAPH1 | diaphanous homolog 1 (Drosophila) | 1729 | ENSG00000131504 |
| 3 | Q06830 | NA | PRDX1 | peroxiredoxin 1 | 5052 | ENSG00000117450 |

  
  

| **Database:Pathway Commons pathway      &nbspName:TNF receptor signaling pathway      &nbspID:DB\_ID:1600** | | | | | | |
| --- | --- | --- | --- | --- | --- | --- |
| C=299; O=3; E=0.93; R=3.23; rawP=0.0660; adjP=0.0669 | | | | | | |
| Index | UserID | Value | Gene Symbol | Gene Name | EntrezGene | Ensembl |
| 1 | P52292 | NA | KPNA2 | karyopherin alpha 2 (RAG cohort 1, importin alpha 1) | 3838 | ENSG00000182481 |
| 2 | P08195 | NA | SLC3A2 | solute carrier family 3 (activators of dibasic and neutral amino acid transport), member 2 | 6520 | ENSG00000168003 |
| 3 | P11387 | NA | TOP1 | topoisomerase (DNA) I | 7150 | ENSG00000198900 |

  
  
  
  

---

WebGestalt is currently developed and maintained by Jing Wang and Bing Zhang at the  Zhang Lab. Other people who have made significant contribution to the project include Dexter Duncan, Stefan Kirov, Zhiao Shi, and Jay Snoddy.  
  
**Funding credits:** NIH/NIAAA (U01 AA016662, U01 AA013512); NIH/NIDA (P01 DA015027); NIH/NIMH (P50 MH078028, P50 MH096972); NIH/NCI (U24 CA159988); NIH/NIGMS (R01 GM088822).
