## Supplementary material for "PGRMC1 phosphorylation and cell plasticity 1: glycolysis, mitochondria, tumor growth": File S8: final_tf_geneset_file_1439190848.html

Anchored HTML File of EIDs

|  |  |
| --- | --- |
|  | WEB-based GEne SeT AnaLysis Toolkit |
|  | ***Translating gene lists into biological insights...*** |

---

  

| **Database:Transcrription Target      &nbspName:hsa\_TTCNRGNNNNTTC\_V$HSF\_Q6      &nbspID:DB\_ID:2424** | | | | | | |
| --- | --- | --- | --- | --- | --- | --- |
| C=149; O=6; E=0.46; R=12.96; rawP=6.84e-06; adjP=0.0006 | | | | | | |
| Index | UserID | Value | Gene Symbol | Gene Name | EntrezGene | Ensembl |
| 1 | Q92598 | NA | HSPH1 | heat shock 105kDa/110kDa protein 1 | 10808 | ENSG00000120694 |
| 2 | Q99832 | NA | CCT7 | chaperonin containing TCP1, subunit 7 (eta) | 10574 | ENSG00000135624 |
| 3 | P50990 | NA | CCT8 | chaperonin containing TCP1, subunit 8 (theta) | 10694 | ENSG00000156261 |
| 4 | O95433 | NA | AHSA1 | AHA1, activator of heat shock 90kDa protein ATPase homolog 1 (yeast) | 10598 | ENSG00000100591 |
| 5 | Q16555 | NA | DPYSL2 | dihydropyrimidinase-like 2 | 1808 | ENSG00000092964 |
| 6 | P50991 | NA | CCT4 | chaperonin containing TCP1, subunit 4 (delta) | 10575 | ENSG00000115484 |

  
  

| **Database:Transcrription Target      &nbspName:hsa\_V$E2F\_Q4\_01      &nbspID:DB\_ID:2296** | | | | | | |
| --- | --- | --- | --- | --- | --- | --- |
| C=233; O=6; E=0.72; R=8.29; rawP=8.47e-05; adjP=0.0011 | | | | | | |
| Index | UserID | Value | Gene Symbol | Gene Name | EntrezGene | Ensembl |
| 1 | P63241 | NA | EIF5A | eukaryotic translation initiation factor 5A | 1984 | ENSG00000132507 |
| 2 | P12268 | NA | IMPDH2 | IMP (inosine 5'-monophosphate) dehydrogenase 2 | 3615 | ENSG00000178035 |
| 3 | P49736 | NA | MCM2 | minichromosome maintenance complex component 2 | 4171 | ENSG00000073111 |
| 4 | P49321 | NA | NASP | nuclear autoantigenic sperm protein (histone-binding) | 4678 | ENSG00000132780 |
| 5 | Q16555 | NA | DPYSL2 | dihydropyrimidinase-like 2 | 1808 | ENSG00000092964 |
| 6 | P11387 | NA | TOP1 | topoisomerase (DNA) I | 7150 | ENSG00000198900 |

  
  

| **Database:Transcrription Target      &nbspName:hsa\_V$HSF2\_01      &nbspID:DB\_ID:1951** | | | | | | |
| --- | --- | --- | --- | --- | --- | --- |
| C=241; O=6; E=0.75; R=8.02; rawP=0.0001; adjP=0.0011 | | | | | | |
| Index | UserID | Value | Gene Symbol | Gene Name | EntrezGene | Ensembl |
| 1 | P49368 | NA | CCT3 | chaperonin containing TCP1, subunit 3 (gamma) | 7203 | ENSG00000163468 |
| 2 | Q92598 | NA | HSPH1 | heat shock 105kDa/110kDa protein 1 | 10808 | ENSG00000120694 |
| 3 | P25685 | NA | DNAJB1 | DnaJ (Hsp40) homolog, subfamily B, member 1 | 3337 | ENSG00000132002 |
| 4 | P07951 | NA | TPM2 | tropomyosin 2 (beta) | 7169 | ENSG00000198467 |
| 5 | P50990 | NA | CCT8 | chaperonin containing TCP1, subunit 8 (theta) | 10694 | ENSG00000156261 |
| 6 | P50991 | NA | CCT4 | chaperonin containing TCP1, subunit 4 (delta) | 10575 | ENSG00000115484 |

  
  

| **Database:Transcrription Target      &nbspName:hsa\_RGAANNTTC\_V$HSF1\_01      &nbspID:DB\_ID:2425** | | | | | | |
| --- | --- | --- | --- | --- | --- | --- |
| C=433; O=8; E=1.34; R=5.95; rawP=5.60e-05; adjP=0.0011 | | | | | | |
| Index | UserID | Value | Gene Symbol | Gene Name | EntrezGene | Ensembl |
| 1 | P49368 | NA | CCT3 | chaperonin containing TCP1, subunit 3 (gamma) | 7203 | ENSG00000163468 |
| 2 | P78371 | NA | CCT2 | chaperonin containing TCP1, subunit 2 (beta) | 10576 | ENSG00000166226 |
| 3 | Q92598 | NA | HSPH1 | heat shock 105kDa/110kDa protein 1 | 10808 | ENSG00000120694 |
| 4 | P25685 | NA | DNAJB1 | DnaJ (Hsp40) homolog, subfamily B, member 1 | 3337 | ENSG00000132002 |
| 5 | P07951 | NA | TPM2 | tropomyosin 2 (beta) | 7169 | ENSG00000198467 |
| 6 | P50990 | NA | CCT8 | chaperonin containing TCP1, subunit 8 (theta) | 10694 | ENSG00000156261 |
| 7 | P07108 | NA | DBI | diazepam binding inhibitor (GABA receptor modulator, acyl-CoA binding protein) | 1622 | ENSG00000155368 |
| 8 | P50991 | NA | CCT4 | chaperonin containing TCP1, subunit 4 (delta) | 10575 | ENSG00000115484 |

  
  

| **Database:Transcrription Target      &nbspName:hsa\_V$E2F\_03      &nbspID:DB\_ID:2158** | | | | | | |
| --- | --- | --- | --- | --- | --- | --- |
| C=243; O=6; E=0.75; R=7.95; rawP=0.0001; adjP=0.0011 | | | | | | |
| Index | UserID | Value | Gene Symbol | Gene Name | EntrezGene | Ensembl |
| 1 | P63241 | NA | EIF5A | eukaryotic translation initiation factor 5A | 1984 | ENSG00000132507 |
| 2 | P17987 | NA | TCP1 | t-complex 1 | 6950 | ENSG00000120438 |
| 3 | P49736 | NA | MCM2 | minichromosome maintenance complex component 2 | 4171 | ENSG00000073111 |
| 4 | P49321 | NA | NASP | nuclear autoantigenic sperm protein (histone-binding) | 4678 | ENSG00000132780 |
| 5 | Q16555 | NA | DPYSL2 | dihydropyrimidinase-like 2 | 1808 | ENSG00000092964 |
| 6 | P11387 | NA | TOP1 | topoisomerase (DNA) I | 7150 | ENSG00000198900 |

  
  

| **Database:Transcrription Target      &nbspName:hsa\_V$E2F\_Q3      &nbspID:DB\_ID:2091** | | | | | | |
| --- | --- | --- | --- | --- | --- | --- |
| C=223; O=6; E=0.69; R=8.66; rawP=6.65e-05; adjP=0.0011 | | | | | | |
| Index | UserID | Value | Gene Symbol | Gene Name | EntrezGene | Ensembl |
| 1 | P17987 | NA | TCP1 | t-complex 1 | 6950 | ENSG00000120438 |
| 2 | P12268 | NA | IMPDH2 | IMP (inosine 5'-monophosphate) dehydrogenase 2 | 3615 | ENSG00000178035 |
| 3 | P49736 | NA | MCM2 | minichromosome maintenance complex component 2 | 4171 | ENSG00000073111 |
| 4 | P49321 | NA | NASP | nuclear autoantigenic sperm protein (histone-binding) | 4678 | ENSG00000132780 |
| 5 | Q16555 | NA | DPYSL2 | dihydropyrimidinase-like 2 | 1808 | ENSG00000092964 |
| 6 | P02786 | NA | TFRC | transferrin receptor (p90, CD71) | 7037 | ENSG00000072274 |

  
  

| **Database:Transcrription Target      &nbspName:hsa\_CACGTG\_V$MYC\_Q2      &nbspID:DB\_ID:2434** | | | | | | |
| --- | --- | --- | --- | --- | --- | --- |
| C=1008; O=12; E=3.13; R=3.83; rawP=5.42e-05; adjP=0.0011 | | | | | | |
| Index | UserID | Value | Gene Symbol | Gene Name | EntrezGene | Ensembl |
| 1 | Q92598 | NA | HSPH1 | heat shock 105kDa/110kDa protein 1 | 10808 | ENSG00000120694 |
| 2 | P34932 | NA | HSPA4 | heat shock 70kDa protein 4 | 3308 | ENSG00000170606 |
| 3 | P07951 | NA | TPM2 | tropomyosin 2 (beta) | 7169 | ENSG00000198467 |
| 4 | P67809 | NA | YBX1 | Y box binding protein 1 | 4904 | ENSG00000065978 |
| 5 | P37802 | NA | TAGLN2 | transgelin 2 | 8407 | ENSG00000158710 |
| 6 | P49736 | NA | MCM2 | minichromosome maintenance complex component 2 | 4171 | ENSG00000073111 |
| 7 | O60610 | NA | DIAPH1 | diaphanous homolog 1 (Drosophila) | 1729 | ENSG00000131504 |
| 8 | Q8TEX9 | NA | IPO4 | importin 4 | 79711 | ENSG00000196497 |
| 9 | P02786 | NA | TFRC | transferrin receptor (p90, CD71) | 7037 | ENSG00000072274 |
| 10 | O00425 | NA | IGF2BP3 | insulin-like growth factor 2 mRNA binding protein 3 | 10643 | ENSG00000136231 |
| 11 | P11586 | NA | MTHFD1 | methylenetetrahydrofolate dehydrogenase (NADP+ dependent) 1, methenyltetrahydrofolate cyclohydrolase, formyltetrahydrofolate synthetase | 4522 | ENSG00000100714 |
| 12 | P11387 | NA | TOP1 | topoisomerase (DNA) I | 7150 | ENSG00000198900 |

  
  

| **Database:Transcrription Target      &nbspName:hsa\_GGGCGGR\_V$SP1\_Q6      &nbspID:DB\_ID:2452** | | | | | | |
| --- | --- | --- | --- | --- | --- | --- |
| C=2863; O=22; E=8.89; R=2.47; rawP=2.74e-05; adjP=0.0011 | | | | | | |
| Index | UserID | Value | Gene Symbol | Gene Name | EntrezGene | Ensembl |
| 1 | P63241 | NA | EIF5A | eukaryotic translation initiation factor 5A | 1984 | ENSG00000132507 |
| 2 | Q16881 | NA | TXNRD1 | thioredoxin reductase 1 | 7296 | ENSG00000198431 |
| 3 | P78371 | NA | CCT2 | chaperonin containing TCP1, subunit 2 (beta) | 10576 | ENSG00000166226 |
| 4 | Q92598 | NA | HSPH1 | heat shock 105kDa/110kDa protein 1 | 10808 | ENSG00000120694 |
| 5 | P02795 | NA | MT2A | metallothionein 2A | 4502 | ENSG00000125148 |
| 6 | P49321 | NA | NASP | nuclear autoantigenic sperm protein (histone-binding) | 4678 | ENSG00000132780 |
| 7 | P07951 | NA | TPM2 | tropomyosin 2 (beta) | 7169 | ENSG00000198467 |
| 8 | P49368 | NA | CCT3 | chaperonin containing TCP1, subunit 3 (gamma) | 7203 | ENSG00000163468 |
| 9 | Q9BQE3 | NA | TUBA1C | tubulin, alpha 1c | 84790 | ENSG00000167553 |
| 10 | O60610 | NA | DIAPH1 | diaphanous homolog 1 (Drosophila) | 1729 | ENSG00000131504 |
| 11 | P68104 | NA | EEF1A1 | eukaryotic translation elongation factor 1 alpha 1 | 1915 | ENSG00000156508 |
| 12 | O00425 | NA | IGF2BP3 | insulin-like growth factor 2 mRNA binding protein 3 | 10643 | ENSG00000136231 |
| 13 | P34932 | NA | HSPA4 | heat shock 70kDa protein 4 | 3308 | ENSG00000170606 |
| 14 | P41091 | NA | EIF2S3 | eukaryotic translation initiation factor 2, subunit 3 gamma, 52kDa | 1968 | ENSG00000130741 |
| 15 | Q16555 | NA | DPYSL2 | dihydropyrimidinase-like 2 | 1808 | ENSG00000092964 |
| 16 | P41567 | NA | EIF1 | eukaryotic translation initiation factor 1 | 10209 | ENSG00000173812 |
| 17 | P67809 | NA | YBX1 | Y box binding protein 1 | 4904 | ENSG00000065978 |
| 18 | P37802 | NA | TAGLN2 | transgelin 2 | 8407 | ENSG00000158710 |
| 19 | Q99832 | NA | CCT7 | chaperonin containing TCP1, subunit 7 (eta) | 10574 | ENSG00000135624 |
| 20 | P18206 | NA | VCL | vinculin | 7414 | ENSG00000035403 |
| 21 | P11387 | NA | TOP1 | topoisomerase (DNA) I | 7150 | ENSG00000198900 |
| 22 | Q9UK76 | NA | HN1 | hematological and neurological expressed 1 | 51155 | ENSG00000189159 |

  
  

| **Database:Transcrription Target      &nbspName:hsa\_SCGGAAGY\_V$ELK1\_02      &nbspID:DB\_ID:2413** | | | | | | |
| --- | --- | --- | --- | --- | --- | --- |
| C=1170; O=12; E=3.63; R=3.30; rawP=0.0002; adjP=0.0018 | | | | | | |
| Index | UserID | Value | Gene Symbol | Gene Name | EntrezGene | Ensembl |
| 1 | P63241 | NA | EIF5A | eukaryotic translation initiation factor 5A | 1984 | ENSG00000132507 |
| 2 | Q92598 | NA | HSPH1 | heat shock 105kDa/110kDa protein 1 | 10808 | ENSG00000120694 |
| 3 | P34932 | NA | HSPA4 | heat shock 70kDa protein 4 | 3308 | ENSG00000170606 |
| 4 | P22102 | NA | GART | phosphoribosylglycinamide formyltransferase, phosphoribosylglycinamide synthetase, phosphoribosylaminoimidazole synthetase | 2618 | ENSG00000159131 |
| 5 | P49321 | NA | NASP | nuclear autoantigenic sperm protein (histone-binding) | 4678 | ENSG00000132780 |
| 6 | P12081 | NA | HARS | histidyl-tRNA synthetase | 3035 | ENSG00000170445 |
| 7 | P56192 | NA | MARS | methionyl-tRNA synthetase | 4141 | ENSG00000166986 |
| 8 | P25685 | NA | DNAJB1 | DnaJ (Hsp40) homolog, subfamily B, member 1 | 3337 | ENSG00000132002 |
| 9 | Q99832 | NA | CCT7 | chaperonin containing TCP1, subunit 7 (eta) | 10574 | ENSG00000135624 |
| 10 | P53701 | NA | HCCS | holocytochrome c synthase | 3052 | ENSG00000004961 |
| 11 | O60610 | NA | DIAPH1 | diaphanous homolog 1 (Drosophila) | 1729 | ENSG00000131504 |
| 12 | Q8TEX9 | NA | IPO4 | importin 4 | 79711 | ENSG00000196497 |

  
  

| **Database:Transcrription Target      &nbspName:hsa\_V$USF\_C      &nbspID:DB\_ID:1999** | | | | | | |
| --- | --- | --- | --- | --- | --- | --- |
| C=274; O=6; E=0.85; R=7.05; rawP=0.0002; adjP=0.0018 | | | | | | |
| Index | UserID | Value | Gene Symbol | Gene Name | EntrezGene | Ensembl |
| 1 | P63241 | NA | EIF5A | eukaryotic translation initiation factor 5A | 1984 | ENSG00000132507 |
| 2 | P67809 | NA | YBX1 | Y box binding protein 1 | 4904 | ENSG00000065978 |
| 3 | Q92598 | NA | HSPH1 | heat shock 105kDa/110kDa protein 1 | 10808 | ENSG00000120694 |
| 4 | P34932 | NA | HSPA4 | heat shock 70kDa protein 4 | 3308 | ENSG00000170606 |
| 5 | Q8TEX9 | NA | IPO4 | importin 4 | 79711 | ENSG00000196497 |
| 6 | P11586 | NA | MTHFD1 | methylenetetrahydrofolate dehydrogenase (NADP+ dependent) 1, methenyltetrahydrofolate cyclohydrolase, formyltetrahydrofolate synthetase | 4522 | ENSG00000100714 |

  
  

| **Database:Transcrription Target      &nbspName:hsa\_V$E2F1\_Q4\_01      &nbspID:DB\_ID:2312** | | | | | | |
| --- | --- | --- | --- | --- | --- | --- |
| C=225; O=5; E=0.70; R=7.15; rawP=0.0007; adjP=0.0059 | | | | | | |
| Index | UserID | Value | Gene Symbol | Gene Name | EntrezGene | Ensembl |
| 1 | P12268 | NA | IMPDH2 | IMP (inosine 5'-monophosphate) dehydrogenase 2 | 3615 | ENSG00000178035 |
| 2 | P49736 | NA | MCM2 | minichromosome maintenance complex component 2 | 4171 | ENSG00000073111 |
| 3 | P49321 | NA | NASP | nuclear autoantigenic sperm protein (histone-binding) | 4678 | ENSG00000132780 |
| 4 | Q16555 | NA | DPYSL2 | dihydropyrimidinase-like 2 | 1808 | ENSG00000092964 |
| 5 | P11387 | NA | TOP1 | topoisomerase (DNA) I | 7150 | ENSG00000198900 |

  
  

| **Database:Transcrription Target      &nbspName:hsa\_CATTGTYY\_V$SOX9\_B1      &nbspID:DB\_ID:2451** | | | | | | |
| --- | --- | --- | --- | --- | --- | --- |
| C=350; O=6; E=1.09; R=5.52; rawP=0.0008; adjP=0.0061 | | | | | | |
| Index | UserID | Value | Gene Symbol | Gene Name | EntrezGene | Ensembl |
| 1 | P49321 | NA | NASP | nuclear autoantigenic sperm protein (histone-binding) | 4678 | ENSG00000132780 |
| 2 | P18206 | NA | VCL | vinculin | 7414 | ENSG00000035403 |
| 3 | Q06830 | NA | PRDX1 | peroxiredoxin 1 | 5052 | ENSG00000117450 |
| 4 | P67936 | NA | TPM4 | tropomyosin 4 | 7171 | ENSG00000167460 |
| 5 | Q9UK76 | NA | HN1 | hematological and neurological expressed 1 | 51155 | ENSG00000189159 |
| 6 | P11387 | NA | TOP1 | topoisomerase (DNA) I | 7150 | ENSG00000198900 |

  
  

| **Database:Transcrription Target      &nbspName:hsa\_V$HSF1\_01      &nbspID:DB\_ID:1949** | | | | | | |
| --- | --- | --- | --- | --- | --- | --- |
| C=258; O=5; E=0.80; R=6.24; rawP=0.0012; adjP=0.0085 | | | | | | |
| Index | UserID | Value | Gene Symbol | Gene Name | EntrezGene | Ensembl |
| 1 | P49368 | NA | CCT3 | chaperonin containing TCP1, subunit 3 (gamma) | 7203 | ENSG00000163468 |
| 2 | Q92598 | NA | HSPH1 | heat shock 105kDa/110kDa protein 1 | 10808 | ENSG00000120694 |
| 3 | P25685 | NA | DNAJB1 | DnaJ (Hsp40) homolog, subfamily B, member 1 | 3337 | ENSG00000132002 |
| 4 | P07951 | NA | TPM2 | tropomyosin 2 (beta) | 7169 | ENSG00000198467 |
| 5 | P07108 | NA | DBI | diazepam binding inhibitor (GABA receptor modulator, acyl-CoA binding protein) | 1622 | ENSG00000155368 |

  
  

| **Database:Transcrription Target      &nbspName:hsa\_TMTCGCGANR\_UNKNOWN      &nbspID:DB\_ID:2370** | | | | | | |
| --- | --- | --- | --- | --- | --- | --- |
| C=156; O=4; E=0.48; R=8.26; rawP=0.0014; adjP=0.0092 | | | | | | |
| Index | UserID | Value | Gene Symbol | Gene Name | EntrezGene | Ensembl |
| 1 | P34932 | NA | HSPA4 | heat shock 70kDa protein 4 | 3308 | ENSG00000170606 |
| 2 | P62906 | NA | RPL10A | ribosomal protein L10a | 4736 | ENSG00000198755 |
| 3 | P50990 | NA | CCT8 | chaperonin containing TCP1, subunit 8 (theta) | 10694 | ENSG00000156261 |
| 4 | Q06830 | NA | PRDX1 | peroxiredoxin 1 | 5052 | ENSG00000117450 |

  
  

| **Database:Transcrription Target      &nbspName:hsa\_V$NMYC\_01      &nbspID:DB\_ID:1875** | | | | | | |
| --- | --- | --- | --- | --- | --- | --- |
| C=268; O=5; E=0.83; R=6.01; rawP=0.0015; adjP=0.0092 | | | | | | |
| Index | UserID | Value | Gene Symbol | Gene Name | EntrezGene | Ensembl |
| 1 | P67809 | NA | YBX1 | Y box binding protein 1 | 4904 | ENSG00000065978 |
| 2 | P07951 | NA | TPM2 | tropomyosin 2 (beta) | 7169 | ENSG00000198467 |
| 3 | Q8TEX9 | NA | IPO4 | importin 4 | 79711 | ENSG00000196497 |
| 4 | P11387 | NA | TOP1 | topoisomerase (DNA) I | 7150 | ENSG00000198900 |
| 5 | P11586 | NA | MTHFD1 | methylenetetrahydrofolate dehydrogenase (NADP+ dependent) 1, methenyltetrahydrofolate cyclohydrolase, formyltetrahydrofolate synthetase | 4522 | ENSG00000100714 |

  
  

| **Database:Transcrription Target      &nbspName:hsa\_GGCNNMSMYNTTG\_UNKNOWN      &nbspID:DB\_ID:2313** | | | | | | |
| --- | --- | --- | --- | --- | --- | --- |
| C=74; O=3; E=0.23; R=13.05; rawP=0.0016; adjP=0.0092 | | | | | | |
| Index | UserID | Value | Gene Symbol | Gene Name | EntrezGene | Ensembl |
| 1 | P67809 | NA | YBX1 | Y box binding protein 1 | 4904 | ENSG00000065978 |
| 2 | P49321 | NA | NASP | nuclear autoantigenic sperm protein (histone-binding) | 4678 | ENSG00000132780 |
| 3 | P68104 | NA | EEF1A1 | eukaryotic translation elongation factor 1 alpha 1 | 1915 | ENSG00000156508 |

  
  

| **Database:Transcrription Target      &nbspName:hsa\_V$HSF\_Q6      &nbspID:DB\_ID:2184** | | | | | | |
| --- | --- | --- | --- | --- | --- | --- |
| C=193; O=4; E=0.60; R=6.67; rawP=0.0031; adjP=0.0168 | | | | | | |
| Index | UserID | Value | Gene Symbol | Gene Name | EntrezGene | Ensembl |
| 1 | P49368 | NA | CCT3 | chaperonin containing TCP1, subunit 3 (gamma) | 7203 | ENSG00000163468 |
| 2 | Q92598 | NA | HSPH1 | heat shock 105kDa/110kDa protein 1 | 10808 | ENSG00000120694 |
| 3 | P25685 | NA | DNAJB1 | DnaJ (Hsp40) homolog, subfamily B, member 1 | 3337 | ENSG00000132002 |
| 4 | P41567 | NA | EIF1 | eukaryotic translation initiation factor 1 | 10209 | ENSG00000173812 |

  
  

| **Database:Transcrription Target      &nbspName:hsa\_TCCCRNNRTGC\_UNKNOWN      &nbspID:DB\_ID:2364** | | | | | | |
| --- | --- | --- | --- | --- | --- | --- |
| C=210; O=4; E=0.65; R=6.13; rawP=0.0041; adjP=0.0210 | | | | | | |
| Index | UserID | Value | Gene Symbol | Gene Name | EntrezGene | Ensembl |
| 1 | P63241 | NA | EIF5A | eukaryotic translation initiation factor 5A | 1984 | ENSG00000132507 |
| 2 | Q92598 | NA | HSPH1 | heat shock 105kDa/110kDa protein 1 | 10808 | ENSG00000120694 |
| 3 | P22102 | NA | GART | phosphoribosylglycinamide formyltransferase, phosphoribosylglycinamide synthetase, phosphoribosylaminoimidazole synthetase | 2618 | ENSG00000159131 |
| 4 | P18206 | NA | VCL | vinculin | 7414 | ENSG00000035403 |

  
  

| **Database:Transcrription Target      &nbspName:hsa\_V$AP1\_Q4      &nbspID:DB\_ID:1977** | | | | | | |
| --- | --- | --- | --- | --- | --- | --- |
| C=262; O=4; E=0.81; R=4.92; rawP=0.0089; adjP=0.0227 | | | | | | |
| Index | UserID | Value | Gene Symbol | Gene Name | EntrezGene | Ensembl |
| 1 | Q9BQE3 | NA | TUBA1C | tubulin, alpha 1c | 84790 | ENSG00000167553 |
| 2 | P37802 | NA | TAGLN2 | transgelin 2 | 8407 | ENSG00000158710 |
| 3 | O60610 | NA | DIAPH1 | diaphanous homolog 1 (Drosophila) | 1729 | ENSG00000131504 |
| 4 | P68104 | NA | EEF1A1 | eukaryotic translation elongation factor 1 alpha 1 | 1915 | ENSG00000156508 |

  
  

| **Database:Transcrription Target      &nbspName:hsa\_V$ATF\_01      &nbspID:DB\_ID:1856** | | | | | | |
| --- | --- | --- | --- | --- | --- | --- |
| C=254; O=4; E=0.79; R=5.07; rawP=0.0080; adjP=0.0227 | | | | | | |
| Index | UserID | Value | Gene Symbol | Gene Name | EntrezGene | Ensembl |
| 1 | P22102 | NA | GART | phosphoribosylglycinamide formyltransferase, phosphoribosylglycinamide synthetase, phosphoribosylaminoimidazole synthetase | 2618 | ENSG00000159131 |
| 2 | P41567 | NA | EIF1 | eukaryotic translation initiation factor 1 | 10209 | ENSG00000173812 |
| 3 | P67936 | NA | TPM4 | tropomyosin 4 | 7171 | ENSG00000167460 |
| 4 | P11387 | NA | TOP1 | topoisomerase (DNA) I | 7150 | ENSG00000198900 |

  
  

| **Database:Transcrription Target      &nbspName:hsa\_V$NRF1\_Q6      &nbspID:DB\_ID:2193** | | | | | | |
| --- | --- | --- | --- | --- | --- | --- |
| C=244; O=4; E=0.76; R=5.28; rawP=0.0070; adjP=0.0227 | | | | | | |
| Index | UserID | Value | Gene Symbol | Gene Name | EntrezGene | Ensembl |
| 1 | P49368 | NA | CCT3 | chaperonin containing TCP1, subunit 3 (gamma) | 7203 | ENSG00000163468 |
| 2 | P63241 | NA | EIF5A | eukaryotic translation initiation factor 5A | 1984 | ENSG00000132507 |
| 3 | P56192 | NA | MARS | methionyl-tRNA synthetase | 4141 | ENSG00000166986 |
| 4 | P41567 | NA | EIF1 | eukaryotic translation initiation factor 1 | 10209 | ENSG00000173812 |

  
  

| **Database:Transcrription Target      &nbspName:hsa\_V$E2F\_Q6\_01      &nbspID:DB\_ID:2297** | | | | | | |
| --- | --- | --- | --- | --- | --- | --- |
| C=237; O=4; E=0.74; R=5.43; rawP=0.0063; adjP=0.0227 | | | | | | |
| Index | UserID | Value | Gene Symbol | Gene Name | EntrezGene | Ensembl |
| 1 | P63241 | NA | EIF5A | eukaryotic translation initiation factor 5A | 1984 | ENSG00000132507 |
| 2 | P49736 | NA | MCM2 | minichromosome maintenance complex component 2 | 4171 | ENSG00000073111 |
| 3 | P49321 | NA | NASP | nuclear autoantigenic sperm protein (histone-binding) | 4678 | ENSG00000132780 |
| 4 | Q16555 | NA | DPYSL2 | dihydropyrimidinase-like 2 | 1808 | ENSG00000092964 |

  
  

| **Database:Transcrription Target      &nbspName:hsa\_V$MYCMAX\_02      &nbspID:DB\_ID:1927** | | | | | | |
| --- | --- | --- | --- | --- | --- | --- |
| C=261; O=4; E=0.81; R=4.93; rawP=0.0088; adjP=0.0227 | | | | | | |
| Index | UserID | Value | Gene Symbol | Gene Name | EntrezGene | Ensembl |
| 1 | Q92598 | NA | HSPH1 | heat shock 105kDa/110kDa protein 1 | 10808 | ENSG00000120694 |
| 2 | P34932 | NA | HSPA4 | heat shock 70kDa protein 4 | 3308 | ENSG00000170606 |
| 3 | P11387 | NA | TOP1 | topoisomerase (DNA) I | 7150 | ENSG00000198900 |
| 4 | P11586 | NA | MTHFD1 | methylenetetrahydrofolate dehydrogenase (NADP+ dependent) 1, methenyltetrahydrofolate cyclohydrolase, formyltetrahydrofolate synthetase | 4522 | ENSG00000100714 |

  
  

| **Database:Transcrription Target      &nbspName:hsa\_GATTGGY\_V$NFY\_Q6\_01      &nbspID:DB\_ID:2440** | | | | | | |
| --- | --- | --- | --- | --- | --- | --- |
| C=1129; O=9; E=3.51; R=2.57; rawP=0.0078; adjP=0.0227 | | | | | | |
| Index | UserID | Value | Gene Symbol | Gene Name | EntrezGene | Ensembl |
| 1 | P22102 | NA | GART | phosphoribosylglycinamide formyltransferase, phosphoribosylglycinamide synthetase, phosphoribosylaminoimidazole synthetase | 2618 | ENSG00000159131 |
| 2 | P49321 | NA | NASP | nuclear autoantigenic sperm protein (histone-binding) | 4678 | ENSG00000132780 |
| 3 | Q16555 | NA | DPYSL2 | dihydropyrimidinase-like 2 | 1808 | ENSG00000092964 |
| 4 | P07108 | NA | DBI | diazepam binding inhibitor (GABA receptor modulator, acyl-CoA binding protein) | 1622 | ENSG00000155368 |
| 5 | P49368 | NA | CCT3 | chaperonin containing TCP1, subunit 3 (gamma) | 7203 | ENSG00000163468 |
| 6 | P17174 | NA | GOT1 | glutamic-oxaloacetic transaminase 1, soluble (aspartate aminotransferase 1) | 2805 | ENSG00000120053 |
| 7 | Q9BQE3 | NA | TUBA1C | tubulin, alpha 1c | 84790 | ENSG00000167553 |
| 8 | P25685 | NA | DNAJB1 | DnaJ (Hsp40) homolog, subfamily B, member 1 | 3337 | ENSG00000132002 |
| 9 | Q99832 | NA | CCT7 | chaperonin containing TCP1, subunit 7 (eta) | 10574 | ENSG00000135624 |

  
  

| **Database:Transcrription Target      &nbspName:hsa\_V$MYCMAX\_01      &nbspID:DB\_ID:1923** | | | | | | |
| --- | --- | --- | --- | --- | --- | --- |
| C=250; O=4; E=0.78; R=5.15; rawP=0.0076; adjP=0.0227 | | | | | | |
| Index | UserID | Value | Gene Symbol | Gene Name | EntrezGene | Ensembl |
| 1 | O60610 | NA | DIAPH1 | diaphanous homolog 1 (Drosophila) | 1729 | ENSG00000131504 |
| 2 | P02786 | NA | TFRC | transferrin receptor (p90, CD71) | 7037 | ENSG00000072274 |
| 3 | P11387 | NA | TOP1 | topoisomerase (DNA) I | 7150 | ENSG00000198900 |
| 4 | P11586 | NA | MTHFD1 | methylenetetrahydrofolate dehydrogenase (NADP+ dependent) 1, methenyltetrahydrofolate cyclohydrolase, formyltetrahydrofolate synthetase | 4522 | ENSG00000100714 |

  
  

| **Database:Transcrription Target      &nbspName:hsa\_RCGCANGCGY\_V$NRF1\_Q6      &nbspID:DB\_ID:2441** | | | | | | |
| --- | --- | --- | --- | --- | --- | --- |
| C=891; O=8; E=2.77; R=2.89; rawP=0.0061; adjP=0.0227 | | | | | | |
| Index | UserID | Value | Gene Symbol | Gene Name | EntrezGene | Ensembl |
| 1 | P49368 | NA | CCT3 | chaperonin containing TCP1, subunit 3 (gamma) | 7203 | ENSG00000163468 |
| 2 | P63241 | NA | EIF5A | eukaryotic translation initiation factor 5A | 1984 | ENSG00000132507 |
| 3 | P56192 | NA | MARS | methionyl-tRNA synthetase | 4141 | ENSG00000166986 |
| 4 | P08195 | NA | SLC3A2 | solute carrier family 3 (activators of dibasic and neutral amino acid transport), member 2 | 6520 | ENSG00000168003 |
| 5 | P22102 | NA | GART | phosphoribosylglycinamide formyltransferase, phosphoribosylglycinamide synthetase, phosphoribosylaminoimidazole synthetase | 2618 | ENSG00000159131 |
| 6 | Q16555 | NA | DPYSL2 | dihydropyrimidinase-like 2 | 1808 | ENSG00000092964 |
| 7 | P41567 | NA | EIF1 | eukaryotic translation initiation factor 1 | 10209 | ENSG00000173812 |
| 8 | P11387 | NA | TOP1 | topoisomerase (DNA) I | 7150 | ENSG00000198900 |

  
  

| **Database:Transcrription Target      &nbspName:hsa\_V$ELK1\_02      &nbspID:DB\_ID:1859** | | | | | | |
| --- | --- | --- | --- | --- | --- | --- |
| C=245; O=4; E=0.76; R=5.26; rawP=0.0071; adjP=0.0227 | | | | | | |
| Index | UserID | Value | Gene Symbol | Gene Name | EntrezGene | Ensembl |
| 1 | P63241 | NA | EIF5A | eukaryotic translation initiation factor 5A | 1984 | ENSG00000132507 |
| 2 | P56192 | NA | MARS | methionyl-tRNA synthetase | 4141 | ENSG00000166986 |
| 3 | P22102 | NA | GART | phosphoribosylglycinamide formyltransferase, phosphoribosylglycinamide synthetase, phosphoribosylaminoimidazole synthetase | 2618 | ENSG00000159131 |
| 4 | Q8TEX9 | NA | IPO4 | importin 4 | 79711 | ENSG00000196497 |

  
  

| **Database:Transcrription Target      &nbspName:hsa\_V$E2F1\_Q3      &nbspID:DB\_ID:2095** | | | | | | |
| --- | --- | --- | --- | --- | --- | --- |
| C=240; O=4; E=0.75; R=5.37; rawP=0.0066; adjP=0.0227 | | | | | | |
| Index | UserID | Value | Gene Symbol | Gene Name | EntrezGene | Ensembl |
| 1 | P17987 | NA | TCP1 | t-complex 1 | 6950 | ENSG00000120438 |
| 2 | P49736 | NA | MCM2 | minichromosome maintenance complex component 2 | 4171 | ENSG00000073111 |
| 3 | P49321 | NA | NASP | nuclear autoantigenic sperm protein (histone-binding) | 4678 | ENSG00000132780 |
| 4 | P02786 | NA | TFRC | transferrin receptor (p90, CD71) | 7037 | ENSG00000072274 |

  
  

| **Database:Transcrription Target      &nbspName:hsa\_V$ERR1\_Q2      &nbspID:DB\_ID:2154** | | | | | | |
| --- | --- | --- | --- | --- | --- | --- |
| C=254; O=4; E=0.79; R=5.07; rawP=0.0080; adjP=0.0227 | | | | | | |
| Index | UserID | Value | Gene Symbol | Gene Name | EntrezGene | Ensembl |
| 1 | P17174 | NA | GOT1 | glutamic-oxaloacetic transaminase 1, soluble (aspartate aminotransferase 1) | 2805 | ENSG00000120053 |
| 2 | Q15056 | NA | EIF4H | eukaryotic translation initiation factor 4H | 7458 | ENSG00000106682 |
| 3 | P18206 | NA | VCL | vinculin | 7414 | ENSG00000035403 |
| 4 | O60610 | NA | DIAPH1 | diaphanous homolog 1 (Drosophila) | 1729 | ENSG00000131504 |

  
  

| **Database:Transcrription Target      &nbspName:hsa\_TCCATTKW\_UNKNOWN      &nbspID:DB\_ID:2363** | | | | | | |
| --- | --- | --- | --- | --- | --- | --- |
| C=231; O=4; E=0.72; R=5.57; rawP=0.0058; adjP=0.0227 | | | | | | |
| Index | UserID | Value | Gene Symbol | Gene Name | EntrezGene | Ensembl |
| 1 | P22102 | NA | GART | phosphoribosylglycinamide formyltransferase, phosphoribosylglycinamide synthetase, phosphoribosylaminoimidazole synthetase | 2618 | ENSG00000159131 |
| 2 | P49321 | NA | NASP | nuclear autoantigenic sperm protein (histone-binding) | 4678 | ENSG00000132780 |
| 3 | O60610 | NA | DIAPH1 | diaphanous homolog 1 (Drosophila) | 1729 | ENSG00000131504 |
| 4 | Q16555 | NA | DPYSL2 | dihydropyrimidinase-like 2 | 1808 | ENSG00000092964 |

  
  

| **Database:Transcrription Target      &nbspName:hsa\_V$AREB6\_01      &nbspID:DB\_ID:2078** | | | | | | |
| --- | --- | --- | --- | --- | --- | --- |
| C=260; O=4; E=0.81; R=4.95; rawP=0.0087; adjP=0.0227 | | | | | | |
| Index | UserID | Value | Gene Symbol | Gene Name | EntrezGene | Ensembl |
| 1 | P00352 | NA | ALDH1A1 | aldehyde dehydrogenase 1 family, member A1 | 216 | ENSG00000165092 |
| 2 | O60610 | NA | DIAPH1 | diaphanous homolog 1 (Drosophila) | 1729 | ENSG00000131504 |
| 3 | O75369 | NA | FLNB | filamin B, beta | 2317 | ENSG00000136068 |
| 4 | Q9UK76 | NA | HN1 | hematological and neurological expressed 1 | 51155 | ENSG00000189159 |

  
  

| **Database:Transcrription Target      &nbspName:hsa\_V$AP1\_Q6\_01      &nbspID:DB\_ID:2301** | | | | | | |
| --- | --- | --- | --- | --- | --- | --- |
| C=259; O=4; E=0.80; R=4.97; rawP=0.0086; adjP=0.0227 | | | | | | |
| Index | UserID | Value | Gene Symbol | Gene Name | EntrezGene | Ensembl |
| 1 | Q9BQE3 | NA | TUBA1C | tubulin, alpha 1c | 84790 | ENSG00000167553 |
| 2 | P37802 | NA | TAGLN2 | transgelin 2 | 8407 | ENSG00000158710 |
| 3 | P18206 | NA | VCL | vinculin | 7414 | ENSG00000035403 |
| 4 | O60610 | NA | DIAPH1 | diaphanous homolog 1 (Drosophila) | 1729 | ENSG00000131504 |

  
  

| **Database:Transcrription Target      &nbspName:hsa\_V$TATA\_01      &nbspID:DB\_ID:2025** | | | | | | |
| --- | --- | --- | --- | --- | --- | --- |
| C=254; O=4; E=0.79; R=5.07; rawP=0.0080; adjP=0.0227 | | | | | | |
| Index | UserID | Value | Gene Symbol | Gene Name | EntrezGene | Ensembl |
| 1 | P49321 | NA | NASP | nuclear autoantigenic sperm protein (histone-binding) | 4678 | ENSG00000132780 |
| 2 | P13639 | NA | EEF2 | eukaryotic translation elongation factor 2 | 1938 | ENSG00000167658 |
| 3 | P68104 | NA | EEF1A1 | eukaryotic translation elongation factor 1 alpha 1 | 1915 | ENSG00000156508 |
| 4 | Q06830 | NA | PRDX1 | peroxiredoxin 1 | 5052 | ENSG00000117450 |

  
  

| **Database:Transcrription Target      &nbspName:hsa\_V$AP1FJ\_Q2      &nbspID:DB\_ID:1963** | | | | | | |
| --- | --- | --- | --- | --- | --- | --- |
| C=261; O=4; E=0.81; R=4.93; rawP=0.0088; adjP=0.0227 | | | | | | |
| Index | UserID | Value | Gene Symbol | Gene Name | EntrezGene | Ensembl |
| 1 | Q9BQE3 | NA | TUBA1C | tubulin, alpha 1c | 84790 | ENSG00000167553 |
| 2 | P37802 | NA | TAGLN2 | transgelin 2 | 8407 | ENSG00000158710 |
| 3 | O60610 | NA | DIAPH1 | diaphanous homolog 1 (Drosophila) | 1729 | ENSG00000131504 |
| 4 | P68104 | NA | EEF1A1 | eukaryotic translation elongation factor 1 alpha 1 | 1915 | ENSG00000156508 |

  
  

| **Database:Transcrription Target      &nbspName:hsa\_GGCNKCCATNK\_UNKNOWN      &nbspID:DB\_ID:2302** | | | | | | |
| --- | --- | --- | --- | --- | --- | --- |
| C=116; O=3; E=0.36; R=8.33; rawP=0.0056; adjP=0.0227 | | | | | | |
| Index | UserID | Value | Gene Symbol | Gene Name | EntrezGene | Ensembl |
| 1 | P67809 | NA | YBX1 | Y box binding protein 1 | 4904 | ENSG00000065978 |
| 2 | P49321 | NA | NASP | nuclear autoantigenic sperm protein (histone-binding) | 4678 | ENSG00000132780 |
| 3 | Q16555 | NA | DPYSL2 | dihydropyrimidinase-like 2 | 1808 | ENSG00000092964 |

  
  

| **Database:Transcrription Target      &nbspName:hsa\_V$E2F1DP1RB\_01      &nbspID:DB\_ID:2235** | | | | | | |
| --- | --- | --- | --- | --- | --- | --- |
| C=226; O=4; E=0.70; R=5.70; rawP=0.0053; adjP=0.0227 | | | | | | |
| Index | UserID | Value | Gene Symbol | Gene Name | EntrezGene | Ensembl |
| 1 | P12268 | NA | IMPDH2 | IMP (inosine 5'-monophosphate) dehydrogenase 2 | 3615 | ENSG00000178035 |
| 2 | P49736 | NA | MCM2 | minichromosome maintenance complex component 2 | 4171 | ENSG00000073111 |
| 3 | P49321 | NA | NASP | nuclear autoantigenic sperm protein (histone-binding) | 4678 | ENSG00000132780 |
| 4 | Q16555 | NA | DPYSL2 | dihydropyrimidinase-like 2 | 1808 | ENSG00000092964 |

  
  

| **Database:Transcrription Target      &nbspName:hsa\_CCAWYNNGAAR\_UNKNOWN      &nbspID:DB\_ID:2060** | | | | | | |
| --- | --- | --- | --- | --- | --- | --- |
| C=141; O=3; E=0.44; R=6.85; rawP=0.0096; adjP=0.0239 | | | | | | |
| Index | UserID | Value | Gene Symbol | Gene Name | EntrezGene | Ensembl |
| 1 | P67809 | NA | YBX1 | Y box binding protein 1 | 4904 | ENSG00000065978 |
| 2 | Q99832 | NA | CCT7 | chaperonin containing TCP1, subunit 7 (eta) | 10574 | ENSG00000135624 |
| 3 | P41567 | NA | EIF1 | eukaryotic translation initiation factor 1 | 10209 | ENSG00000173812 |

  
  

| **Database:Transcrription Target      &nbspName:hsa\_GGGYGTGNY\_UNKNOWN      &nbspID:DB\_ID:2333** | | | | | | |
| --- | --- | --- | --- | --- | --- | --- |
| C=648; O=6; E=2.01; R=2.98; rawP=0.0151; adjP=0.0366 | | | | | | |
| Index | UserID | Value | Gene Symbol | Gene Name | EntrezGene | Ensembl |
| 1 | Q16881 | NA | TXNRD1 | thioredoxin reductase 1 | 7296 | ENSG00000198431 |
| 2 | P25685 | NA | DNAJB1 | DnaJ (Hsp40) homolog, subfamily B, member 1 | 3337 | ENSG00000132002 |
| 3 | Q16555 | NA | DPYSL2 | dihydropyrimidinase-like 2 | 1808 | ENSG00000092964 |
| 4 | P07108 | NA | DBI | diazepam binding inhibitor (GABA receptor modulator, acyl-CoA binding protein) | 1622 | ENSG00000155368 |
| 5 | P41567 | NA | EIF1 | eukaryotic translation initiation factor 1 | 10209 | ENSG00000173812 |
| 6 | O75369 | NA | FLNB | filamin B, beta | 2317 | ENSG00000136068 |

  
  

| **Database:Transcrription Target      &nbspName:hsa\_TTCYRGAA\_UNKNOWN      &nbspID:DB\_ID:2374** | | | | | | |
| --- | --- | --- | --- | --- | --- | --- |
| C=320; O=4; E=0.99; R=4.02; rawP=0.0175; adjP=0.0413 | | | | | | |
| Index | UserID | Value | Gene Symbol | Gene Name | EntrezGene | Ensembl |
| 1 | Q92598 | NA | HSPH1 | heat shock 105kDa/110kDa protein 1 | 10808 | ENSG00000120694 |
| 2 | P50990 | NA | CCT8 | chaperonin containing TCP1, subunit 8 (theta) | 10694 | ENSG00000156261 |
| 3 | P11387 | NA | TOP1 | topoisomerase (DNA) I | 7150 | ENSG00000198900 |
| 4 | P50991 | NA | CCT4 | chaperonin containing TCP1, subunit 4 (delta) | 10575 | ENSG00000115484 |

  
  

| **Database:Transcrription Target      &nbspName:hsa\_TGANTCA\_V$AP1\_C      &nbspID:DB\_ID:2402** | | | | | | |
| --- | --- | --- | --- | --- | --- | --- |
| C=1092; O=8; E=3.39; R=2.36; rawP=0.0193; adjP=0.0444 | | | | | | |
| Index | UserID | Value | Gene Symbol | Gene Name | EntrezGene | Ensembl |
| 1 | P49321 | NA | NASP | nuclear autoantigenic sperm protein (histone-binding) | 4678 | ENSG00000132780 |
| 2 | P07951 | NA | TPM2 | tropomyosin 2 (beta) | 7169 | ENSG00000198467 |
| 3 | Q9BQE3 | NA | TUBA1C | tubulin, alpha 1c | 84790 | ENSG00000167553 |
| 4 | P37802 | NA | TAGLN2 | transgelin 2 | 8407 | ENSG00000158710 |
| 5 | Q15758 | NA | SLC1A5 | solute carrier family 1 (neutral amino acid transporter), member 5 | 6510 | ENSG00000105281 |
| 6 | P18206 | NA | VCL | vinculin | 7414 | ENSG00000035403 |
| 7 | P68104 | NA | EEF1A1 | eukaryotic translation elongation factor 1 alpha 1 | 1915 | ENSG00000156508 |
| 8 | O60610 | NA | DIAPH1 | diaphanous homolog 1 (Drosophila) | 1729 | ENSG00000131504 |

  
  

| **Database:Transcrription Target      &nbspName:hsa\_V$TCF11MAFG\_01      &nbspID:DB\_ID:2040** | | | | | | |
| --- | --- | --- | --- | --- | --- | --- |
| C=202; O=3; E=0.63; R=4.78; rawP=0.0250; adjP=0.0561 | | | | | | |
| Index | UserID | Value | Gene Symbol | Gene Name | EntrezGene | Ensembl |
| 1 | Q16881 | NA | TXNRD1 | thioredoxin reductase 1 | 7296 | ENSG00000198431 |
| 2 | Q9BQE3 | NA | TUBA1C | tubulin, alpha 1c | 84790 | ENSG00000167553 |
| 3 | Q06830 | NA | PRDX1 | peroxiredoxin 1 | 5052 | ENSG00000117450 |

  
  

| **Database:Transcrription Target      &nbspName:hsa\_MGGAAGTG\_V$GABP\_B      &nbspID:DB\_ID:2418** | | | | | | |
| --- | --- | --- | --- | --- | --- | --- |
| C=741; O=6; E=2.30; R=2.61; rawP=0.0272; adjP=0.0596 | | | | | | |
| Index | UserID | Value | Gene Symbol | Gene Name | EntrezGene | Ensembl |
| 1 | P63241 | NA | EIF5A | eukaryotic translation initiation factor 5A | 1984 | ENSG00000132507 |
| 2 | Q92598 | NA | HSPH1 | heat shock 105kDa/110kDa protein 1 | 10808 | ENSG00000120694 |
| 3 | Q99832 | NA | CCT7 | chaperonin containing TCP1, subunit 7 (eta) | 10574 | ENSG00000135624 |
| 4 | P49321 | NA | NASP | nuclear autoantigenic sperm protein (histone-binding) | 4678 | ENSG00000132780 |
| 5 | O60610 | NA | DIAPH1 | diaphanous homolog 1 (Drosophila) | 1729 | ENSG00000131504 |
| 6 | P02786 | NA | TFRC | transferrin receptor (p90, CD71) | 7037 | ENSG00000072274 |

  
  

| **Database:Transcrription Target      &nbspName:hsa\_V$MYCMAX\_B      &nbspID:DB\_ID:2051** | | | | | | |
| --- | --- | --- | --- | --- | --- | --- |
| C=257; O=3; E=0.80; R=3.76; rawP=0.0458; adjP=0.0638 | | | | | | |
| Index | UserID | Value | Gene Symbol | Gene Name | EntrezGene | Ensembl |
| 1 | P63241 | NA | EIF5A | eukaryotic translation initiation factor 5A | 1984 | ENSG00000132507 |
| 2 | O60610 | NA | DIAPH1 | diaphanous homolog 1 (Drosophila) | 1729 | ENSG00000131504 |
| 3 | Q9UK76 | NA | HN1 | hematological and neurological expressed 1 | 51155 | ENSG00000189159 |

  
  

| **Database:Transcrription Target      &nbspName:hsa\_V$YY1\_Q6      &nbspID:DB\_ID:2268** | | | | | | |
| --- | --- | --- | --- | --- | --- | --- |
| C=236; O=3; E=0.73; R=4.09; rawP=0.0371; adjP=0.0638 | | | | | | |
| Index | UserID | Value | Gene Symbol | Gene Name | EntrezGene | Ensembl |
| 1 | P67809 | NA | YBX1 | Y box binding protein 1 | 4904 | ENSG00000065978 |
| 2 | P49321 | NA | NASP | nuclear autoantigenic sperm protein (histone-binding) | 4678 | ENSG00000132780 |
| 3 | P68104 | NA | EEF1A1 | eukaryotic translation elongation factor 1 alpha 1 | 1915 | ENSG00000156508 |

  
  

| **Database:Transcrription Target      &nbspName:hsa\_V$ICSBP\_Q6      &nbspID:DB\_ID:2210** | | | | | | |
| --- | --- | --- | --- | --- | --- | --- |
| C=245; O=3; E=0.76; R=3.94; rawP=0.0407; adjP=0.0638 | | | | | | |
| Index | UserID | Value | Gene Symbol | Gene Name | EntrezGene | Ensembl |
| 1 | P63241 | NA | EIF5A | eukaryotic translation initiation factor 5A | 1984 | ENSG00000132507 |
| 2 | P00352 | NA | ALDH1A1 | aldehyde dehydrogenase 1 family, member A1 | 216 | ENSG00000165092 |
| 3 | P11387 | NA | TOP1 | topoisomerase (DNA) I | 7150 | ENSG00000198900 |

  
  

| **Database:Transcrription Target      &nbspName:hsa\_V$HLF\_01      &nbspID:DB\_ID:2030** | | | | | | |
| --- | --- | --- | --- | --- | --- | --- |
| C=250; O=3; E=0.78; R=3.86; rawP=0.0428; adjP=0.0638 | | | | | | |
| Index | UserID | Value | Gene Symbol | Gene Name | EntrezGene | Ensembl |
| 1 | P13639 | NA | EEF2 | eukaryotic translation elongation factor 2 | 1938 | ENSG00000167658 |
| 2 | P00352 | NA | ALDH1A1 | aldehyde dehydrogenase 1 family, member A1 | 216 | ENSG00000165092 |
| 3 | P11387 | NA | TOP1 | topoisomerase (DNA) I | 7150 | ENSG00000198900 |

  
  

| **Database:Transcrription Target      &nbspName:hsa\_V$HNF4\_Q6      &nbspID:DB\_ID:2331** | | | | | | |
| --- | --- | --- | --- | --- | --- | --- |
| C=253; O=3; E=0.79; R=3.82; rawP=0.0441; adjP=0.0638 | | | | | | |
| Index | UserID | Value | Gene Symbol | Gene Name | EntrezGene | Ensembl |
| 1 | Q9BQE3 | NA | TUBA1C | tubulin, alpha 1c | 84790 | ENSG00000167553 |
| 2 | P37802 | NA | TAGLN2 | transgelin 2 | 8407 | ENSG00000158710 |
| 3 | Q8TEX9 | NA | IPO4 | importin 4 | 79711 | ENSG00000196497 |

  
  

| **Database:Transcrription Target      &nbspName:hsa\_V$AREB6\_02      &nbspID:DB\_ID:2079** | | | | | | |
| --- | --- | --- | --- | --- | --- | --- |
| C=249; O=3; E=0.77; R=3.88; rawP=0.0424; adjP=0.0638 | | | | | | |
| Index | UserID | Value | Gene Symbol | Gene Name | EntrezGene | Ensembl |
| 1 | Q9BQE3 | NA | TUBA1C | tubulin, alpha 1c | 84790 | ENSG00000167553 |
| 2 | P37802 | NA | TAGLN2 | transgelin 2 | 8407 | ENSG00000158710 |
| 3 | O75369 | NA | FLNB | filamin B, beta | 2317 | ENSG00000136068 |

  
  

| **Database:Transcrription Target      &nbspName:hsa\_V$OCT\_Q6      &nbspID:DB\_ID:2271** | | | | | | |
| --- | --- | --- | --- | --- | --- | --- |
| C=252; O=3; E=0.78; R=3.83; rawP=0.0436; adjP=0.0638 | | | | | | |
| Index | UserID | Value | Gene Symbol | Gene Name | EntrezGene | Ensembl |
| 1 | P00352 | NA | ALDH1A1 | aldehyde dehydrogenase 1 family, member A1 | 216 | ENSG00000165092 |
| 2 | Q16555 | NA | DPYSL2 | dihydropyrimidinase-like 2 | 1808 | ENSG00000092964 |
| 3 | P11387 | NA | TOP1 | topoisomerase (DNA) I | 7150 | ENSG00000198900 |

  
  

| **Database:Transcrription Target      &nbspName:hsa\_RYTTCCTG\_V$ETS2\_B      &nbspID:DB\_ID:2415** | | | | | | |
| --- | --- | --- | --- | --- | --- | --- |
| C=1060; O=7; E=3.29; R=2.13; rawP=0.0455; adjP=0.0638 | | | | | | |
| Index | UserID | Value | Gene Symbol | Gene Name | EntrezGene | Ensembl |
| 1 | P63241 | NA | EIF5A | eukaryotic translation initiation factor 5A | 1984 | ENSG00000132507 |
| 2 | P37802 | NA | TAGLN2 | transgelin 2 | 8407 | ENSG00000158710 |
| 3 | Q15758 | NA | SLC1A5 | solute carrier family 1 (neutral amino acid transporter), member 5 | 6510 | ENSG00000105281 |
| 4 | P49321 | NA | NASP | nuclear autoantigenic sperm protein (histone-binding) | 4678 | ENSG00000132780 |
| 5 | P07951 | NA | TPM2 | tropomyosin 2 (beta) | 7169 | ENSG00000198467 |
| 6 | P18206 | NA | VCL | vinculin | 7414 | ENSG00000035403 |
| 7 | O75369 | NA | FLNB | filamin B, beta | 2317 | ENSG00000136068 |

  
  

| **Database:Transcrription Target      &nbspName:hsa\_V$NGFIC\_01      &nbspID:DB\_ID:2018** | | | | | | |
| --- | --- | --- | --- | --- | --- | --- |
| C=250; O=3; E=0.78; R=3.86; rawP=0.0428; adjP=0.0638 | | | | | | |
| Index | UserID | Value | Gene Symbol | Gene Name | EntrezGene | Ensembl |
| 1 | P63241 | NA | EIF5A | eukaryotic translation initiation factor 5A | 1984 | ENSG00000132507 |
| 2 | O60610 | NA | DIAPH1 | diaphanous homolog 1 (Drosophila) | 1729 | ENSG00000131504 |
| 3 | P67936 | NA | TPM4 | tropomyosin 4 | 7171 | ENSG00000167460 |

  
  

| **Database:Transcrription Target      &nbspName:hsa\_V$MAX\_01      &nbspID:DB\_ID:1924** | | | | | | |
| --- | --- | --- | --- | --- | --- | --- |
| C=256; O=3; E=0.80; R=3.77; rawP=0.0454; adjP=0.0638 | | | | | | |
| Index | UserID | Value | Gene Symbol | Gene Name | EntrezGene | Ensembl |
| 1 | P02786 | NA | TFRC | transferrin receptor (p90, CD71) | 7037 | ENSG00000072274 |
| 2 | P11387 | NA | TOP1 | topoisomerase (DNA) I | 7150 | ENSG00000198900 |
| 3 | P11586 | NA | MTHFD1 | methylenetetrahydrofolate dehydrogenase (NADP+ dependent) 1, methenyltetrahydrofolate cyclohydrolase, formyltetrahydrofolate synthetase | 4522 | ENSG00000100714 |

  
  

| **Database:Transcrription Target      &nbspName:hsa\_V$ATF3\_Q6      &nbspID:DB\_ID:2156** | | | | | | |
| --- | --- | --- | --- | --- | --- | --- |
| C=243; O=3; E=0.75; R=3.97; rawP=0.0399; adjP=0.0638 | | | | | | |
| Index | UserID | Value | Gene Symbol | Gene Name | EntrezGene | Ensembl |
| 1 | P13639 | NA | EEF2 | eukaryotic translation elongation factor 2 | 1938 | ENSG00000167658 |
| 2 | P41567 | NA | EIF1 | eukaryotic translation initiation factor 1 | 10209 | ENSG00000173812 |
| 3 | P11387 | NA | TOP1 | topoisomerase (DNA) I | 7150 | ENSG00000198900 |

  
  

| **Database:Transcrription Target      &nbspName:hsa\_V$ATF1\_Q6      &nbspID:DB\_ID:2205** | | | | | | |
| --- | --- | --- | --- | --- | --- | --- |
| C=228; O=3; E=0.71; R=4.24; rawP=0.0340; adjP=0.0638 | | | | | | |
| Index | UserID | Value | Gene Symbol | Gene Name | EntrezGene | Ensembl |
| 1 | P13639 | NA | EEF2 | eukaryotic translation elongation factor 2 | 1938 | ENSG00000167658 |
| 2 | P67936 | NA | TPM4 | tropomyosin 4 | 7171 | ENSG00000167460 |
| 3 | P11387 | NA | TOP1 | topoisomerase (DNA) I | 7150 | ENSG00000198900 |

  
  

| **Database:Transcrription Target      &nbspName:hsa\_V$PAX4\_03      &nbspID:DB\_ID:2066** | | | | | | |
| --- | --- | --- | --- | --- | --- | --- |
| C=246; O=3; E=0.76; R=3.93; rawP=0.0411; adjP=0.0638 | | | | | | |
| Index | UserID | Value | Gene Symbol | Gene Name | EntrezGene | Ensembl |
| 1 | P12268 | NA | IMPDH2 | IMP (inosine 5'-monophosphate) dehydrogenase 2 | 3615 | ENSG00000178035 |
| 2 | P62906 | NA | RPL10A | ribosomal protein L10a | 4736 | ENSG00000198755 |
| 3 | Q16555 | NA | DPYSL2 | dihydropyrimidinase-like 2 | 1808 | ENSG00000092964 |

  
  

| **Database:Transcrription Target      &nbspName:hsa\_V$CREB\_02      &nbspID:DB\_ID:1918** | | | | | | |
| --- | --- | --- | --- | --- | --- | --- |
| C=248; O=3; E=0.77; R=3.89; rawP=0.0419; adjP=0.0638 | | | | | | |
| Index | UserID | Value | Gene Symbol | Gene Name | EntrezGene | Ensembl |
| 1 | P63241 | NA | EIF5A | eukaryotic translation initiation factor 5A | 1984 | ENSG00000132507 |
| 2 | P37802 | NA | TAGLN2 | transgelin 2 | 8407 | ENSG00000158710 |
| 3 | P13639 | NA | EEF2 | eukaryotic translation elongation factor 2 | 1938 | ENSG00000167658 |

  
  

| **Database:Transcrription Target      &nbspName:hsa\_V$E2F\_Q3\_01      &nbspID:DB\_ID:2295** | | | | | | |
| --- | --- | --- | --- | --- | --- | --- |
| C=231; O=3; E=0.72; R=4.18; rawP=0.0351; adjP=0.0638 | | | | | | |
| Index | UserID | Value | Gene Symbol | Gene Name | EntrezGene | Ensembl |
| 1 | P49736 | NA | MCM2 | minichromosome maintenance complex component 2 | 4171 | ENSG00000073111 |
| 2 | P49321 | NA | NASP | nuclear autoantigenic sperm protein (histone-binding) | 4678 | ENSG00000132780 |
| 3 | P11387 | NA | TOP1 | topoisomerase (DNA) I | 7150 | ENSG00000198900 |

  
  

| **Database:Transcrription Target      &nbspName:hsa\_V$AP1\_01      &nbspID:DB\_ID:2160** | | | | | | |
| --- | --- | --- | --- | --- | --- | --- |
| C=256; O=3; E=0.80; R=3.77; rawP=0.0454; adjP=0.0638 | | | | | | |
| Index | UserID | Value | Gene Symbol | Gene Name | EntrezGene | Ensembl |
| 1 | P37802 | NA | TAGLN2 | transgelin 2 | 8407 | ENSG00000158710 |
| 2 | P18206 | NA | VCL | vinculin | 7414 | ENSG00000035403 |
| 3 | P68104 | NA | EEF1A1 | eukaryotic translation elongation factor 1 alpha 1 | 1915 | ENSG00000156508 |

  
  

| **Database:Transcrription Target      &nbspName:hsa\_V$HIF1\_Q3      &nbspID:DB\_ID:2273** | | | | | | |
| --- | --- | --- | --- | --- | --- | --- |
| C=225; O=3; E=0.70; R=4.29; rawP=0.0329; adjP=0.0638 | | | | | | |
| Index | UserID | Value | Gene Symbol | Gene Name | EntrezGene | Ensembl |
| 1 | P67809 | NA | YBX1 | Y box binding protein 1 | 4904 | ENSG00000065978 |
| 2 | P02786 | NA | TFRC | transferrin receptor (p90, CD71) | 7037 | ENSG00000072274 |
| 3 | O00425 | NA | IGF2BP3 | insulin-like growth factor 2 mRNA binding protein 3 | 10643 | ENSG00000136231 |

  
  

| **Database:Transcrription Target      &nbspName:hsa\_V$USF\_Q6\_01      &nbspID:DB\_ID:2272** | | | | | | |
| --- | --- | --- | --- | --- | --- | --- |
| C=237; O=3; E=0.74; R=4.08; rawP=0.0375; adjP=0.0638 | | | | | | |
| Index | UserID | Value | Gene Symbol | Gene Name | EntrezGene | Ensembl |
| 1 | P63241 | NA | EIF5A | eukaryotic translation initiation factor 5A | 1984 | ENSG00000132507 |
| 2 | P67809 | NA | YBX1 | Y box binding protein 1 | 4904 | ENSG00000065978 |
| 3 | O60610 | NA | DIAPH1 | diaphanous homolog 1 (Drosophila) | 1729 | ENSG00000131504 |

  
  

| **Database:Transcrription Target      &nbspName:hsa\_V$BACH1\_01      &nbspID:DB\_ID:2146** | | | | | | |
| --- | --- | --- | --- | --- | --- | --- |
| C=256; O=3; E=0.80; R=3.77; rawP=0.0454; adjP=0.0638 | | | | | | |
| Index | UserID | Value | Gene Symbol | Gene Name | EntrezGene | Ensembl |
| 1 | P37802 | NA | TAGLN2 | transgelin 2 | 8407 | ENSG00000158710 |
| 2 | O60610 | NA | DIAPH1 | diaphanous homolog 1 (Drosophila) | 1729 | ENSG00000131504 |
| 3 | P68104 | NA | EEF1A1 | eukaryotic translation elongation factor 1 alpha 1 | 1915 | ENSG00000156508 |

  
  

| **Database:Transcrription Target      &nbspName:hsa\_V$SOX9\_B1      &nbspID:DB\_ID:2076** | | | | | | |
| --- | --- | --- | --- | --- | --- | --- |
| C=232; O=3; E=0.72; R=4.16; rawP=0.0355; adjP=0.0638 | | | | | | |
| Index | UserID | Value | Gene Symbol | Gene Name | EntrezGene | Ensembl |
| 1 | P49321 | NA | NASP | nuclear autoantigenic sperm protein (histone-binding) | 4678 | ENSG00000132780 |
| 2 | Q16555 | NA | DPYSL2 | dihydropyrimidinase-like 2 | 1808 | ENSG00000092964 |
| 3 | P11387 | NA | TOP1 | topoisomerase (DNA) I | 7150 | ENSG00000198900 |

  
  

| **Database:Transcrription Target      &nbspName:hsa\_V$NERF\_Q2      &nbspID:DB\_ID:2163** | | | | | | |
| --- | --- | --- | --- | --- | --- | --- |
| C=243; O=3; E=0.75; R=3.97; rawP=0.0399; adjP=0.0638 | | | | | | |
| Index | UserID | Value | Gene Symbol | Gene Name | EntrezGene | Ensembl |
| 1 | P18206 | NA | VCL | vinculin | 7414 | ENSG00000035403 |
| 2 | P02786 | NA | TFRC | transferrin receptor (p90, CD71) | 7037 | ENSG00000072274 |
| 3 | Q9UK76 | NA | HN1 | hematological and neurological expressed 1 | 51155 | ENSG00000189159 |

  
  

| **Database:Transcrription Target      &nbspName:hsa\_V$NF1\_Q6      &nbspID:DB\_ID:1982** | | | | | | |
| --- | --- | --- | --- | --- | --- | --- |
| C=255; O=3; E=0.79; R=3.79; rawP=0.0449; adjP=0.0638 | | | | | | |
| Index | UserID | Value | Gene Symbol | Gene Name | EntrezGene | Ensembl |
| 1 | P67809 | NA | YBX1 | Y box binding protein 1 | 4904 | ENSG00000065978 |
| 2 | P17174 | NA | GOT1 | glutamic-oxaloacetic transaminase 1, soluble (aspartate aminotransferase 1) | 2805 | ENSG00000120053 |
| 3 | Q06830 | NA | PRDX1 | peroxiredoxin 1 | 5052 | ENSG00000117450 |

  
  

| **Database:Transcrription Target      &nbspName:hsa\_V$STAT1\_02      &nbspID:DB\_ID:2143** | | | | | | |
| --- | --- | --- | --- | --- | --- | --- |
| C=248; O=3; E=0.77; R=3.89; rawP=0.0419; adjP=0.0638 | | | | | | |
| Index | UserID | Value | Gene Symbol | Gene Name | EntrezGene | Ensembl |
| 1 | P56192 | NA | MARS | methionyl-tRNA synthetase | 4141 | ENSG00000166986 |
| 2 | P49321 | NA | NASP | nuclear autoantigenic sperm protein (histone-binding) | 4678 | ENSG00000132780 |
| 3 | P41091 | NA | EIF2S3 | eukaryotic translation initiation factor 2, subunit 3 gamma, 52kDa | 1968 | ENSG00000130741 |

  
  

| **Database:Transcrription Target      &nbspName:hsa\_V$HIF1\_Q5      &nbspID:DB\_ID:2120** | | | | | | |
| --- | --- | --- | --- | --- | --- | --- |
| C=242; O=3; E=0.75; R=3.99; rawP=0.0395; adjP=0.0638 | | | | | | |
| Index | UserID | Value | Gene Symbol | Gene Name | EntrezGene | Ensembl |
| 1 | P67809 | NA | YBX1 | Y box binding protein 1 | 4904 | ENSG00000065978 |
| 2 | P41567 | NA | EIF1 | eukaryotic translation initiation factor 1 | 10209 | ENSG00000173812 |
| 3 | O00425 | NA | IGF2BP3 | insulin-like growth factor 2 mRNA binding protein 3 | 10643 | ENSG00000136231 |

  
  

| **Database:Transcrription Target      &nbspName:hsa\_V$BACH2\_01      &nbspID:DB\_ID:2141** | | | | | | |
| --- | --- | --- | --- | --- | --- | --- |
| C=263; O=3; E=0.82; R=3.67; rawP=0.0485; adjP=0.0652 | | | | | | |
| Index | UserID | Value | Gene Symbol | Gene Name | EntrezGene | Ensembl |
| 1 | Q9BQE3 | NA | TUBA1C | tubulin, alpha 1c | 84790 | ENSG00000167553 |
| 2 | P37802 | NA | TAGLN2 | transgelin 2 | 8407 | ENSG00000158710 |
| 3 | P68104 | NA | EEF1A1 | eukaryotic translation elongation factor 1 alpha 1 | 1915 | ENSG00000156508 |

  
  

| **Database:Transcrription Target      &nbspName:hsa\_V$CACCCBINDINGFACTOR\_Q6      &nbspID:DB\_ID:2220** | | | | | | |
| --- | --- | --- | --- | --- | --- | --- |
| C=264; O=3; E=0.82; R=3.66; rawP=0.0489; adjP=0.0652 | | | | | | |
| Index | UserID | Value | Gene Symbol | Gene Name | EntrezGene | Ensembl |
| 1 | P63241 | NA | EIF5A | eukaryotic translation initiation factor 5A | 1984 | ENSG00000132507 |
| 2 | P68104 | NA | EEF1A1 | eukaryotic translation elongation factor 1 alpha 1 | 1915 | ENSG00000156508 |
| 3 | P41567 | NA | EIF1 | eukaryotic translation initiation factor 1 | 10209 | ENSG00000173812 |

  
  

| **Database:Transcrription Target      &nbspName:hsa\_V$CREB\_Q2      &nbspID:DB\_ID:1968** | | | | | | |
| --- | --- | --- | --- | --- | --- | --- |
| C=261; O=3; E=0.81; R=3.70; rawP=0.0476; adjP=0.0652 | | | | | | |
| Index | UserID | Value | Gene Symbol | Gene Name | EntrezGene | Ensembl |
| 1 | P37802 | NA | TAGLN2 | transgelin 2 | 8407 | ENSG00000158710 |
| 2 | P41567 | NA | EIF1 | eukaryotic translation initiation factor 1 | 10209 | ENSG00000173812 |
| 3 | P67936 | NA | TPM4 | tropomyosin 4 | 7171 | ENSG00000167460 |

  
  

| **Database:Transcrription Target      &nbspName:hsa\_V$EGR\_Q6      &nbspID:DB\_ID:2283** | | | | | | |
| --- | --- | --- | --- | --- | --- | --- |
| C=270; O=3; E=0.84; R=3.58; rawP=0.0517; adjP=0.0679 | | | | | | |
| Index | UserID | Value | Gene Symbol | Gene Name | EntrezGene | Ensembl |
| 1 | P34932 | NA | HSPA4 | heat shock 70kDa protein 4 | 3308 | ENSG00000170606 |
| 2 | O60610 | NA | DIAPH1 | diaphanous homolog 1 (Drosophila) | 1729 | ENSG00000131504 |
| 3 | P67936 | NA | TPM4 | tropomyosin 4 | 7171 | ENSG00000167460 |

  
  

| **Database:Transcrription Target      &nbspName:hsa\_AACTTT\_UNKNOWN      &nbspID:DB\_ID:1851** | | | | | | |
| --- | --- | --- | --- | --- | --- | --- |
| C=1841; O=10; E=5.72; R=1.75; rawP=0.0565; adjP=0.0732 | | | | | | |
| Index | UserID | Value | Gene Symbol | Gene Name | EntrezGene | Ensembl |
| 1 | Q92598 | NA | HSPH1 | heat shock 105kDa/110kDa protein 1 | 10808 | ENSG00000120694 |
| 2 | Q9BQE3 | NA | TUBA1C | tubulin, alpha 1c | 84790 | ENSG00000167553 |
| 3 | P12268 | NA | IMPDH2 | IMP (inosine 5'-monophosphate) dehydrogenase 2 | 3615 | ENSG00000178035 |
| 4 | P53701 | NA | HCCS | holocytochrome c synthase | 3052 | ENSG00000004961 |
| 5 | P18206 | NA | VCL | vinculin | 7414 | ENSG00000035403 |
| 6 | P00352 | NA | ALDH1A1 | aldehyde dehydrogenase 1 family, member A1 | 216 | ENSG00000165092 |
| 7 | O60610 | NA | DIAPH1 | diaphanous homolog 1 (Drosophila) | 1729 | ENSG00000131504 |
| 8 | Q8TEX9 | NA | IPO4 | importin 4 | 79711 | ENSG00000196497 |
| 9 | P11387 | NA | TOP1 | topoisomerase (DNA) I | 7150 | ENSG00000198900 |
| 10 | Q9UK76 | NA | HN1 | hematological and neurological expressed 1 | 51155 | ENSG00000189159 |

  
  
  
  

---

WebGestalt is currently developed and maintained by Jing Wang and Bing Zhang at the  Zhang Lab. Other people who have made significant contribution to the project include Dexter Duncan, Stefan Kirov, Zhiao Shi, and Jay Snoddy.  
  
**Funding credits:** NIH/NIAAA (U01 AA016662, U01 AA013512); NIH/NIDA (P01 DA015027); NIH/NIMH (P50 MH078028, P50 MH096972); NIH/NCI (U24 CA159988); NIH/NIGMS (R01 GM088822).
