## Supplementary material for "PGRMC1 phosphorylation and cell plasticity 1: glycolysis, mitochondria, tumor growth": File S8: final_wiki_geneset_file_1439190848.html

Anchored HTML File of EIDs

|  |  |
| --- | --- |
|  | WEB-based GEne SeT AnaLysis Toolkit |
|  | ***Translating gene lists into biological insights...*** |

---

  

| **Database:Wikipathways pathway      &nbspName:Translation Factors      &nbspID:WP107** | | | | | | |
| --- | --- | --- | --- | --- | --- | --- |
| C=51; O=6; E=0.16; R=37.88; rawP=1.14e-08; adjP=9.12e-08 | | | | | | |
| Index | UserID | Value | Gene Symbol | Gene Name | EntrezGene | Ensembl |
| 1 | P63241 | NA | EIF5A | eukaryotic translation initiation factor 5A | 1984 | ENSG00000132507 |
| 2 | P13639 | NA | EEF2 | eukaryotic translation elongation factor 2 | 1938 | ENSG00000167658 |
| 3 | P41091 | NA | EIF2S3 | eukaryotic translation initiation factor 2, subunit 3 gamma, 52kDa | 1968 | ENSG00000130741 |
| 4 | Q15056 | NA | EIF4H | eukaryotic translation initiation factor 4H | 7458 | ENSG00000106682 |
| 5 | P68104 | NA | EEF1A1 | eukaryotic translation elongation factor 1 alpha 1 | 1915 | ENSG00000156508 |
| 6 | P41567 | NA | EIF1 | eukaryotic translation initiation factor 1 | 10209 | ENSG00000173812 |

  
  

| **Database:Wikipathways pathway      &nbspName:Parkin-Ubiquitin Proteasomal System pathway      &nbspID:WP2359** | | | | | | |
| --- | --- | --- | --- | --- | --- | --- |
| C=70; O=4; E=0.22; R=18.40; rawP=6.64e-05; adjP=0.0003 | | | | | | |
| Index | UserID | Value | Gene Symbol | Gene Name | EntrezGene | Ensembl |
| 1 | Q9BQE3 | NA | TUBA1C | tubulin, alpha 1c | 84790 | ENSG00000167553 |
| 2 | P34932 | NA | HSPA4 | heat shock 70kDa protein 4 | 3308 | ENSG00000170606 |
| 3 | Q13885 | NA | TUBB2A | tubulin, beta 2A class IIa | 7280 | ENSG00000137267 |
| 4 | P68371 | NA | TUBB4B | tubulin, beta 4B class IVb | 10383 | ENSG00000188229 |

  
  

| **Database:Wikipathways pathway      &nbspName:Lymphocyte TarBase      &nbspID:WP2004** | | | | | | |
| --- | --- | --- | --- | --- | --- | --- |
| C=533; O=8; E=1.66; R=4.83; rawP=0.0002; adjP=0.0005 | | | | | | |
| Index | UserID | Value | Gene Symbol | Gene Name | EntrezGene | Ensembl |
| 1 | Q16881 | NA | TXNRD1 | thioredoxin reductase 1 | 7296 | ENSG00000198431 |
| 2 | P25685 | NA | DNAJB1 | DnaJ (Hsp40) homolog, subfamily B, member 1 | 3337 | ENSG00000132002 |
| 3 | P07951 | NA | TPM2 | tropomyosin 2 (beta) | 7169 | ENSG00000198467 |
| 4 | P00491 | NA | PNP | purine nucleoside phosphorylase | 4860 | ENSG00000198805 |
| 5 | Q6PIU2 | NA | NCEH1 | neutral cholesterol ester hydrolase 1 | 57552 | ENSG00000144959 |
| 6 | P12081 | NA | HARS | histidyl-tRNA synthetase | 3035 | ENSG00000170445 |
| 7 | Q8TEX9 | NA | IPO4 | importin 4 | 79711 | ENSG00000196497 |
| 8 | P67936 | NA | TPM4 | tropomyosin 4 | 7171 | ENSG00000167460 |

  
  

| **Database:Wikipathways pathway      &nbspName:Pathogenic Escherichia coli infection      &nbspID:WP2272** | | | | | | |
| --- | --- | --- | --- | --- | --- | --- |
| C=64; O=3; E=0.20; R=15.09; rawP=0.0010; adjP=0.0020 | | | | | | |
| Index | UserID | Value | Gene Symbol | Gene Name | EntrezGene | Ensembl |
| 1 | Q9BQE3 | NA | TUBA1C | tubulin, alpha 1c | 84790 | ENSG00000167553 |
| 2 | Q13885 | NA | TUBB2A | tubulin, beta 2A class IIa | 7280 | ENSG00000137267 |
| 3 | P68371 | NA | TUBB4B | tubulin, beta 4B class IVb | 10383 | ENSG00000188229 |

  
  

| **Database:Wikipathways pathway      &nbspName:Muscle cell TarBase      &nbspID:WP2005** | | | | | | |
| --- | --- | --- | --- | --- | --- | --- |
| C=424; O=6; E=1.32; R=4.56; rawP=0.0020; adjP=0.0032 | | | | | | |
| Index | UserID | Value | Gene Symbol | Gene Name | EntrezGene | Ensembl |
| 1 | Q16881 | NA | TXNRD1 | thioredoxin reductase 1 | 7296 | ENSG00000198431 |
| 2 | P25685 | NA | DNAJB1 | DnaJ (Hsp40) homolog, subfamily B, member 1 | 3337 | ENSG00000132002 |
| 3 | P07951 | NA | TPM2 | tropomyosin 2 (beta) | 7169 | ENSG00000198467 |
| 4 | Q6PIU2 | NA | NCEH1 | neutral cholesterol ester hydrolase 1 | 57552 | ENSG00000144959 |
| 5 | Q8TEX9 | NA | IPO4 | importin 4 | 79711 | ENSG00000196497 |
| 6 | P67936 | NA | TPM4 | tropomyosin 4 | 7171 | ENSG00000167460 |

  
  

| **Database:Wikipathways pathway      &nbspName:Epithelium TarBase      &nbspID:WP2002** | | | | | | |
| --- | --- | --- | --- | --- | --- | --- |
| C=340; O=5; E=1.06; R=4.73; rawP=0.0041; adjP=0.0055 | | | | | | |
| Index | UserID | Value | Gene Symbol | Gene Name | EntrezGene | Ensembl |
| 1 | P25685 | NA | DNAJB1 | DnaJ (Hsp40) homolog, subfamily B, member 1 | 3337 | ENSG00000132002 |
| 2 | P00491 | NA | PNP | purine nucleoside phosphorylase | 4860 | ENSG00000198805 |
| 3 | P12081 | NA | HARS | histidyl-tRNA synthetase | 3035 | ENSG00000170445 |
| 4 | Q8TEX9 | NA | IPO4 | importin 4 | 79711 | ENSG00000196497 |
| 5 | P67936 | NA | TPM4 | tropomyosin 4 | 7171 | ENSG00000167460 |

  
  

| **Database:Wikipathways pathway      &nbspName:Squamous cell TarBase      &nbspID:WP2006** | | | | | | |
| --- | --- | --- | --- | --- | --- | --- |
| C=150; O=3; E=0.47; R=6.44; rawP=0.0114; adjP=0.0130 | | | | | | |
| Index | UserID | Value | Gene Symbol | Gene Name | EntrezGene | Ensembl |
| 1 | Q16881 | NA | TXNRD1 | thioredoxin reductase 1 | 7296 | ENSG00000198431 |
| 2 | P25685 | NA | DNAJB1 | DnaJ (Hsp40) homolog, subfamily B, member 1 | 3337 | ENSG00000132002 |
| 3 | P00491 | NA | PNP | purine nucleoside phosphorylase | 4860 | ENSG00000198805 |

  
  

| **Database:Wikipathways pathway      &nbspName:Leukocyte TarBase      &nbspID:WP2003** | | | | | | |
| --- | --- | --- | --- | --- | --- | --- |
| C=160; O=3; E=0.50; R=6.04; rawP=0.0135; adjP=0.0135 | | | | | | |
| Index | UserID | Value | Gene Symbol | Gene Name | EntrezGene | Ensembl |
| 1 | Q16881 | NA | TXNRD1 | thioredoxin reductase 1 | 7296 | ENSG00000198431 |
| 2 | P25685 | NA | DNAJB1 | DnaJ (Hsp40) homolog, subfamily B, member 1 | 3337 | ENSG00000132002 |
| 3 | P00491 | NA | PNP | purine nucleoside phosphorylase | 4860 | ENSG00000198805 |

  
  
  
  

---

WebGestalt is currently developed and maintained by Jing Wang and Bing Zhang at the  Zhang Lab. Other people who have made significant contribution to the project include Dexter Duncan, Stefan Kirov, Zhiao Shi, and Jay Snoddy.  
  
**Funding credits:** NIH/NIAAA (U01 AA016662, U01 AA013512); NIH/NIDA (P01 DA015027); NIH/NIMH (P50 MH078028, P50 MH096972); NIH/NCI (U24 CA159988); NIH/NIGMS (R01 GM088822).
